# Three ALS genes regulate expression of the MHC class II antigen presentation pathway

**DOI:** 10.1101/2022.05.11.489680

**Authors:** Binkai Chi, Muhammet M. Öztürk, Christina L. Paraggio, Claudia E. Leonard, Maria E. Sanita, Mahtab Dastpak, Jeremy D. O’Connell, Jordan A. Coady, Jiuchun Zhang, Steven P. Gygi, Rodrigo Lopez-Gonzalez, Robin Reed

## Abstract

Here we report that the major histocompatibility complex II (MHC II) antigen presentation pathway is regulated by the ALS-causative genes, FUS, TAF15, or MATR3. Of >6000 proteins detected by quantitative mass spectrometry, the subunits of the MHC II heterodimer, HLA-DR, were the top 2 downregulated proteins in HeLa knock outs (KO) of these ALS genes, but not the related gene, EWSR1. Moreover, CD74, which is the 3^rd^ essential component of HLA-DR, was downregulated in the 3 KOs. We show that the downregulations are due to loss of CIITA, a transcription factor dedicated to expression of MHC II genes. Thus, our results reveal the 1^st^ shared cellular pathway regulated by multiple ALS genes, and this pathway is ALS genes -> CIITA -> MHC II genes. We obtained the same results in HMC3 cells, a microglia cell line, showing that loss of the MHC II pathway extends to an ALS-relevant cell type in the central nervous system (CNS). Furthermore, the MHC II pathway is downregulated in hematopoietic progenitor cells (HPCs) bearing the ALS FUS^R495X^ mutation. This observation may be highly significant to ALS pathogenesis as HPCs give rise to a multitude of CNS-specific and systemic immune cells, both of which have known or suspected roles in ALS. Together, our data raise the possibility that loss of the MHC II pathway in a large range of immune cells results in global failure of the immune system to protect motor neurons from damage that leads to the disease. Consequently, CIITA and the other genes in the MHC II pathway may be important new therapeutic targets for the disease.

## Introduction

Amyotrophic lateral sclerosis (ALS) is a rapidly progressive fatal motor neuron disease with no effective treatment. Heritable ALS is caused by mutations in ∼30 genes, and many cellular pathways are disrupted as a consequence. It is a daunting prospect to understand how so many pathways with no obvious connection to one another culminate in the same disease (Taylor et al. 2016; Mejzini et al. 2019; Kim et al. 2020). These pathways include protein homeostasis, gene expression, mitochondrial function, intracellular transport, endoplasmic reticulum stress, and several others (Taylor et al. 2016; Mejzini et al. 2019; Kim et al. 2020). Thus, a major challenge in ALS is to decipher how and when the disrupted pathways contribute to the progression of ALS, as different pathways are known or are likely to contribute at different stages of the disease (Brites and Vaz 2014; Hooten et al. 2015; Cherry et al. 2014; Gerbino et al. 2020; Rudnick et al. 2017). ALS is even more complex because although motor neurons are the cell type directly impacted, supporting cells in the central nervous system (CNS), such as microglia, oligodendrocytes, and astrocytes also play critical roles (Benarroch 2021; Yamanaka and Komine 2018; Clarke and Patani 2020). Unraveling the contributions of these glial cells to ALS is a key challenge, with models positing both detrimental and protective roles (Blackburn et al. 2009; Geloso et al. 2017; Ridler 2018; Spiller et al. 2018; Rossi et al. 2018; Béland et al. 2020). Furthermore many types of systemic immune cells throughout the body play important roles in ALS, by for example, migrating into the CNS and secreting cytokines or neurotrophic factors that can be pro- or anti-inflammatory (Chiu et al. 2008; Banerjee et al. 2008; Liu et al. 2007; Tanabe and Yamashita 2018; Ortega et al. 2020; Vitkovic et al. 2001; Ousman and Kubes 2012). The data suggest that the immune cells are protective early in the disease course, but then become detrimental as the disease progresses, and proinflammatory cytokines are thought to contribute to disease progression (Béland et al. 2020).

Among the important insights into ALS disease mechanisms was the observation that >1/3 of the ALS genes encodes RNA/DNA binding proteins (Kapeli et al. 2017), most of which have known roles in gene expression. We recently characterized the proteome of a cellular complex we termed the RNAP polymerase II (RNAP II)/U1 snRNP machinery (Chi et al. 2018b), and we discovered that this machinery is a central hub for most of the RNA/DNA binding ALS proteins. The machinery also houses several spinal muscular atrophy (SMA)-causative proteins, and these also play roles in gene expression (Chi et al. 2018b). In prior studies, we obtained additional evidence that ALS and SMA are linked to one another at the molecular level through direct interactions of proteins involved in each disease (Yamazaki et al. 2012; Chi et al. 2018a). Thus, the finding that ALS/SMA proteins share the same machinery is not surprising considering that the numerous steps in gene expression are extensively coupled to one other both physically and functionally (Maniatis and Reed 2002; Reed and Hurt 2002; Rosonina and Blencowe 2002). The coupling involves virtually all steps in gene expression including transcription, capping, splicing, 3’ end formation, mRNA export, and mRNA degradation, and proteins that function in these coupled processes are components of the RNAP II/U1 snRNP machinery (Chi et al. 2018b). Thus, a critical question is whether the ALS proteins associated with this machinery have downstream roles in different cellular pathways or converge on a common pathway(s).

Among the ALS genes in the RNAP II/U1 snRNP machinery are FUS, TAF15, and MATR3 (Chi et al. 2018b). These proteins are structurally similar to one another and to EWSR1, a putative ALS protein (Schwartz et al. 2015). FUS, TAF15 and EWSR1 comprise the FET family of proteins which were 1^st^ discovered for their roles in transcription when it was found that the transcription activation domains of the FET family members fuse to other genes to generate potent oncogenic fusion proteins (Bertolotti et al. 1999; Shing et al. 2003; Kovar 2011). More recently, FET family members were reported to play roles in other steps of gene expression, such as splicing, though these roles may be indirect via coupling of transcription to splicing and other gene expression steps (Schwartz et al. 2015). Recently, we generated CRISPR-knock out (KO) lines (Chi et al. 2018a) and siRNA knockdowns (KDs) of the FET family members and MATR3 in HeLa cells and in a microglia-like cell line (HMC3s), respectively. Strikingly, we found that the MHC class II antigen presentation pathway was strongly downregulated in FUS, TAF15 or MATR3 KO/KDs. We furthermore discovered that loss of the MHC II pathway in the ALS KO/KDs was due to downregulation of CIITA, the master transcriptional regulator of the pathway.

Importantly, the MHC II pathway was also downregulated in hematopoietic progenitor cells (HPCs) bearing an ALS patient mutation, providing evidence for a direct link between the disease and the MHC II pathway. Indeed, this pathway may be highly pertinent to ALS because it is essential in all MHC II cells, which are a major component of the immune system, playing key roles both directly in the CNS and indirectly via the systemic circulatory system. Collectively, our data are consistent with a model in which the ALS genes regulate CIITA which in turn regulates the MHC II pathway. Our data furthermore suggest CIITA, with its tight restriction to the MHC II pathway, as well as the other genes in the MHC II pathway, as therapeutic targets for multiple forms of ALS.

## Results and Discussion

### Multiple ALS genes regulate expression of immune response pathways

To investigate the functions of the ALS-associated FET family members and MATR3, we used quantitative mass spectrometry to analyze protein expression in our 4 KO HeLa lines. Over 6,000 total proteins were identified (Table S1). Examination of the significantly dysregulated proteins (p-value <0.05, Table S2) by Gene Set Enrichment Analysis (GSEA) revealed striking differences in the GSEA negative (Table S3) versus positive (Table S4) data for the FUS, TAF15, and MATR3 KOs. Specifically, immune pathways were highly enriched in the GSEA negative (Fig. 1A, Table S3) but not in the GSEA positive data (Fig. 1B, Table S4). The opposite was observed in the EWSR1 KO. Immune pathways were highly enriched in the GSEA positive (Fig. 1B, Table S4) but not detected in the GSEA negative (Fig 1A, Table S4) data. Six of the immune pathways in the GSEA negative data were shared among the FUS, TAF15, and MATR3 KOs, and 2 of these were shared with the EWSR1 KO but were in the GSEA positive data (Fig. 1C). The observation that multiple immune pathways are shared in the GSEA negative data of FUS, TAF15 and MATR3 KOs raises the possibility that these 3 ALS proteins impact the disease via downregulation of immune pathways.

**Figure 1.**
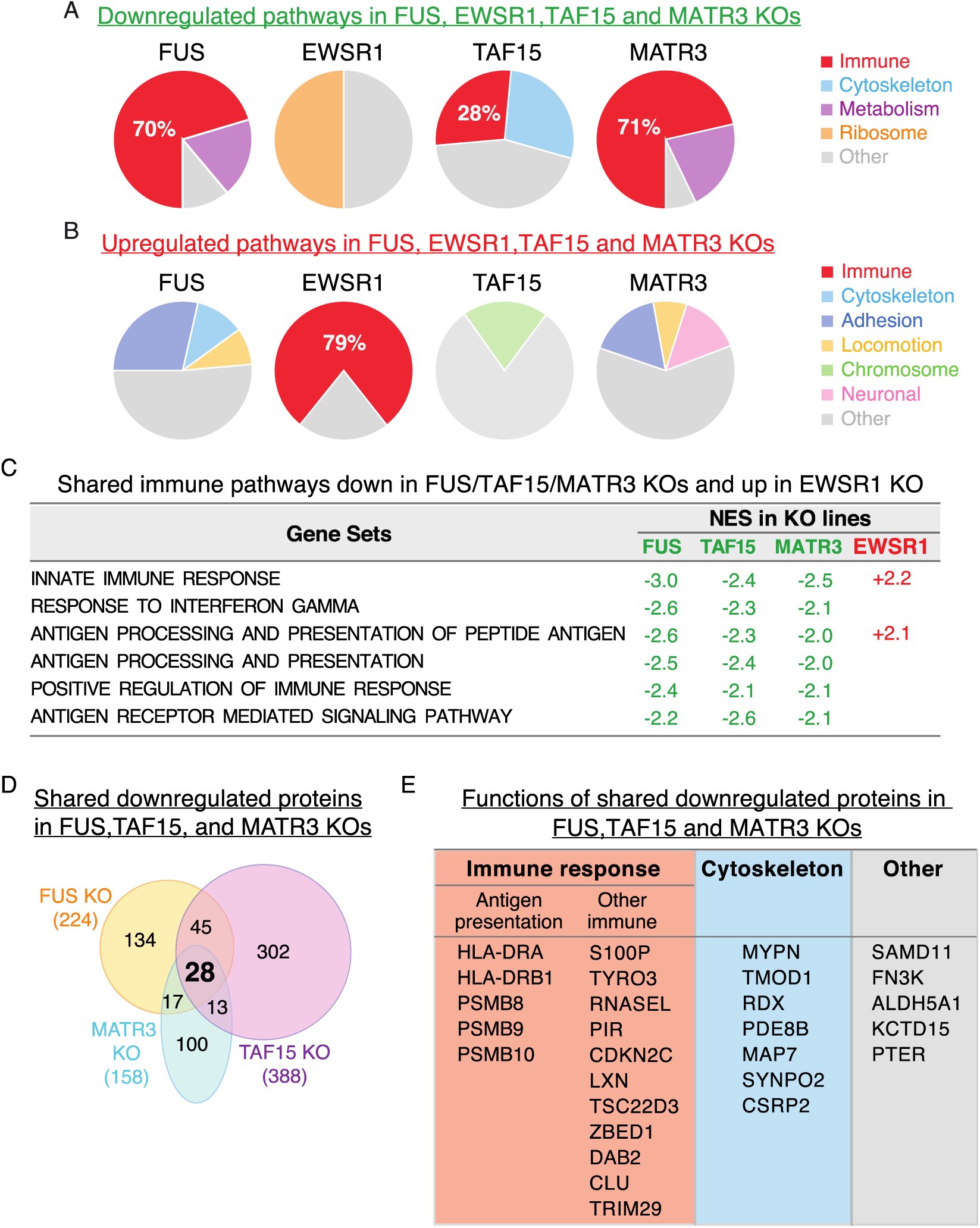
Immune pathways are top downregulated in FUS, TAF15, and MATR3 ALS gene knockout HeLa cells. A. Pie charts showing percentage of downregulated immune pathways in red in each KO. B. Immune pathways (shown in red) are top upregulated in EWSR1 KO. Other categories of downregulated pathways in each KO are indicated. C. Immune pathways shared among all 4 KO lines. Green and red font shows Nominal Enrichment Score (NES) of down or upregulated pathways, respectively. D. Venn diagram showing the 28 shared downregulated proteins in FUS, TAF15, and MATR3 KOs. E. List of the 28 shared proteins showing their general functions.

We next analyzed the data at the level of individual proteins (Fig. 1D). Of the significantly downregulated proteins in the FUS, TAF15 and MATR3 KOs, expression of 28 proteins was decreased in all 3 lines (Fig. 1D and see Supplemental Table S5). Among these, more than half play roles in the innate immune response, consistent with the GSEA results (Fig. 1E). Notably, we observed downregulation of multiple proteins that function in antigen presentation in all 3 KOs (Fig. 1E). These include HLA-DRA and HLA-DRB1, which play key roles in MHC class II antigen presentation (Shackelford et al. 1982; Karakikes et al. 2012) (see below). In addition, all 3 immunoproteasome-specific proteases, PSMBs 8-10, which function in MHC class I antigen presentation (Basler et al. 2013; McCarthy and Weinberg 2015; Rock et al. 2004) were downregulated. Other downregulated proteins involved in antigen presentation include DAB2 and CLU (Figliuolo da Paz et al. 2020; Podleśny-Drabiniok et al. 2020), both of which play roles in phagocytosis (Fig. 1E). Finally, additional proteins that function in immunity were downregulated, such as RNASEL and S100P (Fig. 1E) (Paludan et al. 2021; Kozlyuk et al. 2019). Together, these data indicate that FUS, TAF15 and MATR3 each play a key role in regulating the immune response.

### Regulation of MHC II antigen presentation genes by FUS, TAF15 and MATR3

The most striking of the downregulated immune proteins were HLA-DRA and HLA-DRB1, which were top downregulated proteins in all 3 KOs (Fig. 2A, B) but unchanged in the EWSR1 KO (Supplementary Table S5). HLA-DRA and HLA-DRB1 comprise the HLA-DR heterodimer, which is a cell surface receptor that presents antigens to T cells. These interactions initiate a wide range of immune responses including production of cytokines and antibodies (Yu et al. 1980; Shackelford et al. 1982; Parker 1993). The observation that *both* components of a heterodimer are downregulated and to such a great extent in the 3 KOs strongly indicates that HLA-DR regulation is an important new finding in the biology of FUS, TAF15 and MATR3 ALS genes.

**Figure 2.**
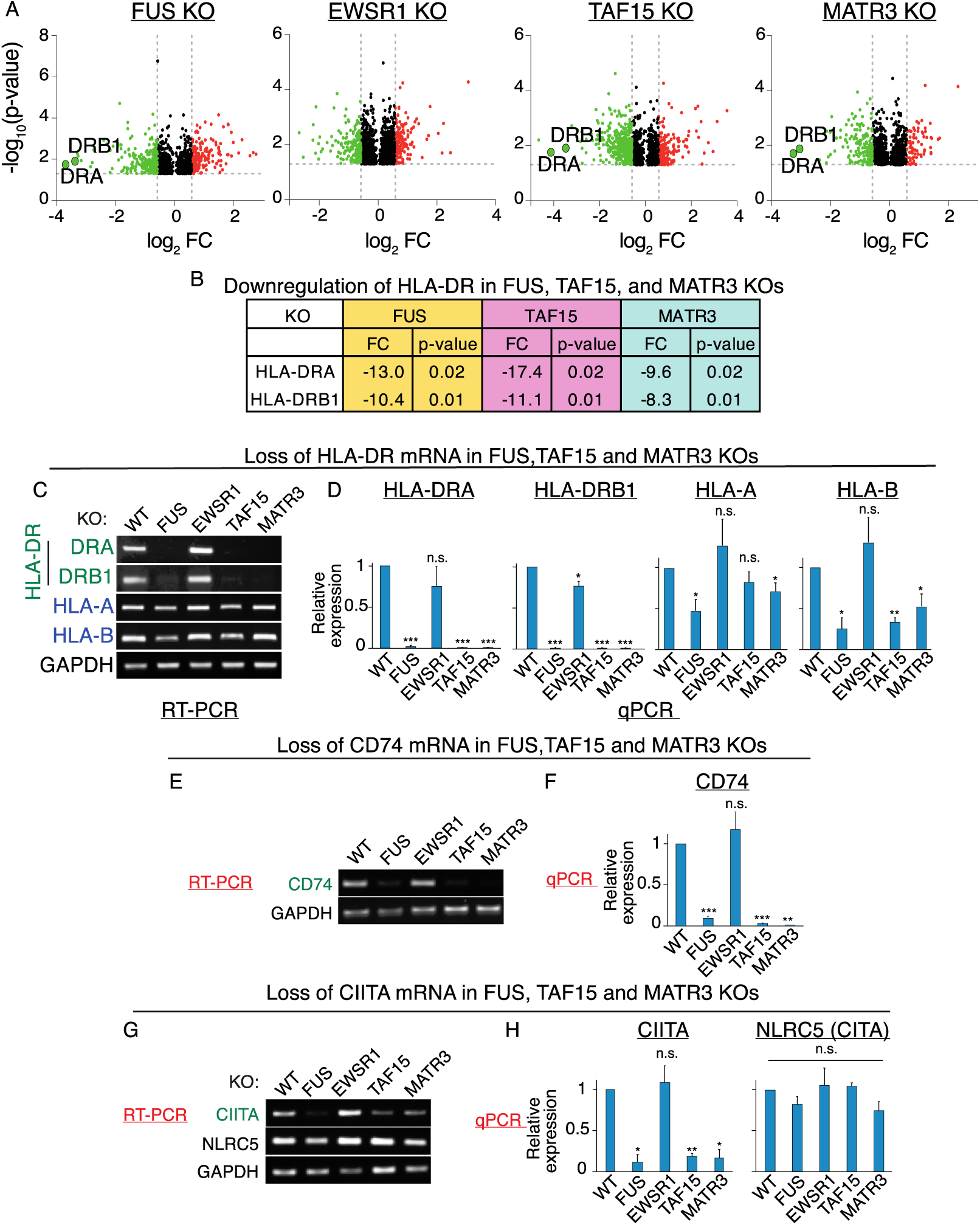
Downregulation of MHC II pathway genes in FUS, TAF15, and MATR3 KO HeLa cells. A. Volcano plots of proteins in HeLa KO lines. HLA-DRA and -DRB1 are indicated. B. Fold change and p-values of downregulated HLA-DRA and -DRB1 in the indicated KO lines. C, D. RT-PCR (C) and qPCR (D) from total RNA isolated from the indicated WT and KO lines for the HLA mRNAs shown. GAPDH was used as an internal control. E, F. Same as D and E, except CD74 was assayed by RT-PCR (E) and qPCR (F). G, E, Same as E and F except CIITA and NLRC5 were assayed by RT-PCR and qPCR (H).

To investigate whether mRNA levels of HLA-DRA and HLA-DRB1 were affected in the KOs, we carried out RT-PCR using total RNA from the wild type (WT) and KO HeLa lines. As shown in Fig. 2C, HLA-DRA and HLA-DRB1 mRNA levels were strongly reduced in FUS, TAF15 and MATR3 KOs but unaffected in the EWSR1 KO. In contrast, mRNA levels of MHC class I genes, such as HLA-A or B, were not significantly affected in any of the KOs (Fig. 2C). These results were confirmed by qPCR (Fig. 2D).

Based on our results with the HLA-DR heterodimer, we further examined the MHC II pathway, and the story became even more compelling. Specifically, RT-PCR revealed that CD74 mRNA levels were strongly decreased in the FUS, TAF15 and MATR3 KOs, but not affected in the EWSR1 KO (Fig. 2E), and qPCR confirmed these results (Fig. 2F). This is a particularly noteworthy result as CD74, which was previously known as the ‘HLA-DR gamma chain’, plays a critical role in MHC II antigen presentation by stabilizing HLA-DR immediately after its synthesis and then chaperoning it to the endosomal system for antigen processing and binding to HLA-DR (Karakikes et al. 2012; Schröder 2016). We conclude that 3 of the key components for MHC II antigen presentation are downregulated at the mRNA level in FUS, TAF15 and MATR3 KOs. We did not detect CD74 in our proteomic analysis, even in WT cells, possibly for technical reasons.

We next investigated mechanisms for downregulation of the HLA-DR heterodimer and CD74 in the KO lines. FUS, TAF15 and MATR3 have known roles in transcription (Bertolotti et al. 1999; Shing et al. 2003; Hibino et al. 2000), and we did not detect any splicing defects in the MHC II mRNAs. This result was not unexpected as different splicing factors recognize distinct sequence elements within pre-mRNAs and are therefore highly unlikely to mis-splice the same pre-mRNAs. Thus, we next examined expression of the transcription factor CIITA in the KO lines. CIITA is unique among transcription factors because it is tightly restricted to regulating expression of genes in the MHC II pathway (LeibundGut-Landmann et al. 2004; Reith et al. 2005). Strikingly, RT-PCR revealed that CIITA mRNA levels were decreased in FUS, TAF15, and MATR3, but not EWSR1, KOs (Fig. 2G). In contrast, the MHC Class I transcription regulator NLRC5 (Meissner et al. 2012) was not affected in any of the KOs (Fig. 2G). qPCR confirmed these results (Fig. 2H). To further examine the role of CIITA in the ALS KOs, we carried out addback studies by transfecting the FUS KO with myc-tagged CIITA or a myc-tagged negative control plasmid. As shown in Supplementary Fig. S1, the levels of the MHC II mRNAs were efficiently restored only by myc-tagged CIITA. We conclude that FUS regulates CIITA which in turn regulates expression of the MHC II pathway.

### Regulation of MHC II pathway by ALS genes in HMC3 microglia cell line

To gain insight into the potential importance of the MHC II pathway in ALS, we examined it in an ALS-relevant microglia cell type known as HMC3 (<u>H</u>uman <u>M</u>icroglial <u>C</u>lone 3) cells (Fig. 3). This line came from SV40-transformed embryonic microglial cells, and it was recently authenticated by the ATCC, which was our source for the line (ATCC®CRL-3304). The line was authenticated by several criteria including morphological characteristics, karyotyping, retention of antigenic features, and expression of the microglia markers IBA1 and CD68, as well as other specific microglial signatures such as P2RY12 and TMEM119 (Dello Russo et al. 2018).

**Figure 3.**
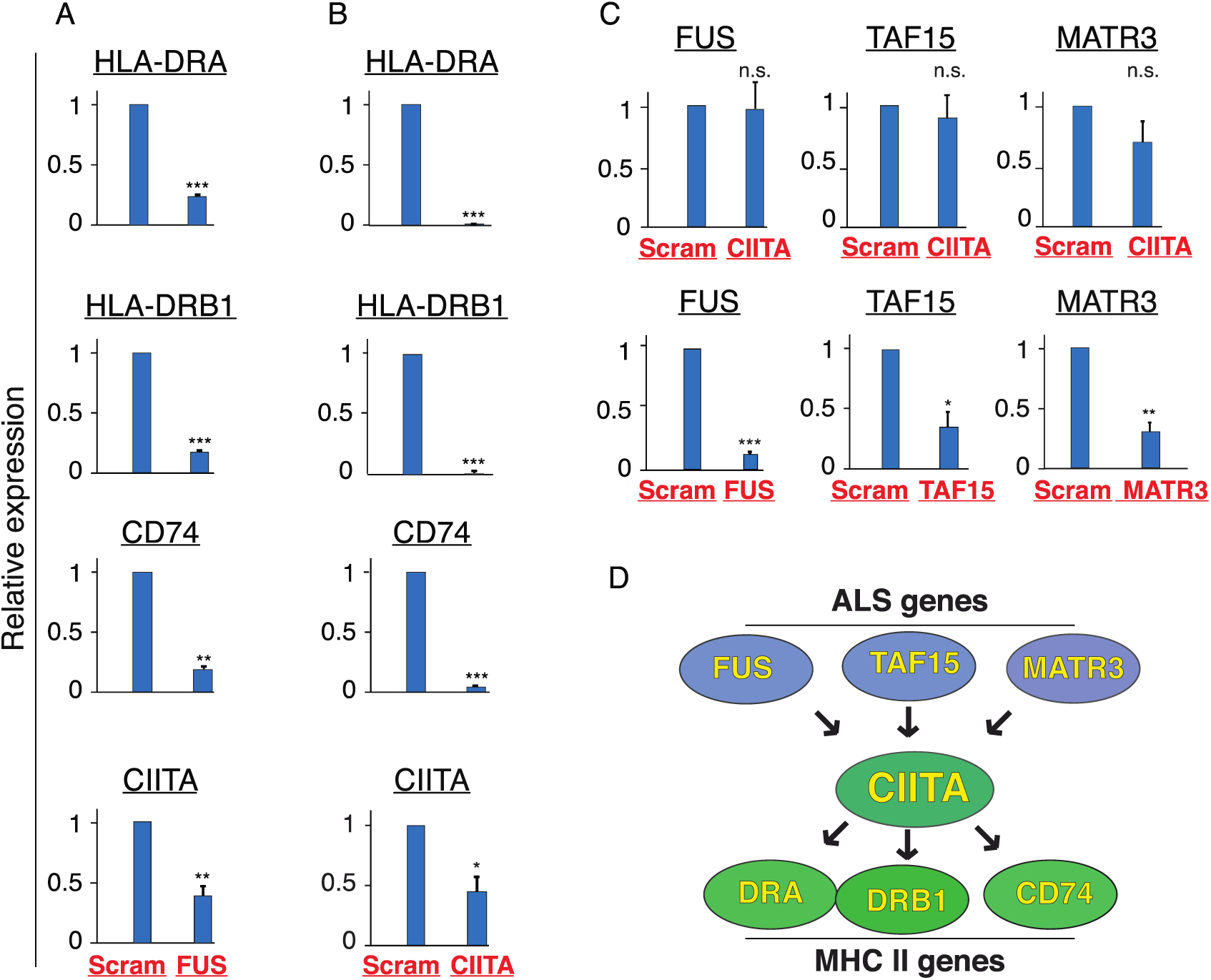
ALS genes are upstream in the MHC II pathway and regulate CIITA expression in HMC3 cells. A-C. HMC3 cells were knocked down using the indicated siRNAs (labeled at the bottom of the panels in red). Scrambled siRNAs were used as a negative control. mRNA levels for each MHC II gene in FUS KD (A) or CIITA KD (B) were assayed by qPCR. C. ALS gene mRNA levels are not affected by CIITA KD but are affected by their cognate siRNA. D. Model for regulation of MHC Class II antigen presentation pathway by ALS genes.

To assay the MHC II pathway in HMC3 cells, we used siRNA knockdowns (KDs) for targeting the ALS genes and CIITA. Scrambled siRNA was used as a negative control. In contrast to the microglia marker genes, our MHC II genes of interest, such as the HLA-DR heterodimer, are known to be expressed only at very low levels in HMC3s (Etemad et al. 2012). However, these genes are also known to be inducible by IFNγ in natural microglia as well as in HMC3s (Wijdeven et al. 2018). Based on this knowledge, we titrated IFNγ levels in HMC3s (Supplementary Fig. S2). This analysis confirmed low-levels of expression of HLA-DR mRNAs in the absence of IFNγ, and a dramatic increase it its levels with 3 ng/ml IFNγ consistent with previous reports (Giroux et al. 2003; Dello Russo et al. 2018). Slightly lower levels of HLA-DR mRNAs were detected when IFNγ levels were further increased (Supplementary Fig. S2). Thus, for the knockdown studies below, 3 ng/ml IFNγ was added to the culture media (see Methods). As shown in Fig. 3A, FUS KD resulted in decreased levels of the MHC II mRNAs, including both components of HLA-DR as well as CD74. CIITA KD also led to a significant decrease in the levels of the MHC II mRNAs (Fig. 3B). Similar results were obtained with the TAF15 and MATR3 KDs (Supplementary Fig. S3).

To further determine whether the ALS genes are upstream of CIITA in the MHC II pathway, we knocked down CIITA and then assayed the mRNA levels of FUS, TAF15 and MATR3 by qPCR. As shown in (Fig. 3C), neither mRNAs were affected by the CIITA KD whereas the targeting siRNAs knocked down their cognate mRNAs, consistent with our HeLa KO data showing that these ALS genes regulate CIITA mRNA levels and are thus upstream of CIITA in the MHC II pathway. There are several important implications of this set of findings regarding the MHC II pathway. As stated in the Introduction, one of the challenges in the ALS field is to understand how so many ALS-causative genes with distinct functions lead to the same disease. To our knowledge, our data provide the 1^st^ example in which multiple ALS genes regulate the same pathway (see schematic in Fig. 3D). Regulation of this pathway by ALS genes is especially noteworthy in a motor neuron disease because both the CNS and systemic MHC II antigen presentation pathways normally play critical roles in the CNS by protecting the neurons from damage.

### Evidence linking downregulation of the MHC II pathway to ALS

Although our findings contribute an important new facet to the biology of ALS genes via their role in regulating the MHC II pathway, a critical question is whether downregulation of these genes plays a role in ALS pathogenesis. To investigate this possibility, we 1^st^ examined patient spinal cords. However, we observed unacceptable levels of variability in both control and ALS subjects, possibly due to inherent difficulties with postmortem tissue. As another approach for determining whether the MHC II pathway plays a role in ALS, we CRISPR-edited human ES cells to harbor a heterozygous FUS^R495X^ mutation, generating 2 independent lines. This mutation causes a severe form of the disease because most of the C-terminal NLS is truncated due to the mutation (Bosco et al. 2010). Sanger sequencing and westerns showed that we successfully inserted the R495X mutation into one allele of the ES cells, and the truncated R495X and WT proteins are expressed at similar levels in the mutant lines (Fig. 4A and B, respectively). We next differentiated the FUS^R495X^ ES cells into hematopoietic progenitor cells (HPCs). We chose HPCs because they differentiate into many different MHC II cell types (Pouzolles et al. 2016; Lee and Hong 2020; Ferrari et al. 2021), and as mentioned in the Introduction, MHC II cell types such as microglia in the CNS and systemic immune cells are known or are thought to play important roles in ALS. As shown in Fig. 4C-E, we found that both the mutant (MT) and WT ES cells were efficiently differentiated into HPCs as revealed by cell-type specific markers. We next carried out qPCR to assay for the MHC II pathway genes. Excitingly, these data revealed a significant decrease in the levels of all the MHC II genes in the FUS^R495X^ HPCs but not in the FUS^R495X^ ES cells (Fig. 4F-I, Supplementary Fig. S4). These results are particularly noteworthy because of the sheer number of CNS and systemic immune cells expected to have disrupted expression of the MHC II pathway due to ALS-mutant HPCs. The systemic immune cells that arise from HPCs include dendritic cells, macrophages, monocytes, as well as CD4+ and CD8+ T cells, and all these cell types have been found in the brains of ALS patients (Engelhardt et al. 1993; Zondler et al. 2016; Graves et al. 2004; Holmøy 2008). The systemic cells enter the CNS as spinal cord infiltrates, and interactions between glial cells and the infiltrating MHC II cells are thought to be one mechanism by which they contribute to ALS pathogenesis (Puentes et al. 2016; Henkel et al. 2004; Antel et al. 2020; Béland et al. 2020; Liu and Wang 2017; Malaspina et al. 2015). Consistent with our results on the ALS genes reported here, C9orf72, mutation of which is the most common genetic cause of ALS, is required for proper systemic as well as CNS immune responses (Atanasio et al. 2016; O’Rourke et al. 2016). Previous studies of HIV patients shed further light on the importance of the systemic immune system in ALS. These studies reported that HIV patients are more likely to exhibit an ALS-like syndrome, and this condition improves after anti-HIV treatment (Satin and Bayat 2021; Bowen et al. 2016; MacGowan et al. 2001; Quevedo-Ramirez et al. 2020; Verma and Berger 2006). Relative to the studies of motor neurons and glial cells in ALS, much less work has been done on the systemic aspect of ALS. Downregulation of the MHC II pathway by mutant ALS genes provides a plausible explanation for why ALS is not only a CNS disease but also a systemic disease. Identifying therapeutics that are efficacious at specific stages of ALS in which the CNS/systemic immune system is impacted, either positively or negatively, would be invaluable for making advances in the disease.

**Figure 4.**
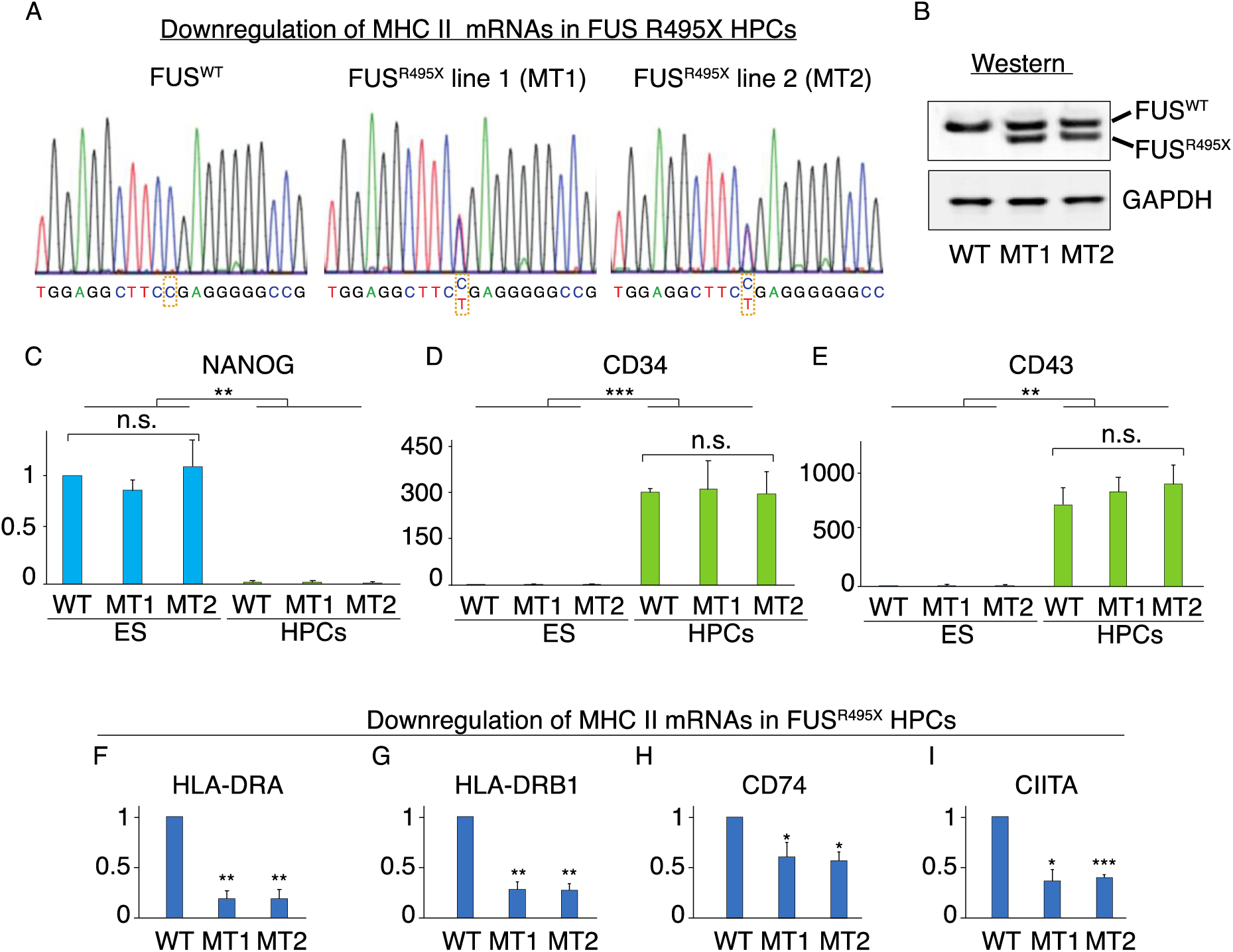
Linking downregulation of MHC II pathway to ALS. A. Sanger sequencing showing heterozygous FUS^R495X^ mutation in two independent ES lines. B. Westerns showing WT FUS and truncated FUS in MT lines. C-E. qPCRs of total RNAs from ES cells and HPCs confirming HPC differentiation. Pluripotency marker NANOG was expressed in ES cells but not in HPCs whereas HPC markers CD34 and CD43 were not expressed in ES cells but were expressed in HPCs. F-I. qPCR of total RNA isolated from HPCs to assay expression levels of MHC II pathway genes.

## Materials and Methods

### Plasmids and Antibodies

The myc-CIITA plasmid was obtained from Addgene (Plasmid #14650). The myc-PK plasmid was a gift from Dr. Gideon Dreyfuss (Siomi and Dreyfuss 1995). The GAPDH antibody was from Proteintech (Cat No. 60004-1-Ig). The FUS antibody was described (Yamazaki et al. 2012).

### Cell Culture

HeLa cells were grown in Dulbecco′s Modified Eagle′s Medium (DMEM, Invitrogen) plus 10% fetal bovine serum (FBS, Gibco) and 1% penicillin-streptomycin (Invitrogen). Generation of HeLa KO lines was described (Chi et al. 2018a). HMC3 cells (ATCC^®^ CRL-3304™) were cultured in EMEM (ATCC^®^ 30-2003™) plus 10% FBS (ATCC^®^ 30-2020™) following the ATCC manual. The H9 human ES cell line was cultured using the mTeSR1 medium (STEMCELL Technologies) plus 1% penicillin-streptomycin on hESC-Qualified Matrigel (Corning)-coated tissue culture plates. All cells were cultured in a 37°C humidified 5% CO_2_ incubator.

### siRNA Knockdown and Plasmid Transfection

HMC3 cells were cultured in a 6 well plate at 70% confluency for siRNA knockdown. ON-TARGETplus human siRNAs targeting FUS, TAF15, MATR3 (Uemura et al. 2017), CIITA or non-targeting siRNA (Horizon Discovery) were transfected into HMC3 cells using Lipofectamine RNAiMAX (Invitrogen) following manufacturer’s instructions. After 48 hr, the medium was supplemented with 3 ng/ml of IFNγ (Sigma). Cells were then incubated for 24 hr before harvesting for RNA extraction.

### Quantitative PCR

Total RNA was isolated from cells using Trizol (Invitrogen) according to the manufacturer’s manual. cDNA was synthesized using the UltraScript 2.0 cDNA Synthesis kit (PCR Biosystems), and qPCR was performed with gene-specific primer sets (Supplemental Table S6) and PowerUp™ SYBR™ Green Master Mix (Applied Biosystems). All qPCR analyses were carried out on an Applied Biosystems QuantStudio 7 Flex Cycler (Thermo Fisher Scientific), and relative expression values were calculated using the comparative C^T^ method.

### Whole-cell Proteomics and Gene Set Enrichment Analysis (GSEA)

For whole-cell proteomic analysis, wild type (WT) and KO HeLa cells were cultured in 150 mm dishes. Cells were harvested at 90% confluency, and whole cell lysates were used for quantitative mass spectrometry. Digested peptides were labeled by tandem mass tag (McAlister et al. 2012) for MS3 analysis using an Orbitrap Fusion mass spectrometer coupled to a Proxeon EASY-nLC 1000 liquid chromatography pump (Thermo Fisher Scientific). GSEA analyses were preformed using https://www.gsea-msigdb.org/gsea/index.jsp (version 4.0.3, gene set database C5.bp.v7.0). Pre-ranked gene lists with p-value less than 0.05 from the quantitative mass spectrometry data were used for GSEA.

### CRISPR Editing FUS^R495X^ in ES cells

Human ES cells (H9, WiCell Institute) were cultured in E8 medium (Chen et al. 2011) on Matrigel-coated tissue culture plates with daily medium change. The SpCas9 expression plasmid pET-Cas9-NLS-6xHis (Addgene plasmid # 62933) was transformed into *Rosetta™(DE3) pLysS* Competent Cells (Novagen). SpCas9 protein was purified as described (Zuris et al. 2015). The sgRNA was generated using the *GeneArt Precision gRNA Synthesis Kit* (Thermo Fisher Scientific) according to the manufacturer’s instructions. To create H9 cells harboring a heterozygous R495X mutation in FUS, 0.6 μg sgRNA targeting sequence GGGACCGTGGAGGCTTCCGA was incubated with 3 μg SpCas9 protein for 10 minutes at room temperature and electroporated into 2×10^5^ H9 cells along with a ssDNA oligo (tcgtcgtggtggcagaggaggctatgatcgaggcggctaccggggccg-cggcggggaccgtggaggcttcTgaggTggccggggtggtggggacagaggtggctttggccctggcaagatggattccaggtaagactttaaat). Mutants were identified by Illumina MiSeq and further confirmed by Sanger sequencing and westerns.

### HPC Differentiation

WT and MT ES lines were differentiated into HPCs using the STEMdiff Hematopoietic Kit (STEMCELL Technologies). To prepare for differentiation, ES cells were detached using ReLeSR (STEMCELL Technologies) and plated on Matrigel-coated 6 well plates to achieve ∼40 attached colonies/well 24 hr after seeding. Medium was changed according to the manufacturer’s manual. Supernatant HPCs were harvested at day 12 for Westerns or RNA extraction.

## Acknowledgments

We thank WiCell Research Institute for the ES cell line (WA09 (H9)). This work was funded by an NIH grant (NIGMS GM122524) and a Harvard Brain Science Initiative ALS Seed Grant to R.R. We are grateful to Dr. Oleg Butovsky for important input on the study.

## Author Contributions

BC and RR conceived the project. BC, MMO, CLP, CEL, MES, and MD carried out assays of the MHC II genes, JDO and SPG performed quantitative mass spectrometry, JAC participated in early stages of the work. JZ established the FUS^R495X^ ES lines. RL-G provided valuable advice and protocols for the study, and all authors contributed to the manuscript.

