## Supplementary Figures for "Three ALS genes regulate expression of the MHC class II antigen presentation pathway"

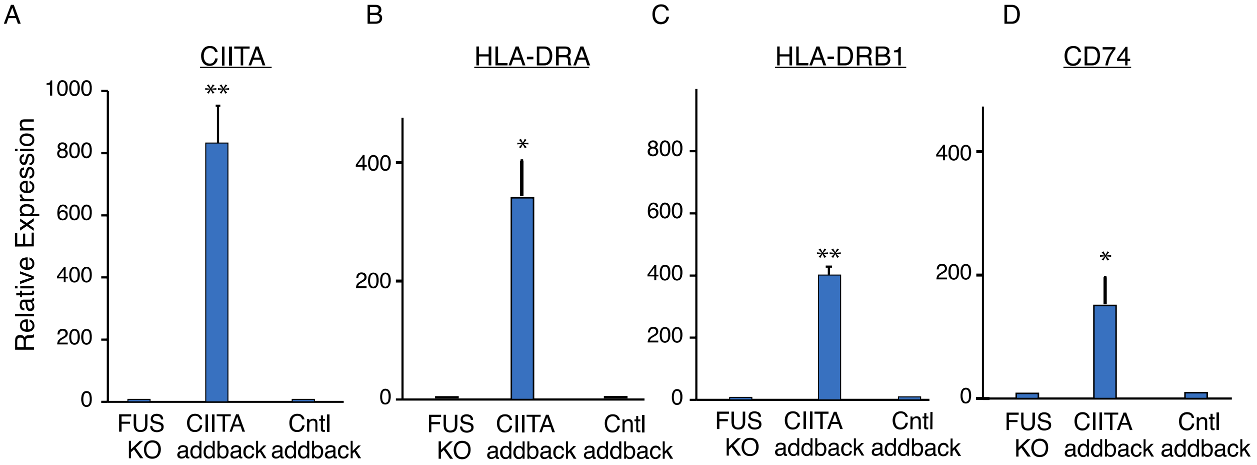


**Supplementary Figure S1. Addback of CIITA to FUS KO restores expression of MHC II genes.** A-D qPCR of the indicated MHC II mRNAs in the FUS KO or the indicated addbacks to the FUS KO. For addbacks, 10 ng of myc-CIITA or myc-PK plasmid was transfected into FUS KO cells cultured in 1 well of 6 well plate using Lipofectamine 3000 transfection kit (Invitrogen).

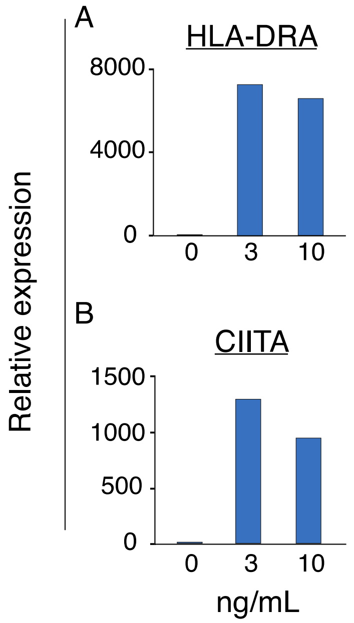


**Supplementary Figure S2. Titration of IFNγ in HMC3 cells to determine optimal levels for expression of MHC II pathway genes.** A, B. qPCR of HLA-DRA (A) and CIITA (B) mRNA levels in HMC3 cells treated with the indicated amounts of IFNγ.

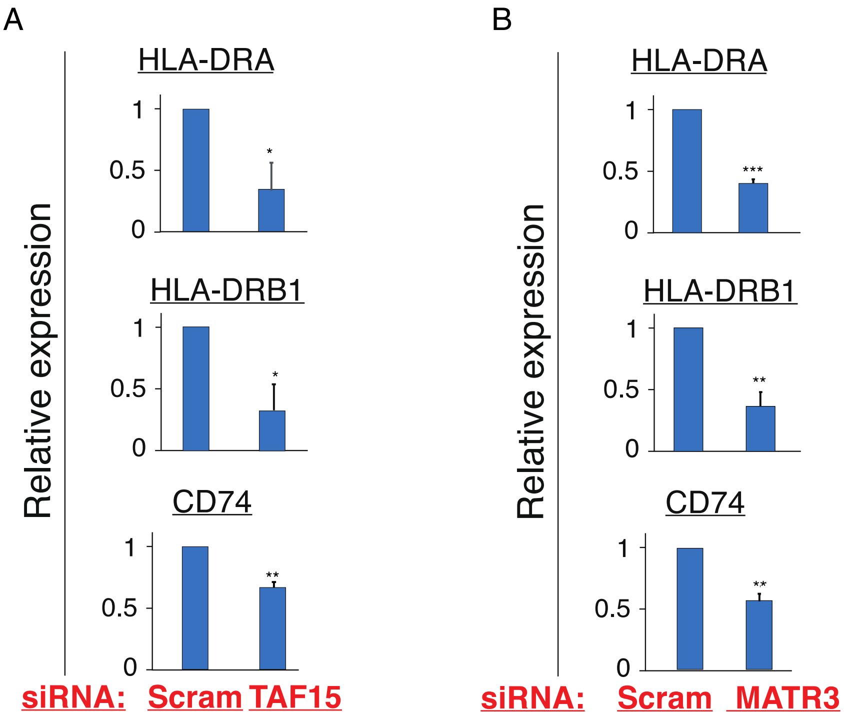


**Supplementary Figure S3. MHC II pathway is downregulated in TAF15 or MATR3 KD HMC3 cells.** mRNA levels were assayed by qPCR.

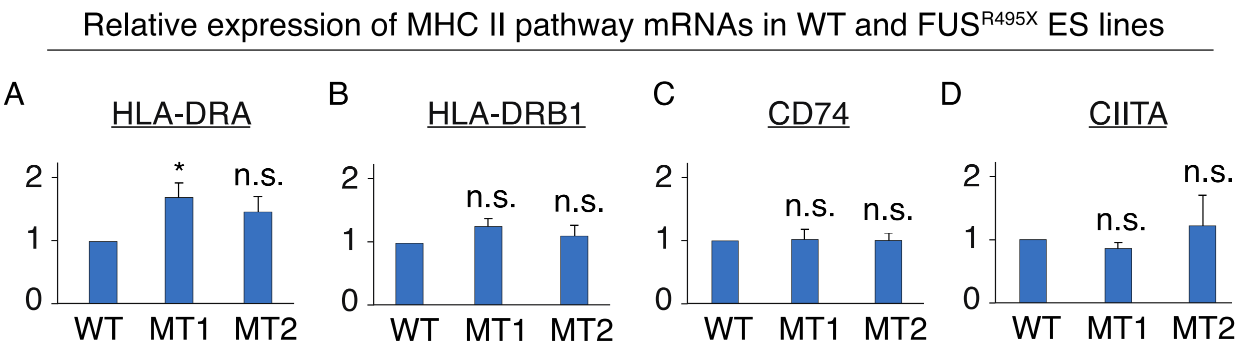


**Supplementary Figure S4. MHC II pathway is unaffected in FUS MT ES lines.** A-D. qPCR of mRNAs levels of MHC II pathway mRNAs in WT and FUS MT ES lines.
