## Supplementary Table S1 for "Three ALS genes regulate expression of the MHC class II antigen presentation pathway"

**Table S1. Total quantitative mass spectrometry of wild type (WT) and FUS, EWSR1, TAF15 and MATR3 knockout (KO) HeLa cell lines.** Duplicates of each line were used. Gene symbol, peptide number (#), relative amount of each protein, and mean value are shown.

| Gene Symbol | Peptide # | WT | | FUS KO | | EWSR1 KO | | TAF15 KO | | MATR3 KO | | Mean value | | | | |
|---|---|---|---|---|---|---|---|---|---|---|---|---|---|---|---|---|
| | | Repeat 1 | Repeat 2 | Repeat 1 | Repeat 2 | Repeat 1 | Repeat 2 | Repeat 1 | Repeat 2 | Repeat 1 | Repeat 2 | WT | FUS KO | EWSR1 KO | TAF15 KO | MATR3 KO |
| A2M | 5 | 12.86 | 11.43 | 20.91 | 20.55 | 6.28 | 5.73 | 2.94 | 2.61 | 8.51 | 8.20 | 12.14 | 20.73 | 6.01 | 2.77 | 8.35 |
| AAAS | 9 | 8.59 | 9.41 | 9.72 | 10.31 | 9.16 | 8.78 | 11.50 | 13.79 | 9.21 | 9.55 | 9.00 | 10.01 | 8.97 | 12.64 | 9.38 |
| AACS | 11 | 10.30 | 11.61 | 8.76 | 9.03 | 11.20 | 11.99 | 7.91 | 8.23 | 10.54 | 10.44 | 10.96 | 8.89 | 11.60 | 8.07 | 10.49 |
| AAGAB | 1 | 8.63 | 9.04 | 10.44 | 8.16 | 13.35 | 11.21 | 10.37 | 6.15 | 11.18 | 11.48 | 8.84 | 9.30 | 12.28 | 8.26 | 11.33 |
| AAK1 | 6 | 9.15 | 10.97 | 12.46 | 12.40 | 10.59 | 10.78 | 8.10 | 7.50 | 9.08 | 8.97 | 10.06 | 12.43 | 10.68 | 7.80 | 9.02 |
| AAMDC | 4 | 10.61 | 10.95 | 10.13 | 9.94 | 9.23 | 10.48 | 8.31 | 7.93 | 11.53 | 10.88 | 10.78 | 10.04 | 9.86 | 8.12 | 11.20 |
| AAMP | 5 | 10.50 | 10.69 | 11.09 | 10.40 | 8.83 | 8.96 | 10.39 | 9.46 | 10.22 | 9.46 | 10.59 | 10.75 | 8.89 | 9.92 | 9.84 |
| AAR2 | 2 | 11.14 | 13.08 | 8.16 | 7.99 | 7.68 | 9.00 | 9.45 | 10.75 | 11.80 | 10.94 | 12.11 | 8.08 | 8.34 | 10.10 | 11.37 |
| AARS | 74 | 9.61 | 10.67 | 7.97 | 7.66 | 12.35 | 12.80 | 9.45 | 9.06 | 10.34 | 10.11 | 10.14 | 7.81 | 12.58 | 9.25 | 10.22 |
| AARS2 | 14 | 11.46 | 9.26 | 8.51 | 10.22 | 9.50 | 8.92 | 10.36 | 9.99 | 11.08 | 10.69 | 10.36 | 9.37 | 9.21 | 10.18 | 10.88 |
| AARSD1 | 13 | 11.10 | 12.10 | 12.12 | 12.10 | 7.14 | 7.15 | 7.65 | 7.53 | 11.82 | 11.28 | 11.60 | 12.11 | 7.15 | 7.59 | 11.55 |
| AASDHPPT | 16 | 8.85 | 10.31 | 11.49 | 10.41 | 8.50 | 8.48 | 12.01 | 10.79 | 9.76 | 9.40 | 9.58 | 10.95 | 8.49 | 11.40 | 9.58 |
| AASS | 3 | 12.77 | 11.07 | 16.98 | 19.99 | 11.79 | 11.28 | 2.90 | 2.44 | 5.32 | 5.46 | 11.92 | 18.49 | 11.54 | 2.67 | 5.39 |
| AATF | 17 | 9.62 | 10.10 | 9.97 | 9.94 | 8.36 | 8.31 | 12.07 | 10.73 | 10.89 | 10.02 | 9.86 | 9.95 | 8.33 | 11.40 | 10.45 |
| ABCA13 | 4 | 9.65 | 10.18 | 10.17 | 10.66 | 10.04 | 10.49 | 9.08 | 9.08 | 10.43 | 10.23 | 9.91 | 10.41 | 10.26 | 9.08 | 10.33 |
| ABCB1 | 2 | 5.29 | 4.24 | 13.36 | 20.67 | 10.73 | 10.54 | 11.61 | 10.91 | 5.86 | 6.79 | 4.77 | 17.01 | 10.64 | 11.26 | 6.33 |
| ABCB10 | 1 | 13.03 | 5.94 | 5.65 | 7.43 | 10.76 | 11.25 | 9.64 | 9.62 | 11.92 | 14.76 | 9.49 | 6.54 | 11.01 | 9.63 | 13.34 |
| ABCB4 | 1 | 16.28 | 8.72 | 5.46 | 7.06 | 10.23 | 11.66 | 11.10 | 12.00 | 8.99 | 8.50 | 12.50 | 6.26 | 10.95 | 11.55 | 8.74 |
| ABCB6 | 8 | 11.79 | 7.80 | 8.27 | 10.17 | 11.67 | 9.82 | 7.65 | 6.92 | 12.13 | 13.79 | 9.79 | 9.22 | 10.74 | 7.28 | 12.96 |
| ABCB7 | 9 | 11.58 | 7.12 | 7.60 | 11.30 | 10.64 | 9.67 | 11.14 | 10.79 | 9.07 | 11.09 | 9.35 | 9.45 | 10.16 | 10.96 | 10.08 |
| ABCC1 | 10 | 11.77 | 9.05 | 8.29 | 9.94 | 9.07 | 9.39 | 9.13 | 7.98 | 12.18 | 13.20 | 10.41 | 9.11 | 9.23 | 8.56 | 12.69 |
| ABCC4 | 6 | 10.66 | 7.64 | 11.65 | 14.23 | 9.86 | 9.39 | 12.01 | 9.26 | 7.40 | 7.90 | 9.15 | 12.94 | 9.63 | 10.63 | 7.65 |
| ABCD1 | 12 | 11.40 | 6.19 | 6.22 | 8.26 | 12.85 | 11.11 | 10.62 | 11.21 | 9.85 | 12.29 | 8.79 | 7.24 | 11.98 | 10.91 | 11.07 |
| ABCD3 | 12 | 13.43 | 7.05 | 6.61 | 8.93 | 9.19 | 8.20 | 8.99 | 10.49 | 12.17 | 14.94 | 10.24 | 7.77 | 8.70 | 9.74 | 13.56 |
| ABCE1 | 36 | 9.23 | 9.88 | 10.81 | 9.61 | 9.15 | 9.51 | 10.71 | 11.87 | 9.69 | 9.53 | 9.55 | 10.21 | 9.33 | 11.29 | 9.61 |
| ABCF1 | 36 | 9.21 | 9.73 | 12.99 | 10.75 | 9.45 | 9.01 | 9.99 | 8.43 | 10.31 | 10.13 | 9.47 | 11.87 | 9.23 | 9.21 | 10.22 |
| ABCF2 | 45 | 9.09 | 10.59 | 11.94 | 11.62 | 7.22 | 7.34 | 11.08 | 12.08 | 9.33 | 9.71 | 9.84 | 11.78 | 7.28 | 11.58 | 9.52 |
| ABCF3 | 21 | 11.24 | 10.13 | 10.75 | 10.43 | 8.88 | 9.32 | 10.29 | 9.47 | 10.78 | 8.70 | 10.69 | 10.59 | 9.10 | 9.88 | 9.74 |
| ABHD10 | 8 | 10.78 | 9.31 | 10.77 | 10.99 | 10.66 | 9.68 | 9.80 | 8.85 | 9.34 | 9.84 | 10.04 | 10.88 | 10.17 | 9.32 | 9.59 |
| ABHD11 | 5 | 9.52 | 9.09 | 11.02 | 12.05 | 8.47 | 9.01 | 11.09 | 10.23 | 9.67 | 9.84 | 9.31 | 11.54 | 8.74 | 10.66 | 9.76 |
| ABHD12 | 2 | 13.22 | 6.84 | 6.72 | 11.78 | 11.23 | 10.84 | 8.94 | 8.52 | 9.81 | 12.09 | 10.03 | 9.25 | 11.04 | 8.73 | 10.95 |
| ABHD14B | 6 | 9.49 | 10.35 | 9.93 | 7.61 | 12.39 | 10.91 | 7.59 | 6.79 | 12.18 | 12.75 | 9.92 | 8.77 | 11.65 | 7.19 | 12.47 |
| ABHD16A | 2 | 11.07 | 9.86 | 9.66 | 9.88 | 9.09 | 9.99 | 9.30 | 12.80 | 8.81 | 9.54 | 10.46 | 9.77 | 9.54 | 11.05 | 9.18 |
| ABHD17A | 1 | 11.02 | 8.93 | 6.76 | 8.20 | 13.44 | 12.16 | 9.48 | 8.83 | 10.21 | 10.96 | 9.98 | 7.48 | 12.80 | 9.16 | 10.59 |
| ABHD4 | 1 | 9.13 | 11.00 | 9.00 | 12.25 | 10.08 | 11.94 | 8.76 | 6.77 | 9.74 | 11.33 | 10.07 | 10.62 | 11.01 | 7.77 | 10.53 |
| ABHD5 | 2 | 7.83 | 7.74 | 11.12 | 10.16 | 13.10 | 13.09 | 9.99 | 9.56 | 8.77 | 8.65 | 7.78 | 10.64 | 13.09 | 9.77 | 8.71 |
| ABHD6 | 2 | 10.10 | 7.12 | 9.09 | 9.33 | 10.84 | 9.87 | 10.20 | 12.99 | 11.08 | 9.38 | 8.61 | 9.21 | 10.35 | 11.59 | 10.23 |
| ABI1 | 9 | 9.42 | 8.75 | 11.63 | 10.83 | 11.19 | 11.26 | 9.35 | 7.94 | 9.73 | 9.89 | 9.09 | 11.23 | 11.22 | 8.65 | 9.81 |

| | | | | | | | | | | | | | | | | |
|---|---|---|---|---|---|---|---|---|---|---|---|---|---|---|---|---|
| ABI2 | 9 | 9.65 | 10.32 | 10.98 | 10.99 | 10.10 | 10.28 | 9.26 | 8.63 | 9.76 | 10.04 | 9.98 | 10.99 | 10.19 | 8.95 | 9.90 |
| ABL1 | 3 | 8.55 | 9.06 | 10.15 | 11.24 | 10.16 | 10.93 | 9.88 | 9.90 | 10.77 | 9.35 | 8.81 | 10.70 | 10.55 | 9.89 | 10.06 |
| ABL2 | 3 | 8.91 | 9.77 | 8.68 | 8.56 | 9.92 | 12.59 | 12.04 | 10.93 | 9.48 | 9.11 | 9.34 | 8.62 | 11.26 | 11.49 | 9.30 |
| ABLIM1 | 12 | 9.18 | 10.28 | 13.15 | 12.85 | 12.75 | 13.24 | 5.68 | 5.59 | 9.04 | 8.24 | 9.73 | 13.00 | 13.00 | 5.63 | 8.64 |
| ABR | 10 | 10.35 | 10.46 | 15.55 | 13.23 | 6.81 | 6.86 | 7.52 | 7.11 | 11.46 | 10.65 | 10.40 | 14.39 | 6.83 | 7.32 | 11.05 |
| ABT1 | 6 | 9.75 | 9.99 | 10.86 | 11.01 | 8.17 | 8.46 | 12.10 | 10.75 | 9.48 | 9.42 | 9.87 | 10.94 | 8.32 | 11.42 | 9.45 |
| ABTB2 | 1 | 9.01 | 8.79 | 12.84 | 13.31 | 8.84 | 8.56 | 8.34 | 7.06 | 10.66 | 12.57 | 8.90 | 13.08 | 8.70 | 7.70 | 11.61 |
| ACAA1 | 6 | 10.50 | 11.05 | 10.85 | 10.30 | 7.46 | 7.91 | 7.86 | 11.74 | 11.18 | 11.15 | 10.78 | 10.58 | 7.68 | 9.80 | 11.16 |
| ACAA2 | 24 | 11.10 | 9.71 | 9.82 | 10.80 | 11.39 | 10.73 | 8.46 | 7.83 | 9.91 | 10.26 | 10.41 | 10.31 | 11.06 | 8.14 | 10.08 |
| ACACA | 65 | 11.46 | 11.02 | 10.99 | 11.14 | 7.15 | 7.26 | 8.93 | 8.61 | 11.78 | 11.66 | 11.24 | 11.06 | 7.21 | 8.77 | 11.72 |
| ACAD10 | 1 | 10.09 | 8.71 | 8.35 | 8.61 | 10.60 | 10.74 | 12.92 | 11.47 | 8.97 | 9.55 | 9.40 | 8.48 | 10.67 | 12.20 | 9.26 |
| ACAD11 | 1 | 9.22 | 10.22 | 11.51 | 9.40 | 9.31 | 10.46 | 12.35 | 11.21 | 7.43 | 8.90 | 9.72 | 10.45 | 9.88 | 11.78 | 8.16 |
| ACAD8 | 1 | 12.08 | 10.64 | 10.64 | 10.27 | 9.28 | 3.68 | 9.66 | 10.28 | 12.26 | 11.21 | 11.36 | 10.45 | 6.48 | 9.97 | 11.74 |
| ACAD9 | 11 | 14.06 | 12.66 | 8.50 | 10.42 | 11.15 | 11.82 | 11.87 | 11.86 | 3.86 | 3.79 | 13.36 | 9.46 | 11.49 | 11.87 | 3.83 |
| ACADL | 1 | 10.30 | 12.65 | 4.61 | 3.71 | 19.04 | 22.24 | 6.27 | 6.68 | 8.08 | 6.42 | 11.47 | 4.16 | 20.64 | 6.47 | 7.25 |
| ACADM | 23 | 12.77 | 11.68 | 8.02 | 8.55 | 9.17 | 8.79 | 10.32 | 9.51 | 10.46 | 10.73 | 12.23 | 8.29 | 8.98 | 9.91 | 10.60 |
| ACADS | 1 | 13.07 | 10.92 | 6.71 | 5.38 | 14.73 | 13.69 | 10.17 | 9.07 | 8.11 | 8.15 | 11.99 | 6.04 | 14.21 | 9.62 | 8.13 |
| ACADVL | 32 | 13.06 | 11.64 | 7.20 | 8.04 | 11.21 | 10.96 | 8.12 | 7.61 | 10.97 | 11.20 | 12.35 | 7.62 | 11.09 | 7.87 | 11.08 |
| ACAP2 | 13 | 10.55 | 11.42 | 9.90 | 9.40 | 10.41 | 11.39 | 8.90 | 8.71 | 9.93 | 9.39 | 10.99 | 9.65 | 10.90 | 8.81 | 9.66 |
| ACAT1 | 17 | 10.74 | 9.28 | 8.97 | 9.44 | 12.23 | 11.24 | 9.80 | 10.67 | 8.88 | 8.76 | 10.01 | 9.20 | 11.73 | 10.23 | 8.82 |
| ACAT2 | 9 | 10.53 | 11.64 | 12.30 | 11.38 | 11.92 | 11.94 | 7.15 | 6.11 | 8.61 | 8.42 | 11.09 | 11.84 | 11.93 | 6.63 | 8.51 |
| ACBD3 | 12 | 8.53 | 10.67 | 9.18 | 8.88 | 12.41 | 12.33 | 9.61 | 8.80 | 9.90 | 9.69 | 9.60 | 9.03 | 12.37 | 9.20 | 9.80 |
| ACBD5 | 2 | 11.02 | 7.90 | 8.19 | 10.23 | 8.75 | 8.92 | 6.99 | 6.30 | 14.92 | 16.78 | 9.46 | 9.21 | 8.84 | 6.64 | 15.85 |
| ACBD6 | 7 | 9.18 | 9.89 | 8.19 | 8.00 | 13.44 | 12.69 | 11.27 | 9.55 | 9.30 | 8.50 | 9.54 | 8.10 | 13.06 | 10.41 | 8.90 |
| ACD | 1 | 9.95 | 10.57 | 8.60 | 7.49 | 10.75 | 11.31 | 10.29 | 9.38 | 11.58 | 10.08 | 10.26 | 8.05 | 11.03 | 9.83 | 10.83 |
| ACIN1 | 52 | 9.34 | 10.83 | 9.84 | 9.19 | 10.13 | 10.49 | 11.56 | 10.94 | 9.02 | 8.66 | 10.09 | 9.51 | 10.31 | 11.25 | 8.84 |
| ACLY | 197 | 9.17 | 9.81 | 13.99 | 12.58 | 7.72 | 7.74 | 5.37 | 5.42 | 14.24 | 13.94 | 9.49 | 13.28 | 7.73 | 5.40 | 14.09 |
| ACN9 | 1 | 10.73 | 9.41 | 10.33 | 9.33 | 10.09 | 9.85 | 11.83 | 10.40 | 9.08 | 8.94 | 10.07 | 9.83 | 9.97 | 11.12 | 9.01 |
| ACO1 | 46 | 8.28 | 9.22 | 12.85 | 12.02 | 8.16 | 7.99 | 8.51 | 8.10 | 12.54 | 12.32 | 8.75 | 12.43 | 8.08 | 8.31 | 12.43 |
| ACO2 | 33 | 12.01 | 9.72 | 9.98 | 11.42 | 9.64 | 8.56 | 9.49 | 8.43 | 10.27 | 10.49 | 10.86 | 10.70 | 9.10 | 8.96 | 10.38 |
| ACOT11 | 1 | 11.24 | 11.89 | 5.76 | 6.11 | 14.88 | 11.67 | 9.27 | 11.69 | 9.67 | 7.82 | 11.56 | 5.94 | 13.28 | 10.48 | 8.74 |
| ACOT13 | 7 | 9.59 | 8.49 | 11.55 | 12.73 | 7.98 | 7.19 | 13.39 | 12.70 | 7.91 | 8.47 | 9.04 | 12.14 | 7.58 | 13.04 | 8.19 |
| ACOT2 | 6 | 9.74 | 8.73 | 13.46 | 14.64 | 8.33 | 7.16 | 7.76 | 6.66 | 11.53 | 11.99 | 9.23 | 14.05 | 7.75 | 7.21 | 11.76 |
| ACOT7 | 26 | 10.09 | 11.24 | 8.68 | 8.05 | 8.30 | 8.41 | 10.26 | 12.06 | 11.54 | 11.37 | 10.67 | 8.37 | 8.35 | 11.16 | 11.45 |
| ACOT8 | 6 | 10.24 | 10.22 | 8.62 | 8.67 | 11.96 | 12.25 | 8.96 | 7.94 | 10.35 | 10.78 | 10.23 | 8.65 | 12.11 | 8.45 | 10.56 |
| ACOT9 | 14 | 9.86 | 8.82 | 8.15 | 9.22 | 11.35 | 10.54 | 11.13 | 10.53 | 9.86 | 10.54 | 9.34 | 8.68 | 10.94 | 10.83 | 10.20 |
| ACOX1 | 14 | 10.82 | 10.94 | 10.43 | 8.84 | 11.29 | 11.44 | 7.85 | 7.62 | 10.54 | 10.23 | 10.88 | 9.63 | 11.37 | 7.74 | 10.39 |
| ACOX3 | 6 | 11.44 | 10.18 | 10.91 | 11.17 | 7.23 | 7.17 | 4.34 | 3.87 | 15.65 | 18.03 | 10.81 | 11.04 | 7.20 | 4.11 | 16.84 |
| ACP1 | 10 | 9.59 | 9.74 | 9.43 | 8.08 | 10.69 | 10.17 | 10.21 | 15.35 | 8.72 | 8.02 | 9.66 | 8.75 | 10.43 | 12.78 | 8.37 |
| ACP2 | 5 | 9.98 | 8.12 | 10.82 | 11.04 | 13.76 | 12.31 | 8.00 | 9.28 | 8.35 | 8.35 | 9.05 | 10.93 | 13.03 | 8.64 | 8.35 |
| ACSF3 | 5 | 11.44 | 9.30 | 6.78 | 7.49 | 12.53 | 11.67 | 10.79 | 9.65 | 9.85 | 10.51 | 10.37 | 7.13 | 12.10 | 10.22 | 10.18 |
| ACSL1 | 7 | 9.56 | 10.81 | 8.95 | 8.38 | 11.24 | 10.98 | 7.72 | 11.07 | 11.01 | 10.28 | 10.18 | 8.67 | 11.11 | 9.40 | 10.65 |

| | | | | | | | | | | | | | | | | |
|---|---|---|---|---|---|---|---|---|---|---|---|---|---|---|---|---|
| ACSL3 | 15 | 10.92 | 8.19 | 7.85 | 9.48 | 8.70 | 7.84 | 10.29 | 11.57 | 11.63 | 13.53 | 9.55 | 8.67 | 8.27 | 10.93 | 12.58 |
| ACSL4 | 28 | 6.23 | 6.18 | 11.22 | 9.99 | 9.55 | 9.16 | 15.66 | 17.20 | 7.51 | 7.31 | 6.20 | 10.61 | 9.35 | 16.43 | 7.41 |
| ACSS2 | 5 | 10.00 | 11.04 | 10.09 | 10.07 | 11.87 | 13.27 | 8.30 | 8.20 | 8.27 | 8.90 | 10.52 | 10.08 | 12.57 | 8.25 | 8.58 |
| ACTA2 | 356 | 9.18 | 12.73 | 12.84 | 12.84 | 8.22 | 8.23 | 7.16 | 6.68 | 11.18 | 10.92 | 10.96 | 12.84 | 8.22 | 6.92 | 11.05 |
| ACTB | 126 | 9.44 | 10.76 | 13.95 | 9.10 | 9.78 | 8.36 | 8.16 | 7.19 | 11.93 | 11.34 | 10.10 | 11.52 | 9.07 | 7.67 | 11.63 |
| ACTG1 | 3 | 6.90 | 7.46 | 13.61 | 6.14 | 8.06 | 6.76 | 18.75 | 12.47 | 9.48 | 10.38 | 7.18 | 9.87 | 7.41 | 15.61 | 9.93 |
| ACTL6A | 13 | 9.90 | 10.73 | 10.37 | 9.54 | 10.93 | 11.30 | 10.06 | 9.07 | 9.24 | 8.85 | 10.32 | 9.95 | 11.12 | 9.56 | 9.05 |
| ACTN1 | 79 | 8.68 | 9.13 | 15.96 | 13.82 | 6.97 | 6.91 | 5.11 | 8.18 | 12.98 | 12.26 | 8.90 | 14.89 | 6.94 | 6.65 | 12.62 |
| ACTN4 | 234 | 9.34 | 11.25 | 13.32 | 11.85 | 8.80 | 8.75 | 6.89 | 6.86 | 11.68 | 11.25 | 10.30 | 12.59 | 8.78 | 6.88 | 11.47 |
| ACTR10 | 7 | 9.17 | 10.02 | 11.08 | 9.20 | 11.92 | 10.73 | 8.67 | 8.96 | 10.05 | 10.20 | 9.60 | 10.14 | 11.32 | 8.81 | 10.12 |
| ACTR1A | 20 | 9.03 | 9.89 | 11.04 | 10.09 | 10.74 | 10.99 | 8.41 | 9.84 | 10.15 | 9.82 | 9.46 | 10.56 | 10.86 | 9.13 | 9.98 |
| ACTR1B | 1 | 10.54 | 9.68 | 5.76 | 5.45 | 18.60 | 14.85 | 9.18 | 8.63 | 10.08 | 7.22 | 10.11 | 5.60 | 16.73 | 8.91 | 8.65 |
| ACTR2 | 27 | 8.54 | 8.79 | 13.35 | 11.47 | 9.63 | 9.25 | 8.26 | 8.82 | 10.94 | 10.96 | 8.67 | 12.41 | 9.44 | 8.54 | 10.95 |
| ACTR3 | 53 | 8.70 | 9.22 | 13.30 | 11.64 | 9.29 | 9.46 | 8.38 | 8.36 | 10.95 | 10.71 | 8.96 | 12.47 | 9.37 | 8.37 | 10.83 |
| ACTR5 | 3 | 8.92 | 10.21 | 10.03 | 9.74 | 10.25 | 10.52 | 10.51 | 9.67 | 10.53 | 9.62 | 9.56 | 9.89 | 10.39 | 10.09 | 10.07 |
| ACTR6 | 1 | 11.93 | 10.37 | 8.51 | 9.76 | 9.33 | 10.64 | 10.10 | 10.64 | 9.85 | 8.88 | 11.15 | 9.14 | 9.98 | 10.37 | 9.36 |
| ACTR8 | 8 | 9.44 | 9.98 | 10.16 | 9.46 | 9.62 | 10.17 | 10.61 | 10.36 | 9.95 | 10.23 | 9.71 | 9.81 | 9.90 | 10.49 | 10.09 |
| ACY1 | 10 | 9.26 | 9.65 | 8.81 | 8.13 | 11.64 | 11.84 | 9.03 | 8.65 | 11.51 | 11.49 | 9.45 | 8.47 | 11.74 | 8.84 | 11.50 |
| ACYP1 | 5 | 12.00 | 12.19 | 9.38 | 8.78 | 11.95 | 12.15 | 12.21 | 10.88 | 5.43 | 5.03 | 12.09 | 9.08 | 12.05 | 11.54 | 5.23 |
| ACYP2 | 2 | 7.70 | 7.72 | 5.04 | 5.22 | 17.48 | 16.52 | 12.73 | 15.62 | 6.03 | 5.95 | 7.71 | 5.13 | 17.00 | 14.17 | 5.99 |
| ADA | 6 | 8.47 | 9.44 | 10.69 | 9.64 | 10.15 | 10.15 | 12.94 | 11.76 | 8.29 | 8.47 | 8.96 | 10.17 | 10.15 | 12.35 | 8.38 |
| ADAL | 2 | 10.66 | 11.71 | 9.11 | 10.13 | 7.33 | 7.29 | 7.78 | 7.27 | 14.80 | 13.91 | 11.19 | 9.62 | 7.31 | 7.53 | 14.35 |
| ADAM10 | 6 | 10.38 | 6.52 | 8.01 | 12.13 | 9.99 | 10.02 | 7.68 | 7.54 | 12.23 | 15.50 | 8.45 | 10.07 | 10.00 | 7.61 | 13.87 |
| ADAM17 | 2 | 10.54 | 7.07 | 8.79 | 13.89 | 8.36 | 8.65 | 8.27 | 6.90 | 12.58 | 14.95 | 8.80 | 11.34 | 8.51 | 7.58 | 13.76 |
| ADAM9 | 3 | 11.11 | 7.21 | 7.94 | 10.57 | 8.58 | 8.32 | 10.23 | 9.40 | 12.81 | 13.81 | 9.16 | 9.26 | 8.45 | 9.82 | 13.31 |
| ADAR | 58 | 8.64 | 9.32 | 10.47 | 9.70 | 11.78 | 11.88 | 9.99 | 11.41 | 8.83 | 7.98 | 8.98 | 10.08 | 11.83 | 10.70 | 8.40 |
| ADARB1 | 2 | 9.47 | 10.53 | 6.50 | 5.69 | 15.84 | 17.15 | 13.73 | 13.77 | 3.83 | 3.50 | 10.00 | 6.09 | 16.49 | 13.75 | 3.67 |
| ADAT1 | 2 | 10.05 | 11.43 | 8.74 | 8.17 | 11.31 | 11.65 | 8.64 | 9.51 | 10.26 | 10.24 | 10.74 | 8.46 | 11.48 | 9.07 | 10.25 |
| ADAT2 | 2 | 9.50 | 9.09 | 14.91 | 13.82 | 9.80 | 9.31 | 13.58 | 11.79 | 4.27 | 3.92 | 9.30 | 14.37 | 9.55 | 12.68 | 4.10 |
| ADCK3 | 2 | 9.61 | 9.79 | 10.01 | 9.40 | 11.34 | 9.08 | 10.33 | 11.08 | 8.82 | 10.54 | 9.70 | 9.71 | 10.21 | 10.70 | 9.68 |
| ADCK4 | 1 | 13.10 | 10.89 | 5.38 | 7.92 | 9.08 | 12.78 | 9.82 | 12.12 | 9.18 | 9.74 | 12.00 | 6.65 | 10.93 | 10.97 | 9.46 |
| ADCY9 | 1 | 11.64 | 9.92 | 10.64 | 13.74 | 8.60 | 10.13 | 6.07 | 4.78 | 12.38 | 12.10 | 10.78 | 12.19 | 9.36 | 5.43 | 12.24 |
| ADD1 | 9 | 8.45 | 8.61 | 14.99 | 12.84 | 9.76 | 9.16 | 8.31 | 7.50 | 10.29 | 10.08 | 8.53 | 13.92 | 9.46 | 7.90 | 10.19 |
| ADD3 | 15 | 7.98 | 8.36 | 13.67 | 13.13 | 9.52 | 9.89 | 9.04 | 8.51 | 10.10 | 9.79 | 8.17 | 13.40 | 9.71 | 8.78 | 9.95 |
| ADH5 | 33 | 9.33 | 10.38 | 10.42 | 10.14 | 9.67 | 10.10 | 9.66 | 12.25 | 9.01 | 9.04 | 9.85 | 10.28 | 9.88 | 10.95 | 9.03 |
| ADI1 | 4 | 6.82 | 9.34 | 11.88 | 10.23 | 13.64 | 13.14 | 10.36 | 7.82 | 8.15 | 8.60 | 8.08 | 11.05 | 13.39 | 9.09 | 8.37 |
| ADIRF | 4 | 6.46 | 6.12 | 1.42 | 2.83 | 30.63 | 38.71 | 2.67 | 2.03 | 4.82 | 4.31 | 6.29 | 2.13 | 34.67 | 2.35 | 4.57 |
| ADK | 20 | 8.55 | 9.54 | 10.59 | 8.11 | 10.58 | 9.51 | 10.84 | 17.47 | 7.51 | 7.29 | 9.05 | 9.35 | 10.04 | 14.16 | 7.40 |
| ADNP | 37 | 9.12 | 9.84 | 11.72 | 10.12 | 9.60 | 9.53 | 10.92 | 10.00 | 9.64 | 9.51 | 9.48 | 10.92 | 9.57 | 10.46 | 9.57 |
| ADNP2 | 2 | 8.23 | 9.96 | 12.87 | 13.87 | 7.78 | 9.71 | 8.89 | 9.21 | 9.56 | 9.92 | 9.09 | 13.37 | 8.75 | 9.05 | 9.74 |
| ADO | 5 | 8.07 | 8.23 | 11.80 | 11.24 | 10.91 | 11.66 | 10.86 | 10.33 | 9.11 | 7.78 | 8.15 | 11.52 | 11.29 | 10.59 | 8.45 |
| ADPGK | 2 | 9.92 | 7.83 | 9.22 | 5.87 | 11.64 | 9.28 | 11.45 | 8.80 | 13.37 | 12.62 | 8.87 | 7.55 | 10.46 | 10.12 | 13.00 |

| | | | | | | | | | | | | | | | | |
|---|---|---|---|---|---|---|---|---|---|---|---|---|---|---|---|---|
| ADPRHL2 | 5 | 10.72 | 11.77 | 9.27 | 9.73 | 8.60 | 8.80 | 8.94 | 7.79 | 12.37 | 12.02 | 11.24 | 9.50 | 8.70 | 8.36 | 12.19 |
| ADRBK1 | 7 | 9.88 | 10.58 | 8.92 | 8.90 | 10.28 | 11.62 | 10.57 | 10.16 | 10.10 | 9.00 | 10.23 | 8.91 | 10.95 | 10.36 | 9.55 |
| ADRM1 | 15 | 9.13 | 9.55 | 8.27 | 7.99 | 10.27 | 9.98 | 9.93 | 16.51 | 9.51 | 8.86 | 9.34 | 8.13 | 10.13 | 13.22 | 9.18 |
| ADSL | 17 | 8.70 | 9.25 | 16.16 | 14.98 | 6.85 | 6.58 | 9.01 | 8.22 | 10.24 | 10.02 | 8.97 | 15.57 | 6.71 | 8.61 | 10.13 |
| ADSS | 19 | 10.08 | 10.23 | 8.17 | 7.33 | 11.34 | 11.15 | 10.41 | 14.23 | 8.97 | 8.09 | 10.15 | 7.75 | 11.24 | 12.32 | 8.53 |
| ADSSL1 | 8 | 9.21 | 10.84 | 6.86 | 7.02 | 10.93 | 11.49 | 14.80 | 13.39 | 7.89 | 7.57 | 10.03 | 6.94 | 11.21 | 14.10 | 7.73 |
| AEBP2 | 1 | 11.57 | 12.54 | 10.43 | 9.22 | 8.30 | 8.41 | 10.81 | 8.68 | 10.40 | 9.63 | 12.06 | 9.82 | 8.36 | 9.75 | 10.01 |
| AFAP1 | 10 | 9.49 | 10.36 | 15.32 | 15.20 | 10.09 | 10.78 | 3.34 | 3.17 | 11.11 | 11.13 | 9.93 | 15.26 | 10.43 | 3.26 | 11.12 |
| AFF4 | 4 | 7.99 | 9.29 | 13.03 | 9.41 | 10.50 | 9.92 | 10.62 | 9.46 | 10.43 | 9.36 | 8.64 | 11.22 | 10.21 | 10.04 | 9.89 |
| AFG3L2 | 14 | 12.84 | 7.19 | 7.62 | 10.85 | 10.07 | 8.84 | 9.12 | 8.54 | 11.65 | 13.27 | 10.02 | 9.24 | 9.46 | 8.83 | 12.46 |
| AFTPH | 2 | 10.54 | 11.00 | 11.08 | 10.32 | 9.29 | 8.67 | 8.29 | 8.67 | 11.61 | 10.54 | 10.77 | 10.70 | 8.98 | 8.48 | 11.07 |
| AGFG1 | 13 | 9.90 | 10.49 | 9.97 | 9.51 | 10.71 | 10.41 | 10.09 | 8.58 | 10.37 | 9.98 | 10.19 | 9.74 | 10.56 | 9.33 | 10.17 |
| AGFG2 | 1 | 9.24 | 9.93 | 11.57 | 10.53 | 8.60 | 8.12 | 8.82 | 6.67 | 12.02 | 14.50 | 9.59 | 11.05 | 8.36 | 7.74 | 13.26 |
| AGGF1 | 4 | 9.37 | 11.13 | 8.99 | 8.69 | 11.18 | 11.73 | 10.13 | 9.11 | 9.70 | 9.98 | 10.25 | 8.84 | 11.46 | 9.62 | 9.84 |
| AGK | 5 | 12.48 | 7.40 | 7.35 | 11.64 | 7.61 | 6.97 | 10.74 | 10.50 | 11.91 | 13.39 | 9.94 | 9.50 | 7.29 | 10.62 | 12.65 |
| AGL | 22 | 12.00 | 12.11 | 7.87 | 7.48 | 10.75 | 10.78 | 9.50 | 9.13 | 10.46 | 9.93 | 12.05 | 7.67 | 10.76 | 9.31 | 10.20 |
| AGMAT | 1 | 11.91 | 10.37 | 7.59 | 7.42 | 9.16 | 8.54 | 15.31 | 14.08 | 7.51 | 8.11 | 11.14 | 7.50 | 8.85 | 14.70 | 7.81 |
| AGO1 | 5 | 9.93 | 11.25 | 8.71 | 8.88 | 9.26 | 8.81 | 11.04 | 10.60 | 10.37 | 11.16 | 10.59 | 8.79 | 9.03 | 10.82 | 10.76 |
| AGO2 | 8 | 8.70 | 8.92 | 10.11 | 9.96 | 10.16 | 9.89 | 12.10 | 10.76 | 9.37 | 10.03 | 8.81 | 10.03 | 10.03 | 11.43 | 9.70 |
| AGO4 | 4 | 9.92 | 10.61 | 9.34 | 8.93 | 9.41 | 10.66 | 10.67 | 9.94 | 10.69 | 9.85 | 10.26 | 9.13 | 10.03 | 10.31 | 10.27 |
| AGPAT3 | 3 | 14.87 | 8.80 | 5.81 | 7.76 | 12.32 | 10.00 | 8.10 | 7.94 | 10.55 | 13.84 | 11.84 | 6.79 | 11.16 | 8.02 | 12.19 |
| AGPAT5 | 3 | 10.26 | 8.99 | 7.88 | 10.22 | 7.19 | 7.87 | 13.26 | 12.39 | 10.54 | 11.39 | 9.62 | 9.05 | 7.53 | 12.83 | 10.97 |
| AGPAT6 | 1 | 12.77 | 5.59 | 6.58 | 10.99 | 8.05 | 7.61 | 12.08 | 10.37 | 11.60 | 14.36 | 9.18 | 8.79 | 7.83 | 11.22 | 12.98 |
| AGPAT9 | 1 | 10.86 | 5.75 | 9.74 | 14.06 | 4.86 | 4.17 | 15.43 | 13.39 | 10.04 | 11.69 | 8.31 | 11.90 | 4.52 | 14.41 | 10.87 |
| AGPS | 29 | 9.41 | 9.32 | 9.65 | 9.16 | 10.01 | 10.24 | 10.32 | 11.23 | 10.38 | 10.28 | 9.36 | 9.40 | 10.12 | 10.78 | 10.33 |
| AGRN | 5 | 9.70 | 8.48 | 8.25 | 7.37 | 12.77 | 11.86 | 11.11 | 10.44 | 9.93 | 10.09 | 9.09 | 7.81 | 12.31 | 10.78 | 10.01 |
| AGTPBP1 | 2 | 8.38 | 8.30 | 11.80 | 11.59 | 13.34 | 12.89 | 8.39 | 7.98 | 8.74 | 8.59 | 8.34 | 11.70 | 13.11 | 8.19 | 8.66 |
| AGTRAP | 2 | 8.57 | 7.84 | 8.08 | 9.00 | 8.31 | 10.83 | 13.13 | 15.35 | 9.62 | 9.27 | 8.21 | 8.54 | 9.57 | 14.24 | 9.45 |
| AHCTF1 | 28 | 10.02 | 9.52 | 7.93 | 8.95 | 10.00 | 9.68 | 10.81 | 11.05 | 10.52 | 11.53 | 9.77 | 8.44 | 9.84 | 10.93 | 11.03 |
| AHCY | 58 | 8.89 | 10.19 | 11.42 | 10.94 | 10.80 | 11.14 | 9.51 | 9.79 | 8.47 | 8.86 | 9.54 | 11.18 | 10.97 | 9.65 | 8.67 |
| AHCYL1 | 20 | 10.68 | 11.10 | 10.25 | 10.41 | 7.68 | 7.75 | 9.38 | 12.17 | 10.58 | 10.00 | 10.89 | 10.33 | 7.72 | 10.78 | 10.29 |
| AHCYL2 | 12 | 10.80 | 10.42 | 10.30 | 9.26 | 8.83 | 8.69 | 9.79 | 12.95 | 9.81 | 9.15 | 10.61 | 9.78 | 8.76 | 11.37 | 9.48 |
| AHDC1 | 1 | 19.24 | 9.46 | 7.39 | 8.09 | 7.49 | 9.53 | 7.30 | 7.01 | 13.93 | 10.57 | 14.35 | 7.74 | 8.51 | 7.15 | 12.25 |
| AHI1 | 2 | 11.45 | 10.57 | 6.45 | 8.05 | 10.28 | 9.42 | 12.13 | 12.09 | 10.22 | 9.36 | 11.01 | 7.25 | 9.85 | 12.11 | 9.79 |
| AHNAK | 683 | 9.04 | 10.10 | 11.33 | 10.84 | 9.68 | 9.99 | 7.81 | 7.03 | 12.47 | 11.73 | 9.57 | 11.08 | 9.84 | 7.42 | 12.10 |
| AHNAK2 | 192 | 11.22 | 12.40 | 6.87 | 6.61 | 15.03 | 15.58 | 3.43 | 3.05 | 13.31 | 12.49 | 11.81 | 6.74 | 15.31 | 3.24 | 12.90 |
| AHRR | 1 | 10.84 | 10.87 | 6.93 | 7.91 | 7.87 | 9.24 | 13.67 | 12.79 | 9.02 | 10.85 | 10.86 | 7.42 | 8.55 | 13.23 | 9.93 |
| AHSA1 | 37 | 8.82 | 9.20 | 12.04 | 10.55 | 9.63 | 8.91 | 9.92 | 10.95 | 10.12 | 9.86 | 9.01 | 11.29 | 9.27 | 10.43 | 9.99 |
| AIDA | 7 | 9.90 | 9.64 | 9.00 | 9.13 | 12.39 | 12.57 | 9.02 | 8.08 | 10.20 | 10.07 | 9.77 | 9.06 | 12.48 | 8.55 | 10.14 |
| AIFM1 | 31 | 9.74 | 8.06 | 9.66 | 9.28 | 11.19 | 10.70 | 11.57 | 12.48 | 8.75 | 8.56 | 8.90 | 9.47 | 10.94 | 12.02 | 8.66 |
| AIFM2 | 8 | 9.67 | 10.08 | 12.03 | 11.87 | 10.27 | 10.47 | 9.08 | 8.59 | 9.19 | 8.75 | 9.88 | 11.95 | 10.37 | 8.84 | 8.97 |
| AIMP1 | 28 | 9.67 | 10.07 | 10.63 | 9.33 | 10.14 | 9.48 | 10.02 | 11.38 | 10.04 | 9.25 | 9.87 | 9.98 | 9.81 | 10.70 | 9.64 |

| | | | | | | | | | | | | | | | | |
|---|---|---|---|---|---|---|---|---|---|---|---|---|---|---|---|---|
| AIMP2 | 11 | 9.37 | 10.06 | 10.01 | 9.21 | 10.78 | 10.83 | 9.50 | 12.07 | 9.45 | 8.73 | 9.71 | 9.61 | 10.80 | 10.78 | 9.09 |
| AIP | 14 | 9.18 | 9.95 | 8.72 | 7.78 | 12.76 | 12.76 | 10.01 | 12.27 | 8.42 | 8.15 | 9.56 | 8.25 | 12.76 | 11.14 | 8.28 |
| AJUBA | 5 | 10.10 | 10.44 | 12.23 | 12.03 | 12.18 | 12.05 | 6.31 | 5.47 | 9.99 | 9.22 | 10.27 | 12.13 | 12.11 | 5.89 | 9.60 |
| AK1 | 17 | 9.11 | 11.46 | 10.50 | 9.93 | 11.29 | 11.98 | 8.25 | 9.21 | 9.03 | 9.25 | 10.28 | 10.21 | 11.64 | 8.73 | 9.14 |
| AK2 | 36 | 8.88 | 8.19 | 9.25 | 9.06 | 9.62 | 9.75 | 11.53 | 14.81 | 9.72 | 9.20 | 8.53 | 9.15 | 9.69 | 13.17 | 9.46 |
| AK3 | 22 | 10.76 | 9.60 | 9.97 | 10.58 | 9.76 | 9.33 | 8.57 | 7.62 | 11.87 | 11.94 | 10.18 | 10.28 | 9.54 | 8.09 | 11.90 |
| AK4 | 10 | 10.99 | 10.17 | 7.60 | 8.06 | 7.57 | 6.76 | 12.28 | 10.44 | 13.01 | 13.13 | 10.58 | 7.83 | 7.16 | 11.36 | 13.07 |
| AK6 | 6 | 10.51 | 10.92 | 8.69 | 6.71 | 9.56 | 8.62 | 11.97 | 9.77 | 11.75 | 11.50 | 10.71 | 7.70 | 9.09 | 10.87 | 11.63 |
| AKAP1 | 3 | 11.92 | 7.91 | 8.26 | 10.48 | 9.87 | 8.39 | 10.46 | 9.26 | 11.76 | 11.68 | 9.92 | 9.37 | 9.13 | 9.86 | 11.72 |
| AKAP10 | 2 | 9.93 | 11.96 | 10.02 | 12.42 | 7.87 | 8.89 | 8.37 | 7.70 | 11.27 | 11.57 | 10.95 | 11.22 | 8.38 | 8.04 | 11.42 |
| AKAP11 | 3 | 8.50 | 10.34 | 8.05 | 7.86 | 13.46 | 13.35 | 12.01 | 11.65 | 7.41 | 7.35 | 9.42 | 7.96 | 13.41 | 11.83 | 7.38 |
| AKAP12 | 87 | 5.41 | 6.06 | 21.37 | 19.21 | 4.52 | 4.45 | 13.72 | 13.01 | 6.33 | 5.93 | 5.74 | 20.29 | 4.48 | 13.36 | 6.13 |
| AKAP13 | 31 | 8.28 | 9.36 | 12.02 | 11.39 | 9.99 | 10.07 | 9.38 | 8.70 | 10.61 | 10.20 | 8.82 | 11.71 | 10.03 | 9.04 | 10.41 |
| AKAP17A | 8 | 8.97 | 10.32 | 9.44 | 9.72 | 10.71 | 11.85 | 10.45 | 10.04 | 9.30 | 9.20 | 9.64 | 9.58 | 11.28 | 10.24 | 9.25 |
| AKAP2 | 49 | 9.08 | 10.10 | 15.82 | 14.92 | 5.79 | 6.01 | 8.07 | 7.03 | 11.92 | 11.26 | 9.59 | 15.37 | 5.90 | 7.55 | 11.59 |
| AKAP8 | 20 | 9.35 | 9.79 | 10.54 | 10.49 | 8.81 | 9.17 | 12.15 | 11.35 | 9.54 | 8.81 | 9.57 | 10.51 | 8.99 | 11.75 | 9.18 |
| AKAP8L | 14 | 9.05 | 10.03 | 11.91 | 10.92 | 7.94 | 8.31 | 12.02 | 13.21 | 8.67 | 7.94 | 9.54 | 11.41 | 8.13 | 12.61 | 8.31 |
| AKAP9 | 42 | 10.17 | 11.27 | 10.04 | 9.95 | 10.26 | 10.66 | 9.35 | 9.09 | 9.92 | 9.28 | 10.72 | 9.99 | 10.46 | 9.22 | 9.60 |
| AKR1A1 | 30 | 10.71 | 11.92 | 8.33 | 7.91 | 9.72 | 9.68 | 10.53 | 10.63 | 10.43 | 10.16 | 11.31 | 8.12 | 9.70 | 10.58 | 10.29 |
| AKR1B1 | 34 | 9.81 | 9.96 | 16.68 | 15.61 | 7.00 | 6.91 | 9.06 | 9.01 | 8.00 | 7.95 | 9.89 | 16.15 | 6.95 | 9.04 | 7.97 |
| AKR1C2 | 36 | 13.57 | 15.03 | 7.33 | 5.70 | 9.73 | 8.98 | 8.78 | 8.03 | 11.64 | 11.23 | 14.30 | 6.51 | 9.35 | 8.40 | 11.43 |
| AKR1C3 | 75 | 11.03 | 16.85 | 6.65 | 5.72 | 11.56 | 11.38 | 8.52 | 7.65 | 10.09 | 10.53 | 13.94 | 6.19 | 11.47 | 8.09 | 10.31 |
| AKR7A2 | 10 | 10.72 | 11.88 | 7.04 | 6.29 | 11.91 | 11.71 | 9.24 | 9.89 | 10.95 | 10.38 | 11.30 | 6.67 | 11.81 | 9.56 | 10.66 |
| AKT1 | 6 | 8.76 | 9.87 | 9.14 | 8.98 | 10.89 | 11.56 | 12.84 | 12.02 | 8.17 | 7.77 | 9.31 | 9.06 | 11.22 | 12.43 | 7.97 |
| AKT1S1 | 6 | 9.64 | 10.33 | 11.23 | 10.10 | 10.61 | 10.47 | 9.38 | 7.95 | 10.23 | 10.05 | 9.99 | 10.67 | 10.54 | 8.67 | 10.14 |
| AKT2 | 12 | 9.55 | 10.40 | 9.03 | 8.30 | 9.77 | 10.14 | 11.39 | 13.29 | 9.08 | 9.05 | 9.98 | 8.66 | 9.96 | 12.34 | 9.06 |
| ALAD | 4 | 9.41 | 11.12 | 11.49 | 10.12 | 10.71 | 11.71 | 7.94 | 6.62 | 11.15 | 9.71 | 10.27 | 10.81 | 11.21 | 7.28 | 10.43 |
| ALCAM | 13 | 10.78 | 8.47 | 12.60 | 14.73 | 7.60 | 7.33 | 9.96 | 9.81 | 9.25 | 9.47 | 9.63 | 13.66 | 7.47 | 9.88 | 9.36 |
| ALDH16A1 | 8 | 8.35 | 9.05 | 6.55 | 6.69 | 12.99 | 13.67 | 9.64 | 16.53 | 8.46 | 8.08 | 8.70 | 6.62 | 13.33 | 13.09 | 8.27 |
| ALDH18A1 | 44 | 10.83 | 9.27 | 7.94 | 8.89 | 9.85 | 9.06 | 11.72 | 13.28 | 9.56 | 9.61 | 10.05 | 8.41 | 9.45 | 12.50 | 9.59 |
| ALDH1A2 | 6 | 11.35 | 10.61 | 10.55 | 11.92 | 7.82 | 7.27 | 5.46 | 6.35 | 14.30 | 14.36 | 10.98 | 11.23 | 7.55 | 5.91 | 14.33 |
| ALDH1A3 | 2 | 11.04 | 13.77 | 12.41 | 15.17 | 6.63 | 7.97 | 3.53 | 2.82 | 13.47 | 13.19 | 12.40 | 13.79 | 7.30 | 3.17 | 13.33 |
| ALDH1B1 | 26 | 11.28 | 9.25 | 10.54 | 11.23 | 8.32 | 7.23 | 4.90 | 4.25 | 16.12 | 16.88 | 10.26 | 10.89 | 7.77 | 4.57 | 16.50 |
| ALDH1L2 | 1 | 7.75 | 8.49 | 9.51 | 9.94 | 8.80 | 9.03 | 13.03 | 11.98 | 9.80 | 11.67 | 8.12 | 9.72 | 8.92 | 12.50 | 10.74 |
| ALDH2 | 11 | 10.60 | 8.87 | 8.68 | 9.50 | 10.88 | 9.37 | 11.23 | 10.38 | 10.04 | 10.45 | 9.74 | 9.09 | 10.12 | 10.80 | 10.24 |
| ALDH3A1 | 13 | 15.66 | 17.81 | 1.93 | 1.48 | 11.08 | 11.09 | 2.85 | 2.73 | 17.63 | 17.74 | 16.73 | 1.71 | 11.08 | 2.79 | 17.69 |
| ALDH3A2 | 9 | 16.27 | 12.60 | 6.21 | 7.03 | 10.04 | 9.58 | 6.65 | 6.54 | 11.95 | 13.13 | 14.43 | 6.62 | 9.81 | 6.59 | 12.54 |
| ALDH3B1 | 1 | 11.31 | 12.31 | 8.60 | 8.33 | 10.74 | 11.21 | 8.66 | 6.10 | 11.19 | 11.56 | 11.81 | 8.47 | 10.98 | 7.38 | 11.38 |
| ALDH4A1 | 2 | 7.84 | 10.87 | 13.11 | 13.66 | 10.26 | 12.82 | 5.46 | 5.49 | 9.96 | 10.53 | 9.36 | 13.39 | 11.54 | 5.47 | 10.25 |
| ALDH5A1 | 2 | 14.35 | 12.62 | 5.62 | 6.94 | 18.28 | 18.02 | 7.25 | 6.80 | 4.82 | 5.33 | 13.48 | 6.28 | 18.15 | 7.02 | 5.07 |
| ALDH6A1 | 5 | 12.05 | 9.72 | 6.36 | 7.41 | 12.68 | 11.65 | 12.40 | 11.78 | 7.77 | 8.18 | 10.88 | 6.88 | 12.16 | 12.09 | 7.98 |
| ALDH7A1 | 29 | 9.39 | 10.13 | 7.31 | 6.87 | 11.56 | 11.82 | 11.38 | 12.14 | 9.61 | 9.79 | 9.76 | 7.09 | 11.69 | 11.76 | 9.70 |

| | | | | | | | | | | | | | | | | |
|---|---|---|---|---|---|---|---|---|---|---|---|---|---|---|---|---|
| ALDH9A1 | 24 | 9.68 | 10.06 | 8.64 | 8.85 | 15.34 | 15.15 | 7.97 | 7.45 | 8.73 | 8.13 | 9.87 | 8.75 | 15.25 | 7.71 | 8.43 |
| ALDOA | 149 | 8.69 | 10.85 | 11.42 | 9.98 | 10.41 | 10.17 | 7.88 | 12.07 | 9.45 | 9.09 | 9.77 | 10.70 | 10.29 | 9.97 | 9.27 |
| ALDOC | 11 | 13.19 | 13.92 | 16.45 | 16.17 | 8.09 | 8.19 | 4.40 | 4.85 | 7.63 | 7.10 | 13.55 | 16.31 | 8.14 | 4.62 | 7.37 |
| ALG1 | 6 | 12.53 | 6.27 | 8.19 | 13.72 | 9.24 | 8.38 | 8.28 | 7.67 | 11.00 | 14.70 | 9.40 | 10.96 | 8.81 | 7.98 | 12.85 |
| ALG13 | 4 | 6.97 | 11.41 | 8.30 | 6.99 | 14.48 | 14.75 | 11.72 | 11.01 | 7.08 | 7.28 | 9.19 | 7.65 | 14.61 | 11.37 | 7.18 |
| ALG2 | 4 | 8.93 | 8.57 | 11.28 | 10.41 | 10.39 | 10.53 | 8.22 | 9.00 | 11.13 | 11.53 | 8.75 | 10.85 | 10.46 | 8.61 | 11.33 |
| ALG5 | 3 | 12.63 | 8.33 | 8.78 | 12.25 | 10.60 | 9.84 | 8.82 | 9.53 | 9.58 | 9.64 | 10.48 | 10.52 | 10.22 | 9.17 | 9.61 |
| ALKBH1 | 1 | 11.62 | 10.36 | 9.51 | 9.52 | 8.43 | 8.60 | 8.80 | 10.16 | 11.61 | 11.39 | 10.99 | 9.52 | 8.52 | 9.48 | 11.50 |
| ALKBH2 | 1 | 10.89 | 9.61 | 9.47 | 8.41 | 9.69 | 8.14 | 12.35 | 9.78 | 10.99 | 10.66 | 10.25 | 8.94 | 8.92 | 11.06 | 10.83 |
| ALKBH4 | 1 | 11.71 | 12.53 | 8.30 | 7.60 | 9.06 | 9.77 | 10.45 | 10.99 | 10.12 | 9.46 | 12.12 | 7.95 | 9.42 | 10.72 | 9.79 |
| ALKBH5 | 8 | 9.60 | 10.50 | 11.07 | 9.50 | 8.23 | 8.79 | 10.31 | 11.02 | 10.55 | 10.42 | 10.05 | 10.28 | 8.51 | 10.67 | 10.48 |
| ALKBH8 | 1 | 9.31 | 7.89 | 14.44 | 12.76 | 5.10 | 6.84 | 9.80 | 9.81 | 13.11 | 10.94 | 8.60 | 13.60 | 5.97 | 9.80 | 12.02 |
| ALPL | 2 | 11.38 | 11.72 | 15.58 | 20.77 | 2.73 | 2.63 | 13.28 | 12.33 | 4.83 | 4.75 | 11.55 | 18.17 | 2.68 | 12.80 | 4.79 |
| ALYREF | 28 | 10.99 | 11.17 | 12.04 | 9.43 | 8.98 | 8.39 | 9.19 | 7.24 | 10.75 | 11.82 | 11.08 | 10.73 | 8.68 | 8.22 | 11.29 |
| AMACR | 2 | 9.89 | 10.47 | 8.82 | 9.59 | 12.80 | 14.48 | 8.77 | 8.80 | 8.13 | 8.24 | 10.18 | 9.21 | 13.64 | 8.78 | 8.19 |
| AMD1 | 3 | 9.45 | 10.43 | 9.25 | 9.64 | 9.16 | 10.28 | 10.70 | 9.98 | 10.26 | 10.83 | 9.94 | 9.45 | 9.72 | 10.34 | 10.55 |
| AMMECR1 | 3 | 8.22 | 9.79 | 9.39 | 8.89 | 11.49 | 12.45 | 10.39 | 11.36 | 9.65 | 8.38 | 9.01 | 9.14 | 11.97 | 10.87 | 9.01 |
| AMPD2 | 17 | 9.69 | 10.81 | 10.52 | 9.88 | 7.07 | 7.22 | 11.29 | 11.18 | 11.35 | 10.98 | 10.25 | 10.20 | 7.14 | 11.24 | 11.17 |
| ANAPC1 | 12 | 9.98 | 10.17 | 8.36 | 9.26 | 10.79 | 11.64 | 10.30 | 10.20 | 9.45 | 9.86 | 10.07 | 8.81 | 11.21 | 10.25 | 9.65 |
| ANAPC10 | 3 | 9.56 | 9.76 | 9.44 | 9.87 | 9.98 | 10.78 | 12.50 | 10.85 | 8.93 | 8.33 | 9.66 | 9.66 | 10.38 | 11.68 | 8.63 |
| ANAPC16 | 2 | 9.87 | 11.61 | 12.44 | 10.37 | 9.24 | 9.98 | 9.82 | 7.31 | 10.02 | 9.33 | 10.74 | 11.41 | 9.61 | 8.56 | 9.68 |
| ANAPC2 | 10 | 10.66 | 10.16 | 8.40 | 9.05 | 10.62 | 10.90 | 9.92 | 10.75 | 9.68 | 9.87 | 10.41 | 8.72 | 10.76 | 10.33 | 9.77 |
| ANAPC4 | 5 | 11.16 | 10.71 | 8.43 | 8.53 | 9.72 | 10.94 | 9.96 | 11.32 | 9.71 | 9.52 | 10.93 | 8.48 | 10.33 | 10.64 | 9.61 |
| ANAPC5 | 9 | 9.24 | 10.04 | 9.41 | 9.88 | 10.96 | 11.21 | 11.00 | 9.74 | 9.56 | 8.98 | 9.64 | 9.64 | 11.08 | 10.37 | 9.27 |
| ANAPC7 | 17 | 9.38 | 11.51 | 11.24 | 10.91 | 9.17 | 9.07 | 9.94 | 9.82 | 9.37 | 9.58 | 10.44 | 11.08 | 9.12 | 9.88 | 9.47 |
| ANGEL2 | 1 | 10.88 | 11.32 | 3.88 | 3.98 | 10.26 | 12.13 | 14.00 | 15.26 | 9.59 | 8.70 | 11.10 | 3.93 | 11.19 | 14.63 | 9.15 |
| ANK3 | 2 | 9.57 | 8.79 | 8.99 | 9.73 | 6.72 | 7.87 | 17.77 | 16.97 | 6.91 | 6.69 | 9.18 | 9.36 | 7.29 | 17.37 | 6.80 |
| ANKFY1 | 20 | 11.12 | 11.75 | 9.32 | 8.41 | 10.86 | 10.64 | 8.57 | 9.37 | 10.22 | 9.72 | 11.44 | 8.87 | 10.75 | 8.97 | 9.97 |
| ANKH | 3 | 9.58 | 17.75 | 5.16 | 7.80 | 9.71 | 8.96 | 10.63 | 10.14 | 9.23 | 11.05 | 13.66 | 6.48 | 9.33 | 10.38 | 10.14 |
| ANKHD1 | 29 | 9.77 | 11.13 | 9.18 | 9.51 | 9.89 | 10.41 | 10.75 | 10.01 | 9.79 | 9.56 | 10.45 | 9.34 | 10.15 | 10.38 | 9.68 |
| ANKLE2 | 17 | 10.75 | 8.67 | 11.14 | 12.03 | 8.87 | 7.99 | 9.28 | 8.09 | 11.11 | 12.07 | 9.71 | 11.58 | 8.43 | 8.69 | 11.59 |
| ANKMY2 | 7 | 10.01 | 10.88 | 11.56 | 11.11 | 8.54 | 8.57 | 10.21 | 8.92 | 10.40 | 9.81 | 10.44 | 11.33 | 8.56 | 9.56 | 10.10 |
| ANKRD13A | 4 | 10.47 | 12.15 | 13.80 | 13.89 | 8.26 | 8.48 | 7.01 | 6.95 | 9.19 | 9.80 | 11.31 | 13.84 | 8.37 | 6.98 | 9.50 |
| ANKRD13C | 1 | 9.55 | 11.01 | 8.94 | 8.35 | 10.79 | 10.13 | 11.44 | 7.48 | 11.72 | 10.60 | 10.28 | 8.64 | 10.46 | 9.46 | 11.16 |
| ANKRD17 | 14 | 9.12 | 10.45 | 11.09 | 10.41 | 9.84 | 10.11 | 9.98 | 9.22 | 10.08 | 9.70 | 9.78 | 10.75 | 9.98 | 9.60 | 9.89 |
| ANKRD26 | 2 | 8.92 | 11.20 | 8.91 | 8.48 | 8.89 | 10.51 | 7.66 | 6.77 | 13.94 | 14.71 | 10.06 | 8.70 | 9.70 | 7.22 | 14.32 |
| ANKRD28 | 9 | 8.01 | 9.33 | 9.48 | 8.64 | 8.99 | 8.98 | 10.20 | 18.25 | 9.33 | 8.80 | 8.67 | 9.06 | 8.99 | 14.22 | 9.06 |
| ANKRD31 | 2 | 9.80 | 9.38 | 10.13 | 10.94 | 10.89 | 10.14 | 9.99 | 10.10 | 9.54 | 9.11 | 9.59 | 10.53 | 10.51 | 10.04 | 9.32 |
| ANKRD39 | 1 | 8.23 | 8.95 | 9.95 | 6.80 | 10.34 | 10.78 | 14.54 | 10.80 | 11.66 | 7.95 | 8.59 | 8.37 | 10.56 | 12.67 | 9.81 |
| ANKRD44 | 3 | 11.29 | 9.31 | 10.16 | 10.80 | 10.13 | 11.62 | 9.36 | 9.09 | 9.27 | 8.96 | 10.30 | 10.48 | 10.88 | 9.22 | 9.12 |
| ANKRD50 | 3 | 8.83 | 8.34 | 10.97 | 11.74 | 11.47 | 11.55 | 10.16 | 9.07 | 9.24 | 8.63 | 8.58 | 11.35 | 11.51 | 9.61 | 8.94 |
| ANKRD52 | 6 | 10.16 | 10.89 | 9.76 | 9.79 | 10.70 | 10.31 | 8.71 | 7.80 | 11.56 | 10.32 | 10.53 | 9.78 | 10.50 | 8.25 | 10.94 |

| | | | | | | | | | | | | | | | | |
|---|---|---|---|---|---|---|---|---|---|---|---|---|---|---|---|---|
| ANKS1A | 5 | 9.53 | 10.25 | 11.06 | 9.59 | 8.64 | 8.67 | 10.55 | 11.69 | 10.49 | 9.54 | 9.89 | 10.33 | 8.65 | 11.12 | 10.01 |
| ANKZF1 | 16 | 8.80 | 10.29 | 7.95 | 7.78 | 11.22 | 11.15 | 12.44 | 11.61 | 9.68 | 9.08 | 9.54 | 7.87 | 11.18 | 12.02 | 9.38 |
| ANLN | 50 | 8.75 | 9.92 | 12.57 | 12.56 | 6.60 | 6.68 | 8.34 | 8.65 | 13.04 | 12.89 | 9.34 | 12.56 | 6.64 | 8.49 | 12.96 |
| ANO10 | 6 | 13.60 | 8.74 | 9.02 | 13.87 | 9.15 | 8.65 | 8.44 | 8.53 | 9.11 | 10.89 | 11.17 | 11.44 | 8.90 | 8.49 | 10.00 |
| ANO6 | 1 | 10.51 | 9.28 | 10.33 | 14.32 | 6.35 | 6.14 | 8.92 | 8.16 | 12.37 | 13.63 | 9.90 | 12.32 | 6.24 | 8.54 | 13.00 |
| ANP32A | 17 | 7.75 | 7.57 | 9.51 | 8.03 | 9.46 | 9.12 | 9.05 | 19.88 | 10.11 | 9.52 | 7.66 | 8.77 | 9.29 | 14.47 | 9.81 |
| ANP32B | 19 | 8.47 | 9.07 | 9.63 | 8.61 | 10.27 | 9.95 | 9.58 | 16.38 | 9.32 | 8.73 | 8.77 | 9.12 | 10.11 | 12.98 | 9.02 |
| ANP32E | 6 | 9.05 | 10.07 | 9.96 | 9.94 | 11.16 | 11.48 | 11.05 | 10.73 | 8.05 | 8.52 | 9.56 | 9.95 | 11.32 | 10.89 | 8.28 |
| ANPEP | 3 | 3.30 | 3.18 | 17.80 | 20.43 | 3.51 | 3.72 | 8.33 | 7.25 | 16.33 | 16.15 | 3.24 | 19.12 | 3.62 | 7.79 | 16.24 |
| ANTXR1 | 1 | 12.17 | 7.58 | 10.85 | 11.38 | 10.37 | 10.28 | 5.02 | 6.04 | 13.20 | 13.12 | 9.87 | 11.11 | 10.33 | 5.53 | 13.16 |
| ANXA1 | 80 | 8.31 | 9.79 | 9.07 | 8.76 | 8.62 | 8.45 | 9.93 | 13.82 | 11.84 | 11.40 | 9.05 | 8.91 | 8.54 | 11.88 | 11.62 |
| ANXA10 | 8 | 11.11 | 11.27 | 3.79 | 3.68 | 16.39 | 16.26 | 14.46 | 14.93 | 4.34 | 3.76 | 11.19 | 3.73 | 16.32 | 14.70 | 4.05 |
| ANXA11 | 28 | 10.84 | 11.89 | 9.62 | 9.14 | 11.17 | 11.46 | 8.79 | 8.76 | 9.36 | 8.96 | 11.37 | 9.38 | 11.32 | 8.78 | 9.16 |
| ANXA2 | 177 | 9.02 | 13.54 | 10.97 | 10.89 | 9.79 | 9.85 | 6.87 | 7.22 | 10.93 | 10.90 | 11.28 | 10.93 | 9.82 | 7.05 | 10.92 |
| ANXA3 | 47 | 9.80 | 9.98 | 11.13 | 10.24 | 12.71 | 12.24 | 3.67 | 3.20 | 14.02 | 13.01 | 9.89 | 10.68 | 12.48 | 3.44 | 13.51 |
| ANXA4 | 17 | 9.32 | 10.22 | 8.93 | 8.60 | 12.66 | 12.53 | 9.51 | 9.18 | 9.70 | 9.34 | 9.77 | 8.77 | 12.59 | 9.35 | 9.52 |
| ANXA5 | 78 | 9.15 | 10.44 | 11.05 | 10.16 | 10.35 | 9.98 | 8.66 | 10.19 | 10.14 | 9.88 | 9.79 | 10.61 | 10.16 | 9.43 | 10.01 |
| ANXA6 | 68 | 12.47 | 13.14 | 13.12 | 11.94 | 8.15 | 8.21 | 6.72 | 6.47 | 9.94 | 9.83 | 12.80 | 12.53 | 8.18 | 6.60 | 9.89 |
| ANXA7 | 20 | 9.07 | 9.65 | 11.29 | 9.65 | 11.81 | 10.64 | 10.08 | 8.76 | 9.34 | 9.73 | 9.36 | 10.47 | 11.22 | 9.42 | 9.53 |
| ANXA8 | 16 | 11.94 | 13.01 | 7.53 | 7.20 | 15.06 | 15.14 | 4.35 | 4.23 | 11.13 | 10.41 | 12.47 | 7.37 | 15.10 | 4.29 | 10.77 |
| AP1B1 | 17 | 10.84 | 11.20 | 9.90 | 9.44 | 9.60 | 9.81 | 9.34 | 9.28 | 10.59 | 10.00 | 11.02 | 9.67 | 9.70 | 9.31 | 10.30 |
| AP1G1 | 16 | 10.16 | 10.13 | 7.83 | 7.34 | 11.80 | 11.27 | 8.79 | 12.39 | 10.03 | 10.26 | 10.14 | 7.59 | 11.54 | 10.59 | 10.15 |
| AP1M1 | 15 | 10.39 | 10.92 | 9.41 | 8.27 | 11.06 | 10.98 | 9.43 | 8.38 | 10.59 | 10.57 | 10.65 | 8.84 | 11.02 | 8.90 | 10.58 |
| AP1S1 | 2 | 9.87 | 10.62 | 10.63 | 10.86 | 9.65 | 9.40 | 7.71 | 7.45 | 12.08 | 11.74 | 10.25 | 10.74 | 9.52 | 7.58 | 11.91 |
| AP1S3 | 1 | 10.67 | 11.51 | 8.03 | 7.51 | 13.65 | 13.30 | 9.00 | 7.30 | 8.54 | 10.48 | 11.09 | 7.77 | 13.47 | 8.15 | 9.51 |
| AP2A1 | 42 | 9.87 | 10.53 | 8.09 | 7.98 | 11.81 | 11.78 | 9.33 | 11.77 | 9.54 | 9.30 | 10.20 | 8.03 | 11.79 | 10.55 | 9.42 |
| AP2A2 | 8 | 9.25 | 9.84 | 10.42 | 9.90 | 9.58 | 10.10 | 10.27 | 9.62 | 10.57 | 10.46 | 9.55 | 10.16 | 9.84 | 9.94 | 10.51 |
| AP2B1 | 48 | 9.94 | 9.87 | 9.31 | 7.77 | 11.57 | 11.11 | 8.74 | 12.67 | 9.65 | 9.38 | 9.90 | 8.54 | 11.34 | 10.70 | 9.52 |
| AP2M1 | 16 | 10.19 | 10.83 | 8.23 | 8.80 | 11.22 | 12.36 | 9.72 | 9.53 | 9.75 | 9.37 | 10.51 | 8.51 | 11.79 | 9.63 | 9.56 |
| AP2S1 | 9 | 9.41 | 10.79 | 8.85 | 8.94 | 10.92 | 11.51 | 9.34 | 10.21 | 10.00 | 10.02 | 10.10 | 8.90 | 11.22 | 9.77 | 10.01 |
| AP3B1 | 37 | 8.83 | 9.56 | 10.78 | 10.51 | 11.39 | 11.58 | 8.85 | 8.51 | 9.90 | 10.10 | 9.19 | 10.64 | 11.49 | 8.68 | 10.00 |
| AP3D1 | 28 | 9.34 | 9.61 | 10.36 | 10.79 | 10.98 | 11.32 | 8.34 | 8.51 | 10.39 | 10.35 | 9.48 | 10.57 | 11.15 | 8.43 | 10.37 |
| AP3M1 | 18 | 9.65 | 10.22 | 10.93 | 10.46 | 11.20 | 10.89 | 8.51 | 8.26 | 10.05 | 9.84 | 9.93 | 10.70 | 11.05 | 8.38 | 9.94 |
| AP3S1 | 8 | 8.94 | 9.03 | 11.42 | 9.96 | 10.85 | 10.16 | 9.58 | 8.10 | 11.06 | 10.89 | 8.98 | 10.69 | 10.51 | 8.84 | 10.98 |
| AP4B1 | 3 | 9.68 | 10.69 | 8.36 | 8.43 | 10.41 | 9.74 | 9.74 | 10.68 | 11.08 | 11.19 | 10.19 | 8.40 | 10.07 | 10.21 | 11.14 |
| AP4E1 | 9 | 9.90 | 10.79 | 8.58 | 8.70 | 10.43 | 11.14 | 9.73 | 8.65 | 10.98 | 11.10 | 10.35 | 8.64 | 10.79 | 9.19 | 11.04 |
| AP4M1 | 1 | 8.96 | 10.47 | 9.57 | 8.71 | 10.40 | 10.51 | 10.52 | 9.86 | 10.57 | 10.43 | 9.71 | 9.14 | 10.45 | 10.19 | 10.50 |
| AP5Z1 | 2 | 10.70 | 10.82 | 9.37 | 10.51 | 10.35 | 9.80 | 9.28 | 8.23 | 10.73 | 10.22 | 10.76 | 9.94 | 10.08 | 8.75 | 10.48 |
| APAF1 | 4 | 10.49 | 10.78 | 9.20 | 9.26 | 10.31 | 9.65 | 9.64 | 10.56 | 10.24 | 9.86 | 10.63 | 9.23 | 9.98 | 10.10 | 10.05 |
| APBB1 | 1 | 10.51 | 10.93 | 12.82 | 13.22 | 8.78 | 9.21 | 5.39 | 4.87 | 12.06 | 12.20 | 10.72 | 13.02 | 9.00 | 5.13 | 12.13 |
| APBB2 | 2 | 7.96 | 9.00 | 14.04 | 14.66 | 7.40 | 8.68 | 6.89 | 7.91 | 11.70 | 11.76 | 8.48 | 14.35 | 8.04 | 7.40 | 11.73 |
| APC | 3 | 8.28 | 12.35 | 8.93 | 12.29 | 8.59 | 11.16 | 8.62 | 10.99 | 8.72 | 10.06 | 10.31 | 10.61 | 9.88 | 9.80 | 9.39 |

| | | | | | | | | | | | | | | | | |
|---|---|---|---|---|---|---|---|---|---|---|---|---|---|---|---|---|
| APEH | 32 | 10.99 | 11.35 | 8.53 | 7.67 | 11.18 | 11.12 | 8.32 | 7.59 | 11.82 | 11.43 | 11.17 | 8.10 | 11.15 | 7.96 | 11.62 |
| APEX1 | 24 | 6.84 | 7.22 | 10.58 | 8.90 | 9.69 | 8.86 | 9.51 | 21.13 | 8.67 | 8.61 | 7.03 | 9.74 | 9.27 | 15.32 | 8.64 |
| APEX2 | 2 | 10.03 | 10.40 | 8.60 | 9.01 | 10.78 | 10.52 | 11.55 | 9.70 | 10.40 | 9.02 | 10.22 | 8.80 | 10.65 | 10.62 | 9.71 |
| API5 | 21 | 9.39 | 9.39 | 11.07 | 10.06 | 9.84 | 9.72 | 10.22 | 10.12 | 10.09 | 10.10 | 9.39 | 10.56 | 9.78 | 10.17 | 10.10 |
| APIP | 9 | 9.71 | 10.30 | 9.79 | 9.14 | 9.33 | 9.66 | 12.39 | 11.37 | 9.34 | 8.97 | 10.00 | 9.46 | 9.49 | 11.88 | 9.16 |
| APLP2 | 1 | 10.97 | 7.50 | 10.81 | 14.13 | 8.56 | 8.81 | 8.53 | 8.64 | 10.06 | 11.99 | 9.24 | 12.47 | 8.69 | 8.58 | 11.02 |
| APMAP | 10 | 12.07 | 6.25 | 7.79 | 11.02 | 9.84 | 8.46 | 9.70 | 9.36 | 11.47 | 14.04 | 9.16 | 9.41 | 9.15 | 9.53 | 12.75 |
| APOA1BP | 6 | 9.18 | 10.09 | 10.31 | 7.04 | 15.09 | 13.09 | 10.60 | 8.34 | 7.99 | 8.27 | 9.64 | 8.68 | 14.09 | 9.47 | 8.13 |
| APOB | 4 | 9.02 | 7.89 | 10.29 | 11.56 | 8.20 | 8.56 | 8.31 | 8.76 | 13.64 | 13.76 | 8.46 | 10.93 | 8.38 | 8.54 | 13.70 |
| APOL2 | 8 | 13.21 | 14.48 | 12.92 | 12.55 | 7.88 | 7.77 | 6.45 | 5.90 | 9.54 | 9.31 | 13.84 | 12.73 | 7.82 | 6.18 | 9.42 |
| APOO | 2 | 12.46 | 6.75 | 7.10 | 8.04 | 11.18 | 9.88 | 13.67 | 12.06 | 8.87 | 9.98 | 9.61 | 7.57 | 10.53 | 12.86 | 9.43 |
| APOOL | 2 | 14.00 | 7.13 | 8.74 | 13.23 | 9.65 | 7.76 | 9.33 | 8.07 | 9.79 | 12.31 | 10.56 | 10.98 | 8.71 | 8.70 | 11.05 |
| APP | 3 | 10.18 | 9.38 | 8.21 | 10.97 | 9.42 | 9.53 | 8.53 | 8.36 | 12.33 | 13.08 | 9.78 | 9.59 | 9.47 | 8.45 | 12.70 |
| APPL1 | 19 | 10.45 | 11.25 | 8.14 | 7.65 | 12.40 | 12.59 | 9.77 | 8.73 | 9.60 | 9.42 | 10.85 | 7.89 | 12.49 | 9.25 | 9.51 |
| APPL2 | 6 | 11.00 | 11.20 | 11.63 | 10.56 | 9.18 | 9.19 | 10.43 | 9.28 | 8.82 | 8.71 | 11.10 | 11.10 | 9.19 | 9.85 | 8.77 |
| APRT | 7 | 10.17 | 10.94 | 7.44 | 7.02 | 12.93 | 12.93 | 8.12 | 7.12 | 11.78 | 11.53 | 10.55 | 7.23 | 12.93 | 7.62 | 11.66 |
| APTX | 6 | 10.29 | 10.62 | 12.41 | 11.35 | 7.73 | 8.03 | 7.82 | 7.04 | 12.75 | 11.96 | 10.46 | 11.88 | 7.88 | 7.43 | 12.36 |
| AQR | 24 | 9.76 | 9.38 | 9.34 | 10.24 | 9.48 | 9.26 | 9.95 | 10.45 | 11.09 | 11.06 | 9.57 | 9.79 | 9.37 | 10.20 | 11.07 |
| ARAF | 2 | 9.13 | 9.23 | 7.36 | 7.00 | 12.78 | 11.98 | 13.05 | 13.11 | 8.31 | 8.03 | 9.18 | 7.18 | 12.38 | 13.08 | 8.17 |
| ARAP1 | 12 | 9.39 | 9.91 | 11.30 | 10.19 | 9.22 | 8.99 | 10.05 | 9.02 | 11.21 | 10.71 | 9.65 | 10.74 | 9.11 | 9.54 | 10.96 |
| ARAP3 | 2 | 9.92 | 12.08 | 8.92 | 7.48 | 8.70 | 9.10 | 12.34 | 11.86 | 10.03 | 9.56 | 11.00 | 8.20 | 8.90 | 12.10 | 9.80 |
| ARCN1 | 52 | 8.82 | 10.72 | 10.71 | 10.59 | 9.88 | 10.52 | 9.40 | 10.32 | 9.41 | 9.64 | 9.77 | 10.65 | 10.20 | 9.86 | 9.52 |
| ARF3 | 24 | 10.67 | 8.99 | 8.69 | 9.33 | 10.61 | 10.82 | 9.63 | 9.27 | 10.82 | 11.17 | 9.83 | 9.01 | 10.72 | 9.45 | 11.00 |
| ARF4 | 28 | 9.96 | 9.68 | 9.10 | 9.83 | 9.71 | 9.88 | 8.79 | 9.77 | 11.48 | 11.78 | 9.82 | 9.47 | 9.80 | 9.28 | 11.63 |
| ARF5 | 4 | 9.60 | 9.25 | 8.51 | 9.52 | 9.48 | 9.85 | 10.85 | 11.17 | 10.60 | 11.17 | 9.42 | 9.02 | 9.66 | 11.01 | 10.89 |
| ARF6 | 10 | 9.38 | 7.91 | 8.86 | 10.87 | 9.84 | 10.27 | 11.09 | 10.32 | 10.65 | 10.80 | 8.64 | 9.87 | 10.06 | 10.70 | 10.73 |
| ARFGAP1 | 18 | 8.41 | 9.77 | 13.10 | 12.99 | 8.88 | 9.08 | 9.59 | 9.20 | 9.49 | 9.49 | 9.09 | 13.05 | 8.98 | 9.40 | 9.49 |
| ARFGAP2 | 16 | 9.89 | 10.28 | 9.60 | 8.86 | 9.90 | 9.98 | 7.28 | 10.84 | 12.06 | 11.31 | 10.08 | 9.23 | 9.94 | 9.06 | 11.69 |
| ARFGAP3 | 12 | 8.48 | 9.90 | 12.53 | 12.08 | 7.81 | 8.00 | 11.33 | 10.57 | 9.78 | 9.50 | 9.19 | 12.31 | 7.91 | 10.95 | 9.64 |
| ARFGEF1 | 15 | 10.22 | 9.64 | 9.70 | 10.38 | 9.80 | 9.94 | 9.99 | 9.76 | 10.18 | 10.41 | 9.93 | 10.04 | 9.87 | 9.87 | 10.29 |
| ARFGEF2 | 7 | 8.69 | 8.53 | 8.72 | 9.52 | 10.99 | 10.20 | 9.63 | 10.80 | 11.22 | 11.68 | 8.61 | 9.12 | 10.60 | 10.22 | 11.45 |
| ARFIP1 | 13 | 9.58 | 10.80 | 11.31 | 11.02 | 10.79 | 11.48 | 9.83 | 9.07 | 8.04 | 8.09 | 10.19 | 11.16 | 11.13 | 9.45 | 8.06 |
| ARFIP2 | 8 | 10.01 | 10.45 | 10.91 | 10.32 | 9.84 | 9.55 | 10.03 | 9.24 | 10.04 | 9.61 | 10.23 | 10.61 | 9.70 | 9.64 | 9.82 |
| ARFRP1 | 1 | 9.59 | 11.28 | 9.62 | 9.44 | 8.75 | 10.31 | 9.31 | 8.50 | 12.18 | 11.03 | 10.43 | 9.53 | 9.53 | 8.90 | 11.60 |
| ARGLU1 | 7 | 9.96 | 11.02 | 8.47 | 7.83 | 10.28 | 10.45 | 12.05 | 10.51 | 10.29 | 9.14 | 10.49 | 8.15 | 10.36 | 11.28 | 9.72 |
| ARHGAP1 | 15 | 9.22 | 9.78 | 13.77 | 12.50 | 10.06 | 9.76 | 7.72 | 7.19 | 10.06 | 9.94 | 9.50 | 13.13 | 9.91 | 7.46 | 10.00 |
| ARHGAP10 | 3 | 8.85 | 9.95 | 11.85 | 11.90 | 10.43 | 10.97 | 10.11 | 9.90 | 7.67 | 8.36 | 9.40 | 11.88 | 10.70 | 10.00 | 8.02 |
| ARHGAP11A | 2 | 8.90 | 10.02 | 10.01 | 8.99 | 9.74 | 9.77 | 10.66 | 10.32 | 10.87 | 10.72 | 9.46 | 9.50 | 9.76 | 10.49 | 10.79 |
| ARHGAP12 | 2 | 8.13 | 8.34 | 11.14 | 14.41 | 7.78 | 7.99 | 12.03 | 12.16 | 9.36 | 8.65 | 8.24 | 12.78 | 7.89 | 12.09 | 9.01 |
| ARHGAP17 | 20 | 11.11 | 12.23 | 12.38 | 11.72 | 10.68 | 11.04 | 6.26 | 5.87 | 9.44 | 9.27 | 11.67 | 12.05 | 10.86 | 6.07 | 9.35 |
| ARHGAP18 | 9 | 8.45 | 9.73 | 13.43 | 13.50 | 7.55 | 7.73 | 8.61 | 8.24 | 11.20 | 11.56 | 9.09 | 13.47 | 7.64 | 8.42 | 11.38 |
| ARHGAP21 | 18 | 9.26 | 10.68 | 8.81 | 8.60 | 12.99 | 13.66 | 9.65 | 9.28 | 8.28 | 8.78 | 9.97 | 8.71 | 13.33 | 9.46 | 8.53 |

| | | | | | | | | | | | | | | | | |
|---|---|---|---|---|---|---|---|---|---|---|---|---|---|---|---|---|
| ARHGAP23 | 2 | 11.90 | 11.01 | 10.78 | 14.67 | 7.37 | 7.77 | 6.97 | 6.58 | 11.26 | 11.71 | 11.45 | 12.72 | 7.57 | 6.77 | 11.48 |
| ARHGAP29 | 3 | 10.64 | 10.04 | 12.94 | 12.98 | 6.32 | 6.46 | 7.92 | 8.71 | 12.48 | 11.52 | 10.34 | 12.96 | 6.39 | 8.32 | 12.00 |
| ARHGAP35 | 15 | 9.99 | 10.85 | 10.80 | 10.21 | 9.23 | 9.09 | 11.11 | 9.80 | 9.20 | 9.71 | 10.42 | 10.50 | 9.16 | 10.46 | 9.46 |
| ARHGAP40 | 2 | 14.10 | 10.75 | 5.67 | 6.49 | 10.57 | 10.85 | 9.82 | 10.90 | 11.29 | 9.57 | 12.43 | 6.08 | 10.71 | 10.36 | 10.43 |
| ARHGAP42 | 1 | 9.27 | 10.13 | 9.76 | 9.79 | 16.82 | 16.40 | 6.47 | 5.86 | 7.36 | 8.14 | 9.70 | 9.78 | 16.61 | 6.16 | 7.75 |
| ARHGAP5 | 14 | 9.19 | 9.30 | 11.73 | 9.67 | 11.05 | 10.88 | 9.06 | 8.27 | 10.59 | 10.27 | 9.24 | 10.70 | 10.96 | 8.66 | 10.43 |
| ARHGDIA | 17 | 8.76 | 10.45 | 12.73 | 12.97 | 9.09 | 9.87 | 8.22 | 7.08 | 10.57 | 10.27 | 9.61 | 12.85 | 9.48 | 7.65 | 10.42 |
| ARHGDIB | 1 | 3.05 | 3.55 | 22.27 | 19.86 | 16.54 | 13.24 | 2.50 | 1.65 | 10.08 | 7.26 | 3.30 | 21.07 | 14.89 | 2.07 | 8.67 |
| ARHGEF1 | 24 | 9.23 | 9.91 | 9.23 | 8.40 | 9.77 | 9.69 | 11.58 | 11.36 | 10.66 | 10.17 | 9.57 | 8.81 | 9.73 | 11.47 | 10.41 |
| ARHGEF10 | 11 | 9.33 | 10.08 | 10.43 | 10.42 | 9.26 | 9.73 | 8.71 | 9.05 | 11.66 | 11.33 | 9.71 | 10.43 | 9.49 | 8.88 | 11.49 |
| ARHGEF10L | 1 | 11.30 | 11.24 | 8.79 | 8.28 | 8.09 | 10.41 | 8.14 | 11.03 | 11.66 | 11.05 | 11.27 | 8.54 | 9.25 | 9.59 | 11.36 |
| ARHGEF11 | 4 | 7.96 | 10.62 | 9.94 | 9.97 | 13.15 | 13.23 | 9.30 | 8.69 | 8.39 | 8.72 | 9.29 | 9.96 | 13.19 | 9.00 | 8.56 |
| ARHGEF12 | 4 | 9.17 | 10.78 | 8.92 | 9.16 | 9.74 | 11.55 | 8.80 | 12.75 | 9.69 | 9.45 | 9.98 | 9.04 | 10.64 | 10.77 | 9.57 |
| ARHGEF16 | 4 | 9.40 | 9.21 | 6.20 | 6.49 | 8.96 | 9.46 | 16.64 | 15.37 | 8.91 | 9.37 | 9.30 | 6.35 | 9.21 | 16.00 | 9.14 |
| ARHGEF17 | 5 | 10.03 | 11.68 | 10.34 | 10.48 | 11.27 | 11.59 | 7.35 | 6.26 | 10.65 | 10.35 | 10.85 | 10.41 | 11.43 | 6.81 | 10.50 |
| ARHGEF18 | 8 | 8.90 | 9.21 | 10.44 | 10.38 | 9.65 | 9.47 | 10.57 | 10.33 | 10.74 | 10.31 | 9.05 | 10.41 | 9.56 | 10.45 | 10.52 |
| ARHGEF2 | 21 | 8.89 | 9.81 | 7.62 | 7.48 | 12.33 | 12.84 | 11.23 | 10.97 | 9.62 | 9.22 | 9.35 | 7.55 | 12.58 | 11.10 | 9.42 |
| ARHGEF26 | 1 | 10.58 | 11.46 | 10.97 | 11.62 | 11.32 | 9.63 | 7.92 | 6.04 | 10.00 | 10.45 | 11.02 | 11.30 | 10.48 | 6.98 | 10.22 |
| ARHGEF28 | 1 | 7.07 | 8.98 | 10.31 | 10.74 | 11.33 | 12.09 | 6.80 | 6.43 | 13.37 | 12.87 | 8.02 | 10.53 | 11.71 | 6.62 | 13.12 |
| ARHGEF40 | 8 | 10.86 | 11.79 | 9.13 | 8.96 | 10.23 | 10.09 | 7.64 | 6.96 | 12.23 | 12.11 | 11.32 | 9.05 | 10.16 | 7.30 | 12.17 |
| ARHGEF7 | 16 | 10.49 | 11.22 | 9.54 | 9.73 | 10.73 | 11.16 | 10.13 | 9.29 | 8.98 | 8.74 | 10.86 | 9.63 | 10.94 | 9.71 | 8.86 |
| ARID1A | 27 | 8.94 | 11.02 | 9.10 | 8.66 | 9.91 | 10.48 | 11.51 | 10.82 | 9.91 | 9.64 | 9.98 | 8.88 | 10.20 | 11.16 | 9.78 |
| ARID1B | 13 | 9.43 | 10.26 | 10.24 | 9.63 | 10.90 | 11.51 | 9.69 | 9.44 | 9.55 | 9.34 | 9.84 | 9.94 | 11.21 | 9.57 | 9.45 |
| ARID2 | 10 | 9.67 | 10.61 | 9.88 | 10.03 | 10.48 | 10.99 | 12.85 | 12.06 | 6.99 | 6.43 | 10.14 | 9.96 | 10.74 | 12.45 | 6.71 |
| ARID4B | 12 | 10.18 | 10.60 | 8.95 | 7.07 | 11.04 | 11.73 | 11.77 | 10.16 | 9.38 | 9.13 | 10.39 | 8.01 | 11.39 | 10.97 | 9.25 |
| ARID5B | 1 | 8.65 | 10.92 | 13.32 | 10.78 | 8.35 | 8.88 | 11.49 | 11.50 | 8.67 | 7.44 | 9.79 | 12.05 | 8.61 | 11.49 | 8.05 |
| ARIH1 | 5 | 10.11 | 10.04 | 9.61 | 8.73 | 9.44 | 9.46 | 10.18 | 8.24 | 12.44 | 11.77 | 10.07 | 9.17 | 9.45 | 9.21 | 12.10 |
| ARIH2 | 5 | 7.72 | 7.92 | 7.58 | 6.25 | 8.26 | 7.15 | 8.56 | 30.23 | 8.29 | 8.04 | 7.82 | 6.91 | 7.70 | 19.40 | 8.17 |
| ARL1 | 4 | 10.44 | 9.20 | 7.98 | 7.42 | 12.13 | 9.17 | 11.46 | 12.67 | 8.93 | 10.60 | 9.82 | 7.70 | 10.65 | 12.06 | 9.76 |
| ARL14EP | 2 | 10.07 | 10.63 | 10.52 | 9.11 | 11.81 | 11.25 | 9.37 | 10.41 | 8.41 | 8.42 | 10.35 | 9.82 | 11.53 | 9.89 | 8.41 |
| ARL2 | 11 | 9.84 | 10.50 | 9.23 | 9.38 | 9.61 | 10.20 | 10.18 | 10.08 | 10.68 | 10.28 | 10.17 | 9.31 | 9.90 | 10.13 | 10.48 |
| ARL3 | 6 | 9.52 | 11.40 | 7.78 | 6.58 | 12.48 | 12.81 | 10.31 | 9.25 | 10.10 | 9.75 | 10.46 | 7.18 | 12.65 | 9.78 | 9.93 |
| ARL6 | 2 | 8.79 | 10.08 | 10.73 | 10.20 | 9.39 | 10.98 | 10.23 | 11.25 | 9.26 | 9.11 | 9.43 | 10.46 | 10.18 | 10.74 | 9.18 |
| ARL6IP1 | 2 | 13.64 | 7.88 | 7.56 | 11.50 | 8.56 | 8.22 | 8.65 | 8.78 | 11.47 | 13.71 | 10.76 | 9.53 | 8.39 | 8.72 | 12.59 |
| ARL6IP4 | 4 | 9.70 | 9.96 | 9.36 | 7.90 | 10.27 | 10.33 | 12.09 | 10.91 | 10.67 | 8.80 | 9.83 | 8.63 | 10.30 | 11.50 | 9.73 |
| ARL6IP5 | 9 | 12.48 | 6.98 | 7.05 | 9.93 | 10.01 | 9.24 | 8.03 | 7.13 | 13.27 | 15.88 | 9.73 | 8.49 | 9.62 | 7.58 | 14.57 |
| ARL8B | 11 | 13.14 | 7.25 | 6.99 | 10.77 | 9.91 | 9.56 | 9.16 | 9.58 | 11.32 | 12.31 | 10.20 | 8.88 | 9.74 | 9.37 | 11.82 |
| ARMC1 | 5 | 10.01 | 10.94 | 9.52 | 9.40 | 9.56 | 10.72 | 11.48 | 11.36 | 8.26 | 8.75 | 10.47 | 9.46 | 10.14 | 11.42 | 8.51 |
| ARMC10 | 1 | 10.69 | 6.47 | 11.12 | 14.88 | 5.49 | 5.27 | 11.21 | 11.57 | 10.42 | 12.88 | 8.58 | 13.00 | 5.38 | 11.39 | 11.65 |
| ARMC6 | 14 | 9.71 | 10.40 | 11.56 | 11.40 | 7.67 | 8.07 | 10.66 | 10.01 | 10.52 | 10.02 | 10.05 | 11.48 | 7.87 | 10.33 | 10.27 |
| ARMC8 | 3 | 9.77 | 10.28 | 7.77 | 8.55 | 10.57 | 11.89 | 9.56 | 9.34 | 11.02 | 11.24 | 10.03 | 8.16 | 11.23 | 9.45 | 11.13 |
| ARMC9 | 2 | 11.30 | 10.97 | 14.34 | 14.04 | 7.98 | 7.35 | 6.92 | 6.55 | 10.48 | 10.06 | 11.13 | 14.19 | 7.67 | 6.74 | 10.27 |

| | | | | | | | | | | | | | | | | |
|---|---|---|---|---|---|---|---|---|---|---|---|---|---|---|---|---|
| ARMCX2 | 6 | 9.99 | 7.20 | 12.40 | 15.74 | 7.61 | 6.85 | 14.12 | 12.88 | 6.19 | 7.02 | 8.60 | 14.07 | 7.23 | 13.50 | 6.60 |
| ARMCX3 | 13 | 10.00 | 8.41 | 8.92 | 10.61 | 12.34 | 11.22 | 11.14 | 10.98 | 8.05 | 8.35 | 9.20 | 9.76 | 11.78 | 11.06 | 8.20 |
| ARMCX4 | 1 | 9.09 | 11.43 | 11.54 | 12.31 | 8.52 | 10.06 | 9.98 | 12.62 | 6.89 | 7.57 | 10.26 | 11.92 | 9.29 | 11.30 | 7.23 |
| ARNT | 3 | 9.63 | 9.71 | 8.87 | 7.76 | 9.88 | 10.46 | 9.91 | 16.27 | 9.87 | 7.64 | 9.67 | 8.31 | 10.17 | 13.09 | 8.75 |
| ARNTL | 1 | 8.70 | 9.22 | 10.35 | 9.31 | 9.88 | 10.29 | 12.61 | 9.80 | 9.49 | 10.35 | 8.96 | 9.83 | 10.08 | 11.21 | 9.92 |
| ARPC1A | 18 | 7.78 | 8.87 | 13.70 | 12.89 | 8.85 | 8.96 | 8.17 | 10.13 | 10.15 | 10.49 | 8.32 | 13.30 | 8.91 | 9.15 | 10.32 |
| ARPC1B | 18 | 8.15 | 10.34 | 12.39 | 11.17 | 9.58 | 9.59 | 8.41 | 9.03 | 10.43 | 10.89 | 9.25 | 11.78 | 9.59 | 8.72 | 10.66 |
| ARPC2 | 32 | 9.12 | 8.90 | 13.87 | 10.94 | 9.72 | 8.98 | 8.36 | 8.76 | 10.92 | 10.43 | 9.01 | 12.41 | 9.35 | 8.56 | 10.68 |
| ARPC3 | 10 | 8.50 | 9.49 | 13.01 | 12.91 | 9.12 | 9.36 | 8.18 | 8.32 | 10.88 | 10.22 | 9.00 | 12.96 | 9.24 | 8.25 | 10.55 |
| ARPC4 | 21 | 9.05 | 9.78 | 12.04 | 11.05 | 9.95 | 9.79 | 7.78 | 8.35 | 11.18 | 11.03 | 9.41 | 11.55 | 9.87 | 8.07 | 11.10 |
| ARPC5 | 16 | 9.43 | 10.10 | 12.91 | 11.73 | 10.58 | 10.11 | 7.01 | 6.96 | 11.19 | 9.97 | 9.77 | 12.32 | 10.35 | 6.99 | 10.58 |
| ARPC5L | 15 | 8.19 | 9.82 | 11.85 | 12.81 | 8.46 | 9.23 | 8.97 | 8.88 | 10.73 | 11.08 | 9.01 | 12.33 | 8.84 | 8.92 | 10.90 |
| ARPIN | 5 | 8.96 | 10.33 | 7.34 | 7.16 | 12.71 | 13.23 | 11.05 | 9.54 | 10.52 | 9.17 | 9.65 | 7.25 | 12.97 | 10.29 | 9.84 |
| ARPP19 | 5 | 9.84 | 10.91 | 11.01 | 10.70 | 10.26 | 10.02 | 9.19 | 5.53 | 10.63 | 11.91 | 10.37 | 10.85 | 10.14 | 7.36 | 11.27 |
| ARRB2 | 3 | 9.77 | 11.51 | 10.56 | 11.47 | 8.16 | 9.15 | 9.92 | 9.69 | 9.23 | 10.55 | 10.64 | 11.02 | 8.65 | 9.80 | 9.89 |
| ARRDC1 | 2 | 9.51 | 10.38 | 9.29 | 9.76 | 12.23 | 11.96 | 7.40 | 7.50 | 10.89 | 11.08 | 9.94 | 9.53 | 12.10 | 7.45 | 10.98 |
| ARSA | 2 | 11.15 | 11.05 | 11.55 | 9.49 | 10.01 | 9.85 | 11.47 | 10.21 | 7.75 | 7.46 | 11.10 | 10.52 | 9.93 | 10.84 | 7.61 |
| ARSB | 4 | 11.87 | 9.96 | 7.18 | 7.45 | 14.28 | 14.95 | 11.30 | 9.64 | 8.64 | 4.72 | 10.92 | 7.32 | 14.62 | 10.47 | 6.68 |
| ASAH1 | 6 | 11.35 | 11.61 | 11.20 | 11.55 | 9.88 | 10.35 | 7.64 | 7.39 | 9.86 | 9.16 | 11.48 | 11.37 | 10.12 | 7.52 | 9.51 |
| ASAP1 | 6 | 9.54 | 10.17 | 11.40 | 11.23 | 8.03 | 9.32 | 12.16 | 9.80 | 9.44 | 8.92 | 9.85 | 11.32 | 8.67 | 10.98 | 9.18 |
| ASAP2 | 1 | 8.45 | 7.59 | 16.69 | 13.41 | 11.58 | 7.19 | 10.26 | 5.23 | 9.54 | 10.07 | 8.02 | 15.05 | 9.38 | 7.74 | 9.80 |
| ASB1 | 1 | 8.52 | 10.43 | 11.66 | 12.26 | 9.14 | 8.22 | 7.83 | 8.28 | 12.03 | 11.63 | 9.48 | 11.96 | 8.68 | 8.06 | 11.83 |
| ASB6 | 3 | 10.02 | 10.51 | 10.67 | 6.85 | 9.59 | 8.20 | 11.17 | 7.99 | 13.60 | 11.40 | 10.27 | 8.76 | 8.90 | 9.58 | 12.50 |
| ASCC1 | 7 | 10.78 | 10.77 | 10.82 | 10.09 | 9.84 | 9.70 | 9.66 | 9.13 | 9.82 | 9.39 | 10.78 | 10.46 | 9.77 | 9.39 | 9.61 |
| ASCC2 | 5 | 10.85 | 11.88 | 10.67 | 10.30 | 9.23 | 9.30 | 9.55 | 10.06 | 9.04 | 9.11 | 11.37 | 10.48 | 9.26 | 9.81 | 9.08 |
| ASCC3 | 34 | 11.31 | 11.28 | 9.92 | 10.48 | 9.22 | 9.56 | 10.59 | 11.00 | 8.40 | 8.25 | 11.29 | 10.20 | 9.39 | 10.79 | 8.32 |
| ASF1A | 5 | 10.59 | 11.41 | 9.01 | 9.14 | 11.21 | 11.66 | 9.09 | 8.90 | 9.42 | 9.57 | 11.00 | 9.07 | 11.44 | 8.99 | 9.50 |
| ASF1B | 2 | 13.80 | 13.44 | 7.49 | 9.94 | 7.77 | 8.58 | 10.86 | 10.08 | 10.05 | 7.99 | 13.62 | 8.71 | 8.18 | 10.47 | 9.02 |
| ASH2L | 12 | 8.85 | 10.51 | 10.10 | 9.61 | 10.71 | 11.38 | 10.94 | 10.44 | 8.70 | 8.78 | 9.68 | 9.85 | 11.04 | 10.69 | 8.74 |
| ASL | 9 | 9.69 | 9.91 | 12.40 | 11.07 | 6.80 | 6.96 | 6.69 | 5.76 | 15.82 | 14.90 | 9.80 | 11.73 | 6.88 | 6.22 | 15.36 |
| ASMTL | 4 | 9.33 | 10.36 | 7.69 | 7.20 | 10.58 | 11.69 | 13.44 | 12.81 | 8.53 | 8.38 | 9.85 | 7.44 | 11.13 | 13.12 | 8.45 |
| ASNA1 | 18 | 9.24 | 9.69 | 11.00 | 9.81 | 11.76 | 11.48 | 8.25 | 8.26 | 10.38 | 10.12 | 9.47 | 10.40 | 11.62 | 8.26 | 10.25 |
| ASNS | 23 | 8.00 | 8.74 | 11.19 | 8.66 | 7.27 | 6.64 | 8.81 | 7.86 | 16.53 | 16.30 | 8.37 | 9.93 | 6.95 | 8.34 | 16.42 |
| ASNSD1 | 7 | 9.02 | 10.37 | 12.89 | 11.09 | 8.53 | 8.90 | 12.12 | 11.42 | 8.09 | 7.57 | 9.70 | 11.99 | 8.71 | 11.77 | 7.83 |
| ASPH | 28 | 12.60 | 6.59 | 4.94 | 6.66 | 9.95 | 8.00 | 12.18 | 13.43 | 11.14 | 14.52 | 9.59 | 5.80 | 8.97 | 12.80 | 12.83 |
| ASPM | 5 | 10.02 | 10.06 | 10.11 | 9.61 | 10.63 | 10.41 | 9.66 | 9.70 | 9.92 | 9.88 | 10.04 | 9.86 | 10.52 | 9.68 | 9.90 |
| ASPSCR1 | 3 | 9.86 | 10.92 | 9.93 | 7.49 | 11.70 | 11.19 | 9.67 | 7.71 | 11.09 | 10.44 | 10.39 | 8.71 | 11.45 | 8.69 | 10.76 |
| ASS1 | 10 | 7.64 | 8.08 | 2.34 | 2.11 | 15.41 | 16.38 | 10.27 | 9.37 | 14.45 | 13.95 | 7.86 | 2.22 | 15.89 | 9.82 | 14.20 |
| ASUN | 22 | 9.90 | 10.65 | 10.26 | 9.98 | 9.35 | 9.87 | 9.47 | 8.86 | 11.00 | 10.66 | 10.28 | 10.12 | 9.61 | 9.16 | 10.83 |
| ASXL1 | 1 | 10.62 | 11.07 | 8.94 | 10.43 | 10.02 | 9.08 | 9.51 | 9.22 | 10.58 | 10.52 | 10.85 | 9.69 | 9.55 | 9.37 | 10.55 |
| ASXL2 | 1 | 9.00 | 12.38 | 8.56 | 9.96 | 8.85 | 11.96 | 11.04 | 7.77 | 11.14 | 9.35 | 10.69 | 9.26 | 10.40 | 9.41 | 10.24 |
| ATAD1 | 7 | 10.24 | 7.50 | 9.21 | 10.78 | 12.13 | 11.49 | 7.78 | 8.26 | 10.32 | 12.28 | 8.87 | 10.00 | 11.81 | 8.02 | 11.30 |

| | | | | | | | | | | | | | | | | |
|---|---|---|---|---|---|---|---|---|---|---|---|---|---|---|---|---|
| ATAD2 | 26 | 9.61 | 11.17 | 9.48 | 8.94 | 9.77 | 10.00 | 10.40 | 9.45 | 10.45 | 10.72 | 10.39 | 9.21 | 9.89 | 9.93 | 10.59 |
| ATAD2B | 4 | 9.70 | 10.57 | 8.15 | 8.48 | 11.27 | 11.87 | 10.95 | 9.68 | 9.81 | 9.52 | 10.14 | 8.31 | 11.57 | 10.32 | 9.67 |
| ATAD3A | 15 | 13.84 | 8.58 | 7.52 | 10.26 | 7.00 | 7.15 | 9.39 | 8.57 | 12.79 | 14.90 | 11.21 | 8.89 | 7.08 | 8.98 | 13.84 |
| ATAD3B | 22 | 14.29 | 8.20 | 6.90 | 9.64 | 7.28 | 6.59 | 10.22 | 9.53 | 12.82 | 14.52 | 11.25 | 8.27 | 6.94 | 9.87 | 13.67 |
| ATAD5 | 1 | 10.15 | 10.82 | 9.01 | 10.55 | 5.76 | 7.13 | 12.83 | 11.29 | 10.99 | 11.45 | 10.49 | 9.78 | 6.45 | 12.06 | 11.22 |
| ATE1 | 2 | 9.17 | 10.29 | 9.24 | 9.35 | 12.94 | 12.97 | 9.62 | 7.68 | 10.02 | 8.72 | 9.73 | 9.30 | 12.95 | 8.65 | 9.37 |
| ATF1 | 4 | 9.82 | 11.20 | 9.38 | 9.36 | 8.90 | 8.83 | 12.52 | 11.71 | 9.66 | 8.62 | 10.51 | 9.37 | 8.86 | 12.12 | 9.14 |
| ATF3 | 1 | 11.93 | 12.32 | 10.00 | 7.61 | 10.62 | 10.32 | 6.22 | 5.67 | 13.38 | 11.93 | 12.12 | 8.80 | 10.47 | 5.95 | 12.65 |
| ATF6 | 1 | 10.62 | 6.95 | 7.94 | 10.47 | 13.15 | 11.35 | 8.76 | 8.87 | 10.66 | 11.23 | 8.78 | 9.21 | 12.25 | 8.81 | 10.95 |
| ATF7 | 5 | 9.89 | 11.06 | 9.12 | 8.57 | 10.49 | 10.31 | 11.34 | 9.84 | 9.85 | 9.56 | 10.47 | 8.84 | 10.40 | 10.59 | 9.70 |
| ATF7IP | 5 | 9.33 | 10.33 | 11.20 | 10.44 | 10.90 | 10.31 | 10.90 | 9.07 | 9.25 | 8.28 | 9.83 | 10.82 | 10.60 | 9.99 | 8.76 |
| ATG101 | 4 | 9.81 | 9.90 | 10.66 | 11.17 | 9.65 | 10.32 | 9.36 | 9.03 | 10.46 | 9.64 | 9.86 | 10.92 | 9.98 | 9.19 | 10.05 |
| ATG12 | 4 | 8.62 | 10.47 | 12.44 | 11.52 | 9.14 | 9.39 | 10.11 | 9.23 | 9.68 | 9.39 | 9.55 | 11.98 | 9.26 | 9.67 | 9.54 |
| ATG13 | 1 | 10.84 | 9.76 | 10.79 | 10.98 | 9.72 | 10.53 | 8.99 | 9.96 | 9.37 | 9.04 | 10.30 | 10.88 | 10.13 | 9.48 | 9.21 |
| ATG16L1 | 8 | 10.15 | 10.68 | 11.34 | 9.26 | 9.86 | 9.26 | 10.06 | 8.52 | 10.46 | 10.41 | 10.41 | 10.30 | 9.56 | 9.29 | 10.44 |
| ATG2B | 8 | 9.53 | 9.93 | 11.08 | 10.55 | 9.42 | 9.58 | 9.42 | 9.34 | 10.67 | 10.48 | 9.73 | 10.82 | 9.50 | 9.38 | 10.57 |
| ATG3 | 4 | 9.83 | 10.45 | 11.13 | 9.08 | 10.26 | 10.37 | 10.58 | 8.84 | 10.20 | 9.26 | 10.14 | 10.10 | 10.31 | 9.71 | 9.73 |
| ATG4B | 2 | 10.20 | 9.08 | 10.03 | 8.49 | 9.62 | 9.83 | 13.00 | 11.81 | 9.18 | 8.74 | 9.64 | 9.26 | 9.73 | 12.41 | 8.96 |
| ATG5 | 7 | 9.39 | 9.68 | 11.31 | 10.99 | 9.22 | 9.39 | 10.06 | 8.83 | 10.85 | 10.28 | 9.54 | 11.15 | 9.31 | 9.44 | 10.56 |
| ATG7 | 7 | 11.24 | 11.30 | 10.12 | 9.76 | 9.25 | 8.67 | 11.48 | 13.92 | 7.18 | 7.08 | 11.27 | 9.94 | 8.96 | 12.70 | 7.13 |
| ATG9A | 7 | 9.40 | 8.19 | 10.86 | 10.44 | 11.71 | 10.60 | 8.35 | 7.91 | 11.19 | 11.35 | 8.80 | 10.65 | 11.16 | 8.13 | 11.27 |
| ATIC | 65 | 8.92 | 9.95 | 10.84 | 10.24 | 10.59 | 10.54 | 9.44 | 10.34 | 9.76 | 9.38 | 9.44 | 10.54 | 10.56 | 9.89 | 9.57 |
| ATL2 | 9 | 10.14 | 6.76 | 9.57 | 13.26 | 8.63 | 8.20 | 11.55 | 11.04 | 9.82 | 11.04 | 8.45 | 11.41 | 8.41 | 11.29 | 10.43 |
| ATL3 | 19 | 11.38 | 6.83 | 7.43 | 11.02 | 9.94 | 8.57 | 9.29 | 9.58 | 11.63 | 14.32 | 9.10 | 9.23 | 9.26 | 9.44 | 12.98 |
| ATM | 3 | 8.91 | 9.24 | 10.38 | 11.49 | 10.41 | 9.83 | 11.58 | 9.06 | 8.89 | 10.21 | 9.08 | 10.93 | 10.12 | 10.32 | 9.55 |
| ATN1 | 2 | 9.16 | 11.20 | 10.29 | 11.41 | 10.48 | 12.40 | 9.04 | 7.22 | 9.43 | 9.36 | 10.18 | 10.85 | 11.44 | 8.13 | 9.40 |
| ATOX1 | 5 | 4.28 | 9.67 | 14.40 | 11.27 | 10.12 | 8.61 | 15.33 | 12.00 | 6.64 | 7.68 | 6.98 | 12.83 | 9.37 | 13.66 | 7.16 |
| ATP11C | 2 | 10.05 | 8.36 | 8.52 | 10.87 | 8.46 | 8.53 | 12.33 | 13.29 | 9.47 | 10.11 | 9.21 | 9.69 | 8.50 | 12.81 | 9.79 |
| ATP13A1 | 3 | 11.32 | 6.04 | 6.75 | 10.12 | 11.71 | 11.15 | 10.71 | 9.56 | 10.34 | 12.31 | 8.68 | 8.43 | 11.43 | 10.14 | 11.32 |
| ATP1A1 | 47 | 12.64 | 9.63 | 8.51 | 9.70 | 8.09 | 7.88 | 10.32 | 9.02 | 11.85 | 12.36 | 11.13 | 9.11 | 7.98 | 9.67 | 12.11 |
| ATP1B1 | 7 | 11.85 | 8.93 | 10.23 | 12.39 | 9.03 | 9.17 | 7.32 | 6.27 | 11.88 | 12.93 | 10.39 | 11.31 | 9.10 | 6.80 | 12.40 |
| ATP1B2 | 1 | 11.37 | 11.64 | 9.38 | 9.51 | 9.77 | 11.32 | 7.78 | 10.50 | 10.05 | 8.69 | 11.50 | 9.44 | 10.54 | 9.14 | 9.37 |
| ATP1B3 | 16 | 12.67 | 9.73 | 8.03 | 9.00 | 7.51 | 7.14 | 12.78 | 10.81 | 10.64 | 11.66 | 11.20 | 8.52 | 7.33 | 11.80 | 11.15 |
| ATP2A2 | 32 | 12.99 | 6.37 | 7.89 | 12.33 | 8.56 | 7.37 | 10.07 | 10.08 | 10.87 | 13.47 | 9.68 | 10.11 | 7.97 | 10.07 | 12.17 |
| ATP2A3 | 6 | 10.97 | 6.30 | 9.01 | 12.65 | 8.10 | 7.00 | 9.21 | 13.73 | 10.19 | 12.82 | 8.64 | 10.83 | 7.55 | 11.47 | 11.50 |
| ATP2B1 | 16 | 9.84 | 8.24 | 9.29 | 11.49 | 10.42 | 10.13 | 10.51 | 9.50 | 9.90 | 10.67 | 9.04 | 10.39 | 10.28 | 10.01 | 10.28 |
| ATP2B4 | 10 | 12.35 | 9.24 | 5.74 | 7.81 | 13.07 | 13.43 | 7.50 | 6.79 | 11.79 | 12.29 | 10.79 | 6.77 | 13.25 | 7.14 | 12.04 |
| ATP2C1 | 5 | 11.05 | 7.33 | 9.98 | 12.31 | 8.71 | 7.96 | 8.41 | 7.66 | 12.95 | 13.64 | 9.19 | 11.15 | 8.33 | 8.03 | 13.29 |
| ATP5A1 | 96 | 12.16 | 9.57 | 9.42 | 11.20 | 11.38 | 10.29 | 8.17 | 7.51 | 10.03 | 10.27 | 10.87 | 10.31 | 10.84 | 7.84 | 10.15 |
| ATP5B | 86 | 12.10 | 9.58 | 9.53 | 9.81 | 11.91 | 10.79 | 8.35 | 7.57 | 10.06 | 10.31 | 10.84 | 9.67 | 11.35 | 7.96 | 10.18 |
| ATP5C1 | 23 | 11.50 | 8.75 | 9.37 | 9.67 | 11.56 | 10.46 | 9.74 | 9.64 | 9.36 | 9.94 | 10.13 | 9.52 | 11.01 | 9.69 | 9.65 |
| ATP5D | 5 | 10.93 | 9.26 | 9.57 | 9.56 | 11.77 | 11.35 | 9.92 | 7.49 | 10.05 | 10.11 | 10.10 | 9.56 | 11.56 | 8.71 | 10.08 |

| | | | | | | | | | | | | | | | | |
|---|---|---|---|---|---|---|---|---|---|---|---|---|---|---|---|---|
| ATP5E | 6 | 7.30 | 5.43 | 7.48 | 4.88 | 9.72 | 7.58 | 7.71 | 35.86 | 7.39 | 6.66 | 6.37 | 6.18 | 8.65 | 21.79 | 7.02 |
| ATP5F1 | 15 | 15.59 | 7.57 | 7.17 | 9.36 | 11.14 | 9.17 | 7.96 | 6.51 | 11.84 | 13.69 | 11.58 | 8.27 | 10.16 | 7.23 | 12.76 |
| ATP5G2 | 2 | 13.43 | 6.38 | 10.04 | 11.22 | 9.08 | 7.56 | 10.29 | 11.11 | 10.50 | 10.39 | 9.90 | 10.63 | 8.32 | 10.70 | 10.44 |
| ATP5H | 9 | 15.91 | 8.21 | 7.60 | 11.89 | 10.74 | 9.24 | 7.29 | 5.91 | 11.11 | 12.09 | 12.06 | 9.75 | 9.99 | 6.60 | 11.60 |
| ATP5I | 4 | 16.19 | 8.07 | 8.25 | 11.02 | 11.39 | 9.52 | 6.99 | 7.10 | 10.85 | 10.62 | 12.13 | 9.64 | 10.46 | 7.05 | 10.73 |
| ATP5J | 3 | 13.82 | 9.93 | 10.42 | 12.64 | 8.07 | 9.09 | 10.32 | 4.19 | 8.47 | 13.06 | 11.87 | 11.53 | 8.58 | 7.26 | 10.77 |
| ATP5J2-PTC[illegible] | 5 | 10.55 | 7.15 | 9.64 | 12.13 | 7.72 | 7.25 | 12.87 | 12.46 | 9.49 | 10.74 | 8.85 | 10.89 | 7.48 | 12.66 | 10.12 |
| ATP5L | 4 | 16.71 | 8.13 | 7.28 | 11.27 | 10.77 | 8.93 | 7.12 | 6.75 | 10.92 | 12.12 | 12.42 | 9.28 | 9.85 | 6.93 | 11.52 |
| ATP5O | 15 | 10.26 | 9.14 | 10.08 | 10.07 | 11.95 | 10.46 | 8.60 | 9.86 | 9.52 | 10.06 | 9.70 | 10.07 | 11.21 | 9.23 | 9.79 |
| ATP6AP1 | 6 | 12.02 | 7.32 | 6.94 | 10.92 | 11.34 | 10.21 | 9.96 | 9.90 | 10.02 | 11.38 | 9.67 | 8.93 | 10.77 | 9.93 | 10.70 |
| ATP6AP2 | 5 | 10.50 | 7.94 | 7.49 | 9.49 | 10.90 | 10.73 | 12.34 | 11.81 | 8.70 | 10.10 | 9.22 | 8.49 | 10.81 | 12.08 | 9.40 |
| ATP6V0A1 | 5 | 10.25 | 8.54 | 10.56 | 10.75 | 9.99 | 9.79 | 8.74 | 8.60 | 11.11 | 11.66 | 9.40 | 10.66 | 9.89 | 8.67 | 11.39 |
| ATP6V0A2 | 1 | 9.86 | 7.62 | 8.97 | 8.63 | 11.22 | 10.41 | 11.72 | 11.04 | 11.69 | 8.84 | 8.74 | 8.80 | 10.82 | 11.38 | 10.27 |
| ATP6V0D1 | 5 | 10.41 | 10.77 | 9.50 | 10.74 | 12.37 | 13.06 | 6.37 | 7.11 | 9.73 | 9.94 | 10.59 | 10.12 | 12.72 | 6.74 | 9.83 |
| ATP6V1A | 45 | 10.10 | 10.55 | 10.89 | 9.39 | 12.44 | 12.27 | 7.89 | 7.67 | 9.58 | 9.22 | 10.32 | 10.14 | 12.35 | 7.78 | 9.40 |
| ATP6V1B2 | 23 | 9.95 | 10.48 | 10.89 | 10.55 | 9.87 | 10.70 | 9.75 | 8.93 | 9.66 | 9.22 | 10.21 | 10.72 | 10.29 | 9.34 | 9.44 |
| ATP6V1C1 | 16 | 9.66 | 10.58 | 10.54 | 9.84 | 10.11 | 10.32 | 10.35 | 9.93 | 9.35 | 9.33 | 10.12 | 10.19 | 10.21 | 10.14 | 9.34 |
| ATP6V1D | 10 | 10.57 | 11.27 | 9.32 | 9.14 | 11.89 | 12.22 | 8.68 | 7.51 | 10.05 | 9.36 | 10.92 | 9.23 | 12.06 | 8.09 | 9.70 |
| ATP6V1E1 | 19 | 9.76 | 10.61 | 11.10 | 11.20 | 10.03 | 10.01 | 9.43 | 8.60 | 9.82 | 9.43 | 10.19 | 11.15 | 10.02 | 9.02 | 9.63 |
| ATP6V1F | 4 | 10.88 | 10.52 | 10.03 | 8.72 | 12.30 | 11.95 | 9.26 | 4.62 | 10.76 | 10.95 | 10.70 | 9.38 | 12.13 | 6.94 | 10.86 |
| ATP6V1G1 | 4 | 9.36 | 9.61 | 10.47 | 9.87 | 10.02 | 9.84 | 9.13 | 12.53 | 9.79 | 9.36 | 9.49 | 10.17 | 9.93 | 10.83 | 9.58 |
| ATP6V1H | 9 | 10.73 | 10.26 | 9.50 | 9.70 | 10.52 | 10.66 | 9.37 | 8.93 | 10.14 | 10.19 | 10.50 | 9.60 | 10.59 | 9.15 | 10.16 |
| ATP7A | 4 | 8.65 | 6.89 | 7.70 | 8.48 | 11.44 | 10.48 | 15.83 | 14.87 | 7.62 | 8.03 | 7.77 | 8.09 | 10.96 | 15.35 | 7.82 |
| ATPAF1 | 2 | 11.08 | 10.03 | 7.33 | 7.72 | 9.18 | 8.90 | 13.32 | 13.97 | 9.35 | 9.10 | 10.55 | 7.53 | 9.04 | 13.65 | 9.23 |
| ATPAF2 | 1 | 12.87 | 9.60 | 12.18 | 9.54 | 9.77 | 9.26 | 10.51 | 9.10 | 8.64 | 8.54 | 11.23 | 10.86 | 9.51 | 9.81 | 8.59 |
| ATPIF1 | 6 | 10.88 | 10.98 | 9.07 | 8.77 | 11.58 | 11.83 | 11.13 | 8.29 | 8.36 | 9.12 | 10.93 | 8.92 | 11.71 | 9.71 | 8.74 |
| ATR | 10 | 9.75 | 10.37 | 9.65 | 11.20 | 9.74 | 10.71 | 9.81 | 9.75 | 9.61 | 9.42 | 10.06 | 10.42 | 10.23 | 9.78 | 9.52 |
| ATRAID | 2 | 9.38 | 7.73 | 8.78 | 12.04 | 10.42 | 10.09 | 10.47 | 10.24 | 9.72 | 11.12 | 8.56 | 10.41 | 10.25 | 10.36 | 10.42 |
| ATRIP | 5 | 10.24 | 10.79 | 10.02 | 10.52 | 10.01 | 9.50 | 10.54 | 9.05 | 10.19 | 9.12 | 10.52 | 10.27 | 9.76 | 9.80 | 9.66 |
| ATRX | 29 | 8.89 | 10.25 | 8.67 | 8.37 | 12.20 | 12.49 | 11.11 | 12.30 | 8.05 | 7.66 | 9.57 | 8.52 | 12.35 | 11.71 | 7.85 |
| ATXN10 | 23 | 9.91 | 10.86 | 11.87 | 11.85 | 12.08 | 12.33 | 7.05 | 7.47 | 8.31 | 8.27 | 10.39 | 11.86 | 12.21 | 7.26 | 8.29 |
| ATXN2 | 10 | 9.84 | 11.12 | 10.66 | 10.29 | 8.81 | 9.12 | 9.74 | 11.06 | 9.78 | 9.58 | 10.48 | 10.47 | 8.96 | 10.40 | 9.68 |
| ATXN2L | 44 | 9.54 | 10.55 | 11.37 | 10.41 | 9.52 | 9.83 | 9.74 | 8.75 | 10.30 | 10.01 | 10.04 | 10.89 | 9.67 | 9.24 | 10.15 |
| ATXN3 | 4 | 9.31 | 10.34 | 11.04 | 10.47 | 10.37 | 10.78 | 10.88 | 8.72 | 9.15 | 8.96 | 9.83 | 10.75 | 10.57 | 9.80 | 9.05 |
| ATXN7 | 2 | 8.87 | 10.08 | 9.73 | 7.99 | 12.25 | 12.33 | 9.78 | 8.26 | 10.41 | 10.29 | 9.47 | 8.86 | 12.29 | 9.02 | 10.35 |
| ATXN7L3 | 1 | 8.44 | 9.23 | 10.45 | 10.11 | 8.27 | 9.53 | 12.18 | 10.87 | 10.76 | 10.16 | 8.83 | 10.28 | 8.90 | 11.52 | 10.46 |
| ATXN7L3B | 1 | 9.94 | 9.82 | 10.70 | 9.82 | 9.42 | 10.05 | 12.68 | 9.42 | 9.05 | 9.10 | 9.88 | 10.26 | 9.74 | 11.05 | 9.08 |
| AUH | 1 | 8.45 | 9.92 | 9.06 | 11.45 | 12.74 | 13.42 | 12.53 | 9.56 | 6.41 | 6.47 | 9.18 | 10.26 | 13.08 | 11.04 | 6.44 |
| AUP1 | 6 | 12.40 | 6.58 | 7.82 | 11.54 | 10.06 | 8.58 | 9.51 | 8.28 | 11.05 | 14.18 | 9.49 | 9.68 | 9.32 | 8.89 | 12.61 |
| AURKA | 9 | 9.22 | 9.29 | 10.01 | 8.66 | 11.16 | 10.79 | 10.41 | 10.62 | 10.26 | 9.57 | 9.25 | 9.34 | 10.98 | 10.52 | 9.91 |
| AURKAIP1 | 1 | 11.47 | 9.28 | 9.01 | 12.16 | 8.98 | 10.17 | 10.64 | 8.81 | 10.12 | 9.36 | 10.38 | 10.58 | 9.57 | 9.73 | 9.74 |
| AURKB | 3 | 8.34 | 9.61 | 11.65 | 11.97 | 7.37 | 7.67 | 11.28 | 10.74 | 10.50 | 10.87 | 8.98 | 11.81 | 7.52 | 11.01 | 10.68 |

| | | | | | | | | | | | | | | | | |
|---|---|---|---|---|---|---|---|---|---|---|---|---|---|---|---|---|
| AVEN | 3 | 9.89 | 10.55 | 10.84 | 8.97 | 10.62 | 10.57 | 9.13 | 9.00 | 10.87 | 9.57 | 10.22 | 9.90 | 10.59 | 9.06 | 10.22 |
| AVL9 | 5 | 8.78 | 12.06 | 11.93 | 11.67 | 8.85 | 8.47 | 10.11 | 8.22 | 9.95 | 9.96 | 10.42 | 11.80 | 8.66 | 9.17 | 9.95 |
| AXIN1 | 2 | 8.96 | 11.03 | 11.95 | 11.76 | 11.41 | 12.37 | 7.39 | 6.88 | 9.22 | 9.04 | 9.99 | 11.85 | 11.89 | 7.13 | 9.13 |
| AXL | 7 | 12.39 | 9.91 | 6.14 | 6.67 | 9.56 | 9.99 | 9.10 | 8.23 | 13.51 | 14.50 | 11.15 | 6.40 | 9.78 | 8.67 | 14.00 |
| AZI1 | 13 | 11.28 | 13.32 | 10.20 | 9.97 | 10.37 | 10.67 | 7.13 | 6.57 | 10.26 | 10.22 | 12.30 | 10.09 | 10.52 | 6.85 | 10.24 |
| B2M | 4 | 10.35 | 9.64 | 6.58 | 3.57 | 19.76 | 16.24 | 6.96 | 3.96 | 11.53 | 11.42 | 9.99 | 5.08 | 18.00 | 5.46 | 11.47 |
| B3GALT6 | 1 | 12.97 | 7.61 | 8.09 | 8.12 | 8.52 | 7.50 | 12.52 | 11.47 | 11.41 | 11.80 | 10.29 | 8.10 | 8.01 | 11.99 | 11.60 |
| B3GALTL | 1 | 10.75 | 9.20 | 8.69 | 9.55 | 16.55 | 16.92 | 7.18 | 7.04 | 6.93 | 7.19 | 9.98 | 9.12 | 16.73 | 7.11 | 7.06 |
| B3GAT3 | 2 | 9.46 | 7.98 | 8.45 | 8.67 | 10.98 | 10.78 | 10.45 | 13.78 | 9.66 | 9.79 | 8.72 | 8.56 | 10.88 | 12.11 | 9.73 |
| B4GALT1 | 14 | 11.00 | 7.61 | 8.26 | 10.41 | 9.83 | 8.99 | 8.71 | 8.63 | 12.65 | 13.92 | 9.30 | 9.33 | 9.41 | 8.67 | 13.29 |
| B4GALT4 | 1 | 14.20 | 9.25 | 5.00 | 6.19 | 14.97 | 13.98 | 10.08 | 11.30 | 7.06 | 7.98 | 11.72 | 5.59 | 14.48 | 10.69 | 7.52 |
| BABAM1 | 9 | 8.88 | 10.54 | 10.81 | 10.39 | 11.02 | 11.23 | 10.31 | 10.50 | 8.18 | 8.15 | 9.71 | 10.60 | 11.12 | 10.40 | 8.17 |
| BAD | 6 | 10.62 | 11.68 | 11.81 | 11.70 | 11.14 | 11.68 | 6.69 | 5.58 | 9.82 | 9.28 | 11.15 | 11.75 | 11.41 | 6.14 | 9.55 |
| BAG1 | 8 | 8.53 | 9.12 | 5.72 | 5.60 | 14.17 | 13.39 | 8.62 | 10.91 | 12.40 | 11.52 | 8.83 | 5.66 | 13.78 | 9.77 | 11.96 |
| BAG2 | 24 | 10.26 | 10.61 | 10.65 | 9.18 | 10.76 | 10.31 | 8.14 | 10.87 | 9.96 | 9.25 | 10.44 | 9.91 | 10.54 | 9.51 | 9.60 |
| BAG3 | 37 | 10.25 | 11.26 | 12.41 | 12.34 | 8.94 | 9.42 | 6.94 | 6.39 | 11.06 | 10.99 | 10.75 | 12.37 | 9.18 | 6.66 | 11.03 |
| BAG4 | 3 | 10.29 | 10.99 | 9.97 | 10.29 | 9.38 | 9.47 | 12.10 | 8.12 | 9.89 | 9.50 | 10.64 | 10.13 | 9.43 | 10.11 | 9.69 |
| BAG5 | 6 | 9.79 | 10.46 | 9.00 | 9.39 | 11.24 | 11.65 | 9.91 | 9.69 | 9.82 | 9.06 | 10.13 | 9.19 | 11.45 | 9.80 | 9.44 |
| BAG6 | 22 | 9.63 | 10.62 | 10.13 | 9.65 | 9.88 | 10.22 | 10.07 | 10.03 | 10.01 | 9.76 | 10.12 | 9.89 | 10.05 | 10.05 | 9.88 |
| BAHCC1 | 2 | 20.39 | 10.89 | 7.06 | 9.02 | 8.16 | 9.73 | 7.87 | 7.78 | 10.68 | 8.41 | 15.64 | 8.04 | 8.95 | 7.83 | 9.54 |
| BAIAP2 | 12 | 11.39 | 11.87 | 11.15 | 11.14 | 8.20 | 8.45 | 7.88 | 6.73 | 11.78 | 11.41 | 11.63 | 11.15 | 8.32 | 7.31 | 11.59 |
| BAIAP2L1 | 11 | 9.85 | 11.12 | 13.99 | 13.38 | 7.34 | 7.56 | 6.27 | 5.45 | 12.67 | 12.37 | 10.48 | 13.69 | 7.45 | 5.86 | 12.52 |
| BAK1 | 1 | 10.76 | 9.23 | 8.84 | 9.84 | 9.51 | 10.23 | 8.92 | 10.26 | 10.92 | 11.49 | 10.00 | 9.34 | 9.87 | 9.59 | 11.21 |
| BANF1 | 15 | 9.48 | 8.41 | 9.95 | 10.58 | 10.43 | 10.30 | 9.26 | 11.94 | 9.58 | 10.07 | 8.95 | 10.27 | 10.36 | 10.60 | 9.83 |
| BAP1 | 2 | 9.56 | 9.80 | 9.85 | 9.37 | 10.07 | 10.20 | 11.28 | 9.97 | 10.39 | 9.51 | 9.68 | 9.61 | 10.13 | 10.63 | 9.95 |
| BAP18 | 2 | 8.89 | 9.07 | 15.28 | 15.32 | 6.52 | 6.50 | 7.98 | 7.18 | 11.99 | 11.26 | 8.98 | 15.30 | 6.51 | 7.58 | 11.63 |
| BARD1 | 3 | 9.98 | 10.66 | 11.56 | 11.62 | 8.09 | 8.06 | 8.79 | 8.69 | 11.24 | 11.31 | 10.32 | 11.59 | 8.08 | 8.74 | 11.27 |
| BASP1 | 50 | 13.58 | 10.98 | 5.95 | 11.20 | 9.63 | 13.32 | 9.37 | 3.99 | 10.11 | 11.88 | 12.28 | 8.57 | 11.47 | 6.68 | 10.99 |
| BAX | 9 | 9.52 | 10.00 | 9.02 | 9.01 | 9.21 | 9.64 | 11.62 | 10.92 | 11.14 | 9.92 | 9.76 | 9.01 | 9.43 | 11.27 | 10.53 |
| BAZ1A | 18 | 9.38 | 10.17 | 10.06 | 10.67 | 8.52 | 8.97 | 11.55 | 11.27 | 9.64 | 9.79 | 9.77 | 10.36 | 8.74 | 11.41 | 9.71 |
| BAZ1B | 34 | 10.91 | 9.56 | 8.82 | 9.84 | 8.71 | 8.39 | 11.54 | 10.41 | 9.69 | 12.14 | 10.24 | 9.33 | 8.55 | 10.97 | 10.92 |
| BAZ2A | 7 | 9.41 | 9.50 | 10.87 | 11.67 | 9.40 | 9.66 | 10.31 | 9.54 | 9.80 | 9.85 | 9.45 | 11.27 | 9.53 | 9.92 | 9.83 |
| BAZ2B | 3 | 9.49 | 10.69 | 12.16 | 11.63 | 10.84 | 11.08 | 8.07 | 7.72 | 9.22 | 9.09 | 10.09 | 11.90 | 10.96 | 7.90 | 9.15 |
| BBX | 3 | 8.68 | 10.58 | 10.23 | 10.12 | 11.24 | 10.34 | 11.13 | 10.63 | 7.31 | 9.75 | 9.63 | 10.17 | 10.79 | 10.88 | 8.53 |
| BCAM | 1 | 10.65 | 9.87 | 8.87 | 13.50 | 6.07 | 5.73 | 9.30 | 9.52 | 14.45 | 12.05 | 10.26 | 11.18 | 5.90 | 9.41 | 13.25 |
| BCAP29 | 3 | 10.15 | 5.52 | 9.76 | 16.25 | 7.92 | 6.98 | 8.63 | 8.63 | 10.46 | 15.69 | 7.84 | 13.00 | 7.45 | 8.63 | 13.08 |
| BCAP31 | 22 | 10.48 | 5.85 | 7.11 | 10.36 | 12.02 | 10.15 | 11.92 | 11.45 | 9.11 | 11.54 | 8.16 | 8.74 | 11.09 | 11.68 | 10.33 |
| BCAR1 | 13 | 10.77 | 11.96 | 8.36 | 7.87 | 9.82 | 10.45 | 8.95 | 8.46 | 11.59 | 11.77 | 11.36 | 8.12 | 10.13 | 8.70 | 11.68 |
| BCAR3 | 6 | 8.13 | 8.98 | 10.36 | 9.94 | 7.33 | 7.36 | 12.99 | 11.90 | 11.82 | 11.21 | 8.56 | 10.15 | 7.34 | 12.44 | 11.51 |
| BCAS2 | 16 | 9.43 | 10.84 | 10.83 | 10.43 | 9.94 | 9.95 | 10.90 | 10.18 | 8.87 | 8.62 | 10.14 | 10.63 | 9.94 | 10.54 | 8.75 |
| BCAS3 | 1 | 9.97 | 9.51 | 9.27 | 9.94 | 9.31 | 9.79 | 9.26 | 19.54 | 7.07 | 6.32 | 9.74 | 9.61 | 9.55 | 14.40 | 6.70 |
| BCAT1 | 9 | 5.25 | 5.21 | 12.28 | 11.35 | 4.17 | 3.92 | 12.32 | 21.31 | 12.04 | 12.15 | 5.23 | 11.81 | 4.05 | 16.81 | 12.09 |

| | | | | | | | | | | | | | | | | |
|---|---|---|---|---|---|---|---|---|---|---|---|---|---|---|---|---|
| BCAT2 | 11 | 10.44 | 9.18 | 10.40 | 10.55 | 10.49 | 10.07 | 11.15 | 9.55 | 9.01 | 9.17 | 9.81 | 10.47 | 10.28 | 10.35 | 9.09 |
| BCCIP | 15 | 8.95 | 10.29 | 11.79 | 10.43 | 8.74 | 9.14 | 10.98 | 10.38 | 9.81 | 9.51 | 9.62 | 11.11 | 8.94 | 10.68 | 9.66 |
| BCKDHA | 8 | 11.52 | 10.86 | 9.33 | 10.17 | 9.60 | 8.43 | 9.64 | 8.55 | 10.58 | 11.31 | 11.19 | 9.75 | 9.02 | 9.10 | 10.95 |
| BCKDHB | 2 | 11.19 | 11.65 | 9.73 | 10.98 | 9.06 | 8.73 | 7.88 | 7.59 | 11.08 | 12.11 | 11.42 | 10.35 | 8.90 | 7.74 | 11.59 |
| BCKDK | 2 | 9.34 | 10.66 | 10.37 | 11.08 | 9.28 | 8.66 | 10.46 | 8.35 | 10.71 | 11.08 | 10.00 | 10.72 | 8.97 | 9.41 | 10.90 |
| BCL10 | 3 | 10.32 | 11.32 | 12.43 | 12.39 | 7.07 | 7.56 | 9.75 | 8.95 | 10.17 | 10.05 | 10.82 | 12.41 | 7.32 | 9.35 | 10.11 |
| BCL2L12 | 1 | 12.16 | 9.83 | 5.23 | 7.51 | 7.31 | 7.49 | 12.92 | 10.80 | 12.03 | 14.72 | 11.00 | 6.37 | 7.40 | 11.86 | 13.37 |
| BCL3 | 5 | 8.55 | 9.86 | 8.34 | 7.99 | 8.88 | 9.39 | 16.25 | 14.05 | 8.81 | 7.88 | 9.21 | 8.16 | 9.14 | 15.15 | 8.35 |
| BCL7A | 4 | 9.58 | 9.82 | 10.91 | 11.27 | 12.18 | 12.56 | 9.73 | 7.02 | 8.69 | 8.23 | 9.70 | 11.09 | 12.37 | 8.37 | 8.46 |
| BCL7B | 1 | 11.06 | 9.44 | 10.97 | 10.11 | 8.95 | 9.41 | 12.88 | 7.38 | 9.76 | 10.05 | 10.25 | 10.54 | 9.18 | 10.13 | 9.90 |
| BCL7C | 4 | 9.49 | 10.42 | 9.14 | 8.94 | 11.51 | 12.12 | 10.87 | 8.88 | 9.75 | 8.88 | 9.96 | 9.04 | 11.82 | 9.87 | 9.31 |
| BCL9 | 1 | 8.26 | 9.30 | 11.19 | 10.92 | 9.48 | 9.82 | 13.58 | 11.59 | 8.80 | 7.05 | 8.78 | 11.06 | 9.65 | 12.58 | 7.93 |
| BCL9L | 4 | 8.65 | 10.22 | 11.28 | 10.95 | 9.52 | 10.77 | 10.17 | 10.37 | 8.60 | 9.46 | 9.43 | 11.11 | 10.15 | 10.27 | 9.03 |
| BCLAF1 | 34 | 8.60 | 9.88 | 10.03 | 8.45 | 9.46 | 9.08 | 10.95 | 15.81 | 9.25 | 8.50 | 9.24 | 9.24 | 9.27 | 13.38 | 8.87 |
| BCOR | 2 | 8.06 | 10.13 | 10.83 | 10.18 | 10.61 | 11.34 | 10.93 | 8.91 | 9.48 | 9.54 | 9.09 | 10.51 | 10.97 | 9.92 | 9.51 |
| BCR | 12 | 9.66 | 10.40 | 10.53 | 10.36 | 7.78 | 8.29 | 9.45 | 8.57 | 12.54 | 12.41 | 10.03 | 10.45 | 8.04 | 9.01 | 12.48 |
| BCS1L | 8 | 10.92 | 8.39 | 8.77 | 10.30 | 9.89 | 9.00 | 11.01 | 9.79 | 11.05 | 10.88 | 9.66 | 9.53 | 9.45 | 10.40 | 10.96 |
| BDH2 | 5 | 9.74 | 10.47 | 12.34 | 11.86 | 10.94 | 11.17 | 9.89 | 11.84 | 5.90 | 5.85 | 10.10 | 12.10 | 11.06 | 10.87 | 5.87 |
| BECN1 | 4 | 9.87 | 10.86 | 11.18 | 11.34 | 9.11 | 9.43 | 8.91 | 7.82 | 11.22 | 10.25 | 10.37 | 11.26 | 9.27 | 8.36 | 10.74 |
| BEGAIN | 6 | 11.19 | 12.04 | 5.56 | 5.10 | 16.11 | 16.53 | 7.57 | 8.15 | 9.12 | 8.64 | 11.61 | 5.33 | 16.32 | 7.86 | 8.88 |
| BEND3 | 1 | 8.81 | 10.74 | 9.03 | 7.37 | 7.05 | 8.60 | 13.14 | 14.06 | 10.92 | 10.29 | 9.77 | 8.20 | 7.83 | 13.60 | 10.60 |
| BET1 | 2 | 11.14 | 6.92 | 10.64 | 6.25 | 7.79 | 6.41 | 15.42 | 9.90 | 11.47 | 14.04 | 9.03 | 8.45 | 7.10 | 12.66 | 12.76 |
| BICC1 | 1 | 8.61 | 9.90 | 14.30 | 14.66 | 6.05 | 5.10 | 9.75 | 10.17 | 11.35 | 10.11 | 9.25 | 14.48 | 5.58 | 9.96 | 10.73 |
| BICD1 | 5 | 9.79 | 9.83 | 11.25 | 10.01 | 9.77 | 8.72 | 10.01 | 8.74 | 11.58 | 10.30 | 9.81 | 10.63 | 9.25 | 9.38 | 10.94 |
| BICD2 | 22 | 8.97 | 10.68 | 11.09 | 10.57 | 9.00 | 9.68 | 8.44 | 7.90 | 12.03 | 11.65 | 9.82 | 10.83 | 9.34 | 8.17 | 11.84 |
| BID | 6 | 9.71 | 10.36 | 14.63 | 11.64 | 7.15 | 6.81 | 11.38 | 10.19 | 9.36 | 8.76 | 10.04 | 13.14 | 6.98 | 10.79 | 9.06 |
| BIN1 | 17 | 9.83 | 10.86 | 14.01 | 11.27 | 8.46 | 8.21 | 9.33 | 7.50 | 10.72 | 9.79 | 10.35 | 12.64 | 8.33 | 8.42 | 10.26 |
| BIN3 | 6 | 11.01 | 11.68 | 7.49 | 8.05 | 11.80 | 12.89 | 10.19 | 9.11 | 9.09 | 8.68 | 11.34 | 7.77 | 12.35 | 9.65 | 8.89 |
| BIRC2 | 7 | 9.24 | 10.13 | 11.53 | 10.84 | 10.42 | 10.83 | 7.92 | 8.71 | 10.26 | 10.11 | 9.69 | 11.19 | 10.63 | 8.31 | 10.19 |
| BIRC6 | 19 | 9.80 | 9.94 | 9.97 | 9.88 | 11.31 | 9.96 | 10.64 | 10.28 | 9.21 | 9.01 | 9.87 | 9.92 | 10.64 | 10.46 | 9.11 |
| BLM | 11 | 8.87 | 9.20 | 11.28 | 11.32 | 8.09 | 8.33 | 11.91 | 11.06 | 9.97 | 9.97 | 9.03 | 11.30 | 8.21 | 11.48 | 9.97 |
| BLMH | 24 | 10.23 | 10.54 | 12.81 | 10.27 | 9.18 | 8.65 | 8.12 | 11.68 | 9.57 | 8.94 | 10.39 | 11.54 | 8.92 | 9.90 | 9.26 |
| BLOC1S1 | 2 | 9.12 | 9.45 | 11.49 | 10.04 | 12.17 | 12.86 | 9.62 | 8.90 | 8.04 | 8.30 | 9.29 | 10.77 | 12.52 | 9.26 | 8.17 |
| BLOC1S2 | 6 | 11.16 | 11.91 | 9.31 | 8.80 | 11.94 | 11.87 | 9.55 | 7.27 | 9.55 | 8.65 | 11.53 | 9.05 | 11.90 | 8.41 | 9.10 |
| BLOC1S3 | 3 | 9.04 | 10.40 | 10.32 | 10.40 | 10.54 | 11.90 | 10.32 | 9.10 | 9.49 | 8.51 | 9.72 | 10.36 | 11.22 | 9.71 | 9.00 |
| BLVRA | 12 | 11.77 | 12.70 | 8.92 | 8.46 | 10.65 | 10.66 | 8.77 | 8.43 | 10.09 | 9.55 | 12.24 | 8.69 | 10.65 | 8.60 | 9.82 |
| BLVRB | 14 | 11.05 | 13.13 | 5.95 | 5.48 | 12.14 | 12.58 | 9.66 | 9.28 | 10.50 | 10.22 | 12.09 | 5.71 | 12.36 | 9.47 | 10.36 |
| BLZF1 | 1 | 8.99 | 12.52 | 10.00 | 10.71 | 9.23 | 12.22 | 7.64 | 7.37 | 8.91 | 12.41 | 10.75 | 10.35 | 10.72 | 7.50 | 10.66 |
| BMP1 | 3 | 9.63 | 6.52 | 9.46 | 10.56 | 6.06 | 6.41 | 16.72 | 15.35 | 10.08 | 9.20 | 8.08 | 10.01 | 6.23 | 16.04 | 9.64 |
| BMP2K | 1 | 7.35 | 8.64 | 11.77 | 10.63 | 9.10 | 10.13 | 9.29 | 8.67 | 13.62 | 10.80 | 7.99 | 11.20 | 9.62 | 8.98 | 12.21 |
| BMS1 | 19 | 10.26 | 10.21 | 10.61 | 10.92 | 8.59 | 8.44 | 12.07 | 10.89 | 8.77 | 9.26 | 10.23 | 10.76 | 8.51 | 11.48 | 9.01 |
| BNIP1 | 2 | 11.28 | 7.18 | 9.30 | 13.24 | 9.15 | 7.89 | 9.73 | 9.46 | 10.25 | 12.52 | 9.23 | 11.27 | 8.52 | 9.60 | 11.38 |

| | | | | | | | | | | | | | | | | |
|---|---|---|---|---|---|---|---|---|---|---|---|---|---|---|---|---|
| BOD1 | 5 | 8.62 | 10.53 | 10.03 | 11.39 | 8.40 | 9.75 | 11.73 | 11.10 | 8.80 | 9.65 | 9.57 | 10.71 | 9.07 | 11.41 | 9.23 |
| BOD1L1 | 36 | 9.61 | 10.83 | 9.65 | 9.03 | 11.44 | 11.70 | 10.02 | 9.01 | 9.45 | 9.26 | 10.22 | 9.34 | 11.57 | 9.52 | 9.36 |
| BOLA1 | 5 | 9.96 | 10.01 | 9.49 | 9.65 | 11.42 | 11.52 | 10.26 | 8.41 | 10.09 | 9.21 | 9.98 | 9.57 | 11.47 | 9.34 | 9.65 |
| BOLA2 | 8 | 8.22 | 8.73 | 12.58 | 11.17 | 8.87 | 9.82 | 10.28 | 13.78 | 8.77 | 7.78 | 8.48 | 11.87 | 9.35 | 12.03 | 8.27 |
| BOLA3 | 4 | 10.30 | 9.41 | 11.71 | 11.54 | 8.64 | 7.98 | 10.90 | 9.78 | 10.11 | 9.64 | 9.85 | 11.62 | 8.31 | 10.34 | 9.88 |
| BOP1 | 20 | 8.76 | 9.87 | 12.07 | 11.43 | 7.36 | 7.33 | 11.21 | 11.13 | 10.43 | 10.40 | 9.31 | 11.75 | 7.35 | 11.17 | 10.42 |
| BPHL | 2 | 12.25 | 11.29 | 5.82 | 6.70 | 12.76 | 12.21 | 10.66 | 9.89 | 9.47 | 8.95 | 11.77 | 6.26 | 12.48 | 10.27 | 9.21 |
| BPNT1 | 12 | 10.69 | 10.34 | 8.24 | 7.30 | 11.75 | 12.12 | 8.50 | 11.09 | 10.13 | 9.85 | 10.52 | 7.77 | 11.93 | 9.80 | 9.99 |
| BPTF | 30 | 9.90 | 10.59 | 10.88 | 10.49 | 9.44 | 9.60 | 9.79 | 9.57 | 10.19 | 9.55 | 10.24 | 10.69 | 9.52 | 9.68 | 9.87 |
| BRAF | 8 | 9.58 | 10.14 | 8.28 | 8.32 | 11.59 | 12.24 | 10.99 | 11.33 | 8.64 | 8.90 | 9.86 | 8.30 | 11.91 | 11.16 | 8.77 |
| BRAP | 4 | 8.73 | 10.46 | 10.60 | 10.69 | 9.97 | 11.25 | 9.45 | 8.53 | 10.26 | 10.06 | 9.60 | 10.64 | 10.61 | 8.99 | 10.16 |
| BRAT1 | 8 | 9.22 | 10.54 | 9.34 | 9.72 | 9.36 | 9.21 | 10.04 | 10.72 | 10.85 | 11.00 | 9.88 | 9.53 | 9.29 | 10.38 | 10.93 |
| BRCA1 | 3 | 10.05 | 11.30 | 11.69 | 12.15 | 8.29 | 8.89 | 8.60 | 8.36 | 10.66 | 9.99 | 10.68 | 11.92 | 8.59 | 8.48 | 10.33 |
| BRCC3 | 8 | 11.52 | 11.62 | 9.34 | 9.69 | 10.46 | 12.14 | 9.83 | 9.85 | 8.04 | 7.51 | 11.57 | 9.52 | 11.30 | 9.84 | 7.78 |
| BRD1 | 3 | 8.05 | 10.50 | 12.72 | 12.12 | 8.23 | 8.75 | 10.70 | 9.65 | 9.57 | 9.71 | 9.28 | 12.42 | 8.49 | 10.17 | 9.64 |
| BRD2 | 11 | 9.17 | 10.46 | 9.64 | 8.74 | 9.71 | 10.10 | 12.75 | 11.70 | 8.91 | 8.81 | 9.81 | 9.19 | 9.90 | 12.23 | 8.86 |
| BRD3 | 14 | 8.83 | 9.26 | 7.62 | 7.29 | 12.56 | 13.10 | 12.94 | 11.93 | 8.47 | 8.01 | 9.04 | 7.46 | 12.83 | 12.44 | 8.24 |
| BRD4 | 25 | 8.61 | 9.27 | 10.21 | 9.67 | 9.14 | 9.29 | 12.73 | 11.60 | 9.93 | 9.55 | 8.94 | 9.94 | 9.22 | 12.17 | 9.74 |
| BRD7 | 2 | 10.75 | 11.91 | 7.83 | 7.66 | 11.52 | 12.75 | 11.52 | 10.88 | 8.00 | 7.18 | 11.33 | 7.75 | 12.13 | 11.20 | 7.59 |
| BRD8 | 7 | 10.16 | 10.21 | 10.26 | 9.60 | 10.19 | 10.81 | 11.46 | 10.66 | 8.76 | 7.88 | 10.19 | 9.93 | 10.50 | 11.06 | 8.32 |
| BRD9 | 3 | 8.44 | 9.90 | 9.72 | 9.26 | 8.99 | 9.93 | 13.74 | 12.65 | 9.15 | 8.24 | 9.17 | 9.49 | 9.46 | 13.19 | 8.70 |
| BRE | 3 | 8.43 | 11.56 | 9.81 | 11.21 | 12.18 | 13.13 | 9.82 | 10.49 | 6.50 | 6.87 | 9.99 | 10.51 | 12.66 | 10.15 | 6.68 |
| BRF1 | 1 | 10.80 | 11.29 | 7.61 | 9.31 | 11.52 | 12.49 | 9.92 | 9.70 | 8.61 | 8.75 | 11.05 | 8.46 | 12.01 | 9.81 | 8.68 |
| BRF2 | 1 | 10.59 | 10.50 | 10.36 | 10.49 | 9.86 | 8.97 | 10.07 | 11.24 | 8.20 | 9.73 | 10.55 | 10.42 | 9.41 | 10.66 | 8.97 |
| BRI3BP | 2 | 12.46 | 7.34 | 8.47 | 8.40 | 11.49 | 8.63 | 10.82 | 9.50 | 9.98 | 12.89 | 9.90 | 8.44 | 10.06 | 10.16 | 11.44 |
| BRIP1 | 2 | 10.84 | 11.77 | 12.67 | 12.01 | 8.14 | 8.33 | 7.89 | 6.49 | 11.29 | 10.57 | 11.30 | 12.34 | 8.24 | 7.19 | 10.93 |
| BRIX1 | 19 | 8.47 | 8.82 | 8.17 | 8.08 | 7.42 | 7.18 | 13.04 | 21.15 | 8.57 | 9.09 | 8.65 | 8.13 | 7.30 | 17.09 | 8.83 |
| BRK1 | 3 | 10.66 | 9.59 | 7.93 | 11.20 | 12.17 | 12.15 | 7.61 | 4.71 | 12.02 | 11.95 | 10.12 | 9.57 | 12.16 | 6.16 | 11.99 |
| BRMS1 | 1 | 9.86 | 10.05 | 10.81 | 10.14 | 8.66 | 12.46 | 12.40 | 8.88 | 8.98 | 7.77 | 9.96 | 10.47 | 10.56 | 10.64 | 8.37 |
| BROX | 13 | 9.14 | 9.23 | 6.73 | 6.26 | 16.46 | 16.40 | 9.07 | 9.79 | 8.77 | 8.16 | 9.18 | 6.50 | 16.43 | 9.43 | 8.46 |
| BRPF1 | 7 | 9.41 | 10.21 | 10.34 | 9.79 | 8.74 | 8.92 | 11.50 | 11.03 | 9.87 | 10.19 | 9.81 | 10.07 | 8.83 | 11.27 | 10.03 |
| BSDC1 | 5 | 9.87 | 12.04 | 10.58 | 9.13 | 9.91 | 10.60 | 9.19 | 7.27 | 11.03 | 10.38 | 10.96 | 9.86 | 10.25 | 8.23 | 10.70 |
| BSG | 17 | 12.42 | 9.33 | 8.27 | 8.13 | 9.04 | 8.61 | 11.68 | 9.15 | 11.67 | 11.70 | 10.88 | 8.20 | 8.82 | 10.42 | 11.69 |
| BST2 | 10 | 12.28 | 8.13 | 9.08 | 9.48 | 9.59 | 8.81 | 12.33 | 10.33 | 10.37 | 9.59 | 10.20 | 9.28 | 9.20 | 11.33 | 9.98 |
| BTAF1 | 12 | 9.53 | 9.74 | 9.16 | 9.78 | 9.87 | 9.98 | 10.72 | 10.44 | 10.16 | 10.60 | 9.64 | 9.47 | 9.93 | 10.58 | 10.38 |
| BTBD3 | 1 | 9.06 | 10.25 | 9.39 | 8.72 | 12.25 | 11.88 | 9.72 | 10.22 | 9.36 | 9.15 | 9.66 | 9.05 | 12.07 | 9.97 | 9.25 |
| BTF3 | 17 | 8.27 | 10.37 | 12.51 | 11.08 | 9.19 | 9.18 | 10.66 | 9.99 | 9.22 | 9.53 | 9.32 | 11.80 | 9.18 | 10.32 | 9.37 |
| BTF3L4 | 10 | 9.27 | 10.74 | 8.65 | 7.94 | 9.71 | 8.95 | 11.67 | 10.49 | 11.38 | 11.20 | 10.01 | 8.29 | 9.33 | 11.08 | 11.29 |
| BUB1 | 4 | 8.94 | 8.83 | 11.53 | 10.23 | 9.38 | 9.76 | 11.16 | 10.71 | 9.54 | 9.91 | 8.89 | 10.88 | 9.57 | 10.94 | 9.73 |
| BUB1B | 29 | 9.26 | 9.79 | 11.02 | 11.01 | 9.39 | 9.68 | 9.76 | 9.33 | 10.48 | 10.28 | 9.52 | 11.01 | 9.54 | 9.54 | 10.38 |
| BUB3 | 23 | 9.05 | 10.71 | 10.48 | 9.76 | 10.17 | 10.15 | 9.85 | 10.61 | 9.81 | 9.43 | 9.88 | 10.12 | 10.16 | 10.23 | 9.62 |
| BUD13 | 7 | 9.09 | 11.21 | 10.23 | 10.80 | 9.89 | 11.29 | 9.60 | 9.07 | 9.35 | 9.48 | 10.15 | 10.51 | 10.59 | 9.33 | 9.41 |

| | | | | | | | | | | | | | | | | |
|---|---|---|---|---|---|---|---|---|---|---|---|---|---|---|---|---|
| BUD31 | 5 | 9.23 | 9.37 | 9.64 | 8.84 | 9.05 | 8.56 | 9.33 | 17.99 | 8.88 | 9.14 | 9.30 | 9.24 | 8.80 | 13.66 | 9.01 |
| BYSL | 14 | 9.93 | 9.70 | 10.30 | 10.29 | 8.07 | 8.53 | 11.88 | 11.00 | 10.37 | 9.94 | 9.81 | 10.29 | 8.30 | 11.44 | 10.15 |
| BZW1 | 38 | 8.39 | 9.05 | 7.43 | 7.13 | 13.33 | 13.19 | 12.60 | 12.70 | 8.25 | 7.94 | 8.72 | 7.28 | 13.26 | 12.65 | 8.09 |
| BZW2 | 33 | 9.48 | 10.42 | 11.78 | 11.35 | 7.18 | 7.18 | 9.35 | 9.54 | 11.87 | 11.84 | 9.95 | 11.56 | 7.18 | 9.45 | 11.86 |
| C10orf118 | 1 | 10.92 | 10.07 | 9.66 | 12.29 | 10.57 | 11.34 | 9.61 | 7.82 | 8.50 | 9.21 | 10.50 | 10.98 | 10.95 | 8.72 | 8.86 |
| C10orf32 | 3 | 10.11 | 11.12 | 9.98 | 9.01 | 12.17 | 12.02 | 9.77 | 8.06 | 9.60 | 8.16 | 10.62 | 9.49 | 12.09 | 8.91 | 8.88 |
| C10orf67 | 1 | 9.07 | 9.07 | 11.23 | 8.78 | 12.30 | 10.61 | 8.28 | 8.70 | 12.46 | 9.50 | 9.07 | 10.01 | 11.45 | 8.49 | 10.98 |
| C10orf76 | 2 | 9.86 | 11.99 | 10.28 | 7.89 | 11.49 | 9.84 | 9.08 | 9.26 | 9.98 | 10.33 | 10.93 | 9.09 | 10.66 | 9.17 | 10.16 |
| C11orf48 | 3 | 9.63 | 10.48 | 10.34 | 10.32 | 8.93 | 9.59 | 11.54 | 11.30 | 8.60 | 9.26 | 10.05 | 10.33 | 9.26 | 11.42 | 8.93 |
| C11orf49 | 1 | 10.39 | 12.76 | 7.96 | 12.42 | 11.09 | 13.05 | 7.31 | 10.65 | 7.22 | 7.16 | 11.57 | 10.19 | 12.07 | 8.98 | 7.19 |
| C11orf54 | 4 | 11.45 | 10.53 | 7.40 | 7.32 | 9.48 | 9.76 | 8.49 | 16.08 | 9.80 | 9.69 | 10.99 | 7.36 | 9.62 | 12.29 | 9.75 |
| C11orf57 | 1 | 10.61 | 13.47 | 8.07 | 9.40 | 8.24 | 10.39 | 10.67 | 9.71 | 12.17 | 7.25 | 12.04 | 8.74 | 9.32 | 10.19 | 9.71 |
| C11orf68 | 5 | 8.74 | 8.92 | 17.50 | 11.35 | 7.69 | 7.02 | 7.74 | 6.10 | 12.64 | 12.29 | 8.83 | 14.43 | 7.36 | 6.92 | 12.47 |
| C11orf73 | 3 | 8.62 | 10.08 | 10.38 | 10.45 | 9.42 | 11.50 | 9.92 | 8.54 | 10.91 | 10.18 | 9.35 | 10.41 | 10.46 | 9.23 | 10.55 |
| C12orf10 | 13 | 9.62 | 10.63 | 12.06 | 11.01 | 8.70 | 8.88 | 9.36 | 8.22 | 10.80 | 10.70 | 10.13 | 11.54 | 8.79 | 8.79 | 10.75 |
| C12orf23 | 20 | 9.66 | 10.46 | 10.10 | 9.61 | 10.96 | 10.99 | 10.08 | 9.45 | 9.51 | 9.19 | 10.06 | 9.85 | 10.97 | 9.76 | 9.35 |
| C12orf29 | 1 | 9.41 | 9.22 | 13.22 | 10.00 | 7.25 | 6.37 | 11.92 | 9.65 | 12.13 | 10.82 | 9.31 | 11.61 | 6.81 | 10.79 | 11.47 |
| C12orf43 | 7 | 9.13 | 10.75 | 11.96 | 11.82 | 9.64 | 9.77 | 9.65 | 7.13 | 10.11 | 10.03 | 9.94 | 11.89 | 9.70 | 8.39 | 10.07 |
| C12orf45 | 7 | 9.23 | 10.57 | 12.24 | 11.74 | 8.00 | 8.22 | 10.31 | 8.84 | 10.56 | 10.28 | 9.90 | 11.99 | 8.11 | 9.58 | 10.42 |
| C12orf57 | 2 | 9.92 | 10.61 | 7.91 | 8.39 | 12.07 | 13.35 | 9.66 | 4.99 | 11.61 | 11.49 | 10.26 | 8.15 | 12.71 | 7.33 | 11.55 |
| C14orf119 | 4 | 8.81 | 10.53 | 12.15 | 12.79 | 9.22 | 9.80 | 11.20 | 5.11 | 10.23 | 10.17 | 9.67 | 12.47 | 9.51 | 8.15 | 10.20 |
| C14orf142 | 2 | 9.97 | 11.55 | 10.14 | 9.25 | 12.14 | 11.56 | 10.85 | 3.59 | 11.61 | 9.34 | 10.76 | 9.70 | 11.85 | 7.22 | 10.47 |
| C14orf166 | 23 | 9.29 | 10.06 | 12.47 | 11.24 | 8.35 | 8.52 | 10.07 | 9.91 | 10.34 | 9.74 | 9.68 | 11.85 | 8.44 | 9.99 | 10.04 |
| C15orf39 | 2 | 7.85 | 8.25 | 11.20 | 10.56 | 9.35 | 9.53 | 11.10 | 10.90 | 10.58 | 10.69 | 8.05 | 10.88 | 9.44 | 11.00 | 10.64 |
| C15orf40 | 4 | 8.72 | 10.33 | 12.08 | 13.07 | 7.64 | 8.28 | 9.55 | 9.69 | 10.29 | 10.36 | 9.52 | 12.58 | 7.96 | 9.62 | 10.33 |
| C15orf52 | 2 | 7.43 | 8.91 | 16.72 | 16.61 | 5.33 | 6.63 | 5.10 | 4.55 | 14.50 | 14.23 | 8.17 | 16.66 | 5.98 | 4.82 | 14.36 |
| C15orf57 | 2 | 8.95 | 11.28 | 11.67 | 10.34 | 8.97 | 9.69 | 9.07 | 4.81 | 12.99 | 12.24 | 10.12 | 11.01 | 9.33 | 6.94 | 12.61 |
| C16orf13 | 1 | 8.76 | 10.89 | 7.25 | 9.12 | 9.73 | 13.81 | 9.63 | 9.66 | 12.81 | 8.34 | 9.83 | 8.18 | 11.77 | 9.64 | 10.58 |
| C16orf58 | 2 | 14.65 | 9.37 | 5.45 | 8.70 | 8.65 | 8.65 | 8.84 | 7.66 | 12.70 | 15.34 | 12.01 | 7.07 | 8.65 | 8.25 | 14.02 |
| C16orf62 | 8 | 10.60 | 9.67 | 9.15 | 10.04 | 12.70 | 13.06 | 9.48 | 9.38 | 8.24 | 7.69 | 10.13 | 9.59 | 12.88 | 9.43 | 7.96 |
| C16orf80 | 3 | 10.18 | 11.62 | 8.60 | 7.20 | 10.81 | 11.08 | 10.85 | 9.45 | 9.98 | 10.23 | 10.90 | 7.90 | 10.94 | 10.15 | 10.10 |
| C16orf87 | 3 | 12.60 | 12.02 | 7.64 | 7.57 | 8.77 | 8.96 | 10.09 | 9.13 | 11.00 | 12.21 | 12.31 | 7.61 | 8.87 | 9.61 | 11.61 |
| C17orf59 | 1 | 7.84 | 11.17 | 12.29 | 9.50 | 10.40 | 8.10 | 10.23 | 9.60 | 10.06 | 10.82 | 9.50 | 10.89 | 9.25 | 9.91 | 10.44 |
| C17orf62 | 1 | 12.79 | 7.06 | 6.02 | 11.12 | 10.07 | 10.01 | 9.92 | 9.00 | 10.92 | 13.09 | 9.93 | 8.57 | 10.04 | 9.46 | 12.00 |
| C17orf75 | 2 | 10.10 | 10.61 | 10.53 | 11.11 | 9.33 | 10.48 | 10.43 | 10.29 | 8.79 | 8.33 | 10.36 | 10.82 | 9.90 | 10.36 | 8.56 |
| C17orf85 | 12 | 9.92 | 11.47 | 10.61 | 9.96 | 9.67 | 9.37 | 10.43 | 8.50 | 10.27 | 9.81 | 10.69 | 10.28 | 9.52 | 9.46 | 10.04 |
| C18orf25 | 5 | 9.41 | 10.54 | 12.54 | 11.40 | 10.37 | 10.78 | 8.02 | 6.56 | 10.94 | 9.43 | 9.98 | 11.97 | 10.57 | 7.29 | 10.19 |
| C18orf32 | 1 | 12.25 | 6.89 | 9.93 | 13.74 | 12.00 | 10.99 | 6.19 | 6.13 | 9.45 | 12.44 | 9.57 | 11.83 | 11.49 | 6.16 | 10.95 |
| C18orf8 | 2 | 10.25 | 10.83 | 10.03 | 11.30 | 10.59 | 11.15 | 6.97 | 7.56 | 10.99 | 10.33 | 10.54 | 10.66 | 10.87 | 7.26 | 10.66 |
| C19orf10 | 14 | 9.27 | 9.69 | 10.07 | 9.51 | 13.22 | 12.45 | 10.10 | 9.09 | 8.36 | 8.23 | 9.48 | 9.79 | 12.84 | 9.60 | 8.30 |
| C19orf43 | 5 | 9.76 | 10.25 | 10.83 | 9.64 | 11.74 | 11.63 | 9.24 | 7.36 | 10.32 | 9.23 | 10.01 | 10.24 | 11.68 | 8.30 | 9.77 |
| C19orf52 | 2 | 9.65 | 9.16 | 11.63 | 9.98 | 10.15 | 9.53 | 10.18 | 10.40 | 9.24 | 10.07 | 9.40 | 10.81 | 9.84 | 10.29 | 9.66 |

| | | | | | | | | | | | | | | | | |
|---|---|---|---|---|---|---|---|---|---|---|---|---|---|---|---|---|
| C19orf53 | 7 | 9.21 | 9.09 | 9.44 | 11.80 | 9.95 | 9.72 | 8.77 | 8.67 | 9.81 | 13.54 | 9.15 | 10.62 | 9.83 | 8.72 | 11.67 |
| C19orf66 | 1 | 10.04 | 11.07 | 7.16 | 7.06 | 13.67 | 14.00 | 11.87 | 10.64 | 7.20 | 7.28 | 10.56 | 7.11 | 13.84 | 11.26 | 7.24 |
| C1D | 2 | 10.37 | 12.54 | 8.07 | 8.40 | 10.77 | 11.60 | 9.49 | 9.42 | 9.98 | 9.37 | 11.45 | 8.24 | 11.18 | 9.46 | 9.67 |
| C1orf109 | 1 | 8.41 | 10.65 | 9.38 | 9.62 | 9.44 | 11.23 | 10.09 | 9.51 | 10.78 | 10.90 | 9.53 | 9.50 | 10.34 | 9.80 | 10.84 |
| C1orf112 | 2 | 10.91 | 10.31 | 9.72 | 9.26 | 10.83 | 9.04 | 11.07 | 10.05 | 9.41 | 9.39 | 10.61 | 9.49 | 9.94 | 10.56 | 9.40 |
| C1orf122 | 2 | 10.12 | 9.26 | 8.86 | 8.47 | 10.21 | 10.92 | 10.56 | 11.62 | 10.27 | 9.70 | 9.69 | 8.66 | 10.57 | 11.09 | 9.99 |
| C1orf123 | 7 | 10.39 | 11.84 | 6.86 | 6.47 | 11.04 | 10.81 | 9.88 | 12.90 | 10.12 | 9.69 | 11.11 | 6.67 | 10.92 | 11.39 | 9.91 |
| C1orf131 | 4 | 9.81 | 10.72 | 8.26 | 7.63 | 11.01 | 11.41 | 11.22 | 10.83 | 9.54 | 9.58 | 10.26 | 7.95 | 11.21 | 11.02 | 9.56 |
| C1orf174 | 1 | 9.70 | 12.07 | 9.72 | 8.00 | 7.60 | 8.10 | 12.63 | 12.35 | 10.43 | 9.40 | 10.89 | 8.86 | 7.85 | 12.49 | 9.92 |
| C1orf198 | 9 | 14.01 | 16.62 | 14.26 | 13.66 | 6.24 | 6.37 | 3.87 | 3.69 | 10.92 | 10.36 | 15.32 | 13.96 | 6.30 | 3.78 | 10.64 |
| C1orf50 | 2 | 11.93 | 10.37 | 8.69 | 8.53 | 10.77 | 10.96 | 8.26 | 6.74 | 11.43 | 12.31 | 11.15 | 8.61 | 10.87 | 7.50 | 11.87 |
| C1orf52 | 7 | 10.31 | 11.89 | 11.37 | 11.79 | 7.89 | 9.41 | 11.37 | 4.18 | 11.12 | 10.66 | 11.10 | 11.58 | 8.65 | 7.78 | 10.89 |
| C1orf85 | 1 | 9.04 | 8.92 | 9.25 | 8.68 | 16.31 | 15.05 | 6.71 | 8.81 | 9.15 | 8.08 | 8.98 | 8.97 | 15.68 | 7.76 | 8.61 |
| C1QBP | 24 | 10.56 | 12.42 | 10.86 | 12.58 | 7.63 | 7.16 | 10.36 | 8.43 | 9.85 | 10.16 | 11.49 | 11.72 | 7.40 | 9.39 | 10.01 |
| C20orf24 | 1 | 10.08 | 5.17 | 5.85 | 8.86 | 10.95 | 7.26 | 13.19 | 10.64 | 12.27 | 15.73 | 7.62 | 7.35 | 9.11 | 11.92 | 14.00 |
| C20orf27 | 1 | 8.61 | 10.48 | 11.24 | 8.65 | 9.60 | 8.15 | 11.69 | 11.31 | 10.71 | 9.56 | 9.55 | 9.94 | 8.87 | 11.50 | 10.13 |
| C21orf33 | 7 | 12.48 | 10.56 | 6.87 | 7.15 | 12.75 | 11.68 | 9.56 | 8.33 | 10.24 | 10.38 | 11.52 | 7.01 | 12.21 | 8.94 | 10.31 |
| C21orf59 | 7 | 10.19 | 11.54 | 8.42 | 8.20 | 10.89 | 11.46 | 10.87 | 10.07 | 9.38 | 8.99 | 10.86 | 8.31 | 11.18 | 10.47 | 9.18 |
| C2CD5 | 1 | 8.98 | 11.09 | 13.67 | 8.38 | 10.12 | 10.11 | 13.49 | 9.23 | 7.44 | 7.50 | 10.03 | 11.02 | 10.11 | 11.36 | 7.47 |
| C2orf43 | 4 | 11.16 | 9.92 | 7.00 | 7.09 | 10.65 | 11.55 | 10.12 | 10.49 | 11.68 | 10.34 | 10.54 | 7.05 | 11.10 | 10.30 | 11.01 |
| C2orf47 | 4 | 11.53 | 9.47 | 9.41 | 11.09 | 9.43 | 8.94 | 9.85 | 10.09 | 9.34 | 10.85 | 10.50 | 10.25 | 9.18 | 9.97 | 10.09 |
| C2orf49 | 6 | 9.59 | 10.68 | 9.78 | 8.62 | 10.45 | 10.37 | 10.72 | 10.22 | 9.61 | 9.96 | 10.14 | 9.20 | 10.41 | 10.47 | 9.79 |
| C2orf69 | 3 | 9.63 | 11.10 | 7.99 | 10.16 | 9.74 | 11.06 | 9.57 | 9.73 | 10.31 | 10.70 | 10.37 | 9.08 | 10.40 | 9.65 | 10.51 |
| C2orf76 | 1 | 9.38 | 9.86 | 7.85 | 7.32 | 11.03 | 15.14 | 8.01 | 13.33 | 9.99 | 8.09 | 9.62 | 7.59 | 13.09 | 10.67 | 9.04 |
| C3orf17 | 4 | 9.66 | 8.92 | 9.86 | 9.67 | 8.23 | 8.57 | 12.19 | 10.93 | 11.56 | 10.41 | 9.29 | 9.77 | 8.40 | 11.56 | 10.98 |
| C3orf38 | 5 | 9.71 | 10.83 | 8.53 | 12.10 | 9.74 | 10.60 | 10.17 | 10.72 | 8.75 | 8.86 | 10.27 | 10.31 | 10.17 | 10.45 | 8.80 |
| C4A | 1 | 7.95 | 8.00 | 15.26 | 14.04 | 7.97 | 8.31 | 8.62 | 8.98 | 11.01 | 9.87 | 7.97 | 14.65 | 8.14 | 8.80 | 10.44 |
| C4orf27 | 6 | 10.78 | 11.40 | 9.53 | 9.02 | 9.87 | 9.81 | 11.12 | 9.54 | 9.71 | 9.22 | 11.09 | 9.27 | 9.84 | 10.33 | 9.46 |
| C5orf22 | 3 | 9.66 | 10.80 | 11.47 | 10.86 | 10.27 | 9.64 | 8.67 | 8.19 | 10.70 | 9.73 | 10.23 | 11.17 | 9.96 | 8.43 | 10.22 |
| C5orf24 | 3 | 10.03 | 9.74 | 9.01 | 7.17 | 10.43 | 10.70 | 13.44 | 11.09 | 9.47 | 8.90 | 9.88 | 8.09 | 10.57 | 12.27 | 9.19 |
| C5orf34 | 1 | 10.34 | 11.57 | 8.10 | 12.89 | 8.14 | 8.62 | 9.44 | 10.50 | 9.62 | 10.76 | 10.96 | 10.49 | 8.38 | 9.97 | 10.19 |
| C5orf51 | 3 | 9.50 | 9.40 | 7.59 | 7.33 | 12.29 | 11.74 | 12.53 | 10.71 | 9.90 | 9.00 | 9.45 | 7.46 | 12.02 | 11.62 | 9.45 |
| C6orf106 | 1 | 10.68 | 11.25 | 9.61 | 9.53 | 11.78 | 11.19 | 8.99 | 8.16 | 10.24 | 8.58 | 10.96 | 9.57 | 11.48 | 8.57 | 9.41 |
| C6orf211 | 12 | 8.89 | 9.44 | 11.51 | 11.35 | 11.57 | 11.59 | 8.39 | 8.22 | 9.44 | 9.60 | 9.17 | 11.43 | 11.58 | 8.30 | 9.52 |
| C6orf57 | 1 | 14.85 | 11.52 | 8.83 | 7.39 | 12.38 | 11.29 | 13.52 | 6.95 | 6.65 | 6.63 | 13.18 | 8.11 | 11.84 | 10.24 | 6.64 |
| C7orf25 | 3 | 10.28 | 10.40 | 11.37 | 10.94 | 9.18 | 8.87 | 9.26 | 8.84 | 10.29 | 10.58 | 10.34 | 11.15 | 9.02 | 9.05 | 10.44 |
| C7orf50 | 9 | 10.27 | 10.71 | 11.20 | 10.99 | 7.37 | 7.34 | 9.14 | 12.45 | 10.31 | 10.22 | 10.49 | 11.10 | 7.35 | 10.79 | 10.27 |
| C7orf55 | 2 | 10.19 | 8.80 | 13.81 | 14.09 | 8.87 | 9.02 | 8.74 | 7.58 | 9.69 | 9.21 | 9.50 | 13.95 | 8.95 | 8.16 | 9.45 |
| C8orf33 | 5 | 9.94 | 10.17 | 9.98 | 8.87 | 10.96 | 11.14 | 9.38 | 9.39 | 9.95 | 10.22 | 10.05 | 9.43 | 11.05 | 9.39 | 10.08 |
| C8orf59 | 5 | 9.58 | 11.36 | 9.32 | 7.87 | 9.81 | 9.79 | 11.30 | 9.92 | 10.11 | 10.95 | 10.47 | 8.59 | 9.80 | 10.61 | 10.53 |
| C9orf114 | 6 | 9.52 | 10.67 | 12.01 | 12.18 | 8.47 | 8.87 | 9.47 | 8.83 | 10.18 | 9.80 | 10.09 | 12.10 | 8.67 | 9.15 | 9.99 |
| C9orf142 | 10 | 9.20 | 10.72 | 6.11 | 6.35 | 13.80 | 14.53 | 8.28 | 7.57 | 11.85 | 11.60 | 9.96 | 6.23 | 14.16 | 7.93 | 11.72 |

| | | | | | | | | | | | | | | | | |
|---|---|---|---|---|---|---|---|---|---|---|---|---|---|---|---|---|
| C9orf16 | 2 | 9.31 | 12.14 | 9.57 | 9.75 | 10.17 | 13.68 | 8.93 | 3.84 | 11.74 | 10.86 | 10.73 | 9.66 | 11.92 | 6.38 | 11.30 |
| C9orf40 | 3 | 10.96 | 12.10 | 7.63 | 7.33 | 11.25 | 11.47 | 9.70 | 5.56 | 12.54 | 11.45 | 11.53 | 7.48 | 11.36 | 7.63 | 11.99 |
| C9orf41 | 1 | 10.77 | 8.86 | 10.19 | 8.03 | 10.13 | 8.14 | 11.77 | 9.60 | 12.00 | 10.50 | 9.82 | 9.11 | 9.14 | 10.69 | 11.25 |
| C9orf64 | 14 | 9.11 | 9.82 | 10.10 | 9.20 | 9.62 | 9.02 | 7.36 | 11.49 | 12.20 | 12.07 | 9.46 | 9.65 | 9.32 | 9.42 | 12.13 |
| C9orf78 | 13 | 9.09 | 10.61 | 11.33 | 9.55 | 10.03 | 9.67 | 10.73 | 9.33 | 10.06 | 9.59 | 9.85 | 10.44 | 9.85 | 10.03 | 9.82 |
| C9orf89 | 1 | 8.40 | 9.95 | 9.23 | 13.42 | 10.22 | 9.73 | 9.96 | 8.61 | 10.82 | 9.65 | 9.18 | 11.33 | 9.98 | 9.29 | 10.23 |
| CA13 | 3 | 10.06 | 10.74 | 11.21 | 9.99 | 9.32 | 8.63 | 10.78 | 9.73 | 9.87 | 9.68 | 10.40 | 10.60 | 8.97 | 10.25 | 9.77 |
| CA2 | 13 | 6.82 | 8.10 | 5.53 | 4.42 | 13.84 | 13.69 | 19.05 | 16.67 | 6.92 | 4.96 | 7.46 | 4.98 | 13.76 | 17.86 | 5.94 |
| CAAP1 | 1 | 9.49 | 10.37 | 10.40 | 10.33 | 10.50 | 10.76 | 9.14 | 8.69 | 10.58 | 9.75 | 9.93 | 10.37 | 10.63 | 8.91 | 10.16 |
| CAB39 | 21 | 9.77 | 10.14 | 7.87 | 7.19 | 8.93 | 8.58 | 16.91 | 16.58 | 7.29 | 6.73 | 9.96 | 7.53 | 8.76 | 16.74 | 7.01 |
| CABIN1 | 1 | 7.19 | 9.77 | 8.38 | 10.36 | 7.55 | 6.69 | 9.99 | 21.39 | 9.70 | 8.97 | 8.48 | 9.37 | 7.12 | 15.69 | 9.34 |
| CABLES1 | 1 | 10.83 | 12.24 | 11.40 | 9.98 | 11.07 | 11.48 | 6.48 | 6.34 | 10.46 | 9.74 | 11.53 | 10.69 | 11.27 | 6.41 | 10.10 |
| CACNA2D1 | 13 | 11.55 | 9.15 | 11.05 | 13.08 | 8.52 | 8.93 | 8.94 | 7.79 | 10.22 | 10.77 | 10.35 | 12.07 | 8.73 | 8.36 | 10.49 |
| CACTIN | 9 | 8.77 | 9.47 | 11.64 | 11.11 | 9.19 | 9.11 | 10.81 | 10.05 | 10.08 | 9.78 | 9.12 | 11.38 | 9.15 | 10.43 | 9.93 |
| CACUL1 | 5 | 9.33 | 10.77 | 9.58 | 10.80 | 8.23 | 10.06 | 10.45 | 11.28 | 9.69 | 9.81 | 10.05 | 10.19 | 9.15 | 10.86 | 9.75 |
| CACYBP | 42 | 9.90 | 9.84 | 10.03 | 9.54 | 11.28 | 11.57 | 9.73 | 9.83 | 9.28 | 9.02 | 9.87 | 9.78 | 11.42 | 9.78 | 9.15 |
| CAD | 98 | 10.69 | 10.66 | 7.31 | 7.08 | 10.83 | 10.65 | 10.15 | 13.05 | 9.86 | 9.74 | 10.67 | 7.19 | 10.74 | 11.60 | 9.80 |
| CALCOCO1 | 2 | 9.77 | 10.35 | 10.33 | 9.23 | 11.04 | 10.33 | 8.85 | 10.40 | 9.05 | 10.66 | 10.06 | 9.78 | 10.68 | 9.63 | 9.86 |
| CALCOCO2 | 8 | 9.93 | 10.09 | 10.29 | 9.33 | 9.59 | 9.24 | 8.04 | 10.63 | 12.00 | 10.87 | 10.01 | 9.81 | 9.41 | 9.33 | 11.43 |
| CALD1 | 66 | 10.01 | 10.82 | 17.73 | 13.53 | 5.88 | 5.61 | 6.48 | 5.27 | 12.59 | 12.07 | 10.42 | 15.63 | 5.75 | 5.88 | 12.33 |
| CALM1 | 42 | 11.41 | 12.68 | 11.17 | 9.16 | 10.54 | 10.94 | 8.19 | 6.02 | 9.92 | 9.97 | 12.04 | 10.16 | 10.74 | 7.10 | 9.95 |
| CALR | 97 | 9.73 | 8.41 | 8.87 | 9.15 | 13.07 | 12.46 | 10.87 | 10.58 | 8.40 | 8.46 | 9.07 | 9.01 | 12.76 | 10.73 | 8.43 |
| CALU | 41 | 9.83 | 9.44 | 9.80 | 8.60 | 9.75 | 9.26 | 12.48 | 9.98 | 10.47 | 10.38 | 9.63 | 9.20 | 9.51 | 11.23 | 10.43 |
| CAMK1 | 4 | 11.39 | 11.85 | 11.68 | 11.77 | 9.12 | 9.51 | 6.55 | 5.87 | 11.50 | 10.76 | 11.62 | 11.73 | 9.32 | 6.21 | 11.13 |
| CAMK1D | 8 | 11.00 | 10.93 | 12.54 | 13.38 | 8.39 | 8.60 | 7.15 | 6.83 | 11.04 | 10.15 | 10.97 | 12.96 | 8.49 | 6.99 | 10.60 |
| CAMK2D | 8 | 9.30 | 11.23 | 11.82 | 10.28 | 11.88 | 12.40 | 6.36 | 5.88 | 10.56 | 10.28 | 10.26 | 11.05 | 12.14 | 6.12 | 10.42 |
| CAMK2G | 8 | 8.47 | 10.32 | 11.06 | 9.98 | 9.81 | 10.38 | 8.43 | 8.97 | 11.10 | 11.46 | 9.40 | 10.52 | 10.10 | 8.70 | 11.28 |
| CAMK4 | 2 | 9.14 | 10.92 | 13.94 | 13.97 | 10.61 | 11.35 | 4.68 | 3.67 | 10.64 | 11.09 | 10.03 | 13.96 | 10.98 | 4.18 | 10.86 |
| CAMKK1 | 8 | 12.97 | 14.23 | 8.86 | 8.56 | 8.09 | 8.43 | 8.07 | 7.70 | 11.67 | 11.42 | 13.60 | 8.71 | 8.26 | 7.88 | 11.54 |
| CAMKK2 | 2 | 9.98 | 10.91 | 9.95 | 9.55 | 7.99 | 8.17 | 13.47 | 12.64 | 8.55 | 8.80 | 10.44 | 9.75 | 8.08 | 13.05 | 8.68 |
| CAMLG | 2 | 10.93 | 8.41 | 7.70 | 8.02 | 10.97 | 9.62 | 10.94 | 8.97 | 11.71 | 12.73 | 9.67 | 7.86 | 10.30 | 9.96 | 12.22 |
| CAMSAP1 | 14 | 10.02 | 10.99 | 9.28 | 8.75 | 10.90 | 10.72 | 10.92 | 9.63 | 9.60 | 9.19 | 10.51 | 9.01 | 10.81 | 10.28 | 9.39 |
| CAMSAP2 | 22 | 8.90 | 10.04 | 8.84 | 8.33 | 12.72 | 12.83 | 10.21 | 9.57 | 9.45 | 9.12 | 9.47 | 8.58 | 12.78 | 9.89 | 9.29 |
| CAND1 | 54 | 8.92 | 8.49 | 9.30 | 8.78 | 11.15 | 10.80 | 9.67 | 14.61 | 9.31 | 8.98 | 8.70 | 9.04 | 10.97 | 12.14 | 9.14 |
| CAND2 | 6 | 10.16 | 11.12 | 9.88 | 11.57 | 9.25 | 9.92 | 8.75 | 8.77 | 10.74 | 9.85 | 10.64 | 10.72 | 9.58 | 8.76 | 10.30 |
| CANX | 64 | 12.53 | 6.43 | 6.93 | 10.33 | 10.28 | 8.60 | 11.15 | 9.97 | 10.37 | 13.41 | 9.48 | 8.63 | 9.44 | 10.56 | 11.89 |
| CAP1 | 58 | 8.99 | 9.65 | 13.07 | 12.05 | 7.50 | 7.41 | 6.76 | 11.47 | 11.68 | 11.43 | 9.32 | 12.56 | 7.45 | 9.11 | 11.55 |
| CAP2 | 9 | 12.06 | 14.11 | 11.99 | 12.10 | 8.20 | 8.87 | 5.37 | 4.98 | 11.10 | 11.22 | 13.08 | 12.05 | 8.54 | 5.18 | 11.16 |
| CAPG | 21 | 7.73 | 9.43 | 6.62 | 6.30 | 16.49 | 16.82 | 10.29 | 9.38 | 8.74 | 8.20 | 8.58 | 6.46 | 16.66 | 9.84 | 8.47 |
| CAPN1 | 31 | 10.69 | 11.50 | 11.21 | 11.05 | 12.67 | 12.70 | 8.46 | 8.21 | 7.01 | 6.50 | 11.09 | 11.13 | 12.68 | 8.34 | 6.75 |
| CAPN2 | 30 | 8.42 | 8.17 | 7.71 | 7.41 | 10.61 | 10.25 | 9.92 | 16.33 | 10.72 | 10.45 | 8.29 | 7.56 | 10.43 | 13.12 | 10.59 |
| CAPN7 | 6 | 10.70 | 10.52 | 8.51 | 8.89 | 10.40 | 10.37 | 10.50 | 9.53 | 10.64 | 9.94 | 10.61 | 8.70 | 10.39 | 10.02 | 10.29 |

| | | | | | | | | | | | | | | | | |
|---|---|---|---|---|---|---|---|---|---|---|---|---|---|---|---|---|
| CAPNS1 | 22 | 8.49 | 9.07 | 9.83 | 8.53 | 10.64 | 10.51 | 9.22 | 15.75 | 9.06 | 8.91 | 8.78 | 9.18 | 10.57 | 12.48 | 8.98 |
| CAPRIN1 | 34 | 9.28 | 11.17 | 10.69 | 9.59 | 10.04 | 10.42 | 9.40 | 10.65 | 9.28 | 9.48 | 10.22 | 10.14 | 10.23 | 10.02 | 9.38 |
| CAPZA1 | 34 | 10.39 | 10.86 | 10.97 | 9.93 | 9.05 | 8.97 | 8.85 | 7.96 | 11.73 | 11.29 | 10.62 | 10.45 | 9.01 | 8.40 | 11.51 |
| CAPZA2 | 16 | 9.61 | 10.19 | 12.63 | 10.95 | 8.88 | 9.14 | 7.96 | 8.88 | 11.18 | 10.59 | 9.90 | 11.79 | 9.01 | 8.42 | 10.88 |
| CAPZB | 32 | 9.94 | 10.93 | 10.87 | 10.72 | 8.53 | 9.21 | 8.06 | 9.51 | 11.30 | 10.93 | 10.44 | 10.79 | 8.87 | 8.78 | 11.12 |
| CARD6 | 7 | 8.72 | 8.93 | 10.76 | 10.33 | 12.35 | 12.83 | 10.09 | 10.73 | 7.96 | 7.30 | 8.82 | 10.55 | 12.59 | 10.41 | 7.63 |
| CARHSP1 | 2 | 9.29 | 9.75 | 10.50 | 9.63 | 12.79 | 12.06 | 9.14 | 7.68 | 9.87 | 9.28 | 9.52 | 10.07 | 12.43 | 8.41 | 9.58 |
| CARKD | 4 | 11.21 | 11.51 | 8.20 | 8.54 | 11.51 | 11.59 | 10.06 | 9.83 | 9.13 | 8.42 | 11.36 | 8.37 | 11.55 | 9.95 | 8.78 |
| CARM1 | 14 | 9.28 | 9.65 | 11.24 | 10.69 | 8.03 | 8.60 | 10.85 | 10.80 | 10.39 | 10.48 | 9.46 | 10.96 | 8.31 | 10.83 | 10.43 |
| CARS | 34 | 8.30 | 8.81 | 11.61 | 10.61 | 8.41 | 8.58 | 10.96 | 10.66 | 11.25 | 10.81 | 8.55 | 11.11 | 8.49 | 10.81 | 11.03 |
| CARS2 | 10 | 10.07 | 11.31 | 7.60 | 8.35 | 11.75 | 11.39 | 13.26 | 12.38 | 6.88 | 7.01 | 10.69 | 7.97 | 11.57 | 12.82 | 6.95 |
| CASC3 | 6 | 9.83 | 11.09 | 11.09 | 10.32 | 8.85 | 9.17 | 9.83 | 9.01 | 10.89 | 9.91 | 10.46 | 10.71 | 9.01 | 9.42 | 10.40 |
| CASC4 | 6 | 10.94 | 7.30 | 7.00 | 9.51 | 12.73 | 11.96 | 9.02 | 8.00 | 11.68 | 11.86 | 9.12 | 8.26 | 12.35 | 8.51 | 11.77 |
| CASC5 | 15 | 8.99 | 9.86 | 9.37 | 9.25 | 9.23 | 9.47 | 10.49 | 11.60 | 10.93 | 10.81 | 9.43 | 9.31 | 9.35 | 11.05 | 10.87 |
| CASK | 12 | 9.44 | 9.25 | 9.09 | 9.22 | 10.69 | 10.25 | 14.05 | 12.34 | 7.65 | 8.01 | 9.34 | 9.16 | 10.47 | 13.20 | 7.83 |
| CASKIN2 | 2 | 11.63 | 12.80 | 9.64 | 8.67 | 10.54 | 10.08 | 5.76 | 4.95 | 13.29 | 12.63 | 12.21 | 9.16 | 10.31 | 5.35 | 12.96 |
| CASP1 | 1 | 11.19 | 14.58 | 4.14 | 3.57 | 11.91 | 13.82 | 15.05 | 15.63 | 4.94 | 5.17 | 12.89 | 3.85 | 12.86 | 15.34 | 5.06 |
| CASP2 | 1 | 13.75 | 13.62 | 8.57 | 8.65 | 9.36 | 9.08 | 9.96 | 8.47 | 9.20 | 9.34 | 13.69 | 8.61 | 9.22 | 9.21 | 9.27 |
| CASP3 | 4 | 9.09 | 10.21 | 10.25 | 10.63 | 10.67 | 10.97 | 9.43 | 8.94 | 9.71 | 10.10 | 9.65 | 10.44 | 10.82 | 9.18 | 9.90 |
| CASP4 | 3 | 7.23 | 8.72 | 11.92 | 11.34 | 7.10 | 7.53 | 14.79 | 15.54 | 8.07 | 7.75 | 7.97 | 11.63 | 7.32 | 15.17 | 7.91 |
| CASP6 | 3 | 9.94 | 10.36 | 8.77 | 9.05 | 11.53 | 12.45 | 10.06 | 9.34 | 9.16 | 9.33 | 10.15 | 8.91 | 11.99 | 9.70 | 9.25 |
| CASP7 | 4 | 10.73 | 10.39 | 9.42 | 8.24 | 12.38 | 11.76 | 9.34 | 10.28 | 9.23 | 8.23 | 10.56 | 8.83 | 12.07 | 9.81 | 8.73 |
| CASP8 | 11 | 10.61 | 10.83 | 12.41 | 9.91 | 7.94 | 7.72 | 9.70 | 9.55 | 11.07 | 10.27 | 10.72 | 11.16 | 7.83 | 9.63 | 10.67 |
| CASP9 | 1 | 9.15 | 10.06 | 6.18 | 5.33 | 16.06 | 17.71 | 12.37 | 11.81 | 5.34 | 5.99 | 9.61 | 5.76 | 16.88 | 12.09 | 5.66 |
| CAST | 68 | 9.24 | 9.93 | 8.22 | 7.62 | 11.18 | 11.65 | 11.58 | 11.16 | 10.04 | 9.38 | 9.58 | 7.92 | 11.41 | 11.37 | 9.71 |
| CAT | 29 | 9.34 | 11.31 | 8.41 | 7.67 | 12.85 | 13.33 | 9.22 | 8.49 | 9.76 | 9.62 | 10.33 | 8.04 | 13.09 | 8.85 | 9.69 |
| CAV1 | 7 | 9.98 | 8.14 | 11.97 | 17.81 | 3.33 | 3.68 | 4.79 | 4.99 | 15.70 | 19.61 | 9.06 | 14.89 | 3.50 | 4.89 | 17.66 |
| CAV2 | 1 | 13.54 | 8.92 | 13.06 | 14.68 | 4.96 | 4.37 | 5.15 | 4.60 | 14.09 | 16.62 | 11.23 | 13.87 | 4.67 | 4.88 | 15.35 |
| CBFB | 3 | 12.00 | 11.40 | 7.34 | 7.04 | 10.65 | 10.74 | 10.38 | 9.57 | 11.05 | 9.83 | 11.70 | 7.19 | 10.69 | 9.97 | 10.44 |
| CBL | 5 | 9.25 | 10.18 | 11.20 | 10.83 | 9.64 | 10.07 | 9.05 | 8.29 | 11.07 | 10.42 | 9.72 | 11.02 | 9.85 | 8.67 | 10.74 |
| CBLB | 2 | 9.65 | 10.15 | 12.23 | 12.48 | 6.80 | 7.64 | 9.05 | 8.60 | 11.62 | 11.77 | 9.90 | 12.36 | 7.22 | 8.82 | 11.70 |
| CBLL1 | 5 | 9.96 | 11.52 | 11.09 | 10.15 | 9.47 | 10.60 | 9.25 | 8.61 | 9.63 | 9.71 | 10.74 | 10.62 | 10.04 | 8.93 | 9.67 |
| CBR1 | 36 | 8.91 | 10.72 | 9.73 | 8.67 | 11.31 | 11.04 | 8.83 | 11.35 | 9.74 | 9.69 | 9.81 | 9.20 | 11.18 | 10.09 | 9.72 |
| CBR3 | 5 | 11.75 | 13.74 | 8.29 | 7.87 | 14.83 | 13.92 | 5.15 | 4.38 | 10.25 | 9.81 | 12.75 | 8.08 | 14.37 | 4.77 | 10.03 |
| CBR4 | 3 | 11.12 | 9.78 | 9.42 | 9.56 | 9.19 | 12.93 | 9.49 | 9.04 | 9.40 | 10.07 | 10.45 | 9.49 | 11.06 | 9.27 | 9.73 |
| CBS | 17 | 10.24 | 10.68 | 7.96 | 8.26 | 7.29 | 7.90 | 8.33 | 10.63 | 14.72 | 14.00 | 10.46 | 8.11 | 7.59 | 9.48 | 14.36 |
| CBWD1 | 5 | 9.19 | 9.66 | 9.97 | 9.20 | 10.83 | 10.95 | 11.97 | 10.75 | 9.13 | 8.36 | 9.43 | 9.58 | 10.89 | 11.36 | 8.75 |
| CBX1 | 14 | 10.04 | 11.05 | 13.56 | 10.69 | 9.35 | 9.04 | 8.86 | 7.13 | 10.30 | 9.97 | 10.55 | 12.13 | 9.20 | 7.99 | 10.14 |
| CBX2 | 2 | 8.52 | 9.62 | 9.84 | 9.39 | 8.21 | 7.65 | 15.04 | 13.67 | 9.10 | 8.96 | 9.07 | 9.62 | 7.93 | 14.36 | 9.03 |
| CBX3 | 11 | 8.95 | 9.77 | 9.73 | 8.95 | 8.70 | 8.96 | 11.70 | 16.57 | 8.61 | 8.07 | 9.36 | 9.34 | 8.83 | 14.13 | 8.34 |
| CBX4 | 3 | 9.57 | 10.28 | 12.15 | 10.52 | 8.61 | 8.79 | 10.90 | 9.39 | 10.21 | 9.58 | 9.92 | 11.34 | 8.70 | 10.15 | 9.90 |
| CBX5 | 14 | 7.81 | 9.16 | 8.07 | 7.43 | 10.32 | 10.80 | 10.61 | 21.84 | 7.13 | 6.82 | 8.49 | 7.75 | 10.56 | 16.23 | 6.98 |

| | | | | | | | | | | | | | | | | |
|---|---|---|---|---|---|---|---|---|---|---|---|---|---|---|---|---|
| CBX8 | 8 | 8.15 | 10.10 | 12.16 | 11.60 | 9.16 | 9.56 | 11.14 | 10.09 | 8.96 | 9.08 | 9.13 | 11.88 | 9.36 | 10.62 | 9.02 |
| CC2D1A | 25 | 10.23 | 11.81 | 8.82 | 8.53 | 11.18 | 11.77 | 10.16 | 9.73 | 9.16 | 8.61 | 11.02 | 8.67 | 11.47 | 9.95 | 8.88 |
| CC2D1B | 14 | 10.65 | 11.32 | 8.78 | 8.45 | 9.16 | 8.66 | 10.87 | 9.54 | 11.58 | 10.99 | 10.99 | 8.61 | 8.91 | 10.20 | 11.29 |
| CCAR1 | 28 | 9.62 | 10.20 | 9.40 | 8.86 | 10.73 | 10.98 | 11.18 | 10.07 | 9.84 | 9.14 | 9.91 | 9.13 | 10.85 | 10.62 | 9.49 |
| CCAR2 | 46 | 8.86 | 9.66 | 10.19 | 9.78 | 9.36 | 9.62 | 12.03 | 11.64 | 9.48 | 9.39 | 9.26 | 9.98 | 9.49 | 11.84 | 9.43 |
| CCBL1 | 6 | 10.13 | 10.71 | 10.12 | 10.06 | 8.38 | 8.30 | 10.02 | 9.36 | 11.55 | 11.38 | 10.42 | 10.09 | 8.34 | 9.69 | 11.46 |
| CCBL2 | 9 | 9.23 | 10.23 | 8.73 | 8.34 | 9.63 | 9.62 | 11.88 | 11.01 | 10.61 | 10.71 | 9.73 | 8.54 | 9.63 | 11.45 | 10.66 |
| CCDC101 | 4 | 9.60 | 11.32 | 11.50 | 11.12 | 8.80 | 9.45 | 8.83 | 9.25 | 9.98 | 10.16 | 10.46 | 11.31 | 9.13 | 9.04 | 10.07 |
| CCDC102A | 2 | 9.81 | 11.42 | 9.67 | 8.62 | 14.82 | 15.17 | 6.21 | 6.04 | 9.55 | 8.68 | 10.62 | 9.15 | 14.99 | 6.12 | 9.11 |
| CCDC104 | 1 | 10.72 | 15.02 | 9.66 | 9.63 | 8.80 | 11.87 | 9.14 | 7.91 | 8.17 | 9.08 | 12.87 | 9.65 | 10.33 | 8.53 | 8.63 |
| CCDC109B | 1 | 12.92 | 8.25 | 5.94 | 8.46 | 13.26 | 10.35 | 6.58 | 5.84 | 13.43 | 14.96 | 10.58 | 7.20 | 11.81 | 6.21 | 14.20 |
| CCDC115 | 1 | 9.21 | 9.48 | 12.92 | 12.89 | 8.27 | 10.43 | 7.66 | 7.93 | 10.76 | 10.46 | 9.34 | 12.90 | 9.35 | 7.79 | 10.61 |
| CCDC117 | 1 | 13.58 | 11.53 | 10.92 | 8.70 | 9.34 | 8.85 | 9.95 | 9.16 | 9.26 | 8.71 | 12.56 | 9.81 | 9.09 | 9.55 | 8.99 |
| CCDC12 | 6 | 10.70 | 10.88 | 10.61 | 10.52 | 9.37 | 9.43 | 10.21 | 8.54 | 10.03 | 9.71 | 10.79 | 10.57 | 9.40 | 9.37 | 9.87 |
| CCDC124 | 12 | 8.75 | 9.38 | 12.55 | 11.81 | 9.15 | 9.00 | 11.03 | 10.19 | 9.43 | 8.70 | 9.07 | 12.18 | 9.08 | 10.61 | 9.06 |
| CCDC127 | 6 | 11.42 | 10.15 | 9.05 | 10.41 | 9.92 | 11.16 | 8.71 | 8.80 | 9.71 | 10.66 | 10.79 | 9.73 | 10.54 | 8.75 | 10.19 |
| CCDC130 | 1 | 9.75 | 12.90 | 7.47 | 8.32 | 9.50 | 11.37 | 10.79 | 9.31 | 10.70 | 9.89 | 11.33 | 7.89 | 10.44 | 10.05 | 10.29 |
| CCDC132 | 15 | 10.16 | 10.46 | 11.74 | 11.20 | 9.79 | 9.67 | 9.17 | 8.32 | 9.96 | 9.52 | 10.31 | 11.47 | 9.73 | 8.74 | 9.74 |
| CCDC134 | 5 | 9.20 | 10.01 | 12.92 | 13.11 | 8.61 | 9.51 | 9.65 | 8.45 | 9.37 | 9.17 | 9.60 | 13.02 | 9.06 | 9.05 | 9.27 |
| CCDC137 | 4 | 9.71 | 10.21 | 11.88 | 12.00 | 8.00 | 8.28 | 9.64 | 8.97 | 10.62 | 10.67 | 9.96 | 11.94 | 8.14 | 9.31 | 10.65 |
| CCDC14 | 1 | 9.17 | 11.58 | 10.49 | 11.52 | 10.76 | 12.51 | 8.31 | 7.03 | 9.85 | 8.76 | 10.37 | 11.01 | 11.64 | 7.67 | 9.31 |
| CCDC152 | 1 | 8.78 | 9.08 | 9.17 | 9.75 | 12.87 | 12.62 | 11.20 | 8.47 | 10.44 | 7.62 | 8.93 | 9.46 | 12.75 | 9.84 | 9.03 |
| CCDC167 | 3 | 12.20 | 8.03 | 7.61 | 11.18 | 10.80 | 10.12 | 7.51 | 7.11 | 11.49 | 13.95 | 10.12 | 9.39 | 10.46 | 7.31 | 12.72 |
| CCDC168 | 2 | 14.78 | 10.11 | 8.14 | 8.30 | 10.63 | 11.59 | 9.29 | 8.90 | 10.56 | 7.71 | 12.44 | 8.22 | 11.11 | 9.09 | 9.14 |
| CCDC170 | 1 | 9.28 | 10.89 | 9.68 | 10.55 | 9.50 | 10.15 | 9.93 | 9.81 | 10.11 | 10.09 | 10.08 | 10.12 | 9.83 | 9.87 | 10.10 |
| CCDC174 | 2 | 9.84 | 10.30 | 8.17 | 8.94 | 10.46 | 10.24 | 10.15 | 10.28 | 11.43 | 10.19 | 10.07 | 8.55 | 10.35 | 10.21 | 10.81 |
| CCDC18 | 1 | 9.25 | 11.32 | 10.51 | 10.80 | 7.90 | 8.49 | 9.59 | 9.96 | 11.72 | 10.47 | 10.28 | 10.65 | 8.20 | 9.77 | 11.10 |
| CCDC22 | 10 | 9.17 | 10.26 | 10.30 | 9.84 | 11.35 | 12.41 | 9.99 | 9.62 | 8.68 | 8.36 | 9.72 | 10.07 | 11.88 | 9.81 | 8.52 |
| CCDC23 | 1 | 11.75 | 13.85 | 9.53 | 7.59 | 9.98 | 9.96 | 11.74 | 5.59 | 10.98 | 9.02 | 12.80 | 8.56 | 9.97 | 8.67 | 10.00 |
| CCDC25 | 12 | 10.03 | 9.04 | 9.13 | 8.08 | 10.14 | 10.42 | 12.74 | 12.79 | 9.21 | 8.41 | 9.54 | 8.61 | 10.28 | 12.77 | 8.81 |
| CCDC41 | 2 | 10.52 | 11.42 | 11.56 | 12.10 | 7.79 | 10.48 | 10.62 | 10.79 | 7.42 | 7.29 | 10.97 | 11.83 | 9.13 | 10.71 | 7.36 |
| CCDC43 | 10 | 10.78 | 12.32 | 10.28 | 10.84 | 8.81 | 10.08 | 7.98 | 7.39 | 10.44 | 11.08 | 11.55 | 10.56 | 9.44 | 7.69 | 10.76 |
| CCDC47 | 15 | 11.42 | 8.44 | 7.90 | 10.59 | 10.48 | 9.64 | 9.46 | 9.47 | 9.94 | 12.66 | 9.93 | 9.25 | 10.06 | 9.46 | 11.30 |
| CCDC50 | 14 | 10.78 | 12.06 | 12.19 | 11.23 | 11.46 | 12.18 | 5.99 | 5.49 | 9.85 | 8.75 | 11.42 | 11.71 | 11.82 | 5.74 | 9.30 |
| CCDC51 | 5 | 13.10 | 6.79 | 7.47 | 11.38 | 9.35 | 7.73 | 9.07 | 7.96 | 12.96 | 14.18 | 9.95 | 9.43 | 8.54 | 8.51 | 13.57 |
| CCDC53 | 4 | 10.15 | 11.24 | 9.66 | 10.34 | 10.72 | 10.94 | 8.36 | 6.97 | 10.74 | 10.88 | 10.69 | 10.00 | 10.83 | 7.66 | 10.81 |
| CCDC58 | 8 | 9.21 | 8.58 | 11.37 | 10.10 | 9.02 | 9.11 | 11.03 | 12.65 | 9.67 | 9.27 | 8.89 | 10.73 | 9.07 | 11.84 | 9.47 |
| CCDC59 | 5 | 9.74 | 8.35 | 12.37 | 12.09 | 8.87 | 7.78 | 11.98 | 9.69 | 8.52 | 10.63 | 9.04 | 12.23 | 8.32 | 10.83 | 9.57 |
| CCDC6 | 27 | 9.38 | 10.46 | 11.46 | 10.71 | 10.45 | 10.48 | 7.95 | 7.39 | 11.12 | 10.61 | 9.92 | 11.08 | 10.46 | 7.67 | 10.87 |
| CCDC77 | 1 | 10.48 | 11.06 | 10.13 | 9.06 | 10.75 | 9.62 | 11.41 | 13.10 | 6.92 | 7.47 | 10.77 | 9.59 | 10.19 | 12.26 | 7.19 |
| CCDC84 | 1 | 12.19 | 8.90 | 7.59 | 7.98 | 9.40 | 12.84 | 10.53 | 6.91 | 13.02 | 10.64 | 10.55 | 7.79 | 11.12 | 8.72 | 11.83 |
| CCDC85C | 2 | 11.34 | 12.36 | 11.68 | 10.43 | 10.84 | 10.33 | 6.92 | 5.98 | 9.47 | 10.63 | 11.85 | 11.06 | 10.59 | 6.45 | 10.05 |

| | | | | | | | | | | | | | | | | |
|---|---|---|---|---|---|---|---|---|---|---|---|---|---|---|---|---|
| CCDC86 | 16 | 10.11 | 8.98 | 9.63 | 10.05 | 9.01 | 9.07 | 11.71 | 11.83 | 9.47 | 10.14 | 9.55 | 9.84 | 9.04 | 11.77 | 9.80 |
| CCDC88A | 13 | 9.62 | 10.99 | 11.98 | 11.62 | 7.84 | 8.37 | 9.96 | 9.47 | 9.97 | 10.18 | 10.31 | 11.80 | 8.10 | 9.72 | 10.07 |
| CCDC9 | 8 | 9.12 | 11.54 | 10.24 | 10.59 | 10.02 | 10.18 | 10.29 | 8.70 | 9.92 | 9.39 | 10.33 | 10.42 | 10.10 | 9.49 | 9.66 |
| CCDC90B | 1 | 11.98 | 9.83 | 9.68 | 12.36 | 8.83 | 8.44 | 10.25 | 9.63 | 9.27 | 9.74 | 10.91 | 11.02 | 8.63 | 9.94 | 9.50 |
| CCDC91 | 10 | 9.61 | 10.29 | 11.68 | 11.10 | 8.98 | 9.21 | 11.92 | 11.02 | 8.45 | 7.74 | 9.95 | 11.39 | 9.09 | 11.47 | 8.09 |
| CCDC93 | 13 | 10.07 | 10.63 | 10.27 | 9.77 | 11.15 | 11.49 | 9.50 | 9.57 | 9.00 | 8.53 | 10.35 | 10.02 | 11.32 | 9.54 | 8.77 |
| CCDC94 | 2 | 8.69 | 10.52 | 10.20 | 10.69 | 8.94 | 9.97 | 11.05 | 12.51 | 9.46 | 7.96 | 9.61 | 10.45 | 9.46 | 11.78 | 8.71 |
| CCDC97 | 2 | 7.63 | 12.22 | 9.57 | 7.73 | 11.29 | 10.68 | 11.55 | 9.32 | 9.26 | 10.75 | 9.92 | 8.65 | 10.98 | 10.44 | 10.00 |
| CCM2 | 2 | 10.44 | 11.35 | 10.02 | 10.03 | 9.55 | 9.17 | 11.69 | 10.71 | 8.76 | 8.29 | 10.90 | 10.02 | 9.36 | 11.20 | 8.53 |
| CCNA2 | 9 | 8.77 | 9.37 | 10.79 | 9.24 | 10.76 | 10.74 | 9.78 | 10.22 | 10.39 | 9.95 | 9.07 | 10.01 | 10.75 | 10.00 | 10.17 |
| CCNB1 | 16 | 8.26 | 8.93 | 9.52 | 9.08 | 8.84 | 8.86 | 13.59 | 13.86 | 9.57 | 9.48 | 8.60 | 9.30 | 8.85 | 13.73 | 9.53 |
| CCNB2 | 8 | 6.82 | 7.84 | 13.03 | 12.71 | 9.48 | 8.91 | 9.69 | 10.04 | 10.71 | 10.77 | 7.33 | 12.87 | 9.19 | 9.86 | 10.74 |
| CCNH | 3 | 10.09 | 9.84 | 11.36 | 10.20 | 9.90 | 9.47 | 10.10 | 9.14 | 10.53 | 9.38 | 9.96 | 10.78 | 9.69 | 9.62 | 9.95 |
| CCNK | 9 | 9.82 | 10.37 | 10.53 | 9.83 | 9.91 | 10.33 | 9.90 | 9.73 | 9.98 | 9.61 | 10.09 | 10.18 | 10.12 | 9.81 | 9.80 |
| CCNL1 | 3 | 9.17 | 8.16 | 9.82 | 8.88 | 8.56 | 9.72 | 10.69 | 18.18 | 8.72 | 8.09 | 8.67 | 9.35 | 9.14 | 14.44 | 8.40 |
| CCNL2 | 3 | 9.69 | 10.18 | 8.83 | 8.05 | 9.61 | 9.10 | 12.42 | 11.41 | 10.72 | 10.00 | 9.94 | 8.44 | 9.35 | 11.91 | 10.36 |
| CCNT1 | 6 | 8.89 | 9.42 | 11.54 | 9.92 | 10.40 | 10.35 | 10.90 | 9.69 | 10.22 | 8.67 | 9.15 | 10.73 | 10.38 | 10.30 | 9.44 |
| CCPG1 | 5 | 12.87 | 6.66 | 7.15 | 10.99 | 12.54 | 10.30 | 6.66 | 5.50 | 12.41 | 14.90 | 9.77 | 9.07 | 11.42 | 6.08 | 13.66 |
| CCS | 7 | 10.30 | 11.85 | 6.56 | 5.88 | 12.97 | 13.96 | 10.76 | 9.87 | 9.23 | 8.62 | 11.08 | 6.22 | 13.46 | 10.32 | 8.92 |
| CCT2 | 77 | 9.47 | 10.73 | 10.80 | 10.12 | 9.59 | 9.70 | 10.05 | 9.97 | 9.93 | 9.63 | 10.10 | 10.46 | 9.65 | 10.01 | 9.78 |
| CCT3 | 87 | 10.20 | 10.98 | 9.85 | 8.38 | 10.65 | 10.70 | 9.71 | 9.11 | 10.38 | 10.03 | 10.59 | 9.12 | 10.68 | 9.41 | 10.20 |
| CCT4 | 74 | 8.62 | 9.50 | 10.33 | 9.34 | 9.04 | 9.21 | 9.80 | 15.96 | 9.12 | 9.06 | 9.06 | 9.84 | 9.13 | 12.88 | 9.09 |
| CCT5 | 121 | 9.21 | 10.16 | 9.85 | 9.51 | 9.63 | 9.65 | 10.07 | 12.80 | 9.68 | 9.44 | 9.69 | 9.68 | 9.64 | 11.44 | 9.56 |
| CCT6A | 72 | 9.61 | 10.74 | 9.95 | 9.02 | 9.84 | 9.69 | 9.77 | 10.94 | 10.32 | 10.12 | 10.18 | 9.48 | 9.77 | 10.35 | 10.22 |
| CCT7 | 70 | 8.93 | 10.02 | 9.61 | 9.28 | 9.60 | 9.69 | 9.67 | 14.64 | 9.37 | 9.19 | 9.47 | 9.45 | 9.65 | 12.16 | 9.28 |
| CCT8 | 109 | 9.68 | 10.48 | 9.36 | 8.64 | 10.45 | 10.63 | 9.55 | 10.61 | 10.44 | 10.16 | 10.08 | 9.00 | 10.54 | 10.08 | 10.30 |
| CCZ1 | 6 | 9.37 | 10.16 | 13.07 | 10.26 | 8.67 | 8.30 | 8.73 | 7.77 | 11.97 | 11.69 | 9.77 | 11.67 | 8.48 | 8.25 | 11.83 |
| CD109 | 23 | 9.89 | 9.73 | 9.67 | 9.18 | 11.59 | 11.63 | 9.13 | 8.82 | 9.99 | 10.39 | 9.81 | 9.42 | 11.61 | 8.97 | 10.19 |
| CD151 | 2 | 10.65 | 8.72 | 7.03 | 8.20 | 7.12 | 7.49 | 7.26 | 21.78 | 9.99 | 11.76 | 9.69 | 7.62 | 7.30 | 14.52 | 10.87 |
| CD276 | 5 | 9.65 | 7.76 | 10.00 | 12.14 | 10.95 | 10.96 | 8.32 | 7.82 | 11.10 | 11.29 | 8.70 | 11.07 | 10.95 | 8.07 | 11.20 |
| CD2AP | 30 | 7.98 | 8.75 | 14.52 | 12.90 | 9.58 | 9.38 | 9.61 | 8.28 | 9.65 | 9.34 | 8.37 | 13.71 | 9.48 | 8.95 | 9.49 |
| CD2BP2 | 12 | 10.15 | 13.32 | 12.34 | 11.67 | 8.72 | 8.93 | 8.20 | 7.10 | 9.98 | 9.60 | 11.73 | 12.00 | 8.83 | 7.65 | 9.79 |
| CD320 | 1 | 8.16 | 6.62 | 8.11 | 9.34 | 11.52 | 9.87 | 13.60 | 11.39 | 9.29 | 12.10 | 7.39 | 8.73 | 10.69 | 12.50 | 10.69 |
| CD3EAP | 10 | 8.67 | 8.76 | 12.60 | 11.50 | 8.83 | 8.11 | 11.75 | 9.13 | 10.41 | 10.23 | 8.72 | 12.05 | 8.47 | 10.44 | 10.32 |
| CD44 | 20 | 10.17 | 8.21 | 8.29 | 9.37 | 8.02 | 7.89 | 12.16 | 20.51 | 7.51 | 7.87 | 9.19 | 8.83 | 7.96 | 16.33 | 7.69 |
| CD46 | 4 | 9.91 | 7.82 | 7.16 | 8.87 | 15.30 | 14.88 | 7.56 | 6.31 | 10.66 | 11.53 | 8.86 | 8.02 | 15.09 | 6.94 | 11.10 |
| CD47 | 3 | 12.07 | 9.82 | 8.86 | 11.55 | 9.72 | 11.01 | 8.52 | 8.24 | 9.84 | 10.38 | 10.95 | 10.20 | 10.36 | 8.38 | 10.11 |
| CD55 | 8 | 6.67 | 5.34 | 3.72 | 3.41 | 12.94 | 11.15 | 17.97 | 28.12 | 5.60 | 5.09 | 6.00 | 3.57 | 12.04 | 23.04 | 5.34 |
| CD58 | 3 | 10.58 | 8.82 | 8.44 | 10.30 | 8.95 | 9.37 | 12.06 | 10.13 | 10.74 | 10.60 | 9.70 | 9.37 | 9.16 | 11.10 | 10.67 |
| CD59 | 6 | 10.48 | 7.73 | 7.86 | 8.58 | 10.45 | 9.72 | 6.79 | 17.57 | 10.60 | 10.21 | 9.10 | 8.22 | 10.09 | 12.18 | 10.41 |
| CD63 | 4 | 10.78 | 7.22 | 8.64 | 10.67 | 12.58 | 10.60 | 11.10 | 8.26 | 9.71 | 10.45 | 9.00 | 9.65 | 11.59 | 9.68 | 10.08 |
| CD70 | 2 | 8.45 | 6.95 | 7.43 | 9.94 | 8.46 | 7.59 | 5.35 | 5.40 | 19.07 | 21.37 | 7.70 | 8.69 | 8.02 | 5.37 | 20.22 |

| | | | | | | | | | | | | | | | | |
|---|---|---|---|---|---|---|---|---|---|---|---|---|---|---|---|---|
| CD81 | 2 | 9.10 | 7.31 | 14.75 | 18.82 | 5.59 | 6.31 | 7.61 | 6.88 | 11.34 | 12.28 | 8.20 | 16.79 | 5.95 | 7.25 | 11.81 |
| CD9 | 5 | 9.01 | 7.69 | 6.53 | 6.59 | 6.60 | 5.82 | 10.16 | 25.05 | 10.52 | 12.02 | 8.35 | 6.56 | 6.21 | 17.61 | 11.27 |
| CD97 | 11 | 7.86 | 7.75 | 7.91 | 8.30 | 12.84 | 12.66 | 10.83 | 12.28 | 9.80 | 9.78 | 7.81 | 8.10 | 12.75 | 11.55 | 9.79 |
| CD99 | 7 | 11.69 | 9.35 | 5.90 | 8.46 | 15.29 | 14.35 | 9.27 | 4.16 | 10.87 | 10.66 | 10.52 | 7.18 | 14.82 | 6.71 | 10.77 |
| CD99L2 | 3 | 13.28 | 10.23 | 5.98 | 6.81 | 16.66 | 15.36 | 11.96 | 6.03 | 7.11 | 6.59 | 11.75 | 6.40 | 16.01 | 8.99 | 6.85 |
| CDA | 7 | 8.47 | 8.25 | 4.72 | 3.72 | 9.19 | 8.44 | 9.08 | 35.17 | 7.17 | 5.80 | 8.36 | 4.22 | 8.81 | 22.13 | 6.48 |
| CDAN1 | 3 | 11.19 | 12.64 | 8.24 | 8.05 | 12.72 | 10.92 | 7.23 | 7.39 | 10.88 | 10.73 | 11.91 | 8.15 | 11.82 | 7.31 | 10.80 |
| CDC123 | 11 | 9.29 | 10.40 | 10.66 | 10.65 | 9.20 | 9.36 | 10.09 | 11.55 | 9.39 | 9.41 | 9.85 | 10.65 | 9.28 | 10.82 | 9.40 |
| CDC16 | 11 | 10.24 | 10.64 | 10.51 | 9.26 | 9.77 | 10.13 | 10.77 | 10.13 | 9.56 | 8.99 | 10.44 | 9.88 | 9.95 | 10.45 | 9.27 |
| CDC20 | 6 | 9.54 | 9.66 | 11.77 | 10.66 | 7.74 | 7.10 | 10.07 | 9.20 | 12.69 | 11.58 | 9.60 | 11.21 | 7.42 | 9.63 | 12.14 |
| CDC23 | 10 | 9.48 | 10.36 | 10.75 | 10.15 | 9.58 | 9.04 | 10.98 | 11.07 | 9.61 | 8.97 | 9.92 | 10.45 | 9.31 | 11.02 | 9.29 |
| CDC25C | 1 | 11.58 | 12.17 | 5.55 | 8.80 | 9.37 | 11.28 | 11.95 | 12.89 | 7.67 | 8.74 | 11.87 | 7.17 | 10.33 | 12.42 | 8.20 |
| CDC26 | 1 | 13.95 | 11.49 | 11.27 | 10.87 | 8.20 | 11.06 | 9.83 | 6.74 | 9.25 | 7.33 | 12.72 | 11.07 | 9.63 | 8.29 | 8.29 |
| CDC27 | 17 | 10.07 | 10.67 | 10.89 | 11.10 | 8.84 | 9.08 | 9.34 | 10.40 | 9.84 | 9.77 | 10.37 | 11.00 | 8.96 | 9.87 | 9.81 |
| CDC34 | 3 | 8.40 | 11.09 | 10.13 | 9.28 | 9.89 | 9.65 | 11.90 | 10.25 | 9.61 | 9.78 | 9.75 | 9.71 | 9.77 | 11.08 | 9.70 |
| CDC37 | 22 | 8.54 | 9.68 | 13.41 | 12.31 | 9.07 | 8.92 | 9.29 | 9.25 | 10.06 | 9.47 | 9.11 | 12.86 | 9.00 | 9.27 | 9.76 |
| CDC37L1 | 4 | 9.58 | 11.55 | 8.71 | 9.65 | 9.03 | 9.90 | 7.82 | 8.03 | 12.49 | 13.24 | 10.56 | 9.18 | 9.47 | 7.93 | 12.86 |
| CDC40 | 9 | 9.78 | 10.76 | 10.70 | 10.32 | 9.49 | 9.73 | 10.94 | 9.66 | 9.45 | 9.17 | 10.27 | 10.51 | 9.61 | 10.30 | 9.31 |
| CDC42 | 12 | 11.33 | 7.17 | 8.21 | 8.70 | 8.56 | 8.25 | 7.45 | 18.28 | 10.81 | 11.24 | 9.25 | 8.46 | 8.40 | 12.86 | 11.03 |
| CDC42BPA | 14 | 9.32 | 10.39 | 8.91 | 8.35 | 11.78 | 12.32 | 8.26 | 7.80 | 11.66 | 11.21 | 9.86 | 8.63 | 12.05 | 8.03 | 11.43 |
| CDC42BPB | 24 | 9.66 | 10.54 | 8.87 | 8.41 | 11.75 | 12.17 | 9.72 | 8.70 | 10.17 | 10.01 | 10.10 | 8.64 | 11.96 | 9.21 | 10.09 |
| CDC42EP1 | 8 | 10.17 | 10.89 | 13.17 | 13.71 | 6.78 | 7.32 | 7.25 | 6.20 | 12.66 | 11.84 | 10.53 | 13.44 | 7.05 | 6.73 | 12.25 |
| CDC42EP2 | 3 | 10.58 | 11.86 | 10.54 | 12.62 | 8.24 | 9.31 | 6.34 | 7.28 | 11.66 | 11.56 | 11.22 | 11.58 | 8.78 | 6.81 | 11.61 |
| CDC42EP3 | 4 | 9.51 | 11.68 | 10.59 | 9.95 | 8.21 | 8.97 | 8.27 | 6.92 | 12.74 | 13.14 | 10.60 | 10.27 | 8.59 | 7.60 | 12.94 |
| CDC42EP4 | 9 | 10.29 | 11.04 | 10.33 | 9.82 | 7.99 | 8.11 | 12.32 | 11.11 | 9.97 | 9.03 | 10.66 | 10.07 | 8.05 | 11.71 | 9.50 |
| CDC45 | 2 | 11.14 | 10.57 | 10.04 | 11.16 | 5.63 | 5.71 | 11.59 | 10.25 | 12.29 | 11.61 | 10.86 | 10.60 | 5.67 | 10.92 | 11.95 |
| CDC5L | 38 | 9.36 | 9.86 | 11.39 | 10.65 | 9.91 | 9.98 | 10.60 | 10.26 | 9.16 | 8.84 | 9.61 | 11.02 | 9.94 | 10.43 | 9.00 |
| CDC6 | 1 | 9.88 | 9.85 | 13.45 | 12.66 | 9.03 | 9.09 | 7.36 | 6.39 | 11.42 | 10.87 | 9.87 | 13.05 | 9.06 | 6.87 | 11.14 |
| CDC7 | 2 | 10.02 | 10.78 | 9.67 | 10.70 | 8.95 | 8.96 | 11.47 | 10.28 | 9.92 | 9.26 | 10.40 | 10.18 | 8.95 | 10.87 | 9.59 |
| CDC73 | 24 | 9.74 | 10.55 | 9.18 | 8.63 | 11.39 | 11.75 | 9.43 | 8.84 | 10.52 | 9.97 | 10.15 | 8.91 | 11.57 | 9.14 | 10.24 |
| CDCA2 | 4 | 8.97 | 10.38 | 9.31 | 8.74 | 9.37 | 10.10 | 12.27 | 11.76 | 9.52 | 9.60 | 9.67 | 9.02 | 9.73 | 12.01 | 9.56 |
| CDCA3 | 4 | 9.18 | 10.26 | 8.05 | 11.08 | 10.07 | 11.66 | 9.55 | 7.76 | 10.47 | 11.92 | 9.72 | 9.56 | 10.87 | 8.66 | 11.19 |
| CDCA5 | 3 | 8.86 | 10.35 | 8.90 | 8.27 | 10.99 | 10.99 | 10.63 | 11.70 | 9.49 | 9.83 | 9.61 | 8.59 | 10.99 | 11.16 | 9.66 |
| CDCA7L | 5 | 11.45 | 11.51 | 10.58 | 9.30 | 8.76 | 8.24 | 9.19 | 7.82 | 11.95 | 11.20 | 11.48 | 9.94 | 8.50 | 8.51 | 11.57 |
| CDCA8 | 7 | 10.01 | 10.08 | 9.10 | 8.89 | 8.49 | 8.47 | 11.54 | 10.00 | 11.50 | 11.92 | 10.05 | 8.99 | 8.48 | 10.77 | 11.71 |
| CDCP1 | 11 | 9.69 | 8.16 | 9.30 | 11.37 | 5.82 | 6.42 | 5.95 | 5.33 | 18.39 | 19.57 | 8.92 | 10.33 | 6.12 | 5.64 | 18.98 |
| CDH13 | 4 | 11.95 | 9.11 | 10.00 | 10.81 | 8.16 | 7.40 | 13.46 | 10.33 | 9.27 | 9.50 | 10.53 | 10.40 | 7.78 | 11.89 | 9.39 |
| CDH2 | 16 | 11.34 | 8.70 | 12.47 | 15.72 | 7.26 | 7.32 | 4.23 | 4.05 | 14.44 | 14.48 | 10.02 | 14.10 | 7.29 | 4.14 | 14.46 |
| CDIPT | 1 | 9.32 | 5.70 | 10.92 | 3.31 | 14.61 | 6.03 | 10.40 | 5.73 | 11.93 | 22.05 | 7.51 | 7.12 | 10.32 | 8.06 | 16.99 |
| CDK1 | 41 | 7.79 | 8.82 | 11.48 | 8.43 | 9.79 | 9.05 | 14.06 | 12.15 | 9.09 | 9.33 | 8.31 | 9.95 | 9.42 | 13.11 | 9.21 |
| CDK11B | 23 | 11.23 | 11.65 | 10.26 | 9.46 | 9.33 | 9.14 | 10.24 | 9.25 | 10.01 | 9.44 | 11.44 | 9.86 | 9.23 | 9.74 | 9.72 |
| CDK12 | 17 | 9.97 | 10.12 | 11.11 | 10.29 | 9.49 | 9.71 | 9.53 | 8.78 | 10.97 | 10.04 | 10.05 | 10.70 | 9.60 | 9.15 | 10.50 |

| | | | | | | | | | | | | | | | | |
|---|---|---|---|---|---|---|---|---|---|---|---|---|---|---|---|---|
| CDK13 | 9 | 9.35 | 10.30 | 9.85 | 10.38 | 8.45 | 9.26 | 11.58 | 10.84 | 10.14 | 9.85 | 9.83 | 10.11 | 8.86 | 11.21 | 10.00 |
| CDK16 | 6 | 11.19 | 9.50 | 7.47 | 8.36 | 12.44 | 11.93 | 10.21 | 9.13 | 9.99 | 9.77 | 10.35 | 7.91 | 12.18 | 9.67 | 9.88 |
| CDK17 | 2 | 8.40 | 9.30 | 12.24 | 11.61 | 8.18 | 9.18 | 11.40 | 10.33 | 9.55 | 9.83 | 8.85 | 11.92 | 8.68 | 10.86 | 9.69 |
| CDK2 | 17 | 9.30 | 10.29 | 11.36 | 10.42 | 11.02 | 11.10 | 7.75 | 8.39 | 10.11 | 10.27 | 9.79 | 10.89 | 11.06 | 8.07 | 10.19 |
| CDK2AP1 | 2 | 8.20 | 9.74 | 11.27 | 9.81 | 10.53 | 9.34 | 12.49 | 10.04 | 8.93 | 9.64 | 8.97 | 10.54 | 9.94 | 11.27 | 9.29 |
| CDK3 | 3 | 9.51 | 9.39 | 10.71 | 9.61 | 9.30 | 9.01 | 11.28 | 11.15 | 10.19 | 9.84 | 9.45 | 10.16 | 9.16 | 11.22 | 10.01 |
| CDK4 | 8 | 9.38 | 9.78 | 10.94 | 9.98 | 9.97 | 9.54 | 10.25 | 9.89 | 10.23 | 10.04 | 9.58 | 10.46 | 9.76 | 10.07 | 10.14 |
| CDK5 | 11 | 10.12 | 10.05 | 11.32 | 10.52 | 8.42 | 8.23 | 10.34 | 9.17 | 11.40 | 10.45 | 10.08 | 10.92 | 8.32 | 9.75 | 10.92 |
| CDK5RAP1 | 3 | 10.23 | 9.91 | 13.19 | 11.26 | 8.92 | 7.83 | 10.09 | 8.46 | 10.09 | 10.00 | 10.07 | 12.23 | 8.38 | 9.28 | 10.04 |
| CDK5RAP2 | 14 | 10.21 | 11.01 | 9.03 | 8.79 | 10.47 | 10.37 | 9.48 | 10.00 | 10.44 | 10.20 | 10.61 | 8.91 | 10.42 | 9.74 | 10.32 |
| CDK5RAP3 | 13 | 9.98 | 10.52 | 8.93 | 7.73 | 12.17 | 11.36 | 9.45 | 10.20 | 10.25 | 9.42 | 10.25 | 8.33 | 11.76 | 9.82 | 9.84 |
| CDK6 | 13 | 7.79 | 7.99 | 13.79 | 13.62 | 7.48 | 7.42 | 10.98 | 10.32 | 10.37 | 10.26 | 7.89 | 13.70 | 7.45 | 10.65 | 10.31 |
| CDK7 | 5 | 9.75 | 10.71 | 11.42 | 11.39 | 9.03 | 9.49 | 9.65 | 9.06 | 9.98 | 9.54 | 10.23 | 11.41 | 9.26 | 9.35 | 9.76 |
| CDK9 | 7 | 8.29 | 8.99 | 9.45 | 9.16 | 9.47 | 9.93 | 10.10 | 15.54 | 9.74 | 9.34 | 8.64 | 9.30 | 9.70 | 12.82 | 9.54 |
| CDKAL1 | 2 | 11.25 | 8.84 | 8.39 | 10.46 | 9.04 | 8.48 | 12.36 | 12.97 | 8.70 | 9.50 | 10.05 | 9.43 | 8.76 | 12.67 | 9.10 |
| CDKN1A | 2 | 10.23 | 10.90 | 9.22 | 11.84 | 8.00 | 8.01 | 10.12 | 12.01 | 10.74 | 8.92 | 10.57 | 10.53 | 8.01 | 11.06 | 9.83 |
| CDKN2A | 22 | 9.65 | 10.22 | 10.76 | 10.03 | 7.03 | 6.89 | 8.07 | 9.25 | 14.12 | 13.99 | 9.93 | 10.40 | 6.96 | 8.66 | 14.06 |
| CDKN2AIP | 17 | 9.91 | 12.94 | 9.56 | 9.65 | 12.39 | 13.08 | 6.71 | 6.42 | 9.81 | 9.53 | 11.42 | 9.61 | 12.74 | 6.57 | 9.67 |
| CDKN2AIPNL | 2 | 9.23 | 9.77 | 13.98 | 14.38 | 5.34 | 6.37 | 14.10 | 4.64 | 11.82 | 10.38 | 9.50 | 14.18 | 5.85 | 9.37 | 11.10 |
| CDKN2C | 4 | 12.65 | 14.61 | 7.57 | 7.79 | 12.43 | 13.34 | 7.01 | 7.87 | 8.49 | 8.24 | 13.63 | 7.68 | 12.89 | 7.44 | 8.36 |
| CDR2 | 3 | 11.45 | 11.95 | 12.39 | 11.74 | 8.10 | 9.56 | 6.75 | 5.03 | 11.54 | 11.49 | 11.70 | 12.07 | 8.83 | 5.89 | 11.51 |
| CDR2L | 1 | 11.75 | 13.15 | 9.81 | 9.85 | 9.60 | 10.85 | 6.62 | 5.22 | 11.50 | 11.64 | 12.45 | 9.83 | 10.23 | 5.92 | 11.57 |
| CDS2 | 1 | 12.58 | 7.42 | 8.25 | 11.86 | 8.15 | 8.44 | 9.57 | 7.32 | 11.04 | 15.38 | 10.00 | 10.06 | 8.29 | 8.44 | 13.21 |
| CDT1 | 2 | 9.17 | 13.45 | 9.78 | 8.07 | 10.32 | 9.89 | 10.84 | 9.18 | 10.35 | 8.95 | 11.31 | 8.93 | 10.10 | 10.01 | 9.65 |
| CDV3 | 30 | 9.62 | 10.93 | 12.00 | 11.08 | 9.66 | 9.72 | 8.94 | 7.44 | 10.64 | 9.98 | 10.27 | 11.54 | 9.69 | 8.19 | 10.31 |
| CDYL | 8 | 9.33 | 10.35 | 11.04 | 10.65 | 9.23 | 9.33 | 11.58 | 11.52 | 9.03 | 7.94 | 9.84 | 10.85 | 9.28 | 11.55 | 8.49 |
| CEBPB | 6 | 7.22 | 8.95 | 8.07 | 8.36 | 6.95 | 5.90 | 15.60 | 28.40 | 5.44 | 5.11 | 8.09 | 8.22 | 6.42 | 22.00 | 5.28 |
| CEBPZ | 31 | 9.69 | 9.46 | 10.77 | 10.20 | 9.47 | 9.25 | 11.26 | 10.19 | 10.11 | 9.60 | 9.58 | 10.48 | 9.36 | 10.73 | 9.85 |
| CECR5 | 8 | 10.89 | 9.89 | 9.58 | 11.47 | 6.42 | 6.25 | 12.26 | 13.07 | 9.85 | 10.32 | 10.39 | 10.53 | 6.33 | 12.66 | 10.09 |
| CELF1 | 24 | 8.85 | 9.24 | 11.05 | 10.43 | 9.89 | 10.04 | 10.52 | 11.01 | 9.84 | 9.12 | 9.04 | 10.74 | 9.96 | 10.77 | 9.48 |
| CENPB | 6 | 8.40 | 8.88 | 10.18 | 10.37 | 9.03 | 9.38 | 12.02 | 11.37 | 9.80 | 10.56 | 8.64 | 10.28 | 9.21 | 11.70 | 10.18 |
| CENPC | 1 | 10.15 | 10.60 | 9.17 | 9.67 | 8.79 | 9.25 | 11.65 | 11.53 | 10.29 | 8.91 | 10.37 | 9.42 | 9.02 | 11.59 | 9.60 |
| CENPE | 11 | 9.43 | 10.36 | 8.78 | 9.04 | 10.72 | 11.03 | 10.79 | 10.45 | 9.87 | 9.53 | 9.89 | 8.91 | 10.88 | 10.62 | 9.70 |
| CENPF | 75 | 8.97 | 9.99 | 8.55 | 8.26 | 12.14 | 12.52 | 11.38 | 10.64 | 9.09 | 8.45 | 9.48 | 8.40 | 12.33 | 11.01 | 8.77 |
| CENPH | 5 | 10.46 | 10.57 | 10.41 | 10.31 | 7.44 | 7.77 | 12.56 | 11.29 | 9.76 | 9.41 | 10.52 | 10.36 | 7.61 | 11.93 | 9.58 |
| CENPI | 3 | 10.79 | 10.22 | 10.60 | 8.76 | 9.28 | 8.34 | 11.68 | 10.53 | 9.92 | 9.88 | 10.51 | 9.68 | 8.81 | 11.10 | 9.90 |
| CENPK | 1 | 10.14 | 13.19 | 8.95 | 10.12 | 8.51 | 10.98 | 9.84 | 10.29 | 8.70 | 9.28 | 11.66 | 9.54 | 9.75 | 10.07 | 8.99 |
| CENPL | 1 | 9.22 | 9.71 | 9.20 | 10.83 | 9.41 | 8.69 | 10.38 | 10.58 | 11.18 | 10.79 | 9.47 | 10.02 | 9.05 | 10.48 | 10.99 |
| CENPM | 1 | 12.10 | 11.89 | 11.57 | 10.72 | 8.33 | 7.13 | 8.80 | 9.50 | 9.60 | 10.34 | 12.00 | 11.15 | 7.73 | 9.15 | 9.97 |
| CENPO | 1 | 9.19 | 10.70 | 10.79 | 12.38 | 6.74 | 7.30 | 10.77 | 11.94 | 10.83 | 9.35 | 9.95 | 11.58 | 7.02 | 11.36 | 10.09 |
| CENPU | 2 | 10.00 | 9.70 | 11.45 | 10.18 | 9.05 | 8.48 | 12.24 | 9.62 | 9.72 | 9.56 | 9.85 | 10.81 | 8.77 | 10.93 | 9.64 |
| CENPV | 3 | 9.94 | 11.38 | 7.33 | 7.03 | 13.44 | 13.01 | 9.87 | 8.30 | 9.73 | 9.98 | 10.66 | 7.18 | 13.22 | 9.08 | 9.85 |

| | | | | | | | | | | | | | | | | |
|---|---|---|---|---|---|---|---|---|---|---|---|---|---|---|---|---|
| CEP152 | 2 | 8.81 | 9.65 | 10.38 | 9.77 | 9.96 | 10.73 | 10.43 | 9.89 | 10.31 | 10.05 | 9.23 | 10.08 | 10.35 | 10.16 | 10.18 |
| CEP170 | 33 | 8.45 | 9.89 | 9.70 | 9.16 | 10.30 | 10.48 | 11.35 | 10.25 | 10.22 | 10.21 | 9.17 | 9.43 | 10.39 | 10.80 | 10.21 |
| CEP170B | 3 | 8.48 | 9.88 | 10.10 | 9.15 | 10.87 | 10.41 | 11.49 | 9.31 | 10.22 | 10.10 | 9.18 | 9.62 | 10.64 | 10.40 | 10.16 |
| CEP192 | 1 | 11.09 | 10.03 | 13.18 | 7.03 | 10.86 | 12.73 | 9.39 | 6.48 | 10.84 | 8.37 | 10.56 | 10.10 | 11.80 | 7.94 | 9.61 |
| CEP250 | 2 | 11.47 | 10.01 | 8.06 | 9.00 | 10.75 | 12.33 | 10.03 | 8.90 | 9.73 | 9.73 | 10.74 | 8.53 | 11.54 | 9.46 | 9.73 |
| CEP350 | 2 | 10.38 | 9.90 | 8.51 | 10.81 | 11.71 | 13.58 | 8.93 | 7.64 | 9.67 | 8.86 | 10.14 | 9.66 | 12.64 | 8.29 | 9.26 |
| CEP41 | 8 | 9.59 | 11.04 | 12.22 | 11.06 | 6.75 | 7.26 | 14.36 | 13.51 | 7.66 | 6.56 | 10.32 | 11.64 | 7.00 | 13.93 | 7.11 |
| CEP44 | 2 | 10.87 | 11.16 | 10.84 | 10.50 | 11.36 | 11.56 | 8.72 | 8.46 | 8.22 | 8.29 | 11.02 | 10.67 | 11.46 | 8.59 | 8.26 |
| CEP55 | 9 | 8.48 | 10.16 | 12.60 | 11.28 | 9.86 | 9.71 | 11.11 | 9.81 | 8.56 | 8.44 | 9.32 | 11.94 | 9.78 | 10.46 | 8.50 |
| CEP72 | 2 | 8.96 | 11.23 | 7.91 | 7.92 | 11.61 | 11.25 | 14.16 | 12.17 | 6.66 | 8.12 | 10.10 | 7.91 | 11.43 | 13.17 | 7.39 |
| CEP78 | 2 | 11.18 | 10.88 | 10.20 | 9.86 | 9.01 | 9.10 | 10.76 | 8.45 | 10.68 | 9.86 | 11.03 | 10.03 | 9.06 | 9.61 | 10.27 |
| CEP85 | 6 | 10.39 | 11.28 | 10.14 | 9.92 | 7.94 | 7.36 | 11.14 | 9.06 | 11.22 | 11.55 | 10.83 | 10.03 | 7.65 | 10.10 | 11.38 |
| CEP97 | 6 | 8.28 | 10.75 | 8.88 | 10.18 | 12.30 | 13.12 | 9.41 | 8.60 | 9.09 | 9.40 | 9.51 | 9.53 | 12.71 | 9.00 | 9.25 |
| CERS6 | 2 | 11.36 | 6.77 | 11.03 | 16.44 | 7.40 | 7.11 | 8.74 | 7.59 | 10.02 | 13.52 | 9.07 | 13.74 | 7.25 | 8.17 | 11.77 |
| CETN2 | 5 | 10.75 | 11.53 | 10.72 | 9.15 | 11.54 | 11.18 | 10.55 | 6.90 | 9.48 | 8.20 | 11.14 | 9.93 | 11.36 | 8.72 | 8.84 |
| CETN3 | 3 | 11.66 | 12.89 | 8.92 | 8.84 | 8.87 | 9.54 | 10.35 | 9.31 | 9.37 | 10.24 | 12.27 | 8.88 | 9.21 | 9.83 | 9.81 |
| CFDP1 | 9 | 10.90 | 11.11 | 6.94 | 6.48 | 11.41 | 10.90 | 12.20 | 10.31 | 10.45 | 9.28 | 11.01 | 6.71 | 11.16 | 11.26 | 9.86 |
| CFL1 | 82 | 7.62 | 11.84 | 12.06 | 11.86 | 8.52 | 8.42 | 7.40 | 13.36 | 9.58 | 9.34 | 9.73 | 11.96 | 8.47 | 10.38 | 9.46 |
| CFL2 | 14 | 11.01 | 10.86 | 10.33 | 10.40 | 12.01 | 12.71 | 6.07 | 5.83 | 11.06 | 9.71 | 10.93 | 10.36 | 12.36 | 5.95 | 10.39 |
| CGA | 6 | 3.65 | 3.54 | 4.21 | 4.43 | 29.86 | 30.23 | 8.90 | 7.90 | 4.14 | 3.13 | 3.60 | 4.32 | 30.05 | 8.40 | 3.64 |
| CGGBP1 | 7 | 9.64 | 10.79 | 9.29 | 8.20 | 10.47 | 10.60 | 10.68 | 9.47 | 10.82 | 10.04 | 10.21 | 8.75 | 10.54 | 10.07 | 10.43 |
| CGN | 1 | 11.18 | 10.69 | 20.46 | 21.86 | 2.10 | 2.65 | 3.31 | 2.24 | 13.31 | 12.21 | 10.93 | 21.16 | 2.38 | 2.77 | 12.76 |
| CGREF1 | 4 | 13.46 | 8.03 | 7.68 | 8.11 | 5.91 | 6.15 | 15.23 | 11.47 | 15.37 | 8.58 | 10.75 | 7.90 | 6.03 | 13.35 | 11.97 |
| CHAC2 | 8 | 10.72 | 11.82 | 9.24 | 9.21 | 10.45 | 10.94 | 9.41 | 8.90 | 9.72 | 9.59 | 11.27 | 9.22 | 10.69 | 9.15 | 9.66 |
| CHAF1A | 13 | 9.68 | 11.17 | 11.08 | 10.76 | 8.98 | 9.37 | 9.96 | 9.70 | 9.68 | 9.63 | 10.43 | 10.92 | 9.17 | 9.83 | 9.65 |
| CHAF1B | 8 | 9.31 | 10.08 | 11.47 | 10.69 | 9.73 | 9.65 | 10.03 | 9.35 | 10.11 | 9.58 | 9.70 | 11.08 | 9.69 | 9.69 | 9.84 |
| CHAMP1 | 24 | 9.04 | 10.68 | 10.84 | 10.11 | 11.04 | 11.48 | 10.17 | 9.23 | 9.19 | 8.22 | 9.86 | 10.47 | 11.26 | 9.70 | 8.70 |
| CHCHD1 | 1 | 10.75 | 9.22 | 9.95 | 9.91 | 11.57 | 10.87 | 10.49 | 9.19 | 9.41 | 8.63 | 9.98 | 9.93 | 11.22 | 9.84 | 9.02 |
| CHCHD2 | 7 | 7.81 | 7.88 | 11.75 | 13.33 | 7.22 | 7.47 | 8.17 | 18.44 | 8.89 | 9.04 | 7.85 | 12.54 | 7.34 | 13.30 | 8.96 |
| CHCHD3 | 19 | 10.06 | 8.78 | 12.46 | 12.00 | 9.19 | 8.97 | 10.58 | 9.13 | 9.61 | 9.24 | 9.42 | 12.23 | 9.08 | 9.85 | 9.43 |
| CHCHD4 | 2 | 11.11 | 10.03 | 10.42 | 10.06 | 11.87 | 12.12 | 12.61 | 8.26 | 6.78 | 6.72 | 10.57 | 10.24 | 12.00 | 10.44 | 6.75 |
| CHCHD5 | 1 | 8.40 | 8.57 | 9.33 | 9.70 | 14.36 | 13.96 | 10.50 | 9.08 | 7.89 | 8.21 | 8.49 | 9.51 | 14.16 | 9.79 | 8.05 |
| CHD1 | 17 | 9.22 | 9.93 | 10.01 | 9.89 | 7.77 | 7.73 | 14.17 | 12.81 | 9.48 | 9.00 | 9.57 | 9.95 | 7.75 | 13.49 | 9.24 |
| CHD1L | 7 | 10.57 | 12.13 | 8.07 | 7.85 | 9.26 | 9.48 | 10.42 | 9.90 | 10.99 | 11.32 | 11.35 | 7.96 | 9.37 | 10.16 | 11.16 |
| CHD2 | 12 | 9.79 | 10.32 | 10.54 | 8.92 | 10.10 | 9.40 | 11.00 | 9.90 | 10.29 | 9.75 | 10.06 | 9.73 | 9.75 | 10.45 | 10.02 |
| CHD3 | 33 | 9.91 | 10.23 | 10.77 | 10.64 | 9.93 | 10.18 | 10.21 | 9.43 | 9.50 | 9.20 | 10.07 | 10.71 | 10.06 | 9.82 | 9.35 |
| CHD4 | 55 | 8.97 | 9.61 | 10.37 | 10.03 | 9.87 | 9.77 | 11.02 | 12.18 | 9.26 | 8.91 | 9.29 | 10.20 | 9.82 | 11.60 | 9.08 |
| CHD5 | 6 | 8.74 | 9.33 | 11.38 | 10.96 | 10.57 | 8.96 | 11.62 | 9.10 | 8.78 | 10.55 | 9.04 | 11.17 | 9.77 | 10.36 | 9.66 |
| CHD7 | 5 | 9.53 | 10.85 | 10.27 | 10.45 | 9.11 | 10.10 | 10.65 | 10.65 | 9.61 | 8.78 | 10.19 | 10.36 | 9.61 | 10.65 | 9.19 |
| CHD8 | 9 | 8.20 | 9.89 | 10.84 | 10.12 | 10.20 | 10.60 | 10.84 | 10.26 | 9.54 | 9.52 | 9.04 | 10.48 | 10.40 | 10.55 | 9.53 |
| CHD9 | 5 | 10.55 | 17.17 | 6.08 | 7.86 | 9.58 | 9.83 | 8.48 | 8.19 | 10.86 | 11.40 | 13.86 | 6.97 | 9.71 | 8.34 | 11.13 |
| CHEK1 | 8 | 8.53 | 9.96 | 12.89 | 11.82 | 9.15 | 9.27 | 10.63 | 10.54 | 8.81 | 8.40 | 9.25 | 12.35 | 9.21 | 10.59 | 8.60 |

| | | | | | | | | | | | | | | | | |
|---|---|---|---|---|---|---|---|---|---|---|---|---|---|---|---|---|
| CHEK2 | 3 | 10.05 | 11.81 | 10.12 | 11.20 | 6.48 | 6.29 | 11.56 | 11.35 | 11.44 | 9.71 | 10.93 | 10.66 | 6.39 | 11.45 | 10.57 |
| CHERP | 19 | 9.41 | 10.34 | 11.02 | 10.54 | 8.87 | 8.95 | 11.17 | 10.28 | 10.01 | 9.41 | 9.87 | 10.78 | 8.91 | 10.73 | 9.71 |
| CHID1 | 2 | 9.34 | 8.73 | 7.93 | 9.17 | 12.23 | 13.61 | 11.84 | 12.20 | 7.27 | 7.69 | 9.03 | 8.55 | 12.92 | 12.02 | 7.48 |
| CHKA | 2 | 9.68 | 10.33 | 12.27 | 10.67 | 7.54 | 7.91 | 11.09 | 13.50 | 8.75 | 8.26 | 10.01 | 11.47 | 7.72 | 12.29 | 8.51 |
| CHM | 2 | 9.27 | 10.07 | 8.34 | 9.49 | 10.37 | 12.43 | 8.69 | 10.75 | 10.44 | 10.16 | 9.67 | 8.91 | 11.40 | 9.72 | 10.30 |
| CHML | 5 | 11.03 | 11.63 | 8.49 | 8.82 | 13.79 | 14.47 | 6.85 | 7.63 | 8.80 | 8.50 | 11.33 | 8.65 | 14.13 | 7.24 | 8.65 |
| CHMP1A | 12 | 11.78 | 11.93 | 8.76 | 8.70 | 10.87 | 11.02 | 8.90 | 8.11 | 10.63 | 9.29 | 11.86 | 8.73 | 10.94 | 8.50 | 9.96 |
| CHMP1B | 14 | 9.98 | 11.00 | 8.57 | 8.56 | 15.00 | 15.76 | 7.44 | 6.85 | 8.32 | 8.52 | 10.49 | 8.57 | 15.38 | 7.15 | 8.42 |
| CHMP2A | 19 | 7.97 | 10.08 | 9.66 | 9.83 | 11.37 | 12.30 | 10.60 | 10.14 | 9.07 | 8.99 | 9.02 | 9.74 | 11.83 | 10.37 | 9.03 |
| CHMP2B | 13 | 10.41 | 11.38 | 9.89 | 9.62 | 9.65 | 9.76 | 9.89 | 9.41 | 10.33 | 9.66 | 10.90 | 9.75 | 9.71 | 9.65 | 9.99 |
| CHMP3 | 2 | 8.78 | 11.22 | 9.07 | 11.51 | 11.32 | 12.55 | 7.56 | 7.94 | 9.83 | 10.21 | 10.00 | 10.29 | 11.93 | 7.75 | 10.02 |
| CHMP4A | 10 | 10.75 | 11.27 | 7.94 | 8.09 | 12.42 | 13.44 | 10.32 | 8.65 | 8.87 | 8.24 | 11.01 | 8.01 | 12.93 | 9.49 | 8.56 |
| CHMP4B | 18 | 9.18 | 10.35 | 11.21 | 10.82 | 10.59 | 10.49 | 10.08 | 9.87 | 9.17 | 8.24 | 9.77 | 11.01 | 10.54 | 9.98 | 8.70 |
| CHMP4C | 4 | 9.97 | 11.20 | 8.42 | 8.92 | 11.75 | 11.80 | 9.61 | 10.67 | 8.63 | 9.02 | 10.59 | 8.67 | 11.77 | 10.14 | 8.83 |
| CHMP5 | 6 | 9.48 | 11.88 | 9.87 | 10.34 | 9.77 | 11.28 | 7.64 | 6.79 | 11.10 | 11.84 | 10.68 | 10.11 | 10.52 | 7.22 | 11.47 |
| CHMP6 | 1 | 10.67 | 8.95 | 14.17 | 15.64 | 8.47 | 8.46 | 6.70 | 5.77 | 10.66 | 10.53 | 9.81 | 14.90 | 8.47 | 6.23 | 10.59 |
| CHMP7 | 12 | 8.99 | 9.99 | 10.01 | 10.76 | 8.82 | 9.51 | 12.15 | 11.81 | 9.09 | 8.86 | 9.49 | 10.39 | 9.16 | 11.98 | 8.98 |
| CHN1 | 4 | 10.59 | 11.51 | 9.55 | 9.69 | 11.02 | 11.60 | 6.82 | 6.70 | 11.58 | 10.94 | 11.05 | 9.62 | 11.31 | 6.76 | 11.26 |
| CHORDC1 | 32 | 10.08 | 10.94 | 11.44 | 10.13 | 8.03 | 8.08 | 10.77 | 11.57 | 9.67 | 9.29 | 10.51 | 10.79 | 8.05 | 11.17 | 9.48 |
| CHP1 | 10 | 8.33 | 8.47 | 13.52 | 12.54 | 8.61 | 9.00 | 11.55 | 9.79 | 9.14 | 9.05 | 8.40 | 13.03 | 8.80 | 10.67 | 9.09 |
| CHPF2 | 1 | 11.42 | 8.05 | 10.09 | 13.33 | 8.56 | 8.47 | 10.78 | 12.75 | 7.87 | 8.67 | 9.74 | 11.71 | 8.52 | 11.76 | 8.27 |
| CHRAC1 | 4 | 9.33 | 9.80 | 12.08 | 11.36 | 9.02 | 9.40 | 9.88 | 9.41 | 10.32 | 9.40 | 9.57 | 11.72 | 9.21 | 9.64 | 9.86 |
| CHRDL1 | 2 | 8.98 | 9.45 | 9.96 | 10.40 | 10.70 | 10.08 | 12.82 | 9.01 | 8.87 | 9.73 | 9.21 | 10.18 | 10.39 | 10.91 | 9.30 |
| CHST14 | 2 | 10.71 | 6.02 | 5.77 | 7.81 | 11.67 | 10.17 | 10.76 | 11.28 | 12.53 | 13.28 | 8.36 | 6.79 | 10.92 | 11.02 | 12.90 |
| CHTF18 | 6 | 9.73 | 13.51 | 8.94 | 9.06 | 9.57 | 9.79 | 9.40 | 9.70 | 9.89 | 10.40 | 11.62 | 9.00 | 9.68 | 9.55 | 10.15 |
| CHTF8 | 2 | 9.59 | 11.26 | 7.18 | 7.08 | 13.25 | 12.97 | 9.96 | 11.61 | 8.77 | 8.32 | 10.42 | 7.13 | 13.11 | 10.79 | 8.55 |
| CHTOP | 8 | 9.93 | 9.77 | 8.66 | 8.90 | 11.59 | 11.63 | 9.87 | 9.03 | 9.87 | 10.75 | 9.85 | 8.78 | 11.61 | 9.45 | 10.31 |
| CHUK | 7 | 9.87 | 10.06 | 9.63 | 9.43 | 9.82 | 9.34 | 10.88 | 10.35 | 10.48 | 10.14 | 9.97 | 9.53 | 9.58 | 10.62 | 10.31 |
| CHURC1 | 3 | 8.72 | 10.64 | 13.03 | 12.35 | 9.07 | 9.37 | 7.45 | 7.53 | 11.46 | 10.39 | 9.68 | 12.69 | 9.22 | 7.49 | 10.93 |
| CHURC1-FNT | 3 | 8.12 | 9.49 | 12.98 | 11.74 | 9.56 | 9.59 | 11.58 | 10.00 | 8.82 | 8.13 | 8.81 | 12.36 | 9.57 | 10.79 | 8.47 |
| CIAO1 | 1 | 3.00 | 2.58 | 5.47 | 0.00 | 4.78 | 2.79 | 3.82 | 71.46 | 2.98 | 3.11 | 2.79 | 2.73 | 3.79 | 37.64 | 3.05 |
| CIAPIN1 | 13 | 9.97 | 10.51 | 6.98 | 6.40 | 10.62 | 10.64 | 7.90 | 17.91 | 9.73 | 9.34 | 10.24 | 6.69 | 10.63 | 12.90 | 9.54 |
| CIC | 8 | 9.88 | 9.89 | 9.45 | 10.78 | 9.54 | 10.21 | 10.07 | 9.30 | 10.53 | 10.37 | 9.88 | 10.11 | 9.87 | 9.68 | 10.45 |
| CIR1 | 3 | 6.01 | 9.47 | 11.99 | 11.88 | 9.76 | 10.23 | 11.70 | 9.69 | 9.47 | 9.81 | 7.74 | 11.93 | 9.99 | 10.70 | 9.64 |
| CIRBP | 3 | 10.03 | 10.18 | 12.00 | 10.95 | 10.02 | 9.66 | 11.09 | 9.11 | 9.03 | 7.92 | 10.11 | 11.47 | 9.84 | 10.10 | 8.48 |
| CIRH1A | 15 | 9.14 | 9.81 | 10.02 | 10.37 | 7.68 | 8.49 | 11.01 | 11.54 | 10.82 | 11.12 | 9.47 | 10.19 | 8.09 | 11.27 | 10.97 |
| CISD1 | 8 | 11.20 | 6.87 | 7.31 | 9.44 | 11.10 | 9.73 | 12.38 | 11.75 | 9.34 | 10.87 | 9.03 | 8.38 | 10.42 | 12.07 | 10.10 |
| CISD2 | 3 | 10.78 | 6.00 | 7.42 | 10.07 | 9.97 | 8.11 | 10.44 | 14.59 | 10.33 | 12.29 | 8.39 | 8.74 | 9.04 | 12.52 | 11.31 |
| CIT | 20 | 8.62 | 9.38 | 10.81 | 10.40 | 10.51 | 10.09 | 9.22 | 8.21 | 11.66 | 11.11 | 9.00 | 10.61 | 10.30 | 8.71 | 11.38 |
| CIZ1 | 6 | 10.00 | 10.85 | 8.92 | 9.16 | 9.45 | 10.39 | 11.05 | 11.00 | 9.80 | 9.38 | 10.43 | 9.04 | 9.92 | 11.02 | 9.59 |
| CKAP2 | 10 | 8.78 | 10.17 | 10.24 | 10.32 | 13.59 | 13.47 | 9.19 | 8.14 | 7.88 | 8.21 | 9.48 | 10.28 | 13.53 | 8.67 | 8.04 |
| CKAP4 | 33 | 10.97 | 6.90 | 7.70 | 9.23 | 11.49 | 9.27 | 13.05 | 10.74 | 9.49 | 11.15 | 8.93 | 8.47 | 10.38 | 11.90 | 10.32 |

| | | | | | | | | | | | | | | | | |
|---|---|---|---|---|---|---|---|---|---|---|---|---|---|---|---|---|
| CKAP5 | 77 | 8.49 | 9.14 | 12.26 | 11.49 | 9.73 | 9.11 | 10.54 | 10.14 | 9.46 | 9.66 | 8.81 | 11.88 | 9.42 | 10.34 | 9.56 |
| CKB | 53 | 9.60 | 10.57 | 11.91 | 10.90 | 10.66 | 10.35 | 7.77 | 6.83 | 10.73 | 10.68 | 10.08 | 11.41 | 10.51 | 7.30 | 10.70 |
| CKS1B | 3 | 7.62 | 11.18 | 12.95 | 7.56 | 10.60 | 10.71 | 11.33 | 8.25 | 9.89 | 9.91 | 9.40 | 10.26 | 10.65 | 9.79 | 9.90 |
| CKS2 | 3 | 8.91 | 8.20 | 12.61 | 5.95 | 12.34 | 9.29 | 12.44 | 8.19 | 11.83 | 10.23 | 8.55 | 9.28 | 10.82 | 10.31 | 11.03 |
| CLASP1 | 27 | 9.31 | 10.01 | 10.89 | 10.87 | 10.02 | 10.50 | 9.00 | 8.80 | 10.47 | 10.12 | 9.66 | 10.88 | 10.26 | 8.90 | 10.30 |
| CLASP2 | 22 | 9.44 | 10.37 | 11.10 | 10.69 | 10.35 | 10.20 | 9.61 | 8.87 | 9.98 | 9.38 | 9.91 | 10.90 | 10.28 | 9.24 | 9.68 |
| CLASRP | 2 | 7.93 | 10.73 | 9.46 | 8.95 | 9.35 | 10.30 | 12.59 | 11.82 | 9.37 | 9.49 | 9.33 | 9.21 | 9.83 | 12.21 | 9.43 |
| CLCC1 | 2 | 16.84 | 8.60 | 5.98 | 9.41 | 8.96 | 7.69 | 9.57 | 9.66 | 9.89 | 13.39 | 12.72 | 7.70 | 8.33 | 9.61 | 11.64 |
| CLCN3 | 3 | 10.83 | 8.86 | 9.56 | 11.73 | 11.39 | 10.55 | 8.78 | 8.32 | 9.15 | 10.84 | 9.85 | 10.64 | 10.97 | 8.55 | 9.99 |
| CLCN7 | 1 | 11.90 | 7.29 | 7.79 | 12.03 | 11.79 | 11.00 | 8.11 | 8.41 | 9.70 | 11.97 | 9.60 | 9.91 | 11.40 | 8.26 | 10.84 |
| CLDN4 | 1 | 14.96 | 12.24 | 6.92 | 8.85 | 3.49 | 3.61 | 9.91 | 8.36 | 14.67 | 16.99 | 13.60 | 7.88 | 3.55 | 9.14 | 15.83 |
| CLEC16A | 5 | 10.40 | 11.18 | 10.63 | 10.52 | 10.97 | 10.68 | 9.65 | 8.32 | 9.10 | 8.53 | 10.79 | 10.58 | 10.83 | 8.98 | 8.82 |
| CLGN | 3 | 13.93 | 6.31 | 6.78 | 11.46 | 14.00 | 10.96 | 9.34 | 9.34 | 8.44 | 9.44 | 10.12 | 9.12 | 12.48 | 9.34 | 8.94 |
| CLIC1 | 32 | 6.72 | 6.18 | 10.43 | 7.59 | 7.28 | 7.23 | 7.82 | 31.31 | 8.10 | 7.34 | 6.45 | 9.01 | 7.25 | 19.57 | 7.72 |
| CLIC2 | 5 | 11.91 | 13.05 | 3.86 | 3.97 | 18.23 | 18.35 | 12.81 | 12.38 | 2.85 | 2.59 | 12.48 | 3.91 | 18.29 | 12.60 | 2.72 |
| CLIC3 | 5 | 10.20 | 11.87 | 5.82 | 4.80 | 20.74 | 19.83 | 3.90 | 3.28 | 10.10 | 9.46 | 11.03 | 5.31 | 20.29 | 3.59 | 9.78 |
| CLIC4 | 19 | 9.49 | 10.36 | 12.37 | 8.07 | 9.75 | 8.13 | 7.81 | 9.89 | 12.25 | 11.88 | 9.93 | 10.22 | 8.94 | 8.85 | 12.07 |
| CLINT1 | 18 | 9.59 | 10.08 | 11.57 | 10.44 | 9.58 | 9.20 | 10.07 | 9.30 | 10.40 | 9.76 | 9.83 | 11.01 | 9.39 | 9.69 | 10.08 |
| CLIP1 | 61 | 10.01 | 11.39 | 9.22 | 8.33 | 11.89 | 12.24 | 8.47 | 7.69 | 10.58 | 10.18 | 10.70 | 8.77 | 12.07 | 8.08 | 10.38 |
| CLIP2 | 2 | 9.84 | 10.14 | 6.97 | 7.23 | 6.59 | 6.96 | 14.58 | 14.59 | 11.88 | 11.24 | 9.99 | 7.10 | 6.77 | 14.58 | 11.56 |
| CLK1 | 4 | 8.35 | 8.31 | 10.85 | 10.95 | 8.38 | 8.70 | 12.43 | 11.03 | 10.32 | 10.67 | 8.33 | 10.90 | 8.54 | 11.73 | 10.50 |
| CLK3 | 2 | 9.35 | 10.30 | 8.33 | 10.66 | 9.00 | 9.66 | 9.50 | 10.29 | 11.08 | 11.82 | 9.83 | 9.50 | 9.33 | 9.89 | 11.45 |
| CLMN | 1 | 17.24 | 10.06 | 5.01 | 7.83 | 11.13 | 9.75 | 9.52 | 9.21 | 11.20 | 9.05 | 13.65 | 6.42 | 10.44 | 9.37 | 10.13 |
| CLN5 | 3 | 11.62 | 11.04 | 14.62 | 12.49 | 10.92 | 10.85 | 7.75 | 6.09 | 7.24 | 7.37 | 11.33 | 13.56 | 10.88 | 6.92 | 7.31 |
| CLN6 | 1 | 8.07 | 6.14 | 8.35 | 8.54 | 11.19 | 8.34 | 8.96 | 7.17 | 14.47 | 18.77 | 7.11 | 8.44 | 9.77 | 8.07 | 16.62 |
| CLNS1A | 17 | 8.37 | 9.17 | 10.50 | 10.37 | 10.17 | 10.45 | 10.02 | 10.27 | 10.35 | 10.33 | 8.77 | 10.43 | 10.31 | 10.14 | 10.34 |
| CLOCK | 1 | 8.45 | 10.11 | 11.19 | 11.27 | 8.94 | 11.01 | 9.55 | 10.31 | 10.47 | 8.71 | 9.28 | 11.23 | 9.97 | 9.93 | 9.59 |
| CLP1 | 4 | 9.94 | 9.62 | 9.76 | 9.72 | 9.27 | 9.69 | 11.37 | 11.25 | 10.06 | 9.32 | 9.78 | 9.74 | 9.48 | 11.31 | 9.69 |
| CLPB | 15 | 9.68 | 8.99 | 13.45 | 13.43 | 10.04 | 10.00 | 9.62 | 8.84 | 7.90 | 8.06 | 9.33 | 13.44 | 10.02 | 9.23 | 7.98 |
| CLPP | 14 | 9.13 | 8.08 | 10.93 | 11.91 | 10.88 | 10.28 | 10.72 | 9.69 | 9.06 | 9.31 | 8.60 | 11.42 | 10.58 | 10.21 | 9.18 |
| CLPTM1 | 7 | 13.73 | 6.79 | 7.74 | 11.84 | 9.52 | 9.29 | 9.22 | 8.81 | 10.59 | 12.45 | 10.26 | 9.79 | 9.41 | 9.02 | 11.52 |
| CLPTM1L | 9 | 14.24 | 7.14 | 5.88 | 8.77 | 10.92 | 10.01 | 10.42 | 9.09 | 10.36 | 13.18 | 10.69 | 7.32 | 10.46 | 9.75 | 11.77 |
| CLPX | 25 | 10.92 | 9.53 | 9.03 | 10.57 | 9.87 | 9.40 | 9.25 | 8.61 | 11.37 | 11.46 | 10.23 | 9.80 | 9.63 | 8.93 | 11.41 |
| CLSPN | 4 | 10.47 | 12.75 | 8.98 | 8.40 | 8.67 | 9.69 | 9.86 | 8.06 | 12.19 | 10.93 | 11.61 | 8.69 | 9.18 | 8.96 | 11.56 |
| CLSTN1 | 1 | 11.94 | 7.30 | 9.33 | 10.53 | 8.40 | 7.57 | 7.68 | 7.59 | 14.49 | 15.17 | 9.62 | 9.93 | 7.99 | 7.63 | 14.83 |
| CLTA | 17 | 9.11 | 9.53 | 12.69 | 11.25 | 8.33 | 8.21 | 7.22 | 12.73 | 10.71 | 10.21 | 9.32 | 11.97 | 8.27 | 9.97 | 10.46 |
| CLTB | 13 | 10.20 | 11.15 | 8.65 | 7.97 | 13.06 | 12.53 | 8.75 | 8.57 | 9.99 | 9.13 | 10.68 | 8.31 | 12.79 | 8.66 | 9.56 |
| CLTC | 130 | 9.67 | 9.07 | 10.78 | 10.06 | 9.73 | 9.02 | 7.58 | 13.09 | 10.70 | 10.30 | 9.37 | 10.42 | 9.37 | 10.33 | 10.50 |
| CLU | 15 | 12.28 | 11.08 | 5.20 | 5.71 | 17.55 | 17.44 | 7.90 | 7.55 | 7.61 | 7.67 | 11.68 | 5.46 | 17.50 | 7.73 | 7.64 |
| CLUH | 32 | 10.19 | 10.83 | 10.07 | 10.10 | 8.40 | 8.32 | 10.02 | 10.32 | 10.98 | 10.75 | 10.51 | 10.09 | 8.36 | 10.17 | 10.87 |
| CLYBL | 2 | 10.85 | 9.88 | 8.38 | 7.96 | 8.50 | 8.29 | 19.08 | 16.94 | 5.59 | 4.53 | 10.36 | 8.17 | 8.40 | 18.01 | 5.06 |
| CMAS | 9 | 9.38 | 10.14 | 9.01 | 8.11 | 14.71 | 14.57 | 9.40 | 8.49 | 8.08 | 8.11 | 9.76 | 8.56 | 14.64 | 8.95 | 8.09 |

| | | | | | | | | | | | | | | | | |
|---|---|---|---|---|---|---|---|---|---|---|---|---|---|---|---|---|
| CMBL | 13 | 12.43 | 12.50 | 7.31 | 6.45 | 15.45 | 14.97 | 6.17 | 13.08 | 6.09 | 5.55 | 12.46 | 6.88 | 15.21 | 9.62 | 5.82 |
| CMC2 | 1 | 11.96 | 10.67 | 10.60 | 9.31 | 10.18 | 10.99 | 10.42 | 7.88 | 8.34 | 9.64 | 11.32 | 9.96 | 10.59 | 9.15 | 8.99 |
| CMPK1 | 23 | 10.68 | 11.41 | 9.03 | 8.90 | 7.97 | 8.06 | 11.91 | 11.09 | 10.45 | 10.49 | 11.04 | 8.97 | 8.02 | 11.50 | 10.47 |
| CMSS1 | 3 | 8.65 | 9.37 | 12.98 | 12.20 | 8.47 | 8.24 | 13.07 | 11.84 | 7.58 | 7.59 | 9.01 | 12.59 | 8.36 | 12.46 | 7.59 |
| CMTM6 | 1 | 11.23 | 8.00 | 9.36 | 12.13 | 8.59 | 8.32 | 8.90 | 8.35 | 13.38 | 11.74 | 9.62 | 10.74 | 8.45 | 8.63 | 12.56 |
| CMTR1 | 11 | 9.68 | 9.99 | 11.19 | 11.26 | 8.51 | 9.06 | 10.09 | 9.73 | 10.47 | 10.02 | 9.84 | 11.23 | 8.78 | 9.91 | 10.24 |
| CNBP | 6 | 4.10 | 2.69 | 3.58 | 2.25 | 5.55 | 4.70 | 3.81 | 67.53 | 3.23 | 2.55 | 3.40 | 2.91 | 5.13 | 35.67 | 2.89 |
| CNDP2 | 15 | 9.35 | 10.57 | 13.73 | 13.13 | 12.19 | 12.31 | 5.12 | 4.90 | 9.41 | 9.30 | 9.96 | 13.43 | 12.25 | 5.01 | 9.35 |
| CNIH4 | 1 | 7.67 | 5.97 | 7.52 | 7.61 | 10.13 | 10.91 | 10.68 | 13.40 | 11.79 | 14.31 | 6.82 | 7.57 | 10.52 | 12.04 | 13.05 |
| CNN1 | 7 | 9.90 | 10.81 | 31.37 | 29.92 | 1.33 | 1.90 | 1.48 | 1.19 | 6.21 | 5.89 | 10.35 | 30.65 | 1.62 | 1.34 | 6.05 |
| CNN2 | 18 | 9.54 | 10.87 | 14.60 | 14.52 | 8.15 | 8.62 | 4.29 | 4.32 | 12.49 | 12.59 | 10.21 | 14.56 | 8.39 | 4.30 | 12.54 |
| CNN3 | 27 | 11.78 | 13.15 | 15.10 | 14.02 | 5.16 | 5.31 | 6.26 | 6.50 | 11.55 | 11.17 | 12.46 | 14.56 | 5.23 | 6.38 | 11.36 |
| CNNM3 | 3 | 9.61 | 9.57 | 11.83 | 10.92 | 8.58 | 8.39 | 9.40 | 8.97 | 11.39 | 11.34 | 9.59 | 11.37 | 8.49 | 9.18 | 11.37 |
| CNNM4 | 1 | 9.22 | 7.59 | 10.73 | 12.03 | 9.31 | 7.98 | 10.81 | 9.39 | 11.68 | 11.27 | 8.41 | 11.38 | 8.64 | 10.10 | 11.47 |
| CNOT1 | 47 | 9.84 | 10.03 | 8.74 | 9.33 | 10.39 | 10.47 | 10.69 | 10.88 | 9.83 | 9.81 | 9.93 | 9.03 | 10.43 | 10.78 | 9.82 |
| CNOT10 | 9 | 9.37 | 10.75 | 8.16 | 8.66 | 10.81 | 11.70 | 9.91 | 10.86 | 9.80 | 9.99 | 10.06 | 8.41 | 11.25 | 10.38 | 9.89 |
| CNOT11 | 4 | 9.81 | 10.22 | 9.19 | 9.34 | 10.36 | 11.28 | 11.23 | 10.03 | 9.71 | 8.82 | 10.02 | 9.26 | 10.82 | 10.63 | 9.27 |
| CNOT2 | 10 | 10.00 | 10.39 | 11.87 | 9.09 | 9.75 | 8.97 | 11.22 | 8.74 | 10.31 | 9.66 | 10.19 | 10.48 | 9.36 | 9.98 | 9.98 |
| CNOT3 | 8 | 9.68 | 10.43 | 11.00 | 10.86 | 9.33 | 9.68 | 10.95 | 9.95 | 9.26 | 8.86 | 10.06 | 10.93 | 9.51 | 10.45 | 9.06 |
| CNOT4 | 7 | 9.56 | 10.39 | 12.26 | 11.41 | 8.63 | 9.29 | 9.73 | 9.31 | 9.60 | 9.81 | 9.98 | 11.84 | 8.96 | 9.52 | 9.71 |
| CNOT6L | 2 | 9.89 | 9.42 | 9.38 | 9.00 | 9.44 | 9.50 | 11.93 | 11.15 | 10.22 | 10.08 | 9.65 | 9.19 | 9.47 | 11.54 | 10.15 |
| CNOT7 | 1 | 8.15 | 8.28 | 10.54 | 8.29 | 11.20 | 9.10 | 12.63 | 11.32 | 9.97 | 10.52 | 8.21 | 9.42 | 10.15 | 11.97 | 10.24 |
| CNOT8 | 2 | 10.50 | 11.83 | 8.89 | 7.60 | 12.00 | 11.18 | 9.16 | 9.49 | 10.27 | 9.07 | 11.17 | 8.25 | 11.59 | 9.32 | 9.67 |
| CNP | 23 | 11.42 | 9.92 | 10.54 | 11.48 | 9.25 | 9.22 | 8.19 | 7.31 | 10.98 | 11.70 | 10.67 | 11.01 | 9.23 | 7.75 | 11.34 |
| CNPY2 | 9 | 9.62 | 9.37 | 8.47 | 8.24 | 12.87 | 12.61 | 11.45 | 8.94 | 9.37 | 9.08 | 9.50 | 8.35 | 12.74 | 10.19 | 9.22 |
| CNPY3 | 9 | 8.97 | 8.49 | 10.93 | 11.64 | 12.53 | 12.94 | 10.02 | 8.82 | 7.74 | 7.93 | 8.73 | 11.29 | 12.73 | 9.42 | 7.83 |
| CNPY4 | 3 | 11.46 | 10.04 | 8.12 | 7.27 | 10.26 | 9.81 | 12.78 | 7.60 | 11.72 | 10.93 | 10.75 | 7.70 | 10.04 | 10.19 | 11.32 |
| CNTLN | 3 | 13.29 | 11.03 | 8.65 | 8.95 | 10.24 | 11.17 | 8.70 | 8.69 | 9.98 | 9.29 | 12.16 | 8.80 | 10.70 | 8.70 | 9.64 |
| CNTNAP1 | 2 | 15.20 | 9.30 | 5.73 | 7.58 | 10.43 | 10.77 | 7.18 | 6.75 | 13.00 | 14.06 | 12.25 | 6.65 | 10.60 | 6.97 | 13.53 |
| COA1 | 1 | 10.58 | 7.72 | 8.75 | 17.24 | 4.68 | 4.15 | 15.10 | 16.78 | 6.08 | 8.91 | 9.15 | 13.00 | 4.42 | 15.94 | 7.50 |
| COA3 | 5 | 13.08 | 8.04 | 7.85 | 11.69 | 9.40 | 8.89 | 8.59 | 8.09 | 11.40 | 12.98 | 10.56 | 9.77 | 9.14 | 8.34 | 12.19 |
| COA6 | 1 | 9.08 | 12.82 | 7.27 | 8.97 | 12.34 | 15.42 | 7.78 | 9.50 | 7.45 | 9.38 | 10.95 | 8.12 | 13.88 | 8.64 | 8.42 |
| COASY | 11 | 9.93 | 10.87 | 8.86 | 8.75 | 10.97 | 11.40 | 7.11 | 6.71 | 12.91 | 12.49 | 10.40 | 8.80 | 11.19 | 6.91 | 12.70 |
| COBL | 20 | 13.30 | 14.63 | 8.77 | 8.36 | 11.36 | 11.92 | 5.79 | 5.23 | 10.61 | 10.03 | 13.97 | 8.56 | 11.64 | 5.51 | 10.32 |
| COBLL1 | 6 | 12.75 | 14.67 | 6.71 | 6.21 | 14.45 | 13.61 | 6.03 | 5.94 | 10.10 | 9.53 | 13.71 | 6.46 | 14.03 | 5.99 | 9.81 |
| COG1 | 9 | 10.73 | 10.66 | 9.67 | 10.59 | 10.97 | 10.80 | 8.87 | 8.95 | 9.50 | 9.27 | 10.69 | 10.13 | 10.88 | 8.91 | 9.39 |
| COG2 | 9 | 10.75 | 11.09 | 11.00 | 10.36 | 11.41 | 10.95 | 8.63 | 7.89 | 9.16 | 8.77 | 10.92 | 10.68 | 11.18 | 8.26 | 8.97 |
| COG3 | 10 | 10.53 | 11.60 | 10.10 | 10.53 | 10.96 | 10.89 | 8.90 | 7.66 | 9.55 | 9.29 | 11.07 | 10.31 | 10.93 | 8.28 | 9.42 |
| COG4 | 7 | 12.20 | 11.05 | 8.92 | 9.17 | 11.37 | 11.62 | 7.65 | 7.55 | 10.56 | 9.91 | 11.63 | 9.04 | 11.50 | 7.60 | 10.24 |
| COG5 | 10 | 9.59 | 10.67 | 11.03 | 11.50 | 9.42 | 9.58 | 9.84 | 9.02 | 9.78 | 9.56 | 10.13 | 11.27 | 9.50 | 9.43 | 9.67 |
| COG6 | 7 | 10.60 | 11.25 | 10.11 | 10.83 | 11.20 | 11.83 | 8.77 | 8.66 | 8.33 | 8.40 | 10.93 | 10.47 | 11.52 | 8.72 | 8.37 |
| COG7 | 9 | 9.74 | 10.85 | 10.74 | 10.44 | 10.18 | 10.57 | 9.17 | 9.53 | 9.29 | 9.49 | 10.30 | 10.59 | 10.37 | 9.35 | 9.39 |

| | | | | | | | | | | | | | | | | |
|---|---|---|---|---|---|---|---|---|---|---|---|---|---|---|---|---|
| COG8 | 3 | 9.83 | 11.14 | 9.39 | 9.75 | 11.22 | 12.19 | 9.11 | 9.40 | 8.83 | 9.14 | 10.49 | 9.57 | 11.70 | 9.26 | 8.98 |
| COIL | 9 | 9.04 | 10.83 | 13.18 | 12.88 | 8.15 | 8.63 | 9.20 | 8.83 | 9.80 | 9.46 | 9.94 | 13.03 | 8.39 | 9.01 | 9.63 |
| COL12A1 | 17 | 13.07 | 9.90 | 8.93 | 10.41 | 7.62 | 7.09 | 10.63 | 10.42 | 10.65 | 11.28 | 11.48 | 9.67 | 7.36 | 10.52 | 10.96 |
| COL1A1 | 5 | 7.76 | 6.56 | 23.71 | 26.23 | 2.88 | 2.77 | 3.49 | 2.53 | 11.81 | 12.24 | 7.16 | 24.97 | 2.83 | 3.01 | 12.03 |
| COL3A1 | 4 | 12.96 | 11.42 | 6.43 | 6.60 | 5.41 | 5.32 | 11.61 | 10.68 | 14.35 | 15.23 | 12.19 | 6.51 | 5.37 | 11.15 | 14.79 |
| COL4A2 | 3 | 10.29 | 7.62 | 14.80 | 17.46 | 6.38 | 6.07 | 11.19 | 9.62 | 7.60 | 8.96 | 8.95 | 16.13 | 6.23 | 10.41 | 8.28 |
| COL4A3BP | 11 | 9.93 | 10.40 | 11.34 | 8.95 | 9.34 | 8.96 | 11.07 | 9.42 | 10.30 | 10.29 | 10.16 | 10.14 | 9.15 | 10.25 | 10.30 |
| COL5A1 | 17 | 11.45 | 8.63 | 10.09 | 10.82 | 6.97 | 6.57 | 9.95 | 9.03 | 12.51 | 13.97 | 10.04 | 10.45 | 6.77 | 9.49 | 13.24 |
| COL5A2 | 1 | 7.78 | 6.67 | 18.06 | 20.96 | 4.24 | 4.58 | 6.63 | 5.56 | 13.72 | 11.81 | 7.23 | 19.51 | 4.41 | 6.09 | 12.76 |
| COL7A1 | 9 | 14.01 | 10.83 | 6.62 | 7.38 | 7.59 | 6.80 | 12.04 | 11.30 | 11.00 | 12.41 | 12.42 | 7.00 | 7.20 | 11.67 | 11.71 |
| COLGALT1 | 27 | 8.74 | 6.50 | 8.96 | 9.35 | 10.21 | 9.49 | 14.14 | 13.23 | 9.36 | 10.03 | 7.62 | 9.16 | 9.85 | 13.69 | 9.69 |
| COMMD1 | 2 | 9.75 | 11.42 | 12.83 | 13.55 | 8.64 | 9.87 | 8.05 | 8.20 | 8.89 | 8.81 | 10.59 | 13.19 | 9.25 | 8.13 | 8.85 |
| COMMD10 | 6 | 9.49 | 10.68 | 10.79 | 10.60 | 12.10 | 12.72 | 9.46 | 8.78 | 7.95 | 7.43 | 10.08 | 10.69 | 12.41 | 9.12 | 7.69 |
| COMMD2 | 2 | 8.99 | 9.19 | 11.77 | 9.69 | 10.38 | 11.80 | 8.85 | 7.85 | 11.31 | 10.16 | 9.09 | 10.73 | 11.09 | 8.35 | 10.74 |
| COMMD3-BM | 4 | 8.86 | 9.87 | 11.40 | 10.97 | 10.58 | 11.01 | 10.52 | 10.05 | 8.42 | 8.31 | 9.37 | 11.18 | 10.80 | 10.29 | 8.36 |
| COMMD4 | 5 | 9.17 | 10.68 | 12.17 | 11.28 | 11.09 | 10.60 | 8.33 | 8.32 | 9.27 | 9.09 | 9.92 | 11.73 | 10.84 | 8.32 | 9.18 |
| COMMD5 | 3 | 8.80 | 10.63 | 8.84 | 9.46 | 10.93 | 13.15 | 10.65 | 10.60 | 8.03 | 8.91 | 9.71 | 9.15 | 12.04 | 10.62 | 8.47 |
| COMMD6 | 4 | 10.07 | 9.87 | 12.44 | 12.10 | 9.98 | 10.03 | 9.08 | 7.44 | 9.84 | 9.16 | 9.97 | 12.27 | 10.00 | 8.26 | 9.50 |
| COMMD7 | 5 | 9.46 | 10.76 | 10.89 | 10.75 | 13.07 | 13.35 | 8.44 | 8.07 | 8.35 | 6.85 | 10.11 | 10.82 | 13.21 | 8.26 | 7.60 |
| COMMD8 | 3 | 9.66 | 10.09 | 11.80 | 11.93 | 10.14 | 10.01 | 9.02 | 8.78 | 9.13 | 9.44 | 9.88 | 11.86 | 10.08 | 8.90 | 9.29 |
| COMMD9 | 5 | 10.68 | 10.64 | 13.18 | 13.20 | 10.72 | 10.37 | 6.73 | 5.93 | 9.46 | 9.10 | 10.66 | 13.19 | 10.54 | 6.33 | 9.28 |
| COMT | 13 | 11.42 | 7.93 | 8.73 | 10.35 | 5.81 | 5.25 | 10.98 | 13.43 | 11.98 | 14.11 | 9.67 | 9.54 | 5.53 | 12.21 | 13.05 |
| COPA | 65 | 9.53 | 9.67 | 9.84 | 9.36 | 10.45 | 10.53 | 9.58 | 11.33 | 9.87 | 9.84 | 9.60 | 9.60 | 10.49 | 10.46 | 9.86 |
| COPB1 | 51 | 9.58 | 9.73 | 10.27 | 10.24 | 10.16 | 10.16 | 9.94 | 9.77 | 10.05 | 10.09 | 9.65 | 10.26 | 10.16 | 9.86 | 10.07 |
| COPB2 | 53 | 9.66 | 10.24 | 10.58 | 10.19 | 10.12 | 9.81 | 9.79 | 9.85 | 9.94 | 9.84 | 9.95 | 10.38 | 9.96 | 9.82 | 9.89 |
| COPE | 12 | 10.12 | 10.40 | 10.49 | 10.68 | 9.83 | 10.61 | 9.31 | 9.70 | 9.35 | 9.53 | 10.26 | 10.58 | 10.22 | 9.50 | 9.44 |
| COPG1 | 30 | 8.48 | 8.42 | 9.44 | 7.81 | 9.53 | 9.18 | 8.70 | 19.69 | 9.69 | 9.07 | 8.45 | 8.62 | 9.35 | 14.20 | 9.38 |
| COPG2 | 11 | 10.40 | 11.39 | 11.20 | 11.24 | 11.70 | 11.80 | 10.94 | 11.05 | 5.18 | 5.11 | 10.89 | 11.22 | 11.75 | 10.99 | 5.14 |
| COPRS | 4 | 10.31 | 11.86 | 12.85 | 10.15 | 9.19 | 9.29 | 8.21 | 4.74 | 12.56 | 10.84 | 11.09 | 11.50 | 9.24 | 6.47 | 11.70 |
| COPS2 | 24 | 10.12 | 10.44 | 11.26 | 10.19 | 10.05 | 9.64 | 9.15 | 8.06 | 10.84 | 10.25 | 10.28 | 10.73 | 9.85 | 8.60 | 10.54 |
| COPS3 | 21 | 9.55 | 10.55 | 11.57 | 10.63 | 9.14 | 9.33 | 8.51 | 8.81 | 10.65 | 11.26 | 10.05 | 11.10 | 9.24 | 8.66 | 10.95 |
| COPS4 | 27 | 9.44 | 10.65 | 12.41 | 9.41 | 11.04 | 10.34 | 8.79 | 7.32 | 10.31 | 10.29 | 10.05 | 10.91 | 10.69 | 8.06 | 10.30 |
| COPS5 | 22 | 9.81 | 11.16 | 10.71 | 10.33 | 10.31 | 10.65 | 8.95 | 8.63 | 9.78 | 9.67 | 10.48 | 10.52 | 10.48 | 8.79 | 9.73 |
| COPS6 | 11 | 9.30 | 9.07 | 10.28 | 8.53 | 9.42 | 9.33 | 7.60 | 18.35 | 9.67 | 8.44 | 9.18 | 9.41 | 9.38 | 12.98 | 9.06 |
| COPS7A | 12 | 10.80 | 11.81 | 9.48 | 9.70 | 10.90 | 11.71 | 7.79 | 7.47 | 10.37 | 9.96 | 11.31 | 9.59 | 11.31 | 7.63 | 10.16 |
| COPS7B | 7 | 9.33 | 10.45 | 12.60 | 11.57 | 9.08 | 9.61 | 9.19 | 8.79 | 10.11 | 9.26 | 9.89 | 12.09 | 9.35 | 8.99 | 9.69 |
| COPS8 | 12 | 9.98 | 10.89 | 11.80 | 11.48 | 9.21 | 9.52 | 9.23 | 7.75 | 10.30 | 9.84 | 10.43 | 11.64 | 9.36 | 8.49 | 10.07 |
| COPZ1 | 5 | 9.58 | 10.94 | 11.59 | 11.14 | 9.61 | 10.33 | 8.98 | 8.53 | 9.78 | 9.52 | 10.26 | 11.36 | 9.97 | 8.75 | 9.65 |
| COQ3 | 1 | 11.52 | 9.93 | 8.81 | 9.91 | 9.51 | 11.38 | 11.74 | 9.89 | 8.69 | 8.63 | 10.72 | 9.36 | 10.44 | 10.81 | 8.66 |
| COQ5 | 2 | 12.12 | 11.15 | 7.72 | 8.80 | 10.52 | 10.81 | 12.27 | 11.17 | 7.45 | 7.98 | 11.64 | 8.26 | 10.67 | 11.72 | 7.71 |
| COQ6 | 3 | 11.37 | 10.27 | 8.22 | 9.86 | 9.64 | 9.19 | 11.04 | 10.19 | 10.33 | 9.91 | 10.82 | 9.04 | 9.41 | 10.61 | 10.12 |
| COQ7 | 1 | 11.97 | 11.09 | 8.56 | 9.47 | 11.33 | 10.30 | 10.74 | 11.54 | 7.26 | 7.76 | 11.53 | 9.01 | 10.81 | 11.14 | 7.51 |

| | | | | | | | | | | | | | | | | |
|---|---|---|---|---|---|---|---|---|---|---|---|---|---|---|---|---|
| COQ9 | 6 | 11.79 | 10.53 | 6.91 | 6.77 | 12.60 | 12.52 | 9.12 | 9.04 | 10.39 | 10.31 | 11.16 | 6.84 | 12.56 | 9.08 | 10.35 |
| CORO1B | 14 | 9.70 | 10.23 | 14.03 | 13.60 | 10.44 | 10.32 | 6.86 | 6.32 | 9.63 | 8.88 | 9.96 | 13.81 | 10.38 | 6.59 | 9.26 |
| CORO1C | 27 | 7.88 | 8.42 | 17.41 | 14.89 | 7.68 | 7.54 | 7.47 | 7.67 | 10.75 | 10.28 | 8.15 | 16.15 | 7.61 | 7.57 | 10.52 |
| CORO2A | 8 | 11.85 | 12.28 | 9.18 | 8.27 | 7.81 | 7.99 | 7.23 | 7.32 | 14.74 | 13.31 | 12.06 | 8.73 | 7.90 | 7.28 | 14.03 |
| CORO7 | 10 | 9.77 | 9.89 | 10.49 | 10.92 | 11.28 | 11.00 | 9.11 | 8.13 | 9.71 | 9.70 | 9.83 | 10.70 | 11.14 | 8.62 | 9.71 |
| COTL1 | 23 | 9.51 | 10.28 | 13.93 | 12.42 | 8.59 | 8.76 | 5.34 | 5.34 | 13.14 | 12.69 | 9.89 | 13.17 | 8.68 | 5.34 | 12.91 |
| COX10 | 1 | 13.12 | 7.88 | 8.19 | 11.93 | 7.48 | 7.71 | 8.20 | 8.14 | 11.78 | 15.57 | 10.50 | 10.06 | 7.59 | 8.17 | 13.68 |
| COX15 | 5 | 15.02 | 6.69 | 7.00 | 9.92 | 8.53 | 7.90 | 8.23 | 8.82 | 12.75 | 15.16 | 10.85 | 8.46 | 8.22 | 8.52 | 13.95 |
| COX16 | 1 | 10.08 | 10.20 | 10.62 | 10.15 | 9.95 | 11.55 | 9.72 | 7.54 | 10.45 | 9.74 | 10.14 | 10.38 | 10.75 | 8.63 | 10.09 |
| COX19 | 1 | 9.71 | 11.76 | 12.37 | 11.27 | 8.76 | 9.39 | 8.70 | 8.79 | 9.87 | 9.39 | 10.73 | 11.82 | 9.07 | 8.74 | 9.63 |
| COX4I1 | 11 | 13.75 | 6.80 | 6.43 | 9.33 | 10.65 | 9.12 | 10.85 | 9.83 | 11.22 | 12.03 | 10.28 | 7.88 | 9.88 | 10.34 | 11.62 |
| COX5A | 11 | 12.62 | 9.68 | 7.83 | 9.90 | 12.22 | 11.65 | 10.35 | 4.29 | 10.60 | 10.86 | 11.15 | 8.86 | 11.93 | 7.32 | 10.73 |
| COX5B | 9 | 11.62 | 8.69 | 9.24 | 10.55 | 11.09 | 10.77 | 11.49 | 8.05 | 9.40 | 9.12 | 10.16 | 9.89 | 10.93 | 9.77 | 9.26 |
| COX6A1 | 1 | 7.77 | 4.83 | 13.05 | 3.31 | 13.22 | 11.93 | 12.63 | 12.67 | 8.26 | 12.34 | 6.30 | 8.18 | 12.57 | 12.65 | 10.30 |
| COX6B1 | 5 | 10.03 | 8.96 | 9.16 | 8.99 | 10.78 | 11.74 | 12.93 | 12.03 | 8.09 | 7.29 | 9.49 | 9.07 | 11.26 | 12.48 | 7.69 |
| COX6C | 3 | 13.77 | 6.57 | 6.70 | 10.38 | 9.77 | 8.91 | 12.59 | 11.13 | 9.82 | 10.35 | 10.17 | 8.54 | 9.34 | 11.86 | 10.09 |
| COX7A2 | 2 | 11.03 | 9.19 | 8.51 | 9.98 | 10.12 | 11.53 | 10.50 | 10.13 | 8.79 | 10.23 | 10.11 | 9.24 | 10.83 | 10.31 | 9.51 |
| COX7A2L | 1 | 16.11 | 9.61 | 9.72 | 10.25 | 12.24 | 10.14 | 11.04 | 7.57 | 6.49 | 6.83 | 12.86 | 9.99 | 11.19 | 9.31 | 6.66 |
| COX7B | 1 | 17.66 | 5.80 | 4.66 | 8.52 | 11.77 | 10.25 | 11.04 | 8.69 | 9.49 | 12.11 | 11.73 | 6.59 | 11.01 | 9.87 | 10.80 |
| COX7C | 4 | 11.98 | 7.97 | 8.63 | 6.36 | 11.59 | 8.15 | 11.37 | 12.03 | 12.20 | 9.72 | 9.97 | 7.50 | 9.87 | 11.70 | 10.96 |
| CPD | 5 | 12.91 | 9.67 | 7.89 | 9.73 | 7.85 | 7.96 | 11.98 | 11.97 | 9.79 | 10.26 | 11.29 | 8.81 | 7.90 | 11.98 | 10.03 |
| CPEB3 | 1 | 9.18 | 9.22 | 9.08 | 11.91 | 10.70 | 10.83 | 9.28 | 9.90 | 10.91 | 8.98 | 9.20 | 10.50 | 10.77 | 9.59 | 9.95 |
| CPEB4 | 7 | 9.73 | 10.68 | 7.66 | 7.41 | 15.07 | 14.44 | 8.95 | 8.12 | 8.87 | 9.05 | 10.21 | 7.53 | 14.76 | 8.54 | 8.96 |
| CPM | 8 | 9.35 | 7.61 | 1.39 | 1.07 | 9.94 | 10.59 | 28.10 | 25.52 | 3.25 | 3.18 | 8.48 | 1.23 | 10.26 | 26.81 | 3.22 |
| CPNE1 | 16 | 9.17 | 9.90 | 8.22 | 7.79 | 11.01 | 10.98 | 9.59 | 14.59 | 9.61 | 9.14 | 9.54 | 8.00 | 10.99 | 12.09 | 9.38 |
| CPNE2 | 1 | 6.34 | 15.18 | 6.66 | 6.29 | 9.08 | 24.02 | 5.33 | 4.53 | 12.72 | 9.84 | 10.76 | 6.48 | 16.55 | 4.93 | 11.28 |
| CPNE3 | 19 | 8.30 | 8.40 | 7.98 | 6.76 | 9.14 | 9.36 | 11.90 | 22.13 | 8.38 | 7.66 | 8.35 | 7.37 | 9.25 | 17.01 | 8.02 |
| CPNE8 | 11 | 10.77 | 11.29 | 8.31 | 8.09 | 10.58 | 11.35 | 10.35 | 9.31 | 9.89 | 10.05 | 11.03 | 8.20 | 10.97 | 9.83 | 9.97 |
| CPOX | 8 | 9.02 | 8.46 | 11.66 | 10.07 | 7.75 | 7.98 | 11.53 | 11.37 | 11.14 | 11.02 | 8.74 | 10.87 | 7.87 | 11.45 | 11.08 |
| CPPED1 | 6 | 9.78 | 11.13 | 13.52 | 14.52 | 8.79 | 10.16 | 8.60 | 7.44 | 8.13 | 7.93 | 10.46 | 14.02 | 9.47 | 8.02 | 8.03 |
| CPS1 | 140 | 14.85 | 11.46 | 4.33 | 5.07 | 16.18 | 14.35 | 8.74 | 10.11 | 7.31 | 7.60 | 13.16 | 4.70 | 15.26 | 9.42 | 7.45 |
| CPSF1 | 23 | 9.42 | 9.87 | 10.53 | 10.79 | 9.03 | 8.93 | 10.90 | 11.63 | 9.69 | 9.20 | 9.65 | 10.66 | 8.98 | 11.27 | 9.45 |
| CPSF2 | 23 | 9.90 | 10.57 | 9.91 | 10.11 | 9.96 | 10.49 | 10.94 | 10.96 | 8.63 | 8.54 | 10.24 | 10.01 | 10.23 | 10.95 | 8.58 |
| CPSF3 | 10 | 10.61 | 9.96 | 10.08 | 8.96 | 10.63 | 11.00 | 10.58 | 10.07 | 9.42 | 8.69 | 10.29 | 9.52 | 10.82 | 10.33 | 9.05 |
| CPSF3L | 4 | 9.59 | 10.64 | 9.90 | 9.42 | 10.04 | 9.93 | 11.39 | 10.51 | 9.39 | 9.20 | 10.11 | 9.66 | 9.99 | 10.95 | 9.29 |
| CPSF4 | 2 | 9.16 | 10.08 | 14.70 | 9.16 | 10.02 | 9.97 | 11.92 | 9.63 | 7.86 | 7.50 | 9.62 | 11.93 | 10.00 | 10.78 | 7.68 |
| CPSF6 | 11 | 9.77 | 10.48 | 9.99 | 8.95 | 9.94 | 10.45 | 11.65 | 10.39 | 9.49 | 8.90 | 10.13 | 9.47 | 10.19 | 11.02 | 9.19 |
| CPSF7 | 11 | 9.52 | 11.56 | 7.92 | 7.74 | 11.37 | 11.68 | 10.47 | 9.49 | 10.20 | 10.04 | 10.54 | 7.83 | 11.53 | 9.98 | 10.12 |
| CPT1A | 14 | 12.40 | 7.21 | 6.30 | 8.81 | 11.45 | 10.03 | 7.83 | 7.81 | 12.60 | 15.57 | 9.80 | 7.55 | 10.74 | 7.82 | 14.09 |
| CPT2 | 11 | 13.02 | 10.38 | 7.19 | 8.66 | 10.24 | 9.66 | 10.82 | 9.95 | 9.51 | 10.58 | 11.70 | 7.93 | 9.95 | 10.38 | 10.05 |
| CPZ | 1 | 10.37 | 11.87 | 6.10 | 15.37 | 8.49 | 11.52 | 8.45 | 13.20 | 7.54 | 7.09 | 11.12 | 10.73 | 10.00 | 10.83 | 7.32 |
| CRABP2 | 8 | 6.98 | 8.62 | 5.13 | 4.73 | 3.06 | 3.44 | 10.78 | 15.88 | 21.13 | 20.24 | 7.80 | 4.93 | 3.25 | 13.33 | 20.68 |

| | | | | | | | | | | | | | | | | |
|---|---|---|---|---|---|---|---|---|---|---|---|---|---|---|---|---|
| CRADD | 2 | 9.80 | 10.32 | 11.59 | 10.23 | 11.64 | 11.03 | 10.41 | 9.13 | 7.34 | 8.51 | 10.06 | 10.91 | 11.33 | 9.77 | 7.93 |
| CRAT | 5 | 12.49 | 10.44 | 6.04 | 6.04 | 12.97 | 11.61 | 9.37 | 9.16 | 10.41 | 11.47 | 11.46 | 6.04 | 12.29 | 9.27 | 10.94 |
| CRBN | 1 | 9.24 | 12.90 | 10.16 | 7.73 | 11.79 | 12.18 | 8.94 | 10.78 | 7.05 | 9.22 | 11.07 | 8.95 | 11.99 | 9.86 | 8.13 |
| CRCP | 1 | 8.54 | 11.43 | 12.87 | 11.87 | 9.24 | 9.68 | 11.41 | 5.68 | 9.93 | 9.34 | 9.99 | 12.37 | 9.46 | 8.55 | 9.64 |
| CREB1 | 3 | 9.41 | 10.73 | 9.88 | 9.09 | 10.50 | 10.37 | 11.99 | 10.42 | 9.14 | 8.48 | 10.07 | 9.49 | 10.43 | 11.20 | 8.81 |
| CREBBP | 11 | 8.38 | 9.86 | 12.83 | 11.62 | 9.60 | 9.59 | 9.97 | 9.60 | 9.07 | 9.48 | 9.12 | 12.23 | 9.60 | 9.78 | 9.27 |
| CRELD2 | 3 | 10.63 | 10.56 | 9.53 | 9.80 | 13.16 | 12.44 | 8.04 | 7.39 | 9.24 | 9.22 | 10.59 | 9.66 | 12.80 | 7.71 | 9.23 |
| CRIP1 | 5 | 15.77 | 15.77 | 4.81 | 5.98 | 10.74 | 11.14 | 5.23 | 5.69 | 12.83 | 12.04 | 15.77 | 5.39 | 10.94 | 5.46 | 12.44 |
| CRIP2 | 10 | 11.25 | 10.72 | 9.67 | 7.11 | 12.40 | 11.16 | 2.96 | 7.86 | 13.96 | 12.91 | 10.99 | 8.39 | 11.78 | 5.41 | 13.44 |
| CRIPT | 2 | 9.31 | 9.85 | 10.63 | 10.39 | 10.66 | 11.40 | 10.70 | 10.46 | 8.81 | 7.80 | 9.58 | 10.51 | 11.03 | 10.58 | 8.31 |
| CRK | 16 | 9.61 | 10.75 | 12.07 | 11.54 | 8.50 | 8.98 | 8.48 | 7.48 | 11.53 | 11.06 | 10.18 | 11.80 | 8.74 | 7.98 | 11.29 |
| CRKL | 13 | 9.76 | 10.12 | 13.16 | 11.72 | 7.36 | 7.22 | 10.42 | 9.76 | 10.58 | 9.91 | 9.94 | 12.44 | 7.29 | 10.09 | 10.25 |
| CRLF1 | 4 | 10.35 | 9.33 | 5.08 | 6.21 | 14.55 | 15.57 | 6.85 | 7.63 | 12.38 | 12.05 | 9.84 | 5.65 | 15.06 | 7.24 | 12.22 |
| CRLF3 | 6 | 10.09 | 10.85 | 12.39 | 12.85 | 7.43 | 7.61 | 8.43 | 8.14 | 11.02 | 11.19 | 10.47 | 12.62 | 7.52 | 8.28 | 11.11 |
| CRMP1 | 20 | 8.29 | 9.21 | 11.88 | 11.74 | 9.49 | 9.50 | 8.45 | 7.81 | 12.11 | 11.52 | 8.75 | 11.81 | 9.50 | 8.13 | 11.81 |
| CRNKL1 | 9 | 9.78 | 9.79 | 10.56 | 9.99 | 10.82 | 11.07 | 9.20 | 8.24 | 10.75 | 9.81 | 9.78 | 10.27 | 10.94 | 8.72 | 10.28 |
| CROCC | 6 | 9.99 | 10.49 | 11.96 | 11.15 | 9.44 | 9.29 | 8.27 | 7.44 | 11.57 | 10.40 | 10.24 | 11.56 | 9.36 | 7.86 | 10.98 |
| CRTAP | 16 | 9.04 | 8.18 | 8.12 | 8.53 | 10.10 | 10.10 | 12.40 | 11.83 | 10.80 | 10.91 | 8.61 | 8.33 | 10.10 | 12.11 | 10.85 |
| CRTC1 | 1 | 9.63 | 10.74 | 10.84 | 11.33 | 8.08 | 9.10 | 11.42 | 12.03 | 8.20 | 8.62 | 10.19 | 11.08 | 8.59 | 11.73 | 8.41 |
| CRTC3 | 2 | 12.93 | 10.23 | 7.70 | 7.50 | 11.12 | 11.28 | 10.69 | 9.66 | 10.42 | 8.47 | 11.58 | 7.60 | 11.20 | 10.18 | 9.44 |
| CRYAB | 8 | 14.01 | 16.29 | 12.17 | 10.61 | 10.40 | 9.84 | 0.74 | 0.46 | 12.77 | 12.72 | 15.15 | 11.39 | 10.12 | 0.60 | 12.75 |
| CRYBG3 | 2 | 11.06 | 15.01 | 9.17 | 10.25 | 9.17 | 7.88 | 7.32 | 6.73 | 12.46 | 10.95 | 13.04 | 9.71 | 8.53 | 7.02 | 11.71 |
| CRYZ | 16 | 10.72 | 10.54 | 9.67 | 8.82 | 6.98 | 6.55 | 10.80 | 9.40 | 13.30 | 13.24 | 10.63 | 9.24 | 6.76 | 10.10 | 13.27 |
| CRYZL1 | 9 | 10.85 | 11.88 | 7.30 | 7.39 | 12.11 | 12.39 | 8.43 | 7.83 | 10.95 | 10.86 | 11.37 | 7.35 | 12.25 | 8.13 | 10.91 |
| CS | 44 | 10.87 | 8.89 | 10.18 | 10.67 | 10.50 | 9.53 | 9.91 | 10.62 | 9.32 | 9.52 | 9.88 | 10.42 | 10.01 | 10.26 | 9.42 |
| CSDC2 | 1 | 12.78 | 15.20 | 15.83 | 14.91 | 4.85 | 4.30 | 4.94 | 4.41 | 11.16 | 11.61 | 13.99 | 15.37 | 4.58 | 4.67 | 11.39 |
| CSDE1 | 49 | 9.66 | 11.09 | 10.69 | 10.11 | 8.67 | 8.88 | 10.28 | 11.72 | 9.62 | 9.28 | 10.37 | 10.40 | 8.78 | 11.00 | 9.45 |
| CSE1L | 67 | 9.56 | 9.28 | 10.13 | 10.06 | 9.55 | 9.39 | 10.81 | 11.72 | 9.78 | 9.73 | 9.42 | 10.10 | 9.47 | 11.26 | 9.75 |
| CSK | 10 | 8.59 | 9.28 | 10.57 | 9.95 | 9.80 | 9.56 | 9.91 | 9.03 | 11.54 | 11.77 | 8.94 | 10.26 | 9.68 | 9.47 | 11.66 |
| CSNK1A1 | 9 | 9.17 | 9.75 | 9.46 | 9.04 | 10.23 | 10.31 | 10.61 | 10.52 | 10.51 | 10.42 | 9.46 | 9.25 | 10.27 | 10.56 | 10.46 |
| CSNK1D | 1 | 10.39 | 11.90 | 11.19 | 11.54 | 8.97 | 9.34 | 8.40 | 8.80 | 9.53 | 9.94 | 11.15 | 11.36 | 9.15 | 8.60 | 9.74 |
| CSNK1E | 4 | 9.61 | 10.71 | 12.88 | 8.74 | 10.10 | 8.95 | 11.26 | 9.02 | 9.49 | 9.24 | 10.16 | 10.81 | 9.52 | 10.14 | 9.37 |
| CSNK1G3 | 5 | 10.32 | 8.67 | 9.32 | 10.98 | 8.16 | 8.28 | 11.78 | 10.10 | 11.05 | 11.33 | 9.50 | 10.15 | 8.22 | 10.94 | 11.19 |
| CSNK2A1 | 8 | 8.90 | 9.68 | 11.27 | 9.47 | 9.95 | 9.99 | 10.37 | 10.75 | 9.96 | 9.66 | 9.29 | 10.37 | 9.97 | 10.56 | 9.81 |
| CSNK2A2 | 9 | 9.77 | 10.55 | 9.76 | 9.01 | 9.62 | 10.10 | 10.47 | 9.34 | 10.49 | 10.89 | 10.16 | 9.39 | 9.86 | 9.91 | 10.69 |
| CSNK2B | 17 | 7.66 | 8.47 | 10.28 | 8.16 | 9.73 | 9.06 | 9.06 | 19.85 | 9.07 | 8.67 | 8.07 | 9.22 | 9.39 | 14.45 | 8.87 |
| CSPG4 | 17 | 8.78 | 7.95 | 4.48 | 5.09 | 12.54 | 13.05 | 8.25 | 7.24 | 16.16 | 16.45 | 8.37 | 4.78 | 12.79 | 7.75 | 16.31 |
| CSRP1 | 28 | 9.88 | 10.65 | 11.96 | 10.11 | 12.06 | 11.68 | 3.84 | 6.71 | 12.01 | 11.10 | 10.26 | 11.03 | 11.87 | 5.28 | 11.56 |
| CSRP2 | 3 | 16.79 | 17.61 | 9.24 | 7.81 | 7.23 | 6.40 | 10.49 | 8.58 | 8.48 | 7.37 | 17.20 | 8.52 | 6.82 | 9.54 | 7.92 |
| CSTB | 18 | 10.52 | 10.36 | 7.82 | 6.85 | 13.69 | 13.64 | 8.58 | 7.43 | 10.61 | 10.50 | 10.44 | 7.33 | 13.67 | 8.01 | 10.55 |
| CSTF1 | 10 | 9.50 | 9.26 | 11.02 | 10.53 | 10.18 | 10.10 | 9.61 | 10.03 | 10.09 | 9.68 | 9.38 | 10.77 | 10.14 | 9.82 | 9.89 |
| CSTF2 | 11 | 9.78 | 10.52 | 9.81 | 9.36 | 10.71 | 10.97 | 11.60 | 10.43 | 8.70 | 8.11 | 10.15 | 9.59 | 10.84 | 11.02 | 8.40 |

| | | | | | | | | | | | | | | | | |
|---|---|---|---|---|---|---|---|---|---|---|---|---|---|---|---|---|
| CSTF2T | 18 | 9.50 | 10.85 | 11.58 | 11.18 | 9.88 | 9.77 | 9.54 | 8.91 | 9.43 | 9.37 | 10.17 | 11.38 | 9.83 | 9.22 | 9.40 |
| CSTF3 | 17 | 9.05 | 9.57 | 11.12 | 11.26 | 9.02 | 9.40 | 10.08 | 10.85 | 10.45 | 9.20 | 9.31 | 11.19 | 9.21 | 10.46 | 9.82 |
| CTAGE5 | 2 | 11.42 | 6.71 | 7.22 | 9.58 | 11.63 | 9.98 | 11.48 | 10.91 | 9.75 | 11.34 | 9.06 | 8.40 | 10.80 | 11.19 | 10.54 |
| CTBP1 | 9 | 8.78 | 9.85 | 10.39 | 9.78 | 9.61 | 10.45 | 11.14 | 10.82 | 9.56 | 9.61 | 9.31 | 10.09 | 10.03 | 10.98 | 9.59 |
| CTBP2 | 8 | 8.43 | 9.60 | 10.50 | 10.24 | 9.89 | 10.24 | 11.20 | 10.37 | 9.80 | 9.73 | 9.01 | 10.37 | 10.07 | 10.79 | 9.76 |
| CTCF | 3 | 10.32 | 10.79 | 9.17 | 9.32 | 10.19 | 10.77 | 10.80 | 10.11 | 9.05 | 9.48 | 10.55 | 9.24 | 10.48 | 10.46 | 9.27 |
| CTDP1 | 7 | 8.63 | 10.30 | 13.57 | 12.66 | 8.93 | 9.17 | 8.48 | 7.48 | 10.70 | 10.07 | 9.47 | 13.12 | 9.05 | 7.98 | 10.39 |
| CTDSPL2 | 7 | 9.19 | 10.67 | 8.88 | 8.65 | 10.97 | 10.99 | 9.77 | 9.37 | 10.92 | 10.59 | 9.93 | 8.76 | 10.98 | 9.57 | 10.75 |
| CTGF | 1 | 10.26 | 9.10 | 18.63 | 22.46 | 5.54 | 6.45 | 4.45 | 3.29 | 9.88 | 9.94 | 9.68 | 20.54 | 5.99 | 3.87 | 9.91 |
| CTH | 3 | 8.37 | 9.32 | 9.84 | 10.82 | 9.83 | 8.29 | 11.66 | 9.36 | 10.75 | 11.75 | 8.84 | 10.33 | 9.06 | 10.51 | 11.25 |
| CTHRC1 | 1 | 5.92 | 31.72 | 5.92 | 12.71 | 6.41 | 5.75 | 6.71 | 7.44 | 7.05 | 10.38 | 18.82 | 9.31 | 6.08 | 7.07 | 8.71 |
| CTIF | 4 | 10.56 | 11.37 | 13.88 | 12.41 | 9.57 | 8.26 | 7.47 | 5.80 | 10.59 | 10.11 | 10.96 | 13.14 | 8.91 | 6.63 | 10.35 |
| CTNNA1 | 68 | 10.23 | 11.06 | 11.67 | 11.60 | 9.67 | 9.90 | 7.04 | 8.66 | 10.22 | 9.94 | 10.65 | 11.64 | 9.78 | 7.85 | 10.08 |
| CTNNAL1 | 3 | 11.86 | 11.28 | 11.00 | 10.32 | 10.18 | 9.48 | 8.03 | 8.29 | 10.36 | 9.20 | 11.57 | 10.66 | 9.83 | 8.16 | 9.78 |
| CTNNB1 | 32 | 11.66 | 11.03 | 13.05 | 12.60 | 7.88 | 7.74 | 7.16 | 6.18 | 11.23 | 11.47 | 11.34 | 12.83 | 7.81 | 6.67 | 11.35 |
| CTNNBL1 | 23 | 8.89 | 9.63 | 10.47 | 10.36 | 9.82 | 10.17 | 11.48 | 11.16 | 9.21 | 8.82 | 9.26 | 10.41 | 10.00 | 11.32 | 9.01 |
| CTNND1 | 41 | 9.13 | 9.94 | 13.85 | 13.75 | 8.12 | 8.10 | 6.57 | 8.47 | 11.15 | 10.92 | 9.53 | 13.80 | 8.11 | 7.52 | 11.04 |
| CTPS1 | 31 | 11.29 | 11.98 | 8.83 | 7.64 | 8.15 | 7.99 | 9.22 | 13.57 | 10.77 | 10.57 | 11.63 | 8.24 | 8.07 | 11.39 | 10.67 |
| CTPS2 | 10 | 9.77 | 11.00 | 7.60 | 7.75 | 11.55 | 12.02 | 11.32 | 13.93 | 7.63 | 7.44 | 10.38 | 7.68 | 11.78 | 12.62 | 7.53 |
| CTR9 | 25 | 9.10 | 10.08 | 10.72 | 9.43 | 10.66 | 10.67 | 9.64 | 9.09 | 10.15 | 10.47 | 9.59 | 10.07 | 10.66 | 9.37 | 10.31 |
| CTSA | 7 | 8.35 | 8.24 | 12.30 | 9.60 | 15.30 | 13.25 | 9.54 | 8.20 | 7.65 | 7.57 | 8.30 | 10.95 | 14.27 | 8.87 | 7.61 |
| CTSB | 6 | 5.72 | 6.09 | 7.63 | 6.44 | 7.32 | 7.89 | 15.61 | 32.92 | 5.58 | 4.79 | 5.90 | 7.04 | 7.61 | 24.26 | 5.19 |
| CTSC | 9 | 8.18 | 8.49 | 6.84 | 5.49 | 16.17 | 15.92 | 14.72 | 17.31 | 3.57 | 3.30 | 8.33 | 6.17 | 16.05 | 16.01 | 3.44 |
| CTSL | 4 | 8.80 | 8.68 | 9.14 | 9.15 | 10.75 | 11.01 | 13.51 | 11.45 | 9.05 | 8.46 | 8.74 | 9.15 | 10.88 | 12.48 | 8.76 |
| CTSZ | 7 | 9.41 | 9.61 | 11.75 | 11.54 | 12.54 | 13.30 | 7.97 | 8.52 | 7.81 | 7.54 | 9.51 | 11.65 | 12.92 | 8.25 | 7.67 |
| CTTN | 34 | 9.39 | 9.60 | 14.80 | 11.01 | 9.72 | 9.28 | 8.69 | 6.99 | 10.59 | 9.92 | 9.50 | 12.90 | 9.50 | 7.84 | 10.26 |
| CTTNBP2NL | 24 | 10.20 | 11.24 | 12.50 | 11.34 | 8.75 | 8.52 | 8.50 | 7.48 | 10.98 | 10.50 | 10.72 | 11.92 | 8.63 | 7.99 | 10.74 |
| CTU1 | 2 | 8.73 | 9.10 | 12.19 | 10.72 | 8.10 | 8.55 | 12.02 | 11.19 | 10.17 | 9.23 | 8.91 | 11.45 | 8.33 | 11.61 | 9.70 |
| CTU2 | 2 | 8.76 | 9.64 | 10.69 | 11.28 | 9.09 | 8.97 | 11.18 | 10.25 | 10.11 | 10.04 | 9.20 | 10.98 | 9.03 | 10.72 | 10.07 |
| CUEDC1 | 2 | 10.84 | 13.45 | 12.19 | 10.89 | 7.75 | 9.10 | 6.23 | 5.05 | 14.84 | 9.67 | 12.15 | 11.54 | 8.42 | 5.64 | 12.25 |
| CUEDC2 | 5 | 10.22 | 11.14 | 6.48 | 5.57 | 13.07 | 12.67 | 10.80 | 9.51 | 10.75 | 9.80 | 10.68 | 6.02 | 12.87 | 10.15 | 10.28 |
| CUL1 | 22 | 9.30 | 9.79 | 11.67 | 10.32 | 7.46 | 7.45 | 11.37 | 13.26 | 9.66 | 9.73 | 9.54 | 10.99 | 7.46 | 12.31 | 9.69 |
| CUL2 | 16 | 8.92 | 10.15 | 11.65 | 11.27 | 9.63 | 9.97 | 9.93 | 9.92 | 9.13 | 9.43 | 9.54 | 11.46 | 9.80 | 9.93 | 9.28 |
| CUL3 | 25 | 8.81 | 9.71 | 10.80 | 9.83 | 10.53 | 10.58 | 10.22 | 9.47 | 10.20 | 9.86 | 9.26 | 10.31 | 10.55 | 9.85 | 10.03 |
| CUL4A | 26 | 10.20 | 11.00 | 9.53 | 9.36 | 10.96 | 11.27 | 9.84 | 9.82 | 9.00 | 9.01 | 10.60 | 9.45 | 11.12 | 9.83 | 9.01 |
| CUL4B | 46 | 8.54 | 9.72 | 9.77 | 8.96 | 12.85 | 12.77 | 9.98 | 9.32 | 9.04 | 9.04 | 9.13 | 9.36 | 12.81 | 9.65 | 9.04 |
| CUL5 | 9 | 9.80 | 9.88 | 10.46 | 9.06 | 10.80 | 10.00 | 10.33 | 10.10 | 10.24 | 9.32 | 9.84 | 9.76 | 10.40 | 10.22 | 9.78 |
| CUTA | 13 | 9.84 | 10.41 | 10.25 | 10.66 | 11.52 | 12.61 | 9.88 | 5.40 | 9.66 | 9.77 | 10.12 | 10.46 | 12.07 | 7.64 | 9.71 |
| CUTC | 5 | 8.68 | 10.29 | 11.30 | 10.24 | 8.84 | 9.30 | 11.38 | 13.91 | 8.02 | 8.05 | 9.48 | 10.77 | 9.07 | 12.64 | 8.03 |
| CUX1 | 25 | 10.84 | 11.98 | 9.46 | 10.05 | 10.04 | 10.41 | 8.98 | 8.49 | 9.98 | 9.77 | 11.41 | 9.76 | 10.22 | 8.74 | 9.88 |
| CWC15 | 6 | 10.12 | 10.45 | 10.54 | 9.80 | 10.79 | 10.32 | 11.39 | 8.63 | 9.50 | 8.46 | 10.29 | 10.17 | 10.55 | 10.01 | 8.98 |
| CWC22 | 6 | 9.54 | 10.00 | 10.28 | 10.48 | 9.08 | 9.39 | 11.08 | 11.01 | 9.71 | 9.44 | 9.77 | 10.38 | 9.24 | 11.04 | 9.57 |

| | | | | | | | | | | | | | | | | |
|---|---|---|---|---|---|---|---|---|---|---|---|---|---|---|---|---|
| CWC25 | 5 | 8.72 | 10.62 | 13.16 | 12.89 | 7.46 | 7.03 | 9.56 | 9.67 | 10.57 | 10.33 | 9.67 | 13.02 | 7.25 | 9.61 | 10.45 |
| CWC27 | 10 | 9.11 | 10.75 | 10.83 | 9.83 | 9.69 | 9.04 | 12.20 | 11.78 | 9.05 | 7.71 | 9.93 | 10.33 | 9.37 | 11.99 | 8.38 |
| CWF19L1 | 8 | 9.60 | 10.50 | 9.58 | 9.87 | 10.20 | 11.14 | 10.84 | 10.81 | 9.08 | 8.37 | 10.05 | 9.73 | 10.67 | 10.82 | 8.73 |
| CXADR | 4 | 9.25 | 8.95 | 8.47 | 8.55 | 9.57 | 9.05 | 9.28 | 9.08 | 12.78 | 15.04 | 9.10 | 8.51 | 9.31 | 9.18 | 13.91 |
| CXCR4 | 1 | 9.78 | 7.40 | 9.29 | 10.66 | 7.15 | 7.08 | 14.21 | 16.36 | 8.12 | 9.94 | 8.59 | 9.98 | 7.11 | 15.28 | 9.03 |
| CXorf56 | 8 | 9.70 | 10.28 | 10.33 | 10.32 | 9.31 | 9.90 | 11.84 | 11.39 | 8.75 | 8.18 | 9.99 | 10.32 | 9.60 | 11.62 | 8.47 |
| CXorf57 | 1 | 10.02 | 11.45 | 9.89 | 12.31 | 7.71 | 10.91 | 9.58 | 10.64 | 8.49 | 9.00 | 10.74 | 11.10 | 9.31 | 10.11 | 8.75 |
| CXXC1 | 2 | 9.67 | 9.02 | 9.32 | 8.47 | 11.76 | 12.19 | 10.89 | 10.47 | 8.75 | 9.46 | 9.34 | 8.90 | 11.98 | 10.68 | 9.10 |
| CYB5A | 4 | 10.80 | 9.86 | 12.07 | 10.83 | 13.37 | 12.15 | 7.20 | 4.97 | 9.23 | 9.51 | 10.33 | 11.45 | 12.76 | 6.09 | 9.37 |
| CYB5B | 3 | 9.47 | 4.91 | 6.97 | 9.54 | 9.65 | 7.76 | 14.54 | 11.03 | 11.62 | 14.53 | 7.19 | 8.26 | 8.70 | 12.78 | 13.07 |
| CYB5R1 | 9 | 10.62 | 8.45 | 9.40 | 10.79 | 11.76 | 11.18 | 8.12 | 7.39 | 10.62 | 11.65 | 9.54 | 10.10 | 11.47 | 7.76 | 11.14 |
| CYB5R3 | 26 | 11.35 | 5.87 | 11.81 | 13.40 | 8.10 | 6.41 | 8.95 | 7.19 | 12.06 | 14.87 | 8.61 | 12.60 | 7.26 | 8.07 | 13.47 |
| CYC1 | 11 | 12.99 | 6.65 | 6.93 | 10.80 | 10.79 | 8.61 | 11.36 | 9.91 | 9.89 | 12.06 | 9.82 | 8.87 | 9.70 | 10.64 | 10.97 |
| CYCS | 15 | 6.60 | 8.37 | 10.99 | 9.37 | 10.40 | 10.13 | 14.65 | 14.08 | 7.43 | 7.97 | 7.49 | 10.18 | 10.27 | 14.37 | 7.70 |
| CYFIP1 | 9 | 9.49 | 10.29 | 11.63 | 11.46 | 8.59 | 8.31 | 9.19 | 9.16 | 10.69 | 11.17 | 9.89 | 11.55 | 8.45 | 9.18 | 10.93 |
| CYFIP2 | 15 | 9.75 | 9.59 | 9.12 | 9.45 | 12.22 | 12.57 | 8.36 | 8.67 | 9.88 | 10.39 | 9.67 | 9.28 | 12.40 | 8.52 | 10.14 |
| CYHR1 | 1 | 10.39 | 10.48 | 11.46 | 9.86 | 11.59 | 11.30 | 9.68 | 7.81 | 9.88 | 7.53 | 10.44 | 10.66 | 11.45 | 8.75 | 8.71 |
| CYLD | 1 | 11.14 | 9.51 | 7.72 | 7.18 | 14.28 | 14.21 | 7.45 | 8.26 | 10.18 | 10.07 | 10.33 | 7.45 | 14.25 | 7.86 | 10.12 |
| CYP51A1 | 5 | 12.07 | 6.54 | 9.31 | 14.70 | 8.43 | 7.27 | 10.11 | 9.18 | 10.09 | 12.30 | 9.30 | 12.01 | 7.85 | 9.64 | 11.19 |
| CYR61 | 3 | 11.76 | 10.33 | 14.29 | 15.65 | 6.45 | 5.88 | 4.15 | 3.54 | 13.79 | 14.17 | 11.05 | 14.97 | 6.17 | 3.84 | 13.98 |
| CYTH1 | 3 | 11.21 | 10.92 | 15.11 | 13.12 | 8.22 | 8.73 | 6.20 | 5.45 | 11.01 | 10.02 | 11.07 | 14.11 | 8.48 | 5.83 | 10.52 |
| CYTH3 | 3 | 11.17 | 11.18 | 13.66 | 13.17 | 7.05 | 7.30 | 5.20 | 3.70 | 14.17 | 13.40 | 11.17 | 13.42 | 7.18 | 4.45 | 13.78 |
| DAAM1 | 1 | 10.08 | 8.72 | 10.87 | 10.96 | 9.82 | 7.76 | 8.97 | 4.47 | 13.75 | 14.59 | 9.40 | 10.92 | 8.79 | 6.72 | 14.17 |
| DAB2 | 31 | 15.60 | 18.16 | 4.71 | 4.35 | 14.63 | 14.86 | 7.38 | 6.72 | 6.99 | 6.59 | 16.88 | 4.53 | 14.74 | 7.05 | 6.79 |
| DAB2IP | 1 | 8.00 | 10.74 | 11.79 | 12.53 | 8.16 | 7.13 | 8.98 | 8.30 | 13.79 | 10.59 | 9.37 | 12.16 | 7.64 | 8.64 | 12.19 |
| DAD1 | 4 | 12.44 | 7.13 | 7.63 | 10.23 | 7.17 | 6.21 | 10.24 | 10.01 | 12.52 | 16.40 | 9.79 | 8.93 | 6.69 | 10.13 | 14.46 |
| DAG1 | 11 | 11.83 | 11.01 | 9.19 | 10.13 | 11.11 | 10.76 | 5.50 | 4.66 | 13.09 | 12.72 | 11.42 | 9.66 | 10.94 | 5.08 | 12.90 |
| DAGLB | 2 | 11.96 | 8.43 | 8.00 | 10.21 | 8.54 | 8.89 | 8.39 | 8.82 | 14.10 | 12.66 | 10.19 | 9.10 | 8.72 | 8.61 | 13.38 |
| DAK | 13 | 8.86 | 9.60 | 7.04 | 6.87 | 14.66 | 14.84 | 9.02 | 8.22 | 10.55 | 10.33 | 9.23 | 6.96 | 14.75 | 8.62 | 10.44 |
| DAP | 14 | 11.34 | 10.76 | 9.60 | 7.61 | 13.06 | 12.86 | 9.54 | 7.14 | 9.33 | 8.75 | 11.05 | 8.61 | 12.96 | 8.34 | 9.04 |
| DAP3 | 12 | 11.60 | 9.44 | 10.47 | 11.54 | 9.28 | 8.19 | 10.04 | 8.72 | 10.15 | 10.57 | 10.52 | 11.00 | 8.73 | 9.38 | 10.36 |
| DAPK3 | 11 | 9.39 | 10.17 | 11.97 | 12.02 | 7.51 | 7.79 | 7.46 | 9.35 | 12.22 | 12.12 | 9.78 | 11.99 | 7.65 | 8.41 | 12.17 |
| DARS | 68 | 9.11 | 9.67 | 10.54 | 10.43 | 9.90 | 10.25 | 10.70 | 11.74 | 8.88 | 8.77 | 9.39 | 10.49 | 10.07 | 11.22 | 8.83 |
| DARS2 | 9 | 11.17 | 9.28 | 10.20 | 10.76 | 9.35 | 8.42 | 9.50 | 8.95 | 11.14 | 11.22 | 10.22 | 10.48 | 8.89 | 9.23 | 11.18 |
| DAXX | 4 | 9.42 | 10.09 | 11.80 | 10.96 | 9.99 | 9.46 | 10.58 | 9.53 | 9.54 | 8.62 | 9.76 | 11.38 | 9.73 | 10.06 | 9.08 |
| DAZAP1 | 9 | 7.45 | 7.77 | 8.24 | 7.19 | 8.68 | 8.18 | 9.53 | 27.81 | 8.03 | 7.11 | 7.61 | 7.72 | 8.43 | 18.67 | 7.57 |
| DBI | 13 | 9.18 | 10.11 | 12.52 | 10.89 | 9.62 | 11.92 | 11.35 | 5.97 | 9.34 | 9.09 | 9.65 | 11.71 | 10.77 | 8.66 | 9.22 |
| DBN1 | 31 | 8.95 | 10.24 | 13.87 | 13.14 | 9.36 | 9.76 | 7.50 | 6.86 | 10.43 | 9.89 | 9.59 | 13.51 | 9.56 | 7.18 | 10.16 |
| DBNL | 15 | 10.62 | 12.39 | 12.03 | 11.73 | 7.31 | 7.36 | 7.93 | 7.14 | 11.93 | 11.56 | 11.50 | 11.88 | 7.34 | 7.53 | 11.75 |
| DBR1 | 7 | 10.08 | 10.38 | 9.68 | 9.07 | 9.75 | 10.25 | 10.55 | 10.56 | 10.22 | 9.46 | 10.23 | 9.38 | 10.00 | 10.55 | 9.84 |
| DBT | 13 | 11.30 | 10.32 | 10.08 | 10.89 | 9.85 | 9.67 | 10.11 | 9.21 | 9.34 | 9.24 | 10.81 | 10.48 | 9.76 | 9.66 | 9.29 |
| DCAF11 | 5 | 9.63 | 11.82 | 9.98 | 9.02 | 12.34 | 11.56 | 9.09 | 7.10 | 9.67 | 9.79 | 10.72 | 9.50 | 11.95 | 8.10 | 9.73 |

| | | | | | | | | | | | | | | | | |
|---|---|---|---|---|---|---|---|---|---|---|---|---|---|---|---|---|
| DCAF12 | 2 | 10.67 | 9.58 | 8.77 | 9.70 | 9.96 | 9.67 | 9.10 | 7.85 | 13.40 | 11.28 | 10.12 | 9.23 | 9.82 | 8.48 | 12.34 |
| DCAF13 | 9 | 9.00 | 9.72 | 10.37 | 9.68 | 7.66 | 8.10 | 11.72 | 15.73 | 9.03 | 8.98 | 9.36 | 10.03 | 7.88 | 13.72 | 9.01 |
| DCAF6 | 1 | 8.02 | 9.39 | 9.85 | 10.10 | 9.71 | 11.30 | 10.49 | 6.66 | 12.55 | 11.93 | 8.71 | 9.98 | 10.50 | 8.57 | 12.24 |
| DCAF7 | 4 | 10.31 | 9.64 | 10.64 | 8.51 | 9.28 | 9.65 | 11.00 | 9.25 | 11.20 | 10.54 | 9.97 | 9.57 | 9.46 | 10.12 | 10.87 |
| DCAF8 | 2 | 8.85 | 9.70 | 8.68 | 8.58 | 14.17 | 14.12 | 10.20 | 9.42 | 8.28 | 8.00 | 9.28 | 8.63 | 14.14 | 9.81 | 8.14 |
| DCBLD1 | 2 | 12.14 | 9.70 | 14.02 | 15.58 | 7.06 | 6.03 | 7.33 | 6.85 | 11.36 | 9.94 | 10.92 | 14.80 | 6.55 | 7.09 | 10.65 |
| DCDC1 | 1 | 9.43 | 11.11 | 11.11 | 15.66 | 6.75 | 11.23 | 5.93 | 0.96 | 13.53 | 14.29 | 10.27 | 13.38 | 8.99 | 3.45 | 13.91 |
| DCK | 6 | 9.92 | 10.34 | 8.99 | 8.37 | 11.03 | 10.92 | 12.43 | 11.82 | 7.94 | 8.24 | 10.13 | 8.68 | 10.97 | 12.13 | 8.09 |
| DCLK1 | 2 | 4.19 | 4.71 | 21.24 | 19.09 | 4.82 | 4.99 | 17.10 | 15.62 | 4.30 | 3.96 | 4.45 | 20.16 | 4.90 | 16.36 | 4.13 |
| DCLK2 | 3 | 11.24 | 13.68 | 12.32 | 11.42 | 11.60 | 11.06 | 4.72 | 4.16 | 10.29 | 9.50 | 12.46 | 11.87 | 11.33 | 4.44 | 9.90 |
| DCP1A | 7 | 9.04 | 9.85 | 10.06 | 9.18 | 10.06 | 10.11 | 11.78 | 10.09 | 9.95 | 9.87 | 9.44 | 9.62 | 10.08 | 10.93 | 9.91 |
| DCP1B | 1 | 9.79 | 9.72 | 12.99 | 10.96 | 10.59 | 9.56 | 11.91 | 9.33 | 8.14 | 7.01 | 9.75 | 11.97 | 10.07 | 10.62 | 7.58 |
| DCPS | 9 | 9.21 | 9.98 | 7.91 | 7.19 | 13.43 | 13.51 | 10.70 | 9.34 | 9.50 | 9.23 | 9.60 | 7.55 | 13.47 | 10.02 | 9.37 |
| DCTD | 5 | 8.67 | 9.61 | 8.99 | 8.59 | 9.69 | 10.22 | 8.69 | 14.71 | 10.72 | 10.10 | 9.14 | 8.79 | 9.96 | 11.70 | 10.41 |
| DCTN1 | 40 | 8.62 | 10.52 | 11.37 | 10.77 | 10.67 | 10.97 | 8.39 | 8.23 | 10.38 | 10.08 | 9.57 | 11.07 | 10.82 | 8.31 | 10.23 |
| DCTN2 | 30 | 9.42 | 10.77 | 11.55 | 10.78 | 10.79 | 10.75 | 8.37 | 7.56 | 10.17 | 9.83 | 10.09 | 11.17 | 10.77 | 7.97 | 10.00 |
| DCTN3 | 11 | 9.84 | 10.46 | 11.27 | 10.66 | 10.60 | 10.73 | 8.19 | 7.71 | 10.33 | 10.21 | 10.15 | 10.97 | 10.67 | 7.95 | 10.27 |
| DCTN4 | 9 | 9.59 | 10.19 | 10.92 | 10.26 | 10.40 | 10.77 | 9.43 | 8.67 | 10.02 | 9.75 | 9.89 | 10.59 | 10.59 | 9.05 | 9.89 |
| DCTN5 | 8 | 10.43 | 10.71 | 10.19 | 10.08 | 10.97 | 11.28 | 8.28 | 8.58 | 10.11 | 9.38 | 10.57 | 10.13 | 11.12 | 8.43 | 9.74 |
| DCTN6 | 2 | 10.74 | 10.72 | 10.40 | 10.57 | 11.52 | 10.92 | 8.79 | 7.66 | 9.96 | 8.71 | 10.73 | 10.49 | 11.22 | 8.23 | 9.33 |
| DCTPP1 | 10 | 10.32 | 11.13 | 11.87 | 11.99 | 9.61 | 10.18 | 9.60 | 5.41 | 10.24 | 9.64 | 10.72 | 11.93 | 9.90 | 7.51 | 9.94 |
| DCUN1D1 | 6 | 9.65 | 11.06 | 9.23 | 8.71 | 11.95 | 11.70 | 9.51 | 9.08 | 9.11 | 10.01 | 10.35 | 8.97 | 11.82 | 9.29 | 9.56 |
| DCUN1D5 | 3 | 9.82 | 9.73 | 10.86 | 10.66 | 8.59 | 9.17 | 12.19 | 10.30 | 9.90 | 8.78 | 9.77 | 10.76 | 8.88 | 11.24 | 9.34 |
| DCXR | 6 | 11.20 | 10.71 | 9.10 | 9.36 | 8.36 | 8.38 | 9.25 | 8.87 | 12.29 | 12.49 | 10.96 | 9.23 | 8.37 | 9.06 | 12.39 |
| DDA1 | 5 | 9.76 | 11.03 | 9.61 | 8.99 | 8.96 | 9.21 | 10.08 | 9.42 | 11.67 | 11.27 | 10.40 | 9.30 | 9.08 | 9.75 | 11.47 |
| DDAH1 | 13 | 11.49 | 11.80 | 13.85 | 10.90 | 9.55 | 9.35 | 5.23 | 5.23 | 11.43 | 11.17 | 11.65 | 12.38 | 9.45 | 5.23 | 11.30 |
| DDAH2 | 7 | 9.46 | 10.10 | 14.78 | 13.45 | 8.79 | 8.55 | 8.52 | 7.61 | 10.09 | 8.66 | 9.78 | 14.11 | 8.67 | 8.07 | 9.38 |
| DDB1 | 79 | 10.00 | 10.16 | 9.42 | 7.78 | 10.79 | 10.38 | 9.34 | 10.17 | 11.19 | 10.77 | 10.08 | 8.60 | 10.58 | 9.75 | 10.98 |
| DDB2 | 13 | 9.05 | 9.87 | 8.82 | 8.04 | 8.37 | 8.35 | 6.78 | 7.10 | 17.01 | 16.60 | 9.46 | 8.43 | 8.36 | 6.94 | 16.81 |
| DDHD1 | 1 | 8.79 | 10.97 | 9.26 | 10.30 | 6.83 | 6.15 | 14.42 | 14.60 | 10.48 | 8.22 | 9.88 | 9.78 | 6.49 | 14.51 | 9.35 |
| DDHD2 | 3 | 10.18 | 10.66 | 9.14 | 8.71 | 11.11 | 11.15 | 10.80 | 10.61 | 9.10 | 8.55 | 10.42 | 8.92 | 11.13 | 10.70 | 8.82 |
| DDI2 | 12 | 9.58 | 10.64 | 10.31 | 9.53 | 8.52 | 8.74 | 12.74 | 11.53 | 9.17 | 9.23 | 10.11 | 9.92 | 8.63 | 12.14 | 9.20 |
| DDOST | 13 | 13.72 | 6.53 | 8.52 | 11.93 | 7.46 | 6.26 | 10.06 | 9.91 | 11.43 | 14.17 | 10.13 | 10.22 | 6.86 | 9.99 | 12.80 |
| DDRGK1 | 4 | 12.44 | 6.26 | 6.37 | 10.57 | 10.42 | 9.39 | 11.19 | 10.12 | 10.49 | 12.76 | 9.35 | 8.47 | 9.90 | 10.66 | 11.62 |
| DDT | 9 | 9.21 | 9.88 | 9.74 | 8.50 | 8.60 | 7.98 | 8.25 | 17.11 | 10.78 | 9.94 | 9.54 | 9.12 | 8.29 | 12.68 | 10.36 |
| DDX1 | 67 | 10.30 | 11.15 | 11.21 | 10.43 | 8.26 | 8.47 | 9.84 | 11.05 | 9.77 | 9.52 | 10.73 | 10.82 | 8.36 | 10.45 | 9.64 |
| DDX10 | 8 | 8.63 | 9.45 | 10.86 | 11.00 | 9.07 | 8.64 | 12.84 | 11.49 | 9.37 | 8.65 | 9.04 | 10.93 | 8.85 | 12.17 | 9.01 |
| DDX11 | 6 | 9.49 | 10.26 | 8.32 | 7.72 | 10.14 | 9.66 | 11.94 | 11.35 | 10.77 | 10.36 | 9.88 | 8.02 | 9.90 | 11.64 | 10.57 |
| DDX17 | 45 | 9.20 | 10.06 | 9.60 | 8.70 | 10.04 | 10.16 | 11.92 | 12.55 | 9.03 | 8.73 | 9.63 | 9.15 | 10.10 | 12.24 | 8.88 |
| DDX18 | 46 | 9.03 | 9.13 | 12.11 | 10.98 | 8.78 | 8.59 | 10.29 | 10.18 | 10.14 | 10.78 | 9.08 | 11.54 | 8.69 | 10.24 | 10.46 |
| DDX19A | 7 | 9.07 | 10.20 | 8.34 | 7.78 | 12.51 | 11.88 | 10.36 | 9.30 | 10.56 | 10.00 | 9.64 | 8.06 | 12.19 | 9.83 | 10.28 |
| DDX19B | 31 | 9.59 | 10.73 | 8.64 | 8.10 | 11.10 | 11.60 | 9.99 | 10.53 | 10.08 | 9.65 | 10.16 | 8.37 | 11.35 | 10.26 | 9.86 |

| | | | | | | | | | | | | | | | | |
|---|---|---|---|---|---|---|---|---|---|---|---|---|---|---|---|---|
| DDX20 | 20 | 9.58 | 10.10 | 11.31 | 9.10 | 9.45 | 9.14 | 11.71 | 10.49 | 9.51 | 9.61 | 9.84 | 10.20 | 9.30 | 11.10 | 9.56 |
| DDX21 | 52 | 8.73 | 8.92 | 11.62 | 11.03 | 7.79 | 7.80 | 11.11 | 11.63 | 10.04 | 11.32 | 8.83 | 11.33 | 7.80 | 11.37 | 10.68 |
| DDX23 | 48 | 9.37 | 10.08 | 11.51 | 10.86 | 9.62 | 10.02 | 10.26 | 9.73 | 9.44 | 9.12 | 9.73 | 11.18 | 9.82 | 10.00 | 9.28 |
| DDX24 | 22 | 9.14 | 9.45 | 9.36 | 9.79 | 8.77 | 8.91 | 12.31 | 11.89 | 9.75 | 10.63 | 9.30 | 9.57 | 8.84 | 12.10 | 10.19 |
| DDX27 | 42 | 9.11 | 10.02 | 10.71 | 10.10 | 8.57 | 8.83 | 10.85 | 12.69 | 9.58 | 9.54 | 9.56 | 10.41 | 8.70 | 11.77 | 9.56 |
| DDX28 | 2 | 9.47 | 9.15 | 9.76 | 9.84 | 9.38 | 8.88 | 10.42 | 12.16 | 9.73 | 11.21 | 9.31 | 9.80 | 9.13 | 11.29 | 10.47 |
| DDX31 | 10 | 9.82 | 9.58 | 8.58 | 9.72 | 8.21 | 8.39 | 11.68 | 11.33 | 10.74 | 11.95 | 9.70 | 9.15 | 8.30 | 11.51 | 11.34 |
| DDX39A | 15 | 9.16 | 9.25 | 9.13 | 9.24 | 10.25 | 9.93 | 12.04 | 13.31 | 8.81 | 8.89 | 9.20 | 9.18 | 10.09 | 12.67 | 8.85 |
| DDX39B | 34 | 10.03 | 9.72 | 8.82 | 7.97 | 9.56 | 9.28 | 11.57 | 15.21 | 8.97 | 8.86 | 9.88 | 8.40 | 9.42 | 13.39 | 8.92 |
| DDX3X | 79 | 9.56 | 10.51 | 9.46 | 9.09 | 9.57 | 9.81 | 11.99 | 12.50 | 8.91 | 8.61 | 10.03 | 9.27 | 9.69 | 12.24 | 8.76 |
| DDX41 | 10 | 8.83 | 9.09 | 7.69 | 7.36 | 11.88 | 12.72 | 12.90 | 11.56 | 9.13 | 8.84 | 8.96 | 7.52 | 12.30 | 12.23 | 8.98 |
| DDX42 | 37 | 9.61 | 10.31 | 11.71 | 10.74 | 9.15 | 9.21 | 9.33 | 8.81 | 10.80 | 10.33 | 9.96 | 11.22 | 9.18 | 9.07 | 10.57 |
| DDX46 | 74 | 9.21 | 9.71 | 10.48 | 10.12 | 10.13 | 10.33 | 10.62 | 12.06 | 8.91 | 8.42 | 9.46 | 10.30 | 10.23 | 11.34 | 8.67 |
| DDX47 | 23 | 8.93 | 9.29 | 11.98 | 11.53 | 7.66 | 7.68 | 11.15 | 10.70 | 10.47 | 10.62 | 9.11 | 11.75 | 7.67 | 10.92 | 10.55 |
| DDX49 | 4 | 8.97 | 11.62 | 9.88 | 10.16 | 8.10 | 8.06 | 11.78 | 12.52 | 9.01 | 9.91 | 10.29 | 10.02 | 8.08 | 12.15 | 9.46 |
| DDX5 | 75 | 9.30 | 11.11 | 10.26 | 9.61 | 9.09 | 9.13 | 10.80 | 11.93 | 9.48 | 9.27 | 10.21 | 9.94 | 9.11 | 11.37 | 9.38 |
| DDX50 | 21 | 9.58 | 9.83 | 10.45 | 9.93 | 9.54 | 9.30 | 11.76 | 10.77 | 9.56 | 9.27 | 9.70 | 10.19 | 9.42 | 11.27 | 9.42 |
| DDX51 | 12 | 9.44 | 9.95 | 10.12 | 9.66 | 9.63 | 9.82 | 11.08 | 11.97 | 9.29 | 9.05 | 9.69 | 9.89 | 9.72 | 11.52 | 9.17 |
| DDX52 | 7 | 9.66 | 10.75 | 10.25 | 12.12 | 7.78 | 8.54 | 10.18 | 10.02 | 9.95 | 10.75 | 10.21 | 11.18 | 8.16 | 10.10 | 10.35 |
| DDX54 | 24 | 9.31 | 9.65 | 9.95 | 9.80 | 8.74 | 8.59 | 12.55 | 12.91 | 9.20 | 9.29 | 9.48 | 9.87 | 8.67 | 12.73 | 9.25 |
| DDX55 | 7 | 9.35 | 10.04 | 8.64 | 10.09 | 8.21 | 8.93 | 11.35 | 12.00 | 10.27 | 11.11 | 9.69 | 9.37 | 8.57 | 11.68 | 10.69 |
| DDX56 | 14 | 9.43 | 9.69 | 10.65 | 10.48 | 7.75 | 7.94 | 11.64 | 12.78 | 9.79 | 9.85 | 9.56 | 10.57 | 7.84 | 12.21 | 9.82 |
| DDX58 | 4 | 7.75 | 9.44 | 13.04 | 13.22 | 9.06 | 11.60 | 7.69 | 7.45 | 10.96 | 9.80 | 8.59 | 13.13 | 10.33 | 7.57 | 10.38 |
| DDX59 | 5 | 9.68 | 11.12 | 6.97 | 6.32 | 12.99 | 13.37 | 11.97 | 11.17 | 8.75 | 7.67 | 10.40 | 6.64 | 13.18 | 11.57 | 8.21 |
| DDX6 | 29 | 9.75 | 9.31 | 11.78 | 11.44 | 9.56 | 9.82 | 9.94 | 9.67 | 9.56 | 9.16 | 9.53 | 11.61 | 9.69 | 9.81 | 9.36 |
| DDX60 | 1 | 8.94 | 9.54 | 6.55 | 3.62 | 15.58 | 16.09 | 11.42 | 8.35 | 10.70 | 9.22 | 9.24 | 5.08 | 15.84 | 9.88 | 9.96 |
| DECR1 | 17 | 10.71 | 9.15 | 7.82 | 8.42 | 9.95 | 9.66 | 13.91 | 12.72 | 8.92 | 8.74 | 9.93 | 8.12 | 9.81 | 13.31 | 8.83 |
| DECR2 | 3 | 10.26 | 11.51 | 9.73 | 10.52 | 12.05 | 12.16 | 9.05 | 7.94 | 8.01 | 8.77 | 10.88 | 10.12 | 12.11 | 8.49 | 8.39 |
| DEGS1 | 2 | 11.76 | 7.25 | 8.69 | 7.81 | 10.74 | 9.34 | 9.28 | 6.80 | 12.43 | 15.90 | 9.51 | 8.25 | 10.04 | 8.04 | 14.16 |
| DEK | 30 | 8.59 | 9.53 | 11.03 | 9.61 | 10.47 | 10.36 | 11.81 | 11.52 | 8.47 | 8.60 | 9.06 | 10.32 | 10.42 | 11.67 | 8.54 |
| DENND1A | 2 | 10.22 | 9.31 | 8.38 | 9.86 | 11.94 | 11.23 | 10.94 | 8.41 | 9.64 | 10.07 | 9.76 | 9.12 | 11.58 | 9.68 | 9.86 |
| DENND3 | 1 | 16.67 | 9.36 | 10.12 | 11.28 | 8.86 | 10.18 | 9.14 | 8.98 | 9.28 | 6.13 | 13.01 | 10.70 | 9.52 | 9.06 | 7.70 |
| DENND4A | 1 | 5.91 | 9.70 | 12.87 | 10.66 | 9.52 | 9.60 | 12.96 | 11.90 | 7.90 | 9.00 | 7.80 | 11.77 | 9.56 | 12.43 | 8.45 |
| DENND4C | 12 | 11.62 | 11.39 | 8.16 | 9.17 | 9.39 | 9.96 | 8.24 | 8.33 | 12.17 | 11.57 | 11.51 | 8.66 | 9.67 | 8.28 | 11.87 |
| DENND5B | 1 | 15.76 | 14.16 | 5.14 | 5.39 | 8.27 | 9.49 | 8.33 | 8.24 | 13.87 | 11.36 | 14.96 | 5.26 | 8.88 | 8.29 | 12.61 |
| DENND6A | 2 | 10.51 | 10.19 | 8.27 | 8.97 | 11.55 | 12.31 | 11.66 | 11.08 | 8.46 | 7.00 | 10.35 | 8.62 | 11.93 | 11.37 | 7.73 |
| DENR | 8 | 9.44 | 10.01 | 9.72 | 9.71 | 10.27 | 10.78 | 11.03 | 10.20 | 9.54 | 9.30 | 9.72 | 9.72 | 10.53 | 10.62 | 9.42 |
| DEPDC7 | 1 | 10.73 | 12.06 | 11.67 | 10.22 | 11.31 | 10.70 | 8.02 | 7.08 | 9.14 | 9.07 | 11.40 | 10.94 | 11.01 | 7.55 | 9.10 |
| DERA | 12 | 8.81 | 9.34 | 8.15 | 7.22 | 15.28 | 15.00 | 9.61 | 9.42 | 8.72 | 8.45 | 9.07 | 7.69 | 15.14 | 9.51 | 8.59 |
| DERL1 | 3 | 9.91 | 8.29 | 10.26 | 12.84 | 11.03 | 8.38 | 10.02 | 7.46 | 9.49 | 12.32 | 9.10 | 11.55 | 9.71 | 8.74 | 10.91 |
| DESI1 | 3 | 10.08 | 10.05 | 14.01 | 12.94 | 7.70 | 7.03 | 9.42 | 7.78 | 10.88 | 10.11 | 10.07 | 13.47 | 7.37 | 8.60 | 10.50 |
| DESI2 | 2 | 9.64 | 11.12 | 8.87 | 9.67 | 10.77 | 11.70 | 11.27 | 10.59 | 7.75 | 8.62 | 10.38 | 9.27 | 11.23 | 10.93 | 8.19 |

| | | | | | | | | | | | | | | | | |
|---|---|---|---|---|---|---|---|---|---|---|---|---|---|---|---|---|
| DFFA | 14 | 9.48 | 11.21 | 10.55 | 9.63 | 7.30 | 7.01 | 10.33 | 12.59 | 10.87 | 11.02 | 10.35 | 10.09 | 7.16 | 11.46 | 10.94 |
| DFNA5 | 11 | 12.57 | 13.07 | 11.78 | 12.54 | 11.42 | 12.33 | 3.58 | 3.57 | 9.67 | 9.47 | 12.82 | 12.16 | 11.87 | 3.57 | 9.57 |
| DGCR14 | 6 | 10.56 | 11.13 | 12.01 | 11.87 | 7.99 | 8.16 | 9.31 | 8.40 | 10.57 | 10.01 | 10.84 | 11.94 | 8.08 | 8.85 | 10.29 |
| DGCR8 | 4 | 9.00 | 10.36 | 11.50 | 11.05 | 8.89 | 9.06 | 10.76 | 10.13 | 9.63 | 9.62 | 9.68 | 11.27 | 8.98 | 10.44 | 9.62 |
| DGKA | 8 | 9.54 | 10.15 | 8.48 | 8.32 | 11.69 | 12.06 | 5.98 | 5.50 | 14.61 | 13.68 | 9.84 | 8.40 | 11.87 | 5.74 | 14.15 |
| DGKH | 1 | 10.36 | 11.22 | 11.23 | 11.75 | 10.85 | 11.29 | 7.65 | 6.27 | 9.77 | 9.59 | 10.79 | 11.49 | 11.07 | 6.96 | 9.68 |
| DGUOK | 2 | 9.95 | 8.95 | 8.56 | 7.68 | 8.71 | 8.51 | 7.14 | 7.72 | 16.03 | 16.77 | 9.45 | 8.12 | 8.61 | 7.43 | 16.40 |
| DHCR24 | 2 | 15.46 | 7.36 | 8.42 | 12.80 | 6.25 | 6.34 | 8.47 | 8.20 | 10.65 | 16.05 | 11.41 | 10.61 | 6.30 | 8.33 | 13.35 |
| DHCR7 | 6 | 14.36 | 7.49 | 9.57 | 13.33 | 9.60 | 7.67 | 8.17 | 6.97 | 10.33 | 12.51 | 10.92 | 11.45 | 8.64 | 7.57 | 11.42 |
| DHFR | 18 | 11.06 | 12.16 | 8.90 | 7.90 | 8.50 | 8.28 | 11.13 | 9.71 | 11.37 | 10.99 | 11.61 | 8.40 | 8.39 | 10.42 | 11.18 |
| DHODH | 5 | 13.62 | 8.66 | 7.44 | 9.08 | 9.38 | 8.92 | 8.17 | 7.36 | 13.24 | 14.13 | 11.14 | 8.26 | 9.15 | 7.76 | 13.69 |
| DHPS | 9 | 8.04 | 8.71 | 12.96 | 12.46 | 7.97 | 7.64 | 13.11 | 12.47 | 8.47 | 8.17 | 8.38 | 12.71 | 7.80 | 12.79 | 8.32 |
| DHRS1 | 2 | 10.17 | 9.81 | 10.01 | 10.07 | 11.52 | 11.81 | 8.55 | 8.04 | 10.51 | 9.50 | 9.99 | 10.04 | 11.67 | 8.30 | 10.01 |
| DHRS4 | 6 | 11.09 | 11.27 | 6.68 | 7.29 | 10.71 | 11.57 | 9.70 | 9.36 | 10.70 | 11.64 | 11.18 | 6.98 | 11.14 | 9.53 | 11.17 |
| DHRS7 | 5 | 10.48 | 6.92 | 11.07 | 11.50 | 11.43 | 8.22 | 9.36 | 6.30 | 11.03 | 13.71 | 8.70 | 11.28 | 9.83 | 7.83 | 12.37 |
| DHX15 | 57 | 9.09 | 9.99 | 10.02 | 6.85 | 11.12 | 9.71 | 10.95 | 12.83 | 9.95 | 9.49 | 9.54 | 8.43 | 10.41 | 11.89 | 9.72 |
| DHX16 | 22 | 8.62 | 10.49 | 10.95 | 9.58 | 9.90 | 9.76 | 10.78 | 10.54 | 9.87 | 9.51 | 9.56 | 10.26 | 9.83 | 10.66 | 9.69 |
| DHX29 | 22 | 9.18 | 10.66 | 9.26 | 9.05 | 9.53 | 9.84 | 12.05 | 11.23 | 9.75 | 9.46 | 9.92 | 9.15 | 9.68 | 11.64 | 9.61 |
| DHX30 | 26 | 10.60 | 8.85 | 7.84 | 8.37 | 11.43 | 10.24 | 10.17 | 11.80 | 10.23 | 10.48 | 9.72 | 8.10 | 10.83 | 10.99 | 10.35 |
| DHX32 | 2 | 8.17 | 9.06 | 11.76 | 11.28 | 9.18 | 8.98 | 10.94 | 11.54 | 10.13 | 8.96 | 8.62 | 11.52 | 9.08 | 11.24 | 9.54 |
| DHX33 | 10 | 9.85 | 10.19 | 10.52 | 10.74 | 7.93 | 8.40 | 10.95 | 10.21 | 11.04 | 10.16 | 10.02 | 10.63 | 8.17 | 10.58 | 10.60 |
| DHX34 | 12 | 9.26 | 9.24 | 11.68 | 11.56 | 8.74 | 8.34 | 10.19 | 9.55 | 10.74 | 10.70 | 9.25 | 11.62 | 8.54 | 9.87 | 10.72 |
| DHX35 | 1 | 7.89 | 9.50 | 10.06 | 10.87 | 10.52 | 8.38 | 10.03 | 11.86 | 9.61 | 11.28 | 8.69 | 10.46 | 9.45 | 10.95 | 10.45 |
| DHX36 | 16 | 9.83 | 10.43 | 10.39 | 9.93 | 9.02 | 9.38 | 10.28 | 10.34 | 10.49 | 9.89 | 10.13 | 10.16 | 9.20 | 10.31 | 10.19 |
| DHX37 | 11 | 9.01 | 8.92 | 9.89 | 10.05 | 8.64 | 8.27 | 10.73 | 16.15 | 9.06 | 9.27 | 8.97 | 9.97 | 8.46 | 13.44 | 9.16 |
| DHX38 | 17 | 10.08 | 11.15 | 9.09 | 8.48 | 10.56 | 10.73 | 9.05 | 8.60 | 11.63 | 10.63 | 10.61 | 8.79 | 10.65 | 8.83 | 11.13 |
| DHX40 | 6 | 10.29 | 12.38 | 10.71 | 10.68 | 8.90 | 9.31 | 9.47 | 8.79 | 9.73 | 9.75 | 11.34 | 10.69 | 9.11 | 9.13 | 9.74 |
| DHX57 | 7 | 10.09 | 10.80 | 10.21 | 11.00 | 8.85 | 8.64 | 9.76 | 9.78 | 10.71 | 10.16 | 10.44 | 10.60 | 8.75 | 9.77 | 10.43 |
| DHX8 | 26 | 9.54 | 10.79 | 11.52 | 9.96 | 9.91 | 9.04 | 9.80 | 9.17 | 10.35 | 9.93 | 10.16 | 10.74 | 9.47 | 9.48 | 10.14 |
| DHX9 | 89 | 8.06 | 8.48 | 10.50 | 8.96 | 11.72 | 10.69 | 11.10 | 12.55 | 8.73 | 9.22 | 8.27 | 9.73 | 11.20 | 11.83 | 8.97 |
| DIABLO | 9 | 10.65 | 9.60 | 10.93 | 9.60 | 10.52 | 10.78 | 9.27 | 8.92 | 9.81 | 9.92 | 10.13 | 10.26 | 10.65 | 9.10 | 9.86 |
| DIAPH1 | 60 | 9.64 | 10.52 | 10.18 | 9.94 | 10.28 | 10.60 | 9.34 | 9.32 | 10.19 | 9.99 | 10.08 | 10.06 | 10.44 | 9.33 | 10.09 |
| DIAPH2 | 15 | 8.43 | 9.54 | 9.40 | 9.67 | 10.34 | 11.17 | 12.26 | 11.52 | 8.77 | 8.91 | 8.98 | 9.53 | 10.76 | 11.89 | 8.84 |
| DIAPH3 | 18 | 11.00 | 11.83 | 11.11 | 10.98 | 10.53 | 10.29 | 9.20 | 8.64 | 8.40 | 8.00 | 11.41 | 11.05 | 10.41 | 8.92 | 8.20 |
| DICER1 | 4 | 10.28 | 10.15 | 9.50 | 9.74 | 9.94 | 9.99 | 12.16 | 10.51 | 9.21 | 8.52 | 10.22 | 9.62 | 9.97 | 11.33 | 8.86 |
| DIDO1 | 46 | 8.57 | 9.78 | 11.19 | 11.06 | 10.09 | 10.76 | 10.58 | 9.68 | 9.32 | 8.97 | 9.17 | 11.12 | 10.43 | 10.13 | 9.15 |
| DIEXF | 12 | 9.28 | 9.55 | 9.73 | 10.29 | 9.60 | 10.38 | 10.22 | 10.46 | 10.39 | 10.10 | 9.41 | 10.01 | 9.99 | 10.34 | 10.25 |
| DIMT1 | 15 | 9.02 | 10.25 | 8.74 | 10.21 | 7.92 | 8.81 | 13.02 | 13.34 | 9.20 | 9.49 | 9.63 | 9.47 | 8.36 | 13.18 | 9.35 |
| DIP2B | 23 | 8.98 | 9.26 | 13.72 | 14.24 | 9.51 | 9.46 | 7.32 | 6.98 | 10.46 | 10.07 | 9.12 | 13.98 | 9.48 | 7.15 | 10.27 |
| DIS3 | 30 | 9.91 | 10.20 | 9.34 | 8.34 | 11.62 | 11.46 | 11.79 | 10.76 | 8.42 | 8.18 | 10.05 | 8.84 | 11.54 | 11.27 | 8.30 |
| DIS3L2 | 7 | 10.48 | 11.19 | 11.14 | 10.51 | 12.72 | 12.02 | 8.69 | 8.24 | 7.67 | 7.35 | 10.83 | 10.82 | 12.37 | 8.46 | 7.51 |
| DIXDC1 | 1 | 9.17 | 10.06 | 8.43 | 7.67 | 7.47 | 6.83 | 11.35 | 13.03 | 12.52 | 13.46 | 9.62 | 8.05 | 7.15 | 12.19 | 12.99 |

| | | | | | | | | | | | | | | | | |
|---|---|---|---|---|---|---|---|---|---|---|---|---|---|---|---|---|
| DKC1 | 28 | 9.08 | 9.47 | 9.52 | 8.68 | 9.42 | 9.53 | 12.66 | 13.09 | 9.48 | 9.07 | 9.28 | 9.10 | 9.47 | 12.88 | 9.28 |
| DLAT | 16 | 11.99 | 9.64 | 8.45 | 9.88 | 10.94 | 10.36 | 9.72 | 8.97 | 10.03 | 10.03 | 10.81 | 9.17 | 10.65 | 9.34 | 10.03 |
| DLD | 26 | 10.72 | 9.11 | 10.91 | 11.60 | 8.76 | 8.04 | 9.60 | 9.97 | 10.64 | 10.65 | 9.92 | 11.25 | 8.40 | 9.78 | 10.64 |
| DLG1 | 10 | 9.04 | 10.28 | 10.82 | 10.72 | 11.08 | 11.16 | 8.71 | 7.90 | 10.25 | 10.04 | 9.66 | 10.77 | 11.12 | 8.30 | 10.15 |
| DLG3 | 2 | 8.37 | 9.94 | 8.16 | 6.68 | 13.27 | 13.31 | 13.00 | 11.61 | 7.91 | 7.75 | 9.16 | 7.42 | 13.29 | 12.30 | 7.83 |
| DLG5 | 3 | 9.28 | 9.46 | 12.19 | 12.40 | 11.75 | 11.72 | 7.72 | 6.88 | 10.06 | 8.55 | 9.37 | 12.30 | 11.73 | 7.30 | 9.31 |
| DLGAP4 | 3 | 8.89 | 8.86 | 13.81 | 12.64 | 10.05 | 10.07 | 9.68 | 8.23 | 9.17 | 8.59 | 8.88 | 13.22 | 10.06 | 8.96 | 8.88 |
| DLGAP5 | 11 | 9.61 | 9.70 | 9.43 | 9.57 | 9.36 | 9.78 | 9.59 | 10.59 | 11.21 | 11.15 | 9.65 | 9.50 | 9.57 | 10.09 | 11.18 |
| DLST | 23 | 11.36 | 9.57 | 9.30 | 11.29 | 9.77 | 9.51 | 9.39 | 8.68 | 10.50 | 10.63 | 10.47 | 10.29 | 9.64 | 9.04 | 10.56 |
| DLX3 | 1 | 26.21 | 9.31 | 3.93 | 8.19 | 8.13 | 13.15 | 7.75 | 3.48 | 10.77 | 9.08 | 17.76 | 6.06 | 10.64 | 5.61 | 9.92 |
| DMAP1 | 8 | 9.00 | 10.02 | 11.47 | 10.95 | 9.99 | 10.31 | 10.13 | 9.27 | 8.85 | 10.02 | 9.51 | 11.21 | 10.15 | 9.70 | 9.43 |
| DMBT1 | 1 | 5.06 | 4.26 | 12.76 | 13.65 | 5.73 | 6.50 | 16.58 | 14.50 | 9.73 | 11.22 | 4.66 | 13.21 | 6.12 | 15.54 | 10.47 |
| DMD | 8 | 10.28 | 11.32 | 10.89 | 12.28 | 8.42 | 9.63 | 6.42 | 7.77 | 11.76 | 11.23 | 10.80 | 11.59 | 9.02 | 7.09 | 11.50 |
| DMXL1 | 6 | 10.17 | 11.46 | 8.80 | 9.41 | 9.46 | 9.99 | 9.31 | 9.80 | 10.85 | 10.75 | 10.82 | 9.10 | 9.72 | 9.55 | 10.80 |
| DMXL2 | 1 | 10.67 | 12.00 | 8.65 | 9.15 | 10.26 | 8.96 | 8.14 | 9.36 | 10.40 | 12.40 | 11.34 | 8.90 | 9.61 | 8.75 | 11.40 |
| DNAAF2 | 3 | 9.11 | 9.25 | 10.59 | 10.84 | 8.68 | 8.86 | 12.42 | 10.65 | 9.57 | 10.02 | 9.18 | 10.72 | 8.77 | 11.53 | 9.80 |
| DNAH9 | 2 | 10.18 | 11.19 | 9.64 | 9.29 | 10.47 | 10.85 | 10.30 | 8.93 | 9.71 | 9.45 | 10.68 | 9.46 | 10.66 | 9.61 | 9.58 |
| DNAJA1 | 34 | 10.83 | 12.28 | 9.71 | 9.75 | 8.58 | 9.04 | 7.98 | 8.66 | 11.58 | 11.59 | 11.56 | 9.73 | 8.81 | 8.32 | 11.58 |
| DNAJA2 | 15 | 10.33 | 10.93 | 8.75 | 8.45 | 10.00 | 10.15 | 10.73 | 10.86 | 10.18 | 9.62 | 10.63 | 8.60 | 10.07 | 10.80 | 9.90 |
| DNAJA3 | 11 | 10.79 | 9.41 | 9.54 | 10.11 | 10.07 | 9.33 | 10.89 | 9.78 | 10.07 | 10.01 | 10.10 | 9.82 | 9.70 | 10.34 | 10.04 |
| DNAJA4 | 5 | 11.14 | 11.96 | 9.67 | 8.74 | 11.36 | 10.60 | 7.18 | 6.10 | 11.87 | 11.38 | 11.55 | 9.20 | 10.98 | 6.64 | 11.62 |
| DNAJB1 | 30 | 8.76 | 9.80 | 11.86 | 10.89 | 8.84 | 9.02 | 10.30 | 11.64 | 9.57 | 9.32 | 9.28 | 11.38 | 8.93 | 10.97 | 9.44 |
| DNAJB11 | 18 | 10.21 | 9.22 | 9.50 | 9.67 | 11.89 | 11.81 | 10.38 | 8.79 | 9.34 | 9.17 | 9.72 | 9.59 | 11.85 | 9.59 | 9.26 |
| DNAJB12 | 8 | 11.91 | 9.17 | 9.95 | 11.26 | 10.33 | 9.78 | 8.77 | 7.78 | 10.51 | 10.56 | 10.54 | 10.60 | 10.06 | 8.27 | 10.53 |
| DNAJB14 | 2 | 9.97 | 8.66 | 10.17 | 10.39 | 10.76 | 9.38 | 11.18 | 7.90 | 10.44 | 11.16 | 9.31 | 10.28 | 10.07 | 9.54 | 10.80 |
| DNAJB2 | 3 | 10.68 | 10.03 | 11.41 | 11.69 | 10.21 | 9.88 | 7.27 | 6.09 | 11.60 | 11.15 | 10.35 | 11.55 | 10.05 | 6.68 | 11.38 |
| DNAJB4 | 11 | 12.92 | 13.64 | 8.93 | 9.04 | 9.92 | 10.50 | 5.58 | 5.01 | 12.51 | 11.96 | 13.28 | 8.98 | 10.21 | 5.30 | 12.24 |
| DNAJB6 | 7 | 9.85 | 11.36 | 11.04 | 10.88 | 7.70 | 8.25 | 9.22 | 9.23 | 11.06 | 11.41 | 10.61 | 10.96 | 7.97 | 9.22 | 11.24 |
| DNAJC1 | 2 | 9.92 | 6.44 | 9.97 | 12.10 | 10.22 | 8.47 | 11.34 | 11.31 | 8.90 | 11.32 | 8.18 | 11.04 | 9.35 | 11.33 | 10.11 |
| DNAJC10 | 7 | 10.54 | 8.48 | 7.62 | 8.80 | 10.76 | 10.78 | 11.94 | 11.65 | 9.32 | 10.09 | 9.51 | 8.21 | 10.77 | 11.80 | 9.70 |
| DNAJC11 | 10 | 11.14 | 7.65 | 8.73 | 12.58 | 8.29 | 7.66 | 10.11 | 10.00 | 10.68 | 13.16 | 9.39 | 10.65 | 7.98 | 10.05 | 11.92 |
| DNAJC12 | 3 | 8.29 | 9.90 | 5.63 | 5.99 | 17.47 | 20.87 | 12.20 | 9.62 | 5.57 | 4.46 | 9.10 | 5.81 | 19.17 | 10.91 | 5.01 |
| DNAJC13 | 36 | 9.56 | 9.96 | 11.53 | 11.22 | 10.41 | 10.62 | 8.02 | 8.13 | 10.32 | 10.23 | 9.76 | 11.38 | 10.52 | 8.08 | 10.27 |
| DNAJC15 | 2 | 12.83 | 10.03 | 7.80 | 11.00 | 9.62 | 9.42 | 8.44 | 7.88 | 9.52 | 13.46 | 11.43 | 9.40 | 9.52 | 8.16 | 11.49 |
| DNAJC16 | 2 | 12.10 | 8.24 | 6.49 | 8.77 | 9.17 | 7.16 | 9.21 | 9.49 | 13.43 | 15.94 | 10.17 | 7.63 | 8.17 | 9.35 | 14.69 |
| DNAJC17 | 8 | 8.89 | 9.42 | 11.23 | 10.48 | 8.42 | 7.50 | 10.88 | 9.61 | 11.80 | 11.76 | 9.16 | 10.86 | 7.96 | 10.25 | 11.78 |
| DNAJC19 | 6 | 9.70 | 9.20 | 10.84 | 11.58 | 9.70 | 9.96 | 11.08 | 10.24 | 8.66 | 9.04 | 9.45 | 11.21 | 9.83 | 10.66 | 8.85 |
| DNAJC2 | 16 | 8.90 | 10.10 | 11.34 | 10.46 | 8.75 | 9.12 | 11.52 | 9.99 | 10.18 | 9.63 | 9.50 | 10.90 | 8.93 | 10.76 | 9.91 |
| DNAJC21 | 17 | 10.07 | 10.45 | 9.39 | 9.11 | 10.80 | 10.77 | 10.98 | 10.64 | 8.67 | 9.12 | 10.26 | 9.25 | 10.78 | 10.81 | 8.90 |
| DNAJC24 | 1 | 9.59 | 11.21 | 10.07 | 9.63 | 12.04 | 11.81 | 11.23 | 6.72 | 9.48 | 8.22 | 10.40 | 9.85 | 11.93 | 8.97 | 8.85 |
| DNAJC25 | 1 | 9.71 | 9.73 | 3.08 | 2.46 | 14.77 | 13.22 | 8.26 | 10.92 | 14.21 | 13.63 | 9.72 | 2.77 | 14.00 | 9.59 | 13.92 |
| DNAJC3 | 18 | 9.56 | 8.25 | 9.00 | 10.33 | 13.91 | 14.21 | 10.02 | 9.63 | 7.72 | 7.37 | 8.90 | 9.66 | 14.06 | 9.83 | 7.55 |

| | | | | | | | | | | | | | | | | |
|---|---|---|---|---|---|---|---|---|---|---|---|---|---|---|---|---|
| DNAJC5 | 1 | 10.92 | 8.31 | 8.21 | 11.94 | 8.92 | 9.28 | 9.87 | 9.86 | 10.87 | 11.81 | 9.62 | 10.07 | 9.10 | 9.87 | 11.34 |
| DNAJC7 | 28 | 10.49 | 11.71 | 13.01 | 13.22 | 8.34 | 8.42 | 9.56 | 9.07 | 8.19 | 7.98 | 11.10 | 13.11 | 8.38 | 9.31 | 8.09 |
| DNAJC8 | 16 | 10.51 | 11.27 | 11.00 | 9.94 | 9.37 | 9.20 | 9.89 | 8.88 | 10.32 | 9.62 | 10.89 | 10.47 | 9.29 | 9.38 | 9.97 |
| DNAJC9 | 11 | 8.88 | 9.66 | 12.95 | 10.19 | 11.23 | 10.40 | 9.47 | 7.68 | 10.01 | 9.53 | 9.27 | 11.57 | 10.82 | 8.57 | 9.77 |
| DNAL1 | 1 | 10.83 | 10.70 | 9.84 | 9.96 | 9.88 | 11.33 | 9.40 | 8.44 | 10.69 | 8.92 | 10.77 | 9.90 | 10.61 | 8.92 | 9.81 |
| DNASE2 | 5 | 8.00 | 7.87 | 8.41 | 8.89 | 13.56 | 14.05 | 11.03 | 16.70 | 6.27 | 5.23 | 7.94 | 8.65 | 13.80 | 13.87 | 5.75 |
| DNM1 | 4 | 14.92 | 15.86 | 11.67 | 9.91 | 7.14 | 8.16 | 7.45 | 6.87 | 9.67 | 8.34 | 15.39 | 10.79 | 7.65 | 7.16 | 9.01 |
| DNM1L | 41 | 10.05 | 10.88 | 9.67 | 9.16 | 10.15 | 10.33 | 10.50 | 10.69 | 9.63 | 8.95 | 10.46 | 9.41 | 10.24 | 10.60 | 9.29 |
| DNM2 | 26 | 9.68 | 10.36 | 11.12 | 10.42 | 9.59 | 9.76 | 9.32 | 8.60 | 10.67 | 10.46 | 10.02 | 10.77 | 9.68 | 8.96 | 10.56 |
| DNM3 | 17 | 9.77 | 10.60 | 11.65 | 9.83 | 9.63 | 9.69 | 9.43 | 8.31 | 10.67 | 10.43 | 10.19 | 10.74 | 9.66 | 8.87 | 10.55 |
| DNMBP | 17 | 11.02 | 11.16 | 6.41 | 6.14 | 14.81 | 14.84 | 9.78 | 8.84 | 8.77 | 8.23 | 11.09 | 6.28 | 14.83 | 9.31 | 8.50 |
| DNMT1 | 59 | 9.33 | 9.87 | 10.01 | 9.57 | 10.05 | 10.22 | 10.51 | 11.91 | 9.21 | 9.33 | 9.60 | 9.79 | 10.14 | 11.21 | 9.27 |
| DNPEP | 8 | 8.77 | 8.73 | 12.59 | 11.93 | 10.45 | 9.28 | 10.21 | 7.85 | 9.73 | 10.45 | 8.75 | 12.26 | 9.87 | 9.03 | 10.09 |
| DNPH1 | 9 | 8.62 | 10.26 | 9.19 | 8.82 | 11.75 | 11.74 | 11.02 | 10.65 | 8.99 | 8.96 | 9.44 | 9.01 | 11.75 | 10.83 | 8.97 |
| DNTTIP1 | 5 | 8.43 | 8.90 | 10.42 | 9.18 | 8.83 | 8.52 | 11.02 | 16.89 | 9.14 | 8.67 | 8.66 | 9.80 | 8.67 | 13.95 | 8.91 |
| DNTTIP2 | 18 | 10.32 | 10.35 | 10.58 | 9.90 | 7.72 | 8.12 | 12.15 | 10.78 | 10.17 | 9.89 | 10.34 | 10.24 | 7.92 | 11.47 | 10.03 |
| DOCK1 | 4 | 9.66 | 10.58 | 11.19 | 12.26 | 8.92 | 9.35 | 10.67 | 10.34 | 8.65 | 8.38 | 10.12 | 11.72 | 9.14 | 10.51 | 8.51 |
| DOCK10 | 10 | 8.78 | 9.02 | 15.13 | 15.04 | 7.20 | 7.94 | 7.93 | 7.62 | 10.80 | 10.55 | 8.90 | 15.08 | 7.57 | 7.77 | 10.67 |
| DOCK11 | 9 | 10.62 | 8.76 | 10.82 | 11.65 | 6.13 | 6.26 | 11.03 | 10.39 | 12.56 | 11.78 | 9.69 | 11.24 | 6.20 | 10.71 | 12.17 |
| DOCK4 | 1 | 5.06 | 5.50 | 15.43 | 14.44 | 6.96 | 6.82 | 18.16 | 14.54 | 6.39 | 6.69 | 5.28 | 14.93 | 6.89 | 16.35 | 6.54 |
| DOCK5 | 19 | 10.45 | 11.42 | 8.08 | 8.82 | 12.19 | 12.83 | 7.64 | 7.58 | 10.48 | 10.51 | 10.94 | 8.45 | 12.51 | 7.61 | 10.50 |
| DOCK7 | 22 | 9.91 | 10.12 | 10.65 | 10.39 | 8.32 | 8.29 | 11.02 | 10.37 | 10.60 | 10.34 | 10.01 | 10.52 | 8.30 | 10.69 | 10.47 |
| DOCK9 | 7 | 9.37 | 10.42 | 9.27 | 10.04 | 10.37 | 11.70 | 8.00 | 8.69 | 10.75 | 11.40 | 9.90 | 9.65 | 11.03 | 8.34 | 11.08 |
| DOHH | 5 | 10.09 | 10.25 | 13.52 | 12.99 | 9.71 | 10.42 | 7.71 | 6.49 | 9.21 | 9.61 | 10.17 | 13.25 | 10.07 | 7.10 | 9.41 |
| DONSON | 1 | 10.57 | 11.39 | 8.80 | 7.93 | 10.67 | 10.98 | 7.67 | 8.38 | 11.31 | 12.30 | 10.98 | 8.36 | 10.82 | 8.02 | 11.81 |
| DPCD | 2 | 10.10 | 11.51 | 8.71 | 9.29 | 8.37 | 9.34 | 10.91 | 10.15 | 11.46 | 10.16 | 10.81 | 9.00 | 8.85 | 10.53 | 10.81 |
| DPF2 | 8 | 9.01 | 9.36 | 10.36 | 9.51 | 9.50 | 9.50 | 10.84 | 15.26 | 8.37 | 8.29 | 9.18 | 9.94 | 9.50 | 13.05 | 8.33 |
| DPH1 | 3 | 9.80 | 9.54 | 10.38 | 10.63 | 7.47 | 7.61 | 11.45 | 12.03 | 10.67 | 10.42 | 9.67 | 10.50 | 7.54 | 11.74 | 10.55 |
| DPH2 | 7 | 9.96 | 9.95 | 11.11 | 11.24 | 6.35 | 6.63 | 11.57 | 10.62 | 11.98 | 10.58 | 9.96 | 11.17 | 6.49 | 11.10 | 11.28 |
| DPH5 | 7 | 11.54 | 11.53 | 9.43 | 9.57 | 6.43 | 7.26 | 11.22 | 11.22 | 11.13 | 10.66 | 11.53 | 9.50 | 6.84 | 11.22 | 10.90 |
| DPH6 | 5 | 8.90 | 9.78 | 10.59 | 10.55 | 10.09 | 11.15 | 10.78 | 10.21 | 8.97 | 8.97 | 9.34 | 10.57 | 10.62 | 10.49 | 8.97 |
| DPM1 | 12 | 11.23 | 9.67 | 10.35 | 10.25 | 11.05 | 10.08 | 9.31 | 8.11 | 10.04 | 9.89 | 10.45 | 10.30 | 10.57 | 8.71 | 9.97 |
| DPM3 | 1 | 11.79 | 7.61 | 6.77 | 12.29 | 9.68 | 8.57 | 7.85 | 8.50 | 11.11 | 15.84 | 9.70 | 9.53 | 9.13 | 8.17 | 13.47 |
| DPP3 | 21 | 8.79 | 9.18 | 11.42 | 10.46 | 13.82 | 13.39 | 9.38 | 8.13 | 7.89 | 7.53 | 8.99 | 10.94 | 13.61 | 8.76 | 7.71 |
| DPP7 | 7 | 10.90 | 10.57 | 6.07 | 5.80 | 15.78 | 16.05 | 10.07 | 9.24 | 7.75 | 7.77 | 10.73 | 5.94 | 15.91 | 9.66 | 7.76 |
| DPP8 | 4 | 9.13 | 10.49 | 10.45 | 9.50 | 10.81 | 10.84 | 7.66 | 8.86 | 11.19 | 11.06 | 9.81 | 9.98 | 10.83 | 8.26 | 11.13 |
| DPP9 | 9 | 8.47 | 9.09 | 10.65 | 8.88 | 10.34 | 10.37 | 10.50 | 11.01 | 10.63 | 10.06 | 8.78 | 9.77 | 10.36 | 10.76 | 10.34 |
| DPYD | 16 | 6.66 | 6.92 | 7.69 | 7.44 | 5.60 | 5.98 | 20.67 | 20.04 | 9.60 | 9.40 | 6.79 | 7.57 | 5.79 | 20.35 | 9.50 |
| DPYSL2 | 35 | 10.77 | 11.92 | 9.54 | 9.24 | 10.78 | 10.93 | 7.59 | 6.71 | 11.43 | 11.09 | 11.34 | 9.39 | 10.86 | 7.15 | 11.26 |
| DPYSL3 | 55 | 5.85 | 6.75 | 12.12 | 11.55 | 8.24 | 8.77 | 8.65 | 10.48 | 13.75 | 13.84 | 6.30 | 11.84 | 8.50 | 9.57 | 13.79 |
| DR1 | 3 | 10.59 | 11.47 | 10.05 | 10.60 | 8.72 | 9.17 | 10.01 | 8.13 | 10.55 | 10.72 | 11.03 | 10.32 | 8.94 | 9.07 | 10.63 |
| DRAP1 | 10 | 9.32 | 9.88 | 9.78 | 9.60 | 8.18 | 8.43 | 9.66 | 14.21 | 10.62 | 10.32 | 9.60 | 9.69 | 8.30 | 11.94 | 10.47 |

| | | | | | | | | | | | | | | | | |
|---|---|---|---|---|---|---|---|---|---|---|---|---|---|---|---|---|
| DRG1 | 24 | 9.41 | 10.17 | 11.14 | 10.33 | 8.64 | 8.57 | 11.29 | 13.63 | 8.56 | 8.24 | 9.79 | 10.74 | 8.61 | 12.46 | 8.40 |
| DRG2 | 10 | 10.12 | 11.00 | 12.20 | 12.00 | 10.16 | 10.52 | 9.97 | 10.06 | 7.18 | 6.79 | 10.56 | 12.10 | 10.34 | 10.01 | 6.99 |
| DROSHA | 11 | 10.84 | 10.39 | 9.44 | 9.23 | 10.39 | 10.18 | 11.08 | 9.54 | 9.98 | 8.93 | 10.62 | 9.33 | 10.29 | 10.31 | 9.45 |
| DSCC1 | 2 | 10.00 | 13.53 | 10.97 | 8.41 | 8.01 | 8.39 | 9.72 | 9.78 | 11.27 | 9.93 | 11.76 | 9.69 | 8.20 | 9.75 | 10.60 |
| DSG2 | 41 | 10.33 | 8.39 | 9.47 | 11.30 | 9.17 | 9.03 | 9.61 | 9.61 | 11.46 | 11.62 | 9.36 | 10.39 | 9.10 | 9.61 | 11.54 |
| DSN1 | 5 | 10.27 | 11.92 | 8.80 | 8.60 | 11.19 | 11.76 | 9.12 | 8.35 | 10.14 | 9.85 | 11.09 | 8.70 | 11.48 | 8.74 | 10.00 |
| DSP | 104 | 8.35 | 8.84 | 9.53 | 9.29 | 13.81 | 14.08 | 7.61 | 7.23 | 10.97 | 10.29 | 8.59 | 9.41 | 13.95 | 7.42 | 10.63 |
| DST | 88 | 12.18 | 11.64 | 8.99 | 9.34 | 9.62 | 9.61 | 6.15 | 6.04 | 13.22 | 13.22 | 11.91 | 9.17 | 9.61 | 6.10 | 13.22 |
| DSTN | 17 | 10.82 | 11.26 | 12.07 | 11.15 | 9.62 | 9.27 | 4.73 | 9.78 | 10.81 | 10.49 | 11.04 | 11.61 | 9.44 | 7.26 | 10.65 |
| DSTYK | 1 | 13.00 | 11.91 | 5.31 | 7.07 | 10.64 | 12.91 | 8.22 | 8.70 | 10.41 | 11.82 | 12.46 | 6.19 | 11.78 | 8.46 | 11.11 |
| DTD1 | 3 | 7.36 | 8.42 | 13.43 | 12.55 | 11.37 | 12.00 | 11.20 | 9.88 | 7.00 | 6.79 | 7.89 | 12.99 | 11.69 | 10.54 | 6.89 |
| DTL | 3 | 9.26 | 10.32 | 8.25 | 8.68 | 12.54 | 13.46 | 10.32 | 10.61 | 7.65 | 8.91 | 9.79 | 8.46 | 13.00 | 10.47 | 8.28 |
| DTNA | 1 | 13.93 | 10.84 | 7.31 | 13.56 | 8.14 | 11.05 | 9.99 | 6.46 | 9.75 | 8.98 | 12.39 | 10.44 | 9.59 | 8.22 | 9.36 |
| DTNBP1 | 1 | 9.50 | 11.72 | 10.96 | 10.19 | 12.92 | 11.01 | 7.78 | 9.33 | 8.68 | 7.90 | 10.61 | 10.58 | 11.96 | 8.56 | 8.29 |
| DTWD1 | 1 | 7.73 | 5.61 | 9.38 | 7.51 | 11.09 | 9.90 | 14.56 | 14.34 | 11.07 | 8.82 | 6.67 | 8.44 | 10.49 | 14.45 | 9.95 |
| DTX3L | 3 | 10.58 | 13.17 | 6.14 | 5.09 | 18.31 | 20.23 | 8.89 | 7.81 | 5.50 | 4.28 | 11.88 | 5.62 | 19.27 | 8.35 | 4.89 |
| DTYMK | 25 | 9.70 | 9.94 | 11.19 | 9.81 | 10.37 | 9.59 | 9.42 | 11.69 | 9.24 | 9.05 | 9.82 | 10.50 | 9.98 | 10.55 | 9.15 |
| DUS2 | 4 | 10.60 | 10.38 | 7.58 | 7.65 | 9.88 | 10.59 | 10.32 | 11.63 | 11.31 | 10.05 | 10.49 | 7.62 | 10.24 | 10.98 | 10.68 |
| DUS3L | 7 | 8.66 | 9.95 | 9.99 | 10.05 | 11.73 | 12.13 | 9.75 | 8.90 | 9.67 | 9.17 | 9.31 | 10.02 | 11.93 | 9.33 | 9.42 |
| DUSP11 | 1 | 8.91 | 10.36 | 10.05 | 9.10 | 10.69 | 7.34 | 9.25 | 13.79 | 10.68 | 9.83 | 9.64 | 9.58 | 9.01 | 11.52 | 10.26 |
| DUSP12 | 8 | 8.44 | 8.84 | 10.66 | 10.38 | 10.90 | 10.76 | 11.24 | 10.41 | 9.20 | 9.17 | 8.64 | 10.52 | 10.83 | 10.83 | 9.18 |
| DUSP14 | 2 | 10.67 | 10.35 | 13.83 | 14.19 | 7.28 | 7.01 | 5.04 | 5.04 | 14.03 | 12.57 | 10.51 | 14.01 | 7.14 | 5.04 | 13.30 |
| DUSP23 | 9 | 6.81 | 6.58 | 11.00 | 10.61 | 10.69 | 10.74 | 14.16 | 17.65 | 6.06 | 5.70 | 6.69 | 10.81 | 10.71 | 15.91 | 5.88 |
| DUSP3 | 8 | 10.72 | 11.00 | 13.31 | 12.89 | 8.33 | 8.60 | 7.50 | 6.96 | 10.40 | 10.30 | 10.86 | 13.10 | 8.47 | 7.23 | 10.35 |
| DUT | 24 | 8.70 | 8.97 | 8.36 | 7.08 | 10.36 | 10.12 | 10.59 | 11.78 | 12.08 | 11.97 | 8.84 | 7.72 | 10.24 | 11.18 | 12.02 |
| DVL1 | 2 | 9.31 | 9.13 | 9.25 | 8.79 | 8.77 | 8.68 | 12.17 | 10.80 | 11.88 | 11.23 | 9.22 | 9.02 | 8.72 | 11.48 | 11.56 |
| DVL2 | 6 | 10.13 | 10.87 | 13.99 | 12.26 | 8.13 | 8.23 | 7.59 | 6.85 | 10.67 | 11.29 | 10.50 | 13.12 | 8.18 | 7.22 | 10.98 |
| DXO | 2 | 9.43 | 9.14 | 10.76 | 10.31 | 10.82 | 11.83 | 9.67 | 9.02 | 10.17 | 8.86 | 9.28 | 10.54 | 11.33 | 9.34 | 9.51 |
| DYM | 1 | 8.43 | 9.85 | 9.95 | 11.79 | 10.99 | 11.85 | 9.91 | 8.43 | 10.67 | 8.13 | 9.14 | 10.87 | 11.42 | 9.17 | 9.40 |
| DYNC1H1 | 296 | 9.62 | 9.55 | 9.99 | 9.87 | 10.86 | 10.63 | 8.72 | 8.85 | 10.86 | 11.04 | 9.59 | 9.93 | 10.74 | 8.78 | 10.95 |
| DYNC1I2 | 17 | 10.26 | 10.91 | 11.29 | 10.36 | 10.38 | 10.30 | 8.24 | 7.37 | 10.77 | 10.11 | 10.59 | 10.83 | 10.34 | 7.80 | 10.44 |
| DYNC1LI1 | 34 | 9.51 | 10.21 | 10.71 | 10.44 | 10.07 | 10.34 | 9.61 | 8.64 | 10.33 | 10.13 | 9.86 | 10.58 | 10.20 | 9.13 | 10.23 |
| DYNC1LI2 | 18 | 9.99 | 10.63 | 9.86 | 9.62 | 11.58 | 11.68 | 8.07 | 7.48 | 10.71 | 10.37 | 10.31 | 9.74 | 11.63 | 7.78 | 10.54 |
| DYNC2LI1 | 1 | 12.28 | 13.74 | 7.85 | 6.89 | 9.62 | 11.24 | 3.64 | 3.24 | 16.38 | 15.13 | 13.01 | 7.37 | 10.43 | 3.44 | 15.76 |
| DYNLL1 | 8 | 10.70 | 11.07 | 9.73 | 10.28 | 10.42 | 10.47 | 8.37 | 9.54 | 9.89 | 9.53 | 10.89 | 10.01 | 10.45 | 8.95 | 9.71 |
| DYNLL2 | 1 | 10.35 | 9.87 | 13.08 | 11.65 | 8.68 | 7.67 | 10.67 | 8.69 | 10.76 | 8.59 | 10.11 | 12.37 | 8.17 | 9.68 | 9.67 |
| DYNLRB1 | 13 | 9.95 | 9.87 | 9.94 | 10.25 | 11.72 | 12.84 | 8.25 | 7.84 | 9.66 | 9.67 | 9.91 | 10.10 | 12.28 | 8.05 | 9.67 |
| DYNLT1 | 1 | 7.96 | 9.29 | 14.57 | 12.97 | 7.33 | 7.61 | 6.93 | 6.80 | 12.54 | 14.00 | 8.63 | 13.77 | 7.47 | 6.86 | 13.27 |
| DYNLT3 | 1 | 3.61 | 1.47 | 4.21 | 0.63 | 7.65 | 4.40 | 4.41 | 66.96 | 4.10 | 2.55 | 2.54 | 2.42 | 6.03 | 35.69 | 3.33 |
| DYRK1A | 2 | 8.85 | 11.54 | 9.83 | 9.96 | 11.07 | 11.40 | 9.51 | 8.57 | 9.53 | 9.74 | 10.19 | 9.89 | 11.24 | 9.04 | 9.64 |
| DYRK1B | 2 | 10.00 | 11.36 | 7.69 | 9.35 | 10.63 | 10.46 | 10.92 | 11.79 | 9.22 | 8.58 | 10.68 | 8.52 | 10.55 | 11.35 | 8.90 |
| EAF1 | 4 | 9.95 | 9.95 | 10.16 | 8.93 | 8.98 | 9.74 | 12.47 | 10.75 | 9.00 | 10.07 | 9.95 | 9.54 | 9.36 | 11.61 | 9.54 |

| | | | | | | | | | | | | | | | | |
|---|---|---|---|---|---|---|---|---|---|---|---|---|---|---|---|---|
| EAPP | 7 | 10.46 | 11.45 | 9.22 | 9.83 | 10.13 | 10.77 | 8.87 | 8.06 | 9.84 | 11.37 | 10.96 | 9.52 | 10.45 | 8.46 | 10.61 |
| EARS2 | 3 | 11.91 | 10.08 | 8.15 | 9.88 | 10.00 | 10.34 | 9.02 | 8.69 | 11.05 | 10.88 | 11.00 | 9.01 | 10.17 | 8.85 | 10.97 |
| EBF3 | 1 | 10.08 | 12.56 | 10.46 | 8.74 | 9.16 | 9.26 | 7.70 | 7.43 | 12.39 | 12.21 | 11.32 | 9.60 | 9.21 | 7.57 | 12.30 |
| EBNA1BP2 | 25 | 8.57 | 10.06 | 8.90 | 9.08 | 7.38 | 7.65 | 13.15 | 16.26 | 9.19 | 9.76 | 9.31 | 8.99 | 7.51 | 14.71 | 9.48 |
| ECD | 8 | 10.38 | 11.17 | 12.85 | 12.58 | 8.11 | 8.34 | 8.19 | 8.23 | 10.29 | 9.87 | 10.77 | 12.71 | 8.22 | 8.21 | 10.08 |
| ECE1 | 10 | 10.94 | 10.43 | 9.32 | 12.38 | 7.71 | 7.54 | 7.80 | 7.90 | 12.22 | 13.76 | 10.68 | 10.85 | 7.62 | 7.85 | 12.99 |
| ECE2 | 1 | 8.24 | 9.94 | 9.18 | 9.60 | 9.36 | 10.18 | 14.41 | 13.43 | 7.91 | 7.75 | 9.09 | 9.39 | 9.77 | 13.92 | 7.83 |
| ECH1 | 15 | 11.47 | 9.40 | 8.36 | 8.99 | 9.11 | 8.76 | 10.29 | 13.29 | 10.13 | 10.22 | 10.43 | 8.67 | 8.93 | 11.79 | 10.17 |
| ECHDC1 | 12 | 9.08 | 8.71 | 12.43 | 13.09 | 9.14 | 9.62 | 10.58 | 10.40 | 8.47 | 8.48 | 8.89 | 12.76 | 9.38 | 10.49 | 8.48 |
| ECHS1 | 30 | 10.97 | 10.21 | 10.05 | 10.58 | 10.15 | 9.32 | 10.31 | 9.23 | 9.42 | 9.76 | 10.59 | 10.31 | 9.73 | 9.77 | 9.59 |
| ECI1 | 5 | 11.63 | 10.45 | 8.63 | 9.11 | 10.26 | 9.82 | 9.67 | 9.47 | 10.28 | 10.67 | 11.04 | 8.87 | 10.04 | 9.57 | 10.48 |
| ECI2 | 8 | 11.00 | 10.65 | 8.85 | 8.95 | 11.59 | 11.38 | 10.28 | 9.28 | 9.00 | 9.01 | 10.82 | 8.90 | 11.49 | 9.78 | 9.01 |
| ECM29 | 56 | 9.39 | 9.47 | 8.76 | 9.09 | 10.63 | 11.01 | 10.03 | 10.72 | 10.44 | 10.46 | 9.43 | 8.93 | 10.82 | 10.37 | 10.45 |
| ECSIT | 2 | 12.01 | 10.52 | 9.77 | 11.88 | 9.78 | 9.74 | 11.48 | 10.93 | 6.75 | 7.15 | 11.26 | 10.82 | 9.76 | 11.20 | 6.95 |
| ECT2 | 6 | 9.43 | 9.88 | 9.80 | 9.84 | 10.54 | 11.15 | 10.03 | 9.33 | 9.97 | 10.03 | 9.65 | 9.82 | 10.85 | 9.68 | 10.00 |
| EDC3 | 10 | 9.30 | 10.25 | 11.77 | 10.18 | 10.07 | 9.77 | 9.59 | 7.95 | 10.76 | 10.36 | 9.77 | 10.98 | 9.92 | 8.77 | 10.56 |
| EDC4 | 27 | 10.94 | 11.96 | 8.98 | 9.01 | 10.15 | 10.78 | 9.12 | 9.13 | 10.26 | 9.66 | 11.45 | 9.00 | 10.47 | 9.12 | 9.96 |
| EDEM3 | 4 | 10.18 | 9.09 | 9.32 | 10.30 | 10.50 | 10.06 | 11.67 | 10.33 | 8.92 | 9.63 | 9.64 | 9.81 | 10.28 | 11.00 | 9.28 |
| EDF1 | 17 | 9.51 | 10.28 | 11.15 | 9.92 | 10.28 | 10.11 | 9.57 | 8.44 | 10.51 | 10.24 | 9.89 | 10.54 | 10.19 | 9.01 | 10.37 |
| EDIL3 | 8 | 7.00 | 5.82 | 18.27 | 18.03 | 5.27 | 4.76 | 11.57 | 10.56 | 9.14 | 9.57 | 6.41 | 18.15 | 5.02 | 11.06 | 9.36 |
| EDRF1 | 2 | 10.25 | 10.85 | 9.64 | 9.79 | 8.60 | 9.17 | 9.89 | 10.27 | 10.95 | 10.60 | 10.55 | 9.71 | 8.89 | 10.08 | 10.77 |
| EEA1 | 55 | 9.52 | 10.16 | 10.81 | 10.18 | 8.95 | 9.29 | 10.61 | 9.77 | 10.64 | 10.07 | 9.84 | 10.49 | 9.12 | 10.19 | 10.35 |
| EED | 8 | 10.14 | 11.05 | 11.51 | 9.14 | 9.84 | 9.19 | 11.20 | 9.03 | 9.93 | 8.97 | 10.59 | 10.33 | 9.52 | 10.11 | 9.45 |
| EEF1A1 | 159 | 8.98 | 12.87 | 10.44 | 10.93 | 9.35 | 9.89 | 9.48 | 9.71 | 9.10 | 9.24 | 10.93 | 10.69 | 9.62 | 9.60 | 9.17 |
| EEF1A2 | 121 | 8.98 | 12.45 | 12.22 | 9.04 | 10.62 | 9.34 | 9.62 | 7.78 | 9.55 | 10.40 | 10.72 | 10.63 | 9.98 | 8.70 | 9.98 |
| EEF1B2 | 42 | 9.93 | 10.16 | 13.55 | 12.54 | 9.00 | 9.06 | 8.93 | 6.67 | 10.52 | 9.64 | 10.05 | 13.04 | 9.03 | 7.80 | 10.08 |
| EEF1D | 60 | 10.53 | 10.37 | 11.73 | 11.37 | 8.46 | 8.65 | 8.99 | 9.05 | 10.76 | 10.10 | 10.45 | 11.55 | 8.55 | 9.02 | 10.43 |
| EEF1E1 | 13 | 9.13 | 10.10 | 9.60 | 10.57 | 10.63 | 11.39 | 11.45 | 12.30 | 7.37 | 7.46 | 9.61 | 10.09 | 11.01 | 11.88 | 7.42 |
| EEF1G | 67 | 9.23 | 11.55 | 12.19 | 11.26 | 8.91 | 9.15 | 8.66 | 8.86 | 10.16 | 10.03 | 10.39 | 11.72 | 9.03 | 8.76 | 10.10 |
| EEF2 | 207 | 9.18 | 9.71 | 11.17 | 9.97 | 10.38 | 10.04 | 9.29 | 10.95 | 9.82 | 9.49 | 9.44 | 10.57 | 10.21 | 10.12 | 9.65 |
| EEF2K | 7 | 10.17 | 11.27 | 10.58 | 10.28 | 10.74 | 10.30 | 9.72 | 8.76 | 9.36 | 8.83 | 10.72 | 10.43 | 10.52 | 9.24 | 9.09 |
| EEFSEC | 8 | 9.16 | 10.23 | 9.45 | 9.74 | 11.78 | 12.66 | 12.13 | 12.93 | 6.09 | 5.83 | 9.69 | 9.60 | 12.22 | 12.53 | 5.96 |
| EFHD2 | 17 | 8.69 | 9.27 | 8.78 | 7.77 | 6.47 | 6.40 | 15.40 | 16.33 | 10.53 | 10.35 | 8.98 | 8.28 | 6.44 | 15.87 | 10.44 |
| EFNA5 | 1 | 10.84 | 10.23 | 9.26 | 11.98 | 7.60 | 9.76 | 8.88 | 8.47 | 9.03 | 13.97 | 10.53 | 10.62 | 8.68 | 8.67 | 11.50 |
| EFR3A | 3 | 11.74 | 9.27 | 7.09 | 8.78 | 8.78 | 9.42 | 11.16 | 10.52 | 11.70 | 11.54 | 10.50 | 7.94 | 9.10 | 10.84 | 11.62 |
| EFTUD1 | 20 | 9.69 | 10.40 | 10.18 | 9.50 | 11.96 | 11.92 | 8.56 | 7.90 | 10.28 | 9.61 | 10.05 | 9.84 | 11.94 | 8.23 | 9.95 |
| EFTUD2 | 70 | 9.56 | 10.21 | 13.01 | 10.57 | 8.83 | 8.56 | 9.15 | 9.72 | 10.42 | 9.97 | 9.88 | 11.79 | 8.70 | 9.44 | 10.20 |
| EGFR | 14 | 9.65 | 7.52 | 8.02 | 9.10 | 9.38 | 9.48 | 12.99 | 13.23 | 9.86 | 10.77 | 8.59 | 8.56 | 9.43 | 13.11 | 10.31 |
| EGLN1 | 3 | 9.19 | 9.67 | 9.29 | 7.72 | 10.23 | 10.99 | 11.81 | 12.15 | 9.31 | 9.65 | 9.43 | 8.50 | 10.61 | 11.98 | 9.48 |
| EHBP1 | 8 | 9.15 | 10.18 | 11.01 | 10.05 | 9.46 | 9.95 | 11.95 | 11.14 | 8.81 | 8.31 | 9.66 | 10.53 | 9.70 | 11.54 | 8.56 |
| EHBP1L1 | 29 | 8.93 | 9.54 | 11.62 | 11.20 | 10.37 | 10.80 | 8.14 | 7.17 | 11.67 | 10.56 | 9.24 | 11.41 | 10.58 | 7.65 | 11.11 |
| EHD1 | 36 | 9.62 | 10.18 | 12.62 | 11.25 | 9.69 | 9.52 | 7.57 | 6.49 | 11.73 | 11.32 | 9.90 | 11.94 | 9.61 | 7.03 | 11.52 |

| | | | | | | | | | | | | | | | | |
|---|---|---|---|---|---|---|---|---|---|---|---|---|---|---|---|---|
| EHD2 | 41 | 10.06 | 11.18 | 12.03 | 11.32 | 7.43 | 7.17 | 7.75 | 7.16 | 12.83 | 13.06 | 10.62 | 11.68 | 7.30 | 7.46 | 12.94 |
| EHD4 | 28 | 7.63 | 8.09 | 16.81 | 15.96 | 6.68 | 6.77 | 8.49 | 8.15 | 10.77 | 10.65 | 7.86 | 16.39 | 6.73 | 8.32 | 10.71 |
| EHHADH | 2 | 11.96 | 12.93 | 7.62 | 8.46 | 11.60 | 11.47 | 9.20 | 9.76 | 8.77 | 8.23 | 12.44 | 8.04 | 11.54 | 9.48 | 8.50 |
| EHMT1 | 10 | 9.76 | 10.55 | 10.61 | 9.83 | 9.82 | 10.53 | 11.01 | 9.69 | 9.46 | 8.73 | 10.15 | 10.22 | 10.17 | 10.35 | 9.10 |
| EHMT2 | 10 | 9.26 | 10.19 | 9.97 | 9.97 | 9.94 | 10.19 | 11.20 | 10.91 | 9.47 | 8.90 | 9.73 | 9.97 | 10.06 | 11.06 | 9.18 |
| EI24 | 1 | 11.51 | 5.86 | 7.85 | 11.87 | 9.17 | 8.19 | 10.79 | 10.30 | 10.54 | 13.90 | 8.69 | 9.86 | 8.68 | 10.55 | 12.22 |
| EID2 | 1 | 13.22 | 10.96 | 8.14 | 5.81 | 12.12 | 12.66 | 10.63 | 11.28 | 7.26 | 7.91 | 12.09 | 6.98 | 12.39 | 10.96 | 7.58 |
| EIF1 | 9 | 9.34 | 9.98 | 9.93 | 8.73 | 9.25 | 9.36 | 9.80 | 13.42 | 10.32 | 9.86 | 9.66 | 9.33 | 9.31 | 11.61 | 10.09 |
| EIF1AD | 8 | 8.96 | 10.44 | 12.91 | 11.59 | 8.96 | 8.72 | 10.45 | 7.57 | 10.39 | 10.01 | 9.70 | 12.25 | 8.84 | 9.01 | 10.20 |
| EIF1AX | 13 | 8.46 | 9.65 | 8.52 | 7.95 | 10.56 | 11.17 | 12.19 | 15.37 | 8.28 | 7.84 | 9.05 | 8.24 | 10.87 | 13.78 | 8.06 |
| EIF2A | 37 | 9.06 | 10.28 | 10.91 | 10.47 | 9.43 | 9.44 | 9.90 | 11.25 | 9.53 | 9.74 | 9.67 | 10.69 | 9.43 | 10.58 | 9.63 |
| EIF2AK2 | 19 | 9.88 | 10.74 | 9.27 | 8.85 | 11.41 | 11.52 | 10.27 | 9.42 | 9.67 | 8.96 | 10.31 | 9.06 | 11.46 | 9.85 | 9.32 |
| EIF2AK4 | 12 | 9.55 | 10.20 | 9.19 | 9.20 | 8.95 | 9.29 | 9.79 | 9.87 | 12.13 | 11.83 | 9.87 | 9.20 | 9.12 | 9.83 | 11.98 |
| EIF2B1 | 18 | 10.05 | 9.98 | 10.24 | 8.97 | 10.33 | 10.06 | 9.51 | 10.37 | 10.47 | 10.01 | 10.02 | 9.61 | 10.20 | 9.94 | 10.24 |
| EIF2B2 | 14 | 9.74 | 10.88 | 10.41 | 9.11 | 9.76 | 9.79 | 9.91 | 9.78 | 10.31 | 10.32 | 10.31 | 9.76 | 9.77 | 9.84 | 10.32 |
| EIF2B3 | 22 | 10.90 | 11.61 | 10.86 | 9.91 | 7.34 | 7.43 | 10.67 | 10.11 | 10.80 | 10.36 | 11.26 | 10.39 | 7.39 | 10.39 | 10.58 |
| EIF2B4 | 15 | 10.30 | 10.87 | 10.81 | 10.38 | 8.67 | 8.28 | 10.79 | 10.41 | 9.74 | 9.75 | 10.58 | 10.60 | 8.48 | 10.60 | 9.74 |
| EIF2B5 | 7 | 10.45 | 11.21 | 10.23 | 10.22 | 8.06 | 9.20 | 10.85 | 10.34 | 9.70 | 9.75 | 10.83 | 10.22 | 8.63 | 10.60 | 9.72 |
| EIF2D | 13 | 9.73 | 10.58 | 8.68 | 8.44 | 10.39 | 10.79 | 10.51 | 11.23 | 10.18 | 9.47 | 10.16 | 8.56 | 10.59 | 10.87 | 9.83 |
| EIF2S1 | 55 | 8.99 | 9.61 | 9.27 | 9.07 | 10.09 | 10.44 | 10.12 | 14.02 | 9.32 | 9.07 | 9.30 | 9.17 | 10.26 | 12.07 | 9.19 |
| EIF2S2 | 40 | 9.23 | 9.58 | 9.84 | 9.13 | 10.21 | 10.44 | 10.16 | 13.38 | 9.30 | 8.72 | 9.40 | 9.48 | 10.33 | 11.77 | 9.01 |
| EIF2S3 | 41 | 8.80 | 9.52 | 9.51 | 8.12 | 10.82 | 10.53 | 10.58 | 14.33 | 9.04 | 8.75 | 9.16 | 8.81 | 10.67 | 12.46 | 8.90 |
| EIF3A | 107 | 9.77 | 10.44 | 10.40 | 9.75 | 9.62 | 9.71 | 9.91 | 10.60 | 10.02 | 9.79 | 10.10 | 10.07 | 9.67 | 10.25 | 9.90 |
| EIF3B | 61 | 9.96 | 10.69 | 10.97 | 9.68 | 8.28 | 8.74 | 10.41 | 11.13 | 10.15 | 9.98 | 10.32 | 10.33 | 8.51 | 10.77 | 10.07 |
| EIF3C | 41 | 8.96 | 10.03 | 10.70 | 9.78 | 9.38 | 9.48 | 10.59 | 11.26 | 9.93 | 9.90 | 9.49 | 10.24 | 9.43 | 10.92 | 9.92 |
| EIF3D | 28 | 10.01 | 10.55 | 10.80 | 10.27 | 8.68 | 9.11 | 10.65 | 10.46 | 9.92 | 9.55 | 10.28 | 10.53 | 8.89 | 10.56 | 9.74 |
| EIF3E | 49 | 9.58 | 10.42 | 10.52 | 10.33 | 9.25 | 9.29 | 10.49 | 9.95 | 10.21 | 9.97 | 10.00 | 10.42 | 9.27 | 10.22 | 10.09 |
| EIF3F | 42 | 8.78 | 10.16 | 11.74 | 10.72 | 9.70 | 9.81 | 10.11 | 9.22 | 9.78 | 9.99 | 9.47 | 11.23 | 9.75 | 9.66 | 9.88 |
| EIF3G | 22 | 9.50 | 10.59 | 10.58 | 10.17 | 8.95 | 8.93 | 9.59 | 13.31 | 9.02 | 9.37 | 10.04 | 10.38 | 8.94 | 11.45 | 9.19 |
| EIF3H | 30 | 9.41 | 10.99 | 10.20 | 10.26 | 9.44 | 9.65 | 9.84 | 10.33 | 10.00 | 9.87 | 10.20 | 10.23 | 9.55 | 10.08 | 9.94 |
| EIF3I | 29 | 8.53 | 10.33 | 12.23 | 10.34 | 8.95 | 8.47 | 10.77 | 10.94 | 9.80 | 9.64 | 9.43 | 11.29 | 8.71 | 10.85 | 9.72 |
| EIF3J | 24 | 9.60 | 10.82 | 10.80 | 9.74 | 10.03 | 10.27 | 9.32 | 6.08 | 11.94 | 11.41 | 10.21 | 10.27 | 10.15 | 7.70 | 11.67 |
| EIF3K | 9 | 8.28 | 9.49 | 9.71 | 8.67 | 9.14 | 8.93 | 9.65 | 18.39 | 9.33 | 8.41 | 8.89 | 9.19 | 9.03 | 14.02 | 8.87 |
| EIF3L | 39 | 9.73 | 10.02 | 10.34 | 9.65 | 9.25 | 9.13 | 10.04 | 10.91 | 10.25 | 10.68 | 9.88 | 10.00 | 9.19 | 10.47 | 10.46 |
| EIF3M | 21 | 8.91 | 9.75 | 10.89 | 9.31 | 9.94 | 9.13 | 10.00 | 12.67 | 9.98 | 9.41 | 9.33 | 10.10 | 9.54 | 11.34 | 9.70 |
| EIF4A1 | 60 | 9.54 | 10.24 | 12.28 | 10.26 | 9.40 | 8.76 | 10.19 | 9.05 | 10.23 | 10.04 | 9.89 | 11.27 | 9.08 | 9.62 | 10.14 |
| EIF4A2 | 52 | 9.32 | 10.33 | 14.51 | 12.56 | 7.82 | 7.68 | 9.32 | 9.34 | 9.57 | 9.55 | 9.82 | 13.53 | 7.75 | 9.33 | 9.56 |
| EIF4A3 | 65 | 9.81 | 10.47 | 11.80 | 10.77 | 8.92 | 9.11 | 9.96 | 9.57 | 9.80 | 9.79 | 10.14 | 11.29 | 9.01 | 9.77 | 9.79 |
| EIF4B | 49 | 9.25 | 10.54 | 11.76 | 11.22 | 10.42 | 10.74 | 8.83 | 7.95 | 9.94 | 9.35 | 9.90 | 11.49 | 10.58 | 8.39 | 9.65 |
| EIF4E | 9 | 8.24 | 9.08 | 12.91 | 9.64 | 11.41 | 10.16 | 8.33 | 10.33 | 10.15 | 9.76 | 8.66 | 11.27 | 10.78 | 9.33 | 9.95 |
| EIF4E2 | 7 | 9.14 | 10.77 | 10.85 | 8.09 | 10.57 | 9.79 | 10.92 | 9.33 | 10.24 | 10.31 | 9.96 | 9.47 | 10.18 | 10.12 | 10.27 |
| EIF4EBP1 | 5 | 9.92 | 9.75 | 7.87 | 12.28 | 7.80 | 9.56 | 13.98 | 7.15 | 10.75 | 10.93 | 9.84 | 10.08 | 8.68 | 10.57 | 10.84 |

| | | | | | | | | | | | | | | | | |
|---|---|---|---|---|---|---|---|---|---|---|---|---|---|---|---|---|
| EIF4EBP2 | 1 | 11.16 | 12.71 | 6.26 | 8.11 | 14.58 | 18.19 | 8.14 | 3.58 | 9.27 | 8.00 | 11.94 | 7.18 | 16.39 | 5.86 | 8.64 |
| EIF4ENIF1 | 3 | 10.77 | 12.73 | 8.78 | 8.73 | 9.05 | 8.79 | 11.54 | 11.51 | 8.90 | 9.20 | 11.75 | 8.75 | 8.92 | 11.53 | 9.05 |
| EIF4G1 | 110 | 9.76 | 10.60 | 9.23 | 8.81 | 10.58 | 10.75 | 10.03 | 10.60 | 10.04 | 9.60 | 10.18 | 9.02 | 10.66 | 10.32 | 9.82 |
| EIF4G2 | 61 | 9.62 | 10.13 | 10.43 | 9.64 | 9.18 | 9.46 | 10.67 | 10.04 | 10.53 | 10.31 | 9.87 | 10.03 | 9.32 | 10.35 | 10.42 |
| EIF4G3 | 27 | 10.26 | 11.91 | 8.88 | 8.41 | 9.24 | 9.63 | 12.39 | 11.25 | 9.09 | 8.96 | 11.09 | 8.64 | 9.43 | 11.82 | 9.02 |
| EIF4H | 25 | 9.96 | 10.96 | 10.68 | 10.30 | 8.48 | 8.62 | 9.91 | 10.98 | 10.39 | 9.72 | 10.46 | 10.49 | 8.55 | 10.45 | 10.06 |
| EIF5 | 26 | 7.52 | 9.14 | 10.62 | 9.23 | 8.44 | 8.09 | 11.15 | 15.27 | 10.35 | 10.21 | 8.33 | 9.92 | 8.26 | 13.21 | 10.28 |
| EIF5A | 51 | 9.18 | 13.60 | 12.36 | 12.96 | 7.62 | 7.69 | 8.51 | 7.92 | 10.03 | 10.13 | 11.39 | 12.66 | 7.65 | 8.21 | 10.08 |
| EIF5B | 58 | 8.73 | 9.54 | 9.13 | 8.16 | 9.52 | 9.63 | 11.59 | 17.36 | 8.29 | 8.05 | 9.13 | 8.64 | 9.58 | 14.48 | 8.17 |
| EIF6 | 9 | 8.82 | 10.29 | 11.89 | 11.94 | 7.84 | 8.36 | 9.01 | 11.49 | 10.32 | 10.06 | 9.55 | 11.91 | 8.10 | 10.25 | 10.19 |
| ELAC2 | 26 | 9.95 | 10.09 | 11.32 | 11.58 | 7.44 | 7.56 | 8.51 | 12.64 | 10.47 | 10.45 | 10.02 | 11.45 | 7.50 | 10.57 | 10.46 |
| ELAVL1 | 26 | 9.26 | 10.38 | 11.19 | 9.91 | 10.72 | 10.73 | 10.58 | 8.96 | 9.34 | 8.94 | 9.82 | 10.55 | 10.73 | 9.77 | 9.14 |
| ELF1 | 1 | 9.04 | 11.23 | 11.87 | 9.73 | 11.16 | 11.28 | 11.61 | 9.44 | 8.49 | 6.15 | 10.13 | 10.80 | 11.22 | 10.52 | 7.32 |
| ELF2 | 2 | 11.06 | 10.38 | 10.59 | 10.41 | 9.90 | 9.42 | 11.12 | 8.51 | 9.41 | 9.20 | 10.72 | 10.50 | 9.66 | 9.81 | 9.31 |
| ELL | 1 | 11.02 | 8.55 | 9.00 | 8.00 | 10.53 | 10.60 | 12.17 | 11.84 | 9.46 | 8.83 | 9.79 | 8.50 | 10.56 | 12.01 | 9.15 |
| ELL2 | 5 | 8.00 | 9.21 | 10.63 | 9.56 | 7.56 | 7.86 | 14.87 | 14.49 | 8.73 | 9.09 | 8.61 | 10.10 | 7.71 | 14.68 | 8.91 |
| ELMO2 | 10 | 9.72 | 10.85 | 10.77 | 10.41 | 10.85 | 11.22 | 8.82 | 8.14 | 9.74 | 9.48 | 10.28 | 10.59 | 11.04 | 8.48 | 9.61 |
| ELMOD2 | 2 | 11.65 | 6.87 | 9.75 | 12.08 | 8.72 | 6.46 | 11.72 | 10.26 | 10.83 | 11.66 | 9.26 | 10.91 | 7.59 | 10.99 | 11.24 |
| ELMSAN1 | 4 | 9.30 | 9.81 | 9.61 | 10.16 | 8.87 | 10.22 | 10.48 | 12.47 | 9.64 | 9.45 | 9.55 | 9.89 | 9.54 | 11.47 | 9.54 |
| ELOVL1 | 1 | 13.00 | 7.56 | 6.62 | 12.06 | 7.59 | 5.68 | 8.14 | 9.84 | 13.12 | 16.39 | 10.28 | 9.34 | 6.63 | 8.99 | 14.75 |
| ELP2 | 4 | 9.64 | 9.54 | 11.07 | 9.20 | 10.11 | 8.49 | 9.01 | 9.09 | 12.22 | 11.63 | 9.59 | 10.14 | 9.30 | 9.05 | 11.93 |
| ELP3 | 9 | 8.35 | 15.09 | 8.90 | 10.28 | 7.96 | 7.71 | 11.20 | 11.08 | 9.36 | 10.06 | 11.72 | 9.59 | 7.84 | 11.14 | 9.71 |
| ELP4 | 4 | 9.77 | 11.25 | 10.79 | 10.12 | 9.99 | 10.07 | 10.87 | 9.81 | 8.69 | 8.63 | 10.51 | 10.46 | 10.03 | 10.34 | 8.66 |
| ELP5 | 1 | 12.12 | 12.20 | 9.47 | 9.96 | 8.79 | 10.16 | 9.45 | 9.92 | 9.46 | 8.48 | 12.16 | 9.72 | 9.48 | 9.68 | 8.97 |
| ELP6 | 8 | 9.51 | 11.47 | 10.29 | 10.04 | 10.06 | 9.86 | 10.38 | 9.91 | 9.40 | 9.08 | 10.49 | 10.17 | 9.96 | 10.14 | 9.24 |
| EMB | 2 | 11.54 | 8.34 | 16.21 | 18.49 | 7.57 | 7.41 | 5.33 | 5.10 | 10.40 | 9.63 | 9.94 | 17.35 | 7.49 | 5.21 | 10.01 |
| EMC1 | 16 | 11.56 | 7.80 | 7.24 | 10.30 | 10.42 | 9.79 | 10.82 | 10.85 | 9.87 | 11.34 | 9.68 | 8.77 | 10.10 | 10.84 | 10.61 |
| EMC10 | 4 | 12.03 | 8.40 | 7.93 | 10.49 | 10.55 | 10.09 | 10.13 | 9.47 | 10.29 | 10.61 | 10.22 | 9.21 | 10.32 | 9.80 | 10.45 |
| EMC2 | 12 | 9.72 | 10.67 | 9.88 | 9.49 | 10.67 | 11.49 | 10.39 | 9.35 | 9.46 | 8.88 | 10.20 | 9.68 | 11.08 | 9.87 | 9.17 |
| EMC3 | 4 | 11.20 | 7.98 | 8.12 | 10.39 | 10.55 | 10.05 | 10.12 | 9.64 | 10.10 | 11.84 | 9.59 | 9.26 | 10.30 | 9.88 | 10.97 |
| EMC4 | 3 | 10.36 | 8.99 | 7.89 | 9.54 | 12.02 | 10.34 | 10.02 | 10.50 | 10.19 | 10.15 | 9.67 | 8.72 | 11.18 | 10.26 | 10.17 |
| EMC7 | 6 | 10.91 | 6.90 | 8.43 | 10.87 | 10.33 | 8.96 | 10.72 | 8.82 | 10.59 | 13.46 | 8.91 | 9.65 | 9.65 | 9.77 | 12.03 |
| EMC8 | 7 | 9.34 | 11.59 | 9.23 | 9.21 | 10.15 | 10.55 | 11.20 | 10.37 | 9.15 | 9.21 | 10.47 | 9.22 | 10.35 | 10.79 | 9.18 |
| EMC9 | 2 | 10.82 | 11.12 | 9.08 | 10.78 | 11.44 | 12.97 | 6.86 | 6.83 | 9.23 | 10.86 | 10.97 | 9.93 | 12.21 | 6.85 | 10.05 |
| EMD | 10 | 12.63 | 5.88 | 6.51 | 9.58 | 10.56 | 8.76 | 12.09 | 9.97 | 10.75 | 13.28 | 9.25 | 8.04 | 9.66 | 11.03 | 12.01 |
| EME1 | 1 | 11.30 | 10.27 | 11.98 | 11.99 | 7.44 | 7.77 | 9.93 | 7.10 | 11.18 | 11.02 | 10.79 | 11.99 | 7.61 | 8.52 | 11.10 |
| EMG1 | 12 | 10.11 | 9.31 | 10.87 | 10.58 | 8.20 | 8.76 | 9.93 | 13.95 | 9.31 | 8.97 | 9.71 | 10.72 | 8.48 | 11.94 | 9.14 |
| EML1 | 1 | 10.43 | 10.55 | 3.61 | 6.16 | 10.90 | 12.94 | 12.49 | 20.51 | 6.49 | 5.93 | 10.49 | 4.88 | 11.92 | 16.50 | 6.21 |
| EML2 | 9 | 10.04 | 10.95 | 7.83 | 7.58 | 11.55 | 11.58 | 10.81 | 11.08 | 9.41 | 9.16 | 10.49 | 7.71 | 11.57 | 10.94 | 9.29 |
| EML3 | 8 | 9.10 | 9.30 | 10.20 | 9.57 | 10.46 | 10.86 | 9.97 | 9.30 | 11.00 | 10.24 | 9.20 | 9.88 | 10.66 | 9.64 | 10.62 |
| EML4 | 14 | 8.62 | 9.22 | 11.89 | 11.41 | 10.09 | 10.25 | 8.33 | 8.18 | 10.95 | 11.06 | 8.92 | 11.65 | 10.17 | 8.26 | 11.00 |
| EMSY | 5 | 9.00 | 9.48 | 11.12 | 9.36 | 9.77 | 9.90 | 11.97 | 11.35 | 9.47 | 8.57 | 9.24 | 10.24 | 9.84 | 11.66 | 9.02 |

| | | | | | | | | | | | | | | | | |
|---|---|---|---|---|---|---|---|---|---|---|---|---|---|---|---|---|
| ENAH | 27 | 7.98 | 10.48 | 14.90 | 13.57 | 9.28 | 9.19 | 8.03 | 6.89 | 9.63 | 10.05 | 9.23 | 14.24 | 9.24 | 7.46 | 9.84 |
| ENDOD1 | 1 | 11.75 | 8.64 | 11.35 | 16.30 | 6.92 | 6.36 | 7.64 | 10.11 | 9.50 | 11.44 | 10.19 | 13.83 | 6.64 | 8.87 | 10.47 |
| ENDOG | 3 | 9.81 | 9.31 | 7.85 | 7.16 | 8.82 | 9.21 | 14.81 | 12.67 | 10.39 | 9.97 | 9.56 | 7.50 | 9.02 | 13.74 | 10.18 |
| ENG | 7 | 12.30 | 10.48 | 9.36 | 9.73 | 9.29 | 8.45 | 8.44 | 7.56 | 11.95 | 12.43 | 11.39 | 9.55 | 8.87 | 8.00 | 12.19 |
| ENGASE | 3 | 10.19 | 8.93 | 10.15 | 10.68 | 9.11 | 9.25 | 11.09 | 12.70 | 8.41 | 9.48 | 9.56 | 10.42 | 9.18 | 11.90 | 8.95 |
| ENO1 | 223 | 9.84 | 14.82 | 9.48 | 8.84 | 8.17 | 8.20 | 10.21 | 10.04 | 10.25 | 10.16 | 12.33 | 9.16 | 8.19 | 10.12 | 10.20 |
| ENO2 | 11 | 9.98 | 10.42 | 6.77 | 4.87 | 12.58 | 11.36 | 8.29 | 17.10 | 9.59 | 9.05 | 10.20 | 5.82 | 11.97 | 12.69 | 9.32 |
| ENO3 | 49 | 9.75 | 10.94 | 5.02 | 4.00 | 13.17 | 13.12 | 5.76 | 14.94 | 11.89 | 11.41 | 10.35 | 4.51 | 13.15 | 10.35 | 11.65 |
| ENOPH1 | 5 | 10.51 | 11.07 | 9.13 | 8.26 | 10.25 | 10.67 | 10.45 | 9.17 | 10.36 | 10.13 | 10.79 | 8.70 | 10.46 | 9.81 | 10.24 |
| ENOSF1 | 2 | 12.23 | 13.04 | 12.64 | 13.34 | 5.97 | 6.14 | 6.77 | 5.74 | 12.22 | 11.90 | 12.64 | 12.99 | 6.06 | 6.26 | 12.06 |
| ENSA | 25 | 8.88 | 9.81 | 11.35 | 11.62 | 12.10 | 12.91 | 9.53 | 6.06 | 8.88 | 8.86 | 9.34 | 11.49 | 12.51 | 7.80 | 8.87 |
| ENTHD2 | 2 | 9.98 | 10.68 | 10.84 | 11.28 | 9.75 | 9.07 | 9.44 | 9.31 | 10.30 | 9.35 | 10.33 | 11.06 | 9.41 | 9.38 | 9.83 |
| ENTPD5 | 1 | 8.56 | 7.14 | 4.11 | 6.80 | 15.04 | 15.27 | 9.99 | 9.10 | 12.59 | 11.41 | 7.85 | 5.45 | 15.15 | 9.55 | 12.00 |
| ENY2 | 3 | 9.64 | 10.51 | 9.20 | 9.11 | 9.61 | 9.51 | 11.80 | 11.35 | 9.95 | 9.32 | 10.08 | 9.16 | 9.56 | 11.57 | 9.63 |
| EOGT | 2 | 8.85 | 7.42 | 9.85 | 11.32 | 9.28 | 9.04 | 10.70 | 10.38 | 11.14 | 12.02 | 8.14 | 10.58 | 9.16 | 10.54 | 11.58 |
| EP300 | 3 | 9.37 | 10.51 | 11.96 | 10.68 | 8.08 | 8.22 | 11.15 | 10.75 | 10.29 | 9.00 | 9.94 | 11.32 | 8.15 | 10.95 | 9.65 |
| EP400 | 10 | 9.62 | 10.50 | 9.86 | 9.49 | 10.19 | 10.23 | 10.99 | 10.16 | 9.64 | 9.31 | 10.06 | 9.67 | 10.21 | 10.58 | 9.48 |
| EPB41 | 17 | 12.38 | 13.76 | 9.91 | 9.27 | 9.28 | 9.79 | 9.48 | 8.44 | 9.04 | 8.64 | 13.07 | 9.59 | 9.54 | 8.96 | 8.84 |
| EPB41L1 | 31 | 8.43 | 9.24 | 13.20 | 11.82 | 11.48 | 11.38 | 3.12 | 2.73 | 14.57 | 14.02 | 8.84 | 12.51 | 11.43 | 2.92 | 14.30 |
| EPB41L2 | 47 | 9.18 | 10.38 | 12.98 | 12.02 | 8.21 | 8.31 | 8.96 | 10.17 | 10.20 | 9.58 | 9.78 | 12.50 | 8.26 | 9.56 | 9.89 |
| EPDR1 | 2 | 11.82 | 12.19 | 15.94 | 12.70 | 5.33 | 4.06 | 9.23 | 8.15 | 10.64 | 9.96 | 12.00 | 14.32 | 4.69 | 8.69 | 10.30 |
| EPHA2 | 7 | 8.57 | 6.89 | 8.68 | 10.72 | 5.04 | 5.61 | 6.43 | 6.39 | 20.35 | 21.32 | 7.73 | 9.70 | 5.32 | 6.41 | 20.83 |
| EPHA5 | 2 | 11.61 | 11.26 | 8.00 | 11.53 | 8.28 | 8.22 | 7.02 | 8.39 | 12.37 | 13.32 | 11.44 | 9.76 | 8.25 | 7.71 | 12.85 |
| EPHA7 | 3 | 11.01 | 8.29 | 5.09 | 5.82 | 2.44 | 2.32 | 1.45 | 0.83 | 30.53 | 32.21 | 9.65 | 5.45 | 2.38 | 1.14 | 31.37 |
| EPHB4 | 2 | 10.62 | 9.17 | 7.69 | 9.81 | 8.26 | 7.93 | 9.12 | 9.36 | 13.38 | 14.66 | 9.89 | 8.75 | 8.09 | 9.24 | 14.02 |
| EPHB6 | 2 | 10.39 | 8.93 | 9.68 | 10.55 | 8.80 | 9.06 | 10.95 | 11.31 | 10.29 | 10.04 | 9.66 | 10.11 | 8.93 | 11.13 | 10.17 |
| EPHX1 | 15 | 15.48 | 8.49 | 5.27 | 7.36 | 11.60 | 9.75 | 8.68 | 8.10 | 11.26 | 14.01 | 11.99 | 6.31 | 10.68 | 8.39 | 12.63 |
| EPN1 | 11 | 9.18 | 10.30 | 9.28 | 8.67 | 11.30 | 11.98 | 12.68 | 11.49 | 7.96 | 7.16 | 9.74 | 8.98 | 11.64 | 12.08 | 7.56 |
| EPN2 | 5 | 9.57 | 9.37 | 11.53 | 12.11 | 8.23 | 8.13 | 10.45 | 8.76 | 10.93 | 10.91 | 9.47 | 11.82 | 8.18 | 9.61 | 10.92 |
| EPPK1 | 212 | 9.36 | 10.19 | 11.03 | 10.48 | 12.04 | 12.27 | 5.23 | 5.75 | 12.11 | 11.55 | 9.77 | 10.76 | 12.16 | 5.49 | 11.83 |
| EPRS | 118 | 9.34 | 9.69 | 9.01 | 8.24 | 11.70 | 11.86 | 9.42 | 11.61 | 9.91 | 9.24 | 9.51 | 8.63 | 11.78 | 10.51 | 9.57 |
| EPS15 | 24 | 9.05 | 10.65 | 11.39 | 10.93 | 9.39 | 9.64 | 10.03 | 9.45 | 9.95 | 9.53 | 9.85 | 11.16 | 9.52 | 9.74 | 9.74 |
| EPS15L1 | 25 | 9.67 | 10.94 | 9.87 | 9.96 | 9.20 | 9.56 | 10.05 | 9.80 | 10.78 | 10.18 | 10.30 | 9.91 | 9.38 | 9.92 | 10.48 |
| EPS8 | 12 | 10.35 | 11.30 | 14.20 | 14.90 | 7.07 | 7.08 | 6.97 | 6.14 | 11.47 | 10.52 | 10.82 | 14.55 | 7.08 | 6.56 | 10.99 |
| EPS8L2 | 14 | 8.94 | 10.81 | 8.26 | 8.49 | 12.39 | 13.01 | 7.41 | 6.88 | 12.16 | 11.67 | 9.87 | 8.37 | 12.70 | 7.14 | 11.92 |
| ERAL1 | 9 | 11.69 | 9.73 | 11.47 | 12.72 | 8.21 | 7.29 | 8.60 | 8.24 | 10.73 | 11.32 | 10.71 | 12.09 | 7.75 | 8.42 | 11.02 |
| ERAP1 | 4 | 11.30 | 9.18 | 8.28 | 8.99 | 12.54 | 11.22 | 8.11 | 11.77 | 8.37 | 10.25 | 10.24 | 8.64 | 11.88 | 9.94 | 9.31 |
| ERBB2IP | 17 | 11.57 | 10.09 | 9.64 | 11.30 | 9.66 | 9.38 | 9.12 | 8.20 | 10.32 | 10.73 | 10.83 | 10.47 | 9.52 | 8.66 | 10.52 |
| ERC1 | 51 | 8.82 | 9.88 | 11.74 | 11.08 | 10.51 | 10.84 | 9.88 | 9.04 | 9.38 | 8.84 | 9.35 | 11.41 | 10.67 | 9.46 | 9.11 |
| ERCC1 | 3 | 10.11 | 11.98 | 10.85 | 9.59 | 9.66 | 9.69 | 9.69 | 9.72 | 9.75 | 8.96 | 11.05 | 10.22 | 9.68 | 9.70 | 9.36 |
| ERCC2 | 9 | 9.76 | 9.41 | 9.95 | 10.23 | 9.13 | 9.79 | 9.45 | 9.82 | 11.28 | 11.19 | 9.58 | 10.09 | 9.46 | 9.64 | 11.23 |
| ERCC3 | 7 | 9.03 | 9.54 | 11.25 | 10.41 | 9.29 | 9.45 | 9.29 | 9.73 | 10.90 | 11.12 | 9.28 | 10.83 | 9.37 | 9.51 | 11.01 |

| | | | | | | | | | | | | | | | | |
|---|---|---|---|---|---|---|---|---|---|---|---|---|---|---|---|---|
| ERCC4 | 5 | 9.10 | 10.13 | 10.38 | 9.95 | 10.53 | 10.76 | 9.96 | 11.05 | 8.99 | 9.17 | 9.61 | 10.16 | 10.64 | 10.50 | 9.08 |
| ERCC5 | 10 | 9.31 | 10.27 | 9.22 | 9.09 | 11.50 | 11.93 | 10.87 | 11.08 | 8.78 | 7.94 | 9.79 | 9.15 | 11.71 | 10.98 | 8.36 |
| ERCC6 | 3 | 9.83 | 10.89 | 10.66 | 9.74 | 9.44 | 10.25 | 9.32 | 9.13 | 10.56 | 10.19 | 10.36 | 10.20 | 9.85 | 9.22 | 10.37 |
| ERCC6L | 9 | 7.75 | 8.24 | 11.53 | 11.00 | 9.18 | 9.18 | 14.01 | 12.80 | 8.50 | 7.82 | 7.99 | 11.26 | 9.18 | 13.40 | 8.16 |
| ERF | 6 | 8.80 | 9.82 | 12.86 | 11.07 | 8.72 | 8.45 | 9.24 | 8.19 | 11.62 | 11.22 | 9.31 | 11.96 | 8.58 | 8.72 | 11.42 |
| ERGIC1 | 9 | 10.68 | 6.24 | 8.69 | 9.39 | 11.18 | 8.76 | 12.37 | 10.26 | 9.96 | 12.47 | 8.46 | 9.04 | 9.97 | 11.31 | 11.21 |
| ERGIC2 | 2 | 10.53 | 6.79 | 8.37 | 11.90 | 10.15 | 9.94 | 9.46 | 10.47 | 9.70 | 12.68 | 8.66 | 10.14 | 10.05 | 9.97 | 11.19 |
| ERGIC3 | 5 | 10.55 | 7.56 | 9.51 | 11.98 | 9.79 | 9.35 | 10.25 | 9.71 | 9.99 | 11.30 | 9.06 | 10.74 | 9.57 | 9.98 | 10.65 |
| ERH | 13 | 9.08 | 9.31 | 11.16 | 7.46 | 10.62 | 10.39 | 10.88 | 11.45 | 9.88 | 9.78 | 9.19 | 9.31 | 10.51 | 11.16 | 9.83 |
| ERI1 | 4 | 8.48 | 8.72 | 11.16 | 10.27 | 8.59 | 8.76 | 14.43 | 13.57 | 8.57 | 7.44 | 8.60 | 10.72 | 8.67 | 14.00 | 8.00 |
| ERI3 | 5 | 10.27 | 11.50 | 8.19 | 7.91 | 9.92 | 10.36 | 9.83 | 10.08 | 10.83 | 11.11 | 10.88 | 8.05 | 10.14 | 9.96 | 10.97 |
| ERICH1 | 2 | 8.73 | 9.29 | 12.79 | 11.61 | 9.79 | 8.51 | 12.22 | 11.11 | 8.21 | 7.74 | 9.01 | 12.20 | 9.15 | 11.67 | 7.98 |
| ERLEC1 | 2 | 7.35 | 16.92 | 7.24 | 14.20 | 9.71 | 9.41 | 9.28 | 9.25 | 7.31 | 9.34 | 12.13 | 10.72 | 9.56 | 9.26 | 8.32 |
| ERLIN1 | 8 | 12.82 | 8.23 | 9.30 | 12.45 | 12.51 | 11.28 | 13.27 | 12.38 | 3.86 | 3.90 | 10.52 | 10.88 | 11.90 | 12.82 | 3.88 |
| ERLIN2 | 13 | 10.40 | 8.76 | 8.18 | 9.61 | 11.77 | 10.65 | 12.90 | 12.33 | 7.69 | 7.72 | 9.58 | 8.90 | 11.21 | 12.61 | 7.71 |
| ERMP1 | 4 | 13.96 | 8.09 | 6.62 | 9.84 | 9.57 | 9.77 | 9.46 | 8.46 | 10.65 | 13.59 | 11.03 | 8.23 | 9.67 | 8.96 | 12.12 |
| ERO1L | 15 | 8.53 | 7.47 | 8.20 | 8.85 | 10.16 | 10.04 | 12.71 | 14.85 | 9.32 | 9.86 | 8.00 | 8.52 | 10.10 | 13.78 | 9.59 |
| ERP29 | 36 | 9.12 | 9.47 | 8.67 | 8.77 | 11.38 | 11.61 | 9.39 | 8.72 | 11.40 | 11.46 | 9.30 | 8.72 | 11.50 | 9.06 | 11.43 |
| ERP44 | 16 | 9.76 | 8.18 | 8.11 | 8.51 | 10.70 | 10.33 | 9.67 | 14.97 | 9.85 | 9.91 | 8.97 | 8.31 | 10.52 | 12.32 | 9.88 |
| ESD | 9 | 9.56 | 10.51 | 11.92 | 11.21 | 13.00 | 12.44 | 8.89 | 8.36 | 7.16 | 6.95 | 10.03 | 11.57 | 12.72 | 8.62 | 7.06 |
| ESF1 | 16 | 9.26 | 10.03 | 11.87 | 11.16 | 8.08 | 8.62 | 11.55 | 10.64 | 9.76 | 9.03 | 9.64 | 11.51 | 8.35 | 11.10 | 9.40 |
| ESPL1 | 2 | 8.21 | 8.99 | 9.62 | 9.78 | 8.75 | 8.73 | 11.07 | 11.76 | 10.77 | 12.32 | 8.60 | 9.70 | 8.74 | 11.41 | 11.55 |
| ESYT1 | 30 | 11.79 | 8.59 | 8.05 | 9.44 | 9.37 | 9.17 | 7.79 | 7.06 | 13.95 | 14.80 | 10.19 | 8.75 | 9.27 | 7.42 | 14.38 |
| ESYT2 | 12 | 11.65 | 7.76 | 6.24 | 8.43 | 9.31 | 8.77 | 12.28 | 11.99 | 10.88 | 12.69 | 9.70 | 7.33 | 9.04 | 12.13 | 11.79 |
| ETF1 | 32 | 9.29 | 10.00 | 9.83 | 8.90 | 9.72 | 10.01 | 10.26 | 12.03 | 10.01 | 9.94 | 9.65 | 9.37 | 9.86 | 11.15 | 9.97 |
| ETFA | 20 | 9.95 | 9.40 | 7.91 | 8.81 | 10.19 | 9.73 | 10.07 | 15.84 | 8.86 | 9.24 | 9.68 | 8.36 | 9.96 | 12.95 | 9.05 |
| ETFB | 26 | 11.05 | 10.36 | 8.64 | 9.19 | 10.93 | 10.05 | 11.11 | 10.69 | 9.04 | 8.94 | 10.70 | 8.92 | 10.49 | 10.90 | 8.99 |
| ETFDH | 4 | 11.70 | 10.12 | 8.09 | 9.93 | 10.88 | 10.25 | 9.96 | 9.44 | 9.45 | 10.17 | 10.91 | 9.01 | 10.57 | 9.70 | 9.81 |
| ETHE1 | 3 | 8.12 | 6.94 | 10.03 | 10.85 | 12.50 | 12.43 | 10.68 | 8.88 | 9.49 | 10.07 | 7.53 | 10.44 | 12.47 | 9.78 | 9.78 |
| ETNK1 | 1 | 11.48 | 11.81 | 10.15 | 9.28 | 11.02 | 8.51 | 10.23 | 8.34 | 10.64 | 8.54 | 11.65 | 9.71 | 9.76 | 9.28 | 9.59 |
| EVI5 | 1 | 9.31 | 10.05 | 10.84 | 10.29 | 9.82 | 10.99 | 11.25 | 9.99 | 8.57 | 8.88 | 9.68 | 10.57 | 10.41 | 10.62 | 8.73 |
| EVPL | 23 | 11.06 | 12.26 | 9.03 | 9.01 | 11.89 | 12.30 | 5.56 | 5.26 | 12.01 | 11.62 | 11.66 | 9.02 | 12.09 | 5.41 | 11.82 |
| EWSR1 | 6 | 11.45 | 12.61 | 14.48 | 12.86 | 0.87 | 0.74 | 11.43 | 15.10 | 10.30 | 10.16 | 12.03 | 13.67 | 0.80 | 13.27 | 10.23 |
| EXD2 | 1 | 12.47 | 8.46 | 10.09 | 12.75 | 8.67 | 8.48 | 11.41 | 10.30 | 8.82 | 8.55 | 10.46 | 11.42 | 8.57 | 10.86 | 8.69 |
| EXOC1 | 17 | 9.46 | 10.06 | 11.35 | 10.72 | 10.32 | 11.01 | 9.31 | 9.17 | 9.78 | 8.81 | 9.76 | 11.04 | 10.66 | 9.24 | 9.30 |
| EXOC2 | 21 | 10.37 | 10.63 | 11.76 | 11.63 | 10.65 | 11.44 | 9.97 | 9.76 | 7.02 | 6.77 | 10.50 | 11.70 | 11.04 | 9.87 | 6.89 |
| EXOC3 | 15 | 9.78 | 10.09 | 11.65 | 11.42 | 10.78 | 10.31 | 9.33 | 8.95 | 8.86 | 8.82 | 9.94 | 11.54 | 10.54 | 9.14 | 8.84 |
| EXOC4 | 21 | 9.90 | 10.30 | 10.80 | 10.53 | 10.82 | 10.60 | 9.01 | 9.27 | 9.44 | 9.33 | 10.10 | 10.66 | 10.71 | 9.14 | 9.38 |
| EXOC5 | 8 | 10.31 | 10.67 | 11.00 | 11.25 | 10.49 | 11.19 | 8.65 | 8.68 | 8.86 | 8.90 | 10.49 | 11.13 | 10.84 | 8.66 | 8.88 |
| EXOC6 | 3 | 9.52 | 10.32 | 11.94 | 11.74 | 12.78 | 11.67 | 7.93 | 7.23 | 8.53 | 8.33 | 9.92 | 11.84 | 12.23 | 7.58 | 8.43 |
| EXOC6B | 5 | 10.53 | 10.96 | 11.60 | 11.31 | 9.23 | 8.70 | 9.55 | 9.55 | 9.33 | 9.23 | 10.74 | 11.46 | 8.97 | 9.55 | 9.28 |
| EXOC7 | 20 | 10.17 | 11.18 | 11.97 | 12.48 | 9.80 | 10.16 | 8.68 | 8.14 | 8.96 | 8.45 | 10.68 | 12.22 | 9.98 | 8.41 | 8.71 |

| | | | | | | | | | | | | | | | | |
|---|---|---|---|---|---|---|---|---|---|---|---|---|---|---|---|---|
| EXOC8 | 11 | 9.30 | 10.98 | 11.02 | 11.28 | 11.08 | 11.77 | 8.92 | 8.60 | 8.36 | 8.68 | 10.14 | 11.15 | 11.42 | 8.76 | 8.52 |
| EXOG | 2 | 14.23 | 7.00 | 4.50 | 7.26 | 15.86 | 12.72 | 8.53 | 6.98 | 10.06 | 12.86 | 10.61 | 5.88 | 14.29 | 7.76 | 11.46 |
| EXOSC1 | 5 | 9.87 | 10.51 | 9.30 | 9.01 | 10.73 | 11.55 | 9.27 | 9.04 | 10.51 | 10.21 | 10.19 | 9.15 | 11.14 | 9.16 | 10.36 |
| EXOSC10 | 32 | 10.35 | 11.20 | 8.23 | 8.02 | 11.13 | 11.43 | 9.68 | 9.11 | 10.74 | 10.10 | 10.78 | 8.12 | 11.28 | 9.39 | 10.42 |
| EXOSC2 | 13 | 9.59 | 10.86 | 8.99 | 8.68 | 10.65 | 11.17 | 9.97 | 9.51 | 10.51 | 10.06 | 10.23 | 8.84 | 10.91 | 9.74 | 10.29 |
| EXOSC3 | 8 | 10.61 | 11.33 | 8.49 | 7.72 | 10.82 | 11.34 | 9.14 | 8.77 | 11.09 | 10.68 | 10.97 | 8.10 | 11.08 | 8.96 | 10.89 |
| EXOSC4 | 11 | 9.31 | 10.50 | 9.32 | 9.59 | 10.35 | 11.20 | 9.64 | 9.57 | 10.32 | 10.20 | 9.91 | 9.45 | 10.77 | 9.60 | 10.26 |
| EXOSC5 | 7 | 10.59 | 10.90 | 8.60 | 8.53 | 11.06 | 11.34 | 9.48 | 9.56 | 10.15 | 9.80 | 10.74 | 8.57 | 11.20 | 9.52 | 9.97 |
| EXOSC6 | 7 | 10.17 | 11.32 | 8.04 | 7.42 | 11.65 | 12.25 | 9.65 | 8.49 | 10.74 | 10.27 | 10.74 | 7.73 | 11.95 | 9.07 | 10.51 |
| EXOSC7 | 11 | 9.27 | 10.29 | 8.99 | 9.09 | 10.31 | 11.25 | 9.81 | 10.76 | 10.22 | 10.01 | 9.78 | 9.04 | 10.78 | 10.29 | 10.11 |
| EXOSC8 | 8 | 10.62 | 10.59 | 9.30 | 9.27 | 10.96 | 11.55 | 9.11 | 8.34 | 10.53 | 9.73 | 10.61 | 9.29 | 11.26 | 8.72 | 10.13 |
| EXOSC9 | 9 | 9.03 | 9.72 | 7.88 | 7.65 | 10.92 | 11.51 | 9.52 | 16.06 | 8.87 | 8.84 | 9.38 | 7.76 | 11.22 | 12.79 | 8.86 |
| EYA3 | 3 | 8.92 | 9.46 | 9.96 | 10.49 | 8.19 | 8.18 | 11.03 | 10.94 | 11.69 | 11.14 | 9.19 | 10.23 | 8.18 | 10.99 | 11.42 |
| EYA4 | 2 | 9.64 | 11.42 | 10.26 | 11.04 | 10.64 | 11.62 | 7.46 | 8.47 | 10.30 | 9.15 | 10.53 | 10.65 | 11.13 | 7.97 | 9.72 |
| EZH2 | 5 | 10.44 | 11.07 | 8.47 | 7.89 | 8.25 | 7.81 | 12.27 | 11.98 | 11.11 | 10.73 | 10.75 | 8.18 | 8.03 | 12.13 | 10.92 |
| EZR | 141 | 9.07 | 10.35 | 12.07 | 11.65 | 9.50 | 9.54 | 7.22 | 6.90 | 11.98 | 11.72 | 9.71 | 11.86 | 9.52 | 7.06 | 11.85 |
| F3 | 2 | 8.65 | 7.38 | 6.24 | 8.48 | 6.52 | 7.06 | 2.86 | 2.63 | 23.94 | 26.25 | 8.01 | 7.36 | 6.79 | 2.74 | 25.10 |
| F8A1 | 3 | 10.28 | 9.96 | 8.63 | 8.82 | 12.20 | 12.66 | 9.60 | 8.91 | 9.33 | 9.62 | 10.12 | 8.72 | 12.43 | 9.26 | 9.48 |
| FAAP100 | 3 | 9.90 | 10.95 | 12.68 | 12.66 | 8.58 | 8.67 | 7.96 | 7.60 | 10.96 | 10.03 | 10.43 | 12.67 | 8.63 | 7.78 | 10.49 |
| FABP3 | 2 | 6.63 | 7.23 | 28.44 | 26.93 | 5.89 | 6.14 | 3.42 | 3.10 | 5.97 | 6.25 | 6.93 | 27.69 | 6.01 | 3.26 | 6.11 |
| FABP5 | 12 | 6.98 | 7.96 | 9.50 | 8.77 | 7.33 | 7.38 | 11.83 | 26.21 | 7.66 | 6.38 | 7.47 | 9.13 | 7.36 | 19.02 | 7.02 |
| FADD | 4 | 7.91 | 10.49 | 10.23 | 9.81 | 10.33 | 10.78 | 10.29 | 9.67 | 10.29 | 10.19 | 9.20 | 10.02 | 10.56 | 9.98 | 10.24 |
| FADS1 | 3 | 12.32 | 6.94 | 8.54 | 11.19 | 10.78 | 9.57 | 9.40 | 7.93 | 10.86 | 12.49 | 9.63 | 9.86 | 10.17 | 8.66 | 11.67 |
| FADS2 | 4 | 16.36 | 9.26 | 11.70 | 19.24 | 6.67 | 6.14 | 4.76 | 4.49 | 9.44 | 11.94 | 12.81 | 15.47 | 6.41 | 4.62 | 10.69 |
| FAF1 | 22 | 10.68 | 10.94 | 10.08 | 9.00 | 8.24 | 8.18 | 11.89 | 10.63 | 10.33 | 10.03 | 10.81 | 9.54 | 8.21 | 11.26 | 10.18 |
| FAF2 | 9 | 11.59 | 7.04 | 7.99 | 10.38 | 9.19 | 8.35 | 9.94 | 9.52 | 12.14 | 13.86 | 9.31 | 9.18 | 8.77 | 9.73 | 13.00 |
| FAH | 9 | 9.14 | 9.75 | 11.69 | 9.90 | 10.62 | 10.17 | 9.07 | 7.82 | 11.13 | 10.73 | 9.44 | 10.79 | 10.39 | 8.44 | 10.93 |
| FAHD1 | 8 | 10.26 | 9.06 | 10.11 | 9.68 | 10.90 | 10.63 | 7.61 | 10.94 | 10.67 | 10.15 | 9.66 | 9.89 | 10.77 | 9.27 | 10.41 |
| FAHD2A | 8 | 11.70 | 11.13 | 10.46 | 10.87 | 7.85 | 7.25 | 10.84 | 9.70 | 9.94 | 10.26 | 11.42 | 10.67 | 7.55 | 10.27 | 10.10 |
| FAIM | 5 | 12.15 | 12.56 | 7.72 | 7.09 | 12.17 | 12.72 | 10.01 | 8.89 | 8.56 | 8.13 | 12.36 | 7.40 | 12.44 | 9.45 | 8.35 |
| FAM102B | 3 | 11.09 | 9.92 | 8.46 | 7.77 | 10.01 | 11.13 | 13.51 | 7.45 | 11.40 | 9.27 | 10.51 | 8.11 | 10.57 | 10.48 | 10.34 |
| FAM103A1 | 2 | 8.45 | 9.60 | 13.59 | 12.85 | 10.17 | 11.05 | 7.83 | 5.09 | 10.28 | 11.09 | 9.02 | 13.22 | 10.61 | 6.46 | 10.69 |
| FAM105A | 2 | 9.74 | 5.65 | 9.97 | 16.74 | 11.53 | 9.26 | 4.84 | 4.85 | 11.90 | 15.52 | 7.70 | 13.35 | 10.39 | 4.85 | 13.71 |
| FAM105B | 9 | 9.21 | 9.69 | 8.33 | 7.91 | 10.98 | 11.17 | 9.07 | 10.83 | 11.64 | 11.16 | 9.45 | 8.12 | 11.08 | 9.95 | 11.40 |
| FAM107B | 12 | 7.26 | 8.58 | 13.78 | 12.43 | 10.02 | 10.31 | 11.94 | 9.67 | 8.07 | 7.94 | 7.92 | 13.11 | 10.16 | 10.81 | 8.00 |
| FAM111A | 1 | 10.11 | 12.62 | 5.58 | 5.03 | 14.22 | 12.99 | 7.72 | 9.89 | 9.40 | 12.45 | 11.36 | 5.30 | 13.61 | 8.80 | 10.92 |
| FAM111B | 5 | 10.07 | 10.63 | 9.44 | 9.65 | 10.88 | 11.07 | 10.24 | 9.70 | 9.34 | 8.99 | 10.35 | 9.54 | 10.97 | 9.97 | 9.17 |
| FAM114A1 | 12 | 11.20 | 11.50 | 8.29 | 9.04 | 9.97 | 10.87 | 8.88 | 8.61 | 11.18 | 10.46 | 11.35 | 8.66 | 10.42 | 8.74 | 10.82 |
| FAM114A2 | 12 | 9.99 | 10.85 | 8.69 | 8.34 | 10.11 | 10.95 | 9.54 | 9.02 | 11.41 | 11.10 | 10.42 | 8.51 | 10.53 | 9.28 | 11.25 |
| FAM115A | 7 | 11.32 | 12.18 | 12.93 | 13.26 | 9.29 | 9.69 | 7.96 | 7.46 | 7.90 | 7.99 | 11.75 | 13.10 | 9.49 | 7.71 | 7.95 |
| FAM117B | 3 | 7.91 | 8.80 | 10.50 | 9.97 | 9.26 | 9.27 | 16.46 | 15.05 | 6.41 | 6.36 | 8.36 | 10.24 | 9.27 | 15.75 | 6.39 |
| FAM118B | 1 | 8.46 | 9.79 | 12.13 | 12.42 | 9.73 | 9.13 | 9.61 | 9.20 | 9.90 | 9.62 | 9.13 | 12.28 | 9.43 | 9.40 | 9.76 |

| | | | | | | | | | | | | | | | | |
|---|---|---|---|---|---|---|---|---|---|---|---|---|---|---|---|---|
| FAM120A | 23 | 9.02 | 9.89 | 9.42 | 9.43 | 12.45 | 12.50 | 10.22 | 10.23 | 8.34 | 8.49 | 9.45 | 9.43 | 12.47 | 10.23 | 8.42 |
| FAM120B | 2 | 9.13 | 9.95 | 10.45 | 9.05 | 9.65 | 8.84 | 8.37 | 8.42 | 13.32 | 12.81 | 9.54 | 9.75 | 9.25 | 8.39 | 13.07 |
| FAM120C | 2 | 7.47 | 7.59 | 6.41 | 6.85 | 9.60 | 10.12 | 21.57 | 19.39 | 5.45 | 5.56 | 7.53 | 6.63 | 9.86 | 20.48 | 5.50 |
| FAM122A | 1 | 9.97 | 10.64 | 10.39 | 11.84 | 10.03 | 12.54 | 9.05 | 7.35 | 7.49 | 10.70 | 10.30 | 11.12 | 11.29 | 8.20 | 9.10 |
| FAM127C | 3 | 10.34 | 11.77 | 7.62 | 7.89 | 13.81 | 14.94 | 8.41 | 8.32 | 8.48 | 8.42 | 11.05 | 7.76 | 14.38 | 8.37 | 8.45 |
| FAM129A | 5 | 9.23 | 10.40 | 12.16 | 13.26 | 9.21 | 10.24 | 7.27 | 6.86 | 11.10 | 10.27 | 9.81 | 12.71 | 9.73 | 7.06 | 10.68 |
| FAM129B | 40 | 8.41 | 8.65 | 10.87 | 10.16 | 9.99 | 10.04 | 10.26 | 10.83 | 10.67 | 10.13 | 8.53 | 10.51 | 10.01 | 10.55 | 10.40 |
| FAM133A | 3 | 9.83 | 9.79 | 11.27 | 10.37 | 9.72 | 9.12 | 11.51 | 12.53 | 8.02 | 7.85 | 9.81 | 10.82 | 9.42 | 12.02 | 7.93 |
| FAM133B | 2 | 9.64 | 9.96 | 13.59 | 11.00 | 6.12 | 5.67 | 11.97 | 10.68 | 11.39 | 10.00 | 9.80 | 12.29 | 5.89 | 11.32 | 10.69 |
| FAM134A | 1 | 8.62 | 5.32 | 7.76 | 11.85 | 10.28 | 10.04 | 9.76 | 8.69 | 11.99 | 15.67 | 6.97 | 9.80 | 10.16 | 9.23 | 13.83 |
| FAM136A | 5 | 9.02 | 8.35 | 10.65 | 9.97 | 9.13 | 8.67 | 11.01 | 19.23 | 7.12 | 6.85 | 8.68 | 10.31 | 8.90 | 15.12 | 6.99 |
| FAM160A2 | 2 | 9.55 | 9.76 | 9.92 | 9.18 | 10.97 | 10.73 | 9.11 | 7.88 | 11.68 | 11.20 | 9.66 | 9.55 | 10.85 | 8.50 | 11.44 |
| FAM160B1 | 4 | 8.02 | 9.35 | 10.77 | 11.12 | 13.54 | 14.21 | 6.97 | 6.72 | 9.45 | 9.87 | 8.68 | 10.94 | 13.87 | 6.84 | 9.66 |
| FAM162A | 10 | 11.09 | 6.31 | 7.27 | 10.72 | 8.46 | 7.23 | 13.36 | 12.67 | 10.17 | 12.72 | 8.70 | 8.99 | 7.84 | 13.02 | 11.45 |
| FAM172A | 3 | 11.04 | 11.46 | 7.96 | 7.68 | 12.56 | 11.13 | 12.00 | 9.33 | 8.90 | 7.95 | 11.25 | 7.82 | 11.84 | 10.66 | 8.43 |
| FAM173B | 3 | 12.93 | 7.58 | 6.59 | 8.71 | 9.56 | 8.69 | 11.09 | 10.65 | 11.50 | 12.69 | 10.26 | 7.65 | 9.13 | 10.87 | 12.10 |
| FAM175B | 9 | 9.10 | 9.61 | 12.04 | 9.56 | 12.07 | 11.84 | 9.78 | 8.26 | 8.88 | 8.88 | 9.35 | 10.80 | 11.95 | 9.02 | 8.88 |
| FAM177A1 | 9 | 10.78 | 10.77 | 9.50 | 8.98 | 12.01 | 12.47 | 7.02 | 6.61 | 11.03 | 10.81 | 10.78 | 9.24 | 12.24 | 6.82 | 10.92 |
| FAM184A | 3 | 9.93 | 11.86 | 8.63 | 9.58 | 9.58 | 11.12 | 7.26 | 6.18 | 13.27 | 12.59 | 10.90 | 9.11 | 10.35 | 6.72 | 12.93 |
| FAM188A | 2 | 10.30 | 11.05 | 11.51 | 12.36 | 7.56 | 7.60 | 8.85 | 8.57 | 11.38 | 10.81 | 10.68 | 11.93 | 7.58 | 8.71 | 11.10 |
| FAM192A | 11 | 10.86 | 11.98 | 8.05 | 8.00 | 11.12 | 11.66 | 10.10 | 7.36 | 10.99 | 9.87 | 11.42 | 8.03 | 11.39 | 8.73 | 10.43 |
| FAM195B | 2 | 13.40 | 12.73 | 9.32 | 9.30 | 8.17 | 9.15 | 5.91 | 5.37 | 14.06 | 12.60 | 13.06 | 9.31 | 8.66 | 5.64 | 13.33 |
| FAM203A | 6 | 10.49 | 11.43 | 7.49 | 6.54 | 11.95 | 12.48 | 11.84 | 10.75 | 8.67 | 8.35 | 10.96 | 7.02 | 12.22 | 11.30 | 8.51 |
| FAM206A | 1 | 9.33 | 11.10 | 12.53 | 11.49 | 7.29 | 8.29 | 10.32 | 8.27 | 10.58 | 10.79 | 10.21 | 12.01 | 7.79 | 9.30 | 10.69 |
| FAM207A | 9 | 10.23 | 11.14 | 10.45 | 9.93 | 8.87 | 9.06 | 11.13 | 10.32 | 9.21 | 9.66 | 10.68 | 10.19 | 8.96 | 10.72 | 9.43 |
| FAM208A | 17 | 10.25 | 11.00 | 9.25 | 9.16 | 10.42 | 11.04 | 9.44 | 9.12 | 10.26 | 10.06 | 10.63 | 9.20 | 10.73 | 9.28 | 10.16 |
| FAM208B | 2 | 13.81 | 11.52 | 8.44 | 10.66 | 9.50 | 8.88 | 10.04 | 9.05 | 8.78 | 9.32 | 12.66 | 9.55 | 9.19 | 9.55 | 9.05 |
| FAM213A | 3 | 10.64 | 8.13 | 7.98 | 9.85 | 12.18 | 12.06 | 13.78 | 11.72 | 6.94 | 6.72 | 9.39 | 8.91 | 12.12 | 12.75 | 6.83 |
| FAM213B | 1 | 10.44 | 11.82 | 9.02 | 8.80 | 8.40 | 8.69 | 11.55 | 10.88 | 10.03 | 10.37 | 11.13 | 8.91 | 8.54 | 11.22 | 10.20 |
| FAM219A | 2 | 10.31 | 6.98 | 8.90 | 8.49 | 10.14 | 9.76 | 7.23 | 17.56 | 10.61 | 10.02 | 8.65 | 8.70 | 9.95 | 12.39 | 10.31 |
| FAM21A | 27 | 9.98 | 11.07 | 10.51 | 9.88 | 10.90 | 11.39 | 8.52 | 7.88 | 10.34 | 9.52 | 10.52 | 10.19 | 11.15 | 8.20 | 9.93 |
| FAM21C | 5 | 11.10 | 10.97 | 9.53 | 10.02 | 10.88 | 11.79 | 8.73 | 8.78 | 9.75 | 8.46 | 11.04 | 9.77 | 11.34 | 8.75 | 9.10 |
| FAM32A | 1 | 5.10 | 4.66 | 14.22 | 4.98 | 12.95 | 9.56 | 11.99 | 10.72 | 11.84 | 13.98 | 4.88 | 9.60 | 11.26 | 11.35 | 12.91 |
| FAM3C | 7 | 10.22 | 6.94 | 7.21 | 10.04 | 7.48 | 6.71 | 15.19 | 15.56 | 9.87 | 10.79 | 8.58 | 8.62 | 7.09 | 15.38 | 10.33 |
| FAM45A | 2 | 10.09 | 9.60 | 8.19 | 8.19 | 11.78 | 11.80 | 9.94 | 9.61 | 10.66 | 10.13 | 9.85 | 8.19 | 11.79 | 9.78 | 10.40 |
| FAM46A | 1 | 11.87 | 11.29 | 8.73 | 6.35 | 16.19 | 14.30 | 10.70 | 7.68 | 6.28 | 6.61 | 11.58 | 7.54 | 15.25 | 9.19 | 6.45 |
| FAM49B | 18 | 11.03 | 10.31 | 10.45 | 8.89 | 10.05 | 9.50 | 10.50 | 8.55 | 10.45 | 10.28 | 10.67 | 9.67 | 9.78 | 9.52 | 10.36 |
| FAM50A | 25 | 9.28 | 10.65 | 8.83 | 8.86 | 9.94 | 11.00 | 12.07 | 11.80 | 8.85 | 8.72 | 9.96 | 8.85 | 10.47 | 11.93 | 8.78 |
| FAM65A | 3 | 9.84 | 10.60 | 7.83 | 7.64 | 11.00 | 11.32 | 10.95 | 9.42 | 11.26 | 10.16 | 10.22 | 7.73 | 11.16 | 10.18 | 10.71 |
| FAM76A | 2 | 9.48 | 11.14 | 8.00 | 7.35 | 9.15 | 9.32 | 12.60 | 12.51 | 10.22 | 10.23 | 10.31 | 7.67 | 9.23 | 12.55 | 10.23 |
| FAM83D | 8 | 9.40 | 10.21 | 9.79 | 9.80 | 9.94 | 10.46 | 9.47 | 8.46 | 11.30 | 11.18 | 9.80 | 9.80 | 10.20 | 8.97 | 11.24 |
| FAM83G | 2 | 9.31 | 10.40 | 11.43 | 10.98 | 7.70 | 8.50 | 9.33 | 9.10 | 11.75 | 11.51 | 9.85 | 11.20 | 8.10 | 9.22 | 11.63 |

| | | | | | | | | | | | | | | | | |
|---|---|---|---|---|---|---|---|---|---|---|---|---|---|---|---|---|
| FAM83H | 12 | 10.61 | 11.61 | 8.88 | 8.90 | 9.43 | 10.49 | 9.15 | 8.76 | 11.38 | 10.79 | 11.11 | 8.89 | 9.96 | 8.96 | 11.08 |
| FAM89B | 1 | 9.51 | 11.57 | 8.76 | 7.83 | 12.20 | 12.91 | 9.33 | 6.62 | 10.42 | 10.84 | 10.54 | 8.30 | 12.56 | 7.97 | 10.63 |
| FAM91A1 | 6 | 9.48 | 9.39 | 11.13 | 10.73 | 9.25 | 9.84 | 11.24 | 11.39 | 9.11 | 8.42 | 9.44 | 10.93 | 9.55 | 11.32 | 8.77 |
| FAM96A | 2 | 9.01 | 9.66 | 10.58 | 9.44 | 11.62 | 10.42 | 8.54 | 9.48 | 10.45 | 10.80 | 9.34 | 10.01 | 11.02 | 9.01 | 10.62 |
| FAM96B | 3 | 9.65 | 11.18 | 11.21 | 12.24 | 9.16 | 9.96 | 9.07 | 3.75 | 12.01 | 11.77 | 10.41 | 11.72 | 9.56 | 6.41 | 11.89 |
| FAM98A | 13 | 9.24 | 9.71 | 13.27 | 11.03 | 7.92 | 7.56 | 9.20 | 12.56 | 9.78 | 9.73 | 9.47 | 12.15 | 7.74 | 10.88 | 9.75 |
| FAM98B | 13 | 8.16 | 10.27 | 13.19 | 10.30 | 8.87 | 8.54 | 10.57 | 10.38 | 9.83 | 9.89 | 9.21 | 11.75 | 8.71 | 10.47 | 9.86 |
| FANCA | 1 | 9.13 | 9.19 | 7.29 | 5.57 | 11.38 | 11.72 | 9.27 | 8.09 | 12.20 | 16.18 | 9.16 | 6.43 | 11.55 | 8.68 | 14.19 |
| FANCB | 2 | 9.68 | 12.70 | 9.10 | 9.56 | 9.41 | 11.10 | 11.29 | 9.60 | 8.42 | 9.15 | 11.19 | 9.33 | 10.25 | 10.44 | 8.78 |
| FANCD2 | 25 | 10.00 | 9.73 | 7.52 | 7.52 | 9.74 | 9.32 | 8.82 | 9.32 | 13.60 | 14.44 | 9.86 | 7.52 | 9.53 | 9.07 | 14.02 |
| FANCE | 1 | 7.53 | 14.02 | 8.57 | 6.76 | 10.07 | 13.80 | 7.12 | 7.27 | 13.30 | 11.57 | 10.77 | 7.66 | 11.94 | 7.19 | 12.44 |
| FANCG | 1 | 11.08 | 12.61 | 5.76 | 8.78 | 11.49 | 10.30 | 8.95 | 8.11 | 10.98 | 11.95 | 11.84 | 7.27 | 10.90 | 8.53 | 11.46 |
| FANCI | 30 | 10.15 | 10.18 | 7.83 | 8.15 | 9.04 | 9.55 | 9.89 | 9.90 | 12.49 | 12.81 | 10.17 | 7.99 | 9.30 | 9.90 | 12.65 |
| FAR1 | 3 | 12.24 | 6.70 | 7.57 | 11.20 | 7.79 | 6.93 | 10.28 | 10.67 | 12.06 | 14.56 | 9.47 | 9.39 | 7.36 | 10.48 | 13.31 |
| FARP1 | 11 | 11.99 | 12.20 | 11.93 | 12.09 | 8.63 | 9.11 | 5.93 | 4.98 | 11.64 | 11.49 | 12.09 | 12.01 | 8.87 | 5.46 | 11.56 |
| FARP2 | 4 | 9.42 | 10.00 | 9.21 | 10.20 | 11.38 | 12.33 | 9.49 | 9.29 | 9.41 | 9.28 | 9.71 | 9.70 | 11.85 | 9.39 | 9.34 |
| FARS2 | 1 | 10.05 | 9.26 | 9.79 | 11.39 | 9.48 | 8.98 | 11.38 | 11.15 | 9.46 | 9.06 | 9.66 | 10.59 | 9.23 | 11.26 | 9.26 |
| FARSA | 19 | 9.91 | 10.08 | 9.62 | 8.88 | 9.69 | 10.24 | 11.63 | 10.65 | 9.69 | 9.62 | 9.99 | 9.25 | 9.96 | 11.14 | 9.66 |
| FARSB | 49 | 8.76 | 9.59 | 10.01 | 9.08 | 10.03 | 10.03 | 11.52 | 11.21 | 10.02 | 9.73 | 9.18 | 9.55 | 10.03 | 11.37 | 9.88 |
| FAS | 3 | 8.17 | 7.01 | 9.13 | 12.21 | 8.47 | 8.98 | 7.15 | 6.88 | 14.77 | 17.24 | 7.59 | 10.67 | 8.73 | 7.01 | 16.00 |
| FASN | 179 | 10.86 | 11.33 | 9.04 | 8.03 | 8.67 | 8.45 | 8.82 | 12.39 | 11.22 | 11.19 | 11.09 | 8.53 | 8.56 | 10.61 | 11.20 |
| FASTKD1 | 1 | 9.93 | 11.11 | 8.94 | 11.09 | 7.84 | 9.58 | 8.71 | 8.18 | 11.22 | 13.41 | 10.52 | 10.01 | 8.71 | 8.44 | 12.32 |
| FASTKD2 | 8 | 11.84 | 9.81 | 8.79 | 10.57 | 8.77 | 8.07 | 10.73 | 10.22 | 10.76 | 10.43 | 10.82 | 9.68 | 8.42 | 10.48 | 10.60 |
| FASTKD3 | 3 | 10.78 | 7.38 | 7.16 | 9.46 | 9.11 | 9.09 | 14.07 | 13.47 | 10.17 | 9.31 | 9.08 | 8.31 | 9.10 | 13.77 | 9.74 |
| FASTKD5 | 5 | 10.72 | 8.79 | 9.18 | 11.41 | 9.34 | 9.41 | 9.37 | 9.47 | 10.60 | 11.70 | 9.75 | 10.30 | 9.38 | 9.42 | 11.15 |
| FAT1 | 3 | 8.92 | 7.14 | 8.12 | 9.56 | 9.11 | 8.53 | 16.65 | 15.12 | 8.21 | 8.65 | 8.03 | 8.84 | 8.82 | 15.88 | 8.43 |
| FAU | 6 | 10.29 | 10.76 | 10.13 | 10.13 | 10.51 | 10.03 | 9.52 | 8.90 | 9.78 | 9.96 | 10.52 | 10.13 | 10.27 | 9.21 | 9.87 |
| FBL | 35 | 8.95 | 9.56 | 10.32 | 9.79 | 8.57 | 8.56 | 12.25 | 12.70 | 9.73 | 9.57 | 9.25 | 10.05 | 8.57 | 12.47 | 9.65 |
| FBLIM1 | 1 | 8.95 | 9.17 | 25.72 | 25.55 | 3.15 | 3.27 | 3.39 | 2.20 | 10.07 | 8.51 | 9.06 | 25.64 | 3.21 | 2.80 | 9.29 |
| FBLN1 | 3 | 9.78 | 8.69 | 11.76 | 13.00 | 7.69 | 8.01 | 7.10 | 6.18 | 13.92 | 13.87 | 9.24 | 12.38 | 7.85 | 6.64 | 13.89 |
| FBN1 | 3 | 8.99 | 6.83 | 9.40 | 10.62 | 12.53 | 11.99 | 11.92 | 10.82 | 7.98 | 8.93 | 7.91 | 10.01 | 12.26 | 11.37 | 8.45 |
| FBN2 | 1 | 9.41 | 7.72 | 12.23 | 13.66 | 8.20 | 8.04 | 14.59 | 12.37 | 6.40 | 7.39 | 8.56 | 12.94 | 8.12 | 13.48 | 6.90 |
| FBRS | 1 | 7.97 | 9.19 | 10.78 | 11.94 | 8.31 | 9.00 | 11.59 | 11.05 | 9.03 | 11.14 | 8.58 | 11.36 | 8.66 | 11.32 | 10.08 |
| FBXL18 | 2 | 9.53 | 10.42 | 11.51 | 12.49 | 8.48 | 9.18 | 8.22 | 8.25 | 10.81 | 11.10 | 9.98 | 12.00 | 8.83 | 8.24 | 10.95 |
| FBXO11 | 3 | 9.77 | 10.77 | 10.14 | 9.80 | 9.47 | 8.84 | 11.56 | 11.21 | 9.28 | 9.15 | 10.27 | 9.97 | 9.15 | 11.38 | 9.22 |
| FBXO22 | 10 | 9.54 | 10.21 | 10.16 | 9.35 | 11.99 | 11.66 | 9.42 | 9.24 | 9.38 | 9.06 | 9.87 | 9.75 | 11.82 | 9.33 | 9.22 |
| FBXO28 | 4 | 10.76 | 10.63 | 8.79 | 9.06 | 11.19 | 11.77 | 9.84 | 9.08 | 9.56 | 9.32 | 10.70 | 8.93 | 11.48 | 9.46 | 9.44 |
| FBXO3 | 7 | 10.02 | 9.86 | 10.54 | 9.46 | 12.54 | 12.35 | 9.59 | 9.84 | 8.18 | 7.63 | 9.94 | 10.00 | 12.44 | 9.71 | 7.90 |
| FBXO30 | 3 | 8.40 | 11.34 | 12.77 | 12.39 | 9.17 | 9.03 | 10.83 | 10.68 | 7.91 | 7.49 | 9.87 | 12.58 | 9.10 | 10.76 | 7.70 |
| FBXO38 | 6 | 9.71 | 10.87 | 8.87 | 7.40 | 11.26 | 11.05 | 11.26 | 10.54 | 9.70 | 9.35 | 10.29 | 8.13 | 11.16 | 10.90 | 9.52 |
| FBXO4 | 3 | 8.84 | 9.04 | 6.63 | 5.43 | 10.43 | 11.29 | 16.34 | 14.78 | 8.56 | 8.68 | 8.94 | 6.03 | 10.86 | 15.56 | 8.62 |
| FBXO5 | 1 | 9.09 | 10.45 | 6.97 | 8.92 | 10.18 | 13.59 | 9.76 | 11.81 | 8.81 | 10.42 | 9.77 | 7.95 | 11.88 | 10.78 | 9.62 |

| | | | | | | | | | | | | | | | | |
|---|---|---|---|---|---|---|---|---|---|---|---|---|---|---|---|---|
| FBXO7 | 5 | 10.48 | 10.92 | 10.01 | 9.38 | 8.97 | 8.87 | 11.23 | 11.53 | 9.11 | 9.50 | 10.70 | 9.70 | 8.92 | 11.38 | 9.30 |
| FBXW11 | 5 | 11.28 | 10.87 | 8.27 | 8.14 | 10.62 | 10.20 | 11.03 | 11.31 | 9.17 | 9.11 | 11.08 | 8.21 | 10.41 | 11.17 | 9.14 |
| FBXW5 | 1 | 10.32 | 11.52 | 9.31 | 8.81 | 9.10 | 9.92 | 11.20 | 9.13 | 10.25 | 10.43 | 10.92 | 9.06 | 9.51 | 10.17 | 10.34 |
| FBXW9 | 3 | 8.73 | 9.94 | 9.37 | 9.61 | 9.26 | 10.52 | 11.42 | 11.22 | 9.74 | 10.18 | 9.34 | 9.49 | 9.89 | 11.32 | 9.96 |
| FCF1 | 1 | 9.53 | 10.16 | 9.79 | 10.77 | 7.86 | 7.98 | 12.08 | 12.30 | 9.46 | 10.08 | 9.85 | 10.28 | 7.92 | 12.19 | 9.77 |
| FCHO2 | 6 | 9.72 | 10.56 | 8.98 | 9.20 | 10.19 | 11.11 | 11.14 | 11.04 | 8.88 | 9.20 | 10.14 | 9.09 | 10.65 | 11.09 | 9.04 |
| FDFT1 | 2 | 12.10 | 10.87 | 11.72 | 13.13 | 10.78 | 11.37 | 7.06 | 7.24 | 7.82 | 7.90 | 11.49 | 12.43 | 11.08 | 7.15 | 7.86 |
| FDPS | 20 | 9.77 | 10.68 | 10.34 | 9.47 | 13.09 | 12.62 | 6.92 | 7.75 | 9.60 | 9.75 | 10.23 | 9.91 | 12.85 | 7.34 | 9.67 |
| FDX1 | 2 | 9.20 | 9.89 | 7.11 | 8.74 | 9.93 | 11.84 | 11.92 | 14.53 | 8.25 | 8.57 | 9.55 | 7.93 | 10.89 | 13.23 | 8.41 |
| FDX1L | 2 | 10.03 | 9.75 | 12.15 | 14.04 | 9.94 | 9.59 | 8.75 | 5.44 | 9.94 | 10.37 | 9.89 | 13.10 | 9.76 | 7.10 | 10.15 |
| FDXR | 8 | 11.62 | 10.78 | 9.76 | 10.71 | 9.59 | 9.15 | 6.97 | 6.53 | 12.80 | 12.10 | 11.20 | 10.23 | 9.37 | 6.75 | 12.45 |
| FECH | 8 | 13.81 | 11.56 | 8.85 | 10.54 | 11.13 | 10.02 | 5.99 | 5.24 | 11.03 | 11.83 | 12.69 | 9.70 | 10.58 | 5.61 | 11.43 |
| FEN1 | 32 | 9.60 | 9.50 | 12.18 | 9.43 | 11.29 | 10.70 | 10.07 | 8.42 | 9.57 | 9.25 | 9.55 | 10.80 | 10.99 | 9.24 | 9.41 |
| FER | 4 | 8.78 | 12.39 | 9.89 | 9.86 | 11.17 | 11.83 | 9.89 | 9.55 | 8.41 | 8.23 | 10.58 | 9.88 | 11.50 | 9.72 | 8.32 |
| FER1L4 | 2 | 10.72 | 11.56 | 10.80 | 10.45 | 9.64 | 9.38 | 8.99 | 7.68 | 11.03 | 9.77 | 11.14 | 10.62 | 9.51 | 8.33 | 10.40 |
| FERMT1 | 6 | 10.55 | 11.15 | 11.00 | 11.09 | 7.72 | 7.85 | 10.66 | 10.84 | 9.81 | 9.33 | 10.85 | 11.04 | 7.79 | 10.75 | 9.57 |
| FERMT2 | 43 | 10.18 | 10.71 | 12.66 | 10.38 | 9.22 | 8.26 | 6.60 | 9.03 | 11.67 | 11.28 | 10.45 | 11.52 | 8.74 | 7.82 | 11.48 |
| FEZF1 | 2 | 9.40 | 12.94 | 10.75 | 9.98 | 11.14 | 12.03 | 6.95 | 6.45 | 10.81 | 9.55 | 11.17 | 10.37 | 11.58 | 6.70 | 10.18 |
| FGD4 | 3 | 9.98 | 10.45 | 10.24 | 10.17 | 13.25 | 14.18 | 4.91 | 4.86 | 11.41 | 10.57 | 10.21 | 10.20 | 13.71 | 4.88 | 10.99 |
| FGF2 | 7 | 6.45 | 7.94 | 11.63 | 10.85 | 8.17 | 8.16 | 11.15 | 10.30 | 12.15 | 13.20 | 7.20 | 11.24 | 8.17 | 10.73 | 12.67 |
| FGFR1OP | 4 | 9.69 | 11.40 | 7.99 | 7.32 | 13.22 | 12.97 | 9.56 | 7.76 | 10.18 | 9.93 | 10.54 | 7.65 | 13.09 | 8.66 | 10.05 |
| FH | 41 | 11.60 | 11.14 | 6.20 | 6.66 | 13.72 | 13.50 | 10.52 | 9.99 | 8.49 | 8.17 | 11.37 | 6.43 | 13.61 | 10.26 | 8.33 |
| FHAD1 | 1 | 7.08 | 9.30 | 10.63 | 11.47 | 11.62 | 11.83 | 8.95 | 9.73 | 9.66 | 9.74 | 8.19 | 11.05 | 11.72 | 9.34 | 9.70 |
| FHIT | 3 | 9.52 | 11.46 | 9.84 | 9.72 | 8.80 | 9.10 | 18.01 | 17.77 | 2.95 | 2.82 | 10.49 | 9.78 | 8.95 | 17.89 | 2.89 |
| FHL1 | 7 | 8.20 | 8.26 | 13.40 | 12.17 | 6.57 | 6.19 | 7.90 | 17.83 | 9.82 | 9.67 | 8.23 | 12.78 | 6.38 | 12.86 | 9.75 |
| FHL2 | 5 | 8.16 | 8.82 | 8.11 | 8.01 | 9.65 | 10.13 | 5.74 | 25.00 | 8.46 | 7.92 | 8.49 | 8.06 | 9.89 | 15.37 | 8.19 |
| FHL3 | 7 | 8.55 | 9.60 | 13.78 | 13.40 | 6.03 | 6.25 | 8.04 | 12.18 | 11.58 | 10.60 | 9.07 | 13.59 | 6.14 | 10.11 | 11.09 |
| FHOD1 | 14 | 9.73 | 10.10 | 8.36 | 7.64 | 9.52 | 9.29 | 11.05 | 9.82 | 12.59 | 11.90 | 9.92 | 8.00 | 9.41 | 10.43 | 12.24 |
| FIBP | 6 | 8.97 | 9.55 | 10.42 | 10.45 | 10.69 | 11.18 | 9.96 | 10.03 | 9.12 | 9.63 | 9.26 | 10.43 | 10.94 | 9.99 | 9.38 |
| FIGNL1 | 8 | 10.56 | 10.97 | 11.13 | 10.61 | 10.35 | 10.25 | 8.84 | 8.53 | 9.36 | 9.41 | 10.77 | 10.87 | 10.30 | 8.68 | 9.38 |
| FIP1L1 | 15 | 9.31 | 10.00 | 11.62 | 10.58 | 9.75 | 9.90 | 11.70 | 10.01 | 8.97 | 8.17 | 9.66 | 11.10 | 9.82 | 10.86 | 8.57 |
| FIS1 | 4 | 9.06 | 9.02 | 14.70 | 12.46 | 8.96 | 7.72 | 10.44 | 8.49 | 9.31 | 9.83 | 9.04 | 13.58 | 8.34 | 9.47 | 9.57 |
| FKBP10 | 32 | 10.09 | 8.27 | 8.43 | 8.74 | 11.87 | 11.30 | 10.23 | 9.00 | 11.09 | 10.98 | 9.18 | 8.58 | 11.59 | 9.61 | 11.04 |
| FKBP11 | 4 | 13.54 | 7.08 | 11.14 | 12.26 | 9.68 | 8.02 | 6.52 | 5.30 | 11.08 | 15.37 | 10.31 | 11.70 | 8.85 | 5.91 | 13.23 |
| FKBP14 | 1 | 8.88 | 10.09 | 11.48 | 12.93 | 8.43 | 8.74 | 9.40 | 9.37 | 10.25 | 10.43 | 9.48 | 12.21 | 8.59 | 9.39 | 10.34 |
| FKBP15 | 13 | 9.67 | 11.67 | 9.06 | 9.20 | 11.37 | 12.31 | 7.79 | 8.03 | 10.45 | 10.44 | 10.67 | 9.13 | 11.84 | 7.91 | 10.45 |
| FKBP1A | 5 | 7.74 | 9.64 | 12.07 | 11.43 | 9.22 | 10.05 | 10.25 | 9.62 | 10.27 | 9.72 | 8.69 | 11.75 | 9.64 | 9.93 | 9.99 |
| FKBP2 | 8 | 8.18 | 10.02 | 10.73 | 4.99 | 11.42 | 8.76 | 8.59 | 23.29 | 7.83 | 6.20 | 9.10 | 7.86 | 10.09 | 15.94 | 7.02 |
| FKBP3 | 19 | 8.93 | 10.11 | 10.92 | 9.42 | 11.98 | 11.50 | 10.19 | 9.72 | 8.75 | 8.49 | 9.52 | 10.17 | 11.74 | 9.96 | 8.62 |
| FKBP4 | 98 | 8.95 | 9.55 | 10.27 | 9.00 | 9.59 | 9.37 | 10.46 | 12.73 | 10.21 | 9.89 | 9.25 | 9.63 | 9.48 | 11.59 | 10.05 |
| FKBP5 | 22 | 11.52 | 11.51 | 9.07 | 7.98 | 13.02 | 13.01 | 7.50 | 6.56 | 10.25 | 9.58 | 11.52 | 8.53 | 13.01 | 7.03 | 9.91 |
| FKBP7 | 3 | 10.78 | 9.99 | 7.43 | 6.66 | 7.25 | 6.90 | 18.04 | 15.16 | 8.60 | 9.20 | 10.38 | 7.05 | 7.08 | 16.60 | 8.90 |

| | | | | | | | | | | | | | | | | |
|---|---|---|---|---|---|---|---|---|---|---|---|---|---|---|---|---|
| FKBP8 | 13 | 9.93 | 8.13 | 9.70 | 9.15 | 9.48 | 8.30 | 12.00 | 12.93 | 9.82 | 10.56 | 9.03 | 9.43 | 8.89 | 12.46 | 10.19 |
| FKBP9 | 16 | 10.29 | 8.55 | 7.42 | 8.36 | 10.36 | 9.80 | 11.57 | 15.37 | 9.10 | 9.18 | 9.42 | 7.89 | 10.08 | 13.47 | 9.14 |
| FKBPL | 1 | 10.33 | 12.66 | 9.30 | 10.52 | 10.22 | 10.68 | 9.64 | 9.27 | 8.35 | 9.03 | 11.50 | 9.91 | 10.45 | 9.45 | 8.69 |
| FLAD1 | 9 | 9.58 | 9.83 | 10.70 | 11.21 | 9.77 | 9.83 | 9.74 | 9.44 | 10.07 | 9.84 | 9.70 | 10.95 | 9.80 | 9.59 | 9.95 |
| FLCN | 1 | 9.44 | 10.11 | 11.36 | 10.56 | 10.30 | 8.28 | 8.70 | 8.11 | 10.96 | 12.18 | 9.77 | 10.96 | 9.29 | 8.41 | 11.57 |
| FLII | 49 | 9.50 | 10.08 | 11.28 | 11.01 | 9.49 | 9.44 | 8.13 | 8.48 | 11.33 | 11.24 | 9.79 | 11.15 | 9.47 | 8.30 | 11.29 |
| FLNA | 541 | 9.28 | 10.37 | 11.03 | 9.41 | 11.26 | 10.84 | 7.12 | 8.80 | 11.20 | 10.71 | 9.82 | 10.22 | 11.05 | 7.96 | 10.95 |
| FLNB | 291 | 8.09 | 8.68 | 12.79 | 11.01 | 11.95 | 11.51 | 5.41 | 5.91 | 12.57 | 12.07 | 8.39 | 11.90 | 11.73 | 5.66 | 12.32 |
| FLNC | 144 | 7.73 | 8.22 | 18.08 | 15.64 | 6.59 | 6.29 | 5.19 | 6.36 | 13.29 | 12.61 | 7.97 | 16.86 | 6.44 | 5.77 | 12.95 |
| FLOT1 | 19 | 11.43 | 10.04 | 9.19 | 10.39 | 11.60 | 11.91 | 9.58 | 8.88 | 8.43 | 8.54 | 10.74 | 9.79 | 11.76 | 9.23 | 8.48 |
| FLOT2 | 12 | 12.59 | 9.07 | 9.47 | 11.44 | 11.03 | 11.23 | 9.27 | 9.28 | 8.26 | 8.37 | 10.83 | 10.45 | 11.13 | 9.28 | 8.32 |
| FLVCR1 | 1 | 12.87 | 9.56 | 3.61 | 3.70 | 13.55 | 12.51 | 13.37 | 10.82 | 9.85 | 10.15 | 11.21 | 3.66 | 13.03 | 12.09 | 10.00 |
| FLYWCH1 | 3 | 9.68 | 10.89 | 9.29 | 8.02 | 10.80 | 11.06 | 12.72 | 11.41 | 8.15 | 7.99 | 10.28 | 8.66 | 10.93 | 12.07 | 8.07 |
| FLYWCH2 | 5 | 8.89 | 9.93 | 8.27 | 7.57 | 14.93 | 14.80 | 10.83 | 9.47 | 7.98 | 7.33 | 9.41 | 7.92 | 14.87 | 10.15 | 7.66 |
| FMNL1 | 28 | 9.14 | 8.93 | 10.46 | 10.53 | 9.16 | 8.91 | 8.23 | 7.94 | 13.54 | 13.17 | 9.03 | 10.49 | 9.04 | 8.08 | 13.36 |
| FMNL2 | 2 | 6.36 | 7.20 | 10.55 | 11.54 | 7.03 | 7.30 | 14.66 | 13.04 | 11.10 | 11.23 | 6.78 | 11.04 | 7.16 | 13.85 | 11.16 |
| FMR1 | 14 | 8.53 | 9.96 | 8.37 | 9.09 | 11.00 | 12.39 | 12.22 | 12.02 | 8.31 | 8.11 | 9.24 | 8.73 | 11.70 | 12.12 | 8.21 |
| FN1 | 33 | 6.14 | 5.70 | 28.27 | 29.76 | 2.76 | 2.89 | 3.70 | 3.08 | 8.77 | 8.92 | 5.92 | 29.02 | 2.83 | 3.39 | 8.85 |
| FN3K | 4 | 14.28 | 15.59 | 6.29 | 5.35 | 15.63 | 14.42 | 7.24 | 6.16 | 7.72 | 7.34 | 14.93 | 5.82 | 15.02 | 6.70 | 7.53 |
| FN3KRP | 7 | 10.99 | 11.47 | 11.00 | 10.36 | 10.70 | 10.88 | 6.56 | 5.63 | 11.72 | 10.69 | 11.23 | 10.68 | 10.79 | 6.09 | 11.21 |
| FNBP1 | 11 | 9.44 | 10.42 | 10.84 | 10.43 | 12.37 | 13.41 | 6.21 | 5.81 | 10.96 | 10.11 | 9.93 | 10.64 | 12.89 | 6.01 | 10.53 |
| FNBP1L | 10 | 11.18 | 10.85 | 13.50 | 12.80 | 8.94 | 8.94 | 6.44 | 5.71 | 11.23 | 10.41 | 11.01 | 13.15 | 8.94 | 6.07 | 10.82 |
| FNBP4 | 14 | 8.76 | 9.80 | 11.40 | 11.32 | 10.62 | 11.03 | 10.13 | 9.59 | 8.78 | 8.57 | 9.28 | 11.36 | 10.83 | 9.86 | 8.67 |
| FNDC3A | 8 | 12.70 | 8.17 | 7.36 | 10.89 | 12.68 | 10.83 | 10.53 | 9.67 | 8.23 | 8.93 | 10.43 | 9.13 | 11.76 | 10.10 | 8.58 |
| FNDC3B | 6 | 11.28 | 6.51 | 8.67 | 12.26 | 9.07 | 7.59 | 7.75 | 6.75 | 13.41 | 16.71 | 8.89 | 10.47 | 8.33 | 7.25 | 15.06 |
| FNTA | 6 | 9.30 | 10.06 | 11.43 | 10.13 | 8.71 | 8.93 | 12.03 | 10.37 | 9.53 | 9.50 | 9.68 | 10.78 | 8.82 | 11.20 | 9.51 |
| FOCAD | 10 | 11.31 | 10.83 | 9.52 | 10.04 | 9.43 | 9.58 | 9.13 | 8.85 | 10.73 | 10.58 | 11.07 | 9.78 | 9.50 | 8.99 | 10.65 |
| FOLR1 | 2 | 10.37 | 10.43 | 5.07 | 6.11 | 13.95 | 15.88 | 12.62 | 12.42 | 5.69 | 7.47 | 10.40 | 5.59 | 14.91 | 12.52 | 6.58 |
| FOSB | 1 | 9.33 | 11.57 | 3.55 | 2.89 | 7.74 | 8.72 | 20.88 | 13.13 | 12.32 | 9.85 | 10.45 | 3.22 | 8.23 | 17.00 | 11.09 |
| FOSL1 | 2 | 5.36 | 6.20 | 13.02 | 13.45 | 5.25 | 4.77 | 17.86 | 12.30 | 11.64 | 10.15 | 5.78 | 13.23 | 5.01 | 15.08 | 10.89 |
| FOSL2 | 4 | 8.96 | 9.03 | 10.47 | 9.96 | 11.00 | 11.47 | 12.51 | 10.76 | 8.16 | 7.68 | 8.99 | 10.21 | 11.24 | 11.64 | 7.92 |
| FOXC2 | 1 | 8.54 | 8.97 | 12.13 | 9.98 | 13.52 | 11.79 | 7.51 | 6.54 | 10.80 | 10.22 | 8.75 | 11.06 | 12.66 | 7.03 | 10.51 |
| FOXJ3 | 3 | 10.11 | 10.89 | 10.19 | 10.36 | 7.07 | 6.98 | 12.77 | 11.33 | 10.78 | 9.52 | 10.50 | 10.27 | 7.02 | 12.05 | 10.15 |
| FOXK1 | 14 | 9.21 | 9.79 | 12.07 | 12.45 | 7.32 | 7.78 | 10.97 | 10.78 | 10.07 | 9.56 | 9.50 | 12.26 | 7.55 | 10.88 | 9.82 |
| FOXK2 | 6 | 9.69 | 11.13 | 10.81 | 11.01 | 8.51 | 8.37 | 10.68 | 10.08 | 9.91 | 9.81 | 10.41 | 10.91 | 8.44 | 10.38 | 9.86 |
| FOXM1 | 1 | 8.24 | 8.46 | 9.73 | 9.79 | 9.66 | 10.84 | 10.82 | 12.35 | 10.84 | 9.27 | 8.35 | 9.76 | 10.25 | 11.58 | 10.05 |
| FOXP1 | 7 | 8.69 | 9.71 | 10.72 | 10.35 | 8.54 | 9.61 | 12.01 | 10.78 | 10.00 | 9.59 | 9.20 | 10.54 | 9.08 | 11.40 | 9.79 |
| FOXP4 | 4 | 9.63 | 10.90 | 9.69 | 9.31 | 9.13 | 9.37 | 12.01 | 11.01 | 9.92 | 9.03 | 10.27 | 9.50 | 9.25 | 11.51 | 9.47 |
| FOXRED1 | 7 | 10.95 | 9.40 | 9.37 | 11.45 | 9.17 | 8.60 | 9.51 | 9.30 | 10.63 | 11.62 | 10.18 | 10.41 | 8.88 | 9.41 | 11.12 |
| FPGS | 6 | 9.50 | 10.72 | 6.68 | 7.53 | 12.05 | 12.77 | 8.58 | 8.42 | 11.71 | 12.05 | 10.11 | 7.10 | 12.41 | 8.50 | 11.88 |
| FRA10AC1 | 1 | 8.74 | 10.34 | 11.99 | 10.96 | 8.36 | 8.04 | 13.41 | 10.03 | 9.46 | 8.68 | 9.54 | 11.47 | 8.20 | 11.72 | 9.07 |
| FRG1 | 8 | 5.93 | 10.66 | 10.84 | 9.11 | 9.02 | 8.90 | 11.11 | 16.99 | 8.28 | 9.15 | 8.29 | 9.98 | 8.96 | 14.05 | 8.72 |

| | | | | | | | | | | | | | | | | |
|---|---|---|---|---|---|---|---|---|---|---|---|---|---|---|---|---|
| FRMPD1 | 1 | 10.59 | 9.02 | 14.05 | 13.37 | 6.01 | 6.03 | 4.43 | 5.03 | 15.60 | 15.87 | 9.81 | 13.71 | 6.02 | 4.73 | 15.73 |
| FRYL | 11 | 9.66 | 9.13 | 10.73 | 11.80 | 9.20 | 9.38 | 7.99 | 7.88 | 11.90 | 12.32 | 9.40 | 11.27 | 9.29 | 7.94 | 12.11 |
| FSCN1 | 87 | 7.60 | 8.69 | 12.54 | 11.20 | 6.69 | 6.62 | 9.96 | 15.03 | 10.79 | 10.88 | 8.14 | 11.87 | 6.66 | 12.50 | 10.84 |
| FSIP2 | 4 | 9.38 | 11.13 | 10.44 | 10.00 | 10.08 | 10.78 | 9.75 | 9.29 | 9.86 | 9.28 | 10.26 | 10.22 | 10.43 | 9.52 | 9.57 |
| FSTL1 | 4 | 12.14 | 11.52 | 13.28 | 13.66 | 7.63 | 8.11 | 3.85 | 3.11 | 13.58 | 13.13 | 11.83 | 13.47 | 7.87 | 3.48 | 13.35 |
| FSTL4 | 2 | 4.39 | 2.89 | 6.58 | 5.25 | 5.61 | 5.12 | 32.66 | 31.24 | 2.61 | 3.66 | 3.64 | 5.91 | 5.36 | 31.95 | 3.14 |
| FTH1 | 8 | 10.14 | 10.85 | 10.23 | 8.03 | 11.83 | 11.74 | 12.58 | 7.61 | 8.62 | 8.37 | 10.50 | 9.13 | 11.78 | 10.09 | 8.50 |
| FTL | 6 | 11.25 | 12.25 | 6.72 | 5.03 | 9.62 | 8.80 | 17.63 | 11.89 | 8.75 | 8.05 | 11.75 | 5.88 | 9.21 | 14.76 | 8.40 |
| FTO | 12 | 11.97 | 12.74 | 7.92 | 7.32 | 10.51 | 10.44 | 10.46 | 10.07 | 9.50 | 9.06 | 12.36 | 7.62 | 10.48 | 10.27 | 9.28 |
| FTSJ1 | 1 | 8.46 | 9.18 | 10.73 | 11.58 | 8.67 | 10.34 | 9.49 | 9.67 | 10.57 | 11.31 | 8.82 | 11.16 | 9.50 | 9.58 | 10.94 |
| FTSJ2 | 2 | 11.75 | 9.77 | 8.77 | 10.08 | 8.17 | 9.36 | 10.48 | 9.49 | 10.47 | 11.66 | 10.76 | 9.42 | 8.77 | 9.99 | 11.06 |
| FTSJ3 | 27 | 9.70 | 10.31 | 10.84 | 11.16 | 7.90 | 8.10 | 11.33 | 11.75 | 9.37 | 9.53 | 10.00 | 11.00 | 8.00 | 11.54 | 9.45 |
| FUBP1 | 75 | 10.14 | 11.29 | 8.27 | 7.64 | 10.54 | 10.62 | 10.83 | 11.55 | 9.84 | 9.28 | 10.72 | 7.95 | 10.58 | 11.19 | 9.56 |
| FUBP3 | 34 | 8.68 | 9.81 | 11.11 | 10.27 | 10.07 | 10.29 | 10.20 | 10.12 | 10.02 | 9.44 | 9.24 | 10.69 | 10.18 | 10.16 | 9.73 |
| FUCA2 | 5 | 10.45 | 8.88 | 9.20 | 8.54 | 13.18 | 11.99 | 11.69 | 10.27 | 7.34 | 8.47 | 9.66 | 8.87 | 12.58 | 10.98 | 7.90 |
| FUNDC2 | 4 | 13.90 | 7.38 | 6.27 | 8.90 | 12.14 | 10.26 | 11.36 | 10.43 | 9.08 | 10.29 | 10.64 | 7.59 | 11.20 | 10.90 | 9.68 |
| FUS | 28 | 11.06 | 11.34 | 3.09 | 2.46 | 15.56 | 13.55 | 13.13 | 11.81 | 8.98 | 9.03 | 11.20 | 2.78 | 14.55 | 12.47 | 9.00 |
| FUT11 | 2 | 10.82 | 8.68 | 10.63 | 11.05 | 9.57 | 10.49 | 11.23 | 10.51 | 9.01 | 8.01 | 9.75 | 10.84 | 10.03 | 10.87 | 8.51 |
| FXN | 2 | 11.17 | 8.54 | 11.24 | 12.11 | 10.50 | 9.69 | 9.29 | 5.93 | 10.59 | 10.94 | 9.86 | 11.67 | 10.10 | 7.61 | 10.77 |
| FXR1 | 28 | 9.30 | 10.60 | 11.40 | 10.69 | 10.31 | 10.69 | 9.18 | 8.41 | 9.71 | 9.71 | 9.95 | 11.05 | 10.50 | 8.80 | 9.71 |
| FXR2 | 17 | 10.78 | 11.48 | 11.23 | 10.74 | 9.84 | 10.29 | 8.62 | 7.91 | 9.67 | 9.45 | 11.13 | 10.99 | 10.06 | 8.26 | 9.56 |
| FYCO1 | 10 | 10.66 | 11.46 | 9.74 | 9.49 | 10.72 | 10.81 | 8.63 | 7.93 | 10.67 | 9.89 | 11.06 | 9.61 | 10.76 | 8.28 | 10.28 |
| FYTTD1 | 3 | 9.54 | 10.55 | 4.89 | 5.26 | 17.46 | 17.90 | 8.89 | 8.39 | 7.44 | 9.67 | 10.04 | 5.08 | 17.68 | 8.64 | 8.56 |
| FZD10 | 1 | 13.45 | 15.13 | 4.77 | 8.45 | 7.51 | 10.43 | 9.43 | 12.42 | 9.82 | 8.60 | 14.29 | 6.61 | 8.97 | 10.92 | 9.21 |
| FZD6 | 1 | 10.36 | 8.41 | 7.41 | 11.00 | 9.76 | 11.21 | 10.16 | 9.40 | 10.14 | 12.17 | 9.38 | 9.20 | 10.48 | 9.78 | 11.15 |
| G3BP1 | 52 | 9.44 | 11.00 | 10.51 | 9.45 | 10.02 | 10.02 | 11.10 | 10.97 | 8.95 | 8.54 | 10.22 | 9.98 | 10.02 | 11.03 | 8.75 |
| G3BP2 | 23 | 9.34 | 10.70 | 12.16 | 10.68 | 10.32 | 10.36 | 8.63 | 7.44 | 10.39 | 9.98 | 10.02 | 11.42 | 10.34 | 8.03 | 10.18 |
| G6PC3 | 1 | 17.02 | 7.06 | 6.17 | 9.15 | 8.16 | 6.69 | 8.87 | 9.46 | 12.34 | 15.09 | 12.04 | 7.66 | 7.43 | 9.16 | 13.71 |
| G6PD | 70 | 11.63 | 13.18 | 7.94 | 7.03 | 12.75 | 12.73 | 7.37 | 8.66 | 9.33 | 9.38 | 12.40 | 7.49 | 12.74 | 8.02 | 9.36 |
| GAA | 5 | 9.62 | 10.36 | 2.99 | 2.69 | 15.31 | 15.51 | 17.19 | 15.31 | 5.72 | 5.29 | 9.99 | 2.84 | 15.41 | 16.25 | 5.50 |
| GABARAP | 6 | 10.34 | 10.93 | 10.55 | 10.25 | 9.06 | 9.44 | 9.50 | 8.99 | 10.70 | 10.23 | 10.64 | 10.40 | 9.25 | 9.25 | 10.46 |
| GABARAPL2 | 5 | 10.25 | 10.42 | 8.01 | 7.35 | 11.45 | 10.94 | 9.78 | 9.34 | 11.48 | 10.98 | 10.33 | 7.68 | 11.20 | 9.56 | 11.23 |
| GABPA | 8 | 9.75 | 10.09 | 9.44 | 9.38 | 10.19 | 10.81 | 11.75 | 10.90 | 8.99 | 8.69 | 9.92 | 9.41 | 10.50 | 11.33 | 8.84 |
| GABPB1 | 2 | 9.77 | 10.41 | 8.36 | 8.03 | 10.63 | 10.31 | 12.92 | 11.18 | 9.40 | 8.99 | 10.09 | 8.19 | 10.47 | 12.05 | 9.20 |
| GADD45GIP1 | 7 | 9.88 | 9.21 | 9.85 | 10.89 | 10.24 | 10.13 | 11.20 | 9.79 | 9.36 | 9.44 | 9.55 | 10.37 | 10.18 | 10.50 | 9.40 |
| GAK | 15 | 9.73 | 9.80 | 10.29 | 10.35 | 11.04 | 11.29 | 10.61 | 9.63 | 8.70 | 8.56 | 9.77 | 10.32 | 11.16 | 10.12 | 8.63 |
| GALE | 15 | 8.75 | 9.49 | 7.42 | 7.12 | 10.93 | 11.68 | 11.21 | 15.17 | 9.28 | 8.95 | 9.12 | 7.27 | 11.30 | 13.19 | 9.12 |
| GALK1 | 12 | 7.79 | 10.52 | 10.28 | 10.03 | 10.10 | 9.78 | 11.26 | 10.10 | 9.90 | 10.24 | 9.15 | 10.16 | 9.94 | 10.68 | 10.07 |
| GALK2 | 6 | 9.48 | 10.58 | 8.33 | 8.47 | 12.67 | 14.32 | 11.05 | 10.65 | 7.03 | 7.40 | 10.03 | 8.40 | 13.49 | 10.85 | 7.22 |
| GALM | 4 | 10.56 | 9.79 | 11.27 | 11.48 | 9.16 | 10.51 | 10.64 | 9.24 | 8.53 | 8.82 | 10.17 | 11.37 | 9.84 | 9.94 | 8.68 |
| GALNS | 3 | 7.33 | 6.44 | 5.48 | 3.05 | 9.14 | 11.88 | 7.95 | 39.61 | 5.09 | 4.02 | 6.88 | 4.27 | 10.51 | 23.78 | 4.56 |
| GALNT1 | 11 | 11.02 | 7.13 | 9.16 | 11.29 | 9.79 | 8.34 | 6.44 | 6.13 | 14.44 | 16.26 | 9.07 | 10.22 | 9.06 | 6.29 | 15.35 |

| | | | | | | | | | | | | | | | | |
|---|---|---|---|---|---|---|---|---|---|---|---|---|---|---|---|---|
| GALNT2 | 38 | 11.24 | 7.03 | 6.27 | 7.36 | 15.54 | 13.34 | 9.50 | 9.49 | 9.40 | 10.83 | 9.14 | 6.82 | 14.44 | 9.49 | 10.11 |
| GALNT7 | 17 | 13.87 | 8.69 | 9.01 | 10.95 | 11.34 | 10.19 | 7.97 | 8.33 | 9.08 | 10.57 | 11.28 | 9.98 | 10.76 | 8.15 | 9.82 |
| GALT | 2 | 11.92 | 12.88 | 4.19 | 4.26 | 12.70 | 12.93 | 6.53 | 5.86 | 13.94 | 14.80 | 12.40 | 4.22 | 12.82 | 6.19 | 14.37 |
| GAMT | 1 | 9.22 | 11.07 | 9.87 | 8.88 | 12.92 | 11.30 | 9.29 | 7.68 | 9.60 | 10.17 | 10.15 | 9.37 | 12.11 | 8.48 | 9.88 |
| GAN | 2 | 9.84 | 10.72 | 9.44 | 7.81 | 11.68 | 9.70 | 8.87 | 9.19 | 11.62 | 11.12 | 10.28 | 8.62 | 10.69 | 9.03 | 11.37 |
| GANAB | 67 | 10.20 | 7.70 | 8.17 | 9.19 | 10.23 | 9.63 | 12.54 | 13.12 | 9.14 | 10.09 | 8.95 | 8.68 | 9.93 | 12.83 | 9.61 |
| GAPDH | 318 | 9.50 | 12.12 | 11.00 | 10.32 | 9.27 | 9.25 | 9.12 | 9.13 | 10.18 | 10.12 | 10.81 | 10.66 | 9.26 | 9.12 | 10.15 |
| GAPVD1 | 44 | 8.85 | 9.78 | 9.40 | 9.47 | 10.92 | 10.92 | 9.63 | 9.47 | 10.77 | 10.80 | 9.31 | 9.44 | 10.92 | 9.55 | 10.78 |
| GAR1 | 7 | 8.75 | 8.59 | 10.41 | 9.25 | 8.54 | 8.39 | 12.12 | 15.93 | 8.95 | 9.08 | 8.67 | 9.83 | 8.47 | 14.03 | 9.01 |
| GARS | 43 | 9.29 | 9.65 | 13.05 | 11.65 | 8.62 | 8.40 | 10.25 | 9.22 | 10.11 | 9.77 | 9.47 | 12.35 | 8.51 | 9.73 | 9.94 |
| GART | 51 | 9.83 | 10.49 | 8.56 | 8.01 | 10.63 | 10.74 | 9.49 | 11.31 | 10.53 | 10.43 | 10.16 | 8.28 | 10.69 | 10.40 | 10.48 |
| GATAD2A | 19 | 8.38 | 9.84 | 11.52 | 10.33 | 9.09 | 9.21 | 12.49 | 11.07 | 8.95 | 9.12 | 9.11 | 10.92 | 9.15 | 11.78 | 9.03 |
| GATAD2B | 11 | 9.29 | 10.88 | 10.75 | 10.36 | 11.10 | 11.27 | 9.43 | 8.40 | 9.40 | 9.12 | 10.09 | 10.56 | 11.19 | 8.91 | 9.26 |
| GATC | 2 | 10.34 | 11.42 | 9.97 | 9.88 | 9.89 | 9.76 | 11.66 | 8.68 | 9.08 | 9.32 | 10.88 | 9.92 | 9.82 | 10.17 | 9.20 |
| GATSL1 | 1 | 12.48 | 15.11 | 9.34 | 8.67 | 8.52 | 10.96 | 10.86 | 7.68 | 8.46 | 7.92 | 13.79 | 9.00 | 9.74 | 9.27 | 8.19 |
| GBA | 18 | 9.68 | 9.75 | 10.40 | 10.60 | 12.63 | 12.66 | 9.28 | 8.75 | 8.06 | 8.19 | 9.72 | 10.50 | 12.65 | 9.01 | 8.12 |
| GBAS | 2 | 12.65 | 6.78 | 8.02 | 7.99 | 8.84 | 5.90 | 10.39 | 21.78 | 9.33 | 8.32 | 9.71 | 8.01 | 7.37 | 16.08 | 8.83 |
| GBE1 | 25 | 9.63 | 9.78 | 8.63 | 7.50 | 13.03 | 12.75 | 11.66 | 11.59 | 7.64 | 7.78 | 9.70 | 8.07 | 12.89 | 11.63 | 7.71 |
| GBF1 | 30 | 10.29 | 9.93 | 10.51 | 10.38 | 10.04 | 9.81 | 8.41 | 8.28 | 11.29 | 11.05 | 10.11 | 10.45 | 9.92 | 8.35 | 11.17 |
| GBP1 | 25 | 12.37 | 11.15 | 12.88 | 10.58 | 8.91 | 7.93 | 3.12 | 2.47 | 15.30 | 15.29 | 11.76 | 11.73 | 8.42 | 2.80 | 15.29 |
| GBP2 | 11 | 12.60 | 12.75 | 8.84 | 8.32 | 10.30 | 10.62 | 9.58 | 8.77 | 9.22 | 9.02 | 12.67 | 8.58 | 10.46 | 9.17 | 9.12 |
| GCC1 | 3 | 8.33 | 9.83 | 11.81 | 11.70 | 9.75 | 9.64 | 9.60 | 9.49 | 10.30 | 9.54 | 9.08 | 11.76 | 9.70 | 9.54 | 9.92 |
| GCC2 | 27 | 10.12 | 10.82 | 11.05 | 10.89 | 9.69 | 10.26 | 9.07 | 8.16 | 10.34 | 9.60 | 10.47 | 10.97 | 9.98 | 8.61 | 9.97 |
| GCDH | 5 | 9.80 | 8.45 | 11.02 | 12.19 | 7.39 | 6.44 | 12.64 | 11.11 | 10.58 | 10.36 | 9.13 | 11.61 | 6.92 | 11.88 | 10.47 |
| GCFC2 | 1 | 11.88 | 11.68 | 9.77 | 9.07 | 9.00 | 8.91 | 13.12 | 11.19 | 7.73 | 7.65 | 11.78 | 9.42 | 8.95 | 12.15 | 7.69 |
| GCLC | 13 | 12.70 | 12.37 | 9.46 | 9.43 | 8.20 | 8.80 | 9.23 | 9.53 | 10.40 | 9.87 | 12.54 | 9.45 | 8.50 | 9.38 | 10.14 |
| GCLM | 16 | 12.18 | 12.89 | 9.73 | 9.23 | 8.00 | 8.13 | 10.98 | 9.79 | 9.82 | 9.25 | 12.53 | 9.48 | 8.06 | 10.38 | 9.54 |
| GCN1L1 | 111 | 9.63 | 9.78 | 9.54 | 9.97 | 10.87 | 10.85 | 8.96 | 9.04 | 10.72 | 10.66 | 9.70 | 9.75 | 10.86 | 9.00 | 10.69 |
| GCSH | 13 | 10.88 | 8.67 | 8.51 | 8.75 | 10.07 | 9.77 | 9.96 | 14.84 | 9.71 | 8.83 | 9.78 | 8.63 | 9.92 | 12.40 | 9.27 |
| GDA | 16 | 5.41 | 5.95 | 7.72 | 7.87 | 19.38 | 19.27 | 6.98 | 6.21 | 10.85 | 10.36 | 5.68 | 7.80 | 19.32 | 6.59 | 10.61 |
| GDAP1 | 3 | 16.34 | 8.82 | 7.47 | 10.10 | 7.87 | 6.99 | 11.82 | 10.06 | 10.02 | 10.53 | 12.58 | 8.78 | 7.43 | 10.94 | 10.27 |
| GDAP2 | 1 | 10.06 | 11.37 | 11.13 | 11.84 | 8.86 | 9.73 | 8.94 | 8.70 | 9.49 | 9.90 | 10.71 | 11.49 | 9.29 | 8.82 | 9.69 |
| GDE1 | 1 | 9.81 | 7.95 | 9.94 | 11.18 | 10.38 | 8.79 | 8.69 | 9.62 | 12.65 | 11.00 | 8.88 | 10.56 | 9.58 | 9.16 | 11.83 |
| GDI1 | 57 | 9.69 | 9.75 | 8.78 | 8.43 | 12.69 | 12.82 | 9.14 | 10.34 | 9.47 | 8.91 | 9.72 | 8.60 | 12.75 | 9.74 | 9.19 |
| GDI2 | 44 | 9.61 | 9.29 | 9.33 | 8.79 | 10.29 | 9.95 | 9.74 | 14.69 | 9.48 | 8.83 | 9.45 | 9.06 | 10.12 | 12.22 | 9.16 |
| GDPGP1 | 2 | 9.95 | 9.98 | 7.54 | 7.99 | 13.71 | 14.79 | 7.58 | 7.55 | 10.45 | 10.46 | 9.96 | 7.77 | 14.25 | 7.56 | 10.46 |
| GEMIN2 | 3 | 8.57 | 8.93 | 10.89 | 9.15 | 9.72 | 10.57 | 10.21 | 13.32 | 9.01 | 9.63 | 8.75 | 10.02 | 10.14 | 11.77 | 9.32 |
| GEMIN4 | 18 | 10.13 | 10.35 | 10.54 | 10.89 | 8.61 | 9.14 | 10.36 | 10.12 | 10.05 | 9.80 | 10.24 | 10.72 | 8.87 | 10.24 | 9.93 |
| GEMIN5 | 36 | 9.07 | 9.29 | 9.99 | 9.42 | 8.30 | 8.15 | 12.13 | 13.74 | 10.31 | 9.60 | 9.18 | 9.70 | 8.22 | 12.93 | 9.96 |
| GEMIN6 | 2 | 10.74 | 9.12 | 11.11 | 9.84 | 9.37 | 9.25 | 11.86 | 10.81 | 9.24 | 8.65 | 9.93 | 10.48 | 9.31 | 11.34 | 8.95 |
| GEMIN7 | 1 | 9.51 | 9.76 | 11.54 | 9.92 | 9.20 | 8.85 | 12.41 | 11.31 | 9.30 | 8.21 | 9.63 | 10.73 | 9.02 | 11.86 | 8.75 |
| GEMIN8 | 4 | 9.67 | 10.88 | 7.72 | 7.64 | 10.28 | 10.85 | 12.75 | 11.92 | 9.57 | 8.72 | 10.27 | 7.68 | 10.56 | 12.34 | 9.14 |

| | | | | | | | | | | | | | | | | |
|---|---|---|---|---|---|---|---|---|---|---|---|---|---|---|---|---|
| GET4 | 4 | 10.02 | 10.97 | 10.25 | 10.23 | 7.93 | 8.34 | 10.13 | 10.29 | 10.68 | 11.15 | 10.50 | 10.24 | 8.14 | 10.21 | 10.92 |
| GFER | 1 | 8.13 | 8.72 | 14.51 | 14.23 | 9.74 | 7.46 | 11.04 | 7.74 | 8.91 | 9.52 | 8.43 | 14.37 | 8.60 | 9.39 | 9.22 |
| GFM1 | 30 | 11.41 | 9.67 | 8.67 | 9.66 | 8.75 | 8.07 | 10.14 | 12.47 | 10.47 | 10.68 | 10.54 | 9.17 | 8.41 | 11.31 | 10.57 |
| GFM2 | 6 | 10.93 | 9.61 | 10.08 | 11.73 | 8.07 | 7.99 | 10.40 | 9.92 | 10.77 | 10.49 | 10.27 | 10.91 | 8.03 | 10.16 | 10.63 |
| GFPT1 | 33 | 9.82 | 10.69 | 9.58 | 9.35 | 10.14 | 10.85 | 8.72 | 9.57 | 10.56 | 10.73 | 10.25 | 9.46 | 10.49 | 9.15 | 10.65 |
| GFPT2 | 13 | 9.35 | 9.62 | 11.06 | 9.28 | 5.89 | 5.80 | 11.85 | 10.84 | 13.35 | 12.97 | 9.48 | 10.17 | 5.84 | 11.34 | 13.16 |
| GGA1 | 4 | 9.32 | 9.91 | 13.33 | 13.17 | 6.71 | 7.37 | 11.37 | 9.79 | 9.78 | 9.25 | 9.62 | 13.25 | 7.04 | 10.58 | 9.52 |
| GGA2 | 7 | 9.66 | 10.07 | 13.10 | 11.70 | 11.75 | 11.28 | 7.81 | 6.01 | 9.60 | 9.03 | 9.86 | 12.40 | 11.52 | 6.91 | 9.31 |
| GGA3 | 4 | 10.65 | 11.58 | 11.70 | 10.17 | 9.74 | 9.86 | 7.40 | 7.86 | 10.27 | 10.77 | 11.12 | 10.94 | 9.80 | 7.63 | 10.52 |
| GGCT | 15 | 11.30 | 12.00 | 11.47 | 9.82 | 7.99 | 8.35 | 9.46 | 10.12 | 9.96 | 9.53 | 11.65 | 10.64 | 8.17 | 9.79 | 9.74 |
| GGCX | 1 | 12.91 | 5.82 | 5.44 | 11.27 | 8.61 | 8.35 | 7.81 | 9.42 | 12.69 | 17.68 | 9.37 | 8.36 | 8.48 | 8.61 | 15.18 |
| GGH | 20 | 8.42 | 8.79 | 10.10 | 9.43 | 11.83 | 11.17 | 15.14 | 14.95 | 5.22 | 4.96 | 8.61 | 9.76 | 11.50 | 15.04 | 5.09 |
| GGNBP2 | 1 | 10.43 | 11.59 | 9.04 | 9.93 | 10.89 | 10.66 | 9.90 | 6.51 | 11.70 | 9.33 | 11.01 | 9.49 | 10.78 | 8.21 | 10.51 |
| GGPS1 | 4 | 9.21 | 10.19 | 9.39 | 9.38 | 12.84 | 12.76 | 11.04 | 9.45 | 7.94 | 7.78 | 9.70 | 9.39 | 12.80 | 10.24 | 7.86 |
| GGT1 | 4 | 10.01 | 9.45 | 6.49 | 7.39 | 8.97 | 10.18 | 18.15 | 16.62 | 6.23 | 6.52 | 9.73 | 6.94 | 9.57 | 17.38 | 6.37 |
| GHDC | 1 | 9.68 | 7.58 | 9.56 | 13.21 | 9.78 | 9.06 | 7.66 | 8.37 | 12.25 | 12.86 | 8.63 | 11.39 | 9.42 | 8.01 | 12.55 |
| GHITM | 8 | 14.16 | 6.57 | 6.97 | 10.94 | 10.04 | 8.33 | 10.26 | 9.35 | 10.85 | 12.53 | 10.37 | 8.96 | 9.18 | 9.80 | 11.69 |
| GID8 | 6 | 11.32 | 11.81 | 9.02 | 8.37 | 12.03 | 12.15 | 11.20 | 4.67 | 10.58 | 8.84 | 11.57 | 8.70 | 12.09 | 7.93 | 9.71 |
| GIGYF2 | 37 | 9.52 | 10.80 | 10.08 | 9.83 | 10.11 | 10.54 | 10.16 | 9.21 | 10.18 | 9.58 | 10.16 | 9.95 | 10.32 | 9.69 | 9.88 |
| GINS1 | 4 | 9.93 | 11.42 | 10.02 | 9.18 | 10.07 | 9.99 | 11.07 | 10.61 | 8.90 | 8.82 | 10.67 | 9.60 | 10.03 | 10.84 | 8.86 |
| GINS2 | 3 | 10.39 | 12.82 | 10.31 | 9.38 | 8.19 | 8.36 | 8.84 | 9.40 | 11.52 | 10.79 | 11.61 | 9.84 | 8.27 | 9.12 | 11.16 |
| GINS3 | 7 | 10.04 | 11.37 | 10.01 | 9.41 | 10.23 | 10.30 | 11.11 | 10.23 | 8.75 | 8.56 | 10.70 | 9.71 | 10.26 | 10.67 | 8.66 |
| GINS4 | 8 | 12.19 | 13.04 | 8.85 | 8.29 | 7.95 | 8.23 | 10.16 | 8.36 | 11.64 | 11.30 | 12.61 | 8.57 | 8.09 | 9.26 | 11.47 |
| GIPC1 | 9 | 10.06 | 11.24 | 11.46 | 11.28 | 9.59 | 9.24 | 7.20 | 6.09 | 12.49 | 11.34 | 10.65 | 11.37 | 9.41 | 6.65 | 11.92 |
| GIPC3 | 3 | 9.22 | 10.24 | 6.61 | 7.74 | 16.78 | 18.25 | 6.91 | 4.81 | 10.67 | 8.78 | 9.73 | 7.17 | 17.52 | 5.86 | 9.72 |
| GIT1 | 18 | 10.38 | 11.80 | 10.39 | 9.91 | 10.39 | 10.30 | 9.05 | 7.79 | 9.96 | 10.04 | 11.09 | 10.15 | 10.34 | 8.42 | 10.00 |
| GIT2 | 9 | 10.99 | 10.28 | 9.06 | 9.98 | 10.55 | 10.57 | 10.58 | 9.91 | 8.79 | 9.29 | 10.64 | 9.52 | 10.56 | 10.24 | 9.04 |
| GK | 2 | 13.68 | 8.11 | 6.03 | 8.26 | 14.72 | 11.98 | 10.39 | 11.13 | 6.68 | 9.03 | 10.89 | 7.14 | 13.35 | 10.76 | 7.85 |
| GLA | 6 | 8.90 | 9.59 | 8.82 | 8.09 | 13.87 | 13.46 | 13.05 | 11.39 | 6.51 | 6.31 | 9.25 | 8.45 | 13.67 | 12.22 | 6.41 |
| GLB1 | 7 | 8.70 | 8.38 | 10.08 | 8.24 | 16.73 | 15.31 | 12.95 | 11.23 | 4.28 | 4.11 | 8.54 | 9.16 | 16.02 | 12.09 | 4.19 |
| GLCCI1 | 3 | 9.23 | 10.81 | 10.43 | 10.03 | 12.12 | 11.99 | 8.86 | 7.74 | 9.18 | 9.61 | 10.02 | 10.23 | 12.05 | 8.30 | 9.39 |
| GLE1 | 14 | 10.22 | 10.69 | 11.35 | 9.50 | 9.64 | 9.35 | 11.34 | 9.59 | 9.20 | 9.12 | 10.46 | 10.42 | 9.49 | 10.46 | 9.16 |
| GLG1 | 7 | 8.55 | 5.90 | 6.15 | 8.73 | 12.20 | 10.88 | 12.43 | 15.66 | 9.70 | 9.80 | 7.23 | 7.44 | 11.54 | 14.04 | 9.75 |
| GLIPR1L1 | 4 | 18.24 | 11.78 | 9.43 | 7.59 | 9.72 | 11.10 | 6.12 | 5.48 | 11.49 | 9.05 | 15.01 | 8.51 | 10.41 | 5.80 | 10.27 |
| GLIPR2 | 6 | 12.13 | 11.97 | 11.84 | 13.70 | 4.56 | 4.63 | 3.93 | 4.02 | 15.83 | 17.39 | 12.05 | 12.77 | 4.59 | 3.98 | 16.61 |
| GLMN | 10 | 10.29 | 10.88 | 8.90 | 8.82 | 10.40 | 10.35 | 12.51 | 12.20 | 7.58 | 8.08 | 10.59 | 8.86 | 10.37 | 12.35 | 7.83 |
| GLO1 | 18 | 8.47 | 8.96 | 9.48 | 8.82 | 9.41 | 9.00 | 10.77 | 18.60 | 8.33 | 8.17 | 8.71 | 9.15 | 9.20 | 14.69 | 8.25 |
| GLOD4 | 17 | 10.34 | 11.41 | 11.33 | 10.43 | 9.61 | 9.43 | 8.97 | 8.32 | 10.29 | 9.88 | 10.87 | 10.88 | 9.52 | 8.65 | 10.08 |
| GLRX | 5 | 9.12 | 8.47 | 6.58 | 7.13 | 11.42 | 11.98 | 10.97 | 20.73 | 6.98 | 6.62 | 8.79 | 6.86 | 11.70 | 15.85 | 6.80 |
| GLRX3 | 21 | 9.36 | 9.67 | 11.72 | 10.55 | 9.87 | 9.26 | 10.58 | 11.68 | 8.81 | 8.50 | 9.51 | 11.13 | 9.56 | 11.13 | 8.66 |
| GLRX5 | 4 | 9.97 | 8.34 | 9.75 | 10.04 | 10.52 | 10.17 | 10.54 | 10.89 | 9.80 | 9.99 | 9.16 | 9.89 | 10.35 | 10.71 | 9.90 |
| GLS | 28 | 12.06 | 9.30 | 8.74 | 10.48 | 6.44 | 6.13 | 5.34 | 6.54 | 16.76 | 18.21 | 10.68 | 9.61 | 6.28 | 5.94 | 17.49 |

| | | | | | | | | | | | | | | | | |
|---|---|---|---|---|---|---|---|---|---|---|---|---|---|---|---|---|
| GLTP | 4 | 9.56 | 10.85 | 10.17 | 9.99 | 10.46 | 10.71 | 10.45 | 10.18 | 8.96 | 8.67 | 10.21 | 10.08 | 10.59 | 10.31 | 8.81 |
| GLTPD1 | 1 | 9.84 | 11.31 | 8.30 | 8.65 | 9.00 | 9.79 | 9.92 | 12.45 | 10.44 | 10.29 | 10.57 | 8.48 | 9.40 | 11.19 | 10.36 |
| GLTSCR2 | 10 | 9.20 | 10.69 | 9.29 | 9.48 | 8.83 | 8.81 | 13.30 | 11.75 | 9.52 | 9.13 | 9.95 | 9.38 | 8.82 | 12.52 | 9.32 |
| GLUD1 | 32 | 9.57 | 8.19 | 9.41 | 10.70 | 8.90 | 8.28 | 12.49 | 12.62 | 9.79 | 10.06 | 8.88 | 10.05 | 8.59 | 12.55 | 9.93 |
| GLUL | 7 | 12.48 | 12.12 | 8.63 | 7.22 | 13.20 | 12.07 | 6.39 | 5.50 | 12.24 | 10.16 | 12.30 | 7.92 | 12.64 | 5.94 | 11.20 |
| GLYR1 | 7 | 9.83 | 10.85 | 9.55 | 10.71 | 10.90 | 11.53 | 10.48 | 8.80 | 8.97 | 8.37 | 10.34 | 10.13 | 11.21 | 9.64 | 8.67 |
| GM2A | 3 | 9.01 | 9.35 | 10.12 | 9.37 | 13.81 | 13.95 | 11.26 | 9.12 | 7.15 | 6.85 | 9.18 | 9.75 | 13.88 | 10.19 | 7.00 |
| GMDS | 12 | 9.11 | 8.34 | 12.12 | 6.55 | 13.44 | 11.26 | 16.02 | 12.21 | 5.53 | 5.41 | 8.73 | 9.33 | 12.35 | 14.12 | 5.47 |
| GMEB1 | 1 | 8.60 | 9.64 | 9.46 | 6.63 | 9.05 | 8.90 | 16.45 | 13.26 | 9.27 | 8.76 | 9.12 | 8.04 | 8.97 | 14.85 | 9.02 |
| GMEB2 | 1 | 9.81 | 9.99 | 9.90 | 9.84 | 8.91 | 9.17 | 12.17 | 12.47 | 9.09 | 8.66 | 9.90 | 9.87 | 9.04 | 12.32 | 8.87 |
| GMFB | 9 | 9.71 | 10.83 | 10.06 | 10.12 | 11.34 | 12.19 | 8.46 | 7.93 | 9.52 | 9.84 | 10.27 | 10.09 | 11.76 | 8.20 | 9.68 |
| GMIP | 3 | 11.72 | 9.09 | 8.36 | 10.33 | 8.38 | 9.92 | 10.13 | 11.23 | 14.01 | 6.82 | 10.41 | 9.35 | 9.15 | 10.68 | 10.42 |
| GMNN | 4 | 9.31 | 9.98 | 11.26 | 10.39 | 11.20 | 11.00 | 9.40 | 7.94 | 10.31 | 9.22 | 9.65 | 10.82 | 11.10 | 8.67 | 9.76 |
| GMPPA | 8 | 10.36 | 11.73 | 10.86 | 10.49 | 9.32 | 9.59 | 7.82 | 7.95 | 11.08 | 10.80 | 11.04 | 10.68 | 9.45 | 7.88 | 10.94 |
| GMPPB | 8 | 11.55 | 12.16 | 9.51 | 9.29 | 9.71 | 9.43 | 7.64 | 7.93 | 11.57 | 11.21 | 11.85 | 9.40 | 9.57 | 7.79 | 11.39 |
| GMPR2 | 4 | 7.98 | 8.52 | 19.24 | 19.72 | 10.62 | 10.88 | 8.14 | 7.38 | 3.85 | 3.66 | 8.25 | 19.48 | 10.75 | 7.76 | 3.76 |
| GMPS | 40 | 9.62 | 10.57 | 9.35 | 8.23 | 10.29 | 10.17 | 11.75 | 12.44 | 8.90 | 8.68 | 10.10 | 8.79 | 10.23 | 12.09 | 8.79 |
| GNA11 | 9 | 10.77 | 9.03 | 8.98 | 11.35 | 8.35 | 9.02 | 9.74 | 9.96 | 11.08 | 11.73 | 9.90 | 10.16 | 8.68 | 9.85 | 11.40 |
| GNA12 | 4 | 10.88 | 9.98 | 10.39 | 11.94 | 7.42 | 8.03 | 9.87 | 9.78 | 10.62 | 11.09 | 10.43 | 11.16 | 7.72 | 9.83 | 10.85 |
| GNA13 | 5 | 11.20 | 10.01 | 10.45 | 13.12 | 7.14 | 8.06 | 7.81 | 8.46 | 11.19 | 12.56 | 10.61 | 11.79 | 7.60 | 8.13 | 11.87 |
| GNAI2 | 12 | 10.59 | 8.06 | 9.03 | 8.83 | 9.36 | 9.26 | 8.13 | 11.89 | 12.27 | 12.56 | 9.33 | 8.93 | 9.31 | 10.01 | 12.42 |
| GNAI3 | 5 | 11.64 | 9.71 | 8.64 | 9.36 | 7.57 | 8.21 | 9.89 | 10.43 | 12.08 | 12.45 | 10.68 | 9.00 | 7.89 | 10.16 | 12.27 |
| GNAS | 18 | 10.81 | 9.72 | 10.34 | 11.84 | 8.40 | 9.07 | 8.13 | 7.71 | 11.81 | 12.16 | 10.27 | 11.09 | 8.73 | 7.92 | 11.99 |
| GNB1 | 4 | 8.68 | 9.16 | 6.21 | 5.68 | 7.19 | 6.94 | 8.19 | 30.19 | 9.59 | 8.18 | 8.92 | 5.94 | 7.06 | 19.19 | 8.89 |
| GNB1L | 2 | 8.76 | 9.76 | 12.49 | 11.87 | 8.05 | 7.17 | 13.74 | 14.06 | 7.08 | 7.01 | 9.26 | 12.18 | 7.61 | 13.90 | 7.05 |
| GNB2 | 8 | 9.00 | 9.05 | 9.73 | 6.95 | 9.77 | 9.13 | 8.54 | 18.27 | 9.62 | 9.94 | 9.03 | 8.34 | 9.45 | 13.41 | 9.78 |
| GNB2L1 | 50 | 8.89 | 9.41 | 9.53 | 8.72 | 9.31 | 9.25 | 9.86 | 17.76 | 8.85 | 8.42 | 9.15 | 9.12 | 9.28 | 13.81 | 8.63 |
| GNB4 | 11 | 10.46 | 10.84 | 10.19 | 10.35 | 9.15 | 9.78 | 9.10 | 8.77 | 10.81 | 10.56 | 10.65 | 10.27 | 9.46 | 8.93 | 10.68 |
| GNB5 | 1 | 10.67 | 11.37 | 8.73 | 7.56 | 11.85 | 11.50 | 9.02 | 8.33 | 10.51 | 10.45 | 11.02 | 8.15 | 11.68 | 8.67 | 10.48 |
| GNE | 23 | 9.98 | 10.83 | 10.37 | 10.05 | 10.57 | 10.64 | 7.83 | 7.33 | 11.29 | 11.10 | 10.40 | 10.21 | 10.61 | 7.58 | 11.20 |
| GNG12 | 4 | 10.33 | 7.71 | 10.90 | 14.33 | 6.79 | 6.50 | 8.44 | 8.71 | 12.33 | 13.97 | 9.02 | 12.61 | 6.64 | 8.58 | 13.15 |
| GNGT1 | 1 | 9.94 | 8.62 | 8.44 | 7.76 | 9.23 | 8.16 | 14.64 | 6.12 | 13.03 | 14.07 | 9.28 | 8.10 | 8.69 | 10.38 | 13.55 |
| GNL1 | 11 | 9.40 | 10.16 | 10.78 | 10.42 | 11.26 | 11.13 | 9.49 | 8.54 | 9.75 | 9.07 | 9.78 | 10.60 | 11.19 | 9.02 | 9.41 |
| GNL2 | 18 | 8.58 | 8.81 | 9.03 | 9.11 | 7.42 | 7.26 | 13.77 | 16.80 | 9.20 | 10.03 | 8.70 | 9.07 | 7.34 | 15.28 | 9.62 |
| GNL3 | 29 | 9.07 | 8.49 | 8.48 | 8.21 | 7.61 | 7.37 | 13.28 | 18.99 | 9.62 | 8.87 | 8.78 | 8.35 | 7.49 | 16.14 | 9.24 |
| GNL3L | 15 | 9.50 | 10.29 | 9.39 | 9.60 | 8.82 | 9.48 | 13.14 | 12.71 | 8.63 | 8.44 | 9.89 | 9.49 | 9.15 | 12.92 | 8.53 |
| GNPAT | 4 | 10.53 | 9.84 | 6.33 | 6.13 | 14.88 | 13.25 | 10.22 | 11.02 | 8.30 | 9.50 | 10.19 | 6.23 | 14.07 | 10.62 | 8.90 |
| GNPDA1 | 21 | 9.11 | 9.97 | 9.29 | 9.30 | 10.44 | 11.04 | 10.77 | 12.45 | 8.97 | 8.65 | 9.54 | 9.29 | 10.74 | 11.61 | 8.81 |
| GNPDA2 | 4 | 9.68 | 10.00 | 9.85 | 9.96 | 12.12 | 12.95 | 8.47 | 7.62 | 9.89 | 9.46 | 9.84 | 9.90 | 12.54 | 8.04 | 9.67 |
| GNPNAT1 | 12 | 8.37 | 9.09 | 8.55 | 7.97 | 11.02 | 10.11 | 9.83 | 18.23 | 8.34 | 8.48 | 8.73 | 8.26 | 10.57 | 14.03 | 8.41 |
| GNS | 11 | 10.10 | 10.21 | 13.66 | 13.44 | 12.33 | 13.01 | 6.52 | 6.10 | 7.56 | 7.07 | 10.16 | 13.55 | 12.67 | 6.31 | 7.31 |
| GOLGA1 | 12 | 9.50 | 10.53 | 8.34 | 8.25 | 12.99 | 13.22 | 9.70 | 8.47 | 9.82 | 9.19 | 10.01 | 8.29 | 13.11 | 9.08 | 9.50 |

| | | | | | | | | | | | | | | | | |
|---|---|---|---|---|---|---|---|---|---|---|---|---|---|---|---|---|
| GOLGA2 | 27 | 9.38 | 11.18 | 10.29 | 9.47 | 11.24 | 11.41 | 8.92 | 8.00 | 10.11 | 10.01 | 10.28 | 9.88 | 11.32 | 8.46 | 10.06 |
| GOLGA3 | 30 | 9.05 | 10.71 | 10.28 | 9.73 | 10.71 | 11.08 | 10.08 | 8.87 | 9.98 | 9.51 | 9.88 | 10.01 | 10.90 | 9.48 | 9.75 |
| GOLGA4 | 55 | 10.18 | 10.93 | 10.00 | 9.62 | 10.22 | 10.31 | 9.46 | 8.49 | 10.66 | 10.12 | 10.56 | 9.81 | 10.27 | 8.98 | 10.39 |
| GOLGA5 | 1 | 11.83 | 7.61 | 8.62 | 6.68 | 11.56 | 10.52 | 11.55 | 8.44 | 12.21 | 10.99 | 9.72 | 7.65 | 11.04 | 9.99 | 11.60 |
| GOLGB1 | 62 | 11.93 | 8.84 | 8.87 | 10.60 | 10.45 | 9.76 | 9.84 | 9.36 | 9.91 | 10.44 | 10.39 | 9.73 | 10.10 | 9.60 | 10.18 |
| GOLIM4 | 19 | 11.32 | 10.39 | 7.82 | 10.73 | 9.62 | 10.58 | 8.48 | 8.68 | 10.52 | 11.85 | 10.86 | 9.28 | 10.10 | 8.58 | 11.18 |
| GOLM1 | 7 | 10.67 | 7.14 | 7.60 | 9.31 | 10.62 | 10.11 | 12.84 | 12.12 | 9.60 | 9.98 | 8.90 | 8.46 | 10.37 | 12.48 | 9.79 |
| GOLPH3 | 11 | 9.53 | 10.22 | 7.89 | 7.64 | 11.14 | 11.91 | 8.75 | 15.96 | 8.72 | 8.24 | 9.87 | 7.77 | 11.53 | 12.35 | 8.48 |
| GOLPH3L | 2 | 11.11 | 12.47 | 12.60 | 11.49 | 11.32 | 10.85 | 7.76 | 6.93 | 7.86 | 7.62 | 11.79 | 12.04 | 11.08 | 7.34 | 7.74 |
| GON4L | 4 | 9.62 | 10.35 | 10.67 | 10.48 | 10.21 | 10.39 | 9.20 | 9.07 | 10.10 | 9.91 | 9.99 | 10.58 | 10.30 | 9.13 | 10.00 |
| GOPC | 17 | 10.65 | 11.56 | 10.15 | 9.02 | 10.18 | 10.23 | 9.40 | 9.14 | 10.03 | 9.63 | 11.10 | 9.59 | 10.20 | 9.27 | 9.83 |
| GORAB | 3 | 8.11 | 9.18 | 9.62 | 8.49 | 13.32 | 14.87 | 11.19 | 9.69 | 7.76 | 7.77 | 8.65 | 9.05 | 14.10 | 10.44 | 7.77 |
| GORASP1 | 2 | 10.63 | 11.20 | 9.52 | 9.96 | 10.76 | 11.08 | 9.43 | 7.34 | 9.64 | 10.44 | 10.91 | 9.74 | 10.92 | 8.38 | 10.04 |
| GORASP2 | 14 | 9.31 | 10.24 | 10.86 | 10.13 | 11.20 | 11.16 | 9.89 | 9.19 | 9.07 | 8.95 | 9.77 | 10.49 | 11.18 | 9.54 | 9.01 |
| GOSR1 | 4 | 11.30 | 8.00 | 9.67 | 12.25 | 9.66 | 9.25 | 8.70 | 8.00 | 10.76 | 12.41 | 9.65 | 10.96 | 9.45 | 8.35 | 11.58 |
| GOSR2 | 2 | 10.28 | 9.42 | 10.41 | 13.22 | 8.25 | 9.55 | 7.45 | 8.42 | 10.33 | 12.68 | 9.85 | 11.82 | 8.90 | 7.93 | 11.51 |
| GOT1 | 19 | 11.09 | 11.73 | 8.85 | 7.56 | 12.37 | 12.48 | 8.08 | 8.08 | 10.25 | 9.51 | 11.41 | 8.20 | 12.42 | 8.08 | 9.88 |
| GOT2 | 38 | 11.04 | 10.27 | 8.35 | 8.68 | 11.35 | 10.88 | 9.85 | 9.52 | 9.92 | 10.13 | 10.66 | 8.52 | 11.12 | 9.69 | 10.03 |
| GPAA1 | 3 | 12.78 | 7.47 | 7.40 | 10.30 | 11.17 | 10.16 | 8.69 | 7.65 | 11.41 | 12.96 | 10.13 | 8.85 | 10.66 | 8.17 | 12.19 |
| GPALPP1 | 4 | 11.48 | 13.66 | 12.11 | 8.51 | 11.21 | 9.76 | 10.47 | 6.13 | 8.71 | 7.96 | 12.57 | 10.31 | 10.49 | 8.30 | 8.33 |
| GPAM | 4 | 8.79 | 12.70 | 9.41 | 9.40 | 9.96 | 10.23 | 8.95 | 10.69 | 10.04 | 9.84 | 10.74 | 9.41 | 10.09 | 9.82 | 9.94 |
| GPATCH1 | 2 | 9.35 | 13.47 | 9.06 | 9.72 | 8.98 | 10.35 | 11.40 | 9.67 | 9.02 | 8.99 | 11.41 | 9.39 | 9.66 | 10.54 | 9.01 |
| GPATCH11 | 5 | 10.27 | 11.42 | 10.99 | 10.54 | 9.79 | 9.97 | 9.51 | 8.21 | 10.06 | 9.24 | 10.84 | 10.77 | 9.88 | 8.86 | 9.65 |
| GPATCH2 | 1 | 8.97 | 9.35 | 8.75 | 7.01 | 13.09 | 14.75 | 10.90 | 9.77 | 8.53 | 8.88 | 9.16 | 7.88 | 13.92 | 10.34 | 8.70 |
| GPATCH4 | 24 | 9.77 | 10.60 | 11.19 | 11.28 | 9.34 | 9.76 | 9.44 | 9.67 | 9.79 | 9.16 | 10.18 | 11.24 | 9.55 | 9.56 | 9.47 |
| GPATCH8 | 11 | 9.36 | 9.73 | 10.41 | 10.16 | 9.81 | 9.99 | 11.28 | 10.43 | 9.53 | 9.30 | 9.54 | 10.29 | 9.90 | 10.85 | 9.41 |
| GPC1 | 15 | 11.70 | 10.38 | 8.88 | 8.84 | 8.39 | 8.34 | 9.40 | 9.33 | 12.41 | 12.31 | 11.04 | 8.86 | 8.37 | 9.37 | 12.36 |
| GPD1L | 9 | 10.08 | 11.11 | 9.91 | 9.90 | 10.53 | 10.88 | 9.22 | 9.04 | 9.80 | 9.52 | 10.60 | 9.91 | 10.71 | 9.13 | 9.66 |
| GPD2 | 34 | 12.78 | 9.41 | 7.31 | 8.31 | 10.98 | 11.11 | 9.97 | 9.42 | 9.96 | 10.73 | 11.10 | 7.81 | 11.04 | 9.70 | 10.35 |
| GPHN | 15 | 10.33 | 11.30 | 11.01 | 10.62 | 9.48 | 9.69 | 10.84 | 9.96 | 8.52 | 8.27 | 10.81 | 10.81 | 9.58 | 10.40 | 8.39 |
| GPI | 83 | 9.62 | 10.44 | 8.41 | 8.18 | 9.42 | 9.49 | 12.75 | 13.56 | 9.04 | 9.09 | 10.03 | 8.30 | 9.45 | 13.16 | 9.06 |
| GPKOW | 21 | 9.99 | 10.80 | 9.48 | 8.84 | 10.58 | 10.51 | 11.97 | 10.43 | 8.94 | 8.44 | 10.40 | 9.16 | 10.55 | 11.20 | 8.69 |
| GPN1 | 3 | 9.76 | 9.45 | 12.18 | 8.85 | 9.73 | 9.56 | 9.10 | 8.40 | 11.42 | 11.56 | 9.60 | 10.51 | 9.64 | 8.75 | 11.49 |
| GPN2 | 1 | 10.67 | 11.86 | 10.70 | 11.25 | 7.54 | 7.89 | 8.24 | 7.90 | 11.76 | 12.19 | 11.27 | 10.97 | 7.72 | 8.07 | 11.97 |
| GPR107 | 2 | 12.38 | 7.83 | 10.37 | 11.34 | 10.50 | 9.01 | 8.69 | 9.65 | 10.31 | 9.92 | 10.10 | 10.85 | 9.76 | 9.17 | 10.11 |
| GPR158 | 1 | 5.33 | 4.17 | 15.74 | 12.02 | 11.42 | 4.53 | 16.79 | 8.26 | 7.73 | 14.02 | 4.75 | 13.88 | 7.98 | 12.52 | 10.87 |
| GPR180 | 1 | 12.81 | 6.80 | 7.20 | 10.04 | 8.71 | 7.96 | 10.81 | 11.32 | 10.75 | 13.60 | 9.81 | 8.62 | 8.33 | 11.06 | 12.18 |
| GPR56 | 5 | 13.21 | 13.19 | 5.69 | 4.81 | 12.59 | 12.30 | 5.35 | 5.28 | 13.24 | 14.33 | 13.20 | 5.25 | 12.44 | 5.32 | 13.78 |
| GPR89B | 2 | 11.50 | 6.45 | 7.20 | 10.53 | 12.16 | 10.88 | 9.85 | 9.24 | 10.08 | 12.11 | 8.97 | 8.87 | 11.52 | 9.55 | 11.09 |
| GPRC5A | 2 | 12.22 | 7.85 | 7.00 | 8.02 | 11.61 | 10.30 | 9.49 | 7.48 | 13.02 | 13.02 | 10.03 | 7.51 | 10.95 | 8.48 | 13.02 |
| GPRIN1 | 4 | 10.21 | 9.44 | 9.43 | 10.32 | 6.82 | 7.05 | 14.73 | 13.40 | 9.45 | 9.15 | 9.82 | 9.87 | 6.93 | 14.07 | 9.30 |
| GPS1 | 18 | 9.90 | 10.38 | 9.98 | 9.14 | 9.41 | 9.50 | 8.35 | 13.46 | 9.84 | 10.04 | 10.14 | 9.56 | 9.45 | 10.90 | 9.94 |

| | | | | | | | | | | | | | | | | |
|---|---|---|---|---|---|---|---|---|---|---|---|---|---|---|---|---|
| GPS2 | 5 | 8.72 | 16.83 | 9.19 | 12.10 | 8.28 | 8.36 | 9.69 | 9.74 | 8.25 | 8.82 | 12.78 | 10.65 | 8.32 | 9.72 | 8.54 |
| GPSM1 | 3 | 11.05 | 12.13 | 9.05 | 8.45 | 10.76 | 10.95 | 10.38 | 9.36 | 9.11 | 8.76 | 11.59 | 8.75 | 10.85 | 9.87 | 8.93 |
| GPX1 | 13 | 8.88 | 9.30 | 8.43 | 9.42 | 12.36 | 12.48 | 12.97 | 12.12 | 7.16 | 6.88 | 9.09 | 8.92 | 12.42 | 12.54 | 7.02 |
| GPX4 | 13 | 10.32 | 11.02 | 10.39 | 10.04 | 10.77 | 10.99 | 9.38 | 12.35 | 7.53 | 7.21 | 10.67 | 10.21 | 10.88 | 10.86 | 7.37 |
| GPX8 | 8 | 13.09 | 7.22 | 8.66 | 13.12 | 11.45 | 10.26 | 8.04 | 7.17 | 9.15 | 11.82 | 10.16 | 10.89 | 10.86 | 7.61 | 10.49 |
| GRAMD1A | 1 | 5.39 | 5.95 | 8.17 | 12.67 | 11.28 | 11.61 | 10.37 | 8.33 | 12.89 | 13.35 | 5.67 | 10.42 | 11.45 | 9.35 | 13.12 |
| GRB2 | 14 | 10.26 | 11.57 | 12.00 | 11.97 | 8.98 | 9.18 | 7.73 | 7.18 | 10.44 | 10.69 | 10.91 | 11.98 | 9.08 | 7.46 | 10.57 |
| GRHPR | 18 | 9.54 | 10.94 | 10.47 | 9.31 | 9.59 | 9.27 | 7.37 | 7.61 | 12.89 | 13.01 | 10.24 | 9.89 | 9.43 | 7.49 | 12.95 |
| GRIPAP1 | 21 | 9.84 | 11.40 | 8.24 | 7.69 | 14.25 | 14.77 | 8.86 | 8.61 | 8.50 | 7.83 | 10.62 | 7.97 | 14.51 | 8.73 | 8.17 |
| GRN | 2 | 9.64 | 7.90 | 11.50 | 13.00 | 11.31 | 10.21 | 8.34 | 4.76 | 11.82 | 11.52 | 8.77 | 12.25 | 10.76 | 6.55 | 11.67 |
| GRPEL1 | 20 | 10.02 | 8.50 | 10.65 | 11.55 | 8.41 | 7.73 | 11.59 | 10.64 | 10.78 | 10.14 | 9.26 | 11.10 | 8.07 | 11.12 | 10.46 |
| GRPEL2 | 1 | 11.14 | 8.49 | 7.83 | 8.74 | 8.69 | 7.00 | 14.15 | 12.31 | 11.19 | 10.46 | 9.81 | 8.29 | 7.84 | 13.23 | 10.83 |
| GRSF1 | 24 | 10.38 | 8.77 | 9.89 | 10.94 | 8.67 | 8.26 | 12.18 | 10.79 | 9.90 | 10.23 | 9.57 | 10.41 | 8.47 | 11.48 | 10.06 |
| GRWD1 | 15 | 8.58 | 8.78 | 10.15 | 9.40 | 9.07 | 8.53 | 10.40 | 17.45 | 9.07 | 8.57 | 8.68 | 9.77 | 8.80 | 13.93 | 8.82 |
| GSDMC | 1 | 9.75 | 14.35 | 9.47 | 12.26 | 9.54 | 12.20 | 10.11 | 1.56 | 10.78 | 9.97 | 12.05 | 10.87 | 10.87 | 5.84 | 10.37 |
| GSE1 | 8 | 11.53 | 12.16 | 6.76 | 6.77 | 11.07 | 11.81 | 9.25 | 8.66 | 11.12 | 10.87 | 11.84 | 6.76 | 11.44 | 8.95 | 10.99 |
| GSK3A | 10 | 9.76 | 9.97 | 10.35 | 9.70 | 9.55 | 9.75 | 11.45 | 11.38 | 9.27 | 8.81 | 9.86 | 10.03 | 9.65 | 11.42 | 9.04 |
| GSK3B | 9 | 8.62 | 11.54 | 11.30 | 10.54 | 10.93 | 10.88 | 8.24 | 8.06 | 9.56 | 10.33 | 10.08 | 10.92 | 10.90 | 8.15 | 9.94 |
| GSKIP | 1 | 11.02 | 11.32 | 11.84 | 10.46 | 11.02 | 10.60 | 9.28 | 4.82 | 10.15 | 9.48 | 11.17 | 11.15 | 10.81 | 7.05 | 9.81 |
| GSN | 31 | 12.91 | 14.90 | 10.18 | 9.07 | 6.43 | 6.37 | 2.08 | 1.83 | 18.23 | 18.00 | 13.90 | 9.62 | 6.40 | 1.95 | 18.12 |
| GSPT1 | 62 | 9.49 | 10.52 | 10.67 | 9.78 | 9.61 | 9.60 | 9.45 | 10.32 | 10.45 | 10.10 | 10.01 | 10.22 | 9.61 | 9.89 | 10.28 |
| GSR | 22 | 11.07 | 12.70 | 8.37 | 9.34 | 10.04 | 10.50 | 8.96 | 9.05 | 9.94 | 10.03 | 11.89 | 8.85 | 10.27 | 9.01 | 9.98 |
| GSS | 26 | 10.46 | 11.27 | 12.57 | 9.33 | 9.87 | 9.11 | 7.66 | 6.15 | 11.88 | 11.72 | 10.86 | 10.95 | 9.49 | 6.90 | 11.80 |
| GSTCD | 3 | 9.30 | 10.36 | 12.64 | 9.67 | 8.93 | 8.25 | 11.63 | 9.27 | 10.32 | 9.63 | 9.83 | 11.15 | 8.59 | 10.45 | 9.98 |
| GSTK1 | 22 | 8.68 | 8.77 | 10.59 | 9.05 | 11.19 | 11.49 | 9.95 | 10.14 | 9.73 | 10.41 | 8.72 | 9.82 | 11.34 | 10.04 | 10.07 |
| GSTM1 | 12 | 10.46 | 10.56 | 8.54 | 7.83 | 8.59 | 8.19 | 10.89 | 14.16 | 10.91 | 9.88 | 10.51 | 8.19 | 8.39 | 12.53 | 10.39 |
| GSTM3 | 30 | 11.40 | 11.20 | 10.81 | 8.04 | 6.82 | 6.38 | 8.05 | 15.49 | 11.18 | 10.63 | 11.30 | 9.42 | 6.60 | 11.77 | 10.91 |
| GSTO1 | 30 | 8.48 | 9.22 | 9.61 | 9.56 | 12.63 | 13.51 | 9.24 | 11.49 | 8.03 | 8.24 | 8.85 | 9.58 | 13.07 | 10.36 | 8.14 |
| GSTP1 | 67 | 9.55 | 10.80 | 11.63 | 10.91 | 10.02 | 9.81 | 7.66 | 10.37 | 9.68 | 9.58 | 10.17 | 11.27 | 9.92 | 9.02 | 9.63 |
| GSTT2 | 3 | 9.63 | 10.23 | 14.32 | 12.94 | 7.93 | 7.52 | 9.73 | 9.18 | 9.28 | 9.24 | 9.93 | 13.63 | 7.73 | 9.45 | 9.26 |
| GSTZ1 | 5 | 11.97 | 12.00 | 12.37 | 11.11 | 7.52 | 7.18 | 9.82 | 8.90 | 9.89 | 9.25 | 11.98 | 11.74 | 7.35 | 9.36 | 9.57 |
| GTF2A1 | 1 | 9.10 | 10.75 | 9.80 | 10.76 | 10.31 | 11.64 | 11.53 | 9.90 | 7.80 | 8.40 | 9.93 | 10.28 | 10.97 | 10.72 | 8.10 |
| GTF2A2 | 1 | 9.50 | 10.66 | 10.22 | 9.79 | 10.13 | 10.85 | 11.10 | 10.57 | 8.75 | 8.42 | 10.08 | 10.01 | 10.49 | 10.83 | 8.59 |
| GTF2B | 7 | 9.80 | 11.22 | 10.04 | 9.77 | 8.52 | 9.07 | 11.50 | 10.23 | 10.32 | 9.55 | 10.51 | 9.90 | 8.79 | 10.86 | 9.93 |
| GTF2E1 | 12 | 10.44 | 11.62 | 10.85 | 10.78 | 8.47 | 8.95 | 9.77 | 9.56 | 9.88 | 9.66 | 11.03 | 10.82 | 8.71 | 9.67 | 9.77 |
| GTF2E2 | 10 | 10.12 | 10.54 | 11.33 | 10.68 | 8.69 | 8.42 | 10.46 | 9.19 | 10.03 | 10.52 | 10.33 | 11.01 | 8.56 | 9.83 | 10.28 |
| GTF2F1 | 12 | 8.39 | 10.77 | 16.42 | 11.91 | 9.37 | 9.07 | 8.30 | 7.22 | 10.28 | 8.28 | 9.58 | 14.17 | 9.22 | 7.76 | 9.28 |
| GTF2F2 | 22 | 9.85 | 10.66 | 13.14 | 11.22 | 10.31 | 10.58 | 8.53 | 7.36 | 9.31 | 9.04 | 10.25 | 12.18 | 10.45 | 7.94 | 9.18 |
| GTF2H1 | 5 | 9.48 | 11.01 | 10.48 | 10.29 | 9.45 | 9.44 | 9.96 | 8.51 | 10.27 | 11.09 | 10.25 | 10.39 | 9.44 | 9.24 | 10.68 |
| GTF2H2 | 6 | 10.95 | 10.33 | 10.10 | 9.83 | 8.34 | 8.68 | 9.48 | 10.25 | 11.37 | 10.66 | 10.64 | 9.97 | 8.51 | 9.86 | 11.01 |
| GTF2H3 | 3 | 9.39 | 9.57 | 11.09 | 10.92 | 9.18 | 9.23 | 9.85 | 9.92 | 11.12 | 9.73 | 9.48 | 11.01 | 9.21 | 9.88 | 10.42 |
| GTF2H4 | 5 | 8.70 | 9.32 | 10.62 | 11.05 | 9.20 | 9.40 | 9.51 | 10.37 | 10.73 | 11.09 | 9.01 | 10.84 | 9.30 | 9.94 | 10.91 |

| | | | | | | | | | | | | | | | | |
|---|---|---|---|---|---|---|---|---|---|---|---|---|---|---|---|---|
| GTF2I | 66 | 9.29 | 10.17 | 11.29 | 10.65 | 8.92 | 9.26 | 11.66 | 11.08 | 9.10 | 8.57 | 9.73 | 10.97 | 9.09 | 11.37 | 8.84 |
| GTF2IRD1 | 2 | 11.15 | 10.72 | 8.30 | 10.61 | 8.97 | 10.02 | 9.60 | 9.70 | 10.14 | 10.80 | 10.93 | 9.45 | 9.50 | 9.65 | 10.47 |
| GTF3C1 | 34 | 9.22 | 9.39 | 10.18 | 10.23 | 10.40 | 10.40 | 10.01 | 10.22 | 10.17 | 9.79 | 9.30 | 10.20 | 10.40 | 10.11 | 9.98 |
| GTF3C2 | 11 | 8.77 | 9.48 | 10.21 | 9.57 | 10.47 | 10.48 | 9.76 | 11.55 | 9.95 | 9.75 | 9.12 | 9.89 | 10.48 | 10.65 | 9.85 |
| GTF3C3 | 13 | 9.40 | 9.14 | 9.96 | 10.07 | 10.66 | 10.42 | 10.63 | 9.67 | 10.26 | 9.79 | 9.27 | 10.02 | 10.54 | 10.15 | 10.03 |
| GTF3C4 | 23 | 8.44 | 9.07 | 10.68 | 10.92 | 9.72 | 9.56 | 9.75 | 10.77 | 10.42 | 10.67 | 8.76 | 10.80 | 9.64 | 10.26 | 10.54 |
| GTF3C5 | 7 | 10.29 | 10.65 | 11.42 | 10.05 | 10.35 | 10.64 | 9.72 | 8.66 | 9.39 | 8.84 | 10.47 | 10.74 | 10.49 | 9.19 | 9.12 |
| GTPBP1 | 12 | 10.37 | 11.48 | 11.44 | 11.06 | 9.04 | 9.04 | 9.69 | 9.91 | 9.15 | 8.82 | 10.92 | 11.25 | 9.04 | 9.80 | 8.98 |
| GTPBP10 | 5 | 11.18 | 8.84 | 8.51 | 9.98 | 9.23 | 8.55 | 12.33 | 11.69 | 9.72 | 9.97 | 10.01 | 9.24 | 8.89 | 12.01 | 9.85 |
| GTPBP2 | 1 | 6.77 | 7.30 | 13.42 | 11.67 | 11.80 | 8.59 | 13.38 | 5.96 | 9.29 | 11.82 | 7.04 | 12.54 | 10.20 | 9.67 | 10.55 |
| GTPBP3 | 3 | 10.86 | 9.44 | 8.82 | 9.95 | 7.97 | 7.14 | 12.31 | 12.02 | 10.73 | 10.76 | 10.15 | 9.39 | 7.56 | 12.16 | 10.74 |
| GTPBP4 | 26 | 10.60 | 8.94 | 9.50 | 9.90 | 8.41 | 7.98 | 11.09 | 11.63 | 10.28 | 11.66 | 9.77 | 9.70 | 8.20 | 11.36 | 10.97 |
| GTPBP6 | 8 | 10.78 | 7.88 | 8.82 | 9.99 | 10.26 | 8.67 | 13.05 | 11.82 | 9.31 | 9.42 | 9.33 | 9.40 | 9.46 | 12.43 | 9.37 |
| GTPBP8 | 2 | 10.89 | 11.06 | 7.81 | 10.37 | 11.46 | 11.28 | 9.59 | 9.51 | 9.33 | 8.71 | 10.97 | 9.09 | 11.37 | 9.55 | 9.02 |
| GTSE1 | 6 | 9.54 | 10.02 | 14.16 | 13.45 | 7.56 | 7.67 | 10.87 | 10.12 | 8.48 | 8.13 | 9.78 | 13.81 | 7.61 | 10.49 | 8.31 |
| GUF1 | 1 | 10.87 | 8.50 | 11.16 | 14.76 | 7.03 | 6.94 | 10.01 | 9.17 | 11.03 | 10.52 | 9.69 | 12.96 | 6.98 | 9.59 | 10.78 |
| GUK1 | 7 | 9.47 | 9.78 | 6.14 | 6.48 | 15.28 | 15.38 | 7.81 | 7.12 | 11.20 | 11.34 | 9.62 | 6.31 | 15.33 | 7.46 | 11.27 |
| GULP1 | 6 | 9.62 | 10.53 | 11.28 | 10.95 | 10.52 | 10.87 | 9.15 | 9.59 | 9.09 | 8.40 | 10.07 | 11.11 | 10.69 | 9.37 | 8.75 |
| GUSB | 3 | 10.22 | 10.93 | 5.47 | 5.77 | 15.49 | 16.21 | 11.23 | 10.09 | 7.28 | 7.31 | 10.57 | 5.62 | 15.85 | 10.66 | 7.29 |
| GVINP1 | 3 | 9.19 | 11.43 | 8.88 | 8.78 | 10.81 | 11.95 | 9.24 | 8.82 | 11.19 | 9.69 | 10.31 | 8.83 | 11.38 | 9.03 | 10.44 |
| GYG1 | 9 | 10.57 | 11.24 | 9.46 | 8.56 | 8.85 | 8.73 | 12.83 | 11.55 | 9.00 | 9.22 | 10.90 | 9.01 | 8.79 | 12.19 | 9.11 |
| GYS1 | 12 | 9.24 | 9.90 | 8.88 | 8.34 | 11.05 | 10.95 | 10.56 | 10.11 | 10.45 | 10.52 | 9.57 | 8.61 | 11.00 | 10.33 | 10.49 |
| H1F0 | 6 | 11.52 | 8.80 | 12.03 | 14.53 | 7.05 | 6.97 | 4.81 | 4.36 | 11.46 | 18.47 | 10.16 | 13.28 | 7.01 | 4.59 | 14.97 |
| H1FX | 11 | 10.40 | 10.15 | 8.73 | 9.22 | 11.24 | 11.18 | 8.61 | 7.46 | 10.34 | 12.65 | 10.28 | 8.98 | 11.21 | 8.04 | 11.49 |
| H2AFV | 17 | 9.22 | 9.91 | 10.45 | 9.75 | 11.38 | 10.80 | 10.58 | 9.86 | 8.89 | 9.16 | 9.57 | 10.10 | 11.09 | 10.22 | 9.02 |
| H2AFX | 39 | 9.07 | 11.64 | 9.23 | 9.87 | 9.36 | 9.56 | 11.68 | 11.25 | 9.00 | 9.35 | 10.36 | 9.55 | 9.46 | 11.46 | 9.17 |
| H2AFY | 33 | 8.60 | 9.16 | 7.60 | 6.79 | 11.38 | 10.54 | 10.05 | 19.70 | 7.95 | 8.22 | 8.88 | 7.20 | 10.96 | 14.87 | 8.08 |
| H2AFY2 | 1 | 13.41 | 13.72 | 4.48 | 4.57 | 10.66 | 14.18 | 10.38 | 10.22 | 9.69 | 8.69 | 13.57 | 4.52 | 12.42 | 10.30 | 9.19 |
| H6PD | 1 | 8.56 | 8.56 | 5.77 | 5.77 | 9.83 | 11.17 | 17.61 | 17.31 | 7.34 | 8.08 | 8.56 | 5.77 | 10.50 | 17.46 | 7.71 |
| HABP4 | 1 | 9.04 | 10.06 | 14.19 | 13.34 | 11.66 | 12.54 | 6.00 | 5.35 | 8.77 | 9.06 | 9.55 | 13.76 | 12.10 | 5.67 | 8.91 |
| HACL1 | 9 | 8.55 | 8.85 | 14.42 | 14.74 | 7.39 | 7.58 | 8.00 | 7.21 | 11.37 | 11.90 | 8.70 | 14.58 | 7.48 | 7.60 | 11.63 |
| HADH | 21 | 11.24 | 9.73 | 7.05 | 6.94 | 13.70 | 12.68 | 11.67 | 10.23 | 8.47 | 8.29 | 10.48 | 7.00 | 13.19 | 10.95 | 8.38 |
| HADHA | 44 | 11.74 | 9.43 | 8.44 | 10.18 | 11.45 | 10.27 | 9.75 | 8.84 | 9.57 | 10.34 | 10.58 | 9.31 | 10.86 | 9.29 | 9.95 |
| HADHB | 31 | 11.64 | 9.57 | 9.00 | 10.67 | 11.33 | 10.54 | 9.92 | 8.89 | 9.01 | 9.43 | 10.61 | 9.84 | 10.94 | 9.40 | 9.22 |
| HAGH | 7 | 10.64 | 10.82 | 8.06 | 7.26 | 15.87 | 15.67 | 6.78 | 7.13 | 9.07 | 8.71 | 10.73 | 7.66 | 15.77 | 6.95 | 8.89 |
| HARS | 39 | 10.19 | 10.53 | 10.52 | 9.68 | 10.07 | 10.02 | 9.79 | 10.37 | 9.71 | 9.12 | 10.36 | 10.10 | 10.04 | 10.08 | 9.41 |
| HARS2 | 2 | 10.20 | 9.40 | 9.24 | 12.15 | 8.93 | 8.95 | 11.44 | 10.66 | 9.34 | 9.68 | 9.80 | 10.70 | 8.94 | 11.05 | 9.51 |
| HAT1 | 28 | 9.92 | 9.91 | 9.43 | 8.22 | 9.33 | 9.05 | 8.93 | 15.36 | 10.21 | 9.64 | 9.92 | 8.82 | 9.19 | 12.14 | 9.93 |
| HAUS1 | 3 | 10.34 | 11.97 | 9.77 | 9.51 | 10.09 | 11.00 | 8.76 | 8.49 | 10.12 | 9.96 | 11.15 | 9.64 | 10.55 | 8.62 | 10.04 |
| HAUS2 | 4 | 9.23 | 9.57 | 11.74 | 11.11 | 9.53 | 8.55 | 9.37 | 8.19 | 11.47 | 11.23 | 9.40 | 11.43 | 9.04 | 8.78 | 11.35 |
| HAUS3 | 4 | 9.34 | 11.28 | 9.67 | 9.92 | 9.53 | 10.92 | 10.30 | 9.96 | 9.88 | 9.20 | 10.31 | 9.79 | 10.23 | 10.13 | 9.54 |
| HAUS4 | 8 | 11.44 | 11.87 | 6.94 | 9.14 | 9.65 | 10.24 | 9.35 | 10.63 | 10.60 | 10.15 | 11.65 | 8.04 | 9.94 | 9.99 | 10.37 |

| | | | | | | | | | | | | | | | | |
|---|---|---|---|---|---|---|---|---|---|---|---|---|---|---|---|---|
| HAUS5 | 6 | 10.60 | 11.03 | 9.12 | 9.09 | 9.92 | 9.92 | 10.02 | 9.03 | 10.54 | 10.74 | 10.81 | 9.11 | 9.92 | 9.52 | 10.64 |
| HAUS6 | 8 | 10.11 | 10.42 | 9.94 | 8.92 | 9.57 | 8.91 | 9.86 | 8.87 | 11.74 | 11.66 | 10.27 | 9.43 | 9.24 | 9.36 | 11.70 |
| HAUS7 | 2 | 9.64 | 10.15 | 10.52 | 9.27 | 8.72 | 9.27 | 10.55 | 9.84 | 10.74 | 11.30 | 9.89 | 9.90 | 9.00 | 10.19 | 11.02 |
| HAUS8 | 3 | 9.40 | 10.83 | 10.49 | 10.05 | 8.91 | 9.84 | 9.86 | 9.77 | 11.06 | 9.79 | 10.11 | 10.27 | 9.37 | 9.82 | 10.43 |
| HAX1 | 5 | 9.26 | 9.92 | 10.24 | 10.09 | 11.77 | 11.81 | 10.28 | 9.49 | 8.28 | 8.87 | 9.59 | 10.16 | 11.79 | 9.88 | 8.57 |
| HBA1 | 6 | 5.42 | 6.29 | 11.24 | 10.57 | 7.63 | 8.10 | 22.12 | 19.09 | 4.78 | 4.77 | 5.85 | 10.91 | 7.86 | 20.60 | 4.77 |
| HBD | 2 | 2.58 | 2.66 | 9.84 | 10.16 | 4.46 | 4.18 | 31.70 | 30.39 | 2.04 | 1.97 | 2.62 | 10.00 | 4.32 | 31.05 | 2.01 |
| HBS1L | 30 | 10.22 | 10.99 | 10.09 | 9.62 | 10.54 | 10.66 | 9.97 | 9.26 | 9.44 | 9.22 | 10.60 | 9.85 | 10.60 | 9.62 | 9.33 |
| HCCS | 7 | 6.52 | 6.60 | 6.82 | 7.86 | 8.13 | 9.23 | 9.76 | 33.04 | 5.66 | 6.39 | 6.56 | 7.34 | 8.68 | 21.40 | 6.02 |
| HCFC1 | 58 | 9.02 | 9.98 | 10.86 | 9.83 | 10.97 | 10.97 | 10.66 | 9.88 | 8.99 | 8.85 | 9.50 | 10.35 | 10.97 | 10.27 | 8.92 |
| hCG_1988162 | 2 | 12.31 | 11.59 | 8.67 | 9.75 | 7.28 | 8.04 | 11.29 | 9.18 | 10.98 | 10.92 | 11.95 | 9.21 | 7.66 | 10.23 | 10.95 |
| HDAC1 | 13 | 9.40 | 10.45 | 10.34 | 8.63 | 10.24 | 9.26 | 11.90 | 11.03 | 9.68 | 9.08 | 9.92 | 9.48 | 9.75 | 11.46 | 9.38 |
| HDAC2 | 18 | 9.43 | 10.04 | 10.88 | 9.64 | 10.26 | 10.45 | 10.79 | 9.86 | 9.28 | 9.37 | 9.73 | 10.26 | 10.36 | 10.32 | 9.32 |
| HDAC3 | 2 | 9.76 | 9.99 | 10.77 | 10.22 | 9.34 | 9.39 | 11.34 | 11.35 | 9.12 | 8.71 | 9.88 | 10.50 | 9.37 | 11.34 | 8.91 |
| HDAC4 | 4 | 9.04 | 10.41 | 9.78 | 9.62 | 9.75 | 11.22 | 12.20 | 11.94 | 7.96 | 8.10 | 9.72 | 9.70 | 10.48 | 12.07 | 8.03 |
| HDAC6 | 5 | 9.83 | 10.97 | 7.84 | 7.90 | 12.37 | 12.73 | 8.94 | 9.54 | 10.24 | 9.64 | 10.40 | 7.87 | 12.55 | 9.24 | 9.94 |
| HDAC7 | 4 | 8.90 | 10.39 | 13.62 | 13.94 | 8.05 | 8.33 | 7.98 | 7.87 | 10.37 | 10.56 | 9.64 | 13.78 | 8.19 | 7.93 | 10.47 |
| HDDC2 | 3 | 10.21 | 12.40 | 12.33 | 9.97 | 8.50 | 8.21 | 8.74 | 8.28 | 11.09 | 10.27 | 11.30 | 11.15 | 8.35 | 8.51 | 10.68 |
| HDGF | 59 | 8.60 | 9.25 | 10.24 | 8.99 | 12.21 | 11.62 | 10.52 | 11.73 | 8.90 | 7.94 | 8.93 | 9.62 | 11.92 | 11.12 | 8.42 |
| HDGFRP2 | 15 | 9.25 | 11.72 | 10.06 | 9.25 | 11.36 | 12.02 | 9.65 | 8.06 | 9.46 | 9.18 | 10.49 | 9.65 | 11.69 | 8.85 | 9.32 |
| HDGFRP3 | 4 | 7.19 | 7.33 | 19.82 | 18.76 | 6.31 | 6.46 | 5.33 | 4.76 | 12.28 | 11.77 | 7.26 | 19.29 | 6.39 | 5.04 | 12.02 |
| HDHD1 | 3 | 9.10 | 10.11 | 9.95 | 9.13 | 10.93 | 12.59 | 11.48 | 9.97 | 9.02 | 7.72 | 9.61 | 9.54 | 11.76 | 10.72 | 8.37 |
| HDHD2 | 3 | 11.95 | 13.70 | 8.98 | 8.82 | 11.90 | 12.44 | 6.27 | 5.98 | 9.73 | 10.24 | 12.83 | 8.90 | 12.17 | 6.12 | 9.98 |
| HDHD3 | 2 | 10.42 | 9.85 | 10.61 | 8.86 | 10.77 | 9.41 | 11.03 | 9.45 | 10.78 | 8.81 | 10.13 | 9.74 | 10.09 | 10.24 | 9.80 |
| HDLBP | 91 | 10.15 | 10.98 | 10.98 | 10.73 | 10.54 | 10.88 | 8.96 | 8.24 | 9.48 | 9.06 | 10.56 | 10.86 | 10.71 | 8.60 | 9.27 |
| HEATR1 | 43 | 9.13 | 8.44 | 9.31 | 9.90 | 8.63 | 8.64 | 12.15 | 12.44 | 10.18 | 11.18 | 8.79 | 9.60 | 8.63 | 12.29 | 10.68 |
| HEATR2 | 15 | 9.92 | 10.02 | 9.58 | 10.06 | 8.75 | 9.06 | 10.23 | 10.69 | 10.94 | 10.75 | 9.97 | 9.82 | 8.91 | 10.46 | 10.84 |
| HEATR3 | 6 | 10.47 | 11.01 | 8.87 | 8.77 | 9.61 | 9.81 | 10.83 | 10.60 | 10.15 | 9.88 | 10.74 | 8.82 | 9.71 | 10.71 | 10.01 |
| HEATR5A | 6 | 12.02 | 11.33 | 9.46 | 9.52 | 9.66 | 9.86 | 7.76 | 8.02 | 11.25 | 11.13 | 11.67 | 9.49 | 9.76 | 7.89 | 11.19 |
| HEATR5B | 4 | 10.25 | 9.43 | 9.67 | 9.91 | 9.93 | 10.75 | 8.52 | 8.79 | 11.66 | 11.09 | 9.84 | 9.79 | 10.34 | 8.65 | 11.38 |
| HEATR6 | 2 | 10.80 | 9.64 | 12.74 | 12.83 | 9.03 | 8.60 | 7.18 | 6.03 | 12.20 | 10.96 | 10.22 | 12.79 | 8.81 | 6.60 | 11.58 |
| HEBP1 | 11 | 10.48 | 10.78 | 10.86 | 11.26 | 9.90 | 11.16 | 8.80 | 8.32 | 9.27 | 9.16 | 10.63 | 11.06 | 10.53 | 8.56 | 9.22 |
| HEBP2 | 5 | 8.76 | 9.85 | 11.25 | 10.33 | 10.89 | 11.59 | 12.64 | 9.85 | 7.53 | 7.29 | 9.31 | 10.79 | 11.24 | 11.25 | 7.41 |
| HECTD1 | 27 | 8.96 | 10.09 | 9.44 | 9.57 | 10.42 | 10.75 | 11.08 | 10.69 | 9.62 | 9.37 | 9.53 | 9.51 | 10.58 | 10.89 | 9.50 |
| HECTD3 | 7 | 10.86 | 11.80 | 10.65 | 11.06 | 7.86 | 8.75 | 9.19 | 8.38 | 11.44 | 10.01 | 11.33 | 10.86 | 8.30 | 8.78 | 10.73 |
| HECTD4 | 1 | 8.71 | 10.32 | 9.38 | 10.84 | 12.43 | 11.87 | 8.42 | 8.48 | 9.89 | 9.65 | 9.52 | 10.11 | 12.15 | 8.45 | 9.77 |
| HELLS | 23 | 10.62 | 11.41 | 10.93 | 11.13 | 9.17 | 9.29 | 9.19 | 9.14 | 9.61 | 9.52 | 11.01 | 11.03 | 9.23 | 9.16 | 9.56 |
| HELZ | 9 | 10.44 | 10.95 | 12.48 | 11.37 | 9.37 | 8.79 | 8.28 | 7.56 | 10.61 | 10.16 | 10.69 | 11.92 | 9.08 | 7.92 | 10.38 |
| HERC1 | 8 | 10.56 | 10.76 | 9.01 | 9.77 | 10.02 | 10.28 | 8.61 | 8.77 | 10.93 | 11.30 | 10.66 | 9.39 | 10.15 | 8.69 | 11.11 |
| HERC2 | 8 | 10.67 | 9.69 | 8.85 | 9.80 | 11.29 | 10.73 | 9.54 | 10.06 | 9.93 | 9.44 | 10.18 | 9.32 | 11.01 | 9.80 | 9.69 |
| HERC4 | 11 | 10.22 | 10.47 | 10.04 | 9.58 | 11.25 | 11.48 | 8.36 | 8.36 | 10.29 | 9.95 | 10.34 | 9.81 | 11.36 | 8.36 | 10.12 |
| HEXA | 18 | 10.40 | 10.08 | 9.77 | 9.04 | 11.76 | 11.63 | 9.85 | 9.82 | 8.92 | 8.73 | 10.24 | 9.40 | 11.69 | 9.83 | 8.83 |

| | | | | | | | | | | | | | | | | |
|---|---|---|---|---|---|---|---|---|---|---|---|---|---|---|---|---|
| HEXB | 23 | 9.84 | 10.02 | 7.18 | 6.97 | 14.67 | 14.93 | 13.68 | 12.42 | 5.31 | 4.97 | 9.93 | 7.08 | 14.80 | 13.05 | 5.14 |
| HEXIM1 | 8 | 10.55 | 12.26 | 9.14 | 8.50 | 10.76 | 11.78 | 10.61 | 6.43 | 10.51 | 9.46 | 11.41 | 8.82 | 11.27 | 8.52 | 9.99 |
| HGF | 1 | 0.98 | 0.40 | 1.49 | 0.20 | 2.43 | 0.30 | 5.00 | 88.24 | 0.40 | 0.54 | 0.69 | 0.85 | 1.37 | 46.62 | 0.47 |
| HGS | 19 | 9.99 | 11.59 | 11.96 | 11.80 | 8.59 | 8.95 | 8.07 | 7.48 | 10.89 | 10.67 | 10.79 | 11.88 | 8.77 | 7.78 | 10.78 |
| HIBADH | 10 | 12.55 | 11.09 | 9.00 | 10.07 | 7.88 | 7.85 | 10.81 | 10.02 | 10.19 | 10.54 | 11.82 | 9.54 | 7.87 | 10.41 | 10.36 |
| HIBCH | 8 | 12.42 | 10.01 | 10.64 | 11.08 | 11.90 | 10.91 | 10.73 | 10.40 | 6.12 | 5.80 | 11.21 | 10.86 | 11.41 | 10.56 | 5.96 |
| HIF1AN | 4 | 10.19 | 10.87 | 11.92 | 10.26 | 10.85 | 10.65 | 8.37 | 7.62 | 9.67 | 9.60 | 10.53 | 11.09 | 10.75 | 8.00 | 9.63 |
| HINT1 | 22 | 9.44 | 10.16 | 8.49 | 7.60 | 11.26 | 11.24 | 10.23 | 15.20 | 8.21 | 8.17 | 9.80 | 8.04 | 11.25 | 12.71 | 8.19 |
| HINT2 | 7 | 11.05 | 9.06 | 8.39 | 9.03 | 10.71 | 10.52 | 7.02 | 5.68 | 14.26 | 14.27 | 10.06 | 8.71 | 10.62 | 6.35 | 14.27 |
| HINT3 | 2 | 10.53 | 10.77 | 10.49 | 10.96 | 9.98 | 9.32 | 7.96 | 8.14 | 11.25 | 10.60 | 10.65 | 10.73 | 9.65 | 8.05 | 10.92 |
| HIP1 | 2 | 11.28 | 11.81 | 7.34 | 7.41 | 6.57 | 6.32 | 13.24 | 11.08 | 13.06 | 11.88 | 11.55 | 7.38 | 6.45 | 12.16 | 12.47 |
| HIP1R | 3 | 10.38 | 10.75 | 8.18 | 8.00 | 9.04 | 10.43 | 11.09 | 12.24 | 10.20 | 9.70 | 10.56 | 8.09 | 9.74 | 11.66 | 9.95 |
| HIRA | 1 | 9.81 | 10.51 | 10.43 | 10.56 | 8.94 | 8.72 | 9.96 | 10.39 | 9.87 | 10.80 | 10.16 | 10.49 | 8.83 | 10.18 | 10.33 |
| HIRIP3 | 10 | 10.02 | 10.93 | 11.17 | 10.20 | 9.69 | 9.67 | 9.31 | 7.97 | 11.03 | 10.00 | 10.47 | 10.68 | 9.68 | 8.64 | 10.52 |
| HIST1H1A | 1 | 13.67 | 8.17 | 12.24 | 20.23 | 7.72 | 7.25 | 3.76 | 3.10 | 7.82 | 16.04 | 10.92 | 16.23 | 7.49 | 3.43 | 11.93 |
| HIST1H1B | 36 | 15.94 | 8.14 | 6.65 | 10.63 | 8.70 | 7.31 | 7.82 | 7.13 | 9.24 | 18.43 | 12.04 | 8.64 | 8.01 | 7.48 | 13.83 |
| HIST1H1C | 9 | 16.12 | 8.61 | 7.57 | 11.79 | 8.62 | 6.69 | 7.04 | 6.08 | 8.95 | 18.53 | 12.37 | 9.68 | 7.65 | 6.56 | 13.74 |
| HIST1H1D | 38 | 12.98 | 9.21 | 8.03 | 11.58 | 8.73 | 8.09 | 7.88 | 7.34 | 9.17 | 16.98 | 11.10 | 9.81 | 8.41 | 7.61 | 13.08 |
| HIST1H1E | 9 | 13.75 | 8.93 | 8.08 | 11.88 | 8.28 | 8.34 | 7.12 | 6.24 | 9.51 | 17.86 | 11.34 | 9.98 | 8.31 | 6.68 | 13.68 |
| HIST1H3A | 21 | 18.68 | 6.69 | 5.84 | 12.98 | 7.84 | 6.20 | 5.73 | 5.37 | 8.54 | 22.14 | 12.68 | 9.41 | 7.02 | 5.55 | 15.34 |
| HIST1H4A | 66 | 18.63 | 6.60 | 6.54 | 12.88 | 8.23 | 6.26 | 5.59 | 5.00 | 8.10 | 22.15 | 12.62 | 9.71 | 7.25 | 5.30 | 15.12 |
| HIST2H2AA3 | 3 | 8.05 | 12.21 | 8.84 | 9.47 | 9.18 | 8.85 | 11.98 | 11.96 | 9.05 | 10.42 | 10.13 | 9.16 | 9.01 | 11.97 | 9.73 |
| HIST2H2AB | 2 | 4.76 | 9.55 | 12.68 | 9.53 | 11.87 | 11.08 | 12.47 | 11.16 | 7.34 | 9.57 | 7.15 | 11.10 | 11.48 | 11.81 | 8.45 |
| HIST2H2BC | 18 | 8.14 | 10.57 | 8.68 | 9.06 | 10.24 | 10.29 | 12.08 | 12.52 | 8.63 | 9.79 | 9.36 | 8.87 | 10.26 | 12.30 | 9.21 |
| HIST2H2BF | 111 | 9.63 | 11.14 | 9.50 | 8.31 | 10.18 | 9.60 | 12.41 | 10.73 | 9.20 | 9.30 | 10.39 | 8.91 | 9.89 | 11.57 | 9.25 |
| HJURP | 3 | 9.22 | 10.19 | 9.40 | 9.61 | 8.18 | 8.16 | 14.32 | 12.39 | 9.29 | 9.23 | 9.71 | 9.50 | 8.17 | 13.35 | 9.26 |
| HK1 | 45 | 10.01 | 8.83 | 11.18 | 10.63 | 10.30 | 9.97 | 9.31 | 9.41 | 10.21 | 10.14 | 9.42 | 10.90 | 10.14 | 9.36 | 10.18 |
| HK2 | 25 | 7.35 | 7.46 | 10.50 | 9.98 | 7.96 | 7.65 | 14.11 | 17.98 | 8.65 | 8.35 | 7.40 | 10.24 | 7.80 | 16.05 | 8.50 |
| HLA-A | 11 | 12.85 | 10.27 | 5.52 | 6.72 | 12.80 | 13.13 | 7.24 | 6.14 | 12.60 | 12.74 | 11.56 | 6.12 | 12.96 | 6.69 | 12.67 |
| HLA-B | 4 | 12.67 | 9.87 | 3.13 | 2.08 | 19.51 | 17.04 | 5.23 | 3.51 | 13.01 | 13.95 | 11.27 | 2.61 | 18.27 | 4.37 | 13.48 |
| HLA-C | 13 | 13.44 | 10.42 | 4.34 | 5.42 | 14.76 | 14.75 | 6.45 | 5.66 | 12.52 | 12.22 | 11.93 | 4.88 | 14.76 | 6.06 | 12.37 |
| HLA-DPA1 | 1 | 15.62 | 11.32 | 5.23 | 3.59 | 21.80 | 20.63 | 4.79 | 2.64 | 8.41 | 5.98 | 13.47 | 4.41 | 21.22 | 3.71 | 7.19 |
| HLA-DRA | 2 | 21.82 | 16.94 | 1.63 | 1.37 | 26.18 | 25.81 | 1.32 | 0.91 | 2.40 | 1.63 | 19.38 | 1.50 | 26.00 | 1.11 | 2.02 |
| HLA-DRB1 | 2 | 20.76 | 16.93 | 1.85 | 1.78 | 25.44 | 25.30 | 1.82 | 1.58 | 2.41 | 2.12 | 18.85 | 1.82 | 25.37 | 1.70 | 2.27 |
| HLCS | 3 | 9.81 | 10.44 | 7.54 | 7.50 | 12.06 | 12.79 | 11.84 | 10.52 | 9.13 | 8.38 | 10.12 | 7.52 | 12.43 | 11.18 | 8.75 |
| HLTF | 16 | 10.20 | 10.92 | 9.51 | 9.54 | 8.64 | 9.11 | 14.78 | 13.90 | 7.03 | 6.36 | 10.56 | 9.53 | 8.87 | 14.34 | 6.69 |
| HM13 | 11 | 13.72 | 7.65 | 7.10 | 11.81 | 8.57 | 8.38 | 9.57 | 8.57 | 10.67 | 13.96 | 10.68 | 9.45 | 8.47 | 9.07 | 12.32 |
| HMBS | 5 | 10.10 | 10.92 | 10.21 | 9.73 | 10.63 | 10.10 | 10.14 | 8.91 | 10.14 | 9.13 | 10.51 | 9.97 | 10.36 | 9.52 | 9.63 |
| HMCES | 4 | 11.56 | 11.79 | 9.72 | 8.78 | 11.22 | 10.25 | 8.62 | 7.51 | 10.67 | 9.87 | 11.68 | 9.25 | 10.73 | 8.07 | 10.27 |
| HMG20A | 7 | 9.10 | 10.14 | 11.99 | 11.95 | 8.27 | 8.88 | 10.59 | 9.73 | 9.85 | 9.49 | 9.62 | 11.97 | 8.58 | 10.16 | 9.67 |
| HMG20B | 2 | 9.33 | 9.78 | 9.73 | 9.93 | 12.04 | 12.74 | 9.92 | 8.96 | 9.41 | 8.17 | 9.55 | 9.83 | 12.39 | 9.44 | 8.79 |
| HMGA1 | 14 | 8.65 | 8.47 | 12.58 | 12.41 | 9.11 | 9.02 | 10.42 | 9.42 | 9.71 | 10.21 | 8.56 | 12.50 | 9.06 | 9.92 | 9.96 |

| | | | | | | | | | | | | | | | | |
|---|---|---|---|---|---|---|---|---|---|---|---|---|---|---|---|---|
| HMGB1 | 29 | 9.14 | 10.47 | 12.16 | 10.80 | 10.13 | 9.92 | 10.92 | 11.45 | 7.54 | 7.48 | 9.80 | 11.48 | 10.03 | 11.18 | 7.51 |
| HMGB2 | 8 | 8.40 | 9.83 | 10.89 | 6.34 | 12.75 | 11.33 | 12.72 | 8.97 | 9.18 | 9.59 | 9.11 | 8.61 | 12.04 | 10.85 | 9.38 |
| HMGB3 | 13 | 7.75 | 9.72 | 8.70 | 7.84 | 11.41 | 11.33 | 13.66 | 12.73 | 8.10 | 8.75 | 8.74 | 8.27 | 11.37 | 13.19 | 8.43 |
| HMGCL | 4 | 12.38 | 11.17 | 8.27 | 9.42 | 9.12 | 9.78 | 9.85 | 8.99 | 10.00 | 11.02 | 11.77 | 8.85 | 9.45 | 9.42 | 10.51 |
| HMGCS1 | 16 | 9.96 | 10.73 | 14.18 | 14.09 | 10.28 | 10.51 | 8.40 | 7.61 | 7.12 | 7.12 | 10.35 | 14.14 | 10.40 | 8.00 | 7.12 |
| HMGN1 | 8 | 11.17 | 11.29 | 7.54 | 7.27 | 12.45 | 12.40 | 9.93 | 6.96 | 10.17 | 10.81 | 11.23 | 7.40 | 12.43 | 8.45 | 10.49 |
| HMGN2 | 5 | 11.36 | 17.11 | 8.64 | 9.10 | 9.48 | 8.84 | 8.56 | 7.03 | 9.17 | 10.72 | 14.24 | 8.87 | 9.16 | 7.79 | 9.95 |
| HMGN3 | 11 | 10.42 | 14.77 | 10.20 | 10.47 | 10.82 | 11.24 | 8.06 | 6.40 | 8.16 | 9.47 | 12.59 | 10.34 | 11.03 | 7.23 | 8.81 |
| HMGN4 | 5 | 5.94 | 11.17 | 11.73 | 9.17 | 11.28 | 9.55 | 11.12 | 7.79 | 9.37 | 12.90 | 8.55 | 10.45 | 10.41 | 9.45 | 11.13 |
| HMGN5 | 30 | 10.09 | 12.19 | 10.73 | 10.54 | 15.06 | 15.76 | 8.25 | 6.10 | 5.87 | 5.42 | 11.14 | 10.64 | 15.41 | 7.17 | 5.64 |
| HMHA1 | 7 | 9.58 | 9.26 | 12.84 | 11.75 | 7.94 | 7.58 | 7.83 | 6.74 | 13.33 | 13.14 | 9.42 | 12.30 | 7.76 | 7.29 | 13.24 |
| HMMR | 17 | 9.28 | 10.99 | 8.48 | 8.82 | 9.71 | 10.09 | 12.43 | 11.77 | 9.29 | 9.13 | 10.13 | 8.65 | 9.90 | 12.10 | 9.21 |
| HMOX1 | 6 | 17.58 | 14.68 | 8.60 | 8.55 | 8.85 | 8.10 | 6.89 | 5.34 | 10.25 | 11.15 | 16.13 | 8.58 | 8.48 | 6.12 | 10.70 |
| HMOX2 | 8 | 12.14 | 7.08 | 8.98 | 11.08 | 9.64 | 7.84 | 10.40 | 11.05 | 9.66 | 12.11 | 9.61 | 10.03 | 8.74 | 10.73 | 10.89 |
| HN1 | 17 | 10.46 | 11.50 | 10.90 | 13.20 | 9.07 | 9.70 | 6.89 | 3.80 | 11.98 | 12.50 | 10.98 | 12.05 | 9.39 | 5.34 | 12.24 |
| HN1L | 26 | 10.91 | 12.17 | 11.64 | 8.85 | 10.69 | 10.99 | 7.78 | 6.54 | 10.58 | 9.85 | 11.54 | 10.24 | 10.84 | 7.16 | 10.22 |
| HNMT | 4 | 9.45 | 9.89 | 8.46 | 8.32 | 23.53 | 22.60 | 6.90 | 6.43 | 2.55 | 1.86 | 9.67 | 8.39 | 23.07 | 6.66 | 2.21 |
| HNRNPA0 | 15 | 8.66 | 9.43 | 9.95 | 9.42 | 8.66 | 8.44 | 11.64 | 17.20 | 8.32 | 8.29 | 9.04 | 9.68 | 8.55 | 14.42 | 8.31 |
| HNRNPA1 | 83 | 8.81 | 9.57 | 10.52 | 9.64 | 11.21 | 11.17 | 10.93 | 11.67 | 8.49 | 7.99 | 9.19 | 10.08 | 11.19 | 11.30 | 8.24 |
| HNRNPA2B1 | 115 | 9.15 | 10.74 | 11.54 | 6.62 | 11.20 | 10.19 | 12.00 | 9.53 | 9.52 | 9.52 | 9.94 | 9.08 | 10.70 | 10.76 | 9.52 |
| HNRNPA3 | 41 | 10.40 | 10.72 | 9.86 | 7.35 | 11.04 | 10.86 | 11.71 | 10.18 | 9.10 | 8.78 | 10.56 | 8.61 | 10.95 | 10.94 | 8.94 |
| HNRNPAB | 31 | 8.58 | 9.90 | 9.91 | 8.05 | 10.58 | 9.38 | 13.48 | 11.99 | 9.04 | 9.10 | 9.24 | 8.98 | 9.98 | 12.74 | 9.07 |
| HNRNPC | 64 | 8.89 | 10.87 | 9.95 | 9.37 | 10.44 | 10.83 | 11.27 | 10.23 | 9.13 | 9.03 | 9.88 | 9.66 | 10.63 | 10.75 | 9.08 |
| HNRNPD | 29 | 9.61 | 9.97 | 11.91 | 7.57 | 11.67 | 10.63 | 10.62 | 9.02 | 9.66 | 9.33 | 9.79 | 9.74 | 11.15 | 9.82 | 9.49 |
| HNRNPDL | 38 | 8.79 | 9.74 | 10.60 | 8.22 | 11.16 | 10.21 | 9.49 | 12.85 | 9.70 | 9.24 | 9.27 | 9.41 | 10.69 | 11.17 | 9.47 |
| HNRNPF | 22 | 8.20 | 9.16 | 11.02 | 9.75 | 9.63 | 9.45 | 10.34 | 14.78 | 8.73 | 8.93 | 8.68 | 10.38 | 9.54 | 12.56 | 8.83 |
| HNRNPH1 | 30 | 9.63 | 11.01 | 9.29 | 8.73 | 9.53 | 9.99 | 11.60 | 11.45 | 9.56 | 9.22 | 10.32 | 9.01 | 9.76 | 11.52 | 9.39 |
| HNRNPH2 | 33 | 8.76 | 10.97 | 8.86 | 8.87 | 10.63 | 10.92 | 10.35 | 14.30 | 8.30 | 8.04 | 9.87 | 8.87 | 10.78 | 12.32 | 8.17 |
| HNRNPH3 | 17 | 9.47 | 10.32 | 9.49 | 8.16 | 11.16 | 11.46 | 11.06 | 9.79 | 9.66 | 9.42 | 9.90 | 8.83 | 11.31 | 10.42 | 9.54 |
| HNRNPK | 78 | 9.69 | 11.68 | 9.71 | 9.17 | 10.31 | 10.35 | 10.54 | 10.99 | 9.07 | 8.49 | 10.69 | 9.44 | 10.33 | 10.76 | 8.78 |
| HNRNPL | 72 | 8.45 | 9.77 | 10.21 | 8.21 | 10.55 | 10.30 | 11.30 | 13.05 | 9.27 | 8.91 | 9.11 | 9.21 | 10.42 | 12.17 | 9.09 |
| HNRNPLL | 15 | 9.28 | 9.69 | 9.11 | 9.35 | 11.27 | 11.98 | 9.92 | 11.29 | 9.12 | 8.99 | 9.48 | 9.23 | 11.62 | 10.61 | 9.06 |
| HNRNPM | 145 | 8.66 | 10.27 | 10.99 | 10.13 | 9.64 | 9.81 | 10.89 | 11.66 | 9.12 | 8.83 | 9.46 | 10.56 | 9.72 | 11.28 | 8.97 |
| HNRNPR | 69 | 8.82 | 9.10 | 9.34 | 7.70 | 9.47 | 9.27 | 10.68 | 18.63 | 8.68 | 8.30 | 8.96 | 8.52 | 9.37 | 14.66 | 8.49 |
| HNRNPU | 112 | 9.25 | 9.84 | 8.24 | 7.80 | 11.23 | 11.28 | 10.50 | 15.00 | 8.59 | 8.27 | 9.54 | 8.02 | 11.26 | 12.75 | 8.43 |
| HNRNPUL1 | 21 | 9.12 | 9.37 | 10.84 | 8.44 | 10.01 | 9.37 | 12.59 | 12.32 | 9.13 | 8.81 | 9.24 | 9.64 | 9.69 | 12.46 | 8.97 |
| HNRNPUL2 | 59 | 8.88 | 9.43 | 10.38 | 9.79 | 10.42 | 10.63 | 10.29 | 12.28 | 9.18 | 8.70 | 9.16 | 10.09 | 10.53 | 11.29 | 8.94 |
| HOMER3 | 3 | 7.98 | 8.82 | 13.87 | 11.48 | 8.88 | 8.43 | 11.59 | 10.30 | 10.21 | 8.45 | 8.40 | 12.67 | 8.65 | 10.94 | 9.33 |
| HOOK1 | 8 | 12.98 | 14.51 | 8.65 | 8.89 | 13.25 | 14.86 | 4.05 | 3.76 | 9.67 | 9.38 | 13.75 | 8.77 | 14.05 | 3.90 | 9.53 |
| HOOK2 | 1 | 8.97 | 9.41 | 12.52 | 12.41 | 11.34 | 11.43 | 9.00 | 7.67 | 9.16 | 8.11 | 9.19 | 12.46 | 11.38 | 8.33 | 8.64 |
| HOOK3 | 13 | 9.71 | 10.66 | 10.35 | 10.62 | 9.91 | 10.80 | 9.45 | 8.96 | 10.03 | 9.51 | 10.19 | 10.49 | 10.35 | 9.20 | 9.77 |
| HOXC10 | 5 | 10.99 | 11.34 | 6.83 | 7.35 | 9.92 | 10.70 | 11.99 | 10.64 | 11.32 | 8.91 | 11.17 | 7.09 | 10.31 | 11.32 | 10.12 |

| | | | | | | | | | | | | | | | | |
|---|---|---|---|---|---|---|---|---|---|---|---|---|---|---|---|---|
| HP1BP3 | 23 | 13.49 | 10.44 | 8.41 | 10.33 | 8.64 | 8.04 | 8.47 | 7.40 | 10.48 | 14.29 | 11.97 | 9.37 | 8.34 | 7.93 | 12.38 |
| HPCAL1 | 21 | 11.32 | 12.23 | 8.27 | 7.53 | 11.04 | 11.64 | 8.80 | 8.11 | 10.47 | 10.59 | 11.78 | 7.90 | 11.34 | 8.45 | 10.53 |
| HPD | 5 | 9.63 | 10.06 | 6.70 | 6.46 | 16.70 | 15.96 | 2.89 | 2.51 | 14.33 | 14.76 | 9.85 | 6.58 | 16.33 | 2.70 | 14.54 |
| HPDL | 6 | 9.66 | 9.03 | 7.93 | 9.06 | 9.50 | 9.32 | 11.60 | 11.83 | 11.11 | 10.97 | 9.34 | 8.49 | 9.41 | 11.72 | 11.04 |
| HPRT1 | 20 | 8.36 | 10.03 | 8.33 | 7.78 | 13.19 | 13.16 | 11.36 | 10.45 | 8.73 | 8.63 | 9.19 | 8.05 | 13.18 | 10.90 | 8.68 |
| HPS3 | 4 | 9.75 | 10.10 | 9.65 | 9.01 | 10.50 | 11.34 | 10.36 | 10.75 | 9.38 | 9.15 | 9.92 | 9.33 | 10.92 | 10.56 | 9.27 |
| HPS5 | 2 | 9.33 | 10.18 | 9.64 | 8.33 | 9.95 | 9.89 | 7.75 | 10.79 | 12.16 | 11.98 | 9.75 | 8.98 | 9.92 | 9.27 | 12.07 |
| HPS6 | 5 | 9.66 | 10.24 | 9.80 | 10.48 | 10.32 | 10.85 | 10.39 | 9.15 | 9.66 | 9.45 | 9.95 | 10.14 | 10.58 | 9.77 | 9.55 |
| HRSP12 | 6 | 6.46 | 6.94 | 7.91 | 7.46 | 7.82 | 7.77 | 15.22 | 25.64 | 7.17 | 7.61 | 6.70 | 7.68 | 7.79 | 20.43 | 7.39 |
| HS1BP3 | 3 | 9.65 | 10.29 | 9.65 | 9.75 | 15.19 | 15.95 | 8.99 | 7.70 | 6.52 | 6.31 | 9.97 | 9.70 | 15.57 | 8.34 | 6.41 |
| HSBP1 | 5 | 13.00 | 12.39 | 3.48 | 13.57 | 9.75 | 15.48 | 5.43 | 1.67 | 10.78 | 14.44 | 12.70 | 8.52 | 12.62 | 3.55 | 12.61 |
| HSCB | 3 | 11.45 | 11.04 | 8.85 | 9.53 | 4.85 | 5.67 | 11.89 | 10.89 | 13.18 | 12.64 | 11.25 | 9.19 | 5.26 | 11.39 | 12.91 |
| HSD17B10 | 22 | 9.58 | 8.10 | 8.75 | 9.04 | 11.66 | 10.63 | 10.55 | 14.68 | 8.58 | 8.43 | 8.84 | 8.90 | 11.14 | 12.62 | 8.50 |
| HSD17B11 | 8 | 9.39 | 7.05 | 8.07 | 10.11 | 15.23 | 14.32 | 8.97 | 9.38 | 7.96 | 9.53 | 8.22 | 9.09 | 14.77 | 9.17 | 8.75 |
| HSD17B12 | 5 | 12.44 | 7.07 | 8.69 | 12.12 | 10.75 | 10.20 | 8.84 | 8.87 | 9.24 | 11.78 | 9.75 | 10.41 | 10.47 | 8.85 | 10.51 |
| HSD17B4 | 50 | 9.35 | 9.97 | 9.62 | 8.98 | 13.24 | 13.41 | 9.63 | 9.53 | 8.38 | 7.88 | 9.66 | 9.30 | 13.33 | 9.58 | 8.13 |
| HSDL1 | 2 | 11.55 | 10.68 | 6.70 | 7.06 | 11.71 | 12.81 | 9.04 | 8.31 | 10.92 | 11.22 | 11.12 | 6.88 | 12.26 | 8.68 | 11.07 |
| HSDL2 | 17 | 11.65 | 10.78 | 8.76 | 9.14 | 10.99 | 10.29 | 7.50 | 6.50 | 12.11 | 12.29 | 11.21 | 8.95 | 10.64 | 7.00 | 12.20 |
| HSF1 | 10 | 9.15 | 9.83 | 10.44 | 10.14 | 9.26 | 9.44 | 11.65 | 10.54 | 9.90 | 9.64 | 9.49 | 10.29 | 9.35 | 11.10 | 9.77 |
| HSP90AA1 | 278 | 9.54 | 11.57 | 10.52 | 9.50 | 10.24 | 10.39 | 9.79 | 9.37 | 9.74 | 9.35 | 10.56 | 10.01 | 10.31 | 9.58 | 9.54 |
| HSP90AA4P | 1 | 12.93 | 10.39 | 11.51 | 8.88 | 10.09 | 10.60 | 8.94 | 8.54 | 9.59 | 8.53 | 11.66 | 10.19 | 10.34 | 8.74 | 9.06 |
| HSP90AB1 | 130 | 8.86 | 10.30 | 10.75 | 9.76 | 8.73 | 8.59 | 9.97 | 12.83 | 10.25 | 9.95 | 9.58 | 10.26 | 8.66 | 11.40 | 10.10 |
| HSP90B1 | 164 | 9.89 | 8.18 | 8.97 | 10.02 | 11.23 | 10.71 | 12.24 | 11.28 | 8.50 | 8.98 | 9.03 | 9.50 | 10.97 | 11.76 | 8.74 |
| HSPA12A | 4 | 12.50 | 12.11 | 13.43 | 13.05 | 8.47 | 8.89 | 6.29 | 5.34 | 10.44 | 9.49 | 12.30 | 13.24 | 8.68 | 5.81 | 9.97 |
| HSPA13 | 7 | 11.11 | 7.52 | 8.62 | 9.42 | 10.85 | 10.96 | 10.73 | 10.26 | 12.17 | 8.36 | 9.31 | 9.02 | 10.90 | 10.50 | 10.27 |
| HSPA14 | 15 | 8.50 | 9.31 | 10.22 | 10.19 | 10.16 | 10.31 | 11.61 | 10.81 | 9.75 | 9.15 | 8.90 | 10.20 | 10.23 | 11.21 | 9.45 |
| HSPA1A | 129 | 9.65 | 12.36 | 10.10 | 10.07 | 9.14 | 9.33 | 8.85 | 9.53 | 10.66 | 10.32 | 11.00 | 10.08 | 9.23 | 9.19 | 10.49 |
| HSPA1L | 4 | 10.35 | 10.52 | 10.09 | 9.16 | 10.73 | 11.43 | 8.18 | 7.40 | 11.14 | 11.01 | 10.43 | 9.62 | 11.08 | 7.79 | 11.08 |
| HSPA2 | 4 | 10.33 | 12.22 | 8.18 | 8.26 | 10.62 | 11.17 | 6.60 | 5.91 | 13.43 | 13.27 | 11.28 | 8.22 | 10.90 | 6.26 | 13.35 |
| HSPA4 | 120 | 9.13 | 9.66 | 10.12 | 9.74 | 8.92 | 9.05 | 11.16 | 14.13 | 9.31 | 8.78 | 9.39 | 9.93 | 8.98 | 12.65 | 9.05 |
| HSPA4L | 41 | 10.07 | 11.19 | 11.93 | 10.61 | 10.16 | 9.68 | 8.35 | 7.63 | 10.24 | 10.15 | 10.63 | 11.27 | 9.92 | 7.99 | 10.19 |
| HSPA5 | 180 | 9.78 | 9.67 | 9.26 | 10.14 | 10.68 | 10.29 | 10.80 | 9.72 | 9.62 | 10.05 | 9.73 | 9.70 | 10.48 | 10.26 | 9.83 |
| HSPA8 | 250 | 9.11 | 11.17 | 11.11 | 10.75 | 9.70 | 9.61 | 9.97 | 11.07 | 8.95 | 8.56 | 10.14 | 10.93 | 9.66 | 10.52 | 8.75 |
| HSPA9 | 111 | 9.38 | 9.05 | 8.42 | 9.33 | 9.20 | 8.59 | 12.01 | 14.85 | 9.48 | 9.69 | 9.21 | 8.88 | 8.89 | 13.43 | 9.59 |
| HSPB1 | 87 | 9.01 | 12.34 | 17.13 | 16.52 | 6.17 | 6.25 | 5.20 | 4.55 | 11.33 | 11.50 | 10.67 | 16.82 | 6.21 | 4.88 | 11.41 |
| HSPB11 | 1 | 11.86 | 12.11 | 8.23 | 7.85 | 10.10 | 11.60 | 10.29 | 10.92 | 8.67 | 8.36 | 11.99 | 8.04 | 10.85 | 10.61 | 8.51 |
| HSPB8 | 10 | 10.70 | 12.66 | 10.75 | 10.54 | 10.69 | 11.27 | 6.12 | 5.07 | 11.20 | 11.00 | 11.68 | 10.64 | 10.98 | 5.60 | 11.10 |
| HSPBP1 | 18 | 9.29 | 10.98 | 8.29 | 7.71 | 10.37 | 11.13 | 11.79 | 11.89 | 9.49 | 9.07 | 10.14 | 8.00 | 10.75 | 11.84 | 9.28 |
| HSPD1 | 262 | 9.70 | 9.22 | 9.93 | 10.91 | 9.40 | 8.59 | 10.26 | 12.54 | 9.64 | 9.81 | 9.46 | 10.42 | 8.99 | 11.40 | 9.72 |
| HSPE1 | 65 | 8.79 | 9.65 | 11.28 | 11.14 | 9.48 | 9.28 | 11.14 | 10.28 | 9.53 | 9.44 | 9.22 | 11.21 | 9.38 | 10.71 | 9.48 |
| HSPG2 | 8 | 12.37 | 9.89 | 10.12 | 9.85 | 6.32 | 6.16 | 9.95 | 9.17 | 12.87 | 13.29 | 11.13 | 9.99 | 6.24 | 9.56 | 13.08 |
| HSPH1 | 74 | 8.99 | 9.60 | 11.91 | 10.86 | 10.47 | 10.30 | 8.70 | 14.02 | 7.69 | 7.46 | 9.30 | 11.38 | 10.39 | 11.36 | 7.57 |

| | | | | | | | | | | | | | | | | |
|---|---|---|---|---|---|---|---|---|---|---|---|---|---|---|---|---|
| HTATIP2 | 25 | 10.84 | 11.16 | 8.39 | 7.47 | 11.28 | 11.56 | 11.79 | 10.24 | 8.84 | 8.44 | 11.00 | 7.93 | 11.42 | 11.01 | 8.64 |
| HTATSF1 | 37 | 9.80 | 10.48 | 7.96 | 7.64 | 12.29 | 12.89 | 12.70 | 12.28 | 7.23 | 6.73 | 10.14 | 7.80 | 12.59 | 12.49 | 6.98 |
| HTRA1 | 5 | 4.22 | 4.63 | 12.02 | 13.14 | 4.74 | 4.95 | 21.93 | 21.41 | 6.83 | 6.13 | 4.43 | 12.58 | 4.85 | 21.67 | 6.48 |
| HTRA2 | 5 | 9.46 | 8.61 | 12.81 | 11.27 | 8.07 | 8.17 | 10.76 | 9.47 | 10.46 | 10.92 | 9.03 | 12.04 | 8.12 | 10.12 | 10.69 |
| HTT | 10 | 9.65 | 9.51 | 8.98 | 9.64 | 11.65 | 11.60 | 9.23 | 9.47 | 10.18 | 10.08 | 9.58 | 9.31 | 11.63 | 9.35 | 10.13 |
| HUS1 | 3 | 10.19 | 10.36 | 10.23 | 9.39 | 9.95 | 9.84 | 11.67 | 12.00 | 8.48 | 7.87 | 10.28 | 9.81 | 9.89 | 11.84 | 8.18 |
| HUWE1 | 100 | 8.94 | 9.61 | 8.68 | 8.29 | 11.44 | 11.31 | 11.50 | 11.99 | 9.11 | 9.14 | 9.27 | 8.48 | 11.38 | 11.74 | 9.12 |
| HYI | 2 | 7.50 | 7.90 | 10.01 | 11.26 | 8.81 | 9.15 | 7.97 | 8.18 | 14.76 | 14.46 | 7.70 | 10.64 | 8.98 | 8.07 | 14.61 |
| HYOU1 | 44 | 9.76 | 8.16 | 8.66 | 9.60 | 12.41 | 11.07 | 10.90 | 10.04 | 9.08 | 10.32 | 8.96 | 9.13 | 11.74 | 10.47 | 9.70 |
| HYPK | 13 | 9.37 | 10.28 | 10.09 | 9.63 | 10.29 | 10.68 | 10.54 | 8.34 | 10.57 | 10.19 | 9.83 | 9.86 | 10.48 | 9.44 | 10.38 |
| IAH1 | 2 | 8.11 | 11.69 | 10.55 | 8.86 | 12.27 | 12.87 | 9.16 | 7.92 | 9.42 | 9.16 | 9.90 | 9.71 | 12.57 | 8.54 | 9.29 |
| IARS | 70 | 8.54 | 8.95 | 10.57 | 10.02 | 10.15 | 10.26 | 9.33 | 10.08 | 11.05 | 11.06 | 8.74 | 10.29 | 10.21 | 9.70 | 11.05 |
| IARS2 | 28 | 10.62 | 8.72 | 6.95 | 7.16 | 12.79 | 11.61 | 11.21 | 10.00 | 10.28 | 10.66 | 9.67 | 7.05 | 12.20 | 10.61 | 10.47 |
| IBTK | 5 | 8.80 | 9.41 | 10.57 | 7.92 | 9.66 | 9.63 | 13.32 | 10.56 | 10.05 | 10.07 | 9.10 | 9.25 | 9.64 | 11.94 | 10.06 |
| ICAM1 | 2 | 11.71 | 10.02 | 11.16 | 11.93 | 7.63 | 7.97 | 9.62 | 7.93 | 10.98 | 11.07 | 10.86 | 11.54 | 7.80 | 8.78 | 11.02 |
| ICMT | 3 | 13.17 | 6.00 | 6.55 | 10.81 | 8.23 | 6.42 | 10.01 | 9.29 | 12.68 | 16.87 | 9.58 | 8.68 | 7.32 | 9.65 | 14.77 |
| ICT1 | 8 | 9.88 | 9.20 | 9.92 | 10.45 | 9.95 | 9.97 | 10.99 | 9.63 | 9.99 | 10.03 | 9.54 | 10.18 | 9.96 | 10.31 | 10.01 |
| ID1 | 3 | 9.69 | 11.16 | 8.46 | 7.06 | 12.47 | 12.05 | 11.38 | 10.01 | 9.14 | 8.58 | 10.43 | 7.76 | 12.26 | 10.70 | 8.86 |
| ID3 | 1 | 12.61 | 12.54 | 7.10 | 9.03 | 9.31 | 11.17 | 7.81 | 7.29 | 13.34 | 9.82 | 12.57 | 8.07 | 10.24 | 7.55 | 11.58 |
| IDE | 42 | 10.13 | 10.75 | 10.45 | 9.21 | 11.34 | 11.07 | 9.42 | 8.85 | 9.52 | 9.26 | 10.44 | 9.83 | 11.20 | 9.14 | 9.39 |
| IDH1 | 38 | 9.92 | 10.37 | 9.42 | 8.21 | 11.88 | 11.74 | 9.76 | 11.62 | 8.80 | 8.27 | 10.15 | 8.82 | 11.81 | 10.69 | 8.54 |
| IDH2 | 17 | 12.05 | 11.19 | 9.48 | 11.27 | 13.16 | 12.25 | 7.32 | 7.15 | 7.87 | 8.26 | 11.62 | 10.37 | 12.71 | 7.24 | 8.07 |
| IDH3A | 23 | 11.59 | 9.53 | 8.06 | 7.61 | 11.46 | 9.86 | 10.44 | 9.11 | 11.06 | 11.27 | 10.56 | 7.83 | 10.66 | 9.78 | 11.17 |
| IDH3B | 22 | 10.40 | 9.02 | 8.58 | 9.93 | 9.65 | 9.07 | 9.98 | 9.43 | 11.63 | 12.31 | 9.71 | 9.25 | 9.36 | 9.71 | 11.97 |
| IDH3G | 9 | 11.26 | 9.62 | 6.89 | 8.17 | 11.16 | 10.92 | 10.27 | 10.23 | 10.51 | 10.97 | 10.44 | 7.53 | 11.04 | 10.25 | 10.74 |
| IDI1 | 4 | 10.04 | 10.22 | 9.07 | 7.38 | 10.45 | 9.82 | 6.74 | 21.13 | 7.80 | 7.35 | 10.13 | 8.23 | 10.13 | 13.94 | 7.57 |
| IER3IP1 | 1 | 8.89 | 6.61 | 8.63 | 8.33 | 9.65 | 7.90 | 10.34 | 18.44 | 8.64 | 12.56 | 7.75 | 8.48 | 8.77 | 14.39 | 10.60 |
| IFI16 | 24 | 5.81 | 6.03 | 6.40 | 6.16 | 11.81 | 12.13 | 21.16 | 22.93 | 3.95 | 3.62 | 5.92 | 6.28 | 11.97 | 22.04 | 3.78 |
| IFI30 | 1 | 10.56 | 10.25 | 12.13 | 6.76 | 10.74 | 9.32 | 16.24 | 8.99 | 7.28 | 7.73 | 10.40 | 9.44 | 10.03 | 12.61 | 7.51 |
| IFI35 | 2 | 7.54 | 10.49 | 17.19 | 10.37 | 12.87 | 10.97 | 6.45 | 5.02 | 7.96 | 11.14 | 9.02 | 13.78 | 11.92 | 5.73 | 9.55 |
| IFIT3 | 1 | 8.13 | 10.08 | 4.00 | 3.21 | 16.91 | 21.71 | 10.43 | 10.22 | 7.99 | 7.31 | 9.11 | 3.61 | 19.31 | 10.33 | 7.65 |
| IFIT5 | 3 | 10.92 | 11.77 | 8.99 | 7.01 | 12.00 | 10.29 | 10.10 | 8.68 | 10.47 | 9.78 | 11.35 | 8.00 | 11.14 | 9.39 | 10.12 |
| IFRD1 | 3 | 10.39 | 10.60 | 10.66 | 10.15 | 6.42 | 7.54 | 9.03 | 8.98 | 12.78 | 13.45 | 10.50 | 10.40 | 6.98 | 9.01 | 13.11 |
| IFRD2 | 1 | 7.92 | 9.13 | 6.95 | 8.47 | 10.06 | 11.61 | 11.19 | 12.23 | 11.14 | 11.32 | 8.52 | 7.71 | 10.83 | 11.71 | 11.23 |
| IFT122 | 2 | 11.10 | 11.99 | 9.51 | 9.41 | 8.63 | 8.23 | 7.82 | 7.88 | 12.85 | 12.56 | 11.55 | 9.46 | 8.43 | 7.85 | 12.71 |
| IFT172 | 2 | 11.25 | 10.99 | 10.58 | 10.21 | 10.69 | 12.04 | 10.78 | 8.58 | 8.39 | 6.49 | 11.12 | 10.40 | 11.36 | 9.68 | 7.44 |
| IFT20 | 1 | 9.78 | 11.86 | 11.63 | 10.54 | 7.90 | 9.22 | 11.38 | 9.23 | 9.51 | 8.95 | 10.82 | 11.08 | 8.56 | 10.31 | 9.23 |
| IFT27 | 2 | 6.43 | 6.16 | 4.71 | 3.62 | 6.84 | 4.95 | 9.25 | 45.57 | 6.82 | 5.63 | 6.30 | 4.17 | 5.89 | 27.41 | 6.23 |
| IFT57 | 1 | 11.03 | 11.57 | 11.05 | 10.53 | 9.93 | 8.49 | 11.44 | 10.07 | 8.94 | 6.97 | 11.30 | 10.79 | 9.21 | 10.75 | 7.95 |
| IFT74 | 3 | 11.23 | 13.03 | 7.68 | 5.50 | 12.61 | 10.59 | 10.50 | 9.43 | 9.66 | 9.77 | 12.13 | 6.59 | 11.60 | 9.97 | 9.71 |
| IFT81 | 5 | 10.74 | 11.50 | 9.37 | 9.29 | 10.72 | 10.85 | 8.66 | 7.85 | 10.52 | 10.52 | 11.12 | 9.33 | 10.78 | 8.26 | 10.52 |
| IGBP1 | 12 | 10.57 | 10.72 | 9.48 | 8.73 | 11.32 | 11.50 | 11.18 | 10.13 | 8.64 | 7.72 | 10.65 | 9.10 | 11.41 | 10.66 | 8.18 |

| | | | | | | | | | | | | | | | | |
|---|---|---|---|---|---|---|---|---|---|---|---|---|---|---|---|---|
| IGF1R | 5 | 10.17 | 8.11 | 9.30 | 10.72 | 10.55 | 10.73 | 9.70 | 9.14 | 10.88 | 10.69 | 9.14 | 10.01 | 10.64 | 9.42 | 10.79 |
| IGF2BP1 | 26 | 8.43 | 8.82 | 12.06 | 11.79 | 9.96 | 9.98 | 9.09 | 9.25 | 10.57 | 10.05 | 8.63 | 11.92 | 9.97 | 9.17 | 10.31 |
| IGF2BP2 | 7 | 9.10 | 9.97 | 14.58 | 14.12 | 7.99 | 7.87 | 8.22 | 7.65 | 10.66 | 9.83 | 9.54 | 14.35 | 7.93 | 7.93 | 10.24 |
| IGF2BP3 | 31 | 10.74 | 11.92 | 13.26 | 12.92 | 6.50 | 6.97 | 8.35 | 7.62 | 10.96 | 10.75 | 11.33 | 13.09 | 6.74 | 7.98 | 10.85 |
| IGF2R | 30 | 10.38 | 7.85 | 7.84 | 9.25 | 13.09 | 12.30 | 8.95 | 9.63 | 9.99 | 10.72 | 9.12 | 8.54 | 12.70 | 9.29 | 10.35 |
| IGFBP7 | 8 | 8.91 | 8.14 | 15.36 | 14.79 | 8.28 | 8.08 | 5.80 | 4.82 | 12.91 | 12.91 | 8.52 | 15.07 | 8.18 | 5.31 | 12.91 |
| IGHMBP2 | 8 | 9.15 | 10.21 | 10.49 | 10.54 | 9.35 | 9.87 | 10.57 | 10.38 | 9.75 | 9.68 | 9.68 | 10.52 | 9.61 | 10.48 | 9.72 |
| IGSF3 | 6 | 10.85 | 8.88 | 9.01 | 9.74 | 9.57 | 9.31 | 7.41 | 6.54 | 14.17 | 14.54 | 9.86 | 9.37 | 9.44 | 6.97 | 14.35 |
| IK | 26 | 10.30 | 10.68 | 10.79 | 10.17 | 8.51 | 8.89 | 8.95 | 8.76 | 11.80 | 11.17 | 10.49 | 10.48 | 8.70 | 8.85 | 11.48 |
| IKBIP | 17 | 10.05 | 6.53 | 8.39 | 10.97 | 8.64 | 8.24 | 10.00 | 9.60 | 13.01 | 14.58 | 8.29 | 9.68 | 8.44 | 9.80 | 13.80 |
| IKBKAP | 28 | 10.65 | 11.00 | 8.91 | 7.95 | 10.37 | 10.37 | 9.08 | 9.76 | 11.31 | 10.60 | 10.82 | 8.43 | 10.37 | 9.42 | 10.96 |
| IKBKB | 10 | 9.48 | 9.87 | 8.32 | 8.57 | 12.03 | 12.13 | 11.04 | 11.20 | 8.92 | 8.45 | 9.68 | 8.44 | 12.08 | 11.12 | 8.68 |
| IKBKG | 15 | 10.95 | 11.29 | 9.38 | 9.08 | 10.78 | 10.72 | 9.93 | 8.81 | 9.92 | 9.15 | 11.12 | 9.23 | 10.75 | 9.37 | 9.54 |
| IL18 | 16 | 9.48 | 9.65 | 14.85 | 12.24 | 12.73 | 12.06 | 2.47 | 1.99 | 12.46 | 12.07 | 9.56 | 13.54 | 12.39 | 2.23 | 12.27 |
| IL1RAP | 2 | 9.07 | 7.29 | 8.21 | 9.97 | 8.36 | 7.79 | 7.13 | 20.20 | 11.46 | 10.52 | 8.18 | 9.09 | 8.07 | 13.66 | 10.99 |
| IL4I1 | 1 | 8.44 | 11.35 | 11.66 | 12.40 | 10.13 | 10.00 | 7.52 | 6.25 | 11.12 | 11.14 | 9.89 | 12.03 | 10.06 | 6.88 | 11.13 |
| IL6ST | 2 | 10.28 | 9.30 | 7.66 | 10.46 | 10.59 | 9.68 | 9.54 | 9.59 | 10.47 | 12.43 | 9.79 | 9.06 | 10.14 | 9.56 | 11.45 |
| ILF2 | 23 | 9.92 | 10.07 | 10.25 | 10.16 | 10.13 | 10.48 | 10.26 | 10.33 | 9.28 | 9.13 | 10.00 | 10.20 | 10.31 | 10.29 | 9.20 |
| ILF3 | 84 | 8.48 | 9.76 | 11.88 | 10.79 | 9.72 | 10.02 | 10.21 | 10.38 | 9.43 | 9.33 | 9.12 | 11.34 | 9.87 | 10.29 | 9.38 |
| ILK | 36 | 9.06 | 9.87 | 13.78 | 13.23 | 7.72 | 7.73 | 7.13 | 8.52 | 11.88 | 11.08 | 9.47 | 13.51 | 7.72 | 7.82 | 11.48 |
| ILKAP | 13 | 9.21 | 10.25 | 9.80 | 9.71 | 9.15 | 9.93 | 11.04 | 10.62 | 10.17 | 10.12 | 9.73 | 9.76 | 9.54 | 10.83 | 10.14 |
| ILVBL | 12 | 8.88 | 7.46 | 10.29 | 11.74 | 11.21 | 10.46 | 9.87 | 10.84 | 9.29 | 9.95 | 8.17 | 11.02 | 10.84 | 10.36 | 9.62 |
| IMMT | 48 | 12.46 | 6.81 | 9.62 | 12.78 | 9.05 | 7.84 | 10.33 | 8.97 | 10.53 | 11.62 | 9.63 | 11.20 | 8.44 | 9.65 | 11.08 |
| IMP3 | 7 | 9.01 | 9.39 | 10.64 | 9.92 | 8.51 | 8.87 | 12.24 | 12.44 | 9.32 | 9.67 | 9.20 | 10.28 | 8.69 | 12.34 | 9.50 |
| IMP4 | 10 | 9.41 | 9.82 | 11.01 | 10.01 | 8.51 | 8.61 | 12.05 | 11.24 | 9.80 | 9.54 | 9.62 | 10.51 | 8.56 | 11.65 | 9.67 |
| IMPA1 | 7 | 10.57 | 10.76 | 10.86 | 9.04 | 8.27 | 7.94 | 11.91 | 9.42 | 10.55 | 10.69 | 10.67 | 9.95 | 8.10 | 10.66 | 10.62 |
| IMPA2 | 6 | 10.59 | 11.72 | 3.87 | 3.87 | 15.11 | 16.13 | 9.19 | 9.11 | 10.29 | 10.13 | 11.16 | 3.87 | 15.62 | 9.15 | 10.21 |
| IMPACT | 1 | 9.52 | 11.42 | 9.18 | 9.42 | 11.50 | 11.15 | 7.94 | 8.78 | 9.79 | 11.30 | 10.47 | 9.30 | 11.32 | 8.36 | 10.55 |
| IMPAD1 | 6 | 11.30 | 7.99 | 8.51 | 10.47 | 9.98 | 9.83 | 17.01 | 18.52 | 3.11 | 3.30 | 9.64 | 9.49 | 9.90 | 17.76 | 3.21 |
| IMPDH1 | 18 | 9.35 | 10.30 | 10.80 | 10.27 | 7.86 | 7.76 | 11.40 | 11.03 | 10.82 | 10.41 | 9.83 | 10.54 | 7.81 | 11.21 | 10.61 |
| IMPDH2 | 49 | 8.79 | 9.45 | 10.23 | 9.08 | 9.49 | 9.34 | 9.85 | 13.94 | 10.22 | 9.59 | 9.12 | 9.66 | 9.42 | 11.90 | 9.91 |
| IMUP | 3 | 11.74 | 12.17 | 8.42 | 9.50 | 10.83 | 13.31 | 9.73 | 2.12 | 10.10 | 12.09 | 11.95 | 8.96 | 12.07 | 5.93 | 11.09 |
| INA | 12 | 8.45 | 9.42 | 15.24 | 15.21 | 3.69 | 3.82 | 14.65 | 14.04 | 7.91 | 7.55 | 8.94 | 15.23 | 3.76 | 14.35 | 7.73 |
| INCENP | 10 | 9.09 | 10.41 | 9.68 | 9.71 | 8.30 | 8.78 | 11.91 | 10.89 | 10.51 | 10.72 | 9.75 | 9.70 | 8.54 | 11.40 | 10.61 |
| INF2 | 26 | 9.30 | 9.17 | 6.98 | 7.50 | 11.49 | 11.41 | 9.36 | 10.75 | 12.06 | 11.99 | 9.23 | 7.24 | 11.45 | 10.06 | 12.02 |
| ING1 | 2 | 8.47 | 10.03 | 9.20 | 10.16 | 9.90 | 11.32 | 12.39 | 11.98 | 8.43 | 8.13 | 9.25 | 9.68 | 10.61 | 12.18 | 8.28 |
| ING3 | 3 | 9.24 | 12.03 | 9.92 | 10.34 | 10.34 | 9.78 | 10.06 | 9.89 | 8.41 | 9.98 | 10.64 | 10.13 | 10.06 | 9.98 | 9.20 |
| ING4 | 2 | 9.08 | 11.42 | 10.54 | 10.66 | 9.49 | 11.39 | 10.07 | 9.72 | 8.83 | 8.82 | 10.25 | 10.60 | 10.44 | 9.89 | 8.82 |
| ING5 | 2 | 8.11 | 9.24 | 13.74 | 12.27 | 8.93 | 8.47 | 11.15 | 8.98 | 9.34 | 9.76 | 8.68 | 13.01 | 8.70 | 10.06 | 9.55 |
| INIP | 2 | 9.89 | 10.08 | 11.93 | 9.77 | 10.48 | 10.49 | 9.42 | 8.34 | 10.31 | 9.28 | 9.99 | 10.85 | 10.48 | 8.88 | 9.80 |
| INO80 | 2 | 9.35 | 8.24 | 9.27 | 11.13 | 9.38 | 8.93 | 10.65 | 11.25 | 10.28 | 11.51 | 8.80 | 10.20 | 9.15 | 10.95 | 10.90 |
| INO80B | 3 | 9.22 | 9.19 | 11.03 | 11.85 | 9.31 | 9.59 | 9.89 | 10.42 | 10.25 | 9.25 | 9.21 | 11.44 | 9.45 | 10.16 | 9.75 |

| | | | | | | | | | | | | | | | | |
|---|---|---|---|---|---|---|---|---|---|---|---|---|---|---|---|---|
| INO80C | 1 | 12.94 | 9.95 | 10.85 | 10.61 | 9.90 | 9.42 | 9.22 | 7.63 | 9.69 | 9.79 | 11.44 | 10.73 | 9.66 | 8.42 | 9.74 |
| INPP1 | 6 | 8.33 | 7.82 | 10.61 | 9.72 | 10.03 | 9.22 | 11.23 | 16.37 | 8.43 | 8.24 | 8.08 | 10.17 | 9.63 | 13.80 | 8.33 |
| INPP4A | 1 | 9.67 | 9.94 | 10.24 | 12.09 | 10.12 | 11.26 | 8.92 | 7.54 | 10.05 | 10.17 | 9.80 | 11.17 | 10.69 | 8.23 | 10.11 |
| INPP4B | 1 | 5.86 | 7.63 | 19.45 | 17.02 | 8.93 | 9.47 | 5.99 | 12.87 | 7.58 | 5.19 | 6.75 | 18.24 | 9.20 | 9.43 | 6.39 |
| INPP5B | 2 | 11.03 | 8.87 | 7.60 | 6.26 | 14.06 | 13.74 | 10.22 | 11.82 | 8.65 | 7.77 | 9.95 | 6.93 | 13.90 | 11.02 | 8.21 |
| INPP5D | 2 | 9.14 | 10.22 | 11.20 | 11.16 | 10.38 | 10.69 | 9.32 | 9.40 | 9.21 | 9.28 | 9.68 | 11.18 | 10.54 | 9.36 | 9.24 |
| INPPL1 | 30 | 9.58 | 10.59 | 10.12 | 10.49 | 11.92 | 12.79 | 8.96 | 9.01 | 8.44 | 8.10 | 10.08 | 10.30 | 12.35 | 8.99 | 8.27 |
| INTS1 | 9 | 10.37 | 9.68 | 9.12 | 8.23 | 10.09 | 10.10 | 11.54 | 12.79 | 9.09 | 8.99 | 10.03 | 8.67 | 10.09 | 12.16 | 9.04 |
| INTS10 | 4 | 9.31 | 10.24 | 9.29 | 9.53 | 9.14 | 8.98 | 12.20 | 12.31 | 9.75 | 9.25 | 9.78 | 9.41 | 9.06 | 12.26 | 9.50 |
| INTS12 | 5 | 9.41 | 10.96 | 10.30 | 9.75 | 10.01 | 10.91 | 10.47 | 9.98 | 9.12 | 9.08 | 10.18 | 10.02 | 10.46 | 10.23 | 9.10 |
| INTS2 | 2 | 8.85 | 9.73 | 8.75 | 10.99 | 11.06 | 11.28 | 10.33 | 10.04 | 9.49 | 9.49 | 9.29 | 9.87 | 11.17 | 10.18 | 9.49 |
| INTS3 | 15 | 9.68 | 9.71 | 9.57 | 9.03 | 11.79 | 11.33 | 9.22 | 9.17 | 10.22 | 10.29 | 9.69 | 9.30 | 11.56 | 9.20 | 10.25 |
| INTS4 | 3 | 9.99 | 9.14 | 9.25 | 9.10 | 10.53 | 10.77 | 11.87 | 12.00 | 8.72 | 8.63 | 9.57 | 9.17 | 10.65 | 11.94 | 8.67 |
| INTS5 | 5 | 9.55 | 9.14 | 10.22 | 9.87 | 9.85 | 9.99 | 12.08 | 10.77 | 9.14 | 9.40 | 9.35 | 10.04 | 9.92 | 11.42 | 9.27 |
| INTS6 | 9 | 9.14 | 9.97 | 10.43 | 9.76 | 10.52 | 10.27 | 11.29 | 10.80 | 8.98 | 8.83 | 9.56 | 10.09 | 10.40 | 11.05 | 8.90 |
| INTS7 | 4 | 9.30 | 10.51 | 8.91 | 9.66 | 11.85 | 12.30 | 9.25 | 9.09 | 9.71 | 9.42 | 9.90 | 9.28 | 12.07 | 9.17 | 9.57 |
| INTS9 | 5 | 10.31 | 9.94 | 11.02 | 10.69 | 9.37 | 9.81 | 10.93 | 10.60 | 8.72 | 8.61 | 10.12 | 10.86 | 9.59 | 10.76 | 8.66 |
| IP6K1 | 4 | 9.82 | 11.05 | 9.36 | 9.27 | 9.58 | 10.43 | 9.60 | 9.74 | 10.21 | 10.94 | 10.43 | 9.32 | 10.01 | 9.67 | 10.57 |
| IPO11 | 15 | 10.15 | 9.96 | 9.86 | 8.78 | 9.18 | 8.99 | 11.18 | 10.87 | 10.69 | 10.34 | 10.06 | 9.32 | 9.08 | 11.02 | 10.52 |
| IPO13 | 1 | 9.96 | 10.55 | 8.28 | 7.48 | 9.22 | 9.77 | 9.95 | 11.99 | 11.14 | 11.67 | 10.26 | 7.88 | 9.49 | 10.97 | 11.40 |
| IPO4 | 18 | 8.20 | 9.46 | 8.72 | 8.78 | 9.73 | 10.57 | 9.21 | 14.01 | 10.88 | 10.45 | 8.83 | 8.75 | 10.15 | 11.61 | 10.67 |
| IPO5 | 68 | 9.81 | 9.94 | 10.26 | 9.69 | 9.48 | 9.40 | 9.44 | 13.02 | 9.47 | 9.48 | 9.88 | 9.97 | 9.44 | 11.23 | 9.48 |
| IPO7 | 39 | 8.99 | 9.39 | 10.87 | 10.58 | 8.30 | 8.11 | 10.28 | 11.66 | 10.81 | 11.00 | 9.19 | 10.73 | 8.21 | 10.97 | 10.90 |
| IPO8 | 16 | 9.30 | 9.52 | 9.33 | 9.99 | 9.63 | 9.87 | 11.25 | 12.12 | 9.57 | 9.43 | 9.41 | 9.66 | 9.75 | 11.69 | 9.50 |
| IPO9 | 29 | 10.18 | 10.87 | 7.47 | 7.32 | 14.96 | 14.93 | 9.05 | 8.82 | 8.38 | 8.04 | 10.52 | 7.39 | 14.94 | 8.94 | 8.21 |
| IPPK | 2 | 9.69 | 10.75 | 12.20 | 10.61 | 8.89 | 8.66 | 7.78 | 8.27 | 12.26 | 10.90 | 10.22 | 11.41 | 8.77 | 8.03 | 11.58 |
| IQGAP1 | 156 | 8.63 | 9.27 | 10.00 | 9.78 | 11.84 | 11.84 | 7.60 | 7.43 | 11.96 | 11.66 | 8.95 | 9.89 | 11.84 | 7.52 | 11.81 |
| IQGAP2 | 49 | 11.84 | 13.05 | 6.41 | 5.40 | 7.80 | 7.61 | 10.43 | 10.30 | 13.79 | 13.38 | 12.44 | 5.90 | 7.70 | 10.37 | 13.59 |
| IQGAP3 | 6 | 9.91 | 10.18 | 11.13 | 12.42 | 9.57 | 9.03 | 7.69 | 7.33 | 11.63 | 11.10 | 10.05 | 11.78 | 9.30 | 7.51 | 11.37 |
| IQSEC1 | 2 | 10.45 | 13.32 | 6.78 | 5.72 | 10.83 | 14.74 | 9.78 | 8.07 | 10.55 | 9.75 | 11.89 | 6.25 | 12.78 | 8.92 | 10.15 |
| IRAK1 | 2 | 9.25 | 10.76 | 7.12 | 8.48 | 10.71 | 11.55 | 11.97 | 12.46 | 8.76 | 8.94 | 10.01 | 7.80 | 11.13 | 12.22 | 8.85 |
| IRAK4 | 4 | 8.03 | 10.37 | 9.60 | 10.21 | 9.85 | 10.77 | 11.13 | 10.97 | 9.68 | 9.40 | 9.20 | 9.90 | 10.31 | 11.05 | 9.54 |
| IREB2 | 10 | 8.05 | 8.69 | 13.73 | 12.93 | 10.07 | 9.95 | 9.08 | 7.71 | 10.47 | 9.33 | 8.37 | 13.33 | 10.01 | 8.40 | 9.90 |
| IRF2 | 1 | 11.77 | 9.89 | 10.60 | 9.34 | 10.79 | 12.02 | 9.53 | 9.02 | 9.87 | 7.17 | 10.83 | 9.97 | 11.41 | 9.28 | 8.52 |
| IRF2BP1 | 5 | 8.01 | 9.61 | 8.40 | 7.81 | 8.86 | 8.91 | 12.21 | 20.22 | 7.89 | 8.09 | 8.81 | 8.11 | 8.89 | 16.21 | 7.99 |
| IRF2BP2 | 10 | 8.64 | 9.83 | 9.54 | 8.78 | 9.60 | 9.78 | 14.10 | 12.46 | 9.25 | 8.01 | 9.24 | 9.16 | 9.69 | 13.28 | 8.63 |
| IRF2BPL | 10 | 8.50 | 9.32 | 12.91 | 12.72 | 8.23 | 8.51 | 11.48 | 11.12 | 8.86 | 8.34 | 8.91 | 12.82 | 8.37 | 11.30 | 8.60 |
| IRF3 | 2 | 9.38 | 10.78 | 7.68 | 8.04 | 10.25 | 12.23 | 11.96 | 12.05 | 9.10 | 8.54 | 10.08 | 7.86 | 11.24 | 12.01 | 8.82 |
| IRGQ | 11 | 9.37 | 10.43 | 11.90 | 11.80 | 9.48 | 9.33 | 9.99 | 8.64 | 9.74 | 9.32 | 9.90 | 11.85 | 9.40 | 9.32 | 9.53 |
| IRS1 | 1 | 7.58 | 8.90 | 12.15 | 10.99 | 7.79 | 6.88 | 11.76 | 13.78 | 9.00 | 11.17 | 8.24 | 11.57 | 7.34 | 12.77 | 10.08 |
| IRS2 | 3 | 7.83 | 8.63 | 11.40 | 10.52 | 12.31 | 11.57 | 13.87 | 12.05 | 6.83 | 4.98 | 8.23 | 10.96 | 11.94 | 12.96 | 5.90 |
| ISCA1 | 3 | 10.70 | 9.14 | 8.84 | 10.12 | 7.94 | 8.45 | 11.19 | 10.13 | 12.03 | 11.46 | 9.92 | 9.48 | 8.20 | 10.66 | 11.74 |

| | | | | | | | | | | | | | | | | |
|---|---|---|---|---|---|---|---|---|---|---|---|---|---|---|---|---|
| ISCA2 | 7 | 9.52 | 9.27 | 11.04 | 11.62 | 10.77 | 10.20 | 9.57 | 8.17 | 10.00 | 9.85 | 9.39 | 11.33 | 10.49 | 8.87 | 9.93 |
| ISCU | 3 | 10.17 | 8.21 | 11.30 | 10.92 | 10.81 | 9.75 | 10.02 | 8.89 | 10.63 | 9.30 | 9.19 | 11.11 | 10.28 | 9.46 | 9.96 |
| ISG15 | 17 | 9.01 | 9.11 | 8.33 | 8.15 | 10.18 | 10.21 | 10.61 | 10.83 | 11.90 | 11.67 | 9.06 | 8.24 | 10.20 | 10.72 | 11.79 |
| ISG20L2 | 3 | 8.18 | 9.43 | 10.52 | 11.13 | 9.39 | 10.13 | 9.66 | 9.57 | 10.38 | 11.62 | 8.80 | 10.82 | 9.76 | 9.61 | 11.00 |
| ISOC1 | 8 | 9.11 | 10.05 | 7.94 | 7.21 | 12.17 | 12.95 | 7.50 | 7.37 | 12.90 | 12.81 | 9.58 | 7.58 | 12.56 | 7.43 | 12.86 |
| IST1 | 14 | 10.95 | 11.03 | 7.89 | 7.25 | 11.94 | 11.31 | 8.95 | 11.48 | 9.96 | 9.24 | 10.99 | 7.57 | 11.62 | 10.22 | 9.60 |
| ISY1 | 13 | 9.42 | 10.52 | 11.99 | 12.12 | 8.91 | 9.40 | 9.22 | 8.21 | 10.41 | 9.79 | 9.97 | 12.05 | 9.16 | 8.72 | 10.10 |
| ITCH | 10 | 10.76 | 9.59 | 11.35 | 9.58 | 9.63 | 9.56 | 8.95 | 8.07 | 11.69 | 10.82 | 10.17 | 10.46 | 9.59 | 8.51 | 11.26 |
| ITFG1 | 1 | 14.27 | 9.10 | 4.99 | 6.83 | 12.91 | 11.31 | 10.03 | 10.71 | 9.53 | 10.31 | 11.69 | 5.91 | 12.11 | 10.37 | 9.92 |
| ITFG3 | 3 | 12.92 | 9.86 | 6.65 | 7.89 | 9.69 | 9.57 | 8.39 | 7.79 | 13.69 | 13.56 | 11.39 | 7.27 | 9.63 | 8.09 | 13.62 |
| ITGA1 | 14 | 12.18 | 9.68 | 12.08 | 13.56 | 7.24 | 7.00 | 7.82 | 6.53 | 12.01 | 11.91 | 10.93 | 12.82 | 7.12 | 7.17 | 11.96 |
| ITGA11 | 26 | 11.57 | 8.71 | 10.08 | 11.95 | 5.08 | 4.98 | 2.66 | 2.58 | 20.32 | 22.07 | 10.14 | 11.01 | 5.03 | 2.62 | 21.19 |
| ITGA2 | 12 | 5.02 | 4.13 | 6.21 | 6.63 | 6.49 | 6.45 | 24.06 | 30.29 | 5.23 | 5.49 | 4.58 | 6.42 | 6.47 | 27.17 | 5.36 |
| ITGA3 | 17 | 10.67 | 8.46 | 9.20 | 11.54 | 8.96 | 9.59 | 6.95 | 6.24 | 13.73 | 14.65 | 9.56 | 10.37 | 9.27 | 6.60 | 14.19 |
| ITGA5 | 14 | 8.24 | 7.18 | 11.05 | 12.51 | 5.71 | 5.79 | 7.95 | 7.06 | 16.86 | 17.65 | 7.71 | 11.78 | 5.75 | 7.50 | 17.26 |
| ITGA6 | 29 | 9.98 | 7.76 | 7.19 | 8.69 | 12.07 | 12.05 | 11.80 | 10.88 | 9.67 | 9.90 | 8.87 | 7.94 | 12.06 | 11.34 | 9.79 |
| ITGAV | 25 | 9.33 | 6.79 | 13.20 | 15.76 | 8.83 | 8.44 | 9.09 | 8.34 | 9.61 | 10.62 | 8.06 | 14.48 | 8.63 | 8.71 | 10.12 |
| ITGB1 | 42 | 8.50 | 6.71 | 10.07 | 12.03 | 8.47 | 7.71 | 8.98 | 11.32 | 12.77 | 13.44 | 7.60 | 11.05 | 8.09 | 10.15 | 13.10 |
| ITGB1BP1 | 1 | 9.54 | 12.57 | 10.27 | 9.76 | 8.85 | 9.35 | 10.74 | 9.77 | 9.36 | 9.78 | 11.05 | 10.01 | 9.10 | 10.26 | 9.57 |
| ITGB4 | 6 | 10.02 | 9.03 | 7.21 | 7.62 | 12.35 | 13.07 | 6.77 | 7.27 | 13.65 | 13.01 | 9.53 | 7.42 | 12.71 | 7.02 | 13.33 |
| ITIH2 | 1 | 5.94 | 5.59 | 16.11 | 17.83 | 4.88 | 4.81 | 16.44 | 15.02 | 7.43 | 5.95 | 5.77 | 16.97 | 4.84 | 15.73 | 6.69 |
| ITPA | 7 | 9.04 | 9.69 | 11.52 | 10.39 | 10.88 | 10.18 | 8.85 | 10.52 | 9.75 | 9.19 | 9.37 | 10.95 | 10.53 | 9.68 | 9.47 |
| ITPK1 | 3 | 7.18 | 8.68 | 16.07 | 13.40 | 8.06 | 6.87 | 10.24 | 7.83 | 10.70 | 10.96 | 7.93 | 14.74 | 7.46 | 9.04 | 10.83 |
| ITPR1 | 23 | 12.89 | 6.99 | 5.42 | 8.48 | 16.10 | 14.30 | 8.19 | 8.57 | 8.47 | 10.59 | 9.94 | 6.95 | 15.20 | 8.38 | 9.53 |
| ITPR3 | 24 | 11.48 | 6.05 | 5.24 | 7.58 | 12.43 | 10.66 | 9.63 | 9.21 | 12.05 | 15.68 | 8.77 | 6.41 | 11.54 | 9.42 | 13.86 |
| ITPRIP | 2 | 11.50 | 7.63 | 9.03 | 12.99 | 8.35 | 8.00 | 9.09 | 8.88 | 11.19 | 13.33 | 9.57 | 11.01 | 8.18 | 8.98 | 12.26 |
| ITSN1 | 18 | 12.61 | 13.12 | 8.41 | 8.65 | 11.12 | 11.86 | 6.97 | 7.19 | 10.48 | 9.58 | 12.87 | 8.53 | 11.49 | 7.08 | 10.03 |
| ITSN2 | 4 | 9.91 | 10.09 | 10.60 | 9.98 | 10.64 | 9.99 | 8.68 | 10.71 | 10.08 | 9.31 | 10.00 | 10.29 | 10.32 | 9.70 | 9.70 |
| IVD | 11 | 9.81 | 8.77 | 11.79 | 12.73 | 10.09 | 9.41 | 7.15 | 6.42 | 11.75 | 12.07 | 9.29 | 12.26 | 9.75 | 6.78 | 11.91 |
| IVNS1ABP | 7 | 7.83 | 9.65 | 11.69 | 10.67 | 7.97 | 7.96 | 10.77 | 13.10 | 10.09 | 10.28 | 8.74 | 11.18 | 7.96 | 11.94 | 10.19 |
| IWS1 | 14 | 9.62 | 11.18 | 11.58 | 10.35 | 9.50 | 9.09 | 9.27 | 8.38 | 10.54 | 10.48 | 10.40 | 10.96 | 9.30 | 8.83 | 10.51 |
| JADE2 | 2 | 9.52 | 10.34 | 7.83 | 8.60 | 11.24 | 13.62 | 11.40 | 9.69 | 10.05 | 7.71 | 9.93 | 8.22 | 12.43 | 10.54 | 8.88 |
| JADE3 | 1 | 14.71 | 10.72 | 7.40 | 6.09 | 10.49 | 10.95 | 10.61 | 12.51 | 10.10 | 6.42 | 12.71 | 6.75 | 10.72 | 11.56 | 8.26 |
| JAGN1 | 1 | 13.05 | 5.86 | 5.90 | 8.72 | 7.39 | 7.34 | 11.23 | 10.23 | 15.59 | 14.70 | 9.45 | 7.31 | 7.36 | 10.73 | 15.15 |
| JAK1 | 2 | 13.18 | 11.92 | 8.92 | 8.85 | 8.69 | 9.05 | 8.59 | 8.37 | 11.81 | 10.62 | 12.55 | 8.88 | 8.87 | 8.48 | 11.21 |
| JMJD1C | 4 | 9.49 | 9.93 | 11.01 | 10.05 | 9.41 | 9.69 | 11.43 | 10.86 | 9.47 | 8.66 | 9.71 | 10.53 | 9.55 | 11.15 | 9.06 |
| JMJD6 | 7 | 11.20 | 11.48 | 10.55 | 11.54 | 8.10 | 9.20 | 8.45 | 8.16 | 10.60 | 10.72 | 11.34 | 11.04 | 8.65 | 8.30 | 10.66 |
| JPH2 | 1 | 13.39 | 9.56 | 13.92 | 15.72 | 5.51 | 4.41 | 3.50 | 2.71 | 14.09 | 17.19 | 11.48 | 14.82 | 4.96 | 3.10 | 15.64 |
| JUN | 1 | 10.17 | 11.36 | 9.82 | 8.88 | 5.47 | 5.89 | 13.16 | 10.25 | 13.25 | 11.77 | 10.76 | 9.35 | 5.68 | 11.70 | 12.51 |
| JUNB | 9 | 8.71 | 8.65 | 10.31 | 8.98 | 9.93 | 10.12 | 14.17 | 12.31 | 8.85 | 7.97 | 8.68 | 9.65 | 10.02 | 13.24 | 8.41 |
| JUND | 5 | 9.64 | 10.40 | 7.76 | 7.21 | 10.65 | 11.07 | 11.89 | 11.76 | 9.58 | 10.04 | 10.02 | 7.48 | 10.86 | 11.82 | 9.81 |
| JUP | 44 | 8.70 | 9.14 | 13.25 | 13.70 | 9.72 | 10.01 | 6.67 | 6.36 | 11.32 | 11.12 | 8.92 | 13.47 | 9.87 | 6.52 | 11.22 |

| | | | | | | | | | | | | | | | | |
|---|---|---|---|---|---|---|---|---|---|---|---|---|---|---|---|---|
| KALRN | 1 | 11.08 | 11.11 | 10.12 | 11.21 | 8.99 | 10.95 | 10.04 | 6.70 | 9.33 | 10.47 | 11.09 | 10.67 | 9.97 | 8.37 | 9.90 |
| KANK1 | 8 | 11.73 | 12.16 | 11.99 | 11.10 | 9.11 | 9.39 | 6.06 | 5.03 | 12.18 | 11.25 | 11.94 | 11.54 | 9.25 | 5.55 | 11.71 |
| KANK2 | 10 | 11.41 | 12.61 | 11.31 | 11.15 | 8.72 | 8.91 | 5.99 | 5.17 | 12.53 | 12.20 | 12.01 | 11.23 | 8.81 | 5.58 | 12.37 |
| KANSL3 | 1 | 8.45 | 11.29 | 10.93 | 10.81 | 10.68 | 10.81 | 9.49 | 8.21 | 8.28 | 11.05 | 9.87 | 10.87 | 10.75 | 8.85 | 9.66 |
| KARS | 49 | 10.10 | 10.72 | 9.02 | 7.26 | 11.72 | 11.28 | 9.91 | 9.50 | 10.30 | 10.20 | 10.41 | 8.14 | 11.50 | 9.71 | 10.25 |
| KAT2A | 2 | 9.81 | 9.47 | 9.41 | 9.62 | 9.31 | 10.65 | 11.38 | 8.85 | 12.00 | 9.50 | 9.64 | 9.52 | 9.98 | 10.11 | 10.75 |
| KAT5 | 1 | 8.63 | 9.74 | 8.31 | 7.08 | 9.04 | 11.68 | 12.00 | 16.79 | 7.95 | 8.78 | 9.19 | 7.70 | 10.36 | 14.39 | 8.36 |
| KAT7 | 5 | 10.01 | 12.68 | 10.19 | 10.26 | 9.10 | 9.98 | 9.31 | 9.20 | 9.78 | 9.49 | 11.34 | 10.23 | 9.54 | 9.25 | 9.64 |
| KAT8 | 2 | 9.51 | 9.51 | 10.31 | 9.28 | 10.42 | 9.20 | 10.49 | 10.70 | 10.82 | 9.74 | 9.51 | 9.80 | 9.81 | 10.59 | 10.28 |
| KATNA1 | 3 | 9.66 | 10.48 | 10.50 | 9.32 | 9.81 | 10.00 | 10.39 | 8.73 | 11.09 | 10.02 | 10.07 | 9.91 | 9.90 | 9.56 | 10.55 |
| KATNAL1 | 1 | 11.25 | 12.83 | 10.14 | 10.41 | 10.47 | 11.25 | 8.13 | 7.63 | 9.19 | 8.70 | 12.04 | 10.27 | 10.86 | 7.88 | 8.95 |
| KATNB1 | 4 | 9.56 | 8.36 | 8.33 | 7.66 | 9.96 | 9.66 | 8.99 | 19.91 | 8.78 | 8.78 | 8.96 | 7.99 | 9.81 | 14.45 | 8.78 |
| KBTBD2 | 1 | 8.17 | 9.62 | 10.82 | 11.33 | 9.05 | 9.79 | 10.77 | 10.11 | 9.84 | 10.50 | 8.89 | 11.08 | 9.42 | 10.44 | 10.17 |
| KBTBD4 | 1 | 9.76 | 10.11 | 11.54 | 10.02 | 10.28 | 10.02 | 8.57 | 8.66 | 10.60 | 10.44 | 9.93 | 10.78 | 10.15 | 8.62 | 10.52 |
| KBTBD7 | 1 | 11.27 | 11.72 | 10.34 | 7.29 | 14.07 | 11.92 | 8.77 | 8.49 | 7.91 | 8.21 | 11.50 | 8.82 | 13.00 | 8.63 | 8.06 |
| KCMF1 | 1 | 9.89 | 10.15 | 10.41 | 10.68 | 9.46 | 10.72 | 9.65 | 10.19 | 10.03 | 8.81 | 10.02 | 10.55 | 10.09 | 9.92 | 9.42 |
| KCNAB2 | 5 | 11.02 | 12.04 | 4.47 | 4.31 | 10.51 | 11.22 | 12.08 | 11.77 | 11.40 | 11.19 | 11.53 | 4.39 | 10.86 | 11.93 | 11.30 |
| KCNH4 | 1 | 11.53 | 10.65 | 8.65 | 10.25 | 11.05 | 11.17 | 8.03 | 3.33 | 13.56 | 11.79 | 11.09 | 9.45 | 11.11 | 5.68 | 12.68 |
| KCNN4 | 1 | 10.76 | 8.87 | 7.91 | 4.10 | 11.31 | 8.65 | 13.53 | 10.71 | 10.52 | 13.63 | 9.82 | 6.00 | 9.98 | 12.12 | 12.08 |
| KCTD10 | 4 | 10.54 | 10.96 | 11.14 | 10.96 | 8.95 | 9.20 | 8.04 | 7.46 | 11.41 | 11.34 | 10.75 | 11.05 | 9.07 | 7.75 | 11.37 |
| KCTD12 | 16 | 11.36 | 11.58 | 18.57 | 16.41 | 5.32 | 5.47 | 6.15 | 5.50 | 9.80 | 9.84 | 11.47 | 17.49 | 5.40 | 5.82 | 9.82 |
| KCTD15 | 4 | 14.06 | 14.76 | 8.86 | 8.78 | 11.26 | 11.68 | 6.05 | 5.40 | 9.53 | 9.63 | 14.41 | 8.82 | 11.47 | 5.72 | 9.58 |
| KCTD3 | 3 | 10.70 | 10.59 | 8.49 | 7.80 | 13.01 | 12.31 | 9.39 | 8.29 | 9.39 | 10.03 | 10.65 | 8.15 | 12.66 | 8.84 | 9.71 |
| KCTD5 | 2 | 11.67 | 12.20 | 11.55 | 11.78 | 7.86 | 8.09 | 8.71 | 8.35 | 10.28 | 9.51 | 11.94 | 11.67 | 7.97 | 8.53 | 9.90 |
| KCTD7 | 1 | 10.02 | 10.48 | 11.44 | 10.43 | 8.45 | 8.05 | 6.30 | 7.83 | 13.82 | 13.18 | 10.25 | 10.93 | 8.25 | 7.07 | 13.50 |
| KCTD9 | 2 | 12.20 | 12.07 | 7.59 | 8.35 | 10.27 | 10.75 | 9.56 | 8.72 | 10.05 | 10.43 | 12.14 | 7.97 | 10.51 | 9.14 | 10.24 |
| KDELC1 | 9 | 9.40 | 8.64 | 7.32 | 7.89 | 12.11 | 11.56 | 14.70 | 13.40 | 7.43 | 7.56 | 9.02 | 7.60 | 11.83 | 14.05 | 7.50 |
| KDELC2 | 5 | 10.40 | 9.02 | 9.75 | 10.65 | 10.10 | 9.22 | 9.40 | 8.49 | 11.39 | 11.58 | 9.71 | 10.20 | 9.66 | 8.95 | 11.48 |
| KDM1A | 23 | 9.56 | 9.72 | 8.76 | 8.34 | 10.71 | 10.98 | 11.39 | 11.30 | 9.65 | 9.58 | 9.64 | 8.55 | 10.85 | 11.35 | 9.61 |
| KDM1B | 6 | 9.54 | 10.39 | 10.66 | 10.83 | 9.08 | 9.13 | 10.65 | 9.38 | 9.91 | 10.44 | 9.96 | 10.74 | 9.10 | 10.01 | 10.17 |
| KDM2A | 16 | 7.89 | 10.10 | 10.47 | 10.53 | 9.50 | 9.26 | 11.19 | 12.01 | 9.66 | 9.38 | 9.00 | 10.50 | 9.38 | 11.60 | 9.52 |
| KDM3A | 2 | 11.04 | 11.26 | 8.58 | 9.47 | 10.70 | 10.76 | 10.80 | 10.79 | 7.75 | 8.86 | 11.15 | 9.03 | 10.73 | 10.79 | 8.31 |
| KDM3B | 17 | 9.70 | 10.59 | 10.77 | 9.98 | 8.84 | 9.27 | 11.63 | 10.66 | 9.45 | 9.12 | 10.14 | 10.37 | 9.06 | 11.14 | 9.28 |
| KDM4A | 4 | 9.90 | 9.78 | 10.55 | 10.07 | 9.07 | 9.24 | 11.66 | 10.21 | 10.29 | 9.22 | 9.84 | 10.31 | 9.16 | 10.94 | 9.76 |
| KDM5A | 5 | 10.12 | 10.41 | 9.96 | 9.58 | 9.73 | 10.16 | 10.24 | 9.47 | 10.16 | 10.18 | 10.26 | 9.77 | 9.94 | 9.86 | 10.17 |
| KDM5C | 8 | 8.58 | 9.00 | 9.06 | 8.31 | 11.74 | 11.93 | 14.11 | 12.69 | 7.21 | 7.38 | 8.79 | 8.69 | 11.84 | 13.40 | 7.29 |
| KDSR | 1 | 9.99 | 6.60 | 7.62 | 9.91 | 14.65 | 9.64 | 9.78 | 9.93 | 9.58 | 12.29 | 8.29 | 8.77 | 12.15 | 9.86 | 10.93 |
| KHDRBS1 | 21 | 10.40 | 11.03 | 11.15 | 10.03 | 9.03 | 9.38 | 10.66 | 9.36 | 9.64 | 9.30 | 10.72 | 10.59 | 9.21 | 10.01 | 9.47 |
| KHDRBS3 | 5 | 11.50 | 11.81 | 7.91 | 8.17 | 13.05 | 13.88 | 10.12 | 11.25 | 6.19 | 6.12 | 11.65 | 8.04 | 13.46 | 10.69 | 6.16 |
| KHNYN | 1 | 10.32 | 11.65 | 7.17 | 7.61 | 10.50 | 11.39 | 10.94 | 9.53 | 11.84 | 9.04 | 10.98 | 7.39 | 10.95 | 10.24 | 10.44 |
| KHSRP | 47 | 9.21 | 9.73 | 11.25 | 10.18 | 9.94 | 9.98 | 10.00 | 10.64 | 9.83 | 9.24 | 9.47 | 10.72 | 9.96 | 10.32 | 9.54 |
| KIAA0020 | 22 | 8.88 | 9.22 | 10.76 | 10.96 | 7.22 | 7.51 | 11.02 | 10.48 | 11.54 | 12.40 | 9.05 | 10.86 | 7.36 | 10.75 | 11.97 |

| | | | | | | | | | | | | | | | | |
|---|---|---|---|---|---|---|---|---|---|---|---|---|---|---|---|---|
| KIAA0101 | 3 | 10.58 | 10.48 | 8.04 | 7.89 | 11.64 | 10.65 | 9.78 | 8.18 | 11.65 | 11.10 | 10.53 | 7.97 | 11.15 | 8.98 | 11.38 |
| KIAA0196 | 14 | 10.14 | 10.29 | 8.98 | 10.17 | 10.37 | 10.68 | 9.44 | 8.85 | 10.58 | 10.52 | 10.21 | 9.57 | 10.52 | 9.15 | 10.55 |
| KIAA0319L | 2 | 11.15 | 9.50 | 7.08 | 9.49 | 9.38 | 11.82 | 12.25 | 11.24 | 7.77 | 10.31 | 10.33 | 8.29 | 10.60 | 11.74 | 9.04 |
| KIAA0391 | 3 | 10.31 | 8.96 | 12.06 | 14.74 | 8.73 | 8.60 | 8.68 | 8.75 | 9.22 | 9.93 | 9.64 | 13.40 | 8.67 | 8.72 | 9.58 |
| KIAA0754 | 4 | 9.49 | 9.28 | 10.10 | 11.51 | 9.02 | 9.85 | 10.94 | 9.89 | 10.12 | 9.80 | 9.39 | 10.81 | 9.44 | 10.41 | 9.96 |
| KIAA0907 | 4 | 8.91 | 14.01 | 9.58 | 9.90 | 8.57 | 8.63 | 11.26 | 10.30 | 9.11 | 9.75 | 11.46 | 9.74 | 8.60 | 10.78 | 9.43 |
| KIAA0930 | 1 | 9.69 | 11.94 | 12.35 | 10.11 | 6.75 | 6.20 | 11.42 | 11.14 | 9.98 | 10.41 | 10.82 | 11.23 | 6.48 | 11.28 | 10.20 |
| KIAA0947 | 8 | 7.98 | 9.01 | 7.76 | 7.33 | 11.38 | 11.05 | 15.26 | 14.00 | 8.23 | 8.00 | 8.50 | 7.55 | 11.22 | 14.63 | 8.11 |
| KIAA1033 | 13 | 10.07 | 9.74 | 9.13 | 9.98 | 10.74 | 11.17 | 8.82 | 8.62 | 11.09 | 10.64 | 9.90 | 9.56 | 10.96 | 8.72 | 10.86 |
| KIAA1143 | 9 | 10.45 | 11.38 | 9.77 | 9.68 | 9.74 | 10.14 | 10.85 | 8.01 | 10.00 | 9.98 | 10.92 | 9.72 | 9.94 | 9.43 | 9.99 |
| KIAA1161 | 1 | 12.09 | 8.65 | 7.74 | 11.02 | 9.87 | 9.34 | 4.89 | 3.60 | 14.70 | 18.09 | 10.37 | 9.38 | 9.60 | 4.25 | 16.40 |
| KIAA1191 | 2 | 11.18 | 9.73 | 7.56 | 8.35 | 8.56 | 8.93 | 12.86 | 12.88 | 10.51 | 9.44 | 10.45 | 7.96 | 8.75 | 12.87 | 9.97 |
| KIAA1211 | 3 | 12.92 | 14.07 | 9.27 | 8.01 | 12.85 | 12.92 | 9.60 | 8.66 | 5.87 | 5.83 | 13.50 | 8.64 | 12.88 | 9.13 | 5.85 |
| KIAA1217 | 16 | 9.33 | 11.98 | 12.03 | 11.84 | 10.56 | 10.71 | 7.50 | 5.79 | 10.47 | 9.81 | 10.65 | 11.93 | 10.64 | 6.64 | 10.14 |
| KIAA1279 | 14 | 10.12 | 10.54 | 9.13 | 9.08 | 11.49 | 11.35 | 9.47 | 9.06 | 9.74 | 10.01 | 10.33 | 9.11 | 11.42 | 9.27 | 9.88 |
| KIAA1429 | 14 | 9.80 | 10.19 | 10.31 | 9.87 | 8.91 | 8.96 | 10.85 | 11.00 | 10.20 | 9.91 | 10.00 | 10.09 | 8.94 | 10.92 | 10.05 |
| KIAA1468 | 8 | 10.78 | 11.37 | 10.01 | 10.87 | 10.17 | 11.23 | 6.95 | 7.06 | 11.00 | 10.57 | 11.08 | 10.44 | 10.70 | 7.01 | 10.78 |
| KIAA1522 | 14 | 9.71 | 11.24 | 10.33 | 10.00 | 9.15 | 9.30 | 8.16 | 7.39 | 12.30 | 12.43 | 10.47 | 10.16 | 9.23 | 7.77 | 12.36 |
| KIAA1524 | 12 | 10.34 | 11.20 | 8.35 | 8.65 | 9.21 | 9.34 | 10.37 | 9.81 | 11.43 | 11.30 | 10.77 | 8.50 | 9.27 | 10.09 | 11.36 |
| KIAA1598 | 25 | 10.20 | 11.33 | 7.70 | 7.29 | 16.00 | 15.54 | 10.12 | 8.88 | 6.81 | 6.13 | 10.77 | 7.49 | 15.77 | 9.50 | 6.47 |
| KIAA1671 | 9 | 10.26 | 11.75 | 10.71 | 10.61 | 7.48 | 8.23 | 7.18 | 6.54 | 13.72 | 13.53 | 11.00 | 10.66 | 7.86 | 6.86 | 13.62 |
| KIAA2013 | 2 | 12.44 | 7.11 | 7.96 | 10.17 | 7.12 | 6.51 | 12.03 | 10.98 | 11.59 | 14.10 | 9.77 | 9.07 | 6.81 | 11.50 | 12.84 |
| KIAA2018 | 2 | 13.27 | 11.10 | 9.02 | 9.40 | 8.29 | 10.35 | 8.72 | 8.73 | 11.23 | 9.90 | 12.18 | 9.21 | 9.32 | 8.73 | 10.56 |
| KIDINS220 | 4 | 10.36 | 8.07 | 6.97 | 9.32 | 9.48 | 9.13 | 8.80 | 8.36 | 13.95 | 15.56 | 9.21 | 8.15 | 9.30 | 8.58 | 14.76 |
| KIF11 | 32 | 9.27 | 9.70 | 10.27 | 10.17 | 9.70 | 9.97 | 10.33 | 9.59 | 10.57 | 10.43 | 9.49 | 10.22 | 9.84 | 9.96 | 10.50 |
| KIF13A | 13 | 10.85 | 11.44 | 9.01 | 8.49 | 11.54 | 11.73 | 9.44 | 8.82 | 9.23 | 9.44 | 11.15 | 8.75 | 11.64 | 9.13 | 9.34 |
| KIF13B | 4 | 10.31 | 10.66 | 8.92 | 11.03 | 8.63 | 10.26 | 8.96 | 10.34 | 10.57 | 10.32 | 10.48 | 9.98 | 9.45 | 9.65 | 10.44 |
| KIF14 | 31 | 9.10 | 9.79 | 9.43 | 9.07 | 11.01 | 11.04 | 9.97 | 9.37 | 10.73 | 10.48 | 9.45 | 9.25 | 11.03 | 9.67 | 10.61 |
| KIF15 | 24 | 9.66 | 10.61 | 9.10 | 8.83 | 10.33 | 10.35 | 11.28 | 10.46 | 9.77 | 9.62 | 10.14 | 8.96 | 10.34 | 10.87 | 9.70 |
| KIF18A | 2 | 6.97 | 8.92 | 12.10 | 14.65 | 8.82 | 9.04 | 11.35 | 10.24 | 8.56 | 9.35 | 7.95 | 13.38 | 8.93 | 10.79 | 8.95 |
| KIF18B | 2 | 8.99 | 11.30 | 11.00 | 10.91 | 8.62 | 8.18 | 9.19 | 9.59 | 10.82 | 11.41 | 10.14 | 10.95 | 8.40 | 9.39 | 11.11 |
| KIF1A | 1 | 15.12 | 13.23 | 6.71 | 7.26 | 9.23 | 12.38 | 10.05 | 8.84 | 8.92 | 8.26 | 14.18 | 6.99 | 10.81 | 9.44 | 8.59 |
| KIF1B | 19 | 10.92 | 11.98 | 10.66 | 10.74 | 8.85 | 9.14 | 9.56 | 9.41 | 9.47 | 9.27 | 11.45 | 10.70 | 9.00 | 9.48 | 9.37 |
| KIF1C | 7 | 10.47 | 11.49 | 12.66 | 11.30 | 8.49 | 9.26 | 8.09 | 7.28 | 10.96 | 10.02 | 10.98 | 11.98 | 8.88 | 7.68 | 10.49 |
| KIF20A | 29 | 8.39 | 9.25 | 9.17 | 9.09 | 10.19 | 10.47 | 11.36 | 10.58 | 10.93 | 10.57 | 8.82 | 9.13 | 10.33 | 10.97 | 10.75 |
| KIF21A | 18 | 11.11 | 11.83 | 9.33 | 8.64 | 11.53 | 11.40 | 9.13 | 8.40 | 9.40 | 9.22 | 11.47 | 8.99 | 11.47 | 8.77 | 9.31 |
| KIF22 | 15 | 9.01 | 9.51 | 10.61 | 10.29 | 9.06 | 9.04 | 11.51 | 10.43 | 10.19 | 10.35 | 9.26 | 10.45 | 9.05 | 10.97 | 10.27 |
| KIF23 | 20 | 9.13 | 9.59 | 10.40 | 10.81 | 9.70 | 10.00 | 9.54 | 8.66 | 11.35 | 10.81 | 9.36 | 10.61 | 9.85 | 9.10 | 11.08 |
| KIF2A | 21 | 9.95 | 10.17 | 10.36 | 9.79 | 8.73 | 8.86 | 12.13 | 11.17 | 9.70 | 9.14 | 10.06 | 10.07 | 8.80 | 11.65 | 9.42 |
| KIF2C | 18 | 9.79 | 10.10 | 8.81 | 8.51 | 8.27 | 8.41 | 12.31 | 11.98 | 11.09 | 10.73 | 9.95 | 8.66 | 8.34 | 12.15 | 10.91 |
| KIF3A | 3 | 10.33 | 11.03 | 10.86 | 10.40 | 9.14 | 9.67 | 9.16 | 9.82 | 10.14 | 9.44 | 10.68 | 10.63 | 9.40 | 9.49 | 9.79 |
| KIF3B | 3 | 11.16 | 12.20 | 10.12 | 10.16 | 8.73 | 10.15 | 9.28 | 9.54 | 9.44 | 9.22 | 11.68 | 10.14 | 9.44 | 9.41 | 9.33 |

| | | | | | | | | | | | | | | | | |
|---|---|---|---|---|---|---|---|---|---|---|---|---|---|---|---|---|
| KIF4A | 38 | 9.68 | 10.11 | 10.08 | 9.16 | 11.68 | 11.89 | 11.14 | 9.92 | 8.48 | 7.86 | 9.90 | 9.62 | 11.79 | 10.53 | 8.17 |
| KIF5A | 7 | 9.08 | 9.56 | 11.80 | 11.19 | 9.39 | 10.54 | 8.77 | 8.73 | 10.19 | 10.75 | 9.32 | 11.49 | 9.97 | 8.75 | 10.47 |
| KIF5B | 59 | 9.40 | 10.16 | 11.68 | 10.85 | 9.60 | 9.47 | 8.87 | 9.26 | 10.59 | 10.13 | 9.78 | 11.26 | 9.53 | 9.06 | 10.36 |
| KIFAP3 | 2 | 10.50 | 10.59 | 10.74 | 10.82 | 9.95 | 9.30 | 9.26 | 9.34 | 10.59 | 8.90 | 10.55 | 10.78 | 9.62 | 9.30 | 9.74 |
| KIFC1 | 16 | 8.95 | 10.28 | 10.38 | 11.08 | 10.09 | 10.25 | 9.20 | 8.44 | 10.80 | 10.53 | 9.61 | 10.73 | 10.17 | 8.82 | 10.67 |
| KIFC3 | 2 | 13.14 | 10.22 | 9.38 | 9.86 | 9.24 | 10.27 | 9.40 | 11.22 | 8.76 | 8.51 | 11.68 | 9.62 | 9.75 | 10.31 | 8.63 |
| KIN | 11 | 9.86 | 10.98 | 11.40 | 11.31 | 8.09 | 8.67 | 10.33 | 9.24 | 10.20 | 9.93 | 10.42 | 11.35 | 8.38 | 9.78 | 10.06 |
| KIRREL | 2 | 7.82 | 10.11 | 11.67 | 13.29 | 8.68 | 8.11 | 7.23 | 6.12 | 13.31 | 13.67 | 8.96 | 12.48 | 8.39 | 6.67 | 13.49 |
| KKLC1 | 1 | 8.47 | 6.54 | 11.73 | 16.41 | 11.17 | 12.13 | 7.91 | 7.39 | 7.54 | 10.70 | 7.50 | 14.07 | 11.65 | 7.65 | 9.12 |
| KLC1 | 23 | 10.28 | 11.22 | 11.45 | 10.76 | 9.22 | 9.10 | 8.03 | 7.06 | 11.85 | 11.03 | 10.75 | 11.11 | 9.16 | 7.54 | 11.44 |
| KLC2 | 26 | 9.63 | 10.62 | 10.21 | 10.10 | 10.08 | 10.49 | 8.99 | 8.44 | 10.74 | 10.70 | 10.12 | 10.16 | 10.28 | 8.72 | 10.72 |
| KLC4 | 9 | 9.03 | 13.08 | 9.41 | 9.43 | 9.32 | 9.23 | 8.80 | 12.38 | 9.69 | 9.62 | 11.06 | 9.42 | 9.28 | 10.59 | 9.66 |
| KLF13 | 1 | 8.05 | 9.29 | 9.99 | 9.74 | 10.37 | 9.56 | 9.72 | 9.74 | 11.67 | 11.88 | 8.67 | 9.86 | 9.96 | 9.73 | 11.78 |
| KLF16 | 2 | 8.31 | 9.93 | 10.97 | 10.75 | 8.84 | 7.99 | 11.37 | 10.30 | 10.93 | 10.62 | 9.12 | 10.86 | 8.41 | 10.83 | 10.77 |
| KLF4 | 1 | 9.88 | 11.75 | 4.50 | 2.63 | 17.04 | 16.82 | 11.49 | 9.64 | 8.81 | 7.45 | 10.81 | 3.56 | 16.93 | 10.57 | 8.13 |
| KLF5 | 4 | 7.97 | 9.39 | 8.81 | 8.12 | 10.22 | 10.18 | 14.74 | 13.50 | 8.83 | 8.25 | 8.68 | 8.46 | 10.20 | 14.12 | 8.54 |
| KLHDC4 | 4 | 8.46 | 9.66 | 12.26 | 9.78 | 10.91 | 10.31 | 9.55 | 9.41 | 10.06 | 9.59 | 9.06 | 11.02 | 10.61 | 9.48 | 9.83 |
| KLHL11 | 3 | 10.80 | 10.18 | 9.94 | 9.05 | 9.76 | 9.57 | 9.76 | 9.61 | 12.07 | 9.27 | 10.49 | 9.49 | 9.66 | 9.69 | 10.67 |
| KLHL13 | 7 | 10.89 | 11.22 | 7.81 | 7.11 | 12.60 | 12.53 | 9.71 | 9.39 | 9.12 | 9.64 | 11.05 | 7.46 | 12.56 | 9.55 | 9.38 |
| KLHL18 | 1 | 10.92 | 10.88 | 11.53 | 10.96 | 7.24 | 7.95 | 8.69 | 7.64 | 12.48 | 11.71 | 10.90 | 11.25 | 7.60 | 8.16 | 12.09 |
| KLHL21 | 2 | 12.00 | 12.80 | 10.87 | 9.59 | 8.57 | 6.26 | 6.70 | 6.26 | 13.70 | 13.25 | 12.40 | 10.23 | 7.42 | 6.48 | 13.47 |
| KLHL42 | 1 | 9.32 | 9.27 | 9.76 | 9.35 | 11.81 | 12.20 | 9.72 | 10.81 | 9.02 | 8.75 | 9.29 | 9.55 | 12.01 | 10.26 | 8.89 |
| KLHL5 | 1 | 9.38 | 8.60 | 11.53 | 15.17 | 8.77 | 10.13 | 5.73 | 6.86 | 11.12 | 12.72 | 8.99 | 13.35 | 9.45 | 6.30 | 11.92 |
| KMT2A | 1 | 8.17 | 11.41 | 10.00 | 10.31 | 11.34 | 9.97 | 11.06 | 8.75 | 9.47 | 9.53 | 9.79 | 10.15 | 10.65 | 9.91 | 9.50 |
| KMT2B | 1 | 8.80 | 9.38 | 9.79 | 8.58 | 9.37 | 10.22 | 13.47 | 12.87 | 9.38 | 8.12 | 9.09 | 9.19 | 9.80 | 13.17 | 8.75 |
| KNOP1 | 10 | 9.15 | 10.16 | 11.68 | 10.97 | 9.40 | 9.18 | 10.26 | 9.28 | 10.00 | 9.92 | 9.66 | 11.32 | 9.29 | 9.77 | 9.96 |
| KNSTRN | 11 | 9.65 | 10.85 | 10.50 | 10.64 | 8.52 | 8.57 | 8.36 | 10.32 | 11.48 | 11.12 | 10.25 | 10.57 | 8.54 | 9.34 | 11.30 |
| KNTC1 | 15 | 10.86 | 10.29 | 8.13 | 8.91 | 10.38 | 10.38 | 7.97 | 7.81 | 12.78 | 12.50 | 10.58 | 8.52 | 10.38 | 7.89 | 12.64 |
| KPNA1 | 20 | 9.71 | 10.36 | 11.29 | 10.57 | 9.95 | 10.26 | 9.90 | 9.23 | 9.58 | 9.15 | 10.03 | 10.93 | 10.10 | 9.57 | 9.37 |
| KPNA2 | 32 | 9.24 | 10.56 | 11.51 | 11.28 | 8.22 | 8.72 | 9.20 | 9.25 | 11.12 | 10.90 | 9.90 | 11.39 | 8.47 | 9.23 | 11.01 |
| KPNA3 | 4 | 9.91 | 10.54 | 13.46 | 12.09 | 9.56 | 9.82 | 8.90 | 7.90 | 8.95 | 8.88 | 10.22 | 12.77 | 9.69 | 8.40 | 8.91 |
| KPNA4 | 18 | 9.15 | 9.68 | 12.63 | 10.09 | 9.42 | 8.87 | 9.61 | 10.87 | 9.92 | 9.75 | 9.42 | 11.36 | 9.14 | 10.24 | 9.84 |
| KPNA6 | 11 | 9.52 | 12.45 | 9.89 | 9.08 | 8.01 | 8.20 | 9.43 | 13.97 | 9.75 | 9.69 | 10.99 | 9.49 | 8.10 | 11.70 | 9.72 |
| KPNB1 | 74 | 9.63 | 10.18 | 10.69 | 10.56 | 8.94 | 9.05 | 8.10 | 11.49 | 10.69 | 10.68 | 9.90 | 10.63 | 9.00 | 9.79 | 10.68 |
| KRAS | 3 | 10.29 | 8.93 | 11.85 | 12.65 | 9.08 | 8.48 | 10.10 | 7.85 | 10.32 | 10.46 | 9.61 | 12.25 | 8.78 | 8.97 | 10.39 |
| KRI1 | 17 | 8.29 | 9.40 | 10.52 | 10.68 | 9.80 | 10.27 | 11.48 | 10.89 | 9.39 | 9.28 | 8.84 | 10.60 | 10.04 | 11.19 | 9.33 |
| KRR1 | 9 | 9.31 | 9.25 | 10.28 | 9.63 | 8.52 | 7.96 | 13.27 | 11.35 | 9.39 | 11.03 | 9.28 | 9.96 | 8.24 | 12.31 | 10.21 |
| KSR1 | 2 | 7.17 | 7.73 | 18.22 | 18.73 | 7.01 | 7.30 | 11.93 | 11.24 | 5.50 | 5.18 | 7.45 | 18.47 | 7.16 | 11.59 | 5.34 |
| KTI12 | 6 | 10.21 | 10.44 | 9.31 | 9.07 | 8.72 | 9.26 | 10.90 | 10.48 | 11.10 | 10.52 | 10.32 | 9.19 | 8.99 | 10.69 | 10.81 |
| KTN1 | 77 | 13.00 | 8.41 | 8.73 | 11.31 | 10.70 | 9.39 | 8.71 | 8.15 | 9.84 | 11.78 | 10.70 | 10.02 | 10.04 | 8.43 | 10.81 |
| KYNU | 24 | 8.91 | 9.21 | 5.41 | 4.41 | 14.73 | 13.73 | 16.25 | 17.65 | 5.09 | 4.62 | 9.06 | 4.91 | 14.23 | 16.95 | 4.85 |
| L1CAM | 26 | 11.65 | 9.78 | 4.11 | 4.55 | 12.19 | 12.07 | 10.45 | 11.04 | 11.86 | 12.30 | 10.71 | 4.33 | 12.13 | 10.75 | 12.08 |

| | | | | | | | | | | | | | | | | |
|---|---|---|---|---|---|---|---|---|---|---|---|---|---|---|---|---|
| L1RE1 | 3 | 9.78 | 8.96 | 9.74 | 9.83 | 15.81 | 14.66 | 5.90 | 5.47 | 10.05 | 9.79 | 9.37 | 9.78 | 15.24 | 5.69 | 9.92 |
| L2HGDH | 3 | 11.43 | 9.46 | 7.56 | 8.87 | 11.60 | 11.30 | 11.94 | 10.20 | 9.42 | 8.22 | 10.44 | 8.22 | 11.45 | 11.07 | 8.82 |
| L3HYPDH | 4 | 6.55 | 7.45 | 16.18 | 15.37 | 10.30 | 11.05 | 6.54 | 7.32 | 9.55 | 9.69 | 7.00 | 15.77 | 10.67 | 6.93 | 9.62 |
| L3MBTL2 | 2 | 8.81 | 9.19 | 11.88 | 12.84 | 8.65 | 8.94 | 9.97 | 10.73 | 9.77 | 9.23 | 9.00 | 12.36 | 8.80 | 10.35 | 9.50 |
| L3MBTL3 | 3 | 8.04 | 7.95 | 13.42 | 11.86 | 6.62 | 6.32 | 16.68 | 13.95 | 7.48 | 7.68 | 8.00 | 12.64 | 6.47 | 15.31 | 7.58 |
| LACTB | 5 | 8.47 | 6.64 | 16.34 | 16.55 | 6.11 | 7.30 | 8.83 | 8.68 | 11.03 | 10.05 | 7.56 | 16.45 | 6.70 | 8.76 | 10.54 |
| LACTB2 | 10 | 9.87 | 10.74 | 9.23 | 9.14 | 9.56 | 9.21 | 13.14 | 12.04 | 8.44 | 8.62 | 10.30 | 9.19 | 9.39 | 12.59 | 8.53 |
| LAGE3 | 2 | 8.46 | 9.88 | 11.44 | 11.20 | 10.48 | 9.98 | 9.61 | 8.42 | 10.53 | 10.00 | 9.17 | 11.32 | 10.23 | 9.01 | 10.27 |
| LAMA4 | 2 | 8.59 | 7.25 | 15.95 | 16.75 | 4.47 | 4.17 | 11.29 | 10.16 | 9.90 | 11.47 | 7.92 | 16.35 | 4.32 | 10.72 | 10.68 |
| LAMA5 | 9 | 10.25 | 8.14 | 8.99 | 9.94 | 10.21 | 10.51 | 10.87 | 10.54 | 10.14 | 10.42 | 9.19 | 9.46 | 10.36 | 10.70 | 10.28 |
| LAMB1 | 25 | 5.59 | 4.43 | 15.45 | 18.34 | 4.38 | 4.15 | 14.50 | 13.55 | 9.42 | 10.19 | 5.01 | 16.90 | 4.26 | 14.02 | 9.81 |
| LAMB2 | 1 | 16.51 | 11.27 | 3.62 | 3.29 | 16.95 | 17.73 | 7.37 | 5.20 | 9.00 | 9.07 | 13.89 | 3.45 | 17.34 | 6.29 | 9.04 |
| LAMB3 | 2 | 12.77 | 10.02 | 6.45 | 5.62 | 15.00 | 15.85 | 6.79 | 7.91 | 9.85 | 9.73 | 11.39 | 6.04 | 15.43 | 7.35 | 9.79 |
| LAMC1 | 24 | 8.14 | 6.20 | 11.53 | 13.88 | 8.43 | 7.73 | 12.07 | 11.32 | 9.95 | 10.76 | 7.17 | 12.70 | 8.08 | 11.69 | 10.36 |
| LAMP1 | 10 | 11.17 | 8.60 | 9.81 | 9.78 | 12.63 | 12.14 | 8.69 | 7.31 | 9.73 | 10.14 | 9.89 | 9.79 | 12.39 | 8.00 | 9.93 |
| LAMP2 | 4 | 8.39 | 8.65 | 8.32 | 8.40 | 11.57 | 9.74 | 9.69 | 17.66 | 8.74 | 8.85 | 8.52 | 8.36 | 10.65 | 13.67 | 8.80 |
| LAMTOR1 | 4 | 10.58 | 8.21 | 9.01 | 10.69 | 10.75 | 9.72 | 10.24 | 9.22 | 10.43 | 11.15 | 9.40 | 9.85 | 10.24 | 9.73 | 10.79 |
| LAMTOR2 | 6 | 9.10 | 10.03 | 10.63 | 11.38 | 11.10 | 11.17 | 9.03 | 9.11 | 9.43 | 9.02 | 9.57 | 11.00 | 11.14 | 9.07 | 9.23 |
| LAMTOR3 | 5 | 10.79 | 10.86 | 9.10 | 7.63 | 11.96 | 11.59 | 9.95 | 8.08 | 11.00 | 9.03 | 10.83 | 8.37 | 11.78 | 9.01 | 10.02 |
| LAMTOR4 | 2 | 9.83 | 9.56 | 10.25 | 10.44 | 10.36 | 10.95 | 10.36 | 9.67 | 9.27 | 9.32 | 9.69 | 10.35 | 10.65 | 10.01 | 9.29 |
| LAMTOR5 | 1 | 9.21 | 9.14 | 15.47 | 3.47 | 14.31 | 10.23 | 12.50 | 3.24 | 12.08 | 10.34 | 9.18 | 9.47 | 12.27 | 7.87 | 11.21 |
| LANCL1 | 10 | 8.81 | 8.85 | 9.50 | 8.35 | 11.62 | 11.06 | 9.73 | 17.93 | 7.02 | 7.13 | 8.83 | 8.93 | 11.34 | 13.83 | 7.07 |
| LANCL2 | 5 | 9.46 | 9.62 | 9.91 | 9.04 | 11.32 | 11.92 | 9.30 | 10.53 | 9.63 | 9.27 | 9.54 | 9.48 | 11.62 | 9.92 | 9.45 |
| LAP3 | 28 | 9.38 | 9.56 | 8.14 | 7.88 | 13.04 | 12.90 | 10.41 | 12.10 | 8.43 | 8.17 | 9.47 | 8.01 | 12.97 | 11.25 | 8.30 |
| LARP1 | 59 | 9.12 | 9.60 | 10.06 | 8.98 | 9.39 | 9.44 | 12.76 | 11.28 | 9.74 | 9.62 | 9.36 | 9.52 | 9.41 | 12.02 | 9.68 |
| LARP1B | 1 | 8.38 | 10.15 | 11.36 | 11.62 | 11.38 | 11.63 | 10.03 | 9.15 | 7.81 | 8.52 | 9.26 | 11.49 | 11.50 | 9.59 | 8.16 |
| LARP4 | 13 | 9.50 | 10.04 | 10.98 | 10.39 | 9.45 | 9.31 | 9.31 | 12.63 | 9.81 | 8.58 | 9.77 | 10.68 | 9.38 | 10.97 | 9.20 |
| LARP4B | 13 | 9.37 | 10.50 | 10.65 | 10.12 | 9.55 | 9.72 | 11.14 | 10.40 | 9.33 | 9.23 | 9.93 | 10.39 | 9.63 | 10.77 | 9.28 |
| LARP7 | 20 | 9.54 | 9.73 | 10.47 | 10.82 | 10.37 | 10.88 | 10.48 | 9.64 | 9.04 | 9.04 | 9.63 | 10.64 | 10.63 | 10.06 | 9.04 |
| LARS | 73 | 8.95 | 9.36 | 11.25 | 9.00 | 10.30 | 9.66 | 10.84 | 10.70 | 10.07 | 9.87 | 9.16 | 10.13 | 9.98 | 10.77 | 9.97 |
| LARS2 | 14 | 12.58 | 10.61 | 8.85 | 9.55 | 10.35 | 9.47 | 9.48 | 8.98 | 9.96 | 10.18 | 11.60 | 9.20 | 9.91 | 9.23 | 10.07 |
| LAS1L | 12 | 9.77 | 10.34 | 9.25 | 8.36 | 10.00 | 10.07 | 11.35 | 11.43 | 9.89 | 9.55 | 10.06 | 8.81 | 10.03 | 11.39 | 9.72 |
| LASP1 | 24 | 9.98 | 11.60 | 16.30 | 16.01 | 8.51 | 8.86 | 4.04 | 3.61 | 10.77 | 10.32 | 10.79 | 16.15 | 8.69 | 3.82 | 10.55 |
| LATS1 | 1 | 9.64 | 10.93 | 9.87 | 13.16 | 8.98 | 9.82 | 9.57 | 10.18 | 8.40 | 9.46 | 10.29 | 11.52 | 9.40 | 9.87 | 8.93 |
| LBR | 6 | 13.52 | 6.85 | 6.94 | 10.74 | 9.43 | 8.03 | 10.59 | 9.51 | 10.76 | 13.64 | 10.18 | 8.84 | 8.73 | 10.05 | 12.20 |
| LCLAT1 | 5 | 11.65 | 6.05 | 9.06 | 14.79 | 8.02 | 7.38 | 10.04 | 8.56 | 10.81 | 13.63 | 8.85 | 11.92 | 7.70 | 9.30 | 12.22 |
| LCMT1 | 8 | 10.18 | 10.01 | 10.19 | 7.87 | 12.75 | 11.59 | 10.87 | 8.60 | 9.26 | 8.68 | 10.09 | 9.03 | 12.17 | 9.74 | 8.97 |
| LCMT2 | 4 | 9.57 | 9.85 | 6.97 | 6.67 | 13.16 | 12.48 | 9.00 | 8.73 | 11.89 | 11.68 | 9.71 | 6.82 | 12.82 | 8.86 | 11.78 |
| LDHA | 107 | 9.05 | 13.48 | 9.18 | 9.35 | 10.06 | 9.79 | 10.50 | 10.59 | 9.01 | 9.00 | 11.27 | 9.26 | 9.92 | 10.55 | 9.00 |
| LDHB | 87 | 8.87 | 11.25 | 11.26 | 10.50 | 10.54 | 10.33 | 9.38 | 9.45 | 9.28 | 9.15 | 10.06 | 10.88 | 10.44 | 9.41 | 9.21 |
| LDLR | 4 | 7.89 | 5.82 | 7.58 | 9.66 | 10.05 | 9.23 | 13.97 | 12.92 | 10.48 | 12.40 | 6.86 | 8.62 | 9.64 | 13.45 | 11.44 |
| LDLRAP1 | 1 | 10.06 | 11.78 | 8.79 | 7.40 | 9.75 | 10.52 | 13.16 | 11.58 | 8.57 | 8.38 | 10.92 | 8.10 | 10.14 | 12.37 | 8.47 |

| | | | | | | | | | | | | | | | | |
|---|---|---|---|---|---|---|---|---|---|---|---|---|---|---|---|---|
| LEMD2 | 3 | 10.91 | 6.28 | 9.08 | 13.06 | 9.37 | 7.06 | 9.33 | 8.72 | 11.21 | 14.98 | 8.59 | 11.07 | 8.21 | 9.03 | 13.10 |
| LEMD3 | 6 | 12.49 | 6.81 | 7.48 | 11.88 | 10.47 | 9.08 | 8.24 | 7.62 | 10.94 | 14.99 | 9.65 | 9.68 | 9.78 | 7.93 | 12.97 |
| LENG1 | 2 | 9.91 | 10.05 | 11.33 | 11.31 | 8.85 | 8.90 | 11.95 | 11.13 | 8.00 | 8.57 | 9.98 | 11.32 | 8.87 | 11.54 | 8.29 |
| LENG8 | 4 | 9.45 | 10.57 | 10.64 | 10.83 | 9.29 | 9.76 | 10.33 | 9.10 | 10.31 | 9.71 | 10.01 | 10.74 | 9.53 | 9.72 | 10.01 |
| LENG9 | 2 | 7.55 | 9.67 | 9.84 | 9.74 | 11.16 | 11.34 | 11.46 | 10.59 | 9.10 | 9.54 | 8.61 | 9.79 | 11.25 | 11.03 | 9.32 |
| LEO1 | 8 | 9.76 | 10.60 | 10.40 | 10.37 | 10.21 | 10.44 | 10.03 | 8.29 | 10.26 | 9.64 | 10.18 | 10.39 | 10.32 | 9.16 | 9.95 |
| LEPRE1 | 18 | 10.10 | 8.23 | 7.77 | 8.50 | 9.68 | 8.78 | 13.10 | 11.66 | 10.69 | 11.48 | 9.17 | 8.13 | 9.23 | 12.38 | 11.09 |
| LEPREL1 | 10 | 8.37 | 7.68 | 9.12 | 10.50 | 6.05 | 6.02 | 15.01 | 14.03 | 11.22 | 11.99 | 8.03 | 9.81 | 6.04 | 14.52 | 11.60 |
| LEPREL2 | 7 | 12.88 | 11.16 | 6.22 | 7.35 | 3.13 | 2.79 | 13.46 | 13.04 | 14.73 | 15.25 | 12.02 | 6.79 | 2.96 | 13.25 | 14.99 |
| LEPREL4 | 8 | 10.84 | 9.57 | 8.13 | 8.00 | 7.14 | 7.13 | 12.51 | 11.26 | 12.45 | 12.96 | 10.21 | 8.07 | 7.14 | 11.88 | 12.71 |
| LETM1 | 25 | 11.83 | 6.55 | 7.51 | 11.57 | 8.73 | 7.54 | 11.26 | 11.29 | 11.05 | 12.66 | 9.19 | 9.54 | 8.14 | 11.28 | 11.85 |
| LGALS1 | 45 | 7.25 | 8.90 | 14.96 | 14.19 | 6.75 | 6.66 | 7.48 | 9.67 | 12.41 | 11.73 | 8.07 | 14.58 | 6.70 | 8.57 | 12.07 |
| LGALS3 | 16 | 9.22 | 10.33 | 8.25 | 7.62 | 10.70 | 10.57 | 10.02 | 10.82 | 11.25 | 11.22 | 9.78 | 7.93 | 10.63 | 10.42 | 11.24 |
| LGALS3BP | 12 | 12.66 | 10.45 | 8.17 | 8.51 | 7.58 | 7.38 | 11.00 | 10.14 | 11.50 | 12.60 | 11.56 | 8.34 | 7.48 | 10.57 | 12.05 |
| LGALS8 | 4 | 13.11 | 11.18 | 8.09 | 9.05 | 12.64 | 12.90 | 8.42 | 7.50 | 8.76 | 8.33 | 12.15 | 8.57 | 12.77 | 7.96 | 8.55 |
| LGALSL | 3 | 9.33 | 9.93 | 13.35 | 13.07 | 9.55 | 9.44 | 7.41 | 6.23 | 10.46 | 11.23 | 9.63 | 13.21 | 9.49 | 6.82 | 10.84 |
| LGMN | 2 | 8.71 | 10.10 | 13.79 | 13.39 | 9.50 | 9.63 | 9.97 | 8.77 | 7.64 | 8.49 | 9.41 | 13.59 | 9.56 | 9.37 | 8.06 |
| LIG1 | 20 | 9.32 | 10.20 | 8.88 | 8.47 | 10.90 | 11.10 | 11.33 | 10.48 | 9.87 | 9.46 | 9.76 | 8.67 | 11.00 | 10.90 | 9.66 |
| LIG3 | 30 | 10.33 | 10.95 | 10.53 | 10.13 | 9.11 | 9.08 | 10.34 | 9.75 | 10.16 | 9.64 | 10.64 | 10.33 | 9.09 | 10.04 | 9.90 |
| LIG4 | 10 | 9.31 | 10.45 | 12.87 | 12.58 | 8.92 | 9.29 | 10.13 | 9.53 | 8.51 | 8.42 | 9.88 | 12.72 | 9.10 | 9.83 | 8.46 |
| LIMA1 | 44 | 9.63 | 11.47 | 24.01 | 22.38 | 2.84 | 2.88 | 5.19 | 4.43 | 8.44 | 8.74 | 10.55 | 23.19 | 2.86 | 4.81 | 8.59 |
| LIMCH1 | 54 | 9.28 | 10.53 | 11.46 | 10.79 | 12.64 | 13.01 | 5.64 | 5.20 | 10.85 | 10.61 | 9.91 | 11.12 | 12.82 | 5.42 | 10.73 |
| LIMD1 | 9 | 9.73 | 11.47 | 10.12 | 8.23 | 11.33 | 11.77 | 9.06 | 8.08 | 10.32 | 9.89 | 10.60 | 9.17 | 11.55 | 8.57 | 10.10 |
| LIMK1 | 1 | 10.35 | 8.41 | 11.56 | 13.61 | 7.90 | 8.30 | 9.28 | 7.86 | 11.50 | 11.24 | 9.38 | 12.58 | 8.10 | 8.57 | 11.37 |
| LIMS1 | 10 | 8.36 | 8.23 | 13.49 | 11.04 | 7.21 | 6.78 | 6.79 | 17.06 | 10.87 | 10.16 | 8.30 | 12.26 | 7.00 | 11.93 | 10.51 |
| LIN54 | 4 | 9.74 | 9.52 | 9.73 | 9.04 | 11.41 | 11.19 | 10.86 | 11.13 | 8.93 | 8.45 | 9.63 | 9.39 | 11.30 | 10.99 | 8.69 |
| LIN7C | 6 | 9.68 | 10.88 | 10.97 | 10.34 | 10.11 | 10.18 | 9.96 | 9.72 | 9.26 | 8.90 | 10.28 | 10.66 | 10.15 | 9.84 | 9.08 |
| LIN9 | 4 | 9.79 | 11.00 | 7.90 | 8.52 | 11.33 | 11.57 | 10.70 | 10.80 | 9.54 | 8.83 | 10.39 | 8.21 | 11.45 | 10.75 | 9.19 |
| LIPA | 2 | 8.55 | 10.35 | 12.86 | 13.72 | 11.00 | 11.50 | 9.31 | 9.82 | 6.73 | 6.15 | 9.45 | 13.29 | 11.25 | 9.56 | 6.44 |
| LLGL1 | 9 | 10.29 | 11.62 | 9.82 | 9.64 | 10.45 | 10.40 | 8.06 | 7.60 | 11.42 | 10.69 | 10.96 | 9.73 | 10.43 | 7.83 | 11.05 |
| LLPH | 6 | 9.11 | 10.34 | 10.29 | 10.13 | 8.50 | 8.63 | 12.66 | 12.02 | 9.50 | 8.82 | 9.72 | 10.21 | 8.57 | 12.34 | 9.16 |
| LMAN1 | 26 | 12.27 | 7.04 | 7.40 | 10.48 | 11.22 | 9.94 | 8.63 | 8.59 | 11.08 | 13.35 | 9.65 | 8.94 | 10.58 | 8.61 | 12.22 |
| LMAN2 | 21 | 11.37 | 6.69 | 7.24 | 9.88 | 10.77 | 9.25 | 11.18 | 13.74 | 9.16 | 10.73 | 9.03 | 8.56 | 10.01 | 12.46 | 9.94 |
| LMAN2L | 3 | 13.83 | 8.12 | 9.58 | 11.30 | 11.13 | 10.47 | 8.76 | 9.75 | 8.20 | 8.87 | 10.97 | 10.44 | 10.80 | 9.26 | 8.54 |
| LMCD1 | 1 | 8.95 | 8.57 | 13.99 | 15.17 | 8.33 | 9.04 | 12.13 | 10.93 | 5.74 | 7.15 | 8.76 | 14.58 | 8.69 | 11.53 | 6.44 |
| LMF2 | 3 | 14.96 | 6.85 | 6.76 | 10.41 | 9.36 | 7.20 | 7.85 | 7.28 | 12.83 | 16.50 | 10.90 | 8.58 | 8.28 | 7.57 | 14.66 |
| LMNA | 198 | 8.89 | 10.48 | 10.26 | 9.55 | 11.38 | 11.65 | 8.42 | 8.78 | 10.62 | 9.97 | 9.69 | 9.90 | 11.51 | 8.60 | 10.30 |
| LMNB1 | 62 | 8.98 | 10.24 | 9.15 | 8.34 | 9.66 | 9.58 | 13.10 | 12.30 | 9.60 | 9.05 | 9.61 | 8.74 | 9.62 | 12.70 | 9.33 |
| LMNB2 | 49 | 8.46 | 9.24 | 12.88 | 11.99 | 8.81 | 8.83 | 10.01 | 9.51 | 10.37 | 9.91 | 8.85 | 12.43 | 8.82 | 9.76 | 10.14 |
| LMO4 | 1 | 11.73 | 9.29 | 9.82 | 8.72 | 7.82 | 8.09 | 12.72 | 10.45 | 10.47 | 10.90 | 10.51 | 9.27 | 7.95 | 11.59 | 10.68 |
| LMO7 | 74 | 10.15 | 11.48 | 8.49 | 8.05 | 12.76 | 13.55 | 10.28 | 9.13 | 8.36 | 7.76 | 10.82 | 8.27 | 13.15 | 9.70 | 8.06 |
| LNP | 3 | 12.16 | 6.75 | 7.68 | 11.89 | 8.60 | 7.12 | 10.16 | 10.33 | 10.73 | 14.59 | 9.45 | 9.79 | 7.86 | 10.24 | 12.66 |

| | | | | | | | | | | | | | | | | |
|---|---|---|---|---|---|---|---|---|---|---|---|---|---|---|---|---|
| LNPEP | 12 | 9.70 | 7.12 | 9.06 | 9.83 | 11.28 | 10.00 | 10.01 | 10.05 | 10.93 | 12.03 | 8.41 | 9.44 | 10.64 | 10.03 | 11.48 |
| LOH12CR1 | 1 | 9.81 | 11.40 | 10.17 | 10.20 | 11.38 | 12.26 | 9.62 | 9.31 | 8.14 | 7.70 | 10.60 | 10.18 | 11.82 | 9.47 | 7.92 |
| LONP1 | 32 | 11.18 | 8.74 | 8.70 | 9.88 | 10.37 | 9.53 | 9.57 | 10.75 | 10.57 | 10.70 | 9.96 | 9.29 | 9.95 | 10.16 | 10.64 |
| LONP2 | 4 | 10.34 | 11.25 | 7.38 | 7.55 | 13.86 | 14.77 | 8.11 | 7.56 | 10.03 | 9.15 | 10.80 | 7.46 | 14.32 | 7.84 | 9.59 |
| LOX | 3 | 10.48 | 10.99 | 11.38 | 10.59 | 9.93 | 10.24 | 7.90 | 7.45 | 10.33 | 10.70 | 10.74 | 10.98 | 10.09 | 7.68 | 10.51 |
| LOXL2 | 2 | 8.60 | 6.86 | 8.51 | 9.43 | 8.62 | 4.80 | 19.03 | 15.70 | 8.19 | 10.25 | 7.73 | 8.97 | 6.71 | 17.36 | 9.22 |
| LPCAT1 | 8 | 11.37 | 5.66 | 5.18 | 7.00 | 8.77 | 6.56 | 13.52 | 16.61 | 11.32 | 14.00 | 8.52 | 6.09 | 7.66 | 15.07 | 12.66 |
| LPCAT3 | 3 | 15.23 | 7.75 | 5.17 | 8.43 | 8.90 | 7.36 | 11.89 | 8.90 | 11.64 | 14.74 | 11.49 | 6.80 | 8.13 | 10.40 | 13.19 |
| LPCAT4 | 2 | 10.38 | 5.05 | 7.62 | 11.63 | 8.07 | 8.29 | 11.18 | 10.91 | 11.89 | 14.98 | 7.72 | 9.63 | 8.18 | 11.05 | 13.43 |
| LPGAT1 | 4 | 10.52 | 7.32 | 8.11 | 11.37 | 11.33 | 10.07 | 7.30 | 6.21 | 12.51 | 15.26 | 8.92 | 9.74 | 10.70 | 6.76 | 13.89 |
| LPIN3 | 4 | 8.21 | 11.17 | 10.16 | 11.20 | 9.74 | 10.29 | 9.53 | 8.97 | 10.49 | 10.24 | 9.69 | 10.68 | 10.01 | 9.25 | 10.36 |
| LPL | 7 | 16.03 | 14.39 | 7.14 | 7.17 | 3.87 | 3.85 | 5.67 | 5.04 | 17.18 | 19.66 | 15.21 | 7.15 | 3.86 | 5.35 | 18.42 |
| LPP | 19 | 11.38 | 11.22 | 14.70 | 12.78 | 9.54 | 9.14 | 6.27 | 6.39 | 9.70 | 8.88 | 11.30 | 13.74 | 9.34 | 6.33 | 9.29 |
| LRBA | 29 | 10.11 | 10.21 | 8.11 | 8.67 | 12.49 | 12.65 | 9.74 | 9.66 | 9.16 | 9.20 | 10.16 | 8.39 | 12.57 | 9.70 | 9.18 |
| LRCH1 | 1 | 9.24 | 10.03 | 11.79 | 11.67 | 11.01 | 9.89 | 10.34 | 8.98 | 8.77 | 8.28 | 9.63 | 11.73 | 10.45 | 9.66 | 8.52 |
| LRCH3 | 4 | 9.52 | 10.55 | 10.97 | 9.97 | 8.11 | 8.29 | 11.50 | 10.65 | 10.78 | 9.65 | 10.04 | 10.47 | 8.20 | 11.07 | 10.22 |
| LRIF1 | 1 | 8.91 | 10.66 | 8.15 | 10.78 | 9.33 | 10.11 | 14.54 | 11.39 | 9.37 | 6.76 | 9.78 | 9.47 | 9.72 | 12.96 | 8.06 |
| LRP1 | 16 | 9.89 | 7.88 | 10.66 | 13.47 | 10.70 | 9.96 | 8.15 | 8.42 | 10.02 | 10.85 | 8.88 | 12.06 | 10.33 | 8.29 | 10.44 |
| LRP11 | 1 | 10.59 | 7.80 | 6.13 | 7.22 | 9.79 | 10.72 | 11.32 | 12.82 | 12.13 | 11.48 | 9.19 | 6.67 | 10.25 | 12.07 | 11.81 |
| LRP5 | 2 | 10.32 | 8.91 | 7.15 | 8.91 | 11.32 | 12.35 | 10.62 | 8.75 | 11.00 | 10.66 | 9.62 | 8.03 | 11.84 | 9.69 | 10.83 |
| LRPAP1 | 15 | 8.89 | 9.28 | 8.71 | 9.24 | 11.09 | 11.24 | 12.51 | 12.44 | 8.34 | 8.26 | 9.09 | 8.98 | 11.16 | 12.47 | 8.30 |
| LRPPRC | 86 | 10.67 | 8.34 | 9.12 | 9.77 | 9.65 | 8.45 | 9.80 | 11.80 | 10.83 | 11.57 | 9.51 | 9.45 | 9.05 | 10.80 | 11.20 |
| LRRC14 | 3 | 10.33 | 9.88 | 9.88 | 8.61 | 9.19 | 9.17 | 10.86 | 11.76 | 10.18 | 10.13 | 10.11 | 9.25 | 9.18 | 11.31 | 10.16 |
| LRRC15 | 1 | 8.20 | 9.06 | 18.24 | 25.91 | 6.17 | 9.73 | 5.36 | 4.75 | 7.06 | 5.52 | 8.63 | 22.08 | 7.95 | 5.05 | 6.29 |
| LRRC16A | 13 | 10.07 | 10.85 | 10.49 | 10.08 | 10.08 | 10.29 | 8.81 | 8.94 | 10.71 | 9.68 | 10.46 | 10.29 | 10.18 | 8.88 | 10.20 |
| LRRC20 | 1 | 9.79 | 12.09 | 7.38 | 6.66 | 15.04 | 15.64 | 6.84 | 6.31 | 10.97 | 9.26 | 10.94 | 7.02 | 15.34 | 6.58 | 10.12 |
| LRRC40 | 28 | 10.58 | 11.79 | 10.25 | 9.66 | 7.02 | 7.03 | 10.52 | 10.67 | 11.31 | 11.16 | 11.18 | 9.96 | 7.03 | 10.59 | 11.24 |
| LRRC41 | 3 | 9.46 | 11.45 | 6.40 | 5.99 | 9.10 | 10.21 | 13.38 | 13.12 | 10.90 | 9.97 | 10.46 | 6.20 | 9.66 | 13.25 | 10.44 |
| LRRC42 | 1 | 10.83 | 12.70 | 10.57 | 9.83 | 8.42 | 9.12 | 6.85 | 6.89 | 12.87 | 11.92 | 11.77 | 10.20 | 8.77 | 6.87 | 12.40 |
| LRRC45 | 5 | 9.63 | 11.19 | 11.20 | 10.71 | 9.72 | 10.18 | 7.98 | 7.62 | 10.91 | 10.86 | 10.41 | 10.96 | 9.95 | 7.80 | 10.88 |
| LRRC47 | 27 | 9.09 | 10.63 | 10.84 | 9.44 | 8.78 | 8.83 | 10.26 | 12.85 | 9.71 | 9.58 | 9.86 | 10.14 | 8.80 | 11.56 | 9.64 |
| LRRC57 | 5 | 8.37 | 9.47 | 8.77 | 8.76 | 12.47 | 12.50 | 9.22 | 8.62 | 10.96 | 10.84 | 8.92 | 8.77 | 12.49 | 8.92 | 10.90 |
| LRRC59 | 16 | 12.07 | 6.48 | 9.45 | 13.02 | 6.60 | 5.72 | 8.00 | 13.38 | 11.15 | 14.11 | 9.27 | 11.24 | 6.16 | 10.69 | 12.63 |
| LRRC8A | 7 | 12.24 | 8.82 | 7.35 | 10.03 | 8.60 | 8.62 | 9.83 | 8.88 | 12.51 | 13.12 | 10.53 | 8.69 | 8.61 | 9.36 | 12.81 |
| LRRC8C | 1 | 11.69 | 8.44 | 7.23 | 10.48 | 5.35 | 5.05 | 12.78 | 13.08 | 12.03 | 13.87 | 10.07 | 8.85 | 5.20 | 12.93 | 12.95 |
| LRRC8D | 1 | 13.59 | 8.36 | 9.52 | 8.64 | 8.90 | 8.06 | 8.93 | 10.55 | 11.60 | 11.86 | 10.97 | 9.08 | 8.48 | 9.74 | 11.73 |
| LRRC9 | 1 | 10.87 | 13.63 | 9.70 | 8.92 | 7.86 | 9.87 | 9.87 | 9.30 | 10.50 | 9.49 | 12.25 | 9.31 | 8.86 | 9.58 | 9.99 |
| LRRFIP1 | 40 | 9.02 | 10.15 | 12.35 | 11.82 | 8.75 | 9.01 | 8.72 | 9.16 | 10.74 | 10.28 | 9.58 | 12.09 | 8.88 | 8.94 | 10.51 |
| LRRFIP2 | 13 | 9.70 | 11.66 | 8.79 | 9.21 | 9.48 | 10.10 | 7.84 | 7.14 | 13.29 | 12.80 | 10.68 | 9.00 | 9.79 | 7.49 | 13.05 |
| LRSAM1 | 15 | 9.85 | 10.60 | 12.07 | 11.50 | 8.90 | 9.06 | 7.86 | 6.89 | 11.87 | 11.38 | 10.23 | 11.79 | 8.98 | 7.38 | 11.63 |
| LRWD1 | 7 | 9.74 | 10.16 | 11.24 | 10.95 | 8.39 | 7.38 | 10.91 | 9.93 | 10.79 | 10.51 | 9.95 | 11.10 | 7.89 | 10.42 | 10.65 |
| LSG1 | 19 | 9.78 | 10.16 | 12.59 | 11.57 | 9.09 | 9.16 | 8.83 | 8.13 | 10.59 | 10.11 | 9.97 | 12.08 | 9.12 | 8.48 | 10.35 |

| | | | | | | | | | | | | | | | | |
|---|---|---|---|---|---|---|---|---|---|---|---|---|---|---|---|---|
| LSM1 | 2 | 9.16 | 9.50 | 10.88 | 11.11 | 9.41 | 10.45 | 10.72 | 9.00 | 10.21 | 9.55 | 9.33 | 11.00 | 9.93 | 9.86 | 9.88 |
| LSM10 | 1 | 9.90 | 11.46 | 7.84 | 8.77 | 9.43 | 9.34 | 14.23 | 12.82 | 8.29 | 7.93 | 10.68 | 8.30 | 9.39 | 13.52 | 8.11 |
| LSM12 | 2 | 9.69 | 9.20 | 11.99 | 10.34 | 9.95 | 9.54 | 10.00 | 9.24 | 10.10 | 9.95 | 9.44 | 11.17 | 9.75 | 9.62 | 10.03 |
| LSM14A | 6 | 9.37 | 10.32 | 9.01 | 8.72 | 9.32 | 10.32 | 12.87 | 12.02 | 9.21 | 8.86 | 9.84 | 8.87 | 9.82 | 12.44 | 9.03 |
| LSM14B | 6 | 9.15 | 9.67 | 11.70 | 11.01 | 8.80 | 9.46 | 8.75 | 8.46 | 12.42 | 10.58 | 9.41 | 11.35 | 9.13 | 8.61 | 11.50 |
| LSM2 | 7 | 9.17 | 9.68 | 11.93 | 7.74 | 11.37 | 10.05 | 11.93 | 9.78 | 9.34 | 9.02 | 9.42 | 9.83 | 10.71 | 10.85 | 9.18 |
| LSM3 | 3 | 9.10 | 9.99 | 11.03 | 10.85 | 10.99 | 10.22 | 11.37 | 7.55 | 9.42 | 9.49 | 9.55 | 10.94 | 10.60 | 9.46 | 9.45 |
| LSM4 | 7 | 8.39 | 8.20 | 10.07 | 9.19 | 8.59 | 8.45 | 11.86 | 18.58 | 8.75 | 7.92 | 8.29 | 9.63 | 8.52 | 15.22 | 8.33 |
| LSM6 | 4 | 9.17 | 9.38 | 10.48 | 9.32 | 9.11 | 9.12 | 13.41 | 12.38 | 8.86 | 8.77 | 9.27 | 9.90 | 9.11 | 12.89 | 8.82 |
| LSM7 | 2 | 8.43 | 8.79 | 11.52 | 9.70 | 9.08 | 8.51 | 9.85 | 16.54 | 9.09 | 8.50 | 8.61 | 10.61 | 8.80 | 13.19 | 8.79 |
| LSMD1 | 1 | 8.21 | 9.00 | 12.12 | 8.61 | 11.49 | 9.09 | 11.93 | 7.08 | 11.12 | 11.36 | 8.60 | 10.36 | 10.29 | 9.50 | 11.24 |
| LSS | 6 | 12.86 | 13.35 | 10.53 | 10.39 | 10.57 | 11.01 | 6.05 | 5.73 | 9.81 | 9.70 | 13.11 | 10.46 | 10.79 | 5.89 | 9.76 |
| LTA4H | 46 | 9.15 | 9.46 | 11.07 | 10.38 | 9.19 | 9.12 | 11.10 | 11.74 | 9.58 | 9.22 | 9.30 | 10.72 | 9.15 | 11.42 | 9.40 |
| LTBP1 | 1 | 7.91 | 5.58 | 18.52 | 21.60 | 4.25 | 4.63 | 11.53 | 11.98 | 6.41 | 7.59 | 6.75 | 20.06 | 4.44 | 11.75 | 7.00 |
| LTF | 1 | 9.25 | 6.93 | 16.40 | 14.17 | 9.41 | 6.88 | 9.44 | 7.26 | 8.74 | 11.51 | 8.09 | 15.29 | 8.15 | 8.35 | 10.12 |
| LTN1 | 6 | 10.22 | 9.82 | 7.45 | 8.29 | 10.76 | 10.86 | 9.08 | 9.09 | 12.63 | 11.81 | 10.02 | 7.87 | 10.81 | 9.09 | 12.22 |
| LTV1 | 3 | 9.21 | 10.04 | 9.63 | 9.92 | 8.71 | 9.75 | 12.97 | 12.96 | 8.34 | 8.46 | 9.63 | 9.77 | 9.23 | 12.97 | 8.40 |
| LUC7L | 7 | 10.40 | 10.78 | 9.95 | 8.77 | 11.26 | 11.16 | 9.66 | 8.64 | 10.01 | 9.37 | 10.59 | 9.36 | 11.21 | 9.15 | 9.69 |
| LUC7L2 | 32 | 9.40 | 10.20 | 9.19 | 9.13 | 9.84 | 9.91 | 10.87 | 13.81 | 8.95 | 8.70 | 9.80 | 9.16 | 9.87 | 12.34 | 8.83 |
| LUC7L3 | 25 | 9.49 | 10.79 | 10.31 | 9.33 | 9.33 | 10.00 | 9.98 | 11.43 | 9.74 | 9.61 | 10.14 | 9.82 | 9.66 | 10.71 | 9.68 |
| LUM | 1 | 7.55 | 6.83 | 16.92 | 18.48 | 8.70 | 8.81 | 7.23 | 6.19 | 9.27 | 10.02 | 7.19 | 17.70 | 8.75 | 6.71 | 9.65 |
| LUZP1 | 29 | 9.83 | 11.26 | 12.06 | 11.65 | 7.63 | 8.18 | 7.69 | 7.40 | 12.36 | 11.93 | 10.55 | 11.86 | 7.90 | 7.55 | 12.15 |
| LXN | 4 | 15.88 | 16.30 | 4.70 | 3.81 | 16.62 | 17.17 | 5.34 | 5.47 | 8.12 | 6.59 | 16.09 | 4.25 | 16.90 | 5.40 | 7.36 |
| LYAR | 12 | 9.30 | 9.61 | 10.39 | 10.50 | 7.83 | 7.91 | 11.50 | 10.98 | 10.54 | 11.45 | 9.45 | 10.44 | 7.87 | 11.24 | 11.00 |
| LYN | 3 | 8.26 | 7.80 | 11.22 | 13.02 | 7.78 | 8.71 | 12.57 | 11.07 | 9.44 | 10.11 | 8.03 | 12.12 | 8.25 | 11.82 | 9.78 |
| LYPLA1 | 9 | 10.09 | 10.26 | 8.85 | 9.04 | 9.89 | 10.33 | 12.81 | 12.02 | 8.76 | 7.94 | 10.18 | 8.95 | 10.11 | 12.42 | 8.35 |
| LYPLA2 | 9 | 9.86 | 10.48 | 9.18 | 9.11 | 9.38 | 8.96 | 9.90 | 12.07 | 10.70 | 10.35 | 10.17 | 9.15 | 9.17 | 10.99 | 10.53 |
| LYPLAL1 | 4 | 8.64 | 10.01 | 6.41 | 6.81 | 13.16 | 14.37 | 10.54 | 10.95 | 9.70 | 9.42 | 9.32 | 6.61 | 13.76 | 10.75 | 9.56 |
| LYRM2 | 3 | 11.00 | 10.10 | 9.19 | 9.32 | 12.22 | 12.19 | 9.16 | 7.97 | 9.51 | 9.33 | 10.55 | 9.26 | 12.21 | 8.56 | 9.42 |
| LYRM4 | 2 | 11.38 | 9.39 | 12.72 | 8.28 | 10.58 | 9.60 | 10.59 | 7.51 | 10.18 | 9.78 | 10.38 | 10.50 | 10.09 | 9.05 | 9.98 |
| LYRM7 | 3 | 10.38 | 9.42 | 8.45 | 9.86 | 8.18 | 8.20 | 12.37 | 11.59 | 10.63 | 10.93 | 9.90 | 9.15 | 8.19 | 11.98 | 10.78 |
| LYSMD1 | 1 | 9.57 | 10.00 | 10.66 | 9.30 | 11.03 | 10.92 | 10.12 | 7.01 | 10.89 | 10.50 | 9.78 | 9.98 | 10.97 | 8.56 | 10.70 |
| LYSMD2 | 4 | 9.15 | 11.60 | 10.17 | 9.41 | 10.24 | 9.87 | 8.98 | 6.96 | 12.01 | 11.63 | 10.37 | 9.79 | 10.05 | 7.97 | 11.82 |
| LZIC | 6 | 10.09 | 10.69 | 10.01 | 9.65 | 8.64 | 8.33 | 10.18 | 11.65 | 10.83 | 9.93 | 10.39 | 9.83 | 8.48 | 10.91 | 10.38 |
| LZTFL1 | 12 | 9.97 | 10.99 | 9.74 | 9.25 | 10.76 | 10.85 | 9.01 | 8.43 | 10.82 | 10.16 | 10.48 | 9.50 | 10.81 | 8.72 | 10.49 |
| LZTR1 | 3 | 9.28 | 8.89 | 10.16 | 11.16 | 7.64 | 9.17 | 10.44 | 14.80 | 9.35 | 9.10 | 9.08 | 10.66 | 8.41 | 12.62 | 9.22 |
| LZTS2 | 3 | 9.82 | 11.21 | 9.77 | 9.34 | 10.45 | 11.30 | 9.05 | 8.41 | 10.67 | 9.97 | 10.51 | 9.56 | 10.88 | 8.73 | 10.32 |
| M6PR | 7 | 6.61 | 4.33 | 8.37 | 6.59 | 7.74 | 6.53 | 7.01 | 38.45 | 7.23 | 7.12 | 5.47 | 7.48 | 7.13 | 22.73 | 7.17 |
| MACF1 | 105 | 9.74 | 10.55 | 12.16 | 11.78 | 6.63 | 6.75 | 8.96 | 8.87 | 12.55 | 12.01 | 10.14 | 11.97 | 6.69 | 8.91 | 12.28 |
| MACROD1 | 4 | 9.59 | 7.82 | 7.75 | 6.11 | 5.27 | 3.77 | 14.48 | 24.30 | 10.05 | 10.87 | 8.70 | 6.93 | 4.52 | 19.39 | 10.46 |
| MAD1L1 | 16 | 9.59 | 10.90 | 10.50 | 9.97 | 9.15 | 9.48 | 11.13 | 10.06 | 9.90 | 9.33 | 10.24 | 10.23 | 9.31 | 10.59 | 9.61 |
| MAD2L1 | 9 | 9.88 | 9.36 | 11.09 | 8.22 | 9.36 | 8.22 | 13.23 | 11.69 | 9.31 | 9.64 | 9.62 | 9.65 | 8.79 | 12.46 | 9.48 |

| | | | | | | | | | | | | | | | | |
|---|---|---|---|---|---|---|---|---|---|---|---|---|---|---|---|---|
| MADD | 12 | 9.46 | 10.19 | 9.20 | 9.49 | 10.52 | 10.89 | 11.76 | 11.06 | 9.03 | 8.39 | 9.83 | 9.34 | 10.71 | 11.41 | 8.71 |
| MAEA | 3 | 11.48 | 12.04 | 7.94 | 6.83 | 11.11 | 12.81 | 10.23 | 9.59 | 9.63 | 8.34 | 11.76 | 7.38 | 11.96 | 9.91 | 8.99 |
| MAF1 | 2 | 7.87 | 9.60 | 10.65 | 8.05 | 11.30 | 10.02 | 12.14 | 10.56 | 10.68 | 9.12 | 8.74 | 9.35 | 10.66 | 11.35 | 9.90 |
| MAFF | 1 | 7.35 | 8.66 | 12.72 | 11.48 | 7.13 | 6.02 | 15.66 | 13.04 | 9.57 | 8.38 | 8.00 | 12.10 | 6.58 | 14.35 | 8.97 |
| MAFG | 6 | 9.11 | 9.88 | 11.49 | 10.31 | 8.55 | 8.53 | 12.04 | 13.00 | 8.76 | 8.34 | 9.50 | 10.90 | 8.54 | 12.52 | 8.55 |
| MAFK | 5 | 7.64 | 8.09 | 8.97 | 9.23 | 8.25 | 8.62 | 17.85 | 16.48 | 7.59 | 7.28 | 7.86 | 9.10 | 8.44 | 17.17 | 7.43 |
| MAGED2 | 29 | 9.62 | 10.96 | 10.53 | 10.30 | 10.61 | 11.02 | 10.97 | 10.07 | 8.13 | 7.79 | 10.29 | 10.42 | 10.81 | 10.52 | 7.96 |
| MAGOHB | 17 | 8.19 | 10.80 | 9.91 | 8.94 | 10.13 | 10.52 | 11.53 | 11.49 | 9.16 | 9.34 | 9.49 | 9.42 | 10.32 | 11.51 | 9.25 |
| MAGT1 | 8 | 12.47 | 7.16 | 8.54 | 10.73 | 8.94 | 8.42 | 9.48 | 9.98 | 10.64 | 13.62 | 9.82 | 9.64 | 8.68 | 9.73 | 12.13 |
| MAK | 2 | 11.41 | 11.12 | 10.05 | 10.79 | 10.87 | 10.36 | 8.06 | 6.82 | 9.84 | 10.68 | 11.27 | 10.42 | 10.62 | 7.44 | 10.26 |
| MAK16 | 5 | 9.42 | 8.91 | 10.58 | 8.80 | 8.88 | 8.75 | 13.82 | 11.65 | 10.08 | 9.12 | 9.16 | 9.69 | 8.82 | 12.73 | 9.60 |
| MALSU1 | 2 | 11.28 | 7.88 | 12.56 | 8.77 | 9.57 | 8.37 | 14.16 | 7.91 | 10.26 | 9.24 | 9.58 | 10.67 | 8.97 | 11.03 | 9.75 |
| MALT1 | 6 | 11.86 | 11.97 | 11.33 | 9.67 | 9.60 | 8.80 | 6.34 | 5.45 | 12.99 | 11.99 | 11.92 | 10.50 | 9.20 | 5.89 | 12.49 |
| MAML1 | 4 | 8.83 | 9.61 | 10.69 | 10.81 | 10.13 | 9.99 | 10.96 | 9.77 | 9.52 | 9.70 | 9.22 | 10.75 | 10.06 | 10.37 | 9.61 |
| MAN1A2 | 2 | 10.80 | 7.64 | 9.96 | 11.37 | 9.45 | 7.90 | 10.50 | 10.59 | 10.59 | 11.20 | 9.22 | 10.66 | 8.67 | 10.55 | 10.89 |
| MAN1B1 | 4 | 10.41 | 6.83 | 8.06 | 10.49 | 9.68 | 8.74 | 8.73 | 9.01 | 13.04 | 15.01 | 8.62 | 9.28 | 9.21 | 8.87 | 14.02 |
| MAN2A1 | 5 | 12.19 | 9.08 | 6.65 | 7.69 | 13.68 | 12.34 | 10.62 | 10.32 | 8.27 | 9.16 | 10.64 | 7.17 | 13.01 | 10.47 | 8.72 |
| MAN2B1 | 6 | 11.46 | 11.76 | 7.24 | 7.00 | 12.47 | 13.00 | 9.35 | 8.52 | 9.82 | 9.38 | 11.61 | 7.12 | 12.73 | 8.94 | 9.60 |
| MAN2C1 | 2 | 7.69 | 9.28 | 8.44 | 10.37 | 11.05 | 12.61 | 10.60 | 9.02 | 10.36 | 10.59 | 8.48 | 9.41 | 11.83 | 9.81 | 10.47 |
| MANBA | 2 | 9.62 | 9.94 | 9.87 | 9.29 | 10.33 | 9.98 | 9.76 | 11.66 | 9.88 | 9.65 | 9.78 | 9.58 | 10.16 | 10.71 | 9.77 |
| MANBAL | 2 | 10.89 | 6.33 | 11.66 | 15.45 | 8.13 | 7.26 | 9.72 | 8.84 | 9.51 | 12.20 | 8.61 | 13.55 | 7.69 | 9.28 | 10.86 |
| MANF | 14 | 9.61 | 9.88 | 8.09 | 7.38 | 12.22 | 12.34 | 9.17 | 13.89 | 8.65 | 8.76 | 9.75 | 7.74 | 12.28 | 11.53 | 8.71 |
| MAOB | 5 | 10.22 | 10.16 | 7.38 | 8.61 | 5.10 | 5.98 | 22.10 | 20.22 | 5.37 | 4.84 | 10.19 | 8.00 | 5.54 | 21.16 | 5.11 |
| MAP1A | 12 | 11.87 | 13.65 | 5.34 | 5.43 | 15.04 | 15.50 | 7.75 | 6.81 | 9.72 | 8.89 | 12.76 | 5.39 | 15.27 | 7.28 | 9.31 |
| MAP1B | 127 | 10.82 | 12.05 | 15.21 | 14.18 | 6.21 | 6.34 | 6.14 | 5.53 | 11.93 | 11.59 | 11.43 | 14.69 | 6.28 | 5.83 | 11.76 |
| MAP1LC3B2 | 4 | 11.71 | 11.74 | 9.60 | 9.08 | 9.92 | 9.88 | 6.42 | 5.76 | 12.99 | 12.89 | 11.72 | 9.34 | 9.90 | 6.09 | 12.94 |
| MAP1S | 11 | 8.37 | 10.06 | 12.58 | 12.72 | 9.31 | 9.45 | 9.88 | 9.41 | 9.05 | 9.17 | 9.21 | 12.65 | 9.38 | 9.65 | 9.11 |
| MAP2 | 8 | 17.13 | 18.86 | 13.67 | 12.90 | 8.72 | 8.97 | 4.44 | 4.02 | 5.83 | 5.47 | 17.99 | 13.28 | 8.85 | 4.23 | 5.65 |
| MAP2K1 | 14 | 9.58 | 9.56 | 9.50 | 9.10 | 10.30 | 10.44 | 9.89 | 8.84 | 11.57 | 11.22 | 9.57 | 9.30 | 10.37 | 9.36 | 11.40 |
| MAP2K2 | 30 | 8.36 | 10.49 | 11.31 | 10.35 | 10.45 | 10.44 | 9.43 | 8.83 | 9.77 | 10.57 | 9.42 | 10.83 | 10.44 | 9.13 | 10.17 |
| MAP2K3 | 14 | 9.38 | 9.69 | 13.88 | 12.55 | 8.60 | 8.71 | 7.96 | 8.70 | 10.45 | 10.08 | 9.54 | 13.22 | 8.65 | 8.33 | 10.27 |
| MAP2K4 | 6 | 9.76 | 10.95 | 11.69 | 11.35 | 9.93 | 10.98 | 9.27 | 8.57 | 9.15 | 8.33 | 10.36 | 11.52 | 10.46 | 8.92 | 8.74 |
| MAP2K6 | 4 | 9.62 | 10.59 | 10.08 | 9.21 | 13.07 | 13.03 | 12.37 | 10.75 | 5.58 | 5.70 | 10.11 | 9.65 | 13.05 | 11.56 | 5.64 |
| MAP2K7 | 6 | 8.89 | 10.05 | 9.33 | 9.96 | 10.82 | 12.04 | 10.24 | 10.62 | 9.23 | 8.83 | 9.47 | 9.65 | 11.43 | 10.43 | 9.03 |
| MAP3K1 | 1 | 8.17 | 11.00 | 13.37 | 13.50 | 8.81 | 10.06 | 7.82 | 4.52 | 11.03 | 11.74 | 9.58 | 13.43 | 9.43 | 6.17 | 11.38 |
| MAP3K2 | 7 | 10.25 | 10.93 | 9.74 | 9.92 | 10.72 | 10.48 | 9.03 | 8.61 | 10.70 | 9.62 | 10.59 | 9.83 | 10.60 | 8.82 | 10.16 |
| MAP3K7 | 8 | 10.08 | 10.99 | 10.23 | 11.48 | 9.40 | 9.97 | 9.39 | 9.44 | 9.56 | 9.46 | 10.53 | 10.85 | 9.69 | 9.42 | 9.51 |
| MAP4 | 106 | 9.62 | 10.52 | 11.23 | 10.19 | 10.50 | 10.70 | 8.23 | 7.72 | 10.89 | 10.40 | 10.07 | 10.71 | 10.60 | 7.98 | 10.65 |
| MAP4K2 | 2 | 8.40 | 9.65 | 9.26 | 10.04 | 12.52 | 14.15 | 9.33 | 7.11 | 9.42 | 10.12 | 9.03 | 9.65 | 13.33 | 8.22 | 9.77 |
| MAP4K3 | 1 | 7.17 | 9.31 | 15.93 | 13.38 | 7.72 | 9.54 | 8.71 | 9.25 | 8.81 | 10.16 | 8.24 | 14.66 | 8.63 | 8.98 | 9.49 |
| MAP4K4 | 6 | 9.37 | 11.02 | 10.66 | 9.89 | 10.24 | 10.38 | 9.25 | 7.35 | 11.23 | 10.61 | 10.20 | 10.27 | 10.31 | 8.30 | 10.92 |
| MAP4K5 | 2 | 12.40 | 13.25 | 10.47 | 9.99 | 9.79 | 10.69 | 9.73 | 8.76 | 8.17 | 6.76 | 12.83 | 10.23 | 10.24 | 9.24 | 7.47 |

| | | | | | | | | | | | | | | | | |
|---|---|---|---|---|---|---|---|---|---|---|---|---|---|---|---|---|
| MAP7 | 33 | 12.66 | 13.83 | 7.30 | 6.06 | 13.88 | 13.70 | 8.38 | 7.22 | 8.77 | 8.19 | 13.25 | 6.68 | 13.79 | 7.80 | 8.48 |
| MAP7D1 | 24 | 10.28 | 11.67 | 9.41 | 9.69 | 8.15 | 8.33 | 8.58 | 8.69 | 12.60 | 12.61 | 10.98 | 9.55 | 8.24 | 8.63 | 12.60 |
| MAP7D2 | 8 | 12.22 | 13.48 | 5.08 | 4.69 | 9.18 | 9.36 | 13.16 | 11.72 | 10.56 | 10.54 | 12.85 | 4.88 | 9.27 | 12.44 | 10.55 |
| MAP7D3 | 14 | 9.77 | 10.76 | 9.85 | 9.35 | 11.26 | 11.63 | 10.29 | 9.22 | 9.16 | 8.71 | 10.27 | 9.60 | 11.44 | 9.75 | 8.93 |
| MAPK1 | 22 | 9.98 | 11.40 | 11.59 | 11.12 | 9.23 | 9.35 | 8.37 | 8.84 | 9.92 | 10.20 | 10.69 | 11.36 | 9.29 | 8.60 | 10.06 |
| MAPK14 | 7 | 9.54 | 10.39 | 10.78 | 9.05 | 11.08 | 11.15 | 10.43 | 8.59 | 9.42 | 9.57 | 9.97 | 9.92 | 11.11 | 9.51 | 9.50 |
| MAPK3 | 9 | 9.72 | 10.39 | 11.34 | 11.19 | 10.71 | 10.84 | 8.59 | 8.34 | 9.45 | 9.44 | 10.05 | 11.27 | 10.77 | 8.46 | 9.45 |
| MAPK7 | 2 | 11.11 | 11.78 | 10.57 | 10.65 | 9.29 | 10.60 | 7.12 | 7.51 | 10.96 | 10.40 | 11.45 | 10.61 | 9.95 | 7.32 | 10.68 |
| MAPK8 | 2 | 9.48 | 9.73 | 10.94 | 11.25 | 8.92 | 9.60 | 11.33 | 10.63 | 9.25 | 8.85 | 9.60 | 11.10 | 9.26 | 10.98 | 9.05 |
| MAPK9 | 2 | 9.89 | 11.28 | 10.14 | 9.56 | 11.66 | 11.84 | 10.89 | 9.34 | 8.12 | 7.27 | 10.59 | 9.85 | 11.75 | 10.12 | 7.70 |
| MAPKAP1 | 5 | 10.22 | 9.38 | 8.96 | 8.46 | 10.75 | 10.93 | 10.86 | 10.17 | 10.25 | 10.01 | 9.80 | 8.71 | 10.84 | 10.52 | 10.13 |
| MAPKAPK2 | 20 | 8.21 | 10.14 | 9.84 | 9.75 | 11.14 | 11.45 | 10.94 | 10.28 | 9.33 | 8.92 | 9.17 | 9.80 | 11.29 | 10.61 | 9.12 |
| MAPKAPK3 | 7 | 7.97 | 8.98 | 11.11 | 11.00 | 9.70 | 9.88 | 11.08 | 10.48 | 9.41 | 10.41 | 8.47 | 11.05 | 9.79 | 10.78 | 9.91 |
| MAPRE1 | 27 | 9.35 | 10.05 | 12.04 | 11.21 | 10.05 | 10.17 | 8.44 | 8.39 | 10.34 | 9.96 | 9.70 | 11.63 | 10.11 | 8.42 | 10.15 |
| MAPRE2 | 11 | 8.88 | 9.55 | 13.08 | 11.54 | 11.27 | 11.08 | 8.05 | 7.09 | 9.87 | 9.60 | 9.22 | 12.31 | 11.17 | 7.57 | 9.73 |
| MAPRE3 | 6 | 9.18 | 10.10 | 12.39 | 11.72 | 9.14 | 9.75 | 8.39 | 7.60 | 11.18 | 10.54 | 9.64 | 12.06 | 9.45 | 7.99 | 10.86 |
| MARC1 | 2 | 12.33 | 9.77 | 9.44 | 11.77 | 11.40 | 9.62 | 6.86 | 5.58 | 11.04 | 12.20 | 11.05 | 10.60 | 10.51 | 6.22 | 11.62 |
| MARC5 | 1 | 12.38 | 7.13 | 7.75 | 11.93 | 7.65 | 7.61 | 11.27 | 9.42 | 10.68 | 14.18 | 9.75 | 9.84 | 7.63 | 10.34 | 12.43 |
| MARCKS | 22 | 10.06 | 10.84 | 16.77 | 17.81 | 4.37 | 4.79 | 8.91 | 6.29 | 9.94 | 10.23 | 10.45 | 17.29 | 4.58 | 7.60 | 10.09 |
| MARCKSL1 | 6 | 11.07 | 13.33 | 10.75 | 15.34 | 7.50 | 10.32 | 14.92 | 7.39 | 4.37 | 5.03 | 12.20 | 13.04 | 8.91 | 11.15 | 4.70 |
| MARK1 | 7 | 8.95 | 10.10 | 10.24 | 8.56 | 12.11 | 11.30 | 9.93 | 10.07 | 9.63 | 9.12 | 9.53 | 9.40 | 11.70 | 10.00 | 9.37 |
| MARK2 | 6 | 8.69 | 9.54 | 12.30 | 8.95 | 10.00 | 9.45 | 9.80 | 10.61 | 10.50 | 10.16 | 9.12 | 10.62 | 9.73 | 10.20 | 10.33 |
| MARK3 | 3 | 9.14 | 10.31 | 9.77 | 9.04 | 10.38 | 11.71 | 11.15 | 10.35 | 9.45 | 8.70 | 9.72 | 9.40 | 11.05 | 10.75 | 9.07 |
| MARS | 42 | 8.96 | 9.35 | 9.81 | 9.59 | 10.48 | 10.59 | 10.96 | 14.29 | 8.12 | 7.86 | 9.15 | 9.70 | 10.54 | 12.62 | 7.99 |
| MARS2 | 1 | 11.57 | 8.30 | 8.69 | 9.41 | 11.17 | 11.09 | 9.73 | 9.27 | 10.05 | 10.72 | 9.94 | 9.05 | 11.13 | 9.50 | 10.38 |
| MAST2 | 2 | 10.67 | 10.50 | 10.38 | 9.87 | 10.96 | 11.15 | 10.14 | 9.18 | 8.67 | 8.47 | 10.58 | 10.13 | 11.06 | 9.66 | 8.57 |
| MASTL | 3 | 8.20 | 9.63 | 11.07 | 10.79 | 9.93 | 9.35 | 10.78 | 9.55 | 9.98 | 10.72 | 8.91 | 10.93 | 9.64 | 10.17 | 10.35 |
| MAT2A | 31 | 10.20 | 10.82 | 9.18 | 7.82 | 10.02 | 10.00 | 11.97 | 11.44 | 9.59 | 8.97 | 10.51 | 8.50 | 10.01 | 11.70 | 9.28 |
| MAT2B | 23 | 10.25 | 11.00 | 6.95 | 6.84 | 9.92 | 10.20 | 12.55 | 17.85 | 7.46 | 6.96 | 10.63 | 6.89 | 10.06 | 15.20 | 7.21 |
| MATR3 | 63 | 11.12 | 12.40 | 12.05 | 10.89 | 12.56 | 12.74 | 13.32 | 12.91 | 1.17 | 0.84 | 11.76 | 11.47 | 12.65 | 13.11 | 1.01 |
| MAU2 | 3 | 8.70 | 9.84 | 9.67 | 8.35 | 10.71 | 10.34 | 12.24 | 13.52 | 8.42 | 8.20 | 9.27 | 9.01 | 10.53 | 12.88 | 8.31 |
| MAVS | 4 | 11.35 | 7.46 | 8.70 | 12.97 | 8.94 | 9.00 | 10.38 | 9.61 | 10.77 | 10.82 | 9.41 | 10.83 | 8.97 | 9.99 | 10.80 |
| MAX | 4 | 8.89 | 11.04 | 11.65 | 10.02 | 9.44 | 10.72 | 11.70 | 5.33 | 10.45 | 10.76 | 9.96 | 10.83 | 10.08 | 8.51 | 10.61 |
| MAZ | 1 | 10.70 | 11.22 | 11.97 | 5.00 | 13.02 | 7.96 | 12.67 | 4.29 | 12.67 | 10.51 | 10.96 | 8.48 | 10.49 | 8.48 | 11.59 |
| MB21D1 | 6 | 10.08 | 9.67 | 8.95 | 8.20 | 10.31 | 9.96 | 11.68 | 10.83 | 10.62 | 9.70 | 9.87 | 8.57 | 10.13 | 11.26 | 10.16 |
| MBD1 | 1 | 8.67 | 8.88 | 14.59 | 11.17 | 10.30 | 10.46 | 9.01 | 6.66 | 9.40 | 10.87 | 8.77 | 12.88 | 10.38 | 7.83 | 10.14 |
| MBD2 | 11 | 9.45 | 10.64 | 11.49 | 10.86 | 11.82 | 11.52 | 9.17 | 8.18 | 8.42 | 8.44 | 10.05 | 11.18 | 11.67 | 8.67 | 8.43 |
| MBD3 | 6 | 8.93 | 9.40 | 8.85 | 8.46 | 8.70 | 8.66 | 10.70 | 19.30 | 8.58 | 8.44 | 9.16 | 8.65 | 8.68 | 15.00 | 8.51 |
| MBIP | 1 | 11.25 | 11.17 | 11.73 | 8.44 | 6.54 | 7.57 | 11.73 | 10.40 | 11.21 | 9.96 | 11.21 | 10.09 | 7.06 | 11.07 | 10.58 |
| MBLAC2 | 2 | 9.67 | 9.55 | 9.57 | 8.56 | 8.23 | 8.55 | 13.59 | 12.87 | 10.07 | 9.33 | 9.61 | 9.07 | 8.39 | 13.23 | 9.70 |
| MBNL1 | 6 | 8.84 | 11.04 | 10.17 | 9.04 | 9.35 | 9.95 | 8.90 | 13.57 | 9.72 | 9.43 | 9.94 | 9.61 | 9.65 | 11.23 | 9.57 |
| MBOAT7 | 4 | 10.92 | 6.23 | 5.13 | 6.92 | 13.48 | 12.06 | 9.12 | 9.31 | 10.79 | 16.03 | 8.58 | 6.03 | 12.77 | 9.22 | 13.41 |

| | | | | | | | | | | | | | | | | |
|---|---|---|---|---|---|---|---|---|---|---|---|---|---|---|---|---|
| MCAM | 27 | 10.66 | 8.07 | 9.02 | 10.86 | 10.64 | 10.56 | 6.12 | 6.57 | 13.89 | 13.61 | 9.36 | 9.94 | 10.60 | 6.35 | 13.75 |
| MCAT | 7 | 10.52 | 10.45 | 11.88 | 13.23 | 7.45 | 8.43 | 10.19 | 7.54 | 9.95 | 10.37 | 10.49 | 12.55 | 7.94 | 8.86 | 10.16 |
| MCC | 2 | 10.66 | 10.38 | 13.19 | 13.47 | 9.93 | 9.96 | 5.86 | 5.59 | 11.01 | 9.96 | 10.52 | 13.33 | 9.95 | 5.72 | 10.48 |
| MCCC1 | 5 | 13.65 | 11.35 | 8.79 | 10.60 | 8.65 | 8.39 | 6.92 | 6.85 | 12.36 | 12.43 | 12.50 | 9.69 | 8.52 | 6.89 | 12.39 |
| MCCC2 | 13 | 11.13 | 9.37 | 8.56 | 9.40 | 9.85 | 9.90 | 13.42 | 11.83 | 8.17 | 8.39 | 10.25 | 8.98 | 9.87 | 12.62 | 8.28 |
| MCEE | 4 | 10.23 | 8.81 | 8.88 | 9.31 | 11.88 | 11.25 | 10.49 | 9.16 | 10.32 | 9.66 | 9.52 | 9.09 | 11.57 | 9.83 | 9.99 |
| MCFD2 | 2 | 11.07 | 10.34 | 11.53 | 11.67 | 10.52 | 8.83 | 10.95 | 4.17 | 11.21 | 9.71 | 10.70 | 11.60 | 9.67 | 7.56 | 10.46 |
| MCL1 | 2 | 8.78 | 10.02 | 10.62 | 11.65 | 9.63 | 11.02 | 9.37 | 8.33 | 10.01 | 10.57 | 9.40 | 11.13 | 10.33 | 8.85 | 10.29 |
| MCM2 | 57 | 9.46 | 10.16 | 9.54 | 9.04 | 10.47 | 10.43 | 9.14 | 13.28 | 9.34 | 9.14 | 9.81 | 9.29 | 10.45 | 11.21 | 9.24 |
| MCM3 | 64 | 9.31 | 9.94 | 11.19 | 9.23 | 10.32 | 9.76 | 9.48 | 11.83 | 9.71 | 9.23 | 9.62 | 10.21 | 10.04 | 10.66 | 9.47 |
| MCM3AP | 8 | 10.04 | 11.49 | 10.54 | 13.70 | 7.82 | 8.48 | 7.71 | 8.19 | 10.74 | 11.31 | 10.76 | 12.12 | 8.15 | 7.95 | 11.02 |
| MCM4 | 58 | 8.10 | 10.05 | 10.71 | 9.29 | 10.79 | 10.21 | 9.62 | 12.57 | 9.37 | 9.29 | 9.07 | 10.00 | 10.50 | 11.10 | 9.33 |
| MCM5 | 60 | 9.44 | 10.11 | 9.48 | 8.25 | 10.09 | 9.97 | 9.07 | 14.91 | 9.42 | 9.25 | 9.77 | 8.87 | 10.03 | 11.99 | 9.34 |
| MCM6 | 64 | 9.46 | 10.24 | 10.53 | 8.68 | 11.12 | 10.49 | 9.67 | 10.67 | 9.73 | 9.41 | 9.85 | 9.60 | 10.80 | 10.17 | 9.57 |
| MCM7 | 54 | 9.52 | 10.15 | 9.97 | 9.18 | 10.13 | 10.58 | 9.91 | 11.34 | 9.76 | 9.46 | 9.83 | 9.57 | 10.35 | 10.63 | 9.61 |
| MCMBP | 10 | 8.23 | 9.00 | 10.45 | 10.20 | 10.05 | 10.02 | 11.29 | 11.28 | 9.86 | 9.60 | 8.61 | 10.33 | 10.04 | 11.29 | 9.73 |
| MCRS1 | 2 | 9.83 | 10.25 | 11.56 | 11.21 | 9.65 | 9.68 | 10.05 | 8.65 | 9.84 | 9.27 | 10.04 | 11.39 | 9.67 | 9.35 | 9.55 |
| MCTP2 | 1 | 6.21 | 7.08 | 11.38 | 14.29 | 13.04 | 11.73 | 8.57 | 6.74 | 9.59 | 11.37 | 6.65 | 12.83 | 12.39 | 7.65 | 10.48 |
| MCTS1 | 11 | 7.55 | 7.64 | 9.26 | 6.84 | 10.94 | 10.16 | 11.28 | 20.14 | 8.24 | 7.97 | 7.59 | 8.05 | 10.55 | 15.71 | 8.10 |
| MCU | 8 | 11.61 | 7.57 | 7.99 | 11.91 | 10.92 | 9.29 | 7.64 | 7.44 | 11.67 | 13.94 | 9.59 | 9.95 | 10.11 | 7.54 | 12.81 |
| MCUR1 | 6 | 10.98 | 9.98 | 8.98 | 9.43 | 10.31 | 10.10 | 10.51 | 11.83 | 8.62 | 9.26 | 10.48 | 9.21 | 10.21 | 11.17 | 8.94 |
| MDC1 | 36 | 9.64 | 10.34 | 12.06 | 12.41 | 10.22 | 10.66 | 8.65 | 7.84 | 9.15 | 9.03 | 9.99 | 12.23 | 10.44 | 8.25 | 9.09 |
| MDH1 | 37 | 10.03 | 11.84 | 10.72 | 10.35 | 9.25 | 9.48 | 8.38 | 9.72 | 10.26 | 9.96 | 10.94 | 10.54 | 9.36 | 9.05 | 10.11 |
| MDH2 | 53 | 9.63 | 8.58 | 9.58 | 8.62 | 7.89 | 7.40 | 10.76 | 17.12 | 10.08 | 10.34 | 9.11 | 9.10 | 7.64 | 13.94 | 10.21 |
| MDN1 | 45 | 9.75 | 8.90 | 9.72 | 11.02 | 7.41 | 7.82 | 10.76 | 10.40 | 12.00 | 12.23 | 9.32 | 10.37 | 7.61 | 10.58 | 12.12 |
| MDP1 | 4 | 8.44 | 9.25 | 9.76 | 8.87 | 12.77 | 13.38 | 11.29 | 9.61 | 8.61 | 8.02 | 8.84 | 9.32 | 13.07 | 10.45 | 8.31 |
| ME1 | 18 | 12.12 | 12.38 | 11.00 | 9.60 | 7.91 | 7.67 | 9.39 | 8.21 | 11.00 | 10.72 | 12.25 | 10.30 | 7.79 | 8.80 | 10.86 |
| ME2 | 14 | 12.60 | 11.28 | 8.86 | 9.95 | 11.20 | 11.02 | 6.69 | 6.18 | 10.82 | 11.40 | 11.94 | 9.40 | 11.11 | 6.43 | 11.11 |
| ME3 | 3 | 11.28 | 10.66 | 10.49 | 11.25 | 9.01 | 9.50 | 9.26 | 9.05 | 9.58 | 9.92 | 10.97 | 10.87 | 9.26 | 9.16 | 9.75 |
| MEA1 | 3 | 9.81 | 12.68 | 11.76 | 11.73 | 9.53 | 11.31 | 8.80 | 3.71 | 9.73 | 10.94 | 11.24 | 11.75 | 10.42 | 6.26 | 10.34 |
| MEAF6 | 2 | 6.69 | 11.14 | 8.99 | 8.73 | 10.00 | 12.08 | 13.05 | 8.90 | 9.07 | 11.34 | 8.92 | 8.86 | 11.04 | 10.97 | 10.20 |
| MECP2 | 5 | 10.01 | 8.91 | 9.44 | 11.16 | 10.84 | 10.16 | 12.50 | 11.10 | 6.89 | 9.00 | 9.46 | 10.30 | 10.50 | 11.80 | 7.95 |
| MECR | 1 | 9.89 | 8.09 | 16.35 | 5.06 | 13.94 | 8.20 | 14.44 | 7.28 | 8.31 | 8.43 | 8.99 | 10.71 | 11.07 | 10.86 | 8.37 |
| MED1 | 12 | 9.73 | 10.82 | 11.62 | 11.40 | 9.10 | 9.42 | 10.04 | 8.99 | 9.64 | 9.24 | 10.28 | 11.51 | 9.26 | 9.52 | 9.44 |
| MED10 | 4 | 10.32 | 11.51 | 9.13 | 9.79 | 12.46 | 12.75 | 7.99 | 5.88 | 10.12 | 10.05 | 10.91 | 9.46 | 12.60 | 6.94 | 10.09 |
| MED12 | 14 | 8.63 | 9.52 | 8.75 | 9.18 | 11.69 | 11.36 | 10.65 | 11.70 | 9.21 | 9.31 | 9.08 | 8.96 | 11.53 | 11.17 | 9.26 |
| MED13 | 1 | 10.86 | 10.89 | 14.55 | 8.94 | 8.22 | 7.83 | 9.62 | 7.61 | 11.22 | 10.26 | 10.87 | 11.75 | 8.03 | 8.61 | 10.74 |
| MED13L | 1 | 8.06 | 9.00 | 13.18 | 12.44 | 9.30 | 8.26 | 12.73 | 9.39 | 9.70 | 7.92 | 8.53 | 12.81 | 8.78 | 11.06 | 8.81 |
| MED14 | 2 | 9.44 | 8.78 | 10.14 | 10.09 | 9.53 | 8.32 | 12.40 | 11.29 | 10.90 | 9.11 | 9.11 | 10.11 | 8.93 | 11.85 | 10.01 |
| MED15 | 4 | 11.55 | 10.94 | 10.56 | 10.56 | 9.39 | 9.95 | 9.21 | 8.58 | 10.16 | 9.11 | 11.25 | 10.56 | 9.67 | 8.89 | 9.64 |
| MED16 | 6 | 9.54 | 9.16 | 9.96 | 10.73 | 9.30 | 9.40 | 11.17 | 11.98 | 9.32 | 9.44 | 9.35 | 10.35 | 9.35 | 11.57 | 9.38 |
| MED17 | 5 | 9.02 | 9.51 | 11.09 | 10.85 | 8.51 | 8.32 | 13.14 | 11.68 | 9.04 | 8.83 | 9.26 | 10.97 | 8.42 | 12.41 | 8.94 |

| | | | | | | | | | | | | | | | | |
|---|---|---|---|---|---|---|---|---|---|---|---|---|---|---|---|---|
| MED18 | 4 | 9.80 | 11.60 | 8.42 | 8.40 | 10.60 | 11.26 | 9.62 | 10.53 | 9.67 | 10.09 | 10.70 | 8.41 | 10.93 | 10.08 | 9.88 |
| MED20 | 3 | 12.27 | 10.55 | 8.48 | 8.20 | 9.76 | 10.40 | 9.77 | 10.63 | 10.73 | 9.20 | 11.41 | 8.34 | 10.08 | 10.20 | 9.97 |
| MED22 | 4 | 10.55 | 10.50 | 9.90 | 9.96 | 10.17 | 9.63 | 12.67 | 7.91 | 9.92 | 8.79 | 10.53 | 9.93 | 9.90 | 10.29 | 9.35 |
| MED23 | 5 | 10.56 | 10.30 | 8.92 | 9.40 | 9.01 | 10.02 | 10.21 | 11.50 | 10.24 | 9.84 | 10.43 | 9.16 | 9.51 | 10.86 | 10.04 |
| MED24 | 2 | 9.31 | 9.03 | 11.11 | 12.13 | 8.94 | 8.93 | 11.60 | 11.29 | 8.56 | 9.10 | 9.17 | 11.62 | 8.94 | 11.44 | 8.83 |
| MED26 | 1 | 8.61 | 9.87 | 10.26 | 9.21 | 8.22 | 9.59 | 13.83 | 12.41 | 9.73 | 8.27 | 9.24 | 9.73 | 8.91 | 13.12 | 9.00 |
| MED28 | 1 | 8.29 | 8.21 | 7.65 | 9.54 | 9.17 | 7.42 | 14.30 | 11.97 | 12.73 | 10.73 | 8.25 | 8.59 | 8.29 | 13.13 | 11.73 |
| MED29 | 5 | 10.13 | 11.35 | 9.20 | 8.50 | 9.40 | 9.74 | 12.27 | 10.58 | 9.75 | 9.07 | 10.74 | 8.85 | 9.57 | 11.43 | 9.41 |
| MED31 | 1 | 9.56 | 11.37 | 12.17 | 9.57 | 9.25 | 9.10 | 10.58 | 10.29 | 9.10 | 8.99 | 10.46 | 10.87 | 9.18 | 10.44 | 9.05 |
| MED4 | 6 | 9.87 | 11.01 | 10.59 | 10.94 | 9.87 | 10.06 | 10.70 | 8.99 | 9.13 | 8.85 | 10.44 | 10.77 | 9.96 | 9.85 | 8.99 |
| MED6 | 4 | 9.26 | 9.43 | 10.05 | 10.63 | 7.74 | 9.15 | 13.36 | 12.63 | 8.79 | 8.96 | 9.35 | 10.34 | 8.44 | 13.00 | 8.88 |
| MED7 | 1 | 9.47 | 10.29 | 11.28 | 9.79 | 9.14 | 9.18 | 11.24 | 10.50 | 10.34 | 8.77 | 9.88 | 10.53 | 9.16 | 10.87 | 9.56 |
| MED8 | 5 | 8.47 | 9.78 | 10.44 | 9.88 | 8.07 | 8.71 | 12.73 | 13.60 | 9.42 | 8.89 | 9.12 | 10.16 | 8.39 | 13.16 | 9.16 |
| MED9 | 2 | 9.47 | 12.21 | 11.02 | 12.37 | 8.18 | 8.33 | 10.52 | 4.69 | 9.33 | 13.87 | 10.84 | 11.69 | 8.26 | 7.61 | 11.60 |
| MEF2A | 3 | 7.83 | 9.67 | 13.90 | 13.83 | 10.00 | 9.64 | 9.48 | 8.27 | 8.88 | 8.50 | 8.75 | 13.87 | 9.82 | 8.87 | 8.69 |
| MEF2BNB | 1 | 8.93 | 11.42 | 11.37 | 11.41 | 9.86 | 8.86 | 10.05 | 5.44 | 11.42 | 11.24 | 10.17 | 11.39 | 9.36 | 7.75 | 11.33 |
| MEF2D | 6 | 9.12 | 10.48 | 12.08 | 11.79 | 9.76 | 10.34 | 9.12 | 8.43 | 9.64 | 9.24 | 9.80 | 11.93 | 10.05 | 8.78 | 9.44 |
| MEIS1 | 3 | 11.40 | 10.81 | 8.37 | 7.55 | 13.27 | 13.83 | 9.57 | 9.67 | 8.23 | 7.31 | 11.10 | 7.96 | 13.55 | 9.62 | 7.77 |
| MELK | 2 | 9.94 | 11.77 | 10.14 | 10.89 | 6.83 | 8.56 | 7.77 | 8.46 | 12.27 | 13.36 | 10.86 | 10.52 | 7.69 | 8.12 | 12.82 |
| MEMO1 | 4 | 9.07 | 8.88 | 12.52 | 6.52 | 11.56 | 9.75 | 10.78 | 7.40 | 11.95 | 11.57 | 8.98 | 9.52 | 10.66 | 9.09 | 11.76 |
| MEN1 | 5 | 9.03 | 9.61 | 10.96 | 8.92 | 9.83 | 9.36 | 11.92 | 9.60 | 10.65 | 10.14 | 9.32 | 9.94 | 9.59 | 10.76 | 10.40 |
| MEPCE | 11 | 8.50 | 8.91 | 11.81 | 9.10 | 11.50 | 11.00 | 11.41 | 9.20 | 9.54 | 9.04 | 8.70 | 10.46 | 11.25 | 10.30 | 9.29 |
| MESDC2 | 16 | 9.26 | 9.44 | 9.03 | 9.23 | 12.52 | 13.30 | 9.55 | 8.78 | 9.48 | 9.40 | 9.35 | 9.13 | 12.91 | 9.17 | 9.44 |
| MET | 11 | 11.67 | 9.66 | 6.45 | 8.54 | 13.49 | 13.86 | 5.96 | 5.67 | 11.77 | 12.93 | 10.67 | 7.49 | 13.68 | 5.81 | 12.35 |
| METAP1 | 13 | 9.28 | 9.21 | 11.62 | 8.20 | 10.69 | 10.16 | 11.86 | 10.55 | 9.13 | 9.29 | 9.24 | 9.91 | 10.43 | 11.21 | 9.21 |
| METAP2 | 13 | 9.93 | 11.06 | 9.32 | 9.19 | 12.48 | 12.39 | 9.59 | 10.32 | 8.12 | 7.60 | 10.49 | 9.26 | 12.44 | 9.95 | 7.86 |
| METTL1 | 9 | 8.83 | 9.40 | 11.26 | 8.20 | 9.77 | 9.07 | 11.71 | 10.52 | 10.47 | 10.76 | 9.12 | 9.73 | 9.42 | 11.12 | 10.62 |
| METTL10 | 1 | 7.96 | 12.73 | 10.17 | 9.82 | 8.43 | 8.34 | 13.43 | 10.54 | 10.38 | 8.21 | 10.34 | 9.99 | 8.38 | 11.98 | 9.30 |
| METTL13 | 9 | 8.34 | 9.29 | 11.66 | 11.35 | 11.05 | 11.08 | 9.23 | 8.49 | 10.13 | 9.39 | 8.81 | 11.50 | 11.06 | 8.86 | 9.76 |
| METTL14 | 5 | 9.61 | 9.90 | 10.27 | 9.49 | 9.36 | 9.63 | 11.73 | 10.08 | 10.09 | 9.85 | 9.75 | 9.88 | 9.49 | 10.91 | 9.97 |
| METTL15 | 2 | 9.72 | 9.85 | 9.53 | 9.74 | 9.89 | 10.23 | 12.03 | 10.36 | 9.27 | 9.39 | 9.79 | 9.63 | 10.06 | 11.20 | 9.33 |
| METTL16 | 9 | 9.23 | 11.80 | 11.84 | 11.40 | 8.35 | 8.61 | 9.21 | 8.93 | 9.80 | 10.84 | 10.51 | 11.62 | 8.48 | 9.07 | 10.32 |
| METTL17 | 1 | 8.37 | 9.17 | 11.28 | 12.86 | 10.64 | 10.16 | 7.60 | 5.47 | 13.90 | 10.55 | 8.77 | 12.07 | 10.40 | 6.53 | 12.22 |
| METTL18 | 2 | 9.48 | 9.50 | 8.54 | 7.26 | 11.23 | 10.95 | 12.31 | 11.15 | 10.00 | 9.58 | 9.49 | 7.90 | 11.09 | 11.73 | 9.79 |
| METTL2B | 8 | 9.91 | 10.80 | 12.31 | 12.03 | 8.54 | 8.33 | 8.24 | 7.86 | 11.11 | 10.88 | 10.35 | 12.17 | 8.44 | 8.05 | 10.99 |
| METTL3 | 9 | 10.04 | 11.82 | 9.38 | 8.35 | 9.29 | 9.72 | 10.84 | 10.12 | 10.28 | 10.16 | 10.93 | 8.87 | 9.50 | 10.48 | 10.22 |
| METTL5 | 3 | 9.32 | 10.70 | 10.58 | 10.01 | 9.48 | 10.34 | 10.60 | 10.46 | 9.60 | 8.91 | 10.01 | 10.30 | 9.91 | 10.53 | 9.25 |
| METTL7A | 1 | 12.42 | 6.70 | 5.06 | 6.64 | 13.60 | 12.98 | 10.86 | 12.66 | 8.73 | 10.33 | 9.56 | 5.85 | 13.29 | 11.76 | 9.53 |
| METTL7B | 1 | 13.67 | 6.40 | 5.64 | 8.16 | 9.51 | 10.45 | 10.99 | 11.29 | 18.01 | 5.90 | 10.03 | 6.90 | 9.98 | 11.14 | 11.96 |
| METTL8 | 1 | 10.12 | 8.16 | 10.19 | 10.89 | 9.94 | 7.82 | 9.78 | 8.49 | 13.38 | 11.23 | 9.14 | 10.54 | 8.88 | 9.13 | 12.30 |
| METTL9 | 1 | 10.77 | 10.42 | 9.55 | 10.75 | 9.46 | 10.04 | 10.30 | 9.01 | 9.72 | 10.00 | 10.59 | 10.15 | 9.75 | 9.65 | 9.86 |
| MEX3C | 3 | 9.01 | 10.34 | 12.77 | 12.88 | 7.72 | 8.65 | 8.72 | 8.45 | 10.77 | 10.69 | 9.67 | 12.83 | 8.18 | 8.59 | 10.73 |

| | | | | | | | | | | | | | | | | |
|---|---|---|---|---|---|---|---|---|---|---|---|---|---|---|---|---|
| MFAP1 | 10 | 8.14 | 11.49 | 11.02 | 9.69 | 9.71 | 9.47 | 11.08 | 9.37 | 10.13 | 9.89 | 9.82 | 10.36 | 9.59 | 10.22 | 10.01 |
| MFF | 16 | 13.19 | 12.13 | 6.14 | 6.85 | 11.09 | 11.20 | 10.71 | 9.63 | 9.65 | 9.42 | 12.66 | 6.49 | 11.15 | 10.17 | 9.53 |
| MFN1 | 6 | 11.05 | 11.57 | 10.17 | 10.01 | 10.15 | 10.04 | 8.10 | 8.51 | 9.99 | 10.41 | 11.31 | 10.09 | 10.10 | 8.30 | 10.20 |
| MFN2 | 13 | 10.98 | 11.09 | 8.79 | 9.68 | 9.76 | 10.06 | 10.49 | 10.02 | 9.52 | 9.60 | 11.04 | 9.23 | 9.91 | 10.26 | 9.56 |
| MFSD1 | 1 | 12.79 | 7.43 | 7.36 | 9.94 | 12.46 | 11.61 | 7.58 | 7.37 | 11.35 | 12.12 | 10.11 | 8.65 | 12.04 | 7.48 | 11.73 |
| MFSD6 | 1 | 9.36 | 11.37 | 7.31 | 12.35 | 10.95 | 10.15 | 8.78 | 10.56 | 9.71 | 9.47 | 10.37 | 9.83 | 10.55 | 9.67 | 9.59 |
| MGA | 5 | 8.60 | 9.45 | 9.70 | 9.85 | 10.08 | 10.72 | 10.85 | 10.01 | 10.83 | 9.92 | 9.02 | 9.77 | 10.40 | 10.43 | 10.37 |
| MGAT1 | 1 | 11.95 | 6.48 | 7.95 | 9.89 | 9.08 | 7.55 | 15.74 | 11.78 | 9.77 | 9.81 | 9.22 | 8.92 | 8.31 | 13.76 | 9.79 |
| MGAT2 | 2 | 11.11 | 8.01 | 7.48 | 10.36 | 10.20 | 11.35 | 9.97 | 9.08 | 10.93 | 11.52 | 9.56 | 8.92 | 10.77 | 9.52 | 11.22 |
| MGAT4B | 1 | 9.54 | 12.08 | 6.96 | 9.15 | 10.82 | 10.06 | 8.73 | 9.14 | 12.14 | 11.38 | 10.81 | 8.06 | 10.44 | 8.93 | 11.76 |
| MGAT5 | 1 | 11.08 | 8.01 | 8.93 | 12.16 | 8.98 | 9.21 | 11.02 | 9.22 | 9.57 | 11.84 | 9.54 | 10.54 | 9.09 | 10.12 | 10.70 |
| MGEA5 | 18 | 10.08 | 10.60 | 9.30 | 9.25 | 11.99 | 12.08 | 8.79 | 8.36 | 9.87 | 9.69 | 10.34 | 9.27 | 12.03 | 8.57 | 9.78 |
| MGLL | 1 | 10.01 | 11.54 | 13.01 | 9.35 | 10.93 | 10.79 | 6.73 | 5.95 | 11.33 | 10.35 | 10.78 | 11.18 | 10.86 | 6.34 | 10.84 |
| MGME1 | 7 | 8.92 | 8.84 | 12.19 | 11.74 | 9.16 | 7.72 | 11.03 | 8.25 | 11.17 | 11.00 | 8.88 | 11.96 | 8.44 | 9.64 | 11.08 |
| MGMT | 4 | 7.94 | 11.02 | 5.54 | 4.80 | 16.07 | 15.30 | 13.86 | 21.18 | 2.44 | 1.86 | 9.48 | 5.17 | 15.68 | 17.52 | 2.15 |
| MGST1 | 4 | 5.47 | 2.03 | 3.33 | 1.02 | 7.85 | 4.42 | 4.08 | 61.16 | 5.37 | 5.28 | 3.75 | 2.17 | 6.13 | 32.62 | 5.32 |
| MGST3 | 4 | 12.58 | 7.17 | 8.05 | 7.51 | 13.15 | 10.47 | 7.84 | 7.32 | 12.06 | 13.86 | 9.88 | 7.78 | 11.81 | 7.58 | 12.96 |
| MIA3 | 14 | 11.25 | 7.05 | 7.81 | 10.44 | 11.97 | 11.03 | 11.63 | 11.18 | 8.12 | 9.52 | 9.15 | 9.13 | 11.50 | 11.41 | 8.82 |
| MIB1 | 7 | 9.88 | 10.67 | 12.06 | 11.51 | 9.37 | 9.65 | 8.32 | 7.61 | 10.47 | 10.46 | 10.28 | 11.78 | 9.51 | 7.97 | 10.47 |
| MICA | 2 | 10.36 | 8.73 | 8.84 | 9.65 | 10.11 | 9.56 | 8.15 | 7.29 | 12.93 | 14.39 | 9.55 | 9.25 | 9.83 | 7.72 | 13.66 |
| MICAL1 | 10 | 10.53 | 11.42 | 12.44 | 12.58 | 6.99 | 7.31 | 4.77 | 4.91 | 14.45 | 14.60 | 10.97 | 12.51 | 7.15 | 4.84 | 14.52 |
| MICAL3 | 8 | 10.29 | 11.38 | 11.44 | 11.42 | 7.86 | 8.11 | 10.07 | 8.72 | 10.49 | 10.22 | 10.84 | 11.43 | 7.98 | 9.40 | 10.36 |
| MICALL1 | 11 | 9.61 | 9.67 | 14.95 | 14.57 | 7.83 | 8.19 | 7.68 | 6.72 | 10.91 | 9.85 | 9.64 | 14.76 | 8.01 | 7.20 | 10.38 |
| MICALL2 | 2 | 10.54 | 10.45 | 9.41 | 8.88 | 10.10 | 9.82 | 11.18 | 10.34 | 10.12 | 9.17 | 10.50 | 9.15 | 9.96 | 10.76 | 9.64 |
| MICU1 | 6 | 10.27 | 8.63 | 11.92 | 12.33 | 9.72 | 10.63 | 8.49 | 8.23 | 10.08 | 9.69 | 9.45 | 12.12 | 10.18 | 8.36 | 9.88 |
| MICU2 | 2 | 9.31 | 8.79 | 12.44 | 11.71 | 10.43 | 10.89 | 8.49 | 8.06 | 9.33 | 10.53 | 9.05 | 12.08 | 10.66 | 8.28 | 9.93 |
| MID1 | 3 | 8.47 | 8.83 | 10.51 | 10.22 | 9.12 | 9.71 | 13.64 | 12.94 | 8.52 | 8.05 | 8.65 | 10.36 | 9.41 | 13.29 | 8.29 |
| MID1IP1 | 2 | 10.16 | 10.86 | 9.09 | 8.03 | 10.09 | 9.81 | 12.66 | 9.83 | 9.71 | 9.76 | 10.51 | 8.56 | 9.95 | 11.24 | 9.74 |
| MID2 | 3 | 9.39 | 10.67 | 9.60 | 9.68 | 11.29 | 11.61 | 11.03 | 10.38 | 8.44 | 7.91 | 10.03 | 9.64 | 11.45 | 10.71 | 8.18 |
| MIEF1 | 3 | 8.78 | 8.28 | 15.72 | 15.41 | 5.89 | 6.35 | 10.33 | 10.39 | 9.48 | 9.37 | 8.53 | 15.57 | 6.12 | 10.36 | 9.42 |
| MIEN1 | 4 | 10.62 | 12.28 | 11.29 | 10.82 | 10.72 | 10.96 | 9.40 | 6.81 | 8.58 | 8.52 | 11.45 | 11.06 | 10.84 | 8.10 | 8.55 |
| MIER1 | 11 | 11.79 | 12.05 | 9.47 | 9.11 | 8.84 | 8.80 | 10.57 | 9.62 | 9.97 | 9.78 | 11.92 | 9.29 | 8.82 | 10.10 | 9.87 |
| MIF | 7 | 8.36 | 8.22 | 10.14 | 7.92 | 7.50 | 6.53 | 9.47 | 24.94 | 8.88 | 8.03 | 8.29 | 9.03 | 7.02 | 17.20 | 8.45 |
| MIF4GD | 2 | 11.74 | 11.50 | 7.76 | 8.14 | 11.54 | 12.00 | 8.19 | 8.14 | 10.59 | 10.40 | 11.62 | 7.95 | 11.77 | 8.16 | 10.50 |
| MINA | 5 | 9.36 | 9.59 | 11.26 | 10.66 | 8.24 | 8.23 | 12.48 | 11.74 | 9.54 | 8.90 | 9.47 | 10.96 | 8.24 | 12.11 | 9.22 |
| MINK1 | 17 | 9.65 | 10.00 | 11.28 | 9.91 | 9.99 | 9.77 | 9.25 | 9.04 | 10.55 | 10.55 | 9.82 | 10.59 | 9.88 | 9.14 | 10.55 |
| MINPP1 | 3 | 11.18 | 8.59 | 8.07 | 9.22 | 11.93 | 12.06 | 9.63 | 9.58 | 10.49 | 9.25 | 9.89 | 8.64 | 11.99 | 9.60 | 9.87 |
| MIOS | 2 | 9.36 | 10.36 | 11.75 | 12.62 | 9.10 | 9.25 | 8.79 | 8.41 | 10.02 | 10.35 | 9.86 | 12.19 | 9.18 | 8.60 | 10.18 |
| MIPEP | 8 | 12.20 | 10.90 | 9.49 | 11.63 | 9.30 | 8.71 | 9.34 | 9.03 | 9.62 | 9.78 | 11.55 | 10.56 | 9.01 | 9.18 | 9.70 |
| MIS12 | 2 | 10.62 | 10.38 | 10.76 | 6.59 | 13.02 | 9.99 | 9.37 | 6.96 | 11.03 | 11.27 | 10.50 | 8.68 | 11.51 | 8.16 | 11.15 |
| MIS18BP1 | 3 | 11.69 | 10.47 | 9.39 | 9.05 | 8.76 | 9.36 | 12.53 | 11.92 | 8.95 | 7.89 | 11.08 | 9.22 | 9.06 | 12.23 | 8.42 |
| MISP | 26 | 10.05 | 10.74 | 10.23 | 10.48 | 12.21 | 13.05 | 5.54 | 5.39 | 11.15 | 11.16 | 10.39 | 10.36 | 12.63 | 5.47 | 11.15 |

| | | | | | | | | | | | | | | | | |
|---|---|---|---|---|---|---|---|---|---|---|---|---|---|---|---|---|
| MITF | 2 | 10.79 | 14.57 | 9.10 | 9.96 | 9.13 | 13.45 | 7.69 | 5.15 | 10.92 | 9.24 | 12.68 | 9.53 | 11.29 | 6.42 | 10.08 |
| MKI67 | 76 | 13.65 | 6.42 | 7.08 | 11.15 | 7.98 | 6.49 | 10.81 | 9.62 | 8.31 | 18.49 | 10.04 | 9.11 | 7.24 | 10.21 | 13.40 |
| MKL1 | 3 | 13.01 | 13.08 | 12.54 | 14.13 | 6.21 | 5.79 | 7.21 | 6.20 | 11.28 | 10.55 | 13.05 | 13.33 | 6.00 | 6.70 | 10.92 |
| MKL2 | 6 | 10.86 | 11.19 | 10.65 | 10.53 | 9.93 | 10.54 | 9.26 | 9.44 | 9.98 | 7.63 | 11.02 | 10.59 | 10.24 | 9.35 | 8.80 |
| MKLN1 | 4 | 9.76 | 9.82 | 10.42 | 8.98 | 9.90 | 9.81 | 10.60 | 9.37 | 10.90 | 10.42 | 9.79 | 9.70 | 9.86 | 9.99 | 10.66 |
| MKNK1 | 1 | 13.15 | 13.67 | 9.74 | 9.61 | 6.36 | 7.71 | 11.23 | 10.36 | 8.97 | 9.21 | 13.41 | 9.67 | 7.04 | 10.79 | 9.09 |
| MKRN2 | 6 | 10.24 | 11.37 | 8.88 | 7.94 | 9.20 | 8.95 | 11.59 | 12.58 | 10.00 | 9.24 | 10.81 | 8.41 | 9.08 | 12.09 | 9.62 |
| MLEC | 11 | 13.16 | 6.02 | 7.93 | 11.37 | 11.47 | 8.84 | 9.47 | 9.99 | 9.42 | 12.31 | 9.59 | 9.65 | 10.15 | 9.73 | 10.87 |
| MLF2 | 2 | 11.01 | 10.85 | 11.54 | 10.82 | 8.85 | 9.94 | 8.54 | 7.42 | 11.59 | 9.45 | 10.93 | 11.18 | 9.40 | 7.98 | 10.52 |
| MLH1 | 19 | 10.73 | 11.18 | 9.27 | 8.51 | 9.96 | 9.66 | 9.68 | 9.42 | 10.93 | 10.67 | 10.95 | 8.89 | 9.81 | 9.55 | 10.80 |
| MLKL | 16 | 9.19 | 9.34 | 7.75 | 6.43 | 12.98 | 13.20 | 12.66 | 11.47 | 8.43 | 8.54 | 9.26 | 7.09 | 13.09 | 12.06 | 8.49 |
| MLLT1 | 5 | 8.12 | 8.57 | 8.31 | 7.70 | 10.50 | 9.98 | 8.83 | 23.86 | 6.97 | 7.16 | 8.34 | 8.01 | 10.24 | 16.35 | 7.07 |
| MLLT11 | 2 | 10.86 | 12.02 | 9.85 | 9.75 | 13.47 | 14.28 | 6.66 | 2.41 | 10.43 | 10.27 | 11.44 | 9.80 | 13.87 | 4.54 | 10.35 |
| MLLT3 | 5 | 8.91 | 9.82 | 13.36 | 10.03 | 8.91 | 8.56 | 8.10 | 7.01 | 12.06 | 13.25 | 9.36 | 11.69 | 8.73 | 7.55 | 12.65 |
| MLLT4 | 9 | 9.77 | 10.87 | 12.82 | 13.05 | 12.24 | 13.49 | 5.54 | 4.78 | 8.73 | 8.71 | 10.32 | 12.94 | 12.86 | 5.16 | 8.72 |
| MLST8 | 1 | 8.61 | 11.79 | 11.97 | 6.62 | 13.58 | 8.38 | 10.84 | 6.94 | 9.82 | 11.45 | 10.20 | 9.30 | 10.98 | 8.89 | 10.63 |
| MLTK | 8 | 9.79 | 10.70 | 9.88 | 8.91 | 9.67 | 9.32 | 12.59 | 11.93 | 8.61 | 8.60 | 10.24 | 9.40 | 9.50 | 12.26 | 8.61 |
| MLX | 1 | 9.22 | 13.67 | 7.52 | 7.65 | 9.70 | 9.94 | 10.35 | 11.29 | 10.05 | 10.62 | 11.44 | 7.58 | 9.82 | 10.82 | 10.33 |
| MLYCD | 2 | 11.70 | 8.89 | 8.88 | 8.96 | 12.58 | 11.67 | 8.91 | 8.60 | 10.27 | 9.54 | 10.29 | 8.92 | 12.13 | 8.75 | 9.91 |
| MMAB | 4 | 10.83 | 10.65 | 8.01 | 8.50 | 14.16 | 14.00 | 8.50 | 7.72 | 8.78 | 8.84 | 10.74 | 8.26 | 14.08 | 8.11 | 8.81 |
| MMACHC | 2 | 11.51 | 12.11 | 8.87 | 7.60 | 7.52 | 8.03 | 12.88 | 11.89 | 9.57 | 10.02 | 11.81 | 8.23 | 7.78 | 12.39 | 9.80 |
| MMADHC | 2 | 9.95 | 9.24 | 7.66 | 8.52 | 12.13 | 12.60 | 6.85 | 13.79 | 9.86 | 9.40 | 9.60 | 8.09 | 12.36 | 10.32 | 9.63 |
| MMGT1 | 1 | 12.94 | 6.16 | 7.35 | 13.18 | 9.53 | 9.15 | 9.10 | 9.38 | 9.12 | 14.08 | 9.55 | 10.27 | 9.34 | 9.24 | 11.60 |
| MMRN1 | 2 | 12.35 | 16.69 | 4.28 | 5.30 | 12.92 | 18.21 | 5.61 | 4.27 | 11.86 | 8.51 | 14.52 | 4.79 | 15.56 | 4.94 | 10.19 |
| MMS19 | 11 | 10.13 | 9.35 | 9.81 | 9.93 | 9.15 | 8.94 | 8.09 | 11.53 | 11.05 | 12.02 | 9.74 | 9.87 | 9.05 | 9.81 | 11.54 |
| MMS22L | 2 | 9.55 | 10.07 | 9.50 | 10.21 | 10.00 | 9.24 | 9.65 | 10.11 | 10.82 | 10.85 | 9.81 | 9.86 | 9.62 | 9.88 | 10.83 |
| MMTAG2 | 3 | 7.45 | 10.98 | 9.34 | 8.19 | 10.93 | 11.47 | 9.70 | 8.60 | 11.67 | 11.66 | 9.22 | 8.77 | 11.20 | 9.15 | 11.67 |
| MND1 | 1 | 10.93 | 11.85 | 13.01 | 11.42 | 11.95 | 9.12 | 10.95 | 12.74 | 4.38 | 3.65 | 11.39 | 12.22 | 10.53 | 11.84 | 4.02 |
| MOB1B | 10 | 10.03 | 10.63 | 10.58 | 9.34 | 10.27 | 9.95 | 9.90 | 9.61 | 9.86 | 9.84 | 10.33 | 9.96 | 10.11 | 9.75 | 9.85 |
| MOB4 | 2 | 3.70 | 2.40 | 4.73 | 1.15 | 5.46 | 2.52 | 3.70 | 69.46 | 3.83 | 3.04 | 3.05 | 2.94 | 3.99 | 36.58 | 3.43 |
| MOCOS | 5 | 15.46 | 14.47 | 12.83 | 11.50 | 6.03 | 6.45 | 6.02 | 5.86 | 10.54 | 10.83 | 14.97 | 12.17 | 6.24 | 5.94 | 10.69 |
| MOCS2 | 11 | 9.30 | 10.20 | 8.73 | 8.21 | 11.97 | 12.26 | 11.96 | 11.19 | 8.47 | 7.71 | 9.75 | 8.47 | 12.11 | 11.58 | 8.09 |
| MOCS3 | 9 | 10.20 | 11.73 | 8.80 | 8.96 | 10.51 | 11.74 | 8.30 | 8.18 | 11.02 | 10.57 | 10.96 | 8.88 | 11.12 | 8.24 | 10.80 |
| MOGS | 9 | 11.18 | 6.84 | 8.42 | 11.92 | 8.32 | 8.32 | 9.59 | 9.25 | 11.71 | 14.44 | 9.01 | 10.17 | 8.32 | 9.42 | 13.08 |
| MON1B | 2 | 11.96 | 10.09 | 8.66 | 7.99 | 11.30 | 11.44 | 8.91 | 9.71 | 9.74 | 10.20 | 11.03 | 8.32 | 11.37 | 9.31 | 9.97 |
| MON2 | 7 | 11.03 | 11.61 | 8.58 | 9.69 | 9.20 | 8.94 | 8.89 | 9.40 | 11.42 | 11.23 | 11.32 | 9.14 | 9.07 | 9.15 | 11.32 |
| MORC2 | 13 | 9.06 | 9.40 | 11.80 | 11.93 | 8.00 | 8.50 | 12.24 | 11.01 | 8.67 | 9.38 | 9.23 | 11.87 | 8.25 | 11.63 | 9.03 |
| MORC3 | 3 | 10.83 | 12.69 | 7.97 | 8.85 | 9.49 | 11.45 | 9.69 | 9.43 | 9.93 | 9.68 | 11.76 | 8.41 | 10.47 | 9.56 | 9.81 |
| MORF4L1 | 9 | 10.39 | 10.96 | 9.53 | 8.36 | 10.05 | 10.02 | 10.03 | 9.39 | 10.50 | 10.76 | 10.67 | 8.95 | 10.04 | 9.71 | 10.63 |
| MORF4L2 | 8 | 10.19 | 10.17 | 7.85 | 8.16 | 12.45 | 12.30 | 11.79 | 10.91 | 8.41 | 7.77 | 10.18 | 8.00 | 12.37 | 11.35 | 8.09 |
| MOSPD2 | 2 | 10.79 | 7.12 | 6.22 | 9.29 | 10.85 | 10.21 | 13.45 | 13.30 | 8.75 | 10.02 | 8.96 | 7.76 | 10.53 | 13.37 | 9.38 |
| MOV10 | 23 | 10.14 | 11.28 | 10.31 | 9.99 | 10.90 | 11.11 | 10.11 | 9.34 | 8.48 | 8.36 | 10.71 | 10.15 | 11.01 | 9.72 | 8.42 |

| | | | | | | | | | | | | | | | | |
|---|---|---|---|---|---|---|---|---|---|---|---|---|---|---|---|---|
| MPC2 | 1 | 16.02 | 8.24 | 5.97 | 6.95 | 10.98 | 10.07 | 9.50 | 8.54 | 10.44 | 13.29 | 12.13 | 6.46 | 10.53 | 9.02 | 11.86 |
| MPDZ | 9 | 10.24 | 10.32 | 12.22 | 12.90 | 7.79 | 8.72 | 7.48 | 6.90 | 11.67 | 11.76 | 10.28 | 12.56 | 8.26 | 7.19 | 11.72 |
| MPG | 8 | 9.60 | 10.24 | 10.82 | 10.16 | 9.66 | 10.13 | 9.75 | 8.53 | 11.16 | 9.95 | 9.92 | 10.49 | 9.90 | 9.14 | 10.55 |
| MPHOSPH10 | 10 | 9.13 | 9.74 | 11.37 | 10.54 | 8.39 | 8.55 | 12.02 | 11.02 | 10.12 | 9.11 | 9.44 | 10.95 | 8.47 | 11.52 | 9.61 |
| MPHOSPH6 | 5 | 10.90 | 11.30 | 8.65 | 7.81 | 11.56 | 11.95 | 10.24 | 6.46 | 10.76 | 10.35 | 11.10 | 8.23 | 11.76 | 8.35 | 10.56 |
| MPHOSPH8 | 9 | 9.67 | 10.80 | 10.04 | 10.13 | 10.81 | 11.46 | 9.03 | 9.12 | 9.29 | 9.64 | 10.24 | 10.09 | 11.13 | 9.08 | 9.46 |
| MPI | 14 | 9.49 | 10.11 | 9.03 | 8.90 | 10.17 | 10.58 | 10.21 | 10.10 | 10.66 | 10.74 | 9.80 | 8.97 | 10.38 | 10.15 | 10.70 |
| MPP5 | 7 | 9.93 | 11.09 | 10.99 | 12.09 | 8.91 | 9.39 | 7.62 | 6.69 | 11.37 | 11.92 | 10.51 | 11.54 | 9.15 | 7.15 | 11.64 |
| MPP6 | 13 | 9.94 | 11.31 | 11.06 | 11.15 | 8.42 | 8.39 | 9.59 | 8.51 | 10.81 | 10.82 | 10.63 | 11.11 | 8.40 | 9.05 | 10.82 |
| MPRIP | 42 | 9.69 | 10.57 | 13.67 | 12.69 | 8.98 | 9.10 | 7.65 | 6.94 | 10.75 | 9.95 | 10.13 | 13.18 | 9.04 | 7.30 | 10.35 |
| MPST | 5 | 10.10 | 11.01 | 11.72 | 12.10 | 7.81 | 7.41 | 10.40 | 9.25 | 9.97 | 10.22 | 10.55 | 11.91 | 7.61 | 9.83 | 10.10 |
| MPZL1 | 6 | 11.73 | 9.51 | 8.72 | 10.63 | 9.26 | 9.37 | 7.45 | 6.35 | 13.29 | 13.68 | 10.62 | 9.67 | 9.32 | 6.90 | 13.49 |
| MRC2 | 1 | 18.51 | 11.79 | 4.43 | 5.73 | 4.32 | 3.87 | 7.09 | 8.47 | 16.74 | 19.04 | 15.15 | 5.08 | 4.09 | 7.78 | 17.89 |
| MRE11A | 34 | 9.01 | 10.19 | 9.39 | 9.55 | 9.62 | 10.28 | 13.00 | 12.29 | 8.40 | 8.27 | 9.60 | 9.47 | 9.95 | 12.64 | 8.33 |
| MRFAP1 | 3 | 10.39 | 12.05 | 8.63 | 9.20 | 11.76 | 13.02 | 10.32 | 4.71 | 10.15 | 9.77 | 11.22 | 8.91 | 12.39 | 7.51 | 9.96 |
| MRGBP | 4 | 10.17 | 8.95 | 11.89 | 12.02 | 8.46 | 8.93 | 11.50 | 9.78 | 9.18 | 9.14 | 9.56 | 11.95 | 8.69 | 10.64 | 9.16 |
| MRI1 | 9 | 9.50 | 10.43 | 8.38 | 8.05 | 11.63 | 11.87 | 12.74 | 11.71 | 8.11 | 7.59 | 9.97 | 8.21 | 11.75 | 12.22 | 7.85 |
| MRM1 | 1 | 10.53 | 10.05 | 8.67 | 12.80 | 8.31 | 8.41 | 8.01 | 7.87 | 12.41 | 12.94 | 10.29 | 10.74 | 8.36 | 7.94 | 12.67 |
| MRP63 | 3 | 10.26 | 8.92 | 9.59 | 10.76 | 10.91 | 10.48 | 10.77 | 9.62 | 9.23 | 9.46 | 9.59 | 10.17 | 10.70 | 10.19 | 9.34 |
| MRPL1 | 8 | 10.93 | 8.92 | 8.03 | 9.51 | 11.70 | 10.83 | 11.13 | 10.65 | 8.99 | 9.32 | 9.92 | 8.77 | 11.27 | 10.89 | 9.16 |
| MRPL10 | 3 | 10.81 | 8.80 | 11.77 | 12.36 | 9.92 | 8.48 | 9.47 | 7.66 | 10.33 | 10.42 | 9.80 | 12.06 | 9.20 | 8.56 | 10.37 |
| MRPL11 | 12 | 9.73 | 8.76 | 10.38 | 11.82 | 10.10 | 9.68 | 10.90 | 9.85 | 9.31 | 9.47 | 9.25 | 11.10 | 9.89 | 10.37 | 9.39 |
| MRPL12 | 7 | 10.28 | 8.85 | 13.51 | 12.94 | 8.93 | 8.05 | 9.94 | 7.95 | 10.10 | 9.45 | 9.57 | 13.23 | 8.49 | 8.94 | 9.77 |
| MRPL13 | 5 | 10.18 | 9.20 | 9.55 | 9.80 | 9.88 | 10.47 | 11.35 | 10.69 | 9.27 | 9.61 | 9.69 | 9.67 | 10.17 | 11.02 | 9.44 |
| MRPL14 | 3 | 9.97 | 9.72 | 8.94 | 11.20 | 10.09 | 10.74 | 10.51 | 10.21 | 8.18 | 10.44 | 9.85 | 10.07 | 10.42 | 10.36 | 9.31 |
| MRPL15 | 18 | 10.80 | 8.36 | 9.39 | 11.44 | 10.53 | 9.38 | 11.08 | 9.76 | 9.27 | 9.99 | 9.58 | 10.42 | 9.95 | 10.42 | 9.63 |
| MRPL16 | 5 | 8.64 | 8.35 | 9.40 | 10.90 | 9.34 | 9.46 | 9.72 | 15.72 | 9.14 | 9.33 | 8.49 | 10.15 | 9.40 | 12.72 | 9.24 |
| MRPL17 | 6 | 9.73 | 8.87 | 9.71 | 10.84 | 10.06 | 9.91 | 10.87 | 10.35 | 10.02 | 9.63 | 9.30 | 10.27 | 9.98 | 10.61 | 9.83 |
| MRPL18 | 2 | 11.15 | 10.83 | 7.51 | 10.49 | 9.94 | 9.72 | 11.34 | 10.89 | 8.92 | 9.21 | 10.99 | 9.00 | 9.83 | 11.11 | 9.07 |
| MRPL19 | 12 | 10.31 | 8.96 | 9.57 | 9.98 | 10.43 | 9.96 | 11.28 | 10.51 | 9.38 | 9.62 | 9.63 | 9.78 | 10.20 | 10.89 | 9.50 |
| MRPL2 | 5 | 11.14 | 8.55 | 9.19 | 10.46 | 10.34 | 9.25 | 10.92 | 9.97 | 9.82 | 10.37 | 9.85 | 9.82 | 9.80 | 10.44 | 10.09 |
| MRPL20 | 4 | 10.04 | 8.99 | 9.49 | 12.31 | 9.42 | 9.05 | 10.59 | 9.90 | 9.71 | 10.49 | 9.52 | 10.90 | 9.24 | 10.25 | 10.10 |
| MRPL21 | 5 | 9.23 | 6.69 | 10.15 | 8.40 | 11.56 | 10.64 | 11.79 | 11.26 | 9.70 | 10.58 | 7.96 | 9.27 | 11.10 | 11.53 | 10.14 |
| MRPL22 | 9 | 11.10 | 9.36 | 9.20 | 9.43 | 9.96 | 9.48 | 11.42 | 10.58 | 10.06 | 9.42 | 10.23 | 9.31 | 9.72 | 11.00 | 9.74 |
| MRPL23 | 7 | 9.59 | 9.95 | 8.60 | 9.11 | 10.98 | 11.61 | 11.17 | 10.56 | 9.31 | 9.14 | 9.77 | 8.85 | 11.29 | 10.86 | 9.22 |
| MRPL24 | 2 | 9.93 | 8.90 | 9.59 | 10.61 | 10.94 | 10.68 | 11.08 | 10.33 | 9.09 | 8.86 | 9.41 | 10.10 | 10.81 | 10.70 | 8.98 |
| MRPL27 | 4 | 10.90 | 8.06 | 9.67 | 9.78 | 9.13 | 9.17 | 12.07 | 10.54 | 12.14 | 8.54 | 9.48 | 9.72 | 9.15 | 11.31 | 10.34 |
| MRPL28 | 2 | 10.66 | 8.72 | 10.41 | 8.59 | 11.61 | 10.21 | 11.20 | 9.66 | 9.52 | 9.42 | 9.69 | 9.50 | 10.91 | 10.43 | 9.47 |
| MRPL3 | 10 | 10.04 | 8.92 | 8.84 | 10.51 | 9.91 | 9.56 | 11.37 | 10.32 | 9.82 | 10.72 | 9.48 | 9.68 | 9.73 | 10.84 | 10.27 |
| MRPL30 | 6 | 10.03 | 9.48 | 10.39 | 10.03 | 10.25 | 9.79 | 11.20 | 10.16 | 9.18 | 9.47 | 9.76 | 10.21 | 10.02 | 10.68 | 9.32 |
| MRPL32 | 3 | 7.06 | 6.51 | 7.10 | 7.74 | 7.54 | 7.20 | 7.88 | 34.12 | 7.59 | 7.27 | 6.78 | 7.42 | 7.37 | 21.00 | 7.43 |
| MRPL34 | 3 | 11.73 | 9.51 | 10.10 | 8.89 | 10.13 | 9.71 | 10.78 | 9.19 | 10.10 | 9.86 | 10.62 | 9.50 | 9.92 | 9.98 | 9.98 |

| | | | | | | | | | | | | | | | | |
|---|---|---|---|---|---|---|---|---|---|---|---|---|---|---|---|---|
| MRPL37 | 13 | 11.51 | 8.79 | 8.60 | 10.01 | 10.60 | 9.93 | 10.84 | 10.11 | 10.02 | 9.57 | 10.15 | 9.31 | 10.27 | 10.48 | 9.79 |
| MRPL38 | 9 | 11.15 | 9.20 | 9.33 | 9.97 | 10.45 | 9.30 | 10.28 | 9.72 | 10.32 | 10.27 | 10.18 | 9.65 | 9.87 | 10.00 | 10.30 |
| MRPL39 | 11 | 10.64 | 9.44 | 9.46 | 9.76 | 10.57 | 10.24 | 10.77 | 8.64 | 10.49 | 10.00 | 10.04 | 9.61 | 10.40 | 9.70 | 10.24 |
| MRPL4 | 10 | 9.73 | 8.80 | 9.27 | 11.12 | 10.44 | 9.38 | 11.39 | 11.17 | 9.04 | 9.67 | 9.26 | 10.19 | 9.91 | 11.28 | 9.36 |
| MRPL40 | 6 | 10.07 | 8.90 | 10.35 | 11.69 | 10.02 | 9.70 | 10.21 | 9.32 | 9.94 | 9.81 | 9.48 | 11.02 | 9.86 | 9.76 | 9.87 |
| MRPL41 | 7 | 9.69 | 9.53 | 9.27 | 12.30 | 10.09 | 10.47 | 10.08 | 9.92 | 9.00 | 9.63 | 9.61 | 10.78 | 10.28 | 10.00 | 9.32 |
| MRPL42 | 4 | 11.14 | 8.57 | 10.22 | 10.24 | 10.94 | 9.81 | 11.42 | 9.32 | 9.13 | 9.22 | 9.85 | 10.23 | 10.37 | 10.37 | 9.17 |
| MRPL43 | 5 | 10.54 | 8.20 | 9.88 | 11.00 | 10.70 | 10.12 | 11.38 | 9.80 | 9.34 | 9.04 | 9.37 | 10.44 | 10.41 | 10.59 | 9.19 |
| MRPL44 | 8 | 10.40 | 8.30 | 9.30 | 11.26 | 10.05 | 9.63 | 11.15 | 10.90 | 9.35 | 9.66 | 9.35 | 10.28 | 9.84 | 11.02 | 9.51 |
| MRPL45 | 12 | 11.28 | 9.32 | 9.58 | 10.88 | 10.03 | 9.20 | 10.44 | 9.17 | 10.00 | 10.12 | 10.30 | 10.23 | 9.61 | 9.80 | 10.06 |
| MRPL46 | 8 | 10.51 | 8.49 | 10.28 | 10.84 | 10.50 | 9.81 | 9.63 | 8.55 | 10.79 | 10.60 | 9.50 | 10.56 | 10.15 | 9.09 | 10.70 |
| MRPL47 | 9 | 10.51 | 8.48 | 9.26 | 9.66 | 10.99 | 9.50 | 11.32 | 10.20 | 9.60 | 10.48 | 9.50 | 9.46 | 10.24 | 10.76 | 10.04 |
| MRPL48 | 4 | 9.78 | 9.20 | 10.05 | 9.73 | 10.42 | 10.46 | 10.25 | 10.67 | 9.77 | 9.69 | 9.49 | 9.89 | 10.44 | 10.46 | 9.73 |
| MRPL49 | 4 | 10.40 | 8.83 | 11.02 | 10.73 | 10.40 | 9.97 | 11.11 | 9.62 | 8.89 | 9.02 | 9.62 | 10.88 | 10.19 | 10.37 | 8.95 |
| MRPL50 | 4 | 6.54 | 9.26 | 12.75 | 11.79 | 10.64 | 9.71 | 11.33 | 7.78 | 10.19 | 10.01 | 7.90 | 12.27 | 10.18 | 9.55 | 10.10 |
| MRPL51 | 3 | 9.85 | 9.15 | 9.22 | 11.17 | 9.75 | 9.81 | 10.81 | 10.63 | 9.60 | 10.01 | 9.50 | 10.20 | 9.78 | 10.72 | 9.80 |
| MRPL52 | 1 | 8.73 | 9.43 | 11.57 | 9.79 | 10.85 | 9.52 | 10.56 | 10.22 | 9.42 | 9.92 | 9.08 | 10.68 | 10.18 | 10.39 | 9.67 |
| MRPL53 | 1 | 9.48 | 10.22 | 11.66 | 12.32 | 10.14 | 9.82 | 8.72 | 8.47 | 9.72 | 9.45 | 9.85 | 11.99 | 9.98 | 8.60 | 9.59 |
| MRPL54 | 3 | 9.50 | 8.95 | 10.35 | 12.16 | 9.94 | 10.49 | 9.92 | 9.65 | 9.53 | 9.51 | 9.22 | 11.26 | 10.22 | 9.78 | 9.52 |
| MRPL55 | 3 | 10.44 | 8.55 | 10.18 | 9.08 | 11.94 | 10.16 | 11.38 | 8.49 | 9.67 | 10.12 | 9.49 | 9.63 | 11.05 | 9.94 | 9.90 |
| MRPL9 | 13 | 10.10 | 8.48 | 9.77 | 10.69 | 10.70 | 10.10 | 10.65 | 10.25 | 9.45 | 9.81 | 9.29 | 10.23 | 10.40 | 10.45 | 9.63 |
| MRPS10 | 8 | 10.43 | 9.04 | 10.31 | 11.58 | 9.35 | 9.33 | 11.21 | 10.52 | 9.20 | 9.03 | 9.73 | 10.94 | 9.34 | 10.86 | 9.11 |
| MRPS11 | 6 | 9.56 | 8.84 | 8.94 | 9.62 | 10.12 | 9.98 | 11.06 | 10.75 | 10.02 | 11.12 | 9.20 | 9.28 | 10.05 | 10.90 | 10.57 |
| MRPS12 | 2 | 10.10 | 9.00 | 10.34 | 11.21 | 9.40 | 9.17 | 11.02 | 10.03 | 9.64 | 10.10 | 9.55 | 10.77 | 9.28 | 10.52 | 9.87 |
| MRPS14 | 2 | 10.99 | 9.91 | 8.70 | 9.47 | 10.38 | 10.10 | 10.28 | 10.31 | 10.05 | 9.82 | 10.45 | 9.08 | 10.24 | 10.29 | 9.94 |
| MRPS15 | 6 | 11.31 | 9.60 | 7.63 | 10.41 | 8.61 | 8.72 | 10.53 | 11.16 | 10.19 | 11.84 | 10.46 | 9.02 | 8.67 | 10.84 | 11.01 |
| MRPS16 | 5 | 10.40 | 9.54 | 9.98 | 10.91 | 9.06 | 8.61 | 11.15 | 10.80 | 9.52 | 10.02 | 9.97 | 10.45 | 8.84 | 10.97 | 9.77 |
| MRPS17 | 7 | 10.85 | 9.23 | 9.08 | 10.68 | 8.64 | 8.70 | 11.69 | 11.90 | 9.42 | 9.80 | 10.04 | 9.88 | 8.67 | 11.80 | 9.61 |
| MRPS18A | 3 | 11.11 | 9.31 | 8.98 | 9.56 | 10.19 | 9.82 | 10.75 | 9.96 | 10.39 | 9.93 | 10.21 | 9.27 | 10.01 | 10.36 | 10.16 |
| MRPS18B | 3 | 9.99 | 9.55 | 9.54 | 10.70 | 8.81 | 8.78 | 11.16 | 10.80 | 10.12 | 10.56 | 9.77 | 10.12 | 8.79 | 10.98 | 10.34 |
| MRPS2 | 12 | 10.84 | 9.37 | 10.12 | 11.79 | 9.02 | 8.89 | 10.49 | 9.77 | 9.59 | 10.12 | 10.10 | 10.96 | 8.96 | 10.13 | 9.85 |
| MRPS21 | 1 | 10.97 | 8.77 | 8.06 | 8.73 | 10.54 | 10.17 | 11.86 | 10.83 | 9.90 | 10.19 | 9.87 | 8.39 | 10.35 | 11.34 | 10.04 |
| MRPS22 | 21 | 10.49 | 9.65 | 9.83 | 10.97 | 8.95 | 8.81 | 10.92 | 10.41 | 9.68 | 10.29 | 10.07 | 10.40 | 8.88 | 10.67 | 9.99 |
| MRPS23 | 8 | 10.88 | 9.85 | 10.63 | 10.68 | 9.40 | 9.26 | 10.63 | 9.61 | 9.78 | 9.27 | 10.36 | 10.65 | 9.33 | 10.12 | 9.53 |
| MRPS25 | 6 | 10.15 | 9.04 | 10.79 | 11.71 | 8.17 | 8.07 | 11.81 | 10.67 | 10.15 | 9.45 | 9.59 | 11.25 | 8.12 | 11.24 | 9.80 |
| MRPS26 | 13 | 9.55 | 9.78 | 9.49 | 10.30 | 9.24 | 9.33 | 11.58 | 10.71 | 9.78 | 10.23 | 9.67 | 9.89 | 9.29 | 11.14 | 10.01 |
| MRPS27 | 10 | 11.62 | 9.52 | 8.20 | 9.96 | 9.82 | 8.95 | 10.96 | 10.81 | 9.75 | 10.41 | 10.57 | 9.08 | 9.38 | 10.88 | 10.08 |
| MRPS28 | 9 | 10.68 | 9.48 | 10.80 | 11.89 | 8.78 | 8.48 | 10.89 | 10.43 | 9.26 | 9.32 | 10.08 | 11.35 | 8.63 | 10.66 | 9.29 |
| MRPS30 | 1 | 10.69 | 8.99 | 7.91 | 8.14 | 11.12 | 10.41 | 11.17 | 10.77 | 10.71 | 10.09 | 9.84 | 8.02 | 10.77 | 10.97 | 10.40 |
| MRPS31 | 6 | 10.22 | 9.65 | 11.33 | 11.36 | 10.02 | 9.78 | 10.87 | 9.97 | 8.48 | 8.33 | 9.93 | 11.34 | 9.90 | 10.42 | 8.41 |
| MRPS33 | 2 | 10.89 | 9.18 | 8.65 | 10.09 | 10.59 | 10.46 | 10.63 | 10.15 | 9.18 | 10.17 | 10.04 | 9.37 | 10.52 | 10.39 | 9.67 |
| MRPS34 | 9 | 11.30 | 9.88 | 9.42 | 10.26 | 9.40 | 8.70 | 10.98 | 9.19 | 10.52 | 10.35 | 10.59 | 9.84 | 9.05 | 10.08 | 10.44 |

| | | | | | | | | | | | | | | | | |
|---|---|---|---|---|---|---|---|---|---|---|---|---|---|---|---|---|
| MRPS35 | 7 | 10.59 | 8.69 | 9.34 | 11.05 | 9.17 | 8.62 | 11.23 | 13.03 | 9.00 | 9.29 | 9.64 | 10.19 | 8.90 | 12.13 | 9.14 |
| MRPS36 | 5 | 9.87 | 9.54 | 10.15 | 10.75 | 10.68 | 9.94 | 9.74 | 8.97 | 10.60 | 9.77 | 9.70 | 10.45 | 10.31 | 9.35 | 10.18 |
| MRPS5 | 14 | 10.22 | 8.67 | 8.89 | 10.80 | 9.61 | 8.90 | 11.40 | 10.68 | 10.00 | 10.84 | 9.44 | 9.84 | 9.26 | 11.04 | 10.42 |
| MRPS6 | 4 | 9.51 | 8.09 | 8.54 | 7.79 | 8.71 | 8.40 | 11.16 | 19.71 | 9.38 | 8.70 | 8.80 | 8.17 | 8.56 | 15.44 | 9.04 |
| MRPS7 | 12 | 10.49 | 9.25 | 11.56 | 11.49 | 8.89 | 8.08 | 10.11 | 9.76 | 10.21 | 10.16 | 9.87 | 11.52 | 8.48 | 9.94 | 10.19 |
| MRPS9 | 17 | 10.44 | 9.76 | 11.30 | 11.33 | 8.82 | 8.51 | 9.79 | 8.99 | 10.62 | 10.43 | 10.10 | 11.32 | 8.67 | 9.39 | 10.53 |
| MRRF | 7 | 11.04 | 9.76 | 8.56 | 9.73 | 9.70 | 9.74 | 10.02 | 9.38 | 11.35 | 10.72 | 10.40 | 9.15 | 9.72 | 9.70 | 11.04 |
| MRTO4 | 23 | 8.68 | 9.74 | 10.27 | 8.41 | 7.27 | 7.18 | 11.37 | 17.47 | 9.85 | 9.75 | 9.21 | 9.34 | 7.23 | 14.42 | 9.80 |
| MSANTD2 | 2 | 4.87 | 16.74 | 6.28 | 5.51 | 14.19 | 22.43 | 7.11 | 6.44 | 8.67 | 7.75 | 10.81 | 5.90 | 18.31 | 6.77 | 8.21 |
| MSH2 | 34 | 10.15 | 9.98 | 8.16 | 7.71 | 11.27 | 11.51 | 10.56 | 12.17 | 9.46 | 9.02 | 10.06 | 7.93 | 11.39 | 11.36 | 9.24 |
| MSH3 | 17 | 9.90 | 10.22 | 7.93 | 7.69 | 11.68 | 11.58 | 11.35 | 10.91 | 9.64 | 9.10 | 10.06 | 7.81 | 11.63 | 11.13 | 9.37 |
| MSH6 | 45 | 9.73 | 10.19 | 8.33 | 7.53 | 11.48 | 11.51 | 10.63 | 11.25 | 9.82 | 9.52 | 9.96 | 7.93 | 11.49 | 10.94 | 9.67 |
| MSI2 | 9 | 11.69 | 12.14 | 10.19 | 7.34 | 13.46 | 12.89 | 8.86 | 6.72 | 8.80 | 7.91 | 11.91 | 8.77 | 13.18 | 7.79 | 8.35 |
| MSL1 | 4 | 8.91 | 11.91 | 11.42 | 9.44 | 9.05 | 9.14 | 10.81 | 8.35 | 10.70 | 10.27 | 10.41 | 10.43 | 9.09 | 9.58 | 10.48 |
| MSLN | 2 | 12.35 | 11.14 | 5.20 | 5.28 | 14.97 | 14.65 | 6.05 | 5.56 | 11.90 | 12.89 | 11.74 | 5.24 | 14.81 | 5.81 | 12.40 |
| MSN | 124 | 7.84 | 9.17 | 12.66 | 11.81 | 11.01 | 10.90 | 8.81 | 7.91 | 10.26 | 9.62 | 8.51 | 12.24 | 10.96 | 8.36 | 9.94 |
| MSRA | 2 | 9.37 | 11.56 | 10.29 | 8.35 | 13.64 | 12.67 | 13.29 | 10.75 | 4.95 | 5.13 | 10.46 | 9.32 | 13.15 | 12.02 | 5.04 |
| MSRB2 | 1 | 10.28 | 8.74 | 8.62 | 8.66 | 14.57 | 13.62 | 8.84 | 8.83 | 8.96 | 8.88 | 9.51 | 8.64 | 14.10 | 8.83 | 8.92 |
| MST4 | 6 | 7.81 | 7.97 | 8.39 | 6.99 | 9.63 | 8.85 | 9.86 | 25.59 | 7.72 | 7.21 | 7.89 | 7.69 | 9.24 | 17.72 | 7.46 |
| MSTO1 | 6 | 9.40 | 9.98 | 11.02 | 11.31 | 10.96 | 11.06 | 9.45 | 8.79 | 9.11 | 8.93 | 9.69 | 11.16 | 11.01 | 9.12 | 9.02 |
| MT-ATP6 | 2 | 14.25 | 6.23 | 7.89 | 9.89 | 11.15 | 9.14 | 7.78 | 8.40 | 11.57 | 13.69 | 10.24 | 8.89 | 10.15 | 8.09 | 12.63 |
| MT-CO2 | 4 | 12.95 | 6.43 | 5.84 | 10.23 | 9.58 | 7.93 | 12.30 | 12.97 | 9.90 | 11.86 | 9.69 | 8.04 | 8.76 | 12.63 | 10.88 |
| MT1G | 2 | 0.19 | 0.13 | 0.35 | 0.54 | 1.65 | 0.81 | 0.85 | 94.98 | 0.34 | 0.17 | 0.16 | 0.44 | 1.23 | 47.92 | 0.25 |
| MTA1 | 16 | 9.74 | 10.30 | 9.30 | 8.95 | 9.77 | 10.43 | 11.65 | 10.51 | 9.48 | 9.86 | 10.02 | 9.12 | 10.10 | 11.08 | 9.67 |
| MTA2 | 32 | 9.39 | 10.01 | 11.41 | 10.77 | 10.19 | 9.77 | 10.21 | 9.76 | 8.97 | 9.53 | 9.70 | 11.09 | 9.98 | 9.98 | 9.25 |
| MTA3 | 4 | 10.14 | 10.42 | 9.63 | 9.33 | 10.29 | 11.85 | 9.64 | 9.49 | 10.48 | 8.72 | 10.28 | 9.48 | 11.07 | 9.57 | 9.60 |
| MTAP | 25 | 9.78 | 10.77 | 10.90 | 9.36 | 8.35 | 8.12 | 9.10 | 8.86 | 12.38 | 12.37 | 10.27 | 10.13 | 8.24 | 8.98 | 12.38 |
| MTCH1 | 5 | 16.04 | 7.92 | 7.27 | 10.94 | 7.84 | 7.09 | 7.90 | 7.92 | 12.27 | 14.79 | 11.98 | 9.11 | 7.46 | 7.91 | 13.53 |
| MTCH2 | 8 | 12.41 | 7.39 | 8.88 | 10.34 | 10.56 | 9.32 | 9.84 | 9.67 | 9.80 | 11.78 | 9.90 | 9.61 | 9.94 | 9.75 | 10.79 |
| MTDH | 27 | 10.96 | 6.15 | 7.37 | 10.72 | 9.46 | 8.20 | 13.23 | 12.68 | 9.36 | 11.87 | 8.55 | 9.04 | 8.83 | 12.96 | 10.61 |
| MTERFD1 | 5 | 9.75 | 9.24 | 9.54 | 9.73 | 8.43 | 8.19 | 13.11 | 11.04 | 10.74 | 10.23 | 9.49 | 9.64 | 8.31 | 12.08 | 10.48 |
| MTERFD2 | 2 | 10.59 | 9.56 | 9.22 | 12.47 | 9.75 | 8.46 | 9.94 | 11.08 | 9.63 | 9.31 | 10.07 | 10.84 | 9.11 | 10.51 | 9.47 |
| MTFR1 | 2 | 8.51 | 10.83 | 12.13 | 14.37 | 8.08 | 10.33 | 8.73 | 6.25 | 9.92 | 10.86 | 9.67 | 13.25 | 9.20 | 7.49 | 10.39 |
| MTFR1L | 3 | 11.10 | 12.73 | 7.88 | 10.00 | 8.95 | 11.97 | 8.45 | 6.22 | 12.27 | 10.42 | 11.92 | 8.94 | 10.46 | 7.34 | 11.35 |
| MTFR2 | 1 | 9.45 | 10.25 | 8.46 | 11.91 | 10.93 | 11.46 | 11.26 | 11.37 | 7.53 | 7.40 | 9.85 | 10.19 | 11.19 | 11.31 | 7.46 |
| MTG1 | 1 | 9.78 | 9.27 | 7.68 | 9.92 | 8.62 | 11.01 | 9.39 | 11.14 | 11.88 | 11.31 | 9.53 | 8.80 | 9.82 | 10.27 | 11.59 |
| MTG2 | 3 | 9.80 | 8.59 | 10.47 | 11.07 | 10.11 | 9.99 | 11.00 | 10.28 | 9.37 | 9.32 | 9.20 | 10.77 | 10.05 | 10.64 | 9.34 |
| MTHFD1 | 105 | 9.37 | 10.42 | 9.01 | 8.26 | 11.12 | 11.17 | 8.95 | 10.69 | 10.71 | 10.30 | 9.89 | 8.64 | 11.15 | 9.82 | 10.50 |
| MTHFD1L | 20 | 11.94 | 9.89 | 7.87 | 9.89 | 11.30 | 10.59 | 9.09 | 8.49 | 10.16 | 10.78 | 10.92 | 8.88 | 10.94 | 8.79 | 10.47 |
| MTHFD2 | 13 | 10.58 | 9.91 | 9.93 | 11.45 | 6.99 | 6.82 | 11.13 | 10.58 | 11.20 | 11.42 | 10.25 | 10.69 | 6.90 | 10.86 | 11.31 |
| MTHFS | 3 | 9.58 | 9.60 | 11.33 | 8.42 | 13.27 | 12.94 | 7.60 | 6.95 | 9.98 | 10.32 | 9.59 | 9.87 | 13.11 | 7.28 | 10.15 |
| MTHFSD | 3 | 11.13 | 12.13 | 8.29 | 8.02 | 11.74 | 11.54 | 9.19 | 7.92 | 10.71 | 9.34 | 11.63 | 8.15 | 11.64 | 8.55 | 10.03 |

| | | | | | | | | | | | | | | | | |
|---|---|---|---|---|---|---|---|---|---|---|---|---|---|---|---|---|
| MTIF2 | 7 | 10.95 | 8.68 | 9.58 | 10.24 | 9.87 | 9.62 | 11.71 | 10.95 | 9.16 | 9.25 | 9.81 | 9.91 | 9.74 | 11.33 | 9.20 |
| MTMR1 | 5 | 9.90 | 9.88 | 9.58 | 9.86 | 10.87 | 10.78 | 11.41 | 10.15 | 8.96 | 8.62 | 9.89 | 9.72 | 10.82 | 10.78 | 8.79 |
| MTMR10 | 1 | 9.27 | 10.19 | 11.83 | 11.89 | 9.43 | 9.49 | 7.93 | 7.54 | 10.84 | 11.59 | 9.73 | 11.86 | 9.46 | 7.74 | 11.22 |
| MTMR12 | 12 | 9.69 | 10.57 | 8.65 | 8.23 | 12.59 | 13.38 | 8.45 | 7.72 | 10.73 | 9.99 | 10.13 | 8.44 | 12.99 | 8.09 | 10.36 |
| MTMR14 | 1 | 8.91 | 8.50 | 13.30 | 11.11 | 9.82 | 7.99 | 10.54 | 8.79 | 12.25 | 8.77 | 8.71 | 12.21 | 8.91 | 9.67 | 10.51 |
| MTMR2 | 14 | 9.33 | 9.83 | 11.13 | 10.85 | 11.50 | 11.27 | 9.24 | 8.32 | 9.60 | 8.94 | 9.58 | 10.99 | 11.38 | 8.78 | 9.27 |
| MTMR6 | 5 | 9.19 | 9.95 | 11.83 | 11.66 | 12.35 | 12.15 | 8.36 | 8.82 | 8.06 | 7.64 | 9.57 | 11.74 | 12.25 | 8.59 | 7.85 |
| MTMR9 | 2 | 9.34 | 8.48 | 12.19 | 11.76 | 9.95 | 11.40 | 10.52 | 10.07 | 9.46 | 6.82 | 8.91 | 11.98 | 10.68 | 10.30 | 8.14 |
| MTO1 | 3 | 10.96 | 9.19 | 10.43 | 11.26 | 8.51 | 7.72 | 10.65 | 9.55 | 10.65 | 11.08 | 10.08 | 10.84 | 8.11 | 10.10 | 10.87 |
| MTOR | 29 | 10.40 | 9.48 | 8.75 | 9.58 | 9.98 | 10.00 | 9.72 | 9.52 | 11.49 | 11.08 | 9.94 | 9.17 | 9.99 | 9.62 | 11.29 |
| MTPAP | 5 | 9.46 | 8.43 | 9.92 | 11.89 | 9.16 | 7.41 | 11.22 | 9.94 | 10.76 | 11.80 | 8.95 | 10.91 | 8.29 | 10.58 | 11.28 |
| MTPN | 10 | 10.72 | 11.06 | 11.43 | 12.21 | 8.92 | 9.24 | 9.73 | 4.94 | 11.09 | 10.65 | 10.89 | 11.82 | 9.08 | 7.34 | 10.87 |
| MTR | 17 | 9.83 | 10.45 | 7.42 | 7.19 | 10.98 | 10.89 | 9.91 | 11.70 | 10.80 | 10.84 | 10.14 | 7.31 | 10.93 | 10.80 | 10.82 |
| MTRF1L | 1 | 9.10 | 10.27 | 8.33 | 13.72 | 9.03 | 10.13 | 9.45 | 9.72 | 9.78 | 10.46 | 9.69 | 11.03 | 9.58 | 9.59 | 10.12 |
| MTRR | 12 | 10.05 | 10.05 | 8.77 | 8.70 | 8.07 | 8.40 | 13.29 | 12.62 | 10.19 | 9.85 | 10.05 | 8.74 | 8.23 | 12.96 | 10.02 |
| MTSS1L | 2 | 11.72 | 12.88 | 8.36 | 8.17 | 10.44 | 11.46 | 6.04 | 6.76 | 12.20 | 11.98 | 12.30 | 8.26 | 10.95 | 6.40 | 12.09 |
| MTUS1 | 3 | 10.42 | 11.99 | 7.95 | 8.52 | 13.99 | 13.32 | 11.28 | 10.31 | 6.26 | 5.97 | 11.20 | 8.23 | 13.66 | 10.79 | 6.11 |
| MTX1 | 6 | 11.07 | 8.16 | 11.20 | 12.44 | 10.15 | 9.32 | 8.67 | 8.34 | 9.88 | 10.76 | 9.62 | 11.82 | 9.74 | 8.51 | 10.32 |
| MTX2 | 5 | 8.49 | 9.32 | 12.42 | 10.00 | 10.42 | 10.17 | 11.16 | 9.68 | 9.03 | 9.31 | 8.90 | 11.21 | 10.29 | 10.42 | 9.17 |
| MTX3 | 9 | 9.94 | 10.80 | 9.25 | 8.86 | 9.15 | 8.95 | 11.09 | 10.92 | 10.55 | 10.47 | 10.37 | 9.06 | 9.05 | 11.01 | 10.51 |
| MUC1 | 7 | 8.50 | 7.48 | 3.12 | 3.53 | 23.89 | 26.45 | 6.84 | 5.56 | 7.16 | 7.47 | 7.99 | 3.33 | 25.17 | 6.20 | 7.32 |
| MUC16 | 14 | 15.12 | 14.71 | 3.18 | 2.47 | 16.41 | 16.95 | 5.32 | 4.54 | 10.77 | 10.53 | 14.92 | 2.83 | 16.68 | 4.93 | 10.65 |
| MUL1 | 1 | 10.69 | 9.88 | 9.45 | 9.99 | 9.87 | 9.71 | 11.08 | 7.68 | 11.39 | 10.26 | 10.29 | 9.72 | 9.79 | 9.38 | 10.83 |
| MUT | 8 | 12.25 | 10.24 | 7.53 | 9.89 | 11.30 | 11.07 | 9.90 | 9.81 | 8.94 | 9.07 | 11.25 | 8.71 | 11.18 | 9.85 | 9.00 |
| MVB12A | 3 | 8.64 | 8.85 | 15.12 | 13.53 | 8.71 | 8.69 | 8.70 | 8.89 | 9.60 | 9.27 | 8.75 | 14.33 | 8.70 | 8.80 | 9.44 |
| MVD | 4 | 10.63 | 12.22 | 8.31 | 8.47 | 13.12 | 13.61 | 6.71 | 5.75 | 10.48 | 10.69 | 11.42 | 8.39 | 13.37 | 6.23 | 10.59 |
| MVK | 7 | 11.52 | 11.95 | 10.92 | 10.97 | 10.00 | 10.73 | 7.80 | 6.70 | 9.75 | 9.66 | 11.74 | 10.94 | 10.36 | 7.25 | 9.70 |
| MVP | 36 | 8.30 | 9.01 | 12.87 | 12.38 | 13.62 | 13.69 | 9.22 | 8.21 | 6.62 | 6.08 | 8.66 | 12.62 | 13.65 | 8.72 | 6.35 |
| MXRA7 | 1 | 10.13 | 7.68 | 16.27 | 19.97 | 4.95 | 5.08 | 11.37 | 4.42 | 9.89 | 10.25 | 8.90 | 18.12 | 5.02 | 7.89 | 10.07 |
| MYADM | 3 | 10.16 | 9.06 | 10.11 | 11.89 | 7.56 | 7.04 | 9.60 | 7.36 | 12.11 | 15.11 | 9.61 | 11.00 | 7.30 | 8.48 | 13.61 |
| MYBBP1A | 43 | 9.71 | 9.15 | 11.70 | 12.27 | 7.10 | 6.89 | 9.80 | 9.77 | 11.30 | 12.31 | 9.43 | 11.99 | 6.99 | 9.78 | 11.80 |
| MYBL2 | 1 | 9.07 | 9.80 | 9.99 | 8.70 | 10.20 | 9.70 | 10.56 | 11.27 | 11.04 | 9.66 | 9.44 | 9.34 | 9.95 | 10.91 | 10.35 |
| MYC | 2 | 9.44 | 9.48 | 10.28 | 10.47 | 8.22 | 9.46 | 11.37 | 9.98 | 12.18 | 9.11 | 9.46 | 10.38 | 8.84 | 10.68 | 10.65 |
| MYCBP | 9 | 11.88 | 12.54 | 10.98 | 10.91 | 8.00 | 9.18 | 10.13 | 3.53 | 10.62 | 12.23 | 12.21 | 10.95 | 8.59 | 6.83 | 11.42 |
| MYCBP2 | 24 | 9.76 | 10.29 | 9.95 | 10.71 | 9.24 | 9.90 | 11.41 | 11.28 | 8.73 | 8.73 | 10.02 | 10.33 | 9.57 | 11.34 | 8.73 |
| MYD88 | 3 | 9.80 | 10.64 | 7.80 | 7.81 | 13.26 | 14.14 | 9.94 | 9.23 | 8.58 | 8.79 | 10.22 | 7.81 | 13.70 | 9.58 | 8.68 |
| MYH10 | 93 | 12.88 | 14.50 | 11.20 | 10.37 | 9.89 | 9.88 | 2.91 | 6.33 | 11.28 | 10.78 | 13.69 | 10.78 | 9.88 | 4.62 | 11.03 |
| MYH11 | 21 | 8.44 | 10.43 | 15.72 | 16.41 | 6.14 | 6.12 | 6.28 | 6.08 | 12.51 | 11.86 | 9.44 | 16.06 | 6.13 | 6.18 | 12.19 |
| MYH14 | 21 | 10.60 | 11.08 | 14.96 | 14.84 | 6.62 | 6.78 | 5.84 | 5.67 | 12.18 | 11.43 | 10.84 | 14.90 | 6.70 | 5.76 | 11.80 |
| MYH9 | 384 | 8.87 | 10.04 | 16.24 | 14.70 | 6.37 | 6.19 | 6.14 | 7.81 | 11.99 | 11.63 | 9.46 | 15.47 | 6.28 | 6.98 | 11.81 |
| MYL12B | 40 | 7.81 | 10.81 | 17.20 | 13.83 | 6.92 | 6.68 | 6.73 | 5.50 | 12.04 | 12.48 | 9.31 | 15.52 | 6.80 | 6.11 | 12.26 |
| MYL6 | 22 | 10.00 | 11.32 | 13.52 | 13.99 | 8.31 | 8.31 | 6.36 | 4.51 | 12.07 | 11.61 | 10.66 | 13.76 | 8.31 | 5.43 | 11.84 |

| | | | | | | | | | | | | | | | | |
|---|---|---|---|---|---|---|---|---|---|---|---|---|---|---|---|---|
| MYL6B | 16 | 9.46 | 10.17 | 13.29 | 11.69 | 8.67 | 8.66 | 8.22 | 9.42 | 10.48 | 9.94 | 9.82 | 12.49 | 8.67 | 8.82 | 10.21 |
| MYL9 | 7 | 9.14 | 9.53 | 21.77 | 19.93 | 4.40 | 4.32 | 3.44 | 4.05 | 12.03 | 11.39 | 9.33 | 20.85 | 4.36 | 3.74 | 11.71 |
| MYLK | 6 | 15.62 | 17.43 | 10.77 | 9.55 | 6.61 | 7.33 | 6.48 | 10.26 | 8.30 | 7.64 | 16.53 | 10.16 | 6.97 | 8.37 | 7.97 |
| MYO10 | 25 | 7.69 | 8.30 | 9.84 | 10.13 | 10.73 | 10.98 | 11.21 | 12.17 | 9.53 | 9.41 | 8.00 | 9.98 | 10.86 | 11.69 | 9.47 |
| MYO18A | 18 | 9.79 | 10.91 | 9.55 | 9.41 | 10.49 | 10.53 | 10.04 | 9.10 | 10.60 | 9.58 | 10.35 | 9.48 | 10.51 | 9.57 | 10.09 |
| MYO19 | 6 | 10.43 | 9.22 | 8.56 | 10.17 | 9.75 | 10.11 | 9.81 | 10.35 | 10.63 | 10.97 | 9.82 | 9.36 | 9.93 | 10.08 | 10.80 |
| MYO1B | 35 | 9.92 | 9.73 | 12.56 | 12.70 | 7.03 | 7.04 | 11.73 | 12.79 | 8.21 | 8.29 | 9.82 | 12.63 | 7.04 | 12.26 | 8.25 |
| MYO1C | 45 | 9.77 | 10.76 | 12.35 | 12.48 | 9.33 | 9.37 | 5.31 | 5.10 | 12.78 | 12.74 | 10.27 | 12.42 | 9.35 | 5.21 | 12.76 |
| MYO1E | 38 | 10.25 | 10.85 | 9.52 | 9.27 | 12.56 | 12.81 | 7.40 | 7.02 | 10.20 | 10.13 | 10.55 | 9.39 | 12.68 | 7.21 | 10.16 |
| MYO1H | 1 | 11.52 | 8.42 | 7.60 | 9.01 | 13.85 | 13.21 | 11.87 | 10.20 | 7.97 | 6.36 | 9.97 | 8.31 | 13.53 | 11.03 | 7.16 |
| MYO5A | 6 | 10.75 | 11.13 | 12.84 | 13.19 | 6.78 | 7.06 | 6.13 | 5.21 | 13.97 | 12.92 | 10.94 | 13.02 | 6.92 | 5.67 | 13.45 |
| MYO6 | 7 | 8.81 | 8.76 | 9.94 | 9.30 | 14.49 | 14.25 | 9.45 | 12.29 | 5.94 | 6.76 | 8.79 | 9.62 | 14.37 | 10.87 | 6.35 |
| MYO9B | 12 | 9.95 | 10.24 | 10.36 | 10.36 | 7.05 | 7.41 | 12.56 | 11.70 | 10.17 | 10.21 | 10.09 | 10.36 | 7.23 | 12.13 | 10.19 |
| MYOF | 94 | 13.00 | 8.86 | 9.76 | 13.53 | 9.13 | 8.60 | 6.25 | 6.25 | 11.73 | 12.89 | 10.93 | 11.65 | 8.86 | 6.25 | 12.31 |
| MYOM1 | 4 | 9.63 | 12.67 | 11.27 | 9.59 | 9.33 | 9.70 | 10.63 | 10.05 | 8.94 | 8.20 | 11.15 | 10.43 | 9.52 | 10.34 | 8.57 |
| MYPN | 3 | 13.86 | 16.10 | 5.57 | 5.61 | 17.02 | 16.25 | 6.45 | 5.29 | 7.33 | 6.52 | 14.98 | 5.59 | 16.64 | 5.87 | 6.92 |
| MZT1 | 1 | 10.70 | 10.37 | 9.26 | 10.35 | 9.34 | 12.79 | 9.90 | 4.89 | 12.42 | 10.01 | 10.53 | 9.80 | 11.06 | 7.39 | 11.21 |
| MZT2B | 3 | 9.57 | 10.20 | 10.52 | 10.36 | 10.83 | 11.72 | 9.73 | 9.06 | 9.14 | 8.87 | 9.89 | 10.44 | 11.27 | 9.39 | 9.01 |
| N4BP1 | 3 | 10.99 | 9.01 | 10.54 | 8.80 | 10.46 | 8.65 | 13.24 | 9.20 | 10.10 | 9.02 | 10.00 | 9.67 | 9.56 | 11.22 | 9.56 |
| N6AMT1 | 2 | 9.72 | 10.27 | 7.47 | 6.49 | 16.56 | 15.71 | 9.56 | 7.46 | 8.51 | 8.24 | 10.00 | 6.98 | 16.14 | 8.51 | 8.38 |
| NAA10 | 19 | 9.90 | 10.42 | 10.51 | 10.02 | 9.34 | 9.84 | 11.12 | 10.15 | 9.60 | 9.09 | 10.16 | 10.27 | 9.59 | 10.63 | 9.35 |
| NAA15 | 49 | 9.24 | 9.73 | 10.74 | 9.31 | 9.38 | 9.11 | 10.99 | 11.68 | 10.04 | 9.78 | 9.49 | 10.03 | 9.25 | 11.33 | 9.91 |
| NAA16 | 1 | 10.45 | 9.76 | 10.85 | 13.53 | 8.60 | 10.38 | 10.94 | 9.08 | 6.85 | 9.56 | 10.10 | 12.19 | 9.49 | 10.01 | 8.21 |
| NAA20 | 2 | 10.07 | 10.31 | 10.55 | 10.45 | 10.91 | 9.76 | 9.88 | 9.43 | 9.38 | 9.26 | 10.19 | 10.50 | 10.33 | 9.66 | 9.32 |
| NAA25 | 10 | 9.40 | 10.18 | 9.83 | 9.49 | 10.31 | 10.32 | 10.50 | 11.05 | 9.37 | 9.55 | 9.79 | 9.66 | 10.31 | 10.77 | 9.46 |
| NAA30 | 4 | 9.59 | 9.86 | 10.50 | 10.60 | 9.64 | 10.30 | 9.76 | 8.85 | 10.67 | 10.22 | 9.73 | 10.55 | 9.97 | 9.31 | 10.45 |
| NAA35 | 10 | 9.49 | 10.45 | 9.88 | 10.55 | 9.28 | 10.12 | 10.29 | 9.54 | 10.43 | 9.96 | 9.97 | 10.21 | 9.70 | 9.91 | 10.20 |
| NAA38 | 5 | 9.81 | 10.73 | 9.09 | 9.17 | 10.04 | 11.15 | 11.55 | 10.59 | 9.22 | 8.65 | 10.27 | 9.13 | 10.60 | 11.07 | 8.93 |
| NAA50 | 19 | 9.30 | 9.35 | 10.84 | 9.91 | 8.93 | 8.64 | 10.90 | 13.63 | 9.63 | 8.85 | 9.32 | 10.37 | 8.79 | 12.27 | 9.24 |
| NAB1 | 1 | 10.64 | 12.46 | 8.36 | 9.71 | 11.44 | 11.85 | 7.92 | 10.07 | 8.68 | 8.86 | 11.55 | 9.03 | 11.65 | 9.00 | 8.77 |
| NAB2 | 3 | 9.52 | 11.02 | 8.71 | 8.01 | 8.83 | 9.98 | 14.27 | 13.16 | 8.50 | 8.00 | 10.27 | 8.36 | 9.40 | 13.72 | 8.25 |
| NACA | 30 | 9.49 | 11.08 | 9.40 | 9.74 | 9.23 | 10.17 | 11.07 | 10.82 | 9.54 | 9.46 | 10.29 | 9.57 | 9.70 | 10.95 | 9.50 |
| NACAP1 | 2 | 9.76 | 11.00 | 9.85 | 10.00 | 9.02 | 9.78 | 10.61 | 10.37 | 9.79 | 9.81 | 10.38 | 9.93 | 9.40 | 10.49 | 9.80 |
| NACC1 | 3 | 8.54 | 9.96 | 10.61 | 11.12 | 9.04 | 9.70 | 11.84 | 11.32 | 9.07 | 8.81 | 9.25 | 10.86 | 9.37 | 11.58 | 8.94 |
| NACC2 | 1 | 9.98 | 12.63 | 10.77 | 10.48 | 9.99 | 11.32 | 6.62 | 6.33 | 11.04 | 10.85 | 11.30 | 10.63 | 10.66 | 6.47 | 10.94 |
| NADK | 4 | 10.18 | 11.30 | 10.43 | 9.48 | 8.85 | 8.32 | 10.51 | 10.06 | 10.61 | 10.24 | 10.74 | 9.95 | 8.59 | 10.29 | 10.43 |
| NADK2 | 15 | 9.76 | 8.66 | 8.28 | 9.38 | 10.15 | 10.07 | 12.81 | 11.72 | 9.44 | 9.73 | 9.21 | 8.83 | 10.11 | 12.27 | 9.59 |
| NADSYN1 | 1 | 10.99 | 9.56 | 8.46 | 7.68 | 15.50 | 16.64 | 6.28 | 11.99 | 6.98 | 5.93 | 10.27 | 8.07 | 16.07 | 9.13 | 6.45 |
| NAE1 | 10 | 10.12 | 10.43 | 7.59 | 7.54 | 11.72 | 11.51 | 10.95 | 10.22 | 10.26 | 9.65 | 10.28 | 7.57 | 11.62 | 10.59 | 9.96 |
| NAGA | 1 | 9.30 | 11.90 | 9.30 | 14.02 | 9.17 | 10.71 | 9.14 | 8.10 | 8.68 | 9.67 | 10.60 | 11.66 | 9.94 | 8.62 | 9.18 |
| NAGK | 8 | 10.17 | 11.81 | 10.94 | 10.37 | 9.78 | 10.16 | 7.56 | 7.52 | 10.95 | 10.74 | 10.99 | 10.65 | 9.97 | 7.54 | 10.85 |
| NAGLU | 2 | 10.00 | 9.93 | 11.60 | 11.68 | 12.91 | 11.51 | 7.36 | 7.21 | 9.21 | 8.60 | 9.97 | 11.64 | 12.21 | 7.28 | 8.90 |

| | | | | | | | | | | | | | | | | |
|---|---|---|---|---|---|---|---|---|---|---|---|---|---|---|---|---|
| NAMPT | 29 | 7.66 | 8.05 | 10.41 | 9.34 | 7.51 | 7.51 | 17.38 | 17.43 | 7.57 | 7.15 | 7.86 | 9.87 | 7.51 | 17.40 | 7.36 |
| NANS | 18 | 10.29 | 10.64 | 9.06 | 8.13 | 11.20 | 10.85 | 9.07 | 10.01 | 10.60 | 10.13 | 10.47 | 8.60 | 11.03 | 9.54 | 10.37 |
| NAP1L1 | 33 | 7.95 | 10.15 | 12.59 | 10.84 | 9.38 | 9.35 | 10.19 | 9.53 | 10.21 | 9.80 | 9.05 | 11.72 | 9.36 | 9.86 | 10.01 |
| NAP1L4 | 15 | 7.91 | 8.81 | 9.64 | 7.82 | 10.08 | 10.21 | 9.41 | 19.80 | 8.41 | 7.92 | 8.36 | 8.73 | 10.14 | 14.61 | 8.17 |
| NAPA | 21 | 9.70 | 10.19 | 9.90 | 8.46 | 10.11 | 9.85 | 9.35 | 13.08 | 9.67 | 9.69 | 9.94 | 9.18 | 9.98 | 11.22 | 9.68 |
| NAPG | 19 | 9.66 | 10.89 | 11.54 | 10.71 | 12.19 | 12.57 | 6.66 | 5.94 | 10.22 | 9.63 | 10.28 | 11.12 | 12.38 | 6.30 | 9.93 |
| NAPRT1 | 8 | 10.23 | 11.38 | 4.81 | 4.96 | 16.56 | 16.43 | 7.70 | 7.26 | 10.27 | 10.38 | 10.81 | 4.89 | 16.50 | 7.48 | 10.33 |
| NARF | 6 | 9.96 | 11.23 | 11.56 | 10.81 | 7.96 | 7.99 | 11.84 | 10.63 | 9.10 | 8.91 | 10.60 | 11.19 | 7.98 | 11.24 | 9.00 |
| NARFL | 1 | 10.30 | 8.99 | 10.00 | 11.94 | 8.24 | 10.36 | 9.95 | 8.61 | 10.08 | 11.53 | 9.64 | 10.97 | 9.30 | 9.28 | 10.80 |
| NARG2 | 3 | 8.24 | 9.98 | 9.88 | 10.68 | 10.93 | 10.27 | 12.10 | 11.91 | 8.04 | 7.97 | 9.11 | 10.28 | 10.60 | 12.01 | 8.00 |
| NARS | 32 | 9.11 | 10.17 | 11.90 | 10.78 | 10.63 | 10.09 | 7.08 | 8.46 | 11.32 | 10.47 | 9.64 | 11.34 | 10.36 | 7.77 | 10.90 |
| NARS2 | 1 | 9.47 | 11.21 | 7.32 | 8.71 | 9.97 | 12.28 | 9.83 | 13.08 | 9.60 | 8.52 | 10.34 | 8.02 | 11.12 | 11.46 | 9.06 |
| NASP | 68 | 9.30 | 10.21 | 9.35 | 8.49 | 9.94 | 10.21 | 12.21 | 12.63 | 8.98 | 8.67 | 9.75 | 8.92 | 10.08 | 12.42 | 8.83 |
| NAT1 | 2 | 10.56 | 10.52 | 10.86 | 9.90 | 11.99 | 11.33 | 11.13 | 10.09 | 7.34 | 6.28 | 10.54 | 10.38 | 11.66 | 10.61 | 6.81 |
| NAT10 | 27 | 9.30 | 8.97 | 10.20 | 9.75 | 10.06 | 9.78 | 12.08 | 10.70 | 9.35 | 9.80 | 9.14 | 9.98 | 9.92 | 11.39 | 9.58 |
| NAT6 | 3 | 8.21 | 9.85 | 9.64 | 9.96 | 9.63 | 9.54 | 11.76 | 11.86 | 9.55 | 9.99 | 9.03 | 9.80 | 9.58 | 11.81 | 9.77 |
| NAV1 | 7 | 8.23 | 8.84 | 9.09 | 8.90 | 10.75 | 11.10 | 9.51 | 8.67 | 12.61 | 12.30 | 8.53 | 9.00 | 10.93 | 9.09 | 12.46 |
| NAV2 | 8 | 8.58 | 9.78 | 12.26 | 11.80 | 11.98 | 11.25 | 9.00 | 7.38 | 8.70 | 9.27 | 9.18 | 12.03 | 11.62 | 8.19 | 8.99 |
| NBAS | 9 | 9.76 | 8.90 | 9.60 | 10.97 | 10.49 | 10.83 | 9.46 | 10.25 | 9.87 | 9.86 | 9.33 | 10.29 | 10.66 | 9.86 | 9.87 |
| NBEA | 1 | 9.62 | 8.63 | 8.03 | 7.48 | 11.29 | 11.17 | 9.02 | 19.00 | 8.11 | 7.65 | 9.12 | 7.76 | 11.23 | 14.01 | 7.88 |
| NBEAL1 | 1 | 8.00 | 9.39 | 10.17 | 12.40 | 10.65 | 9.94 | 7.87 | 8.63 | 11.93 | 11.03 | 8.69 | 11.28 | 10.29 | 8.25 | 11.48 |
| NBN | 14 | 9.01 | 10.34 | 9.43 | 9.29 | 8.58 | 9.45 | 14.28 | 13.38 | 8.20 | 8.05 | 9.67 | 9.36 | 9.01 | 13.83 | 8.13 |
| NBR1 | 1 | 11.08 | 10.57 | 11.42 | 11.51 | 11.82 | 12.17 | 7.18 | 6.07 | 10.21 | 7.96 | 10.82 | 11.47 | 12.00 | 6.63 | 9.09 |
| NCAPD2 | 54 | 9.04 | 9.54 | 10.99 | 10.59 | 8.82 | 8.69 | 10.78 | 11.00 | 10.21 | 10.32 | 9.29 | 10.79 | 8.76 | 10.89 | 10.26 |
| NCAPD3 | 11 | 9.78 | 9.57 | 9.98 | 9.63 | 9.22 | 8.67 | 10.47 | 9.67 | 11.20 | 11.79 | 9.68 | 9.81 | 8.95 | 10.07 | 11.50 |
| NCAPG | 27 | 9.58 | 9.29 | 11.12 | 10.06 | 9.21 | 8.46 | 10.80 | 10.14 | 10.39 | 10.95 | 9.44 | 10.59 | 8.84 | 10.47 | 10.67 |
| NCAPG2 | 9 | 10.24 | 9.63 | 9.47 | 9.45 | 8.10 | 7.98 | 10.80 | 10.90 | 11.62 | 11.80 | 9.94 | 9.46 | 8.04 | 10.85 | 11.71 |
| NCAPH | 22 | 9.17 | 9.69 | 13.00 | 11.21 | 8.74 | 8.58 | 10.50 | 9.53 | 9.78 | 9.81 | 9.43 | 12.10 | 8.66 | 10.02 | 9.80 |
| NCAPH2 | 6 | 9.71 | 10.95 | 10.69 | 9.75 | 8.05 | 9.09 | 9.69 | 9.47 | 11.86 | 10.74 | 10.33 | 10.22 | 8.57 | 9.58 | 11.30 |
| NCBP1 | 18 | 9.60 | 8.72 | 9.68 | 6.86 | 11.06 | 9.87 | 9.43 | 12.40 | 11.39 | 10.97 | 9.16 | 8.27 | 10.47 | 10.92 | 11.18 |
| NCBP2 | 4 | 9.13 | 10.22 | 10.29 | 9.46 | 10.05 | 10.75 | 11.03 | 10.50 | 9.67 | 8.90 | 9.67 | 9.88 | 10.40 | 10.77 | 9.28 |
| NCDN | 8 | 10.86 | 11.15 | 10.02 | 10.47 | 8.25 | 8.41 | 8.86 | 9.37 | 11.48 | 11.13 | 11.00 | 10.25 | 8.33 | 9.12 | 11.31 |
| NCEH1 | 2 | 11.65 | 6.37 | 7.37 | 8.44 | 10.71 | 8.36 | 8.73 | 7.46 | 13.87 | 17.03 | 9.01 | 7.90 | 9.54 | 8.09 | 15.45 |
| NCK1 | 12 | 10.20 | 11.66 | 10.26 | 9.82 | 9.59 | 10.16 | 10.08 | 8.89 | 9.57 | 9.78 | 10.93 | 10.04 | 9.87 | 9.48 | 9.67 |
| NCKAP1 | 26 | 9.97 | 9.67 | 10.32 | 10.52 | 10.61 | 10.61 | 9.02 | 8.44 | 10.55 | 10.29 | 9.82 | 10.42 | 10.61 | 8.73 | 10.42 |
| NCKAP5L | 2 | 9.82 | 10.30 | 14.49 | 14.36 | 8.95 | 8.38 | 7.06 | 6.21 | 10.45 | 9.96 | 10.06 | 14.43 | 8.67 | 6.63 | 10.21 |
| NCKIPSD | 8 | 10.72 | 11.78 | 7.37 | 7.18 | 11.78 | 11.46 | 8.85 | 8.48 | 11.32 | 11.05 | 11.25 | 7.28 | 11.62 | 8.67 | 11.18 |
| NCL | 146 | 8.83 | 10.11 | 11.15 | 10.67 | 9.02 | 9.22 | 11.96 | 11.69 | 8.88 | 8.48 | 9.47 | 10.91 | 9.12 | 11.82 | 8.68 |
| NCLN | 13 | 11.40 | 6.71 | 7.96 | 11.86 | 9.45 | 8.53 | 9.87 | 9.36 | 11.19 | 13.67 | 9.06 | 9.91 | 8.99 | 9.62 | 12.43 |
| NCOA2 | 3 | 9.51 | 11.25 | 9.48 | 10.42 | 9.72 | 11.19 | 10.46 | 10.10 | 8.94 | 8.92 | 10.38 | 9.95 | 10.46 | 10.28 | 8.93 |
| NCOA3 | 2 | 9.76 | 9.49 | 11.86 | 10.53 | 8.92 | 10.30 | 11.14 | 9.04 | 10.28 | 8.69 | 9.62 | 11.19 | 9.61 | 10.09 | 9.49 |
| NCOA5 | 20 | 9.33 | 10.37 | 10.54 | 10.75 | 10.23 | 10.52 | 10.85 | 11.01 | 8.42 | 7.97 | 9.85 | 10.65 | 10.37 | 10.93 | 8.20 |

| | | | | | | | | | | | | | | | | |
|---|---|---|---|---|---|---|---|---|---|---|---|---|---|---|---|---|
| NCOA6 | 2 | 7.34 | 18.06 | 9.57 | 10.43 | 9.08 | 9.58 | 9.79 | 8.71 | 8.40 | 9.03 | 12.70 | 10.00 | 9.33 | 9.25 | 8.72 |
| NCOA7 | 11 | 10.33 | 11.21 | 9.30 | 9.30 | 11.87 | 12.03 | 7.51 | 6.91 | 11.09 | 10.44 | 10.77 | 9.30 | 11.95 | 7.21 | 10.77 |
| NCOR1 | 17 | 9.58 | 11.01 | 10.77 | 10.28 | 8.69 | 9.06 | 11.17 | 10.58 | 9.42 | 9.43 | 10.30 | 10.53 | 8.87 | 10.88 | 9.43 |
| NCOR2 | 39 | 9.12 | 10.15 | 13.03 | 12.05 | 9.48 | 9.52 | 9.39 | 8.47 | 9.49 | 9.29 | 9.64 | 12.54 | 9.50 | 8.93 | 9.39 |
| NCR3LG1 | 1 | 13.41 | 8.75 | 7.19 | 8.87 | 14.16 | 12.46 | 7.44 | 7.98 | 8.42 | 11.30 | 11.08 | 8.03 | 13.31 | 7.71 | 9.86 |
| NCS1 | 5 | 10.28 | 10.73 | 13.74 | 12.25 | 6.76 | 7.30 | 9.91 | 8.42 | 10.32 | 10.29 | 10.51 | 13.00 | 7.03 | 9.16 | 10.31 |
| NCSTN | 10 | 11.66 | 8.41 | 8.03 | 10.47 | 12.82 | 12.03 | 8.70 | 7.98 | 9.40 | 10.50 | 10.04 | 9.25 | 12.42 | 8.34 | 9.95 |
| NDC1 | 5 | 12.13 | 6.14 | 8.33 | 11.62 | 7.61 | 6.09 | 11.46 | 11.13 | 10.87 | 14.62 | 9.14 | 9.98 | 6.85 | 11.30 | 12.74 |
| NDC80 | 17 | 10.24 | 11.09 | 11.84 | 11.08 | 11.46 | 11.14 | 8.36 | 7.83 | 8.48 | 8.50 | 10.66 | 11.46 | 11.30 | 8.09 | 8.49 |
| NDE1 | 5 | 8.39 | 9.57 | 10.23 | 10.16 | 10.14 | 11.16 | 9.91 | 8.96 | 10.83 | 10.65 | 8.98 | 10.20 | 10.65 | 9.43 | 10.74 |
| NDEL1 | 3 | 9.90 | 11.11 | 9.20 | 10.19 | 9.85 | 10.81 | 9.91 | 8.97 | 10.09 | 9.99 | 10.50 | 9.69 | 10.33 | 9.44 | 10.04 |
| NDNL2 | 4 | 8.66 | 9.81 | 9.50 | 8.74 | 10.62 | 11.08 | 11.14 | 9.66 | 10.49 | 10.31 | 9.23 | 9.12 | 10.85 | 10.40 | 10.40 |
| NDOR1 | 3 | 9.75 | 10.38 | 7.55 | 6.53 | 16.04 | 15.75 | 8.20 | 6.68 | 10.01 | 9.11 | 10.06 | 7.04 | 15.90 | 7.44 | 9.56 |
| NDRG1 | 8 | 6.98 | 6.20 | 6.00 | 5.15 | 7.00 | 6.28 | 15.74 | 33.63 | 7.13 | 5.90 | 6.59 | 5.58 | 6.64 | 24.68 | 6.51 |
| NDRG2 | 1 | 8.14 | 10.64 | 8.94 | 5.78 | 10.76 | 13.93 | 14.47 | 16.66 | 6.16 | 4.52 | 9.39 | 7.36 | 12.35 | 15.56 | 5.34 |
| NDRG3 | 10 | 9.71 | 11.27 | 9.88 | 10.09 | 8.85 | 9.51 | 7.96 | 13.81 | 9.33 | 9.59 | 10.49 | 9.99 | 9.18 | 10.89 | 9.46 |
| NDUFA10 | 8 | 9.67 | 7.27 | 8.63 | 8.51 | 11.20 | 9.67 | 11.85 | 17.98 | 7.31 | 7.88 | 8.47 | 8.57 | 10.44 | 14.92 | 7.60 |
| NDUFA12 | 1 | 10.08 | 8.09 | 9.08 | 9.96 | 11.63 | 10.82 | 10.38 | 9.74 | 10.00 | 10.22 | 9.09 | 9.52 | 11.22 | 10.06 | 10.11 |
| NDUFA13 | 4 | 11.67 | 6.15 | 9.61 | 13.12 | 10.57 | 8.59 | 12.01 | 10.53 | 8.04 | 9.72 | 8.91 | 11.36 | 9.58 | 11.27 | 8.88 |
| NDUFA2 | 6 | 10.54 | 9.12 | 9.59 | 10.46 | 10.30 | 9.68 | 10.90 | 9.27 | 10.25 | 9.88 | 9.83 | 10.03 | 9.99 | 10.09 | 10.07 |
| NDUFA3 | 3 | 10.70 | 7.46 | 9.64 | 10.46 | 12.14 | 10.59 | 12.02 | 10.52 | 8.54 | 7.95 | 9.08 | 10.05 | 11.36 | 11.27 | 8.24 |
| NDUFA4 | 11 | 11.60 | 7.69 | 7.48 | 9.58 | 9.44 | 8.82 | 12.76 | 14.99 | 8.32 | 9.31 | 9.65 | 8.53 | 9.13 | 13.88 | 8.82 |
| NDUFA5 | 3 | 10.23 | 8.89 | 10.75 | 10.35 | 10.65 | 9.51 | 12.92 | 10.95 | 8.27 | 7.48 | 9.56 | 10.55 | 10.08 | 11.93 | 7.88 |
| NDUFA6 | 7 | 9.35 | 8.43 | 13.55 | 8.73 | 10.97 | 8.87 | 11.92 | 8.72 | 9.62 | 9.84 | 8.89 | 11.14 | 9.92 | 10.32 | 9.73 |
| NDUFA7 | 3 | 8.62 | 8.22 | 10.03 | 11.03 | 11.75 | 11.18 | 12.51 | 13.00 | 6.75 | 6.91 | 8.42 | 10.53 | 11.47 | 12.75 | 6.83 |
| NDUFA8 | 3 | 8.25 | 6.98 | 8.31 | 8.68 | 10.97 | 9.65 | 10.03 | 24.07 | 6.31 | 6.76 | 7.62 | 8.49 | 10.31 | 17.05 | 6.53 |
| NDUFA9 | 9 | 11.82 | 8.53 | 7.91 | 10.33 | 10.43 | 9.81 | 11.91 | 11.07 | 8.71 | 9.48 | 10.17 | 9.12 | 10.12 | 11.49 | 9.09 |
| NDUFAB1 | 3 | 10.41 | 6.95 | 16.81 | 2.34 | 16.99 | 10.44 | 12.94 | 1.35 | 10.68 | 11.09 | 8.68 | 9.58 | 13.72 | 7.14 | 10.89 |
| NDUFAF1 | 5 | 12.11 | 10.09 | 11.44 | 14.20 | 9.79 | 9.68 | 10.70 | 10.20 | 5.69 | 6.11 | 11.10 | 12.82 | 9.74 | 10.45 | 5.90 |
| NDUFAF2 | 11 | 9.50 | 8.57 | 11.01 | 11.13 | 8.79 | 8.02 | 14.71 | 13.24 | 7.56 | 7.46 | 9.03 | 11.07 | 8.41 | 13.98 | 7.51 |
| NDUFAF3 | 5 | 10.64 | 9.14 | 9.10 | 9.56 | 10.85 | 10.97 | 10.77 | 10.37 | 9.32 | 9.28 | 9.89 | 9.33 | 10.91 | 10.57 | 9.30 |
| NDUFAF4 | 5 | 10.51 | 8.06 | 9.30 | 11.51 | 10.08 | 8.97 | 12.02 | 11.12 | 9.03 | 9.39 | 9.29 | 10.40 | 9.53 | 11.57 | 9.21 |
| NDUFAF5 | 1 | 8.63 | 8.83 | 10.54 | 13.44 | 9.93 | 8.62 | 10.99 | 10.28 | 8.94 | 9.80 | 8.73 | 11.99 | 9.28 | 10.63 | 9.37 |
| NDUFAF7 | 7 | 9.25 | 9.58 | 11.45 | 12.87 | 8.56 | 7.63 | 10.94 | 10.72 | 9.15 | 9.86 | 9.41 | 12.16 | 8.09 | 10.83 | 9.50 |
| NDUFB1 | 2 | 8.45 | 8.24 | 9.22 | 8.56 | 11.85 | 8.56 | 11.81 | 11.98 | 8.85 | 12.49 | 8.34 | 8.89 | 10.20 | 11.89 | 10.67 |
| NDUFB10 | 11 | 11.12 | 9.20 | 9.10 | 9.34 | 9.90 | 9.48 | 10.27 | 13.32 | 9.15 | 9.11 | 10.16 | 9.22 | 9.69 | 11.80 | 9.13 |
| NDUFB11 | 6 | 15.50 | 6.85 | 6.60 | 10.20 | 10.80 | 8.80 | 10.76 | 8.41 | 10.25 | 11.84 | 11.17 | 8.40 | 9.80 | 9.58 | 11.04 |
| NDUFB3 | 4 | 12.93 | 8.22 | 8.82 | 9.22 | 10.47 | 8.42 | 10.41 | 10.58 | 10.00 | 10.93 | 10.57 | 9.02 | 9.44 | 10.50 | 10.47 |
| NDUFB4 | 3 | 12.15 | 7.55 | 8.71 | 11.57 | 11.01 | 9.41 | 11.61 | 9.79 | 9.21 | 8.98 | 9.85 | 10.14 | 10.21 | 10.70 | 9.10 |
| NDUFB5 | 4 | 13.89 | 7.29 | 8.18 | 10.66 | 9.10 | 7.37 | 11.02 | 9.57 | 11.11 | 11.83 | 10.59 | 9.42 | 8.23 | 10.29 | 11.47 |
| NDUFB6 | 3 | 14.46 | 7.53 | 7.03 | 11.02 | 8.09 | 7.65 | 9.92 | 8.85 | 11.58 | 13.89 | 10.99 | 9.02 | 7.87 | 9.38 | 12.74 |
| NDUFB8 | 2 | 14.82 | 7.07 | 7.13 | 9.80 | 9.57 | 7.31 | 13.34 | 9.59 | 10.07 | 11.30 | 10.95 | 8.46 | 8.44 | 11.46 | 10.69 |

| | | | | | | | | | | | | | | | | |
|---|---|---|---|---|---|---|---|---|---|---|---|---|---|---|---|---|
| NDUFB9 | 6 | 10.13 | 8.42 | 10.56 | 9.95 | 10.48 | 9.67 | 12.09 | 10.48 | 9.37 | 8.86 | 9.27 | 10.25 | 10.07 | 11.28 | 9.12 |
| NDUFC1 | 1 | 14.59 | 7.57 | 10.79 | 6.23 | 12.25 | 10.43 | 11.27 | 10.25 | 8.92 | 7.70 | 11.08 | 8.51 | 11.34 | 10.76 | 8.31 |
| NDUFS1 | 31 | 10.70 | 9.14 | 8.29 | 10.66 | 10.26 | 9.69 | 11.79 | 12.02 | 8.54 | 8.92 | 9.92 | 9.47 | 9.97 | 11.90 | 8.73 |
| NDUFS2 | 11 | 10.31 | 9.11 | 7.67 | 9.30 | 11.31 | 10.79 | 11.37 | 12.39 | 8.65 | 9.11 | 9.71 | 8.48 | 11.05 | 11.88 | 8.88 |
| NDUFS3 | 10 | 10.05 | 8.22 | 10.07 | 9.46 | 11.42 | 10.01 | 12.67 | 10.64 | 8.67 | 8.79 | 9.14 | 9.77 | 10.71 | 11.66 | 8.73 |
| NDUFS4 | 4 | 9.27 | 8.09 | 9.73 | 11.29 | 11.32 | 10.66 | 13.51 | 12.91 | 6.65 | 6.57 | 8.68 | 10.51 | 10.99 | 13.21 | 6.61 |
| NDUFS5 | 3 | 9.54 | 8.97 | 10.64 | 9.25 | 11.54 | 11.25 | 12.57 | 10.67 | 7.83 | 7.74 | 9.25 | 9.95 | 11.40 | 11.62 | 7.79 |
| NDUFS6 | 5 | 9.88 | 8.38 | 8.55 | 7.78 | 11.27 | 10.68 | 11.48 | 15.50 | 8.00 | 8.47 | 9.13 | 8.16 | 10.98 | 13.49 | 8.24 |
| NDUFS7 | 5 | 10.31 | 8.42 | 8.66 | 9.88 | 10.91 | 10.62 | 12.95 | 11.41 | 8.35 | 8.49 | 9.37 | 9.27 | 10.77 | 12.18 | 8.42 |
| NDUFS8 | 2 | 10.09 | 7.82 | 12.00 | 11.61 | 9.91 | 8.99 | 12.41 | 8.92 | 8.94 | 9.32 | 8.96 | 11.80 | 9.45 | 10.66 | 9.13 |
| NDUFV1 | 12 | 10.53 | 8.87 | 8.69 | 10.37 | 10.84 | 10.30 | 12.35 | 11.36 | 8.25 | 8.43 | 9.70 | 9.53 | 10.57 | 11.86 | 8.34 |
| NDUFV2 | 8 | 10.11 | 8.93 | 8.97 | 11.16 | 10.79 | 10.33 | 11.53 | 11.22 | 8.62 | 8.34 | 9.52 | 10.07 | 10.56 | 11.38 | 8.48 |
| NDUFV3 | 18 | 10.27 | 9.15 | 8.77 | 9.16 | 11.08 | 10.71 | 12.10 | 11.00 | 8.88 | 8.88 | 9.71 | 8.97 | 10.90 | 11.55 | 8.88 |
| NECAP1 | 5 | 11.85 | 13.32 | 7.00 | 6.64 | 10.27 | 11.68 | 8.18 | 8.86 | 11.77 | 10.43 | 12.58 | 6.82 | 10.98 | 8.52 | 11.10 |
| NECAP2 | 7 | 10.82 | 11.48 | 10.18 | 9.83 | 8.72 | 8.84 | 7.67 | 7.22 | 13.32 | 11.92 | 11.15 | 10.01 | 8.78 | 7.45 | 12.62 |
| NEDD1 | 9 | 10.40 | 11.89 | 11.83 | 11.08 | 9.94 | 9.80 | 8.92 | 8.34 | 8.91 | 8.89 | 11.14 | 11.46 | 9.87 | 8.63 | 8.90 |
| NEDD4 | 2 | 9.79 | 10.30 | 13.42 | 11.97 | 9.54 | 9.79 | 5.44 | 5.77 | 11.63 | 12.33 | 10.05 | 12.70 | 9.67 | 5.60 | 11.98 |
| NEDD4L | 4 | 8.12 | 8.73 | 11.74 | 11.71 | 11.59 | 12.09 | 9.53 | 12.11 | 7.11 | 7.27 | 8.43 | 11.72 | 11.84 | 10.82 | 7.19 |
| NEDD8-MDP1 | 6 | 8.43 | 9.81 | 11.17 | 10.51 | 10.66 | 11.17 | 10.26 | 9.82 | 9.38 | 8.79 | 9.12 | 10.84 | 10.92 | 10.04 | 9.09 |
| NEFH | 18 | 8.95 | 10.55 | 11.96 | 12.21 | 7.26 | 7.62 | 10.05 | 9.20 | 11.24 | 10.96 | 9.75 | 12.08 | 7.44 | 9.63 | 11.10 |
| NEGR1 | 2 | 11.19 | 9.54 | 15.84 | 17.65 | 5.36 | 4.98 | 11.97 | 10.79 | 6.58 | 6.09 | 10.37 | 16.75 | 5.17 | 11.38 | 6.33 |
| NEIL2 | 2 | 8.24 | 14.08 | 6.89 | 7.20 | 11.88 | 13.09 | 7.95 | 10.66 | 10.38 | 9.63 | 11.16 | 7.05 | 12.49 | 9.30 | 10.00 |
| NEK4 | 1 | 7.41 | 11.40 | 11.25 | 10.52 | 10.45 | 7.32 | 10.98 | 11.01 | 10.89 | 8.78 | 9.40 | 10.88 | 8.89 | 10.99 | 9.84 |
| NEK6 | 7 | 10.00 | 10.43 | 12.21 | 10.82 | 8.74 | 8.30 | 9.87 | 9.19 | 10.12 | 10.32 | 10.21 | 11.52 | 8.52 | 9.53 | 10.22 |
| NEK7 | 8 | 7.23 | 7.32 | 10.39 | 7.64 | 14.48 | 13.10 | 5.79 | 10.79 | 11.86 | 11.40 | 7.27 | 9.01 | 13.79 | 8.29 | 11.63 |
| NEK9 | 7 | 10.26 | 10.44 | 8.86 | 9.02 | 9.98 | 10.89 | 8.89 | 8.94 | 11.29 | 11.43 | 10.35 | 8.94 | 10.43 | 8.92 | 11.36 |
| NELFA | 12 | 9.19 | 9.83 | 12.25 | 11.08 | 9.84 | 8.91 | 11.34 | 10.69 | 8.72 | 8.14 | 9.51 | 11.67 | 9.38 | 11.01 | 8.43 |
| NELFB | 10 | 9.25 | 9.30 | 10.12 | 10.45 | 9.17 | 9.28 | 10.35 | 11.22 | 10.81 | 10.05 | 9.27 | 10.29 | 9.22 | 10.78 | 10.43 |
| NELFCD | 6 | 9.31 | 9.62 | 11.26 | 10.63 | 9.24 | 9.12 | 11.68 | 11.08 | 8.86 | 9.21 | 9.46 | 10.94 | 9.18 | 11.38 | 9.03 |
| NELFE | 23 | 9.25 | 10.66 | 11.48 | 10.75 | 8.67 | 9.15 | 10.12 | 10.50 | 9.98 | 9.43 | 9.96 | 11.12 | 8.91 | 10.31 | 9.71 |
| NEMF | 18 | 9.14 | 10.20 | 10.55 | 10.84 | 10.32 | 10.86 | 9.57 | 9.26 | 9.52 | 9.74 | 9.67 | 10.70 | 10.59 | 9.41 | 9.63 |
| NENF | 8 | 10.05 | 10.67 | 7.55 | 7.53 | 14.14 | 15.23 | 10.05 | 5.98 | 9.46 | 9.34 | 10.36 | 7.54 | 14.69 | 8.01 | 9.40 |
| NES | 75 | 5.65 | 6.33 | 13.07 | 12.51 | 2.92 | 2.88 | 14.85 | 13.85 | 14.25 | 13.69 | 5.99 | 12.79 | 2.90 | 14.35 | 13.97 |
| NEU1 | 3 | 10.39 | 9.63 | 11.71 | 11.35 | 12.44 | 12.78 | 8.96 | 8.78 | 7.24 | 6.71 | 10.01 | 11.53 | 12.61 | 8.87 | 6.98 |
| NEURL4 | 2 | 9.92 | 11.80 | 10.49 | 10.46 | 9.63 | 8.08 | 11.74 | 10.29 | 8.94 | 8.66 | 10.86 | 10.47 | 8.86 | 11.01 | 8.80 |
| NEXN | 21 | 11.52 | 12.54 | 16.55 | 16.06 | 6.40 | 6.75 | 3.09 | 2.93 | 12.42 | 11.74 | 12.03 | 16.31 | 6.58 | 3.01 | 12.08 |
| NF1 | 11 | 9.72 | 9.73 | 12.75 | 14.25 | 8.28 | 8.54 | 8.72 | 8.50 | 9.72 | 9.80 | 9.73 | 13.50 | 8.41 | 8.61 | 9.76 |
| NF2 | 3 | 9.17 | 9.28 | 11.88 | 11.75 | 8.50 | 8.62 | 9.13 | 8.80 | 11.42 | 11.46 | 9.22 | 11.82 | 8.56 | 8.96 | 11.44 |
| NFAT5 | 1 | 8.23 | 10.05 | 8.50 | 6.42 | 8.37 | 9.82 | 12.49 | 11.21 | 13.33 | 11.58 | 9.14 | 7.46 | 9.09 | 11.85 | 12.46 |
| NFATC4 | 5 | 10.67 | 12.50 | 10.92 | 11.67 | 8.86 | 9.29 | 7.54 | 7.99 | 10.33 | 10.24 | 11.58 | 11.29 | 9.08 | 7.76 | 10.28 |
| NFIA | 5 | 12.34 | 12.51 | 8.37 | 7.56 | 11.62 | 11.08 | 9.96 | 8.40 | 9.93 | 8.23 | 12.43 | 7.97 | 11.35 | 9.18 | 9.08 |
| NFIB | 8 | 10.52 | 11.52 | 8.63 | 8.55 | 11.17 | 11.06 | 6.68 | 6.39 | 12.81 | 12.68 | 11.02 | 8.59 | 11.11 | 6.54 | 12.75 |

| | | | | | | | | | | | | | | | | |
|---|---|---|---|---|---|---|---|---|---|---|---|---|---|---|---|---|
| NFIC | 5 | 9.57 | 9.94 | 10.42 | 10.19 | 10.92 | 11.24 | 10.23 | 9.43 | 9.06 | 9.00 | 9.76 | 10.31 | 11.08 | 9.83 | 9.03 |
| NFIX | 4 | 9.34 | 9.03 | 14.19 | 14.08 | 4.51 | 4.61 | 11.05 | 10.47 | 11.46 | 11.25 | 9.19 | 14.13 | 4.56 | 10.76 | 11.36 |
| NFKB1 | 14 | 8.72 | 10.12 | 11.14 | 10.96 | 9.99 | 10.23 | 9.67 | 9.32 | 9.89 | 9.97 | 9.42 | 11.05 | 10.11 | 9.49 | 9.93 |
| NFKB2 | 14 | 8.56 | 8.43 | 12.45 | 11.50 | 10.71 | 10.80 | 6.67 | 6.05 | 13.15 | 11.68 | 8.49 | 11.97 | 10.75 | 6.36 | 12.42 |
| NFKBIB | 2 | 9.40 | 9.60 | 12.23 | 12.95 | 7.96 | 9.78 | 10.98 | 9.03 | 9.79 | 8.28 | 9.50 | 12.59 | 8.87 | 10.01 | 9.04 |
| NFKBIL1 | 1 | 10.02 | 10.92 | 9.10 | 8.31 | 10.98 | 13.57 | 10.38 | 9.68 | 9.33 | 7.71 | 10.47 | 8.71 | 12.28 | 10.03 | 8.52 |
| NFRKB | 4 | 9.36 | 10.18 | 9.95 | 10.20 | 9.66 | 9.48 | 9.77 | 9.61 | 10.60 | 11.19 | 9.77 | 10.07 | 9.57 | 9.69 | 10.89 |
| NFS1 | 10 | 10.80 | 9.61 | 10.37 | 11.03 | 9.80 | 9.43 | 9.79 | 8.80 | 10.01 | 10.37 | 10.20 | 10.70 | 9.61 | 9.29 | 10.19 |
| NFU1 | 6 | 10.33 | 11.37 | 12.17 | 12.55 | 9.03 | 8.76 | 9.67 | 6.19 | 9.75 | 10.18 | 10.85 | 12.36 | 8.89 | 7.93 | 9.97 |
| NFX1 | 6 | 11.21 | 11.17 | 10.74 | 10.08 | 10.53 | 10.89 | 8.49 | 7.58 | 10.10 | 9.22 | 11.19 | 10.41 | 10.71 | 8.03 | 9.66 |
| NFXL1 | 3 | 12.27 | 7.36 | 7.29 | 11.00 | 8.19 | 7.12 | 11.14 | 10.99 | 11.45 | 13.19 | 9.82 | 9.15 | 7.65 | 11.07 | 12.32 |
| NFYA | 2 | 8.61 | 10.27 | 10.70 | 8.77 | 10.11 | 8.75 | 12.26 | 10.76 | 10.12 | 9.65 | 9.44 | 9.74 | 9.43 | 11.51 | 9.88 |
| NFYB | 2 | 10.32 | 12.45 | 8.78 | 10.86 | 9.03 | 9.67 | 10.84 | 10.02 | 9.22 | 8.81 | 11.39 | 9.82 | 9.35 | 10.43 | 9.01 |
| NFYC | 5 | 10.30 | 11.59 | 9.51 | 9.95 | 9.28 | 10.00 | 11.15 | 11.20 | 8.74 | 8.28 | 10.95 | 9.73 | 9.64 | 11.17 | 8.51 |
| NGDN | 7 | 8.72 | 9.61 | 11.69 | 10.48 | 8.53 | 8.08 | 11.58 | 11.56 | 9.95 | 9.81 | 9.16 | 11.08 | 8.31 | 11.57 | 9.88 |
| NGEF | 3 | 8.19 | 8.93 | 9.20 | 8.82 | 11.16 | 10.50 | 7.51 | 7.85 | 13.99 | 13.85 | 8.56 | 9.01 | 10.83 | 7.68 | 13.92 |
| NGRN | 2 | 9.31 | 9.50 | 9.33 | 10.10 | 8.76 | 8.06 | 11.27 | 10.80 | 11.52 | 11.35 | 9.41 | 9.72 | 8.41 | 11.03 | 11.43 |
| NHEJ1 | 4 | 9.11 | 11.36 | 10.05 | 9.01 | 9.58 | 10.61 | 10.27 | 12.95 | 8.15 | 8.91 | 10.24 | 9.53 | 10.10 | 11.61 | 8.53 |
| NHLRC2 | 6 | 9.29 | 9.53 | 11.08 | 11.87 | 10.01 | 11.12 | 9.24 | 9.41 | 9.46 | 8.98 | 9.41 | 11.48 | 10.57 | 9.33 | 9.22 |
| NHP2 | 7 | 9.25 | 8.97 | 9.45 | 8.29 | 9.12 | 9.00 | 12.56 | 14.65 | 9.28 | 9.44 | 9.11 | 8.87 | 9.06 | 13.60 | 9.36 |
| NHP2L1 | 8 | 8.96 | 8.88 | 11.63 | 8.09 | 8.91 | 7.96 | 10.90 | 16.60 | 9.21 | 8.85 | 8.92 | 9.86 | 8.44 | 13.75 | 9.03 |
| NHS | 1 | 7.47 | 9.54 | 8.00 | 8.94 | 12.20 | 14.29 | 15.15 | 13.94 | 5.59 | 4.89 | 8.50 | 8.47 | 13.25 | 14.55 | 5.24 |
| NHSL1 | 1 | 6.94 | 8.39 | 13.78 | 13.75 | 9.03 | 8.52 | 12.53 | 11.04 | 8.84 | 7.18 | 7.66 | 13.76 | 8.78 | 11.79 | 8.01 |
| NID1 | 1 | 4.61 | 4.78 | 14.76 | 15.04 | 3.69 | 2.65 | 21.74 | 21.05 | 5.24 | 6.44 | 4.70 | 14.90 | 3.17 | 21.39 | 5.84 |
| NIF3L1 | 14 | 9.73 | 10.47 | 9.53 | 10.28 | 10.28 | 10.69 | 10.06 | 10.01 | 9.18 | 9.77 | 10.10 | 9.90 | 10.48 | 10.04 | 9.48 |
| NIFK | 10 | 10.02 | 9.71 | 10.37 | 10.46 | 8.59 | 8.11 | 11.38 | 9.90 | 9.80 | 11.67 | 9.86 | 10.41 | 8.35 | 10.64 | 10.74 |
| NIP7 | 8 | 9.42 | 9.66 | 9.99 | 8.46 | 9.79 | 9.46 | 11.44 | 11.96 | 9.82 | 9.99 | 9.54 | 9.23 | 9.62 | 11.70 | 9.91 |
| NIPBL | 24 | 9.15 | 9.15 | 9.44 | 8.79 | 10.98 | 10.66 | 12.77 | 11.59 | 9.00 | 8.48 | 9.15 | 9.11 | 10.82 | 12.18 | 8.74 |
| NIPSNAP1 | 5 | 9.89 | 10.06 | 10.77 | 11.16 | 8.29 | 7.52 | 11.28 | 10.83 | 9.82 | 10.38 | 9.98 | 10.96 | 7.91 | 11.05 | 10.10 |
| NIPSNAP3A | 4 | 8.94 | 7.70 | 10.15 | 9.60 | 14.71 | 13.77 | 9.07 | 8.97 | 8.52 | 8.58 | 8.32 | 9.87 | 14.24 | 9.02 | 8.55 |
| NIT1 | 3 | 10.06 | 10.57 | 9.12 | 8.90 | 12.02 | 12.05 | 8.23 | 7.61 | 10.94 | 10.50 | 10.32 | 9.01 | 12.04 | 7.92 | 10.72 |
| NIT2 | 14 | 9.26 | 9.51 | 9.00 | 8.28 | 11.92 | 11.42 | 9.82 | 16.04 | 7.54 | 7.21 | 9.38 | 8.64 | 11.67 | 12.93 | 7.38 |
| NKAP | 4 | 8.18 | 9.29 | 6.82 | 6.61 | 12.00 | 12.89 | 13.92 | 14.94 | 8.00 | 7.35 | 8.74 | 6.72 | 12.44 | 14.43 | 7.68 |
| NKIRAS1 | 1 | 10.04 | 11.06 | 8.22 | 8.37 | 10.04 | 9.74 | 10.41 | 11.11 | 10.63 | 10.37 | 10.55 | 8.30 | 9.89 | 10.76 | 10.50 |
| NKIRAS2 | 2 | 9.61 | 11.06 | 11.96 | 12.69 | 10.04 | 10.59 | 9.17 | 8.67 | 7.77 | 8.44 | 10.34 | 12.33 | 10.31 | 8.92 | 8.10 |
| NKRF | 32 | 7.60 | 8.71 | 10.46 | 9.24 | 9.95 | 10.00 | 14.31 | 13.47 | 8.32 | 7.95 | 8.15 | 9.85 | 9.97 | 13.89 | 8.13 |
| NKTR | 7 | 11.04 | 12.08 | 7.54 | 7.45 | 10.41 | 11.47 | 10.20 | 10.05 | 10.27 | 9.50 | 11.56 | 7.49 | 10.94 | 10.12 | 9.88 |
| NLE1 | 10 | 9.61 | 10.25 | 13.09 | 12.35 | 7.10 | 7.05 | 10.89 | 9.92 | 9.57 | 10.17 | 9.93 | 12.72 | 7.07 | 10.41 | 9.87 |
| NLN | 16 | 11.01 | 9.88 | 10.88 | 10.92 | 8.73 | 8.61 | 9.56 | 8.90 | 11.05 | 10.46 | 10.44 | 10.90 | 8.67 | 9.23 | 10.75 |
| NMD3 | 12 | 10.79 | 10.91 | 12.91 | 9.65 | 9.68 | 9.04 | 10.05 | 9.17 | 8.99 | 8.81 | 10.85 | 11.28 | 9.36 | 9.61 | 8.90 |
| NME1 | 76 | 9.08 | 11.45 | 12.91 | 11.05 | 9.27 | 9.00 | 8.05 | 8.73 | 10.16 | 10.30 | 10.26 | 11.98 | 9.13 | 8.39 | 10.23 |
| NME2 | 1 | 20.29 | 9.11 | 9.14 | 9.63 | 8.95 | 10.80 | 7.63 | 6.09 | 11.44 | 6.93 | 14.70 | 9.38 | 9.87 | 6.86 | 9.19 |

| | | | | | | | | | | | | | | | | |
|---|---|---|---|---|---|---|---|---|---|---|---|---|---|---|---|---|
| NME2P1 | 24 | 10.76 | 12.84 | 12.12 | 10.24 | 10.34 | 10.03 | 7.11 | 5.98 | 10.66 | 9.92 | 11.80 | 11.18 | 10.19 | 6.54 | 10.29 |
| NME3 | 3 | 12.92 | 8.86 | 8.57 | 10.55 | 10.05 | 9.60 | 8.74 | 7.99 | 10.69 | 12.04 | 10.89 | 9.56 | 9.82 | 8.36 | 11.36 |
| NME6 | 2 | 9.88 | 10.25 | 10.62 | 10.85 | 8.40 | 7.97 | 10.27 | 10.21 | 10.57 | 10.98 | 10.06 | 10.73 | 8.19 | 10.24 | 10.77 |
| NME7 | 4 | 8.74 | 10.23 | 14.10 | 13.28 | 10.35 | 9.25 | 8.40 | 7.43 | 9.35 | 8.88 | 9.49 | 13.69 | 9.80 | 7.91 | 9.11 |
| NMI | 7 | 12.46 | 13.39 | 13.68 | 13.33 | 13.96 | 14.33 | 4.04 | 4.21 | 5.53 | 5.06 | 12.93 | 13.51 | 14.15 | 4.13 | 5.29 |
| NMNAT1 | 4 | 13.45 | 15.68 | 5.88 | 6.12 | 6.30 | 6.84 | 11.04 | 11.14 | 12.16 | 11.40 | 14.56 | 6.00 | 6.57 | 11.09 | 11.78 |
| NMRAL1 | 6 | 9.86 | 10.95 | 10.49 | 10.69 | 10.51 | 11.15 | 9.48 | 8.60 | 9.02 | 9.25 | 10.40 | 10.59 | 10.83 | 9.04 | 9.13 |
| NMT1 | 19 | 9.40 | 10.64 | 12.57 | 11.98 | 9.12 | 9.20 | 8.50 | 8.07 | 10.44 | 10.07 | 10.02 | 12.28 | 9.16 | 8.28 | 10.26 |
| NNMT | 14 | 7.78 | 8.48 | 8.80 | 5.76 | 11.82 | 10.08 | 8.92 | 23.88 | 7.38 | 7.10 | 8.13 | 7.28 | 10.95 | 16.40 | 7.24 |
| NNT | 62 | 12.91 | 6.18 | 6.42 | 9.79 | 12.17 | 10.04 | 9.73 | 9.63 | 10.67 | 12.46 | 9.54 | 8.11 | 11.11 | 9.68 | 11.57 |
| NO66 | 6 | 9.67 | 11.15 | 8.81 | 8.83 | 11.01 | 11.12 | 10.73 | 9.92 | 9.46 | 9.31 | 10.41 | 8.82 | 11.07 | 10.32 | 9.38 |
| NOA1 | 9 | 10.82 | 9.92 | 9.20 | 10.12 | 9.28 | 9.88 | 9.33 | 8.58 | 11.35 | 11.52 | 10.37 | 9.66 | 9.58 | 8.96 | 11.44 |
| NOB1 | 11 | 9.88 | 10.68 | 8.81 | 8.54 | 8.88 | 8.99 | 11.79 | 10.75 | 10.93 | 10.74 | 10.28 | 8.67 | 8.94 | 11.27 | 10.83 |
| NOC2L | 17 | 10.21 | 9.76 | 8.99 | 9.37 | 8.96 | 9.13 | 11.56 | 11.26 | 10.47 | 10.31 | 9.99 | 9.18 | 9.04 | 11.41 | 10.39 |
| NOC3L | 14 | 10.35 | 9.94 | 10.13 | 9.36 | 8.71 | 8.15 | 11.49 | 9.70 | 10.53 | 11.65 | 10.14 | 9.75 | 8.43 | 10.60 | 11.09 |
| NOC4L | 10 | 9.12 | 10.11 | 9.73 | 9.67 | 8.66 | 9.39 | 12.02 | 12.00 | 9.88 | 9.43 | 9.62 | 9.70 | 9.02 | 12.01 | 9.65 |
| NOL10 | 12 | 9.15 | 10.23 | 10.44 | 9.94 | 8.68 | 9.01 | 11.09 | 11.25 | 10.03 | 10.17 | 9.69 | 10.19 | 8.84 | 11.17 | 10.10 |
| NOL11 | 18 | 8.80 | 9.27 | 10.99 | 10.53 | 7.85 | 7.80 | 11.08 | 13.03 | 10.34 | 10.30 | 9.04 | 10.76 | 7.83 | 12.05 | 10.32 |
| NOL12 | 2 | 9.40 | 10.66 | 12.19 | 10.54 | 9.00 | 8.85 | 10.83 | 9.56 | 9.57 | 9.39 | 10.03 | 11.36 | 8.93 | 10.20 | 9.48 |
| NOL3 | 3 | 9.82 | 10.78 | 8.76 | 7.78 | 12.87 | 12.95 | 9.57 | 4.68 | 11.57 | 11.22 | 10.30 | 8.27 | 12.91 | 7.13 | 11.39 |
| NOL6 | 24 | 9.34 | 9.18 | 10.13 | 10.14 | 8.43 | 8.66 | 10.19 | 9.81 | 12.05 | 12.06 | 9.26 | 10.14 | 8.55 | 10.00 | 12.06 |
| NOL7 | 4 | 10.17 | 10.37 | 11.01 | 10.49 | 8.58 | 8.51 | 11.86 | 10.64 | 9.25 | 9.13 | 10.27 | 10.75 | 8.54 | 11.25 | 9.19 |
| NOL8 | 15 | 9.94 | 10.54 | 10.05 | 9.80 | 10.65 | 11.10 | 11.16 | 10.17 | 8.63 | 7.96 | 10.24 | 9.93 | 10.88 | 10.66 | 8.29 |
| NOL9 | 9 | 9.13 | 10.31 | 9.49 | 9.28 | 9.08 | 9.46 | 11.79 | 11.26 | 10.01 | 10.19 | 9.72 | 9.39 | 9.27 | 11.52 | 10.10 |
| NOLC1 | 48 | 9.62 | 10.29 | 10.73 | 10.24 | 8.78 | 9.00 | 10.53 | 9.45 | 10.89 | 10.50 | 9.95 | 10.48 | 8.89 | 9.99 | 10.69 |
| NOMO1 | 1 | 10.62 | 7.11 | 8.09 | 12.57 | 8.13 | 10.24 | 9.53 | 11.85 | 9.35 | 12.50 | 8.87 | 10.33 | 9.18 | 10.69 | 10.93 |
| NOMO2 | 28 | 10.31 | 6.69 | 8.55 | 10.87 | 10.40 | 9.07 | 11.57 | 11.47 | 10.00 | 11.08 | 8.50 | 9.71 | 9.73 | 11.52 | 10.54 |
| NONO | 90 | 9.82 | 10.74 | 9.11 | 8.94 | 10.42 | 10.59 | 11.35 | 12.51 | 8.46 | 8.07 | 10.28 | 9.03 | 10.50 | 11.93 | 8.26 |
| NOP10 | 2 | 9.22 | 10.66 | 10.72 | 7.85 | 9.22 | 9.42 | 12.06 | 10.99 | 10.38 | 9.48 | 9.94 | 9.29 | 9.32 | 11.52 | 9.93 |
| NOP14 | 17 | 9.18 | 9.93 | 10.40 | 10.31 | 9.40 | 9.01 | 11.09 | 10.29 | 9.91 | 10.49 | 9.55 | 10.35 | 9.21 | 10.69 | 10.20 |
| NOP16 | 9 | 9.85 | 9.86 | 9.51 | 10.28 | 8.31 | 8.35 | 13.08 | 11.68 | 9.12 | 9.97 | 9.86 | 9.89 | 8.33 | 12.38 | 9.55 |
| NOP2 | 33 | 10.20 | 9.75 | 9.61 | 9.79 | 9.23 | 8.79 | 10.42 | 9.65 | 10.52 | 12.05 | 9.97 | 9.70 | 9.01 | 10.04 | 11.28 |
| NOP56 | 25 | 7.88 | 9.29 | 10.41 | 10.41 | 8.53 | 8.91 | 11.13 | 14.66 | 8.89 | 9.89 | 8.58 | 10.41 | 8.72 | 12.89 | 9.39 |
| NOP58 | 24 | 9.45 | 8.78 | 10.26 | 8.04 | 9.19 | 8.52 | 11.69 | 14.41 | 9.94 | 9.72 | 9.11 | 9.15 | 8.86 | 13.05 | 9.83 |
| NOP9 | 8 | 8.55 | 9.17 | 10.38 | 9.90 | 9.91 | 9.92 | 10.96 | 10.94 | 10.25 | 10.03 | 8.86 | 10.14 | 9.91 | 10.95 | 10.14 |
| NOSIP | 13 | 8.87 | 8.87 | 11.03 | 9.77 | 7.83 | 7.70 | 14.07 | 12.25 | 10.16 | 9.44 | 8.87 | 10.40 | 7.77 | 13.16 | 9.80 |
| NOTCH1 | 2 | 10.10 | 11.15 | 9.37 | 13.70 | 8.93 | 9.74 | 8.74 | 9.69 | 9.65 | 8.93 | 10.62 | 11.54 | 9.33 | 9.21 | 9.29 |
| NOTCH2 | 5 | 11.81 | 10.63 | 10.74 | 11.64 | 8.19 | 7.80 | 6.62 | 5.73 | 13.62 | 13.20 | 11.22 | 11.19 | 8.00 | 6.18 | 13.41 |
| NOV | 1 | 10.07 | 5.83 | 6.65 | 8.94 | 8.31 | 8.51 | 18.53 | 19.36 | 6.20 | 7.59 | 7.95 | 7.80 | 8.41 | 18.95 | 6.90 |
| NOVA2 | 3 | 9.84 | 10.33 | 10.66 | 10.54 | 9.46 | 9.15 | 11.81 | 10.49 | 9.19 | 8.53 | 10.09 | 10.60 | 9.31 | 11.15 | 8.86 |
| NPC2 | 5 | 8.00 | 8.67 | 12.11 | 10.80 | 12.50 | 13.57 | 9.17 | 8.38 | 8.52 | 8.28 | 8.34 | 11.45 | 13.04 | 8.77 | 8.40 |
| NPEPL1 | 5 | 9.24 | 9.38 | 7.14 | 6.29 | 13.57 | 13.78 | 10.83 | 10.88 | 9.79 | 9.10 | 9.31 | 6.72 | 13.67 | 10.85 | 9.45 |

| | | | | | | | | | | | | | | | | |
|---|---|---|---|---|---|---|---|---|---|---|---|---|---|---|---|---|
| NPEPPS | 66 | 11.76 | 11.82 | 9.41 | 9.02 | 11.08 | 11.13 | 7.28 | 7.12 | 11.00 | 10.40 | 11.79 | 9.21 | 11.10 | 7.20 | 10.70 |
| NPLOC4 | 20 | 11.22 | 11.17 | 11.62 | 11.30 | 9.33 | 9.50 | 7.91 | 8.33 | 10.15 | 9.47 | 11.19 | 11.46 | 9.42 | 8.12 | 9.81 |
| NPM1 | 87 | 9.16 | 11.36 | 8.95 | 8.82 | 8.37 | 8.63 | 15.29 | 13.97 | 7.95 | 7.49 | 10.26 | 8.88 | 8.50 | 14.63 | 7.72 |
| NPM3 | 2 | 8.56 | 9.39 | 11.16 | 10.92 | 7.80 | 8.34 | 13.04 | 12.10 | 9.59 | 9.11 | 8.97 | 11.04 | 8.07 | 12.57 | 9.35 |
| NPR1 | 2 | 11.38 | 8.48 | 2.70 | 3.02 | 22.94 | 23.21 | 2.16 | 2.03 | 12.04 | 12.05 | 9.93 | 2.86 | 23.07 | 2.09 | 12.05 |
| NPR3 | 9 | 11.85 | 8.85 | 6.88 | 8.28 | 10.20 | 8.85 | 12.57 | 12.02 | 9.99 | 10.51 | 10.35 | 7.58 | 9.53 | 12.29 | 10.25 |
| NPRL3 | 1 | 10.63 | 11.01 | 7.92 | 9.98 | 10.57 | 11.38 | 9.09 | 7.32 | 10.95 | 11.15 | 10.82 | 8.95 | 10.97 | 8.20 | 11.05 |
| NPTN | 5 | 10.69 | 7.37 | 8.29 | 12.11 | 9.69 | 9.76 | 7.45 | 9.19 | 11.95 | 13.51 | 9.03 | 10.20 | 9.72 | 8.32 | 12.73 |
| NPTX1 | 2 | 19.96 | 16.45 | 6.45 | 7.36 | 8.86 | 8.58 | 2.58 | 2.78 | 13.79 | 13.19 | 18.20 | 6.91 | 8.72 | 2.68 | 13.49 |
| NQO1 | 38 | 13.53 | 16.09 | 5.86 | 5.12 | 11.96 | 11.99 | 4.98 | 4.69 | 13.07 | 12.72 | 14.81 | 5.49 | 11.97 | 4.83 | 12.89 |
| NQO2 | 13 | 10.67 | 11.98 | 6.31 | 5.96 | 13.99 | 14.20 | 11.35 | 10.42 | 7.65 | 7.48 | 11.32 | 6.13 | 14.10 | 10.88 | 7.56 |
| NR1H2 | 1 | 9.87 | 11.22 | 8.51 | 8.74 | 10.21 | 10.33 | 9.89 | 13.22 | 8.91 | 9.10 | 10.55 | 8.62 | 10.27 | 11.56 | 9.00 |
| NR2C2 | 3 | 9.18 | 10.09 | 9.94 | 10.18 | 11.63 | 11.29 | 9.61 | 9.41 | 9.72 | 8.96 | 9.63 | 10.06 | 11.46 | 9.51 | 9.34 |
| NR2C2AP | 2 | 7.92 | 9.13 | 9.76 | 9.16 | 11.82 | 12.91 | 10.82 | 9.38 | 9.09 | 10.02 | 8.53 | 9.46 | 12.37 | 10.10 | 9.55 |
| NR2F2 | 1 | 6.90 | 8.29 | 9.68 | 7.77 | 8.12 | 7.74 | 19.48 | 15.61 | 7.31 | 9.09 | 7.59 | 8.72 | 7.93 | 17.55 | 8.20 |
| NR3C1 | 20 | 10.52 | 11.15 | 9.80 | 9.41 | 8.96 | 8.64 | 12.64 | 11.36 | 8.87 | 8.65 | 10.84 | 9.60 | 8.80 | 12.00 | 8.76 |
| NR4A1 | 2 | 9.39 | 9.77 | 7.05 | 8.67 | 12.41 | 12.44 | 14.02 | 14.20 | 6.48 | 5.58 | 9.58 | 7.86 | 12.43 | 14.11 | 6.03 |
| NRAS | 12 | 10.52 | 9.11 | 10.07 | 11.15 | 8.63 | 8.24 | 10.48 | 9.22 | 11.24 | 11.35 | 9.82 | 10.61 | 8.43 | 9.85 | 11.30 |
| NRBF2 | 2 | 9.70 | 10.70 | 11.97 | 11.30 | 10.46 | 10.17 | 8.25 | 8.12 | 9.97 | 9.37 | 10.20 | 11.63 | 10.32 | 8.18 | 9.67 |
| NRBP1 | 7 | 9.95 | 10.87 | 11.84 | 12.08 | 10.06 | 10.11 | 9.14 | 8.70 | 8.99 | 8.27 | 10.41 | 11.96 | 10.08 | 8.92 | 8.63 |
| NRBP2 | 3 | 8.46 | 10.54 | 11.09 | 11.28 | 8.98 | 9.47 | 8.05 | 8.17 | 12.78 | 11.18 | 9.50 | 11.18 | 9.22 | 8.11 | 11.98 |
| NRD1 | 37 | 9.89 | 10.20 | 9.67 | 8.50 | 9.10 | 8.84 | 11.29 | 13.56 | 9.53 | 9.42 | 10.04 | 9.09 | 8.97 | 12.42 | 9.47 |
| NRDE2 | 3 | 9.27 | 9.10 | 9.89 | 11.50 | 9.31 | 10.12 | 10.84 | 10.06 | 9.85 | 10.05 | 9.18 | 10.70 | 9.72 | 10.45 | 9.95 |
| NRF1 | 1 | 9.43 | 10.54 | 10.44 | 10.43 | 8.93 | 8.42 | 11.96 | 12.41 | 8.63 | 8.81 | 9.98 | 10.44 | 8.67 | 12.18 | 8.72 |
| NRP1 | 1 | 12.19 | 11.15 | 8.95 | 9.23 | 5.99 | 7.62 | 12.88 | 11.45 | 11.21 | 9.32 | 11.67 | 9.09 | 6.80 | 12.17 | 10.27 |
| NSA2 | 6 | 8.20 | 7.89 | 8.60 | 7.66 | 6.88 | 6.70 | 11.03 | 26.11 | 8.55 | 8.37 | 8.05 | 8.13 | 6.79 | 18.57 | 8.46 |
| NSD1 | 3 | 10.34 | 10.41 | 8.97 | 8.60 | 10.25 | 9.67 | 12.78 | 11.64 | 8.68 | 8.66 | 10.38 | 8.79 | 9.96 | 12.21 | 8.67 |
| NSDHL | 11 | 11.02 | 7.04 | 9.13 | 12.26 | 10.87 | 9.63 | 11.53 | 10.83 | 7.99 | 9.69 | 9.03 | 10.70 | 10.25 | 11.18 | 8.84 |
| NSF | 36 | 11.32 | 12.58 | 10.72 | 10.17 | 10.00 | 10.20 | 8.09 | 7.39 | 9.85 | 9.69 | 11.95 | 10.45 | 10.10 | 7.74 | 9.77 |
| NSFL1C | 24 | 9.70 | 10.85 | 10.60 | 10.25 | 10.99 | 11.63 | 9.49 | 8.68 | 9.02 | 8.77 | 10.28 | 10.43 | 11.31 | 9.09 | 8.90 |
| NSL1 | 4 | 9.84 | 11.92 | 8.76 | 9.56 | 10.87 | 11.64 | 8.76 | 8.70 | 10.07 | 9.89 | 10.88 | 9.16 | 11.25 | 8.73 | 9.98 |
| NSMAF | 2 | 10.08 | 11.44 | 9.81 | 9.77 | 8.20 | 10.12 | 11.44 | 11.88 | 8.59 | 8.67 | 10.76 | 9.79 | 9.16 | 11.66 | 8.63 |
| NSMCE1 | 1 | 8.16 | 9.76 | 10.53 | 8.26 | 10.75 | 13.78 | 11.75 | 9.09 | 9.76 | 8.16 | 8.96 | 9.39 | 12.27 | 10.42 | 8.96 |
| NSMCE2 | 2 | 10.50 | 11.58 | 8.15 | 8.73 | 9.02 | 10.55 | 11.83 | 10.82 | 9.85 | 8.98 | 11.04 | 8.44 | 9.78 | 11.32 | 9.41 |
| NSMCE4A | 1 | 7.87 | 8.53 | 11.27 | 9.83 | 10.35 | 9.84 | 10.15 | 10.88 | 10.08 | 11.20 | 8.20 | 10.55 | 10.09 | 10.51 | 10.64 |
| NSMF | 1 | 8.52 | 9.79 | 5.70 | 6.31 | 11.17 | 11.99 | 14.34 | 14.01 | 8.77 | 9.39 | 9.16 | 6.01 | 11.58 | 14.17 | 9.08 |
| NSRP1 | 13 | 9.70 | 10.98 | 12.80 | 11.55 | 9.55 | 9.16 | 8.31 | 7.50 | 10.43 | 10.01 | 10.34 | 12.17 | 9.36 | 7.91 | 10.22 |
| NSUN2 | 76 | 8.90 | 9.48 | 8.59 | 7.69 | 9.27 | 9.43 | 10.10 | 16.88 | 10.02 | 9.64 | 9.19 | 8.14 | 9.35 | 13.49 | 9.83 |
| NSUN4 | 1 | 12.04 | 10.76 | 8.97 | 11.60 | 9.11 | 7.80 | 11.13 | 10.52 | 9.32 | 8.75 | 11.40 | 10.28 | 8.45 | 10.82 | 9.04 |
| NSUN5 | 6 | 8.63 | 10.20 | 11.17 | 11.15 | 7.41 | 7.45 | 12.26 | 11.87 | 9.95 | 9.90 | 9.42 | 11.16 | 7.43 | 12.07 | 9.92 |
| NSUN6 | 2 | 10.13 | 11.06 | 13.29 | 12.09 | 9.71 | 9.97 | 11.53 | 9.43 | 6.83 | 5.96 | 10.60 | 12.69 | 9.84 | 10.48 | 6.40 |
| NT5C | 4 | 10.05 | 10.71 | 9.73 | 9.93 | 10.24 | 10.58 | 6.73 | 6.92 | 12.84 | 12.28 | 10.38 | 9.83 | 10.41 | 6.82 | 12.56 |

| | | | | | | | | | | | | | | | | |
|---|---|---|---|---|---|---|---|---|---|---|---|---|---|---|---|---|
| NT5C2 | 8 | 9.97 | 10.57 | 11.19 | 10.72 | 12.66 | 12.34 | 6.87 | 6.63 | 9.67 | 9.40 | 10.27 | 10.95 | 12.50 | 6.75 | 9.53 |
| NT5C3A | 12 | 11.59 | 11.23 | 8.95 | 8.87 | 10.10 | 10.33 | 10.30 | 12.82 | 7.95 | 7.85 | 11.41 | 8.91 | 10.22 | 11.56 | 7.90 |
| NT5DC1 | 9 | 9.08 | 9.53 | 11.56 | 9.52 | 13.63 | 13.40 | 9.24 | 10.54 | 7.00 | 6.49 | 9.31 | 10.54 | 13.52 | 9.89 | 6.74 |
| NT5DC2 | 4 | 11.04 | 10.02 | 10.32 | 8.87 | 9.75 | 8.74 | 9.30 | 7.15 | 12.09 | 12.72 | 10.53 | 9.60 | 9.24 | 8.22 | 12.41 |
| NT5DC3 | 3 | 8.36 | 7.28 | 15.83 | 12.43 | 11.49 | 6.86 | 10.75 | 6.72 | 8.38 | 11.89 | 7.82 | 14.13 | 9.18 | 8.73 | 10.13 |
| NT5E | 5 | 5.95 | 5.07 | 12.57 | 12.84 | 5.10 | 4.68 | 22.31 | 18.45 | 6.26 | 6.76 | 5.51 | 12.71 | 4.89 | 20.38 | 6.51 |
| NTHL1 | 5 | 9.29 | 9.32 | 8.86 | 8.80 | 9.00 | 9.81 | 13.25 | 12.24 | 10.07 | 9.35 | 9.30 | 8.83 | 9.41 | 12.75 | 9.71 |
| NTMT1 | 8 | 9.22 | 9.62 | 14.15 | 12.01 | 8.49 | 8.00 | 10.41 | 9.37 | 9.74 | 8.99 | 9.42 | 13.08 | 8.24 | 9.89 | 9.37 |
| NTPCR | 6 | 9.25 | 10.02 | 10.53 | 9.92 | 11.59 | 11.38 | 9.75 | 8.68 | 9.47 | 9.40 | 9.63 | 10.23 | 11.49 | 9.22 | 9.44 |
| NUB1 | 5 | 9.99 | 11.27 | 10.87 | 11.41 | 7.91 | 8.57 | 10.14 | 10.87 | 9.47 | 9.50 | 10.63 | 11.14 | 8.24 | 10.50 | 9.48 |
| NUBP1 | 5 | 10.11 | 11.26 | 9.58 | 9.51 | 12.51 | 12.51 | 9.09 | 8.66 | 8.61 | 8.17 | 10.68 | 9.54 | 12.51 | 8.88 | 8.39 |
| NUBP2 | 7 | 8.69 | 10.36 | 9.95 | 9.63 | 11.60 | 11.33 | 8.64 | 10.17 | 9.36 | 10.27 | 9.53 | 9.79 | 11.47 | 9.40 | 9.82 |
| NUBPL | 2 | 11.02 | 9.17 | 9.77 | 10.49 | 10.08 | 9.24 | 11.18 | 10.07 | 9.41 | 9.57 | 10.10 | 10.13 | 9.66 | 10.62 | 9.49 |
| NUCB1 | 29 | 10.34 | 8.86 | 7.94 | 8.87 | 12.54 | 12.53 | 12.35 | 7.00 | 9.64 | 9.91 | 9.60 | 8.40 | 12.54 | 9.68 | 9.78 |
| NUCB2 | 15 | 9.81 | 9.06 | 8.49 | 8.86 | 13.99 | 13.46 | 10.51 | 3.40 | 11.05 | 11.36 | 9.44 | 8.67 | 13.73 | 6.96 | 11.20 |
| NUCKS1 | 11 | 9.36 | 10.08 | 8.06 | 7.50 | 12.61 | 13.08 | 14.54 | 6.05 | 9.55 | 9.17 | 9.72 | 7.78 | 12.85 | 10.29 | 9.36 |
| NUDC | 69 | 11.09 | 12.16 | 9.58 | 8.91 | 8.34 | 8.74 | 10.54 | 9.99 | 10.39 | 10.26 | 11.62 | 9.25 | 8.54 | 10.26 | 10.33 |
| NUDCD1 | 13 | 9.30 | 9.24 | 9.69 | 8.83 | 7.94 | 7.94 | 13.02 | 15.42 | 9.44 | 9.18 | 9.27 | 9.26 | 7.94 | 14.22 | 9.31 |
| NUDCD2 | 6 | 8.88 | 9.80 | 10.38 | 8.93 | 7.60 | 7.90 | 11.29 | 18.41 | 8.78 | 8.03 | 9.34 | 9.66 | 7.75 | 14.85 | 8.40 |
| NUDCD3 | 13 | 9.06 | 9.99 | 11.92 | 11.11 | 8.44 | 8.71 | 10.45 | 10.67 | 9.97 | 9.67 | 9.53 | 11.52 | 8.58 | 10.56 | 9.82 |
| NUDT1 | 6 | 9.14 | 9.89 | 12.50 | 10.42 | 8.77 | 8.76 | 10.19 | 9.09 | 10.97 | 10.27 | 9.51 | 11.46 | 8.77 | 9.64 | 10.62 |
| NUDT12 | 4 | 10.36 | 11.08 | 8.98 | 8.25 | 9.45 | 10.11 | 11.26 | 10.53 | 10.23 | 9.75 | 10.72 | 8.61 | 9.78 | 10.89 | 9.99 |
| NUDT15 | 2 | 10.71 | 10.40 | 14.92 | 8.76 | 11.55 | 10.01 | 11.69 | 8.05 | 7.08 | 6.84 | 10.56 | 11.84 | 10.78 | 9.87 | 6.96 |
| NUDT16 | 4 | 10.07 | 10.40 | 11.66 | 12.39 | 9.38 | 10.00 | 7.71 | 7.94 | 10.05 | 10.40 | 10.23 | 12.03 | 9.69 | 7.83 | 10.22 |
| NUDT16L1 | 5 | 10.59 | 10.75 | 11.43 | 10.90 | 11.62 | 11.94 | 6.79 | 6.71 | 9.64 | 9.63 | 10.67 | 11.17 | 11.78 | 6.75 | 9.63 |
| NUDT19 | 4 | 9.31 | 8.09 | 8.99 | 8.29 | 9.07 | 8.37 | 15.39 | 14.39 | 8.96 | 9.14 | 8.70 | 8.64 | 8.72 | 14.89 | 9.05 |
| NUDT2 | 4 | 9.36 | 10.42 | 10.92 | 9.89 | 11.92 | 12.69 | 9.43 | 9.17 | 7.94 | 8.26 | 9.89 | 10.40 | 12.31 | 9.30 | 8.10 |
| NUDT21 | 22 | 9.93 | 10.92 | 9.18 | 8.62 | 10.55 | 11.05 | 10.62 | 9.89 | 9.73 | 9.50 | 10.43 | 8.90 | 10.80 | 10.26 | 9.61 |
| NUDT3 | 4 | 9.23 | 9.83 | 9.60 | 10.05 | 9.45 | 10.10 | 11.65 | 11.15 | 9.62 | 9.33 | 9.53 | 9.82 | 9.78 | 11.40 | 9.47 |
| NUDT4 | 4 | 9.07 | 9.17 | 10.46 | 10.15 | 9.22 | 8.89 | 14.43 | 11.99 | 8.78 | 7.85 | 9.12 | 10.30 | 9.05 | 13.21 | 8.32 |
| NUDT5 | 18 | 8.88 | 9.58 | 9.89 | 7.80 | 10.82 | 10.28 | 9.24 | 14.63 | 9.59 | 9.30 | 9.23 | 8.84 | 10.55 | 11.93 | 9.45 |
| NUDT9 | 10 | 9.54 | 9.04 | 12.12 | 9.64 | 13.34 | 12.24 | 10.82 | 8.68 | 7.57 | 7.01 | 9.29 | 10.88 | 12.79 | 9.75 | 7.29 |
| NUF2 | 14 | 9.85 | 11.22 | 10.36 | 10.61 | 11.29 | 12.18 | 8.82 | 8.25 | 9.02 | 8.40 | 10.54 | 10.48 | 11.74 | 8.53 | 8.71 |
| NUFIP1 | 2 | 10.68 | 10.20 | 10.58 | 10.39 | 9.16 | 10.16 | 9.83 | 9.98 | 9.40 | 9.61 | 10.44 | 10.48 | 9.66 | 9.91 | 9.51 |
| NUFIP2 | 17 | 8.49 | 9.47 | 11.50 | 10.30 | 10.60 | 10.48 | 9.13 | 10.06 | 10.28 | 9.69 | 8.98 | 10.90 | 10.54 | 9.60 | 9.98 |
| NUMA1 | 125 | 8.78 | 9.63 | 11.26 | 10.36 | 10.67 | 10.32 | 9.92 | 11.28 | 9.25 | 8.54 | 9.20 | 10.81 | 10.49 | 10.60 | 8.90 |
| NUMB | 9 | 9.14 | 9.58 | 11.60 | 10.36 | 10.53 | 10.84 | 9.57 | 8.22 | 10.18 | 9.98 | 9.36 | 10.98 | 10.68 | 8.89 | 10.08 |
| NUMBL | 5 | 9.51 | 11.00 | 12.02 | 11.78 | 10.91 | 10.02 | 8.13 | 7.77 | 9.43 | 9.42 | 10.26 | 11.90 | 10.47 | 7.95 | 9.43 |
| NUP107 | 19 | 9.83 | 10.07 | 9.09 | 8.97 | 9.61 | 9.74 | 10.59 | 11.36 | 10.71 | 10.04 | 9.95 | 9.03 | 9.68 | 10.97 | 10.37 |
| NUP133 | 33 | 9.53 | 10.19 | 9.37 | 8.56 | 10.28 | 10.03 | 10.71 | 12.62 | 9.67 | 9.04 | 9.86 | 8.96 | 10.15 | 11.67 | 9.36 |
| NUP153 | 39 | 8.69 | 9.65 | 11.09 | 10.35 | 9.22 | 9.20 | 10.61 | 12.27 | 9.67 | 9.23 | 9.17 | 10.72 | 9.21 | 11.44 | 9.45 |
| NUP155 | 37 | 10.00 | 9.90 | 9.50 | 8.10 | 10.23 | 9.60 | 11.12 | 10.92 | 10.36 | 10.28 | 9.95 | 8.80 | 9.91 | 11.02 | 10.32 |

| | | | | | | | | | | | | | | | | |
|---|---|---|---|---|---|---|---|---|---|---|---|---|---|---|---|---|
| NUP160 | 17 | 10.19 | 9.91 | 9.47 | 9.68 | 9.34 | 9.14 | 9.63 | 12.54 | 9.94 | 10.16 | 10.05 | 9.58 | 9.24 | 11.08 | 10.05 |
| NUP188 | 18 | 9.96 | 9.65 | 8.41 | 8.51 | 10.84 | 10.46 | 9.66 | 9.75 | 10.96 | 11.81 | 9.81 | 8.46 | 10.65 | 9.70 | 11.38 |
| NUP205 | 30 | 10.91 | 8.75 | 8.33 | 9.36 | 8.78 | 8.44 | 10.37 | 12.48 | 11.02 | 11.56 | 9.83 | 8.85 | 8.61 | 11.42 | 11.29 |
| NUP210 | 14 | 11.89 | 6.35 | 7.48 | 11.71 | 10.64 | 9.44 | 10.81 | 10.62 | 9.12 | 11.95 | 9.12 | 9.59 | 10.04 | 10.71 | 10.53 |
| NUP214 | 56 | 10.33 | 10.75 | 10.27 | 9.51 | 9.91 | 10.00 | 9.53 | 8.93 | 10.71 | 10.05 | 10.54 | 9.89 | 9.96 | 9.23 | 10.38 |
| NUP35 | 12 | 10.88 | 11.50 | 11.14 | 10.73 | 9.25 | 9.33 | 11.52 | 10.83 | 7.64 | 7.18 | 11.19 | 10.94 | 9.29 | 11.17 | 7.41 |
| NUP37 | 6 | 9.76 | 10.88 | 9.90 | 10.03 | 9.35 | 10.17 | 10.74 | 10.05 | 9.43 | 9.69 | 10.32 | 9.96 | 9.76 | 10.40 | 9.56 |
| NUP43 | 6 | 8.72 | 10.14 | 11.71 | 10.90 | 9.88 | 9.53 | 10.13 | 8.81 | 9.81 | 10.39 | 9.43 | 11.31 | 9.70 | 9.47 | 10.10 |
| NUP50 | 18 | 8.58 | 10.03 | 12.99 | 12.34 | 7.53 | 7.67 | 10.86 | 9.85 | 10.35 | 9.80 | 9.30 | 12.67 | 7.60 | 10.35 | 10.08 |
| NUP54 | 23 | 9.05 | 10.40 | 11.50 | 11.44 | 9.02 | 8.94 | 11.61 | 10.41 | 8.71 | 8.92 | 9.72 | 11.47 | 8.98 | 11.01 | 8.81 |
| NUP62 | 12 | 9.59 | 10.29 | 11.56 | 10.66 | 9.65 | 9.72 | 10.55 | 9.34 | 9.10 | 9.56 | 9.94 | 11.11 | 9.68 | 9.94 | 9.33 |
| NUP85 | 17 | 10.14 | 10.41 | 10.67 | 8.66 | 10.32 | 9.84 | 10.02 | 8.87 | 10.32 | 10.75 | 10.28 | 9.67 | 10.08 | 9.44 | 10.54 |
| NUP88 | 18 | 10.19 | 11.17 | 9.88 | 10.06 | 8.89 | 9.79 | 9.76 | 9.61 | 10.24 | 10.40 | 10.68 | 9.97 | 9.34 | 9.69 | 10.32 |
| NUP93 | 48 | 10.30 | 10.64 | 8.73 | 9.29 | 9.61 | 9.73 | 9.95 | 9.79 | 11.07 | 10.89 | 10.47 | 9.01 | 9.67 | 9.87 | 10.98 |
| NUP98 | 54 | 9.62 | 9.91 | 10.14 | 9.51 | 9.77 | 9.68 | 10.58 | 11.92 | 9.57 | 9.30 | 9.77 | 9.83 | 9.72 | 11.25 | 9.43 |
| NUPL1 | 10 | 9.15 | 10.79 | 11.56 | 11.49 | 8.65 | 9.20 | 10.98 | 10.27 | 9.34 | 8.58 | 9.97 | 11.52 | 8.92 | 10.63 | 8.96 |
| NUPL2 | 8 | 9.63 | 11.75 | 11.28 | 10.52 | 8.27 | 8.81 | 10.89 | 10.16 | 9.51 | 9.17 | 10.69 | 10.90 | 8.54 | 10.53 | 9.34 |
| NUSAP1 | 6 | 9.89 | 8.94 | 8.31 | 8.53 | 10.03 | 9.05 | 10.89 | 9.62 | 10.82 | 13.93 | 9.41 | 8.42 | 9.54 | 10.26 | 12.37 |
| NUTF2 | 15 | 9.17 | 10.16 | 9.66 | 9.41 | 11.98 | 11.87 | 9.98 | 8.43 | 9.91 | 9.44 | 9.66 | 9.53 | 11.93 | 9.20 | 9.67 |
| NVL | 22 | 9.09 | 10.26 | 7.27 | 7.19 | 11.40 | 11.94 | 12.42 | 11.60 | 9.55 | 9.28 | 9.67 | 7.23 | 11.67 | 12.01 | 9.42 |
| NXF1 | 24 | 9.10 | 10.01 | 10.43 | 9.21 | 9.94 | 9.80 | 11.88 | 10.27 | 9.66 | 9.71 | 9.55 | 9.82 | 9.87 | 11.07 | 9.69 |
| NXN | 12 | 8.85 | 10.10 | 19.42 | 19.38 | 5.45 | 5.41 | 5.23 | 5.10 | 10.45 | 10.60 | 9.48 | 19.40 | 5.43 | 5.17 | 10.52 |
| NXT1 | 2 | 10.46 | 9.60 | 10.94 | 9.65 | 9.38 | 8.13 | 10.67 | 10.10 | 11.02 | 10.04 | 10.03 | 10.30 | 8.76 | 10.39 | 10.53 |
| OARD1 | 2 | 9.40 | 9.49 | 10.75 | 10.95 | 7.97 | 7.83 | 11.96 | 11.05 | 10.35 | 10.25 | 9.45 | 10.85 | 7.90 | 11.50 | 10.30 |
| OAS3 | 5 | 13.01 | 12.81 | 7.47 | 7.72 | 13.69 | 13.34 | 5.30 | 4.99 | 10.99 | 10.67 | 12.91 | 7.60 | 13.52 | 5.15 | 10.83 |
| OAT | 10 | 10.94 | 9.39 | 8.85 | 10.82 | 11.03 | 10.08 | 10.57 | 9.67 | 9.05 | 9.59 | 10.17 | 9.83 | 10.56 | 10.12 | 9.32 |
| OBSL1 | 2 | 9.48 | 10.56 | 9.64 | 9.62 | 16.25 | 16.69 | 4.28 | 4.79 | 9.24 | 9.44 | 10.02 | 9.63 | 16.47 | 4.53 | 9.34 |
| OCC1 | 1 | 10.90 | 8.67 | 10.80 | 14.86 | 6.36 | 6.80 | 10.02 | 7.32 | 11.86 | 12.42 | 9.79 | 12.83 | 6.58 | 8.67 | 12.14 |
| OCIAD1 | 4 | 11.35 | 8.46 | 9.67 | 10.57 | 11.04 | 10.66 | 9.81 | 8.29 | 10.00 | 10.15 | 9.91 | 10.12 | 10.85 | 9.05 | 10.07 |
| OCIAD2 | 2 | 10.45 | 4.75 | 10.52 | 16.73 | 7.12 | 5.44 | 11.64 | 10.50 | 10.18 | 12.67 | 7.60 | 13.63 | 6.28 | 11.07 | 11.42 |
| OCRL | 7 | 7.95 | 8.39 | 9.43 | 9.28 | 14.12 | 15.05 | 10.04 | 9.23 | 8.44 | 8.06 | 8.17 | 9.35 | 14.59 | 9.64 | 8.25 |
| ODC1 | 1 | 13.40 | 12.66 | 8.64 | 7.60 | 10.73 | 9.52 | 7.01 | 7.25 | 12.61 | 10.58 | 13.03 | 8.12 | 10.13 | 7.13 | 11.60 |
| ODF2 | 5 | 9.68 | 10.46 | 9.70 | 9.26 | 9.31 | 10.03 | 11.42 | 10.61 | 10.58 | 8.96 | 10.07 | 9.48 | 9.67 | 11.01 | 9.77 |
| ODR4 | 1 | 10.11 | 8.87 | 7.69 | 9.72 | 10.86 | 11.26 | 10.92 | 12.13 | 8.77 | 9.68 | 9.49 | 8.70 | 11.06 | 11.52 | 9.22 |
| OFD1 | 5 | 10.96 | 12.43 | 9.42 | 9.22 | 9.00 | 9.57 | 9.47 | 9.10 | 10.66 | 10.17 | 11.69 | 9.32 | 9.29 | 9.29 | 10.41 |
| OGDH | 29 | 12.13 | 9.78 | 8.50 | 10.46 | 10.88 | 9.93 | 8.73 | 9.28 | 9.76 | 10.56 | 10.96 | 9.48 | 10.40 | 9.00 | 10.16 |
| OGFOD1 | 10 | 10.78 | 11.77 | 7.73 | 6.99 | 9.71 | 10.18 | 10.00 | 9.33 | 12.17 | 11.36 | 11.27 | 7.36 | 9.94 | 9.66 | 11.76 |
| OGFOD2 | 1 | 11.11 | 11.31 | 11.20 | 9.67 | 9.60 | 10.84 | 8.44 | 8.15 | 10.19 | 9.48 | 11.21 | 10.44 | 10.22 | 8.30 | 9.84 |
| OGFR | 19 | 10.25 | 10.01 | 9.35 | 8.17 | 8.61 | 9.04 | 10.76 | 12.41 | 12.10 | 9.29 | 10.13 | 8.76 | 8.83 | 11.58 | 10.70 |
| OGT | 29 | 9.48 | 10.05 | 10.06 | 9.93 | 11.58 | 12.22 | 9.62 | 9.04 | 9.21 | 8.79 | 9.77 | 10.00 | 11.90 | 9.33 | 9.00 |
| OLA1 | 42 | 9.32 | 10.67 | 10.18 | 8.72 | 11.01 | 11.06 | 11.56 | 10.05 | 8.77 | 8.66 | 10.00 | 9.45 | 11.04 | 10.81 | 8.71 |
| OMA1 | 1 | 12.26 | 8.02 | 9.10 | 10.89 | 9.76 | 8.25 | 9.95 | 10.61 | 10.00 | 11.15 | 10.14 | 10.00 | 9.01 | 10.28 | 10.58 |

| | | | | | | | | | | | | | | | | |
|---|---|---|---|---|---|---|---|---|---|---|---|---|---|---|---|---|
| OPA1 | 26 | 11.04 | 9.88 | 12.02 | 12.54 | 7.66 | 8.04 | 9.47 | 9.00 | 10.28 | 10.06 | 10.46 | 12.28 | 7.85 | 9.24 | 10.17 |
| OPLAH | 2 | 9.68 | 10.01 | 10.01 | 8.43 | 8.07 | 8.49 | 11.70 | 11.88 | 10.63 | 11.11 | 9.84 | 9.22 | 8.28 | 11.79 | 10.87 |
| OPTN | 13 | 9.89 | 10.59 | 11.00 | 10.32 | 13.64 | 13.95 | 5.23 | 4.75 | 10.70 | 9.93 | 10.24 | 10.66 | 13.79 | 4.99 | 10.31 |
| OR10G3 | 1 | 9.21 | 6.92 | 9.69 | 7.38 | 7.79 | 10.11 | 14.27 | 10.77 | 18.38 | 5.47 | 8.07 | 8.53 | 8.95 | 12.52 | 11.93 |
| ORC1 | 6 | 9.37 | 10.00 | 10.03 | 9.45 | 10.40 | 10.13 | 11.08 | 10.42 | 9.53 | 9.58 | 9.68 | 9.74 | 10.27 | 10.75 | 9.56 |
| ORC2 | 8 | 9.60 | 10.35 | 11.17 | 10.43 | 8.78 | 8.94 | 10.29 | 9.43 | 10.92 | 10.09 | 9.97 | 10.80 | 8.86 | 9.86 | 10.50 |
| ORC3 | 4 | 9.20 | 9.98 | 9.80 | 10.56 | 8.42 | 9.63 | 9.86 | 11.59 | 10.04 | 10.93 | 9.59 | 10.18 | 9.02 | 10.73 | 10.48 |
| ORC4 | 10 | 8.82 | 9.83 | 10.56 | 10.52 | 9.33 | 9.80 | 10.39 | 9.93 | 10.38 | 10.45 | 9.32 | 10.54 | 9.56 | 10.16 | 10.41 |
| ORC5 | 10 | 9.39 | 10.06 | 9.96 | 10.79 | 8.56 | 9.03 | 10.19 | 10.89 | 10.34 | 10.78 | 9.73 | 10.38 | 8.80 | 10.54 | 10.56 |
| ORC6 | 1 | 10.37 | 10.66 | 9.10 | 8.86 | 7.34 | 7.21 | 12.12 | 10.20 | 13.30 | 10.84 | 10.52 | 8.98 | 7.27 | 11.16 | 12.07 |
| ORMDL2 | 1 | 14.15 | 7.15 | 9.48 | 10.74 | 9.71 | 7.94 | 7.62 | 6.92 | 12.92 | 13.37 | 10.65 | 10.11 | 8.82 | 7.27 | 13.15 |
| OS9 | 8 | 10.28 | 7.74 | 8.95 | 11.48 | 11.79 | 12.34 | 11.17 | 10.84 | 7.58 | 7.81 | 9.01 | 10.22 | 12.07 | 11.01 | 7.70 |
| OSBP | 19 | 9.86 | 9.59 | 10.52 | 9.96 | 10.87 | 10.29 | 9.91 | 11.58 | 9.04 | 8.38 | 9.72 | 10.24 | 10.58 | 10.74 | 8.71 |
| OSBP2 | 4 | 9.99 | 10.29 | 6.68 | 6.58 | 10.64 | 11.26 | 11.28 | 11.06 | 11.27 | 10.94 | 10.14 | 6.63 | 10.95 | 11.17 | 11.11 |
| OSBPL10 | 3 | 9.87 | 10.92 | 9.33 | 9.28 | 8.33 | 8.61 | 9.27 | 11.91 | 11.47 | 11.00 | 10.40 | 9.31 | 8.47 | 10.59 | 11.23 |
| OSBPL11 | 9 | 10.59 | 10.64 | 10.90 | 10.63 | 9.33 | 9.63 | 8.89 | 7.60 | 11.56 | 10.22 | 10.61 | 10.77 | 9.48 | 8.25 | 10.89 |
| OSBPL1A | 2 | 9.31 | 8.95 | 13.20 | 15.12 | 7.52 | 8.72 | 9.73 | 9.83 | 8.55 | 9.08 | 9.13 | 14.16 | 8.12 | 9.78 | 8.81 |
| OSBPL2 | 3 | 9.27 | 9.92 | 10.79 | 11.22 | 10.08 | 10.69 | 8.32 | 7.98 | 11.04 | 10.69 | 9.60 | 11.00 | 10.38 | 8.15 | 10.87 |
| OSBPL3 | 9 | 8.78 | 9.28 | 14.27 | 14.50 | 7.60 | 7.83 | 9.06 | 8.21 | 10.51 | 9.97 | 9.03 | 14.38 | 7.72 | 8.63 | 10.24 |
| OSBPL6 | 5 | 7.80 | 8.14 | 9.34 | 9.28 | 12.53 | 13.57 | 7.79 | 7.44 | 12.21 | 11.90 | 7.97 | 9.31 | 13.05 | 7.62 | 12.05 |
| OSBPL8 | 5 | 10.36 | 6.05 | 9.56 | 12.79 | 8.32 | 7.64 | 11.51 | 10.10 | 10.92 | 12.76 | 8.20 | 11.17 | 7.98 | 10.80 | 11.84 |
| OSBPL9 | 13 | 10.35 | 9.40 | 12.27 | 11.22 | 9.80 | 8.31 | 9.32 | 7.60 | 10.92 | 10.79 | 9.88 | 11.75 | 9.06 | 8.46 | 10.86 |
| OSGEP | 5 | 9.50 | 10.02 | 8.74 | 8.61 | 11.72 | 12.54 | 10.60 | 9.52 | 9.85 | 8.88 | 9.76 | 8.68 | 12.13 | 10.06 | 9.37 |
| OSGEPL1 | 2 | 8.67 | 7.71 | 10.29 | 12.56 | 8.70 | 7.97 | 12.20 | 12.17 | 9.18 | 10.56 | 8.19 | 11.42 | 8.33 | 12.18 | 9.87 |
| OSMR | 1 | 7.71 | 11.43 | 8.66 | 15.14 | 11.61 | 8.38 | 10.80 | 10.33 | 8.49 | 7.45 | 9.57 | 11.90 | 10.00 | 10.56 | 7.97 |
| OSTC | 1 | 16.27 | 6.82 | 7.45 | 11.43 | 7.39 | 5.77 | 10.90 | 9.83 | 9.71 | 14.44 | 11.54 | 9.44 | 6.58 | 10.36 | 12.08 |
| OSTF1 | 3 | 9.50 | 10.17 | 8.37 | 7.73 | 14.86 | 15.00 | 7.04 | 6.35 | 10.73 | 10.23 | 9.84 | 8.05 | 14.93 | 6.70 | 10.48 |
| OSTM1 | 1 | 10.72 | 11.22 | 9.71 | 9.46 | 11.87 | 12.16 | 9.68 | 7.11 | 8.51 | 9.55 | 10.97 | 9.58 | 12.02 | 8.40 | 9.03 |
| OTC | 2 | 17.84 | 8.51 | 7.64 | 8.84 | 9.84 | 11.37 | 7.88 | 9.61 | 10.42 | 8.05 | 13.18 | 8.24 | 10.60 | 8.74 | 9.24 |
| OTUB1 | 13 | 9.19 | 10.73 | 12.05 | 10.67 | 10.73 | 11.60 | 8.12 | 8.08 | 9.50 | 9.34 | 9.96 | 11.36 | 11.16 | 8.10 | 9.42 |
| OTUD3 | 1 | 9.78 | 12.35 | 8.97 | 7.31 | 10.64 | 14.35 | 10.00 | 11.00 | 7.70 | 7.88 | 11.07 | 8.14 | 12.50 | 10.50 | 7.79 |
| OTUD4 | 7 | 8.64 | 9.30 | 11.28 | 10.64 | 9.50 | 9.74 | 11.17 | 10.06 | 10.12 | 9.56 | 8.97 | 10.96 | 9.62 | 10.61 | 9.84 |
| OTUD5 | 4 | 11.10 | 11.44 | 9.61 | 9.72 | 11.49 | 11.71 | 9.77 | 8.16 | 8.82 | 8.18 | 11.27 | 9.66 | 11.60 | 8.96 | 8.50 |
| OTUD6B | 8 | 9.06 | 10.31 | 10.11 | 10.08 | 9.21 | 9.18 | 10.88 | 10.81 | 10.27 | 10.10 | 9.68 | 10.09 | 9.20 | 10.84 | 10.19 |
| OTUD7B | 4 | 9.74 | 10.84 | 10.36 | 10.13 | 12.29 | 12.08 | 8.85 | 8.32 | 8.97 | 8.43 | 10.29 | 10.24 | 12.18 | 8.58 | 8.70 |
| OVCA2 | 2 | 9.28 | 10.13 | 11.16 | 12.35 | 10.46 | 10.49 | 7.32 | 7.50 | 10.51 | 10.80 | 9.71 | 11.75 | 10.48 | 7.41 | 10.65 |
| OXA1L | 8 | 12.46 | 6.68 | 7.86 | 10.87 | 8.71 | 7.39 | 10.54 | 9.79 | 12.12 | 13.58 | 9.57 | 9.36 | 8.05 | 10.17 | 12.85 |
| OXCT1 | 51 | 12.14 | 10.20 | 7.07 | 7.20 | 13.35 | 11.81 | 7.82 | 10.19 | 10.20 | 10.01 | 11.17 | 7.14 | 12.58 | 9.01 | 10.10 |
| OXNAD1 | 2 | 10.41 | 9.26 | 8.17 | 9.28 | 9.76 | 9.36 | 10.75 | 10.89 | 11.04 | 11.07 | 9.84 | 8.72 | 9.56 | 10.82 | 11.06 |
| OXR1 | 5 | 9.12 | 10.93 | 10.42 | 10.07 | 10.45 | 10.06 | 12.77 | 11.48 | 7.55 | 7.14 | 10.03 | 10.24 | 10.26 | 12.13 | 7.35 |
| OXSM | 1 | 9.37 | 8.43 | 9.59 | 10.52 | 10.02 | 12.14 | 11.08 | 11.23 | 8.30 | 9.32 | 8.90 | 10.05 | 11.08 | 11.16 | 8.81 |
| OXSR1 | 22 | 9.30 | 10.07 | 10.28 | 10.06 | 11.10 | 11.76 | 8.78 | 8.53 | 10.32 | 9.81 | 9.68 | 10.17 | 11.43 | 8.65 | 10.06 |

| | | | | | | | | | | | | | | | | |
|---|---|---|---|---|---|---|---|---|---|---|---|---|---|---|---|---|
| P4HA1 | 23 | 9.44 | 7.93 | 7.68 | 8.17 | 8.25 | 8.24 | 12.71 | 16.27 | 10.59 | 10.72 | 8.68 | 7.93 | 8.25 | 14.49 | 10.65 |
| P4HA2 | 25 | 11.94 | 9.06 | 8.31 | 7.62 | 9.37 | 8.87 | 11.73 | 14.10 | 9.31 | 9.68 | 10.50 | 7.97 | 9.12 | 12.92 | 9.49 |
| P4HB | 92 | 10.72 | 8.99 | 8.79 | 9.19 | 10.68 | 10.24 | 8.53 | 10.06 | 11.24 | 11.55 | 9.86 | 8.99 | 10.46 | 9.29 | 11.39 |
| P4HTM | 4 | 10.55 | 9.68 | 5.85 | 6.30 | 13.39 | 14.26 | 10.70 | 9.26 | 9.78 | 10.22 | 10.12 | 6.07 | 13.83 | 9.98 | 10.00 |
| PA2G4 | 71 | 8.59 | 9.51 | 9.77 | 9.30 | 9.00 | 9.35 | 11.82 | 15.48 | 8.65 | 8.53 | 9.05 | 9.54 | 9.17 | 13.65 | 8.59 |
| PAAF1 | 6 | 10.80 | 11.71 | 8.95 | 10.47 | 9.19 | 10.26 | 9.97 | 9.42 | 9.61 | 9.62 | 11.25 | 9.71 | 9.72 | 9.70 | 9.62 |
| PABPC1 | 61 | 8.87 | 10.22 | 10.48 | 9.95 | 9.79 | 10.01 | 11.27 | 10.35 | 9.70 | 9.35 | 9.55 | 10.21 | 9.90 | 10.81 | 9.53 |
| PABPC4 | 53 | 9.26 | 10.06 | 9.31 | 8.86 | 9.17 | 8.85 | 11.29 | 13.85 | 9.89 | 9.46 | 9.66 | 9.09 | 9.01 | 12.57 | 9.68 |
| PABPN1 | 15 | 9.50 | 10.32 | 10.42 | 10.29 | 9.36 | 9.54 | 11.10 | 10.01 | 9.73 | 9.72 | 9.91 | 10.35 | 9.45 | 10.56 | 9.73 |
| PACS1 | 12 | 8.51 | 9.80 | 13.31 | 12.77 | 10.68 | 10.97 | 7.49 | 6.86 | 10.09 | 9.53 | 9.15 | 13.04 | 10.82 | 7.17 | 9.81 |
| PACSIN2 | 23 | 9.29 | 9.10 | 17.96 | 12.75 | 6.68 | 6.44 | 9.88 | 7.80 | 10.21 | 9.89 | 9.19 | 15.36 | 6.56 | 8.84 | 10.05 |
| PACSIN3 | 13 | 9.93 | 10.31 | 8.49 | 8.28 | 11.48 | 11.95 | 11.01 | 9.82 | 9.53 | 9.21 | 10.12 | 8.39 | 11.71 | 10.41 | 9.37 |
| PAF1 | 23 | 9.54 | 10.20 | 10.03 | 9.18 | 10.74 | 10.83 | 9.95 | 8.66 | 10.82 | 10.05 | 9.87 | 9.61 | 10.79 | 9.30 | 10.43 |
| PAFAH1B1 | 17 | 11.11 | 12.75 | 12.34 | 12.43 | 8.49 | 8.60 | 7.44 | 6.17 | 10.63 | 10.05 | 11.93 | 12.38 | 8.54 | 6.81 | 10.34 |
| PAFAH1B2 | 4 | 9.14 | 9.17 | 11.88 | 10.15 | 10.28 | 9.60 | 10.56 | 8.74 | 10.64 | 9.84 | 9.16 | 11.02 | 9.94 | 9.65 | 10.24 |
| PAFAH1B3 | 10 | 8.88 | 10.57 | 10.15 | 10.52 | 9.10 | 9.35 | 10.34 | 11.85 | 9.51 | 9.73 | 9.72 | 10.34 | 9.22 | 11.10 | 9.62 |
| PAFAH2 | 1 | 12.77 | 12.15 | 8.19 | 9.18 | 8.42 | 7.73 | 8.42 | 9.15 | 13.89 | 10.10 | 12.46 | 8.69 | 8.07 | 8.78 | 12.00 |
| PAGR1 | 2 | 19.42 | 13.28 | 5.91 | 10.48 | 6.51 | 6.90 | 9.71 | 5.98 | 12.19 | 9.62 | 16.35 | 8.20 | 6.70 | 7.85 | 10.91 |
| PAICS | 25 | 9.41 | 10.07 | 9.06 | 8.79 | 10.18 | 10.47 | 8.83 | 14.44 | 9.36 | 9.40 | 9.74 | 8.92 | 10.32 | 11.63 | 9.38 |
| PAIP1 | 29 | 8.46 | 10.01 | 8.12 | 7.90 | 10.71 | 10.96 | 13.12 | 13.38 | 8.52 | 8.83 | 9.23 | 8.01 | 10.83 | 13.25 | 8.67 |
| PAIP2 | 4 | 10.15 | 13.09 | 8.64 | 9.45 | 9.87 | 11.30 | 12.60 | 5.81 | 9.16 | 9.93 | 11.62 | 9.04 | 10.58 | 9.21 | 9.54 |
| PAK1 | 13 | 9.52 | 10.34 | 12.15 | 12.47 | 8.33 | 8.41 | 10.72 | 10.27 | 8.97 | 8.83 | 9.93 | 12.31 | 8.37 | 10.50 | 8.90 |
| PAK1IP1 | 10 | 9.15 | 9.94 | 10.64 | 11.02 | 8.18 | 8.68 | 10.53 | 11.16 | 10.44 | 10.26 | 9.55 | 10.83 | 8.43 | 10.85 | 10.35 |
| PAK2 | 28 | 9.63 | 10.77 | 11.73 | 10.70 | 9.75 | 9.88 | 9.52 | 9.06 | 9.73 | 9.24 | 10.20 | 11.21 | 9.81 | 9.29 | 9.48 |
| PAK4 | 3 | 8.75 | 9.84 | 11.40 | 10.76 | 8.19 | 8.29 | 11.12 | 10.48 | 10.57 | 10.61 | 9.29 | 11.08 | 8.24 | 10.80 | 10.59 |
| PALLD | 56 | 10.66 | 11.67 | 12.40 | 11.58 | 8.62 | 8.74 | 4.24 | 4.02 | 14.29 | 13.78 | 11.16 | 11.99 | 8.68 | 4.13 | 14.04 |
| PALM | 11 | 14.07 | 11.53 | 5.88 | 6.90 | 14.04 | 14.21 | 7.35 | 5.55 | 10.59 | 9.89 | 12.80 | 6.39 | 14.12 | 6.45 | 10.24 |
| PALM2 | 8 | 9.64 | 10.44 | 14.84 | 14.93 | 4.65 | 5.67 | 8.42 | 8.18 | 11.64 | 11.59 | 10.04 | 14.88 | 5.16 | 8.30 | 11.61 |
| PALMD | 11 | 13.60 | 15.41 | 8.68 | 7.83 | 9.83 | 9.90 | 6.08 | 5.30 | 12.18 | 11.17 | 14.51 | 8.26 | 9.87 | 5.69 | 11.68 |
| PAM | 2 | 8.37 | 6.71 | 11.13 | 13.55 | 7.55 | 8.93 | 11.77 | 12.06 | 9.58 | 10.35 | 7.54 | 12.34 | 8.24 | 11.92 | 9.96 |
| PANK1 | 5 | 11.77 | 10.39 | 10.91 | 9.84 | 10.75 | 7.93 | 10.74 | 10.03 | 8.91 | 8.73 | 11.08 | 10.38 | 9.34 | 10.38 | 8.82 |
| PANK2 | 2 | 9.47 | 10.26 | 11.98 | 12.01 | 8.92 | 9.38 | 9.27 | 9.51 | 8.90 | 10.30 | 9.86 | 11.99 | 9.15 | 9.39 | 9.60 |
| PANK4 | 8 | 9.41 | 10.72 | 11.48 | 10.32 | 8.92 | 8.45 | 10.53 | 9.18 | 10.07 | 10.92 | 10.07 | 10.90 | 8.69 | 9.85 | 10.50 |
| PAPD5 | 5 | 9.90 | 10.08 | 8.89 | 8.93 | 9.24 | 8.51 | 11.76 | 11.55 | 11.08 | 10.06 | 9.99 | 8.91 | 8.87 | 11.65 | 10.57 |
| PAPOLA | 16 | 9.84 | 10.31 | 10.58 | 9.77 | 7.83 | 8.08 | 11.69 | 11.49 | 10.20 | 10.22 | 10.07 | 10.17 | 7.95 | 11.59 | 10.21 |
| PAPOLG | 1 | 8.96 | 12.26 | 9.75 | 9.59 | 8.80 | 10.55 | 8.92 | 9.64 | 10.45 | 11.07 | 10.61 | 9.67 | 9.67 | 9.28 | 10.76 |
| PAPSS1 | 30 | 9.98 | 10.84 | 10.15 | 9.08 | 10.12 | 9.85 | 10.13 | 10.22 | 9.89 | 9.73 | 10.41 | 9.62 | 9.99 | 10.18 | 9.81 |
| PAPSS2 | 20 | 10.19 | 11.23 | 14.50 | 13.86 | 11.64 | 12.20 | 3.79 | 3.25 | 9.74 | 9.61 | 10.71 | 14.18 | 11.92 | 3.52 | 9.67 |
| PARD3 | 14 | 8.12 | 9.07 | 12.86 | 12.38 | 9.27 | 9.75 | 9.04 | 7.96 | 10.88 | 10.67 | 8.60 | 12.62 | 9.51 | 8.50 | 10.77 |
| PARG | 2 | 8.74 | 8.91 | 11.31 | 9.95 | 10.30 | 8.90 | 10.68 | 10.53 | 10.22 | 10.45 | 8.83 | 10.63 | 9.60 | 10.61 | 10.34 |
| PARK7 | 46 | 9.60 | 11.33 | 8.98 | 8.75 | 10.28 | 10.08 | 9.36 | 13.16 | 9.52 | 8.95 | 10.46 | 8.87 | 10.18 | 11.26 | 9.23 |
| PARN | 18 | 9.74 | 10.06 | 11.03 | 9.63 | 10.27 | 10.29 | 10.67 | 9.84 | 9.38 | 9.08 | 9.90 | 10.33 | 10.28 | 10.25 | 9.23 |

| | | | | | | | | | | | | | | | | |
|---|---|---|---|---|---|---|---|---|---|---|---|---|---|---|---|---|
| PARP1 | 93 | 9.18 | 10.40 | 8.08 | 7.23 | 12.20 | 12.20 | 11.25 | 11.60 | 9.16 | 8.70 | 9.79 | 7.65 | 12.20 | 11.43 | 8.93 |
| PARP10 | 1 | 10.80 | 10.42 | 8.08 | 8.45 | 11.55 | 15.10 | 7.76 | 7.41 | 11.25 | 9.18 | 10.61 | 8.26 | 13.33 | 7.59 | 10.22 |
| PARP12 | 5 | 12.15 | 12.67 | 7.68 | 7.86 | 9.47 | 10.03 | 9.32 | 9.15 | 10.94 | 10.73 | 12.41 | 7.77 | 9.75 | 9.23 | 10.84 |
| PARP16 | 1 | 10.29 | 6.49 | 6.52 | 10.22 | 9.09 | 7.70 | 9.82 | 10.30 | 11.96 | 17.61 | 8.39 | 8.37 | 8.39 | 10.06 | 14.78 |
| PARP2 | 2 | 9.87 | 8.43 | 10.69 | 9.48 | 10.54 | 10.32 | 11.07 | 9.80 | 9.66 | 10.14 | 9.15 | 10.09 | 10.43 | 10.43 | 9.90 |
| PARP4 | 15 | 10.22 | 10.36 | 9.67 | 9.20 | 14.56 | 14.88 | 9.43 | 9.55 | 6.05 | 6.08 | 10.29 | 9.44 | 14.72 | 9.49 | 6.06 |
| PARP9 | 2 | 10.47 | 10.01 | 7.25 | 6.44 | 16.35 | 17.90 | 9.53 | 8.92 | 7.47 | 5.65 | 10.24 | 6.84 | 17.13 | 9.22 | 6.56 |
| PARS2 | 1 | 12.25 | 11.42 | 7.92 | 9.14 | 9.23 | 7.85 | 10.45 | 10.34 | 10.76 | 10.64 | 11.83 | 8.53 | 8.54 | 10.39 | 10.70 |
| PARVA | 20 | 10.23 | 10.52 | 13.47 | 13.11 | 8.82 | 9.13 | 6.50 | 6.81 | 10.87 | 10.53 | 10.38 | 13.29 | 8.98 | 6.66 | 10.70 |
| PARVB | 6 | 8.85 | 8.88 | 14.47 | 14.72 | 5.48 | 5.18 | 8.29 | 7.79 | 12.95 | 13.39 | 8.86 | 14.59 | 5.33 | 8.04 | 13.17 |
| PATL1 | 11 | 8.74 | 9.83 | 10.49 | 9.73 | 10.20 | 10.33 | 10.97 | 10.71 | 9.93 | 9.06 | 9.29 | 10.11 | 10.27 | 10.84 | 9.50 |
| PAWR | 11 | 8.41 | 11.31 | 14.48 | 13.33 | 9.05 | 8.40 | 7.27 | 6.05 | 10.74 | 10.95 | 9.86 | 13.91 | 8.72 | 6.66 | 10.85 |
| PAXBP1 | 5 | 9.17 | 10.88 | 8.52 | 9.55 | 9.44 | 9.97 | 10.72 | 12.39 | 9.52 | 9.83 | 10.03 | 9.04 | 9.71 | 11.56 | 9.67 |
| PBDC1 | 7 | 10.99 | 12.13 | 11.39 | 9.15 | 8.55 | 8.86 | 8.01 | 13.39 | 9.21 | 8.32 | 11.56 | 10.27 | 8.71 | 10.70 | 8.76 |
| PBK | 8 | 8.28 | 9.69 | 9.62 | 7.67 | 9.51 | 8.78 | 12.78 | 16.96 | 8.58 | 8.14 | 8.98 | 8.64 | 9.15 | 14.87 | 8.36 |
| PBRM1 | 18 | 9.92 | 11.15 | 9.16 | 7.75 | 11.15 | 11.85 | 10.91 | 10.71 | 8.67 | 8.74 | 10.54 | 8.46 | 11.50 | 10.81 | 8.70 |
| PBX1 | 5 | 9.36 | 10.10 | 10.28 | 9.22 | 12.44 | 12.67 | 9.83 | 9.76 | 8.21 | 8.15 | 9.73 | 9.75 | 12.55 | 9.79 | 8.18 |
| PBXIP1 | 8 | 11.76 | 7.29 | 9.14 | 12.49 | 13.86 | 11.88 | 6.56 | 6.14 | 9.51 | 11.36 | 9.53 | 10.82 | 12.87 | 6.35 | 10.43 |
| PC | 35 | 12.21 | 11.35 | 8.19 | 9.40 | 10.26 | 9.20 | 11.34 | 10.56 | 8.51 | 8.98 | 11.78 | 8.80 | 9.73 | 10.95 | 8.74 |
| PCBD1 | 3 | 10.15 | 10.81 | 10.60 | 10.25 | 14.18 | 14.20 | 5.26 | 4.09 | 10.34 | 10.13 | 10.48 | 10.42 | 14.19 | 4.67 | 10.23 |
| PCBD2 | 3 | 9.93 | 11.40 | 8.62 | 9.40 | 14.50 | 15.10 | 7.20 | 6.16 | 8.98 | 8.71 | 10.66 | 9.01 | 14.80 | 6.68 | 8.84 |
| PCBP1 | 24 | 7.02 | 7.05 | 8.26 | 5.36 | 9.89 | 7.99 | 8.08 | 32.32 | 7.38 | 6.65 | 7.03 | 6.81 | 8.94 | 20.20 | 7.01 |
| PCBP2 | 45 | 9.25 | 9.85 | 9.79 | 9.40 | 9.35 | 9.43 | 10.32 | 14.84 | 9.18 | 8.59 | 9.55 | 9.59 | 9.39 | 12.58 | 8.88 |
| PCBP4 | 4 | 11.11 | 10.68 | 9.85 | 9.34 | 10.34 | 9.53 | 10.58 | 9.43 | 9.97 | 9.16 | 10.90 | 9.59 | 9.94 | 10.00 | 9.57 |
| PCCA | 3 | 12.13 | 10.11 | 8.10 | 8.75 | 9.91 | 9.31 | 11.42 | 12.82 | 8.28 | 9.17 | 11.12 | 8.43 | 9.61 | 12.12 | 8.73 |
| PCCB | 2 | 12.60 | 10.04 | 9.63 | 10.14 | 9.82 | 8.93 | 11.60 | 9.76 | 8.06 | 9.43 | 11.32 | 9.88 | 9.38 | 10.68 | 8.74 |
| PCDH7 | 2 | 7.15 | 6.72 | 9.83 | 9.99 | 9.98 | 9.37 | 17.29 | 14.96 | 7.71 | 7.00 | 6.94 | 9.91 | 9.68 | 16.12 | 7.36 |
| PCF11 | 8 | 8.27 | 9.81 | 9.22 | 9.39 | 9.87 | 10.54 | 12.53 | 10.76 | 10.04 | 9.58 | 9.04 | 9.30 | 10.20 | 11.64 | 9.81 |
| PCGF2 | 4 | 9.31 | 9.59 | 11.60 | 11.29 | 9.31 | 9.99 | 11.14 | 9.79 | 9.40 | 8.59 | 9.45 | 11.44 | 9.65 | 10.46 | 9.00 |
| PCID2 | 14 | 10.78 | 11.21 | 8.56 | 8.64 | 9.93 | 10.71 | 10.01 | 10.76 | 9.35 | 10.06 | 11.00 | 8.60 | 10.32 | 10.38 | 9.70 |
| PCIF1 | 4 | 8.94 | 9.55 | 9.92 | 10.51 | 10.19 | 10.40 | 11.00 | 10.11 | 9.57 | 9.80 | 9.25 | 10.22 | 10.29 | 10.56 | 9.69 |
| PCK2 | 7 | 8.24 | 8.62 | 6.06 | 6.83 | 7.79 | 9.33 | 8.76 | 8.02 | 17.65 | 18.69 | 8.43 | 6.44 | 8.56 | 8.39 | 18.17 |
| PCM1 | 22 | 9.57 | 11.44 | 8.71 | 10.51 | 9.70 | 11.26 | 10.28 | 10.51 | 8.82 | 9.20 | 10.50 | 9.61 | 10.48 | 10.40 | 9.01 |
| PCMT1 | 19 | 9.92 | 11.18 | 10.34 | 9.38 | 10.47 | 10.01 | 8.63 | 10.62 | 9.89 | 9.55 | 10.55 | 9.86 | 10.24 | 9.63 | 9.72 |
| PCNA | 49 | 9.20 | 10.04 | 8.31 | 8.08 | 10.03 | 10.42 | 11.17 | 14.21 | 9.43 | 9.10 | 9.62 | 8.20 | 10.23 | 12.69 | 9.27 |
| PCNP | 17 | 8.94 | 9.78 | 11.00 | 8.97 | 11.17 | 10.51 | 11.55 | 9.49 | 9.53 | 9.06 | 9.36 | 9.98 | 10.84 | 10.52 | 9.29 |
| PCNT | 10 | 11.23 | 11.52 | 8.39 | 8.69 | 10.38 | 10.58 | 10.17 | 9.35 | 10.03 | 9.65 | 11.38 | 8.54 | 10.48 | 9.76 | 9.84 |
| PCYOX1 | 11 | 10.09 | 7.86 | 8.08 | 7.94 | 13.27 | 12.06 | 10.27 | 11.51 | 9.17 | 9.75 | 8.97 | 8.01 | 12.67 | 10.89 | 9.46 |
| PCYT1A | 8 | 10.62 | 11.66 | 13.38 | 12.32 | 9.74 | 9.85 | 7.19 | 6.17 | 9.88 | 9.20 | 11.14 | 12.85 | 9.79 | 6.68 | 9.54 |
| PCYT2 | 8 | 12.21 | 12.82 | 11.16 | 10.71 | 8.70 | 8.61 | 6.15 | 5.57 | 12.49 | 11.59 | 12.52 | 10.93 | 8.65 | 5.86 | 12.04 |
| PDAP1 | 21 | 8.23 | 10.53 | 14.21 | 11.86 | 8.76 | 8.46 | 9.92 | 8.22 | 10.01 | 9.80 | 9.38 | 13.03 | 8.61 | 9.07 | 9.90 |
| PDCD10 | 12 | 9.34 | 9.79 | 10.88 | 10.13 | 10.02 | 10.12 | 10.56 | 10.27 | 9.48 | 9.42 | 9.57 | 10.50 | 10.07 | 10.42 | 9.45 |

| | | | | | | | | | | | | | | | | |
|---|---|---|---|---|---|---|---|---|---|---|---|---|---|---|---|---|
| PDCD11 | 45 | 9.06 | 9.00 | 10.41 | 9.23 | 9.40 | 8.94 | 11.71 | 11.21 | 10.62 | 10.41 | 9.03 | 9.82 | 9.17 | 11.46 | 10.52 |
| PDCD2 | 4 | 9.06 | 12.24 | 11.76 | 12.43 | 6.16 | 5.85 | 10.53 | 9.99 | 11.29 | 10.69 | 10.65 | 12.09 | 6.01 | 10.26 | 10.99 |
| PDCD2L | 5 | 10.18 | 11.34 | 9.55 | 9.45 | 8.76 | 9.15 | 10.64 | 10.25 | 10.38 | 10.31 | 10.76 | 9.50 | 8.95 | 10.45 | 10.34 |
| PDCD4 | 20 | 9.64 | 10.58 | 6.97 | 7.28 | 14.69 | 15.26 | 9.80 | 10.73 | 7.80 | 7.23 | 10.11 | 7.13 | 14.98 | 10.27 | 7.52 |
| PDCD5 | 16 | 9.65 | 10.71 | 9.62 | 9.93 | 10.60 | 10.78 | 11.78 | 8.05 | 9.52 | 9.35 | 10.18 | 9.77 | 10.69 | 9.92 | 9.44 |
| PDCD6 | 17 | 10.09 | 10.54 | 7.25 | 6.13 | 13.58 | 13.87 | 10.91 | 9.70 | 9.00 | 8.93 | 10.32 | 6.69 | 13.72 | 10.31 | 8.96 |
| PDCD6IP | 68 | 9.67 | 10.31 | 11.05 | 10.41 | 10.38 | 10.33 | 8.73 | 8.90 | 10.38 | 9.83 | 9.99 | 10.73 | 10.36 | 8.81 | 10.11 |
| PDCL | 5 | 9.69 | 10.37 | 11.56 | 11.08 | 10.10 | 10.37 | 8.64 | 8.05 | 10.10 | 10.03 | 10.03 | 11.32 | 10.23 | 8.35 | 10.07 |
| PDCL3 | 12 | 10.19 | 10.48 | 11.98 | 11.37 | 8.83 | 8.50 | 8.94 | 8.05 | 11.16 | 10.50 | 10.34 | 11.67 | 8.66 | 8.50 | 10.83 |
| PDE12 | 8 | 7.20 | 10.13 | 10.66 | 10.10 | 10.24 | 9.75 | 10.48 | 9.92 | 9.93 | 11.59 | 8.67 | 10.38 | 9.99 | 10.20 | 10.76 |
| PDE2A | 3 | 8.66 | 9.86 | 12.80 | 11.66 | 8.89 | 9.23 | 8.70 | 9.89 | 10.92 | 9.40 | 9.26 | 12.23 | 9.06 | 9.29 | 10.16 |
| PDE3A | 21 | 9.48 | 9.50 | 10.58 | 10.11 | 10.32 | 10.23 | 9.81 | 9.96 | 10.19 | 9.81 | 9.49 | 10.35 | 10.28 | 9.89 | 10.00 |
| PDE4D | 17 | 10.32 | 10.95 | 7.34 | 6.98 | 12.93 | 13.54 | 11.49 | 10.84 | 7.82 | 7.78 | 10.64 | 7.16 | 13.24 | 11.16 | 7.80 |
| PDE4DIP | 6 | 9.56 | 11.36 | 9.13 | 9.46 | 10.79 | 11.77 | 9.07 | 9.31 | 10.27 | 9.29 | 10.46 | 9.29 | 11.28 | 9.19 | 9.78 |
| PDE6D | 2 | 10.99 | 10.53 | 10.30 | 10.54 | 11.40 | 12.41 | 9.26 | 10.63 | 6.65 | 7.29 | 10.76 | 10.42 | 11.91 | 9.94 | 6.97 |
| PDE8A | 1 | 11.01 | 9.26 | 10.14 | 9.71 | 10.01 | 10.19 | 7.92 | 6.93 | 13.59 | 11.22 | 10.14 | 9.93 | 10.10 | 7.42 | 12.41 |
| PDE8B | 2 | 13.22 | 14.46 | 8.57 | 8.15 | 10.86 | 11.77 | 7.68 | 7.19 | 9.11 | 8.99 | 13.84 | 8.36 | 11.31 | 7.44 | 9.05 |
| PDHA1 | 18 | 10.98 | 9.85 | 9.08 | 10.90 | 11.70 | 11.57 | 9.02 | 8.51 | 9.20 | 9.20 | 10.41 | 9.99 | 11.63 | 8.76 | 9.20 |
| PDHB | 23 | 10.74 | 9.28 | 10.05 | 11.05 | 11.83 | 11.15 | 8.98 | 7.99 | 9.24 | 9.68 | 10.01 | 10.55 | 11.49 | 8.49 | 9.46 |
| PDHX | 8 | 10.84 | 9.54 | 9.43 | 10.86 | 10.51 | 9.95 | 10.21 | 8.60 | 10.06 | 9.99 | 10.19 | 10.15 | 10.23 | 9.41 | 10.02 |
| PDIA3 | 102 | 8.40 | 8.56 | 8.30 | 8.47 | 12.42 | 12.12 | 11.06 | 11.00 | 9.74 | 9.95 | 8.48 | 8.38 | 12.27 | 11.03 | 9.84 |
| PDIA4 | 64 | 9.73 | 8.40 | 10.18 | 10.71 | 9.13 | 8.81 | 12.64 | 11.56 | 9.35 | 9.49 | 9.07 | 10.45 | 8.97 | 12.10 | 9.42 |
| PDIA5 | 1 | 8.45 | 11.21 | 8.50 | 11.49 | 6.97 | 9.29 | 12.38 | 11.03 | 9.98 | 10.70 | 9.83 | 10.00 | 8.13 | 11.71 | 10.34 |
| PDIA6 | 26 | 9.71 | 9.18 | 8.88 | 9.32 | 11.83 | 11.74 | 10.98 | 10.42 | 8.98 | 8.96 | 9.45 | 9.10 | 11.78 | 10.70 | 8.97 |
| PDK1 | 2 | 9.81 | 10.22 | 9.22 | 11.05 | 6.82 | 6.84 | 13.84 | 14.18 | 9.36 | 8.66 | 10.02 | 10.14 | 6.83 | 14.01 | 9.01 |
| PDK2 | 1 | 11.40 | 11.96 | 9.25 | 12.19 | 11.54 | 10.30 | 7.71 | 7.91 | 8.50 | 9.23 | 11.68 | 10.72 | 10.92 | 7.81 | 8.86 |
| PDK3 | 3 | 11.76 | 8.42 | 5.31 | 5.32 | 10.18 | 10.19 | 16.54 | 16.41 | 8.15 | 7.72 | 10.09 | 5.32 | 10.18 | 16.47 | 7.93 |
| PDLIM1 | 15 | 9.53 | 11.04 | 12.39 | 10.88 | 9.09 | 9.35 | 7.46 | 7.21 | 11.67 | 11.38 | 10.29 | 11.63 | 9.22 | 7.33 | 11.53 |
| PDLIM2 | 3 | 8.39 | 10.79 | 12.46 | 10.75 | 10.58 | 10.59 | 6.45 | 6.13 | 12.51 | 11.35 | 9.59 | 11.60 | 10.59 | 6.29 | 11.93 |
| PDLIM3 | 6 | 10.69 | 9.73 | 14.94 | 14.00 | 6.83 | 7.75 | 10.50 | 9.75 | 8.38 | 7.42 | 10.21 | 14.47 | 7.29 | 10.13 | 7.90 |
| PDLIM5 | 50 | 9.79 | 10.72 | 15.06 | 13.84 | 6.90 | 6.97 | 5.14 | 5.79 | 12.99 | 12.81 | 10.25 | 14.45 | 6.93 | 5.46 | 12.90 |
| PDLIM7 | 31 | 9.48 | 10.26 | 13.55 | 11.89 | 8.74 | 8.64 | 7.08 | 6.91 | 12.03 | 11.42 | 9.87 | 12.72 | 8.69 | 7.00 | 11.72 |
| PDP1 | 4 | 7.13 | 6.08 | 12.77 | 14.53 | 7.77 | 7.72 | 12.32 | 12.09 | 9.44 | 10.15 | 6.61 | 13.65 | 7.74 | 12.20 | 9.80 |
| PDPK1 | 4 | 10.48 | 11.06 | 10.36 | 9.38 | 11.07 | 11.70 | 8.18 | 7.89 | 9.80 | 10.08 | 10.77 | 9.87 | 11.38 | 8.04 | 9.94 |
| PDPR | 2 | 9.72 | 8.02 | 7.56 | 9.25 | 10.20 | 10.32 | 11.72 | 10.69 | 10.24 | 12.28 | 8.87 | 8.40 | 10.26 | 11.21 | 11.26 |
| PDRG1 | 3 | 8.41 | 9.62 | 11.95 | 10.95 | 9.23 | 9.55 | 14.35 | 11.81 | 7.56 | 6.57 | 9.01 | 11.45 | 9.39 | 13.08 | 7.06 |
| PDS5A | 44 | 8.76 | 8.95 | 8.70 | 8.62 | 9.90 | 10.00 | 13.40 | 12.87 | 9.39 | 9.41 | 8.86 | 8.66 | 9.95 | 13.14 | 9.40 |
| PDS5B | 43 | 9.57 | 10.16 | 9.59 | 9.59 | 11.80 | 11.61 | 10.26 | 9.91 | 8.68 | 8.84 | 9.87 | 9.59 | 11.70 | 10.08 | 8.76 |
| PDSS2 | 3 | 11.83 | 10.57 | 9.74 | 9.96 | 9.67 | 8.60 | 9.96 | 11.04 | 9.59 | 9.03 | 11.20 | 9.85 | 9.14 | 10.50 | 9.31 |
| PDXDC1 | 28 | 10.06 | 11.09 | 8.75 | 8.59 | 10.81 | 10.95 | 9.15 | 9.18 | 10.95 | 10.48 | 10.57 | 8.67 | 10.88 | 9.17 | 10.71 |
| PDXK | 13 | 10.79 | 12.01 | 10.00 | 7.18 | 11.97 | 11.06 | 6.87 | 5.49 | 12.55 | 12.07 | 11.40 | 8.59 | 11.51 | 6.18 | 12.31 |
| PDXP | 5 | 8.84 | 9.76 | 11.80 | 11.12 | 9.74 | 8.74 | 10.86 | 9.42 | 9.90 | 9.81 | 9.30 | 11.46 | 9.24 | 10.14 | 9.86 |

| | | | | | | | | | | | | | | | | |
|---|---|---|---|---|---|---|---|---|---|---|---|---|---|---|---|---|
| PDZD11 | 4 | 9.43 | 9.56 | 10.08 | 8.63 | 11.12 | 12.05 | 11.17 | 10.82 | 8.84 | 8.30 | 9.50 | 9.36 | 11.58 | 11.00 | 8.57 |
| PDZD8 | 1 | 8.90 | 8.89 | 10.24 | 10.04 | 8.91 | 11.15 | 8.68 | 8.38 | 13.91 | 10.90 | 8.90 | 10.14 | 10.03 | 8.53 | 12.41 |
| PEA15 | 6 | 10.26 | 10.83 | 14.64 | 14.11 | 10.92 | 10.72 | 5.67 | 3.11 | 9.84 | 9.91 | 10.55 | 14.37 | 10.82 | 4.39 | 9.87 |
| PEAK1 | 5 | 10.31 | 10.87 | 13.49 | 14.11 | 8.33 | 8.76 | 5.46 | 4.86 | 11.88 | 11.95 | 10.59 | 13.80 | 8.54 | 5.16 | 11.91 |
| PEBP1 | 38 | 9.05 | 10.26 | 9.90 | 8.59 | 11.90 | 11.31 | 10.20 | 10.84 | 9.26 | 8.70 | 9.65 | 9.24 | 11.60 | 10.52 | 8.98 |
| PECR | 3 | 7.89 | 8.27 | 8.66 | 8.00 | 15.40 | 14.94 | 10.17 | 10.03 | 8.65 | 7.98 | 8.08 | 8.33 | 15.17 | 10.10 | 8.32 |
| PEF1 | 4 | 10.92 | 12.24 | 8.59 | 9.12 | 9.49 | 9.16 | 8.66 | 9.01 | 11.23 | 11.58 | 11.58 | 8.86 | 9.33 | 8.83 | 11.41 |
| PEG10 | 5 | 11.82 | 12.58 | 7.60 | 7.05 | 13.91 | 13.34 | 7.89 | 6.59 | 9.97 | 9.23 | 12.20 | 7.32 | 13.63 | 7.24 | 9.60 |
| PELO | 10 | 8.98 | 9.33 | 8.78 | 8.91 | 9.30 | 9.19 | 9.92 | 18.60 | 8.66 | 8.33 | 9.16 | 8.84 | 9.25 | 14.26 | 8.49 |
| PELP1 | 20 | 9.03 | 9.43 | 10.31 | 9.56 | 10.21 | 9.78 | 9.78 | 9.94 | 10.88 | 11.09 | 9.23 | 9.94 | 9.99 | 9.86 | 10.98 |
| PEO1 | 3 | 11.42 | 11.94 | 7.80 | 10.32 | 8.57 | 9.41 | 10.55 | 9.52 | 10.27 | 10.21 | 11.68 | 9.06 | 8.99 | 10.03 | 10.24 |
| PEPD | 9 | 8.54 | 8.87 | 9.80 | 10.35 | 13.34 | 14.43 | 10.98 | 10.81 | 6.59 | 6.28 | 8.71 | 10.08 | 13.89 | 10.90 | 6.43 |
| PES1 | 17 | 9.87 | 9.90 | 10.88 | 10.66 | 7.48 | 7.70 | 11.68 | 10.56 | 10.26 | 11.01 | 9.88 | 10.77 | 7.59 | 11.12 | 10.64 |
| PET100 | 1 | 10.00 | 11.43 | 9.72 | 10.72 | 10.37 | 11.77 | 10.71 | 9.24 | 8.22 | 7.82 | 10.72 | 10.22 | 11.07 | 9.97 | 8.02 |
| PET112 | 2 | 11.04 | 10.22 | 9.09 | 12.13 | 8.52 | 9.26 | 11.03 | 9.57 | 9.70 | 9.43 | 10.63 | 10.61 | 8.89 | 10.30 | 9.57 |
| PEX1 | 5 | 10.26 | 11.35 | 9.49 | 10.77 | 8.68 | 10.23 | 8.89 | 8.95 | 10.13 | 11.25 | 10.80 | 10.13 | 9.45 | 8.92 | 10.69 |
| PEX11B | 1 | 9.89 | 6.51 | 8.17 | 12.33 | 13.61 | 11.34 | 8.65 | 8.09 | 10.18 | 11.24 | 8.20 | 10.25 | 12.48 | 8.37 | 10.71 |
| PEX14 | 5 | 13.04 | 9.01 | 9.38 | 11.64 | 7.87 | 7.57 | 10.64 | 9.82 | 10.03 | 10.99 | 11.03 | 10.51 | 7.72 | 10.23 | 10.51 |
| PEX19 | 15 | 9.86 | 11.55 | 9.91 | 9.75 | 13.22 | 13.71 | 8.96 | 5.17 | 9.08 | 8.80 | 10.71 | 9.83 | 13.46 | 7.06 | 8.94 |
| PEX3 | 2 | 11.30 | 7.01 | 9.43 | 11.03 | 9.65 | 8.21 | 10.58 | 8.16 | 11.21 | 13.41 | 9.15 | 10.23 | 8.93 | 9.37 | 12.31 |
| PEX5 | 4 | 9.79 | 9.90 | 11.01 | 10.72 | 9.85 | 8.87 | 11.11 | 9.87 | 9.30 | 9.59 | 9.85 | 10.86 | 9.36 | 10.49 | 9.44 |
| PEX6 | 2 | 9.09 | 11.44 | 8.04 | 8.56 | 9.95 | 11.69 | 10.01 | 9.98 | 10.75 | 10.49 | 10.26 | 8.30 | 10.82 | 10.00 | 10.62 |
| PFAS | 40 | 10.24 | 10.91 | 11.77 | 10.61 | 8.88 | 9.05 | 9.01 | 8.99 | 10.62 | 9.93 | 10.57 | 11.19 | 8.97 | 9.00 | 10.27 |
| PFDN1 | 8 | 10.22 | 10.08 | 9.09 | 11.10 | 11.30 | 11.13 | 9.53 | 6.97 | 10.54 | 10.05 | 10.15 | 10.09 | 11.22 | 8.25 | 10.29 |
| PFDN2 | 21 | 8.96 | 10.15 | 9.74 | 9.68 | 11.14 | 11.78 | 10.78 | 10.15 | 9.01 | 8.62 | 9.55 | 9.71 | 11.46 | 10.47 | 8.81 |
| PFDN4 | 4 | 10.15 | 11.45 | 10.36 | 9.84 | 10.69 | 11.28 | 11.84 | 5.34 | 9.93 | 9.11 | 10.80 | 10.10 | 10.99 | 8.59 | 9.52 |
| PFDN5 | 18 | 8.41 | 9.58 | 10.08 | 9.59 | 10.39 | 10.34 | 10.66 | 11.93 | 9.25 | 9.77 | 9.00 | 9.84 | 10.37 | 11.29 | 9.51 |
| PFDN6 | 18 | 9.79 | 10.01 | 8.69 | 9.05 | 10.59 | 10.99 | 11.23 | 10.91 | 9.51 | 9.24 | 9.90 | 8.87 | 10.79 | 11.07 | 9.37 |
| PFKFB2 | 5 | 9.11 | 10.00 | 8.27 | 8.40 | 12.51 | 12.83 | 10.09 | 10.26 | 9.55 | 8.99 | 9.55 | 8.34 | 12.67 | 10.17 | 9.27 |
| PFKFB3 | 1 | 10.94 | 11.34 | 10.24 | 9.01 | 10.67 | 9.32 | 9.50 | 10.87 | 9.30 | 8.79 | 11.14 | 9.63 | 10.00 | 10.19 | 9.04 |
| PFKL | 20 | 9.56 | 10.30 | 8.58 | 7.81 | 11.90 | 11.75 | 10.29 | 10.10 | 9.89 | 9.82 | 9.93 | 8.19 | 11.83 | 10.20 | 9.85 |
| PFKM | 38 | 9.85 | 10.73 | 8.57 | 8.43 | 10.15 | 10.91 | 9.50 | 10.08 | 10.73 | 11.05 | 10.29 | 8.50 | 10.53 | 9.79 | 10.89 |
| PFKP | 59 | 8.58 | 8.77 | 11.17 | 9.52 | 8.25 | 7.94 | 9.13 | 15.57 | 10.60 | 10.47 | 8.67 | 10.34 | 8.10 | 12.35 | 10.54 |
| PFN1 | 81 | 8.32 | 12.54 | 13.89 | 13.07 | 7.83 | 7.52 | 6.38 | 8.23 | 10.84 | 11.39 | 10.43 | 13.48 | 7.67 | 7.30 | 11.11 |
| PFN2 | 9 | 8.36 | 9.40 | 14.45 | 13.13 | 10.63 | 9.86 | 7.11 | 5.97 | 10.62 | 10.47 | 8.88 | 13.79 | 10.24 | 6.54 | 10.55 |
| PGAM1 | 41 | 8.58 | 9.34 | 11.81 | 8.30 | 11.32 | 10.41 | 9.79 | 10.15 | 10.15 | 10.14 | 8.96 | 10.06 | 10.87 | 9.97 | 10.15 |
| PGAM5 | 14 | 9.46 | 7.84 | 9.06 | 9.32 | 8.08 | 8.39 | 9.52 | 21.32 | 8.47 | 8.54 | 8.65 | 9.19 | 8.23 | 15.42 | 8.50 |
| PGD | 51 | 11.86 | 13.77 | 9.30 | 8.39 | 8.69 | 8.66 | 9.12 | 9.11 | 10.58 | 10.51 | 12.82 | 8.85 | 8.67 | 9.12 | 10.55 |
| PGGT1B | 6 | 8.81 | 11.46 | 8.75 | 9.06 | 9.00 | 8.71 | 11.09 | 10.69 | 11.35 | 11.09 | 10.13 | 8.91 | 8.85 | 10.89 | 11.22 |
| PGK1 | 168 | 7.67 | 9.80 | 7.54 | 7.05 | 10.97 | 11.03 | 14.03 | 17.28 | 7.52 | 7.11 | 8.74 | 7.30 | 11.00 | 15.65 | 7.32 |
| PGK2 | 1 | 7.80 | 9.57 | 7.58 | 7.87 | 10.69 | 10.06 | 15.90 | 14.64 | 8.17 | 7.72 | 8.68 | 7.72 | 10.37 | 15.27 | 7.95 |
| PGLS | 10 | 9.48 | 10.68 | 9.25 | 8.51 | 9.49 | 8.86 | 12.90 | 12.21 | 9.54 | 9.08 | 10.08 | 8.88 | 9.18 | 12.56 | 9.31 |

| | | | | | | | | | | | | | | | | |
|---|---|---|---|---|---|---|---|---|---|---|---|---|---|---|---|---|
| PGM1 | 37 | 11.26 | 11.86 | 10.44 | 9.46 | 9.23 | 9.02 | 9.92 | 8.80 | 10.19 | 9.81 | 11.56 | 9.95 | 9.13 | 9.36 | 10.00 |
| PGM2 | 22 | 9.62 | 9.91 | 10.00 | 9.58 | 11.43 | 11.57 | 10.19 | 10.42 | 8.91 | 8.37 | 9.77 | 9.79 | 11.50 | 10.30 | 8.64 |
| PGM2L1 | 12 | 9.79 | 10.33 | 8.20 | 8.52 | 10.21 | 11.28 | 12.72 | 13.19 | 7.93 | 7.84 | 10.06 | 8.36 | 10.74 | 12.95 | 7.88 |
| PGM3 | 17 | 9.98 | 9.81 | 8.67 | 7.72 | 8.94 | 8.77 | 8.93 | 12.79 | 12.55 | 11.86 | 9.89 | 8.19 | 8.85 | 10.86 | 12.21 |
| PGP | 7 | 10.41 | 10.57 | 10.21 | 9.92 | 11.34 | 11.32 | 8.75 | 8.00 | 10.01 | 9.49 | 10.49 | 10.06 | 11.33 | 8.37 | 9.75 |
| PGRMC1 | 8 | 12.07 | 6.32 | 8.21 | 12.83 | 9.79 | 8.32 | 13.97 | 10.33 | 8.02 | 10.14 | 9.19 | 10.52 | 9.06 | 12.15 | 9.08 |
| PGRMC2 | 14 | 10.82 | 6.42 | 8.45 | 12.88 | 9.25 | 8.04 | 8.77 | 9.53 | 11.21 | 14.63 | 8.62 | 10.67 | 8.64 | 9.15 | 12.92 |
| PGS1 | 3 | 10.75 | 10.14 | 10.31 | 10.95 | 8.72 | 8.54 | 9.61 | 8.77 | 10.55 | 11.66 | 10.44 | 10.63 | 8.63 | 9.19 | 11.10 |
| PHACTR2 | 1 | 9.10 | 11.15 | 10.23 | 10.77 | 11.05 | 11.95 | 8.41 | 8.23 | 9.47 | 9.64 | 10.12 | 10.50 | 11.50 | 8.32 | 9.56 |
| PHACTR4 | 6 | 11.13 | 11.41 | 9.15 | 8.33 | 8.90 | 8.91 | 10.68 | 9.37 | 11.52 | 10.60 | 11.27 | 8.74 | 8.91 | 10.03 | 11.06 |
| PHAX | 15 | 9.40 | 10.43 | 10.71 | 9.82 | 9.27 | 9.58 | 10.74 | 9.76 | 10.47 | 9.82 | 9.92 | 10.27 | 9.42 | 10.25 | 10.14 |
| PHB | 36 | 9.00 | 8.75 | 10.52 | 10.66 | 9.71 | 10.05 | 10.29 | 12.82 | 9.02 | 9.17 | 8.88 | 10.59 | 9.88 | 11.56 | 9.10 |
| PHB2 | 42 | 9.62 | 8.99 | 11.34 | 9.88 | 10.26 | 10.48 | 11.36 | 9.26 | 9.41 | 9.40 | 9.30 | 10.61 | 10.37 | 10.31 | 9.40 |
| PHC1 | 1 | 10.38 | 11.20 | 11.05 | 9.86 | 7.53 | 8.75 | 12.36 | 12.50 | 8.32 | 8.04 | 10.79 | 10.45 | 8.14 | 12.43 | 8.18 |
| PHC2 | 3 | 7.86 | 8.05 | 10.87 | 10.28 | 8.62 | 8.94 | 14.89 | 13.13 | 8.66 | 8.71 | 7.96 | 10.57 | 8.78 | 14.01 | 8.68 |
| PHC3 | 2 | 7.73 | 9.84 | 13.61 | 11.64 | 10.19 | 10.51 | 10.13 | 8.85 | 8.43 | 9.07 | 8.78 | 12.62 | 10.35 | 9.49 | 8.75 |
| PHF10 | 3 | 9.68 | 10.49 | 8.95 | 7.81 | 10.02 | 12.13 | 14.17 | 11.14 | 8.29 | 7.31 | 10.09 | 8.38 | 11.08 | 12.65 | 7.80 |
| PHF12 | 1 | 11.69 | 9.67 | 9.32 | 8.27 | 7.85 | 9.51 | 8.96 | 13.99 | 10.82 | 9.92 | 10.68 | 8.79 | 8.68 | 11.47 | 10.37 |
| PHF14 | 8 | 9.20 | 9.83 | 11.68 | 10.48 | 8.48 | 8.15 | 12.26 | 10.13 | 9.94 | 9.86 | 9.52 | 11.08 | 8.31 | 11.19 | 9.90 |
| PHF2 | 2 | 9.35 | 9.02 | 7.95 | 8.20 | 9.79 | 11.70 | 13.42 | 11.96 | 8.51 | 10.10 | 9.18 | 8.07 | 10.75 | 12.69 | 9.31 |
| PHF20L1 | 1 | 10.23 | 10.03 | 9.55 | 9.21 | 9.71 | 9.47 | 12.74 | 9.66 | 10.05 | 9.35 | 10.13 | 9.38 | 9.59 | 11.20 | 9.70 |
| PHF21A | 3 | 8.74 | 9.60 | 10.31 | 9.62 | 11.10 | 11.39 | 12.02 | 10.08 | 8.66 | 8.49 | 9.17 | 9.96 | 11.24 | 11.05 | 8.57 |
| PHF23 | 2 | 9.82 | 10.99 | 10.87 | 12.16 | 8.46 | 9.11 | 9.28 | 9.44 | 10.20 | 9.67 | 10.40 | 11.52 | 8.79 | 9.36 | 9.94 |
| PHF3 | 22 | 8.86 | 10.03 | 9.92 | 9.38 | 10.83 | 10.90 | 11.79 | 10.58 | 9.26 | 8.46 | 9.44 | 9.65 | 10.86 | 11.18 | 8.86 |
| PHF5A | 5 | 8.99 | 10.40 | 10.41 | 10.21 | 8.81 | 9.16 | 9.75 | 13.66 | 9.29 | 9.34 | 9.69 | 10.31 | 8.98 | 11.70 | 9.31 |
| PHF6 | 10 | 8.74 | 9.81 | 9.06 | 8.82 | 10.92 | 11.55 | 12.72 | 14.01 | 7.11 | 7.26 | 9.28 | 8.94 | 11.24 | 13.37 | 7.18 |
| PHF8 | 3 | 9.05 | 8.94 | 7.38 | 7.32 | 10.54 | 11.67 | 15.27 | 14.33 | 8.25 | 7.24 | 9.00 | 7.35 | 11.10 | 14.80 | 7.75 |
| PHGDH | 60 | 9.60 | 10.85 | 9.54 | 8.82 | 8.67 | 8.27 | 10.39 | 14.56 | 9.76 | 9.54 | 10.23 | 9.18 | 8.47 | 12.47 | 9.65 |
| PHIP | 18 | 10.31 | 10.44 | 8.25 | 7.95 | 12.37 | 12.13 | 10.36 | 10.69 | 8.66 | 8.84 | 10.37 | 8.10 | 12.25 | 10.53 | 8.75 |
| PHKA1 | 3 | 9.02 | 10.59 | 9.45 | 9.58 | 11.43 | 9.56 | 11.66 | 10.34 | 8.65 | 9.72 | 9.81 | 9.51 | 10.50 | 11.00 | 9.18 |
| PHKB | 8 | 10.97 | 11.44 | 7.45 | 8.87 | 10.90 | 11.90 | 9.29 | 9.91 | 9.68 | 9.59 | 11.21 | 8.16 | 11.40 | 9.60 | 9.63 |
| PHKG2 | 2 | 10.73 | 9.53 | 9.47 | 9.87 | 11.47 | 12.61 | 7.17 | 7.40 | 10.95 | 10.80 | 10.13 | 9.67 | 12.04 | 7.28 | 10.88 |
| PHLDA2 | 4 | 8.55 | 10.78 | 12.09 | 11.14 | 10.65 | 10.84 | 8.17 | 5.83 | 11.15 | 10.81 | 9.66 | 11.62 | 10.74 | 7.00 | 10.98 |
| PHLDB1 | 15 | 9.39 | 11.08 | 15.40 | 15.31 | 7.92 | 8.21 | 4.75 | 4.70 | 11.95 | 11.29 | 10.24 | 15.35 | 8.06 | 4.72 | 11.62 |
| PHLDB2 | 11 | 8.80 | 9.92 | 15.04 | 14.59 | 7.40 | 7.61 | 9.06 | 8.07 | 9.94 | 9.56 | 9.36 | 14.82 | 7.50 | 8.57 | 9.75 |
| PHPT1 | 8 | 9.65 | 10.32 | 10.93 | 9.68 | 10.91 | 10.56 | 8.35 | 6.31 | 12.18 | 11.11 | 9.98 | 10.31 | 10.74 | 7.33 | 11.64 |
| PHRF1 | 4 | 8.64 | 10.49 | 9.60 | 10.02 | 9.55 | 11.01 | 10.65 | 10.09 | 9.78 | 10.16 | 9.57 | 9.81 | 10.28 | 10.37 | 9.97 |
| PHYH | 2 | 7.93 | 8.68 | 5.12 | 5.71 | 21.22 | 22.04 | 7.47 | 6.73 | 7.73 | 7.38 | 8.31 | 5.41 | 21.63 | 7.10 | 7.55 |
| PI4K2A | 3 | 9.11 | 7.26 | 9.27 | 10.86 | 10.69 | 10.23 | 11.82 | 11.49 | 9.36 | 9.90 | 8.19 | 10.07 | 10.46 | 11.66 | 9.63 |
| PI4KA | 10 | 11.09 | 10.81 | 8.07 | 8.98 | 9.14 | 9.80 | 10.60 | 9.85 | 11.25 | 10.41 | 10.95 | 8.53 | 9.47 | 10.23 | 10.83 |
| PI4KB | 6 | 9.29 | 10.68 | 11.25 | 9.27 | 12.44 | 12.51 | 9.17 | 7.57 | 9.24 | 8.59 | 9.98 | 10.26 | 12.48 | 8.37 | 8.91 |
| PIAS1 | 2 | 10.01 | 10.53 | 9.49 | 8.79 | 9.57 | 10.95 | 10.29 | 12.21 | 9.03 | 9.13 | 10.27 | 9.14 | 10.26 | 11.25 | 9.08 |

| | | | | | | | | | | | | | | | | |
|---|---|---|---|---|---|---|---|---|---|---|---|---|---|---|---|---|
| PIBF1 | 7 | 11.48 | 11.99 | 10.06 | 9.99 | 10.02 | 10.71 | 10.77 | 10.24 | 7.63 | 7.10 | 11.74 | 10.03 | 10.36 | 10.51 | 7.37 |
| PICALM | 18 | 9.03 | 9.10 | 11.79 | 11.49 | 10.59 | 10.89 | 8.73 | 8.00 | 10.48 | 9.89 | 9.07 | 11.64 | 10.74 | 8.37 | 10.19 |
| PIGG | 1 | 9.56 | 12.08 | 10.38 | 10.45 | 9.48 | 11.21 | 7.32 | 8.88 | 10.82 | 9.82 | 10.82 | 10.42 | 10.34 | 8.10 | 10.32 |
| PIGK | 4 | 12.41 | 7.50 | 7.55 | 8.70 | 11.71 | 9.78 | 10.10 | 10.26 | 10.76 | 11.24 | 9.95 | 8.13 | 10.74 | 10.18 | 11.00 |
| PIGS | 9 | 11.68 | 9.77 | 7.99 | 9.14 | 11.74 | 10.45 | 8.90 | 9.14 | 9.93 | 11.25 | 10.72 | 8.56 | 11.10 | 9.02 | 10.59 |
| PIGT | 4 | 11.95 | 7.77 | 6.86 | 9.70 | 11.69 | 11.31 | 10.02 | 9.18 | 10.34 | 11.17 | 9.86 | 8.28 | 11.50 | 9.60 | 10.76 |
| PIH1D1 | 8 | 9.40 | 10.41 | 11.43 | 10.29 | 9.71 | 9.82 | 10.35 | 10.86 | 8.99 | 8.75 | 9.90 | 10.86 | 9.77 | 10.61 | 8.87 |
| PIK3C2A | 18 | 9.29 | 9.89 | 10.63 | 10.82 | 10.14 | 10.63 | 9.79 | 9.17 | 10.08 | 9.56 | 9.59 | 10.73 | 10.39 | 9.48 | 9.82 |
| PIK3C3 | 3 | 9.31 | 10.83 | 10.24 | 10.32 | 9.67 | 10.77 | 8.76 | 9.01 | 10.59 | 10.49 | 10.07 | 10.28 | 10.22 | 8.89 | 10.54 |
| PIK3CB | 1 | 10.75 | 9.76 | 8.66 | 11.51 | 9.71 | 10.90 | 10.02 | 10.80 | 8.22 | 9.67 | 10.25 | 10.08 | 10.31 | 10.41 | 8.94 |
| PIK3R1 | 2 | 8.74 | 9.23 | 14.11 | 14.30 | 8.61 | 8.78 | 9.47 | 8.45 | 8.83 | 9.48 | 8.99 | 14.20 | 8.69 | 8.96 | 9.16 |
| PIK3R4 | 10 | 10.00 | 9.90 | 10.24 | 10.45 | 10.38 | 9.32 | 10.40 | 9.75 | 9.80 | 9.75 | 9.95 | 10.35 | 9.85 | 10.08 | 9.77 |
| PIKFYVE | 3 | 10.24 | 11.19 | 7.82 | 8.77 | 10.52 | 11.56 | 10.55 | 10.25 | 9.95 | 9.15 | 10.72 | 8.29 | 11.04 | 10.40 | 9.55 |
| PIN1 | 8 | 8.07 | 7.23 | 9.03 | 6.24 | 11.48 | 9.55 | 7.35 | 27.67 | 7.10 | 6.29 | 7.65 | 7.63 | 10.52 | 17.51 | 6.69 |
| PIN4 | 10 | 9.34 | 10.51 | 8.59 | 8.14 | 12.90 | 12.89 | 12.50 | 11.55 | 6.91 | 6.67 | 9.93 | 8.36 | 12.90 | 12.03 | 6.79 |
| PINX1 | 6 | 10.29 | 9.87 | 11.26 | 10.33 | 8.71 | 7.67 | 15.28 | 12.45 | 7.13 | 7.02 | 10.08 | 10.79 | 8.19 | 13.86 | 7.08 |
| PIP4K2A | 5 | 7.61 | 10.13 | 11.67 | 10.88 | 10.74 | 10.86 | 9.62 | 9.13 | 10.10 | 9.25 | 8.87 | 11.28 | 10.80 | 9.38 | 9.68 |
| PIP4K2B | 9 | 9.30 | 10.61 | 12.19 | 11.69 | 9.82 | 10.59 | 8.93 | 8.32 | 9.07 | 9.47 | 9.95 | 11.94 | 10.21 | 8.63 | 9.27 |
| PIP4K2C | 11 | 9.12 | 11.34 | 10.60 | 10.01 | 11.66 | 11.92 | 8.89 | 8.70 | 9.21 | 8.54 | 10.23 | 10.31 | 11.79 | 8.80 | 8.88 |
| PIP5K1A | 5 | 10.37 | 10.13 | 10.29 | 7.75 | 9.68 | 9.35 | 6.97 | 12.16 | 12.19 | 11.12 | 10.25 | 9.02 | 9.51 | 9.57 | 11.66 |
| PIP5K1C | 5 | 10.39 | 10.78 | 11.16 | 9.72 | 10.10 | 10.70 | 7.57 | 7.29 | 11.30 | 10.99 | 10.58 | 10.44 | 10.40 | 7.43 | 11.14 |
| PIR | 21 | 13.05 | 13.94 | 7.88 | 7.01 | 13.76 | 13.46 | 8.05 | 7.08 | 8.15 | 7.61 | 13.50 | 7.45 | 13.61 | 7.56 | 7.88 |
| PITHD1 | 5 | 11.79 | 13.11 | 8.22 | 7.48 | 9.58 | 10.21 | 9.48 | 8.10 | 11.19 | 10.85 | 12.45 | 7.85 | 9.89 | 8.79 | 11.02 |
| PITPNA | 12 | 11.56 | 12.26 | 10.19 | 9.61 | 9.02 | 9.57 | 8.20 | 8.41 | 10.60 | 10.58 | 11.91 | 9.90 | 9.29 | 8.30 | 10.59 |
| PITPNB | 20 | 10.54 | 10.85 | 11.36 | 10.19 | 7.51 | 7.81 | 9.71 | 9.14 | 11.71 | 11.19 | 10.69 | 10.77 | 7.66 | 9.42 | 11.45 |
| PITPNC1 | 3 | 7.21 | 7.33 | 11.40 | 10.05 | 8.70 | 9.26 | 12.82 | 13.65 | 9.95 | 9.64 | 7.27 | 10.72 | 8.98 | 13.24 | 9.79 |
| PITPNM1 | 4 | 8.33 | 8.20 | 5.88 | 6.49 | 17.14 | 19.09 | 10.45 | 10.76 | 7.59 | 6.08 | 8.26 | 6.18 | 18.12 | 10.60 | 6.83 |
| PITRM1 | 28 | 10.98 | 8.83 | 8.81 | 10.08 | 12.03 | 11.14 | 10.26 | 9.66 | 8.86 | 9.36 | 9.90 | 9.44 | 11.59 | 9.96 | 9.11 |
| PITX1 | 2 | 11.46 | 13.55 | 4.55 | 4.60 | 11.17 | 10.59 | 14.49 | 14.97 | 7.05 | 7.57 | 12.50 | 4.57 | 10.88 | 14.73 | 7.31 |
| PJA2 | 3 | 9.95 | 11.39 | 8.29 | 8.15 | 10.69 | 12.21 | 11.88 | 12.25 | 7.51 | 7.68 | 10.67 | 8.22 | 11.45 | 12.06 | 7.59 |
| PKM | 289 | 8.52 | 10.84 | 9.36 | 9.21 | 9.95 | 10.05 | 7.55 | 12.17 | 11.31 | 11.05 | 9.68 | 9.28 | 10.00 | 9.86 | 11.18 |
| PKMYT1 | 1 | 10.30 | 6.90 | 10.08 | 11.64 | 7.13 | 3.94 | 10.56 | 10.69 | 13.65 | 15.11 | 8.60 | 10.86 | 5.53 | 10.63 | 14.38 |
| PKN1 | 12 | 10.32 | 11.31 | 8.66 | 8.86 | 11.21 | 11.33 | 9.17 | 9.64 | 9.95 | 9.53 | 10.82 | 8.76 | 11.27 | 9.41 | 9.74 |
| PKN2 | 22 | 10.09 | 10.33 | 9.20 | 8.68 | 8.78 | 8.76 | 10.70 | 13.96 | 9.88 | 9.61 | 10.21 | 8.94 | 8.77 | 12.33 | 9.74 |
| PKP3 | 11 | 10.70 | 12.05 | 6.69 | 6.95 | 12.16 | 13.02 | 9.77 | 9.32 | 10.16 | 9.19 | 11.38 | 6.82 | 12.59 | 9.54 | 9.67 |
| PKP4 | 3 | 9.09 | 10.79 | 12.10 | 12.25 | 8.33 | 9.04 | 6.70 | 6.00 | 12.68 | 13.04 | 9.94 | 12.17 | 8.68 | 6.35 | 12.86 |
| PLA2G16 | 1 | 8.37 | 9.71 | 13.86 | 11.65 | 12.57 | 12.54 | 7.35 | 2.78 | 8.23 | 12.94 | 9.04 | 12.75 | 12.55 | 5.07 | 10.58 |
| PLA2G4A | 35 | 6.66 | 7.18 | 5.99 | 5.39 | 12.65 | 12.33 | 16.48 | 19.26 | 7.18 | 6.89 | 6.92 | 5.69 | 12.49 | 17.87 | 7.03 |
| PLA2G4B | 1 | 10.21 | 10.58 | 15.23 | 15.40 | 6.29 | 7.12 | 5.09 | 6.34 | 11.84 | 11.90 | 10.39 | 15.32 | 6.71 | 5.71 | 11.87 |
| PLAA | 34 | 9.63 | 9.47 | 9.48 | 9.77 | 10.43 | 10.61 | 8.83 | 9.71 | 11.17 | 10.92 | 9.55 | 9.62 | 10.52 | 9.27 | 11.04 |
| PLBD2 | 2 | 10.51 | 11.24 | 12.07 | 11.64 | 11.39 | 11.52 | 8.97 | 8.18 | 6.98 | 7.49 | 10.88 | 11.86 | 11.45 | 8.58 | 7.23 |
| PLCB1 | 1 | 8.66 | 9.48 | 11.90 | 10.11 | 9.39 | 10.18 | 10.85 | 11.00 | 9.78 | 8.65 | 9.07 | 11.01 | 9.78 | 10.93 | 9.22 |

| | | | | | | | | | | | | | | | | |
|---|---|---|---|---|---|---|---|---|---|---|---|---|---|---|---|---|
| PLCB3 | 24 | 9.25 | 10.63 | 11.28 | 11.40 | 9.28 | 9.92 | 7.61 | 7.46 | 11.80 | 11.38 | 9.94 | 11.34 | 9.60 | 7.54 | 11.59 |
| PLCB4 | 1 | 10.75 | 9.44 | 14.77 | 14.31 | 8.09 | 5.14 | 6.11 | 3.92 | 13.87 | 13.61 | 10.10 | 14.54 | 6.61 | 5.01 | 13.74 |
| PLCD3 | 3 | 11.31 | 10.47 | 6.83 | 8.75 | 9.46 | 11.28 | 8.86 | 8.65 | 11.24 | 13.14 | 10.89 | 7.79 | 10.37 | 8.76 | 12.19 |
| PLCG1 | 21 | 10.28 | 10.70 | 10.43 | 10.04 | 9.71 | 10.00 | 10.07 | 9.41 | 9.96 | 9.40 | 10.49 | 10.23 | 9.86 | 9.74 | 9.68 |
| PLCH1 | 2 | 11.33 | 11.07 | 8.64 | 10.75 | 10.00 | 10.46 | 10.42 | 9.82 | 9.04 | 8.47 | 11.20 | 9.69 | 10.23 | 10.12 | 8.75 |
| PLD1 | 2 | 11.31 | 8.61 | 11.02 | 15.18 | 12.24 | 13.49 | 9.01 | 7.72 | 5.63 | 5.77 | 9.96 | 13.10 | 12.87 | 8.37 | 5.70 |
| PLD3 | 7 | 10.98 | 10.87 | 8.71 | 8.92 | 8.65 | 8.57 | 10.82 | 10.11 | 10.91 | 11.45 | 10.93 | 8.82 | 8.61 | 10.46 | 11.18 |
| PLEC | 603 | 8.91 | 9.71 | 10.17 | 9.60 | 10.55 | 10.57 | 8.84 | 9.30 | 11.42 | 10.91 | 9.31 | 9.89 | 10.56 | 9.07 | 11.17 |
| PLEK2 | 7 | 7.20 | 8.08 | 12.32 | 11.06 | 10.81 | 11.23 | 10.42 | 9.78 | 9.57 | 9.54 | 7.64 | 11.69 | 11.02 | 10.10 | 9.55 |
| PLEKHA1 | 3 | 6.09 | 7.63 | 12.14 | 11.74 | 8.61 | 9.20 | 11.44 | 12.23 | 11.33 | 9.58 | 6.86 | 11.94 | 8.91 | 11.84 | 10.45 |
| PLEKHA2 | 15 | 8.64 | 9.61 | 11.38 | 10.86 | 10.60 | 10.88 | 11.68 | 10.67 | 7.92 | 7.77 | 9.12 | 11.12 | 10.74 | 11.17 | 7.84 |
| PLEKHA5 | 26 | 7.66 | 8.52 | 9.65 | 9.09 | 11.28 | 11.34 | 14.20 | 14.03 | 7.25 | 6.98 | 8.09 | 9.37 | 11.31 | 14.12 | 7.11 |
| PLEKHF1 | 1 | 10.15 | 12.38 | 9.89 | 10.64 | 9.79 | 9.99 | 11.02 | 10.35 | 8.18 | 7.61 | 11.27 | 10.27 | 9.89 | 10.68 | 7.89 |
| PLEKHG2 | 2 | 11.05 | 11.10 | 7.29 | 6.86 | 11.04 | 12.27 | 8.82 | 11.92 | 11.14 | 8.50 | 11.08 | 7.07 | 11.66 | 10.37 | 9.82 |
| PLEKHG3 | 5 | 10.82 | 11.42 | 8.45 | 6.98 | 11.52 | 11.52 | 6.95 | 10.27 | 10.86 | 11.22 | 11.12 | 7.71 | 11.52 | 8.61 | 11.04 |
| PLEKHG4 | 1 | 11.75 | 11.01 | 7.09 | 7.24 | 11.06 | 11.09 | 9.92 | 8.91 | 11.03 | 10.90 | 11.38 | 7.17 | 11.08 | 9.42 | 10.96 |
| PLEKHG4B | 5 | 10.77 | 12.17 | 11.68 | 11.96 | 9.62 | 9.80 | 9.91 | 1.76 | 11.10 | 11.24 | 11.47 | 11.82 | 9.71 | 5.83 | 11.17 |
| PLEKHH2 | 1 | 8.70 | 9.50 | 8.75 | 9.34 | 9.98 | 10.35 | 13.39 | 13.21 | 8.62 | 8.16 | 9.10 | 9.05 | 10.17 | 13.30 | 8.39 |
| PLEKHM2 | 2 | 9.01 | 12.48 | 9.36 | 9.93 | 9.42 | 11.57 | 8.28 | 8.44 | 10.73 | 10.78 | 10.75 | 9.65 | 10.49 | 8.36 | 10.76 |
| PLEKHO2 | 3 | 8.89 | 10.00 | 13.58 | 12.65 | 7.51 | 6.94 | 12.84 | 10.92 | 8.76 | 7.90 | 9.45 | 13.12 | 7.23 | 11.88 | 8.33 |
| PLG | 1 | 0.98 | 0.77 | 0.87 | 0.38 | 2.75 | 0.52 | 5.45 | 86.59 | 0.83 | 0.85 | 0.88 | 0.62 | 1.64 | 46.02 | 0.84 |
| PLIN2 | 4 | 8.30 | 9.26 | 6.72 | 5.71 | 10.60 | 10.40 | 16.45 | 16.10 | 8.11 | 8.36 | 8.78 | 6.21 | 10.50 | 16.28 | 8.23 |
| PLIN3 | 26 | 7.71 | 8.74 | 10.44 | 9.45 | 12.62 | 12.82 | 10.90 | 10.65 | 8.47 | 8.19 | 8.22 | 9.95 | 12.72 | 10.78 | 8.33 |
| PLK1 | 14 | 8.26 | 9.08 | 10.76 | 10.59 | 9.84 | 10.06 | 9.80 | 10.31 | 10.54 | 10.78 | 8.67 | 10.67 | 9.95 | 10.05 | 10.66 |
| PLOD1 | 29 | 9.04 | 6.73 | 9.01 | 10.08 | 6.78 | 6.50 | 14.42 | 14.43 | 11.18 | 11.83 | 7.89 | 9.54 | 6.64 | 14.42 | 11.51 |
| PLOD2 | 33 | 13.85 | 9.89 | 9.90 | 10.83 | 7.67 | 6.87 | 8.42 | 7.43 | 12.06 | 13.08 | 11.87 | 10.36 | 7.27 | 7.92 | 12.57 |
| PLOD3 | 27 | 9.01 | 8.01 | 8.31 | 8.81 | 9.33 | 9.05 | 12.72 | 15.02 | 9.40 | 10.35 | 8.51 | 8.56 | 9.19 | 13.87 | 9.87 |
| PLP2 | 6 | 7.50 | 5.48 | 6.48 | 7.71 | 13.14 | 13.06 | 12.75 | 14.29 | 9.51 | 10.08 | 6.49 | 7.09 | 13.10 | 13.52 | 9.79 |
| PLRG1 | 16 | 9.10 | 10.42 | 10.85 | 9.70 | 10.22 | 10.32 | 10.99 | 9.93 | 9.16 | 9.31 | 9.76 | 10.27 | 10.27 | 10.46 | 9.23 |
| PLS1 | 10 | 11.80 | 13.47 | 9.88 | 9.71 | 13.88 | 13.74 | 4.98 | 4.20 | 9.37 | 8.96 | 12.64 | 9.80 | 13.81 | 4.59 | 9.17 |
| PLS3 | 102 | 9.08 | 10.35 | 11.18 | 9.64 | 11.27 | 11.14 | 6.17 | 10.34 | 10.61 | 10.22 | 9.71 | 10.41 | 11.20 | 8.26 | 10.41 |
| PLXDC2 | 3 | 12.05 | 8.81 | 8.02 | 8.76 | 14.97 | 12.99 | 5.61 | 5.32 | 11.36 | 12.11 | 10.43 | 8.39 | 13.98 | 5.46 | 11.74 |
| PLXNA3 | 1 | 9.64 | 8.52 | 9.75 | 13.31 | 10.15 | 10.01 | 10.32 | 10.51 | 9.19 | 8.60 | 9.08 | 11.53 | 10.08 | 10.42 | 8.89 |
| PLXNB2 | 26 | 11.41 | 10.08 | 9.75 | 10.32 | 9.84 | 9.62 | 9.82 | 8.45 | 10.28 | 10.44 | 10.75 | 10.03 | 9.73 | 9.13 | 10.36 |
| PLXND1 | 3 | 10.54 | 9.32 | 10.84 | 13.21 | 7.64 | 7.85 | 6.17 | 5.18 | 14.27 | 14.98 | 9.93 | 12.02 | 7.74 | 5.68 | 14.62 |
| PM20D2 | 4 | 8.86 | 10.46 | 9.02 | 8.75 | 9.34 | 9.88 | 12.27 | 12.14 | 9.60 | 9.68 | 9.66 | 8.89 | 9.61 | 12.21 | 9.64 |
| PMF1 | 1 | 11.79 | 14.48 | 9.28 | 8.62 | 10.12 | 12.44 | 8.46 | 8.84 | 7.85 | 8.13 | 13.13 | 8.95 | 11.28 | 8.65 | 7.99 |
| PML | 10 | 8.45 | 9.53 | 10.88 | 10.53 | 9.37 | 9.90 | 8.54 | 9.33 | 12.27 | 11.20 | 8.99 | 10.70 | 9.63 | 8.94 | 11.74 |
| PMM1 | 6 | 9.04 | 10.00 | 12.19 | 12.80 | 9.22 | 9.47 | 9.43 | 9.25 | 9.45 | 9.15 | 9.52 | 12.50 | 9.35 | 9.34 | 9.30 |
| PMM2 | 16 | 9.20 | 9.96 | 13.90 | 11.82 | 9.90 | 9.82 | 9.33 | 7.85 | 9.39 | 8.83 | 9.58 | 12.86 | 9.86 | 8.59 | 9.11 |
| PMPCA | 20 | 11.29 | 9.69 | 9.85 | 11.42 | 9.09 | 8.53 | 9.43 | 8.51 | 10.60 | 11.60 | 10.49 | 10.63 | 8.81 | 8.97 | 11.10 |
| PMPCB | 14 | 11.91 | 9.77 | 9.28 | 10.92 | 9.37 | 8.75 | 9.69 | 11.02 | 9.52 | 9.77 | 10.84 | 10.10 | 9.06 | 10.35 | 9.64 |

| | | | | | | | | | | | | | | | | |
|---|---|---|---|---|---|---|---|---|---|---|---|---|---|---|---|---|
| PMS1 | 5 | 9.39 | 9.47 | 10.47 | 9.83 | 9.75 | 9.92 | 11.52 | 9.85 | 9.81 | 10.00 | 9.43 | 10.15 | 9.84 | 10.68 | 9.90 |
| PMS2 | 8 | 9.51 | 10.90 | 9.84 | 9.29 | 10.79 | 10.64 | 9.51 | 9.19 | 10.25 | 10.09 | 10.20 | 9.56 | 10.72 | 9.35 | 10.17 |
| PMVK | 8 | 9.42 | 9.80 | 9.42 | 8.86 | 13.61 | 13.92 | 9.47 | 8.32 | 8.81 | 8.37 | 9.61 | 9.14 | 13.76 | 8.89 | 8.59 |
| PNISR | 11 | 9.16 | 10.20 | 9.69 | 9.54 | 9.37 | 9.75 | 12.14 | 11.30 | 9.81 | 9.05 | 9.68 | 9.61 | 9.56 | 11.72 | 9.43 |
| PNKD | 1 | 13.65 | 6.87 | 8.46 | 11.88 | 8.19 | 7.98 | 7.72 | 7.29 | 12.32 | 15.63 | 10.26 | 10.17 | 8.09 | 7.51 | 13.97 |
| PNKP | 12 | 8.75 | 10.20 | 10.85 | 9.53 | 8.88 | 9.16 | 11.85 | 11.16 | 10.46 | 9.17 | 9.47 | 10.19 | 9.02 | 11.50 | 9.82 |
| PNMA3 | 1 | 4.65 | 0.86 | 3.70 | 0.04 | 7.58 | 2.88 | 5.18 | 67.02 | 4.85 | 3.24 | 2.75 | 1.87 | 5.23 | 36.10 | 4.05 |
| PNN | 47 | 8.73 | 9.69 | 11.12 | 9.93 | 10.39 | 10.15 | 10.87 | 10.35 | 9.41 | 9.35 | 9.21 | 10.53 | 10.27 | 10.61 | 9.38 |
| PNO1 | 9 | 8.18 | 8.32 | 10.38 | 9.71 | 8.94 | 8.01 | 12.18 | 16.88 | 8.22 | 9.18 | 8.25 | 10.04 | 8.47 | 14.53 | 8.70 |
| PNP | 37 | 9.37 | 10.03 | 9.22 | 7.88 | 11.97 | 11.75 | 9.60 | 11.50 | 9.48 | 9.18 | 9.70 | 8.55 | 11.86 | 10.55 | 9.33 |
| PNPLA6 | 2 | 11.97 | 6.25 | 9.16 | 11.66 | 10.61 | 9.35 | 9.74 | 8.20 | 10.83 | 12.24 | 9.11 | 10.41 | 9.98 | 8.97 | 11.53 |
| PNPLA8 | 2 | 10.14 | 10.43 | 8.48 | 9.22 | 9.30 | 10.00 | 12.14 | 10.35 | 10.34 | 9.60 | 10.28 | 8.85 | 9.65 | 11.24 | 9.97 |
| PNPO | 5 | 9.61 | 10.73 | 11.80 | 10.74 | 11.47 | 11.18 | 8.39 | 7.18 | 9.99 | 8.91 | 10.17 | 11.27 | 11.32 | 7.78 | 9.45 |
| PNPT1 | 30 | 9.92 | 8.65 | 10.92 | 12.71 | 8.65 | 8.23 | 10.57 | 9.57 | 10.13 | 10.65 | 9.28 | 11.82 | 8.44 | 10.07 | 10.39 |
| POC1A | 3 | 10.40 | 11.67 | 9.50 | 9.25 | 11.48 | 12.98 | 8.96 | 8.18 | 8.60 | 8.96 | 11.04 | 9.37 | 12.23 | 8.57 | 8.78 |
| PODXL | 7 | 12.37 | 8.64 | 9.12 | 11.37 | 10.22 | 9.07 | 5.19 | 7.84 | 12.71 | 13.46 | 10.50 | 10.24 | 9.64 | 6.52 | 13.09 |
| PODXL2 | 2 | 12.64 | 9.31 | 8.55 | 13.53 | 6.36 | 6.60 | 15.67 | 10.92 | 8.29 | 8.14 | 10.98 | 11.04 | 6.48 | 13.30 | 8.21 |
| POFUT1 | 13 | 9.26 | 8.72 | 9.14 | 9.03 | 12.55 | 12.34 | 11.06 | 10.43 | 8.64 | 8.83 | 8.99 | 9.08 | 12.45 | 10.74 | 8.73 |
| POFUT2 | 1 | 11.64 | 9.28 | 12.19 | 13.53 | 10.00 | 10.85 | 9.15 | 8.53 | 7.77 | 7.06 | 10.46 | 12.86 | 10.43 | 8.84 | 7.41 |
| POGK | 3 | 9.71 | 8.98 | 9.22 | 9.08 | 11.44 | 11.54 | 11.44 | 10.60 | 8.83 | 9.17 | 9.34 | 9.15 | 11.49 | 11.02 | 9.00 |
| POGLUT1 | 3 | 8.02 | 6.89 | 7.89 | 5.90 | 17.16 | 14.78 | 11.45 | 8.56 | 9.34 | 10.01 | 7.45 | 6.89 | 15.97 | 10.01 | 9.68 |
| POGZ | 12 | 9.39 | 10.66 | 10.88 | 10.59 | 11.05 | 11.12 | 9.83 | 9.32 | 8.73 | 8.43 | 10.02 | 10.73 | 11.08 | 9.58 | 8.58 |
| POLA1 | 27 | 10.91 | 10.53 | 8.38 | 7.84 | 10.01 | 10.38 | 12.29 | 12.47 | 8.73 | 8.45 | 10.72 | 8.11 | 10.20 | 12.38 | 8.59 |
| POLA2 | 11 | 10.37 | 10.86 | 9.14 | 8.53 | 9.71 | 9.89 | 11.75 | 12.89 | 8.95 | 7.91 | 10.61 | 8.84 | 9.80 | 12.32 | 8.43 |
| POLB | 3 | 8.96 | 10.19 | 8.06 | 8.04 | 9.46 | 10.27 | 14.02 | 11.92 | 9.93 | 9.14 | 9.58 | 8.05 | 9.87 | 12.97 | 9.54 |
| POLD1 | 23 | 9.11 | 9.67 | 8.45 | 8.72 | 9.36 | 9.10 | 12.19 | 12.95 | 10.20 | 10.25 | 9.39 | 8.59 | 9.23 | 12.57 | 10.22 |
| POLD2 | 10 | 10.06 | 10.18 | 9.49 | 7.65 | 9.58 | 9.54 | 12.36 | 10.67 | 10.50 | 9.98 | 10.12 | 8.57 | 9.56 | 11.52 | 10.24 |
| POLD3 | 8 | 9.95 | 10.72 | 9.49 | 8.42 | 9.28 | 9.24 | 12.12 | 10.60 | 10.59 | 9.60 | 10.33 | 8.95 | 9.26 | 11.36 | 10.09 |
| POLD4 | 1 | 9.76 | 14.05 | 8.85 | 8.15 | 10.31 | 11.35 | 8.27 | 5.57 | 12.91 | 10.79 | 11.90 | 8.50 | 10.83 | 6.92 | 11.85 |
| POLDIP2 | 10 | 11.05 | 9.60 | 9.70 | 10.39 | 9.20 | 9.03 | 9.94 | 8.75 | 11.19 | 11.16 | 10.32 | 10.05 | 9.11 | 9.34 | 11.17 |
| POLDIP3 | 26 | 10.17 | 10.82 | 12.47 | 11.64 | 8.05 | 8.27 | 8.97 | 9.19 | 10.27 | 10.15 | 10.50 | 12.05 | 8.16 | 9.08 | 10.21 |
| POLE | 11 | 9.78 | 9.52 | 8.17 | 9.24 | 9.12 | 9.25 | 11.15 | 10.93 | 11.76 | 11.08 | 9.65 | 8.71 | 9.18 | 11.04 | 11.42 |
| POLE2 | 3 | 9.74 | 11.05 | 8.05 | 8.09 | 9.51 | 10.07 | 10.74 | 10.30 | 10.91 | 11.54 | 10.40 | 8.07 | 9.79 | 10.52 | 11.22 |
| POLE3 | 18 | 9.34 | 10.11 | 11.37 | 11.31 | 8.83 | 8.87 | 9.77 | 9.45 | 10.63 | 10.32 | 9.73 | 11.34 | 8.85 | 9.61 | 10.48 |
| POLE4 | 4 | 8.72 | 10.02 | 10.42 | 9.91 | 10.15 | 9.38 | 9.57 | 7.69 | 12.93 | 11.22 | 9.37 | 10.16 | 9.76 | 8.63 | 12.07 |
| POLG | 2 | 9.81 | 9.65 | 9.18 | 10.01 | 8.68 | 8.88 | 9.99 | 11.59 | 10.76 | 11.44 | 9.73 | 9.59 | 8.78 | 10.79 | 11.10 |
| POLG2 | 2 | 8.52 | 8.92 | 11.46 | 10.98 | 9.44 | 8.86 | 10.99 | 9.58 | 10.44 | 10.80 | 8.72 | 11.22 | 9.15 | 10.28 | 10.62 |
| POLR1A | 35 | 8.93 | 9.15 | 11.50 | 11.64 | 8.97 | 8.87 | 12.04 | 11.56 | 8.75 | 8.60 | 9.04 | 11.57 | 8.92 | 11.80 | 8.67 |
| POLR1B | 11 | 9.31 | 9.39 | 10.97 | 11.34 | 8.39 | 9.10 | 12.17 | 11.57 | 8.95 | 8.80 | 9.35 | 11.16 | 8.74 | 11.87 | 8.87 |
| POLR1C | 14 | 9.38 | 9.83 | 11.30 | 10.43 | 9.18 | 9.60 | 9.64 | 13.09 | 9.04 | 8.51 | 9.61 | 10.86 | 9.39 | 11.37 | 8.78 |
| POLR1D | 2 | 10.67 | 9.63 | 12.08 | 11.68 | 10.33 | 9.99 | 9.58 | 7.73 | 9.80 | 8.51 | 10.15 | 11.88 | 10.16 | 8.66 | 9.15 |
| POLR1E | 15 | 8.31 | 8.84 | 11.42 | 11.15 | 8.41 | 8.53 | 11.67 | 11.37 | 10.09 | 10.21 | 8.57 | 11.29 | 8.47 | 11.52 | 10.15 |

| | | | | | | | | | | | | | | | | |
|---|---|---|---|---|---|---|---|---|---|---|---|---|---|---|---|---|
| POLR2A | 55 | 9.38 | 10.05 | 10.37 | 9.55 | 9.78 | 10.11 | 11.71 | 10.56 | 9.37 | 9.13 | 9.71 | 9.96 | 9.94 | 11.13 | 9.25 |
| POLR2B | 28 | 9.30 | 10.20 | 9.70 | 9.50 | 9.99 | 10.11 | 10.64 | 10.58 | 10.13 | 9.84 | 9.75 | 9.60 | 10.05 | 10.61 | 9.99 |
| POLR2C | 13 | 9.36 | 11.10 | 10.34 | 9.71 | 9.67 | 10.70 | 9.89 | 9.22 | 9.79 | 10.22 | 10.23 | 10.02 | 10.19 | 9.55 | 10.01 |
| POLR2D | 6 | 11.46 | 11.59 | 10.64 | 10.00 | 10.08 | 10.14 | 10.36 | 5.39 | 10.23 | 10.11 | 11.52 | 10.32 | 10.11 | 7.87 | 10.17 |
| POLR2E | 6 | 8.73 | 9.94 | 11.11 | 11.69 | 8.72 | 8.78 | 11.51 | 11.52 | 9.24 | 8.76 | 9.33 | 11.40 | 8.75 | 11.52 | 9.00 |
| POLR2F | 2 | 12.04 | 9.21 | 9.84 | 12.01 | 9.77 | 13.14 | 11.66 | 3.57 | 8.29 | 10.47 | 10.62 | 10.92 | 11.46 | 7.62 | 9.38 |
| POLR2G | 4 | 9.55 | 9.98 | 11.13 | 9.99 | 10.26 | 10.17 | 10.39 | 9.70 | 9.39 | 9.44 | 9.76 | 10.56 | 10.21 | 10.05 | 9.41 |
| POLR2H | 4 | 9.70 | 10.72 | 11.47 | 10.56 | 9.51 | 10.07 | 10.47 | 9.36 | 8.83 | 9.31 | 10.21 | 11.01 | 9.79 | 9.92 | 9.07 |
| POLR2I | 2 | 7.56 | 8.91 | 12.88 | 11.48 | 11.53 | 9.82 | 11.91 | 6.80 | 10.00 | 9.10 | 8.24 | 12.18 | 10.67 | 9.36 | 9.55 |
| POLR2J | 3 | 10.30 | 9.88 | 10.47 | 11.02 | 9.87 | 9.92 | 10.34 | 7.98 | 10.10 | 10.12 | 10.09 | 10.75 | 9.89 | 9.16 | 10.11 |
| POLR2L | 5 | 9.11 | 9.88 | 10.64 | 10.07 | 9.85 | 10.35 | 10.41 | 10.71 | 9.12 | 9.84 | 9.50 | 10.36 | 10.10 | 10.56 | 9.48 |
| POLR2M | 2 | 8.80 | 10.12 | 15.66 | 13.41 | 8.22 | 7.86 | 8.53 | 8.92 | 10.14 | 8.33 | 9.46 | 14.53 | 8.04 | 8.73 | 9.24 |
| POLR3A | 10 | 8.69 | 10.28 | 10.16 | 10.52 | 9.82 | 10.67 | 11.38 | 11.08 | 8.61 | 8.79 | 9.49 | 10.34 | 10.25 | 11.23 | 8.70 |
| POLR3B | 10 | 9.50 | 9.42 | 9.47 | 8.81 | 10.76 | 10.41 | 10.82 | 11.58 | 9.83 | 9.41 | 9.46 | 9.14 | 10.58 | 11.20 | 9.62 |
| POLR3C | 8 | 9.58 | 10.76 | 10.52 | 10.39 | 10.22 | 10.50 | 9.77 | 9.22 | 9.83 | 9.21 | 10.17 | 10.45 | 10.36 | 9.50 | 9.52 |
| POLR3D | 6 | 9.78 | 10.47 | 8.79 | 8.61 | 9.96 | 10.48 | 10.88 | 12.69 | 9.51 | 8.84 | 10.12 | 8.70 | 10.22 | 11.78 | 9.17 |
| POLR3E | 7 | 8.88 | 10.29 | 8.65 | 9.14 | 10.67 | 10.96 | 11.78 | 10.54 | 9.33 | 9.76 | 9.58 | 8.90 | 10.82 | 11.16 | 9.54 |
| POLR3F | 2 | 9.47 | 8.46 | 9.76 | 10.29 | 9.93 | 9.42 | 11.51 | 10.89 | 10.59 | 9.67 | 8.96 | 10.03 | 9.68 | 11.20 | 10.13 |
| POLR3H | 1 | 10.83 | 9.70 | 10.51 | 11.16 | 9.06 | 9.58 | 9.18 | 9.65 | 9.72 | 10.61 | 10.26 | 10.83 | 9.32 | 9.42 | 10.17 |
| POLRMT | 7 | 10.10 | 8.64 | 11.83 | 14.51 | 7.85 | 7.67 | 10.34 | 10.02 | 8.98 | 10.04 | 9.37 | 13.17 | 7.76 | 10.18 | 9.51 |
| POM121 | 4 | 11.89 | 8.90 | 8.91 | 10.36 | 8.58 | 9.14 | 10.47 | 10.15 | 10.47 | 11.14 | 10.39 | 9.63 | 8.86 | 10.31 | 10.80 |
| POM121C | 3 | 9.97 | 10.10 | 8.25 | 10.54 | 10.04 | 9.87 | 9.58 | 8.56 | 10.98 | 12.11 | 10.04 | 9.39 | 9.95 | 9.07 | 11.55 |
| POMP | 2 | 7.56 | 8.23 | 6.29 | 6.32 | 13.26 | 13.83 | 19.31 | 11.53 | 6.95 | 6.72 | 7.90 | 6.31 | 13.55 | 15.42 | 6.83 |
| PON2 | 9 | 14.64 | 5.85 | 5.77 | 6.90 | 10.91 | 9.09 | 8.85 | 17.09 | 9.73 | 11.18 | 10.24 | 6.33 | 10.00 | 12.97 | 10.45 |
| POP1 | 14 | 8.44 | 9.68 | 11.27 | 11.80 | 7.77 | 8.01 | 11.56 | 10.47 | 10.07 | 10.93 | 9.06 | 11.54 | 7.89 | 11.01 | 10.50 |
| POP4 | 3 | 9.61 | 10.44 | 10.48 | 10.73 | 9.04 | 8.21 | 10.96 | 10.56 | 10.33 | 9.64 | 10.03 | 10.60 | 8.62 | 10.76 | 9.99 |
| POP5 | 1 | 8.77 | 11.88 | 7.87 | 13.26 | 9.26 | 10.02 | 9.10 | 10.55 | 9.59 | 9.71 | 10.32 | 10.57 | 9.64 | 9.82 | 9.65 |
| POP7 | 3 | 9.00 | 9.26 | 10.34 | 10.21 | 9.04 | 10.06 | 11.05 | 10.37 | 10.88 | 9.78 | 9.13 | 10.28 | 9.55 | 10.71 | 10.33 |
| POR | 15 | 11.85 | 6.64 | 8.47 | 12.14 | 7.76 | 6.97 | 12.72 | 13.23 | 9.09 | 11.13 | 9.24 | 10.30 | 7.37 | 12.97 | 10.11 |
| POU2F1 | 3 | 6.27 | 10.07 | 11.79 | 11.12 | 11.28 | 11.61 | 11.65 | 11.08 | 7.29 | 7.83 | 8.17 | 11.46 | 11.45 | 11.36 | 7.56 |
| PPA1 | 32 | 9.36 | 9.72 | 9.43 | 8.44 | 10.64 | 10.27 | 10.02 | 14.22 | 9.16 | 8.74 | 9.54 | 8.94 | 10.45 | 12.12 | 8.95 |
| PPA2 | 16 | 9.61 | 9.03 | 9.23 | 8.73 | 12.26 | 11.41 | 12.23 | 10.59 | 8.24 | 8.66 | 9.32 | 8.98 | 11.84 | 11.41 | 8.45 |
| PPAN | 11 | 9.75 | 9.47 | 10.42 | 10.50 | 7.88 | 8.27 | 10.52 | 12.89 | 9.61 | 10.70 | 9.61 | 10.46 | 8.07 | 11.70 | 10.15 |
| PPAP2C | 1 | 11.63 | 7.70 | 6.98 | 9.52 | 12.51 | 13.07 | 11.59 | 11.84 | 7.76 | 7.40 | 9.66 | 8.25 | 12.79 | 11.71 | 7.58 |
| PPAT | 22 | 9.80 | 10.41 | 11.06 | 10.37 | 9.04 | 9.29 | 9.81 | 9.62 | 10.32 | 10.27 | 10.11 | 10.72 | 9.17 | 9.71 | 10.29 |
| PPCS | 11 | 9.52 | 10.12 | 7.42 | 6.75 | 13.13 | 12.75 | 10.78 | 10.36 | 9.78 | 9.38 | 9.82 | 7.09 | 12.94 | 10.57 | 9.58 |
| PPFIA1 | 19 | 9.65 | 10.54 | 10.20 | 10.02 | 10.01 | 10.89 | 9.07 | 8.19 | 10.73 | 10.69 | 10.09 | 10.11 | 10.45 | 8.63 | 10.71 |
| PPFIBP1 | 27 | 8.30 | 9.39 | 14.88 | 12.85 | 7.77 | 7.71 | 9.00 | 8.09 | 11.24 | 10.77 | 8.85 | 13.86 | 7.74 | 8.54 | 11.00 |
| PPHLN1 | 3 | 9.91 | 10.95 | 10.00 | 9.30 | 11.44 | 11.11 | 10.35 | 9.33 | 9.18 | 8.43 | 10.43 | 9.65 | 11.28 | 9.84 | 8.81 |
| PPIA | 85 | 9.29 | 12.54 | 9.71 | 9.51 | 8.52 | 8.63 | 9.64 | 13.64 | 9.54 | 8.99 | 10.91 | 9.61 | 8.58 | 11.64 | 9.27 |
| PPIB | 51 | 8.48 | 8.90 | 9.33 | 8.67 | 11.85 | 11.69 | 8.69 | 12.38 | 10.07 | 9.94 | 8.69 | 9.00 | 11.77 | 10.53 | 10.01 |
| PPIC | 8 | 10.25 | 11.54 | 10.16 | 10.03 | 11.92 | 12.38 | 8.52 | 8.17 | 8.56 | 8.47 | 10.89 | 10.10 | 12.15 | 8.34 | 8.52 |

| | | | | | | | | | | | | | | | | |
|---|---|---|---|---|---|---|---|---|---|---|---|---|---|---|---|---|
| PPID | 27 | 9.00 | 9.65 | 11.07 | 10.47 | 9.50 | 9.46 | 12.10 | 10.95 | 9.10 | 8.70 | 9.32 | 10.77 | 9.48 | 11.52 | 8.90 |
| PPIE | 15 | 10.98 | 11.34 | 10.79 | 10.23 | 8.20 | 8.41 | 9.62 | 8.89 | 11.29 | 10.24 | 11.16 | 10.51 | 8.31 | 9.26 | 10.76 |
| PPIF | 8 | 10.88 | 9.26 | 11.84 | 11.77 | 8.30 | 7.69 | 11.27 | 10.17 | 9.68 | 9.14 | 10.07 | 11.80 | 8.00 | 10.72 | 9.41 |
| PPIG | 4 | 8.94 | 9.78 | 9.83 | 9.59 | 10.10 | 10.06 | 11.50 | 10.33 | 10.36 | 9.49 | 9.36 | 9.71 | 10.08 | 10.92 | 9.93 |
| PPIH | 16 | 8.26 | 8.53 | 8.48 | 7.18 | 8.15 | 7.48 | 10.56 | 25.33 | 8.10 | 7.91 | 8.40 | 7.83 | 7.82 | 17.94 | 8.01 |
| PPIL1 | 15 | 10.15 | 10.95 | 9.63 | 9.51 | 9.66 | 9.71 | 9.68 | 11.41 | 10.01 | 9.28 | 10.55 | 9.57 | 9.69 | 10.54 | 9.65 |
| PPIL2 | 8 | 10.84 | 11.54 | 12.40 | 10.83 | 7.69 | 7.36 | 9.46 | 8.62 | 10.93 | 10.32 | 11.19 | 11.62 | 7.52 | 9.04 | 10.63 |
| PPIL3 | 9 | 8.21 | 8.76 | 8.95 | 7.33 | 13.68 | 13.00 | 11.61 | 10.28 | 9.44 | 8.74 | 8.48 | 8.14 | 13.34 | 10.94 | 9.09 |
| PPIL4 | 7 | 9.26 | 11.22 | 10.85 | 9.56 | 9.72 | 9.78 | 9.80 | 9.21 | 10.72 | 9.88 | 10.24 | 10.21 | 9.75 | 9.51 | 10.30 |
| PPIP5K2 | 10 | 7.93 | 10.13 | 11.15 | 11.32 | 10.33 | 10.63 | 9.98 | 8.13 | 10.02 | 10.38 | 9.03 | 11.24 | 10.48 | 9.05 | 10.20 |
| PPL | 56 | 11.28 | 12.99 | 9.77 | 9.58 | 13.01 | 13.75 | 2.89 | 2.99 | 12.15 | 11.59 | 12.13 | 9.68 | 13.38 | 2.94 | 11.87 |
| PPM1A | 9 | 10.53 | 10.06 | 10.53 | 10.04 | 10.82 | 10.92 | 11.81 | 10.11 | 7.89 | 7.28 | 10.29 | 10.29 | 10.87 | 10.96 | 7.58 |
| PPM1B | 8 | 9.52 | 9.98 | 11.29 | 11.57 | 10.52 | 10.98 | 9.05 | 8.30 | 9.37 | 9.42 | 9.75 | 11.43 | 10.75 | 8.68 | 9.40 |
| PPM1D | 1 | 11.31 | 11.80 | 9.93 | 10.80 | 9.65 | 11.70 | 8.36 | 7.97 | 9.40 | 9.10 | 11.55 | 10.37 | 10.68 | 8.16 | 9.25 |
| PPM1F | 5 | 11.70 | 11.08 | 9.53 | 10.27 | 8.09 | 9.41 | 9.15 | 7.95 | 11.70 | 11.13 | 11.39 | 9.90 | 8.75 | 8.55 | 11.41 |
| PPM1G | 33 | 8.98 | 9.37 | 10.82 | 9.74 | 8.61 | 8.74 | 10.45 | 13.84 | 9.83 | 9.62 | 9.17 | 10.28 | 8.68 | 12.15 | 9.73 |
| PPME1 | 32 | 11.75 | 12.13 | 9.37 | 9.49 | 10.36 | 11.28 | 7.93 | 9.15 | 9.25 | 9.29 | 11.94 | 9.43 | 10.82 | 8.54 | 9.27 |
| PPOX | 3 | 10.98 | 10.40 | 7.40 | 8.03 | 12.12 | 12.44 | 8.30 | 8.22 | 10.53 | 11.58 | 10.69 | 7.72 | 12.28 | 8.26 | 11.06 |
| PPP1CA | 12 | 8.71 | 9.71 | 12.71 | 11.68 | 9.17 | 9.78 | 9.07 | 9.72 | 9.98 | 9.45 | 9.21 | 12.20 | 9.48 | 9.40 | 9.72 |
| PPP1CB | 8 | 10.03 | 9.27 | 9.13 | 7.92 | 10.57 | 10.20 | 8.56 | 15.65 | 10.25 | 8.42 | 9.65 | 8.52 | 10.39 | 12.10 | 9.34 |
| PPP1CC | 17 | 8.12 | 8.08 | 9.45 | 7.80 | 8.82 | 8.05 | 8.12 | 25.30 | 8.50 | 7.77 | 8.10 | 8.63 | 8.43 | 16.71 | 8.13 |
| PPP1R10 | 20 | 9.65 | 10.38 | 9.39 | 8.94 | 11.04 | 11.72 | 9.21 | 8.66 | 10.48 | 10.53 | 10.01 | 9.17 | 11.38 | 8.94 | 10.50 |
| PPP1R11 | 3 | 10.25 | 9.88 | 8.65 | 7.57 | 11.60 | 11.17 | 11.82 | 8.32 | 10.97 | 9.78 | 10.06 | 8.11 | 11.38 | 10.07 | 10.37 |
| PPP1R12A | 50 | 9.87 | 11.20 | 13.88 | 13.26 | 8.86 | 9.06 | 6.64 | 6.22 | 10.81 | 10.20 | 10.53 | 13.57 | 8.96 | 6.43 | 10.50 |
| PPP1R12B | 2 | 10.22 | 13.08 | 7.75 | 8.32 | 15.73 | 17.57 | 11.82 | 10.01 | 3.02 | 2.48 | 11.65 | 8.03 | 16.65 | 10.92 | 2.75 |
| PPP1R12C | 5 | 10.39 | 12.39 | 8.47 | 8.41 | 10.80 | 11.31 | 9.68 | 8.79 | 9.87 | 9.89 | 11.39 | 8.44 | 11.05 | 9.24 | 9.88 |
| PPP1R13L | 30 | 10.29 | 10.68 | 11.97 | 10.67 | 9.95 | 9.62 | 6.87 | 6.06 | 12.52 | 11.37 | 10.49 | 11.32 | 9.78 | 6.46 | 11.95 |
| PPP1R14B | 3 | 9.46 | 10.32 | 12.01 | 10.31 | 10.47 | 11.45 | 10.82 | 5.35 | 10.38 | 9.41 | 9.89 | 11.16 | 10.96 | 8.08 | 9.90 |
| PPP1R18 | 15 | 8.77 | 9.55 | 12.19 | 11.40 | 9.81 | 9.96 | 7.70 | 6.77 | 12.13 | 11.74 | 9.16 | 11.79 | 9.89 | 7.23 | 11.93 |
| PPP1R2 | 6 | 10.13 | 12.07 | 10.67 | 9.94 | 9.77 | 10.38 | 12.04 | 7.61 | 8.79 | 8.61 | 11.10 | 10.30 | 10.07 | 9.83 | 8.70 |
| PPP1R21 | 3 | 9.74 | 10.96 | 9.23 | 9.37 | 12.29 | 13.48 | 8.28 | 7.50 | 9.88 | 9.27 | 10.35 | 9.30 | 12.89 | 7.89 | 9.58 |
| PPP1R35 | 1 | 10.29 | 12.26 | 9.79 | 9.79 | 7.09 | 8.02 | 9.11 | 12.74 | 9.65 | 11.26 | 11.28 | 9.79 | 7.56 | 10.92 | 10.45 |
| PPP1R7 | 16 | 10.22 | 10.83 | 9.96 | 9.10 | 12.24 | 12.17 | 9.17 | 8.11 | 9.47 | 8.74 | 10.53 | 9.53 | 12.20 | 8.64 | 9.10 |
| PPP1R8 | 12 | 9.44 | 11.31 | 9.39 | 9.54 | 9.62 | 9.78 | 10.88 | 11.72 | 9.00 | 9.32 | 10.38 | 9.46 | 9.70 | 11.30 | 9.16 |
| PPP1R9A | 4 | 9.85 | 11.50 | 14.18 | 14.17 | 8.42 | 8.73 | 7.06 | 6.81 | 9.46 | 9.82 | 10.68 | 14.17 | 8.57 | 6.94 | 9.64 |
| PPP1R9B | 12 | 10.31 | 10.65 | 11.02 | 10.28 | 8.67 | 8.80 | 11.61 | 9.75 | 9.67 | 9.24 | 10.48 | 10.65 | 8.73 | 10.68 | 9.46 |
| PPP2CA | 14 | 8.68 | 9.87 | 9.26 | 8.39 | 9.94 | 10.62 | 9.79 | 14.93 | 9.29 | 9.23 | 9.27 | 8.83 | 10.28 | 12.36 | 9.26 |
| PPP2CB | 1 | 9.77 | 11.07 | 10.93 | 10.15 | 9.09 | 9.55 | 10.36 | 8.78 | 10.89 | 9.40 | 10.42 | 10.54 | 9.32 | 9.57 | 10.15 |
| PPP2R1A | 31 | 9.29 | 9.07 | 9.77 | 7.97 | 10.49 | 10.04 | 9.98 | 15.42 | 9.38 | 8.57 | 9.18 | 8.87 | 10.27 | 12.70 | 8.98 |
| PPP2R1B | 15 | 7.66 | 8.57 | 9.03 | 8.42 | 8.51 | 8.43 | 8.96 | 23.96 | 8.49 | 7.96 | 8.12 | 8.73 | 8.47 | 16.46 | 8.22 |
| PPP2R2A | 21 | 9.40 | 9.51 | 9.10 | 8.54 | 10.08 | 10.38 | 10.94 | 14.24 | 9.23 | 8.59 | 9.45 | 8.82 | 10.23 | 12.59 | 8.91 |
| PPP2R2D | 2 | 10.27 | 9.61 | 11.21 | 8.68 | 10.42 | 9.09 | 11.20 | 8.94 | 10.06 | 10.51 | 9.94 | 9.95 | 9.76 | 10.07 | 10.29 |

| | | | | | | | | | | | | | | | | |
|---|---|---|---|---|---|---|---|---|---|---|---|---|---|---|---|---|
| PPP2R4 | 19 | 9.38 | 10.12 | 10.10 | 9.76 | 11.58 | 11.13 | 8.48 | 8.27 | 10.68 | 10.51 | 9.75 | 9.93 | 11.35 | 8.37 | 10.59 |
| PPP2R5A | 4 | 10.03 | 10.65 | 8.13 | 7.76 | 12.67 | 12.22 | 8.36 | 9.00 | 10.11 | 11.08 | 10.34 | 7.94 | 12.44 | 8.68 | 10.59 |
| PPP2R5C | 6 | 9.97 | 9.89 | 9.34 | 8.20 | 11.07 | 10.81 | 10.47 | 10.45 | 10.21 | 9.59 | 9.93 | 8.77 | 10.94 | 10.46 | 9.90 |
| PPP2R5D | 23 | 9.40 | 9.76 | 10.81 | 10.09 | 10.38 | 10.06 | 9.61 | 9.51 | 9.97 | 10.41 | 9.58 | 10.45 | 10.22 | 9.56 | 10.19 |
| PPP2R5E | 4 | 9.11 | 9.99 | 9.88 | 9.68 | 9.05 | 9.06 | 10.35 | 11.18 | 10.54 | 11.16 | 9.55 | 9.78 | 9.06 | 10.76 | 10.85 |
| PPP3CA | 23 | 10.13 | 10.55 | 10.22 | 9.59 | 10.24 | 10.40 | 9.83 | 9.19 | 10.28 | 9.56 | 10.34 | 9.91 | 10.32 | 9.51 | 9.92 |
| PPP3CB | 8 | 10.44 | 11.89 | 9.32 | 7.26 | 10.57 | 10.37 | 9.68 | 9.92 | 10.46 | 10.09 | 11.17 | 8.29 | 10.47 | 9.80 | 10.27 |
| PPP3CC | 1 | 11.97 | 10.09 | 8.33 | 8.60 | 10.12 | 10.33 | 9.80 | 9.47 | 11.38 | 9.91 | 11.03 | 8.46 | 10.23 | 9.63 | 10.64 |
| PPP3R1 | 8 | 10.32 | 10.65 | 11.13 | 10.53 | 10.73 | 10.36 | 9.87 | 6.48 | 10.30 | 9.64 | 10.49 | 10.83 | 10.54 | 8.18 | 9.97 |
| PPP4C | 3 | 7.41 | 7.76 | 7.57 | 6.98 | 9.03 | 8.87 | 7.39 | 29.56 | 7.94 | 7.50 | 7.58 | 7.28 | 8.95 | 18.48 | 7.72 |
| PPP4R1 | 5 | 10.09 | 9.87 | 11.89 | 11.47 | 10.21 | 9.54 | 6.58 | 6.38 | 11.96 | 12.00 | 9.98 | 11.68 | 9.88 | 6.48 | 11.98 |
| PPP4R2 | 10 | 9.59 | 10.56 | 9.50 | 9.35 | 10.41 | 11.23 | 10.12 | 9.49 | 9.96 | 9.80 | 10.07 | 9.42 | 10.82 | 9.81 | 9.88 |
| PPP5C | 24 | 9.38 | 10.04 | 10.55 | 7.13 | 10.47 | 9.42 | 11.49 | 14.29 | 8.80 | 8.43 | 9.71 | 8.84 | 9.95 | 12.89 | 8.61 |
| PPP6C | 12 | 10.31 | 11.01 | 10.52 | 9.31 | 9.92 | 9.78 | 9.79 | 10.01 | 9.69 | 9.68 | 10.66 | 9.92 | 9.85 | 9.90 | 9.68 |
| PPP6R1 | 10 | 8.41 | 9.45 | 9.70 | 9.30 | 9.13 | 8.88 | 10.96 | 15.32 | 9.64 | 9.21 | 8.93 | 9.50 | 9.01 | 13.14 | 9.42 |
| PPP6R2 | 1 | 9.05 | 10.90 | 10.18 | 12.07 | 7.86 | 9.82 | 10.07 | 8.80 | 10.91 | 10.35 | 9.97 | 11.12 | 8.84 | 9.44 | 10.63 |
| PPP6R3 | 14 | 10.45 | 10.68 | 10.25 | 9.49 | 10.36 | 10.48 | 10.40 | 10.12 | 8.98 | 8.78 | 10.57 | 9.87 | 10.42 | 10.26 | 8.88 |
| PPT1 | 9 | 10.68 | 10.91 | 12.46 | 10.95 | 9.00 | 8.73 | 10.10 | 9.71 | 8.82 | 8.64 | 10.79 | 11.71 | 8.87 | 9.90 | 8.73 |
| PPT2 | 5 | 9.36 | 9.73 | 9.49 | 9.96 | 11.91 | 12.29 | 10.69 | 10.19 | 8.40 | 7.97 | 9.55 | 9.72 | 12.10 | 10.44 | 8.19 |
| PPWD1 | 17 | 9.73 | 10.35 | 8.96 | 9.05 | 8.75 | 9.27 | 11.64 | 11.29 | 10.61 | 10.35 | 10.04 | 9.00 | 9.01 | 11.47 | 10.48 |
| PQBP1 | 12 | 9.84 | 10.30 | 10.11 | 9.50 | 9.84 | 10.36 | 12.14 | 9.97 | 8.98 | 8.94 | 10.07 | 9.81 | 10.10 | 11.06 | 8.96 |
| PRAF2 | 2 | 10.76 | 7.06 | 7.79 | 10.93 | 10.60 | 8.59 | 10.83 | 10.58 | 10.68 | 12.18 | 8.91 | 9.36 | 9.59 | 10.71 | 11.43 |
| PRC1 | 17 | 9.18 | 9.53 | 8.67 | 8.08 | 10.85 | 11.29 | 10.26 | 9.82 | 11.39 | 10.93 | 9.36 | 8.38 | 11.07 | 10.04 | 11.16 |
| PRCC | 17 | 8.75 | 9.29 | 11.72 | 9.51 | 11.16 | 11.14 | 10.47 | 9.17 | 9.53 | 9.25 | 9.02 | 10.62 | 11.15 | 9.82 | 9.39 |
| PRCP | 5 | 9.10 | 8.83 | 10.20 | 9.62 | 10.60 | 10.22 | 13.96 | 13.71 | 6.99 | 6.77 | 8.96 | 9.91 | 10.41 | 13.83 | 6.88 |
| PRDX1 | 117 | 11.36 | 16.21 | 7.99 | 8.26 | 8.70 | 8.66 | 9.19 | 8.93 | 10.37 | 10.34 | 13.78 | 8.12 | 8.68 | 9.06 | 10.36 |
| PRDX2 | 35 | 9.39 | 11.27 | 7.29 | 6.83 | 12.69 | 13.09 | 11.55 | 12.59 | 7.77 | 7.52 | 10.33 | 7.06 | 12.89 | 12.07 | 7.65 |
| PRDX3 | 15 | 10.87 | 9.31 | 10.01 | 11.18 | 10.91 | 10.62 | 9.02 | 8.30 | 9.64 | 10.13 | 10.09 | 10.60 | 10.77 | 8.66 | 9.89 |
| PRDX4 | 29 | 10.57 | 10.98 | 7.00 | 6.95 | 9.51 | 8.82 | 10.91 | 17.90 | 8.78 | 8.58 | 10.78 | 6.98 | 9.17 | 14.40 | 8.68 |
| PRDX5 | 31 | 8.62 | 9.04 | 9.69 | 9.16 | 11.75 | 11.50 | 9.61 | 12.34 | 9.34 | 8.94 | 8.83 | 9.43 | 11.63 | 10.97 | 9.14 |
| PRDX6 | 53 | 8.80 | 10.72 | 10.01 | 10.01 | 13.05 | 13.60 | 8.95 | 8.89 | 7.87 | 8.10 | 9.76 | 10.01 | 13.33 | 8.92 | 7.99 |
| PREB | 5 | 11.81 | 5.72 | 7.83 | 11.01 | 9.36 | 7.60 | 10.83 | 9.82 | 11.00 | 15.00 | 8.77 | 9.42 | 8.48 | 10.33 | 13.00 |
| PRELID1 | 1 | 9.80 | 10.80 | 8.98 | 9.54 | 7.98 | 9.21 | 11.36 | 11.23 | 10.75 | 10.35 | 10.30 | 9.26 | 8.60 | 11.29 | 10.55 |
| PREP | 35 | 8.98 | 9.07 | 12.91 | 11.52 | 7.09 | 6.92 | 13.20 | 12.94 | 8.69 | 8.68 | 9.02 | 12.22 | 7.01 | 13.07 | 8.68 |
| PREPL | 11 | 9.66 | 10.40 | 12.84 | 12.00 | 9.43 | 9.62 | 7.91 | 7.33 | 10.26 | 10.54 | 10.03 | 12.42 | 9.52 | 7.62 | 10.40 |
| PREX1 | 1 | 9.16 | 10.55 | 9.78 | 8.22 | 13.19 | 11.69 | 10.03 | 9.44 | 9.44 | 8.48 | 9.86 | 9.00 | 12.44 | 9.74 | 8.96 |
| PRIM1 | 8 | 9.83 | 10.73 | 9.43 | 9.01 | 9.98 | 9.92 | 11.37 | 11.39 | 9.38 | 8.95 | 10.28 | 9.22 | 9.95 | 11.38 | 9.16 |
| PRIM2 | 14 | 9.85 | 10.77 | 9.48 | 8.55 | 9.54 | 9.50 | 11.72 | 11.44 | 10.00 | 9.16 | 10.31 | 9.01 | 9.52 | 11.58 | 9.58 |
| PRKAA1 | 10 | 8.74 | 9.72 | 12.55 | 10.78 | 10.31 | 10.41 | 9.51 | 8.96 | 9.53 | 9.48 | 9.23 | 11.66 | 10.36 | 9.24 | 9.51 |
| PRKAB1 | 1 | 9.73 | 9.17 | 10.57 | 11.37 | 13.10 | 11.44 | 9.23 | 8.65 | 8.62 | 8.13 | 9.45 | 10.97 | 12.27 | 8.94 | 8.37 |
| PRKAB2 | 2 | 8.84 | 9.09 | 16.36 | 16.40 | 9.80 | 9.74 | 8.03 | 7.60 | 6.96 | 7.18 | 8.96 | 16.38 | 9.77 | 7.82 | 7.07 |
| PRKACA | 1 | 14.82 | 12.55 | 6.72 | 7.20 | 10.65 | 11.64 | 6.03 | 6.12 | 12.21 | 12.05 | 13.68 | 6.96 | 11.14 | 6.07 | 12.13 |

| | | | | | | | | | | | | | | | | |
|---|---|---|---|---|---|---|---|---|---|---|---|---|---|---|---|---|
| PRKACB | 14 | 12.27 | 11.95 | 9.59 | 9.23 | 8.83 | 9.25 | 7.73 | 7.87 | 11.67 | 11.60 | 12.11 | 9.41 | 9.04 | 7.80 | 11.63 |
| PRKAG1 | 6 | 9.88 | 10.06 | 10.89 | 10.10 | 11.36 | 11.36 | 9.36 | 10.23 | 8.56 | 8.20 | 9.97 | 10.50 | 11.36 | 9.80 | 8.38 |
| PRKAR1A | 22 | 11.92 | 13.11 | 11.21 | 10.48 | 9.07 | 9.01 | 5.32 | 6.52 | 11.94 | 11.42 | 12.51 | 10.84 | 9.04 | 5.92 | 11.68 |
| PRKAR1B | 7 | 11.81 | 13.20 | 8.50 | 8.86 | 9.48 | 9.89 | 7.42 | 6.84 | 11.96 | 12.04 | 12.51 | 8.68 | 9.69 | 7.13 | 12.00 |
| PRKAR2A | 26 | 10.10 | 10.35 | 9.69 | 9.73 | 9.81 | 10.42 | 10.19 | 9.47 | 10.22 | 10.01 | 10.23 | 9.71 | 10.11 | 9.83 | 10.12 |
| PRKCA | 15 | 9.83 | 10.73 | 13.02 | 11.78 | 8.11 | 7.97 | 9.37 | 9.80 | 9.91 | 9.48 | 10.28 | 12.40 | 8.04 | 9.59 | 9.70 |
| PRKCD | 5 | 9.90 | 9.82 | 10.26 | 11.40 | 9.02 | 9.13 | 10.37 | 9.83 | 10.07 | 10.20 | 9.86 | 10.83 | 9.07 | 10.10 | 10.14 |
| PRKCDBP | 11 | 8.29 | 9.86 | 15.08 | 14.76 | 6.06 | 6.06 | 6.90 | 6.33 | 13.11 | 13.55 | 9.07 | 14.92 | 6.06 | 6.62 | 13.33 |
| PRKCG | 6 | 9.72 | 10.39 | 11.75 | 12.27 | 8.12 | 8.20 | 9.70 | 9.71 | 10.10 | 10.03 | 10.06 | 12.01 | 8.16 | 9.71 | 10.06 |
| PRKCI | 1 | 8.59 | 10.21 | 12.19 | 7.75 | 11.64 | 7.15 | 11.02 | 11.42 | 10.32 | 9.70 | 9.40 | 9.97 | 9.40 | 11.22 | 10.01 |
| PRKCSH | 39 | 9.43 | 8.17 | 9.24 | 9.46 | 10.99 | 10.44 | 12.77 | 11.27 | 8.97 | 9.26 | 8.80 | 9.35 | 10.71 | 12.02 | 9.12 |
| PRKD2 | 1 | 6.96 | 9.70 | 10.69 | 9.50 | 10.26 | 11.70 | 12.74 | 10.45 | 8.58 | 9.40 | 8.33 | 10.10 | 10.98 | 11.60 | 8.99 |
| PRKDC | 257 | 8.97 | 8.40 | 8.16 | 8.74 | 9.82 | 9.47 | 12.68 | 15.77 | 8.86 | 9.12 | 8.68 | 8.45 | 9.65 | 14.22 | 8.99 |
| PRKRA | 6 | 9.12 | 10.24 | 8.24 | 6.59 | 11.74 | 11.19 | 13.19 | 11.10 | 9.11 | 9.49 | 9.68 | 7.42 | 11.47 | 12.14 | 9.30 |
| PRKRIP1 | 2 | 11.03 | 10.39 | 13.74 | 12.91 | 9.23 | 7.87 | 9.09 | 7.88 | 9.92 | 7.95 | 10.71 | 13.32 | 8.55 | 8.48 | 8.93 |
| PRKRIR | 3 | 9.18 | 9.60 | 11.24 | 11.60 | 8.36 | 8.27 | 11.27 | 11.37 | 10.34 | 8.79 | 9.39 | 11.42 | 8.31 | 11.32 | 9.56 |
| PRMT1 | 20 | 8.42 | 7.96 | 10.67 | 7.80 | 7.97 | 6.64 | 11.07 | 21.77 | 9.18 | 8.51 | 8.19 | 9.23 | 7.31 | 16.42 | 8.85 |
| PRMT10 | 3 | 11.00 | 11.03 | 10.43 | 8.82 | 8.78 | 9.46 | 10.17 | 9.66 | 11.12 | 9.52 | 11.01 | 9.62 | 9.12 | 9.92 | 10.32 |
| PRMT3 | 9 | 8.93 | 10.78 | 10.44 | 9.28 | 10.42 | 10.63 | 10.40 | 10.70 | 8.77 | 9.66 | 9.85 | 9.86 | 10.53 | 10.55 | 9.22 |
| PRMT5 | 22 | 9.28 | 9.18 | 9.74 | 9.04 | 9.55 | 9.20 | 9.44 | 14.66 | 10.35 | 9.55 | 9.23 | 9.39 | 9.37 | 12.05 | 9.95 |
| PRMT6 | 5 | 10.33 | 11.80 | 9.85 | 9.04 | 9.63 | 9.03 | 9.86 | 9.97 | 10.25 | 10.25 | 11.07 | 9.44 | 9.33 | 9.92 | 10.25 |
| PRMT7 | 2 | 10.63 | 12.01 | 6.65 | 7.70 | 12.06 | 13.01 | 9.72 | 10.35 | 9.00 | 8.87 | 11.32 | 7.17 | 12.53 | 10.03 | 8.94 |
| PROCR | 1 | 9.83 | 9.24 | 12.17 | 12.03 | 8.52 | 7.74 | 9.78 | 8.59 | 10.03 | 12.07 | 9.53 | 12.10 | 8.13 | 9.19 | 11.05 |
| PROSC | 7 | 8.69 | 9.00 | 8.77 | 6.50 | 8.54 | 8.02 | 13.19 | 17.55 | 9.72 | 10.04 | 8.84 | 7.63 | 8.28 | 15.37 | 9.88 |
| PROSER1 | 1 | 11.45 | 11.21 | 11.96 | 9.51 | 12.62 | 11.04 | 10.26 | 8.18 | 7.06 | 6.70 | 11.33 | 10.73 | 11.83 | 9.22 | 6.88 |
| PROSER2 | 1 | 10.95 | 10.88 | 13.23 | 12.94 | 8.32 | 10.66 | 6.07 | 5.12 | 11.54 | 10.29 | 10.92 | 13.08 | 9.49 | 5.59 | 10.92 |
| PRPF18 | 2 | 9.65 | 10.83 | 10.37 | 10.80 | 8.30 | 9.04 | 10.07 | 9.38 | 10.65 | 10.92 | 10.24 | 10.58 | 8.67 | 9.72 | 10.79 |
| PRPF19 | 28 | 9.50 | 10.14 | 11.03 | 10.52 | 9.88 | 10.06 | 11.32 | 10.74 | 8.73 | 8.07 | 9.82 | 10.78 | 9.97 | 11.03 | 8.40 |
| PRPF3 | 24 | 9.79 | 10.55 | 9.90 | 10.12 | 10.33 | 10.61 | 9.40 | 8.81 | 9.91 | 10.56 | 10.17 | 10.01 | 10.47 | 9.11 | 10.24 |
| PRPF31 | 15 | 9.47 | 9.97 | 11.46 | 10.30 | 9.27 | 9.14 | 11.90 | 10.37 | 9.30 | 8.82 | 9.72 | 10.88 | 9.21 | 11.14 | 9.06 |
| PRPF38A | 6 | 10.87 | 9.69 | 9.51 | 6.15 | 10.26 | 8.60 | 10.69 | 13.84 | 10.59 | 9.79 | 10.28 | 7.83 | 9.43 | 12.26 | 10.19 |
| PRPF38B | 7 | 8.85 | 9.03 | 9.19 | 8.78 | 9.60 | 9.17 | 12.60 | 13.19 | 9.85 | 9.73 | 8.94 | 8.99 | 9.39 | 12.90 | 9.79 |
| PRPF39 | 3 | 9.41 | 9.54 | 8.08 | 8.53 | 11.32 | 11.21 | 11.67 | 11.43 | 9.91 | 8.89 | 9.48 | 8.31 | 11.27 | 11.55 | 9.40 |
| PRPF4 | 19 | 9.06 | 10.20 | 9.93 | 8.77 | 10.15 | 10.19 | 9.61 | 11.11 | 10.73 | 10.25 | 9.63 | 9.35 | 10.17 | 10.36 | 10.49 |
| PRPF40A | 22 | 9.41 | 9.90 | 11.09 | 9.55 | 10.06 | 9.50 | 11.26 | 10.12 | 9.57 | 9.52 | 9.65 | 10.32 | 9.78 | 10.69 | 9.55 |
| PRPF4B | 19 | 9.42 | 10.49 | 10.35 | 8.99 | 11.04 | 10.72 | 10.37 | 9.75 | 9.65 | 9.22 | 9.96 | 9.67 | 10.88 | 10.06 | 9.43 |
| PRPF6 | 44 | 9.57 | 10.15 | 11.69 | 10.39 | 9.93 | 9.57 | 9.64 | 9.10 | 10.07 | 9.89 | 9.86 | 11.04 | 9.75 | 9.37 | 9.98 |
| PRPF8 | 100 | 9.83 | 9.97 | 11.08 | 10.81 | 8.92 | 8.78 | 9.76 | 9.78 | 10.46 | 10.61 | 9.90 | 10.95 | 8.85 | 9.77 | 10.53 |
| PRPS1 | 10 | 8.47 | 8.87 | 8.11 | 7.14 | 12.47 | 12.79 | 10.28 | 16.51 | 7.62 | 7.76 | 8.67 | 7.62 | 12.63 | 13.39 | 7.69 |
| PRPS2 | 25 | 8.80 | 9.33 | 8.62 | 8.37 | 12.04 | 12.32 | 11.86 | 13.64 | 7.56 | 7.46 | 9.07 | 8.50 | 12.18 | 12.75 | 7.51 |
| PRPSAP1 | 17 | 9.76 | 10.37 | 12.87 | 11.73 | 9.27 | 8.94 | 7.33 | 8.40 | 10.64 | 10.69 | 10.07 | 12.30 | 9.11 | 7.86 | 10.66 |
| PRPSAP2 | 11 | 11.33 | 11.78 | 12.05 | 12.19 | 8.61 | 8.79 | 7.27 | 7.19 | 10.40 | 10.38 | 11.56 | 12.12 | 8.70 | 7.23 | 10.39 |

| | | | | | | | | | | | | | | | | |
|---|---|---|---|---|---|---|---|---|---|---|---|---|---|---|---|---|
| PRR11 | 3 | 9.11 | 9.20 | 14.50 | 16.10 | 7.10 | 7.15 | 7.68 | 7.24 | 11.02 | 10.89 | 9.15 | 15.30 | 7.13 | 7.46 | 10.96 |
| PRR12 | 1 | 7.83 | 10.41 | 8.87 | 8.97 | 12.88 | 11.35 | 13.33 | 9.49 | 9.56 | 7.31 | 9.12 | 8.92 | 12.12 | 11.41 | 8.44 |
| PRR14 | 1 | 9.67 | 11.37 | 11.00 | 11.17 | 10.44 | 11.54 | 8.64 | 7.80 | 9.20 | 9.16 | 10.52 | 11.09 | 10.99 | 8.22 | 9.18 |
| PRR22 | 2 | 14.86 | 12.28 | 11.27 | 10.37 | 6.66 | 5.31 | 13.18 | 13.98 | 7.01 | 5.08 | 13.57 | 10.82 | 5.98 | 13.58 | 6.05 |
| PRR5L | 2 | 10.09 | 10.60 | 9.89 | 9.84 | 10.15 | 10.19 | 8.21 | 8.18 | 11.10 | 11.75 | 10.35 | 9.86 | 10.17 | 8.19 | 11.42 |
| PRRC1 | 13 | 9.72 | 10.30 | 9.02 | 8.87 | 10.94 | 10.79 | 11.25 | 10.60 | 9.54 | 8.95 | 10.01 | 8.95 | 10.87 | 10.93 | 9.25 |
| PRRC2A | 26 | 10.02 | 10.94 | 10.96 | 10.69 | 8.76 | 9.24 | 10.69 | 10.13 | 9.51 | 9.06 | 10.48 | 10.83 | 9.00 | 10.41 | 9.28 |
| PRRC2B | 22 | 9.80 | 11.02 | 11.03 | 10.60 | 9.14 | 9.33 | 9.78 | 9.16 | 10.37 | 9.78 | 10.41 | 10.82 | 9.23 | 9.47 | 10.07 |
| PRRC2C | 84 | 9.28 | 10.44 | 10.44 | 9.93 | 10.02 | 10.44 | 10.31 | 9.42 | 9.96 | 9.77 | 9.86 | 10.18 | 10.23 | 9.86 | 9.86 |
| PRSS21 | 3 | 9.51 | 7.66 | 6.88 | 6.24 | 9.89 | 9.30 | 9.80 | 21.82 | 9.03 | 9.88 | 8.58 | 6.56 | 9.59 | 15.81 | 9.46 |
| PRSS23 | 2 | 5.73 | 4.62 | 14.58 | 15.18 | 5.44 | 6.18 | 15.23 | 13.26 | 10.10 | 9.68 | 5.18 | 14.88 | 5.81 | 14.24 | 9.89 |
| PRUNE | 5 | 10.11 | 11.06 | 10.90 | 8.96 | 10.46 | 10.19 | 10.62 | 7.86 | 9.68 | 10.14 | 10.59 | 9.93 | 10.32 | 9.24 | 9.91 |
| PRUNE2 | 12 | 10.06 | 11.77 | 8.10 | 7.77 | 10.88 | 11.33 | 4.10 | 3.33 | 16.74 | 15.91 | 10.92 | 7.94 | 11.11 | 3.71 | 16.33 |
| PSAP | 18 | 11.28 | 11.45 | 10.04 | 11.06 | 11.58 | 12.48 | 11.66 | 2.66 | 8.61 | 9.17 | 11.37 | 10.55 | 12.03 | 7.16 | 8.89 |
| PSAT1 | 45 | 7.03 | 10.87 | 10.90 | 9.42 | 7.41 | 7.32 | 9.37 | 9.14 | 13.67 | 14.89 | 8.95 | 10.16 | 7.37 | 9.25 | 14.28 |
| PSEN1 | 1 | 8.67 | 8.11 | 9.49 | 13.05 | 10.25 | 11.13 | 9.69 | 7.58 | 10.19 | 11.85 | 8.39 | 11.27 | 10.69 | 8.64 | 11.02 |
| PSIP1 | 42 | 10.28 | 11.47 | 9.12 | 8.66 | 9.24 | 10.03 | 8.71 | 9.30 | 11.55 | 11.64 | 10.88 | 8.89 | 9.63 | 9.01 | 11.59 |
| PSMA1 | 32 | 9.82 | 10.87 | 9.74 | 8.46 | 11.42 | 11.54 | 10.54 | 9.70 | 8.99 | 8.92 | 10.35 | 9.10 | 11.48 | 10.12 | 8.95 |
| PSMA2 | 14 | 9.70 | 10.96 | 10.07 | 8.92 | 11.10 | 11.14 | 10.69 | 9.71 | 9.12 | 8.59 | 10.33 | 9.50 | 11.12 | 10.20 | 8.85 |
| PSMA3 | 22 | 10.17 | 10.54 | 9.74 | 8.44 | 11.67 | 11.58 | 10.85 | 9.41 | 8.85 | 8.75 | 10.35 | 9.09 | 11.63 | 10.13 | 8.80 |
| PSMA4 | 26 | 10.07 | 9.79 | 9.05 | 8.10 | 10.82 | 10.77 | 9.54 | 14.68 | 8.80 | 8.37 | 9.93 | 8.57 | 10.80 | 12.11 | 8.59 |
| PSMA5 | 20 | 9.25 | 10.51 | 11.11 | 9.89 | 10.54 | 10.48 | 10.25 | 10.84 | 8.86 | 8.26 | 9.88 | 10.50 | 10.51 | 10.55 | 8.56 |
| PSMA6 | 27 | 10.06 | 10.67 | 10.08 | 6.87 | 12.01 | 10.94 | 10.79 | 10.94 | 8.99 | 8.64 | 10.37 | 8.48 | 11.47 | 10.87 | 8.82 |
| PSMA7 | 25 | 10.16 | 10.52 | 10.24 | 9.55 | 10.85 | 10.78 | 10.39 | 9.62 | 9.15 | 8.73 | 10.34 | 9.89 | 10.82 | 10.01 | 8.94 |
| PSMB1 | 24 | 9.40 | 10.40 | 10.22 | 9.22 | 11.43 | 10.79 | 11.00 | 9.18 | 9.24 | 9.12 | 9.90 | 9.72 | 11.11 | 10.09 | 9.18 |
| PSMB10 | 1 | 15.99 | 17.21 | 3.95 | 3.89 | 18.63 | 18.61 | 4.96 | 3.70 | 6.99 | 6.07 | 16.60 | 3.92 | 18.62 | 4.33 | 6.53 |
| PSMB2 | 14 | 9.66 | 10.16 | 8.79 | 7.50 | 11.22 | 10.21 | 10.08 | 15.23 | 8.51 | 8.66 | 9.91 | 8.15 | 10.71 | 12.65 | 8.58 |
| PSMB3 | 18 | 10.06 | 10.92 | 8.96 | 7.86 | 11.53 | 10.99 | 10.36 | 12.01 | 8.73 | 8.60 | 10.49 | 8.41 | 11.26 | 11.18 | 8.66 |
| PSMB4 | 24 | 10.31 | 11.27 | 9.70 | 8.69 | 11.20 | 11.66 | 9.84 | 9.32 | 8.99 | 9.02 | 10.79 | 9.20 | 11.43 | 9.58 | 9.00 |
| PSMB5 | 20 | 10.27 | 10.82 | 10.04 | 8.75 | 10.31 | 10.55 | 11.11 | 9.73 | 9.27 | 9.16 | 10.54 | 9.39 | 10.43 | 10.42 | 9.22 |
| PSMB6 | 9 | 10.66 | 11.71 | 9.64 | 9.59 | 10.59 | 10.91 | 9.94 | 9.23 | 9.18 | 8.54 | 11.18 | 9.62 | 10.75 | 9.59 | 8.86 |
| PSMB7 | 20 | 9.65 | 10.60 | 8.68 | 8.33 | 11.27 | 11.59 | 11.94 | 11.31 | 8.45 | 8.19 | 10.12 | 8.50 | 11.43 | 11.62 | 8.32 |
| PSMB8 | 7 | 11.78 | 12.49 | 5.75 | 4.82 | 19.11 | 18.52 | 6.64 | 6.03 | 7.96 | 6.90 | 12.13 | 5.28 | 18.82 | 6.33 | 7.43 |
| PSMB9 | 1 | 13.86 | 14.06 | 3.74 | 2.43 | 25.79 | 24.53 | 3.36 | 2.92 | 5.65 | 3.67 | 13.96 | 3.09 | 25.16 | 3.14 | 4.66 |
| PSMC1 | 34 | 10.49 | 9.77 | 9.65 | 10.19 | 9.04 | 8.99 | 10.47 | 11.52 | 10.03 | 9.86 | 10.13 | 9.92 | 9.01 | 10.99 | 9.95 |
| PSMC2 | 36 | 9.50 | 10.40 | 11.41 | 10.63 | 9.64 | 9.61 | 10.40 | 9.17 | 9.77 | 9.47 | 9.95 | 11.02 | 9.62 | 9.78 | 9.62 |
| PSMC3 | 52 | 8.81 | 10.19 | 10.36 | 9.42 | 10.30 | 10.42 | 11.07 | 11.55 | 9.14 | 8.73 | 9.50 | 9.89 | 10.36 | 11.31 | 8.94 |
| PSMC3IP | 1 | 12.34 | 12.97 | 11.83 | 11.25 | 8.66 | 8.86 | 10.83 | 10.67 | 6.70 | 5.88 | 12.66 | 11.54 | 8.76 | 10.75 | 6.29 |
| PSMC4 | 29 | 10.03 | 9.92 | 10.03 | 8.32 | 9.86 | 9.62 | 10.65 | 12.17 | 10.03 | 9.37 | 9.97 | 9.18 | 9.74 | 11.41 | 9.70 |
| PSMC5 | 46 | 9.83 | 10.38 | 10.31 | 9.40 | 9.73 | 9.94 | 10.51 | 10.58 | 9.81 | 9.51 | 10.11 | 9.85 | 9.83 | 10.55 | 9.66 |
| PSMC6 | 36 | 9.22 | 9.84 | 10.14 | 8.78 | 10.25 | 10.20 | 10.90 | 12.26 | 9.41 | 8.99 | 9.53 | 9.46 | 10.22 | 11.58 | 9.20 |
| PSMD1 | 57 | 9.10 | 10.11 | 9.94 | 9.68 | 9.84 | 10.18 | 10.10 | 10.88 | 10.14 | 10.02 | 9.60 | 9.81 | 10.01 | 10.49 | 10.08 |

| | | | | | | | | | | | | | | | | |
|---|---|---|---|---|---|---|---|---|---|---|---|---|---|---|---|---|
| PSMD10 | 15 | 9.56 | 10.63 | 6.96 | 5.68 | 14.69 | 12.97 | 13.06 | 11.31 | 7.44 | 7.70 | 10.09 | 6.32 | 13.83 | 12.19 | 7.57 |
| PSMD11 | 35 | 9.00 | 9.72 | 12.15 | 9.26 | 10.18 | 9.58 | 9.95 | 9.98 | 10.14 | 10.04 | 9.36 | 10.70 | 9.88 | 9.96 | 10.09 |
| PSMD12 | 34 | 9.05 | 9.33 | 10.13 | 9.03 | 9.42 | 9.35 | 9.03 | 14.83 | 10.20 | 9.64 | 9.19 | 9.58 | 9.38 | 11.93 | 9.92 |
| PSMD13 | 26 | 9.37 | 9.41 | 10.31 | 8.85 | 9.90 | 9.79 | 9.72 | 12.74 | 10.17 | 9.75 | 9.39 | 9.58 | 9.84 | 11.23 | 9.96 |
| PSMD14 | 24 | 9.52 | 10.07 | 10.95 | 9.67 | 9.99 | 9.82 | 10.08 | 9.27 | 10.11 | 10.51 | 9.80 | 10.31 | 9.91 | 9.67 | 10.31 |
| PSMD2 | 49 | 9.36 | 10.21 | 11.00 | 10.27 | 9.97 | 10.06 | 10.17 | 9.41 | 9.95 | 9.61 | 9.78 | 10.63 | 10.02 | 9.79 | 9.78 |
| PSMD3 | 42 | 9.90 | 10.03 | 10.09 | 9.13 | 10.03 | 9.78 | 10.26 | 9.34 | 10.83 | 10.61 | 9.97 | 9.61 | 9.90 | 9.80 | 10.72 |
| PSMD4 | 21 | 9.51 | 10.43 | 9.98 | 9.58 | 10.16 | 10.60 | 9.99 | 11.08 | 9.43 | 9.25 | 9.97 | 9.78 | 10.38 | 10.53 | 9.34 |
| PSMD5 | 31 | 10.01 | 10.54 | 10.11 | 9.45 | 9.64 | 9.46 | 9.39 | 8.73 | 11.68 | 10.99 | 10.28 | 9.78 | 9.55 | 9.06 | 11.34 |
| PSMD6 | 26 | 9.68 | 10.32 | 9.67 | 9.06 | 10.15 | 9.96 | 10.28 | 9.60 | 10.81 | 10.47 | 10.00 | 9.36 | 10.05 | 9.94 | 10.64 |
| PSMD7 | 17 | 9.74 | 10.19 | 11.25 | 8.98 | 10.08 | 9.54 | 10.28 | 9.32 | 10.18 | 10.45 | 9.96 | 10.11 | 9.81 | 9.80 | 10.32 |
| PSMD8 | 15 | 10.03 | 10.49 | 9.83 | 8.03 | 10.12 | 10.04 | 9.78 | 10.92 | 10.51 | 10.25 | 10.26 | 8.93 | 10.08 | 10.35 | 10.38 |
| PSMD9 | 10 | 5.82 | 9.11 | 12.59 | 12.03 | 11.75 | 11.21 | 9.49 | 10.14 | 8.37 | 9.49 | 7.47 | 12.31 | 11.48 | 9.81 | 8.93 |
| PSME1 | 20 | 10.35 | 11.48 | 9.60 | 9.24 | 13.12 | 13.88 | 9.43 | 8.89 | 7.24 | 6.78 | 10.91 | 9.42 | 13.50 | 9.16 | 7.01 |
| PSME2 | 13 | 12.23 | 9.35 | 7.47 | 7.01 | 11.25 | 11.60 | 8.46 | 18.34 | 7.57 | 6.73 | 10.79 | 7.24 | 11.42 | 13.40 | 7.15 |
| PSME3 | 17 | 9.75 | 10.35 | 11.57 | 11.07 | 8.64 | 8.70 | 8.98 | 11.27 | 9.88 | 9.78 | 10.05 | 11.32 | 8.67 | 10.13 | 9.83 |
| PSME4 | 16 | 9.14 | 10.26 | 11.10 | 12.36 | 7.88 | 8.18 | 10.48 | 11.40 | 9.35 | 9.86 | 9.70 | 11.73 | 8.03 | 10.94 | 9.61 |
| PSMF1 | 5 | 10.56 | 11.94 | 10.66 | 9.38 | 9.13 | 9.16 | 10.25 | 9.55 | 9.85 | 9.51 | 11.25 | 10.02 | 9.15 | 9.90 | 9.68 |
| PSMG1 | 10 | 10.28 | 11.22 | 8.47 | 8.51 | 11.42 | 11.76 | 9.76 | 9.24 | 9.66 | 9.68 | 10.75 | 8.49 | 11.59 | 9.50 | 9.67 |
| PSMG2 | 7 | 9.53 | 10.88 | 8.94 | 7.96 | 11.39 | 11.39 | 9.01 | 11.73 | 10.05 | 9.12 | 10.20 | 8.45 | 11.39 | 10.37 | 9.59 |
| PSMG3 | 7 | 8.48 | 9.78 | 10.84 | 10.09 | 10.00 | 10.36 | 11.13 | 11.02 | 9.02 | 9.27 | 9.13 | 10.46 | 10.18 | 11.08 | 9.15 |
| PSMG4 | 1 | 16.20 | 9.53 | 7.48 | 8.19 | 9.22 | 8.99 | 10.78 | 10.18 | 10.82 | 8.62 | 12.87 | 7.84 | 9.11 | 10.48 | 9.72 |
| PSPC1 | 27 | 8.43 | 9.47 | 16.10 | 15.19 | 9.49 | 10.20 | 9.15 | 8.19 | 7.07 | 6.71 | 8.95 | 15.65 | 9.85 | 8.67 | 6.89 |
| PSPH | 14 | 8.25 | 8.03 | 14.29 | 7.83 | 8.43 | 7.22 | 10.02 | 7.22 | 14.08 | 14.63 | 8.14 | 11.06 | 7.82 | 8.62 | 14.35 |
| PSTPIP2 | 5 | 9.37 | 10.65 | 18.82 | 17.49 | 9.31 | 9.84 | 6.03 | 5.45 | 6.77 | 6.27 | 10.01 | 18.16 | 9.58 | 5.74 | 6.52 |
| PTAR1 | 5 | 9.43 | 12.06 | 8.77 | 9.71 | 11.15 | 11.28 | 8.83 | 9.02 | 9.64 | 10.12 | 10.74 | 9.24 | 11.21 | 8.93 | 9.88 |
| PTBP1 | 14 | 8.65 | 9.59 | 11.16 | 8.10 | 11.79 | 10.76 | 12.61 | 9.81 | 9.05 | 8.49 | 9.12 | 9.63 | 11.27 | 11.21 | 8.77 |
| PTBP2 | 3 | 10.00 | 11.36 | 11.78 | 10.43 | 9.52 | 8.45 | 12.22 | 10.91 | 8.10 | 7.25 | 10.68 | 11.10 | 8.98 | 11.56 | 7.67 |
| PTBP3 | 15 | 9.36 | 10.14 | 9.73 | 9.62 | 10.89 | 10.87 | 10.71 | 11.46 | 8.63 | 8.60 | 9.75 | 9.68 | 10.88 | 11.08 | 8.61 |
| PTCD2 | 2 | 10.55 | 8.32 | 6.76 | 8.40 | 8.69 | 8.21 | 12.57 | 12.54 | 11.49 | 12.46 | 9.43 | 7.58 | 8.45 | 12.56 | 11.98 |
| PTCD3 | 13 | 11.93 | 9.11 | 8.84 | 10.51 | 10.02 | 9.12 | 11.48 | 11.00 | 8.95 | 9.04 | 10.52 | 9.67 | 9.57 | 11.24 | 8.99 |
| PTDSS2 | 1 | 12.92 | 5.53 | 10.09 | 13.09 | 6.83 | 7.81 | 9.25 | 9.76 | 10.55 | 14.18 | 9.23 | 11.59 | 7.32 | 9.51 | 12.36 |
| PTEN | 3 | 8.97 | 10.61 | 9.97 | 10.04 | 11.08 | 11.70 | 8.70 | 8.21 | 10.11 | 10.61 | 9.79 | 10.01 | 11.39 | 8.45 | 10.36 |
| PTER | 2 | 14.63 | 16.03 | 7.51 | 7.33 | 8.18 | 8.68 | 9.23 | 9.39 | 9.79 | 9.22 | 15.33 | 7.42 | 8.43 | 9.31 | 9.51 |
| PTGES | 2 | 9.93 | 5.00 | 7.63 | 11.59 | 12.36 | 9.82 | 14.91 | 14.57 | 6.30 | 7.91 | 7.46 | 9.61 | 11.09 | 14.74 | 7.10 |
| PTGES2 | 17 | 9.78 | 9.10 | 12.51 | 12.32 | 7.98 | 8.59 | 10.34 | 9.33 | 10.03 | 10.01 | 9.44 | 12.41 | 8.29 | 9.84 | 10.02 |
| PTGES3 | 15 | 8.92 | 9.35 | 10.64 | 9.34 | 9.09 | 8.51 | 10.13 | 15.32 | 9.69 | 9.00 | 9.14 | 9.99 | 8.80 | 12.72 | 9.35 |
| PTGFRN | 13 | 10.56 | 8.78 | 11.46 | 13.02 | 8.55 | 8.72 | 10.54 | 9.43 | 9.35 | 9.59 | 9.67 | 12.24 | 8.63 | 9.98 | 9.47 |
| PTGR1 | 26 | 9.50 | 9.89 | 10.52 | 9.22 | 12.27 | 11.94 | 9.54 | 8.89 | 9.39 | 8.82 | 9.70 | 9.87 | 12.11 | 9.21 | 9.11 |
| PTGR2 | 1 | 9.12 | 12.02 | 9.53 | 9.28 | 9.23 | 9.74 | 11.94 | 9.23 | 9.41 | 10.49 | 10.57 | 9.41 | 9.48 | 10.58 | 9.95 |
| PTGS1 | 3 | 4.50 | 4.47 | 3.53 | 3.00 | 7.18 | 8.65 | 23.46 | 24.91 | 9.41 | 10.89 | 4.48 | 3.26 | 7.92 | 24.19 | 10.15 |
| PTK2 | 17 | 9.31 | 9.45 | 9.47 | 9.41 | 9.41 | 9.34 | 11.58 | 10.53 | 10.93 | 10.56 | 9.38 | 9.44 | 9.38 | 11.05 | 10.75 |

| | | | | | | | | | | | | | | | | |
|---|---|---|---|---|---|---|---|---|---|---|---|---|---|---|---|---|
| PTK7 | 3 | 11.57 | 12.02 | 7.63 | 9.97 | 7.46 | 7.80 | 7.80 | 6.32 | 13.92 | 15.52 | 11.79 | 8.80 | 7.63 | 7.06 | 14.72 |
| PTMA | 27 | 8.86 | 10.95 | 11.89 | 11.62 | 9.98 | 10.08 | 12.69 | 4.84 | 10.16 | 8.91 | 9.91 | 11.76 | 10.03 | 8.77 | 9.54 |
| PTMS | 19 | 9.21 | 11.92 | 11.58 | 11.80 | 10.00 | 10.22 | 9.57 | 3.18 | 11.37 | 11.15 | 10.56 | 11.69 | 10.11 | 6.37 | 11.26 |
| PTOV1 | 1 | 10.45 | 10.33 | 10.66 | 10.00 | 9.85 | 11.65 | 9.87 | 10.19 | 8.33 | 8.67 | 10.39 | 10.33 | 10.75 | 10.03 | 8.50 |
| PTP4A1 | 3 | 10.69 | 10.36 | 9.35 | 8.41 | 11.55 | 11.14 | 9.93 | 9.18 | 9.57 | 9.81 | 10.53 | 8.88 | 11.35 | 9.55 | 9.69 |
| PTPLAD1 | 8 | 12.91 | 6.12 | 6.04 | 10.47 | 9.30 | 7.99 | 7.45 | 7.03 | 14.33 | 18.35 | 9.52 | 8.25 | 8.65 | 7.24 | 16.34 |
| PTPLB | 1 | 16.31 | 7.26 | 6.16 | 11.02 | 9.81 | 9.55 | 5.40 | 6.50 | 12.37 | 15.63 | 11.78 | 8.59 | 9.68 | 5.95 | 14.00 |
| PTPMT1 | 3 | 12.33 | 9.32 | 10.39 | 9.01 | 9.09 | 10.33 | 8.70 | 8.58 | 11.21 | 11.04 | 10.83 | 9.70 | 9.71 | 8.64 | 11.12 |
| PTPN1 | 16 | 12.14 | 6.91 | 9.63 | 11.62 | 8.26 | 7.25 | 9.94 | 10.16 | 11.12 | 12.96 | 9.52 | 10.63 | 7.76 | 10.05 | 12.04 |
| PTPN11 | 24 | 9.08 | 9.86 | 11.69 | 10.73 | 10.54 | 10.38 | 10.46 | 9.42 | 9.04 | 8.79 | 9.47 | 11.21 | 10.46 | 9.94 | 8.91 |
| PTPN12 | 19 | 8.70 | 9.49 | 12.93 | 12.84 | 8.17 | 8.62 | 9.40 | 8.57 | 10.86 | 10.41 | 9.09 | 12.89 | 8.39 | 8.99 | 10.64 |
| PTPN2 | 4 | 11.09 | 9.63 | 9.58 | 8.67 | 9.72 | 9.53 | 10.14 | 9.13 | 11.33 | 11.19 | 10.36 | 9.12 | 9.63 | 9.64 | 11.26 |
| PTPN23 | 26 | 9.90 | 10.43 | 9.16 | 9.52 | 10.83 | 11.38 | 9.01 | 8.16 | 10.74 | 10.86 | 10.17 | 9.34 | 11.10 | 8.59 | 10.80 |
| PTPRE | 1 | 10.54 | 10.53 | 8.40 | 11.47 | 7.78 | 9.27 | 9.73 | 9.21 | 11.65 | 11.43 | 10.53 | 9.94 | 8.52 | 9.47 | 11.54 |
| PTPRF | 27 | 8.15 | 8.99 | 15.84 | 16.03 | 4.07 | 3.85 | 6.60 | 6.14 | 14.94 | 15.41 | 8.57 | 15.93 | 3.96 | 6.37 | 15.17 |
| PTPRJ | 7 | 9.20 | 8.80 | 9.07 | 10.13 | 8.19 | 7.76 | 11.46 | 9.86 | 12.59 | 12.94 | 9.00 | 9.60 | 7.98 | 10.66 | 12.77 |
| PTPRK | 3 | 8.38 | 6.79 | 13.94 | 10.33 | 12.02 | 7.73 | 10.13 | 7.72 | 10.30 | 12.64 | 7.59 | 12.14 | 9.88 | 8.93 | 11.47 |
| PTRF | 27 | 9.57 | 10.09 | 14.80 | 14.23 | 6.02 | 6.10 | 5.76 | 5.29 | 14.16 | 13.98 | 9.83 | 14.51 | 6.06 | 5.53 | 14.07 |
| PTRH1 | 4 | 9.40 | 8.95 | 10.97 | 12.38 | 9.13 | 9.46 | 7.53 | 7.09 | 12.49 | 12.60 | 9.18 | 11.67 | 9.29 | 7.31 | 12.54 |
| PTRH2 | 5 | 11.38 | 8.87 | 9.78 | 11.60 | 7.36 | 8.04 | 10.33 | 7.80 | 12.04 | 12.81 | 10.12 | 10.69 | 7.70 | 9.07 | 12.43 |
| PTRHD1 | 5 | 9.18 | 10.49 | 10.89 | 10.92 | 13.10 | 14.02 | 7.25 | 7.03 | 8.69 | 8.42 | 9.83 | 10.90 | 13.56 | 7.14 | 8.56 |
| PTS | 2 | 11.09 | 10.57 | 11.97 | 11.27 | 11.12 | 10.39 | 5.27 | 4.53 | 11.97 | 11.82 | 10.83 | 11.62 | 10.76 | 4.90 | 11.89 |
| PUF60 | 52 | 9.13 | 9.79 | 10.24 | 9.71 | 9.60 | 9.52 | 11.69 | 11.67 | 9.49 | 9.16 | 9.46 | 9.97 | 9.56 | 11.68 | 9.32 |
| PUM1 | 25 | 9.79 | 10.54 | 9.90 | 9.97 | 8.90 | 9.28 | 11.33 | 10.63 | 9.98 | 9.67 | 10.17 | 9.94 | 9.09 | 10.98 | 9.82 |
| PUM2 | 10 | 8.84 | 9.42 | 10.82 | 10.81 | 9.30 | 10.18 | 10.39 | 13.16 | 8.91 | 8.16 | 9.13 | 10.81 | 9.74 | 11.78 | 8.53 |
| PURA | 9 | 9.09 | 9.96 | 8.33 | 8.52 | 11.10 | 11.10 | 12.42 | 11.58 | 9.03 | 8.87 | 9.52 | 8.43 | 11.10 | 12.00 | 8.95 |
| PURB | 5 | 9.84 | 10.42 | 10.20 | 10.05 | 8.30 | 8.72 | 10.96 | 10.22 | 10.86 | 10.43 | 10.13 | 10.12 | 8.51 | 10.59 | 10.64 |
| PUS1 | 9 | 8.30 | 8.23 | 10.17 | 11.27 | 7.70 | 8.11 | 14.32 | 14.27 | 8.54 | 9.09 | 8.26 | 10.72 | 7.90 | 14.30 | 8.81 |
| PUS3 | 2 | 9.03 | 10.98 | 10.85 | 11.23 | 9.22 | 8.46 | 9.61 | 9.62 | 11.26 | 9.74 | 10.00 | 11.04 | 8.84 | 9.61 | 10.50 |
| PUS7 | 27 | 8.99 | 9.17 | 12.02 | 10.49 | 7.06 | 6.94 | 13.48 | 13.68 | 9.44 | 8.73 | 9.08 | 11.26 | 7.00 | 13.58 | 9.09 |
| PUS7L | 5 | 10.57 | 10.51 | 10.98 | 9.63 | 10.02 | 10.04 | 11.35 | 10.64 | 8.68 | 7.58 | 10.54 | 10.31 | 10.03 | 11.00 | 8.13 |
| PVR | 5 | 9.82 | 8.35 | 10.79 | 13.21 | 7.60 | 7.87 | 11.21 | 9.59 | 10.57 | 11.00 | 9.08 | 12.00 | 7.74 | 10.40 | 10.78 |
| PVRL2 | 7 | 11.15 | 9.13 | 10.34 | 12.55 | 7.80 | 7.98 | 10.16 | 9.12 | 10.87 | 10.91 | 10.14 | 11.44 | 7.89 | 9.64 | 10.89 |
| PWP1 | 9 | 8.20 | 10.48 | 13.36 | 12.68 | 9.06 | 9.76 | 9.73 | 9.90 | 8.39 | 8.43 | 9.34 | 13.02 | 9.41 | 9.82 | 8.41 |
| PWP2 | 11 | 9.29 | 9.54 | 10.39 | 10.20 | 8.44 | 8.27 | 10.47 | 9.87 | 11.81 | 11.72 | 9.41 | 10.29 | 8.36 | 10.17 | 11.76 |
| PWWP2A | 2 | 10.00 | 10.25 | 10.23 | 9.11 | 11.61 | 11.91 | 11.05 | 8.36 | 8.77 | 8.71 | 10.13 | 9.67 | 11.76 | 9.70 | 8.74 |
| PXDC1 | 1 | 8.33 | 12.58 | 13.33 | 9.99 | 8.56 | 9.91 | 9.26 | 9.03 | 9.25 | 9.76 | 10.46 | 11.66 | 9.24 | 9.14 | 9.50 |
| PXDN | 7 | 10.34 | 7.43 | 10.24 | 11.83 | 10.16 | 9.27 | 10.88 | 10.02 | 9.60 | 10.23 | 8.89 | 11.03 | 9.71 | 10.45 | 9.91 |
| PXK | 3 | 8.15 | 8.20 | 10.99 | 11.15 | 8.72 | 8.05 | 11.54 | 13.12 | 10.05 | 10.03 | 8.18 | 11.07 | 8.38 | 12.33 | 10.04 |
| PXN | 20 | 8.90 | 9.71 | 12.87 | 11.81 | 9.49 | 9.45 | 7.36 | 6.76 | 11.96 | 11.69 | 9.31 | 12.34 | 9.47 | 7.06 | 11.82 |
| PYCARD | 1 | 9.67 | 10.55 | 7.90 | 6.44 | 20.83 | 19.02 | 4.36 | 3.25 | 9.66 | 8.33 | 10.11 | 7.17 | 19.93 | 3.80 | 8.99 |
| PYCR1 | 10 | 8.95 | 8.36 | 10.58 | 11.01 | 7.63 | 7.16 | 8.42 | 7.51 | 14.83 | 15.54 | 8.66 | 10.80 | 7.40 | 7.97 | 15.19 |

| | | | | | | | | | | | | | | | | |
|---|---|---|---|---|---|---|---|---|---|---|---|---|---|---|---|---|
| PYCR2 | 11 | 10.96 | 9.85 | 6.47 | 6.92 | 12.01 | 11.36 | 10.61 | 11.49 | 10.11 | 10.22 | 10.40 | 6.70 | 11.69 | 11.05 | 10.16 |
| PYCRL | 5 | 9.34 | 12.10 | 11.11 | 10.08 | 10.08 | 10.44 | 9.24 | 8.11 | 9.55 | 9.95 | 10.72 | 10.59 | 10.26 | 8.68 | 9.75 |
| PYGB | 21 | 9.05 | 9.35 | 10.66 | 10.27 | 12.12 | 12.34 | 8.79 | 8.76 | 9.51 | 9.16 | 9.20 | 10.47 | 12.23 | 8.77 | 9.33 |
| PYGL | 61 | 10.36 | 10.88 | 8.56 | 7.12 | 9.34 | 8.85 | 11.95 | 10.72 | 11.23 | 10.99 | 10.62 | 7.84 | 9.09 | 11.34 | 11.11 |
| PYGO2 | 3 | 8.82 | 9.50 | 10.89 | 10.89 | 10.97 | 11.53 | 9.60 | 8.93 | 9.82 | 9.05 | 9.16 | 10.89 | 11.25 | 9.26 | 9.43 |
| PYROXD1 | 1 | 9.94 | 9.80 | 12.49 | 9.74 | 10.25 | 9.74 | 8.65 | 8.17 | 11.46 | 9.74 | 9.87 | 11.12 | 10.00 | 8.41 | 10.60 |
| PYROXD2 | 1 | 13.15 | 9.66 | 6.08 | 6.66 | 12.84 | 13.89 | 7.93 | 6.67 | 11.50 | 11.62 | 11.41 | 6.37 | 13.37 | 7.30 | 11.56 |
| PZP | 2 | 10.16 | 9.58 | 21.62 | 20.56 | 7.52 | 7.57 | 3.16 | 3.45 | 8.18 | 8.20 | 9.87 | 21.09 | 7.55 | 3.31 | 8.19 |
| QARS | 64 | 9.59 | 10.10 | 10.86 | 9.03 | 10.79 | 10.21 | 8.82 | 8.50 | 11.06 | 11.03 | 9.85 | 9.95 | 10.50 | 8.66 | 11.05 |
| QDPR | 9 | 9.33 | 9.74 | 9.05 | 9.05 | 10.37 | 10.41 | 9.50 | 8.73 | 12.15 | 11.68 | 9.53 | 9.05 | 10.39 | 9.12 | 11.91 |
| QIL1 | 5 | 8.46 | 9.86 | 11.30 | 9.88 | 10.48 | 10.50 | 10.98 | 9.40 | 9.87 | 9.28 | 9.16 | 10.59 | 10.49 | 10.19 | 9.57 |
| QKI | 12 | 8.33 | 9.59 | 11.37 | 11.06 | 9.67 | 10.24 | 9.68 | 10.45 | 9.76 | 9.85 | 8.96 | 11.22 | 9.96 | 10.07 | 9.80 |
| QPRT | 1 | 13.50 | 19.44 | 15.35 | 13.87 | 8.01 | 7.67 | 3.07 | 1.81 | 7.53 | 9.75 | 16.47 | 14.61 | 7.84 | 2.44 | 8.64 |
| QRICH1 | 6 | 9.56 | 10.63 | 9.82 | 9.52 | 10.05 | 9.98 | 10.54 | 10.50 | 10.01 | 9.38 | 10.10 | 9.67 | 10.02 | 10.52 | 9.69 |
| QRSL1 | 1 | 11.72 | 10.65 | 9.11 | 10.50 | 9.90 | 8.77 | 10.78 | 11.08 | 9.37 | 8.12 | 11.19 | 9.80 | 9.34 | 10.93 | 8.74 |
| QSER1 | 12 | 8.95 | 9.58 | 11.40 | 10.33 | 8.38 | 8.35 | 11.61 | 10.48 | 10.70 | 10.23 | 9.27 | 10.87 | 8.36 | 11.04 | 10.46 |
| QSOX2 | 10 | 9.27 | 6.29 | 10.32 | 12.11 | 9.68 | 8.62 | 14.26 | 13.41 | 7.85 | 8.20 | 7.78 | 11.21 | 9.15 | 13.83 | 8.03 |
| QTRT1 | 3 | 9.96 | 9.42 | 7.69 | 8.98 | 9.76 | 10.89 | 9.96 | 12.19 | 10.87 | 10.29 | 9.69 | 8.33 | 10.32 | 11.08 | 10.58 |
| QTRTD1 | 8 | 8.75 | 9.90 | 9.22 | 9.16 | 10.07 | 9.49 | 11.65 | 10.65 | 11.07 | 10.03 | 9.32 | 9.19 | 9.78 | 11.15 | 10.55 |
| R3HCC1 | 2 | 8.20 | 7.90 | 10.36 | 7.58 | 10.86 | 11.04 | 13.50 | 12.96 | 8.91 | 8.69 | 8.05 | 8.97 | 10.95 | 13.23 | 8.80 |
| R3HDM1 | 4 | 9.44 | 10.71 | 9.01 | 9.09 | 11.56 | 12.06 | 11.26 | 10.73 | 8.63 | 7.51 | 10.08 | 9.05 | 11.81 | 10.99 | 8.07 |
| RAB10 | 11 | 12.17 | 8.08 | 7.90 | 10.28 | 9.40 | 9.32 | 9.76 | 10.10 | 10.78 | 12.20 | 10.13 | 9.09 | 9.36 | 9.93 | 11.49 |
| RAB11A | 1 | 6.49 | 6.86 | 10.60 | 10.72 | 11.76 | 8.26 | 8.07 | 7.63 | 11.93 | 17.68 | 6.67 | 10.66 | 10.01 | 7.85 | 14.80 |
| RAB11B | 22 | 10.90 | 7.58 | 9.16 | 10.11 | 11.22 | 10.21 | 8.86 | 8.44 | 11.40 | 12.12 | 9.24 | 9.64 | 10.72 | 8.65 | 11.76 |
| RAB11FIP1 | 10 | 8.80 | 9.23 | 11.28 | 11.05 | 9.57 | 9.78 | 11.09 | 10.30 | 9.53 | 9.38 | 9.01 | 11.16 | 9.68 | 10.69 | 9.45 |
| RAB11FIP5 | 7 | 7.57 | 8.26 | 11.14 | 10.15 | 9.42 | 8.90 | 13.16 | 10.59 | 10.78 | 10.03 | 7.92 | 10.65 | 9.16 | 11.88 | 10.40 |
| RAB12 | 9 | 10.73 | 8.20 | 9.64 | 11.74 | 9.66 | 9.39 | 8.63 | 10.09 | 10.55 | 11.37 | 9.47 | 10.69 | 9.52 | 9.36 | 10.96 |
| RAB13 | 9 | 11.54 | 8.68 | 8.24 | 8.98 | 12.34 | 12.68 | 9.52 | 8.95 | 9.14 | 9.94 | 10.11 | 8.61 | 12.51 | 9.24 | 9.54 |
| RAB14 | 22 | 11.07 | 7.39 | 8.02 | 9.75 | 10.57 | 9.86 | 8.40 | 9.02 | 12.21 | 13.69 | 9.23 | 8.89 | 10.21 | 8.71 | 12.95 |
| RAB18 | 7 | 12.68 | 7.92 | 9.44 | 11.46 | 8.56 | 7.70 | 8.36 | 8.05 | 12.47 | 13.35 | 10.30 | 10.45 | 8.13 | 8.21 | 12.91 |
| RAB1A | 19 | 10.63 | 7.88 | 9.48 | 10.86 | 9.75 | 9.18 | 9.39 | 10.41 | 10.80 | 11.61 | 9.26 | 10.17 | 9.47 | 9.90 | 11.21 |
| RAB1B | 13 | 11.38 | 7.61 | 9.63 | 12.13 | 9.94 | 10.20 | 8.97 | 8.97 | 10.51 | 10.66 | 9.49 | 10.88 | 10.07 | 8.97 | 10.58 |
| RAB21 | 10 | 11.34 | 7.95 | 8.35 | 10.09 | 10.10 | 9.86 | 9.38 | 8.99 | 11.15 | 12.80 | 9.64 | 9.22 | 9.98 | 9.18 | 11.98 |
| RAB22A | 5 | 10.32 | 7.88 | 7.89 | 9.36 | 11.30 | 11.12 | 9.75 | 9.36 | 11.10 | 11.92 | 9.10 | 8.62 | 11.21 | 9.56 | 11.51 |
| RAB23 | 5 | 9.22 | 6.96 | 11.22 | 14.70 | 7.90 | 7.80 | 8.26 | 8.20 | 13.00 | 12.75 | 8.09 | 12.96 | 7.85 | 8.23 | 12.88 |
| RAB24 | 2 | 9.93 | 9.73 | 9.11 | 9.63 | 8.67 | 9.26 | 10.46 | 9.81 | 10.90 | 12.49 | 9.83 | 9.37 | 8.97 | 10.14 | 11.69 |
| RAB27A | 5 | 11.43 | 8.36 | 7.94 | 8.86 | 11.32 | 10.48 | 9.31 | 9.32 | 10.86 | 12.13 | 9.90 | 8.40 | 10.90 | 9.32 | 11.49 |
| RAB27B | 1 | 11.07 | 7.59 | 5.62 | 8.38 | 12.21 | 12.17 | 14.72 | 14.63 | 6.55 | 7.06 | 9.33 | 7.00 | 12.19 | 14.68 | 6.81 |
| RAB2A | 6 | 10.97 | 7.08 | 7.31 | 10.67 | 8.84 | 8.79 | 11.96 | 12.78 | 9.70 | 11.89 | 9.03 | 8.99 | 8.82 | 12.37 | 10.80 |
| RAB2B | 8 | 10.57 | 8.06 | 8.33 | 9.90 | 9.31 | 9.03 | 10.94 | 11.56 | 10.48 | 11.82 | 9.31 | 9.12 | 9.17 | 11.25 | 11.15 |
| RAB30 | 1 | 12.75 | 7.80 | 9.73 | 11.40 | 9.66 | 8.73 | 7.79 | 8.67 | 10.56 | 12.90 | 10.27 | 10.57 | 9.20 | 8.23 | 11.73 |
| RAB31 | 8 | 8.99 | 7.04 | 7.71 | 7.57 | 12.94 | 11.04 | 10.24 | 15.89 | 8.96 | 9.64 | 8.02 | 7.64 | 11.99 | 13.06 | 9.30 |

| | | | | | | | | | | | | | | | | |
|---|---|---|---|---|---|---|---|---|---|---|---|---|---|---|---|---|
| RAB32 | 8 | 10.24 | 8.07 | 9.21 | 11.13 | 9.61 | 9.44 | 7.00 | 10.41 | 11.54 | 13.35 | 9.15 | 10.17 | 9.52 | 8.71 | 12.44 |
| RAB33B | 1 | 12.93 | 9.47 | 7.29 | 13.03 | 8.31 | 8.05 | 7.36 | 7.91 | 12.71 | 12.95 | 11.20 | 10.16 | 8.18 | 7.63 | 12.83 |
| RAB34 | 8 | 12.42 | 9.20 | 10.90 | 9.83 | 9.43 | 9.41 | 8.83 | 7.67 | 10.70 | 11.61 | 10.81 | 10.36 | 9.42 | 8.25 | 11.16 |
| RAB35 | 4 | 10.72 | 8.82 | 9.60 | 13.00 | 8.30 | 8.29 | 8.22 | 8.29 | 11.58 | 13.18 | 9.77 | 11.30 | 8.30 | 8.25 | 12.38 |
| RAB3D | 2 | 6.44 | 3.00 | 5.63 | 4.71 | 5.76 | 3.77 | 4.70 | 50.88 | 7.41 | 7.69 | 4.72 | 5.17 | 4.76 | 27.79 | 7.55 |
| RAB3GAP1 | 18 | 9.36 | 10.22 | 10.57 | 10.14 | 10.91 | 10.58 | 8.95 | 8.47 | 10.40 | 10.40 | 9.79 | 10.36 | 10.74 | 8.71 | 10.40 |
| RAB3GAP2 | 38 | 9.79 | 9.75 | 8.92 | 9.17 | 11.52 | 11.42 | 8.98 | 8.73 | 11.01 | 10.70 | 9.77 | 9.05 | 11.47 | 8.85 | 10.86 |
| RAB3IL1 | 2 | 7.99 | 8.24 | 12.96 | 13.40 | 8.79 | 9.92 | 9.46 | 8.21 | 10.52 | 10.50 | 8.12 | 13.18 | 9.35 | 8.84 | 10.51 |
| RAB4A | 6 | 8.96 | 8.04 | 8.19 | 8.37 | 12.09 | 11.30 | 11.82 | 11.82 | 9.21 | 10.22 | 8.50 | 8.28 | 11.69 | 11.82 | 9.71 |
| RAB5A | 8 | 10.00 | 8.81 | 7.27 | 8.59 | 10.23 | 10.59 | 9.73 | 9.81 | 11.79 | 13.18 | 9.40 | 7.93 | 10.41 | 9.77 | 12.48 |
| RAB5B | 6 | 11.26 | 8.52 | 10.26 | 9.16 | 11.99 | 10.18 | 8.96 | 6.89 | 11.04 | 11.76 | 9.89 | 9.71 | 11.08 | 7.92 | 11.40 |
| RAB5C | 12 | 11.62 | 8.29 | 10.56 | 11.32 | 9.39 | 8.62 | 8.48 | 7.85 | 11.40 | 12.48 | 9.95 | 10.94 | 9.00 | 8.17 | 11.94 |
| RAB6A | 14 | 10.50 | 7.00 | 8.57 | 10.84 | 9.67 | 8.61 | 10.83 | 10.57 | 10.89 | 12.53 | 8.75 | 9.70 | 9.14 | 10.70 | 11.71 |
| RAB7A | 17 | 11.99 | 7.67 | 9.06 | 11.56 | 9.01 | 7.99 | 9.88 | 9.13 | 11.34 | 12.38 | 9.83 | 10.31 | 8.50 | 9.50 | 11.86 |
| RAB8A | 12 | 11.70 | 8.04 | 9.17 | 10.76 | 9.32 | 8.57 | 9.58 | 11.37 | 10.37 | 11.11 | 9.87 | 9.96 | 8.95 | 10.47 | 10.74 |
| RAB8B | 2 | 13.34 | 8.67 | 11.07 | 8.13 | 9.68 | 8.67 | 11.14 | 8.65 | 10.44 | 10.22 | 11.00 | 9.60 | 9.17 | 9.90 | 10.33 |
| RAB9A | 4 | 11.08 | 7.44 | 6.74 | 8.53 | 13.85 | 13.49 | 11.32 | 11.51 | 7.66 | 8.38 | 9.26 | 7.64 | 13.67 | 11.42 | 8.02 |
| RABAC1 | 1 | 9.55 | 6.05 | 7.68 | 10.44 | 12.18 | 10.60 | 10.96 | 11.72 | 10.00 | 10.82 | 7.80 | 9.06 | 11.39 | 11.34 | 10.41 |
| RABEP1 | 23 | 10.70 | 11.60 | 11.87 | 11.56 | 8.22 | 8.31 | 9.19 | 8.38 | 10.21 | 9.96 | 11.15 | 11.71 | 8.26 | 8.78 | 10.09 |
| RABEP2 | 15 | 10.23 | 11.96 | 11.21 | 10.91 | 11.60 | 12.30 | 7.10 | 5.67 | 9.84 | 9.18 | 11.09 | 11.06 | 11.95 | 6.38 | 9.51 |
| RABEPK | 4 | 9.36 | 10.06 | 11.19 | 10.32 | 9.11 | 9.02 | 11.85 | 10.79 | 9.41 | 8.89 | 9.71 | 10.75 | 9.06 | 11.32 | 9.15 |
| RABGAP1 | 24 | 10.26 | 10.87 | 10.27 | 9.81 | 11.46 | 11.60 | 8.99 | 8.98 | 8.91 | 8.83 | 10.57 | 10.04 | 11.53 | 8.98 | 8.87 |
| RABGAP1L | 3 | 12.02 | 11.98 | 8.70 | 8.52 | 9.28 | 9.85 | 11.89 | 10.65 | 9.19 | 7.92 | 12.00 | 8.61 | 9.57 | 11.27 | 8.55 |
| RABGEF1 | 11 | 11.33 | 11.02 | 11.35 | 11.13 | 9.78 | 9.81 | 8.61 | 8.25 | 9.82 | 8.89 | 11.18 | 11.24 | 9.80 | 8.43 | 9.36 |
| RABGGTA | 8 | 10.11 | 10.70 | 8.43 | 8.51 | 14.27 | 14.82 | 8.07 | 7.39 | 9.09 | 8.61 | 10.40 | 8.47 | 14.55 | 7.73 | 8.85 |
| RABGGTB | 8 | 10.24 | 10.55 | 8.37 | 8.72 | 9.72 | 10.37 | 10.48 | 11.23 | 9.98 | 10.34 | 10.40 | 8.54 | 10.04 | 10.86 | 10.16 |
| RABIF | 1 | 9.35 | 11.55 | 7.36 | 9.51 | 14.06 | 15.68 | 6.68 | 6.36 | 10.05 | 9.40 | 10.45 | 8.44 | 14.87 | 6.52 | 9.72 |
| RABL2A | 1 | 11.51 | 10.50 | 10.70 | 11.61 | 8.82 | 9.16 | 10.08 | 9.24 | 10.03 | 8.35 | 11.01 | 11.16 | 8.99 | 9.66 | 9.19 |
| RABL3 | 8 | 9.04 | 10.03 | 9.79 | 8.66 | 12.81 | 12.45 | 10.26 | 10.78 | 8.28 | 7.90 | 9.53 | 9.22 | 12.63 | 10.52 | 8.09 |
| RABL6 | 7 | 9.49 | 11.03 | 7.72 | 7.39 | 13.22 | 14.57 | 9.26 | 10.33 | 8.63 | 8.37 | 10.26 | 7.55 | 13.90 | 9.79 | 8.50 |
| RAC1 | 17 | 6.91 | 5.21 | 10.40 | 8.01 | 7.63 | 6.59 | 6.72 | 29.75 | 9.25 | 9.53 | 6.06 | 9.20 | 7.11 | 18.24 | 9.39 |
| RACGAP1 | 21 | 9.13 | 9.85 | 10.54 | 9.67 | 10.66 | 10.16 | 10.26 | 9.70 | 9.97 | 10.07 | 9.49 | 10.11 | 10.41 | 9.98 | 10.02 |
| RAD1 | 1 | 11.43 | 10.64 | 9.57 | 7.76 | 10.10 | 10.73 | 10.99 | 11.94 | 8.25 | 8.59 | 11.04 | 8.66 | 10.42 | 11.46 | 8.42 |
| RAD17 | 3 | 9.49 | 9.13 | 7.74 | 7.05 | 10.66 | 9.05 | 11.51 | 16.28 | 9.34 | 9.75 | 9.31 | 7.39 | 9.85 | 13.90 | 9.55 |
| RAD18 | 6 | 9.38 | 10.77 | 9.50 | 9.29 | 8.79 | 9.78 | 11.67 | 10.98 | 10.45 | 9.39 | 10.08 | 9.39 | 9.29 | 11.33 | 9.92 |
| RAD21 | 25 | 8.50 | 9.46 | 9.04 | 8.43 | 9.95 | 9.68 | 11.68 | 16.31 | 8.78 | 8.19 | 8.98 | 8.73 | 9.81 | 13.99 | 8.48 |
| RAD23A | 6 | 10.03 | 10.75 | 7.42 | 7.20 | 11.87 | 11.83 | 12.04 | 10.34 | 9.15 | 9.38 | 10.39 | 7.31 | 11.85 | 11.19 | 9.26 |
| RAD23B | 29 | 9.38 | 10.80 | 9.58 | 9.57 | 11.18 | 10.99 | 9.63 | 8.90 | 10.00 | 9.97 | 10.09 | 9.58 | 11.09 | 9.27 | 9.98 |
| RAD50 | 63 | 9.18 | 9.78 | 9.50 | 8.63 | 9.86 | 10.01 | 13.30 | 12.41 | 8.93 | 8.40 | 9.48 | 9.07 | 9.93 | 12.86 | 8.67 |
| RAD51AP1 | 3 | 9.49 | 11.21 | 9.65 | 9.62 | 7.94 | 7.73 | 10.05 | 9.20 | 13.23 | 11.89 | 10.35 | 9.63 | 7.83 | 9.62 | 12.56 |
| RAD52 | 1 | 10.93 | 13.37 | 6.92 | 9.37 | 10.74 | 11.48 | 9.28 | 9.92 | 9.15 | 8.84 | 12.15 | 8.14 | 11.11 | 9.60 | 8.99 |
| RAD54B | 1 | 9.61 | 9.72 | 5.84 | 8.25 | 9.15 | 8.15 | 15.06 | 12.36 | 11.37 | 10.50 | 9.66 | 7.05 | 8.65 | 13.71 | 10.93 |

| | | | | | | | | | | | | | | | | |
|---|---|---|---|---|---|---|---|---|---|---|---|---|---|---|---|---|
| RAE1 | 15 | 9.85 | 10.43 | 10.71 | 10.34 | 9.13 | 9.72 | 11.02 | 10.60 | 9.20 | 9.01 | 10.14 | 10.52 | 9.42 | 10.81 | 9.11 |
| RAET1G | 2 | 10.25 | 9.07 | 4.66 | 5.43 | 20.51 | 21.14 | 6.91 | 6.83 | 7.73 | 7.47 | 9.66 | 5.05 | 20.82 | 6.87 | 7.60 |
| RAF1 | 3 | 9.53 | 10.48 | 9.73 | 8.84 | 11.36 | 9.98 | 10.50 | 8.65 | 10.48 | 10.44 | 10.01 | 9.28 | 10.67 | 9.58 | 10.46 |
| RAI1 | 7 | 10.33 | 10.95 | 11.26 | 11.13 | 8.00 | 8.37 | 10.41 | 9.34 | 10.30 | 9.92 | 10.64 | 11.19 | 8.18 | 9.87 | 10.11 |
| RAI14 | 32 | 9.83 | 10.89 | 14.13 | 12.72 | 8.72 | 8.63 | 7.55 | 6.58 | 10.71 | 10.25 | 10.36 | 13.42 | 8.68 | 7.06 | 10.48 |
| RALA | 8 | 9.65 | 8.10 | 11.90 | 11.19 | 7.23 | 6.72 | 11.36 | 8.90 | 11.52 | 13.42 | 8.87 | 11.55 | 6.98 | 10.13 | 12.47 |
| RALB | 8 | 10.58 | 8.46 | 10.70 | 11.73 | 7.68 | 7.40 | 10.54 | 9.67 | 11.31 | 11.94 | 9.52 | 11.22 | 7.54 | 10.10 | 11.62 |
| RALBP1 | 4 | 8.39 | 9.97 | 12.01 | 11.65 | 9.80 | 10.63 | 9.75 | 9.85 | 9.07 | 8.87 | 9.18 | 11.83 | 10.22 | 9.80 | 8.97 |
| RALGAPA1 | 5 | 10.23 | 10.47 | 8.17 | 8.10 | 13.17 | 12.82 | 11.68 | 11.15 | 7.46 | 6.74 | 10.35 | 8.14 | 13.00 | 11.42 | 7.10 |
| RALGAPB | 4 | 10.82 | 10.48 | 8.77 | 10.52 | 11.39 | 12.18 | 12.56 | 12.70 | 5.46 | 5.12 | 10.65 | 9.65 | 11.78 | 12.63 | 5.29 |
| RALGPS2 | 1 | 7.81 | 8.09 | 12.46 | 8.93 | 10.55 | 12.30 | 12.14 | 9.51 | 8.81 | 9.40 | 7.95 | 10.70 | 11.43 | 10.83 | 9.10 |
| RALY | 33 | 9.63 | 10.32 | 10.89 | 11.08 | 9.46 | 10.05 | 10.57 | 9.99 | 9.03 | 8.98 | 9.98 | 10.98 | 9.76 | 10.28 | 9.00 |
| RAN | 47 | 9.06 | 10.69 | 10.24 | 8.74 | 9.94 | 9.30 | 10.08 | 12.93 | 9.62 | 9.41 | 9.87 | 9.49 | 9.62 | 11.50 | 9.52 |
| RANBP1 | 6 | 10.20 | 11.50 | 11.38 | 11.33 | 7.86 | 8.04 | 11.42 | 6.76 | 11.08 | 10.44 | 10.85 | 11.35 | 7.95 | 9.09 | 10.76 |
| RANBP2 | 122 | 9.42 | 9.89 | 10.98 | 11.13 | 9.20 | 9.18 | 10.67 | 10.81 | 9.52 | 9.20 | 9.65 | 11.06 | 9.19 | 10.74 | 9.36 |
| RANBP3 | 19 | 9.62 | 10.39 | 10.59 | 9.39 | 10.53 | 10.28 | 10.13 | 8.83 | 10.33 | 9.91 | 10.00 | 9.99 | 10.41 | 9.48 | 10.12 |
| RANBP6 | 6 | 9.87 | 10.13 | 6.35 | 7.72 | 12.09 | 12.11 | 10.27 | 10.85 | 10.27 | 10.34 | 10.00 | 7.03 | 12.10 | 10.56 | 10.31 |
| RANBP9 | 4 | 10.11 | 12.02 | 7.83 | 8.06 | 10.36 | 11.54 | 10.27 | 9.36 | 10.61 | 9.85 | 11.06 | 7.94 | 10.95 | 9.82 | 10.23 |
| RANGAP1 | 40 | 8.16 | 9.06 | 14.34 | 13.98 | 8.05 | 8.20 | 9.44 | 10.84 | 8.85 | 9.08 | 8.61 | 14.16 | 8.13 | 10.14 | 8.96 |
| RANGRF | 1 | 10.89 | 10.72 | 8.32 | 8.43 | 11.95 | 13.10 | 11.08 | 6.42 | 9.99 | 9.11 | 10.80 | 8.37 | 12.53 | 8.75 | 9.55 |
| RAP1A | 10 | 12.32 | 8.31 | 9.97 | 12.21 | 8.54 | 8.32 | 8.77 | 8.14 | 11.34 | 12.09 | 10.32 | 11.09 | 8.43 | 8.46 | 11.71 |
| RAP1B | 6 | 9.78 | 7.86 | 8.87 | 12.23 | 9.37 | 9.53 | 8.13 | 9.25 | 11.59 | 13.39 | 8.82 | 10.55 | 9.45 | 8.69 | 12.49 |
| RAP1GAP | 1 | 12.65 | 13.95 | 5.92 | 5.02 | 6.27 | 6.36 | 16.24 | 12.22 | 9.82 | 11.54 | 13.30 | 5.47 | 6.32 | 14.23 | 10.68 |
| RAP1GDS1 | 17 | 9.49 | 10.48 | 9.42 | 9.39 | 11.75 | 12.19 | 8.97 | 9.01 | 9.81 | 9.49 | 9.98 | 9.41 | 11.97 | 8.99 | 9.65 |
| RAP2A | 4 | 11.42 | 9.26 | 7.94 | 9.86 | 9.99 | 10.63 | 10.73 | 10.45 | 9.92 | 9.79 | 10.34 | 8.90 | 10.31 | 10.59 | 9.86 |
| RAPGEF1 | 1 | 14.16 | 12.27 | 6.85 | 8.37 | 9.57 | 11.15 | 10.02 | 8.64 | 10.36 | 8.61 | 13.22 | 7.61 | 10.36 | 9.33 | 9.49 |
| RAPGEF2 | 4 | 13.82 | 9.16 | 9.68 | 8.59 | 11.68 | 12.20 | 8.31 | 8.12 | 9.49 | 8.95 | 11.49 | 9.14 | 11.94 | 8.21 | 9.22 |
| RAPGEF6 | 9 | 9.83 | 9.75 | 8.72 | 8.64 | 11.03 | 11.07 | 11.59 | 10.46 | 9.77 | 9.15 | 9.79 | 8.68 | 11.05 | 11.02 | 9.46 |
| RAPH1 | 20 | 9.48 | 9.21 | 11.82 | 12.10 | 10.05 | 10.52 | 8.52 | 8.31 | 10.51 | 9.49 | 9.34 | 11.96 | 10.28 | 8.42 | 10.00 |
| RARS | 54 | 9.27 | 9.56 | 9.41 | 9.30 | 9.53 | 9.46 | 10.00 | 14.62 | 9.48 | 9.38 | 9.41 | 9.36 | 9.50 | 12.31 | 9.43 |
| RARS2 | 5 | 10.45 | 10.99 | 8.53 | 9.77 | 10.13 | 10.22 | 9.51 | 9.73 | 10.41 | 10.26 | 10.72 | 9.15 | 10.18 | 9.62 | 10.33 |
| RASA1 | 8 | 8.09 | 8.27 | 11.67 | 12.31 | 9.78 | 9.57 | 10.46 | 10.73 | 9.89 | 9.24 | 8.18 | 11.99 | 9.67 | 10.59 | 9.56 |
| RASA3 | 1 | 6.61 | 8.68 | 14.32 | 12.38 | 10.68 | 8.48 | 11.64 | 10.81 | 7.56 | 8.82 | 7.65 | 13.35 | 9.58 | 11.23 | 8.19 |
| RASAL2 | 11 | 8.61 | 9.32 | 12.09 | 12.22 | 8.89 | 9.18 | 9.40 | 9.21 | 10.92 | 10.16 | 8.96 | 12.15 | 9.04 | 9.31 | 10.54 |
| RASSF1 | 3 | 10.23 | 11.48 | 8.12 | 7.83 | 11.57 | 13.32 | 9.86 | 9.37 | 9.42 | 8.79 | 10.86 | 7.98 | 12.45 | 9.61 | 9.11 |
| RASSF7 | 1 | 11.83 | 14.05 | 7.64 | 7.08 | 9.48 | 9.88 | 8.63 | 7.58 | 11.74 | 12.10 | 12.94 | 7.36 | 9.68 | 8.11 | 11.92 |
| RASSF8 | 9 | 8.30 | 9.01 | 12.28 | 12.70 | 8.78 | 9.71 | 10.69 | 9.55 | 9.68 | 9.29 | 8.66 | 12.49 | 9.25 | 10.12 | 9.48 |
| RAVER1 | 14 | 8.50 | 9.53 | 11.09 | 9.94 | 11.31 | 10.99 | 9.76 | 8.65 | 10.47 | 9.76 | 9.01 | 10.51 | 11.15 | 9.21 | 10.12 |
| RAVER2 | 1 | 9.15 | 12.46 | 9.34 | 8.04 | 9.68 | 11.27 | 10.90 | 9.01 | 9.77 | 10.38 | 10.80 | 8.69 | 10.48 | 9.95 | 10.08 |
| RB1 | 2 | 9.65 | 10.93 | 8.42 | 9.00 | 12.41 | 13.31 | 11.33 | 9.31 | 7.32 | 8.32 | 10.29 | 8.71 | 12.86 | 10.32 | 7.82 |
| RB1CC1 | 3 | 7.83 | 7.55 | 9.15 | 8.97 | 9.17 | 7.74 | 13.05 | 22.68 | 7.04 | 6.82 | 7.69 | 9.06 | 8.45 | 17.87 | 6.93 |
| RBBP4 | 26 | 10.99 | 11.36 | 8.65 | 8.16 | 9.95 | 10.15 | 10.38 | 10.71 | 10.28 | 9.37 | 11.18 | 8.40 | 10.05 | 10.55 | 9.82 |

| | | | | | | | | | | | | | | | | |
|---|---|---|---|---|---|---|---|---|---|---|---|---|---|---|---|---|
| RBBP5 | 13 | 9.53 | 10.45 | 9.27 | 8.51 | 10.34 | 10.70 | 10.75 | 11.47 | 9.82 | 9.15 | 9.99 | 8.89 | 10.52 | 11.11 | 9.49 |
| RBBP6 | 20 | 9.17 | 11.16 | 9.05 | 8.39 | 11.41 | 11.81 | 10.39 | 9.51 | 9.90 | 9.21 | 10.17 | 8.72 | 11.61 | 9.95 | 9.55 |
| RBBP7 | 16 | 9.36 | 9.09 | 10.21 | 7.97 | 12.18 | 11.44 | 10.16 | 13.94 | 8.00 | 7.65 | 9.23 | 9.09 | 11.81 | 12.05 | 7.82 |
| RBBP8 | 1 | 8.70 | 12.18 | 10.92 | 9.58 | 10.87 | 10.45 | 7.93 | 6.87 | 12.30 | 10.19 | 10.44 | 10.25 | 10.66 | 7.40 | 11.25 |
| RBFOX2 | 4 | 8.96 | 10.82 | 10.91 | 10.93 | 7.68 | 7.84 | 10.69 | 10.13 | 11.39 | 10.65 | 9.89 | 10.92 | 7.76 | 10.41 | 11.02 |
| RBM10 | 24 | 9.50 | 10.27 | 10.40 | 9.83 | 9.81 | 9.83 | 11.65 | 10.33 | 9.48 | 8.89 | 9.89 | 10.11 | 9.82 | 10.99 | 9.19 |
| RBM12 | 25 | 9.43 | 9.89 | 10.38 | 10.43 | 9.54 | 9.77 | 10.93 | 10.23 | 9.90 | 9.50 | 9.66 | 10.41 | 9.65 | 10.58 | 9.70 |
| RBM12B | 18 | 9.28 | 10.64 | 10.29 | 9.39 | 10.11 | 10.49 | 12.30 | 11.16 | 8.60 | 7.74 | 9.96 | 9.84 | 10.30 | 11.73 | 8.17 |
| RBM14 | 39 | 8.76 | 10.02 | 10.85 | 9.01 | 10.22 | 9.91 | 11.23 | 12.24 | 9.00 | 8.76 | 9.39 | 9.93 | 10.06 | 11.74 | 8.88 |
| RBM15 | 25 | 9.39 | 10.61 | 9.19 | 9.06 | 10.86 | 11.31 | 10.95 | 10.36 | 9.32 | 8.94 | 10.00 | 9.12 | 11.08 | 10.66 | 9.13 |
| RBM15B | 8 | 9.13 | 9.66 | 10.26 | 8.91 | 10.25 | 9.81 | 10.26 | 9.92 | 11.07 | 10.73 | 9.39 | 9.58 | 10.03 | 10.09 | 10.90 |
| RBM17 | 21 | 8.72 | 10.38 | 11.31 | 10.62 | 9.10 | 9.23 | 10.81 | 10.29 | 10.09 | 9.45 | 9.55 | 10.96 | 9.17 | 10.55 | 9.77 |
| RBM19 | 10 | 9.69 | 9.87 | 12.22 | 11.35 | 7.48 | 7.88 | 11.45 | 10.43 | 10.38 | 9.27 | 9.78 | 11.78 | 7.68 | 10.94 | 9.82 |
| RBM22 | 13 | 10.27 | 10.23 | 8.95 | 8.01 | 9.06 | 9.34 | 11.25 | 14.49 | 9.53 | 8.87 | 10.25 | 8.48 | 9.20 | 12.87 | 9.20 |
| RBM23 | 3 | 9.88 | 11.58 | 13.03 | 12.43 | 8.37 | 8.43 | 8.21 | 7.50 | 9.96 | 10.60 | 10.73 | 12.73 | 8.40 | 7.86 | 10.28 |
| RBM25 | 31 | 9.87 | 10.27 | 10.16 | 8.89 | 9.24 | 8.97 | 11.17 | 11.25 | 10.60 | 9.58 | 10.07 | 9.52 | 9.11 | 11.21 | 10.09 |
| RBM26 | 23 | 10.11 | 11.09 | 9.84 | 9.26 | 9.70 | 9.74 | 11.65 | 10.75 | 9.18 | 8.68 | 10.60 | 9.55 | 9.72 | 11.20 | 8.93 |
| RBM27 | 18 | 9.15 | 9.78 | 10.00 | 9.11 | 8.55 | 8.99 | 13.00 | 11.76 | 10.40 | 9.28 | 9.46 | 9.55 | 8.77 | 12.38 | 9.84 |
| RBM28 | 20 | 11.05 | 8.87 | 9.57 | 11.75 | 6.67 | 6.58 | 10.59 | 12.13 | 9.28 | 13.51 | 9.96 | 10.66 | 6.63 | 11.36 | 11.40 |
| RBM3 | 9 | 8.05 | 8.30 | 9.86 | 7.60 | 15.04 | 14.52 | 12.92 | 10.15 | 7.13 | 6.43 | 8.18 | 8.73 | 14.78 | 11.53 | 6.78 |
| RBM33 | 4 | 9.21 | 10.96 | 11.63 | 12.17 | 8.27 | 8.99 | 9.90 | 9.76 | 9.57 | 9.54 | 10.09 | 11.90 | 8.63 | 9.83 | 9.56 |
| RBM34 | 11 | 10.70 | 10.25 | 7.79 | 8.49 | 10.53 | 10.56 | 11.10 | 10.30 | 9.54 | 10.75 | 10.47 | 8.14 | 10.54 | 10.70 | 10.15 |
| RBM39 | 32 | 7.30 | 7.71 | 11.46 | 8.88 | 10.78 | 10.35 | 11.26 | 14.32 | 8.78 | 9.16 | 7.51 | 10.17 | 10.56 | 12.79 | 8.97 |
| RBM4 | 17 | 9.82 | 10.93 | 10.63 | 9.43 | 10.69 | 10.70 | 9.96 | 8.60 | 9.79 | 9.46 | 10.38 | 10.03 | 10.69 | 9.28 | 9.62 |
| RBM41 | 2 | 10.01 | 10.88 | 11.15 | 11.68 | 10.53 | 10.91 | 12.92 | 11.93 | 5.50 | 4.50 | 10.45 | 11.41 | 10.72 | 12.42 | 5.00 |
| RBM42 | 9 | 9.69 | 10.40 | 10.28 | 9.56 | 9.57 | 9.59 | 11.65 | 10.68 | 9.16 | 9.41 | 10.04 | 9.92 | 9.58 | 11.16 | 9.29 |
| RBM45 | 3 | 9.96 | 11.39 | 9.29 | 8.56 | 10.57 | 10.81 | 11.46 | 11.01 | 8.86 | 8.08 | 10.67 | 8.92 | 10.69 | 11.24 | 8.47 |
| RBM47 | 3 | 8.89 | 8.37 | 6.70 | 8.04 | 13.91 | 13.06 | 13.56 | 13.32 | 7.15 | 6.99 | 8.63 | 7.37 | 13.48 | 13.44 | 7.07 |
| RBM4B | 1 | 10.72 | 11.46 | 10.11 | 7.84 | 10.51 | 9.68 | 12.21 | 7.96 | 10.55 | 8.96 | 11.09 | 8.98 | 10.09 | 10.08 | 9.76 |
| RBM5 | 13 | 9.99 | 10.28 | 10.99 | 10.32 | 9.07 | 8.81 | 10.86 | 9.92 | 10.26 | 9.49 | 10.13 | 10.66 | 8.94 | 10.39 | 9.88 |
| RBM6 | 15 | 9.45 | 11.14 | 9.76 | 9.28 | 9.55 | 10.17 | 11.81 | 11.22 | 9.16 | 8.47 | 10.29 | 9.52 | 9.86 | 11.51 | 8.81 |
| RBM7 | 2 | 7.83 | 10.61 | 12.01 | 9.23 | 10.59 | 9.06 | 10.88 | 8.89 | 10.76 | 10.15 | 9.22 | 10.62 | 9.82 | 9.89 | 10.45 |
| RBM8A | 13 | 10.41 | 11.28 | 8.84 | 8.59 | 10.22 | 10.39 | 11.17 | 9.99 | 9.72 | 9.39 | 10.85 | 8.72 | 10.30 | 10.58 | 9.56 |
| RBMS1 | 3 | 9.01 | 10.04 | 12.63 | 10.06 | 8.20 | 8.59 | 9.67 | 8.41 | 12.11 | 11.28 | 9.52 | 11.35 | 8.40 | 9.04 | 11.70 |
| RBMS2 | 5 | 8.84 | 9.94 | 13.17 | 12.77 | 8.06 | 8.11 | 8.07 | 6.90 | 12.26 | 11.88 | 9.39 | 12.97 | 8.09 | 7.49 | 12.07 |
| RBMX | 34 | 9.53 | 10.83 | 9.17 | 8.59 | 11.01 | 11.49 | 11.93 | 10.85 | 8.40 | 8.19 | 10.18 | 8.88 | 11.25 | 11.39 | 8.30 |
| RBMX2 | 4 | 10.07 | 10.47 | 10.10 | 9.06 | 10.31 | 10.84 | 10.72 | 10.43 | 9.80 | 8.21 | 10.27 | 9.58 | 10.58 | 10.57 | 9.00 |
| RBMXL3 | 4 | 9.88 | 10.81 | 9.35 | 11.23 | 10.33 | 11.15 | 8.73 | 8.25 | 10.17 | 10.10 | 10.35 | 10.29 | 10.74 | 8.49 | 10.13 |
| RBPJ | 5 | 9.18 | 9.83 | 11.57 | 9.29 | 9.40 | 9.33 | 12.16 | 10.54 | 9.04 | 9.65 | 9.50 | 10.43 | 9.37 | 11.35 | 9.35 |
| RBPMS | 4 | 6.72 | 8.52 | 17.08 | 16.88 | 9.53 | 8.41 | 9.45 | 6.84 | 7.33 | 9.22 | 7.62 | 16.98 | 8.97 | 8.15 | 8.27 |
| RC3H1 | 2 | 7.71 | 9.53 | 9.61 | 11.55 | 9.16 | 10.71 | 11.41 | 10.96 | 9.21 | 10.15 | 8.62 | 10.58 | 9.94 | 11.18 | 9.68 |
| RC3H2 | 4 | 9.74 | 11.65 | 8.33 | 8.26 | 11.78 | 13.41 | 9.63 | 9.27 | 9.26 | 8.68 | 10.69 | 8.29 | 12.59 | 9.45 | 8.97 |

| | | | | | | | | | | | | | | | | |
|---|---|---|---|---|---|---|---|---|---|---|---|---|---|---|---|---|
| RCC1 | 20 | 8.68 | 9.58 | 10.03 | 10.01 | 7.35 | 7.95 | 11.92 | 15.86 | 9.30 | 9.31 | 9.13 | 10.02 | 7.65 | 13.89 | 9.31 |
| RCC2 | 40 | 8.51 | 9.07 | 8.62 | 7.63 | 8.60 | 8.45 | 12.57 | 19.12 | 8.95 | 8.49 | 8.79 | 8.13 | 8.52 | 15.84 | 8.72 |
| RCCD1 | 2 | 9.82 | 11.19 | 7.53 | 6.89 | 10.89 | 10.78 | 11.29 | 10.48 | 11.14 | 9.99 | 10.50 | 7.21 | 10.83 | 10.89 | 10.57 |
| RCL1 | 9 | 9.19 | 9.99 | 10.76 | 10.82 | 8.63 | 8.80 | 11.57 | 11.82 | 9.07 | 9.35 | 9.59 | 10.79 | 8.71 | 11.69 | 9.21 |
| RCN1 | 26 | 10.31 | 9.85 | 12.68 | 10.56 | 11.37 | 11.42 | 10.00 | 6.60 | 8.53 | 8.68 | 10.08 | 11.62 | 11.40 | 8.30 | 8.61 |
| RCN2 | 17 | 8.66 | 8.94 | 9.02 | 8.56 | 13.59 | 13.41 | 9.63 | 7.52 | 10.56 | 10.10 | 8.80 | 8.79 | 13.50 | 8.58 | 10.33 |
| RCN3 | 2 | 4.87 | 4.52 | 7.74 | 7.21 | 6.57 | 5.92 | 30.90 | 21.27 | 5.77 | 5.24 | 4.69 | 7.47 | 6.25 | 26.08 | 5.50 |
| RCOR1 | 16 | 9.66 | 10.22 | 8.92 | 8.36 | 10.62 | 10.72 | 11.87 | 10.64 | 9.60 | 9.39 | 9.94 | 8.64 | 10.67 | 11.25 | 9.50 |
| RCOR3 | 4 | 10.19 | 10.95 | 8.83 | 7.84 | 12.07 | 11.97 | 10.84 | 9.57 | 9.05 | 8.69 | 10.57 | 8.34 | 12.02 | 10.20 | 8.87 |
| RDH11 | 6 | 12.28 | 7.58 | 8.68 | 13.41 | 8.10 | 7.77 | 9.51 | 9.88 | 10.22 | 12.56 | 9.93 | 11.05 | 7.94 | 9.69 | 11.39 |
| RDH13 | 6 | 9.10 | 8.52 | 11.38 | 10.38 | 10.05 | 9.45 | 13.26 | 11.85 | 7.84 | 8.17 | 8.81 | 10.88 | 9.75 | 12.55 | 8.01 |
| RDH14 | 7 | 9.44 | 9.21 | 8.98 | 9.70 | 9.93 | 10.60 | 11.50 | 11.30 | 9.09 | 10.24 | 9.33 | 9.34 | 10.27 | 11.40 | 9.67 |
| RDX | 43 | 12.69 | 13.27 | 7.28 | 6.69 | 13.77 | 13.83 | 7.62 | 7.94 | 8.81 | 8.10 | 12.98 | 6.98 | 13.80 | 7.78 | 8.46 |
| RECQL | 52 | 9.01 | 9.20 | 10.05 | 9.46 | 9.60 | 9.63 | 10.92 | 12.78 | 9.86 | 9.49 | 9.11 | 9.76 | 9.61 | 11.85 | 9.67 |
| RECQL4 | 1 | 10.99 | 11.05 | 6.65 | 6.61 | 8.15 | 9.09 | 12.87 | 12.08 | 12.01 | 10.50 | 11.02 | 6.63 | 8.62 | 12.48 | 11.25 |
| RECQL5 | 2 | 9.66 | 9.72 | 11.23 | 9.54 | 9.64 | 10.24 | 8.96 | 7.77 | 12.08 | 11.15 | 9.69 | 10.38 | 9.94 | 8.37 | 11.61 |
| REEP4 | 1 | 10.93 | 6.60 | 7.09 | 10.29 | 8.53 | 7.92 | 14.19 | 11.92 | 11.17 | 11.35 | 8.77 | 8.69 | 8.22 | 13.06 | 11.26 |
| REEP5 | 7 | 12.58 | 7.09 | 8.10 | 11.20 | 11.76 | 9.79 | 11.31 | 10.23 | 8.09 | 9.84 | 9.83 | 9.65 | 10.78 | 10.77 | 8.97 |
| REL | 5 | 9.00 | 9.96 | 11.29 | 8.91 | 12.70 | 11.30 | 8.43 | 10.58 | 8.93 | 8.89 | 9.48 | 10.10 | 12.00 | 9.51 | 8.91 |
| RELA | 11 | 9.09 | 10.51 | 11.88 | 11.94 | 9.30 | 9.63 | 9.22 | 9.57 | 9.16 | 9.71 | 9.80 | 11.91 | 9.46 | 9.39 | 9.43 |
| REPIN1 | 2 | 10.14 | 8.82 | 10.30 | 12.18 | 7.06 | 6.68 | 12.11 | 13.00 | 9.01 | 10.71 | 9.48 | 11.24 | 6.87 | 12.55 | 9.86 |
| REPS1 | 3 | 8.78 | 9.59 | 11.66 | 11.06 | 10.37 | 10.68 | 10.74 | 9.83 | 8.88 | 8.42 | 9.18 | 11.36 | 10.53 | 10.28 | 8.65 |
| RER1 | 5 | 11.81 | 7.49 | 6.93 | 8.19 | 10.62 | 8.92 | 11.07 | 9.45 | 11.41 | 14.11 | 9.65 | 7.56 | 9.77 | 10.26 | 12.76 |
| RETSAT | 4 | 11.54 | 7.90 | 7.20 | 9.78 | 9.40 | 9.25 | 8.33 | 7.72 | 12.62 | 16.25 | 9.72 | 8.49 | 9.33 | 8.03 | 14.44 |
| REXO2 | 7 | 9.28 | 9.26 | 13.62 | 12.22 | 8.80 | 8.90 | 10.09 | 8.67 | 10.01 | 9.14 | 9.27 | 12.92 | 8.85 | 9.38 | 9.58 |
| REXO4 | 15 | 9.55 | 10.01 | 11.11 | 11.05 | 8.17 | 8.20 | 11.03 | 9.88 | 10.39 | 10.60 | 9.78 | 11.08 | 8.19 | 10.45 | 10.50 |
| RFC1 | 22 | 10.33 | 10.54 | 10.06 | 10.23 | 8.50 | 8.62 | 10.92 | 10.12 | 9.83 | 10.85 | 10.44 | 10.14 | 8.56 | 10.52 | 10.34 |
| RFC2 | 17 | 10.11 | 10.38 | 9.85 | 8.64 | 9.65 | 9.72 | 11.03 | 10.36 | 10.44 | 9.82 | 10.25 | 9.25 | 9.68 | 10.70 | 10.13 |
| RFC3 | 7 | 9.74 | 10.75 | 9.90 | 9.77 | 9.62 | 10.48 | 10.60 | 10.60 | 9.28 | 9.27 | 10.25 | 9.83 | 10.05 | 10.60 | 9.27 |
| RFC4 | 14 | 9.32 | 10.37 | 9.95 | 9.71 | 9.17 | 9.21 | 11.16 | 12.34 | 9.38 | 9.40 | 9.84 | 9.83 | 9.19 | 11.75 | 9.39 |
| RFC5 | 21 | 9.97 | 10.69 | 9.46 | 8.73 | 9.63 | 9.78 | 10.60 | 11.37 | 10.18 | 9.60 | 10.33 | 9.09 | 9.70 | 10.99 | 9.89 |
| RFK | 1 | 11.39 | 12.72 | 12.64 | 5.79 | 12.69 | 9.19 | 9.67 | 4.86 | 11.31 | 9.74 | 12.05 | 9.21 | 10.94 | 7.27 | 10.53 |
| RFTN1 | 1 | 11.12 | 9.87 | 27.53 | 28.02 | 2.54 | 1.99 | 1.45 | 1.17 | 8.13 | 8.18 | 10.49 | 27.77 | 2.27 | 1.31 | 8.16 |
| RFWD2 | 1 | 7.81 | 9.79 | 10.10 | 10.15 | 12.13 | 12.51 | 10.49 | 8.02 | 8.26 | 10.75 | 8.80 | 10.12 | 12.32 | 9.26 | 9.50 |
| RFX1 | 7 | 9.50 | 10.76 | 9.50 | 9.54 | 10.69 | 10.86 | 11.02 | 10.73 | 8.45 | 8.95 | 10.13 | 9.52 | 10.77 | 10.87 | 8.70 |
| RFX5 | 2 | 9.53 | 11.06 | 9.25 | 10.05 | 12.34 | 12.86 | 8.74 | 9.12 | 8.81 | 8.23 | 10.30 | 9.65 | 12.60 | 8.93 | 8.52 |
| RFX7 | 1 | 7.96 | 11.38 | 12.93 | 10.43 | 9.00 | 11.60 | 6.83 | 7.51 | 12.38 | 9.98 | 9.67 | 11.68 | 10.30 | 7.17 | 11.18 |
| RGS10 | 2 | 7.83 | 9.27 | 14.20 | 13.58 | 8.22 | 8.24 | 9.61 | 8.85 | 10.08 | 10.14 | 8.55 | 13.89 | 8.23 | 9.23 | 10.11 |
| RGS19 | 1 | 9.30 | 10.79 | 8.63 | 14.42 | 7.26 | 10.10 | 8.62 | 8.98 | 10.56 | 11.34 | 10.05 | 11.53 | 8.68 | 8.80 | 10.95 |
| RGS3 | 1 | 11.71 | 12.93 | 7.57 | 5.94 | 11.45 | 10.43 | 2.42 | 2.37 | 17.48 | 17.71 | 12.32 | 6.76 | 10.94 | 2.39 | 17.60 |
| RHEB | 19 | 9.60 | 10.31 | 13.93 | 13.55 | 6.78 | 6.68 | 8.59 | 7.87 | 11.16 | 11.53 | 9.95 | 13.74 | 6.73 | 8.23 | 11.35 |
| RHOA | 6 | 10.79 | 7.03 | 9.02 | 8.53 | 8.11 | 6.57 | 7.57 | 20.82 | 10.74 | 10.82 | 8.91 | 8.77 | 7.34 | 14.19 | 10.78 |

| | | | | | | | | | | | | | | | | |
|---|---|---|---|---|---|---|---|---|---|---|---|---|---|---|---|---|
| RHOB | 6 | 10.77 | 8.84 | 10.09 | 11.04 | 9.50 | 9.65 | 8.81 | 8.50 | 10.87 | 11.92 | 9.81 | 10.57 | 9.58 | 8.65 | 11.39 |
| RHOF | 1 | 10.82 | 7.24 | 6.95 | 8.99 | 9.68 | 9.60 | 8.95 | 8.11 | 14.44 | 15.24 | 9.03 | 7.97 | 9.64 | 8.53 | 14.84 |
| RHOG | 10 | 10.37 | 7.94 | 11.47 | 15.03 | 7.83 | 7.75 | 8.85 | 7.85 | 10.77 | 12.14 | 9.16 | 13.25 | 7.79 | 8.35 | 11.45 |
| RHOT1 | 4 | 13.89 | 8.01 | 8.85 | 11.70 | 8.07 | 7.11 | 9.05 | 8.17 | 11.65 | 13.50 | 10.95 | 10.28 | 7.59 | 8.61 | 12.58 |
| RHOT2 | 6 | 12.83 | 8.12 | 7.25 | 9.68 | 9.85 | 8.15 | 8.96 | 9.00 | 12.45 | 13.72 | 10.47 | 8.46 | 9.00 | 8.98 | 13.08 |
| RIC8A | 7 | 9.99 | 9.52 | 10.49 | 9.03 | 10.38 | 10.08 | 8.49 | 11.70 | 10.39 | 9.92 | 9.76 | 9.76 | 10.23 | 10.10 | 10.16 |
| RIC8B | 1 | 9.40 | 8.38 | 10.09 | 8.91 | 9.69 | 10.81 | 11.91 | 11.02 | 10.60 | 9.19 | 8.89 | 9.50 | 10.25 | 11.47 | 9.90 |
| RICTOR | 13 | 8.76 | 9.49 | 8.77 | 8.31 | 11.06 | 10.87 | 11.34 | 11.83 | 9.92 | 9.65 | 9.13 | 8.54 | 10.96 | 11.58 | 9.79 |
| RIF1 | 50 | 8.79 | 8.44 | 10.40 | 11.57 | 8.96 | 9.05 | 11.50 | 11.03 | 10.06 | 10.20 | 8.62 | 10.98 | 9.00 | 11.26 | 10.13 |
| RILPL1 | 6 | 10.42 | 11.76 | 8.87 | 8.70 | 6.84 | 7.11 | 13.49 | 11.54 | 10.80 | 10.46 | 11.09 | 8.78 | 6.98 | 12.52 | 10.63 |
| RIMKLB | 1 | 7.36 | 7.08 | 15.12 | 14.67 | 16.23 | 14.63 | 5.25 | 3.76 | 7.44 | 8.44 | 7.22 | 14.90 | 15.43 | 4.50 | 7.94 |
| RIN1 | 1 | 6.22 | 6.44 | 12.60 | 11.65 | 8.82 | 8.17 | 11.01 | 10.20 | 12.96 | 11.94 | 6.33 | 12.12 | 8.49 | 10.60 | 12.45 |
| RING1 | 6 | 10.09 | 11.50 | 8.52 | 8.22 | 10.00 | 12.36 | 10.49 | 9.46 | 9.57 | 9.78 | 10.80 | 8.37 | 11.18 | 9.97 | 9.67 |
| RINT1 | 5 | 9.80 | 9.98 | 10.92 | 10.57 | 9.64 | 9.19 | 9.78 | 9.56 | 10.72 | 9.83 | 9.89 | 10.75 | 9.41 | 9.67 | 10.27 |
| RIOK1 | 3 | 9.80 | 10.38 | 10.91 | 10.80 | 7.92 | 8.28 | 11.98 | 11.65 | 9.35 | 8.94 | 10.09 | 10.85 | 8.10 | 11.82 | 9.14 |
| RIOK2 | 6 | 8.91 | 9.47 | 9.31 | 9.15 | 9.07 | 9.06 | 14.95 | 13.49 | 8.28 | 8.31 | 9.19 | 9.23 | 9.06 | 14.22 | 8.29 |
| RIOK3 | 3 | 10.79 | 11.90 | 9.90 | 8.42 | 10.85 | 10.23 | 8.56 | 7.45 | 11.05 | 10.85 | 11.35 | 9.16 | 10.54 | 8.01 | 10.95 |
| RIPK1 | 7 | 9.83 | 10.82 | 10.95 | 10.35 | 10.94 | 11.43 | 9.75 | 10.00 | 7.72 | 8.20 | 10.32 | 10.65 | 11.19 | 9.88 | 7.96 |
| RIPK2 | 3 | 9.57 | 9.50 | 11.55 | 11.55 | 9.32 | 9.48 | 11.09 | 9.96 | 8.59 | 9.40 | 9.54 | 11.55 | 9.40 | 10.52 | 8.99 |
| RIT1 | 1 | 7.77 | 7.90 | 9.46 | 5.17 | 18.30 | 13.16 | 7.37 | 7.65 | 10.31 | 12.91 | 7.84 | 7.31 | 15.73 | 7.51 | 11.61 |
| RLF | 2 | 8.87 | 10.61 | 9.37 | 9.33 | 7.16 | 8.80 | 12.71 | 12.02 | 10.31 | 10.82 | 9.74 | 9.35 | 7.98 | 12.36 | 10.57 |
| RLIM | 1 | 7.84 | 9.92 | 8.12 | 9.21 | 11.97 | 13.14 | 10.72 | 9.12 | 10.91 | 9.03 | 8.88 | 8.67 | 12.56 | 9.92 | 9.97 |
| RMDN1 | 10 | 10.37 | 9.93 | 11.05 | 11.58 | 6.79 | 7.48 | 10.10 | 9.82 | 11.40 | 11.48 | 10.15 | 11.32 | 7.13 | 9.96 | 11.44 |
| RMDN3 | 13 | 10.04 | 8.70 | 9.09 | 9.67 | 10.83 | 10.96 | 8.86 | 8.18 | 11.78 | 11.89 | 9.37 | 9.38 | 10.90 | 8.52 | 11.84 |
| RMI1 | 5 | 9.86 | 10.28 | 9.84 | 9.04 | 10.98 | 11.20 | 8.95 | 8.52 | 10.64 | 10.69 | 10.07 | 9.44 | 11.09 | 8.73 | 10.66 |
| RMND1 | 1 | 10.85 | 9.19 | 10.51 | 11.46 | 9.78 | 9.95 | 7.70 | 8.36 | 12.89 | 9.31 | 10.02 | 10.99 | 9.87 | 8.03 | 11.10 |
| RMND5A | 3 | 11.18 | 11.32 | 8.23 | 7.33 | 11.86 | 11.39 | 10.73 | 10.69 | 8.55 | 8.70 | 11.25 | 7.78 | 11.63 | 10.71 | 8.63 |
| RNASEH1 | 1 | 11.97 | 9.79 | 10.26 | 8.30 | 8.63 | 10.14 | 10.84 | 10.20 | 11.34 | 8.53 | 10.88 | 9.28 | 9.38 | 10.52 | 9.93 |
| RNASEH2A | 8 | 9.03 | 10.31 | 11.16 | 10.88 | 10.42 | 11.45 | 11.60 | 11.03 | 7.08 | 7.04 | 9.67 | 11.02 | 10.93 | 11.32 | 7.06 |
| RNASEH2B | 6 | 9.07 | 10.02 | 12.01 | 11.91 | 11.55 | 11.98 | 10.52 | 10.62 | 6.32 | 6.00 | 9.54 | 11.96 | 11.76 | 10.57 | 6.16 |
| RNASEH2C | 4 | 8.24 | 10.16 | 13.15 | 12.41 | 11.63 | 11.50 | 10.33 | 9.25 | 6.84 | 6.49 | 9.20 | 12.78 | 11.56 | 9.79 | 6.66 |
| RNASEL | 1 | 10.90 | 10.55 | 5.92 | 5.40 | 22.39 | 20.85 | 7.41 | 5.73 | 6.16 | 4.68 | 10.73 | 5.66 | 21.62 | 6.57 | 5.42 |
| RNASET2 | 2 | 12.35 | 12.27 | 7.27 | 8.08 | 14.48 | 14.55 | 9.66 | 7.62 | 7.24 | 6.47 | 12.31 | 7.68 | 14.51 | 8.64 | 6.86 |
| RND3 | 1 | 10.08 | 10.15 | 14.42 | 14.84 | 7.42 | 7.09 | 4.48 | 4.15 | 13.25 | 14.12 | 10.11 | 14.63 | 7.26 | 4.31 | 13.68 |
| RNF113A | 6 | 8.63 | 10.51 | 6.46 | 6.16 | 12.87 | 12.67 | 13.91 | 12.92 | 8.29 | 7.57 | 9.57 | 6.31 | 12.77 | 13.42 | 7.93 |
| RNF114 | 4 | 9.90 | 10.34 | 9.41 | 7.96 | 8.87 | 8.78 | 9.38 | 15.62 | 9.76 | 9.98 | 10.12 | 8.68 | 8.83 | 12.50 | 9.87 |
| RNF123 | 7 | 9.97 | 10.62 | 8.78 | 9.11 | 11.25 | 11.82 | 9.96 | 8.67 | 10.09 | 9.73 | 10.29 | 8.94 | 11.54 | 9.31 | 9.91 |
| RNF126 | 1 | 9.37 | 9.61 | 10.29 | 10.02 | 8.81 | 8.46 | 12.26 | 9.56 | 11.84 | 9.78 | 9.49 | 10.15 | 8.63 | 10.91 | 10.81 |
| RNF13 | 1 | 10.47 | 8.90 | 8.04 | 12.35 | 10.44 | 11.27 | 6.98 | 6.48 | 12.01 | 13.06 | 9.69 | 10.19 | 10.85 | 6.73 | 12.54 |
| RNF139 | 1 | 2.79 | 16.85 | 4.47 | 3.49 | 15.06 | 27.41 | 3.42 | 2.96 | 13.20 | 10.34 | 9.82 | 3.98 | 21.23 | 3.19 | 11.77 |
| RNF14 | 3 | 10.63 | 11.18 | 10.84 | 9.71 | 7.72 | 8.47 | 10.07 | 9.88 | 10.59 | 10.92 | 10.91 | 10.28 | 8.09 | 9.97 | 10.75 |
| RNF169 | 6 | 7.96 | 9.57 | 11.87 | 10.66 | 10.31 | 9.91 | 11.47 | 11.00 | 8.79 | 8.46 | 8.76 | 11.27 | 10.11 | 11.23 | 8.62 |

| | | | | | | | | | | | | | | | | |
|---|---|---|---|---|---|---|---|---|---|---|---|---|---|---|---|---|
| RNF181 | 1 | 9.47 | 11.76 | 10.68 | 10.02 | 11.69 | 12.44 | 10.09 | 5.43 | 9.28 | 9.13 | 10.62 | 10.35 | 12.07 | 7.76 | 9.21 |
| RNF185 | 1 | 12.46 | 7.54 | 7.40 | 12.61 | 8.45 | 7.88 | 9.01 | 8.83 | 11.20 | 14.63 | 10.00 | 10.00 | 8.16 | 8.92 | 12.91 |
| RNF187 | 1 | 9.12 | 10.88 | 6.75 | 8.81 | 11.40 | 12.94 | 11.43 | 10.61 | 8.29 | 9.75 | 10.00 | 7.78 | 12.17 | 11.02 | 9.02 |
| RNF20 | 33 | 9.85 | 10.86 | 10.15 | 10.23 | 9.76 | 10.36 | 9.54 | 9.09 | 10.15 | 10.01 | 10.36 | 10.19 | 10.06 | 9.31 | 10.08 |
| RNF213 | 44 | 11.00 | 10.57 | 8.61 | 9.48 | 11.12 | 11.51 | 9.16 | 9.39 | 9.48 | 9.68 | 10.78 | 9.05 | 11.32 | 9.27 | 9.58 |
| RNF214 | 6 | 9.75 | 10.64 | 11.41 | 11.12 | 9.87 | 9.73 | 9.37 | 7.88 | 10.45 | 9.77 | 10.19 | 11.27 | 9.80 | 8.63 | 10.11 |
| RNF219 | 4 | 10.06 | 12.51 | 9.79 | 10.03 | 10.04 | 12.53 | 10.58 | 9.43 | 7.56 | 7.46 | 11.29 | 9.91 | 11.29 | 10.00 | 7.51 |
| RNF220 | 2 | 10.31 | 11.94 | 10.34 | 10.17 | 9.45 | 9.11 | 10.66 | 10.43 | 9.11 | 8.49 | 11.12 | 10.25 | 9.28 | 10.55 | 8.80 |
| RNF25 | 9 | 10.06 | 10.57 | 11.56 | 10.33 | 9.97 | 9.76 | 9.15 | 8.05 | 10.65 | 9.90 | 10.31 | 10.94 | 9.87 | 8.60 | 10.28 |
| RNF40 | 38 | 9.94 | 10.89 | 10.47 | 9.94 | 9.69 | 10.33 | 9.57 | 9.13 | 10.18 | 9.86 | 10.41 | 10.21 | 10.01 | 9.35 | 10.02 |
| RNFT2 | 1 | 12.48 | 7.85 | 7.08 | 7.30 | 14.97 | 12.54 | 12.09 | 10.95 | 6.42 | 8.32 | 10.16 | 7.19 | 13.75 | 11.52 | 7.37 |
| RNGTT | 5 | 9.40 | 11.72 | 9.42 | 9.92 | 9.91 | 10.31 | 10.87 | 11.13 | 8.91 | 8.42 | 10.56 | 9.67 | 10.11 | 11.00 | 8.66 |
| RNH1 | 15 | 8.65 | 8.54 | 8.92 | 7.78 | 9.68 | 9.20 | 9.37 | 20.80 | 8.86 | 8.21 | 8.59 | 8.35 | 9.44 | 15.08 | 8.54 |
| RNMT | 11 | 9.98 | 9.50 | 10.93 | 10.67 | 9.60 | 10.29 | 9.14 | 8.07 | 11.03 | 10.78 | 9.74 | 10.80 | 9.94 | 8.61 | 10.90 |
| RNMTL1 | 4 | 10.84 | 10.03 | 10.94 | 12.01 | 8.98 | 8.15 | 9.17 | 7.14 | 11.38 | 11.36 | 10.43 | 11.47 | 8.57 | 8.15 | 11.37 |
| RNPC3 | 1 | 8.43 | 9.84 | 10.68 | 9.74 | 10.61 | 11.44 | 11.69 | 9.59 | 8.16 | 9.81 | 9.14 | 10.21 | 11.03 | 10.64 | 8.99 |
| RNPEP | 34 | 8.69 | 9.84 | 7.89 | 6.97 | 12.99 | 12.96 | 9.36 | 10.08 | 10.69 | 10.53 | 9.27 | 7.43 | 12.98 | 9.72 | 10.61 |
| RNPS1 | 7 | 9.26 | 9.47 | 10.46 | 9.90 | 10.12 | 9.51 | 11.10 | 10.42 | 10.36 | 9.40 | 9.37 | 10.18 | 9.81 | 10.76 | 9.88 |
| ROBO1 | 18 | 11.89 | 9.60 | 8.39 | 9.16 | 10.50 | 10.19 | 10.34 | 8.70 | 10.51 | 10.73 | 10.74 | 8.77 | 10.34 | 9.52 | 10.62 |
| ROCK1 | 34 | 9.78 | 11.08 | 12.38 | 11.85 | 10.19 | 10.08 | 7.51 | 6.60 | 10.59 | 9.93 | 10.43 | 12.11 | 10.14 | 7.06 | 10.26 |
| ROCK2 | 72 | 10.67 | 11.61 | 11.78 | 11.65 | 9.89 | 10.32 | 5.72 | 5.35 | 11.62 | 11.40 | 11.14 | 11.72 | 10.10 | 5.53 | 11.51 |
| RP2 | 3 | 11.47 | 9.60 | 8.15 | 8.65 | 12.74 | 13.19 | 9.86 | 7.58 | 9.68 | 9.10 | 10.53 | 8.40 | 12.96 | 8.72 | 9.39 |
| RPA1 | 38 | 10.45 | 10.74 | 8.47 | 8.04 | 10.41 | 10.33 | 9.51 | 13.14 | 9.59 | 9.32 | 10.59 | 8.26 | 10.37 | 11.33 | 9.45 |
| RPA2 | 7 | 9.87 | 10.95 | 8.42 | 7.63 | 10.61 | 11.26 | 9.93 | 12.25 | 9.50 | 9.58 | 10.41 | 8.02 | 10.93 | 11.09 | 9.54 |
| RPA3 | 9 | 9.67 | 10.22 | 8.62 | 8.03 | 10.88 | 10.22 | 10.36 | 13.59 | 9.31 | 9.11 | 9.94 | 8.33 | 10.55 | 11.98 | 9.21 |
| RPAP1 | 4 | 10.00 | 10.69 | 8.97 | 9.28 | 9.91 | 9.77 | 9.09 | 8.78 | 12.19 | 11.30 | 10.35 | 9.13 | 9.84 | 8.94 | 11.75 |
| RPAP3 | 28 | 9.52 | 10.97 | 12.25 | 11.17 | 8.66 | 8.66 | 10.65 | 9.67 | 9.21 | 9.24 | 10.25 | 11.71 | 8.66 | 10.16 | 9.23 |
| RPE | 6 | 8.65 | 9.71 | 9.81 | 8.41 | 10.94 | 10.68 | 9.09 | 11.20 | 11.00 | 10.51 | 9.18 | 9.11 | 10.81 | 10.14 | 10.75 |
| RPF1 | 6 | 8.50 | 10.01 | 10.47 | 10.73 | 7.57 | 7.89 | 12.38 | 11.02 | 10.00 | 11.43 | 9.25 | 10.60 | 7.73 | 11.70 | 10.71 |
| RPF2 | 8 | 10.85 | 9.19 | 9.46 | 9.89 | 7.76 | 7.69 | 12.17 | 10.56 | 10.11 | 12.33 | 10.02 | 9.67 | 7.72 | 11.36 | 11.22 |
| RPIA | 6 | 7.77 | 9.79 | 12.96 | 12.76 | 7.07 | 7.30 | 12.35 | 12.05 | 8.53 | 9.43 | 8.78 | 12.86 | 7.19 | 12.20 | 8.98 |
| RPL10 | 24 | 8.65 | 9.61 | 9.35 | 8.85 | 8.86 | 8.77 | 9.07 | 18.34 | 8.98 | 9.51 | 9.13 | 9.10 | 8.82 | 13.70 | 9.24 |
| RPL10A | 27 | 8.62 | 11.92 | 11.17 | 10.07 | 9.18 | 9.04 | 9.90 | 10.70 | 9.08 | 10.32 | 10.27 | 10.62 | 9.11 | 10.30 | 9.70 |
| RPL11 | 20 | 9.53 | 10.25 | 10.69 | 9.74 | 9.47 | 9.59 | 10.18 | 11.59 | 9.65 | 9.31 | 9.89 | 10.22 | 9.53 | 10.88 | 9.48 |
| RPL12 | 28 | 7.28 | 12.53 | 10.78 | 10.01 | 9.30 | 9.32 | 9.67 | 12.50 | 9.20 | 9.42 | 9.91 | 10.40 | 9.31 | 11.08 | 9.31 |
| RPL13 | 45 | 9.99 | 10.91 | 10.07 | 10.38 | 9.49 | 10.03 | 9.37 | 9.49 | 9.90 | 10.36 | 10.45 | 10.22 | 9.76 | 9.43 | 10.13 |
| RPL13A | 26 | 9.42 | 9.96 | 10.75 | 10.52 | 9.13 | 9.36 | 9.88 | 11.04 | 9.77 | 10.17 | 9.69 | 10.64 | 9.25 | 10.46 | 9.97 |
| RPL14 | 10 | 7.57 | 7.10 | 11.73 | 8.59 | 8.81 | 7.12 | 8.05 | 21.19 | 8.81 | 11.04 | 7.33 | 10.16 | 7.96 | 14.62 | 9.93 |
| RPL15 | 39 | 9.67 | 10.33 | 9.98 | 9.84 | 9.59 | 9.55 | 10.50 | 10.09 | 10.09 | 10.36 | 10.00 | 9.91 | 9.57 | 10.30 | 10.22 |
| RPL17 | 38 | 9.94 | 10.60 | 10.98 | 10.69 | 9.50 | 9.07 | 9.24 | 8.32 | 10.15 | 11.51 | 10.27 | 10.83 | 9.28 | 8.78 | 10.83 |
| RPL18 | 21 | 9.70 | 9.85 | 9.88 | 9.62 | 9.39 | 8.93 | 9.51 | 12.95 | 9.67 | 10.50 | 9.77 | 9.75 | 9.16 | 11.23 | 10.09 |
| RPL18A | 19 | 10.21 | 10.36 | 9.98 | 10.37 | 9.62 | 9.79 | 9.69 | 10.12 | 9.42 | 10.44 | 10.28 | 10.17 | 9.71 | 9.91 | 9.93 |

| | | | | | | | | | | | | | | | | |
|---|---|---|---|---|---|---|---|---|---|---|---|---|---|---|---|---|
| RPL19 | 18 | 9.48 | 10.27 | 9.89 | 10.20 | 9.97 | 10.13 | 9.63 | 9.43 | 9.99 | 11.00 | 9.88 | 10.05 | 10.05 | 9.53 | 10.49 |
| RPL21 | 22 | 10.11 | 9.88 | 10.92 | 10.01 | 9.71 | 9.44 | 10.00 | 9.06 | 10.05 | 10.83 | 10.00 | 10.47 | 9.57 | 9.53 | 10.44 |
| RPL22 | 8 | 10.43 | 10.27 | 9.67 | 9.99 | 8.83 | 9.14 | 10.27 | 11.06 | 10.05 | 10.27 | 10.35 | 9.83 | 8.99 | 10.66 | 10.16 |
| RPL22L1 | 2 | 5.54 | 5.59 | 13.31 | 13.10 | 21.88 | 24.05 | 4.48 | 4.43 | 3.87 | 3.75 | 5.56 | 13.21 | 22.96 | 4.46 | 3.81 |
| RPL23 | 31 | 9.40 | 9.95 | 9.94 | 9.86 | 8.81 | 8.43 | 8.85 | 15.79 | 9.30 | 9.68 | 9.67 | 9.90 | 8.62 | 12.32 | 9.49 |
| RPL23A | 33 | 10.71 | 10.56 | 10.82 | 9.81 | 9.73 | 9.24 | 9.74 | 8.56 | 10.36 | 10.46 | 10.64 | 10.31 | 9.49 | 9.15 | 10.41 |
| RPL24 | 32 | 9.92 | 10.19 | 10.28 | 10.09 | 9.45 | 9.28 | 8.46 | 11.12 | 10.18 | 11.03 | 10.05 | 10.18 | 9.37 | 9.79 | 10.61 |
| RPL26 | 19 | 10.00 | 10.48 | 10.72 | 10.45 | 9.49 | 9.66 | 8.91 | 7.87 | 10.32 | 12.10 | 10.24 | 10.58 | 9.58 | 8.39 | 11.21 |
| RPL26L1 | 1 | 3.98 | 10.09 | 11.46 | 9.59 | 13.42 | 12.56 | 11.31 | 8.83 | 8.26 | 10.52 | 7.03 | 10.53 | 12.99 | 10.07 | 9.39 |
| RPL27 | 26 | 10.37 | 10.71 | 11.16 | 10.13 | 9.43 | 9.30 | 9.31 | 8.28 | 10.34 | 10.96 | 10.54 | 10.64 | 9.36 | 8.80 | 10.65 |
| RPL27A | 19 | 9.42 | 9.48 | 10.88 | 7.42 | 9.41 | 8.38 | 9.13 | 16.18 | 9.78 | 9.91 | 9.45 | 9.15 | 8.90 | 12.66 | 9.84 |
| RPL28 | 22 | 9.89 | 11.54 | 9.91 | 10.36 | 9.09 | 9.27 | 8.92 | 10.39 | 9.70 | 10.95 | 10.72 | 10.13 | 9.18 | 9.65 | 10.32 |
| RPL29 | 16 | 9.66 | 10.06 | 12.06 | 9.93 | 9.96 | 9.70 | 9.53 | 7.94 | 10.32 | 10.85 | 9.86 | 10.99 | 9.83 | 8.73 | 10.58 |
| RPL3 | 55 | 10.10 | 9.74 | 9.75 | 9.78 | 9.26 | 9.14 | 9.51 | 13.52 | 9.61 | 9.57 | 9.92 | 9.77 | 9.20 | 11.52 | 9.59 |
| RPL30 | 18 | 7.30 | 8.95 | 9.57 | 8.62 | 9.31 | 9.73 | 9.64 | 18.23 | 9.03 | 9.62 | 8.13 | 9.10 | 9.52 | 13.93 | 9.32 |
| RPL31 | 32 | 10.98 | 11.86 | 9.49 | 10.52 | 9.50 | 9.68 | 8.91 | 8.57 | 9.89 | 10.61 | 11.42 | 10.01 | 9.59 | 8.74 | 10.25 |
| RPL32 | 24 | 9.50 | 9.54 | 10.60 | 9.83 | 9.47 | 9.18 | 9.74 | 13.27 | 9.43 | 9.43 | 9.52 | 10.22 | 9.33 | 11.51 | 9.43 |
| RPL34 | 21 | 10.48 | 10.29 | 10.61 | 10.81 | 9.08 | 8.87 | 8.86 | 10.18 | 9.93 | 10.89 | 10.38 | 10.71 | 8.98 | 9.52 | 10.41 |
| RPL35 | 6 | 10.83 | 9.90 | 9.99 | 10.45 | 9.01 | 10.16 | 8.82 | 9.42 | 10.27 | 11.15 | 10.36 | 10.22 | 9.59 | 9.12 | 10.71 |
| RPL35A | 22 | 11.08 | 10.71 | 9.46 | 10.24 | 9.10 | 9.72 | 9.51 | 9.78 | 9.90 | 10.50 | 10.89 | 9.85 | 9.41 | 9.64 | 10.20 |
| RPL36 | 18 | 9.16 | 9.61 | 10.70 | 10.88 | 9.91 | 9.47 | 9.71 | 9.63 | 10.05 | 10.87 | 9.39 | 10.79 | 9.69 | 9.67 | 10.46 |
| RPL36A | 12 | 6.25 | 12.43 | 12.28 | 11.78 | 8.82 | 8.83 | 8.99 | 9.84 | 9.21 | 11.58 | 9.34 | 12.03 | 8.82 | 9.42 | 10.39 |
| RPL36AL | 1 | 10.99 | 10.74 | 12.01 | 11.16 | 9.29 | 9.32 | 7.05 | 5.71 | 11.34 | 12.39 | 10.87 | 11.58 | 9.31 | 6.38 | 11.86 |
| RPL37 | 6 | 9.95 | 10.94 | 9.72 | 10.01 | 9.58 | 9.91 | 9.92 | 9.41 | 10.31 | 10.26 | 10.44 | 9.86 | 9.75 | 9.66 | 10.29 |
| RPL37A | 15 | 6.94 | 10.71 | 14.00 | 10.91 | 9.08 | 8.79 | 9.11 | 10.36 | 9.00 | 11.10 | 8.82 | 12.45 | 8.93 | 9.74 | 10.05 |
| RPL38 | 15 | 9.77 | 10.42 | 10.27 | 11.72 | 9.28 | 10.17 | 9.23 | 9.44 | 9.90 | 9.81 | 10.09 | 11.00 | 9.72 | 9.34 | 9.85 |
| RPL39 | 2 | 9.09 | 9.65 | 10.88 | 11.31 | 9.32 | 9.68 | 10.14 | 9.46 | 10.43 | 10.04 | 9.37 | 11.09 | 9.50 | 9.80 | 10.24 |
| RPL4 | 64 | 9.11 | 12.44 | 10.05 | 10.38 | 9.29 | 9.37 | 9.36 | 11.00 | 9.44 | 9.55 | 10.78 | 10.22 | 9.33 | 10.18 | 9.50 |
| RPL5 | 73 | 9.31 | 10.05 | 11.36 | 10.14 | 9.20 | 8.92 | 10.38 | 10.41 | 10.12 | 10.12 | 9.68 | 10.75 | 9.06 | 10.39 | 10.12 |
| RPL6 | 41 | 9.57 | 10.06 | 11.23 | 8.90 | 10.82 | 10.26 | 10.10 | 8.82 | 10.08 | 10.16 | 9.81 | 10.06 | 10.54 | 9.46 | 10.12 |
| RPL7 | 40 | 9.37 | 10.15 | 11.76 | 10.78 | 9.62 | 9.30 | 9.81 | 9.56 | 9.43 | 10.22 | 9.76 | 11.27 | 9.46 | 9.69 | 9.83 |
| RPL7A | 35 | 9.87 | 9.65 | 9.51 | 10.20 | 9.03 | 9.22 | 8.83 | 13.36 | 9.55 | 10.78 | 9.76 | 9.85 | 9.13 | 11.10 | 10.16 |
| RPL7L1 | 4 | 11.68 | 8.27 | 9.67 | 12.76 | 8.27 | 7.61 | 12.48 | 9.75 | 8.03 | 11.48 | 9.97 | 11.21 | 7.94 | 11.12 | 9.76 |
| RPL8 | 20 | 9.69 | 10.46 | 10.75 | 10.51 | 9.57 | 9.44 | 8.87 | 9.36 | 10.37 | 10.97 | 10.07 | 10.63 | 9.50 | 9.12 | 10.67 |
| RPL9 | 28 | 9.54 | 9.78 | 9.83 | 9.33 | 9.29 | 9.22 | 9.89 | 14.53 | 9.35 | 9.24 | 9.66 | 9.58 | 9.25 | 12.21 | 9.29 |
| RPLP0 | 39 | 9.25 | 9.71 | 9.76 | 9.50 | 9.18 | 9.67 | 9.43 | 15.30 | 9.28 | 8.91 | 9.48 | 9.63 | 9.43 | 12.36 | 9.09 |
| RPLP1 | 2 | 5.22 | 4.12 | 7.31 | 4.61 | 7.76 | 5.62 | 6.00 | 47.10 | 7.11 | 5.16 | 4.67 | 5.96 | 6.69 | 26.55 | 6.13 |
| RPLP2 | 36 | 9.11 | 14.16 | 10.48 | 11.11 | 9.52 | 9.54 | 9.77 | 5.77 | 10.28 | 10.25 | 11.64 | 10.79 | 9.53 | 7.77 | 10.27 |
| RPN1 | 39 | 12.80 | 6.22 | 8.30 | 9.71 | 10.44 | 7.76 | 10.58 | 8.17 | 11.39 | 14.64 | 9.51 | 9.00 | 9.10 | 9.37 | 13.02 |
| RPN2 | 12 | 14.34 | 6.28 | 8.44 | 10.63 | 7.42 | 5.20 | 9.98 | 10.41 | 12.59 | 14.70 | 10.31 | 9.54 | 6.31 | 10.20 | 13.65 |
| RPP14 | 1 | 9.35 | 9.85 | 10.10 | 7.81 | 9.87 | 10.06 | 11.59 | 11.62 | 9.70 | 10.04 | 9.60 | 8.96 | 9.97 | 11.61 | 9.87 |
| RPP21 | 1 | 7.80 | 10.41 | 10.60 | 7.99 | 9.30 | 9.91 | 11.89 | 10.39 | 10.17 | 11.53 | 9.10 | 9.29 | 9.61 | 11.14 | 10.85 |

| | | | | | | | | | | | | | | | | |
|---|---|---|---|---|---|---|---|---|---|---|---|---|---|---|---|---|
| RPP25 | 3 | 9.19 | 9.30 | 7.46 | 8.57 | 9.85 | 10.42 | 11.84 | 11.82 | 10.67 | 10.89 | 9.24 | 8.02 | 10.13 | 11.83 | 10.78 |
| RPP25L | 4 | 9.27 | 12.07 | 10.74 | 10.48 | 8.18 | 8.70 | 10.24 | 9.80 | 10.54 | 9.99 | 10.67 | 10.61 | 8.44 | 10.02 | 10.26 |
| RPP30 | 8 | 9.51 | 10.09 | 9.76 | 10.03 | 9.64 | 9.56 | 11.08 | 11.21 | 9.62 | 9.50 | 9.80 | 9.90 | 9.60 | 11.14 | 9.56 |
| RPP38 | 8 | 9.84 | 10.01 | 10.05 | 10.58 | 8.95 | 9.62 | 10.69 | 9.90 | 9.80 | 10.55 | 9.93 | 10.31 | 9.29 | 10.30 | 10.17 |
| RPP40 | 6 | 8.25 | 8.39 | 9.59 | 7.55 | 9.33 | 7.78 | 9.63 | 21.56 | 9.60 | 8.31 | 8.32 | 8.57 | 8.56 | 15.60 | 8.96 |
| RPRD1A | 5 | 9.56 | 10.49 | 11.48 | 11.11 | 12.06 | 12.36 | 8.56 | 8.34 | 7.94 | 8.10 | 10.03 | 11.30 | 12.21 | 8.45 | 8.02 |
| RPRD1B | 7 | 8.63 | 9.02 | 11.71 | 10.70 | 10.65 | 10.19 | 10.94 | 10.69 | 8.71 | 8.76 | 8.82 | 11.21 | 10.42 | 10.82 | 8.74 |
| RPRD2 | 20 | 8.43 | 9.88 | 9.45 | 9.14 | 12.43 | 12.70 | 10.65 | 10.10 | 8.76 | 8.46 | 9.16 | 9.29 | 12.56 | 10.38 | 8.61 |
| RPS10 | 31 | 9.46 | 10.66 | 10.74 | 10.55 | 9.39 | 9.72 | 10.55 | 9.93 | 9.62 | 9.39 | 10.06 | 10.64 | 9.55 | 10.24 | 9.51 |
| RPS11 | 31 | 10.16 | 10.24 | 10.12 | 9.89 | 9.29 | 9.02 | 9.64 | 11.95 | 9.52 | 10.17 | 10.20 | 10.00 | 9.15 | 10.79 | 9.85 |
| RPS12 | 22 | 8.04 | 9.36 | 8.22 | 8.05 | 8.82 | 8.54 | 8.74 | 24.00 | 8.39 | 7.84 | 8.70 | 8.14 | 8.68 | 16.37 | 8.12 |
| RPS13 | 25 | 10.29 | 10.21 | 10.25 | 10.62 | 9.40 | 9.65 | 9.60 | 8.84 | 10.18 | 10.96 | 10.25 | 10.43 | 9.53 | 9.22 | 10.57 |
| RPS14 | 17 | 10.68 | 10.71 | 10.29 | 10.23 | 9.57 | 9.53 | 10.31 | 9.52 | 9.66 | 9.50 | 10.70 | 10.26 | 9.55 | 9.91 | 9.58 |
| RPS15 | 29 | 8.90 | 10.33 | 9.63 | 10.77 | 9.26 | 10.27 | 10.24 | 10.37 | 9.38 | 10.84 | 9.61 | 10.20 | 9.77 | 10.31 | 10.11 |
| RPS15A | 33 | 8.79 | 9.91 | 10.56 | 9.33 | 10.49 | 10.40 | 10.78 | 10.38 | 9.72 | 9.64 | 9.35 | 9.94 | 10.44 | 10.58 | 9.68 |
| RPS16 | 26 | 9.42 | 10.57 | 10.38 | 9.73 | 9.52 | 10.26 | 10.57 | 10.43 | 9.36 | 9.76 | 9.99 | 10.05 | 9.89 | 10.50 | 9.56 |
| RPS17L | 22 | 8.54 | 8.88 | 12.06 | 9.67 | 9.67 | 9.54 | 10.36 | 12.21 | 9.36 | 9.70 | 8.71 | 10.87 | 9.61 | 11.29 | 9.53 |
| RPS18 | 29 | 10.26 | 10.17 | 10.21 | 10.45 | 9.81 | 9.96 | 10.15 | 9.15 | 9.70 | 10.13 | 10.22 | 10.33 | 9.88 | 9.65 | 9.91 |
| RPS19 | 37 | 9.18 | 10.50 | 10.79 | 10.61 | 10.06 | 10.11 | 10.43 | 9.20 | 9.52 | 9.59 | 9.84 | 10.70 | 10.08 | 9.81 | 9.55 |
| RPS19BP1 | 1 | 10.63 | 10.84 | 7.99 | 9.25 | 9.64 | 8.46 | 11.25 | 10.66 | 10.71 | 10.58 | 10.73 | 8.62 | 9.05 | 10.95 | 10.64 |
| RPS2 | 45 | 9.79 | 9.49 | 9.92 | 9.00 | 9.55 | 9.38 | 10.32 | 13.51 | 9.43 | 9.61 | 9.64 | 9.46 | 9.47 | 11.91 | 9.52 |
| RPS20 | 19 | 8.73 | 10.82 | 10.36 | 8.79 | 9.39 | 9.04 | 8.95 | 17.32 | 8.38 | 8.22 | 9.78 | 9.58 | 9.21 | 13.13 | 8.30 |
| RPS21 | 12 | 8.92 | 8.81 | 12.32 | 9.74 | 10.02 | 9.77 | 9.91 | 11.98 | 9.66 | 8.87 | 8.87 | 11.03 | 9.90 | 10.94 | 9.26 |
| RPS23 | 14 | 8.86 | 10.85 | 11.33 | 10.85 | 9.75 | 9.69 | 9.59 | 8.71 | 9.62 | 10.75 | 9.85 | 11.09 | 9.72 | 9.15 | 10.19 |
| RPS24 | 19 | 9.74 | 10.97 | 10.16 | 10.46 | 9.30 | 9.81 | 10.10 | 9.56 | 9.65 | 10.26 | 10.35 | 10.31 | 9.55 | 9.83 | 9.95 |
| RPS25 | 25 | 9.86 | 10.35 | 10.36 | 10.26 | 9.45 | 9.89 | 9.98 | 9.60 | 10.10 | 10.15 | 10.10 | 10.31 | 9.67 | 9.79 | 10.13 |
| RPS26P11 | 13 | 10.91 | 10.60 | 9.39 | 10.57 | 9.52 | 9.86 | 9.97 | 9.51 | 10.06 | 9.60 | 10.75 | 9.98 | 9.69 | 9.74 | 9.83 |
| RPS27 | 3 | 9.71 | 10.14 | 12.67 | 6.99 | 11.90 | 10.70 | 10.91 | 7.49 | 10.39 | 9.10 | 9.92 | 9.83 | 11.30 | 9.20 | 9.74 |
| RPS27A | 55 | 9.72 | 11.09 | 10.25 | 8.60 | 9.86 | 9.21 | 10.98 | 12.57 | 8.92 | 8.78 | 10.41 | 9.43 | 9.54 | 11.78 | 8.85 |
| RPS27L | 16 | 8.99 | 8.53 | 11.48 | 8.31 | 10.65 | 9.82 | 10.85 | 12.29 | 9.83 | 9.25 | 8.76 | 9.89 | 10.24 | 11.57 | 9.54 |
| RPS28 | 18 | 9.31 | 10.79 | 10.00 | 10.08 | 9.79 | 9.92 | 9.95 | 10.63 | 9.98 | 9.53 | 10.05 | 10.04 | 9.86 | 10.29 | 9.76 |
| RPS29 | 1 | 4.01 | 4.09 | 6.61 | 1.26 | 8.71 | 4.65 | 5.67 | 54.67 | 5.49 | 4.82 | 4.05 | 3.94 | 6.68 | 30.17 | 5.16 |
| RPS3 | 76 | 9.64 | 10.40 | 10.31 | 9.63 | 9.81 | 10.00 | 10.61 | 11.59 | 9.20 | 8.81 | 10.02 | 9.97 | 9.91 | 11.10 | 9.01 |
| RPS3A | 45 | 9.62 | 9.28 | 10.16 | 9.43 | 9.52 | 9.25 | 9.75 | 13.47 | 9.72 | 9.80 | 9.45 | 9.79 | 9.39 | 11.61 | 9.76 |
| RPS4X | 43 | 9.28 | 9.83 | 10.54 | 9.47 | 9.78 | 9.58 | 10.76 | 11.49 | 9.31 | 9.96 | 9.55 | 10.00 | 9.68 | 11.13 | 9.64 |
| RPS5 | 14 | 7.47 | 7.68 | 12.81 | 11.20 | 9.21 | 8.33 | 9.95 | 17.20 | 8.21 | 7.95 | 7.57 | 12.00 | 8.77 | 13.58 | 8.08 |
| RPS6 | 29 | 10.10 | 10.08 | 9.14 | 9.02 | 9.15 | 9.50 | 9.07 | 13.81 | 9.90 | 10.23 | 10.09 | 9.08 | 9.33 | 11.44 | 10.06 |
| RPS6KA1 | 26 | 11.16 | 11.93 | 8.79 | 7.72 | 10.23 | 9.96 | 10.27 | 9.55 | 10.16 | 10.23 | 11.55 | 8.25 | 10.09 | 9.91 | 10.20 |
| RPS6KA3 | 15 | 8.41 | 10.10 | 8.77 | 9.15 | 12.45 | 14.39 | 10.94 | 11.12 | 7.10 | 7.58 | 9.25 | 8.96 | 13.42 | 11.03 | 7.34 |
| RPS6KA4 | 4 | 8.89 | 9.34 | 8.69 | 8.98 | 10.83 | 11.67 | 11.51 | 11.52 | 9.07 | 9.50 | 9.11 | 8.83 | 11.25 | 11.52 | 9.29 |
| RPS6KA5 | 3 | 10.01 | 8.81 | 9.61 | 9.09 | 11.50 | 12.24 | 12.28 | 11.48 | 7.84 | 7.13 | 9.41 | 9.35 | 11.87 | 11.88 | 7.49 |
| RPS6KA6 | 6 | 9.30 | 9.70 | 9.93 | 7.70 | 13.02 | 12.09 | 11.58 | 9.88 | 8.34 | 8.47 | 9.50 | 8.81 | 12.56 | 10.73 | 8.41 |

| | | | | | | | | | | | | | | | | |
|---|---|---|---|---|---|---|---|---|---|---|---|---|---|---|---|---|
| RPS6KB1 | 7 | 9.14 | 11.89 | 10.67 | 10.74 | 8.03 | 8.59 | 10.50 | 10.57 | 9.69 | 10.18 | 10.51 | 10.71 | 8.31 | 10.54 | 9.93 |
| RPS6KB2 | 3 | 9.47 | 10.53 | 10.07 | 9.68 | 10.18 | 11.02 | 9.36 | 8.42 | 11.93 | 9.34 | 10.00 | 9.88 | 10.60 | 8.89 | 10.63 |
| RPS6KC1 | 1 | 11.16 | 13.00 | 8.11 | 8.17 | 9.17 | 10.66 | 10.36 | 8.27 | 11.28 | 9.82 | 12.08 | 8.14 | 9.91 | 9.32 | 10.55 |
| RPS7 | 29 | 9.15 | 9.95 | 11.46 | 11.18 | 9.66 | 9.63 | 10.35 | 9.72 | 9.36 | 9.55 | 9.55 | 11.32 | 9.64 | 10.03 | 9.45 |
| RPS8 | 30 | 9.35 | 9.39 | 10.33 | 9.54 | 9.28 | 9.39 | 9.74 | 14.14 | 9.58 | 9.27 | 9.37 | 9.93 | 9.33 | 11.94 | 9.42 |
| RPS9 | 38 | 10.04 | 10.49 | 10.19 | 10.60 | 9.33 | 9.74 | 9.81 | 9.26 | 9.63 | 10.91 | 10.26 | 10.39 | 9.54 | 9.54 | 10.27 |
| RPSA | 57 | 9.23 | 10.83 | 9.86 | 8.87 | 9.88 | 9.80 | 10.46 | 12.23 | 9.48 | 9.35 | 10.03 | 9.37 | 9.84 | 11.34 | 9.42 |
| RPTOR | 7 | 10.04 | 11.48 | 9.28 | 9.71 | 9.89 | 10.12 | 8.62 | 8.31 | 11.01 | 11.55 | 10.76 | 9.50 | 10.00 | 8.46 | 11.28 |
| RPUSD2 | 9 | 8.51 | 9.33 | 9.94 | 9.80 | 10.08 | 10.12 | 11.49 | 10.66 | 10.23 | 9.85 | 8.92 | 9.87 | 10.10 | 11.08 | 10.04 |
| RPUSD3 | 2 | 10.58 | 9.70 | 9.02 | 11.85 | 7.92 | 8.66 | 11.33 | 10.75 | 10.08 | 10.11 | 10.14 | 10.44 | 8.29 | 11.04 | 10.09 |
| RPUSD4 | 3 | 10.45 | 8.81 | 9.51 | 11.48 | 8.42 | 8.40 | 11.21 | 10.37 | 10.72 | 10.63 | 9.63 | 10.49 | 8.41 | 10.79 | 10.67 |
| RQCD1 | 5 | 9.86 | 10.54 | 9.45 | 9.01 | 10.02 | 10.01 | 10.77 | 10.53 | 10.05 | 9.75 | 10.20 | 9.23 | 10.02 | 10.65 | 9.90 |
| RRAGB | 5 | 10.75 | 11.16 | 7.34 | 8.15 | 10.20 | 10.58 | 7.86 | 8.14 | 13.15 | 12.67 | 10.96 | 7.75 | 10.39 | 8.00 | 12.91 |
| RRAGC | 7 | 10.20 | 11.01 | 8.58 | 8.83 | 9.57 | 10.17 | 8.80 | 9.62 | 11.64 | 11.56 | 10.61 | 8.71 | 9.87 | 9.21 | 11.60 |
| RRAS | 8 | 10.02 | 8.45 | 10.09 | 12.63 | 7.28 | 7.90 | 7.94 | 7.58 | 13.83 | 14.27 | 9.24 | 11.36 | 7.59 | 7.76 | 14.05 |
| RRAS2 | 3 | 10.21 | 7.98 | 9.62 | 11.59 | 7.72 | 7.95 | 9.65 | 8.08 | 13.32 | 13.88 | 9.10 | 10.61 | 7.83 | 8.86 | 13.60 |
| RRBP1 | 110 | 8.61 | 8.20 | 12.81 | 12.62 | 9.74 | 9.40 | 9.24 | 8.72 | 10.36 | 10.31 | 8.40 | 12.72 | 9.57 | 8.98 | 10.33 |
| RREB1 | 1 | 7.24 | 8.04 | 10.48 | 9.48 | 11.08 | 9.51 | 13.90 | 12.64 | 8.38 | 9.25 | 7.64 | 9.98 | 10.30 | 13.27 | 8.82 |
| RRM1 | 70 | 10.38 | 10.88 | 8.64 | 8.11 | 11.60 | 11.61 | 9.02 | 11.40 | 9.21 | 9.15 | 10.63 | 8.38 | 11.61 | 10.21 | 9.18 |
| RRM2 | 12 | 9.81 | 11.93 | 7.97 | 7.94 | 9.49 | 9.40 | 8.85 | 10.96 | 11.90 | 11.75 | 10.87 | 7.95 | 9.44 | 9.91 | 11.83 |
| RRM2B | 4 | 11.31 | 11.00 | 8.07 | 8.03 | 10.84 | 10.69 | 11.14 | 10.09 | 9.49 | 9.34 | 11.16 | 8.05 | 10.77 | 10.61 | 9.42 |
| RRN3 | 9 | 10.05 | 10.10 | 10.40 | 10.91 | 9.49 | 10.02 | 8.11 | 8.64 | 11.15 | 11.13 | 10.07 | 10.66 | 9.75 | 8.38 | 11.14 |
| RRP1 | 14 | 9.57 | 10.97 | 8.76 | 9.34 | 8.96 | 9.02 | 11.33 | 11.11 | 10.25 | 10.70 | 10.27 | 9.05 | 8.99 | 11.22 | 10.47 |
| RRP12 | 27 | 9.81 | 8.95 | 9.39 | 9.38 | 8.13 | 7.66 | 11.13 | 15.40 | 9.44 | 10.71 | 9.38 | 9.38 | 7.90 | 13.26 | 10.07 |
| RRP15 | 13 | 9.31 | 10.29 | 10.88 | 10.58 | 9.66 | 9.31 | 12.26 | 9.90 | 9.17 | 8.65 | 9.80 | 10.73 | 9.48 | 11.08 | 8.91 |
| RRP1B | 10 | 10.89 | 10.81 | 8.88 | 9.23 | 9.46 | 8.92 | 9.01 | 8.80 | 11.74 | 12.26 | 10.85 | 9.06 | 9.19 | 8.90 | 12.00 |
| RRP36 | 7 | 9.66 | 10.41 | 11.42 | 12.07 | 8.31 | 8.30 | 9.42 | 9.24 | 10.34 | 10.84 | 10.04 | 11.75 | 8.30 | 9.33 | 10.59 |
| RRP7A | 11 | 8.50 | 10.49 | 11.11 | 10.31 | 7.85 | 8.47 | 9.87 | 10.65 | 11.29 | 11.45 | 9.50 | 10.71 | 8.16 | 10.26 | 11.37 |
| RRP8 | 16 | 8.60 | 10.05 | 9.83 | 10.42 | 9.46 | 9.78 | 10.02 | 10.48 | 10.49 | 10.88 | 9.32 | 10.12 | 9.62 | 10.25 | 10.69 |
| RRP9 | 11 | 9.62 | 10.06 | 9.91 | 9.32 | 8.65 | 9.10 | 11.25 | 9.92 | 11.40 | 10.78 | 9.84 | 9.61 | 8.88 | 10.58 | 11.09 |
| RRS1 | 17 | 9.51 | 9.58 | 10.23 | 10.24 | 7.85 | 8.03 | 12.95 | 12.57 | 9.59 | 9.46 | 9.54 | 10.23 | 7.94 | 12.76 | 9.53 |
| RSBN1 | 1 | 9.49 | 10.58 | 8.92 | 9.29 | 7.64 | 7.18 | 14.40 | 13.73 | 9.35 | 9.43 | 10.03 | 9.10 | 7.41 | 14.06 | 9.39 |
| RSBN1L | 12 | 10.11 | 11.03 | 10.90 | 12.05 | 7.75 | 7.84 | 10.29 | 10.00 | 10.05 | 9.98 | 10.57 | 11.48 | 7.79 | 10.15 | 10.02 |
| RSF1 | 26 | 9.10 | 10.58 | 12.04 | 11.59 | 9.73 | 10.16 | 9.45 | 9.30 | 9.24 | 8.81 | 9.84 | 11.81 | 9.95 | 9.38 | 9.03 |
| RSL1D1 | 31 | 9.43 | 9.21 | 9.82 | 10.44 | 7.78 | 8.07 | 10.54 | 15.37 | 9.24 | 10.10 | 9.32 | 10.13 | 7.93 | 12.95 | 9.67 |
| RSL24D1 | 6 | 9.54 | 9.16 | 10.80 | 11.05 | 8.36 | 8.11 | 11.94 | 10.69 | 10.73 | 9.62 | 9.35 | 10.93 | 8.23 | 11.31 | 10.17 |
| RSRC1 | 5 | 9.70 | 10.62 | 10.51 | 9.78 | 9.61 | 10.08 | 11.09 | 9.42 | 9.50 | 9.70 | 10.16 | 10.14 | 9.84 | 10.26 | 9.60 |
| RSRC2 | 7 | 8.92 | 9.71 | 8.38 | 7.30 | 10.29 | 10.21 | 11.16 | 15.99 | 9.48 | 8.56 | 9.31 | 7.84 | 10.25 | 13.58 | 9.02 |
| RSU1 | 20 | 9.04 | 9.99 | 15.61 | 15.10 | 7.38 | 7.73 | 6.68 | 6.36 | 11.24 | 10.89 | 9.51 | 15.35 | 7.55 | 6.52 | 11.06 |
| RTCA | 9 | 8.89 | 10.85 | 8.96 | 9.53 | 9.23 | 10.88 | 10.56 | 11.54 | 9.38 | 10.18 | 9.87 | 9.24 | 10.06 | 11.05 | 9.78 |
| RTCB | 53 | 10.62 | 11.29 | 12.06 | 11.00 | 7.79 | 7.84 | 10.16 | 9.92 | 9.95 | 9.37 | 10.95 | 11.53 | 7.82 | 10.04 | 9.66 |
| RTEL1 | 2 | 9.37 | 10.55 | 9.34 | 9.31 | 8.72 | 8.75 | 12.13 | 10.86 | 10.52 | 10.46 | 9.96 | 9.32 | 8.73 | 11.49 | 10.49 |

| | | | | | | | | | | | | | | | | |
|---|---|---|---|---|---|---|---|---|---|---|---|---|---|---|---|---|
| RTF1 | 25 | 8.91 | 10.27 | 9.75 | 9.57 | 9.72 | 10.05 | 10.26 | 9.36 | 11.33 | 10.79 | 9.59 | 9.66 | 9.88 | 9.81 | 11.06 |
| RTFDC1 | 11 | 9.17 | 10.42 | 10.61 | 10.13 | 10.19 | 10.59 | 10.61 | 9.83 | 9.47 | 8.97 | 9.80 | 10.37 | 10.39 | 10.22 | 9.22 |
| RTN3 | 7 | 13.69 | 9.15 | 8.32 | 9.70 | 9.25 | 8.12 | 7.01 | 5.84 | 13.54 | 15.37 | 11.42 | 9.01 | 8.69 | 6.43 | 14.46 |
| RTN4 | 11 | 11.64 | 6.89 | 9.00 | 9.01 | 10.26 | 7.80 | 9.36 | 7.78 | 13.07 | 15.19 | 9.26 | 9.00 | 9.03 | 8.57 | 14.13 |
| RTN4IP1 | 1 | 11.57 | 9.98 | 7.66 | 9.17 | 9.57 | 9.66 | 10.38 | 10.90 | 10.52 | 10.60 | 10.77 | 8.42 | 9.61 | 10.64 | 10.56 |
| RTTN | 2 | 10.10 | 9.97 | 10.57 | 9.54 | 9.35 | 6.76 | 10.15 | 9.24 | 12.75 | 11.57 | 10.04 | 10.06 | 8.06 | 9.69 | 12.16 |
| RUFY1 | 17 | 8.58 | 9.15 | 8.71 | 8.36 | 13.71 | 13.51 | 11.27 | 10.42 | 8.50 | 7.79 | 8.86 | 8.54 | 13.61 | 10.84 | 8.14 |
| RUFY2 | 2 | 8.06 | 10.10 | 9.77 | 9.23 | 11.50 | 13.13 | 8.88 | 8.64 | 10.29 | 10.39 | 9.08 | 9.50 | 12.32 | 8.76 | 10.34 |
| RUFY3 | 1 | 9.40 | 12.37 | 8.69 | 14.96 | 7.19 | 9.68 | 8.16 | 9.48 | 9.29 | 10.76 | 10.89 | 11.83 | 8.44 | 8.82 | 10.03 |
| RUNX1 | 1 | 11.14 | 12.34 | 8.89 | 9.06 | 8.65 | 9.92 | 11.52 | 9.87 | 9.11 | 9.51 | 11.74 | 8.97 | 9.28 | 10.69 | 9.31 |
| RUNX3 | 1 | 10.53 | 12.50 | 11.19 | 12.17 | 6.99 | 6.54 | 10.76 | 10.67 | 10.88 | 7.77 | 11.51 | 11.68 | 6.77 | 10.71 | 9.33 |
| RUVBL1 | 49 | 8.54 | 9.04 | 9.97 | 8.75 | 10.63 | 10.44 | 10.27 | 14.70 | 8.96 | 8.69 | 8.79 | 9.36 | 10.54 | 12.49 | 8.82 |
| RUVBL2 | 51 | 8.16 | 9.18 | 9.98 | 9.25 | 10.54 | 10.64 | 10.48 | 12.90 | 9.49 | 9.39 | 8.67 | 9.62 | 10.59 | 11.69 | 9.44 |
| RUXGL | 8 | 9.35 | 10.29 | 10.21 | 9.80 | 11.07 | 10.68 | 10.11 | 9.11 | 9.55 | 9.82 | 9.82 | 10.01 | 10.87 | 9.61 | 9.68 |
| RWDD1 | 4 | 10.73 | 11.75 | 12.35 | 12.22 | 10.74 | 11.02 | 10.07 | 8.51 | 6.32 | 6.29 | 11.24 | 12.29 | 10.88 | 9.29 | 6.30 |
| RWDD2B | 2 | 11.10 | 11.37 | 11.18 | 9.02 | 10.56 | 8.74 | 8.43 | 5.89 | 11.16 | 12.54 | 11.24 | 10.10 | 9.65 | 7.16 | 11.85 |
| RWDD4 | 2 | 9.71 | 11.23 | 10.30 | 8.82 | 9.20 | 9.12 | 10.48 | 9.39 | 11.02 | 10.73 | 10.47 | 9.56 | 9.16 | 9.94 | 10.88 |
| RXRA | 2 | 10.52 | 9.89 | 7.85 | 6.77 | 7.54 | 7.58 | 16.16 | 17.89 | 7.86 | 7.95 | 10.20 | 7.31 | 7.56 | 17.02 | 7.91 |
| RXRB | 3 | 10.85 | 9.54 | 9.42 | 8.88 | 9.55 | 8.87 | 12.72 | 11.92 | 9.51 | 8.73 | 10.20 | 9.15 | 9.21 | 12.32 | 9.12 |
| RYBP | 3 | 9.24 | 10.05 | 10.14 | 11.24 | 8.87 | 10.08 | 12.61 | 12.01 | 8.00 | 7.76 | 9.65 | 10.69 | 9.47 | 12.31 | 7.88 |
| S100A10 | 11 | 8.05 | 8.82 | 10.16 | 9.93 | 10.45 | 10.92 | 12.03 | 11.07 | 9.48 | 9.10 | 8.44 | 10.04 | 10.68 | 11.55 | 9.29 |
| S100A11 | 12 | 8.27 | 8.99 | 8.73 | 8.33 | 14.46 | 14.57 | 10.35 | 8.65 | 9.08 | 8.56 | 8.63 | 8.53 | 14.52 | 9.50 | 8.82 |
| S100A13 | 17 | 8.95 | 9.85 | 7.29 | 6.77 | 14.22 | 14.48 | 13.52 | 11.05 | 7.18 | 6.69 | 9.40 | 7.03 | 14.35 | 12.28 | 6.94 |
| S100A14 | 2 | 9.22 | 7.67 | 1.19 | 0.69 | 14.71 | 13.91 | 24.88 | 12.37 | 7.78 | 7.57 | 8.45 | 0.94 | 14.31 | 18.62 | 7.67 |
| S100A16 | 5 | 8.53 | 8.82 | 5.75 | 3.59 | 16.52 | 13.73 | 15.52 | 11.01 | 8.49 | 8.03 | 8.68 | 4.67 | 15.13 | 13.27 | 8.26 |
| S100A2 | 5 | 8.05 | 8.99 | 4.46 | 4.75 | 9.64 | 10.18 | 22.58 | 8.21 | 11.39 | 11.75 | 8.52 | 4.61 | 9.91 | 15.40 | 11.57 |
| S100A3 | 2 | 9.85 | 10.85 | 4.10 | 4.16 | 10.38 | 9.21 | 14.59 | 7.72 | 14.22 | 14.92 | 10.35 | 4.13 | 9.79 | 11.15 | 14.57 |
| S100A4 | 24 | 8.53 | 11.96 | 2.41 | 2.09 | 14.95 | 14.89 | 14.25 | 11.99 | 9.42 | 9.52 | 10.24 | 2.25 | 14.92 | 13.12 | 9.47 |
| S100A6 | 29 | 9.55 | 12.08 | 9.41 | 11.59 | 11.68 | 12.91 | 8.86 | 2.79 | 10.66 | 10.46 | 10.82 | 10.50 | 12.29 | 5.83 | 10.56 |
| S100A7 | 1 | 17.64 | 10.07 | 12.87 | 9.19 | 8.33 | 7.70 | 5.06 | 4.26 | 10.64 | 14.24 | 13.85 | 11.03 | 8.01 | 4.66 | 12.44 |
| S100P | 2 | 12.32 | 13.06 | 2.81 | 2.34 | 28.66 | 30.07 | 1.70 | 1.04 | 4.28 | 3.73 | 12.69 | 2.57 | 29.36 | 1.37 | 4.01 |
| SAAL1 | 7 | 9.86 | 9.53 | 11.98 | 11.60 | 8.23 | 8.32 | 9.97 | 10.23 | 10.32 | 9.95 | 9.69 | 11.79 | 8.28 | 10.10 | 10.14 |
| SAC3D1 | 1 | 8.84 | 10.52 | 9.56 | 10.62 | 10.45 | 10.59 | 10.81 | 8.69 | 10.33 | 9.59 | 9.68 | 10.09 | 10.52 | 9.75 | 9.96 |
| SACM1L | 8 | 12.56 | 6.89 | 6.93 | 8.43 | 10.47 | 8.27 | 10.99 | 8.94 | 11.97 | 14.55 | 9.72 | 7.68 | 9.37 | 9.96 | 13.26 |
| SACS | 7 | 8.45 | 8.50 | 16.14 | 16.80 | 8.78 | 8.39 | 8.81 | 8.22 | 8.09 | 7.82 | 8.48 | 16.47 | 8.59 | 8.52 | 7.95 |
| SAE1 | 31 | 9.26 | 9.69 | 9.23 | 8.59 | 8.39 | 8.14 | 12.49 | 17.11 | 8.79 | 8.32 | 9.47 | 8.91 | 8.27 | 14.80 | 8.55 |
| SAFB | 20 | 9.01 | 9.27 | 11.66 | 11.01 | 9.87 | 9.77 | 11.06 | 10.57 | 9.29 | 8.49 | 9.14 | 11.34 | 9.82 | 10.82 | 8.89 |
| SAFB2 | 26 | 8.80 | 9.62 | 10.23 | 9.79 | 10.08 | 10.09 | 10.89 | 13.12 | 9.05 | 8.33 | 9.21 | 10.01 | 10.08 | 12.00 | 8.69 |
| SAMD1 | 5 | 8.88 | 11.19 | 8.81 | 8.54 | 10.87 | 11.86 | 11.56 | 10.73 | 8.91 | 8.65 | 10.04 | 8.67 | 11.36 | 11.15 | 8.78 |
| SAMD11 | 5 | 14.81 | 14.88 | 4.13 | 4.07 | 16.77 | 16.32 | 10.15 | 7.53 | 5.99 | 5.37 | 14.84 | 4.10 | 16.54 | 8.84 | 5.68 |
| SAMD4B | 3 | 10.42 | 10.34 | 8.72 | 7.79 | 11.67 | 11.93 | 12.63 | 11.12 | 7.81 | 7.56 | 10.38 | 8.26 | 11.80 | 11.87 | 7.69 |
| SAMD9 | 7 | 11.03 | 10.44 | 6.93 | 7.51 | 11.89 | 12.31 | 9.16 | 8.19 | 11.33 | 11.20 | 10.74 | 7.22 | 12.10 | 8.67 | 11.27 |

| | | | | | | | | | | | | | | | | |
|---|---|---|---|---|---|---|---|---|---|---|---|---|---|---|---|---|
| SAMHD1 | 22 | 11.50 | 12.01 | 10.19 | 10.28 | 11.86 | 12.30 | 6.68 | 6.35 | 9.46 | 9.37 | 11.75 | 10.24 | 12.08 | 6.52 | 9.41 |
| SAMM50 | 11 | 13.33 | 6.66 | 8.52 | 13.51 | 8.09 | 7.08 | 9.67 | 9.45 | 10.62 | 13.08 | 9.99 | 11.01 | 7.58 | 9.56 | 11.85 |
| SAP130 | 5 | 9.82 | 10.01 | 10.02 | 10.74 | 10.05 | 10.27 | 11.33 | 10.17 | 9.19 | 8.41 | 9.91 | 10.38 | 10.16 | 10.75 | 8.80 |
| SAP18 | 12 | 9.45 | 10.87 | 9.59 | 9.84 | 10.63 | 11.84 | 10.35 | 10.30 | 8.47 | 8.66 | 10.16 | 9.71 | 11.23 | 10.33 | 8.57 |
| SAP30 | 2 | 9.88 | 10.42 | 9.71 | 9.30 | 10.18 | 10.17 | 10.88 | 9.48 | 10.19 | 9.79 | 10.15 | 9.51 | 10.18 | 10.18 | 9.99 |
| SAP30BP | 8 | 10.08 | 10.75 | 10.90 | 10.81 | 8.72 | 8.83 | 10.35 | 9.51 | 10.46 | 9.59 | 10.41 | 10.86 | 8.77 | 9.93 | 10.03 |
| SAPCD2 | 1 | 11.25 | 8.87 | 7.19 | 7.26 | 9.75 | 8.64 | 10.60 | 9.86 | 14.41 | 12.17 | 10.06 | 7.23 | 9.20 | 10.23 | 13.29 |
| SAR1A | 6 | 8.50 | 8.13 | 12.77 | 14.39 | 9.57 | 9.35 | 9.45 | 8.47 | 9.57 | 9.81 | 8.31 | 13.58 | 9.46 | 8.96 | 9.69 |
| SAR1B | 15 | 9.03 | 9.73 | 9.99 | 9.54 | 10.74 | 9.78 | 10.87 | 9.67 | 10.12 | 10.52 | 9.38 | 9.77 | 10.26 | 10.27 | 10.32 |
| SARNP | 14 | 9.48 | 10.06 | 9.72 | 10.31 | 10.77 | 11.84 | 10.38 | 9.73 | 9.09 | 8.64 | 9.77 | 10.02 | 11.30 | 10.05 | 8.87 |
| SARS | 25 | 9.77 | 11.16 | 10.41 | 9.69 | 7.90 | 8.37 | 10.05 | 11.55 | 10.69 | 10.41 | 10.47 | 10.05 | 8.14 | 10.80 | 10.55 |
| SARS2 | 8 | 11.85 | 10.12 | 7.50 | 8.75 | 9.31 | 8.25 | 11.20 | 10.85 | 11.06 | 11.11 | 10.98 | 8.13 | 8.78 | 11.03 | 11.09 |
| SART1 | 39 | 9.48 | 10.23 | 11.47 | 10.78 | 9.23 | 9.38 | 10.58 | 9.53 | 9.83 | 9.49 | 9.85 | 11.13 | 9.30 | 10.06 | 9.66 |
| SART3 | 31 | 8.65 | 10.07 | 11.14 | 10.37 | 9.63 | 9.86 | 10.17 | 10.28 | 10.08 | 9.75 | 9.36 | 10.76 | 9.74 | 10.22 | 9.91 |
| SASH1 | 2 | 10.83 | 11.04 | 8.08 | 9.72 | 11.55 | 11.96 | 7.40 | 7.21 | 10.62 | 11.57 | 10.94 | 8.90 | 11.76 | 7.31 | 11.10 |
| SASS6 | 7 | 9.76 | 10.46 | 9.83 | 10.07 | 8.59 | 8.48 | 11.54 | 10.80 | 10.57 | 9.91 | 10.11 | 9.95 | 8.53 | 11.17 | 10.24 |
| SBDS | 32 | 9.12 | 9.61 | 11.42 | 10.57 | 9.27 | 9.31 | 8.36 | 13.23 | 9.66 | 9.44 | 9.37 | 11.00 | 9.29 | 10.79 | 9.55 |
| SBF1 | 7 | 9.59 | 9.76 | 9.85 | 10.35 | 8.89 | 9.17 | 11.32 | 11.13 | 10.31 | 9.64 | 9.67 | 10.10 | 9.03 | 11.22 | 9.98 |
| SBF2 | 1 | 10.32 | 12.11 | 11.90 | 9.85 | 10.10 | 9.69 | 9.81 | 8.38 | 9.44 | 8.39 | 11.22 | 10.87 | 9.90 | 9.10 | 8.92 |
| SBNO1 | 14 | 8.50 | 9.74 | 10.61 | 11.91 | 8.63 | 8.96 | 10.89 | 10.93 | 9.80 | 10.03 | 9.12 | 11.26 | 8.79 | 10.91 | 9.91 |
| SBNO2 | 1 | 7.04 | 9.06 | 7.26 | 6.54 | 10.69 | 12.03 | 14.45 | 16.95 | 7.92 | 8.04 | 8.05 | 6.90 | 11.36 | 15.70 | 7.98 |
| SC5D | 2 | 13.00 | 10.08 | 8.74 | 10.92 | 11.38 | 10.04 | 7.74 | 5.55 | 10.93 | 11.61 | 11.54 | 9.83 | 10.71 | 6.65 | 11.27 |
| SCAF1 | 5 | 8.72 | 11.02 | 9.10 | 10.45 | 9.07 | 10.14 | 10.73 | 11.12 | 9.34 | 10.31 | 9.87 | 9.77 | 9.61 | 10.93 | 9.83 |
| SCAF11 | 18 | 8.55 | 10.13 | 10.56 | 10.54 | 9.66 | 10.36 | 11.41 | 10.83 | 9.00 | 8.97 | 9.34 | 10.55 | 10.01 | 11.12 | 8.99 |
| SCAF4 | 8 | 10.66 | 12.03 | 6.58 | 6.14 | 11.46 | 12.77 | 10.11 | 9.49 | 10.90 | 9.84 | 11.34 | 6.36 | 12.12 | 9.80 | 10.37 |
| SCAF8 | 17 | 9.52 | 10.20 | 9.53 | 9.08 | 11.04 | 11.27 | 10.26 | 9.95 | 9.71 | 9.43 | 9.86 | 9.30 | 11.16 | 10.11 | 9.57 |
| SCAI | 5 | 9.98 | 9.86 | 9.99 | 9.53 | 11.46 | 10.66 | 9.48 | 9.00 | 10.06 | 9.98 | 9.92 | 9.76 | 11.06 | 9.24 | 10.02 |
| SCAMP1 | 5 | 9.89 | 8.27 | 6.73 | 8.54 | 9.12 | 9.07 | 17.18 | 15.90 | 7.18 | 8.12 | 9.08 | 7.63 | 9.09 | 16.54 | 7.65 |
| SCAMP2 | 2 | 10.33 | 8.18 | 7.26 | 10.29 | 11.43 | 12.03 | 8.42 | 7.74 | 11.94 | 12.39 | 9.25 | 8.77 | 11.73 | 8.08 | 12.16 |
| SCAMP3 | 10 | 9.44 | 7.89 | 9.48 | 10.27 | 13.36 | 12.73 | 9.14 | 8.44 | 9.60 | 9.66 | 8.67 | 9.88 | 13.04 | 8.79 | 9.63 |
| SCAMP4 | 3 | 9.83 | 8.12 | 8.48 | 11.50 | 10.13 | 9.99 | 8.46 | 9.17 | 11.43 | 12.89 | 8.98 | 9.99 | 10.06 | 8.81 | 12.16 |
| SCAPER | 1 | 9.83 | 10.32 | 9.38 | 9.77 | 9.81 | 10.59 | 10.90 | 10.47 | 9.53 | 9.41 | 10.07 | 9.57 | 10.20 | 10.69 | 9.47 |
| SCARB1 | 7 | 13.21 | 8.93 | 7.19 | 9.13 | 9.53 | 9.61 | 8.50 | 7.82 | 12.24 | 13.85 | 11.07 | 8.16 | 9.57 | 8.16 | 13.04 |
| SCARB2 | 4 | 9.79 | 6.50 | 8.09 | 9.93 | 11.94 | 11.43 | 12.20 | 11.82 | 8.53 | 9.78 | 8.14 | 9.01 | 11.69 | 12.01 | 9.15 |
| SCCPDH | 5 | 12.60 | 9.59 | 7.08 | 8.09 | 13.09 | 12.00 | 8.22 | 7.81 | 10.50 | 11.01 | 11.10 | 7.59 | 12.55 | 8.02 | 10.76 |
| SCD | 4 | 13.77 | 6.08 | 5.19 | 9.84 | 10.33 | 10.17 | 9.16 | 9.40 | 11.38 | 14.67 | 9.92 | 7.52 | 10.25 | 9.28 | 13.03 |
| SCFD1 | 29 | 9.97 | 10.81 | 9.54 | 9.36 | 10.71 | 10.98 | 8.99 | 8.30 | 10.83 | 10.50 | 10.39 | 9.45 | 10.84 | 8.65 | 10.67 |
| SCFD2 | 2 | 9.55 | 10.02 | 10.70 | 10.71 | 11.05 | 10.48 | 9.80 | 8.59 | 9.62 | 9.50 | 9.78 | 10.70 | 10.77 | 9.19 | 9.56 |
| SCLT1 | 2 | 10.48 | 11.14 | 9.51 | 10.23 | 10.12 | 11.22 | 11.88 | 10.75 | 7.61 | 7.06 | 10.81 | 9.87 | 10.67 | 11.32 | 7.33 |
| SCLY | 5 | 11.56 | 11.86 | 13.61 | 13.16 | 10.41 | 10.79 | 11.72 | 10.88 | 3.21 | 2.79 | 11.71 | 13.39 | 10.60 | 11.30 | 3.00 |
| SCML2 | 5 | 9.67 | 11.12 | 10.04 | 8.22 | 11.46 | 12.18 | 9.54 | 8.07 | 10.37 | 9.32 | 10.40 | 9.13 | 11.82 | 8.80 | 9.85 |
| SCNM1 | 2 | 9.86 | 10.98 | 11.12 | 9.83 | 10.28 | 9.78 | 11.02 | 9.00 | 9.09 | 9.05 | 10.42 | 10.47 | 10.03 | 10.01 | 9.07 |

| | | | | | | | | | | | | | | | | |
|---|---|---|---|---|---|---|---|---|---|---|---|---|---|---|---|---|
| SCO1 | 4 | 13.90 | 6.91 | 9.94 | 14.51 | 6.85 | 6.96 | 8.42 | 8.51 | 10.58 | 13.41 | 10.41 | 12.22 | 6.91 | 8.47 | 12.00 |
| SCO2 | 4 | 12.63 | 7.77 | 8.97 | 12.45 | 8.78 | 8.92 | 9.35 | 9.55 | 10.35 | 11.24 | 10.20 | 10.71 | 8.85 | 9.45 | 10.79 |
| SCP2 | 17 | 9.47 | 10.35 | 10.75 | 10.33 | 9.51 | 9.74 | 10.04 | 9.54 | 10.37 | 9.90 | 9.91 | 10.54 | 9.63 | 9.79 | 10.13 |
| SCPEP1 | 11 | 9.66 | 10.67 | 12.17 | 11.94 | 12.38 | 12.75 | 7.36 | 7.15 | 8.24 | 7.68 | 10.17 | 12.06 | 12.57 | 7.25 | 7.96 |
| SCRIB | 38 | 10.67 | 9.44 | 9.04 | 9.98 | 8.43 | 8.84 | 11.45 | 11.56 | 10.46 | 10.14 | 10.05 | 9.51 | 8.64 | 11.51 | 10.30 |
| SCRN1 | 19 | 9.06 | 10.30 | 14.17 | 13.21 | 8.63 | 8.68 | 7.30 | 7.34 | 10.64 | 10.67 | 9.68 | 13.69 | 8.66 | 7.32 | 10.65 |
| SCRN2 | 4 | 11.63 | 12.08 | 11.20 | 10.52 | 9.83 | 9.72 | 7.32 | 6.73 | 10.80 | 10.17 | 11.85 | 10.86 | 9.78 | 7.03 | 10.48 |
| SCRN3 | 3 | 10.40 | 11.39 | 8.65 | 9.51 | 11.49 | 12.48 | 7.18 | 7.06 | 11.73 | 10.11 | 10.90 | 9.08 | 11.99 | 7.12 | 10.92 |
| SCYL1 | 20 | 10.33 | 10.84 | 10.01 | 9.70 | 10.43 | 10.71 | 9.49 | 8.97 | 10.07 | 9.46 | 10.58 | 9.86 | 10.57 | 9.23 | 9.76 |
| SCYL2 | 13 | 9.77 | 10.44 | 10.09 | 10.40 | 10.53 | 10.67 | 9.61 | 8.89 | 10.05 | 9.57 | 10.10 | 10.24 | 10.60 | 9.25 | 9.81 |
| SDAD1 | 11 | 9.26 | 9.10 | 10.26 | 10.99 | 7.98 | 7.97 | 10.98 | 13.78 | 9.70 | 9.98 | 9.18 | 10.62 | 7.97 | 12.38 | 9.84 |
| SDC1 | 3 | 12.55 | 11.57 | 7.79 | 9.01 | 7.53 | 6.81 | 6.11 | 5.43 | 17.13 | 16.06 | 12.06 | 8.40 | 7.17 | 5.77 | 16.59 |
| SDC2 | 7 | 11.90 | 10.40 | 10.10 | 12.03 | 10.75 | 11.18 | 3.74 | 3.49 | 13.31 | 13.11 | 11.15 | 11.06 | 10.96 | 3.61 | 13.21 |
| SDCBP | 7 | 10.43 | 10.61 | 7.54 | 8.10 | 12.70 | 13.30 | 8.74 | 8.00 | 10.48 | 10.09 | 10.52 | 7.82 | 13.00 | 8.37 | 10.29 |
| SDCCAG3 | 3 | 10.02 | 10.99 | 8.27 | 7.87 | 11.84 | 11.73 | 9.70 | 10.26 | 10.09 | 9.23 | 10.50 | 8.07 | 11.78 | 9.98 | 9.66 |
| SDCCAG8 | 1 | 10.25 | 12.69 | 6.25 | 8.63 | 13.29 | 12.75 | 9.92 | 9.78 | 8.82 | 7.63 | 11.47 | 7.44 | 13.02 | 9.85 | 8.23 |
| SDE2 | 5 | 9.65 | 9.84 | 8.84 | 8.18 | 11.78 | 12.80 | 11.66 | 9.68 | 9.13 | 8.45 | 9.74 | 8.51 | 12.29 | 10.67 | 8.79 |
| SDF2 | 4 | 10.04 | 9.09 | 13.74 | 9.38 | 12.71 | 11.09 | 8.09 | 6.37 | 9.87 | 9.62 | 9.57 | 11.56 | 11.90 | 7.23 | 9.74 |
| SDF2L1 | 6 | 10.17 | 9.61 | 9.68 | 8.89 | 11.00 | 10.03 | 10.89 | 12.72 | 8.53 | 8.49 | 9.89 | 9.29 | 10.51 | 11.81 | 8.51 |
| SDF4 | 14 | 11.05 | 9.02 | 8.26 | 8.92 | 9.46 | 8.98 | 15.98 | 13.88 | 6.99 | 7.44 | 10.04 | 8.59 | 9.22 | 14.93 | 7.21 |
| SDHA | 39 | 10.54 | 9.45 | 7.69 | 7.83 | 10.79 | 9.76 | 9.71 | 11.91 | 10.47 | 11.85 | 10.00 | 7.76 | 10.28 | 10.81 | 11.16 |
| SDHAF2 | 2 | 12.97 | 11.42 | 11.51 | 10.51 | 12.09 | 11.83 | 8.26 | 3.83 | 9.16 | 8.42 | 12.20 | 11.01 | 11.96 | 6.05 | 8.79 |
| SDHB | 15 | 10.16 | 9.03 | 9.40 | 9.11 | 8.71 | 8.08 | 14.16 | 11.73 | 9.49 | 10.14 | 9.59 | 9.26 | 8.39 | 12.94 | 9.81 |
| SDHC | 1 | 3.68 | 4.19 | 3.79 | 4.06 | 4.18 | 4.06 | 4.06 | 63.59 | 4.66 | 3.73 | 3.94 | 3.92 | 4.12 | 33.82 | 4.20 |
| SDPR | 1 | 8.88 | 10.64 | 12.96 | 13.42 | 7.91 | 7.47 | 6.90 | 5.28 | 13.74 | 12.81 | 9.76 | 13.19 | 7.69 | 6.09 | 13.27 |
| SDR39U1 | 3 | 10.13 | 7.98 | 7.65 | 8.13 | 14.34 | 12.84 | 10.40 | 9.40 | 9.80 | 9.33 | 9.05 | 7.89 | 13.59 | 9.90 | 9.57 |
| SDSL | 3 | 10.69 | 13.58 | 12.21 | 13.10 | 8.98 | 9.03 | 7.57 | 7.53 | 8.56 | 8.74 | 12.13 | 12.66 | 9.01 | 7.55 | 8.65 |
| SEC11A | 8 | 11.68 | 7.65 | 8.12 | 11.22 | 8.98 | 9.47 | 9.75 | 8.94 | 11.62 | 12.59 | 9.66 | 9.67 | 9.22 | 9.34 | 12.10 |
| SEC11C | 2 | 12.42 | 10.34 | 9.14 | 10.29 | 12.30 | 14.78 | 8.65 | 8.45 | 6.98 | 6.63 | 11.38 | 9.72 | 13.54 | 8.55 | 6.81 |
| SEC13 | 9 | 7.73 | 9.12 | 9.28 | 8.50 | 9.95 | 9.85 | 8.56 | 20.73 | 8.34 | 7.94 | 8.42 | 8.89 | 9.90 | 14.65 | 8.14 |
| SEC14L1 | 5 | 9.19 | 9.75 | 12.00 | 12.42 | 8.66 | 9.71 | 7.68 | 7.43 | 11.73 | 11.42 | 9.47 | 12.21 | 9.19 | 7.55 | 11.58 |
| SEC14L2 | 1 | 9.63 | 10.03 | 17.45 | 17.71 | 4.01 | 5.29 | 7.65 | 6.76 | 11.59 | 9.89 | 9.83 | 17.58 | 4.65 | 7.20 | 10.74 |
| SEC16A | 45 | 9.17 | 10.41 | 10.24 | 10.12 | 11.73 | 11.91 | 8.89 | 8.16 | 9.85 | 9.53 | 9.79 | 10.18 | 11.82 | 8.52 | 9.69 |
| SEC22B | 20 | 12.91 | 8.06 | 7.42 | 10.35 | 9.31 | 8.46 | 8.86 | 8.48 | 11.84 | 14.31 | 10.48 | 8.89 | 8.88 | 8.67 | 13.08 |
| SEC23A | 16 | 9.92 | 9.72 | 9.56 | 10.34 | 9.61 | 10.01 | 9.57 | 11.22 | 10.60 | 9.45 | 9.82 | 9.95 | 9.81 | 10.40 | 10.03 |
| SEC23B | 16 | 8.98 | 9.58 | 9.19 | 8.55 | 10.22 | 10.26 | 10.35 | 15.13 | 9.12 | 8.63 | 9.28 | 8.87 | 10.24 | 12.74 | 8.87 |
| SEC23IP | 27 | 9.82 | 10.63 | 11.36 | 10.48 | 9.16 | 9.02 | 9.87 | 10.72 | 9.77 | 9.16 | 10.22 | 10.92 | 9.09 | 10.30 | 9.47 |
| SEC24A | 10 | 9.43 | 10.31 | 8.80 | 8.64 | 8.90 | 9.36 | 11.27 | 15.00 | 9.25 | 9.05 | 9.87 | 8.72 | 9.13 | 13.14 | 9.15 |
| SEC24B | 14 | 8.52 | 9.39 | 10.13 | 9.97 | 11.60 | 11.07 | 10.31 | 10.12 | 9.53 | 9.37 | 8.96 | 10.05 | 11.33 | 10.22 | 9.45 |
| SEC24C | 25 | 10.10 | 10.20 | 10.82 | 8.88 | 10.74 | 10.59 | 9.02 | 8.74 | 10.50 | 10.40 | 10.15 | 9.85 | 10.67 | 8.88 | 10.45 |
| SEC24D | 17 | 10.10 | 10.80 | 10.44 | 10.23 | 9.78 | 9.95 | 9.64 | 9.39 | 9.93 | 9.75 | 10.45 | 10.33 | 9.86 | 9.51 | 9.84 |
| SEC31A | 40 | 10.77 | 10.87 | 9.82 | 9.43 | 11.05 | 11.07 | 9.26 | 9.60 | 9.19 | 8.94 | 10.82 | 9.62 | 11.06 | 9.43 | 9.07 |

| | | | | | | | | | | | | | | | | |
|---|---|---|---|---|---|---|---|---|---|---|---|---|---|---|---|---|
| SEC61A1 | 5 | 14.82 | 6.90 | 6.31 | 10.88 | 8.51 | 8.14 | 9.94 | 9.02 | 10.12 | 15.35 | 10.86 | 8.59 | 8.33 | 9.48 | 12.74 |
| SEC61B | 3 | 13.01 | 6.83 | 8.30 | 11.56 | 9.00 | 7.94 | 9.66 | 8.55 | 10.55 | 14.61 | 9.92 | 9.93 | 8.47 | 9.10 | 12.58 |
| SEC61G | 1 | 9.68 | 10.15 | 9.84 | 6.07 | 9.87 | 9.44 | 10.83 | 15.68 | 9.72 | 8.72 | 9.92 | 7.95 | 9.66 | 13.26 | 9.22 |
| SEC62 | 4 | 13.64 | 6.05 | 8.28 | 12.87 | 9.47 | 7.29 | 9.99 | 7.89 | 11.65 | 12.86 | 9.84 | 10.58 | 8.38 | 8.94 | 12.26 |
| SEC63 | 17 | 12.84 | 6.04 | 7.13 | 11.07 | 9.73 | 9.00 | 10.70 | 11.29 | 9.90 | 12.30 | 9.44 | 9.10 | 9.37 | 11.00 | 11.10 |
| SECISBP2L | 1 | 9.49 | 9.68 | 8.78 | 9.00 | 8.83 | 11.96 | 12.06 | 12.25 | 8.67 | 9.28 | 9.59 | 8.89 | 10.39 | 12.16 | 8.98 |
| SEH1L | 11 | 11.11 | 11.38 | 9.95 | 9.88 | 9.27 | 9.92 | 9.25 | 9.34 | 10.08 | 9.82 | 11.24 | 9.92 | 9.60 | 9.30 | 9.95 |
| SEL1L | 4 | 10.53 | 5.94 | 8.84 | 12.87 | 12.16 | 10.21 | 10.15 | 8.98 | 9.82 | 10.50 | 8.23 | 10.86 | 11.19 | 9.56 | 10.16 |
| SELENBP1 | 26 | 10.56 | 11.58 | 6.51 | 6.41 | 18.23 | 18.55 | 6.12 | 5.61 | 8.42 | 8.01 | 11.07 | 6.46 | 18.39 | 5.86 | 8.22 |
| SELH | 2 | 9.47 | 10.50 | 11.64 | 10.89 | 11.58 | 12.23 | 10.76 | 10.07 | 6.50 | 6.36 | 9.98 | 11.26 | 11.90 | 10.42 | 6.43 |
| SELK | 2 | 12.80 | 8.91 | 7.36 | 7.23 | 10.46 | 10.01 | 10.42 | 10.72 | 11.45 | 10.64 | 10.86 | 7.29 | 10.24 | 10.57 | 11.04 |
| SELM | 1 | 11.16 | 12.39 | 6.03 | 6.53 | 10.78 | 9.64 | 14.98 | 10.52 | 8.60 | 9.36 | 11.78 | 6.28 | 10.21 | 12.75 | 8.98 |
| SELO | 1 | 9.00 | 9.68 | 16.40 | 14.29 | 7.77 | 7.97 | 9.70 | 7.22 | 10.42 | 7.56 | 9.34 | 15.35 | 7.87 | 8.46 | 8.99 |
| SELRC1 | 5 | 10.08 | 9.16 | 10.91 | 10.02 | 8.31 | 8.38 | 13.78 | 11.82 | 8.96 | 8.58 | 9.62 | 10.46 | 8.34 | 12.80 | 8.77 |
| SEMA3B | 3 | 9.53 | 8.35 | 5.57 | 6.27 | 11.60 | 12.54 | 14.42 | 13.66 | 9.13 | 8.93 | 8.94 | 5.92 | 12.07 | 14.04 | 9.03 |
| SEMA3C | 2 | 9.23 | 9.52 | 13.50 | 12.46 | 8.62 | 6.99 | 8.83 | 8.05 | 11.09 | 11.72 | 9.38 | 12.98 | 7.80 | 8.44 | 11.40 |
| SEMA4B | 1 | 8.43 | 8.96 | 8.22 | 7.94 | 8.75 | 9.79 | 15.95 | 13.62 | 9.85 | 8.49 | 8.69 | 8.08 | 9.27 | 14.78 | 9.17 |
| SENP1 | 3 | 9.68 | 9.93 | 9.80 | 9.28 | 10.13 | 9.32 | 12.10 | 10.75 | 8.98 | 10.02 | 9.80 | 9.54 | 9.73 | 11.43 | 9.50 |
| SENP3 | 4 | 9.25 | 10.72 | 10.32 | 9.82 | 8.87 | 9.07 | 9.65 | 11.36 | 10.82 | 10.12 | 9.99 | 10.07 | 8.97 | 10.51 | 10.47 |
| SENP6 | 5 | 9.05 | 9.69 | 11.05 | 10.11 | 8.51 | 8.36 | 12.82 | 11.36 | 9.94 | 9.11 | 9.37 | 10.58 | 8.43 | 12.09 | 9.53 |
| SEPHS1 | 9 | 9.81 | 10.34 | 11.17 | 8.42 | 11.26 | 9.89 | 11.60 | 9.17 | 9.28 | 9.06 | 10.07 | 9.80 | 10.57 | 10.39 | 9.17 |
| SEPHS2 | 1 | 9.42 | 10.77 | 9.00 | 10.24 | 11.51 | 10.29 | 10.67 | 9.75 | 8.31 | 10.04 | 10.09 | 9.62 | 10.90 | 10.21 | 9.18 |
| SEPT10 | 16 | 10.15 | 9.70 | 9.30 | 9.07 | 9.71 | 10.33 | 10.45 | 10.66 | 10.52 | 10.12 | 9.92 | 9.18 | 10.02 | 10.55 | 10.32 |
| SEPT11 | 17 | 9.70 | 10.39 | 12.28 | 11.93 | 8.63 | 8.87 | 8.71 | 8.08 | 11.03 | 10.37 | 10.05 | 12.11 | 8.75 | 8.39 | 10.70 |
| SEPT2 | 30 | 9.42 | 10.61 | 12.46 | 11.33 | 9.40 | 9.13 | 9.05 | 8.37 | 10.31 | 9.93 | 10.01 | 11.90 | 9.27 | 8.71 | 10.12 |
| SEPT6 | 6 | 10.27 | 11.47 | 11.67 | 10.19 | 10.47 | 10.62 | 7.67 | 6.41 | 10.89 | 10.34 | 10.87 | 10.93 | 10.55 | 7.04 | 10.62 |
| SEPT7 | 36 | 9.44 | 10.52 | 10.88 | 11.30 | 8.87 | 9.65 | 8.72 | 9.70 | 10.43 | 10.49 | 9.98 | 11.09 | 9.26 | 9.21 | 10.46 |
| SEPT8 | 12 | 9.91 | 10.98 | 11.09 | 10.58 | 9.29 | 9.77 | 9.05 | 8.59 | 10.45 | 10.28 | 10.45 | 10.84 | 9.53 | 8.82 | 10.36 |
| SEPT9 | 56 | 9.97 | 10.63 | 11.11 | 10.68 | 9.17 | 9.46 | 8.54 | 8.26 | 11.23 | 10.94 | 10.30 | 10.90 | 9.32 | 8.40 | 11.08 |
| SEPW1 | 1 | 8.44 | 11.24 | 9.18 | 10.39 | 11.70 | 14.24 | 8.48 | 10.23 | 6.99 | 9.11 | 9.84 | 9.79 | 12.97 | 9.35 | 8.05 |
| SERBP1 | 62 | 10.64 | 11.25 | 9.45 | 8.91 | 8.81 | 9.11 | 11.42 | 10.23 | 10.36 | 9.82 | 10.95 | 9.18 | 8.96 | 10.82 | 10.09 |
| SERF2 | 5 | 8.77 | 9.45 | 8.56 | 8.71 | 11.08 | 11.13 | 10.19 | 8.75 | 11.82 | 11.54 | 9.11 | 8.64 | 11.10 | 9.47 | 11.68 |
| SERINC1 | 2 | 10.15 | 8.88 | 10.74 | 8.01 | 10.55 | 10.05 | 9.39 | 7.12 | 11.12 | 13.99 | 9.52 | 9.37 | 10.30 | 8.25 | 12.55 |
| SERPINB1 | 17 | 9.01 | 8.83 | 8.28 | 7.83 | 13.81 | 13.47 | 12.57 | 11.53 | 7.50 | 7.18 | 8.92 | 8.05 | 13.64 | 12.05 | 7.34 |
| SERPINB3 | 4 | 3.86 | 3.15 | 3.76 | 3.36 | 16.74 | 15.95 | 25.40 | 21.89 | 3.27 | 2.62 | 3.50 | 3.56 | 16.34 | 23.64 | 2.95 |
| SERPINB5 | 37 | 9.73 | 10.48 | 10.03 | 8.64 | 18.87 | 18.16 | 2.94 | 2.54 | 9.44 | 9.16 | 10.11 | 9.34 | 18.51 | 2.74 | 9.30 |
| SERPINB6 | 43 | 7.97 | 8.65 | 7.47 | 7.17 | 13.53 | 13.63 | 12.94 | 15.48 | 6.66 | 6.50 | 8.31 | 7.32 | 13.58 | 14.21 | 6.58 |
| SERPINB8 | 8 | 8.90 | 9.74 | 11.18 | 11.17 | 14.22 | 14.76 | 5.96 | 7.53 | 8.51 | 8.02 | 9.32 | 11.18 | 14.49 | 6.75 | 8.27 |
| SERPINB9 | 6 | 8.94 | 10.20 | 7.16 | 7.57 | 17.87 | 18.45 | 8.13 | 7.89 | 6.99 | 6.79 | 9.57 | 7.36 | 18.16 | 8.01 | 6.89 |
| SERPINE1 | 3 | 6.44 | 7.22 | 18.53 | 23.74 | 5.00 | 4.31 | 6.28 | 5.19 | 11.28 | 12.01 | 6.83 | 21.14 | 4.65 | 5.73 | 11.65 |
| SERPINE2 | 8 | 3.39 | 3.28 | 21.34 | 24.26 | 3.35 | 3.59 | 11.04 | 12.36 | 8.49 | 8.90 | 3.33 | 22.80 | 3.47 | 11.70 | 8.70 |
| SERPINH1 | 76 | 7.68 | 7.30 | 14.09 | 14.44 | 10.18 | 9.75 | 7.82 | 7.60 | 10.43 | 10.69 | 7.49 | 14.27 | 9.97 | 7.71 | 10.56 |

| | | | | | | | | | | | | | | | | |
|---|---|---|---|---|---|---|---|---|---|---|---|---|---|---|---|---|
| SERPINI1 | 1 | 9.85 | 9.92 | 12.87 | 16.15 | 9.46 | 10.27 | 5.74 | 6.06 | 9.76 | 9.91 | 9.89 | 14.51 | 9.86 | 5.90 | 9.84 |
| SESTD1 | 7 | 9.56 | 10.24 | 10.03 | 10.76 | 10.98 | 11.63 | 7.47 | 7.62 | 11.19 | 10.52 | 9.90 | 10.40 | 11.30 | 7.54 | 10.85 |
| SET | 54 | 8.54 | 11.08 | 11.76 | 10.67 | 9.76 | 9.63 | 9.93 | 9.13 | 9.89 | 9.62 | 9.81 | 11.21 | 9.69 | 9.53 | 9.75 |
| SETD1A | 5 | 8.83 | 10.32 | 10.33 | 10.04 | 11.01 | 11.21 | 11.02 | 10.36 | 8.50 | 8.37 | 9.58 | 10.18 | 11.11 | 10.69 | 8.43 |
| SETD2 | 7 | 9.57 | 10.36 | 10.15 | 8.46 | 10.84 | 10.00 | 11.49 | 10.12 | 9.45 | 9.57 | 9.96 | 9.31 | 10.42 | 10.80 | 9.51 |
| SETD3 | 14 | 9.42 | 9.97 | 10.13 | 8.64 | 10.77 | 10.79 | 9.85 | 9.34 | 10.89 | 10.18 | 9.70 | 9.39 | 10.78 | 9.60 | 10.54 |
| SETD7 | 9 | 11.04 | 11.66 | 9.06 | 7.55 | 12.56 | 12.84 | 8.48 | 7.13 | 10.34 | 9.34 | 11.35 | 8.31 | 12.70 | 7.81 | 9.84 |
| SETD8 | 1 | 9.60 | 9.44 | 8.29 | 8.39 | 9.53 | 10.34 | 11.58 | 10.35 | 11.30 | 11.19 | 9.52 | 8.34 | 9.94 | 10.97 | 11.24 |
| SETDB1 | 7 | 9.06 | 10.29 | 10.51 | 9.79 | 11.99 | 12.47 | 9.61 | 8.40 | 9.26 | 8.63 | 9.67 | 10.15 | 12.23 | 9.01 | 8.94 |
| SETMAR | 2 | 9.86 | 10.54 | 7.48 | 7.28 | 11.46 | 11.96 | 12.58 | 11.71 | 8.74 | 8.39 | 10.20 | 7.38 | 11.71 | 12.14 | 8.56 |
| SETX | 8 | 8.11 | 10.89 | 12.11 | 12.01 | 9.03 | 9.49 | 8.72 | 8.34 | 10.33 | 10.98 | 9.50 | 12.06 | 9.26 | 8.53 | 10.65 |
| SF1 | 25 | 9.52 | 10.27 | 9.97 | 9.70 | 9.81 | 10.55 | 11.17 | 10.35 | 9.50 | 9.15 | 9.90 | 9.84 | 10.18 | 10.76 | 9.33 |
| SF3A1 | 51 | 9.91 | 11.10 | 10.50 | 10.17 | 8.71 | 9.39 | 10.17 | 10.33 | 9.88 | 9.84 | 10.51 | 10.33 | 9.05 | 10.25 | 9.86 |
| SF3A2 | 14 | 9.94 | 10.65 | 10.91 | 9.77 | 9.25 | 9.01 | 10.65 | 10.52 | 9.69 | 9.61 | 10.29 | 10.34 | 9.13 | 10.58 | 9.65 |
| SF3A3 | 21 | 10.03 | 10.84 | 10.56 | 10.45 | 8.90 | 8.90 | 10.74 | 9.88 | 9.80 | 9.90 | 10.43 | 10.51 | 8.90 | 10.31 | 9.85 |
| SF3B1 | 54 | 9.99 | 10.23 | 9.14 | 8.61 | 11.00 | 10.92 | 10.62 | 10.38 | 9.72 | 9.38 | 10.11 | 8.88 | 10.96 | 10.50 | 9.55 |
| SF3B14 | 9 | 9.81 | 10.45 | 10.01 | 8.77 | 10.50 | 10.11 | 10.73 | 10.16 | 9.81 | 9.64 | 10.13 | 9.39 | 10.31 | 10.45 | 9.73 |
| SF3B2 | 65 | 8.97 | 10.57 | 11.27 | 9.68 | 10.42 | 10.40 | 10.93 | 9.62 | 9.29 | 8.85 | 9.77 | 10.47 | 10.41 | 10.27 | 9.07 |
| SF3B3 | 61 | 9.94 | 10.77 | 8.80 | 8.72 | 10.71 | 10.97 | 10.15 | 10.93 | 9.62 | 9.39 | 10.36 | 8.76 | 10.84 | 10.54 | 9.50 |
| SF3B4 | 3 | 8.35 | 9.71 | 11.82 | 12.23 | 9.80 | 9.86 | 10.45 | 10.09 | 9.20 | 8.51 | 9.03 | 12.02 | 9.83 | 10.27 | 8.86 |
| SF3B5 | 4 | 10.94 | 10.52 | 10.61 | 7.69 | 11.93 | 12.38 | 10.95 | 4.51 | 10.25 | 10.21 | 10.73 | 9.15 | 12.16 | 7.73 | 10.23 |
| SFMBT1 | 2 | 10.45 | 10.34 | 9.95 | 10.02 | 8.86 | 9.73 | 10.80 | 9.54 | 10.17 | 10.13 | 10.40 | 9.98 | 9.30 | 10.17 | 10.15 |
| SFN | 43 | 10.38 | 11.91 | 7.69 | 7.14 | 10.08 | 10.06 | 8.44 | 9.29 | 12.66 | 12.35 | 11.14 | 7.42 | 10.07 | 8.86 | 12.51 |
| SFPQ | 93 | 10.14 | 11.25 | 9.44 | 8.73 | 9.80 | 9.96 | 11.44 | 10.69 | 9.60 | 8.96 | 10.69 | 9.09 | 9.88 | 11.06 | 9.28 |
| SFSWAP | 8 | 9.34 | 10.70 | 10.44 | 10.59 | 9.07 | 9.82 | 10.24 | 10.01 | 9.84 | 9.95 | 10.02 | 10.52 | 9.45 | 10.13 | 9.89 |
| SFT2D2 | 1 | 12.40 | 7.94 | 6.15 | 9.07 | 16.77 | 14.06 | 9.16 | 8.64 | 6.99 | 8.81 | 10.17 | 7.61 | 15.41 | 8.90 | 7.90 |
| SFT2D3 | 1 | 10.02 | 7.70 | 7.71 | 8.58 | 10.28 | 10.46 | 8.23 | 9.18 | 12.56 | 15.27 | 8.86 | 8.15 | 10.37 | 8.71 | 13.92 |
| SFXN1 | 12 | 10.06 | 9.51 | 9.32 | 9.69 | 10.43 | 10.54 | 10.24 | 9.58 | 10.04 | 10.59 | 9.78 | 9.50 | 10.49 | 9.91 | 10.32 |
| SFXN3 | 13 | 10.18 | 7.46 | 9.72 | 10.96 | 11.33 | 9.77 | 9.70 | 8.74 | 10.92 | 11.23 | 8.82 | 10.34 | 10.55 | 9.22 | 11.07 |
| SGK1 | 2 | 8.13 | 8.75 | 8.34 | 8.47 | 12.94 | 12.88 | 11.65 | 10.27 | 9.66 | 8.93 | 8.44 | 8.40 | 12.91 | 10.96 | 9.29 |
| SGK223 | 1 | 7.67 | 10.66 | 11.38 | 14.12 | 8.56 | 8.57 | 10.28 | 9.66 | 8.60 | 10.50 | 9.16 | 12.75 | 8.57 | 9.97 | 9.55 |
| SGPL1 | 3 | 12.28 | 7.13 | 9.44 | 14.12 | 7.79 | 7.98 | 7.79 | 8.02 | 11.86 | 13.60 | 9.70 | 11.78 | 7.88 | 7.91 | 12.73 |
| SGSH | 2 | 11.97 | 12.90 | 12.06 | 9.93 | 9.63 | 9.77 | 10.61 | 10.39 | 6.51 | 6.23 | 12.44 | 10.99 | 9.70 | 10.50 | 6.37 |
| SGTA | 14 | 8.44 | 9.44 | 12.28 | 10.09 | 10.22 | 9.87 | 9.87 | 10.37 | 9.92 | 9.51 | 8.94 | 11.18 | 10.04 | 10.12 | 9.72 |
| SGTB | 2 | 9.83 | 11.15 | 9.40 | 10.48 | 10.59 | 9.77 | 9.22 | 7.27 | 11.06 | 11.22 | 10.49 | 9.94 | 10.18 | 8.25 | 11.14 |
| SH2B1 | 2 | 7.31 | 8.88 | 12.82 | 12.60 | 9.65 | 7.79 | 11.51 | 9.79 | 10.85 | 8.79 | 8.10 | 12.71 | 8.72 | 10.65 | 9.82 |
| SH2D4A | 18 | 9.67 | 10.97 | 12.15 | 10.77 | 9.72 | 9.91 | 9.52 | 8.75 | 9.25 | 9.30 | 10.32 | 11.46 | 9.82 | 9.13 | 9.27 |
| SH3BGRL | 16 | 9.02 | 10.31 | 7.56 | 7.23 | 17.15 | 17.33 | 10.13 | 8.74 | 6.53 | 5.98 | 9.67 | 7.40 | 17.24 | 9.44 | 6.25 |
| SH3BGRL3 | 14 | 8.49 | 9.42 | 12.66 | 12.20 | 8.70 | 9.28 | 10.27 | 9.25 | 9.99 | 9.74 | 8.96 | 12.43 | 8.99 | 9.76 | 9.86 |
| SH3BP4 | 8 | 9.51 | 9.88 | 17.49 | 18.22 | 5.60 | 5.65 | 9.33 | 8.05 | 8.00 | 8.28 | 9.69 | 17.85 | 5.62 | 8.69 | 8.14 |
| SH3BP5 | 3 | 11.45 | 11.86 | 10.25 | 9.04 | 10.99 | 11.63 | 6.97 | 6.65 | 10.77 | 10.39 | 11.65 | 9.64 | 11.31 | 6.81 | 10.58 |
| SH3BP5L | 2 | 8.15 | 8.36 | 13.67 | 12.60 | 10.16 | 9.89 | 9.76 | 9.96 | 9.37 | 8.09 | 8.25 | 13.13 | 10.02 | 9.86 | 8.73 |

| | | | | | | | | | | | | | | | | |
|---|---|---|---|---|---|---|---|---|---|---|---|---|---|---|---|---|
| SH3D19 | 6 | 9.76 | 11.14 | 10.81 | 10.85 | 10.07 | 10.94 | 9.68 | 9.37 | 8.86 | 8.51 | 10.45 | 10.83 | 10.51 | 9.52 | 8.68 |
| SH3GL1 | 24 | 9.92 | 10.74 | 11.41 | 10.53 | 9.75 | 9.89 | 8.30 | 8.94 | 10.60 | 9.93 | 10.33 | 10.97 | 9.82 | 8.62 | 10.26 |
| SH3GL2 | 1 | 13.55 | 13.87 | 6.90 | 6.97 | 14.05 | 12.98 | 6.12 | 5.34 | 10.68 | 9.52 | 13.71 | 6.94 | 13.52 | 5.73 | 10.10 |
| SH3GLB1 | 7 | 10.06 | 11.12 | 10.13 | 8.91 | 9.41 | 9.31 | 9.69 | 8.60 | 11.55 | 11.23 | 10.59 | 9.52 | 9.36 | 9.14 | 11.39 |
| SH3GLB2 | 10 | 9.85 | 10.76 | 9.58 | 9.52 | 12.48 | 12.85 | 8.19 | 8.04 | 9.79 | 8.94 | 10.31 | 9.55 | 12.67 | 8.11 | 9.36 |
| SH3KBP1 | 12 | 4.87 | 5.35 | 13.41 | 11.78 | 12.58 | 12.77 | 10.51 | 10.07 | 9.78 | 8.86 | 5.11 | 12.60 | 12.68 | 10.29 | 9.32 |
| SH3PXD2A | 4 | 10.16 | 11.24 | 10.96 | 11.32 | 8.93 | 9.06 | 8.89 | 8.87 | 10.61 | 9.96 | 10.70 | 11.14 | 8.99 | 8.88 | 10.28 |
| SH3PXD2B | 25 | 11.29 | 11.93 | 9.68 | 9.38 | 10.72 | 10.91 | 9.58 | 8.94 | 9.18 | 8.41 | 11.61 | 9.53 | 10.81 | 9.26 | 8.79 |
| SH3RF1 | 3 | 11.39 | 11.65 | 9.78 | 9.33 | 12.70 | 12.18 | 7.04 | 5.98 | 10.48 | 9.46 | 11.52 | 9.56 | 12.44 | 6.51 | 9.97 |
| SHB | 1 | 9.32 | 9.81 | 12.54 | 13.46 | 6.05 | 6.18 | 9.31 | 7.52 | 12.34 | 13.47 | 9.56 | 13.00 | 6.11 | 8.41 | 12.91 |
| SHC1 | 11 | 9.12 | 10.15 | 11.06 | 11.19 | 10.67 | 11.16 | 7.99 | 7.32 | 10.83 | 10.51 | 9.63 | 11.13 | 10.91 | 7.65 | 10.67 |
| SHCBP1 | 6 | 12.75 | 13.84 | 11.08 | 8.89 | 13.37 | 13.08 | 9.33 | 8.07 | 4.83 | 4.77 | 13.29 | 9.98 | 13.22 | 8.70 | 4.80 |
| SHMT1 | 14 | 11.84 | 11.95 | 10.69 | 10.19 | 8.42 | 8.64 | 8.26 | 7.62 | 11.60 | 10.78 | 11.90 | 10.44 | 8.53 | 7.94 | 11.19 |
| SHMT2 | 45 | 10.47 | 8.66 | 8.27 | 8.57 | 10.02 | 9.01 | 9.79 | 10.24 | 12.33 | 12.65 | 9.56 | 8.42 | 9.51 | 10.02 | 12.49 |
| SHOC2 | 6 | 12.06 | 11.46 | 9.05 | 9.08 | 10.75 | 11.18 | 8.14 | 7.46 | 11.02 | 9.80 | 11.76 | 9.07 | 10.97 | 7.80 | 10.41 |
| SHPK | 2 | 10.76 | 12.87 | 14.41 | 15.55 | 7.71 | 7.92 | 4.90 | 4.30 | 10.76 | 10.84 | 11.81 | 14.98 | 7.81 | 4.60 | 10.80 |
| SHQ1 | 2 | 9.93 | 10.56 | 9.12 | 8.89 | 9.98 | 10.00 | 10.94 | 11.29 | 8.96 | 10.33 | 10.24 | 9.01 | 9.99 | 11.12 | 9.64 |
| SHROOM1 | 1 | 8.80 | 9.68 | 9.48 | 9.55 | 12.35 | 8.94 | 11.08 | 8.87 | 9.77 | 11.47 | 9.24 | 9.52 | 10.64 | 9.97 | 10.62 |
| SIAE | 1 | 7.77 | 8.79 | 6.14 | 8.96 | 12.10 | 13.90 | 16.05 | 17.10 | 4.15 | 5.04 | 8.28 | 7.55 | 13.00 | 16.57 | 4.59 |
| SIGMAR1 | 1 | 13.17 | 5.20 | 8.80 | 6.75 | 9.91 | 5.89 | 8.56 | 5.54 | 14.95 | 21.24 | 9.18 | 7.78 | 7.90 | 7.05 | 18.09 |
| SIK3 | 2 | 10.70 | 9.90 | 10.33 | 11.24 | 10.15 | 11.38 | 7.65 | 8.46 | 9.35 | 10.85 | 10.30 | 10.78 | 10.77 | 8.05 | 10.10 |
| SIKE1 | 1 | 11.11 | 11.05 | 11.00 | 9.34 | 10.03 | 9.78 | 11.49 | 6.04 | 10.93 | 9.23 | 11.08 | 10.17 | 9.91 | 8.77 | 10.08 |
| SIL1 | 4 | 9.87 | 8.33 | 7.67 | 7.36 | 12.45 | 12.22 | 12.67 | 12.49 | 8.78 | 8.16 | 9.10 | 7.52 | 12.33 | 12.58 | 8.47 |
| SIN3A | 17 | 8.55 | 9.92 | 9.79 | 9.97 | 10.17 | 10.75 | 10.49 | 10.50 | 10.02 | 9.86 | 9.23 | 9.88 | 10.46 | 10.49 | 9.94 |
| SIN3B | 2 | 9.13 | 9.98 | 11.19 | 8.59 | 10.54 | 9.37 | 11.93 | 10.21 | 10.29 | 8.78 | 9.56 | 9.89 | 9.96 | 11.07 | 9.53 |
| SIPA1 | 4 | 8.69 | 9.70 | 14.12 | 14.10 | 8.86 | 9.06 | 7.46 | 7.36 | 10.49 | 10.15 | 9.19 | 14.11 | 8.96 | 7.41 | 10.32 |
| SIPA1L1 | 10 | 8.83 | 9.74 | 8.73 | 8.69 | 11.56 | 11.67 | 12.92 | 12.13 | 8.02 | 7.70 | 9.29 | 8.71 | 11.61 | 12.53 | 7.86 |
| SIRPA | 2 | 9.16 | 10.10 | 13.94 | 17.13 | 8.33 | 9.54 | 7.15 | 6.58 | 9.29 | 8.78 | 9.63 | 15.53 | 8.93 | 6.87 | 9.04 |
| SIRT1 | 4 | 9.78 | 10.15 | 8.88 | 8.02 | 12.07 | 11.83 | 12.32 | 10.81 | 8.18 | 7.97 | 9.97 | 8.45 | 11.95 | 11.57 | 8.07 |
| SIRT2 | 3 | 9.28 | 10.53 | 9.46 | 9.81 | 8.37 | 9.09 | 10.89 | 10.06 | 10.95 | 11.54 | 9.91 | 9.64 | 8.73 | 10.48 | 11.25 |
| SIRT5 | 2 | 9.48 | 9.78 | 10.90 | 11.33 | 8.48 | 8.71 | 9.51 | 9.02 | 12.03 | 10.76 | 9.63 | 11.11 | 8.59 | 9.27 | 11.40 |
| SIRT6 | 3 | 9.86 | 10.73 | 10.45 | 9.59 | 8.88 | 10.02 | 9.64 | 8.78 | 11.43 | 10.61 | 10.29 | 10.02 | 9.45 | 9.21 | 11.02 |
| SKA2 | 2 | 10.13 | 11.42 | 11.86 | 12.86 | 7.96 | 9.30 | 8.90 | 7.51 | 9.87 | 10.19 | 10.78 | 12.36 | 8.63 | 8.20 | 10.03 |
| SKA3 | 2 | 10.09 | 12.32 | 13.64 | 11.25 | 8.34 | 8.43 | 8.81 | 7.77 | 9.12 | 10.23 | 11.20 | 12.45 | 8.39 | 8.29 | 9.67 |
| SKIV2L | 29 | 10.06 | 10.35 | 9.60 | 9.10 | 10.92 | 11.08 | 9.14 | 8.70 | 10.53 | 10.51 | 10.21 | 9.35 | 11.00 | 8.92 | 10.52 |
| SKIV2L2 | 54 | 10.40 | 10.55 | 9.51 | 8.97 | 9.85 | 9.88 | 10.95 | 10.45 | 9.82 | 9.62 | 10.48 | 9.24 | 9.86 | 10.70 | 9.72 |
| SKP1 | 20 | 8.39 | 8.10 | 8.28 | 5.96 | 10.21 | 9.57 | 9.23 | 25.58 | 7.62 | 7.07 | 8.25 | 7.12 | 9.89 | 17.40 | 7.34 |
| SKP2 | 4 | 10.33 | 11.22 | 6.90 | 7.05 | 10.05 | 9.96 | 12.06 | 11.79 | 10.51 | 10.15 | 10.77 | 6.97 | 10.00 | 11.92 | 10.33 |
| SLAIN2 | 11 | 9.46 | 10.47 | 10.83 | 10.93 | 9.65 | 10.72 | 9.66 | 8.97 | 9.97 | 9.34 | 9.97 | 10.88 | 10.18 | 9.32 | 9.66 |
| SLBP | 2 | 9.78 | 10.72 | 8.64 | 9.15 | 10.96 | 10.63 | 11.14 | 8.72 | 10.15 | 10.10 | 10.25 | 8.89 | 10.80 | 9.93 | 10.13 |
| SLC12A2 | 8 | 13.35 | 10.01 | 7.89 | 9.71 | 8.24 | 8.81 | 11.15 | 9.56 | 10.24 | 11.04 | 11.68 | 8.80 | 8.52 | 10.36 | 10.64 |
| SLC12A4 | 7 | 12.43 | 9.52 | 6.19 | 7.00 | 13.50 | 13.18 | 7.96 | 8.00 | 10.91 | 11.31 | 10.97 | 6.59 | 13.34 | 7.98 | 11.11 |

| | | | | | | | | | | | | | | | | |
|---|---|---|---|---|---|---|---|---|---|---|---|---|---|---|---|---|
| SLC12A7 | 13 | 12.96 | 9.98 | 1.95 | 1.24 | 5.02 | 4.55 | 28.64 | 25.24 | 5.09 | 5.34 | 11.47 | 1.59 | 4.78 | 26.94 | 5.21 |
| SLC16A1 | 9 | 12.45 | 9.39 | 7.49 | 9.25 | 7.92 | 8.13 | 11.56 | 10.25 | 11.77 | 11.79 | 10.92 | 8.37 | 8.02 | 10.91 | 11.78 |
| SLC16A3 | 8 | 12.71 | 9.58 | 8.35 | 9.55 | 8.51 | 8.10 | 10.44 | 8.86 | 11.77 | 12.13 | 11.14 | 8.95 | 8.30 | 9.65 | 11.95 |
| SLC16A7 | 2 | 11.74 | 8.81 | 8.33 | 9.36 | 9.85 | 9.97 | 12.42 | 10.02 | 9.89 | 9.61 | 10.28 | 8.85 | 9.91 | 11.22 | 9.75 |
| SLC19A1 | 1 | 10.13 | 8.39 | 8.08 | 5.82 | 8.80 | 10.92 | 11.46 | 10.77 | 11.77 | 13.86 | 9.26 | 6.95 | 9.86 | 11.11 | 12.81 |
| SLC1A1 | 1 | 11.12 | 7.82 | 7.48 | 11.34 | 8.01 | 7.24 | 18.54 | 15.84 | 5.95 | 6.66 | 9.47 | 9.41 | 7.63 | 17.19 | 6.30 |
| SLC1A4 | 3 | 12.85 | 10.51 | 6.90 | 5.91 | 11.32 | 11.57 | 7.68 | 6.77 | 12.24 | 14.25 | 11.68 | 6.40 | 11.44 | 7.23 | 13.25 |
| SLC1A5 | 12 | 9.22 | 7.87 | 6.05 | 6.88 | 6.61 | 6.63 | 12.51 | 23.12 | 9.79 | 11.31 | 8.55 | 6.46 | 6.62 | 17.82 | 10.55 |
| SLC20A1 | 2 | 12.08 | 7.99 | 6.89 | 9.35 | 9.74 | 10.31 | 9.21 | 8.93 | 12.69 | 12.83 | 10.03 | 8.12 | 10.02 | 9.07 | 12.76 |
| SLC20A2 | 1 | 9.85 | 7.41 | 11.41 | 10.71 | 9.07 | 8.87 | 12.02 | 10.84 | 10.34 | 9.48 | 8.63 | 11.06 | 8.97 | 11.43 | 9.91 |
| SLC22A23 | 1 | 10.05 | 7.59 | 9.92 | 12.73 | 5.22 | 5.45 | 13.68 | 13.61 | 10.81 | 10.93 | 8.82 | 11.33 | 5.33 | 13.64 | 10.87 |
| SLC22A3 | 1 | 19.61 | 13.02 | 3.53 | 6.03 | 10.30 | 13.64 | 7.21 | 8.71 | 9.12 | 8.84 | 16.31 | 4.78 | 11.97 | 7.96 | 8.98 |
| SLC25A1 | 6 | 14.31 | 7.55 | 7.35 | 9.31 | 8.89 | 7.44 | 9.06 | 8.07 | 12.56 | 15.46 | 10.93 | 8.33 | 8.16 | 8.56 | 14.01 |
| SLC25A10 | 5 | 14.23 | 8.26 | 8.12 | 11.54 | 7.76 | 6.74 | 9.41 | 8.73 | 11.40 | 13.80 | 11.24 | 9.83 | 7.25 | 9.07 | 12.60 |
| SLC25A11 | 11 | 12.95 | 6.47 | 7.16 | 10.05 | 9.55 | 7.85 | 8.10 | 12.08 | 11.49 | 14.31 | 9.71 | 8.60 | 8.70 | 10.09 | 12.90 |
| SLC25A12 | 16 | 13.20 | 7.48 | 8.00 | 12.15 | 8.36 | 7.52 | 9.15 | 9.76 | 11.00 | 13.37 | 10.34 | 10.07 | 7.94 | 9.46 | 12.19 |
| SLC25A13 | 16 | 14.14 | 7.12 | 7.63 | 11.91 | 8.64 | 7.60 | 10.08 | 10.12 | 10.30 | 12.46 | 10.63 | 9.77 | 8.12 | 10.10 | 11.38 |
| SLC25A19 | 2 | 15.71 | 8.39 | 6.87 | 9.52 | 7.90 | 6.54 | 8.56 | 8.62 | 12.42 | 15.47 | 12.05 | 8.20 | 7.22 | 8.59 | 13.95 |
| SLC25A20 | 4 | 13.79 | 8.35 | 5.95 | 9.54 | 10.01 | 8.94 | 7.28 | 7.94 | 12.66 | 15.53 | 11.07 | 7.74 | 9.47 | 7.61 | 14.10 |
| SLC25A22 | 6 | 11.29 | 8.49 | 7.90 | 10.87 | 11.21 | 9.15 | 9.99 | 9.14 | 10.18 | 11.79 | 9.89 | 9.38 | 10.18 | 9.57 | 10.98 |
| SLC25A24 | 21 | 11.82 | 7.15 | 8.12 | 9.03 | 11.27 | 9.37 | 11.56 | 10.58 | 9.96 | 11.15 | 9.48 | 8.58 | 10.32 | 11.07 | 10.55 |
| SLC25A3 | 12 | 13.80 | 7.33 | 6.36 | 9.50 | 10.77 | 9.69 | 9.93 | 10.05 | 10.40 | 12.15 | 10.56 | 7.93 | 10.23 | 9.99 | 11.28 |
| SLC25A32 | 2 | 14.30 | 5.87 | 7.25 | 9.81 | 7.04 | 5.92 | 12.24 | 12.74 | 11.66 | 13.17 | 10.08 | 8.53 | 6.48 | 12.49 | 12.42 |
| SLC25A40 | 1 | 12.81 | 7.68 | 8.17 | 9.70 | 9.28 | 8.68 | 10.01 | 10.21 | 10.81 | 12.66 | 10.24 | 8.93 | 8.98 | 10.11 | 11.73 |
| SLC25A5 | 9 | 11.58 | 7.50 | 6.45 | 8.69 | 12.58 | 10.22 | 12.39 | 12.35 | 8.49 | 9.73 | 9.54 | 7.57 | 11.40 | 12.37 | 9.11 |
| SLC25A6 | 31 | 11.76 | 7.59 | 6.05 | 8.05 | 11.30 | 10.12 | 10.49 | 16.66 | 8.35 | 9.62 | 9.68 | 7.05 | 10.71 | 13.57 | 8.99 |
| SLC26A6 | 1 | 11.45 | 7.78 | 7.73 | 16.74 | 6.76 | 8.84 | 10.98 | 7.66 | 10.21 | 11.85 | 9.61 | 12.24 | 7.80 | 9.32 | 11.03 |
| SLC27A2 | 6 | 13.36 | 7.95 | 5.80 | 7.87 | 15.69 | 14.98 | 8.13 | 7.61 | 8.60 | 10.01 | 10.66 | 6.83 | 15.34 | 7.87 | 9.31 |
| SLC27A4 | 8 | 11.24 | 6.30 | 8.09 | 11.95 | 8.00 | 6.51 | 9.12 | 8.46 | 13.44 | 16.89 | 8.77 | 10.02 | 7.26 | 8.79 | 15.17 |
| SLC29A1 | 1 | 11.62 | 8.99 | 8.77 | 11.92 | 7.93 | 8.97 | 10.43 | 7.08 | 12.47 | 11.82 | 10.31 | 10.34 | 8.45 | 8.75 | 12.15 |
| SLC2A1 | 12 | 9.72 | 8.00 | 12.23 | 14.03 | 4.64 | 4.75 | 7.54 | 6.72 | 15.56 | 16.80 | 8.86 | 13.13 | 4.70 | 7.13 | 16.18 |
| SLC30A1 | 6 | 10.72 | 8.46 | 6.16 | 7.23 | 11.76 | 12.01 | 11.01 | 9.24 | 11.06 | 12.36 | 9.59 | 6.69 | 11.88 | 10.13 | 11.71 |
| SLC30A5 | 1 | 12.32 | 7.41 | 6.16 | 7.52 | 10.54 | 9.78 | 12.01 | 12.91 | 9.59 | 11.76 | 9.86 | 6.84 | 10.16 | 12.46 | 10.68 |
| SLC30A6 | 2 | 15.10 | 7.21 | 6.96 | 8.35 | 10.53 | 8.83 | 11.73 | 9.88 | 9.93 | 11.47 | 11.16 | 7.65 | 9.68 | 10.81 | 10.70 |
| SLC30A7 | 3 | 14.00 | 7.89 | 5.81 | 7.63 | 9.85 | 9.35 | 10.32 | 12.02 | 11.41 | 11.71 | 10.95 | 6.72 | 9.60 | 11.17 | 11.56 |
| SLC35A1 | 1 | 10.69 | 6.86 | 7.47 | 9.72 | 13.52 | 11.50 | 10.76 | 10.02 | 9.77 | 9.69 | 8.78 | 8.59 | 12.51 | 10.39 | 9.73 |
| SLC35B2 | 4 | 12.36 | 7.67 | 8.17 | 9.90 | 11.99 | 11.09 | 9.63 | 9.76 | 9.22 | 10.20 | 10.01 | 9.04 | 11.54 | 9.70 | 9.71 |
| SLC35C1 | 1 | 11.53 | 5.76 | 7.61 | 8.90 | 9.76 | 8.89 | 12.80 | 12.47 | 11.01 | 11.27 | 8.64 | 8.26 | 9.33 | 12.63 | 11.14 |
| SLC35E1 | 1 | 8.18 | 9.04 | 12.57 | 10.72 | 10.51 | 9.47 | 9.78 | 6.45 | 10.94 | 12.36 | 8.61 | 11.64 | 9.99 | 8.12 | 11.65 |
| SLC35F6 | 1 | 10.19 | 7.45 | 6.61 | 9.98 | 12.97 | 10.72 | 11.36 | 12.06 | 8.96 | 9.70 | 8.82 | 8.29 | 11.85 | 11.71 | 9.33 |
| SLC38A1 | 4 | 8.21 | 6.13 | 16.30 | 6.25 | 9.24 | 6.73 | 15.52 | 7.50 | 11.76 | 12.37 | 7.17 | 11.27 | 7.99 | 11.51 | 12.06 |
| SLC38A2 | 1 | 5.67 | 5.16 | 14.58 | 10.56 | 10.92 | 7.03 | 11.90 | 9.51 | 11.53 | 13.14 | 5.41 | 12.57 | 8.98 | 10.70 | 12.34 |

| | | | | | | | | | | | | | | | | |
|---|---|---|---|---|---|---|---|---|---|---|---|---|---|---|---|---|
| SLC38A5 | 4 | 9.38 | 6.67 | 9.53 | 11.98 | 9.14 | 10.07 | 12.88 | 11.68 | 8.72 | 9.95 | 8.03 | 10.76 | 9.61 | 12.28 | 9.33 |
| SLC39A10 | 9 | 12.22 | 10.57 | 8.03 | 8.91 | 10.59 | 10.98 | 9.08 | 7.50 | 11.20 | 10.91 | 11.40 | 8.47 | 10.79 | 8.29 | 11.06 |
| SLC39A14 | 5 | 12.04 | 9.17 | 8.15 | 10.55 | 7.94 | 7.87 | 10.23 | 8.91 | 12.49 | 12.66 | 10.61 | 9.35 | 7.90 | 9.57 | 12.57 |
| SLC39A6 | 2 | 16.36 | 9.50 | 7.84 | 10.43 | 7.99 | 8.68 | 7.53 | 5.62 | 13.56 | 12.50 | 12.93 | 9.13 | 8.34 | 6.57 | 13.03 |
| SLC39A7 | 3 | 11.71 | 6.22 | 8.94 | 14.71 | 8.81 | 7.83 | 7.53 | 7.64 | 11.38 | 15.23 | 8.97 | 11.82 | 8.32 | 7.58 | 13.30 |
| SLC39A8 | 1 | 11.96 | 9.84 | 5.96 | 8.80 | 11.26 | 10.67 | 11.09 | 10.71 | 9.56 | 10.14 | 10.90 | 7.38 | 10.97 | 10.90 | 9.85 |
| SLC3A2 | 74 | 11.88 | 9.93 | 8.88 | 11.26 | 8.53 | 8.59 | 9.47 | 8.41 | 11.42 | 11.62 | 10.91 | 10.07 | 8.56 | 8.94 | 11.52 |
| SLC43A3 | 2 | 12.98 | 9.79 | 7.16 | 8.47 | 7.39 | 8.02 | 8.92 | 8.41 | 13.88 | 14.99 | 11.38 | 7.82 | 7.71 | 8.66 | 14.43 |
| SLC44A1 | 1 | 8.11 | 6.10 | 8.23 | 11.17 | 10.38 | 7.10 | 13.29 | 14.92 | 8.72 | 11.97 | 7.11 | 9.70 | 8.74 | 14.11 | 10.34 |
| SLC4A1AP | 21 | 9.22 | 10.47 | 10.04 | 10.11 | 11.55 | 11.86 | 9.32 | 8.55 | 9.60 | 9.28 | 9.84 | 10.08 | 11.70 | 8.94 | 9.44 |
| SLC4A2 | 7 | 13.53 | 10.34 | 7.92 | 10.23 | 8.31 | 7.77 | 6.97 | 6.11 | 14.09 | 14.73 | 11.94 | 9.07 | 8.04 | 6.54 | 14.41 |
| SLC4A7 | 5 | 13.47 | 9.56 | 6.91 | 9.07 | 6.65 | 6.76 | 12.64 | 10.45 | 12.05 | 12.44 | 11.52 | 7.99 | 6.71 | 11.54 | 12.25 |
| SLC52A2 | 1 | 10.64 | 6.81 | 7.82 | 9.23 | 8.88 | 6.89 | 14.08 | 13.68 | 9.17 | 12.80 | 8.73 | 8.52 | 7.89 | 13.88 | 10.98 |
| SLC5A6 | 1 | 9.00 | 6.28 | 9.15 | 9.59 | 9.36 | 8.23 | 14.01 | 13.91 | 9.84 | 10.63 | 7.64 | 9.37 | 8.79 | 13.96 | 10.24 |
| SLC7A1 | 3 | 13.25 | 9.51 | 9.53 | 11.41 | 9.98 | 9.73 | 8.50 | 8.24 | 9.51 | 10.34 | 11.38 | 10.47 | 9.85 | 8.37 | 9.93 |
| SLC7A11 | 3 | 11.72 | 7.63 | 10.92 | 13.03 | 3.87 | 3.94 | 13.30 | 10.09 | 12.47 | 13.02 | 9.67 | 11.98 | 3.91 | 11.70 | 12.75 |
| SLC7A5 | 7 | 12.94 | 9.94 | 8.19 | 10.21 | 8.00 | 8.62 | 9.12 | 9.45 | 11.25 | 12.29 | 11.44 | 9.20 | 8.31 | 9.28 | 11.77 |
| SLC9A3R1 | 30 | 10.73 | 12.89 | 8.97 | 7.89 | 12.60 | 12.35 | 7.63 | 6.03 | 10.73 | 10.19 | 11.81 | 8.43 | 12.47 | 6.83 | 10.46 |
| SLC9A3R2 | 10 | 8.86 | 10.24 | 8.94 | 7.70 | 11.34 | 12.02 | 11.45 | 9.94 | 9.74 | 9.77 | 9.55 | 8.32 | 11.68 | 10.69 | 9.76 |
| SLC9A6 | 2 | 11.58 | 7.71 | 7.46 | 8.71 | 13.97 | 12.19 | 10.00 | 9.27 | 9.36 | 9.74 | 9.65 | 8.08 | 13.08 | 9.64 | 9.55 |
| SLFN5 | 10 | 8.35 | 8.37 | 17.32 | 17.09 | 10.47 | 10.93 | 9.83 | 9.79 | 3.98 | 3.89 | 8.36 | 17.21 | 10.70 | 9.81 | 3.93 |
| SLIRP | 11 | 9.21 | 9.66 | 11.06 | 12.95 | 9.20 | 9.38 | 9.29 | 9.43 | 9.73 | 10.10 | 9.44 | 12.01 | 9.29 | 9.36 | 9.91 |
| SLK | 38 | 9.03 | 9.94 | 10.05 | 9.49 | 10.99 | 11.05 | 11.41 | 10.34 | 9.21 | 8.48 | 9.49 | 9.77 | 11.02 | 10.87 | 8.85 |
| SLMAP | 11 | 9.62 | 8.28 | 14.63 | 16.26 | 8.46 | 8.43 | 8.59 | 7.86 | 8.71 | 9.16 | 8.95 | 15.45 | 8.44 | 8.22 | 8.93 |
| SLTM | 27 | 10.04 | 10.89 | 9.32 | 9.13 | 10.05 | 10.46 | 9.87 | 9.59 | 10.47 | 10.18 | 10.46 | 9.23 | 10.25 | 9.73 | 10.33 |
| SLU7 | 4 | 9.16 | 10.98 | 10.37 | 9.47 | 9.95 | 9.93 | 12.42 | 10.72 | 8.40 | 8.60 | 10.07 | 9.92 | 9.94 | 11.57 | 8.50 |
| SMAD2 | 9 | 9.18 | 11.14 | 8.85 | 9.23 | 12.31 | 14.12 | 7.93 | 7.70 | 9.85 | 9.69 | 10.16 | 9.04 | 13.22 | 7.81 | 9.77 |
| SMAD3 | 1 | 8.90 | 10.51 | 8.72 | 14.14 | 6.81 | 10.53 | 9.22 | 8.63 | 12.77 | 9.77 | 9.70 | 11.43 | 8.67 | 8.92 | 11.27 |
| SMAD4 | 6 | 10.15 | 10.86 | 10.80 | 9.21 | 11.15 | 10.44 | 9.12 | 8.00 | 10.39 | 9.88 | 10.50 | 10.01 | 10.80 | 8.56 | 10.14 |
| SMAD5 | 5 | 8.38 | 10.68 | 10.17 | 9.69 | 8.82 | 8.60 | 13.66 | 12.79 | 8.81 | 8.40 | 9.53 | 9.93 | 8.71 | 13.23 | 8.60 |
| SMAP | 12 | 9.14 | 10.33 | 11.55 | 10.34 | 11.04 | 10.89 | 10.66 | 7.87 | 9.37 | 8.81 | 9.74 | 10.94 | 10.97 | 9.27 | 9.09 |
| SMAP1 | 4 | 10.34 | 11.63 | 11.58 | 10.99 | 11.31 | 12.33 | 7.86 | 7.49 | 8.64 | 7.84 | 10.98 | 11.29 | 11.82 | 7.68 | 8.24 |
| SMAP2 | 3 | 10.70 | 12.58 | 9.51 | 9.48 | 7.27 | 8.13 | 9.94 | 8.71 | 12.51 | 11.19 | 11.64 | 9.49 | 7.70 | 9.32 | 11.85 |
| SMARCA1 | 34 | 9.05 | 9.68 | 10.13 | 9.66 | 9.68 | 9.91 | 10.51 | 14.03 | 8.75 | 8.60 | 9.36 | 9.89 | 9.79 | 12.27 | 8.68 |
| SMARCA2 | 18 | 10.25 | 11.42 | 7.31 | 7.21 | 12.72 | 13.36 | 7.14 | 6.95 | 11.84 | 11.79 | 10.84 | 7.26 | 13.04 | 7.05 | 11.82 |
| SMARCA4 | 46 | 8.92 | 9.49 | 10.67 | 10.00 | 9.89 | 9.74 | 12.29 | 11.42 | 8.90 | 8.67 | 9.21 | 10.33 | 9.82 | 11.86 | 8.78 |
| SMARCA5 | 37 | 9.01 | 9.47 | 10.41 | 10.69 | 8.80 | 9.12 | 10.96 | 12.06 | 9.73 | 9.75 | 9.24 | 10.55 | 8.96 | 11.51 | 9.74 |
| SMARCAD1 | 8 | 8.86 | 9.81 | 9.62 | 9.09 | 10.16 | 10.44 | 12.21 | 11.25 | 9.14 | 9.42 | 9.33 | 9.36 | 10.30 | 11.73 | 9.28 |
| SMARCAL1 | 7 | 8.95 | 9.19 | 9.68 | 9.98 | 10.64 | 11.10 | 10.46 | 9.71 | 10.61 | 9.68 | 9.07 | 9.83 | 10.87 | 10.08 | 10.15 |
| SMARCB1 | 5 | 9.93 | 9.92 | 9.95 | 9.38 | 9.70 | 9.67 | 12.16 | 11.15 | 9.61 | 8.52 | 9.93 | 9.67 | 9.69 | 11.65 | 9.07 |
| SMARCC1 | 31 | 8.91 | 10.01 | 9.26 | 8.20 | 9.80 | 10.07 | 13.08 | 12.52 | 9.27 | 8.87 | 9.46 | 8.73 | 9.94 | 12.80 | 9.07 |
| SMARCC2 | 38 | 10.00 | 9.94 | 9.71 | 9.71 | 9.96 | 10.57 | 11.11 | 11.77 | 8.89 | 8.34 | 9.97 | 9.71 | 10.26 | 11.44 | 8.62 |

| | | | | | | | | | | | | | | | | |
|---|---|---|---|---|---|---|---|---|---|---|---|---|---|---|---|---|
| SMARCD1 | 13 | 8.81 | 10.36 | 10.17 | 10.38 | 8.96 | 9.29 | 11.90 | 11.08 | 9.41 | 9.65 | 9.59 | 10.27 | 9.12 | 11.49 | 9.53 |
| SMARCD2 | 10 | 9.66 | 9.70 | 9.95 | 10.23 | 10.39 | 10.50 | 11.07 | 10.16 | 9.27 | 9.10 | 9.68 | 10.09 | 10.44 | 10.61 | 9.18 |
| SMARCE1 | 13 | 9.02 | 11.01 | 9.90 | 9.45 | 9.67 | 10.04 | 12.32 | 11.73 | 8.59 | 8.26 | 10.01 | 9.68 | 9.86 | 12.03 | 8.42 |
| SMC1A | 78 | 9.27 | 9.88 | 10.08 | 9.62 | 10.40 | 10.66 | 11.24 | 10.47 | 9.41 | 8.97 | 9.57 | 9.85 | 10.53 | 10.86 | 9.19 |
| SMC2 | 75 | 9.26 | 10.12 | 11.78 | 10.79 | 9.05 | 9.22 | 10.19 | 9.22 | 10.40 | 9.97 | 9.69 | 11.29 | 9.14 | 9.70 | 10.18 |
| SMC3 | 65 | 9.33 | 9.73 | 10.14 | 9.97 | 10.24 | 10.59 | 11.47 | 10.37 | 9.25 | 8.91 | 9.53 | 10.06 | 10.41 | 10.92 | 9.08 |
| SMC4 | 66 | 9.52 | 10.22 | 11.24 | 10.60 | 8.78 | 8.85 | 10.13 | 10.65 | 10.17 | 9.85 | 9.87 | 10.92 | 8.81 | 10.39 | 10.01 |
| SMC5 | 11 | 9.13 | 9.34 | 10.19 | 9.36 | 10.12 | 10.10 | 10.28 | 10.43 | 10.76 | 10.27 | 9.23 | 9.78 | 10.11 | 10.36 | 10.52 |
| SMC6 | 5 | 8.71 | 10.13 | 9.10 | 10.29 | 9.12 | 9.04 | 10.85 | 10.65 | 11.06 | 11.05 | 9.42 | 9.70 | 9.08 | 10.75 | 11.06 |
| SMCHD1 | 47 | 10.18 | 10.87 | 10.17 | 9.50 | 11.34 | 11.36 | 8.59 | 7.86 | 10.19 | 9.93 | 10.52 | 9.84 | 11.35 | 8.23 | 10.06 |
| SMEK1 | 7 | 9.88 | 10.85 | 9.13 | 9.59 | 9.75 | 10.49 | 10.52 | 10.16 | 9.95 | 9.67 | 10.37 | 9.36 | 10.12 | 10.34 | 9.81 |
| SMEK2 | 9 | 9.83 | 10.32 | 9.44 | 8.52 | 9.84 | 10.10 | 11.38 | 10.50 | 9.93 | 10.15 | 10.07 | 8.98 | 9.97 | 10.94 | 10.04 |
| SMEK3P | 1 | 12.75 | 12.95 | 7.03 | 7.10 | 9.32 | 11.62 | 9.41 | 9.94 | 10.61 | 9.25 | 12.85 | 7.07 | 10.47 | 9.68 | 9.93 |
| SMG1 | 7 | 9.78 | 9.82 | 9.67 | 10.00 | 9.07 | 7.24 | 11.18 | 11.05 | 11.29 | 10.91 | 9.80 | 9.84 | 8.15 | 11.12 | 11.10 |
| SMG5 | 4 | 10.22 | 10.36 | 10.33 | 10.48 | 10.15 | 10.59 | 8.89 | 8.61 | 10.54 | 9.83 | 10.29 | 10.41 | 10.37 | 8.75 | 10.18 |
| SMG6 | 2 | 11.18 | 11.88 | 10.83 | 12.86 | 8.91 | 8.80 | 9.58 | 10.38 | 7.45 | 8.14 | 11.53 | 11.84 | 8.85 | 9.98 | 7.79 |
| SMG7 | 1 | 9.24 | 10.94 | 9.57 | 10.02 | 9.90 | 9.77 | 9.59 | 10.40 | 10.11 | 10.47 | 10.09 | 9.79 | 9.84 | 9.99 | 10.29 |
| SMG8 | 7 | 9.64 | 10.98 | 11.79 | 10.90 | 8.89 | 8.71 | 9.51 | 8.29 | 10.68 | 10.60 | 10.31 | 11.35 | 8.80 | 8.90 | 10.64 |
| SMG9 | 5 | 9.34 | 11.16 | 10.34 | 11.09 | 9.27 | 8.83 | 9.78 | 9.08 | 10.78 | 10.33 | 10.25 | 10.71 | 9.05 | 9.43 | 10.56 |
| SMIM12 | 4 | 8.56 | 12.49 | 10.48 | 10.88 | 8.63 | 9.36 | 11.29 | 8.19 | 9.70 | 10.43 | 10.52 | 10.68 | 8.99 | 9.74 | 10.07 |
| SMIM20 | 1 | 12.15 | 8.58 | 8.30 | 10.56 | 10.32 | 11.01 | 9.15 | 8.79 | 10.08 | 11.05 | 10.36 | 9.43 | 10.67 | 8.97 | 10.57 |
| SMIM4 | 1 | 13.44 | 7.96 | 10.54 | 10.68 | 8.56 | 8.07 | 12.46 | 7.80 | 9.70 | 10.78 | 10.70 | 10.61 | 8.32 | 10.13 | 10.24 |
| SMIM7 | 1 | 11.39 | 10.04 | 6.74 | 4.99 | 9.35 | 8.37 | 10.21 | 10.28 | 12.38 | 16.24 | 10.72 | 5.86 | 8.86 | 10.25 | 14.31 |
| SMIM8 | 1 | 12.86 | 8.44 | 9.61 | 11.91 | 10.06 | 9.24 | 8.93 | 7.95 | 10.17 | 10.82 | 10.65 | 10.76 | 9.65 | 8.44 | 10.50 |
| SMN1 | 3 | 10.30 | 11.13 | 9.21 | 9.39 | 9.59 | 9.94 | 12.97 | 11.51 | 8.14 | 7.81 | 10.72 | 9.30 | 9.77 | 12.24 | 7.97 |
| SMNDC1 | 10 | 9.57 | 10.94 | 10.27 | 9.89 | 9.58 | 9.48 | 10.79 | 9.48 | 10.26 | 9.73 | 10.26 | 10.08 | 9.53 | 10.14 | 9.99 |
| SMOC1 | 2 | 8.72 | 7.11 | 8.66 | 9.13 | 11.78 | 10.88 | 16.31 | 13.36 | 7.79 | 6.26 | 7.91 | 8.90 | 11.33 | 14.83 | 7.02 |
| SMPD1 | 1 | 9.12 | 9.32 | 6.21 | 6.39 | 9.41 | 11.66 | 6.41 | 26.22 | 7.15 | 8.11 | 9.22 | 6.30 | 10.53 | 16.32 | 7.63 |
| SMPD4 | 2 | 13.61 | 7.27 | 7.87 | 13.34 | 7.54 | 7.18 | 9.02 | 8.06 | 10.47 | 15.64 | 10.44 | 10.60 | 7.36 | 8.54 | 13.06 |
| SMPDL3A | 1 | 10.76 | 10.00 | 8.41 | 8.96 | 18.55 | 18.69 | 6.44 | 5.44 | 6.49 | 6.27 | 10.38 | 8.68 | 18.62 | 5.94 | 6.38 |
| SMS | 18 | 7.79 | 8.33 | 8.34 | 7.77 | 10.77 | 10.55 | 11.05 | 20.16 | 7.85 | 7.39 | 8.06 | 8.06 | 10.66 | 15.60 | 7.62 |
| SMTN | 3 | 10.00 | 10.75 | 12.81 | 11.90 | 5.96 | 5.59 | 8.59 | 8.34 | 13.27 | 12.79 | 10.38 | 12.35 | 5.78 | 8.47 | 13.03 |
| SMU1 | 21 | 10.23 | 10.96 | 10.87 | 10.05 | 8.59 | 8.68 | 8.39 | 7.95 | 12.06 | 12.23 | 10.60 | 10.46 | 8.63 | 8.17 | 12.14 |
| SMUG1 | 1 | 7.65 | 9.87 | 10.76 | 13.12 | 8.77 | 11.00 | 8.46 | 8.61 | 9.00 | 12.75 | 8.76 | 11.94 | 9.88 | 8.53 | 10.88 |
| SMYD2 | 3 | 8.22 | 9.00 | 9.36 | 9.06 | 11.99 | 12.66 | 10.23 | 10.50 | 9.43 | 9.55 | 8.61 | 9.21 | 12.32 | 10.36 | 9.49 |
| SMYD3 | 8 | 10.41 | 11.40 | 8.48 | 7.42 | 14.19 | 13.41 | 11.49 | 11.63 | 6.07 | 5.50 | 10.91 | 7.95 | 13.80 | 11.56 | 5.79 |
| SMYD4 | 2 | 11.01 | 9.61 | 11.52 | 12.70 | 8.24 | 9.30 | 8.00 | 7.69 | 11.20 | 10.73 | 10.31 | 12.11 | 8.77 | 7.85 | 10.97 |
| SNAP23 | 11 | 9.96 | 8.54 | 12.50 | 13.28 | 9.15 | 9.11 | 7.36 | 8.24 | 11.49 | 10.37 | 9.25 | 12.89 | 9.13 | 7.80 | 10.93 |
| SNAP29 | 6 | 10.21 | 11.47 | 10.88 | 9.75 | 9.64 | 9.36 | 9.43 | 9.47 | 10.17 | 9.62 | 10.84 | 10.31 | 9.50 | 9.45 | 9.90 |
| SNAPC1 | 1 | 9.61 | 11.62 | 13.03 | 7.63 | 9.51 | 8.69 | 7.01 | 5.92 | 13.43 | 13.55 | 10.62 | 10.33 | 9.10 | 6.46 | 13.49 |
| SNAPC2 | 1 | 9.13 | 10.83 | 12.78 | 11.50 | 9.06 | 8.20 | 9.22 | 7.73 | 11.52 | 10.02 | 9.98 | 12.14 | 8.63 | 8.48 | 10.77 |
| SNAPC4 | 7 | 9.17 | 10.06 | 10.10 | 10.93 | 8.25 | 9.00 | 10.08 | 11.15 | 10.41 | 10.84 | 9.62 | 10.52 | 8.63 | 10.61 | 10.62 |

| | | | | | | | | | | | | | | | | |
|---|---|---|---|---|---|---|---|---|---|---|---|---|---|---|---|---|
| SNAPIN | 3 | 9.67 | 11.20 | 10.89 | 9.85 | 11.70 | 12.68 | 8.53 | 8.54 | 8.54 | 8.41 | 10.43 | 10.37 | 12.19 | 8.54 | 8.47 |
| SNCA | 5 | 14.81 | 15.24 | 10.46 | 9.83 | 7.38 | 7.74 | 11.25 | 9.25 | 7.37 | 6.66 | 15.03 | 10.15 | 7.56 | 10.25 | 7.01 |
| SNCG | 15 | 10.36 | 10.91 | 7.29 | 6.89 | 13.99 | 14.28 | 5.22 | 4.51 | 13.63 | 12.92 | 10.63 | 7.09 | 14.13 | 4.86 | 13.28 |
| SND1 | 86 | 10.09 | 10.33 | 10.53 | 9.48 | 10.03 | 9.96 | 9.80 | 11.29 | 9.50 | 8.98 | 10.21 | 10.00 | 10.00 | 10.54 | 9.24 |
| SNF8 | 8 | 10.01 | 11.82 | 10.66 | 9.16 | 12.83 | 13.09 | 7.33 | 6.72 | 9.08 | 9.30 | 10.92 | 9.91 | 12.96 | 7.03 | 9.19 |
| SNIP1 | 2 | 9.62 | 10.52 | 9.12 | 9.29 | 8.04 | 7.49 | 10.83 | 14.36 | 9.38 | 11.35 | 10.07 | 9.21 | 7.76 | 12.59 | 10.37 |
| SNRNP200 | 120 | 10.19 | 9.80 | 10.70 | 10.26 | 9.63 | 9.63 | 9.64 | 9.26 | 10.51 | 10.38 | 9.99 | 10.48 | 9.63 | 9.45 | 10.45 |
| SNRNP27 | 4 | 9.39 | 9.59 | 12.01 | 9.32 | 10.19 | 10.59 | 10.52 | 9.27 | 10.44 | 8.68 | 9.49 | 10.67 | 10.39 | 9.90 | 9.56 |
| SNRNP40 | 15 | 9.45 | 10.92 | 9.86 | 9.54 | 8.78 | 9.16 | 9.59 | 13.13 | 9.72 | 9.84 | 10.19 | 9.70 | 8.97 | 11.36 | 9.78 |
| SNRNP48 | 2 | 9.40 | 9.63 | 9.64 | 9.78 | 11.02 | 9.75 | 10.88 | 8.61 | 10.95 | 10.35 | 9.51 | 9.71 | 10.38 | 9.74 | 10.65 |
| SNRNP70 | 27 | 9.97 | 10.81 | 9.37 | 8.61 | 10.74 | 10.45 | 10.91 | 9.66 | 9.86 | 9.62 | 10.39 | 8.99 | 10.59 | 10.29 | 9.74 |
| SNRPA | 24 | 10.06 | 10.73 | 9.49 | 9.42 | 10.10 | 10.55 | 10.81 | 9.96 | 9.56 | 9.32 | 10.40 | 9.46 | 10.32 | 10.38 | 9.44 |
| SNRPA1 | 22 | 8.62 | 10.38 | 10.52 | 10.09 | 9.98 | 10.70 | 11.00 | 9.98 | 9.42 | 9.30 | 9.50 | 10.31 | 10.34 | 10.49 | 9.36 |
| SNRPB | 15 | 8.82 | 9.55 | 10.93 | 8.71 | 9.62 | 8.48 | 10.47 | 15.01 | 9.04 | 9.37 | 9.18 | 9.82 | 9.05 | 12.74 | 9.20 |
| SNRPB2 | 13 | 9.57 | 10.05 | 11.11 | 9.20 | 10.37 | 9.97 | 11.47 | 9.49 | 9.69 | 9.08 | 9.81 | 10.15 | 10.17 | 10.48 | 9.39 |
| SNRPC | 7 | 9.11 | 11.07 | 9.84 | 9.71 | 9.97 | 11.05 | 10.38 | 9.92 | 9.61 | 9.33 | 10.09 | 9.77 | 10.51 | 10.15 | 9.47 |
| SNRPD1 | 3 | 9.01 | 10.76 | 9.90 | 9.48 | 9.60 | 10.87 | 9.04 | 9.85 | 10.30 | 11.18 | 9.89 | 9.69 | 10.24 | 9.45 | 10.74 |
| SNRPD2 | 21 | 8.71 | 8.82 | 9.13 | 8.80 | 9.69 | 9.52 | 8.81 | 19.66 | 8.47 | 8.38 | 8.77 | 8.96 | 9.61 | 14.24 | 8.43 |
| SNRPD3 | 14 | 9.16 | 8.96 | 9.04 | 7.34 | 8.32 | 8.45 | 9.27 | 23.38 | 8.31 | 7.77 | 9.06 | 8.19 | 8.38 | 16.32 | 8.04 |
| SNRPE | 4 | 9.73 | 10.50 | 9.60 | 8.83 | 11.06 | 10.97 | 9.99 | 10.10 | 10.10 | 9.12 | 10.12 | 9.21 | 11.01 | 10.05 | 9.61 |
| SNRPF | 8 | 7.23 | 9.44 | 10.93 | 9.66 | 10.81 | 10.37 | 10.54 | 12.80 | 8.46 | 9.75 | 8.34 | 10.30 | 10.59 | 11.67 | 9.10 |
| SNTA1 | 1 | 10.29 | 8.89 | 10.43 | 14.05 | 8.10 | 8.23 | 9.38 | 9.65 | 11.26 | 9.72 | 9.59 | 12.24 | 8.17 | 9.52 | 10.49 |
| SNTB2 | 12 | 11.49 | 11.26 | 8.95 | 8.60 | 10.09 | 10.22 | 10.21 | 9.04 | 10.09 | 10.04 | 11.37 | 8.78 | 10.15 | 9.63 | 10.07 |
| SNUPN | 3 | 8.87 | 10.11 | 10.89 | 10.33 | 10.32 | 11.57 | 7.97 | 7.04 | 12.32 | 10.58 | 9.49 | 10.61 | 10.95 | 7.50 | 11.45 |
| SNW1 | 23 | 9.08 | 10.12 | 11.78 | 10.41 | 9.52 | 9.59 | 10.85 | 9.72 | 9.79 | 9.14 | 9.60 | 11.09 | 9.56 | 10.28 | 9.46 |
| SNX1 | 28 | 9.02 | 9.61 | 11.11 | 10.47 | 11.01 | 10.93 | 8.47 | 7.73 | 10.95 | 10.70 | 9.31 | 10.79 | 10.97 | 8.10 | 10.82 |
| SNX12 | 18 | 10.46 | 11.05 | 8.77 | 8.29 | 13.11 | 13.23 | 8.69 | 7.93 | 9.55 | 8.92 | 10.76 | 8.53 | 13.17 | 8.31 | 9.23 |
| SNX15 | 4 | 10.65 | 10.69 | 10.59 | 10.66 | 9.65 | 10.05 | 10.23 | 9.19 | 9.61 | 8.67 | 10.67 | 10.63 | 9.85 | 9.71 | 9.14 |
| SNX17 | 8 | 9.91 | 10.60 | 10.96 | 9.74 | 10.18 | 10.01 | 8.16 | 8.29 | 10.90 | 11.25 | 10.26 | 10.35 | 10.09 | 8.23 | 11.07 |
| SNX18 | 7 | 7.53 | 8.39 | 18.00 | 16.38 | 6.51 | 6.38 | 8.54 | 7.66 | 10.28 | 10.33 | 7.96 | 17.19 | 6.45 | 8.10 | 10.30 |
| SNX2 | 14 | 9.45 | 10.39 | 11.99 | 11.20 | 10.17 | 10.09 | 9.38 | 8.85 | 9.51 | 8.97 | 9.92 | 11.60 | 10.13 | 9.12 | 9.24 |
| SNX24 | 1 | 11.40 | 11.56 | 11.59 | 10.45 | 12.57 | 12.45 | 5.86 | 5.58 | 10.60 | 7.95 | 11.48 | 11.02 | 12.51 | 5.72 | 9.27 |
| SNX27 | 14 | 9.25 | 9.68 | 10.79 | 10.72 | 10.23 | 10.47 | 9.29 | 9.52 | 10.13 | 9.92 | 9.46 | 10.75 | 10.35 | 9.41 | 10.03 |
| SNX29 | 1 | 12.12 | 12.96 | 11.55 | 8.01 | 7.79 | 11.17 | 10.48 | 6.64 | 10.08 | 9.21 | 12.54 | 9.78 | 9.48 | 8.56 | 9.65 |
| SNX3 | 10 | 9.00 | 9.78 | 11.44 | 10.08 | 9.50 | 9.43 | 8.57 | 11.18 | 10.75 | 10.27 | 9.39 | 10.76 | 9.47 | 9.87 | 10.51 |
| SNX33 | 3 | 8.47 | 10.96 | 11.30 | 9.77 | 9.41 | 9.03 | 9.69 | 7.48 | 12.08 | 11.81 | 9.71 | 10.53 | 9.22 | 8.59 | 11.95 |
| SNX4 | 9 | 10.94 | 11.03 | 11.83 | 12.05 | 8.27 | 8.37 | 8.40 | 8.41 | 10.67 | 10.03 | 10.98 | 11.94 | 8.32 | 8.41 | 10.35 |
| SNX5 | 12 | 9.75 | 10.38 | 10.64 | 10.04 | 10.32 | 10.29 | 9.93 | 9.41 | 9.99 | 9.25 | 10.06 | 10.34 | 10.30 | 9.67 | 9.62 |
| SNX6 | 20 | 9.17 | 9.83 | 11.17 | 10.61 | 10.15 | 10.39 | 8.52 | 10.50 | 10.06 | 9.60 | 9.50 | 10.89 | 10.27 | 9.51 | 9.83 |
| SNX7 | 7 | 11.03 | 12.07 | 11.44 | 11.86 | 7.82 | 8.42 | 7.36 | 7.58 | 11.56 | 10.88 | 11.55 | 11.65 | 8.12 | 7.47 | 11.22 |
| SNX8 | 4 | 10.54 | 11.34 | 11.25 | 9.04 | 7.95 | 7.82 | 9.64 | 8.64 | 11.73 | 12.05 | 10.94 | 10.14 | 7.89 | 9.14 | 11.89 |
| SNX9 | 12 | 8.33 | 8.92 | 12.50 | 12.01 | 8.01 | 8.50 | 11.79 | 10.86 | 9.61 | 9.47 | 8.62 | 12.25 | 8.26 | 11.32 | 9.54 |

| | | | | | | | | | | | | | | | | |
|---|---|---|---|---|---|---|---|---|---|---|---|---|---|---|---|---|
| SOAT1 | 4 | 12.27 | 6.31 | 5.65 | 8.33 | 10.11 | 8.52 | 14.44 | 12.61 | 10.41 | 11.36 | 9.29 | 6.99 | 9.32 | 13.52 | 10.89 |
| SOCS6 | 1 | 11.00 | 9.62 | 11.96 | 10.13 | 10.31 | 9.16 | 8.74 | 8.97 | 10.57 | 9.54 | 10.31 | 11.05 | 9.74 | 8.86 | 10.06 |
| SOD1 | 10 | 10.07 | 11.65 | 8.89 | 7.70 | 12.08 | 10.86 | 7.50 | 15.91 | 8.52 | 6.83 | 10.86 | 8.29 | 11.47 | 11.71 | 7.67 |
| SOD2 | 17 | 10.43 | 9.81 | 8.51 | 9.55 | 11.63 | 11.79 | 9.48 | 9.28 | 9.47 | 10.05 | 10.12 | 9.03 | 11.71 | 9.38 | 9.76 |
| SOGA1 | 2 | 9.33 | 9.37 | 9.09 | 8.80 | 9.04 | 9.12 | 11.67 | 11.94 | 11.12 | 10.53 | 9.35 | 8.95 | 9.08 | 11.80 | 10.82 |
| SOGA2 | 8 | 9.55 | 10.90 | 12.04 | 11.45 | 10.46 | 10.45 | 7.85 | 7.29 | 10.46 | 9.55 | 10.22 | 11.74 | 10.45 | 7.57 | 10.01 |
| SON | 41 | 10.29 | 11.29 | 9.24 | 8.55 | 10.70 | 10.78 | 9.78 | 8.86 | 10.54 | 9.96 | 10.79 | 8.90 | 10.74 | 9.32 | 10.25 |
| SORBS2 | 18 | 7.67 | 8.91 | 9.45 | 9.43 | 10.92 | 11.37 | 9.55 | 8.64 | 12.31 | 11.75 | 8.29 | 9.44 | 11.14 | 9.10 | 12.03 |
| SORBS3 | 10 | 10.61 | 11.93 | 9.06 | 8.28 | 11.22 | 11.80 | 7.37 | 6.39 | 11.81 | 11.53 | 11.27 | 8.67 | 11.51 | 6.88 | 11.67 |
| SORD | 17 | 8.75 | 9.01 | 11.10 | 8.66 | 10.37 | 9.76 | 8.09 | 11.10 | 11.87 | 11.28 | 8.88 | 9.88 | 10.07 | 9.60 | 11.57 |
| SOS1 | 4 | 10.78 | 10.97 | 9.41 | 9.58 | 10.73 | 11.16 | 8.86 | 7.82 | 10.43 | 10.25 | 10.87 | 9.50 | 10.95 | 8.34 | 10.34 |
| SOWAHC | 7 | 7.53 | 8.13 | 10.38 | 9.13 | 9.72 | 9.16 | 15.06 | 13.65 | 8.94 | 8.30 | 7.83 | 9.75 | 9.44 | 14.35 | 8.62 |
| SOX9 | 2 | 8.42 | 11.13 | 16.66 | 17.05 | 6.04 | 5.92 | 5.85 | 5.17 | 12.05 | 11.71 | 9.78 | 16.86 | 5.98 | 5.51 | 11.88 |
| SP1 | 3 | 9.19 | 9.96 | 9.93 | 9.08 | 9.95 | 9.79 | 12.86 | 10.86 | 9.75 | 8.64 | 9.57 | 9.50 | 9.87 | 11.86 | 9.19 |
| SP100 | 10 | 9.70 | 10.38 | 13.61 | 13.18 | 8.51 | 8.67 | 11.30 | 10.64 | 7.15 | 6.86 | 10.04 | 13.40 | 8.59 | 10.97 | 7.01 |
| SP2 | 2 | 8.62 | 10.54 | 11.35 | 9.63 | 9.78 | 10.94 | 11.43 | 10.07 | 8.47 | 9.17 | 9.58 | 10.49 | 10.36 | 10.75 | 8.82 |
| SP3 | 2 | 9.41 | 10.48 | 8.59 | 9.26 | 10.06 | 11.11 | 12.82 | 11.39 | 8.52 | 8.36 | 9.94 | 8.92 | 10.59 | 12.11 | 8.44 |
| SP4 | 2 | 9.40 | 11.05 | 9.90 | 11.47 | 7.51 | 9.27 | 12.19 | 12.86 | 7.82 | 8.53 | 10.23 | 10.68 | 8.39 | 12.52 | 8.17 |
| SPAG1 | 5 | 10.69 | 11.53 | 8.51 | 8.30 | 8.72 | 8.93 | 7.65 | 7.67 | 14.77 | 13.23 | 11.11 | 8.41 | 8.82 | 7.66 | 14.00 |
| SPAG5 | 17 | 10.18 | 11.57 | 11.52 | 10.70 | 8.70 | 9.36 | 7.61 | 7.16 | 11.48 | 11.72 | 10.88 | 11.11 | 9.03 | 7.38 | 11.60 |
| SPAG7 | 7 | 10.20 | 11.63 | 11.94 | 11.35 | 10.50 | 10.71 | 8.50 | 7.44 | 9.01 | 8.71 | 10.91 | 11.64 | 10.61 | 7.97 | 8.86 |
| SPAG9 | 41 | 10.98 | 11.43 | 11.10 | 9.98 | 8.24 | 8.14 | 8.70 | 8.37 | 11.89 | 11.18 | 11.20 | 10.54 | 8.19 | 8.53 | 11.53 |
| SPAST | 6 | 10.50 | 9.95 | 9.79 | 10.45 | 11.39 | 11.10 | 10.34 | 9.85 | 8.38 | 8.26 | 10.23 | 10.12 | 11.24 | 10.09 | 8.32 |
| SPATA20 | 2 | 10.95 | 9.30 | 9.31 | 9.32 | 10.38 | 10.36 | 5.71 | 5.28 | 15.03 | 14.35 | 10.13 | 9.32 | 10.37 | 5.50 | 14.69 |
| SPATA5 | 25 | 9.89 | 10.07 | 10.35 | 9.77 | 9.38 | 9.83 | 10.58 | 9.74 | 10.33 | 10.05 | 9.98 | 10.06 | 9.61 | 10.16 | 10.19 |
| SPATA5L1 | 14 | 8.71 | 10.03 | 10.06 | 9.07 | 10.09 | 9.88 | 10.63 | 9.74 | 10.96 | 10.83 | 9.37 | 9.56 | 9.98 | 10.19 | 10.90 |
| SPATS2 | 8 | 11.37 | 11.97 | 7.95 | 7.77 | 11.33 | 11.51 | 10.65 | 9.80 | 9.03 | 8.62 | 11.67 | 7.86 | 11.42 | 10.22 | 8.83 |
| SPATS2L | 17 | 9.72 | 10.08 | 11.20 | 10.43 | 8.11 | 8.21 | 8.65 | 11.33 | 11.34 | 10.92 | 9.90 | 10.81 | 8.16 | 9.99 | 11.13 |
| SPC24 | 5 | 9.64 | 11.85 | 11.45 | 11.54 | 12.77 | 13.32 | 10.45 | 6.79 | 6.13 | 6.06 | 10.75 | 11.50 | 13.04 | 8.62 | 6.09 |
| SPC25 | 9 | 10.09 | 10.82 | 11.09 | 10.16 | 12.61 | 12.91 | 10.37 | 9.21 | 6.64 | 6.12 | 10.45 | 10.62 | 12.76 | 9.79 | 6.38 |
| SPCS1 | 1 | 12.31 | 6.50 | 8.28 | 12.56 | 9.20 | 7.08 | 9.10 | 9.86 | 11.11 | 14.03 | 9.40 | 10.42 | 8.14 | 9.48 | 12.57 |
| SPCS2 | 10 | 11.66 | 6.91 | 8.60 | 13.38 | 8.56 | 8.33 | 8.83 | 8.48 | 11.19 | 14.06 | 9.29 | 10.99 | 8.44 | 8.66 | 12.62 |
| SPCS3 | 4 | 9.53 | 8.44 | 9.75 | 11.34 | 10.08 | 10.98 | 10.33 | 9.90 | 9.63 | 10.01 | 8.98 | 10.55 | 10.53 | 10.12 | 9.82 |
| SPDL1 | 18 | 10.77 | 11.77 | 9.64 | 9.48 | 10.34 | 10.85 | 8.06 | 9.02 | 10.27 | 9.80 | 11.27 | 9.56 | 10.60 | 8.54 | 10.03 |
| SPECC1 | 18 | 8.93 | 9.47 | 11.42 | 11.23 | 13.17 | 13.89 | 6.18 | 5.51 | 10.27 | 9.94 | 9.20 | 11.33 | 13.53 | 5.84 | 10.10 |
| SPECC1L | 18 | 10.72 | 12.26 | 12.18 | 11.47 | 8.13 | 8.53 | 7.77 | 7.29 | 10.88 | 10.78 | 11.49 | 11.82 | 8.33 | 7.53 | 10.83 |
| SPEG | 2 | 10.08 | 10.21 | 11.62 | 12.61 | 9.50 | 9.57 | 7.46 | 6.83 | 11.33 | 10.79 | 10.15 | 12.12 | 9.54 | 7.14 | 11.06 |
| SPEN | 26 | 9.27 | 10.16 | 10.07 | 10.23 | 8.61 | 9.27 | 12.45 | 11.72 | 9.25 | 8.96 | 9.72 | 10.15 | 8.94 | 12.09 | 9.10 |
| SPG11 | 1 | 11.18 | 10.27 | 10.70 | 9.88 | 8.97 | 9.11 | 9.38 | 7.46 | 11.20 | 11.85 | 10.73 | 10.29 | 9.04 | 8.42 | 11.53 |
| SPG20 | 13 | 10.98 | 11.78 | 11.48 | 10.95 | 13.20 | 13.05 | 7.70 | 6.73 | 7.27 | 6.86 | 11.38 | 11.22 | 13.12 | 7.22 | 7.07 |
| SPG21 | 4 | 9.28 | 9.66 | 9.87 | 10.64 | 12.82 | 12.18 | 7.96 | 7.80 | 10.11 | 9.68 | 9.47 | 10.25 | 12.50 | 7.88 | 9.89 |
| SPHK1 | 4 | 9.75 | 10.67 | 11.03 | 12.59 | 9.90 | 10.01 | 6.33 | 6.75 | 11.37 | 11.61 | 10.21 | 11.81 | 9.96 | 6.54 | 11.49 |

| | | | | | | | | | | | | | | | | |
|---|---|---|---|---|---|---|---|---|---|---|---|---|---|---|---|---|
| SPICE1 | 1 | 7.93 | 10.56 | 9.34 | 8.02 | 12.80 | 13.81 | 11.00 | 8.67 | 9.05 | 8.82 | 9.25 | 8.68 | 13.31 | 9.83 | 8.93 |
| SPIN1 | 5 | 11.17 | 11.67 | 8.62 | 9.24 | 10.70 | 11.33 | 8.45 | 7.72 | 10.47 | 10.64 | 11.42 | 8.93 | 11.01 | 8.08 | 10.56 |
| SPNS1 | 2 | 9.94 | 7.37 | 8.93 | 10.97 | 11.39 | 10.24 | 9.00 | 9.60 | 10.47 | 12.07 | 8.66 | 9.95 | 10.82 | 9.30 | 11.27 |
| SPOCK1 | 1 | 5.27 | 9.17 | 12.19 | 15.52 | 6.87 | 9.53 | 9.15 | 7.99 | 12.61 | 11.71 | 7.22 | 13.86 | 8.20 | 8.57 | 12.16 |
| SPPL2A | 1 | 10.04 | 9.42 | 6.99 | 9.76 | 11.60 | 12.17 | 7.52 | 7.90 | 11.38 | 13.21 | 9.73 | 8.37 | 11.89 | 7.71 | 12.30 |
| SPR | 14 | 7.85 | 8.05 | 12.65 | 11.10 | 10.10 | 9.77 | 8.27 | 13.60 | 9.66 | 8.94 | 7.95 | 11.87 | 9.93 | 10.94 | 9.30 |
| SPRYD3 | 1 | 9.11 | 8.24 | 11.53 | 9.12 | 8.89 | 9.12 | 11.95 | 11.46 | 11.16 | 9.44 | 8.67 | 10.32 | 9.01 | 11.70 | 10.30 |
| SPRYD4 | 5 | 9.86 | 8.17 | 12.29 | 13.37 | 7.69 | 7.43 | 6.91 | 6.36 | 13.27 | 14.67 | 9.01 | 12.83 | 7.56 | 6.63 | 13.97 |
| SPRYD7 | 1 | 12.74 | 10.41 | 10.14 | 11.45 | 11.62 | 10.54 | 6.16 | 6.85 | 10.24 | 9.86 | 11.57 | 10.79 | 11.08 | 6.50 | 10.05 |
| SPTAN1 | 250 | 9.84 | 10.38 | 11.85 | 10.79 | 11.70 | 11.70 | 6.53 | 6.34 | 10.71 | 10.16 | 10.11 | 11.32 | 11.70 | 6.44 | 10.44 |
| SPTBN1 | 197 | 9.74 | 10.23 | 11.92 | 10.59 | 11.66 | 11.50 | 6.78 | 6.59 | 10.73 | 10.26 | 9.99 | 11.25 | 11.58 | 6.69 | 10.50 |
| SPTBN2 | 54 | 11.87 | 11.69 | 10.00 | 10.04 | 9.77 | 10.36 | 7.93 | 7.14 | 10.95 | 10.25 | 11.78 | 10.02 | 10.07 | 7.54 | 10.60 |
| SPTBN4 | 11 | 11.10 | 11.18 | 11.15 | 11.72 | 9.87 | 9.86 | 7.09 | 6.82 | 10.78 | 10.43 | 11.14 | 11.43 | 9.87 | 6.95 | 10.60 |
| SPTLC1 | 9 | 10.33 | 8.93 | 10.74 | 11.57 | 9.95 | 9.97 | 9.42 | 9.11 | 9.91 | 10.07 | 9.63 | 11.16 | 9.96 | 9.26 | 9.99 |
| SPTLC2 | 7 | 9.82 | 9.99 | 10.58 | 11.53 | 9.08 | 9.55 | 9.63 | 10.22 | 9.81 | 9.81 | 9.90 | 11.05 | 9.31 | 9.92 | 9.81 |
| SQRDL | 12 | 12.33 | 7.09 | 4.87 | 6.59 | 17.43 | 15.63 | 4.70 | 4.33 | 12.72 | 14.31 | 9.71 | 5.73 | 16.53 | 4.52 | 13.51 |
| SQSTM1 | 18 | 12.58 | 14.92 | 8.79 | 8.27 | 10.72 | 10.81 | 5.01 | 6.59 | 11.12 | 11.20 | 13.75 | 8.53 | 10.76 | 5.80 | 11.16 |
| SRA1 | 7 | 10.45 | 11.08 | 9.50 | 9.53 | 10.35 | 11.00 | 8.57 | 8.14 | 10.79 | 10.60 | 10.76 | 9.51 | 10.67 | 8.36 | 10.70 |
| SRBD1 | 5 | 10.49 | 11.52 | 11.13 | 10.92 | 8.90 | 8.63 | 9.36 | 10.99 | 8.76 | 9.30 | 11.00 | 11.02 | 8.77 | 10.18 | 9.03 |
| SRCAP | 15 | 8.27 | 8.86 | 11.37 | 11.57 | 10.81 | 10.58 | 10.54 | 10.51 | 8.72 | 8.75 | 8.57 | 11.47 | 10.70 | 10.53 | 8.74 |
| SREBF2 | 1 | 9.57 | 11.23 | 8.82 | 9.73 | 12.01 | 10.83 | 9.86 | 7.64 | 11.31 | 8.99 | 10.40 | 9.28 | 11.42 | 8.75 | 10.15 |
| SREK1 | 13 | 8.97 | 10.84 | 9.26 | 8.41 | 11.30 | 11.82 | 10.81 | 10.14 | 9.27 | 9.19 | 9.90 | 8.83 | 11.56 | 10.48 | 9.23 |
| SREK1IP1 | 2 | 10.35 | 10.72 | 9.49 | 9.39 | 7.95 | 9.60 | 12.13 | 11.57 | 9.28 | 9.51 | 10.54 | 9.44 | 8.77 | 11.85 | 9.40 |
| SRFBP1 | 11 | 9.44 | 10.36 | 10.02 | 9.70 | 8.22 | 8.61 | 12.36 | 11.37 | 9.95 | 9.98 | 9.90 | 9.86 | 8.41 | 11.86 | 9.96 |
| SRGAP1 | 6 | 9.71 | 9.67 | 10.53 | 9.38 | 11.36 | 12.13 | 9.41 | 8.94 | 9.84 | 9.02 | 9.69 | 9.96 | 11.74 | 9.18 | 9.43 |
| SRGAP2 | 12 | 9.56 | 10.12 | 9.04 | 9.13 | 13.06 | 13.93 | 8.35 | 8.48 | 9.36 | 8.97 | 9.84 | 9.08 | 13.50 | 8.41 | 9.17 |
| SRI | 29 | 10.16 | 11.22 | 11.55 | 10.60 | 10.02 | 10.20 | 7.31 | 10.45 | 9.39 | 9.11 | 10.69 | 11.08 | 10.11 | 8.88 | 9.25 |
| SRM | 25 | 8.95 | 9.85 | 8.29 | 7.41 | 9.35 | 9.39 | 12.82 | 13.29 | 10.25 | 10.40 | 9.40 | 7.85 | 9.37 | 13.05 | 10.33 |
| SRP14 | 23 | 8.10 | 10.18 | 9.62 | 8.74 | 13.09 | 13.67 | 9.76 | 9.24 | 8.72 | 8.88 | 9.14 | 9.18 | 13.38 | 9.50 | 8.80 |
| SRP19 | 1 | 9.96 | 10.01 | 9.73 | 9.37 | 10.05 | 10.62 | 10.50 | 9.31 | 11.03 | 9.41 | 9.99 | 9.55 | 10.33 | 9.91 | 10.22 |
| SRP54 | 29 | 9.48 | 10.42 | 10.18 | 9.68 | 10.46 | 10.50 | 9.78 | 11.41 | 9.20 | 8.89 | 9.95 | 9.93 | 10.48 | 10.59 | 9.05 |
| SRP68 | 28 | 10.12 | 10.50 | 10.49 | 9.85 | 10.31 | 10.09 | 9.31 | 9.54 | 10.17 | 9.62 | 10.31 | 10.17 | 10.20 | 9.43 | 9.90 |
| SRP72 | 31 | 10.10 | 10.78 | 10.62 | 10.56 | 10.23 | 10.43 | 9.25 | 8.92 | 9.62 | 9.51 | 10.44 | 10.59 | 10.33 | 9.08 | 9.56 |
| SRP9 | 15 | 9.35 | 9.62 | 8.89 | 8.26 | 13.19 | 12.97 | 10.32 | 10.64 | 8.75 | 8.01 | 9.49 | 8.58 | 13.08 | 10.48 | 8.38 |
| SRPK1 | 11 | 9.53 | 11.37 | 8.93 | 8.88 | 9.43 | 9.94 | 11.78 | 11.69 | 9.53 | 8.93 | 10.45 | 8.91 | 9.68 | 11.73 | 9.23 |
| SRPK2 | 14 | 8.92 | 10.95 | 11.86 | 11.91 | 8.13 | 8.46 | 10.07 | 9.80 | 10.02 | 9.87 | 9.93 | 11.89 | 8.30 | 9.94 | 9.94 |
| SRPR | 13 | 11.37 | 6.99 | 8.24 | 11.48 | 9.66 | 8.06 | 10.59 | 10.03 | 10.42 | 13.16 | 9.18 | 9.86 | 8.86 | 10.31 | 11.79 |
| SRPRB | 7 | 12.77 | 6.36 | 7.81 | 12.34 | 9.88 | 7.38 | 10.06 | 9.42 | 10.43 | 13.55 | 9.56 | 10.07 | 8.63 | 9.74 | 11.99 |
| SRRM1 | 14 | 9.24 | 10.04 | 9.19 | 8.32 | 8.50 | 8.66 | 11.93 | 14.66 | 9.91 | 9.56 | 9.64 | 8.75 | 8.58 | 13.29 | 9.74 |
| SRRM2 | 67 | 8.82 | 10.24 | 12.05 | 11.42 | 9.41 | 9.92 | 10.57 | 9.47 | 9.29 | 8.82 | 9.53 | 11.73 | 9.66 | 10.02 | 9.06 |
| SRRT | 50 | 9.14 | 9.70 | 10.59 | 9.59 | 9.02 | 8.96 | 11.34 | 14.45 | 8.91 | 8.30 | 9.42 | 10.09 | 8.99 | 12.89 | 8.61 |
| SRSF1 | 33 | 9.68 | 10.47 | 10.04 | 9.26 | 10.07 | 10.32 | 11.13 | 10.43 | 9.70 | 8.90 | 10.08 | 9.65 | 10.19 | 10.78 | 9.30 |

| | | | | | | | | | | | | | | | | |
|---|---|---|---|---|---|---|---|---|---|---|---|---|---|---|---|---|
| SRSF10 | 14 | 9.50 | 11.35 | 9.29 | 9.03 | 9.63 | 10.25 | 11.35 | 10.71 | 9.60 | 9.28 | 10.42 | 9.16 | 9.94 | 11.03 | 9.44 |
| SRSF11 | 9 | 10.59 | 10.37 | 10.02 | 7.59 | 10.94 | 10.09 | 11.53 | 9.12 | 10.15 | 9.61 | 10.48 | 8.81 | 10.51 | 10.32 | 9.88 |
| SRSF2 | 10 | 9.91 | 11.22 | 9.29 | 9.10 | 9.37 | 10.36 | 11.00 | 9.88 | 10.08 | 9.78 | 10.56 | 9.20 | 9.87 | 10.44 | 9.93 |
| SRSF3 | 21 | 8.46 | 9.33 | 9.58 | 8.96 | 9.84 | 9.56 | 10.01 | 17.97 | 8.24 | 8.06 | 8.90 | 9.27 | 9.70 | 13.99 | 8.15 |
| SRSF4 | 25 | 9.59 | 11.19 | 8.62 | 7.72 | 9.12 | 9.25 | 12.01 | 14.26 | 9.30 | 8.94 | 10.39 | 8.17 | 9.18 | 13.13 | 9.12 |
| SRSF5 | 7 | 10.23 | 10.35 | 8.82 | 8.87 | 9.91 | 9.99 | 11.68 | 11.21 | 9.61 | 9.34 | 10.29 | 8.85 | 9.95 | 11.45 | 9.47 |
| SRSF6 | 9 | 6.59 | 10.73 | 9.94 | 9.21 | 11.24 | 11.51 | 11.46 | 10.46 | 9.16 | 9.69 | 8.66 | 9.58 | 11.38 | 10.96 | 9.43 |
| SRSF7 | 13 | 9.47 | 10.20 | 9.46 | 8.78 | 10.27 | 10.14 | 11.47 | 12.12 | 9.14 | 8.96 | 9.83 | 9.12 | 10.20 | 11.80 | 9.05 |
| SRSF8 | 4 | 9.69 | 10.67 | 11.21 | 9.27 | 9.66 | 9.38 | 11.37 | 9.16 | 10.18 | 9.40 | 10.18 | 10.24 | 9.52 | 10.27 | 9.79 |
| SRSF9 | 11 | 8.16 | 8.91 | 13.58 | 12.96 | 9.31 | 8.50 | 10.89 | 8.82 | 9.40 | 9.47 | 8.54 | 13.27 | 8.90 | 9.85 | 9.44 |
| SRXN1 | 6 | 12.27 | 14.41 | 7.47 | 6.54 | 9.27 | 9.67 | 8.54 | 8.34 | 11.72 | 11.79 | 13.34 | 7.00 | 9.47 | 8.44 | 11.75 |
| SS18 | 1 | 7.90 | 8.07 | 10.45 | 12.16 | 10.85 | 11.57 | 9.29 | 7.64 | 9.46 | 12.61 | 7.98 | 11.30 | 11.21 | 8.47 | 11.04 |
| SS18L1 | 1 | 11.21 | 12.28 | 11.35 | 10.92 | 9.19 | 11.42 | 10.77 | 11.70 | 4.37 | 6.77 | 11.74 | 11.14 | 10.31 | 11.24 | 5.57 |
| SSB | 45 | 9.12 | 10.41 | 10.87 | 10.05 | 9.83 | 10.07 | 11.09 | 10.77 | 8.99 | 8.79 | 9.77 | 10.46 | 9.95 | 10.93 | 8.89 |
| SSBP1 | 15 | 10.88 | 8.99 | 11.04 | 11.50 | 7.05 | 6.65 | 12.17 | 10.90 | 10.73 | 10.09 | 9.94 | 11.27 | 6.85 | 11.53 | 10.41 |
| SSBP2 | 1 | 11.78 | 13.48 | 10.97 | 10.60 | 8.86 | 8.86 | 7.70 | 7.18 | 10.31 | 10.26 | 12.63 | 10.78 | 8.86 | 7.44 | 10.29 |
| SSBP3 | 1 | 10.57 | 6.91 | 8.23 | 7.79 | 12.40 | 8.13 | 14.56 | 10.49 | 10.03 | 10.89 | 8.74 | 8.01 | 10.26 | 12.53 | 10.46 |
| SSFA2 | 28 | 8.37 | 9.57 | 7.27 | 6.60 | 12.34 | 12.74 | 13.84 | 12.73 | 8.47 | 8.06 | 8.97 | 6.93 | 12.54 | 13.29 | 8.27 |
| SSH3 | 4 | 9.49 | 11.20 | 9.07 | 8.90 | 12.71 | 13.20 | 7.02 | 6.62 | 11.14 | 10.65 | 10.34 | 8.98 | 12.96 | 6.82 | 10.89 |
| SSNA1 | 3 | 8.54 | 10.50 | 7.69 | 10.27 | 14.56 | 14.61 | 10.12 | 4.13 | 9.40 | 10.19 | 9.52 | 8.98 | 14.59 | 7.12 | 9.79 |
| SSR1 | 6 | 12.40 | 7.18 | 6.39 | 13.57 | 9.62 | 10.38 | 10.14 | 7.89 | 9.85 | 12.58 | 9.79 | 9.98 | 10.00 | 9.02 | 11.22 |
| SSR3 | 2 | 14.13 | 5.62 | 6.70 | 10.88 | 9.90 | 7.78 | 11.58 | 9.18 | 11.20 | 13.05 | 9.88 | 8.79 | 8.84 | 10.38 | 12.12 |
| SSR4 | 6 | 9.69 | 7.71 | 8.84 | 8.72 | 12.13 | 11.58 | 11.17 | 12.04 | 8.76 | 9.35 | 8.70 | 8.78 | 11.85 | 11.61 | 9.06 |
| SSRP1 | 35 | 8.91 | 11.31 | 10.55 | 9.58 | 9.85 | 9.86 | 9.85 | 9.53 | 10.07 | 10.49 | 10.11 | 10.07 | 9.85 | 9.69 | 10.28 |
| SSSCA1 | 5 | 5.57 | 6.28 | 11.69 | 9.11 | 7.73 | 6.73 | 8.58 | 30.92 | 6.77 | 6.61 | 5.93 | 10.40 | 7.23 | 19.75 | 6.69 |
| SSU72 | 4 | 11.65 | 10.26 | 9.46 | 8.41 | 9.72 | 9.34 | 11.49 | 9.43 | 10.57 | 9.67 | 10.95 | 8.93 | 9.53 | 10.46 | 10.12 |
| ST13 | 41 | 9.67 | 10.42 | 13.75 | 12.38 | 8.36 | 8.26 | 9.20 | 7.91 | 10.37 | 9.68 | 10.04 | 13.06 | 8.31 | 8.56 | 10.03 |
| ST5 | 3 | 9.94 | 10.92 | 10.08 | 11.59 | 8.54 | 8.38 | 9.39 | 8.58 | 11.73 | 10.86 | 10.43 | 10.83 | 8.46 | 8.98 | 11.29 |
| ST6GALNAC1 | 1 | 8.71 | 11.21 | 9.98 | 11.46 | 9.15 | 10.85 | 8.46 | 8.74 | 10.04 | 11.40 | 9.96 | 10.72 | 10.00 | 8.60 | 10.72 |
| STAC2 | 1 | 17.45 | 19.82 | 5.33 | 5.48 | 5.75 | 5.48 | 3.76 | 3.78 | 17.69 | 15.47 | 18.64 | 5.40 | 5.61 | 3.77 | 16.58 |
| STAG1 | 8 | 9.46 | 9.71 | 11.64 | 9.80 | 8.68 | 8.45 | 10.64 | 9.48 | 11.16 | 10.99 | 9.58 | 10.72 | 8.56 | 10.06 | 11.07 |
| STAG2 | 19 | 9.43 | 9.72 | 8.14 | 8.29 | 10.66 | 11.20 | 11.55 | 14.45 | 8.14 | 8.40 | 9.57 | 8.22 | 10.93 | 13.00 | 8.27 |
| STAM | 11 | 9.36 | 10.79 | 11.71 | 10.75 | 10.19 | 10.26 | 8.78 | 8.24 | 10.32 | 9.61 | 10.07 | 11.23 | 10.22 | 8.51 | 9.96 |
| STAM2 | 7 | 7.81 | 12.79 | 12.45 | 12.94 | 7.93 | 9.00 | 6.99 | 6.99 | 11.06 | 12.03 | 10.30 | 12.70 | 8.46 | 6.99 | 11.54 |
| STAMBP | 6 | 10.08 | 11.61 | 10.77 | 11.57 | 8.95 | 9.39 | 8.48 | 8.94 | 10.17 | 10.04 | 10.84 | 11.17 | 9.17 | 8.71 | 10.10 |
| STAMBPL1 | 3 | 10.29 | 10.61 | 8.04 | 7.66 | 11.21 | 11.67 | 7.28 | 7.44 | 13.14 | 12.68 | 10.45 | 7.85 | 11.44 | 7.36 | 12.91 |
| STARD13 | 1 | 8.93 | 10.02 | 13.23 | 12.49 | 9.82 | 9.41 | 7.94 | 6.14 | 10.34 | 11.67 | 9.48 | 12.86 | 9.61 | 7.04 | 11.01 |
| STARD3NL | 2 | 12.67 | 12.07 | 7.76 | 8.01 | 8.51 | 8.86 | 12.00 | 11.32 | 10.14 | 8.66 | 12.37 | 7.89 | 8.68 | 11.66 | 9.40 |
| STARD7 | 1 | 10.79 | 10.31 | 8.98 | 8.53 | 10.27 | 11.80 | 9.49 | 10.17 | 9.81 | 9.85 | 10.55 | 8.75 | 11.04 | 9.83 | 9.83 |
| STAT1 | 46 | 10.58 | 10.74 | 10.61 | 10.44 | 9.92 | 10.07 | 9.16 | 9.15 | 9.78 | 9.54 | 10.66 | 10.53 | 9.99 | 9.16 | 9.66 |
| STAT2 | 3 | 10.48 | 10.40 | 7.86 | 7.91 | 13.51 | 14.61 | 6.28 | 7.15 | 10.92 | 10.89 | 10.44 | 7.88 | 14.06 | 6.72 | 10.90 |
| STAT3 | 28 | 10.54 | 11.05 | 11.67 | 11.30 | 9.54 | 9.71 | 8.24 | 7.93 | 10.17 | 9.85 | 10.79 | 11.49 | 9.63 | 8.08 | 10.01 |

| | | | | | | | | | | | | | | | | |
|---|---|---|---|---|---|---|---|---|---|---|---|---|---|---|---|---|
| STAT5B | 16 | 11.84 | 11.97 | 11.32 | 10.73 | 9.65 | 9.95 | 8.12 | 7.86 | 9.47 | 9.10 | 11.90 | 11.02 | 9.80 | 7.99 | 9.28 |
| STAT6 | 12 | 9.57 | 10.27 | 11.04 | 9.54 | 12.02 | 12.51 | 8.18 | 7.95 | 9.37 | 9.55 | 9.92 | 10.29 | 12.26 | 8.07 | 9.46 |
| STAU1 | 11 | 9.85 | 11.12 | 9.99 | 10.12 | 10.19 | 10.36 | 10.43 | 9.87 | 9.10 | 8.97 | 10.48 | 10.05 | 10.28 | 10.15 | 9.03 |
| STAU2 | 15 | 9.33 | 9.87 | 11.39 | 10.66 | 10.94 | 10.37 | 11.48 | 10.27 | 7.84 | 7.85 | 9.60 | 11.02 | 10.65 | 10.88 | 7.84 |
| STBD1 | 3 | 11.40 | 6.50 | 11.15 | 13.28 | 9.59 | 7.99 | 7.27 | 6.55 | 12.44 | 13.84 | 8.95 | 12.21 | 8.79 | 6.91 | 13.14 |
| STC2 | 3 | 6.08 | 5.82 | 13.75 | 14.70 | 7.57 | 8.09 | 10.04 | 10.30 | 11.64 | 12.01 | 5.95 | 14.22 | 7.83 | 10.17 | 11.82 |
| STEAP3 | 1 | 9.26 | 10.01 | 8.39 | 9.63 | 8.10 | 9.90 | 10.55 | 12.77 | 10.85 | 10.54 | 9.63 | 9.01 | 9.00 | 11.66 | 10.70 |
| STIL | 1 | 9.75 | 9.91 | 10.04 | 13.29 | 9.06 | 5.50 | 9.22 | 9.87 | 11.75 | 11.62 | 9.83 | 11.66 | 7.28 | 9.54 | 11.69 |
| STIM1 | 3 | 11.77 | 7.15 | 8.43 | 10.84 | 10.71 | 9.28 | 10.66 | 9.45 | 10.26 | 11.47 | 9.46 | 9.63 | 9.99 | 10.05 | 10.86 |
| STIP1 | 125 | 8.78 | 9.53 | 12.35 | 10.59 | 9.43 | 8.99 | 10.13 | 11.14 | 9.77 | 9.28 | 9.15 | 11.47 | 9.21 | 10.64 | 9.53 |
| STK10 | 10 | 8.63 | 9.77 | 12.18 | 12.03 | 7.94 | 8.28 | 10.59 | 10.34 | 10.18 | 10.07 | 9.20 | 12.10 | 8.11 | 10.46 | 10.12 |
| STK11IP | 4 | 9.03 | 9.64 | 10.05 | 9.60 | 10.98 | 11.31 | 10.08 | 9.66 | 10.50 | 9.16 | 9.33 | 9.82 | 11.14 | 9.87 | 9.83 |
| STK17A | 1 | 8.08 | 9.49 | 18.14 | 16.20 | 5.41 | 5.86 | 7.16 | 6.30 | 11.80 | 11.57 | 8.79 | 17.17 | 5.63 | 6.73 | 11.69 |
| STK24 | 10 | 10.09 | 10.42 | 11.61 | 10.70 | 9.84 | 9.97 | 10.11 | 8.94 | 9.56 | 8.74 | 10.25 | 11.15 | 9.91 | 9.53 | 9.15 |
| STK25 | 3 | 9.76 | 10.20 | 11.12 | 11.06 | 9.54 | 9.19 | 9.41 | 8.22 | 10.97 | 10.53 | 9.98 | 11.09 | 9.36 | 8.82 | 10.75 |
| STK3 | 18 | 9.29 | 10.87 | 9.04 | 9.10 | 11.19 | 12.58 | 11.42 | 10.75 | 8.14 | 7.62 | 10.08 | 9.07 | 11.88 | 11.09 | 7.88 |
| STK32C | 3 | 9.78 | 9.34 | 12.61 | 12.21 | 9.99 | 9.79 | 8.06 | 7.50 | 10.60 | 10.13 | 9.56 | 12.41 | 9.89 | 7.78 | 10.37 |
| STK33 | 1 | 8.73 | 12.42 | 11.07 | 12.89 | 7.35 | 8.56 | 10.84 | 12.32 | 7.59 | 8.22 | 10.58 | 11.98 | 7.96 | 11.58 | 7.91 |
| STK38 | 3 | 11.03 | 11.17 | 9.94 | 8.62 | 10.04 | 9.63 | 7.44 | 6.63 | 13.32 | 12.17 | 11.10 | 9.28 | 9.84 | 7.04 | 12.74 |
| STK38L | 11 | 8.32 | 9.04 | 18.02 | 16.76 | 7.36 | 7.49 | 5.97 | 6.05 | 10.84 | 10.15 | 8.68 | 17.39 | 7.43 | 6.01 | 10.49 |
| STK39 | 8 | 9.56 | 10.09 | 11.76 | 10.55 | 10.67 | 10.06 | 8.64 | 9.56 | 9.62 | 9.50 | 9.82 | 11.16 | 10.36 | 9.10 | 9.56 |
| STK4 | 9 | 8.87 | 10.13 | 11.59 | 11.55 | 10.18 | 10.89 | 9.77 | 8.95 | 9.05 | 9.03 | 9.50 | 11.57 | 10.53 | 9.36 | 9.04 |
| STMN1 | 25 | 10.01 | 11.75 | 8.47 | 8.75 | 11.36 | 12.16 | 10.16 | 5.51 | 11.23 | 10.59 | 10.88 | 8.61 | 11.76 | 7.83 | 10.91 |
| STOM | 17 | 11.89 | 9.69 | 8.59 | 10.19 | 10.06 | 9.89 | 6.05 | 5.27 | 13.72 | 14.64 | 10.79 | 9.39 | 9.98 | 5.66 | 14.18 |
| STOML2 | 18 | 11.05 | 9.20 | 8.93 | 10.77 | 7.60 | 7.68 | 8.09 | 7.35 | 14.13 | 15.19 | 10.12 | 9.85 | 7.64 | 7.72 | 14.66 |
| STON2 | 1 | 13.17 | 13.98 | 9.03 | 8.43 | 9.07 | 11.11 | 6.87 | 4.74 | 12.87 | 10.72 | 13.58 | 8.73 | 10.09 | 5.81 | 11.79 |
| STRA13 | 1 | 11.32 | 12.18 | 6.96 | 9.04 | 9.65 | 11.49 | 9.88 | 4.70 | 12.46 | 12.31 | 11.75 | 8.00 | 10.57 | 7.29 | 12.39 |
| STRAP | 36 | 9.92 | 10.28 | 11.83 | 9.49 | 9.46 | 8.74 | 10.67 | 10.89 | 9.62 | 9.09 | 10.10 | 10.66 | 9.10 | 10.78 | 9.35 |
| STRBP | 13 | 9.05 | 10.08 | 10.47 | 10.34 | 9.37 | 9.99 | 12.00 | 10.95 | 9.07 | 8.67 | 9.57 | 10.40 | 9.68 | 11.48 | 8.87 |
| STRIP1 | 6 | 10.92 | 10.92 | 9.94 | 9.44 | 7.98 | 7.84 | 10.02 | 10.24 | 11.40 | 11.29 | 10.92 | 9.69 | 7.91 | 10.13 | 11.35 |
| STRN | 11 | 9.64 | 10.60 | 10.57 | 9.89 | 10.57 | 10.74 | 10.37 | 8.84 | 9.86 | 8.91 | 10.12 | 10.23 | 10.66 | 9.60 | 9.39 |
| STRN3 | 10 | 8.99 | 10.12 | 13.06 | 12.01 | 8.56 | 8.52 | 9.76 | 8.25 | 10.48 | 10.25 | 9.55 | 12.54 | 8.54 | 9.00 | 10.37 |
| STRN4 | 6 | 10.29 | 11.84 | 10.12 | 9.56 | 9.17 | 8.87 | 10.74 | 9.64 | 10.13 | 9.64 | 11.07 | 9.84 | 9.02 | 10.19 | 9.88 |
| STS | 2 | 10.56 | 7.68 | 5.55 | 7.43 | 12.44 | 12.95 | 11.98 | 11.87 | 8.37 | 11.17 | 9.12 | 6.49 | 12.70 | 11.92 | 9.77 |
| STT3A | 12 | 14.36 | 6.69 | 7.68 | 11.74 | 7.16 | 6.12 | 10.39 | 9.71 | 11.15 | 15.00 | 10.53 | 9.71 | 6.64 | 10.05 | 13.07 |
| STT3B | 10 | 13.98 | 6.67 | 7.62 | 11.99 | 7.95 | 6.92 | 9.26 | 8.98 | 11.84 | 14.79 | 10.33 | 9.81 | 7.43 | 9.12 | 13.32 |
| STUB1 | 14 | 9.70 | 10.40 | 11.26 | 10.83 | 10.55 | 10.67 | 8.42 | 7.25 | 10.66 | 10.26 | 10.05 | 11.04 | 10.61 | 7.83 | 10.46 |
| STX10 | 1 | 9.77 | 7.67 | 9.91 | 8.52 | 11.07 | 9.11 | 15.39 | 9.04 | 9.00 | 10.52 | 8.72 | 9.21 | 10.09 | 12.21 | 9.76 |
| STX12 | 5 | 11.71 | 9.72 | 8.52 | 9.77 | 10.49 | 9.90 | 9.18 | 8.25 | 11.01 | 11.45 | 10.72 | 9.15 | 10.19 | 8.71 | 11.23 |
| STX16 | 1 | 10.60 | 8.05 | 9.67 | 10.58 | 9.77 | 9.85 | 8.61 | 8.48 | 11.64 | 12.76 | 9.32 | 10.12 | 9.81 | 8.55 | 12.20 |
| STX17 | 1 | 11.77 | 12.13 | 7.91 | 9.31 | 11.05 | 14.23 | 8.48 | 7.34 | 9.68 | 8.11 | 11.95 | 8.61 | 12.64 | 7.91 | 8.90 |
| STX18 | 3 | 11.04 | 6.97 | 7.97 | 10.23 | 8.43 | 8.19 | 8.58 | 15.60 | 10.39 | 12.60 | 9.00 | 9.10 | 8.31 | 12.09 | 11.50 |

| | | | | | | | | | | | | | | | | |
|---|---|---|---|---|---|---|---|---|---|---|---|---|---|---|---|---|
| STX2 | 1 | 11.37 | 12.41 | 9.81 | 11.89 | 7.79 | 8.47 | 6.49 | 8.90 | 10.93 | 11.93 | 11.89 | 10.85 | 8.13 | 7.70 | 11.43 |
| STX3 | 4 | 14.64 | 12.21 | 7.26 | 8.66 | 9.04 | 10.61 | 8.74 | 9.02 | 9.87 | 9.96 | 13.43 | 7.96 | 9.82 | 8.88 | 9.91 |
| STX4 | 7 | 10.50 | 8.18 | 9.26 | 11.72 | 9.35 | 9.25 | 10.43 | 8.93 | 10.68 | 11.70 | 9.34 | 10.49 | 9.30 | 9.68 | 11.19 |
| STX5 | 8 | 11.17 | 8.50 | 8.59 | 10.42 | 10.72 | 10.17 | 9.65 | 8.83 | 10.03 | 11.91 | 9.83 | 9.50 | 10.45 | 9.24 | 10.97 |
| STX6 | 1 | 9.43 | 7.96 | 8.31 | 9.30 | 10.89 | 11.62 | 10.49 | 10.29 | 10.74 | 10.97 | 8.69 | 8.81 | 11.25 | 10.39 | 10.86 |
| STX7 | 6 | 11.03 | 8.62 | 12.16 | 14.15 | 9.81 | 9.10 | 7.31 | 6.88 | 9.92 | 11.03 | 9.82 | 13.15 | 9.45 | 7.10 | 10.47 |
| STX8 | 3 | 11.17 | 8.74 | 9.99 | 12.51 | 9.45 | 9.19 | 9.77 | 9.87 | 9.74 | 9.57 | 9.96 | 11.25 | 9.32 | 9.82 | 9.66 |
| STXBP1 | 22 | 11.28 | 12.27 | 7.32 | 7.16 | 13.48 | 13.74 | 8.46 | 8.16 | 9.29 | 8.85 | 11.78 | 7.24 | 13.61 | 8.31 | 9.07 |
| STXBP2 | 10 | 10.66 | 11.40 | 8.76 | 7.52 | 13.98 | 14.52 | 8.93 | 7.99 | 8.46 | 7.78 | 11.03 | 8.14 | 14.25 | 8.46 | 8.12 |
| STXBP3 | 14 | 9.17 | 9.91 | 9.87 | 10.28 | 9.97 | 10.78 | 10.23 | 11.30 | 9.25 | 9.25 | 9.54 | 10.07 | 10.37 | 10.76 | 9.25 |
| STXBP4 | 7 | 11.32 | 12.43 | 12.72 | 12.24 | 7.63 | 7.66 | 9.05 | 7.74 | 9.81 | 9.40 | 11.87 | 12.48 | 7.65 | 8.39 | 9.60 |
| STXBP5 | 7 | 7.95 | 8.55 | 11.15 | 9.65 | 9.81 | 9.37 | 14.76 | 13.32 | 7.66 | 7.78 | 8.25 | 10.40 | 9.59 | 14.04 | 7.72 |
| SUB1 | 37 | 9.06 | 11.16 | 8.75 | 8.74 | 9.95 | 11.05 | 11.56 | 11.26 | 9.22 | 9.23 | 10.11 | 8.75 | 10.50 | 11.41 | 9.22 |
| SUCLA2 | 22 | 11.82 | 10.04 | 9.16 | 11.13 | 12.23 | 11.70 | 9.93 | 9.75 | 6.90 | 7.35 | 10.93 | 10.14 | 11.96 | 9.84 | 7.13 |
| SUCLG1 | 10 | 11.64 | 9.63 | 9.85 | 8.41 | 12.44 | 10.93 | 10.51 | 7.82 | 9.43 | 9.34 | 10.64 | 9.13 | 11.69 | 9.16 | 9.38 |
| SUCLG2 | 32 | 10.91 | 9.74 | 8.04 | 9.46 | 11.19 | 10.84 | 9.75 | 9.35 | 10.26 | 10.47 | 10.32 | 8.75 | 11.01 | 9.55 | 10.36 |
| SUDS3 | 3 | 8.87 | 10.06 | 9.71 | 9.98 | 10.79 | 11.23 | 11.27 | 10.59 | 8.36 | 9.12 | 9.47 | 9.85 | 11.01 | 10.93 | 8.74 |
| SUFU | 2 | 9.96 | 10.76 | 9.59 | 9.61 | 10.71 | 11.02 | 11.82 | 10.12 | 8.30 | 8.10 | 10.36 | 9.60 | 10.87 | 10.97 | 8.20 |
| SUGP1 | 15 | 9.67 | 10.66 | 9.93 | 9.86 | 10.27 | 10.66 | 10.41 | 9.78 | 9.54 | 9.21 | 10.16 | 9.90 | 10.47 | 10.10 | 9.37 |
| SUGP2 | 14 | 9.69 | 11.08 | 14.74 | 11.62 | 8.95 | 9.33 | 11.20 | 9.82 | 6.79 | 6.79 | 10.38 | 13.18 | 9.14 | 10.51 | 6.79 |
| SUGT1 | 38 | 10.64 | 12.23 | 10.29 | 10.24 | 11.06 | 11.65 | 8.43 | 8.28 | 8.93 | 8.25 | 11.44 | 10.27 | 11.36 | 8.35 | 8.59 |
| SULT1A3 | 3 | 10.76 | 10.94 | 4.85 | 3.63 | 21.45 | 20.62 | 2.92 | 2.31 | 12.05 | 10.47 | 10.85 | 4.24 | 21.03 | 2.62 | 11.26 |
| SUMF1 | 2 | 7.49 | 9.94 | 9.82 | 10.68 | 14.84 | 14.11 | 6.98 | 6.06 | 9.11 | 10.96 | 8.72 | 10.25 | 14.48 | 6.52 | 10.04 |
| SUMF2 | 11 | 11.10 | 11.09 | 10.09 | 10.07 | 7.77 | 6.91 | 11.00 | 10.72 | 10.95 | 10.30 | 11.09 | 10.08 | 7.34 | 10.86 | 10.63 |
| SUMO1 | 10 | 9.33 | 9.74 | 11.40 | 10.39 | 10.45 | 10.42 | 11.60 | 8.73 | 9.14 | 8.80 | 9.54 | 10.89 | 10.44 | 10.16 | 8.97 |
| SUMO2 | 2 | 10.86 | 11.35 | 11.48 | 9.51 | 9.76 | 9.20 | 11.43 | 7.44 | 9.95 | 9.02 | 11.11 | 10.49 | 9.48 | 9.43 | 9.49 |
| SUMO3 | 8 | 9.40 | 10.99 | 10.85 | 9.09 | 10.91 | 10.24 | 11.43 | 7.78 | 9.69 | 9.62 | 10.19 | 9.97 | 10.57 | 9.61 | 9.66 |
| SUN1 | 8 | 12.12 | 6.37 | 7.15 | 11.33 | 7.64 | 7.25 | 9.14 | 8.46 | 13.05 | 17.49 | 9.25 | 9.24 | 7.45 | 8.80 | 15.27 |
| SUN2 | 2 | 11.15 | 5.78 | 11.94 | 19.90 | 6.08 | 5.54 | 6.14 | 5.52 | 11.44 | 16.52 | 8.46 | 15.92 | 5.81 | 5.83 | 13.98 |
| SUOX | 1 | 9.34 | 7.44 | 11.49 | 11.23 | 10.77 | 10.72 | 8.19 | 10.83 | 11.24 | 8.73 | 8.39 | 11.36 | 10.75 | 9.51 | 9.99 |
| SUPT16H | 69 | 9.38 | 9.92 | 10.02 | 9.47 | 9.82 | 9.61 | 9.85 | 11.02 | 10.76 | 10.16 | 9.65 | 9.74 | 9.71 | 10.43 | 10.46 |
| SUPT20H | 4 | 9.68 | 10.43 | 9.39 | 9.10 | 11.04 | 11.68 | 9.18 | 9.79 | 10.03 | 9.66 | 10.06 | 9.25 | 11.36 | 9.48 | 9.85 |
| SUPT5H | 47 | 9.42 | 10.24 | 11.28 | 9.27 | 9.80 | 9.89 | 11.29 | 11.05 | 8.90 | 8.86 | 9.83 | 10.28 | 9.84 | 11.17 | 8.88 |
| SUPT6H | 44 | 10.56 | 11.35 | 11.17 | 10.58 | 8.87 | 9.05 | 9.28 | 9.05 | 10.13 | 9.95 | 10.96 | 10.88 | 8.96 | 9.16 | 10.04 |
| SUPT7L | 3 | 9.37 | 9.88 | 9.67 | 9.08 | 12.03 | 12.53 | 9.35 | 8.42 | 10.45 | 9.21 | 9.63 | 9.37 | 12.28 | 8.89 | 9.83 |
| SUPV3L1 | 10 | 10.09 | 8.29 | 9.81 | 11.70 | 8.83 | 8.05 | 11.89 | 11.42 | 9.75 | 10.17 | 9.19 | 10.75 | 8.44 | 11.65 | 9.96 |
| SURF1 | 3 | 12.09 | 7.83 | 7.89 | 10.59 | 11.57 | 10.75 | 7.06 | 6.19 | 12.49 | 13.54 | 9.96 | 9.24 | 11.16 | 6.63 | 13.01 |
| SURF2 | 4 | 9.60 | 11.72 | 9.08 | 8.61 | 8.04 | 9.07 | 9.85 | 10.23 | 12.14 | 11.67 | 10.66 | 8.84 | 8.55 | 10.04 | 11.91 |
| SURF4 | 7 | 11.09 | 8.26 | 7.70 | 7.72 | 11.79 | 9.59 | 10.20 | 7.39 | 11.18 | 15.08 | 9.68 | 7.71 | 10.69 | 8.80 | 13.13 |
| SURF6 | 12 | 9.27 | 10.16 | 9.57 | 9.55 | 8.41 | 8.53 | 12.23 | 14.03 | 9.37 | 8.88 | 9.71 | 9.56 | 8.47 | 13.13 | 9.13 |
| SUV39H1 | 4 | 6.58 | 8.02 | 7.17 | 6.01 | 9.03 | 10.00 | 10.01 | 30.53 | 6.64 | 6.02 | 7.30 | 6.59 | 9.51 | 20.27 | 6.33 |
| SUV39H2 | 1 | 8.50 | 9.26 | 12.00 | 11.20 | 7.57 | 8.16 | 9.29 | 8.36 | 13.23 | 12.44 | 8.88 | 11.60 | 7.86 | 8.83 | 12.83 |

| | | | | | | | | | | | | | | | | |
|---|---|---|---|---|---|---|---|---|---|---|---|---|---|---|---|---|
| SUZ12 | 10 | 10.55 | 12.05 | 9.09 | 9.62 | 8.06 | 8.50 | 10.89 | 10.47 | 10.67 | 10.09 | 11.30 | 9.35 | 8.28 | 10.68 | 10.38 |
| SVIL | 18 | 11.84 | 12.60 | 14.17 | 13.29 | 7.37 | 7.60 | 7.98 | 7.14 | 9.28 | 8.74 | 12.22 | 13.73 | 7.48 | 7.56 | 9.01 |
| SVIP | 1 | 11.30 | 10.19 | 11.68 | 13.63 | 8.51 | 8.06 | 8.77 | 8.16 | 10.07 | 9.63 | 10.74 | 12.65 | 8.29 | 8.47 | 9.85 |
| SWAP70 | 27 | 9.14 | 10.36 | 11.66 | 10.79 | 11.25 | 11.38 | 8.24 | 7.44 | 10.08 | 9.64 | 9.75 | 11.23 | 11.32 | 7.84 | 9.86 |
| SYAP1 | 19 | 9.50 | 10.89 | 9.61 | 9.52 | 11.49 | 12.55 | 10.58 | 8.89 | 8.49 | 8.48 | 10.19 | 9.57 | 12.02 | 9.73 | 8.49 |
| SYDE1 | 1 | 8.95 | 9.43 | 11.76 | 10.34 | 9.61 | 10.07 | 8.47 | 11.09 | 10.82 | 9.46 | 9.19 | 11.05 | 9.84 | 9.78 | 10.14 |
| SYF2 | 1 | 9.87 | 11.75 | 10.68 | 8.92 | 9.32 | 9.16 | 10.92 | 9.54 | 10.17 | 9.66 | 10.81 | 9.80 | 9.24 | 10.23 | 9.92 |
| SYMPK | 23 | 9.62 | 9.69 | 8.86 | 8.92 | 11.55 | 11.39 | 10.69 | 10.09 | 9.86 | 9.34 | 9.66 | 8.89 | 11.47 | 10.39 | 9.60 |
| SYNC | 4 | 8.92 | 10.59 | 12.53 | 8.20 | 8.63 | 6.71 | 10.65 | 8.01 | 13.24 | 12.49 | 9.76 | 10.37 | 7.67 | 9.33 | 12.87 |
| SYNCRIP | 42 | 9.49 | 9.58 | 10.95 | 8.73 | 9.67 | 9.43 | 11.57 | 11.12 | 9.84 | 9.62 | 9.54 | 9.84 | 9.55 | 11.35 | 9.73 |
| SYNE1 | 6 | 12.19 | 9.58 | 9.52 | 10.23 | 11.61 | 10.83 | 5.86 | 6.38 | 12.01 | 11.78 | 10.88 | 9.88 | 11.22 | 6.12 | 11.90 |
| SYNE2 | 3 | 16.17 | 7.60 | 6.41 | 7.30 | 10.23 | 9.20 | 4.48 | 3.84 | 15.39 | 19.38 | 11.89 | 6.85 | 9.72 | 4.16 | 17.38 |
| SYNGR2 | 1 | 12.61 | 11.51 | 9.04 | 9.12 | 10.77 | 9.82 | 5.45 | 4.97 | 12.18 | 14.54 | 12.06 | 9.08 | 10.29 | 5.21 | 13.36 |
| SYNJ1 | 3 | 11.13 | 11.42 | 8.32 | 8.13 | 12.21 | 11.75 | 7.73 | 6.95 | 11.79 | 10.57 | 11.27 | 8.22 | 11.98 | 7.34 | 11.18 |
| SYNJ2 | 13 | 9.85 | 10.28 | 10.68 | 10.89 | 9.23 | 9.44 | 10.83 | 9.74 | 9.60 | 9.45 | 10.07 | 10.79 | 9.33 | 10.29 | 9.52 |
| SYNJ2BP | 8 | 12.97 | 7.67 | 8.21 | 10.68 | 9.78 | 8.65 | 9.54 | 8.45 | 11.19 | 12.87 | 10.32 | 9.44 | 9.22 | 9.00 | 12.03 |
| SYNM | 22 | 10.46 | 12.03 | 8.12 | 7.84 | 14.11 | 14.55 | 6.64 | 5.97 | 10.15 | 10.13 | 11.24 | 7.98 | 14.33 | 6.30 | 10.14 |
| SYNPO | 11 | 10.48 | 13.31 | 10.71 | 10.44 | 12.88 | 13.72 | 6.22 | 5.50 | 8.50 | 8.25 | 11.89 | 10.57 | 13.30 | 5.86 | 8.38 |
| SYNPO2 | 3 | 20.42 | 25.54 | 7.78 | 7.26 | 5.23 | 5.31 | 4.36 | 3.30 | 9.75 | 11.05 | 22.98 | 7.52 | 5.27 | 3.83 | 10.40 |
| SYNRG | 8 | 9.82 | 11.32 | 10.80 | 10.38 | 10.33 | 10.46 | 8.11 | 8.26 | 10.50 | 10.01 | 10.57 | 10.59 | 10.40 | 8.19 | 10.26 |
| SYPL1 | 3 | 9.81 | 8.77 | 9.51 | 8.54 | 9.15 | 7.73 | 12.13 | 12.76 | 11.37 | 10.23 | 9.29 | 9.03 | 8.44 | 12.45 | 10.80 |
| SYTL4 | 2 | 9.44 | 9.75 | 9.96 | 9.38 | 12.83 | 13.05 | 11.02 | 10.82 | 7.08 | 6.67 | 9.60 | 9.67 | 12.94 | 10.92 | 6.87 |
| SYVN1 | 1 | 11.41 | 7.89 | 8.63 | 11.98 | 9.66 | 9.93 | 9.34 | 10.15 | 9.45 | 11.56 | 9.65 | 10.30 | 9.79 | 9.75 | 10.50 |
| SZRD1 | 10 | 10.46 | 11.75 | 10.74 | 10.20 | 8.43 | 8.57 | 8.77 | 7.87 | 11.77 | 11.45 | 11.10 | 10.47 | 8.50 | 8.32 | 11.61 |
| TAB1 | 9 | 9.52 | 10.75 | 11.31 | 10.89 | 9.76 | 9.35 | 10.03 | 9.00 | 9.43 | 9.97 | 10.13 | 11.10 | 9.55 | 9.51 | 9.70 |
| TAB2 | 4 | 8.77 | 9.99 | 11.53 | 10.25 | 9.02 | 8.69 | 9.63 | 13.06 | 10.42 | 8.63 | 9.38 | 10.89 | 8.86 | 11.35 | 9.53 |
| TAB3 | 1 | 10.11 | 11.28 | 7.37 | 13.23 | 8.48 | 9.65 | 8.68 | 11.71 | 9.71 | 9.79 | 10.69 | 10.30 | 9.06 | 10.19 | 9.75 |
| TACC1 | 15 | 9.62 | 9.97 | 11.23 | 10.94 | 8.47 | 8.65 | 11.49 | 10.63 | 10.55 | 8.43 | 9.80 | 11.09 | 8.56 | 11.06 | 9.49 |
| TACC2 | 11 | 9.64 | 10.03 | 11.85 | 11.60 | 8.99 | 9.39 | 9.88 | 9.06 | 9.95 | 9.61 | 9.83 | 11.73 | 9.19 | 9.47 | 9.78 |
| TACC3 | 16 | 7.98 | 9.61 | 11.31 | 10.73 | 10.00 | 10.04 | 8.52 | 7.86 | 12.23 | 11.72 | 8.79 | 11.02 | 10.02 | 8.19 | 11.97 |
| TACO1 | 6 | 11.16 | 9.49 | 11.74 | 12.41 | 9.26 | 8.35 | 8.39 | 7.01 | 11.23 | 10.96 | 10.32 | 12.07 | 8.81 | 7.70 | 11.10 |
| TACSTD2 | 3 | 13.31 | 12.38 | 7.10 | 8.10 | 8.34 | 10.19 | 6.19 | 4.11 | 15.47 | 14.81 | 12.84 | 7.60 | 9.27 | 5.15 | 15.14 |
| TADA1 | 1 | 8.63 | 10.69 | 8.81 | 8.12 | 12.45 | 11.96 | 10.36 | 8.46 | 9.93 | 10.59 | 9.66 | 8.46 | 12.21 | 9.41 | 10.26 |
| TADA2A | 1 | 10.98 | 11.44 | 10.77 | 8.82 | 8.72 | 7.46 | 10.54 | 8.71 | 11.38 | 11.17 | 11.21 | 9.79 | 8.09 | 9.62 | 11.28 |
| TADA2B | 1 | 8.89 | 10.88 | 7.60 | 10.09 | 9.66 | 12.29 | 10.40 | 9.03 | 9.84 | 11.32 | 9.89 | 8.84 | 10.98 | 9.72 | 10.58 |
| TADA3 | 2 | 9.98 | 9.94 | 13.82 | 12.84 | 8.61 | 8.88 | 7.89 | 7.07 | 10.62 | 10.35 | 9.96 | 13.33 | 8.75 | 7.48 | 10.49 |
| TAF1 | 2 | 10.62 | 9.98 | 10.22 | 9.88 | 8.87 | 9.95 | 11.97 | 10.97 | 9.29 | 8.24 | 10.30 | 10.05 | 9.41 | 11.47 | 8.77 |
| TAF10 | 3 | 8.73 | 10.04 | 12.24 | 11.34 | 10.18 | 10.69 | 9.87 | 8.90 | 8.81 | 9.18 | 9.39 | 11.79 | 10.44 | 9.39 | 9.00 |
| TAF11 | 2 | 11.10 | 10.98 | 9.87 | 9.44 | 8.86 | 10.22 | 10.09 | 8.52 | 10.22 | 10.70 | 11.04 | 9.65 | 9.54 | 9.31 | 10.46 |
| TAF12 | 3 | 9.13 | 11.65 | 8.45 | 8.61 | 9.17 | 10.45 | 10.76 | 10.80 | 10.95 | 10.03 | 10.39 | 8.53 | 9.81 | 10.78 | 10.49 |
| TAF15 | 23 | 12.37 | 12.71 | 15.31 | 12.87 | 12.44 | 11.04 | 1.04 | 0.67 | 10.06 | 11.50 | 12.54 | 14.09 | 11.74 | 0.86 | 10.78 |
| TAF3 | 2 | 9.49 | 10.66 | 9.76 | 9.09 | 8.80 | 9.28 | 10.88 | 10.26 | 10.81 | 10.97 | 10.07 | 9.43 | 9.04 | 10.57 | 10.89 |

| | | | | | | | | | | | | | | | | |
|---|---|---|---|---|---|---|---|---|---|---|---|---|---|---|---|---|
| TAF4 | 8 | 8.84 | 10.34 | 10.24 | 10.96 | 9.09 | 8.99 | 11.34 | 10.83 | 9.84 | 9.52 | 9.59 | 10.60 | 9.04 | 11.09 | 9.68 |
| TAF5 | 4 | 8.73 | 9.41 | 11.30 | 10.50 | 8.40 | 9.18 | 11.40 | 10.42 | 10.39 | 10.28 | 9.07 | 10.90 | 8.79 | 10.91 | 10.33 |
| TAF6 | 8 | 9.87 | 9.87 | 11.71 | 10.09 | 8.58 | 8.44 | 11.67 | 9.61 | 10.17 | 9.99 | 9.87 | 10.90 | 8.51 | 10.64 | 10.08 |
| TAF6L | 2 | 9.78 | 10.14 | 8.80 | 9.50 | 12.32 | 11.72 | 8.98 | 8.68 | 9.61 | 10.49 | 9.96 | 9.15 | 12.02 | 8.83 | 10.05 |
| TAF7 | 5 | 10.17 | 10.85 | 9.07 | 9.05 | 10.82 | 11.51 | 10.43 | 8.25 | 10.03 | 9.81 | 10.51 | 9.06 | 11.16 | 9.34 | 9.92 |
| TAF9B | 8 | 8.91 | 10.02 | 10.08 | 9.41 | 10.05 | 10.30 | 11.66 | 10.67 | 9.89 | 9.00 | 9.46 | 9.75 | 10.18 | 11.17 | 9.45 |
| TAGLN | 86 | 10.54 | 11.59 | 19.53 | 18.98 | 3.72 | 3.70 | 1.70 | 1.52 | 14.42 | 14.30 | 11.06 | 19.26 | 3.71 | 1.61 | 14.36 |
| TAGLN2 | 89 | 7.82 | 10.34 | 10.42 | 10.56 | 11.81 | 12.12 | 9.79 | 9.88 | 8.74 | 8.53 | 9.08 | 10.49 | 11.97 | 9.83 | 8.63 |
| TALDO1 | 49 | 10.36 | 11.89 | 8.82 | 8.26 | 12.91 | 13.24 | 8.26 | 8.06 | 9.24 | 8.94 | 11.13 | 8.54 | 13.08 | 8.16 | 9.09 |
| TAMM41 | 5 | 10.01 | 8.71 | 8.87 | 10.77 | 14.07 | 13.08 | 11.35 | 10.78 | 5.84 | 6.52 | 9.36 | 9.82 | 13.57 | 11.07 | 6.18 |
| TANC1 | 10 | 9.61 | 10.74 | 11.66 | 12.21 | 9.71 | 9.77 | 6.68 | 6.43 | 11.64 | 11.55 | 10.18 | 11.93 | 9.74 | 6.56 | 11.60 |
| TANC2 | 4 | 9.96 | 10.43 | 14.76 | 14.04 | 9.55 | 9.67 | 9.81 | 9.93 | 6.03 | 5.82 | 10.20 | 14.40 | 9.61 | 9.87 | 5.92 |
| TANGO6 | 3 | 11.30 | 11.07 | 9.64 | 9.57 | 8.51 | 9.74 | 9.93 | 10.08 | 10.28 | 9.88 | 11.18 | 9.60 | 9.13 | 10.01 | 10.08 |
| TANK | 3 | 10.59 | 11.56 | 10.47 | 13.03 | 9.01 | 9.63 | 10.22 | 9.99 | 8.04 | 7.45 | 11.07 | 11.75 | 9.32 | 10.11 | 7.75 |
| TAOK1 | 8 | 9.81 | 10.82 | 12.40 | 11.82 | 8.55 | 9.08 | 8.70 | 8.47 | 10.43 | 9.92 | 10.32 | 12.11 | 8.82 | 8.58 | 10.17 |
| TAOK2 | 4 | 9.58 | 10.74 | 9.69 | 10.41 | 10.03 | 9.47 | 9.85 | 8.01 | 10.85 | 11.37 | 10.16 | 10.05 | 9.75 | 8.93 | 11.11 |
| TAOK3 | 3 | 9.66 | 10.23 | 10.51 | 11.02 | 9.23 | 10.57 | 12.31 | 10.45 | 8.41 | 7.61 | 9.94 | 10.77 | 9.90 | 11.38 | 8.01 |
| TAP1 | 1 | 8.66 | 9.15 | 13.79 | 6.48 | 15.26 | 10.55 | 10.63 | 2.97 | 11.76 | 10.77 | 8.90 | 10.13 | 12.90 | 6.80 | 11.26 |
| TAP2 | 1 | 15.88 | 8.18 | 4.86 | 7.01 | 13.50 | 9.85 | 7.49 | 5.86 | 11.67 | 15.70 | 12.03 | 5.93 | 11.68 | 6.67 | 13.68 |
| TAPBP | 1 | 13.57 | 7.88 | 4.93 | 8.48 | 13.96 | 12.64 | 7.77 | 6.92 | 11.12 | 12.73 | 10.72 | 6.71 | 13.30 | 7.34 | 11.93 |
| TARBP1 | 6 | 8.56 | 9.37 | 6.93 | 6.60 | 10.23 | 9.62 | 13.13 | 12.89 | 10.76 | 11.93 | 8.96 | 6.76 | 9.93 | 13.01 | 11.34 |
| TARBP2 | 3 | 8.46 | 9.15 | 11.59 | 10.69 | 9.74 | 10.48 | 11.99 | 9.32 | 9.43 | 9.15 | 8.81 | 11.14 | 10.11 | 10.65 | 9.29 |
| TARDBP | 11 | 10.19 | 10.10 | 10.00 | 9.27 | 10.43 | 10.20 | 11.66 | 9.83 | 9.61 | 8.70 | 10.14 | 9.64 | 10.31 | 10.75 | 9.16 |
| TARS | 119 | 8.69 | 9.61 | 9.08 | 8.26 | 10.55 | 10.48 | 10.07 | 12.65 | 10.43 | 10.17 | 9.15 | 8.67 | 10.52 | 11.36 | 10.30 |
| TARS2 | 15 | 11.35 | 9.56 | 8.94 | 10.27 | 11.72 | 11.04 | 9.98 | 9.48 | 8.71 | 8.97 | 10.45 | 9.60 | 11.38 | 9.73 | 8.84 |
| TATDN1 | 13 | 9.47 | 10.34 | 8.95 | 8.61 | 8.12 | 8.54 | 12.53 | 15.60 | 9.06 | 8.78 | 9.91 | 8.78 | 8.33 | 14.06 | 8.92 |
| TATDN2 | 2 | 8.68 | 10.45 | 10.13 | 9.39 | 10.32 | 12.16 | 10.92 | 9.19 | 9.48 | 9.28 | 9.57 | 9.76 | 11.24 | 10.06 | 9.38 |
| TATDN3 | 1 | 9.91 | 8.25 | 8.79 | 6.53 | 12.19 | 11.45 | 11.94 | 7.42 | 12.48 | 11.05 | 9.08 | 7.66 | 11.82 | 9.68 | 11.76 |
| TAX1BP1 | 8 | 11.60 | 11.68 | 10.32 | 9.87 | 8.51 | 8.42 | 9.43 | 9.18 | 11.28 | 9.71 | 11.64 | 10.10 | 8.46 | 9.31 | 10.49 |
| TAX1BP3 | 3 | 10.10 | 11.77 | 14.51 | 14.08 | 10.77 | 10.28 | 5.58 | 5.57 | 8.53 | 8.82 | 10.93 | 14.30 | 10.52 | 5.57 | 8.67 |
| TBC1D10B | 6 | 9.32 | 10.91 | 10.79 | 11.69 | 10.32 | 11.53 | 7.55 | 6.87 | 10.24 | 10.79 | 10.12 | 11.24 | 10.92 | 7.21 | 10.51 |
| TBC1D13 | 8 | 10.20 | 10.33 | 10.63 | 9.71 | 11.04 | 11.05 | 9.76 | 8.91 | 9.78 | 8.60 | 10.27 | 10.17 | 11.04 | 9.34 | 9.19 |
| TBC1D15 | 13 | 9.87 | 9.91 | 9.39 | 7.60 | 11.46 | 11.14 | 11.66 | 10.10 | 9.61 | 9.26 | 9.89 | 8.50 | 11.30 | 10.88 | 9.43 |
| TBC1D17 | 5 | 10.92 | 11.28 | 7.30 | 6.74 | 13.62 | 14.11 | 9.23 | 8.48 | 9.73 | 8.60 | 11.10 | 7.02 | 13.87 | 8.85 | 9.16 |
| TBC1D2 | 14 | 9.01 | 9.69 | 6.66 | 6.53 | 10.65 | 10.68 | 9.23 | 8.57 | 14.79 | 14.19 | 9.35 | 6.59 | 10.67 | 8.90 | 14.49 |
| TBC1D22A | 1 | 10.16 | 10.51 | 10.68 | 10.86 | 8.87 | 10.01 | 10.70 | 10.63 | 9.21 | 8.36 | 10.34 | 10.77 | 9.44 | 10.67 | 8.78 |
| TBC1D22B | 2 | 9.90 | 11.12 | 9.65 | 9.23 | 10.29 | 11.16 | 8.19 | 8.29 | 10.78 | 11.39 | 10.51 | 9.44 | 10.72 | 8.24 | 11.09 |
| TBC1D23 | 8 | 8.75 | 9.59 | 13.14 | 13.28 | 10.16 | 10.48 | 8.61 | 8.44 | 8.65 | 8.90 | 9.17 | 13.21 | 10.32 | 8.53 | 8.78 |
| TBC1D24 | 3 | 10.89 | 11.58 | 8.62 | 10.18 | 8.88 | 8.78 | 9.43 | 9.48 | 10.14 | 12.03 | 11.23 | 9.40 | 8.83 | 9.45 | 11.08 |
| TBC1D25 | 4 | 8.31 | 9.90 | 10.31 | 10.20 | 10.42 | 10.75 | 11.31 | 10.39 | 9.20 | 9.21 | 9.10 | 10.26 | 10.58 | 10.85 | 9.20 |
| TBC1D2B | 2 | 9.61 | 9.52 | 8.69 | 9.87 | 9.17 | 10.31 | 10.40 | 10.68 | 11.50 | 10.24 | 9.56 | 9.28 | 9.74 | 10.54 | 10.87 |
| TBC1D4 | 15 | 8.36 | 9.05 | 14.42 | 13.51 | 7.09 | 7.28 | 12.05 | 11.24 | 8.52 | 8.48 | 8.70 | 13.97 | 7.19 | 11.64 | 8.50 |

| | | | | | | | | | | | | | | | | |
|---|---|---|---|---|---|---|---|---|---|---|---|---|---|---|---|---|
| TBC1D5 | 8 | 10.23 | 10.81 | 9.75 | 8.62 | 5.86 | 5.88 | 12.77 | 11.80 | 12.43 | 11.86 | 10.52 | 9.18 | 5.87 | 12.28 | 12.15 |
| TBC1D8B | 3 | 10.66 | 11.79 | 14.13 | 13.18 | 8.78 | 8.23 | 7.38 | 4.26 | 10.98 | 10.60 | 11.23 | 13.65 | 8.50 | 5.82 | 10.79 |
| TBC1D9B | 15 | 9.12 | 9.51 | 8.15 | 7.84 | 12.26 | 11.98 | 12.11 | 11.66 | 8.82 | 8.55 | 9.31 | 8.00 | 12.12 | 11.89 | 8.69 |
| TBCA | 19 | 10.07 | 10.94 | 8.82 | 8.24 | 10.85 | 10.67 | 11.99 | 11.02 | 9.17 | 8.23 | 10.50 | 8.53 | 10.76 | 11.51 | 8.70 |
| TBCB | 14 | 9.42 | 10.56 | 10.34 | 11.10 | 9.43 | 10.15 | 9.29 | 10.70 | 9.17 | 9.82 | 9.99 | 10.72 | 9.79 | 10.00 | 9.50 |
| TBCC | 7 | 8.72 | 11.32 | 10.60 | 11.09 | 10.36 | 11.40 | 8.73 | 8.09 | 9.84 | 9.84 | 10.02 | 10.84 | 10.88 | 8.41 | 9.84 |
| TBCD | 14 | 11.18 | 11.79 | 9.24 | 10.05 | 9.46 | 9.77 | 8.19 | 8.50 | 11.00 | 10.82 | 11.48 | 9.65 | 9.62 | 8.35 | 10.91 |
| TBCE | 9 | 11.25 | 12.50 | 8.70 | 8.64 | 11.47 | 11.81 | 8.51 | 8.70 | 9.32 | 9.10 | 11.88 | 8.67 | 11.64 | 8.60 | 9.21 |
| TBK1 | 13 | 9.12 | 9.73 | 9.74 | 9.61 | 11.41 | 11.20 | 9.87 | 9.51 | 9.82 | 9.99 | 9.43 | 9.68 | 11.30 | 9.69 | 9.91 |
| TBL1X | 6 | 9.87 | 9.96 | 11.32 | 10.91 | 9.94 | 10.10 | 10.65 | 9.89 | 8.76 | 8.61 | 9.91 | 11.11 | 10.02 | 10.27 | 8.68 |
| TBL1XR1 | 5 | 9.63 | 10.58 | 11.27 | 10.35 | 9.26 | 9.46 | 11.42 | 10.25 | 8.93 | 8.86 | 10.10 | 10.81 | 9.36 | 10.83 | 8.89 |
| TBL2 | 6 | 10.70 | 6.91 | 9.25 | 12.09 | 7.78 | 6.22 | 11.22 | 11.39 | 11.06 | 13.39 | 8.80 | 10.67 | 7.00 | 11.31 | 12.22 |
| TBL3 | 16 | 9.36 | 9.59 | 10.91 | 10.12 | 8.47 | 8.44 | 10.75 | 9.81 | 11.42 | 11.14 | 9.47 | 10.51 | 8.46 | 10.28 | 11.28 |
| TBPL2 | 1 | 10.24 | 11.47 | 7.64 | 7.36 | 9.52 | 11.39 | 9.93 | 12.10 | 10.40 | 9.95 | 10.86 | 7.50 | 10.45 | 11.02 | 10.18 |
| TBRG4 | 11 | 10.80 | 8.47 | 10.02 | 10.84 | 7.78 | 6.63 | 11.99 | 10.90 | 11.00 | 11.57 | 9.64 | 10.43 | 7.21 | 11.44 | 11.28 |
| TCEA1 | 18 | 8.38 | 10.04 | 10.55 | 10.04 | 8.75 | 8.66 | 12.34 | 15.09 | 8.35 | 7.79 | 9.21 | 10.30 | 8.71 | 13.72 | 8.07 |
| TCEAL1 | 2 | 8.99 | 9.86 | 7.26 | 6.65 | 18.05 | 20.20 | 9.57 | 5.96 | 7.16 | 6.32 | 9.43 | 6.95 | 19.12 | 7.76 | 6.74 |
| TCEAL4 | 12 | 9.54 | 10.91 | 8.14 | 7.16 | 16.06 | 15.68 | 10.68 | 6.41 | 8.01 | 7.40 | 10.23 | 7.65 | 15.87 | 8.55 | 7.70 |
| TCEAL5 | 2 | 12.25 | 14.68 | 5.49 | 10.00 | 13.80 | 19.93 | 6.24 | 2.61 | 7.46 | 7.54 | 13.46 | 7.75 | 16.87 | 4.43 | 7.50 |
| TCEANC2 | 1 | 10.84 | 12.43 | 8.14 | 9.28 | 8.66 | 10.13 | 11.50 | 11.06 | 8.84 | 9.12 | 11.64 | 8.71 | 9.39 | 11.28 | 8.98 |
| TCEB1 | 6 | 10.25 | 9.30 | 10.12 | 8.09 | 11.14 | 11.81 | 10.42 | 7.03 | 10.69 | 11.15 | 9.78 | 9.10 | 11.47 | 8.73 | 10.92 |
| TCEB2 | 13 | 9.63 | 10.75 | 10.04 | 9.84 | 10.47 | 11.07 | 10.45 | 8.74 | 9.66 | 9.35 | 10.19 | 9.94 | 10.77 | 9.59 | 9.51 |
| TCEB3 | 13 | 11.14 | 11.57 | 8.75 | 9.82 | 8.16 | 9.15 | 9.66 | 9.36 | 9.85 | 12.54 | 11.36 | 9.28 | 8.65 | 9.51 | 11.19 |
| TCERG1 | 41 | 9.05 | 9.98 | 9.03 | 8.35 | 10.33 | 10.71 | 12.67 | 12.18 | 9.08 | 8.62 | 9.52 | 8.69 | 10.52 | 12.43 | 8.85 |
| TCF12 | 8 | 10.17 | 11.41 | 9.06 | 9.92 | 9.14 | 10.17 | 9.08 | 8.89 | 11.36 | 10.80 | 10.79 | 9.49 | 9.66 | 8.98 | 11.08 |
| TCF19 | 1 | 10.42 | 10.21 | 9.20 | 10.81 | 9.29 | 8.72 | 6.24 | 5.43 | 11.90 | 17.78 | 10.32 | 10.01 | 9.01 | 5.83 | 14.84 |
| TCF20 | 10 | 8.78 | 10.36 | 13.06 | 12.13 | 7.59 | 7.89 | 11.48 | 10.89 | 9.19 | 8.63 | 9.57 | 12.60 | 7.74 | 11.19 | 8.91 |
| TCF25 | 6 | 11.53 | 12.18 | 8.51 | 8.02 | 10.78 | 11.13 | 9.36 | 8.54 | 9.72 | 10.24 | 11.85 | 8.26 | 10.96 | 8.95 | 9.98 |
| TCF3 | 3 | 8.79 | 10.54 | 10.35 | 10.26 | 9.53 | 9.66 | 12.19 | 11.02 | 8.99 | 8.66 | 9.67 | 10.31 | 9.60 | 11.61 | 8.82 |
| TCHP | 1 | 11.94 | 9.77 | 10.41 | 7.00 | 12.73 | 12.24 | 9.79 | 8.69 | 10.03 | 7.42 | 10.85 | 8.70 | 12.48 | 9.24 | 8.72 |
| TCIRG1 | 3 | 11.80 | 7.54 | 5.23 | 6.91 | 14.65 | 12.80 | 7.23 | 7.24 | 12.11 | 14.49 | 9.67 | 6.07 | 13.72 | 7.24 | 13.30 |
| TCOF1 | 90 | 8.84 | 9.87 | 10.55 | 9.91 | 7.44 | 7.64 | 14.65 | 13.37 | 9.09 | 8.64 | 9.36 | 10.23 | 7.54 | 14.01 | 8.87 |
| TCP1 | 60 | 9.43 | 10.34 | 9.09 | 8.36 | 10.00 | 9.89 | 9.02 | 15.12 | 9.72 | 9.04 | 9.88 | 8.72 | 9.95 | 12.07 | 9.38 |
| TCP11L1 | 9 | 11.03 | 12.45 | 11.84 | 11.06 | 12.16 | 12.33 | 4.97 | 4.54 | 10.16 | 9.46 | 11.74 | 11.45 | 12.24 | 4.76 | 9.81 |
| TDG | 2 | 10.25 | 9.60 | 10.70 | 8.03 | 9.22 | 8.00 | 11.44 | 7.89 | 12.07 | 12.81 | 9.93 | 9.36 | 8.61 | 9.66 | 12.44 |
| TDP1 | 4 | 10.40 | 9.75 | 9.09 | 8.44 | 10.96 | 10.87 | 11.13 | 11.29 | 9.19 | 8.88 | 10.08 | 8.76 | 10.92 | 11.21 | 9.04 |
| TDP2 | 4 | 10.68 | 11.17 | 10.55 | 10.87 | 9.25 | 9.28 | 10.36 | 9.29 | 9.38 | 9.17 | 10.92 | 10.71 | 9.27 | 9.83 | 9.28 |
| TDRD7 | 2 | 12.68 | 13.01 | 9.79 | 10.14 | 12.96 | 12.88 | 5.95 | 5.79 | 8.98 | 7.82 | 12.84 | 9.97 | 12.92 | 5.87 | 8.40 |
| TEAD1 | 4 | 11.08 | 10.56 | 10.24 | 8.87 | 10.21 | 11.01 | 10.39 | 9.57 | 9.22 | 8.86 | 10.82 | 9.55 | 10.61 | 9.98 | 9.04 |
| TECR | 10 | 14.31 | 6.89 | 6.46 | 6.39 | 11.46 | 8.71 | 10.55 | 9.37 | 11.74 | 14.12 | 10.60 | 6.43 | 10.09 | 9.96 | 12.93 |
| TEFM | 2 | 11.66 | 9.96 | 11.81 | 13.78 | 8.06 | 7.80 | 8.08 | 6.43 | 11.33 | 11.08 | 10.81 | 12.80 | 7.93 | 7.26 | 11.21 |
| TELO2 | 14 | 10.88 | 10.39 | 8.92 | 8.85 | 10.14 | 9.99 | 9.09 | 10.47 | 10.71 | 10.56 | 10.64 | 8.89 | 10.06 | 9.78 | 10.63 |

| | | | | | | | | | | | | | | | | |
|---|---|---|---|---|---|---|---|---|---|---|---|---|---|---|---|---|
| TENM1 | 1 | 12.90 | 15.98 | 5.41 | 6.12 | 7.47 | 11.38 | 9.66 | 10.63 | 10.82 | 9.62 | 14.44 | 5.77 | 9.43 | 10.14 | 10.22 |
| TERF1 | 1 | 9.90 | 10.87 | 8.66 | 10.80 | 7.68 | 10.56 | 13.06 | 11.53 | 8.59 | 8.35 | 10.38 | 9.73 | 9.12 | 12.30 | 8.47 |
| TERF2 | 2 | 8.80 | 11.16 | 7.99 | 6.20 | 11.12 | 11.88 | 11.76 | 9.70 | 11.01 | 10.37 | 9.98 | 7.10 | 11.50 | 10.73 | 10.69 |
| TERF2IP | 9 | 11.62 | 12.04 | 7.37 | 6.88 | 10.62 | 10.75 | 9.93 | 9.69 | 10.79 | 10.31 | 11.83 | 7.13 | 10.69 | 9.81 | 10.55 |
| TES | 40 | 8.87 | 9.93 | 14.02 | 13.55 | 6.88 | 7.27 | 8.74 | 9.88 | 10.58 | 10.29 | 9.40 | 13.78 | 7.07 | 9.31 | 10.43 |
| TESC | 1 | 3.78 | 5.11 | 17.83 | 16.19 | 11.30 | 13.82 | 15.63 | 14.63 | 0.98 | 0.73 | 4.44 | 17.01 | 12.56 | 15.13 | 0.86 |
| TET2 | 1 | 12.35 | 10.20 | 10.50 | 10.63 | 8.16 | 7.95 | 8.37 | 9.63 | 11.09 | 11.14 | 11.27 | 10.56 | 8.05 | 9.00 | 11.12 |
| TEX10 | 14 | 9.12 | 9.07 | 9.70 | 10.12 | 9.93 | 9.65 | 9.89 | 10.09 | 10.94 | 11.49 | 9.10 | 9.91 | 9.79 | 9.99 | 11.22 |
| TEX15 | 2 | 10.74 | 8.33 | 8.16 | 9.51 | 11.85 | 11.40 | 10.75 | 9.81 | 9.55 | 9.92 | 9.53 | 8.83 | 11.62 | 10.28 | 9.73 |
| TEX30 | 5 | 9.48 | 10.13 | 9.14 | 8.76 | 10.83 | 10.64 | 11.34 | 11.63 | 8.99 | 9.06 | 9.80 | 8.95 | 10.73 | 11.49 | 9.03 |
| TFAM | 12 | 10.06 | 8.66 | 11.23 | 11.42 | 7.47 | 7.08 | 13.22 | 11.26 | 9.63 | 9.99 | 9.36 | 11.32 | 7.28 | 12.24 | 9.81 |
| TFAP4 | 4 | 8.62 | 9.75 | 9.93 | 9.24 | 8.42 | 9.17 | 14.16 | 12.86 | 8.76 | 9.10 | 9.19 | 9.58 | 8.79 | 13.51 | 8.93 |
| TFB1M | 8 | 10.92 | 9.41 | 10.93 | 12.46 | 8.60 | 8.29 | 8.86 | 8.65 | 10.72 | 11.14 | 10.17 | 11.69 | 8.45 | 8.76 | 10.93 |
| TFB2M | 8 | 10.89 | 8.68 | 8.29 | 7.80 | 10.40 | 8.83 | 12.47 | 10.96 | 10.52 | 11.15 | 9.79 | 8.05 | 9.62 | 11.72 | 10.84 |
| TFCP2 | 5 | 10.19 | 10.33 | 11.80 | 12.45 | 8.17 | 8.05 | 10.82 | 10.73 | 9.03 | 8.43 | 10.26 | 12.13 | 8.11 | 10.77 | 8.73 |
| TFDP1 | 2 | 10.00 | 10.45 | 8.45 | 8.63 | 10.83 | 10.73 | 12.42 | 11.45 | 8.06 | 8.97 | 10.23 | 8.54 | 10.78 | 11.94 | 8.51 |
| TFE3 | 3 | 10.49 | 11.35 | 8.47 | 9.27 | 11.55 | 12.79 | 7.72 | 9.85 | 9.58 | 8.94 | 10.92 | 8.87 | 12.17 | 8.78 | 9.26 |
| TFEB | 3 | 8.42 | 11.85 | 10.57 | 9.66 | 10.69 | 10.76 | 8.95 | 8.72 | 10.66 | 9.73 | 10.14 | 10.12 | 10.72 | 8.83 | 10.19 |
| TFG | 45 | 8.71 | 10.24 | 11.24 | 10.62 | 11.06 | 10.85 | 9.18 | 8.08 | 10.23 | 9.79 | 9.47 | 10.93 | 10.95 | 8.63 | 10.01 |
| TFIP11 | 10 | 9.79 | 9.62 | 9.80 | 9.58 | 9.40 | 9.50 | 11.35 | 10.64 | 10.10 | 10.21 | 9.70 | 9.69 | 9.45 | 11.00 | 10.16 |
| TFPI | 1 | 12.36 | 11.53 | 8.58 | 7.72 | 9.29 | 8.51 | 12.32 | 13.40 | 7.97 | 8.33 | 11.94 | 8.15 | 8.90 | 12.86 | 8.15 |
| TFPT | 1 | 9.10 | 9.48 | 12.31 | 13.97 | 7.48 | 8.29 | 11.32 | 10.35 | 8.51 | 9.19 | 9.29 | 13.14 | 7.89 | 10.83 | 8.85 |
| TFRC | 60 | 10.79 | 7.88 | 9.77 | 12.09 | 9.16 | 8.56 | 9.37 | 9.24 | 11.13 | 12.01 | 9.33 | 10.93 | 8.86 | 9.31 | 11.57 |
| TGFB1 | 1 | 8.42 | 5.92 | 7.54 | 8.73 | 9.51 | 9.10 | 17.08 | 14.56 | 9.99 | 9.14 | 7.17 | 8.14 | 9.31 | 15.82 | 9.56 |
| TGFB1I1 | 7 | 10.26 | 11.58 | 14.55 | 15.02 | 6.68 | 6.63 | 3.50 | 3.29 | 14.31 | 14.18 | 10.92 | 14.78 | 6.65 | 3.40 | 14.25 |
| TGFBI | 14 | 7.36 | 6.61 | 14.77 | 16.76 | 2.61 | 2.54 | 12.76 | 12.72 | 11.36 | 12.51 | 6.98 | 15.76 | 2.57 | 12.74 | 11.93 |
| TGM2 | 37 | 5.61 | 6.02 | 20.54 | 20.48 | 3.33 | 3.26 | 7.75 | 8.19 | 12.52 | 12.30 | 5.81 | 20.51 | 3.30 | 7.97 | 12.41 |
| TGOLN2 | 2 | 11.76 | 9.11 | 8.34 | 11.06 | 8.55 | 8.11 | 12.22 | 8.72 | 10.66 | 11.47 | 10.43 | 9.70 | 8.33 | 10.47 | 11.07 |
| TGS1 | 3 | 8.95 | 9.61 | 10.09 | 6.70 | 8.48 | 8.71 | 12.36 | 14.30 | 10.90 | 9.89 | 9.28 | 8.39 | 8.60 | 13.33 | 10.40 |
| THADA | 3 | 10.34 | 9.20 | 8.07 | 8.74 | 10.67 | 10.22 | 9.26 | 10.29 | 11.06 | 12.14 | 9.77 | 8.40 | 10.45 | 9.78 | 11.60 |
| THAP11 | 2 | 9.34 | 10.46 | 10.81 | 9.27 | 9.64 | 9.27 | 12.14 | 9.34 | 9.77 | 9.95 | 9.90 | 10.04 | 9.45 | 10.74 | 9.86 |
| THBS1 | 18 | 7.33 | 6.08 | 18.73 | 21.39 | 3.81 | 3.76 | 3.59 | 3.15 | 15.83 | 16.34 | 6.70 | 20.06 | 3.78 | 3.37 | 16.08 |
| THEM4 | 2 | 8.95 | 7.33 | 5.13 | 5.23 | 13.24 | 12.77 | 16.36 | 15.69 | 7.51 | 7.79 | 8.14 | 5.18 | 13.00 | 16.03 | 7.65 |
| THEM6 | 3 | 10.60 | 7.45 | 6.22 | 8.24 | 11.82 | 11.02 | 12.09 | 10.92 | 10.52 | 11.13 | 9.02 | 7.23 | 11.42 | 11.50 | 10.82 |
| THG1L | 7 | 8.91 | 8.56 | 11.25 | 10.09 | 7.34 | 6.90 | 10.94 | 17.88 | 9.09 | 9.03 | 8.73 | 10.67 | 7.12 | 14.41 | 9.06 |
| THOC1 | 14 | 9.48 | 10.82 | 10.68 | 10.24 | 11.38 | 11.21 | 9.16 | 8.79 | 9.63 | 8.62 | 10.15 | 10.46 | 11.30 | 8.97 | 9.13 |
| THOC2 | 24 | 9.59 | 10.05 | 9.33 | 9.27 | 11.04 | 10.91 | 10.26 | 11.90 | 8.90 | 8.74 | 9.82 | 9.30 | 10.98 | 11.08 | 8.82 |
| THOC3 | 11 | 10.20 | 11.15 | 9.21 | 8.96 | 10.55 | 11.03 | 10.73 | 11.68 | 8.48 | 8.01 | 10.67 | 9.09 | 10.79 | 11.21 | 8.24 |
| THOC5 | 17 | 10.53 | 11.15 | 10.41 | 10.34 | 10.53 | 11.50 | 9.35 | 8.90 | 8.67 | 8.62 | 10.84 | 10.37 | 11.02 | 9.12 | 8.64 |
| THOC6 | 7 | 9.68 | 9.56 | 11.77 | 8.54 | 10.57 | 9.72 | 10.64 | 8.99 | 10.84 | 9.68 | 9.62 | 10.16 | 10.15 | 9.82 | 10.26 |
| THOC7 | 10 | 10.94 | 10.78 | 10.70 | 9.98 | 11.00 | 10.77 | 9.40 | 7.87 | 10.05 | 8.52 | 10.86 | 10.34 | 10.88 | 8.63 | 9.28 |
| THOP1 | 21 | 10.21 | 9.92 | 10.03 | 8.48 | 10.98 | 10.72 | 10.60 | 11.91 | 9.03 | 8.12 | 10.06 | 9.26 | 10.85 | 11.26 | 8.58 |

| | | | | | | | | | | | | | | | | |
|---|---|---|---|---|---|---|---|---|---|---|---|---|---|---|---|---|
| THRAP3 | 41 | 9.45 | 10.51 | 10.41 | 8.64 | 10.18 | 9.91 | 10.93 | 9.60 | 10.45 | 9.93 | 9.98 | 9.53 | 10.04 | 10.26 | 10.19 |
| THTPA | 2 | 9.72 | 10.76 | 12.30 | 11.91 | 10.73 | 11.18 | 8.40 | 5.48 | 10.13 | 9.39 | 10.24 | 12.11 | 10.95 | 6.94 | 9.76 |
| THUMPD1 | 18 | 10.06 | 9.73 | 10.48 | 10.33 | 9.01 | 9.34 | 9.90 | 12.63 | 9.03 | 9.49 | 9.89 | 10.41 | 9.18 | 11.26 | 9.26 |
| THUMPD2 | 1 | 4.94 | 10.78 | 15.45 | 12.84 | 9.47 | 10.40 | 9.14 | 7.90 | 9.58 | 9.52 | 7.86 | 14.14 | 9.93 | 8.52 | 9.55 |
| THUMPD3 | 13 | 9.86 | 10.39 | 10.56 | 9.32 | 10.44 | 10.02 | 9.35 | 8.80 | 10.85 | 10.40 | 10.12 | 9.94 | 10.23 | 9.08 | 10.63 |
| THY1 | 2 | 9.30 | 6.97 | 14.19 | 12.53 | 1.47 | 1.32 | 6.04 | 5.03 | 20.04 | 23.10 | 8.14 | 13.36 | 1.39 | 5.54 | 21.57 |
| THYN1 | 9 | 9.05 | 10.15 | 9.96 | 9.35 | 11.35 | 11.64 | 10.50 | 11.06 | 8.74 | 8.19 | 9.60 | 9.66 | 11.50 | 10.78 | 8.46 |
| TIA1 | 5 | 9.09 | 8.93 | 9.64 | 7.71 | 12.12 | 10.70 | 13.00 | 10.65 | 9.66 | 8.53 | 9.01 | 8.67 | 11.41 | 11.82 | 9.09 |
| TIAL1 | 17 | 9.94 | 10.49 | 10.49 | 10.10 | 9.87 | 9.88 | 10.47 | 9.39 | 9.78 | 9.59 | 10.22 | 10.29 | 9.88 | 9.93 | 9.68 |
| TICAM2 | 3 | 9.56 | 6.78 | 7.18 | 4.77 | 14.48 | 11.64 | 12.04 | 10.04 | 11.17 | 12.35 | 8.17 | 5.98 | 13.06 | 11.04 | 11.76 |
| TIFA | 1 | 10.38 | 10.33 | 10.86 | 10.99 | 10.33 | 10.81 | 9.65 | 8.97 | 9.71 | 7.97 | 10.36 | 10.92 | 10.57 | 9.31 | 8.84 |
| TIGAR | 12 | 8.96 | 9.50 | 11.67 | 10.24 | 12.84 | 11.84 | 8.96 | 6.92 | 9.59 | 9.49 | 9.23 | 10.95 | 12.34 | 7.94 | 9.54 |
| TIMELESS | 11 | 10.25 | 11.15 | 8.35 | 8.27 | 11.15 | 11.05 | 8.86 | 9.47 | 11.05 | 10.41 | 10.70 | 8.31 | 11.10 | 9.16 | 10.73 |
| TIMM10 | 11 | 10.54 | 9.05 | 10.48 | 10.78 | 10.50 | 10.84 | 11.42 | 7.53 | 9.36 | 9.51 | 9.79 | 10.63 | 10.67 | 9.47 | 9.43 |
| TIMM10B | 1 | 8.94 | 8.53 | 11.93 | 10.52 | 8.98 | 9.76 | 12.08 | 10.54 | 9.73 | 9.01 | 8.73 | 11.22 | 9.37 | 11.31 | 9.37 |
| TIMM13 | 9 | 8.47 | 8.66 | 10.25 | 10.57 | 10.74 | 11.01 | 10.87 | 12.39 | 8.32 | 8.72 | 8.57 | 10.41 | 10.87 | 11.63 | 8.52 |
| TIMM17A | 1 | 8.38 | 8.92 | 9.96 | 11.61 | 8.30 | 7.17 | 8.29 | 8.51 | 11.14 | 17.72 | 8.65 | 10.78 | 7.74 | 8.40 | 14.43 |
| TIMM17B | 1 | 12.18 | 5.93 | 3.90 | 8.03 | 12.91 | 9.39 | 15.34 | 13.86 | 6.91 | 11.55 | 9.06 | 5.97 | 11.15 | 14.60 | 9.23 |
| TIMM21 | 3 | 13.66 | 6.28 | 7.89 | 12.50 | 8.87 | 7.76 | 7.11 | 7.08 | 12.45 | 16.40 | 9.97 | 10.20 | 8.32 | 7.09 | 14.42 |
| TIMM23B | 7 | 12.72 | 6.26 | 7.55 | 9.92 | 9.01 | 8.21 | 11.38 | 10.19 | 11.11 | 13.65 | 9.49 | 8.73 | 8.61 | 10.79 | 12.38 |
| TIMM44 | 19 | 10.65 | 8.76 | 10.02 | 11.52 | 9.94 | 8.96 | 9.37 | 8.87 | 10.90 | 11.00 | 9.71 | 10.77 | 9.45 | 9.12 | 10.95 |
| TIMM50 | 8 | 11.38 | 6.81 | 9.71 | 12.11 | 6.98 | 6.65 | 12.62 | 11.88 | 10.06 | 11.80 | 9.09 | 10.91 | 6.81 | 12.25 | 10.93 |
| TIMM8A | 3 | 8.20 | 8.93 | 10.40 | 12.52 | 11.39 | 12.55 | 12.77 | 3.83 | 8.91 | 10.50 | 8.56 | 11.46 | 11.97 | 8.30 | 9.71 |
| TIMM8B | 4 | 10.05 | 9.22 | 11.07 | 12.19 | 10.65 | 12.19 | 10.72 | 5.59 | 8.62 | 9.69 | 9.63 | 11.63 | 11.42 | 8.16 | 9.16 |
| TIMM9 | 5 | 9.55 | 8.98 | 8.90 | 10.38 | 12.15 | 12.93 | 11.29 | 6.13 | 9.50 | 10.18 | 9.27 | 9.64 | 12.54 | 8.71 | 9.84 |
| TIMP1 | 1 | 4.55 | 3.90 | 5.23 | 5.02 | 14.46 | 13.31 | 19.95 | 22.27 | 5.57 | 5.74 | 4.23 | 5.13 | 13.88 | 21.11 | 5.66 |
| TIMP2 | 3 | 9.79 | 9.40 | 14.64 | 15.23 | 8.44 | 8.62 | 6.18 | 6.08 | 10.91 | 10.71 | 9.60 | 14.93 | 8.53 | 6.13 | 10.81 |
| TIPIN | 3 | 10.46 | 11.58 | 10.14 | 9.98 | 10.70 | 11.14 | 8.02 | 6.10 | 10.95 | 10.93 | 11.02 | 10.06 | 10.92 | 7.06 | 10.94 |
| TIPRL | 6 | 9.36 | 10.06 | 11.03 | 9.31 | 12.42 | 12.38 | 8.86 | 8.32 | 9.32 | 8.93 | 9.71 | 10.17 | 12.40 | 8.59 | 9.12 |
| TJAP1 | 3 | 8.90 | 10.82 | 10.94 | 9.83 | 11.18 | 11.51 | 9.12 | 9.02 | 9.09 | 9.60 | 9.86 | 10.38 | 11.34 | 9.07 | 9.34 |
| TJP1 | 57 | 8.39 | 9.27 | 13.19 | 12.28 | 9.60 | 9.87 | 7.64 | 6.75 | 11.66 | 11.35 | 8.83 | 12.74 | 9.74 | 7.19 | 11.51 |
| TJP2 | 34 | 9.26 | 9.66 | 12.98 | 11.96 | 12.14 | 12.42 | 4.33 | 3.73 | 12.29 | 11.23 | 9.46 | 12.47 | 12.28 | 4.03 | 11.76 |
| TK1 | 9 | 9.13 | 9.95 | 12.91 | 13.25 | 5.60 | 5.66 | 5.33 | 14.81 | 11.44 | 11.93 | 9.54 | 13.08 | 5.63 | 10.07 | 11.68 |
| TKT | 162 | 10.92 | 14.29 | 8.66 | 7.67 | 10.52 | 10.10 | 8.30 | 8.77 | 10.51 | 10.26 | 12.61 | 8.17 | 10.31 | 8.53 | 10.38 |
| TLDC1 | 1 | 9.71 | 10.68 | 8.84 | 9.76 | 7.74 | 9.26 | 7.78 | 8.32 | 14.67 | 13.24 | 10.20 | 9.30 | 8.50 | 8.05 | 13.96 |
| TLE3 | 12 | 8.56 | 9.20 | 12.36 | 11.22 | 9.26 | 9.35 | 10.88 | 10.21 | 9.51 | 9.47 | 8.88 | 11.79 | 9.30 | 10.54 | 9.49 |
| TLK1 | 9 | 9.50 | 10.57 | 11.37 | 10.55 | 10.10 | 9.76 | 9.96 | 9.18 | 9.67 | 9.34 | 10.04 | 10.96 | 9.93 | 9.57 | 9.50 |
| TLK2 | 6 | 11.10 | 11.12 | 10.77 | 11.04 | 8.41 | 8.62 | 9.64 | 8.26 | 11.03 | 10.00 | 11.11 | 10.91 | 8.51 | 8.95 | 10.52 |
| TLN1 | 194 | 9.66 | 10.91 | 13.71 | 12.78 | 6.79 | 6.84 | 5.25 | 6.22 | 14.26 | 13.59 | 10.28 | 13.25 | 6.81 | 5.74 | 13.92 |
| TLN2 | 45 | 9.75 | 10.94 | 11.54 | 11.24 | 8.10 | 8.53 | 6.26 | 8.81 | 12.47 | 12.35 | 10.35 | 11.39 | 8.31 | 7.54 | 12.41 |
| TLR1 | 1 | 5.76 | 6.95 | 18.00 | 14.99 | 7.21 | 7.37 | 13.99 | 10.92 | 8.04 | 6.78 | 6.35 | 16.50 | 7.29 | 12.45 | 7.41 |
| TLX3 | 1 | 10.31 | 12.07 | 6.71 | 7.37 | 13.55 | 13.73 | 8.88 | 8.74 | 9.25 | 9.39 | 11.19 | 7.04 | 13.64 | 8.81 | 9.32 |

| | | | | | | | | | | | | | | | | |
|---|---|---|---|---|---|---|---|---|---|---|---|---|---|---|---|---|
| TM7SF3 | 2 | 10.26 | 8.10 | 7.75 | 8.94 | 12.73 | 11.89 | 7.08 | 10.16 | 11.78 | 11.31 | 9.18 | 8.34 | 12.31 | 8.62 | 11.55 |
| TM9SF1 | 4 | 11.21 | 7.46 | 6.61 | 7.72 | 12.12 | 11.58 | 8.61 | 9.18 | 12.15 | 13.37 | 9.34 | 7.16 | 11.85 | 8.89 | 12.76 |
| TM9SF2 | 3 | 10.99 | 5.90 | 8.15 | 6.99 | 12.94 | 9.67 | 12.22 | 13.27 | 8.52 | 11.35 | 8.45 | 7.57 | 11.31 | 12.75 | 9.93 |
| TM9SF3 | 6 | 10.74 | 7.27 | 7.91 | 10.08 | 10.21 | 9.30 | 10.95 | 10.44 | 11.17 | 11.93 | 9.00 | 9.00 | 9.76 | 10.69 | 11.55 |
| TM9SF4 | 10 | 10.89 | 7.39 | 9.07 | 10.61 | 10.71 | 9.75 | 10.26 | 9.82 | 10.23 | 11.28 | 9.14 | 9.84 | 10.23 | 10.04 | 10.75 |
| TMA16 | 2 | 9.78 | 11.04 | 10.96 | 10.15 | 8.75 | 9.20 | 11.20 | 11.17 | 8.59 | 9.17 | 10.41 | 10.56 | 8.97 | 11.18 | 8.88 |
| TMA7 | 6 | 9.40 | 10.38 | 9.61 | 9.27 | 11.42 | 11.93 | 9.53 | 8.17 | 10.30 | 10.00 | 9.89 | 9.44 | 11.67 | 8.85 | 10.15 |
| TMC1 | 1 | 12.33 | 10.45 | 10.31 | 10.22 | 9.17 | 10.14 | 7.79 | 7.70 | 11.14 | 10.74 | 11.39 | 10.27 | 9.66 | 7.74 | 10.94 |
| TMCO1 | 4 | 13.22 | 6.03 | 7.78 | 12.52 | 9.16 | 7.99 | 10.16 | 10.11 | 10.33 | 12.71 | 9.62 | 10.15 | 8.57 | 10.13 | 11.52 |
| TMCO3 | 1 | 14.85 | 7.36 | 6.33 | 10.89 | 17.03 | 17.23 | 7.71 | 7.17 | 5.88 | 5.55 | 11.10 | 8.61 | 17.13 | 7.44 | 5.71 |
| TMED1 | 2 | 8.85 | 7.11 | 9.11 | 8.22 | 14.66 | 11.45 | 8.22 | 9.05 | 9.74 | 13.60 | 7.98 | 8.67 | 13.05 | 8.63 | 11.67 |
| TMED10 | 10 | 10.62 | 6.45 | 8.18 | 10.10 | 11.80 | 9.89 | 10.22 | 9.30 | 10.44 | 12.99 | 8.54 | 9.14 | 10.84 | 9.76 | 11.72 |
| TMED2 | 9 | 11.39 | 6.35 | 7.13 | 6.76 | 12.54 | 10.12 | 10.64 | 10.67 | 11.28 | 13.13 | 8.87 | 6.94 | 11.33 | 10.66 | 12.20 |
| TMED3 | 1 | 7.88 | 6.25 | 7.34 | 4.94 | 10.00 | 8.08 | 8.88 | 31.39 | 7.98 | 7.26 | 7.06 | 6.14 | 9.04 | 20.13 | 7.62 |
| TMED4 | 9 | 10.92 | 8.62 | 6.55 | 7.15 | 11.25 | 10.94 | 10.36 | 10.14 | 11.36 | 12.70 | 9.77 | 6.85 | 11.10 | 10.25 | 12.03 |
| TMED5 | 6 | 11.03 | 7.93 | 9.40 | 8.76 | 10.27 | 8.53 | 9.25 | 7.55 | 12.74 | 14.53 | 9.48 | 9.08 | 9.40 | 8.40 | 13.64 |
| TMED8 | 6 | 9.61 | 11.57 | 9.52 | 9.92 | 10.81 | 10.30 | 10.79 | 9.80 | 8.96 | 8.72 | 10.59 | 9.72 | 10.55 | 10.29 | 8.84 |
| TMED9 | 10 | 11.12 | 6.56 | 7.08 | 9.82 | 10.66 | 9.33 | 9.50 | 14.04 | 9.94 | 11.96 | 8.84 | 8.45 | 9.99 | 11.77 | 10.95 |
| TMEM106B | 1 | 12.49 | 11.19 | 7.61 | 9.29 | 8.01 | 8.91 | 8.82 | 9.12 | 11.18 | 13.38 | 11.84 | 8.45 | 8.46 | 8.97 | 12.28 |
| TMEM109 | 6 | 11.54 | 5.42 | 9.37 | 15.37 | 7.47 | 6.79 | 9.77 | 9.54 | 10.95 | 13.78 | 8.48 | 12.37 | 7.13 | 9.66 | 12.36 |
| TMEM11 | 5 | 10.52 | 11.28 | 11.45 | 9.30 | 11.20 | 10.25 | 7.18 | 8.16 | 10.57 | 10.09 | 10.90 | 10.38 | 10.73 | 7.67 | 10.33 |
| TMEM115 | 5 | 11.29 | 7.83 | 7.48 | 10.44 | 9.86 | 9.18 | 7.68 | 7.83 | 13.23 | 15.17 | 9.56 | 8.96 | 9.52 | 7.76 | 14.20 |
| TMEM126A | 6 | 10.92 | 8.84 | 11.17 | 10.15 | 11.12 | 9.61 | 8.96 | 6.54 | 10.41 | 12.27 | 9.88 | 10.66 | 10.36 | 7.75 | 11.34 |
| TMEM126B | 1 | 10.46 | 7.46 | 13.93 | 13.66 | 10.39 | 8.97 | 12.09 | 11.03 | 6.33 | 5.68 | 8.96 | 13.79 | 9.68 | 11.56 | 6.00 |
| TMEM132A | 1 | 10.22 | 7.78 | 11.15 | 9.90 | 8.44 | 7.76 | 11.02 | 11.04 | 12.28 | 10.42 | 9.00 | 10.52 | 8.10 | 11.03 | 11.35 |
| TMEM160 | 1 | 8.54 | 6.13 | 9.83 | 12.44 | 7.90 | 6.91 | 13.04 | 9.27 | 12.68 | 13.24 | 7.34 | 11.14 | 7.41 | 11.16 | 12.96 |
| TMEM164 | 1 | 13.80 | 6.99 | 6.45 | 8.47 | 10.62 | 8.68 | 12.97 | 13.31 | 8.61 | 10.11 | 10.39 | 7.46 | 9.65 | 13.14 | 9.36 |
| TMEM165 | 6 | 13.80 | 7.62 | 7.77 | 10.17 | 10.19 | 9.56 | 8.40 | 8.37 | 11.17 | 12.94 | 10.71 | 8.97 | 9.87 | 8.39 | 12.06 |
| TMEM167A | 1 | 10.73 | 8.26 | 6.69 | 8.26 | 11.27 | 8.70 | 9.70 | 14.17 | 9.24 | 12.98 | 9.50 | 7.48 | 9.98 | 11.93 | 11.11 |
| TMEM168 | 2 | 9.82 | 8.17 | 13.39 | 13.56 | 6.76 | 6.45 | 9.89 | 8.57 | 11.70 | 11.69 | 9.00 | 13.47 | 6.61 | 9.23 | 11.70 |
| TMEM179B | 1 | 10.36 | 8.61 | 8.61 | 9.29 | 11.27 | 10.32 | 9.30 | 9.19 | 12.22 | 10.84 | 9.48 | 8.95 | 10.79 | 9.25 | 11.53 |
| TMEM184C | 1 | 9.40 | 7.75 | 9.85 | 13.67 | 9.90 | 9.67 | 10.43 | 10.61 | 8.47 | 10.24 | 8.58 | 11.76 | 9.78 | 10.52 | 9.36 |
| TMEM189 | 1 | 13.58 | 7.29 | 7.45 | 12.13 | 10.18 | 8.57 | 7.56 | 8.23 | 10.78 | 14.22 | 10.43 | 9.79 | 9.38 | 7.90 | 12.50 |
| TMEM192 | 3 | 11.24 | 6.52 | 8.06 | 11.28 | 10.28 | 9.26 | 13.15 | 12.95 | 7.86 | 9.42 | 8.88 | 9.67 | 9.77 | 13.05 | 8.64 |
| TMEM199 | 1 | 14.67 | 7.02 | 7.74 | 13.67 | 7.50 | 6.48 | 6.88 | 6.76 | 12.22 | 17.07 | 10.84 | 10.70 | 6.99 | 6.82 | 14.64 |
| TMEM2 | 2 | 10.08 | 7.35 | 9.24 | 12.30 | 8.81 | 9.34 | 10.00 | 9.12 | 11.24 | 12.53 | 8.72 | 10.77 | 9.07 | 9.56 | 11.88 |
| TMEM201 | 1 | 13.62 | 6.10 | 7.79 | 10.25 | 7.82 | 7.02 | 10.94 | 10.47 | 10.78 | 15.21 | 9.86 | 9.02 | 7.42 | 10.70 | 12.99 |
| TMEM205 | 1 | 11.28 | 7.38 | 11.32 | 12.30 | 11.18 | 9.10 | 10.93 | 8.50 | 8.25 | 9.76 | 9.33 | 11.81 | 10.14 | 9.71 | 9.01 |
| TMEM209 | 2 | 13.23 | 7.99 | 8.25 | 13.51 | 7.68 | 7.76 | 9.29 | 9.33 | 10.73 | 12.24 | 10.61 | 10.88 | 7.72 | 9.31 | 11.49 |
| TMEM214 | 8 | 12.56 | 6.47 | 6.64 | 10.42 | 10.83 | 9.10 | 9.00 | 8.74 | 11.19 | 15.04 | 9.52 | 8.53 | 9.97 | 8.87 | 13.11 |
| TMEM223 | 4 | 12.47 | 7.55 | 8.99 | 12.05 | 8.73 | 7.47 | 9.12 | 8.26 | 11.85 | 13.52 | 10.01 | 10.52 | 8.10 | 8.69 | 12.68 |
| TMEM230 | 1 | 11.38 | 6.33 | 7.04 | 11.36 | 9.92 | 10.86 | 10.24 | 10.15 | 10.06 | 12.66 | 8.85 | 9.20 | 10.39 | 10.19 | 11.36 |

| | | | | | | | | | | | | | | | | |
|---|---|---|---|---|---|---|---|---|---|---|---|---|---|---|---|---|
| TMEM237 | 3 | 11.09 | 7.55 | 9.41 | 13.10 | 7.77 | 7.23 | 9.74 | 9.26 | 12.66 | 12.20 | 9.32 | 11.25 | 7.50 | 9.50 | 12.43 |
| TMEM245 | 1 | 13.88 | 7.29 | 5.72 | 8.81 | 10.18 | 10.82 | 8.63 | 6.75 | 12.66 | 15.27 | 10.58 | 7.26 | 10.50 | 7.69 | 13.97 |
| TMEM259 | 1 | 13.35 | 5.93 | 5.87 | 12.00 | 9.56 | 10.40 | 10.19 | 10.07 | 9.86 | 12.76 | 9.64 | 8.93 | 9.98 | 10.13 | 11.31 |
| TMEM30A | 2 | 11.89 | 8.06 | 7.62 | 9.21 | 10.13 | 9.28 | 11.25 | 11.18 | 10.35 | 11.03 | 9.97 | 8.42 | 9.71 | 11.21 | 10.69 |
| TMEM33 | 7 | 11.72 | 6.02 | 5.75 | 7.60 | 10.71 | 9.20 | 12.50 | 10.30 | 10.50 | 15.70 | 8.87 | 6.67 | 9.96 | 11.40 | 13.10 |
| TMEM43 | 13 | 11.47 | 5.96 | 7.32 | 11.86 | 7.87 | 6.50 | 8.68 | 7.58 | 14.02 | 18.74 | 8.71 | 9.59 | 7.19 | 8.13 | 16.38 |
| TMEM57 | 1 | 9.92 | 8.04 | 8.93 | 11.91 | 10.45 | 6.96 | 14.02 | 10.47 | 8.94 | 10.35 | 8.98 | 10.42 | 8.71 | 12.24 | 9.64 |
| TMEM63A | 1 | 11.24 | 10.11 | 10.63 | 10.76 | 8.49 | 9.95 | 7.52 | 6.81 | 14.90 | 9.59 | 10.67 | 10.69 | 9.22 | 7.16 | 12.25 |
| TMEM65 | 3 | 12.76 | 6.68 | 6.31 | 10.61 | 7.67 | 7.38 | 14.03 | 13.93 | 9.01 | 11.61 | 9.72 | 8.46 | 7.53 | 13.98 | 10.31 |
| TMEM70 | 1 | 12.30 | 8.57 | 6.49 | 8.01 | 8.99 | 8.89 | 12.88 | 14.35 | 9.85 | 9.66 | 10.44 | 7.25 | 8.94 | 13.62 | 9.76 |
| TMEM87A | 6 | 10.86 | 7.53 | 10.37 | 11.03 | 8.61 | 7.86 | 9.41 | 9.17 | 12.46 | 12.71 | 9.19 | 10.70 | 8.24 | 9.29 | 12.58 |
| TMEM9 | 2 | 11.89 | 6.89 | 8.33 | 10.46 | 14.22 | 12.65 | 7.63 | 7.04 | 9.26 | 11.63 | 9.39 | 9.39 | 13.43 | 7.34 | 10.45 |
| TMF1 | 22 | 10.40 | 12.07 | 8.62 | 8.21 | 11.37 | 11.42 | 7.86 | 7.31 | 11.80 | 10.95 | 11.23 | 8.42 | 11.39 | 7.58 | 11.37 |
| TMOD1 | 11 | 12.35 | 13.80 | 8.43 | 8.44 | 14.12 | 14.40 | 6.74 | 7.10 | 7.56 | 7.07 | 13.07 | 8.43 | 14.26 | 6.92 | 7.31 |
| TMOD2 | 1 | 3.70 | 5.48 | 12.72 | 13.28 | 6.83 | 7.09 | 21.72 | 21.24 | 3.77 | 4.18 | 4.59 | 13.00 | 6.96 | 21.48 | 3.97 |
| TMOD3 | 33 | 9.44 | 10.02 | 11.04 | 9.69 | 10.60 | 10.50 | 7.07 | 8.33 | 12.01 | 11.30 | 9.73 | 10.37 | 10.55 | 7.70 | 11.66 |
| TMPO | 31 | 8.80 | 8.58 | 14.80 | 10.55 | 10.45 | 8.85 | 8.22 | 8.20 | 10.61 | 10.93 | 8.69 | 12.68 | 9.65 | 8.21 | 10.77 |
| TMSB10 | 5 | 7.93 | 11.23 | 15.39 | 15.78 | 6.21 | 8.42 | 15.86 | 6.81 | 6.19 | 6.19 | 9.58 | 15.58 | 7.32 | 11.34 | 6.19 |
| TMTC3 | 3 | 10.23 | 7.04 | 10.60 | 16.02 | 7.93 | 6.80 | 10.51 | 9.45 | 9.59 | 11.82 | 8.64 | 13.31 | 7.37 | 9.98 | 10.71 |
| TMUB1 | 1 | 12.61 | 6.44 | 8.78 | 14.14 | 7.81 | 6.40 | 9.36 | 10.32 | 9.81 | 14.31 | 9.53 | 11.46 | 7.10 | 9.84 | 12.06 |
| TMX1 | 12 | 12.61 | 7.31 | 9.27 | 12.53 | 8.71 | 7.33 | 9.93 | 8.93 | 10.85 | 12.53 | 9.96 | 10.90 | 8.02 | 9.43 | 11.69 |
| TMX2 | 3 | 11.95 | 6.88 | 10.64 | 15.80 | 9.32 | 7.67 | 6.28 | 5.58 | 11.40 | 14.50 | 9.41 | 13.22 | 8.49 | 5.93 | 12.95 |
| TMX3 | 6 | 13.19 | 6.07 | 9.94 | 14.63 | 8.39 | 6.70 | 7.44 | 6.83 | 12.17 | 14.64 | 9.63 | 12.29 | 7.54 | 7.14 | 13.40 |
| TNC | 5 | 6.87 | 5.86 | 12.54 | 13.53 | 2.93 | 2.84 | 17.46 | 16.24 | 10.97 | 10.76 | 6.37 | 13.03 | 2.88 | 16.85 | 10.86 |
| TNFAIP2 | 14 | 10.60 | 11.35 | 11.87 | 10.62 | 8.36 | 8.13 | 8.01 | 7.30 | 12.07 | 11.71 | 10.97 | 11.24 | 8.24 | 7.66 | 11.89 |
| TNFAIP8 | 5 | 9.73 | 11.72 | 10.08 | 10.22 | 12.00 | 12.99 | 5.16 | 5.92 | 11.39 | 10.79 | 10.72 | 10.15 | 12.50 | 5.54 | 11.09 |
| TNFRSF10A | 1 | 9.13 | 5.91 | 8.39 | 10.12 | 8.63 | 7.94 | 14.04 | 10.66 | 11.67 | 13.51 | 7.52 | 9.26 | 8.28 | 12.35 | 12.59 |
| TNIP1 | 3 | 8.55 | 9.77 | 10.69 | 10.55 | 9.23 | 10.91 | 10.50 | 10.61 | 9.49 | 9.70 | 9.16 | 10.62 | 10.07 | 10.56 | 9.60 |
| TNK1 | 1 | 11.82 | 12.22 | 9.54 | 10.02 | 10.10 | 11.26 | 7.43 | 7.23 | 10.06 | 10.32 | 12.02 | 9.78 | 10.68 | 7.33 | 10.19 |
| TNKS | 1 | 8.44 | 10.05 | 12.19 | 12.90 | 9.98 | 9.80 | 10.15 | 8.89 | 9.01 | 8.59 | 9.24 | 12.55 | 9.89 | 9.52 | 8.80 |
| TNKS1BP1 | 53 | 9.28 | 10.14 | 11.88 | 11.02 | 10.58 | 10.69 | 7.79 | 7.45 | 10.95 | 10.24 | 9.71 | 11.45 | 10.63 | 7.62 | 10.59 |
| TNPO1 | 50 | 9.60 | 9.86 | 9.20 | 9.25 | 10.15 | 9.82 | 10.36 | 12.37 | 9.72 | 9.67 | 9.73 | 9.23 | 9.99 | 11.36 | 9.70 |
| TNPO2 | 4 | 12.42 | 9.42 | 9.82 | 9.49 | 10.16 | 11.51 | 8.51 | 9.12 | 10.89 | 8.64 | 10.92 | 9.66 | 10.84 | 8.82 | 9.76 |
| TNPO3 | 13 | 9.81 | 9.90 | 11.47 | 11.28 | 8.46 | 8.20 | 10.90 | 10.96 | 9.17 | 9.85 | 9.85 | 11.37 | 8.33 | 10.93 | 9.51 |
| TNRC6A | 5 | 9.49 | 11.88 | 10.90 | 11.46 | 8.02 | 9.02 | 9.46 | 8.52 | 11.04 | 10.21 | 10.69 | 11.18 | 8.52 | 8.99 | 10.62 |
| TNRC6B | 5 | 9.88 | 9.59 | 12.03 | 11.21 | 8.71 | 9.01 | 10.23 | 9.65 | 10.24 | 9.46 | 9.74 | 11.62 | 8.86 | 9.94 | 9.85 |
| TNS3 | 16 | 12.48 | 13.57 | 13.38 | 12.54 | 6.80 | 7.00 | 5.76 | 5.27 | 11.63 | 11.56 | 13.03 | 12.96 | 6.90 | 5.52 | 11.59 |
| TNS4 | 9 | 10.68 | 12.51 | 7.41 | 7.35 | 8.23 | 8.60 | 8.62 | 8.09 | 14.04 | 14.47 | 11.59 | 7.38 | 8.41 | 8.35 | 14.26 |
| TOE1 | 9 | 9.60 | 10.20 | 8.92 | 7.93 | 10.46 | 10.13 | 11.78 | 10.89 | 10.28 | 9.81 | 9.90 | 8.42 | 10.29 | 11.34 | 10.05 |
| TOLLIP | 1 | 9.81 | 10.35 | 11.70 | 12.34 | 9.85 | 10.66 | 8.73 | 9.18 | 8.69 | 8.69 | 10.08 | 12.02 | 10.26 | 8.96 | 8.69 |
| TOM1 | 5 | 10.27 | 13.69 | 9.74 | 11.11 | 8.85 | 10.30 | 8.55 | 9.16 | 9.24 | 9.09 | 11.98 | 10.43 | 9.57 | 8.86 | 9.16 |
| TOM1L1 | 4 | 10.94 | 12.56 | 13.26 | 14.37 | 8.77 | 10.14 | 5.60 | 4.90 | 10.64 | 8.82 | 11.75 | 13.82 | 9.45 | 5.25 | 9.73 |

| | | | | | | | | | | | | | | | | |
|---|---|---|---|---|---|---|---|---|---|---|---|---|---|---|---|---|
| TOM1L2 | 5 | 10.04 | 11.52 | 12.80 | 11.64 | 8.95 | 8.97 | 6.00 | 5.87 | 12.18 | 12.03 | 10.78 | 12.22 | 8.96 | 5.93 | 12.11 |
| TOMM20 | 5 | 12.11 | 9.33 | 8.27 | 10.13 | 9.44 | 8.53 | 10.14 | 9.35 | 11.07 | 11.63 | 10.72 | 9.20 | 8.99 | 9.75 | 11.35 |
| TOMM22 | 7 | 13.29 | 7.76 | 9.61 | 12.95 | 6.67 | 6.13 | 9.41 | 8.79 | 11.84 | 13.55 | 10.53 | 11.28 | 6.40 | 9.10 | 12.70 |
| TOMM34 | 22 | 9.23 | 10.51 | 9.45 | 9.02 | 9.57 | 9.55 | 9.17 | 12.48 | 10.72 | 10.31 | 9.87 | 9.23 | 9.56 | 10.82 | 10.51 |
| TOMM40 | 17 | 11.86 | 6.07 | 7.71 | 12.15 | 7.37 | 6.58 | 12.17 | 12.04 | 10.60 | 13.44 | 8.96 | 9.93 | 6.98 | 12.11 | 12.02 |
| TOMM5 | 5 | 10.49 | 4.86 | 7.20 | 11.16 | 8.31 | 6.57 | 11.53 | 12.77 | 12.64 | 14.48 | 7.67 | 9.18 | 7.44 | 12.15 | 13.56 |
| TOMM6 | 2 | 13.25 | 5.39 | 4.01 | 16.24 | 8.55 | 7.68 | 9.11 | 4.29 | 13.01 | 18.46 | 9.32 | 10.13 | 8.12 | 6.70 | 15.74 |
| TOMM70A | 26 | 11.81 | 6.20 | 9.44 | 13.46 | 8.62 | 7.69 | 10.38 | 9.62 | 10.43 | 12.35 | 9.00 | 11.45 | 8.16 | 10.00 | 11.39 |
| TONSL | 3 | 9.56 | 10.32 | 10.10 | 9.39 | 10.58 | 10.68 | 9.27 | 8.45 | 10.88 | 10.76 | 9.94 | 9.75 | 10.63 | 8.86 | 10.82 |
| TOP1 | 48 | 9.49 | 8.99 | 10.65 | 10.40 | 8.71 | 8.71 | 12.35 | 13.59 | 8.12 | 9.00 | 9.24 | 10.53 | 8.71 | 12.97 | 8.56 |
| TOP2A | 63 | 8.70 | 9.31 | 10.17 | 9.53 | 9.67 | 9.44 | 11.00 | 11.79 | 10.36 | 10.03 | 9.00 | 9.85 | 9.55 | 11.40 | 10.19 |
| TOP2B | 61 | 9.58 | 9.95 | 9.52 | 8.21 | 9.99 | 9.61 | 12.30 | 11.32 | 10.03 | 9.48 | 9.77 | 8.87 | 9.80 | 11.81 | 9.76 |
| TOP3A | 3 | 10.69 | 10.41 | 9.46 | 9.97 | 10.01 | 10.30 | 8.59 | 8.66 | 11.56 | 10.36 | 10.55 | 9.71 | 10.15 | 8.62 | 10.96 |
| TOP3B | 1 | 10.26 | 10.36 | 9.67 | 9.16 | 10.47 | 10.99 | 9.47 | 19.33 | 5.25 | 5.04 | 10.31 | 9.42 | 10.73 | 14.40 | 5.14 |
| TOPBP1 | 9 | 9.89 | 10.08 | 9.51 | 9.49 | 9.95 | 10.19 | 11.65 | 11.30 | 9.09 | 8.84 | 9.99 | 9.50 | 10.07 | 11.48 | 8.97 |
| TOR1A | 6 | 9.79 | 8.60 | 8.60 | 6.71 | 13.59 | 10.75 | 11.78 | 9.64 | 10.24 | 10.30 | 9.19 | 7.65 | 12.17 | 10.71 | 10.27 |
| TOR1AIP1 | 15 | 11.65 | 6.05 | 5.95 | 8.83 | 12.54 | 11.04 | 9.70 | 9.53 | 10.88 | 13.85 | 8.85 | 7.39 | 11.79 | 9.61 | 12.36 |
| TOR1AIP2 | 3 | 12.71 | 6.91 | 7.57 | 9.91 | 10.86 | 9.49 | 9.04 | 8.96 | 11.30 | 13.26 | 9.81 | 8.74 | 10.17 | 9.00 | 12.28 |
| TOR1B | 7 | 9.84 | 9.24 | 8.93 | 10.57 | 11.25 | 11.81 | 8.82 | 9.07 | 9.91 | 10.56 | 9.54 | 9.75 | 11.53 | 8.94 | 10.24 |
| TOR3A | 4 | 10.90 | 10.34 | 9.42 | 9.75 | 9.54 | 9.85 | 8.96 | 8.55 | 11.61 | 11.09 | 10.62 | 9.58 | 9.69 | 8.76 | 11.35 |
| TOR4A | 1 | 8.08 | 5.23 | 9.29 | 13.05 | 14.14 | 9.97 | 8.36 | 8.42 | 10.94 | 12.51 | 6.65 | 11.17 | 12.06 | 8.39 | 11.73 |
| TOX | 1 | 11.64 | 10.14 | 19.83 | 12.95 | 3.78 | 3.66 | 7.89 | 7.68 | 10.85 | 11.58 | 10.89 | 16.39 | 3.72 | 7.79 | 11.21 |
| TOX4 | 8 | 9.73 | 10.78 | 10.46 | 9.89 | 10.66 | 11.21 | 9.45 | 8.50 | 9.74 | 9.58 | 10.25 | 10.17 | 10.94 | 8.98 | 9.66 |
| TP53BP1 | 52 | 9.93 | 10.83 | 10.70 | 9.75 | 10.89 | 10.84 | 8.31 | 7.49 | 11.06 | 10.19 | 10.38 | 10.22 | 10.87 | 7.90 | 10.63 |
| TP53BP2 | 6 | 9.87 | 11.20 | 11.35 | 12.99 | 9.62 | 9.40 | 8.43 | 8.29 | 9.89 | 8.96 | 10.53 | 12.17 | 9.51 | 8.36 | 9.43 |
| TP53I3 | 10 | 10.65 | 11.77 | 14.31 | 13.56 | 9.10 | 9.25 | 5.49 | 5.01 | 10.47 | 10.38 | 11.21 | 13.94 | 9.18 | 5.25 | 10.42 |
| TP53RK | 3 | 8.93 | 9.43 | 11.62 | 9.37 | 11.39 | 11.00 | 10.27 | 10.05 | 9.02 | 8.92 | 9.18 | 10.49 | 11.20 | 10.16 | 8.97 |
| TPBG | 1 | 7.81 | 6.12 | 10.74 | 13.89 | 7.76 | 7.47 | 13.89 | 11.93 | 10.23 | 10.16 | 6.97 | 12.31 | 7.62 | 12.91 | 10.19 |
| TPD52 | 13 | 10.19 | 11.81 | 13.57 | 12.72 | 9.59 | 9.84 | 6.44 | 5.73 | 10.33 | 9.77 | 11.00 | 13.15 | 9.72 | 6.08 | 10.05 |
| TPD52L1 | 5 | 9.22 | 10.53 | 12.11 | 11.44 | 9.48 | 9.67 | 7.54 | 6.94 | 11.59 | 11.49 | 9.88 | 11.77 | 9.57 | 7.24 | 11.54 |
| TPD52L2 | 23 | 8.69 | 10.17 | 12.23 | 11.44 | 9.86 | 10.41 | 8.42 | 7.39 | 10.72 | 10.67 | 9.43 | 11.84 | 10.13 | 7.91 | 10.69 |
| TPI1 | 90 | 9.56 | 12.56 | 9.82 | 8.95 | 8.86 | 8.68 | 10.72 | 13.57 | 8.88 | 8.40 | 11.06 | 9.38 | 8.77 | 12.15 | 8.64 |
| TPM1 | 65 | 11.30 | 13.05 | 12.88 | 12.24 | 9.31 | 9.51 | 5.03 | 2.44 | 12.49 | 11.76 | 12.17 | 12.56 | 9.41 | 3.73 | 12.13 |
| TPM2 | 46 | 9.31 | 12.88 | 13.06 | 11.49 | 8.93 | 8.23 | 6.45 | 5.86 | 12.12 | 11.67 | 11.09 | 12.27 | 8.58 | 6.16 | 11.90 |
| TPM3 | 27 | 9.46 | 10.35 | 11.94 | 10.48 | 10.91 | 11.16 | 8.15 | 6.36 | 11.22 | 9.96 | 9.91 | 11.21 | 11.04 | 7.25 | 10.59 |
| TPM4 | 23 | 9.98 | 12.68 | 13.32 | 13.14 | 9.21 | 8.85 | 8.42 | 3.26 | 10.98 | 10.14 | 11.33 | 13.23 | 9.03 | 5.84 | 10.56 |
| TPMT | 5 | 10.35 | 11.09 | 11.23 | 10.97 | 11.19 | 11.78 | 8.45 | 7.99 | 8.62 | 8.32 | 10.72 | 11.10 | 11.49 | 8.22 | 8.47 |
| TPP1 | 10 | 8.22 | 9.19 | 9.05 | 7.35 | 16.71 | 15.68 | 9.37 | 7.77 | 8.62 | 8.04 | 8.71 | 8.20 | 16.20 | 8.57 | 8.33 |
| TPP2 | 51 | 9.88 | 10.64 | 10.08 | 9.84 | 10.55 | 10.71 | 10.40 | 10.45 | 8.80 | 8.66 | 10.26 | 9.96 | 10.63 | 10.42 | 8.73 |
| TPPP3 | 1 | 9.56 | 11.03 | 1.30 | 0.83 | 9.68 | 11.23 | 16.11 | 15.90 | 11.65 | 12.72 | 10.29 | 1.07 | 10.46 | 16.00 | 12.18 |
| TPR | 121 | 9.36 | 10.31 | 9.90 | 9.32 | 10.00 | 10.13 | 11.30 | 10.92 | 9.62 | 9.15 | 9.84 | 9.61 | 10.06 | 11.11 | 9.39 |
| TPRG1L | 2 | 8.52 | 9.30 | 9.43 | 8.87 | 10.23 | 9.37 | 10.49 | 9.81 | 11.69 | 12.29 | 8.91 | 9.15 | 9.80 | 10.15 | 11.99 |

| | | | | | | | | | | | | | | | | |
|---|---|---|---|---|---|---|---|---|---|---|---|---|---|---|---|---|
| TPRKB | 5 | 8.74 | 9.71 | 9.72 | 10.15 | 11.14 | 10.64 | 10.33 | 9.72 | 9.94 | 9.91 | 9.22 | 9.93 | 10.89 | 10.02 | 9.93 |
| TPRN | 1 | 9.09 | 10.30 | 9.19 | 10.92 | 9.85 | 10.95 | 7.66 | 8.04 | 12.66 | 11.34 | 9.69 | 10.05 | 10.40 | 7.85 | 12.00 |
| TPST1 | 2 | 9.02 | 6.86 | 11.05 | 14.44 | 6.83 | 6.43 | 12.99 | 10.07 | 10.70 | 11.61 | 7.94 | 12.74 | 6.63 | 11.53 | 11.16 |
| TPT1 | 37 | 9.02 | 9.98 | 12.33 | 10.96 | 11.15 | 10.55 | 7.28 | 12.62 | 8.04 | 8.07 | 9.50 | 11.65 | 10.85 | 9.95 | 8.05 |
| TPX2 | 38 | 8.02 | 9.63 | 11.41 | 10.56 | 9.70 | 9.86 | 10.63 | 9.89 | 10.31 | 9.99 | 8.82 | 10.98 | 9.78 | 10.26 | 10.15 |
| TRA2A | 5 | 10.53 | 10.36 | 10.69 | 8.67 | 8.42 | 8.00 | 10.80 | 11.55 | 10.53 | 10.46 | 10.44 | 9.68 | 8.21 | 11.18 | 10.50 |
| TRA2B | 13 | 9.48 | 10.75 | 9.15 | 8.20 | 11.27 | 11.70 | 11.18 | 11.24 | 8.59 | 8.44 | 10.12 | 8.68 | 11.49 | 11.21 | 8.51 |
| TRABD | 4 | 9.96 | 7.01 | 10.63 | 13.47 | 7.65 | 7.91 | 11.23 | 11.77 | 9.51 | 10.86 | 8.48 | 12.05 | 7.78 | 11.50 | 10.19 |
| TRADD | 3 | 10.96 | 10.96 | 9.40 | 9.02 | 11.82 | 10.89 | 9.24 | 7.10 | 10.67 | 9.93 | 10.96 | 9.21 | 11.36 | 8.17 | 10.30 |
| TRAF2 | 8 | 9.67 | 10.24 | 10.37 | 10.26 | 10.18 | 10.55 | 7.40 | 6.43 | 12.48 | 12.41 | 9.95 | 10.32 | 10.36 | 6.92 | 12.45 |
| TRAF3IP1 | 1 | 7.87 | 10.99 | 9.47 | 10.70 | 9.18 | 9.14 | 8.55 | 17.49 | 7.28 | 9.32 | 9.43 | 10.08 | 9.16 | 13.02 | 8.30 |
| TRAF4 | 3 | 10.64 | 11.37 | 9.99 | 8.41 | 11.06 | 11.00 | 8.99 | 7.91 | 10.64 | 9.99 | 11.00 | 9.20 | 11.03 | 8.45 | 10.31 |
| TRAF6 | 3 | 9.72 | 10.26 | 11.61 | 12.61 | 8.96 | 9.68 | 10.09 | 9.86 | 8.76 | 8.45 | 9.99 | 12.11 | 9.32 | 9.97 | 8.60 |
| TRAF7 | 4 | 9.82 | 9.48 | 11.02 | 10.47 | 9.09 | 9.33 | 8.94 | 8.23 | 12.86 | 10.76 | 9.65 | 10.74 | 9.21 | 8.59 | 11.81 |
| TRAFD1 | 1 | 13.64 | 15.99 | 7.08 | 7.15 | 9.97 | 10.67 | 5.55 | 6.32 | 10.84 | 12.79 | 14.81 | 7.11 | 10.32 | 5.93 | 11.82 |
| TRAM1 | 2 | 13.37 | 6.64 | 5.98 | 9.69 | 9.29 | 9.37 | 9.71 | 9.98 | 10.57 | 15.40 | 10.01 | 7.84 | 9.33 | 9.85 | 12.98 |
| TRAP1 | 54 | 10.72 | 9.50 | 9.46 | 10.03 | 10.15 | 9.60 | 10.61 | 9.59 | 9.84 | 10.51 | 10.11 | 9.74 | 9.87 | 10.10 | 10.18 |
| TRAPPC10 | 1 | 10.72 | 12.12 | 10.44 | 8.54 | 11.30 | 12.13 | 10.55 | 11.98 | 6.27 | 5.95 | 11.42 | 9.49 | 11.71 | 11.26 | 6.11 |
| TRAPPC11 | 4 | 8.92 | 10.51 | 11.20 | 9.28 | 11.49 | 11.45 | 8.93 | 8.28 | 10.40 | 9.54 | 9.71 | 10.24 | 11.47 | 8.61 | 9.97 |
| TRAPPC12 | 5 | 10.00 | 10.84 | 10.28 | 9.93 | 11.43 | 12.04 | 8.74 | 7.73 | 9.47 | 9.54 | 10.42 | 10.11 | 11.73 | 8.24 | 9.50 |
| TRAPPC13 | 3 | 10.05 | 11.24 | 8.77 | 9.31 | 11.71 | 12.41 | 8.29 | 8.27 | 9.83 | 10.13 | 10.64 | 9.04 | 12.06 | 8.28 | 9.98 |
| TRAPPC2L | 2 | 9.58 | 10.35 | 9.57 | 8.50 | 13.44 | 13.48 | 9.46 | 8.55 | 8.57 | 8.49 | 9.96 | 9.03 | 13.46 | 9.00 | 8.53 |
| TRAPPC3 | 8 | 9.72 | 11.24 | 9.85 | 9.82 | 9.85 | 10.15 | 9.51 | 9.81 | 10.03 | 10.02 | 10.48 | 9.83 | 10.00 | 9.66 | 10.03 |
| TRAPPC4 | 3 | 9.02 | 11.74 | 11.00 | 8.37 | 9.71 | 11.44 | 8.70 | 8.09 | 11.21 | 10.73 | 10.38 | 9.68 | 10.58 | 8.39 | 10.97 |
| TRAPPC5 | 7 | 10.03 | 10.66 | 10.19 | 10.89 | 11.20 | 11.75 | 8.76 | 8.02 | 9.33 | 9.17 | 10.35 | 10.54 | 11.47 | 8.39 | 9.25 |
| TRAPPC6B | 2 | 11.68 | 10.59 | 9.08 | 8.49 | 11.96 | 13.07 | 8.60 | 8.29 | 9.72 | 8.52 | 11.13 | 8.79 | 12.52 | 8.44 | 9.12 |
| TRAPPC8 | 15 | 10.39 | 10.46 | 10.15 | 9.67 | 12.32 | 12.45 | 7.15 | 8.18 | 9.68 | 9.54 | 10.43 | 9.91 | 12.39 | 7.67 | 9.61 |
| TRAPPC9 | 1 | 8.59 | 8.60 | 11.14 | 9.28 | 10.18 | 8.63 | 10.80 | 10.08 | 10.73 | 11.98 | 8.59 | 10.21 | 9.41 | 10.44 | 11.35 |
| TREX1 | 1 | 11.26 | 12.95 | 7.69 | 9.12 | 11.87 | 11.77 | 6.23 | 7.13 | 10.85 | 11.13 | 12.11 | 8.41 | 11.82 | 6.68 | 10.99 |
| TRIM11 | 4 | 11.54 | 12.84 | 8.67 | 8.37 | 10.42 | 10.03 | 8.13 | 9.44 | 10.68 | 9.88 | 12.19 | 8.52 | 10.23 | 8.78 | 10.28 |
| TRIM14 | 1 | 9.74 | 11.34 | 12.88 | 12.30 | 7.25 | 8.04 | 6.24 | 6.95 | 12.25 | 13.01 | 10.54 | 12.59 | 7.65 | 6.59 | 12.63 |
| TRIM16 | 20 | 12.21 | 12.46 | 9.42 | 8.75 | 9.57 | 9.29 | 4.62 | 4.28 | 14.92 | 14.48 | 12.34 | 9.09 | 9.43 | 4.45 | 14.70 |
| TRIM21 | 5 | 9.93 | 10.89 | 12.38 | 12.43 | 10.95 | 11.64 | 6.85 | 7.12 | 9.37 | 8.46 | 10.41 | 12.40 | 11.29 | 6.98 | 8.91 |
| TRIM24 | 8 | 8.68 | 9.19 | 10.59 | 9.88 | 9.66 | 9.69 | 13.37 | 12.19 | 8.30 | 8.44 | 8.93 | 10.24 | 9.68 | 12.78 | 8.37 |
| TRIM25 | 18 | 8.43 | 8.52 | 11.49 | 10.49 | 11.74 | 11.36 | 9.52 | 9.66 | 9.67 | 9.10 | 8.48 | 10.99 | 11.55 | 9.59 | 9.39 |
| TRIM26 | 4 | 9.62 | 10.57 | 11.09 | 10.90 | 10.31 | 10.62 | 9.22 | 8.29 | 9.90 | 9.49 | 10.10 | 11.00 | 10.46 | 8.75 | 9.69 |
| TRIM27 | 2 | 10.03 | 9.71 | 11.03 | 10.35 | 10.39 | 10.07 | 9.62 | 8.41 | 10.29 | 10.10 | 9.87 | 10.69 | 10.23 | 9.01 | 10.20 |
| TRIM28 | 85 | 8.94 | 9.52 | 10.62 | 9.46 | 9.35 | 9.21 | 12.98 | 14.76 | 7.80 | 7.35 | 9.23 | 10.04 | 9.28 | 13.87 | 7.58 |
| TRIM29 | 3 | 18.28 | 19.48 | 4.69 | 5.29 | 10.83 | 11.78 | 7.31 | 5.62 | 8.77 | 7.93 | 18.88 | 4.99 | 11.31 | 6.47 | 8.35 |
| TRIM3 | 6 | 8.52 | 9.76 | 13.04 | 12.37 | 9.72 | 9.26 | 8.43 | 7.99 | 10.54 | 10.38 | 9.14 | 12.71 | 9.49 | 8.21 | 10.46 |
| TRIM32 | 10 | 9.19 | 10.05 | 13.09 | 13.07 | 8.75 | 9.76 | 7.91 | 7.24 | 10.54 | 10.40 | 9.62 | 13.08 | 9.25 | 7.57 | 10.47 |
| TRIM33 | 10 | 9.70 | 10.49 | 10.12 | 9.91 | 8.50 | 8.74 | 11.96 | 11.16 | 10.02 | 9.39 | 10.10 | 10.01 | 8.62 | 11.56 | 9.71 |

| | | | | | | | | | | | | | | | | |
|---|---|---|---|---|---|---|---|---|---|---|---|---|---|---|---|---|
| TRIM38 | 1 | 11.16 | 11.68 | 9.99 | 10.94 | 11.65 | 14.18 | 8.26 | 6.26 | 8.09 | 7.79 | 11.42 | 10.47 | 12.91 | 7.26 | 7.94 |
| TRIM4 | 2 | 11.46 | 11.19 | 7.36 | 7.40 | 8.73 | 10.05 | 12.30 | 10.05 | 11.47 | 10.01 | 11.32 | 7.38 | 9.39 | 11.17 | 10.74 |
| TRIM47 | 9 | 10.25 | 11.20 | 8.70 | 8.04 | 8.23 | 7.62 | 8.01 | 8.04 | 15.08 | 14.83 | 10.72 | 8.37 | 7.92 | 8.02 | 14.96 |
| TRIM5 | 3 | 10.45 | 11.18 | 12.64 | 12.33 | 10.08 | 10.08 | 7.31 | 6.16 | 9.96 | 9.80 | 10.82 | 12.49 | 10.08 | 6.74 | 9.88 |
| TRIM56 | 5 | 10.28 | 11.13 | 11.66 | 12.19 | 8.48 | 8.83 | 9.66 | 8.01 | 10.33 | 9.43 | 10.71 | 11.93 | 8.66 | 8.84 | 9.88 |
| TRIM65 | 5 | 8.92 | 10.37 | 7.73 | 8.49 | 10.62 | 10.37 | 10.03 | 10.25 | 11.12 | 12.09 | 9.65 | 8.11 | 10.49 | 10.14 | 11.61 |
| TRIM7 | 1 | 10.11 | 12.95 | 6.09 | 4.60 | 11.27 | 12.30 | 12.58 | 11.41 | 9.84 | 8.83 | 11.53 | 5.35 | 11.79 | 12.00 | 9.34 |
| TRIO | 47 | 8.98 | 9.53 | 11.74 | 11.07 | 9.77 | 9.77 | 9.30 | 9.03 | 10.55 | 10.26 | 9.25 | 11.40 | 9.77 | 9.17 | 10.41 |
| TRIOBP | 6 | 10.06 | 10.74 | 12.26 | 11.84 | 9.14 | 9.28 | 7.53 | 6.93 | 11.25 | 10.96 | 10.40 | 12.05 | 9.21 | 7.23 | 11.11 |
| TRIP10 | 18 | 8.55 | 9.51 | 11.46 | 10.93 | 9.74 | 9.82 | 9.02 | 8.08 | 11.73 | 11.16 | 9.03 | 11.19 | 9.78 | 8.55 | 11.44 |
| TRIP11 | 25 | 9.62 | 11.02 | 9.09 | 8.90 | 9.14 | 9.33 | 11.58 | 10.71 | 10.47 | 10.13 | 10.32 | 8.99 | 9.24 | 11.14 | 10.30 |
| TRIP12 | 26 | 8.85 | 9.38 | 12.08 | 12.46 | 9.64 | 9.69 | 8.94 | 8.64 | 9.81 | 10.52 | 9.12 | 12.27 | 9.66 | 8.79 | 10.17 |
| TRIP13 | 27 | 9.93 | 10.62 | 10.02 | 9.66 | 8.43 | 8.54 | 9.80 | 11.21 | 10.98 | 10.82 | 10.27 | 9.84 | 8.49 | 10.50 | 10.90 |
| TRIP4 | 10 | 11.25 | 12.36 | 11.48 | 10.17 | 10.30 | 10.19 | 9.93 | 9.64 | 7.69 | 6.98 | 11.81 | 10.83 | 10.24 | 9.78 | 7.34 |
| TRIP6 | 10 | 10.24 | 10.72 | 10.95 | 8.09 | 9.88 | 9.86 | 9.03 | 7.60 | 12.03 | 11.60 | 10.48 | 9.52 | 9.87 | 8.32 | 11.81 |
| TRIT1 | 6 | 10.65 | 10.29 | 10.39 | 10.78 | 7.35 | 7.51 | 9.62 | 10.52 | 11.49 | 11.39 | 10.47 | 10.59 | 7.43 | 10.07 | 11.44 |
| TRMT1 | 19 | 8.86 | 10.06 | 10.42 | 10.36 | 8.36 | 8.77 | 11.70 | 11.70 | 9.91 | 9.86 | 9.46 | 10.39 | 8.56 | 11.70 | 9.88 |
| TRMT10A | 4 | 9.61 | 10.86 | 14.18 | 12.97 | 8.79 | 9.25 | 9.48 | 8.21 | 8.18 | 8.47 | 10.24 | 13.57 | 9.02 | 8.84 | 8.33 |
| TRMT10C | 17 | 9.76 | 8.15 | 10.31 | 11.35 | 8.94 | 8.72 | 11.01 | 12.86 | 9.17 | 9.73 | 8.96 | 10.83 | 8.83 | 11.94 | 9.45 |
| TRMT11 | 2 | 7.89 | 8.82 | 10.33 | 10.00 | 10.24 | 9.57 | 10.81 | 9.26 | 10.75 | 12.33 | 8.36 | 10.16 | 9.91 | 10.04 | 11.54 |
| TRMT112 | 8 | 9.12 | 9.52 | 11.64 | 8.54 | 11.56 | 10.11 | 10.15 | 9.18 | 10.52 | 9.67 | 9.32 | 10.09 | 10.84 | 9.66 | 10.09 |
| TRMT13 | 3 | 10.28 | 10.89 | 10.20 | 10.13 | 7.08 | 6.65 | 11.46 | 11.74 | 10.83 | 10.75 | 10.58 | 10.16 | 6.86 | 11.60 | 10.79 |
| TRMT1L | 7 | 8.45 | 9.68 | 8.10 | 7.55 | 11.21 | 11.90 | 10.94 | 13.07 | 9.58 | 9.51 | 9.07 | 7.83 | 11.55 | 12.01 | 9.55 |
| TRMT2A | 8 | 10.54 | 11.46 | 10.36 | 11.29 | 6.93 | 7.04 | 10.07 | 10.47 | 11.22 | 10.62 | 11.00 | 10.82 | 6.99 | 10.27 | 10.92 |
| TRMT5 | 9 | 9.41 | 9.99 | 11.29 | 11.18 | 9.35 | 9.56 | 11.02 | 9.87 | 9.37 | 8.97 | 9.70 | 11.24 | 9.45 | 10.45 | 9.17 |
| TRMT6 | 11 | 9.87 | 10.53 | 10.78 | 10.36 | 8.89 | 9.20 | 10.67 | 10.05 | 9.85 | 9.79 | 10.20 | 10.57 | 9.05 | 10.36 | 9.82 |
| TRMT61A | 1 | 8.77 | 11.04 | 10.58 | 9.72 | 9.65 | 10.51 | 9.05 | 9.51 | 10.78 | 10.39 | 9.91 | 10.15 | 10.08 | 9.28 | 10.58 |
| TRMT61B | 3 | 10.79 | 8.74 | 9.09 | 10.39 | 7.91 | 8.44 | 11.19 | 10.39 | 10.57 | 12.50 | 9.77 | 9.74 | 8.17 | 10.79 | 11.54 |
| TRMU | 5 | 11.59 | 9.38 | 9.54 | 11.50 | 7.87 | 7.64 | 11.93 | 11.24 | 9.50 | 9.82 | 10.49 | 10.52 | 7.76 | 11.58 | 9.66 |
| TRNAU1AP | 5 | 10.26 | 11.72 | 9.23 | 8.61 | 10.01 | 10.43 | 9.74 | 9.09 | 10.44 | 10.47 | 10.99 | 8.92 | 10.22 | 9.42 | 10.45 |
| TRNT1 | 17 | 9.00 | 9.61 | 9.11 | 8.76 | 11.90 | 11.70 | 11.50 | 10.80 | 9.07 | 8.55 | 9.31 | 8.93 | 11.80 | 11.15 | 8.81 |
| TROVE2 | 27 | 8.98 | 11.08 | 9.22 | 8.43 | 9.32 | 9.41 | 10.36 | 11.28 | 10.83 | 11.11 | 10.03 | 8.82 | 9.36 | 10.82 | 10.97 |
| TRPM7 | 1 | 8.71 | 11.97 | 8.88 | 8.30 | 9.67 | 12.47 | 10.12 | 10.13 | 11.13 | 8.63 | 10.34 | 8.59 | 11.07 | 10.13 | 9.88 |
| TRPS1 | 9 | 8.60 | 9.10 | 9.21 | 7.55 | 10.46 | 9.99 | 16.24 | 13.57 | 7.86 | 7.43 | 8.85 | 8.38 | 10.22 | 14.90 | 7.64 |
| TRPT1 | 3 | 8.89 | 9.22 | 11.84 | 9.54 | 12.06 | 10.64 | 9.83 | 8.65 | 9.53 | 9.80 | 9.05 | 10.69 | 11.35 | 9.24 | 9.67 |
| TRRAP | 13 | 9.79 | 9.62 | 9.41 | 8.68 | 10.20 | 10.23 | 10.85 | 10.94 | 10.13 | 10.15 | 9.70 | 9.05 | 10.21 | 10.89 | 10.14 |
| TRUB1 | 7 | 9.54 | 10.48 | 9.13 | 8.50 | 10.68 | 10.91 | 9.67 | 9.74 | 10.34 | 11.02 | 10.01 | 8.82 | 10.80 | 9.70 | 10.68 |
| TRUB2 | 3 | 9.74 | 8.95 | 9.13 | 12.35 | 8.21 | 8.16 | 11.51 | 13.23 | 9.57 | 9.14 | 9.35 | 10.74 | 8.19 | 12.37 | 9.36 |
| TSC1 | 5 | 10.11 | 10.62 | 9.06 | 8.67 | 10.71 | 10.98 | 9.32 | 8.92 | 10.56 | 11.04 | 10.37 | 8.87 | 10.84 | 9.12 | 10.80 |
| TSC2 | 4 | 10.61 | 10.86 | 9.18 | 10.22 | 11.17 | 11.35 | 8.08 | 7.73 | 10.82 | 9.96 | 10.74 | 9.70 | 11.26 | 7.91 | 10.39 |
| TSC22D1 | 3 | 9.44 | 11.14 | 10.95 | 14.25 | 9.63 | 11.75 | 7.89 | 6.30 | 8.03 | 10.62 | 10.29 | 12.60 | 10.69 | 7.09 | 9.33 |
| TSC22D2 | 2 | 13.87 | 11.13 | 11.80 | 12.14 | 9.43 | 10.83 | 6.56 | 6.50 | 9.97 | 7.78 | 12.50 | 11.97 | 10.13 | 6.53 | 8.87 |

| | | | | | | | | | | | | | | | | |
|---|---|---|---|---|---|---|---|---|---|---|---|---|---|---|---|---|
| TSC22D3 | 3 | 12.55 | 12.29 | 8.03 | 7.59 | 15.00 | 16.01 | 7.21 | 5.23 | 8.11 | 7.98 | 12.42 | 7.81 | 15.51 | 6.22 | 8.04 |
| TSC22D4 | 1 | 8.18 | 10.01 | 13.35 | 11.58 | 8.91 | 8.53 | 8.64 | 8.06 | 11.73 | 11.01 | 9.10 | 12.47 | 8.72 | 8.35 | 11.37 |
| TSEN15 | 2 | 9.77 | 9.92 | 9.63 | 7.71 | 11.12 | 11.71 | 8.24 | 7.61 | 11.90 | 12.41 | 9.84 | 8.67 | 11.41 | 7.92 | 12.15 |
| TSEN2 | 1 | 9.27 | 10.44 | 8.73 | 9.87 | 10.71 | 10.05 | 10.87 | 8.72 | 11.00 | 10.33 | 9.86 | 9.30 | 10.38 | 9.80 | 10.67 |
| TSEN34 | 10 | 9.64 | 10.63 | 10.27 | 9.73 | 10.66 | 9.76 | 9.79 | 8.26 | 10.52 | 10.72 | 10.14 | 10.00 | 10.21 | 9.03 | 10.62 |
| TSEN54 | 4 | 12.53 | 11.42 | 9.84 | 9.82 | 9.10 | 9.06 | 7.90 | 7.56 | 11.74 | 11.04 | 11.97 | 9.83 | 9.08 | 7.73 | 11.39 |
| TSFM | 14 | 10.52 | 8.57 | 10.24 | 11.47 | 9.52 | 8.78 | 9.65 | 12.05 | 9.50 | 9.71 | 9.54 | 10.85 | 9.15 | 10.85 | 9.61 |
| TSG101 | 16 | 8.11 | 10.26 | 12.57 | 11.32 | 10.09 | 10.02 | 9.84 | 9.03 | 9.60 | 9.16 | 9.19 | 11.95 | 10.06 | 9.43 | 9.38 |
| TSHZ2 | 1 | 12.21 | 11.91 | 7.07 | 7.18 | 9.04 | 8.87 | 16.37 | 15.27 | 6.61 | 5.48 | 12.06 | 7.13 | 8.96 | 15.82 | 6.04 |
| TSKU | 1 | 9.22 | 6.84 | 8.28 | 10.19 | 11.67 | 10.18 | 10.53 | 15.97 | 9.61 | 7.52 | 8.03 | 9.24 | 10.92 | 13.25 | 8.56 |
| TSN | 14 | 9.77 | 10.07 | 9.07 | 8.75 | 11.02 | 11.35 | 8.94 | 11.88 | 9.71 | 9.45 | 9.92 | 8.91 | 11.19 | 10.41 | 9.58 |
| TSNAX | 29 | 9.67 | 10.32 | 8.93 | 8.25 | 12.33 | 12.49 | 9.66 | 9.06 | 9.93 | 9.36 | 9.99 | 8.59 | 12.41 | 9.36 | 9.65 |
| TSPAN3 | 1 | 9.77 | 6.63 | 11.21 | 13.05 | 10.77 | 9.74 | 11.12 | 8.88 | 9.22 | 9.62 | 8.20 | 12.13 | 10.25 | 10.00 | 9.42 |
| TSPYL1 | 3 | 9.91 | 11.10 | 9.90 | 9.54 | 9.18 | 9.41 | 8.28 | 8.36 | 12.14 | 12.19 | 10.50 | 9.72 | 9.30 | 8.32 | 12.16 |
| TSR1 | 25 | 9.56 | 10.56 | 10.69 | 9.46 | 8.53 | 8.54 | 10.88 | 10.57 | 10.30 | 10.90 | 10.06 | 10.07 | 8.54 | 10.72 | 10.60 |
| TSR2 | 2 | 10.39 | 10.24 | 9.26 | 8.87 | 10.16 | 11.80 | 12.25 | 7.57 | 8.94 | 10.51 | 10.32 | 9.06 | 10.98 | 9.91 | 9.73 |
| TSR3 | 3 | 8.40 | 10.73 | 11.36 | 10.65 | 9.28 | 10.01 | 9.20 | 9.53 | 10.63 | 10.21 | 9.56 | 11.00 | 9.65 | 9.36 | 10.42 |
| TSSC1 | 7 | 10.12 | 11.30 | 11.60 | 11.44 | 10.41 | 11.36 | 8.79 | 11.48 | 6.78 | 6.73 | 10.71 | 11.52 | 10.88 | 10.14 | 6.75 |
| TSSC4 | 4 | 9.38 | 10.95 | 13.87 | 13.34 | 8.46 | 8.37 | 8.61 | 6.77 | 10.30 | 9.95 | 10.16 | 13.60 | 8.42 | 7.69 | 10.13 |
| TST | 9 | 12.31 | 10.36 | 8.78 | 6.58 | 9.98 | 8.94 | 12.11 | 10.97 | 10.09 | 9.88 | 11.33 | 7.68 | 9.46 | 11.54 | 9.99 |
| TSTA3 | 16 | 7.96 | 9.18 | 7.80 | 7.40 | 14.77 | 14.46 | 9.56 | 9.14 | 9.76 | 9.98 | 8.57 | 7.60 | 14.61 | 9.35 | 9.87 |
| TSTD1 | 1 | 13.27 | 10.08 | 9.34 | 9.62 | 11.81 | 10.85 | 8.46 | 7.99 | 11.00 | 7.57 | 11.67 | 9.48 | 11.33 | 8.23 | 9.29 |
| TSTD2 | 1 | 9.68 | 10.57 | 11.77 | 11.03 | 7.85 | 7.69 | 11.21 | 8.73 | 11.23 | 10.24 | 10.12 | 11.40 | 7.77 | 9.97 | 10.73 |
| TTC1 | 16 | 9.64 | 11.20 | 10.39 | 9.56 | 9.62 | 9.39 | 10.17 | 12.50 | 8.65 | 8.89 | 10.42 | 9.97 | 9.50 | 11.33 | 8.77 |
| TTC12 | 1 | 11.13 | 13.61 | 8.99 | 8.43 | 9.25 | 10.42 | 10.98 | 10.18 | 8.47 | 8.54 | 12.37 | 8.71 | 9.84 | 10.58 | 8.50 |
| TTC14 | 1 | 8.33 | 9.89 | 12.42 | 9.99 | 9.71 | 9.92 | 10.28 | 9.88 | 9.83 | 9.74 | 9.11 | 11.21 | 9.82 | 10.08 | 9.79 |
| TTC17 | 4 | 8.68 | 9.76 | 9.55 | 9.03 | 10.80 | 12.09 | 10.64 | 8.32 | 10.25 | 10.89 | 9.22 | 9.29 | 11.44 | 9.48 | 10.57 |
| TTC19 | 2 | 11.65 | 10.73 | 10.76 | 10.50 | 9.54 | 10.86 | 8.70 | 12.43 | 7.75 | 7.09 | 11.19 | 10.63 | 10.20 | 10.57 | 7.42 |
| TTC21B | 4 | 9.89 | 10.96 | 10.53 | 9.65 | 9.84 | 8.98 | 8.66 | 9.29 | 11.76 | 10.44 | 10.43 | 10.09 | 9.41 | 8.98 | 11.10 |
| TTC26 | 1 | 11.01 | 8.94 | 8.95 | 8.07 | 8.56 | 8.71 | 12.96 | 11.45 | 14.83 | 6.52 | 9.97 | 8.51 | 8.64 | 12.20 | 10.68 |
| TTC27 | 6 | 10.07 | 10.37 | 11.55 | 11.41 | 8.84 | 9.03 | 9.20 | 9.07 | 9.86 | 10.60 | 10.22 | 11.48 | 8.93 | 9.13 | 10.23 |
| TTC28 | 4 | 11.62 | 13.20 | 8.80 | 8.78 | 8.82 | 9.49 | 10.29 | 10.42 | 9.21 | 9.37 | 12.41 | 8.79 | 9.15 | 10.36 | 9.29 |
| TTC3 | 1 | 10.95 | 11.85 | 9.10 | 9.03 | 9.75 | 12.58 | 9.67 | 9.35 | 8.86 | 8.87 | 11.40 | 9.06 | 11.17 | 9.51 | 8.86 |
| TTC31 | 1 | 10.51 | 11.59 | 10.55 | 8.63 | 11.55 | 10.74 | 7.26 | 8.39 | 10.72 | 10.07 | 11.05 | 9.59 | 11.14 | 7.82 | 10.40 |
| TTC33 | 6 | 8.16 | 8.57 | 10.42 | 9.22 | 8.95 | 8.85 | 11.26 | 17.01 | 8.78 | 8.78 | 8.37 | 9.82 | 8.90 | 14.13 | 8.78 |
| TTC37 | 33 | 10.33 | 10.16 | 9.25 | 9.80 | 10.33 | 10.61 | 9.24 | 9.67 | 10.33 | 10.30 | 10.24 | 9.52 | 10.47 | 9.46 | 10.31 |
| TTC38 | 2 | 9.50 | 8.82 | 15.06 | 13.27 | 7.25 | 7.30 | 13.35 | 11.65 | 7.53 | 6.27 | 9.16 | 14.17 | 7.27 | 12.50 | 6.90 |
| TTC39C | 4 | 10.23 | 11.05 | 9.12 | 8.76 | 14.37 | 14.83 | 6.50 | 6.12 | 9.16 | 9.86 | 10.64 | 8.94 | 14.60 | 6.31 | 9.51 |
| TTC4 | 9 | 10.64 | 11.90 | 9.97 | 9.56 | 7.92 | 7.71 | 10.54 | 10.04 | 10.98 | 10.73 | 11.27 | 9.77 | 7.82 | 10.29 | 10.86 |
| TTC5 | 5 | 9.23 | 9.86 | 11.02 | 11.07 | 10.56 | 10.21 | 9.65 | 8.48 | 10.67 | 9.25 | 9.54 | 11.04 | 10.39 | 9.06 | 9.96 |
| TTC7A | 2 | 8.99 | 10.25 | 8.89 | 9.68 | 10.06 | 10.07 | 9.89 | 8.37 | 12.60 | 11.21 | 9.62 | 9.28 | 10.07 | 9.13 | 11.90 |
| TTC9C | 6 | 9.65 | 10.61 | 10.48 | 9.35 | 10.07 | 10.50 | 10.07 | 9.37 | 10.16 | 9.74 | 10.13 | 9.92 | 10.28 | 9.72 | 9.95 |

| | | | | | | | | | | | | | | | | |
|---|---|---|---|---|---|---|---|---|---|---|---|---|---|---|---|---|
| TTF1 | 3 | 10.46 | 9.48 | 9.80 | 10.35 | 8.79 | 9.13 | 10.43 | 9.82 | 11.60 | 10.13 | 9.97 | 10.08 | 8.96 | 10.13 | 10.86 |
| TTF2 | 11 | 10.19 | 11.20 | 12.00 | 11.63 | 7.47 | 7.62 | 9.15 | 8.64 | 10.89 | 11.20 | 10.70 | 11.82 | 7.55 | 8.89 | 11.05 |
| TTI1 | 11 | 9.22 | 9.45 | 9.06 | 9.46 | 9.51 | 10.12 | 10.70 | 15.26 | 8.79 | 8.43 | 9.33 | 9.26 | 9.82 | 12.98 | 8.61 |
| TTI2 | 6 | 9.64 | 10.06 | 9.46 | 9.97 | 9.73 | 10.50 | 11.44 | 11.19 | 8.97 | 9.05 | 9.85 | 9.72 | 10.11 | 11.32 | 9.01 |
| TTK | 19 | 9.40 | 9.51 | 11.95 | 11.38 | 8.89 | 8.96 | 10.65 | 9.43 | 10.23 | 9.61 | 9.45 | 11.67 | 8.92 | 10.04 | 9.92 |
| TTL | 1 | 10.52 | 9.76 | 10.77 | 10.14 | 11.30 | 10.43 | 9.71 | 9.10 | 9.19 | 9.07 | 10.14 | 10.45 | 10.87 | 9.40 | 9.13 |
| TTLL12 | 23 | 8.17 | 8.70 | 13.70 | 12.43 | 7.16 | 6.89 | 8.97 | 13.04 | 10.37 | 10.56 | 8.44 | 13.07 | 7.03 | 11.00 | 10.47 |
| TTN | 5 | 9.27 | 10.11 | 10.31 | 7.84 | 9.86 | 8.96 | 12.25 | 13.24 | 9.36 | 8.78 | 9.69 | 9.08 | 9.41 | 12.74 | 9.07 |
| TTPAL | 1 | 9.22 | 8.93 | 13.57 | 13.29 | 9.71 | 8.51 | 9.50 | 8.59 | 9.24 | 9.42 | 9.07 | 13.43 | 9.11 | 9.05 | 9.33 |
| TUB | 3 | 11.50 | 13.00 | 10.51 | 9.73 | 8.05 | 10.04 | 5.90 | 5.68 | 12.86 | 12.73 | 12.25 | 10.12 | 9.04 | 5.79 | 12.80 |
| TUBA1B | 190 | 8.92 | 11.61 | 10.91 | 8.73 | 10.42 | 9.74 | 9.28 | 10.18 | 10.16 | 10.04 | 10.27 | 9.82 | 10.08 | 9.73 | 10.10 |
| TUBA1C | 13 | 5.78 | 5.00 | 7.28 | 5.45 | 8.37 | 6.37 | 5.87 | 41.80 | 7.46 | 6.62 | 5.39 | 6.37 | 7.37 | 23.83 | 7.04 |
| TUBA4A | 12 | 9.98 | 10.39 | 11.37 | 10.62 | 11.65 | 11.27 | 6.78 | 8.86 | 9.98 | 9.09 | 10.19 | 11.00 | 11.46 | 7.82 | 9.54 |
| TUBAL3 | 1 | 10.96 | 11.71 | 6.54 | 5.13 | 8.76 | 11.48 | 8.61 | 20.87 | 8.23 | 7.71 | 11.33 | 5.83 | 10.12 | 14.74 | 7.97 |
| TUBB | 24 | 9.00 | 11.96 | 10.92 | 10.53 | 9.21 | 9.11 | 10.09 | 8.92 | 10.23 | 10.03 | 10.48 | 10.72 | 9.16 | 9.50 | 10.13 |
| TUBB2A | 58 | 7.79 | 8.37 | 9.08 | 8.24 | 8.66 | 8.48 | 8.02 | 24.60 | 8.60 | 8.16 | 8.08 | 8.66 | 8.57 | 16.31 | 8.38 |
| TUBB3 | 133 | 8.41 | 10.25 | 10.58 | 10.16 | 9.98 | 9.67 | 9.27 | 12.63 | 9.62 | 9.43 | 9.33 | 10.37 | 9.82 | 10.95 | 9.53 |
| TUBB4A | 1 | 6.98 | 9.92 | 11.39 | 10.75 | 11.27 | 8.79 | 7.19 | 9.24 | 12.69 | 11.76 | 8.45 | 11.07 | 10.03 | 8.22 | 12.23 |
| TUBB4B | 31 | 8.62 | 10.01 | 10.96 | 9.87 | 10.72 | 10.41 | 9.28 | 10.13 | 10.13 | 9.87 | 9.32 | 10.42 | 10.56 | 9.71 | 10.00 |
| TUBB6 | 52 | 8.48 | 9.70 | 12.17 | 10.56 | 11.38 | 11.03 | 6.83 | 7.07 | 11.43 | 11.36 | 9.09 | 11.36 | 11.21 | 6.95 | 11.39 |
| TUBD1 | 2 | 12.58 | 11.65 | 11.42 | 9.15 | 6.87 | 6.62 | 7.18 | 7.20 | 14.10 | 13.22 | 12.12 | 10.29 | 6.74 | 7.19 | 13.66 |
| TUBG1 | 17 | 9.03 | 10.04 | 11.68 | 10.50 | 10.55 | 10.52 | 9.01 | 9.23 | 9.88 | 9.58 | 9.53 | 11.09 | 10.53 | 9.12 | 9.73 |
| TUBGCP2 | 15 | 9.48 | 10.09 | 9.96 | 10.22 | 10.52 | 11.10 | 10.45 | 10.18 | 9.01 | 8.99 | 9.78 | 10.09 | 10.81 | 10.31 | 9.00 |
| TUBGCP3 | 16 | 9.71 | 9.85 | 10.05 | 9.99 | 11.17 | 11.29 | 10.69 | 10.39 | 8.37 | 8.49 | 9.78 | 10.02 | 11.23 | 10.54 | 8.43 |
| TUBGCP4 | 8 | 9.43 | 10.08 | 10.06 | 10.20 | 9.59 | 10.29 | 9.33 | 9.33 | 10.91 | 10.77 | 9.76 | 10.13 | 9.94 | 9.33 | 10.84 |
| TUBGCP5 | 1 | 8.63 | 10.90 | 11.06 | 10.17 | 10.61 | 10.40 | 7.30 | 5.55 | 12.70 | 12.68 | 9.76 | 10.62 | 10.50 | 6.43 | 12.69 |
| TUBGCP6 | 3 | 8.92 | 12.00 | 8.51 | 9.40 | 10.47 | 12.48 | 7.59 | 7.33 | 14.71 | 8.59 | 10.46 | 8.96 | 11.48 | 7.46 | 11.65 |
| TUFM | 55 | 10.51 | 8.89 | 10.43 | 11.45 | 8.74 | 8.25 | 9.02 | 13.11 | 9.54 | 10.06 | 9.70 | 10.94 | 8.50 | 11.07 | 9.80 |
| TUFT1 | 7 | 9.88 | 11.25 | 13.25 | 12.66 | 10.29 | 10.34 | 3.22 | 2.70 | 13.33 | 13.07 | 10.56 | 12.96 | 10.32 | 2.96 | 13.20 |
| TUSC1 | 5 | 9.18 | 10.94 | 10.01 | 9.46 | 9.26 | 9.92 | 8.16 | 8.10 | 13.01 | 11.96 | 10.06 | 9.74 | 9.59 | 8.13 | 12.49 |
| TUSC3 | 4 | 13.64 | 7.67 | 6.93 | 12.32 | 5.95 | 6.67 | 10.06 | 9.39 | 11.83 | 15.55 | 10.65 | 9.63 | 6.31 | 9.73 | 13.69 |
| TUT1 | 7 | 8.94 | 9.61 | 10.19 | 9.62 | 12.53 | 12.16 | 9.71 | 8.84 | 9.88 | 8.53 | 9.27 | 9.90 | 12.34 | 9.27 | 9.21 |
| TWF1 | 28 | 9.72 | 9.52 | 10.73 | 8.29 | 10.98 | 10.41 | 9.84 | 11.61 | 9.78 | 9.10 | 9.62 | 9.51 | 10.70 | 10.73 | 9.44 |
| TWF2 | 18 | 9.30 | 10.81 | 10.96 | 10.32 | 9.01 | 9.48 | 10.24 | 9.64 | 9.80 | 10.43 | 10.06 | 10.64 | 9.24 | 9.94 | 10.12 |
| TWISTNB | 7 | 9.43 | 10.18 | 12.25 | 10.24 | 7.85 | 8.20 | 12.23 | 12.47 | 8.64 | 8.52 | 9.81 | 11.24 | 8.02 | 12.35 | 8.58 |
| TXLNA | 34 | 11.31 | 12.79 | 8.91 | 8.59 | 8.97 | 9.22 | 10.21 | 9.55 | 10.46 | 9.98 | 12.05 | 8.75 | 9.10 | 9.88 | 10.22 |
| TXLNG | 12 | 9.58 | 10.59 | 8.80 | 8.50 | 11.48 | 11.93 | 11.44 | 11.32 | 8.41 | 7.93 | 10.09 | 8.65 | 11.70 | 11.38 | 8.17 |
| TXN | 40 | 9.09 | 11.99 | 8.63 | 7.81 | 10.21 | 9.92 | 11.04 | 11.34 | 9.73 | 10.23 | 10.54 | 8.22 | 10.07 | 11.19 | 9.98 |
| TXN2 | 2 | 13.44 | 10.70 | 9.33 | 12.20 | 7.93 | 9.54 | 8.15 | 6.69 | 10.93 | 11.10 | 12.07 | 10.76 | 8.73 | 7.42 | 11.01 |
| TXNDC12 | 19 | 9.52 | 10.40 | 9.46 | 8.38 | 10.56 | 10.15 | 12.17 | 9.74 | 10.01 | 9.60 | 9.96 | 8.92 | 10.36 | 10.96 | 9.81 |
| TXNDC15 | 1 | 11.94 | 7.57 | 6.69 | 9.83 | 10.73 | 9.52 | 9.81 | 10.12 | 10.36 | 13.44 | 9.75 | 8.26 | 10.13 | 9.96 | 11.90 |
| TXNDC17 | 11 | 9.82 | 10.92 | 10.69 | 9.32 | 10.82 | 10.69 | 9.52 | 7.61 | 10.20 | 10.43 | 10.37 | 10.00 | 10.75 | 8.56 | 10.31 |

| | | | | | | | | | | | | | | | | |
|---|---|---|---|---|---|---|---|---|---|---|---|---|---|---|---|---|
| TXNDC5 | 32 | 7.99 | 7.85 | 13.38 | 12.66 | 9.38 | 9.24 | 9.70 | 9.22 | 10.09 | 10.48 | 7.92 | 13.02 | 9.31 | 9.46 | 10.29 |
| TXNDC9 | 10 | 10.53 | 11.14 | 11.70 | 9.93 | 10.46 | 10.42 | 8.48 | 6.90 | 10.13 | 10.30 | 10.83 | 10.82 | 10.44 | 7.69 | 10.22 |
| TXNL1 | 25 | 9.86 | 10.82 | 12.26 | 11.44 | 10.94 | 10.88 | 8.13 | 7.73 | 9.00 | 8.96 | 10.34 | 11.85 | 10.91 | 7.93 | 8.98 |
| TXNL4A | 9 | 10.28 | 10.79 | 10.32 | 9.95 | 10.44 | 10.50 | 8.48 | 8.03 | 10.76 | 10.46 | 10.53 | 10.13 | 10.47 | 8.25 | 10.61 |
| TXNL4B | 1 | 10.83 | 12.03 | 9.42 | 9.43 | 8.44 | 9.00 | 9.84 | 7.91 | 11.58 | 11.53 | 11.43 | 9.43 | 8.72 | 8.87 | 11.55 |
| TXNRD1 | 57 | 13.05 | 14.13 | 10.95 | 10.29 | 8.94 | 9.24 | 5.88 | 5.41 | 11.30 | 10.82 | 13.59 | 10.62 | 9.09 | 5.64 | 11.06 |
| TXNRD2 | 6 | 11.33 | 12.04 | 8.70 | 10.06 | 11.23 | 13.18 | 10.91 | 11.27 | 5.21 | 6.08 | 11.68 | 9.38 | 12.20 | 11.09 | 5.64 |
| TYK2 | 2 | 11.50 | 10.17 | 7.86 | 8.24 | 11.02 | 11.36 | 8.58 | 8.95 | 11.34 | 10.97 | 10.84 | 8.05 | 11.19 | 8.77 | 11.16 |
| TYMP | 9 | 9.10 | 10.70 | 9.68 | 9.28 | 5.13 | 5.48 | 13.07 | 12.08 | 12.83 | 12.65 | 9.90 | 9.48 | 5.30 | 12.57 | 12.74 |
| TYMS | 13 | 10.96 | 11.55 | 8.82 | 8.37 | 8.41 | 8.50 | 6.66 | 6.05 | 15.40 | 15.27 | 11.26 | 8.60 | 8.46 | 6.35 | 15.33 |
| TYRO3 | 1 | 18.01 | 19.94 | 2.83 | 2.85 | 14.30 | 14.07 | 7.71 | 6.90 | 6.72 | 6.67 | 18.98 | 2.84 | 14.19 | 7.30 | 6.69 |
| TYW3 | 1 | 10.91 | 10.73 | 7.87 | 9.36 | 9.51 | 10.74 | 10.41 | 11.70 | 9.23 | 9.56 | 10.82 | 8.61 | 10.12 | 11.05 | 9.40 |
| TYW5 | 2 | 7.55 | 8.32 | 9.68 | 7.45 | 8.78 | 9.81 | 8.11 | 21.85 | 10.36 | 8.09 | 7.93 | 8.56 | 9.30 | 14.98 | 9.22 |
| U2AF1 | 6 | 7.32 | 6.53 | 8.09 | 5.87 | 8.07 | 7.10 | 8.19 | 33.63 | 7.84 | 7.36 | 6.93 | 6.98 | 7.59 | 20.91 | 7.60 |
| U2AF2 | 24 | 9.20 | 9.86 | 9.88 | 9.19 | 9.26 | 9.25 | 11.26 | 13.91 | 9.33 | 8.86 | 9.53 | 9.54 | 9.26 | 12.58 | 9.10 |
| U2SURP | 32 | 8.89 | 10.02 | 12.47 | 10.16 | 9.22 | 8.86 | 10.75 | 10.46 | 9.82 | 9.35 | 9.46 | 11.31 | 9.04 | 10.60 | 9.58 |
| UACA | 10 | 11.38 | 12.37 | 7.82 | 7.69 | 10.41 | 10.72 | 6.68 | 6.40 | 13.58 | 12.94 | 11.87 | 7.76 | 10.56 | 6.54 | 13.26 |
| UAP1 | 25 | 8.94 | 9.38 | 12.45 | 11.71 | 9.16 | 9.27 | 9.58 | 9.14 | 10.29 | 10.09 | 9.16 | 12.08 | 9.21 | 9.36 | 10.19 |
| UAP1L1 | 7 | 10.96 | 11.70 | 10.24 | 9.79 | 7.65 | 7.68 | 8.89 | 7.80 | 12.60 | 12.67 | 11.33 | 10.02 | 7.67 | 8.35 | 12.64 |
| UBA1 | 112 | 8.44 | 9.12 | 8.82 | 8.13 | 12.25 | 12.33 | 11.53 | 13.63 | 8.15 | 7.61 | 8.78 | 8.47 | 12.29 | 12.58 | 7.88 |
| UBA2 | 35 | 9.58 | 10.07 | 9.46 | 8.77 | 8.23 | 8.40 | 13.23 | 13.66 | 9.61 | 8.99 | 9.83 | 9.12 | 8.31 | 13.45 | 9.30 |
| UBA3 | 12 | 9.53 | 10.36 | 8.15 | 8.22 | 11.21 | 11.38 | 9.93 | 12.28 | 9.42 | 9.53 | 9.94 | 8.18 | 11.30 | 11.10 | 9.47 |
| UBA5 | 10 | 9.21 | 10.04 | 11.73 | 11.62 | 10.35 | 10.95 | 9.07 | 8.88 | 9.25 | 8.89 | 9.63 | 11.67 | 10.65 | 8.97 | 9.07 |
| UBA6 | 46 | 10.23 | 11.01 | 10.14 | 9.79 | 9.40 | 9.57 | 9.84 | 10.48 | 9.98 | 9.56 | 10.62 | 9.96 | 9.49 | 10.16 | 9.77 |
| UBAC1 | 5 | 10.03 | 10.98 | 8.43 | 8.37 | 12.18 | 11.88 | 10.89 | 9.93 | 8.87 | 8.43 | 10.51 | 8.40 | 12.03 | 10.41 | 8.65 |
| UBAP1 | 7 | 9.93 | 9.69 | 9.86 | 8.10 | 10.99 | 10.60 | 10.36 | 8.31 | 11.48 | 10.65 | 9.81 | 8.98 | 10.80 | 9.34 | 11.07 |
| UBAP2 | 22 | 10.75 | 11.89 | 11.44 | 10.78 | 8.11 | 8.49 | 8.29 | 7.38 | 11.64 | 11.24 | 11.32 | 11.11 | 8.30 | 7.83 | 11.44 |
| UBAP2L | 33 | 8.66 | 9.32 | 10.92 | 9.85 | 12.54 | 12.35 | 10.13 | 8.73 | 9.12 | 8.37 | 8.99 | 10.39 | 12.45 | 9.43 | 8.75 |
| UBE2A | 1 | 9.70 | 9.94 | 10.00 | 9.70 | 11.25 | 11.73 | 10.24 | 8.61 | 9.72 | 9.10 | 9.82 | 9.85 | 11.49 | 9.42 | 9.41 |
| UBE2B | 1 | 9.86 | 11.11 | 10.50 | 8.53 | 10.40 | 8.65 | 12.29 | 6.82 | 10.46 | 11.37 | 10.49 | 9.51 | 9.53 | 9.56 | 10.92 |
| UBE2C | 7 | 9.67 | 10.44 | 10.49 | 8.65 | 10.13 | 10.34 | 10.81 | 9.05 | 10.45 | 9.98 | 10.05 | 9.57 | 10.24 | 9.93 | 10.21 |
| UBE2D3 | 3 | 8.95 | 9.89 | 10.16 | 10.03 | 10.68 | 8.88 | 12.12 | 10.16 | 9.74 | 9.40 | 9.42 | 10.09 | 9.78 | 11.14 | 9.57 |
| UBE2D4 | 1 | 8.28 | 9.24 | 10.26 | 9.76 | 9.49 | 9.31 | 14.17 | 12.77 | 8.78 | 7.94 | 8.76 | 10.01 | 9.40 | 13.47 | 8.36 |
| UBE2E1 | 3 | 8.63 | 10.14 | 10.40 | 9.11 | 8.96 | 9.02 | 8.35 | 17.41 | 9.36 | 8.62 | 9.39 | 9.76 | 8.99 | 12.88 | 8.99 |
| UBE2E2 | 1 | 10.87 | 11.90 | 9.17 | 8.70 | 9.58 | 8.97 | 8.91 | 8.86 | 12.40 | 10.64 | 11.38 | 8.93 | 9.28 | 8.89 | 11.52 |
| UBE2E3 | 6 | 8.75 | 9.16 | 12.40 | 11.26 | 9.47 | 10.27 | 10.74 | 9.95 | 8.99 | 9.00 | 8.95 | 11.83 | 9.87 | 10.35 | 9.00 |
| UBE2F | 1 | 8.82 | 9.79 | 14.02 | 13.28 | 7.21 | 7.86 | 8.90 | 8.27 | 10.77 | 11.10 | 9.30 | 13.65 | 7.53 | 8.58 | 10.93 |
| UBE2G1 | 6 | 10.67 | 11.33 | 14.57 | 9.63 | 9.54 | 9.12 | 8.87 | 6.57 | 10.18 | 9.52 | 11.00 | 12.10 | 9.33 | 7.72 | 9.85 |
| UBE2G2 | 1 | 10.72 | 11.46 | 8.25 | 8.03 | 8.00 | 8.69 | 11.92 | 9.91 | 11.55 | 11.48 | 11.09 | 8.14 | 8.34 | 10.91 | 11.52 |
| UBE2H | 9 | 9.65 | 10.52 | 12.95 | 11.92 | 7.80 | 7.98 | 7.04 | 6.15 | 13.61 | 12.37 | 10.09 | 12.44 | 7.89 | 6.60 | 12.99 |
| UBE2I | 15 | 9.21 | 10.34 | 11.29 | 10.14 | 9.61 | 9.86 | 11.15 | 10.37 | 9.20 | 8.83 | 9.77 | 10.72 | 9.74 | 10.76 | 9.01 |
| UBE2J2 | 1 | 12.29 | 8.75 | 8.78 | 10.28 | 8.67 | 8.75 | 10.19 | 10.80 | 10.30 | 11.19 | 10.52 | 9.53 | 8.71 | 10.50 | 10.75 |

| | | | | | | | | | | | | | | | | |
|---|---|---|---|---|---|---|---|---|---|---|---|---|---|---|---|---|
| UBE2K | 10 | 8.68 | 9.99 | 10.03 | 9.89 | 10.10 | 10.52 | 10.79 | 10.56 | 9.92 | 9.53 | 9.33 | 9.96 | 10.31 | 10.67 | 9.72 |
| UBE2L3 | 13 | 9.89 | 10.36 | 12.75 | 10.06 | 8.17 | 7.66 | 9.73 | 11.38 | 10.32 | 9.68 | 10.12 | 11.41 | 7.92 | 10.55 | 10.00 |
| UBE2M | 14 | 9.10 | 9.93 | 10.04 | 8.72 | 9.34 | 9.21 | 13.10 | 13.61 | 8.89 | 8.05 | 9.51 | 9.38 | 9.27 | 13.36 | 8.47 |
| UBE2N | 21 | 8.95 | 9.37 | 10.51 | 8.93 | 9.64 | 9.57 | 10.01 | 13.86 | 9.67 | 9.49 | 9.16 | 9.72 | 9.61 | 11.94 | 9.58 |
| UBE2O | 17 | 10.90 | 11.44 | 11.00 | 11.07 | 8.59 | 8.89 | 9.29 | 8.70 | 10.22 | 9.90 | 11.17 | 11.04 | 8.74 | 8.99 | 10.06 |
| UBE2Q1 | 8 | 9.48 | 9.88 | 10.66 | 8.59 | 12.42 | 11.82 | 10.24 | 8.91 | 9.26 | 8.74 | 9.68 | 9.62 | 12.12 | 9.58 | 9.00 |
| UBE2R2 | 5 | 10.08 | 10.93 | 10.62 | 10.34 | 9.96 | 10.20 | 7.90 | 7.18 | 12.03 | 10.78 | 10.50 | 10.48 | 10.08 | 7.54 | 11.40 |
| UBE2S | 11 | 9.30 | 9.97 | 10.44 | 9.50 | 7.83 | 7.67 | 12.62 | 12.80 | 10.04 | 9.84 | 9.63 | 9.97 | 7.75 | 12.71 | 9.94 |
| UBE2T | 8 | 10.24 | 10.59 | 6.93 | 7.06 | 12.59 | 12.66 | 11.16 | 10.24 | 9.51 | 9.04 | 10.41 | 6.99 | 12.62 | 10.70 | 9.27 |
| UBE2V1 | 17 | 9.20 | 10.22 | 11.52 | 11.01 | 9.41 | 9.56 | 10.26 | 10.06 | 9.46 | 9.31 | 9.71 | 11.26 | 9.48 | 10.16 | 9.39 |
| UBE2V2 | 11 | 9.93 | 10.49 | 9.65 | 8.66 | 10.11 | 10.23 | 12.31 | 10.99 | 9.07 | 8.57 | 10.21 | 9.15 | 10.17 | 11.65 | 8.82 |
| UBE2W | 1 | 8.09 | 9.22 | 10.69 | 11.59 | 9.34 | 11.51 | 10.87 | 10.29 | 9.13 | 9.27 | 8.66 | 11.14 | 10.43 | 10.58 | 9.20 |
| UBE2Z | 18 | 9.47 | 11.09 | 11.91 | 9.94 | 8.21 | 8.13 | 8.48 | 11.23 | 11.17 | 10.38 | 10.28 | 10.92 | 8.17 | 9.85 | 10.78 |
| UBE3A | 16 | 8.64 | 10.17 | 10.62 | 10.56 | 10.03 | 10.72 | 9.78 | 8.84 | 10.40 | 10.23 | 9.41 | 10.59 | 10.38 | 9.31 | 10.31 |
| UBE3B | 2 | 9.87 | 10.87 | 9.41 | 10.74 | 11.64 | 10.95 | 8.95 | 7.80 | 10.02 | 9.75 | 10.37 | 10.08 | 11.30 | 8.37 | 9.88 |
| UBE3C | 6 | 10.09 | 10.53 | 9.54 | 10.94 | 7.15 | 7.62 | 10.24 | 10.87 | 11.15 | 11.86 | 10.31 | 10.24 | 7.38 | 10.56 | 11.50 |
| UBE4A | 7 | 10.44 | 9.22 | 9.18 | 9.35 | 10.42 | 9.96 | 9.96 | 10.98 | 10.12 | 10.38 | 9.83 | 9.27 | 10.19 | 10.47 | 10.25 |
| UBE4B | 23 | 10.59 | 10.63 | 10.27 | 10.24 | 9.42 | 9.37 | 9.23 | 8.80 | 10.71 | 10.74 | 10.61 | 10.25 | 9.40 | 9.02 | 10.72 |
| UBFD1 | 14 | 9.80 | 11.50 | 10.81 | 9.93 | 11.17 | 11.97 | 9.70 | 9.18 | 8.03 | 7.91 | 10.65 | 10.37 | 11.57 | 9.44 | 7.97 |
| UBIAD1 | 1 | 11.48 | 6.35 | 9.06 | 10.05 | 8.98 | 7.03 | 11.04 | 9.73 | 12.34 | 13.95 | 8.91 | 9.56 | 8.00 | 10.38 | 13.15 |
| UBL4A | 7 | 9.12 | 9.39 | 9.63 | 8.45 | 10.27 | 10.26 | 10.88 | 13.98 | 9.28 | 8.75 | 9.25 | 9.04 | 10.26 | 12.43 | 9.01 |
| UBL5 | 3 | 9.35 | 10.88 | 9.24 | 9.49 | 11.20 | 12.90 | 9.50 | 10.01 | 8.80 | 8.62 | 10.11 | 9.37 | 12.05 | 9.76 | 8.71 |
| UBL7 | 4 | 10.24 | 11.64 | 10.77 | 11.26 | 9.23 | 10.55 | 9.19 | 8.27 | 9.10 | 9.75 | 10.94 | 11.01 | 9.89 | 8.73 | 9.43 |
| UBLCP1 | 13 | 9.70 | 10.80 | 10.95 | 11.33 | 8.08 | 8.78 | 10.64 | 10.81 | 9.51 | 9.40 | 10.25 | 11.14 | 8.43 | 10.73 | 9.45 |
| UBN1 | 3 | 9.98 | 10.76 | 10.46 | 11.27 | 8.46 | 9.46 | 9.24 | 9.44 | 10.87 | 10.07 | 10.37 | 10.87 | 8.96 | 9.34 | 10.47 |
| UBQLN1 | 13 | 9.80 | 11.50 | 9.07 | 10.00 | 9.52 | 9.79 | 9.31 | 8.40 | 11.26 | 11.34 | 10.65 | 9.54 | 9.66 | 8.86 | 11.30 |
| UBQLN2 | 39 | 9.67 | 11.06 | 8.10 | 8.03 | 11.99 | 12.32 | 10.34 | 9.62 | 9.64 | 9.23 | 10.36 | 8.06 | 12.15 | 9.98 | 9.44 |
| UBQLN4 | 9 | 8.80 | 10.51 | 10.09 | 8.96 | 10.45 | 11.08 | 10.79 | 12.19 | 8.65 | 8.48 | 9.65 | 9.53 | 10.77 | 11.49 | 8.56 |
| UBR1 | 8 | 10.44 | 10.08 | 8.83 | 8.95 | 8.23 | 8.46 | 8.48 | 8.69 | 13.61 | 14.22 | 10.26 | 8.89 | 8.35 | 8.58 | 13.92 |
| UBR2 | 1 | 9.14 | 9.45 | 11.80 | 11.79 | 9.84 | 9.60 | 10.93 | 9.98 | 8.48 | 9.00 | 9.30 | 11.79 | 9.72 | 10.45 | 8.74 |
| UBR3 | 2 | 1.65 | 1.95 | 12.40 | 8.36 | 8.14 | 2.64 | 42.35 | 16.85 | 2.07 | 3.59 | 1.80 | 10.38 | 5.39 | 29.60 | 2.83 |
| UBR4 | 67 | 9.77 | 9.11 | 9.78 | 9.94 | 9.91 | 9.36 | 10.25 | 11.10 | 10.27 | 10.51 | 9.44 | 9.86 | 9.63 | 10.67 | 10.39 |
| UBR5 | 34 | 8.62 | 8.70 | 7.96 | 8.48 | 9.27 | 9.94 | 14.85 | 15.44 | 8.65 | 8.09 | 8.66 | 8.22 | 9.61 | 15.14 | 8.37 |
| UBR7 | 12 | 10.33 | 11.26 | 9.15 | 9.01 | 10.60 | 10.62 | 9.50 | 8.99 | 10.55 | 9.99 | 10.79 | 9.08 | 10.61 | 9.25 | 10.27 |
| UBTD2 | 1 | 9.62 | 8.89 | 11.13 | 13.31 | 9.68 | 10.00 | 10.39 | 7.86 | 9.32 | 9.80 | 9.25 | 12.22 | 9.84 | 9.12 | 9.56 |
| UBTF | 26 | 8.94 | 10.64 | 11.50 | 11.09 | 8.70 | 9.68 | 10.91 | 9.81 | 9.32 | 9.41 | 9.79 | 11.29 | 9.19 | 10.36 | 9.36 |
| UBXN1 | 9 | 10.23 | 10.83 | 10.51 | 10.04 | 10.24 | 9.95 | 9.42 | 8.42 | 10.31 | 10.06 | 10.53 | 10.28 | 10.09 | 8.92 | 10.18 |
| UBXN4 | 4 | 11.59 | 7.96 | 7.48 | 9.62 | 12.04 | 10.12 | 9.99 | 9.94 | 9.84 | 11.43 | 9.77 | 8.55 | 11.08 | 9.96 | 10.63 |
| UBXN6 | 4 | 8.71 | 11.17 | 10.99 | 11.96 | 8.80 | 9.78 | 9.31 | 9.12 | 10.43 | 9.71 | 9.94 | 11.47 | 9.29 | 9.22 | 10.07 |
| UBXN7 | 10 | 10.48 | 10.21 | 11.00 | 9.27 | 10.66 | 10.68 | 11.14 | 9.55 | 8.76 | 8.24 | 10.34 | 10.14 | 10.67 | 10.35 | 8.50 |
| UCHL3 | 9 | 9.99 | 10.48 | 12.74 | 12.08 | 9.42 | 9.49 | 10.74 | 8.07 | 8.62 | 8.37 | 10.23 | 12.41 | 9.45 | 9.40 | 8.50 |
| UCHL5 | 23 | 8.74 | 10.48 | 8.19 | 6.92 | 12.15 | 10.37 | 11.46 | 10.53 | 10.11 | 11.04 | 9.61 | 7.56 | 11.26 | 11.00 | 10.58 |

| | | | | | | | | | | | | | | | | |
|---|---|---|---|---|---|---|---|---|---|---|---|---|---|---|---|---|
| UCK1 | 3 | 8.51 | 9.46 | 12.36 | 8.79 | 12.30 | 10.03 | 9.78 | 7.61 | 9.78 | 11.38 | 8.99 | 10.58 | 11.16 | 8.69 | 10.58 |
| UCK2 | 6 | 8.63 | 9.88 | 11.66 | 8.78 | 10.33 | 9.72 | 10.49 | 9.40 | 10.50 | 10.62 | 9.26 | 10.22 | 10.03 | 9.94 | 10.56 |
| UCKL1 | 4 | 9.87 | 10.56 | 8.96 | 9.03 | 10.15 | 10.57 | 10.93 | 10.83 | 9.77 | 9.32 | 10.22 | 8.99 | 10.36 | 10.88 | 9.55 |
| UEVLD | 4 | 10.31 | 10.69 | 9.17 | 9.67 | 10.49 | 10.65 | 8.42 | 8.28 | 11.23 | 11.10 | 10.50 | 9.42 | 10.57 | 8.35 | 11.16 |
| UFC1 | 7 | 8.75 | 9.31 | 9.85 | 8.53 | 14.83 | 13.56 | 6.68 | 13.43 | 7.86 | 7.20 | 9.03 | 9.19 | 14.20 | 10.05 | 7.53 |
| UFD1L | 18 | 10.38 | 11.40 | 12.10 | 11.24 | 9.18 | 9.19 | 8.39 | 7.54 | 10.73 | 9.85 | 10.89 | 11.67 | 9.19 | 7.96 | 10.29 |
| UFL1 | 26 | 9.36 | 9.60 | 8.83 | 7.95 | 11.59 | 11.81 | 11.03 | 11.52 | 9.18 | 9.15 | 9.48 | 8.39 | 11.70 | 11.27 | 9.16 |
| UFM1 | 2 | 9.19 | 10.09 | 14.13 | 13.84 | 9.68 | 9.58 | 9.45 | 8.37 | 8.11 | 7.58 | 9.64 | 13.99 | 9.63 | 8.91 | 7.84 |
| UFSP1 | 1 | 9.39 | 10.91 | 9.24 | 11.21 | 9.93 | 9.67 | 9.15 | 9.77 | 10.29 | 10.45 | 10.15 | 10.22 | 9.80 | 9.46 | 10.37 |
| UFSP2 | 7 | 8.46 | 9.53 | 10.78 | 9.85 | 10.49 | 11.10 | 9.83 | 11.44 | 9.44 | 9.10 | 9.00 | 10.31 | 10.79 | 10.63 | 9.27 |
| UGDH | 30 | 9.56 | 11.18 | 10.58 | 9.45 | 11.06 | 11.12 | 9.85 | 9.84 | 8.84 | 8.53 | 10.37 | 10.01 | 11.09 | 9.84 | 8.68 |
| UGGT1 | 41 | 11.26 | 7.73 | 8.35 | 10.06 | 11.11 | 10.22 | 11.33 | 10.99 | 8.90 | 10.06 | 9.50 | 9.20 | 10.66 | 11.16 | 9.48 |
| UGGT2 | 3 | 12.36 | 9.36 | 5.79 | 7.18 | 10.37 | 10.48 | 13.80 | 13.51 | 8.71 | 8.43 | 10.86 | 6.49 | 10.43 | 13.66 | 8.57 |
| UGP2 | 31 | 8.90 | 9.43 | 14.11 | 13.99 | 11.02 | 11.11 | 7.94 | 7.46 | 8.22 | 7.81 | 9.17 | 14.05 | 11.06 | 7.70 | 8.02 |
| UGT8 | 3 | 14.38 | 8.75 | 8.20 | 12.65 | 8.52 | 7.45 | 6.80 | 6.06 | 12.24 | 14.96 | 11.57 | 10.42 | 7.98 | 6.43 | 13.60 |
| UHRF1 | 21 | 8.81 | 9.54 | 10.07 | 9.49 | 10.18 | 10.42 | 12.00 | 12.14 | 8.80 | 8.55 | 9.17 | 9.78 | 10.30 | 12.07 | 8.68 |
| UHRF1BP1L | 2 | 8.79 | 10.05 | 9.90 | 8.83 | 13.25 | 13.12 | 10.00 | 7.57 | 8.91 | 9.58 | 9.42 | 9.36 | 13.19 | 8.78 | 9.24 |
| UHRF2 | 3 | 9.93 | 12.35 | 13.03 | 14.40 | 10.62 | 11.92 | 8.48 | 10.46 | 4.02 | 4.78 | 11.14 | 13.71 | 11.27 | 9.47 | 4.40 |
| UIMC1 | 7 | 8.53 | 11.69 | 8.68 | 8.12 | 9.58 | 10.11 | 12.70 | 11.81 | 9.50 | 9.26 | 10.11 | 8.40 | 9.85 | 12.26 | 9.38 |
| ULK3 | 3 | 8.95 | 10.34 | 6.51 | 6.80 | 12.92 | 12.66 | 8.14 | 11.05 | 11.29 | 11.33 | 9.65 | 6.66 | 12.79 | 9.59 | 11.31 |
| UMPS | 33 | 8.60 | 9.68 | 12.05 | 11.46 | 10.13 | 10.18 | 8.88 | 9.26 | 9.84 | 9.93 | 9.14 | 11.75 | 10.15 | 9.07 | 9.88 |
| UNC119B | 3 | 11.02 | 12.04 | 7.46 | 7.71 | 11.94 | 13.26 | 9.39 | 8.45 | 9.76 | 8.97 | 11.53 | 7.59 | 12.60 | 8.92 | 9.36 |
| UNC13B | 2 | 10.54 | 11.03 | 9.59 | 8.80 | 12.04 | 11.92 | 6.32 | 8.32 | 11.34 | 10.10 | 10.79 | 9.19 | 11.98 | 7.32 | 10.72 |
| UNC13D | 14 | 9.30 | 9.77 | 10.60 | 11.01 | 10.79 | 10.41 | 7.31 | 7.05 | 11.93 | 11.83 | 9.54 | 10.81 | 10.60 | 7.18 | 11.88 |
| UNC45A | 34 | 9.39 | 10.02 | 9.44 | 9.49 | 10.34 | 10.57 | 8.44 | 9.45 | 11.50 | 11.36 | 9.70 | 9.47 | 10.46 | 8.95 | 11.43 |
| UNG | 6 | 8.37 | 11.07 | 10.90 | 10.92 | 8.25 | 8.67 | 10.80 | 10.25 | 10.21 | 10.55 | 9.72 | 10.91 | 8.46 | 10.53 | 10.38 |
| UNK | 3 | 10.25 | 11.69 | 12.05 | 10.16 | 11.68 | 10.17 | 10.02 | 9.27 | 7.47 | 7.22 | 10.97 | 11.11 | 10.92 | 9.65 | 7.35 |
| UPF1 | 53 | 9.95 | 10.29 | 9.36 | 8.94 | 9.78 | 9.76 | 11.28 | 11.83 | 9.65 | 9.15 | 10.12 | 9.15 | 9.77 | 11.55 | 9.40 |
| UPF2 | 10 | 9.79 | 10.85 | 9.62 | 9.92 | 9.13 | 9.56 | 10.50 | 10.74 | 9.98 | 9.92 | 10.32 | 9.77 | 9.35 | 10.62 | 9.95 |
| UPF3B | 9 | 7.85 | 9.49 | 10.25 | 9.28 | 10.75 | 10.81 | 12.29 | 10.24 | 9.41 | 9.65 | 8.67 | 9.76 | 10.78 | 11.27 | 9.53 |
| UPP1 | 2 | 9.62 | 10.80 | 17.91 | 18.50 | 4.25 | 4.26 | 8.98 | 7.68 | 9.26 | 8.74 | 10.21 | 18.21 | 4.26 | 8.33 | 9.00 |
| UPRT | 2 | 9.35 | 11.25 | 7.73 | 8.81 | 10.45 | 10.91 | 12.37 | 11.32 | 8.92 | 8.89 | 10.30 | 8.27 | 10.68 | 11.85 | 8.90 |
| UQCC1 | 1 | 13.93 | 12.76 | 9.63 | 12.97 | 9.94 | 8.46 | 15.11 | 13.88 | 1.36 | 1.97 | 13.35 | 11.30 | 9.20 | 14.50 | 1.66 |
| UQCC2 | 4 | 13.51 | 10.80 | 11.45 | 12.00 | 10.66 | 9.47 | 13.76 | 12.31 | 3.15 | 2.89 | 12.15 | 11.73 | 10.06 | 13.03 | 3.02 |
| UQCR10 | 1 | 11.52 | 12.02 | 8.66 | 7.41 | 10.91 | 10.24 | 10.96 | 9.89 | 8.52 | 9.87 | 11.77 | 8.03 | 10.58 | 10.42 | 9.20 |
| UQCRB | 8 | 12.21 | 10.23 | 9.68 | 10.86 | 12.27 | 11.98 | 12.36 | 10.66 | 4.88 | 4.86 | 11.22 | 10.27 | 12.13 | 11.51 | 4.87 |
| UQCRC1 | 23 | 11.32 | 10.17 | 9.98 | 10.70 | 12.29 | 10.24 | 10.91 | 10.99 | 6.49 | 6.92 | 10.74 | 10.34 | 11.26 | 10.95 | 6.70 |
| UQCRC2 | 21 | 12.52 | 10.61 | 9.45 | 11.15 | 12.22 | 11.52 | 11.06 | 10.30 | 5.44 | 5.72 | 11.56 | 10.30 | 11.87 | 10.68 | 5.58 |
| UQCRFS1 | 11 | 12.29 | 9.27 | 11.06 | 9.21 | 11.27 | 9.77 | 12.78 | 12.18 | 6.48 | 5.69 | 10.78 | 10.14 | 10.52 | 12.48 | 6.09 |
| UQCRQ | 2 | 12.91 | 10.11 | 9.20 | 12.09 | 10.84 | 10.09 | 10.94 | 10.67 | 6.29 | 6.88 | 11.51 | 10.64 | 10.46 | 10.81 | 6.58 |
| URB1 | 10 | 10.22 | 9.67 | 9.00 | 9.54 | 10.16 | 10.06 | 10.50 | 10.52 | 9.91 | 10.44 | 9.94 | 9.27 | 10.11 | 10.51 | 10.17 |
| URB2 | 9 | 10.04 | 9.88 | 9.19 | 10.13 | 10.11 | 10.04 | 10.28 | 10.67 | 9.63 | 10.01 | 9.96 | 9.66 | 10.08 | 10.48 | 9.82 |

| | | | | | | | | | | | | | | | | |
|---|---|---|---|---|---|---|---|---|---|---|---|---|---|---|---|---|
| URGCP | 6 | 10.74 | 11.15 | 10.29 | 10.46 | 7.97 | 8.90 | 9.56 | 9.29 | 10.35 | 11.28 | 10.94 | 10.38 | 8.44 | 9.43 | 10.81 |
| URI1 | 5 | 8.07 | 8.89 | 13.57 | 12.34 | 8.75 | 8.22 | 14.33 | 12.21 | 6.60 | 7.03 | 8.48 | 12.96 | 8.48 | 13.27 | 6.81 |
| URM1 | 3 | 12.75 | 11.71 | 9.60 | 9.66 | 9.00 | 11.23 | 8.46 | 4.47 | 11.58 | 11.54 | 12.23 | 9.63 | 10.11 | 6.46 | 11.56 |
| UROD | 12 | 9.61 | 11.32 | 10.79 | 9.64 | 7.68 | 7.60 | 10.19 | 10.32 | 11.24 | 11.61 | 10.46 | 10.22 | 7.64 | 10.26 | 11.43 |
| UROS | 1 | 9.30 | 11.18 | 8.40 | 8.65 | 12.60 | 13.31 | 9.40 | 7.40 | 10.79 | 8.95 | 10.24 | 8.53 | 12.96 | 8.40 | 9.87 |
| USE1 | 1 | 10.35 | 5.25 | 7.76 | 10.94 | 7.44 | 7.19 | 11.83 | 11.60 | 12.52 | 15.12 | 7.80 | 9.35 | 7.31 | 11.72 | 13.82 |
| USF1 | 1 | 7.99 | 14.06 | 8.27 | 8.97 | 12.45 | 13.11 | 12.04 | 10.94 | 6.36 | 5.82 | 11.02 | 8.62 | 12.78 | 11.49 | 6.09 |
| USF2 | 3 | 8.70 | 10.33 | 8.32 | 7.31 | 9.99 | 8.64 | 11.84 | 16.79 | 8.69 | 9.38 | 9.52 | 7.82 | 9.31 | 14.32 | 9.04 |
| USMG5 | 1 | 10.67 | 10.19 | 9.89 | 9.30 | 10.53 | 9.72 | 9.84 | 8.76 | 9.64 | 11.47 | 10.43 | 9.59 | 10.12 | 9.30 | 10.56 |
| USO1 | 39 | 9.24 | 9.88 | 10.94 | 10.95 | 10.16 | 10.75 | 10.07 | 9.58 | 9.30 | 9.14 | 9.56 | 10.95 | 10.45 | 9.82 | 9.22 |
| USP1 | 11 | 10.59 | 10.95 | 8.97 | 9.05 | 9.18 | 9.22 | 11.18 | 10.00 | 10.30 | 10.56 | 10.77 | 9.01 | 9.20 | 10.59 | 10.43 |
| USP10 | 23 | 10.60 | 10.81 | 9.48 | 8.68 | 10.16 | 10.31 | 9.84 | 9.03 | 10.78 | 10.30 | 10.71 | 9.08 | 10.24 | 9.44 | 10.54 |
| USP11 | 6 | 9.04 | 9.77 | 8.99 | 8.59 | 11.66 | 12.28 | 10.92 | 10.75 | 9.29 | 8.71 | 9.40 | 8.79 | 11.97 | 10.83 | 9.00 |
| USP13 | 11 | 11.18 | 12.29 | 10.15 | 9.72 | 7.50 | 8.04 | 9.19 | 8.36 | 11.87 | 11.70 | 11.74 | 9.94 | 7.77 | 8.77 | 11.79 |
| USP14 | 39 | 9.91 | 10.73 | 11.76 | 11.09 | 10.82 | 10.61 | 7.19 | 8.37 | 10.00 | 9.52 | 10.32 | 11.43 | 10.71 | 7.78 | 9.76 |
| USP15 | 31 | 9.15 | 9.40 | 11.82 | 10.85 | 8.67 | 8.52 | 10.01 | 10.51 | 10.87 | 10.20 | 9.28 | 11.33 | 8.60 | 10.26 | 10.53 |
| USP16 | 2 | 10.57 | 11.25 | 9.78 | 8.84 | 10.33 | 10.22 | 9.67 | 9.24 | 11.05 | 9.05 | 10.91 | 9.31 | 10.28 | 9.45 | 10.05 |
| USP19 | 10 | 9.05 | 9.83 | 10.36 | 9.25 | 10.20 | 10.38 | 10.95 | 10.27 | 10.31 | 9.39 | 9.44 | 9.81 | 10.29 | 10.61 | 9.85 |
| USP22 | 3 | 7.48 | 11.09 | 11.54 | 10.02 | 8.01 | 8.10 | 10.07 | 10.64 | 11.97 | 11.06 | 9.29 | 10.78 | 8.06 | 10.36 | 11.51 |
| USP24 | 18 | 10.40 | 10.38 | 7.03 | 12.91 | 10.52 | 10.77 | 10.52 | 10.53 | 8.63 | 8.31 | 10.39 | 9.97 | 10.64 | 10.52 | 8.47 |
| USP25 | 2 | 9.93 | 11.55 | 10.73 | 9.13 | 8.85 | 9.35 | 9.96 | 9.18 | 10.87 | 10.46 | 10.74 | 9.93 | 9.10 | 9.57 | 10.67 |
| USP28 | 3 | 9.69 | 9.67 | 9.34 | 9.28 | 8.56 | 9.34 | 11.69 | 12.13 | 10.36 | 9.94 | 9.68 | 9.31 | 8.95 | 11.91 | 10.15 |
| USP3 | 4 | 8.25 | 9.31 | 9.36 | 9.67 | 10.02 | 10.35 | 10.00 | 11.38 | 10.71 | 10.95 | 8.78 | 9.52 | 10.18 | 10.69 | 10.83 |
| USP32 | 3 | 10.40 | 10.81 | 10.71 | 8.22 | 10.97 | 9.44 | 8.05 | 7.36 | 11.69 | 12.35 | 10.61 | 9.47 | 10.20 | 7.70 | 12.02 |
| USP34 | 2 | 9.03 | 9.80 | 8.54 | 9.93 | 9.74 | 10.29 | 11.08 | 13.24 | 8.53 | 9.81 | 9.41 | 9.24 | 10.02 | 12.16 | 9.17 |
| USP36 | 6 | 9.89 | 10.10 | 10.29 | 10.06 | 8.33 | 8.31 | 11.74 | 10.32 | 10.80 | 10.14 | 10.00 | 10.18 | 8.32 | 11.03 | 10.47 |
| USP37 | 2 | 9.37 | 10.63 | 8.80 | 8.59 | 12.54 | 12.24 | 10.63 | 10.05 | 8.52 | 8.63 | 10.00 | 8.70 | 12.39 | 10.34 | 8.57 |
| USP39 | 17 | 9.74 | 9.33 | 10.56 | 10.88 | 9.30 | 9.75 | 10.92 | 10.92 | 9.26 | 9.34 | 9.54 | 10.72 | 9.52 | 10.92 | 9.30 |
| USP4 | 9 | 10.04 | 10.57 | 10.18 | 9.95 | 10.52 | 10.90 | 8.62 | 9.59 | 9.80 | 9.84 | 10.30 | 10.07 | 10.71 | 9.10 | 9.82 |
| USP42 | 1 | 9.09 | 10.34 | 8.55 | 8.14 | 11.33 | 10.79 | 10.02 | 12.38 | 9.53 | 9.83 | 9.72 | 8.35 | 11.06 | 11.20 | 9.68 |
| USP47 | 27 | 9.44 | 10.45 | 10.29 | 10.38 | 11.73 | 12.85 | 9.01 | 9.31 | 8.21 | 8.33 | 9.95 | 10.34 | 12.29 | 9.16 | 8.27 |
| USP48 | 17 | 11.69 | 11.37 | 9.20 | 8.57 | 9.50 | 8.91 | 10.18 | 9.66 | 10.54 | 10.39 | 11.53 | 8.89 | 9.20 | 9.92 | 10.46 |
| USP5 | 45 | 9.74 | 10.47 | 12.46 | 11.31 | 9.93 | 10.00 | 8.32 | 7.53 | 10.40 | 9.82 | 10.11 | 11.89 | 9.97 | 7.92 | 10.11 |
| USP6NL | 1 | 8.97 | 9.67 | 9.72 | 10.42 | 13.21 | 13.09 | 8.45 | 8.68 | 8.82 | 8.98 | 9.32 | 10.07 | 13.15 | 8.56 | 8.90 |
| USP7 | 39 | 8.93 | 9.28 | 10.59 | 9.45 | 10.26 | 10.19 | 11.58 | 11.32 | 9.33 | 9.06 | 9.11 | 10.02 | 10.22 | 11.45 | 9.20 |
| USP8 | 18 | 9.28 | 10.06 | 10.45 | 10.04 | 9.93 | 10.37 | 10.03 | 8.86 | 10.57 | 10.41 | 9.67 | 10.24 | 10.15 | 9.45 | 10.49 |
| USP9X | 46 | 9.50 | 9.33 | 9.45 | 9.54 | 11.28 | 11.22 | 10.67 | 10.82 | 9.17 | 9.02 | 9.42 | 9.49 | 11.25 | 10.75 | 9.09 |
| UTP11L | 6 | 11.00 | 9.40 | 10.38 | 11.85 | 7.70 | 7.32 | 10.60 | 9.96 | 9.27 | 12.53 | 10.20 | 11.11 | 7.51 | 10.28 | 10.90 |
| UTP14A | 27 | 9.13 | 10.34 | 9.95 | 9.74 | 8.95 | 8.92 | 12.63 | 12.37 | 9.02 | 8.95 | 9.73 | 9.85 | 8.93 | 12.50 | 8.99 |
| UTP15 | 16 | 8.53 | 9.54 | 10.69 | 10.83 | 8.43 | 8.41 | 11.75 | 12.06 | 9.95 | 9.81 | 9.03 | 10.76 | 8.42 | 11.91 | 9.88 |
| UTP18 | 15 | 9.40 | 10.04 | 10.40 | 10.86 | 8.03 | 8.39 | 11.34 | 10.98 | 10.12 | 10.43 | 9.72 | 10.63 | 8.21 | 11.16 | 10.27 |
| UTP20 | 36 | 9.63 | 8.68 | 10.14 | 11.61 | 7.57 | 7.35 | 11.06 | 11.65 | 10.53 | 11.78 | 9.16 | 10.87 | 7.46 | 11.36 | 11.15 |

| | | | | | | | | | | | | | | | | |
|---|---|---|---|---|---|---|---|---|---|---|---|---|---|---|---|---|
| UTP23 | 1 | 9.15 | 9.76 | 10.11 | 10.72 | 7.82 | 8.29 | 13.28 | 13.13 | 9.51 | 8.22 | 9.46 | 10.41 | 8.05 | 13.21 | 8.87 |
| UTP3 | 8 | 8.94 | 9.79 | 11.44 | 10.82 | 8.49 | 8.45 | 11.61 | 10.16 | 10.06 | 10.24 | 9.36 | 11.13 | 8.47 | 10.88 | 10.15 |
| UTP6 | 13 | 9.64 | 9.94 | 9.71 | 10.32 | 8.28 | 8.82 | 11.26 | 11.48 | 10.20 | 10.34 | 9.79 | 10.02 | 8.55 | 11.37 | 10.27 |
| UTRN | 27 | 9.07 | 9.56 | 10.76 | 11.56 | 11.84 | 12.41 | 7.19 | 6.74 | 10.54 | 10.32 | 9.32 | 11.16 | 12.13 | 6.97 | 10.43 |
| UVRAG | 2 | 10.04 | 11.95 | 9.94 | 10.69 | 9.79 | 10.68 | 9.25 | 9.21 | 9.18 | 9.25 | 11.00 | 10.32 | 10.24 | 9.23 | 9.21 |
| UXS1 | 2 | 9.39 | 7.36 | 10.25 | 11.30 | 9.86 | 9.26 | 11.73 | 9.37 | 10.57 | 10.90 | 8.38 | 10.78 | 9.56 | 10.55 | 10.74 |
| UXT | 4 | 8.38 | 8.27 | 12.33 | 11.33 | 9.29 | 9.15 | 13.56 | 12.62 | 7.99 | 7.08 | 8.32 | 11.83 | 9.22 | 13.09 | 7.53 |
| VAC14 | 7 | 10.95 | 10.52 | 7.67 | 7.30 | 12.20 | 11.50 | 11.64 | 11.72 | 8.47 | 8.03 | 10.74 | 7.49 | 11.85 | 11.68 | 8.25 |
| VAMP2 | 4 | 9.31 | 9.00 | 11.83 | 14.94 | 9.11 | 8.32 | 9.67 | 9.68 | 8.37 | 9.76 | 9.15 | 13.39 | 8.72 | 9.68 | 9.06 |
| VAMP3 | 2 | 11.13 | 10.16 | 10.66 | 11.54 | 8.95 | 7.44 | 11.29 | 10.86 | 9.19 | 8.78 | 10.65 | 11.10 | 8.20 | 11.08 | 8.98 |
| VAMP4 | 1 | 10.86 | 5.72 | 9.58 | 10.47 | 16.24 | 12.97 | 11.26 | 8.15 | 7.53 | 7.22 | 8.29 | 10.02 | 14.61 | 9.70 | 7.38 |
| VAMP7 | 7 | 10.38 | 7.67 | 7.29 | 9.18 | 12.15 | 11.69 | 11.51 | 11.61 | 9.01 | 9.52 | 9.02 | 8.23 | 11.92 | 11.56 | 9.27 |
| VANGL1 | 2 | 11.56 | 8.71 | 7.08 | 11.24 | 10.30 | 9.14 | 12.15 | 9.58 | 8.73 | 11.51 | 10.14 | 9.16 | 9.72 | 10.86 | 10.12 |
| VAPA | 28 | 12.25 | 6.86 | 8.35 | 10.58 | 10.43 | 8.39 | 8.32 | 9.07 | 11.81 | 13.93 | 9.55 | 9.47 | 9.41 | 8.70 | 12.87 |
| VAPB | 14 | 12.57 | 6.49 | 8.48 | 12.44 | 7.08 | 6.42 | 7.49 | 8.85 | 13.14 | 17.06 | 9.53 | 10.46 | 6.75 | 8.17 | 15.10 |
| VARS | 61 | 9.40 | 10.00 | 10.03 | 8.97 | 10.41 | 10.35 | 9.22 | 10.32 | 10.76 | 10.54 | 9.70 | 9.50 | 10.38 | 9.77 | 10.65 |
| VARS2 | 1 | 9.93 | 8.70 | 8.01 | 8.60 | 10.84 | 9.97 | 10.36 | 9.06 | 12.02 | 12.51 | 9.32 | 8.30 | 10.41 | 9.71 | 12.26 |
| VASH2 | 1 | 8.60 | 10.73 | 8.79 | 11.46 | 7.57 | 9.32 | 11.35 | 10.59 | 11.14 | 10.46 | 9.67 | 10.12 | 8.44 | 10.97 | 10.80 |
| VASN | 8 | 13.64 | 12.77 | 8.27 | 9.93 | 8.87 | 9.23 | 5.90 | 5.67 | 12.69 | 13.03 | 13.21 | 9.10 | 9.05 | 5.79 | 12.86 |
| VASP | 17 | 8.14 | 9.02 | 16.79 | 15.46 | 6.61 | 6.07 | 7.86 | 7.54 | 11.66 | 10.84 | 8.58 | 16.13 | 6.34 | 7.70 | 11.25 |
| VAT1 | 21 | 9.05 | 9.67 | 14.83 | 12.25 | 9.66 | 9.18 | 7.90 | 7.56 | 10.14 | 9.76 | 9.36 | 13.54 | 9.42 | 7.73 | 9.95 |
| VAT1L | 1 | 7.52 | 8.27 | 11.47 | 10.74 | 13.01 | 11.23 | 9.22 | 8.10 | 10.04 | 10.40 | 7.90 | 11.10 | 12.12 | 8.66 | 10.22 |
| VAV2 | 11 | 10.32 | 11.39 | 7.70 | 7.58 | 10.72 | 10.80 | 10.44 | 11.50 | 10.00 | 9.54 | 10.86 | 7.64 | 10.76 | 10.97 | 9.77 |
| VBP1 | 21 | 8.92 | 9.60 | 9.09 | 8.15 | 11.57 | 11.48 | 10.76 | 14.09 | 8.46 | 7.89 | 9.26 | 8.62 | 11.53 | 12.42 | 8.17 |
| VCL | 202 | 9.63 | 10.92 | 13.34 | 13.08 | 7.97 | 7.99 | 6.23 | 7.04 | 12.17 | 11.63 | 10.27 | 13.21 | 7.98 | 6.64 | 11.90 |
| VCP | 122 | 10.19 | 11.46 | 10.43 | 10.01 | 8.95 | 9.29 | 7.32 | 9.61 | 11.49 | 11.25 | 10.83 | 10.22 | 9.12 | 8.47 | 11.37 |
| VCPIP1 | 14 | 9.36 | 10.87 | 9.87 | 10.50 | 10.15 | 10.90 | 9.96 | 8.94 | 10.03 | 9.43 | 10.11 | 10.18 | 10.53 | 9.45 | 9.73 |
| VDAC1 | 52 | 10.59 | 6.78 | 9.19 | 11.25 | 9.12 | 8.02 | 11.16 | 12.88 | 9.52 | 11.49 | 8.69 | 10.22 | 8.57 | 12.02 | 10.51 |
| VDAC2 | 22 | 12.96 | 6.42 | 9.62 | 10.08 | 9.67 | 8.04 | 9.59 | 9.45 | 11.13 | 13.03 | 9.69 | 9.85 | 8.86 | 9.52 | 12.08 |
| VDAC3 | 18 | 12.86 | 7.21 | 8.74 | 11.05 | 8.90 | 8.30 | 11.79 | 10.75 | 9.37 | 11.03 | 10.04 | 9.89 | 8.60 | 11.27 | 10.20 |
| VEZT | 1 | 15.73 | 9.12 | 8.35 | 13.44 | 11.42 | 8.46 | 8.69 | 8.22 | 8.33 | 8.25 | 12.42 | 10.90 | 9.94 | 8.45 | 8.29 |
| VIM | 283 | 8.32 | 10.38 | 11.01 | 11.36 | 8.15 | 8.51 | 9.48 | 10.03 | 11.60 | 11.16 | 9.35 | 11.18 | 8.33 | 9.76 | 11.38 |
| VIMP | 3 | 13.09 | 6.06 | 7.13 | 9.94 | 9.94 | 7.94 | 8.51 | 6.73 | 14.14 | 16.51 | 9.58 | 8.54 | 8.94 | 7.62 | 15.33 |
| VIPAS39 | 7 | 9.83 | 10.74 | 8.90 | 9.12 | 10.45 | 11.42 | 9.71 | 9.27 | 9.85 | 10.71 | 10.28 | 9.01 | 10.94 | 9.49 | 10.28 |
| VKORC1 | 2 | 14.86 | 6.87 | 6.76 | 10.35 | 8.64 | 6.78 | 10.09 | 8.39 | 12.63 | 14.63 | 10.86 | 8.55 | 7.71 | 9.24 | 13.63 |
| VKORC1L1 | 1 | 13.74 | 5.72 | 7.84 | 10.84 | 7.72 | 8.46 | 9.06 | 7.01 | 12.37 | 17.25 | 9.73 | 9.34 | 8.09 | 8.03 | 14.81 |
| VLDLR | 1 | 9.54 | 6.55 | 14.85 | 21.84 | 6.74 | 5.91 | 2.91 | 2.17 | 14.00 | 15.50 | 8.05 | 18.34 | 6.32 | 2.54 | 14.75 |
| VMA21 | 2 | 9.55 | 7.78 | 7.37 | 9.58 | 10.48 | 8.86 | 12.41 | 11.72 | 9.95 | 12.30 | 8.66 | 8.48 | 9.67 | 12.07 | 11.13 |
| VPRBP | 11 | 8.66 | 9.49 | 9.71 | 10.22 | 9.23 | 9.33 | 11.65 | 10.78 | 10.56 | 10.36 | 9.07 | 9.97 | 9.28 | 11.21 | 10.46 |
| VPS11 | 10 | 10.03 | 10.68 | 9.93 | 10.11 | 12.72 | 13.23 | 8.12 | 7.88 | 8.75 | 8.56 | 10.35 | 10.02 | 12.98 | 8.00 | 8.65 |
| VPS13A | 9 | 10.98 | 11.33 | 12.07 | 12.45 | 11.53 | 11.90 | 10.19 | 9.55 | 5.04 | 4.96 | 11.15 | 12.26 | 11.72 | 9.87 | 5.00 |
| VPS13B | 2 | 10.05 | 10.75 | 9.62 | 11.56 | 10.34 | 10.63 | 10.30 | 10.60 | 8.15 | 8.00 | 10.40 | 10.59 | 10.49 | 10.45 | 8.08 |

| | | | | | | | | | | | | | | | | |
|---|---|---|---|---|---|---|---|---|---|---|---|---|---|---|---|---|
| VPS13C | 19 | 10.53 | 11.67 | 9.83 | 9.39 | 10.53 | 10.57 | 6.67 | 6.06 | 12.56 | 12.20 | 11.10 | 9.61 | 10.55 | 6.37 | 12.38 |
| VPS13D | 2 | 10.17 | 12.19 | 8.08 | 7.58 | 10.71 | 9.36 | 11.44 | 10.06 | 9.88 | 10.54 | 11.18 | 7.83 | 10.03 | 10.75 | 10.21 |
| VPS16 | 5 | 9.74 | 9.56 | 10.64 | 10.44 | 12.45 | 12.13 | 8.87 | 7.94 | 9.57 | 8.65 | 9.65 | 10.54 | 12.29 | 8.41 | 9.11 |
| VPS18 | 7 | 10.66 | 11.01 | 9.34 | 9.82 | 11.65 | 12.22 | 8.49 | 7.72 | 9.67 | 9.41 | 10.83 | 9.58 | 11.94 | 8.11 | 9.54 |
| VPS25 | 11 | 10.73 | 11.61 | 9.74 | 9.53 | 12.83 | 13.65 | 7.16 | 7.22 | 9.07 | 8.47 | 11.17 | 9.63 | 13.24 | 7.19 | 8.77 |
| VPS26A | 16 | 10.60 | 10.25 | 8.97 | 6.29 | 13.10 | 12.75 | 9.47 | 8.18 | 10.40 | 10.00 | 10.42 | 7.63 | 12.93 | 8.82 | 10.20 |
| VPS26B | 8 | 9.90 | 11.39 | 8.63 | 8.03 | 11.70 | 11.81 | 9.33 | 9.26 | 10.13 | 9.84 | 10.64 | 8.33 | 11.75 | 9.29 | 9.98 |
| VPS28 | 9 | 8.89 | 10.24 | 11.07 | 10.87 | 9.89 | 9.97 | 10.10 | 9.76 | 9.56 | 9.64 | 9.57 | 10.97 | 9.93 | 9.93 | 9.60 |
| VPS29 | 14 | 9.65 | 12.27 | 10.01 | 8.15 | 12.26 | 11.28 | 8.76 | 9.37 | 9.15 | 9.10 | 10.96 | 9.08 | 11.77 | 9.07 | 9.12 |
| VPS33A | 7 | 10.07 | 10.06 | 10.69 | 8.63 | 12.99 | 11.69 | 9.01 | 7.95 | 9.44 | 9.48 | 10.06 | 9.66 | 12.34 | 8.48 | 9.46 |
| VPS33B | 13 | 9.85 | 10.56 | 8.77 | 8.55 | 10.37 | 10.33 | 10.34 | 10.62 | 10.41 | 10.20 | 10.21 | 8.66 | 10.35 | 10.48 | 10.30 |
| VPS35 | 22 | 9.80 | 10.08 | 8.10 | 6.82 | 12.09 | 11.17 | 9.17 | 12.57 | 10.40 | 9.80 | 9.94 | 7.46 | 11.63 | 10.87 | 10.10 |
| VPS36 | 11 | 11.09 | 11.47 | 8.98 | 8.41 | 14.37 | 14.26 | 8.24 | 7.75 | 8.01 | 7.41 | 11.28 | 8.69 | 14.32 | 8.00 | 7.71 |
| VPS37A | 3 | 10.12 | 11.62 | 7.69 | 7.78 | 10.90 | 10.11 | 11.56 | 10.31 | 10.52 | 9.39 | 10.87 | 7.74 | 10.51 | 10.93 | 9.95 |
| VPS37B | 2 | 8.29 | 8.81 | 14.13 | 12.34 | 9.21 | 8.33 | 10.75 | 8.55 | 9.19 | 10.39 | 8.55 | 13.23 | 8.77 | 9.65 | 9.79 |
| VPS37C | 1 | 9.75 | 10.52 | 9.67 | 10.67 | 9.36 | 9.88 | 9.16 | 11.75 | 8.56 | 10.68 | 10.13 | 10.17 | 9.62 | 10.45 | 9.62 |
| VPS39 | 7 | 10.63 | 9.87 | 9.96 | 9.29 | 12.80 | 13.14 | 8.39 | 8.11 | 9.33 | 8.48 | 10.25 | 9.62 | 12.97 | 8.25 | 8.91 |
| VPS41 | 3 | 10.46 | 9.21 | 11.48 | 8.40 | 13.57 | 12.38 | 8.32 | 8.81 | 9.00 | 8.37 | 9.83 | 9.94 | 12.97 | 8.57 | 8.68 |
| VPS45 | 12 | 10.36 | 10.53 | 10.32 | 9.68 | 11.38 | 11.19 | 8.96 | 8.35 | 9.73 | 9.50 | 10.44 | 10.00 | 11.28 | 8.66 | 9.62 |
| VPS4A | 12 | 9.64 | 11.02 | 7.85 | 7.63 | 12.48 | 12.66 | 10.84 | 10.13 | 8.91 | 8.86 | 10.33 | 7.74 | 12.57 | 10.48 | 8.88 |
| VPS4B | 21 | 9.94 | 10.66 | 11.94 | 10.95 | 11.16 | 11.12 | 7.78 | 7.08 | 9.93 | 9.44 | 10.30 | 11.45 | 11.14 | 7.43 | 9.68 |
| VPS51 | 10 | 9.92 | 10.31 | 10.76 | 10.65 | 10.57 | 10.68 | 9.66 | 9.18 | 9.08 | 9.19 | 10.12 | 10.70 | 10.62 | 9.42 | 9.14 |
| VPS52 | 11 | 10.43 | 10.61 | 11.33 | 10.52 | 10.73 | 10.29 | 9.54 | 8.85 | 8.72 | 8.98 | 10.52 | 10.92 | 10.51 | 9.20 | 8.85 |
| VPS53 | 15 | 10.69 | 11.61 | 11.11 | 9.70 | 10.05 | 9.82 | 8.61 | 9.15 | 9.55 | 9.71 | 11.15 | 10.40 | 9.94 | 8.88 | 9.63 |
| VPS72 | 2 | 9.63 | 10.42 | 10.78 | 10.96 | 10.45 | 10.91 | 10.08 | 9.26 | 9.49 | 8.01 | 10.03 | 10.87 | 10.68 | 9.67 | 8.75 |
| VPS8 | 1 | 12.11 | 12.78 | 8.77 | 12.99 | 10.46 | 13.38 | 8.55 | 8.43 | 7.10 | 5.44 | 12.44 | 10.88 | 11.92 | 8.49 | 6.27 |
| VRK1 | 12 | 9.07 | 10.07 | 8.21 | 8.21 | 8.33 | 8.96 | 13.79 | 13.00 | 9.94 | 10.41 | 9.57 | 8.21 | 8.64 | 13.40 | 10.18 |
| VRK2 | 1 | 9.64 | 8.77 | 10.74 | 12.64 | 10.59 | 9.33 | 8.45 | 8.51 | 10.15 | 11.18 | 9.21 | 11.69 | 9.96 | 8.48 | 10.66 |
| VTA1 | 13 | 9.81 | 10.40 | 11.01 | 10.59 | 10.94 | 11.35 | 9.56 | 9.69 | 8.59 | 8.05 | 10.10 | 10.80 | 11.15 | 9.63 | 8.32 |
| VTI1A | 1 | 12.61 | 9.31 | 9.72 | 11.36 | 10.87 | 10.75 | 10.92 | 10.38 | 7.72 | 6.36 | 10.96 | 10.54 | 10.81 | 10.65 | 7.04 |
| VTI1B | 2 | 9.81 | 7.70 | 9.90 | 11.98 | 11.16 | 10.73 | 10.83 | 8.85 | 9.61 | 9.44 | 8.76 | 10.94 | 10.94 | 9.84 | 9.52 |
| VTN | 6 | 3.38 | 3.49 | 12.10 | 12.33 | 4.76 | 4.68 | 29.69 | 23.65 | 3.04 | 2.88 | 3.44 | 12.22 | 4.72 | 26.67 | 2.96 |
| VWA8 | 15 | 11.63 | 9.91 | 8.76 | 10.97 | 11.44 | 10.96 | 9.03 | 8.45 | 9.20 | 9.64 | 10.77 | 9.86 | 11.20 | 8.74 | 9.42 |
| VWA9 | 8 | 9.49 | 10.01 | 9.52 | 9.41 | 10.30 | 11.00 | 10.00 | 9.54 | 10.61 | 10.12 | 9.75 | 9.47 | 10.65 | 9.77 | 10.36 |
| WAC | 12 | 8.94 | 9.06 | 11.17 | 10.06 | 10.22 | 10.15 | 10.88 | 10.16 | 9.85 | 9.51 | 9.00 | 10.62 | 10.18 | 10.52 | 9.68 |
| WAPAL | 19 | 9.21 | 9.60 | 10.76 | 9.94 | 9.57 | 9.71 | 11.19 | 10.34 | 10.16 | 9.50 | 9.41 | 10.35 | 9.64 | 10.77 | 9.83 |
| WARS | 28 | 8.55 | 10.10 | 9.28 | 8.84 | 9.13 | 9.65 | 7.87 | 14.32 | 10.89 | 11.38 | 9.32 | 9.06 | 9.39 | 11.10 | 11.13 |
| WARS2 | 7 | 11.37 | 9.88 | 10.40 | 10.83 | 8.81 | 8.46 | 11.12 | 10.22 | 9.84 | 9.07 | 10.63 | 10.61 | 8.64 | 10.67 | 9.45 |
| WASF1 | 2 | 9.60 | 9.51 | 13.26 | 13.46 | 10.56 | 10.91 | 7.51 | 6.03 | 10.17 | 8.99 | 9.55 | 13.36 | 10.74 | 6.77 | 9.58 |
| WASF2 | 11 | 8.39 | 10.72 | 11.49 | 11.33 | 10.34 | 10.38 | 8.97 | 7.68 | 10.14 | 10.57 | 9.55 | 11.41 | 10.36 | 8.32 | 10.36 |
| WASH2P | 7 | 10.27 | 11.13 | 9.81 | 9.53 | 11.37 | 11.88 | 8.30 | 8.20 | 9.82 | 9.69 | 10.70 | 9.67 | 11.63 | 8.25 | 9.76 |
| WASL | 9 | 9.03 | 10.52 | 11.82 | 11.30 | 9.19 | 9.32 | 10.21 | 9.77 | 9.47 | 9.36 | 9.78 | 11.56 | 9.25 | 9.99 | 9.42 |

| | | | | | | | | | | | | | | | | |
|---|---|---|---|---|---|---|---|---|---|---|---|---|---|---|---|---|
| WBP11 | 31 | 9.16 | 10.00 | 11.75 | 10.53 | 9.35 | 9.07 | 11.30 | 9.67 | 9.99 | 9.20 | 9.58 | 11.14 | 9.21 | 10.48 | 9.59 |
| WBP2 | 5 | 11.12 | 11.82 | 9.57 | 8.77 | 12.86 | 12.12 | 7.13 | 6.13 | 10.42 | 10.05 | 11.47 | 9.17 | 12.49 | 6.63 | 10.23 |
| WBP4 | 3 | 11.04 | 13.05 | 11.51 | 11.42 | 9.93 | 11.06 | 8.80 | 7.85 | 7.99 | 7.34 | 12.04 | 11.47 | 10.50 | 8.33 | 7.67 |
| WBSCR16 | 3 | 11.18 | 9.40 | 9.79 | 9.48 | 9.90 | 9.08 | 10.54 | 10.10 | 10.17 | 10.34 | 10.29 | 9.64 | 9.49 | 10.32 | 10.26 |
| WBSCR22 | 5 | 10.09 | 10.12 | 9.87 | 8.37 | 9.35 | 9.67 | 11.38 | 10.73 | 10.17 | 10.24 | 10.10 | 9.12 | 9.51 | 11.06 | 10.21 |
| WDFY1 | 3 | 9.12 | 10.91 | 10.49 | 10.51 | 12.14 | 12.72 | 6.85 | 6.28 | 9.91 | 11.07 | 10.01 | 10.50 | 12.43 | 6.56 | 10.49 |
| WDFY3 | 2 | 8.49 | 7.59 | 11.63 | 12.62 | 10.08 | 10.56 | 9.15 | 10.17 | 9.76 | 9.95 | 8.04 | 12.12 | 10.32 | 9.66 | 9.85 |
| WDHD1 | 32 | 9.50 | 10.25 | 10.84 | 10.28 | 9.10 | 9.20 | 10.10 | 9.47 | 10.83 | 10.42 | 9.88 | 10.56 | 9.15 | 9.78 | 10.62 |
| WDR1 | 45 | 9.70 | 9.94 | 13.37 | 11.58 | 9.56 | 9.05 | 5.90 | 8.18 | 11.75 | 10.97 | 9.82 | 12.48 | 9.31 | 7.04 | 11.36 |
| WDR11 | 9 | 11.60 | 12.44 | 8.50 | 8.03 | 9.53 | 11.01 | 10.06 | 9.64 | 9.95 | 9.25 | 12.02 | 8.26 | 10.27 | 9.85 | 9.60 |
| WDR12 | 14 | 9.85 | 9.96 | 11.44 | 9.36 | 8.99 | 8.92 | 11.45 | 10.04 | 10.31 | 9.69 | 9.90 | 10.40 | 8.95 | 10.74 | 10.00 |
| WDR13 | 2 | 9.10 | 10.16 | 9.80 | 9.86 | 11.87 | 12.20 | 10.11 | 8.66 | 9.27 | 8.97 | 9.63 | 9.83 | 12.04 | 9.39 | 9.12 |
| WDR18 | 7 | 9.28 | 10.12 | 10.32 | 10.25 | 9.88 | 9.85 | 9.62 | 9.28 | 10.76 | 10.65 | 9.70 | 10.29 | 9.87 | 9.45 | 10.70 |
| WDR19 | 2 | 8.79 | 10.52 | 10.72 | 10.14 | 10.50 | 9.64 | 9.31 | 8.38 | 11.64 | 10.37 | 9.65 | 10.43 | 10.07 | 8.84 | 11.00 |
| WDR25 | 1 | 8.97 | 12.27 | 10.99 | 9.16 | 9.93 | 10.15 | 7.73 | 7.70 | 10.42 | 12.67 | 10.62 | 10.07 | 10.04 | 7.72 | 11.55 |
| WDR26 | 3 | 9.37 | 10.58 | 10.36 | 9.65 | 10.83 | 10.76 | 9.45 | 9.57 | 9.61 | 9.82 | 9.98 | 10.00 | 10.79 | 9.51 | 9.71 |
| WDR3 | 26 | 9.75 | 9.80 | 9.83 | 10.50 | 8.15 | 8.50 | 10.34 | 10.55 | 11.07 | 11.52 | 9.78 | 10.16 | 8.32 | 10.44 | 11.30 |
| WDR33 | 12 | 10.91 | 11.18 | 10.58 | 10.40 | 11.12 | 10.64 | 10.39 | 9.52 | 7.97 | 7.27 | 11.05 | 10.49 | 10.88 | 9.96 | 7.62 |
| WDR34 | 3 | 9.48 | 13.36 | 7.74 | 8.98 | 11.46 | 10.75 | 8.23 | 8.37 | 10.25 | 11.37 | 11.42 | 8.36 | 11.10 | 8.30 | 10.81 |
| WDR35 | 4 | 10.18 | 9.58 | 11.83 | 11.64 | 9.64 | 8.33 | 9.02 | 8.47 | 10.47 | 10.84 | 9.88 | 11.74 | 8.98 | 8.75 | 10.66 |
| WDR36 | 20 | 9.76 | 10.39 | 10.37 | 10.68 | 8.37 | 8.76 | 10.46 | 9.87 | 10.90 | 10.44 | 10.07 | 10.52 | 8.56 | 10.17 | 10.67 |
| WDR37 | 3 | 9.99 | 9.53 | 9.38 | 9.47 | 11.67 | 12.89 | 9.25 | 8.54 | 9.61 | 9.68 | 9.76 | 9.42 | 12.28 | 8.89 | 9.64 |
| WDR4 | 5 | 10.26 | 11.15 | 9.74 | 9.51 | 8.60 | 9.24 | 11.58 | 11.28 | 9.33 | 9.30 | 10.70 | 9.63 | 8.92 | 11.43 | 9.32 |
| WDR41 | 2 | 11.51 | 11.13 | 9.62 | 8.42 | 8.74 | 9.15 | 8.07 | 10.75 | 12.06 | 10.54 | 11.32 | 9.02 | 8.95 | 9.41 | 11.30 |
| WDR43 | 21 | 7.52 | 8.82 | 11.77 | 11.01 | 7.84 | 8.03 | 11.06 | 14.53 | 9.43 | 9.99 | 8.17 | 11.39 | 7.93 | 12.79 | 9.71 |
| WDR44 | 36 | 10.44 | 11.40 | 8.07 | 8.00 | 13.10 | 13.82 | 8.91 | 8.55 | 9.01 | 8.70 | 10.92 | 8.03 | 13.46 | 8.73 | 8.86 |
| WDR45 | 4 | 9.12 | 10.04 | 8.98 | 8.66 | 12.53 | 12.53 | 9.99 | 9.64 | 9.38 | 9.13 | 9.58 | 8.82 | 12.53 | 9.81 | 9.26 |
| WDR45B | 3 | 8.26 | 10.96 | 11.27 | 11.55 | 9.52 | 8.94 | 9.49 | 8.39 | 10.80 | 10.84 | 9.61 | 11.41 | 9.23 | 8.94 | 10.82 |
| WDR46 | 9 | 9.71 | 9.89 | 10.74 | 10.83 | 7.55 | 7.78 | 11.74 | 10.14 | 10.42 | 11.19 | 9.80 | 10.79 | 7.67 | 10.94 | 10.80 |
| WDR48 | 5 | 10.58 | 10.57 | 10.87 | 10.61 | 9.39 | 10.08 | 9.72 | 9.12 | 9.83 | 9.22 | 10.58 | 10.74 | 9.74 | 9.42 | 9.53 |
| WDR5 | 13 | 9.29 | 9.40 | 11.75 | 9.46 | 10.06 | 10.06 | 10.70 | 9.17 | 10.17 | 9.93 | 9.35 | 10.61 | 10.06 | 9.93 | 10.05 |
| WDR55 | 5 | 11.07 | 11.57 | 7.45 | 7.00 | 10.72 | 11.33 | 11.22 | 10.08 | 10.21 | 9.35 | 11.32 | 7.23 | 11.02 | 10.65 | 9.78 |
| WDR59 | 1 | 9.22 | 10.54 | 18.47 | 8.81 | 6.64 | 8.15 | 7.73 | 6.63 | 12.91 | 10.91 | 9.88 | 13.64 | 7.40 | 7.18 | 11.91 |
| WDR6 | 7 | 9.54 | 9.92 | 12.01 | 11.58 | 9.44 | 9.53 | 7.85 | 7.92 | 11.07 | 11.14 | 9.73 | 11.79 | 9.48 | 7.88 | 11.10 |
| WDR61 | 14 | 9.57 | 9.90 | 10.47 | 9.10 | 10.65 | 10.21 | 9.66 | 9.70 | 10.60 | 10.14 | 9.73 | 9.78 | 10.43 | 9.68 | 10.37 |
| WDR62 | 3 | 9.33 | 9.87 | 13.20 | 12.35 | 7.90 | 7.69 | 10.26 | 8.76 | 10.94 | 9.71 | 9.60 | 12.77 | 7.79 | 9.51 | 10.32 |
| WDR7 | 3 | 11.57 | 11.32 | 8.53 | 10.25 | 10.71 | 11.05 | 8.14 | 9.37 | 9.33 | 9.72 | 11.45 | 9.39 | 10.88 | 8.76 | 9.53 |
| WDR70 | 19 | 10.03 | 10.82 | 9.85 | 9.17 | 9.02 | 9.30 | 12.19 | 11.00 | 9.52 | 9.09 | 10.43 | 9.51 | 9.16 | 11.60 | 9.30 |
| WDR73 | 4 | 9.29 | 9.76 | 12.74 | 10.94 | 10.19 | 10.02 | 8.55 | 7.41 | 10.75 | 10.35 | 9.52 | 11.84 | 10.11 | 7.98 | 10.55 |
| WDR74 | 9 | 9.74 | 10.22 | 9.22 | 9.92 | 7.80 | 8.49 | 12.21 | 11.85 | 10.49 | 10.07 | 9.98 | 9.57 | 8.14 | 12.03 | 10.28 |
| WDR75 | 6 | 8.84 | 7.81 | 10.12 | 9.39 | 8.87 | 8.20 | 12.22 | 14.52 | 10.15 | 9.89 | 8.32 | 9.75 | 8.54 | 13.37 | 10.02 |
| WDR76 | 4 | 8.93 | 10.49 | 7.14 | 7.43 | 10.37 | 10.96 | 10.49 | 9.94 | 11.63 | 12.61 | 9.71 | 7.29 | 10.66 | 10.22 | 12.12 |

| | | | | | | | | | | | | | | | | |
|---|---|---|---|---|---|---|---|---|---|---|---|---|---|---|---|---|
| WDR77 | 4 | 10.51 | 10.22 | 9.28 | 9.41 | 9.51 | 9.52 | 10.43 | 10.83 | 10.24 | 10.03 | 10.37 | 9.34 | 9.52 | 10.63 | 10.14 |
| WDR81 | 2 | 7.96 | 11.06 | 11.49 | 11.68 | 9.21 | 9.14 | 9.56 | 8.69 | 10.81 | 10.40 | 9.51 | 11.58 | 9.17 | 9.12 | 10.61 |
| WDR82 | 10 | 9.62 | 9.88 | 10.75 | 8.50 | 11.12 | 10.46 | 10.52 | 8.32 | 10.43 | 10.39 | 9.75 | 9.63 | 10.79 | 9.42 | 10.41 |
| WDR83 | 1 | 9.88 | 11.00 | 11.09 | 12.84 | 7.90 | 8.57 | 10.94 | 12.59 | 6.61 | 8.56 | 10.44 | 11.97 | 8.24 | 11.77 | 7.59 |
| WDR89 | 2 | 9.34 | 10.47 | 9.13 | 8.27 | 10.19 | 11.64 | 11.39 | 10.38 | 9.85 | 9.33 | 9.91 | 8.70 | 10.92 | 10.89 | 9.59 |
| WDR91 | 6 | 10.00 | 10.78 | 10.17 | 10.82 | 8.58 | 9.87 | 9.13 | 9.69 | 10.51 | 10.46 | 10.39 | 10.50 | 9.23 | 9.41 | 10.48 |
| WDR92 | 8 | 10.15 | 10.56 | 13.20 | 11.67 | 7.34 | 7.47 | 10.77 | 10.37 | 9.17 | 9.30 | 10.36 | 12.43 | 7.40 | 10.57 | 9.24 |
| WFS1 | 1 | 12.31 | 7.25 | 10.81 | 13.14 | 11.85 | 8.97 | 7.37 | 6.99 | 9.78 | 11.54 | 9.78 | 11.97 | 10.41 | 7.18 | 10.66 |
| WHSC1 | 7 | 11.15 | 11.67 | 9.23 | 9.36 | 9.39 | 9.44 | 11.27 | 11.05 | 8.34 | 9.09 | 11.41 | 9.29 | 9.41 | 11.16 | 8.72 |
| WHSC1L1 | 10 | 8.39 | 9.60 | 9.14 | 9.87 | 8.84 | 8.90 | 14.85 | 13.92 | 8.39 | 8.10 | 8.99 | 9.51 | 8.87 | 14.39 | 8.24 |
| WIBG | 6 | 11.53 | 12.37 | 13.20 | 11.45 | 13.21 | 13.43 | 10.38 | 8.53 | 3.25 | 2.66 | 11.95 | 12.32 | 13.32 | 9.45 | 2.95 |
| WIPF1 | 12 | 10.84 | 11.99 | 11.33 | 10.79 | 10.56 | 10.20 | 9.61 | 8.58 | 8.36 | 7.75 | 11.41 | 11.06 | 10.38 | 9.09 | 8.05 |
| WIPF2 | 4 | 7.54 | 9.11 | 12.09 | 12.99 | 7.00 | 7.79 | 10.71 | 10.82 | 11.19 | 10.77 | 8.32 | 12.54 | 7.39 | 10.77 | 10.98 |
| WIPI2 | 3 | 10.18 | 9.79 | 12.07 | 11.66 | 8.84 | 8.72 | 9.30 | 8.75 | 10.72 | 9.98 | 9.98 | 11.86 | 8.78 | 9.02 | 10.35 |
| WIZ | 15 | 9.17 | 9.91 | 10.20 | 10.14 | 9.23 | 9.72 | 11.98 | 11.63 | 9.27 | 8.75 | 9.54 | 10.17 | 9.47 | 11.80 | 9.01 |
| WLS | 2 | 11.52 | 8.42 | 9.85 | 15.19 | 6.66 | 6.91 | 12.19 | 11.90 | 8.71 | 8.65 | 9.97 | 12.52 | 6.78 | 12.05 | 8.68 |
| WNK1 | 37 | 10.03 | 10.94 | 10.21 | 9.75 | 10.49 | 10.84 | 9.79 | 9.18 | 9.66 | 9.11 | 10.48 | 9.98 | 10.67 | 9.49 | 9.38 |
| WNT5A | 1 | 11.27 | 7.31 | 8.42 | 10.71 | 7.47 | 10.49 | 12.67 | 10.67 | 14.41 | 6.59 | 9.29 | 9.56 | 8.98 | 11.67 | 10.50 |
| WRAP53 | 1 | 9.70 | 10.98 | 11.76 | 12.42 | 8.09 | 8.90 | 8.50 | 8.37 | 12.04 | 9.25 | 10.34 | 12.09 | 8.49 | 8.44 | 10.65 |
| WRN | 3 | 9.29 | 9.53 | 10.11 | 9.09 | 7.71 | 7.61 | 15.27 | 14.10 | 8.71 | 8.57 | 9.41 | 9.60 | 7.66 | 14.68 | 8.64 |
| WRNIP1 | 10 | 9.80 | 10.88 | 7.86 | 7.28 | 9.11 | 10.12 | 11.54 | 11.04 | 11.61 | 10.77 | 10.34 | 7.57 | 9.62 | 11.29 | 11.19 |
| WTAP | 6 | 9.40 | 11.20 | 10.85 | 10.40 | 9.00 | 9.18 | 10.70 | 9.41 | 10.20 | 9.67 | 10.30 | 10.62 | 9.09 | 10.05 | 9.94 |
| WWC1 | 5 | 9.75 | 11.00 | 14.14 | 13.78 | 7.23 | 7.72 | 6.60 | 5.98 | 12.17 | 11.63 | 10.37 | 13.96 | 7.48 | 6.29 | 11.90 |
| WWC2 | 3 | 12.10 | 13.76 | 11.21 | 10.75 | 8.14 | 8.98 | 6.69 | 5.14 | 12.12 | 11.09 | 12.93 | 10.98 | 8.56 | 5.92 | 11.60 |
| WWP1 | 4 | 10.42 | 11.16 | 7.94 | 11.22 | 9.12 | 11.00 | 9.87 | 10.22 | 9.59 | 9.46 | 10.79 | 9.58 | 10.06 | 10.05 | 9.53 |
| WWP2 | 1 | 11.48 | 10.72 | 9.69 | 9.15 | 11.00 | 10.38 | 9.64 | 9.81 | 8.70 | 9.42 | 11.10 | 9.42 | 10.69 | 9.73 | 9.06 |
| XAB2 | 19 | 9.10 | 10.37 | 10.47 | 10.09 | 9.96 | 9.45 | 9.94 | 9.89 | 10.41 | 10.31 | 9.73 | 10.28 | 9.71 | 9.92 | 10.36 |
| XBP1 | 1 | 9.19 | 7.68 | 11.49 | 0.36 | 15.96 | 11.16 | 12.78 | 5.20 | 14.37 | 11.79 | 8.44 | 5.93 | 13.56 | 8.99 | 13.08 |
| XIAP | 6 | 13.04 | 9.50 | 8.30 | 8.38 | 11.51 | 12.17 | 11.07 | 9.17 | 8.83 | 8.02 | 11.27 | 8.34 | 11.84 | 10.12 | 8.42 |
| XPNPEP1 | 17 | 9.97 | 10.57 | 9.54 | 8.20 | 11.15 | 11.04 | 10.36 | 10.40 | 9.61 | 9.17 | 10.27 | 8.87 | 11.09 | 10.38 | 9.39 |
| XPNPEP3 | 4 | 11.09 | 9.75 | 10.63 | 12.19 | 6.01 | 6.00 | 12.96 | 13.06 | 9.24 | 9.08 | 10.42 | 11.41 | 6.01 | 13.01 | 9.16 |
| XPO1 | 51 | 9.51 | 9.03 | 9.87 | 9.25 | 9.90 | 9.26 | 10.54 | 12.87 | 9.86 | 9.90 | 9.27 | 9.56 | 9.58 | 11.71 | 9.88 |
| XPO4 | 8 | 9.35 | 10.17 | 10.04 | 10.33 | 11.23 | 11.71 | 8.72 | 9.10 | 9.40 | 9.95 | 9.76 | 10.18 | 11.47 | 8.91 | 9.67 |
| XPO5 | 20 | 8.74 | 8.76 | 10.61 | 11.13 | 9.37 | 9.75 | 10.66 | 11.13 | 9.84 | 10.03 | 8.75 | 10.87 | 9.56 | 10.89 | 9.93 |
| XPO6 | 5 | 11.35 | 10.26 | 10.06 | 10.71 | 9.01 | 9.34 | 8.75 | 9.31 | 10.45 | 10.76 | 10.80 | 10.39 | 9.17 | 9.03 | 10.60 |
| XPO7 | 23 | 9.18 | 9.00 | 9.27 | 9.54 | 8.52 | 9.01 | 12.40 | 13.32 | 9.76 | 9.98 | 9.09 | 9.41 | 8.77 | 12.86 | 9.87 |
| XPOT | 25 | 8.52 | 8.60 | 7.93 | 8.35 | 10.32 | 10.81 | 11.55 | 13.56 | 10.11 | 10.26 | 8.56 | 8.14 | 10.56 | 12.56 | 10.19 |
| XPR1 | 1 | 8.12 | 8.76 | 7.00 | 9.97 | 8.91 | 8.84 | 8.19 | 8.72 | 14.10 | 17.39 | 8.44 | 8.48 | 8.88 | 8.45 | 15.74 |
| XRCC1 | 13 | 10.35 | 12.85 | 9.71 | 9.03 | 9.58 | 10.28 | 9.94 | 9.94 | 9.32 | 9.01 | 11.60 | 9.37 | 9.93 | 9.94 | 9.16 |
| XRCC4 | 9 | 8.77 | 9.46 | 14.31 | 12.89 | 8.75 | 8.40 | 11.70 | 10.64 | 8.03 | 7.06 | 9.11 | 13.60 | 8.58 | 11.17 | 7.54 |
| XRCC5 | 101 | 9.53 | 9.80 | 12.12 | 10.60 | 8.93 | 8.60 | 9.41 | 11.70 | 9.79 | 9.52 | 9.67 | 11.36 | 8.76 | 10.56 | 9.65 |
| XRCC6 | 113 | 9.61 | 10.43 | 12.09 | 11.29 | 8.98 | 9.10 | 9.56 | 9.08 | 10.04 | 9.81 | 10.02 | 11.69 | 9.04 | 9.32 | 9.93 |

| | | | | | | | | | | | | | | | | |
|---|---|---|---|---|---|---|---|---|---|---|---|---|---|---|---|---|
| XRN1 | 13 | 10.16 | 10.88 | 8.67 | 8.52 | 10.74 | 11.38 | 11.06 | 10.09 | 9.18 | 9.31 | 10.52 | 8.60 | 11.06 | 10.57 | 9.25 |
| XRN2 | 24 | 9.52 | 9.52 | 10.86 | 10.36 | 9.59 | 9.50 | 11.29 | 10.68 | 9.68 | 9.00 | 9.52 | 10.61 | 9.54 | 10.98 | 9.34 |
| XYLB | 2 | 9.79 | 10.60 | 6.24 | 6.04 | 10.46 | 9.83 | 9.25 | 8.99 | 14.48 | 14.32 | 10.20 | 6.14 | 10.14 | 9.12 | 14.40 |
| YAP1 | 13 | 8.64 | 9.63 | 12.79 | 12.07 | 9.91 | 9.83 | 11.04 | 7.44 | 9.45 | 9.21 | 9.13 | 12.43 | 9.87 | 9.24 | 9.33 |
| YARS | 70 | 9.73 | 10.24 | 11.21 | 10.10 | 6.86 | 6.63 | 9.92 | 10.07 | 12.85 | 12.39 | 9.99 | 10.65 | 6.75 | 10.00 | 12.62 |
| YARS2 | 12 | 9.79 | 8.50 | 10.12 | 12.04 | 9.49 | 9.05 | 10.41 | 9.78 | 10.21 | 10.63 | 9.14 | 11.08 | 9.27 | 10.09 | 10.42 |
| YBEY | 1 | 11.88 | 11.50 | 9.65 | 3.76 | 14.02 | 9.44 | 12.59 | 5.19 | 10.99 | 10.98 | 11.69 | 6.70 | 11.73 | 8.89 | 10.98 |
| YBX1 | 32 | 9.26 | 10.96 | 9.90 | 9.85 | 8.82 | 9.12 | 10.92 | 9.87 | 10.57 | 10.72 | 10.11 | 9.88 | 8.97 | 10.39 | 10.65 |
| YBX3 | 19 | 9.00 | 9.61 | 11.81 | 9.61 | 10.01 | 9.39 | 11.13 | 9.49 | 10.12 | 9.83 | 9.31 | 10.71 | 9.70 | 10.31 | 9.98 |
| YEATS2 | 3 | 9.98 | 12.33 | 9.95 | 9.00 | 7.29 | 7.68 | 11.52 | 10.69 | 10.93 | 10.63 | 11.16 | 9.47 | 7.48 | 11.11 | 10.78 |
| YEATS4 | 1 | 10.87 | 11.91 | 10.66 | 9.93 | 9.06 | 10.41 | 8.26 | 7.61 | 11.39 | 9.89 | 11.39 | 10.29 | 9.73 | 7.93 | 10.64 |
| YES1 | 3 | 10.88 | 9.47 | 11.48 | 13.23 | 8.81 | 8.87 | 9.07 | 8.22 | 10.08 | 9.89 | 10.18 | 12.36 | 8.84 | 8.64 | 9.98 |
| YIF1A | 1 | 16.82 | 11.46 | 7.58 | 9.83 | 7.35 | 7.03 | 9.73 | 7.99 | 12.36 | 9.85 | 14.14 | 8.71 | 7.19 | 8.86 | 11.10 |
| YIF1B | 1 | 11.27 | 8.16 | 8.14 | 6.53 | 10.69 | 9.39 | 12.39 | 10.76 | 10.23 | 12.44 | 9.72 | 7.34 | 10.04 | 11.57 | 11.33 |
| YIPF3 | 1 | 9.60 | 8.25 | 8.04 | 11.46 | 10.84 | 10.96 | 11.31 | 8.27 | 9.11 | 12.17 | 8.92 | 9.75 | 10.90 | 9.79 | 10.64 |
| YIPF4 | 2 | 10.37 | 8.68 | 8.69 | 12.00 | 9.22 | 9.72 | 9.38 | 9.04 | 11.01 | 11.89 | 9.52 | 10.34 | 9.47 | 9.21 | 11.45 |
| YIPF5 | 3 | 11.25 | 7.14 | 7.85 | 10.36 | 9.34 | 9.03 | 11.32 | 10.27 | 10.74 | 12.70 | 9.19 | 9.11 | 9.18 | 10.80 | 11.72 |
| YKT6 | 6 | 11.12 | 8.94 | 10.72 | 12.64 | 7.66 | 7.45 | 9.28 | 9.41 | 11.04 | 11.75 | 10.03 | 11.68 | 7.55 | 9.34 | 11.39 |
| YLPM1 | 35 | 9.14 | 10.39 | 10.86 | 10.06 | 10.17 | 10.25 | 10.29 | 9.11 | 10.17 | 9.55 | 9.77 | 10.46 | 10.21 | 9.70 | 9.86 |
| YME1L1 | 9 | 13.25 | 6.66 | 7.65 | 11.97 | 8.24 | 7.39 | 10.65 | 9.69 | 10.89 | 13.61 | 9.95 | 9.81 | 7.82 | 10.17 | 12.25 |
| YOD1 | 2 | 8.45 | 8.62 | 6.98 | 6.80 | 12.85 | 12.79 | 11.98 | 11.54 | 10.40 | 9.61 | 8.53 | 6.89 | 12.82 | 11.76 | 10.00 |
| YRDC | 6 | 10.92 | 11.50 | 7.72 | 8.15 | 8.53 | 8.39 | 10.83 | 10.55 | 12.01 | 11.41 | 11.21 | 7.93 | 8.46 | 10.69 | 11.71 |
| YTHDC1 | 3 | 8.87 | 10.57 | 9.24 | 9.13 | 9.76 | 11.03 | 11.37 | 10.78 | 9.88 | 9.38 | 9.72 | 9.18 | 10.39 | 11.07 | 9.63 |
| YTHDC2 | 22 | 9.54 | 10.47 | 10.32 | 9.69 | 9.19 | 9.45 | 10.83 | 10.79 | 9.91 | 9.80 | 10.01 | 10.01 | 9.32 | 10.81 | 9.86 |
| YTHDF1 | 8 | 9.35 | 11.15 | 9.86 | 9.45 | 9.75 | 9.89 | 10.52 | 9.67 | 10.69 | 9.68 | 10.25 | 9.65 | 9.82 | 10.10 | 10.18 |
| YTHDF2 | 15 | 9.33 | 11.00 | 9.35 | 9.18 | 9.57 | 9.71 | 11.41 | 11.01 | 9.68 | 9.76 | 10.17 | 9.27 | 9.64 | 11.21 | 9.72 |
| YTHDF3 | 15 | 9.35 | 10.45 | 11.06 | 9.53 | 9.89 | 9.62 | 11.52 | 9.73 | 9.86 | 9.00 | 9.90 | 10.30 | 9.75 | 10.63 | 9.43 |
| YWHAB | 30 | 10.27 | 10.52 | 10.39 | 9.61 | 10.80 | 11.23 | 9.44 | 9.16 | 9.24 | 9.33 | 10.39 | 10.00 | 11.02 | 9.30 | 9.28 |
| YWHAE | 118 | 10.09 | 12.05 | 10.55 | 10.39 | 9.32 | 9.42 | 9.19 | 8.49 | 10.39 | 10.12 | 11.07 | 10.47 | 9.37 | 8.84 | 10.25 |
| YWHAG | 25 | 7.03 | 10.10 | 13.44 | 11.99 | 8.78 | 8.56 | 9.16 | 10.33 | 10.12 | 10.49 | 8.56 | 12.71 | 8.67 | 9.74 | 10.31 |
| YWHAH | 20 | 9.68 | 11.51 | 13.30 | 11.74 | 7.43 | 7.18 | 7.41 | 7.77 | 12.41 | 11.57 | 10.60 | 12.52 | 7.31 | 7.59 | 11.99 |
| YWHAQ | 21 | 7.03 | 9.18 | 8.79 | 7.45 | 9.89 | 9.28 | 8.79 | 24.01 | 8.10 | 7.47 | 8.11 | 8.12 | 9.59 | 16.40 | 7.78 |
| YWHAZ | 42 | 8.83 | 10.40 | 10.91 | 9.73 | 9.53 | 9.25 | 10.59 | 11.54 | 9.82 | 9.39 | 9.62 | 10.32 | 9.39 | 11.06 | 9.61 |
| YY1 | 9 | 9.97 | 9.53 | 8.82 | 7.39 | 8.99 | 8.73 | 11.19 | 18.16 | 8.95 | 8.25 | 9.75 | 8.11 | 8.86 | 14.68 | 8.60 |
| YY1AP1 | 3 | 9.94 | 11.15 | 10.00 | 8.38 | 12.03 | 10.52 | 9.67 | 9.26 | 9.81 | 9.25 | 10.54 | 9.19 | 11.27 | 9.46 | 9.53 |
| ZADH2 | 2 | 9.64 | 9.92 | 11.87 | 12.47 | 8.76 | 7.78 | 11.20 | 9.37 | 9.13 | 9.86 | 9.78 | 12.17 | 8.27 | 10.29 | 9.49 |
| ZBED1 | 2 | 12.09 | 12.50 | 4.47 | 4.60 | 19.30 | 20.15 | 7.10 | 6.53 | 6.49 | 6.76 | 12.29 | 4.54 | 19.73 | 6.82 | 6.63 |
| ZBED4 | 1 | 7.55 | 9.78 | 11.83 | 13.20 | 7.91 | 8.17 | 10.50 | 11.80 | 8.45 | 10.81 | 8.67 | 12.51 | 8.04 | 11.15 | 9.63 |
| ZBTB1 | 2 | 9.74 | 10.51 | 10.32 | 8.88 | 11.26 | 10.70 | 9.45 | 9.02 | 10.28 | 9.83 | 10.13 | 9.60 | 10.98 | 9.24 | 10.05 |
| ZBTB2 | 1 | 10.29 | 11.04 | 10.79 | 10.64 | 11.02 | 8.98 | 10.07 | 9.93 | 9.41 | 7.82 | 10.67 | 10.72 | 10.00 | 10.00 | 8.61 |
| ZBTB21 | 4 | 10.03 | 10.02 | 9.29 | 8.97 | 9.21 | 9.80 | 11.17 | 10.40 | 10.71 | 10.40 | 10.03 | 9.13 | 9.50 | 10.79 | 10.56 |
| ZBTB3 | 1 | 10.36 | 10.85 | 11.52 | 10.48 | 11.53 | 11.48 | 7.56 | 7.07 | 10.71 | 8.43 | 10.61 | 11.00 | 11.50 | 7.32 | 9.57 |

| | | | | | | | | | | | | | | | | |
|---|---|---|---|---|---|---|---|---|---|---|---|---|---|---|---|---|
| ZBTB33 | 7 | 8.53 | 9.61 | 7.51 | 7.32 | 11.31 | 11.57 | 15.86 | 14.90 | 6.89 | 6.50 | 9.07 | 7.42 | 11.44 | 15.38 | 6.69 |
| ZBTB40 | 1 | 9.21 | 11.50 | 9.50 | 7.88 | 10.41 | 10.60 | 12.59 | 11.58 | 8.88 | 7.85 | 10.35 | 8.69 | 10.51 | 12.08 | 8.37 |
| ZBTB7A | 1 | 7.93 | 8.62 | 11.14 | 12.27 | 6.46 | 9.51 | 12.32 | 10.27 | 9.42 | 12.06 | 8.28 | 11.70 | 7.98 | 11.29 | 10.74 |
| ZBTB8OS | 3 | 10.35 | 10.71 | 10.49 | 7.36 | 8.11 | 9.89 | 9.96 | 10.67 | 11.02 | 11.43 | 10.53 | 8.92 | 9.00 | 10.32 | 11.23 |
| ZBTB9 | 2 | 9.17 | 10.23 | 11.03 | 10.28 | 8.83 | 8.86 | 12.50 | 10.41 | 9.82 | 8.88 | 9.70 | 10.65 | 8.85 | 11.45 | 9.35 |
| ZC2HC1A | 6 | 9.67 | 11.13 | 11.65 | 11.23 | 8.54 | 9.10 | 11.96 | 11.08 | 8.08 | 7.57 | 10.40 | 11.44 | 8.82 | 11.52 | 7.82 |
| ZC3H11A | 21 | 8.34 | 9.67 | 10.35 | 10.19 | 10.47 | 10.77 | 11.46 | 10.46 | 9.18 | 9.12 | 9.01 | 10.27 | 10.62 | 10.96 | 9.15 |
| ZC3H13 | 13 | 10.62 | 11.60 | 10.12 | 10.08 | 8.75 | 9.84 | 10.29 | 10.16 | 9.62 | 8.92 | 11.11 | 10.10 | 9.29 | 10.22 | 9.27 |
| ZC3H14 | 29 | 9.76 | 10.42 | 9.70 | 8.87 | 10.66 | 10.79 | 11.48 | 9.84 | 9.52 | 8.97 | 10.09 | 9.28 | 10.72 | 10.66 | 9.24 |
| ZC3H15 | 24 | 9.53 | 9.81 | 11.84 | 11.47 | 9.57 | 9.61 | 11.84 | 10.81 | 7.78 | 7.73 | 9.67 | 11.66 | 9.59 | 11.33 | 7.76 |
| ZC3H18 | 22 | 10.45 | 11.38 | 8.52 | 8.12 | 10.56 | 11.06 | 10.06 | 8.80 | 11.01 | 10.04 | 10.91 | 8.32 | 10.81 | 9.43 | 10.53 |
| ZC3H4 | 16 | 10.00 | 10.10 | 9.99 | 9.05 | 11.27 | 11.12 | 10.82 | 9.32 | 9.56 | 8.77 | 10.05 | 9.52 | 11.20 | 10.07 | 9.17 |
| ZC3H7A | 13 | 9.33 | 9.67 | 12.38 | 10.78 | 10.70 | 9.49 | 10.02 | 8.48 | 9.49 | 9.66 | 9.50 | 11.58 | 10.10 | 9.25 | 9.58 |
| ZC3H7B | 8 | 9.85 | 10.75 | 12.03 | 11.72 | 9.02 | 8.78 | 8.62 | 10.03 | 9.70 | 9.50 | 10.30 | 11.88 | 8.90 | 9.32 | 9.60 |
| ZC3H8 | 5 | 9.11 | 9.75 | 10.51 | 10.12 | 9.67 | 10.40 | 13.08 | 11.67 | 8.42 | 7.26 | 9.43 | 10.31 | 10.04 | 12.38 | 7.84 |
| ZC3HAV1 | 39 | 9.74 | 11.03 | 10.40 | 9.39 | 9.15 | 9.08 | 10.42 | 10.05 | 10.44 | 10.30 | 10.38 | 9.90 | 9.12 | 10.23 | 10.37 |
| ZC3HAV1L | 1 | 9.58 | 10.31 | 13.31 | 6.68 | 10.53 | 7.59 | 11.29 | 5.93 | 12.22 | 12.56 | 9.95 | 9.99 | 9.06 | 8.61 | 12.39 |
| ZC3HC1 | 10 | 8.88 | 11.39 | 10.24 | 9.34 | 10.26 | 10.24 | 9.41 | 9.98 | 9.76 | 10.49 | 10.14 | 9.79 | 10.25 | 9.70 | 10.12 |
| ZCCHC11 | 4 | 11.18 | 10.89 | 9.32 | 9.41 | 8.71 | 8.26 | 11.31 | 10.54 | 10.36 | 10.02 | 11.04 | 9.36 | 8.48 | 10.92 | 10.19 |
| ZCCHC17 | 2 | 8.73 | 11.87 | 9.48 | 10.04 | 9.20 | 9.26 | 10.48 | 12.38 | 8.63 | 9.92 | 10.30 | 9.76 | 9.23 | 11.43 | 9.28 |
| ZCCHC24 | 1 | 7.46 | 11.95 | 10.95 | 10.86 | 9.30 | 10.71 | 12.69 | 12.01 | 7.37 | 6.71 | 9.71 | 10.90 | 10.00 | 12.35 | 7.04 |
| ZCCHC6 | 9 | 8.97 | 9.69 | 10.42 | 10.22 | 10.77 | 11.13 | 10.74 | 9.62 | 9.19 | 9.25 | 9.33 | 10.32 | 10.95 | 10.18 | 9.22 |
| ZCCHC7 | 1 | 10.87 | 10.59 | 10.61 | 8.08 | 8.37 | 7.56 | 10.51 | 10.17 | 11.30 | 11.95 | 10.73 | 9.34 | 7.97 | 10.34 | 11.62 |
| ZCCHC8 | 10 | 8.53 | 9.50 | 12.05 | 11.51 | 8.65 | 9.15 | 10.50 | 9.69 | 10.23 | 10.19 | 9.01 | 11.78 | 8.90 | 10.09 | 10.21 |
| ZCCHC9 | 1 | 8.89 | 10.85 | 8.72 | 12.18 | 6.74 | 7.14 | 13.50 | 14.33 | 8.90 | 8.76 | 9.87 | 10.45 | 6.94 | 13.91 | 8.83 |
| ZDBF2 | 1 | 8.76 | 10.28 | 7.86 | 9.95 | 11.04 | 14.18 | 9.79 | 9.88 | 7.34 | 10.91 | 9.52 | 8.91 | 12.61 | 9.83 | 9.13 |
| ZDHHC17 | 1 | 11.61 | 7.49 | 7.37 | 10.05 | 13.25 | 11.44 | 8.61 | 8.29 | 10.04 | 11.85 | 9.55 | 8.71 | 12.35 | 8.45 | 10.95 |
| ZDHHC5 | 1 | 9.84 | 9.45 | 9.36 | 10.08 | 9.21 | 9.65 | 9.30 | 10.94 | 10.50 | 11.67 | 9.64 | 9.72 | 9.43 | 10.12 | 11.08 |
| ZEB1 | 2 | 8.05 | 9.23 | 13.17 | 12.61 | 8.70 | 8.48 | 11.77 | 10.45 | 9.12 | 8.42 | 8.64 | 12.89 | 8.59 | 11.11 | 8.77 |
| ZFAND5 | 2 | 9.31 | 8.60 | 12.20 | 10.45 | 11.68 | 10.18 | 12.16 | 10.09 | 8.41 | 6.91 | 8.96 | 11.32 | 10.93 | 11.13 | 7.66 |
| ZFAND6 | 2 | 8.16 | 10.07 | 10.72 | 11.66 | 10.95 | 12.65 | 9.10 | 10.16 | 7.82 | 8.70 | 9.12 | 11.19 | 11.80 | 9.63 | 8.26 |
| ZFC3H1 | 18 | 9.26 | 9.27 | 9.11 | 9.46 | 9.45 | 9.29 | 12.93 | 11.79 | 9.89 | 9.53 | 9.27 | 9.29 | 9.37 | 12.36 | 9.71 |
| ZFHX3 | 9 | 11.85 | 11.62 | 8.85 | 8.75 | 10.85 | 12.23 | 8.22 | 7.74 | 10.52 | 9.39 | 11.73 | 8.80 | 11.54 | 7.98 | 9.95 |
| ZFP36L1 | 1 | 10.22 | 9.47 | 12.10 | 10.75 | 8.31 | 7.94 | 10.14 | 9.17 | 11.00 | 10.90 | 9.84 | 11.43 | 8.12 | 9.66 | 10.95 |
| ZFP91 | 2 | 9.67 | 10.03 | 10.27 | 8.97 | 10.15 | 9.32 | 12.99 | 12.08 | 8.52 | 8.01 | 9.85 | 9.62 | 9.73 | 12.54 | 8.26 |
| ZFR | 35 | 9.17 | 10.47 | 10.42 | 9.75 | 9.57 | 9.88 | 10.14 | 11.04 | 9.99 | 9.56 | 9.82 | 10.09 | 9.72 | 10.59 | 9.77 |
| ZFYVE1 | 1 | 10.62 | 11.36 | 12.16 | 12.22 | 8.73 | 8.55 | 9.89 | 9.83 | 8.11 | 8.54 | 10.99 | 12.19 | 8.64 | 9.86 | 8.33 |
| ZFYVE16 | 14 | 10.65 | 11.45 | 9.91 | 9.32 | 9.83 | 10.39 | 10.53 | 9.66 | 9.68 | 8.59 | 11.05 | 9.61 | 10.11 | 10.09 | 9.14 |
| ZFYVE19 | 7 | 9.49 | 10.65 | 8.86 | 8.32 | 11.52 | 11.89 | 7.73 | 6.67 | 12.70 | 12.17 | 10.07 | 8.59 | 11.71 | 7.20 | 12.44 |
| ZFYVE20 | 1 | 9.56 | 11.20 | 10.48 | 9.35 | 11.16 | 10.83 | 8.95 | 7.92 | 10.03 | 10.52 | 10.38 | 9.91 | 10.99 | 8.43 | 10.28 |
| ZFYVE21 | 1 | 7.15 | 6.54 | 13.33 | 5.07 | 15.68 | 8.40 | 9.00 | 16.96 | 8.60 | 9.28 | 6.84 | 9.20 | 12.04 | 12.98 | 8.94 |
| ZFYVE28 | 1 | 9.80 | 9.93 | 9.51 | 6.85 | 10.82 | 11.19 | 10.57 | 10.90 | 10.02 | 10.40 | 9.87 | 8.18 | 11.01 | 10.74 | 10.21 |

| | | | | | | | | | | | | | | | | |
|---|---|---|---|---|---|---|---|---|---|---|---|---|---|---|---|---|
| ZGPAT | 9 | 9.09 | 9.71 | 12.86 | 12.42 | 8.47 | 9.07 | 9.46 | 8.57 | 10.75 | 9.59 | 9.40 | 12.64 | 8.77 | 9.02 | 10.17 |
| ZHX1 | 5 | 10.23 | 9.52 | 10.60 | 9.86 | 9.74 | 9.90 | 11.64 | 9.97 | 9.84 | 8.70 | 9.88 | 10.23 | 9.82 | 10.80 | 9.27 |
| ZKSCAN4 | 2 | 8.56 | 10.67 | 12.12 | 10.76 | 9.62 | 8.78 | 10.26 | 7.92 | 11.10 | 10.19 | 9.62 | 11.44 | 9.20 | 9.09 | 10.65 |
| ZMAT2 | 3 | 7.80 | 12.23 | 10.31 | 10.68 | 9.74 | 11.33 | 9.86 | 10.40 | 8.09 | 9.56 | 10.02 | 10.49 | 10.53 | 10.13 | 8.83 |
| ZMIZ1 | 4 | 9.35 | 10.68 | 10.12 | 10.85 | 6.75 | 7.70 | 14.53 | 13.33 | 8.69 | 7.99 | 10.01 | 10.49 | 7.23 | 13.93 | 8.34 |
| ZMPSTE24 | 13 | 13.07 | 6.76 | 7.14 | 12.01 | 8.51 | 7.96 | 11.26 | 10.56 | 9.91 | 12.81 | 9.92 | 9.57 | 8.24 | 10.91 | 11.36 |
| ZMYM1 | 2 | 10.14 | 11.46 | 9.76 | 10.09 | 7.50 | 8.24 | 11.22 | 10.86 | 10.59 | 10.15 | 10.80 | 9.92 | 7.87 | 11.04 | 10.37 |
| ZMYM2 | 8 | 9.86 | 10.90 | 11.42 | 11.55 | 11.38 | 11.76 | 8.75 | 7.45 | 8.80 | 8.14 | 10.38 | 11.48 | 11.57 | 8.10 | 8.47 |
| ZMYM3 | 6 | 9.23 | 10.26 | 9.39 | 8.96 | 12.56 | 12.94 | 11.37 | 10.41 | 7.52 | 7.35 | 9.74 | 9.17 | 12.75 | 10.89 | 7.44 |
| ZMYM4 | 11 | 9.75 | 10.66 | 8.39 | 8.66 | 9.79 | 9.89 | 12.22 | 11.80 | 9.56 | 9.28 | 10.21 | 8.53 | 9.84 | 12.01 | 9.42 |
| ZMYND11 | 3 | 8.53 | 9.46 | 9.73 | 9.84 | 8.93 | 10.57 | 12.96 | 11.79 | 9.54 | 8.65 | 9.00 | 9.79 | 9.75 | 12.37 | 9.09 |
| ZMYND8 | 3 | 7.99 | 9.48 | 9.96 | 9.84 | 11.69 | 12.02 | 10.99 | 10.41 | 8.92 | 8.70 | 8.73 | 9.90 | 11.86 | 10.70 | 8.81 |
| ZNF106 | 10 | 8.36 | 10.54 | 9.69 | 9.38 | 9.83 | 10.89 | 11.00 | 10.24 | 10.19 | 9.88 | 9.45 | 9.54 | 10.36 | 10.62 | 10.03 |
| ZNF131 | 3 | 8.58 | 9.57 | 7.77 | 7.11 | 11.05 | 11.20 | 15.32 | 14.06 | 7.92 | 7.41 | 9.08 | 7.44 | 11.12 | 14.69 | 7.67 |
| ZNF143 | 2 | 8.64 | 10.89 | 9.06 | 9.58 | 9.81 | 9.77 | 12.65 | 11.94 | 8.43 | 9.24 | 9.77 | 9.32 | 9.79 | 12.29 | 8.84 |
| ZNF148 | 12 | 8.12 | 15.06 | 10.31 | 10.07 | 9.24 | 9.25 | 10.55 | 10.35 | 8.31 | 8.73 | 11.59 | 10.19 | 9.25 | 10.45 | 8.52 |
| ZNF185 | 4 | 6.95 | 7.86 | 6.06 | 6.86 | 9.80 | 8.57 | 21.48 | 18.94 | 7.70 | 5.76 | 7.41 | 6.46 | 9.19 | 20.21 | 6.73 |
| ZNF207 | 17 | 9.49 | 11.27 | 9.84 | 9.36 | 10.52 | 11.51 | 9.04 | 9.56 | 9.81 | 9.60 | 10.38 | 9.60 | 11.02 | 9.30 | 9.71 |
| ZNF217 | 1 | 7.92 | 8.36 | 11.10 | 11.01 | 9.25 | 9.00 | 12.37 | 11.49 | 10.51 | 8.97 | 8.14 | 11.06 | 9.13 | 11.93 | 9.74 |
| ZNF24 | 4 | 9.74 | 10.80 | 13.19 | 10.57 | 10.07 | 10.99 | 7.67 | 6.59 | 11.06 | 9.32 | 10.27 | 11.88 | 10.53 | 7.13 | 10.19 |
| ZNF259 | 13 | 9.38 | 10.18 | 12.90 | 11.99 | 8.96 | 9.01 | 10.37 | 8.87 | 9.45 | 8.88 | 9.78 | 12.44 | 8.99 | 9.62 | 9.16 |
| ZNF277 | 2 | 8.73 | 10.53 | 9.73 | 8.99 | 9.09 | 9.96 | 11.58 | 12.34 | 9.32 | 9.74 | 9.63 | 9.36 | 9.53 | 11.96 | 9.53 |
| ZNF280D | 1 | 9.32 | 10.23 | 10.00 | 9.61 | 10.39 | 10.46 | 11.11 | 11.44 | 8.22 | 9.22 | 9.77 | 9.81 | 10.43 | 11.27 | 8.72 |
| ZNF281 | 2 | 9.34 | 10.16 | 10.93 | 9.17 | 10.08 | 9.83 | 9.81 | 8.46 | 10.93 | 11.29 | 9.75 | 10.05 | 9.95 | 9.14 | 11.11 |
| ZNF318 | 3 | 9.96 | 9.53 | 9.46 | 7.74 | 12.34 | 12.81 | 10.26 | 9.01 | 10.43 | 8.46 | 9.74 | 8.60 | 12.58 | 9.63 | 9.45 |
| ZNF326 | 20 | 9.58 | 10.60 | 12.37 | 11.87 | 7.00 | 7.00 | 13.23 | 11.59 | 8.55 | 8.21 | 10.09 | 12.12 | 7.00 | 12.41 | 8.38 |
| ZNF330 | 2 | 8.87 | 11.42 | 10.44 | 10.47 | 9.01 | 9.38 | 10.24 | 10.54 | 9.84 | 9.80 | 10.15 | 10.46 | 9.19 | 10.39 | 9.82 |
| ZNF346 | 3 | 11.96 | 12.32 | 7.96 | 7.44 | 7.43 | 8.19 | 12.88 | 11.79 | 10.45 | 9.58 | 12.14 | 7.70 | 7.81 | 12.34 | 10.01 |
| ZNF384 | 3 | 9.58 | 10.27 | 12.30 | 9.27 | 10.36 | 10.10 | 11.83 | 8.51 | 9.10 | 8.67 | 9.92 | 10.79 | 10.23 | 10.17 | 8.88 |
| ZNF428 | 2 | 8.56 | 10.59 | 8.98 | 9.89 | 9.44 | 9.40 | 9.45 | 21.22 | 6.87 | 5.58 | 9.58 | 9.44 | 9.42 | 15.34 | 6.23 |
| ZNF444 | 2 | 10.74 | 11.01 | 11.19 | 9.40 | 8.51 | 8.70 | 9.92 | 8.94 | 11.04 | 10.57 | 10.87 | 10.29 | 8.60 | 9.43 | 10.80 |
| ZNF451 | 2 | 9.83 | 9.66 | 9.70 | 10.15 | 9.71 | 9.22 | 12.25 | 13.33 | 7.57 | 8.57 | 9.75 | 9.92 | 9.47 | 12.79 | 8.07 |
| ZNF462 | 1 | 11.52 | 11.59 | 8.44 | 9.33 | 10.92 | 12.28 | 7.40 | 6.61 | 11.98 | 9.93 | 11.56 | 8.88 | 11.60 | 7.01 | 10.95 |
| ZNF511 | 1 | 9.62 | 11.43 | 10.40 | 8.32 | 11.50 | 12.19 | 10.00 | 7.68 | 10.88 | 7.99 | 10.52 | 9.36 | 11.84 | 8.84 | 9.43 |
| ZNF512 | 1 | 16.52 | 10.36 | 8.45 | 12.03 | 9.29 | 9.12 | 6.05 | 5.71 | 8.05 | 14.42 | 13.44 | 10.24 | 9.21 | 5.88 | 11.23 |
| ZNF512B | 1 | 12.07 | 8.04 | 10.14 | 13.88 | 7.95 | 6.99 | 7.22 | 7.24 | 9.79 | 16.68 | 10.05 | 12.01 | 7.47 | 7.23 | 13.24 |
| ZNF532 | 2 | 10.16 | 11.24 | 10.27 | 9.46 | 8.09 | 11.07 | 8.71 | 9.29 | 12.08 | 9.63 | 10.70 | 9.87 | 9.58 | 9.00 | 10.85 |
| ZNF574 | 1 | 7.59 | 9.53 | 8.70 | 8.01 | 10.12 | 10.99 | 12.58 | 13.95 | 9.37 | 9.15 | 8.56 | 8.36 | 10.55 | 13.27 | 9.26 |
| ZNF579 | 1 | 8.78 | 10.00 | 11.79 | 13.39 | 8.05 | 9.79 | 10.28 | 10.00 | 8.94 | 8.99 | 9.39 | 12.59 | 8.92 | 10.14 | 8.96 |
| ZNF592 | 2 | 9.02 | 9.58 | 9.60 | 9.04 | 9.40 | 10.52 | 11.65 | 10.02 | 9.69 | 11.48 | 9.30 | 9.32 | 9.96 | 10.84 | 10.58 |
| ZNF593 | 5 | 9.45 | 10.05 | 9.56 | 8.95 | 8.58 | 9.35 | 12.64 | 11.29 | 10.17 | 9.95 | 9.75 | 9.26 | 8.96 | 11.97 | 10.06 |
| ZNF598 | 17 | 8.99 | 10.12 | 9.89 | 9.07 | 9.32 | 9.44 | 10.20 | 12.22 | 10.57 | 10.19 | 9.55 | 9.48 | 9.38 | 11.21 | 10.38 |

| | | | | | | | | | | | | | | | | |
|---|---|---|---|---|---|---|---|---|---|---|---|---|---|---|---|---|
| ZNF609 | 3 | 10.14 | 11.24 | 10.51 | 9.57 | 9.18 | 9.81 | 9.29 | 8.05 | 11.38 | 10.83 | 10.69 | 10.04 | 9.49 | 8.67 | 11.10 |
| ZNF622 | 17 | 9.06 | 10.84 | 10.75 | 9.85 | 9.26 | 9.58 | 11.23 | 11.70 | 8.97 | 8.78 | 9.95 | 10.30 | 9.42 | 11.46 | 8.87 |
| ZNF629 | 2 | 8.69 | 11.17 | 9.71 | 10.79 | 9.83 | 9.65 | 10.25 | 7.62 | 11.83 | 10.46 | 9.93 | 10.25 | 9.74 | 8.93 | 11.14 |
| ZNF638 | 36 | 9.08 | 10.50 | 9.22 | 8.84 | 10.30 | 11.16 | 12.70 | 12.08 | 8.28 | 7.84 | 9.79 | 9.03 | 10.73 | 12.39 | 8.06 |
| ZNF644 | 2 | 10.60 | 11.01 | 9.71 | 8.44 | 7.89 | 7.56 | 13.64 | 13.36 | 8.54 | 9.25 | 10.81 | 9.08 | 7.72 | 13.50 | 8.90 |
| ZNF654 | 1 | 9.57 | 10.50 | 9.24 | 9.39 | 9.77 | 10.64 | 10.85 | 9.87 | 10.44 | 9.71 | 10.03 | 9.32 | 10.21 | 10.36 | 10.08 |
| ZNF668 | 1 | 8.77 | 10.47 | 10.95 | 9.66 | 9.40 | 10.29 | 12.07 | 9.86 | 8.93 | 9.60 | 9.62 | 10.30 | 9.84 | 10.97 | 9.27 |
| ZNF687 | 4 | 7.89 | 9.56 | 10.39 | 11.17 | 12.28 | 13.03 | 11.02 | 10.20 | 7.28 | 7.17 | 8.73 | 10.78 | 12.65 | 10.61 | 7.22 |
| ZNF706 | 1 | 8.32 | 8.78 | 9.84 | 9.00 | 10.43 | 9.67 | 14.19 | 12.15 | 8.89 | 8.74 | 8.55 | 9.42 | 10.05 | 13.17 | 8.81 |
| ZNF740 | 1 | 9.22 | 8.49 | 10.65 | 9.92 | 9.76 | 10.08 | 12.98 | 13.35 | 6.29 | 9.25 | 8.86 | 10.29 | 9.92 | 13.17 | 7.77 |
| ZNF787 | 1 | 9.26 | 10.36 | 11.82 | 10.28 | 8.05 | 10.21 | 12.76 | 8.58 | 9.11 | 9.57 | 9.81 | 11.05 | 9.13 | 10.67 | 9.34 |
| ZNF800 | 1 | 11.78 | 9.95 | 10.40 | 10.28 | 7.45 | 8.77 | 11.97 | 11.66 | 8.46 | 9.28 | 10.86 | 10.34 | 8.11 | 11.82 | 8.87 |
| ZNF814 | 1 | 7.95 | 10.40 | 7.28 | 5.94 | 21.76 | 7.76 | 8.99 | 4.92 | 9.35 | 15.64 | 9.18 | 6.61 | 14.76 | 6.95 | 12.50 |
| ZNF827 | 1 | 9.07 | 11.32 | 13.93 | 13.13 | 8.26 | 10.65 | 5.57 | 6.72 | 10.74 | 10.60 | 10.19 | 13.53 | 9.45 | 6.15 | 10.67 |
| ZNF830 | 10 | 9.79 | 10.56 | 10.68 | 10.42 | 10.07 | 9.74 | 8.62 | 10.23 | 10.19 | 9.70 | 10.17 | 10.55 | 9.90 | 9.42 | 9.94 |
| ZNFX1 | 1 | 8.81 | 9.62 | 7.48 | 5.91 | 14.00 | 10.98 | 8.17 | 12.79 | 12.15 | 10.10 | 9.22 | 6.69 | 12.49 | 10.48 | 11.12 |
| ZNHIT1 | 1 | 10.86 | 11.18 | 11.28 | 9.50 | 9.72 | 10.36 | 10.06 | 9.50 | 8.92 | 8.63 | 11.02 | 10.39 | 10.04 | 9.78 | 8.77 |
| ZNHIT2 | 7 | 9.58 | 10.96 | 12.18 | 11.84 | 9.03 | 9.67 | 7.46 | 7.57 | 11.51 | 10.19 | 10.27 | 12.01 | 9.35 | 7.51 | 10.85 |
| ZNHIT3 | 1 | 10.13 | 11.33 | 13.53 | 8.53 | 9.57 | 9.60 | 10.64 | 7.37 | 9.87 | 9.44 | 10.73 | 11.03 | 9.58 | 9.00 | 9.65 |
| ZNHIT6 | 8 | 9.64 | 10.41 | 7.95 | 8.19 | 9.94 | 10.19 | 11.28 | 10.04 | 11.21 | 11.15 | 10.03 | 8.07 | 10.06 | 10.66 | 11.18 |
| ZNRF2 | 1 | 10.14 | 12.64 | 8.29 | 8.52 | 12.77 | 11.92 | 8.47 | 9.36 | 8.59 | 9.31 | 11.39 | 8.40 | 12.34 | 8.92 | 8.95 |
| ZRANB2 | 12 | 8.57 | 8.99 | 10.66 | 8.97 | 8.20 | 7.97 | 10.99 | 17.89 | 9.16 | 8.60 | 8.78 | 9.82 | 8.08 | 14.44 | 8.88 |
| ZSWIM8 | 2 | 7.62 | 9.06 | 13.21 | 13.93 | 9.30 | 8.62 | 9.08 | 9.43 | 9.30 | 10.46 | 8.34 | 13.57 | 8.96 | 9.26 | 9.88 |
| ZW10 | 7 | 9.73 | 9.61 | 9.92 | 9.57 | 10.92 | 10.30 | 9.32 | 8.52 | 11.33 | 10.78 | 9.67 | 9.75 | 10.61 | 8.92 | 11.06 |
| ZWILCH | 12 | 10.71 | 11.16 | 9.62 | 8.68 | 10.16 | 10.37 | 8.15 | 7.22 | 12.07 | 11.87 | 10.93 | 9.15 | 10.26 | 7.69 | 11.97 |
| ZWINT | 7 | 11.57 | 12.11 | 10.56 | 9.60 | 9.63 | 9.22 | 9.14 | 7.41 | 10.71 | 10.05 | 11.84 | 10.08 | 9.42 | 8.27 | 10.38 |
| ZYG11B | 1 | 8.42 | 10.89 | 7.07 | 7.23 | 11.23 | 8.69 | 11.61 | 14.79 | 10.44 | 9.62 | 9.65 | 7.15 | 9.96 | 13.20 | 10.03 |
| ZYX | 36 | 9.15 | 10.52 | 16.69 | 14.82 | 6.59 | 6.42 | 5.29 | 5.28 | 12.80 | 12.43 | 9.84 | 15.75 | 6.50 | 5.29 | 12.62 |
| ZZEF1 | 9 | 10.74 | 10.69 | 11.15 | 11.17 | 8.40 | 8.29 | 8.13 | 8.60 | 11.16 | 11.66 | 10.71 | 11.16 | 8.35 | 8.37 | 11.41 |
| ZZZ3 | 3 | 9.51 | 9.22 | 12.99 | 10.36 | 8.34 | 7.57 | 11.85 | 10.18 | 9.35 | 10.63 | 9.37 | 11.68 | 7.96 | 11.01 | 9.99 |
