## Supplementary Table S2 for "Three ALS genes regulate expression of the MHC class II antigen presentation pathway"

**Table S2. Significantly dysregulated proteins in KO lines (p-value < 0.05).** Gene symbol, fold change (FC) and p-value are shown.

**FUS KO**

| Gene Symbol | FC | p-value |
|---|---|---|
| HLA-DRA | -12.96 | 1.8E-02 |
| HLA-DRB1 | -10.37 | 1.2E-02 |
| ALDH3A1 | -9.81 | 5.3E-03 |
| TPPP3 | -9.64 | 7.0E-03 |
| S100A14 | -8.97 | 1.2E-02 |
| SLC12A7 | -7.20 | 2.3E-02 |
| CPM | -6.90 | 1.4E-02 |
| TYRO3 | -6.68 | 3.5E-03 |
| MUC16 | -5.28 | 1.2E-03 |
| S100P | -4.93 | 1.9E-03 |
| S100A4 | -4.55 | 4.3E-02 |
| PSMB9 | -4.52 | 3.7E-03 |
| HLA-B | -4.32 | 2.8E-02 |
| PSMB10 | -4.23 | 2.3E-03 |
| FUS | -4.03 | 1.7E-03 |
| LXN | -3.78 | 1.7E-03 |
| TRIM29 | -3.78 | 2.3E-03 |
| DAB2 | -3.73 | 1.1E-02 |
| SAMD11 | -3.62 | 1.9E-05 |
| ASS1 | -3.53 | 2.0E-03 |
| GAA | -3.52 | 3.1E-03 |
| DNAJC25 | -3.51 | 2.0E-03 |
| NPR1 | -3.47 | 4.0E-02 |
| STAC2 | -3.45 | 8.0E-03 |
| CASP1 | -3.35 | 3.4E-02 |
| FOSB | -3.25 | 2.5E-02 |
| CLIC2 | -3.19 | 4.5E-03 |
| FLVCR1 | -3.06 | 4.5E-02 |
| SYNPO2 | -3.05 | 2.7E-02 |
| KLF4 | -3.03 | 3.1E-02 |
| MMRN1 | -3.03 | 4.9E-02 |
| H2AFY2 | -3.00 | 3.3E-04 |
| ANXA10 | -3.00 | 1.6E-04 |
| ADIRF | -2.96 | 2.9E-02 |
| GALT | -2.94 | 3.4E-03 |
| CRIP1 | -2.92 | 3.2E-03 |
| IMPA2 | -2.89 | 6.0E-03 |
| DENND5B | -2.84 | 6.8E-03 |
| ANGEL2 | -2.83 | 9.7E-04 |
| CPS1 | -2.80 | 4.0E-02 |
| ACADL | -2.76 | 2.9E-02 |
| PITX1 | -2.73 | 1.7E-02 |
| ZBED1 | -2.71 | 7.7E-04 |
| NQO1 | -2.70 | 2.0E-02 |
| MYPN | -2.68 | 1.4E-02 |
| NPTX1 | -2.64 | 2.5E-02 |
| MAP7D2 | -2.63 | 6.7E-03 |
| KCNAB2 | -2.63 | 5.2E-03 |
| FN3K | -2.57 | 7.8E-03 |
| SULT1A3 | -2.56 | 8.6E-03 |
| IFIT3 | -2.52 | 3.5E-02 |
| GPR56 | -2.51 | 3.0E-03 |
| S100A3 | -2.51 | 6.4E-03 |
| TENM1 | -2.50 | 3.2E-02 |
| L1CAM | -2.47 | 2.2E-02 |
| HLA-C | -2.44 | 4.8E-02 |
| RAP1GAP | -2.43 | 1.0E-02 |
| NMNAT1 | -2.43 | 1.7E-02 |
| MUC1 | -2.40 | 1.4E-02 |
| MAP1A | -2.37 | 1.4E-02 |
| PSMB8 | -2.30 | 7.3E-03 |

**EWSR1 KO**

| Gene Symbol | FC | p-value |
|---|---|---|
| EWSR1 | -15.0 | 2.7E-03 |
| CNN1 | -6.41 | 3.7E-03 |
| THY1 | -5.83 | 2.9E-02 |
| RFTN1 | -4.63 | 6.8E-03 |
| CGN | -4.60 | 1.9E-03 |
| SYNPO2 | -4.36 | 2.0E-02 |
| ALPL | -4.31 | 4.2E-04 |
| LEPREL2 | -4.07 | 9.2E-03 |
| EPHA7 | -4.05 | 3.3E-02 |
| LPL | -3.94 | 5.2E-03 |
| CLDN4 | -3.83 | 1.8E-02 |
| LIMA1 | -3.69 | 1.4E-02 |
| STAC2 | -3.32 | 8.3E-03 |
| CSDC2 | -3.06 | 1.7E-02 |
| TAGLN | -2.98 | 5.0E-03 |
| TOX | -2.93 | 1.1E-02 |
| FBLIM1 | -2.82 | 4.6E-04 |
| TGFBI | -2.71 | 7.2E-03 |
| GLIPR2 | -2.62 | 1.4E-04 |
| CAV1 | -2.59 | 2.7E-02 |
| EPDR1 | -2.56 | 8.1E-03 |
| COL1A1 | -2.53 | 1.9E-02 |
| CSRP2 | -2.52 | 3.1E-03 |
| C1orf198 | -2.43 | 2.0E-02 |
| CRABP2 | -2.40 | 3.2E-02 |
| SLC12A7 | -2.40 | 4.7E-02 |
| MOCOS | -2.40 | 3.8E-03 |
| UPP1 | -2.40 | 9.7E-03 |
| CNN3 | -2.38 | 9.0E-03 |
| INA | -2.38 | 8.9E-03 |
| MYLK | -2.37 | 1.0E-02 |
| MARCKS | -2.28 | 5.7E-03 |
| COL3A1 | -2.27 | 1.3E-02 |
| PRR22 | -2.27 | 3.5E-02 |
| NMNAT1 | -2.22 | 2.0E-02 |
| TNC | -2.21 | 2.1E-02 |
| MKL1 | -2.17 | 9.1E-04 |
| GSN | -2.17 | 1.7E-02 |
| PTPRF | -2.17 | 8.7E-03 |
| MYL9 | -2.14 | 1.6E-03 |
| HSCB | -2.14 | 5.8E-03 |
| KCTD12 | -2.12 | 5.0E-04 |
| SEC14L2 | -2.11 | 1.6E-02 |
| RAP1GAP | -2.11 | 8.5E-03 |
| FN1 | -2.10 | 5.4E-03 |
| ENOSF1 | -2.09 | 3.9E-03 |
| NPTX1 | -2.09 | 3.3E-02 |
| NES | -2.07 | 1.2E-02 |
| MAP2 | -2.03 | 9.0E-03 |
| A2M | -2.02 | 1.5E-02 |
| NFIX | -2.01 | 1.3E-03 |
| DNM1 | -2.01 | 7.9E-03 |
| NEGR1 | -2.00 | 2.6E-02 |
| SNCA | -1.99 | 1.4E-03 |
| PALM2 | -1.95 | 1.7E-02 |
| CDC45 | -1.91 | 3.0E-03 |
| MKNK1 | -1.91 | 1.3E-02 |
| HMOX1 | -1.90 | 3.6E-02 |
| JUN | -1.90 | 1.5E-02 |
| SLC2A1 | -1.89 | 4.0E-02 |
| TNS3 | -1.89 | 8.1E-03 |

**TAF15 KO**

| Gene Symbol | FC | p-value |
|---|---|---|
| CRYAB | -25.3 | 6.17E-03 |
| HLA-DRA | -17.4 | 1.75E-02 |
| TAF15 | -14.6 | 4.56E-04 |
| HLA-DRB1 | -11.1 | 1.23E-02 |
| S100P | -9.27 | 1.92E-03 |
| EPHA7 | -8.46 | 2.59E-02 |
| RFTN1 | -8.00 | 4.83E-03 |
| CNN1 | -7.75 | 2.75E-03 |
| GSN | -7.11 | 6.97E-03 |
| TAGLN | -6.87 | 3.14E-03 |
| NPTX1 | -6.79 | 1.26E-02 |
| QPRT | -6.76 | 4.39E-02 |
| SYNPO2 | -6.00 | 1.81E-02 |
| ALDH3A1 | -6.00 | 5.95E-03 |
| RGS3 | -5.15 | 3.73E-03 |
| STAC2 | -4.95 | 6.32E-03 |
| NPR1 | -4.75 | 3.27E-02 |
| AASS | -4.47 | 8.98E-03 |
| PSMB9 | -4.45 | 4.95E-04 |
| A2M | -4.38 | 6.08E-03 |
| IL18 | -4.28 | 1.23E-03 |
| MAP2 | -4.26 | 4.16E-03 |
| GBP1 | -4.20 | 5.84E-03 |
| SULT1A3 | -4.15 | 1.49E-03 |
| PPL | -4.12 | 8.60E-03 |
| C1orf198 | -4.05 | 1.27E-02 |
| NEXN | -4.00 | 3.22E-03 |
| CGN | -3.94 | 5.12E-03 |
| ALDH1A3 | -3.91 | 2.26E-02 |
| ITGA11 | -3.87 | 3.43E-02 |
| PSMB10 | -3.83 | 5.08E-03 |
| DYNC2LI1 | -3.78 | 6.25E-03 |
| SERPINB5 | -3.69 | 3.34E-03 |
| HPD | -3.65 | 1.54E-03 |
| AHNAK2 | -3.64 | 5.21E-03 |
| DFNA5 | -3.59 | 7.33E-04 |
| HSBP1 | -3.58 | 4.06E-02 |
| TUFT1 | -3.57 | 9.24E-03 |
| HOOK1 | -3.52 | 6.19E-03 |
| FSTL1 | -3.40 | 3.35E-03 |
| TPM1 | -3.26 | 3.28E-02 |
| FBLIM1 | -3.24 | 9.25E-03 |
| TGFB1I1 | -3.22 | 7.87E-03 |
| NMI | -3.13 | 2.91E-03 |
| SDC2 | -3.09 | 1.01E-02 |
| CLIC3 | -3.07 | 1.40E-02 |
| NQO1 | -3.06 | 1.62E-02 |
| PAPSS2 | -3.05 | 6.54E-03 |
| AFAP1 | -3.05 | 4.32E-03 |
| GLIPR2 | -3.03 | 1.29E-04 |
| MUC16 | -3.03 | 1.93E-03 |
| EPB41L1 | -3.02 | 5.63E-03 |
| CSDC2 | -2.99 | 1.71E-02 |
| PZP | -2.98 | 2.39E-03 |
| LXN | -2.98 | 4.38E-04 |
| MYH10 | -2.96 | 4.10E-02 |
| PRUNE2 | -2.94 | 1.66E-02 |
| ALDOC | -2.93 | 2.31E-03 |
| F3 | -2.92 | 1.47E-02 |
| TRIM29 | -2.92 | 6.89E-03 |
| ANXA8 | -2.91 | 4.32E-03 |

**MATR3 KO**

| Gene Symbol | FC | p-value |
|---|---|---|
| MATR3 | -11.7 | 3.78E-03 |
| HLA-DRA | -9.62 | 1.97E-02 |
| HLA-DRB1 | -8.31 | 1.32E-02 |
| UQCC1 | -8.03 | 3.20E-03 |
| TESC | -5.19 | 3.35E-02 |
| CLIC2 | -4.59 | 3.61E-03 |
| MGMT | -4.41 | 4.28E-02 |
| HNMT | -4.39 | 2.97E-03 |
| PPP1R12B | -4.23 | 2.57E-02 |
| WIBG | -4.04 | 3.21E-03 |
| UQCC2 | -4.02 | 2.15E-02 |
| SCLY | -3.90 | 8.88E-04 |
| FHIT | -3.63 | 1.60E-02 |
| ACAD9 | -3.49 | 5.34E-03 |
| MAP2 | -3.18 | 5.09E-03 |
| S100P | -3.17 | 2.76E-03 |
| PSMB9 | -3.00 | 1.12E-02 |
| MND1 | -2.84 | 6.27E-03 |
| TYRO3 | -2.83 | 6.10E-03 |
| SHCBP1 | -2.77 | 4.12E-03 |
| ANXA10 | -2.76 | 1.73E-03 |
| ADARB1 | -2.73 | 7.65E-03 |
| ALDH5A1 | -2.66 | 1.14E-02 |
| CPM | -2.63 | 2.62E-02 |
| SAMD11 | -2.61 | 1.16E-03 |
| MARCKSL1 | -2.59 | 2.38E-02 |
| CASP1 | -2.55 | 4.40E-02 |
| PSMB10 | -2.54 | 5.67E-03 |
| UHRF2 | -2.53 | 3.36E-02 |
| DAB2 | -2.49 | 1.61E-02 |
| NMI | -2.44 | 4.67E-03 |
| DTX3L | -2.43 | 3.96E-02 |
| CTSC | -2.42 | 1.83E-03 |
| ALPL | -2.41 | 6.98E-04 |
| ACYP1 | -2.31 | 1.03E-03 |
| KIAA1211 | -2.31 | 5.71E-03 |
| UQCRB | -2.30 | 2.33E-02 |
| ADAT2 | -2.27 | 2.65E-03 |
| TRIM29 | -2.26 | 4.82E-03 |
| PRR22 | -2.24 | 4.29E-02 |
| VPS13A | -2.23 | 8.59E-04 |
| AASS | -2.21 | 1.67E-02 |
| SYNPO2 | -2.21 | 4.14E-02 |
| GMPR2 | -2.20 | 3.99E-03 |
| LXN | -2.19 | 8.16E-03 |
| CSRP2 | -2.17 | 5.50E-03 |
| MYPN | -2.16 | 2.12E-02 |
| SNCA | -2.14 | 2.69E-03 |
| CMBL | -2.14 | 1.65E-03 |
| SLFN5 | -2.13 | 1.13E-04 |
| COPG2 | -2.12 | 7.35E-03 |
| SS18L1 | -2.11 | 4.24E-02 |
| RBM41 | -2.09 | 1.43E-02 |
| MSRA | -2.08 | 3.88E-02 |
| MYLK | -2.07 | 1.24E-02 |
| UQCRC2 | -2.07 | 2.50E-02 |
| TXNRD2 | -2.07 | 8.59E-03 |
| CLYBL | -2.05 | 1.78E-02 |
| GLB1 | -2.04 | 1.73E-03 |
| RALGAPB | -2.01 | 2.00E-03 |
| PSMC3IP | -2.01 | 6.54E-03 |

| | | |
|---|---|---|
| ENO3 | -2.30 | 1.8E-02 |
| MSLN | -2.24 | 8.7E-03 |
| NAPRT1 | -2.21 | 9.4E-03 |
| AKR1C2 | -2.20 | 1.9E-02 |
| BEGAIN | -2.18 | 5.9E-03 |
| ALDH5A1 | -2.15 | 2.2E-02 |
| EML1 | -2.15 | 4.8E-02 |
| FAM111A | -2.14 | 4.2E-02 |
| CLU | -2.14 | 1.1E-02 |
| LPL | -2.13 | 1.0E-02 |
| COBLL1 | -2.12 | 1.8E-02 |
| BLVRB | -2.12 | 2.7E-02 |
| DTX3L | -2.11 | 4.7E-02 |
| IQGAP2 | -2.11 | 1.4E-02 |
| TRAFD1 | -2.08 | 2.3E-02 |
| CLIC3 | -2.08 | 2.8E-02 |
| PTER | -2.07 | 7.9E-03 |
| KIF1A | -2.03 | 1.8E-02 |
| CSRP2 | -2.02 | 8.9E-03 |
| DSTYK | -2.01 | 2.6E-02 |
| PALM | -2.00 | 4.3E-02 |
| FTL | -2.00 | 2.7E-02 |
| ACADS | -1.98 | 4.2E-02 |
| MAP7 | -1.98 | 1.6E-02 |
| CDA | -1.98 | 1.5E-02 |
| FYTTD1 | -1.98 | 1.1E-02 |
| SH3GL2 | -1.98 | 5.9E-04 |
| PRKACA | -1.97 | 2.9E-02 |
| MFF | -1.95 | 1.1E-02 |
| ACOT11 | -1.95 | 4.3E-03 |
| TUBAL3 | -1.94 | 2.1E-02 |
| RAET1G | -1.91 | 2.3E-02 |
| SRXN1 | -1.90 | 3.2E-02 |
| CRAT | -1.90 | 3.4E-02 |
| RNASEL | -1.89 | 3.8E-03 |
| GUSB | -1.88 | 6.0E-03 |
| HMOX1 | -1.88 | 3.5E-02 |
| BPHL | -1.88 | 1.4E-02 |
| SELM | -1.88 | 1.4E-02 |
| COL3A1 | -1.87 | 1.8E-02 |
| FOLR1 | -1.86 | 1.2E-02 |
| RDX | -1.86 | 4.7E-03 |
| S100A2 | -1.85 | 1.6E-02 |
| NQO2 | -1.85 | 1.7E-02 |
| KYNU | -1.85 | 1.5E-02 |
| NECAP1 | -1.85 | 1.7E-02 |
| SMIM7 | -1.83 | 4.8E-02 |
| RGS3 | -1.82 | 3.2E-02 |
| SMEK3P | -1.82 | 3.3E-04 |
| PIR | -1.81 | 1.0E-02 |
| CMBL | -1.81 | 5.9E-03 |
| DPP7 | -1.81 | 2.0E-03 |
| ACTR1B | -1.80 | 1.0E-02 |
| SCAF4 | -1.78 | 2.0E-02 |
| CCS | -1.78 | 2.9E-02 |
| CDKN2C | -1.77 | 2.6E-02 |
| CUEDC2 | -1.77 | 1.9E-02 |
| LEPREL2 | -1.77 | 3.6E-02 |
| FH | -1.77 | 4.3E-03 |
| DNMBP | -1.77 | 1.0E-03 |
| DYNC2LI1 | -1.77 | 2.3E-02 |
| RASSF7 | -1.76 | 4.0E-02 |
| PALMD | -1.76 | 2.5E-02 |
| ENO2 | -1.75 | 4.6E-02 |
| AHNAK2 | -1.75 | 1.4E-02 |
| GSE1 | -1.75 | 3.8E-03 |
| CSPG4 | -1.75 | 2.0E-02 |
| CRLF1 | -1.74 | 3.1E-02 |

| | | |
|---|---|---|
| TYMP | -1.87 | 3.0E-02 |
| MAOB | -1.84 | 9.0E-03 |
| LAMA4 | -1.83 | 3.4E-02 |
| NEXN | -1.83 | 9.5E-03 |
| MAP1B | -1.82 | 1.4E-02 |
| PTER | -1.82 | 1.1E-02 |
| CALD1 | -1.81 | 8.2E-03 |
| TUBD1 | -1.80 | 7.9E-03 |
| SMTN | -1.80 | 8.1E-03 |
| TBC1D5 | -1.79 | 3.9E-03 |
| CYR61 | -1.79 | 2.4E-02 |
| HIP1 | -1.79 | 3.4E-03 |
| THBS1 | -1.77 | 4.3E-02 |
| APOL2 | -1.77 | 1.1E-02 |
| TGM2 | -1.76 | 6.8E-03 |
| NXN | -1.75 | 2.3E-02 |
| BCAM | -1.74 | 9.3E-03 |
| XPNPEP3 | -1.73 | 2.2E-02 |
| SH3BP4 | -1.72 | 2.1E-03 |
| NRP1 | -1.71 | 3.8E-02 |
| GSTM3 | -1.71 | 2.7E-03 |
| CHEK2 | -1.71 | 3.6E-02 |
| RUNX3 | -1.70 | 4.3E-02 |
| TK1 | -1.69 | 1.1E-02 |
| DENND5B | -1.69 | 2.6E-02 |
| DPH5 | -1.68 | 7.6E-03 |
| SDC1 | -1.68 | 1.5E-02 |
| IGF2BP3 | -1.68 | 1.8E-02 |
| TRIM29 | -1.67 | 1.0E-02 |
| ASF1B | -1.67 | 6.6E-03 |
| ALDOC | -1.66 | 4.7E-03 |
| FAM133B | -1.66 | 4.9E-03 |
| PARVB | -1.66 | 1.8E-03 |
| BICC1 | -1.66 | 4.4E-02 |
| CAMKK1 | -1.65 | 1.5E-02 |
| TGFB1I1 | -1.64 | 2.3E-02 |
| CECR5 | -1.64 | 1.5E-02 |
| COL5A2 | -1.64 | 3.9E-02 |
| SVIL | -1.63 | 7.0E-03 |
| FRMPD1 | -1.63 | 4.0E-02 |
| GSTZ1 | -1.63 | 1.4E-03 |
| ATAD5 | -1.63 | 3.4E-02 |
| AKAP2 | -1.63 | 1.9E-02 |
| AARSD1 | -1.62 | 1.2E-02 |
| PTRF | -1.62 | 4.8E-03 |
| CDKN2AIPN | -1.62 | 2.4E-02 |
| GFPT2 | -1.62 | 1.6E-03 |
| ARHGAP29 | -1.62 | 6.1E-03 |
| MYH14 | -1.62 | 3.8E-03 |
| CTGF | -1.62 | 3.8E-02 |
| IQGAP2 | -1.62 | 1.6E-02 |
| LRRC40 | -1.59 | 2.0E-02 |
| RILPL1 | -1.59 | 2.6E-02 |
| MBIP | -1.59 | 1.5E-02 |
| ANO6 | -1.59 | 2.8E-02 |
| MYO5A | -1.58 | 3.4E-03 |
| TRMT2A | -1.57 | 1.3E-02 |
| ME1 | -1.57 | 1.6E-03 |
| CRYZ | -1.57 | 3.6E-03 |
| DBNL | -1.57 | 4.2E-02 |
| ANXA6 | -1.57 | 5.4E-03 |
| SHB | -1.56 | 5.3E-03 |
| hCG_198816 | -1.56 | 1.5E-02 |
| ACACA | -1.56 | 3.1E-03 |
| GINS4 | -1.56 | 9.7E-03 |
| CYTH3 | -1.56 | 9.5E-04 |
| ZNF346 | -1.55 | 9.5E-03 |
| GCLM | -1.55 | 6.5E-03 |

| | | |
|---|---|---|
| CRIP1 | -2.89 | 5.01E-04 |
| ANXA3 | -2.88 | 1.50E-03 |
| CYR61 | -2.88 | 1.15E-02 |
| LPL | -2.84 | 7.85E-03 |
| LASP1 | -2.82 | 1.42E-02 |
| DCLK2 | -2.81 | 2.36E-02 |
| TRIM16 | -2.77 | 7.15E-04 |
| PLS1 | -2.75 | 1.28E-02 |
| PALLD | -2.70 | 5.36E-03 |
| ADIRF | -2.68 | 8.43E-03 |
| CBR3 | -2.67 | 1.74E-02 |
| PYCARD | -2.66 | 1.23E-02 |
| HMOX1 | -2.64 | 2.59E-02 |
| ACOX3 | -2.63 | 9.85E-03 |
| TYRO3 | -2.60 | 7.93E-03 |
| SHPK | -2.57 | 2.23E-02 |
| MYPN | -2.55 | 1.86E-02 |
| PALMD | -2.55 | 1.23E-02 |
| COBL | -2.53 | 7.15E-03 |
| CAP2 | -2.53 | 1.70E-02 |
| KCTD15 | -2.52 | 3.06E-03 |
| MOCOS | -2.52 | 3.12E-03 |
| CYTH3 | -2.51 | 1.23E-02 |
| DNAJB4 | -2.51 | 3.23E-03 |
| OAS3 | -2.51 | 5.53E-04 |
| CTGF | -2.50 | 1.94E-02 |
| TRAFD1 | -2.50 | 1.89E-02 |
| TACSTD2 | -2.50 | 2.11E-02 |
| MYL9 | -2.49 | 4.15E-03 |
| GPR56 | -2.48 | 2.39E-05 |
| TCP11L1 | -2.47 | 1.11E-02 |
| CDH2 | -2.42 | 4.72E-02 |
| TXNRD1 | -2.41 | 5.46E-03 |
| PEA15 | -2.40 | 4.26E-02 |
| CAMK4 | -2.40 | 2.94E-02 |
| DAB2 | -2.39 | 1.76E-02 |
| SH3GL2 | -2.39 | 2.81E-03 |
| COL1A1 | -2.38 | 3.24E-02 |
| CNN2 | -2.37 | 1.25E-02 |
| SQSTM1 | -2.37 | 3.02E-02 |
| TNS3 | -2.36 | 6.26E-03 |
| TJP2 | -2.35 | 4.36E-03 |
| RND3 | -2.34 | 8.49E-04 |
| STON2 | -2.34 | 2.09E-02 |
| FAM195B | -2.32 | 3.35E-03 |
| SYNGR2 | -2.32 | 7.69E-03 |
| COBLL1 | -2.29 | 1.52E-02 |
| VASN | -2.28 | 3.66E-03 |
| CASKIN2 | -2.28 | 1.05E-02 |
| MICAL1 | -2.27 | 5.32E-03 |
| FECH | -2.26 | 2.71E-02 |
| PRKACA | -2.25 | 2.16E-02 |
| DAG1 | -2.25 | 8.52E-03 |
| ALDH1B1 | -2.24 | 3.31E-02 |
| PCBD1 | -2.24 | 1.31E-02 |
| APOL2 | -2.24 | 7.99E-03 |
| TOM1L1 | -2.24 | 1.80E-02 |
| FN3K | -2.23 | 1.05E-02 |
| LSS | -2.23 | 1.62E-03 |
| DDAH1 | -2.23 | 5.61E-04 |
| FARP1 | -2.22 | 5.34E-03 |
| OBSL1 | -2.21 | 1.17E-02 |
| PTS | -2.21 | 5.85E-03 |
| LIMA1 | -2.19 | 2.89E-02 |
| TDRD7 | -2.19 | 6.88E-04 |
| SNCG | -2.19 | 5.98E-03 |
| WWC2 | -2.19 | 2.52E-02 |
| PHLDB1 | -2.17 | 2.27E-02 |

| | | |
|---|---|---|
| TOP3B | -2.00 | 5.04E-04 |
| TSHZ2 | -2.00 | 9.30E-03 |
| MAOB | -2.00 | 2.74E-03 |
| VPS8 | -1.98 | 2.04E-02 |
| FN3K | -1.98 | 8.44E-03 |
| RNASEL | -1.98 | 1.98E-02 |
| HMGN5 | -1.97 | 3.59E-02 |
| SGSH | -1.95 | 6.37E-03 |
| HEXB | -1.93 | 1.61E-03 |
| KHDRBS3 | -1.89 | 8.81E-04 |
| SMYD3 | -1.88 | 1.22E-02 |
| NDUFAF1 | -1.88 | 3.74E-02 |
| HIBCH | -1.88 | 4.96E-02 |
| TRAPPC10 | -1.87 | 1.78E-02 |
| KYNU | -1.87 | 4.40E-03 |
| ZBED1 | -1.86 | 1.85E-03 |
| ALDOC | -1.84 | 5.31E-03 |
| MTUS1 | -1.83 | 2.35E-02 |
| GAA | -1.82 | 8.91E-03 |
| SIAE | -1.80 | 3.18E-02 |
| FAM46A | -1.80 | 4.21E-03 |
| TCEAL5 | -1.80 | 3.91E-02 |
| RNASET2 | -1.79 | 4.98E-03 |
| MAP2K6 | -1.79 | 1.18E-02 |
| TMOD1 | -1.79 | 1.71E-02 |
| RWDD1 | -1.78 | 1.06E-02 |
| TANC2 | -1.72 | 3.70E-03 |
| BDH2 | -1.72 | 7.43E-03 |
| MAP4K5 | -1.72 | 2.28E-02 |
| PIR | -1.71 | 8.39E-03 |
| CNN1 | -1.71 | 1.21E-02 |
| PITX1 | -1.71 | 4.02E-02 |
| DNM1 | -1.71 | 1.58E-02 |
| PARP4 | -1.70 | 2.94E-04 |
| CASP9 | -1.70 | 1.95E-02 |
| GGH | -1.69 | 4.04E-03 |
| EML1 | -1.69 | 4.55E-03 |
| SEC11C | -1.67 | 4.94E-02 |
| KIAA1598 | -1.66 | 2.28E-02 |
| NSUN6 | -1.66 | 2.22E-02 |
| KIF1A | -1.65 | 3.06E-02 |
| PROSER1 | -1.65 | 2.38E-03 |
| CAPN1 | -1.64 | 1.21E-02 |
| SPC25 | -1.64 | 1.19E-02 |
| NEGR1 | -1.64 | 4.30E-02 |
| PSMB8 | -1.63 | 1.80E-02 |
| CDKN2C | -1.63 | 3.33E-02 |
| SMPDL3A | -1.63 | 9.70E-03 |
| EEFSEC | -1.63 | 2.10E-02 |
| ECSIT | -1.62 | 3.03E-02 |
| PTER | -1.61 | 1.65E-02 |
| SPG20 | -1.61 | 1.07E-02 |
| TRIP4 | -1.61 | 2.10E-02 |
| UQCRC1 | -1.60 | 2.21E-02 |
| GMDS | -1.59 | 1.39E-02 |
| PIBF1 | -1.59 | 6.95E-03 |
| NADSYN1 | -1.59 | 4.97E-02 |
| NR4A1 | -1.59 | 1.83E-02 |
| TSSC1 | -1.59 | 2.18E-02 |
| ATG7 | -1.58 | 2.01E-04 |
| HLTF | -1.58 | 1.58E-02 |
| CHCHD4 | -1.57 | 1.95E-02 |
| IFI16 | -1.57 | 8.32E-03 |
| MAP7 | -1.56 | 1.82E-02 |
| PSME1 | -1.56 | 2.36E-02 |
| HS1BP3 | -1.56 | 8.76E-03 |
| FZD10 | -1.55 | 3.92E-02 |
| SELH | -1.55 | 2.07E-02 |

| | | |
|---|---|---|
| RAPGEF1 | -1.74 | 4.4E-02 |
| SELENBP1 | -1.71 | 1.2E-02 |
| OAS3 | -1.70 | 8.7E-04 |
| AKR7A2 | -1.69 | 2.1E-02 |
| ANXA8 | -1.69 | 1.2E-02 |
| TSHZ2 | -1.69 | 1.0E-03 |
| TACSTD2 | -1.69 | 1.6E-02 |
| PKP3 | -1.67 | 2.2E-02 |
| FAIM | -1.67 | 5.8E-03 |
| CASP9 | -1.67 | 2.5E-02 |
| C1orf123 | -1.67 | 2.7E-02 |
| PEG10 | -1.67 | 9.1E-03 |
| P4HTM | -1.67 | 1.4E-02 |
| RECQL4 | -1.66 | 6.6E-05 |
| XYLB | -1.66 | 1.1E-02 |
| TERF2IP | -1.66 | 4.5E-03 |
| G6PD | -1.66 | 3.2E-02 |
| PDE8B | -1.65 | 1.4E-02 |
| C21orf33 | -1.64 | 4.3E-02 |
| ADARB1 | -1.64 | 2.8E-02 |
| CFDP1 | -1.64 | 3.5E-03 |
| GNPAT | -1.64 | 8.1E-03 |
| KCTD15 | -1.63 | 4.0E-03 |
| COBL | -1.63 | 1.6E-02 |
| COQ9 | -1.63 | 2.1E-02 |
| MAN2B1 | -1.63 | 1.8E-03 |
| CBFB | -1.63 | 5.4E-03 |
| STXBP1 | -1.63 | 1.2E-02 |
| MYLK | -1.63 | 2.8E-02 |
| FTO | -1.62 | 1.0E-02 |
| ACADVL | -1.62 | 2.9E-02 |
| C16orf87 | -1.62 | 3.8E-03 |
| HSDL1 | -1.62 | 1.2E-02 |
| SQSTM1 | -1.61 | 4.9E-02 |
| ODC1 | -1.60 | 1.6E-02 |
| RNASET2 | -1.60 | 7.7E-03 |
| DHRS4 | -1.60 | 5.8E-03 |
| C9orf142 | -1.60 | 4.0E-02 |
| PSMD10 | -1.60 | 4.5E-02 |
| PARP12 | -1.60 | 3.5E-03 |
| MAEA | -1.59 | 2.0E-02 |
| CASP2 | -1.59 | 2.2E-04 |
| TSC22D3 | -1.59 | 3.0E-03 |
| TLX3 | -1.59 | 4.8E-02 |
| PLEKHG4 | -1.59 | 7.9E-03 |
| PITHD1 | -1.59 | 2.6E-02 |
| TBC1D17 | -1.58 | 6.8E-03 |
| PRMT7 | -1.58 | 4.1E-02 |
| ZNF346 | -1.58 | 5.1E-03 |
| CBR3 | -1.58 | 4.4E-02 |
| HOXC10 | -1.57 | 6.0E-03 |
| AGL | -1.57 | 2.0E-03 |
| TNS4 | -1.57 | 4.4E-02 |
| HOOK1 | -1.57 | 2.3E-02 |
| STARD3NL | -1.57 | 5.3E-03 |
| PLEKHG2 | -1.57 | 2.8E-03 |
| WDR55 | -1.57 | 6.6E-03 |
| DDX59 | -1.57 | 4.2E-02 |
| HIP1 | -1.56 | 4.2E-03 |
| FAM203A | -1.56 | 2.7E-02 |
| CAMKK1 | -1.56 | 1.7E-02 |
| KIAA1211 | -1.56 | 3.0E-02 |
| NFIA | -1.56 | 8.4E-03 |
| ID3 | -1.56 | 4.3E-02 |
| BAG1 | -1.56 | 8.9E-03 |
| STON2 | -1.56 | 1.1E-02 |
| PYCR2 | -1.55 | 2.5E-02 |
| GLUL | -1.55 | 2.6E-02 |

| | | |
|---|---|---|
| STXBP4 | -1.55 | 1.7E-02 |
| CENPM | -1.55 | 2.0E-02 |
| PLA2G4B | -1.55 | 1.5E-02 |
| PTK7 | -1.55 | 4.5E-03 |
| PAFAH2 | -1.54 | 1.1E-02 |
| TRMT13 | -1.54 | 9.8E-03 |
| MICAL1 | -1.54 | 1.5E-02 |
| DPH2 | -1.53 | 1.7E-03 |
| ADAL | -1.53 | 1.8E-02 |
| EPS8 | -1.53 | 1.6E-02 |
| AKR1C2 | -1.53 | 2.6E-02 |
| CORO2A | -1.53 | 3.2E-03 |
| EIF2B3 | -1.52 | 8.4E-03 |
| ABR | -1.52 | 2.7E-04 |
| MMACHC | -1.52 | 9.4E-03 |
| MACF1 | -1.52 | 1.4E-02 |
| ARHGAP23 | -1.51 | 1.5E-02 |
| ZYX | -1.51 | 4.0E-02 |
| FAHD2A | -1.51 | 1.1E-02 |
| SUMF2 | -1.51 | 1.3E-02 |
| USP13 | -1.51 | 2.3E-02 |
| WWC2 | -1.51 | 4.2E-02 |
| ALDH3A1 | -1.51 | 3.5E-02 |
| TLN1 | -1.51 | 3.1E-02 |
| FAM195B | -1.51 | 1.8E-02 |
| MYH9 | -1.51 | 3.3E-02 |
| IFRD1 | -1.50 | 2.5E-02 |
| FSTL1 | -1.50 | 9.7E-03 |
| HIBADH | -1.50 | 3.3E-02 |
| ACOX3 | -1.50 | 2.9E-02 |
| KCTD5 | -1.50 | 5.2E-03 |
| TXNRD1 | -1.50 | 1.5E-02 |
| FOXJ3 | -1.50 | 1.3E-02 |
| NCS1 | -1.49 | 1.0E-02 |
| CDC42EP1 | -1.49 | 1.6E-02 |
| PPIL2 | -1.49 | 1.1E-02 |
| MTHFD2 | -1.48 | 1.1E-02 |
| CASP2 | -1.48 | 1.2E-03 |
| YARS | -1.48 | 7.5E-03 |
| RHEB | -1.48 | 1.2E-02 |
| PDLIM5 | -1.48 | 1.9E-02 |
| BCL10 | -1.48 | 2.4E-02 |
| UAP1L1 | -1.48 | 1.0E-02 |
| FHL3 | -1.48 | 3.2E-02 |
| PGD | -1.48 | 4.9E-02 |
| AK4 | -1.48 | 2.7E-02 |
| CLIP2 | -1.47 | 5.5E-03 |
| GCLC | -1.47 | 7.1E-03 |
| CEP41 | -1.47 | 5.0E-02 |
| DUSP14 | -1.47 | 3.8E-03 |
| PALMD | -1.47 | 3.6E-02 |
| PDK1 | -1.47 | 4.0E-03 |
| FKBP7 | -1.47 | 1.7E-02 |
| STX2 | -1.46 | 2.6E-02 |
| GPN2 | -1.46 | 2.9E-02 |
| VASN | -1.46 | 1.3E-02 |
| EHD2 | -1.46 | 2.8E-02 |
| ALDH1A2 | -1.45 | 1.8E-02 |
| CTNNB1 | -1.45 | 8.2E-03 |
| ARMC9 | -1.45 | 1.0E-02 |
| UQCC1 | -1.45 | 4.8E-02 |
| PCYT2 | -1.45 | 6.4E-03 |
| ORC6 | -1.45 | 2.5E-03 |
| PSMC3IP | -1.44 | 7.1E-03 |
| AEBP2 | -1.44 | 1.7E-02 |
| CTPS1 | -1.44 | 9.9E-03 |
| ZNF326 | -1.44 | 2.6E-02 |
| TTC4 | -1.44 | 3.2E-02 |

| | | |
|---|---|---|
| EVPL | -2.15 | 9.79E-03 |
| CUEDC1 | -2.15 | 4.52E-02 |
| KANK1 | -2.15 | 7.50E-03 |
| KANK2 | -2.15 | 1.25E-02 |
| DNM1 | -2.15 | 4.45E-03 |
| PCYT2 | -2.14 | 4.02E-03 |
| TP53I3 | -2.13 | 1.03E-02 |
| FABP3 | -2.12 | 8.34E-03 |
| HSPA12A | -2.12 | 6.13E-03 |
| TUB | -2.11 | 1.36E-02 |
| PRKAR1A | -2.11 | 1.59E-02 |
| CDR2L | -2.10 | 2.23E-02 |
| HDHD2 | -2.09 | 1.72E-02 |
| APBB1 | -2.09 | 3.52E-03 |
| FGD4 | -2.09 | 1.97E-03 |
| SDC1 | -2.09 | 8.95E-03 |
| HSPB8 | -2.09 | 3.20E-02 |
| DUSP14 | -2.09 | 8.50E-04 |
| FRMPD1 | -2.07 | 2.63E-02 |
| GLUL | -2.07 | 5.62E-03 |
| PEAK1 | -2.05 | 5.61E-03 |
| OPTN | -2.05 | 6.47E-03 |
| ATF3 | -2.04 | 3.03E-03 |
| MSLN | -2.02 | 1.20E-02 |
| MALT1 | -2.02 | 5.54E-03 |
| ENOSF1 | -2.02 | 1.03E-02 |
| ROCK2 | -2.01 | 7.98E-03 |
| SNX24 | -2.01 | 7.94E-04 |
| GALT | -2.00 | 8.78E-03 |
| MLLT4 | -2.00 | 1.63E-02 |
| TSC22D3 | -2.00 | 2.48E-02 |
| CCDC50 | -1.99 | 1.44E-02 |
| CNDP2 | -1.99 | 1.53E-02 |
| THBS1 | -1.99 | 3.74E-02 |
| ADCY9 | -1.99 | 3.81E-02 |
| CDR2 | -1.98 | 2.29E-02 |
| MYO1C | -1.97 | 1.00E-02 |
| KCTD12 | -1.97 | 3.67E-03 |
| TAX1BP3 | -1.96 | 2.35E-02 |
| MYL6 | -1.96 | 4.43E-02 |
| MAP1B | -1.96 | 1.46E-02 |
| CNN3 | -1.95 | 1.29E-02 |
| DST | -1.95 | 2.21E-03 |
| PROSER2 | -1.95 | 7.85E-03 |
| MKL1 | -1.95 | 6.33E-03 |
| ANXA6 | -1.94 | 3.37E-03 |
| TNFAIP8 | -1.94 | 3.96E-02 |
| MYO5A | -1.93 | 8.83E-03 |
| MAP1LC3B2 | -1.92 | 3.34E-03 |
| ARHGAP17 | -1.92 | 1.08E-02 |
| MTSS1L | -1.92 | 1.30E-02 |
| ALDH5A1 | -1.92 | 1.87E-02 |
| PSMB8 | -1.92 | 6.41E-03 |
| TSC22D2 | -1.91 | 4.87E-02 |
| KLHL21 | -1.91 | 5.95E-03 |
| STOM | -1.91 | 4.81E-02 |
| MISP | -1.90 | 5.11E-03 |
| CYTH1 | -1.90 | 5.76E-03 |
| SELENBP1 | -1.89 | 1.18E-02 |
| TMOD1 | -1.89 | 1.44E-02 |
| MYH14 | -1.88 | 2.56E-03 |
| PDLIM5 | -1.88 | 1.37E-02 |
| CAMK1 | -1.87 | 5.75E-03 |
| ZYX | -1.86 | 2.19E-02 |
| PDE8B | -1.86 | 1.07E-02 |
| ALDH1A2 | -1.86 | 1.27E-02 |
| ME2 | -1.86 | 1.62E-02 |
| CAV1 | -1.85 | 4.58E-02 |

| | | |
|---|---|---|
| CLN5 | -1.55 | 5.50E-03 |
| RNASEH2B | -1.55 | 2.13E-02 |
| NAT1 | -1.55 | 1.98E-02 |
| SH3BGRL | -1.55 | 3.97E-02 |
| PDE6D | -1.54 | 1.06E-02 |
| TSC22D3 | -1.54 | 1.03E-03 |
| CARS2 | -1.54 | 2.66E-02 |
| COQ7 | -1.54 | 1.54E-02 |
| PSTPIP2 | -1.54 | 3.67E-02 |
| RDX | -1.53 | 1.02E-02 |
| GCFC2 | -1.53 | 6.84E-04 |
| SUGP2 | -1.53 | 3.56E-02 |
| CLU | -1.53 | 2.13E-02 |
| TDRD7 | -1.53 | 1.79E-02 |
| PDE8B | -1.53 | 1.65E-02 |
| GGT1 | -1.53 | 8.72E-03 |
| EXOC2 | -1.52 | 2.48E-03 |
| GOLPH3L | -1.52 | 2.78E-02 |
| CPZ | -1.52 | 3.96E-02 |
| NUDT15 | -1.52 | 2.89E-03 |
| TAMM41 | -1.52 | 4.94E-02 |
| DRG2 | -1.51 | 1.76E-02 |
| NUP35 | -1.51 | 1.01E-02 |
| ARID2 | -1.51 | 2.43E-02 |
| ASF1B | -1.51 | 4.79E-02 |
| COQ5 | -1.51 | 1.93E-02 |
| TTC19 | -1.51 | 2.15E-02 |
| KCTD15 | -1.50 | 5.46E-03 |
| PLBD2 | -1.50 | 1.43E-02 |
| CCDC77 | -1.50 | 1.22E-02 |
| NQO2 | -1.50 | 2.96E-02 |
| UNK | -1.49 | 3.83E-02 |
| BRD7 | -1.49 | 3.40E-02 |
| CCDC41 | -1.49 | 1.55E-02 |
| BRCC3 | -1.49 | 4.92E-03 |
| FAIM | -1.48 | 5.58E-03 |
| SMG6 | -1.48 | 1.71E-02 |
| FAM115A | -1.48 | 1.27E-02 |
| EPB41 | -1.48 | 2.78E-02 |
| CASP2 | -1.48 | 4.49E-04 |
| H2AFY2 | -1.48 | 1.42E-02 |
| MKNK1 | -1.47 | 4.35E-03 |
| SCLT1 | -1.47 | 1.49E-02 |
| MAT2B | -1.47 | 1.69E-02 |
| APOL2 | -1.47 | 2.05E-02 |
| TFPI | -1.47 | 1.39E-02 |
| EHHADH | -1.46 | 1.92E-02 |
| PIN4 | -1.46 | 3.46E-02 |
| VPS36 | -1.46 | 9.91E-03 |
| FDFT1 | -1.46 | 2.79E-02 |
| RPL22L1 | -1.46 | 1.48E-03 |
| C19orf66 | -1.46 | 2.34E-02 |
| ARSA | -1.46 | 1.93E-03 |
| RALGAPA1 | -1.46 | 1.33E-02 |
| BCAS3 | -1.45 | 2.05E-02 |
| A2M | -1.45 | 3.52E-02 |
| HMGCS1 | -1.45 | 1.41E-02 |
| HTATSF1 | -1.45 | 1.70E-02 |
| WDR33 | -1.45 | 1.19E-02 |
| GUSB | -1.45 | 1.14E-02 |
| GPX4 | -1.45 | 1.33E-02 |
| CDC25C | -1.45 | 2.66E-02 |
| NT5C3A | -1.44 | 2.75E-03 |
| HOOK1 | -1.44 | 3.23E-02 |
| DIS3L2 | -1.44 | 1.34E-02 |
| GLA | -1.44 | 1.56E-02 |
| IDH2 | -1.44 | 1.75E-02 |
| TRIM38 | -1.44 | 7.31E-03 |

| | | |
|---|---|---|
| TMOD1 | -1.55 | 2.3E-02 |
| CRYZL1 | -1.55 | 1.6E-02 |
| EHHADH | -1.55 | 2.0E-02 |
| NCKIPSD | -1.55 | 1.8E-02 |
| PRDX4 | -1.54 | 2.9E-03 |
| SKP2 | -1.54 | 1.4E-02 |
| NKTR | -1.54 | 1.6E-02 |
| PDCD6 | -1.54 | 2.6E-02 |
| MAT2B | -1.54 | 1.0E-02 |
| C9orf40 | -1.54 | 2.0E-02 |
| PHYH | -1.53 | 2.6E-02 |
| TRIM4 | -1.53 | 1.2E-03 |
| OGFOD1 | -1.53 | 2.4E-02 |
| CIAPIN1 | -1.53 | 1.2E-02 |
| UACA | -1.53 | 1.4E-02 |
| OSBP2 | -1.53 | 2.2E-03 |
| GUK1 | -1.53 | 4.8E-03 |
| NSMF | -1.52 | 4.6E-02 |
| ALKBH4 | -1.52 | 1.6E-02 |
| KCTD9 | -1.52 | 8.5E-03 |
| UNC119B | -1.52 | 1.7E-02 |
| HMGN1 | -1.52 | 1.6E-03 |
| SEMA3B | -1.51 | 4.8E-02 |
| ITSN1 | -1.51 | 4.3E-03 |
| SFN | -1.50 | 4.5E-02 |
| ACYP2 | -1.50 | 1.2E-03 |
| GNB1 | -1.50 | 1.4E-02 |
| SNCG | -1.50 | 9.0E-03 |
| HADH | -1.50 | 4.4E-02 |
| HPD | -1.50 | 5.4E-03 |
| PARP9 | -1.50 | 1.8E-02 |
| C2orf43 | -1.50 | 3.1E-02 |
| C11orf54 | -1.49 | 1.6E-02 |
| SMARCA2 | -1.49 | 2.6E-02 |
| HPCAL1 | -1.49 | 2.2E-02 |
| HSPB11 | -1.49 | 3.3E-03 |
| UBE2T | -1.49 | 3.0E-03 |
| MTSS1L | -1.49 | 2.1E-02 |
| SAMD9 | -1.49 | 1.3E-02 |
| KHNYN | -1.49 | 3.6E-02 |
| AGMAT | -1.49 | 4.2E-02 |
| C19orf66 | -1.48 | 2.2E-02 |
| PDE4D | -1.48 | 1.1E-02 |
| CENPV | -1.48 | 4.2E-02 |
| CAD | -1.48 | 1.1E-03 |
| SPATS2 | -1.48 | 6.8E-03 |
| H6PD | -1.48 | 1.7E-07 |
| FBXO4 | -1.48 | 4.1E-02 |
| KLHL13 | -1.48 | 1.1E-02 |
| SNCA | -1.48 | 6.1E-03 |
| DNAJB4 | -1.48 | 7.0E-03 |
| GBP2 | -1.48 | 4.4E-03 |
| ACADM | -1.48 | 2.3E-02 |
| FKBP7 | -1.47 | 2.7E-02 |
| GINS4 | -1.47 | 1.5E-02 |
| UQCR10 | -1.47 | 3.1E-02 |
| ARHGEF16 | -1.47 | 3.4E-03 |
| TFPI | -1.47 | 2.4E-02 |
| SMPD1 | -1.46 | 2.1E-03 |
| AHRR | -1.46 | 2.0E-02 |
| BRD7 | -1.46 | 2.6E-02 |
| CDAN1 | -1.46 | 3.6E-02 |
| MIF4GD | -1.46 | 3.8E-03 |
| BIN3 | -1.46 | 1.5E-02 |
| APRT | -1.46 | 1.7E-02 |
| RCCD1 | -1.46 | 4.9E-02 |
| WDR11 | -1.45 | 1.6E-02 |
| IST1 | -1.45 | 8.7E-03 |

| | | |
|---|---|---|
| CRIP1 | -1.44 | 1.7E-03 |
| BID | -1.44 | 1.4E-02 |
| AMPD2 | -1.43 | 3.1E-02 |
| KLHL18 | -1.43 | 1.1E-02 |
| LEPREL4 | -1.43 | 4.0E-02 |
| CDKN2A | -1.43 | 9.7E-03 |
| C7orf50 | -1.43 | 4.7E-03 |
| GGCT | -1.43 | 1.2E-02 |
| SSBP2 | -1.43 | 4.8E-02 |
| ASL | -1.42 | 2.1E-03 |
| STARD3NL | -1.42 | 8.9E-03 |
| RMDN1 | -1.42 | 1.8E-02 |
| SNX7 | -1.42 | 2.9E-02 |
| AKR1B1 | -1.42 | 8.4E-04 |
| MYCBP | -1.42 | 3.3E-02 |
| HSPA12A | -1.42 | 6.1E-03 |
| EME1 | -1.42 | 2.7E-02 |
| TTF2 | -1.42 | 2.5E-02 |
| GPRIN1 | -1.42 | 1.9E-02 |
| CEP85 | -1.42 | 2.7E-02 |
| JAK1 | -1.41 | 3.0E-02 |
| AHCYL1 | -1.41 | 4.4E-03 |
| TRIT1 | -1.41 | 4.3E-03 |
| FAM188A | -1.41 | 1.4E-02 |
| BAIAP2L1 | -1.41 | 4.2E-02 |
| ANLN | -1.41 | 4.4E-02 |
| NLE1 | -1.40 | 1.2E-02 |
| NOTCH2 | -1.40 | 3.5E-02 |
| ACAA1 | -1.40 | 1.3E-02 |
| PACSIN2 | -1.40 | 3.5E-03 |
| RTCB | -1.40 | 1.1E-02 |
| ECE1 | -1.40 | 7.5E-03 |
| PDLIM3 | -1.40 | 4.8E-02 |
| ZNF644 | -1.40 | 7.3E-03 |
| WDR92 | -1.40 | 5.5E-03 |
| BAIAP2 | -1.40 | 6.7E-03 |
| UBE3C | -1.40 | 1.2E-02 |
| PITPNB | -1.40 | 5.0E-03 |
| MYO9B | -1.40 | 6.2E-03 |
| MYO1B | -1.40 | 1.1E-03 |
| EFHD2 | -1.39 | 1.3E-02 |
| SHMT1 | -1.39 | 1.4E-03 |
| FERMT1 | -1.39 | 9.8E-03 |
| RND3 | -1.39 | 3.4E-03 |
| MIEF1 | -1.39 | 1.9E-02 |
| CRLF3 | -1.39 | 1.7E-02 |
| C16orf87 | -1.39 | 7.6E-03 |
| WWC1 | -1.39 | 5.0E-02 |
| SNX8 | -1.39 | 1.7E-02 |
| EPHA5 | -1.39 | 3.1E-03 |
| MPST | -1.39 | 2.8E-02 |
| BZW2 | -1.39 | 2.8E-02 |
| MAP7D2 | -1.39 | 3.0E-02 |
| TADA2A | -1.39 | 4.3E-02 |
| MYH10 | -1.38 | 4.2E-02 |
| PRKAR1A | -1.38 | 2.8E-02 |
| DHFR | -1.38 | 2.9E-02 |
| CENPH | -1.38 | 3.5E-03 |
| STRIP1 | -1.38 | 5.7E-04 |
| BAP18 | -1.38 | 1.4E-03 |
| CMPK1 | -1.38 | 1.4E-02 |
| CBS | -1.38 | 1.7E-02 |
| TAX1BP1 | -1.38 | 3.8E-04 |
| BRIP1 | -1.37 | 2.3E-02 |
| CBLB | -1.37 | 3.2E-02 |
| EPB41 | -1.37 | 4.1E-02 |
| IFT122 | -1.37 | 2.4E-02 |
| CASP8 | -1.37 | 3.0E-03 |

| | | |
|---|---|---|
| COTL1 | -1.85 | 7.05E-03 |
| PDXK | -1.84 | 2.97E-02 |
| FN3KRP | -1.84 | 1.03E-02 |
| SPATA20 | -1.84 | 3.21E-02 |
| MCC | -1.84 | 1.64E-03 |
| CCDC85C | -1.84 | 1.62E-02 |
| CFL2 | -1.84 | 8.00E-04 |
| MET | -1.84 | 4.10E-02 |
| MVD | -1.83 | 3.03E-02 |
| NXN | -1.83 | 2.07E-02 |
| CDKN2C | -1.83 | 2.86E-02 |
| ODC1 | -1.83 | 4.34E-03 |
| LIMCH1 | -1.83 | 2.09E-02 |
| PLA2G4B | -1.82 | 1.88E-02 |
| BAD | -1.82 | 2.26E-02 |
| TOM1L2 | -1.82 | 2.26E-02 |
| NOTCH2 | -1.82 | 2.09E-02 |
| ITSN1 | -1.82 | 2.33E-03 |
| MCCC1 | -1.82 | 3.94E-02 |
| UACA | -1.81 | 9.18E-03 |
| FNBP1L | -1.81 | 6.45E-03 |
| TREX1 | -1.81 | 2.99E-02 |
| TPD52 | -1.81 | 3.09E-02 |
| DENND5B | -1.80 | 1.40E-02 |
| SAMHD1 | -1.80 | 3.32E-03 |
| CSRP2 | -1.80 | 1.80E-02 |
| NME2P1 | -1.80 | 4.71E-02 |
| ZBED1 | -1.80 | 4.10E-03 |
| HSPA2 | -1.80 | 3.81E-02 |
| CABLES1 | -1.80 | 1.87E-02 |
| AZI1 | -1.80 | 3.56E-02 |
| NEDD4 | -1.79 | 4.57E-03 |
| TLN1 | -1.79 | 2.89E-02 |
| BAIAP2L1 | -1.79 | 2.56E-02 |
| PIR | -1.78 | 1.21E-02 |
| LPP | -1.78 | 3.95E-04 |
| SYNM | -1.78 | 2.88E-02 |
| EPPK1 | -1.78 | 1.29E-02 |
| PTRF | -1.78 | 6.68E-03 |
| CALD1 | -1.77 | 2.46E-02 |
| TYMS | -1.77 | 7.47E-03 |
| SH3RF1 | -1.77 | 1.17E-02 |
| TCF19 | -1.77 | 8.55E-03 |
| ACLY | -1.76 | 6.08E-03 |
| PRKAR1B | -1.75 | 1.90E-02 |
| MAP1A | -1.75 | 3.21E-02 |
| PAFAH1B1 | -1.75 | 3.89E-02 |
| PLXND1 | -1.75 | 3.23E-02 |
| FN1 | -1.75 | 2.21E-02 |
| SMPDL3A | -1.75 | 1.94E-02 |
| AJUBA | -1.74 | 1.05E-02 |
| PSTPIP2 | -1.74 | 2.61E-02 |
| VPS13C | -1.74 | 1.81E-02 |
| DNAJA4 | -1.74 | 1.87E-02 |
| CCDC102A | -1.73 | 3.09E-02 |
| WBP2 | -1.73 | 1.55E-02 |
| ABLIM1 | -1.73 | 1.77E-02 |
| CAMKK1 | -1.73 | 1.29E-02 |
| DGKA | -1.72 | 8.95E-03 |
| LRRC42 | -1.71 | 3.46E-02 |
| SH3BP5 | -1.71 | 2.88E-03 |
| LRRC15 | -1.71 | 2.09E-02 |
| RPL36AL | -1.70 | 2.21E-02 |
| AKR1C2 | -1.70 | 1.86E-02 |
| CTNNB1 | -1.70 | 1.51E-02 |
| MGLL | -1.70 | 3.56E-02 |
| SSBP2 | -1.70 | 2.82E-02 |
| MAP7 | -1.70 | 2.21E-02 |

| | | |
|---|---|---|
| NEU1 | -1.43 | 2.27E-02 |
| SP100 | -1.43 | 1.44E-02 |
| TANK | -1.43 | 2.76E-02 |
| MEIS1 | -1.43 | 2.57E-02 |
| KBTBD7 | -1.43 | 6.23E-03 |
| MSI2 | -1.43 | 1.91E-02 |
| PINX1 | -1.42 | 5.09E-03 |
| ESD | -1.42 | 2.52E-02 |
| TUBAL3 | -1.42 | 1.79E-02 |
| CAB39 | -1.42 | 1.27E-02 |
| WIPF1 | -1.42 | 3.56E-02 |
| HSPB11 | -1.41 | 3.34E-03 |
| RABGAP1L | -1.40 | 3.26E-02 |
| MOCOS | -1.40 | 1.44E-02 |
| MUC16 | -1.40 | 3.09E-03 |
| FTL | -1.40 | 3.18E-02 |
| TCEAL1 | -1.40 | 4.63E-02 |
| SYTL4 | -1.40 | 8.84E-03 |
| KSR1 | -1.40 | 2.27E-02 |
| DIAPH3 | -1.39 | 1.99E-02 |
| GALK2 | -1.39 | 4.00E-02 |
| GBP2 | -1.39 | 1.29E-03 |
| GNS | -1.39 | 7.77E-03 |
| IFI30 | -1.39 | 8.69E-03 |
| MYO6 | -1.38 | 2.77E-02 |
| DPP7 | -1.38 | 2.98E-03 |
| NT5DC1 | -1.38 | 1.70E-02 |
| CTPS2 | -1.38 | 4.44E-02 |
| ADIRF | -1.38 | 3.04E-02 |
| DNAJC7 | -1.37 | 3.93E-02 |
| FAM120C | -1.37 | 1.49E-03 |
| FH | -1.37 | 8.30E-03 |
| PDE4D | -1.36 | 1.24E-02 |
| MVP | -1.36 | 3.61E-02 |
| STXBP2 | -1.36 | 2.84E-02 |
| PPM1A | -1.36 | 1.95E-02 |
| SH3GL2 | -1.36 | 2.65E-02 |
| ASCC3 | -1.36 | 7.00E-04 |
| SVIL | -1.36 | 2.06E-02 |
| ELP5 | -1.36 | 2.29E-02 |
| S100A13 | -1.36 | 4.08E-02 |
| PEPD | -1.35 | 1.00E-02 |
| COBL | -1.35 | 3.73E-02 |
| GLMN | -1.35 | 1.89E-02 |
| SAMD4B | -1.35 | 2.36E-03 |
| ANK3 | -1.35 | 2.79E-02 |
| SELENBP1 | -1.35 | 3.51E-02 |
| PC | -1.35 | 2.48E-02 |
| PDCD4 | -1.34 | 4.24E-02 |
| SMN1 | -1.34 | 2.54E-02 |
| LSS | -1.34 | 5.43E-03 |
| KDM3A | -1.34 | 3.70E-02 |
| DFNA5 | -1.34 | 6.77E-03 |
| IDI1 | -1.34 | 9.17E-03 |
| FAM172A | -1.34 | 3.20E-02 |
| CEP44 | -1.33 | 2.98E-03 |
| PSMD10 | -1.33 | 4.49E-02 |
| FTO | -1.33 | 1.97E-02 |
| MFF | -1.33 | 2.90E-02 |
| OTUD5 | -1.33 | 1.65E-02 |
| SPATS2 | -1.32 | 1.61E-02 |
| SH3PXD2B | -1.32 | 2.99E-02 |
| ZFYVE1 | -1.32 | 2.50E-02 |
| MTFR2 | -1.32 | 2.77E-02 |
| PHC1 | -1.32 | 2.63E-02 |
| PDK2 | -1.32 | 2.55E-02 |
| GCLM | -1.31 | 2.24E-02 |
| GNB1L | -1.31 | 4.83E-02 |

| | | |
|---|---|---|
| VASN | -1.45 | 4.9E-02 |
| KHDRBS3 | -1.45 | 3.3E-03 |
| TBPL2 | -1.45 | 3.4E-02 |
| LONP2 | -1.45 | 1.8E-02 |
| RMND5A | -1.45 | 1.7E-02 |
| HDHD2 | -1.44 | 4.7E-02 |
| PRKAR1B | -1.44 | 3.3E-02 |
| FAM172A | -1.44 | 5.4E-03 |
| KIAA1598 | -1.44 | 3.2E-02 |
| SDC1 | -1.44 | 4.3E-02 |
| PAFAH2 | -1.43 | 2.3E-02 |
| TCF25 | -1.43 | 1.3E-02 |
| MMACHC | -1.43 | 3.6E-02 |
| VAC14 | -1.43 | 7.6E-03 |
| N6AMT1 | -1.43 | 3.3E-02 |
| TRIM11 | -1.43 | 3.1E-02 |
| NDOR1 | -1.43 | 3.7E-02 |
| AP1S3 | -1.43 | 2.1E-02 |
| DNM1 | -1.43 | 4.4E-02 |
| MTHFSD | -1.43 | 2.2E-02 |
| FAM127C | -1.43 | 4.6E-02 |
| CSTB | -1.42 | 2.4E-02 |
| LCMT2 | -1.42 | 5.1E-03 |
| IPO9 | -1.42 | 1.2E-02 |
| FAM192A | -1.42 | 2.6E-02 |
| RAD23A | -1.42 | 1.4E-02 |
| VAV2 | -1.42 | 2.7E-02 |
| PDCD4 | -1.42 | 2.7E-02 |
| TBC1D2 | -1.42 | 1.6E-02 |
| RRAGB | -1.41 | 1.9E-02 |
| BROX | -1.41 | 8.1E-03 |
| JAK1 | -1.41 | 2.8E-02 |
| YRDC | -1.41 | 1.2E-02 |
| CTPS1 | -1.41 | 3.9E-02 |
| TTC28 | -1.41 | 4.4E-02 |
| BLVRA | -1.41 | 2.1E-02 |
| COQ5 | -1.41 | 4.4E-02 |
| CLIP2 | -1.41 | 4.8E-03 |
| FAM195B | -1.40 | 7.9E-03 |
| HEXB | -1.40 | 2.4E-03 |
| ATPAF1 | -1.40 | 3.3E-02 |
| GGT1 | -1.40 | 3.4E-02 |
| HAGH | -1.40 | 1.7E-02 |
| BCAR1 | -1.40 | 3.7E-02 |
| RPS6KA1 | -1.40 | 3.8E-02 |
| RXRA | -1.40 | 4.3E-02 |
| MEIS1 | -1.40 | 2.5E-02 |
| ALDH3B1 | -1.39 | 2.3E-02 |
| RABGAP1L | -1.39 | 8.0E-04 |
| AKR1A1 | -1.39 | 3.8E-02 |
| GOT1 | -1.39 | 4.7E-02 |
| EXOSC6 | -1.39 | 4.4E-02 |
| STX17 | -1.39 | 4.4E-02 |
| MTR | -1.39 | 1.3E-02 |
| PARS2 | -1.39 | 4.6E-02 |
| HTATIP2 | -1.39 | 2.4E-02 |
| RRM2B | -1.39 | 2.5E-03 |
| NPEPL1 | -1.39 | 2.6E-02 |
| MKNK1 | -1.39 | 5.1E-03 |
| PPCS | -1.39 | 2.6E-02 |
| PPOX | -1.38 | 2.0E-02 |
| DHFR | -1.38 | 4.9E-02 |
| CORO2A | -1.38 | 2.2E-02 |
| SETMAR | -1.38 | 1.5E-02 |
| CETN3 | -1.38 | 3.1E-02 |
| APEH | -1.38 | 2.2E-02 |
| DUS2 | -1.38 | 1.6E-03 |
| TXLNA | -1.38 | 4.9E-02 |

| | | |
|---|---|---|
| SPAG9 | -1.37 | 6.0E-03 |
| C12orf29 | -1.37 | 3.1E-02 |
| DESI1 | -1.37 | 1.5E-02 |
| GGA1 | -1.37 | 2.8E-02 |
| HECTD3 | -1.36 | 4.3E-02 |
| RANBP1 | -1.36 | 4.8E-02 |
| UBAP2 | -1.36 | 3.7E-02 |
| FARP1 | -1.36 | 6.5E-03 |
| CRKL | -1.36 | 5.3E-03 |
| KANK2 | -1.36 | 3.4E-02 |
| ACADM | -1.36 | 3.0E-02 |
| NUDC | -1.36 | 3.3E-02 |
| MICAL3 | -1.36 | 3.6E-02 |
| RSBN1L | -1.36 | 2.7E-02 |
| MED13 | -1.35 | 4.5E-03 |
| RSBN1 | -1.35 | 4.7E-02 |
| VASP | -1.35 | 5.0E-02 |
| TRIM47 | -1.35 | 3.8E-02 |
| MIER1 | -1.35 | 1.8E-03 |
| CDCA7L | -1.35 | 7.5E-03 |
| GNA12 | -1.35 | 3.8E-02 |
| RXRA | -1.35 | 1.4E-02 |
| RABEP1 | -1.35 | 2.4E-02 |
| MYBBP1A | -1.35 | 1.5E-02 |
| RNF14 | -1.35 | 2.7E-02 |
| ZCCHC7 | -1.35 | 2.3E-02 |
| TSHZ2 | -1.35 | 3.1E-03 |
| DIXDC1 | -1.34 | 4.6E-02 |
| PPIE | -1.34 | 5.4E-03 |
| DGCR14 | -1.34 | 1.1E-02 |
| TFPI | -1.34 | 3.3E-02 |
| PRKACB | -1.34 | 7.2E-03 |
| EZH2 | -1.34 | 1.9E-02 |
| TYRO3 | -1.34 | 3.9E-02 |
| ADSL | -1.34 | 1.8E-02 |
| ELAC2 | -1.34 | 1.3E-03 |
| CETN3 | -1.33 | 4.8E-02 |
| TNFAIP2 | -1.33 | 2.0E-02 |
| TYMS | -1.33 | 1.1E-02 |
| UBE2G2 | -1.33 | 3.2E-02 |
| LEPREL1 | -1.33 | 2.9E-02 |
| PRPSAP2 | -1.33 | 7.2E-03 |
| PBDC1 | -1.33 | 4.1E-02 |
| APTX | -1.33 | 7.7E-03 |
| YRDC | -1.33 | 1.2E-02 |
| DNAJC7 | -1.32 | 4.7E-02 |
| CDC42EP4 | -1.32 | 2.1E-02 |
| NCDN | -1.32 | 3.7E-03 |
| MCAT | -1.32 | 3.5E-02 |
| SNX4 | -1.32 | 6.9E-04 |
| TBC1D8B | -1.32 | 5.0E-02 |
| CDKN1A | -1.32 | 1.7E-02 |
| TSEN54 | -1.32 | 3.4E-02 |
| FAF1 | -1.32 | 2.7E-03 |
| IMPA1 | -1.32 | 5.6E-03 |
| GCFC2 | -1.32 | 1.5E-03 |
| TCEB3 | -1.31 | 3.8E-02 |
| JMJD6 | -1.31 | 4.2E-02 |
| ECD | -1.31 | 2.5E-02 |
| DCXR | -1.31 | 9.0E-03 |
| TRIM16 | -1.31 | 4.1E-03 |
| PZP | -1.31 | 1.5E-02 |
| CHORDC1 | -1.31 | 2.9E-02 |
| CYTH1 | -1.31 | 1.3E-02 |
| TLK2 | -1.31 | 1.7E-03 |
| DNTTIP2 | -1.30 | 6.7E-03 |
| TSTD2 | -1.30 | 3.5E-02 |
| SMG6 | -1.30 | 1.7E-02 |

| | | |
|---|---|---|
| ARHGAP23 | -1.69 | 1.07E-02 |
| SCRN2 | -1.69 | 5.88E-03 |
| TTC39C | -1.69 | 1.07E-02 |
| NFIB | -1.69 | 1.33E-02 |
| PEG10 | -1.68 | 2.21E-02 |
| TUBD1 | -1.68 | 8.79E-03 |
| COMMD9 | -1.68 | 8.39E-03 |
| HABP4 | -1.68 | 2.35E-02 |
| SAMD11 | -1.68 | 4.47E-02 |
| SERPINI1 | -1.67 | 1.72E-03 |
| ACAT2 | -1.67 | 2.79E-02 |
| PTK7 | -1.67 | 2.55E-02 |
| RDX | -1.67 | 4.03E-03 |
| PCYT1A | -1.67 | 2.56E-02 |
| ID3 | -1.67 | 2.78E-03 |
| FDXR | -1.66 | 1.11E-02 |
| CORO2A | -1.66 | 2.12E-03 |
| ARMC9 | -1.65 | 3.19E-03 |
| FNBP1 | -1.65 | 1.75E-02 |
| CTIF | -1.65 | 4.30E-02 |
| EPS8 | -1.65 | 2.12E-02 |
| WWC1 | -1.65 | 2.80E-02 |
| ZNF462 | -1.65 | 7.54E-03 |
| L1RE1 | -1.65 | 1.55E-02 |
| CDC42EP2 | -1.65 | 3.09E-02 |
| PTER | -1.65 | 1.35E-02 |
| TNK1 | -1.64 | 2.26E-03 |
| FKBP5 | -1.64 | 1.09E-02 |
| SORBS3 | -1.64 | 3.34E-02 |
| PPP1R12A | -1.64 | 2.77E-02 |
| NPEPPS | -1.64 | 3.42E-04 |
| CLN5 | -1.64 | 3.77E-02 |
| CHN1 | -1.63 | 1.13E-02 |
| NBR1 | -1.63 | 2.04E-02 |
| RNASEL | -1.63 | 4.05E-02 |
| NAPG | -1.63 | 3.06E-02 |
| CDAN1 | -1.63 | 2.41E-02 |
| PPP1R13L | -1.62 | 1.21E-02 |
| ANKRD13A | -1.62 | 3.56E-02 |
| MVK | -1.62 | 1.69E-02 |
| SVIL | -1.62 | 1.46E-02 |
| BAG3 | -1.61 | 1.90E-02 |
| HN1L | -1.61 | 3.84E-02 |
| GNS | -1.61 | 3.15E-03 |
| FDFT1 | -1.61 | 2.00E-02 |
| TRIM5 | -1.61 | 2.65E-02 |
| IDH2 | -1.61 | 1.00E-02 |
| GOLPH3L | -1.60 | 3.05E-02 |
| IGFBP7 | -1.60 | 3.56E-02 |
| KIAA1671 | -1.60 | 3.62E-02 |
| GIPC1 | -1.60 | 3.85E-02 |
| SPTBN4 | -1.60 | 1.13E-03 |
| HSDL2 | -1.60 | 2.37E-02 |
| EMC9 | -1.60 | 1.39E-03 |
| TRIM14 | -1.60 | 4.54E-02 |
| PRPSAP2 | -1.60 | 2.78E-03 |
| PCBD2 | -1.60 | 4.77E-02 |
| BAIAP2 | -1.59 | 2.01E-02 |
| DPYSL2 | -1.59 | 2.86E-02 |
| FAM177A1 | -1.58 | 2.71E-03 |
| SRXN1 | -1.58 | 4.50E-02 |
| KIAA1468 | -1.58 | 5.33E-03 |
| NUDT16L1 | -1.58 | 5.36E-04 |
| STK38 | -1.58 | 1.00E-02 |
| SPG20 | -1.58 | 2.20E-02 |
| ASL | -1.57 | 1.77E-02 |
| SPECC1 | -1.57 | 1.61E-02 |
| GSS | -1.57 | 4.38E-02 |

| | | |
|---|---|---|
| GM2A | -1.31 | 1.03E-02 |
| CHML | -1.31 | 1.55E-02 |
| RAD1 | -1.31 | 2.64E-02 |
| ZMYM3 | -1.31 | 4.74E-02 |
| WHSC1 | -1.31 | 2.75E-02 |
| FAM117B | -1.31 | 4.79E-02 |
| HBD | -1.31 | 7.23E-03 |
| BEGAIN | -1.31 | 3.03E-02 |
| COG6 | -1.31 | 1.59E-02 |
| DNMBP | -1.31 | 1.15E-02 |
| RMND5A | -1.30 | 1.56E-03 |
| ACAT2 | -1.30 | 4.44E-02 |
| PRCP | -1.30 | 6.85E-03 |
| VAC14 | -1.30 | 1.50E-02 |
| IGBP1 | -1.30 | 3.42E-02 |
| SNX29 | -1.30 | 4.13E-02 |
| STXBP1 | -1.30 | 3.82E-02 |
| GPSM1 | -1.30 | 4.26E-02 |
| EEF1E1 | -1.30 | 4.60E-02 |
| PUS7L | -1.30 | 4.88E-02 |
| ANXA6 | -1.30 | 1.36E-02 |
| GPX1 | -1.29 | 1.46E-02 |
| THOC3 | -1.29 | 4.44E-02 |
| CARKD | -1.29 | 2.16E-02 |
| GLRX | -1.29 | 3.28E-02 |
| RXRA | -1.29 | 1.85E-02 |
| FAM203A | -1.29 | 3.84E-02 |
| GPHN | -1.29 | 4.01E-02 |
| VPS13B | -1.29 | 2.27E-02 |
| PPME1 | -1.29 | 5.15E-03 |
| ACYP2 | -1.29 | 5.71E-04 |
| TTC3 | -1.29 | 3.00E-02 |
| CHMP4A | -1.29 | 2.68E-02 |
| ITSN1 | -1.28 | 3.14E-02 |
| IPO9 | -1.28 | 2.64E-02 |
| STAT5B | -1.28 | 5.55E-03 |
| PLCH1 | -1.28 | 1.62E-02 |
| PSAP | -1.28 | 1.40E-02 |
| CCM2 | -1.28 | 4.38E-02 |
| BPHL | -1.28 | 4.28E-02 |
| BIN3 | -1.28 | 2.48E-02 |
| PGM2L1 | -1.28 | 1.55E-02 |
| NUDT9 | -1.27 | 3.30E-02 |
| VPS25 | -1.27 | 4.55E-02 |
| HTATIP2 | -1.27 | 1.17E-02 |
| BRF1 | -1.27 | 1.11E-02 |
| NIT2 | -1.27 | 1.05E-02 |
| PEG10 | -1.27 | 3.90E-02 |
| CRIP1 | -1.27 | 1.37E-02 |
| PRKAB2 | -1.27 | 7.59E-03 |
| APPL2 | -1.27 | 2.55E-03 |
| TPMT | -1.27 | 3.00E-02 |
| SUFU | -1.26 | 3.47E-02 |
| SERPINB6 | -1.26 | 3.89E-02 |
| GBE1 | -1.26 | 2.52E-03 |
| MORF4L2 | -1.26 | 2.24E-02 |
| FBXO3 | -1.26 | 1.90E-02 |
| HUS1 | -1.26 | 2.21E-02 |
| NDC80 | -1.26 | 3.63E-02 |
| ZNHIT1 | -1.26 | 9.08E-03 |
| THOC5 | -1.25 | 1.93E-02 |
| ATXN10 | -1.25 | 4.83E-02 |
| ASCC2 | -1.25 | 4.71E-02 |
| GSTZ1 | -1.25 | 1.71E-02 |
| WDR11 | -1.25 | 4.76E-02 |
| DCK | -1.25 | 1.55E-02 |
| KIF3B | -1.25 | 4.77E-02 |
| ATPIF1 | -1.25 | 2.98E-02 |

| | | |
|---|---|---|
| ALDH7A1 | -1.38 | 2.5E-02 |
| APPL1 | -1.37 | 2.4E-02 |
| NENF | -1.37 | 1.2E-02 |
| CHMP4A | -1.37 | 8.1E-03 |
| PTGS1 | -1.37 | 4.5E-02 |
| SYNJ1 | -1.37 | 3.1E-03 |
| TBCE | -1.37 | 3.6E-02 |
| WRNIP1 | -1.37 | 4.5E-02 |
| EPB41 | -1.36 | 4.5E-02 |
| UBE2G2 | -1.36 | 1.7E-02 |
| EML2 | -1.36 | 2.8E-02 |
| RASSF1 | -1.36 | 4.6E-02 |
| WDR44 | -1.36 | 2.7E-02 |
| NAE1 | -1.36 | 3.3E-03 |
| TRIM16 | -1.36 | 1.2E-02 |
| CHMP1A | -1.36 | 7.2E-04 |
| CARKD | -1.36 | 5.8E-03 |
| TCEAL1 | -1.36 | 4.3E-02 |
| MAP2 | -1.35 | 3.8E-02 |
| CPEB4 | -1.35 | 3.2E-02 |
| BPNT1 | -1.35 | 3.2E-02 |
| EXOSC3 | -1.35 | 3.2E-02 |
| CTPS2 | -1.35 | 4.9E-02 |
| ERI3 | -1.35 | 4.6E-02 |
| AK4 | -1.35 | 2.8E-02 |
| FKBP5 | -1.35 | 3.2E-02 |
| FBXW11 | -1.35 | 5.8E-03 |
| MPHOSPH6 | -1.35 | 2.5E-02 |
| HLCS | -1.35 | 1.5E-02 |
| GABARAPL2 | -1.35 | 1.6E-02 |
| CPNE8 | -1.35 | 9.7E-03 |
| SDCBP | -1.34 | 1.2E-02 |
| JUND | -1.34 | 3.3E-02 |
| S100A13 | -1.34 | 4.5E-02 |
| AP1G1 | -1.34 | 9.2E-03 |
| PITPNM1 | -1.34 | 2.2E-02 |
| IMUP | -1.33 | 3.6E-02 |
| ZFHX3 | -1.33 | 1.8E-03 |
| ACYP1 | -1.33 | 1.1E-02 |
| PCNT | -1.33 | 5.4E-03 |
| STAMBPL1 | -1.33 | 9.0E-03 |
| RBBP4 | -1.33 | 1.2E-02 |
| GID8 | -1.33 | 2.0E-02 |
| SYNGR2 | -1.33 | 3.3E-02 |
| DENND4C | -1.33 | 3.2E-02 |
| GCLC | -1.33 | 2.9E-03 |
| DAK | -1.33 | 2.7E-02 |
| EXOSC10 | -1.33 | 2.6E-02 |
| TARBP1 | -1.33 | 3.7E-02 |
| GPSM1 | -1.32 | 4.4E-02 |
| STAT2 | -1.32 | 3.8E-04 |
| CAB39 | -1.32 | 2.4E-02 |
| POLA1 | -1.32 | 1.5E-02 |
| GCLM | -1.32 | 2.0E-02 |
| FAM65A | -1.32 | 2.4E-02 |
| KIAA0101 | -1.32 | 1.3E-03 |
| HDAC6 | -1.32 | 4.8E-02 |
| SPAG1 | -1.32 | 2.5E-02 |
| ARHGEF10L | -1.32 | 8.5E-03 |
| EZH2 | -1.32 | 2.7E-02 |
| ALDH16A1 | -1.31 | 2.8E-02 |
| DONSON | -1.31 | 4.9E-02 |
| FANCD2 | -1.31 | 3.2E-03 |
| ZC3H18 | -1.31 | 3.6E-02 |
| ADSS | -1.31 | 3.0E-02 |
| FAM114A1 | -1.31 | 2.2E-02 |
| CHML | -1.31 | 1.6E-02 |
| TYMS | -1.31 | 1.9E-02 |

| | | |
|---|---|---|
| PES1 | -1.30 | 2.4E-03 |
| ZCCHC11 | -1.30 | 1.1E-02 |
| DNAJB4 | -1.30 | 2.2E-02 |
| RAI1 | -1.30 | 2.1E-02 |
| PEO1 | -1.30 | 3.2E-02 |
| URGCP | -1.30 | 3.8E-02 |
| SNIP1 | -1.30 | 4.9E-02 |
| PUS7 | -1.30 | 2.6E-03 |
| FASN | -1.30 | 1.0E-02 |
| CHKA | -1.30 | 2.6E-02 |
| MALT1 | -1.30 | 2.1E-02 |
| CDC20 | -1.29 | 2.1E-02 |
| BCAT1 | -1.29 | 1.1E-02 |
| CAMKK2 | -1.29 | 3.7E-02 |
| ADPRHL2 | -1.29 | 4.2E-02 |
| CAMK1D | -1.29 | 2.0E-03 |
| KANK1 | -1.29 | 8.8E-03 |
| GET4 | -1.29 | 4.5E-02 |
| FHL1 | -1.29 | 1.1E-02 |
| PREP | -1.29 | 2.4E-03 |
| ALKBH4 | -1.29 | 3.8E-02 |
| PALLD | -1.29 | 3.9E-02 |
| POLDIP3 | -1.29 | 2.1E-02 |
| GSTT2 | -1.29 | 2.6E-02 |
| PRR11 | -1.28 | 6.0E-04 |
| GTSE1 | -1.28 | 1.3E-02 |
| ZZEF1 | -1.28 | 7.5E-04 |
| DVL2 | -1.28 | 2.5E-02 |
| BCKDHB | -1.28 | 1.3E-02 |
| ACTN1 | -1.28 | 1.4E-02 |
| DPH1 | -1.28 | 4.7E-03 |
| DUSP3 | -1.28 | 6.4E-03 |
| DDX1 | -1.28 | 3.3E-02 |
| SGSH | -1.28 | 2.9E-02 |
| PITPNA | -1.28 | 2.8E-02 |
| CUL1 | -1.28 | 1.4E-02 |
| UBE2L3 | -1.28 | 2.4E-02 |
| DAPK3 | -1.28 | 3.5E-02 |
| WDR46 | -1.28 | 4.7E-03 |
| ARMC6 | -1.28 | 3.2E-02 |
| UBE2H | -1.28 | 3.9E-02 |
| RECQL4 | -1.28 | 3.6E-02 |
| PRKCA | -1.28 | 3.9E-02 |
| BARD1 | -1.28 | 2.3E-02 |
| UBE2O | -1.28 | 1.6E-02 |
| ALYREF | -1.28 | 1.6E-02 |
| RBM19 | -1.27 | 1.1E-02 |
| KIF1B | -1.27 | 4.6E-02 |
| PARP12 | -1.27 | 2.0E-02 |
| TRA2A | -1.27 | 1.0E-02 |
| ZFYVE1 | -1.27 | 2.5E-02 |
| TBC1D24 | -1.27 | 2.0E-02 |
| NECAP2 | -1.27 | 1.9E-02 |
| PLD3 | -1.27 | 7.6E-04 |
| PGM1 | -1.27 | 1.6E-02 |
| PAPOLA | -1.27 | 1.5E-02 |
| TFCP2 | -1.27 | 1.8E-03 |
| WDR41 | -1.27 | 1.4E-02 |
| PHACTR4 | -1.27 | 3.6E-03 |
| ZNF444 | -1.26 | 5.4E-03 |
| GNB1 | -1.26 | 2.0E-02 |
| NIPSNAP1 | -1.26 | 3.5E-02 |
| TTC38 | -1.26 | 3.1E-02 |
| SPAG1 | -1.26 | 3.4E-02 |
| FOXK1 | -1.26 | 3.4E-02 |
| ATG7 | -1.26 | 1.6E-02 |
| CLUH | -1.26 | 2.2E-02 |
| KCTD15 | -1.26 | 1.9E-02 |

| | | |
|---|---|---|
| ARHGAP42 | -1.57 | 2.16E-02 |
| BRIP1 | -1.57 | 3.94E-02 |
| ATP6V0D1 | -1.57 | 1.12E-02 |
| SPTAN1 | -1.57 | 5.83E-03 |
| CAMK1D | -1.57 | 1.70E-03 |
| ACADVL | -1.57 | 2.74E-02 |
| TIMP2 | -1.57 | 3.32E-03 |
| CDC42EP1 | -1.57 | 2.70E-02 |
| MAPK7 | -1.56 | 8.69E-03 |
| CHML | -1.56 | 1.42E-02 |
| SPTBN2 | -1.56 | 9.05E-03 |
| SPHK1 | -1.56 | 1.83E-02 |
| PARVA | -1.56 | 3.34E-03 |
| STAT2 | -1.55 | 1.36E-02 |
| VPS25 | -1.55 | 1.19E-02 |
| TANC1 | -1.55 | 2.49E-02 |
| PRKACB | -1.55 | 1.63E-03 |
| ARHGEF40 | -1.55 | 1.99E-02 |
| DNAJB2 | -1.55 | 3.21E-02 |
| DGKH | -1.55 | 4.21E-02 |
| TSEN54 | -1.55 | 1.79E-02 |
| HEATR6 | -1.55 | 4.75E-02 |
| VCL | -1.55 | 4.12E-02 |
| G6PD | -1.55 | 4.90E-02 |
| SNX7 | -1.55 | 1.65E-02 |
| SEPT6 | -1.54 | 4.77E-02 |
| NSF | -1.54 | 2.82E-02 |
| HAGH | -1.54 | 2.70E-03 |
| PPP4R1 | -1.54 | 1.72E-03 |
| PPP1R9A | -1.54 | 4.62E-02 |
| SMARCA2 | -1.54 | 2.35E-02 |
| SYNJ1 | -1.54 | 1.10E-02 |
| SCRN3 | -1.53 | 1.69E-02 |
| COG4 | -1.53 | 1.99E-02 |
| UBE2H | -1.53 | 3.07E-02 |
| SHCBP1 | -1.53 | 3.13E-02 |
| KATNAL1 | -1.53 | 3.77E-02 |
| AARSD1 | -1.53 | 1.52E-02 |
| ASAH1 | -1.53 | 2.12E-03 |
| SPECC1L | -1.53 | 3.94E-02 |
| FAM115A | -1.52 | 1.49E-02 |
| DMD | -1.52 | 4.90E-02 |
| GMPPB | -1.52 | 6.85E-03 |
| NT5C2 | -1.52 | 8.29E-03 |
| NT5C | -1.52 | 9.08E-03 |
| NCKAP5L | -1.52 | 1.96E-02 |
| MAK | -1.51 | 2.66E-02 |
| CORO1B | -1.51 | 1.23E-02 |
| CLU | -1.51 | 2.39E-02 |
| STX17 | -1.51 | 2.12E-02 |
| DEPDC7 | -1.51 | 4.22E-02 |
| SHOC2 | -1.51 | 1.28E-02 |
| COASY | -1.51 | 2.06E-02 |
| CCDC43 | -1.50 | 4.24E-02 |
| DUSP3 | -1.50 | 6.99E-03 |
| SHMT1 | -1.50 | 6.85E-03 |
| NECAP2 | -1.50 | 1.12E-02 |
| SASH1 | -1.50 | 1.59E-03 |
| PDK2 | -1.50 | 5.88E-03 |
| NCOA7 | -1.49 | 2.16E-02 |
| SPTBN1 | -1.49 | 6.25E-03 |
| TRIM21 | -1.49 | 2.06E-02 |
| TMEM63A | -1.49 | 3.43E-02 |
| STAT5B | -1.49 | 1.42E-03 |
| ADAL | -1.49 | 2.46E-02 |
| CASP2 | -1.49 | 2.67E-02 |
| EPHA5 | -1.48 | 3.41E-02 |
| FLNB | -1.48 | 1.98E-02 |

| | | |
|---|---|---|
| PPM1D | -1.25 | 1.51E-02 |
| POLA1 | -1.25 | 1.20E-02 |
| SAMHD1 | -1.25 | 1.18E-02 |
| LANCL1 | -1.25 | 1.05E-03 |
| ZC3H15 | -1.25 | 5.64E-03 |
| BLVRA | -1.25 | 4.63E-02 |
| AMACR | -1.24 | 2.11E-02 |
| FUS | -1.24 | 4.11E-03 |
| FAM136A | -1.24 | 4.28E-02 |
| PRDX4 | -1.24 | 1.16E-02 |
| AKR1B1 | -1.24 | 1.59E-03 |
| CTSZ | -1.24 | 8.33E-03 |
| ALKBH4 | -1.24 | 4.66E-02 |
| NR3C1 | -1.24 | 2.49E-02 |
| GCLC | -1.24 | 1.64E-02 |
| FAM133A | -1.24 | 2.08E-03 |
| PPT1 | -1.24 | 5.08E-03 |
| FTH1 | -1.24 | 3.35E-02 |
| MAST2 | -1.23 | 4.31E-03 |
| SIRT1 | -1.23 | 1.23E-02 |
| HSPA12A | -1.23 | 4.53E-02 |
| NOL8 | -1.23 | 4.94E-02 |
| NDUFA7 | -1.23 | 1.76E-02 |
| KIF21A | -1.23 | 2.77E-02 |
| PFKFB3 | -1.23 | 2.33E-02 |
| ARL14EP | -1.23 | 2.03E-02 |
| SPAST | -1.23 | 2.06E-02 |
| HSPH1 | -1.23 | 3.35E-02 |
| USP24 | -1.23 | 6.95E-03 |
| ARHGEF7 | -1.23 | 3.51E-02 |
| STAU2 | -1.22 | 2.24E-02 |
| PTOV1 | -1.22 | 8.82E-03 |
| DAP | -1.22 | 3.89E-02 |
| RBM47 | -1.22 | 2.91E-02 |
| NMD3 | -1.22 | 2.97E-03 |
| LYPLA1 | -1.22 | 4.94E-02 |
| MMAB | -1.22 | 2.61E-03 |
| TRIM28 | -1.22 | 4.63E-02 |
| COG2 | -1.22 | 1.73E-02 |
| UBXN7 | -1.22 | 2.45E-02 |
| LPP | -1.22 | 4.04E-02 |
| GTPBP8 | -1.22 | 2.64E-02 |
| SERPINB1 | -1.21 | 1.29E-02 |
| ZNF644 | -1.21 | 4.26E-02 |
| VTA1 | -1.21 | 4.66E-02 |
| ZNF346 | -1.21 | 4.61E-02 |
| MRPL18 | -1.21 | 1.24E-02 |
| FBXW11 | -1.21 | 1.16E-02 |
| DIS3 | -1.21 | 1.13E-02 |
| KIF4A | -1.21 | 4.38E-02 |
| MAN2B1 | -1.21 | 1.74E-02 |
| C17orf75 | -1.21 | 3.54E-02 |
| MVK | -1.21 | 1.13E-02 |
| PRPS2 | -1.21 | 2.84E-02 |
| MIER1 | -1.21 | 6.03E-03 |
| HAGH | -1.21 | 1.21E-02 |
| ASAH1 | -1.21 | 3.42E-02 |
| KCTD5 | -1.21 | 4.86E-02 |
| CMAS | -1.21 | 4.92E-02 |
| MMACHC | -1.21 | 3.26E-02 |
| PZP | -1.21 | 2.80E-02 |
| KDM5C | -1.20 | 2.16E-02 |
| LENG1 | -1.20 | 2.83E-02 |
| UCHL3 | -1.20 | 2.44E-02 |
| CENPM | -1.20 | 3.39E-02 |
| ECI2 | -1.20 | 9.07E-03 |
| WDR7 | -1.20 | 1.42E-02 |
| NSD1 | -1.20 | 4.86E-04 |

| | | |
|---|---|---|
| PCK2 | -1.31 | 4.3E-02 |
| MYD88 | -1.31 | 3.0E-02 |
| RUNX1 | -1.31 | 4.5E-02 |
| KCTD3 | -1.31 | 1.9E-02 |
| HIP1R | -1.31 | 6.7E-03 |
| IPO13 | -1.30 | 4.1E-02 |
| PPP2R5A | -1.30 | 2.2E-02 |
| SDCCAG3 | -1.30 | 4.3E-02 |
| MMAB | -1.30 | 1.1E-02 |
| ZNF462 | -1.30 | 2.7E-02 |
| HTATSF1 | -1.30 | 2.5E-02 |
| FASN | -1.30 | 4.5E-02 |
| DST | -1.30 | 1.3E-02 |
| USP48 | -1.30 | 1.8E-02 |
| VPS36 | -1.30 | 1.7E-02 |
| SHOC2 | -1.30 | 1.2E-02 |
| SNTB2 | -1.30 | 6.5E-03 |
| EVPL | -1.29 | 4.8E-02 |
| CBS | -1.29 | 1.2E-02 |
| RANGRF | -1.29 | 1.7E-03 |
| PHACTR4 | -1.29 | 2.8E-02 |
| ANAPC4 | -1.29 | 8.7E-03 |
| ANKFY1 | -1.29 | 4.4E-02 |
| TDRD7 | -1.29 | 7.0E-03 |
| MSH3 | -1.29 | 7.7E-03 |
| TIMELESS | -1.29 | 3.4E-02 |
| PRKACB | -1.29 | 8.0E-03 |
| RBM34 | -1.29 | 3.1E-02 |
| COG4 | -1.29 | 4.8E-02 |
| PI4KA | -1.28 | 3.6E-02 |
| IKBKAP | -1.28 | 4.3E-02 |
| PARP10 | -1.28 | 1.3E-02 |
| GSTM1 | -1.28 | 2.3E-02 |
| MIER1 | -1.28 | 7.2E-03 |
| NRP1 | -1.28 | 4.1E-02 |
| RPA1 | -1.28 | 1.2E-02 |
| NFIB | -1.28 | 4.0E-02 |
| GDPGP1 | -1.28 | 1.0E-02 |
| GLRX | -1.28 | 4.4E-02 |
| PHIP | -1.28 | 5.3E-03 |
| NT5C3A | -1.28 | 5.3E-03 |
| TXNRD1 | -1.28 | 4.2E-02 |
| NPEPPS | -1.28 | 5.7E-03 |
| SPIN1 | -1.28 | 2.5E-02 |
| PCID2 | -1.28 | 8.1E-03 |
| ARAF | -1.28 | 8.4E-03 |
| KIF21A | -1.28 | 3.8E-02 |
| CC2D1B | -1.28 | 2.4E-02 |
| INF2 | -1.28 | 1.8E-02 |
| BLOC1S2 | -1.27 | 3.1E-02 |
| KIF13A | -1.27 | 2.6E-02 |
| ENDOG | -1.27 | 4.1E-02 |
| UBE2E2 | -1.27 | 4.9E-02 |
| LTN1 | -1.27 | 4.4E-02 |
| EDC4 | -1.27 | 4.1E-02 |
| FANCI | -1.27 | 5.5E-03 |
| RALGAPA1 | -1.27 | 3.2E-03 |
| NKIRAS1 | -1.27 | 4.8E-02 |
| MORF4L2 | -1.27 | 5.1E-03 |
| GOLPH3 | -1.27 | 2.9E-02 |
| AP2A1 | -1.27 | 2.3E-02 |
| URM1 | -1.27 | 3.8E-02 |
| MSH2 | -1.27 | 1.2E-02 |
| RRM1 | -1.27 | 2.5E-02 |
| C2orf76 | -1.27 | 3.0E-02 |
| KIAA1524 | -1.27 | 3.8E-02 |
| C5orf51 | -1.27 | 4.8E-03 |
| PSMG1 | -1.27 | 4.1E-02 |

| | | |
|---|---|---|
| ZWINT | -1.26 | 1.9E-02 |
| RABGAP1L | -1.25 | 1.3E-02 |
| GSTM1 | -1.25 | 9.4E-03 |
| USP48 | -1.25 | 2.0E-02 |
| CAP1 | -1.25 | 3.0E-02 |
| FTSJ3 | -1.25 | 2.4E-02 |
| GTF2H2 | -1.25 | 2.7E-02 |
| CCBL1 | -1.25 | 1.9E-02 |
| EIF2B4 | -1.25 | 2.6E-02 |
| MON2 | -1.25 | 1.9E-02 |
| BCR | -1.25 | 4.6E-02 |
| CAMK1 | -1.25 | 1.7E-02 |
| RIOK1 | -1.25 | 2.8E-02 |
| MPDZ | -1.24 | 4.9E-02 |
| FCF1 | -1.24 | 2.6E-02 |
| UBE2S | -1.24 | 3.2E-02 |
| KCTD7 | -1.24 | 2.2E-02 |
| GYG1 | -1.24 | 2.4E-02 |
| PEAK1 | -1.24 | 2.8E-02 |
| GMPPB | -1.24 | 2.1E-02 |
| CDK11B | -1.24 | 1.1E-02 |
| DST | -1.24 | 1.3E-02 |
| FLNC | -1.24 | 3.3E-02 |
| FAM115A | -1.24 | 4.2E-02 |
| TRIM56 | -1.24 | 4.7E-02 |
| CHD1 | -1.24 | 3.6E-02 |
| NCAPG2 | -1.24 | 2.6E-02 |
| L3MBTL3 | -1.23 | 1.0E-02 |
| DDAH1 | -1.23 | 6.8E-03 |
| PRKCG | -1.23 | 3.0E-02 |
| CC2D1B | -1.23 | 3.8E-02 |
| WDR62 | -1.23 | 2.4E-02 |
| NR3C1 | -1.23 | 2.8E-02 |
| FNBP1L | -1.23 | 6.1E-03 |
| MRM1 | -1.23 | 1.5E-02 |
| KIAA0020 | -1.23 | 1.7E-02 |
| NME6 | -1.23 | 2.2E-02 |
| UBR1 | -1.23 | 1.2E-02 |
| ACLY | -1.23 | 3.2E-02 |
| WRN | -1.23 | 5.4E-03 |
| SMU1 | -1.23 | 3.4E-02 |
| DHX33 | -1.23 | 2.3E-02 |
| THG1L | -1.23 | 2.9E-02 |
| WDR74 | -1.23 | 4.9E-02 |
| LZIC | -1.22 | 2.9E-02 |
| CCDC137 | -1.22 | 2.4E-02 |
| FAM98A | -1.22 | 2.8E-02 |
| SUPT6H | -1.22 | 3.9E-02 |
| TWISTNB | -1.22 | 4.9E-02 |
| EEF1D | -1.22 | 4.2E-03 |
| SERBP1 | -1.22 | 2.8E-02 |
| ANKMY2 | -1.22 | 4.8E-02 |
| MTRR | -1.22 | 7.9E-03 |
| DDX56 | -1.22 | 8.9E-03 |
| RFC1 | -1.22 | 4.4E-03 |
| CEP78 | -1.22 | 6.3E-03 |
| PPT1 | -1.22 | 8.5E-03 |
| LSS | -1.21 | 1.9E-02 |
| HMHA1 | -1.21 | 2.0E-02 |
| STAT5B | -1.21 | 6.1E-03 |
| SPATS2L | -1.21 | 1.1E-02 |
| SCRN2 | -1.21 | 1.2E-02 |
| WHSC1 | -1.21 | 1.7E-02 |
| GBP2 | -1.21 | 6.3E-03 |
| AHCYL2 | -1.21 | 1.2E-02 |
| CDK5 | -1.21 | 3.6E-03 |
| TRIP13 | -1.21 | 3.6E-02 |
| PGS1 | -1.21 | 3.0E-02 |

| | | |
|---|---|---|
| COPS7A | -1.48 | 2.04E-02 |
| HEATR5A | -1.48 | 9.32E-03 |
| JAK1 | -1.48 | 2.36E-02 |
| KIAA1211 | -1.48 | 2.80E-02 |
| BEGAIN | -1.48 | 1.83E-02 |
| NECAP1 | -1.48 | 3.72E-02 |
| BCKDHB | -1.48 | 5.43E-03 |
| SPAG5 | -1.47 | 4.14E-02 |
| DSTYK | -1.47 | 2.15E-02 |
| ZFHX3 | -1.47 | 4.98E-03 |
| IFT122 | -1.47 | 1.43E-02 |
| MPP5 | -1.47 | 4.59E-02 |
| OSTF1 | -1.47 | 2.26E-02 |
| CHMP1B | -1.47 | 2.94E-02 |
| RAI14 | -1.47 | 4.47E-02 |
| SNCA | -1.47 | 4.29E-02 |
| VPS8 | -1.47 | 7.40E-03 |
| MYO1E | -1.46 | 1.12E-02 |
| GRB2 | -1.46 | 3.97E-02 |
| RSU1 | -1.46 | 2.69E-02 |
| EPB41 | -1.46 | 4.15E-02 |
| GGA3 | -1.46 | 2.14E-02 |
| DVL2 | -1.45 | 2.44E-02 |
| SETD7 | -1.45 | 4.11E-02 |
| HNMT | -1.45 | 1.14E-02 |
| C18orf8 | -1.45 | 1.52E-02 |
| SPAG1 | -1.45 | 1.44E-02 |
| ZBTB3 | -1.45 | 1.11E-02 |
| HMCES | -1.45 | 2.41E-02 |
| NAPRT1 | -1.45 | 3.26E-02 |
| UBAP2 | -1.44 | 4.11E-02 |
| STK38L | -1.44 | 1.81E-02 |
| HDGFRP3 | -1.44 | 1.73E-02 |
| TRAF2 | -1.44 | 3.25E-02 |
| TRIOBP | -1.44 | 1.97E-02 |
| DOCK5 | -1.44 | 2.05E-02 |
| YEATS4 | -1.44 | 3.00E-02 |
| CDC6 | -1.44 | 2.56E-02 |
| PITPNA | -1.43 | 1.01E-02 |
| TNFAIP2 | -1.43 | 2.33E-02 |
| DOHH | -1.43 | 3.79E-02 |
| SMAP1 | -1.43 | 3.90E-02 |
| ATXN10 | -1.43 | 2.67E-02 |
| KIF1C | -1.43 | 3.66E-02 |
| MPDZ | -1.43 | 8.68E-03 |
| KLC1 | -1.42 | 4.14E-02 |
| PIP5K1C | -1.42 | 5.67E-03 |
| EHD2 | -1.42 | 3.75E-02 |
| MIF4GD | -1.42 | 1.26E-03 |
| BLVRA | -1.42 | 1.80E-02 |
| ZWILCH | -1.42 | 2.42E-02 |
| TMEM11 | -1.42 | 3.50E-02 |
| ABR | -1.42 | 4.71E-03 |
| SPEG | -1.42 | 1.11E-02 |
| STAMBPL1 | -1.42 | 3.37E-03 |
| IGF2BP3 | -1.42 | 4.00E-02 |
| PAFAH2 | -1.42 | 1.67E-02 |
| CTNNAL1 | -1.42 | 8.46E-03 |
| STXBP1 | -1.42 | 2.19E-02 |
| PPM1D | -1.42 | 8.36E-03 |
| GOT1 | -1.41 | 9.03E-03 |
| SPIN1 | -1.41 | 1.73E-02 |
| VPS36 | -1.41 | 9.10E-03 |
| PDLIM7 | -1.41 | 1.88E-02 |
| EHD1 | -1.41 | 4.19E-02 |
| APTX | -1.41 | 1.92E-02 |
| ACOX1 | -1.41 | 1.79E-03 |
| RAET1G | -1.41 | 4.24E-02 |

| | | |
|---|---|---|
| HINT1 | -1.20 | 4.73E-02 |
| TEAD1 | -1.20 | 3.03E-02 |
| GYG1 | -1.20 | 3.62E-02 |
| VPS11 | -1.20 | 3.81E-02 |
| GBA | -1.20 | 2.15E-03 |
| GGCT | -1.20 | 4.29E-02 |
| MKL1 | -1.20 | 2.88E-02 |
| KIF13A | -1.19 | 2.85E-02 |
| SEC31A | -1.19 | 5.65E-03 |
| N6AMT1 | -1.19 | 3.38E-02 |
| CASK | -1.19 | 1.77E-02 |
| CPSF2 | -1.19 | 4.04E-02 |
| OAS3 | -1.19 | 7.89E-03 |
| ZFP91 | -1.19 | 3.67E-02 |
| SH3BP4 | -1.19 | 2.21E-02 |
| RABGAP1 | -1.19 | 3.13E-02 |
| ETFB | -1.19 | 3.85E-02 |
| MYO1B | -1.19 | 4.03E-03 |
| PRKAG1 | -1.19 | 1.57E-02 |
| PPP6R3 | -1.19 | 8.08E-03 |
| IDH1 | -1.19 | 4.27E-02 |
| VPS52 | -1.19 | 8.89E-03 |
| NDUFS5 | -1.19 | 3.69E-02 |
| TCF25 | -1.19 | 4.56E-02 |
| PHIP | -1.19 | 4.47E-03 |
| KCTD9 | -1.19 | 1.13E-02 |
| RRM2B | -1.18 | 9.58E-03 |
| AGL | -1.18 | 2.08E-02 |
| MRPS31 | -1.18 | 3.54E-02 |
| EXOC5 | -1.18 | 1.25E-02 |
| MLLT1 | -1.18 | 3.50E-02 |
| DDHD2 | -1.18 | 4.87E-02 |
| HNRNPA3 | -1.18 | 1.91E-02 |
| TNK1 | -1.18 | 1.68E-02 |
| RBBP7 | -1.18 | 2.41E-02 |
| KLHL13 | -1.18 | 3.28E-02 |
| TDP2 | -1.18 | 2.49E-02 |
| GTSE1 | -1.18 | 3.79E-02 |
| PSMA3 | -1.18 | 1.50E-02 |
| CDK11B | -1.18 | 3.96E-02 |
| PSMA6 | -1.18 | 4.74E-02 |
| TFCP2 | -1.18 | 3.83E-02 |
| RCN1 | -1.17 | 2.65E-02 |
| INTS9 | -1.17 | 1.69E-02 |
| KCTD12 | -1.17 | 4.80E-03 |
| PPT2 | -1.17 | 4.06E-02 |
| EPDR1 | -1.17 | 4.77E-02 |
| DPP3 | -1.17 | 4.07E-02 |
| FXR2 | -1.16 | 4.95E-02 |
| PHKB | -1.16 | 2.21E-02 |
| VTN | -1.16 | 3.81E-02 |
| SDF2L1 | -1.16 | 3.93E-02 |
| FKBP5 | -1.16 | 4.02E-02 |
| TRAF6 | -1.16 | 4.69E-02 |
| TOLLIP | -1.16 | 3.52E-02 |
| HEXA | -1.16 | 1.74E-02 |
| TUBGCP3 | -1.16 | 4.83E-03 |
| PWWP2A | -1.16 | 8.70E-03 |
| RRM1 | -1.16 | 2.94E-02 |
| EXOC6B | -1.16 | 2.20E-02 |
| SMARCC2 | -1.16 | 3.91E-02 |
| STAG2 | -1.16 | 2.12E-02 |
| PSMA7 | -1.16 | 3.63E-02 |
| PSMA4 | -1.16 | 3.44E-02 |
| PCNT | -1.16 | 2.32E-02 |
| PGM1 | -1.16 | 4.82E-02 |
| PSMB2 | -1.15 | 3.63E-02 |
| MFN2 | -1.15 | 2.44E-03 |

| | | | | | | | | | | | |
|---|---|---|---|---|---|---|---|---|---|---|---|
| PPME1 | -1.27 | 6.3E-03 | LPP | -1.21 | 1.2E-02 | APEH | -1.40 | 1.55E-02 | HEBP1 | -1.15 | 1.21E-02 |
| HMCES | -1.26 | 3.7E-02 | RRP36 | -1.21 | 4.4E-02 | SIRPA | -1.40 | 3.73E-02 | DNAJC21 | -1.15 | 4.47E-02 |
| GMPPB | -1.26 | 1.7E-02 | ST13 | -1.21 | 4.4E-02 | SCPEP1 | -1.40 | 2.98E-02 | TYW3 | -1.15 | 1.72E-02 |
| SNX12 | -1.26 | 2.8E-02 | APPL2 | -1.21 | 2.9E-03 | GMPPA | -1.40 | 4.42E-02 | PDCD6 | -1.15 | 2.70E-02 |
| MANF | -1.26 | 3.4E-02 | GTF2E2 | -1.21 | 1.9E-02 | KCTD5 | -1.40 | 8.56E-03 | FBXO7 | -1.15 | 4.15E-02 |
| C12orf57 | -1.26 | 3.8E-02 | PRMT10 | -1.21 | 3.1E-02 | LLGL1 | -1.40 | 4.72E-02 | COMMD9 | -1.15 | 1.64E-02 |
| RAD17 | -1.26 | 3.9E-02 | IK | -1.21 | 2.2E-02 | LOX | -1.40 | 1.21E-02 | FIGNL1 | -1.15 | 2.15E-02 |
| TTC3 | -1.26 | 3.5E-02 | DOCK7 | -1.21 | 3.8E-03 | PARP10 | -1.40 | 7.43E-03 | MYCBP2 | -1.15 | 3.94E-02 |
| SAMD4B | -1.26 | 4.5E-02 | NUP35 | -1.20 | 2.6E-02 | ARPC5 | -1.40 | 1.43E-02 | CRELD2 | -1.15 | 6.26E-04 |
| MSH6 | -1.26 | 4.8E-02 | MICALL1 | -1.20 | 1.2E-02 | PPME1 | -1.40 | 3.36E-02 | BCL7A | -1.15 | 4.15E-02 |
| MAP1LC3B2 | -1.26 | 1.2E-02 | NOC3L | -1.20 | 3.9E-02 | CRYZL1 | -1.40 | 3.24E-02 | MARS | -1.15 | 3.88E-02 |
| EXOSC5 | -1.25 | 5.2E-03 | ASCC3 | -1.20 | 7.8E-03 | GPN2 | -1.40 | 3.51E-02 | PARP12 | -1.15 | 3.02E-02 |
| GALE | -1.25 | 4.4E-02 | RRS1 | -1.20 | 3.6E-03 | FDPS | -1.39 | 4.26E-02 | PSMB5 | -1.14 | 4.21E-02 |
| MAP4K5 | -1.25 | 3.4E-02 | BMS1 | -1.20 | 2.1E-03 | CHMP1A | -1.39 | 1.40E-02 | C19orf10 | -1.14 | 3.32E-02 |
| LSS | -1.25 | 8.9E-03 | NOL7 | -1.20 | 3.8E-03 | UBE2R2 | -1.39 | 3.38E-02 | TBL1X | -1.14 | 5.40E-03 |
| HSDL2 | -1.25 | 4.1E-02 | LYAR | -1.20 | 9.6E-03 | HPCAL1 | -1.39 | 2.84E-02 | GLE1 | -1.14 | 3.20E-02 |
| POMP | -1.25 | 4.3E-02 | USP36 | -1.20 | 3.9E-03 | NFX1 | -1.39 | 2.00E-02 | LAP3 | -1.14 | 1.69E-02 |
| ARHGEF40 | -1.25 | 4.0E-02 | TTLL12 | -1.20 | 4.2E-02 | ME1 | -1.39 | 2.92E-02 | PEO1 | -1.14 | 3.12E-02 |
| ELP5 | -1.25 | 9.9E-03 | C11orf68 | -1.20 | 5.0E-02 | PHKG2 | -1.39 | 4.33E-02 | COG1 | -1.14 | 8.38E-03 |
| GOT2 | -1.25 | 3.6E-02 | PPP1R9B | -1.20 | 1.1E-02 | FAM129A | -1.39 | 4.71E-02 | ANAPC4 | -1.14 | 3.22E-02 |
| GCFC2 | -1.25 | 2.3E-02 | AFTPH | -1.20 | 4.5E-02 | DENND4C | -1.39 | 1.53E-03 | BLOC1S1 | -1.14 | 3.49E-02 |
| UBAC1 | -1.25 | 4.7E-02 | MTPN | -1.20 | 1.6E-02 | MPRIP | -1.39 | 3.72E-02 | FOSL2 | -1.14 | 4.63E-02 |
| WBP2 | -1.25 | 5.0E-02 | FBXO7 | -1.20 | 1.6E-02 | MAP4K5 | -1.39 | 3.12E-02 | VPS18 | -1.14 | 2.73E-02 |
| FAM83H | -1.25 | 4.8E-02 | NEK6 | -1.20 | 3.2E-02 | KCTD10 | -1.39 | 1.36E-02 | BMS1 | -1.14 | 3.75E-02 |
| RBM8A | -1.24 | 4.2E-02 | EXOC6B | -1.20 | 3.5E-02 | VPS4B | -1.39 | 2.94E-02 | POLR2D | -1.13 | 4.43E-03 |
| ZNHIT6 | -1.24 | 4.0E-02 | HEATR5A | -1.20 | 3.3E-02 | LRSAM1 | -1.39 | 4.33E-02 | FBXO28 | -1.13 | 1.16E-02 |
| FHOD1 | -1.24 | 4.2E-02 | OARD1 | -1.20 | 2.9E-03 | APRT | -1.38 | 4.30E-02 | GAK | -1.13 | 4.78E-03 |
| PLD3 | -1.24 | 3.1E-03 | REXO4 | -1.19 | 2.1E-02 | EPDR1 | -1.38 | 2.84E-02 | PIAS1 | -1.13 | 4.58E-02 |
| YOD1 | -1.24 | 6.0E-03 | DHX57 | -1.19 | 4.4E-02 | GBP2 | -1.38 | 1.33E-02 | SEH1L | -1.13 | 2.01E-02 |
| DNAJA2 | -1.24 | 2.6E-02 | TBCD | -1.19 | 3.2E-02 | ABHD14B | -1.38 | 4.38E-02 | RPS6KA6 | -1.13 | 3.64E-02 |
| KDM3A | -1.24 | 4.3E-02 | HELLS | -1.19 | 4.6E-02 | TBCE | -1.38 | 3.53E-02 | SNTB2 | -1.13 | 8.24E-03 |
| AP2M1 | -1.23 | 4.3E-02 | PDCL3 | -1.19 | 1.7E-02 | NPLOC4 | -1.38 | 4.69E-03 | ME1 | -1.13 | 1.87E-02 |
| MLH1 | -1.23 | 4.3E-02 | KIF2C | -1.19 | 1.1E-02 | LONP2 | -1.38 | 3.04E-02 | PRPS1 | -1.13 | 4.37E-02 |
| CMPK1 | -1.23 | 3.0E-02 | APBB1 | -1.19 | 2.9E-02 | USP32 | -1.38 | 1.83E-02 | RNF213 | -1.13 | 3.76E-02 |
| GABPB1 | -1.23 | 3.4E-02 | AAMP | -1.19 | 4.3E-03 | TBCD | -1.38 | 1.15E-02 | MTMR1 | -1.13 | 2.27E-02 |
| HEATR5A | -1.23 | 2.4E-02 | ITIH2 | -1.19 | 3.5E-02 | CRELD2 | -1.37 | 1.25E-02 | PQBP1 | -1.12 | 4.03E-02 |
| TNK1 | -1.23 | 1.9E-02 | PURB | -1.19 | 4.6E-02 | GNE | -1.37 | 2.90E-02 | EXOC3 | -1.12 | 2.01E-02 |
| WHSC1 | -1.23 | 1.6E-02 | PPAN | -1.19 | 2.4E-02 | RRAGB | -1.37 | 6.84E-03 | ARAF | -1.12 | 2.10E-02 |
| RABGGTA | -1.23 | 2.3E-02 | DENND4C | -1.19 | 2.7E-02 | MLF2 | -1.37 | 3.43E-02 | ARID4B | -1.12 | 4.26E-02 |
| GART | -1.23 | 4.9E-02 | OPLAH | -1.19 | 2.8E-02 | JUP | -1.37 | 1.21E-02 | GABPA | -1.12 | 4.17E-02 |
| ATPIF1 | -1.22 | 6.3E-03 | NPLOC4 | -1.19 | 2.6E-03 | NAGLU | -1.37 | 9.39E-04 | ASCC1 | -1.12 | 3.31E-02 |
| XRN1 | -1.22 | 3.5E-02 | DDX47 | -1.19 | 1.5E-02 | DONSON | -1.37 | 3.17E-02 | DYNLL1 | -1.12 | 4.50E-02 |
| HSCB | -1.22 | 3.5E-02 | SAP30BP | -1.19 | 4.1E-02 | UFD1L | -1.37 | 4.75E-02 | SREK1IP1 | -1.12 | 3.44E-02 |
| PHF8 | -1.22 | 1.5E-03 | ABT1 | -1.19 | 1.5E-02 | JMJD6 | -1.37 | 4.30E-03 | WDR92 | -1.12 | 3.67E-02 |
| CAPN7 | -1.22 | 1.2E-02 | LIMS1 | -1.19 | 2.8E-02 | SEC14L2 | -1.36 | 3.30E-02 | RPA1 | -1.12 | 2.87E-02 |
| IFT122 | -1.22 | 4.3E-02 | NSA2 | -1.19 | 2.0E-02 | DDB2 | -1.36 | 2.96E-02 | MANF | -1.12 | 1.85E-02 |
| ATP1B2 | -1.22 | 5.2E-03 | GNL2 | -1.19 | 1.1E-02 | MFN1 | -1.36 | 1.20E-02 | YWHAB | -1.12 | 1.35E-02 |
| SH3PXD2B | -1.22 | 2.7E-02 | ME3 | -1.19 | 5.0E-02 | ECE1 | -1.36 | 8.23E-03 | NOL7 | -1.12 | 1.18E-02 |
| RRAGC | -1.22 | 4.6E-02 | KCTD10 | -1.18 | 2.1E-02 | TRAPPC8 | -1.36 | 3.34E-02 | RPS14 | -1.12 | 5.62E-03 |
| HEATR3 | -1.22 | 2.0E-02 | SASS6 | -1.18 | 4.7E-02 | UAP1L1 | -1.36 | 4.58E-02 | GMEB2 | -1.12 | 4.81E-02 |
| RABGGTB | -1.22 | 1.6E-02 | CDCA8 | -1.18 | 5.6E-04 | TSC2 | -1.36 | 5.79E-03 | TOPBP1 | -1.11 | 2.30E-02 |
| ECI2 | -1.22 | 8.9E-03 | BRIX1 | -1.18 | 2.4E-02 | ITGB4 | -1.36 | 4.61E-02 | PRPSAP2 | -1.11 | 3.52E-02 |
| BRCC3 | -1.22 | 7.6E-03 | MAP1LC3B2 | -1.18 | 1.9E-04 | DYNC1I2 | -1.36 | 3.58E-02 | TANGO6 | -1.11 | 4.22E-02 |
| DPH5 | -1.21 | 1.1E-03 | NOP16 | -1.18 | 1.4E-04 | AP1S1 | -1.35 | 2.14E-02 | NUDT16L1 | -1.11 | 6.28E-03 |
| BRD3 | -1.21 | 2.7E-02 | AATF | -1.18 | 2.4E-02 | OGFOD2 | -1.35 | 3.61E-03 | DDX39B | -1.11 | 2.85E-02 |
| ASF1A | -1.21 | 4.4E-02 | TNPO3 | -1.18 | 8.0E-03 | HELZ | -1.35 | 2.45E-02 | VPS51 | -1.11 | 4.06E-02 |
| LRBA | -1.21 | 2.5E-02 | BYSL | -1.18 | 2.8E-02 | CDCA7L | -1.35 | 4.91E-02 | LRBA | -1.11 | 2.97E-03 |
| EXOSC9 | -1.21 | 4.8E-02 | UBA2 | -1.18 | 2.9E-02 | IFT81 | -1.35 | 3.56E-02 | MIF4GD | -1.11 | 1.80E-02 |
| PACSIN3 | -1.21 | 1.5E-02 | RRP1B | -1.18 | 2.6E-02 | FXR2 | -1.35 | 2.85E-02 | STAT1 | -1.10 | 2.01E-02 |
| SH3RF1 | -1.21 | 1.7E-02 | NIFK | -1.18 | 3.4E-02 | PTPRF | -1.35 | 4.40E-02 | DPF2 | -1.10 | 4.16E-02 |
| STARD7 | -1.21 | 3.1E-02 | FTO | -1.18 | 3.9E-02 | RABGGTA | -1.35 | 2.71E-02 | USP48 | -1.10 | 2.76E-02 |
| IKBKG | -1.20 | 1.4E-02 | PFAS | -1.18 | 4.3E-02 | PARP12 | -1.34 | 7.26E-03 | CENPH | -1.10 | 3.52E-02 |
| PGM2L1 | -1.20 | 3.2E-02 | TDP2 | -1.18 | 2.1E-02 | KNTC1 | -1.34 | 1.19E-02 | NAA20 | -1.09 | 2.24E-02 |
| PITPNA | -1.20 | 4.8E-02 | CAPZA1 | -1.18 | 2.2E-02 | FAAP100 | -1.34 | 4.11E-02 | CCDC12 | -1.09 | 3.79E-02 |
| FAM45A | -1.20 | 2.1E-02 | UBE2G1 | -1.18 | 5.0E-02 | HSPA1L | -1.34 | 2.24E-02 | MLKL | -1.09 | 1.29E-02 |
| POLA2 | -1.20 | 4.5E-02 | RSL1D1 | -1.18 | 1.7E-02 | PDPK1 | -1.34 | 1.37E-02 | CTSA | -1.09 | 1.08E-02 |
| DENND6A | -1.20 | 4.6E-02 | WDR3 | -1.17 | 1.5E-02 | MICALL1 | -1.34 | 3.61E-02 | CAD | -1.09 | 4.90E-03 |
| RRP1B | -1.20 | 9.9E-03 | SEH1L | -1.17 | 4.3E-02 | IQGAP3 | -1.34 | 7.98E-03 | WRN | -1.09 | 3.05E-02 |

| | | |
|---|---|---|
| FBXO28 | -1.20 | 7.0E-03 |
| TFDP1 | -1.20 | 2.0E-02 |
| TELO2 | -1.20 | 2.0E-02 |
| C4orf27 | -1.20 | 4.6E-02 |
| USP1 | -1.20 | 1.1E-02 |
| GLMN | -1.19 | 2.8E-02 |
| SERBP1 | -1.19 | 4.9E-02 |
| IFT81 | -1.19 | 4.2E-02 |
| DDX41 | -1.19 | 2.1E-02 |
| TRADD | -1.19 | 1.1E-02 |
| BRAF | -1.19 | 3.2E-02 |
| HAUS5 | -1.19 | 1.6E-02 |
| DCXR | -1.19 | 2.5E-02 |
| ATP6V1D | -1.18 | 4.3E-02 |
| ACOT8 | -1.18 | 3.0E-04 |
| LAP3 | -1.18 | 1.1E-02 |
| NSD1 | -1.18 | 1.4E-02 |
| ZCCHC11 | -1.18 | 8.1E-03 |
| SPTBN2 | -1.18 | 2.8E-03 |
| DGKA | -1.17 | 4.5E-02 |
| ORC6 | -1.17 | 1.5E-02 |
| PIBF1 | -1.17 | 2.2E-02 |
| TSC1 | -1.17 | 4.4E-02 |
| DDHD2 | -1.17 | 4.3E-02 |
| OTUD5 | -1.17 | 1.2E-02 |
| FAM177A1 | -1.17 | 2.8E-02 |
| STAG2 | -1.16 | 1.4E-02 |
| TANGO6 | -1.16 | 6.1E-03 |
| TBC1D9B | -1.16 | 3.4E-02 |
| SRSF5 | -1.16 | 2.0E-03 |
| NUP93 | -1.16 | 4.7E-02 |
| APEX2 | -1.16 | 3.6E-02 |
| F8A1 | -1.16 | 1.7E-02 |
| NUP188 | -1.16 | 1.5E-02 |
| NEK9 | -1.16 | 7.1E-03 |
| UBR1 | -1.15 | 1.9E-02 |
| TAX1BP1 | -1.15 | 2.1E-02 |
| HNMT | -1.15 | 3.1E-02 |
| APAF1 | -1.15 | 1.2E-02 |
| MTRR | -1.15 | 7.3E-04 |
| ADRM1 | -1.15 | 4.0E-02 |
| KIF2C | -1.15 | 2.6E-02 |
| SAMHD1 | -1.15 | 2.7E-02 |
| MICALL2 | -1.15 | 3.8E-02 |
| IKBKB | -1.15 | 3.4E-02 |
| C6orf106 | -1.15 | 4.1E-02 |
| NIT1 | -1.15 | 4.2E-02 |
| SOS1 | -1.14 | 8.7E-03 |
| OSTM1 | -1.14 | 3.8E-02 |
| H1FX | -1.14 | 4.2E-02 |
| TAF11 | -1.14 | 2.4E-02 |
| PKN2 | -1.14 | 4.8E-02 |
| EXOSC8 | -1.14 | 2.5E-04 |
| CTCF | -1.14 | 3.5E-02 |
| SETD8 | -1.14 | 6.4E-03 |
| PRPF39 | -1.14 | 3.8E-02 |
| AP1B1 | -1.14 | 4.3E-02 |
| SHMT1 | -1.14 | 2.9E-02 |
| CASP6 | -1.14 | 3.9E-02 |
| MED23 | -1.14 | 4.5E-02 |
| TSR2 | -1.14 | 2.7E-02 |
| DROSHA | -1.14 | 3.5E-02 |
| PHF2 | -1.14 | 3.3E-02 |
| LYPLA1 | -1.14 | 1.1E-02 |
| CNPY2 | -1.14 | 2.2E-02 |
| SKIV2L2 | -1.13 | 4.7E-02 |
| SEH1L | -1.13 | 1.1E-02 |
| KIAA1279 | -1.13 | 2.9E-02 |

| | | |
|---|---|---|
| HAUS4 | -1.17 | 4.2E-02 |
| USP1 | -1.17 | 1.3E-02 |
| SUMO2 | -1.17 | 4.8E-02 |
| SAAL1 | -1.17 | 1.4E-02 |
| SPTBN2 | -1.17 | 3.1E-02 |
| OSBPL3 | -1.17 | 4.1E-02 |
| LIG3 | -1.17 | 3.8E-02 |
| DDX31 | -1.17 | 1.1E-02 |
| DSTN | -1.17 | 2.9E-02 |
| EHD4 | -1.17 | 4.1E-02 |
| RPL36AL | -1.17 | 6.4E-03 |
| NUDCD1 | -1.17 | 4.1E-04 |
| FOCAD | -1.16 | 2.4E-02 |
| BLMH | -1.16 | 4.2E-02 |
| PKN2 | -1.16 | 6.9E-03 |
| HACL1 | -1.16 | 2.2E-02 |
| TAF6 | -1.16 | 2.9E-03 |
| PSME3 | -1.16 | 4.5E-02 |
| NMD3 | -1.16 | 4.5E-02 |
| EPN2 | -1.16 | 7.3E-03 |
| ATF3 | -1.16 | 2.2E-02 |
| CDC27 | -1.16 | 4.8E-02 |
| RPL34 | -1.16 | 9.9E-03 |
| NF1 | -1.16 | 9.5E-03 |
| PARVA | -1.16 | 2.3E-02 |
| KCTD9 | -1.15 | 2.2E-02 |
| NOL11 | -1.15 | 3.6E-02 |
| GEMIN4 | -1.15 | 4.1E-02 |
| RPL22 | -1.15 | 1.6E-02 |
| SDAD1 | -1.15 | 4.5E-03 |
| MINA | -1.15 | 8.9E-03 |
| KTI12 | -1.15 | 4.5E-02 |
| OGFR | -1.15 | 3.4E-02 |
| CCDC12 | -1.15 | 4.8E-03 |
| C7orf25 | -1.15 | 1.5E-02 |
| RNF114 | -1.15 | 2.9E-02 |
| MBLAC2 | -1.15 | 1.9E-02 |
| SAE1 | -1.15 | 4.0E-02 |
| TCF19 | -1.15 | 4.9E-02 |
| TACC1 | -1.14 | 2.4E-02 |
| TTC27 | -1.14 | 1.9E-02 |
| KIF2A | -1.14 | 1.0E-02 |
| NOSIP | -1.14 | 3.3E-03 |
| RABGEF1 | -1.14 | 1.2E-02 |
| POLR2D | -1.14 | 2.8E-03 |
| TADA3 | -1.14 | 1.3E-02 |
| WIPI2 | -1.14 | 2.8E-02 |
| CAB39 | -1.14 | 4.2E-02 |
| HDGFRP3 | -1.14 | 1.4E-02 |
| RSL24D1 | -1.14 | 3.9E-02 |
| AP1B1 | -1.14 | 2.5E-02 |
| ASAH1 | -1.13 | 3.8E-02 |
| RBM5 | -1.13 | 2.6E-02 |
| DDX21 | -1.13 | 8.6E-03 |
| CNNM3 | -1.13 | 7.4E-03 |
| PRKRIR | -1.13 | 3.6E-02 |
| SPTBN4 | -1.13 | 1.0E-03 |
| UBE4B | -1.13 | 7.0E-04 |
| STK38 | -1.13 | 2.9E-02 |
| CLTA | -1.13 | 4.2E-02 |
| PWP2 | -1.13 | 1.9E-02 |
| TIMP2 | -1.13 | 3.8E-02 |
| RPL27 | -1.13 | 2.3E-02 |
| LENG1 | -1.13 | 4.5E-03 |
| IMP4 | -1.12 | 3.8E-02 |
| STAT3 | -1.12 | 4.9E-02 |
| RPL23A | -1.12 | 4.7E-02 |
| SIRT5 | -1.12 | 3.3E-02 |

| | | |
|---|---|---|
| UTRN | -1.34 | 1.95E-02 |
| GCLC | -1.34 | 4.95E-03 |
| TRIM47 | -1.34 | 2.99E-02 |
| VPS18 | -1.34 | 2.30E-02 |
| FYCO1 | -1.34 | 3.46E-02 |
| KLHL18 | -1.34 | 3.48E-02 |
| NFKB2 | -1.34 | 2.11E-02 |
| STAT3 | -1.34 | 1.18E-02 |
| ARRDC1 | -1.33 | 2.94E-02 |
| KBTBD7 | -1.33 | 8.48E-03 |
| CAPN1 | -1.33 | 2.31E-02 |
| TADA3 | -1.33 | 2.63E-02 |
| UNC13D | -1.33 | 1.26E-02 |
| SMEK3P | -1.33 | 7.97E-03 |
| AAMDC | -1.33 | 9.07E-03 |
| KCTD9 | -1.33 | 1.96E-02 |
| ATP6V1A | -1.33 | 9.76E-03 |
| DYNC1LI2 | -1.33 | 2.78E-02 |
| RABGEF1 | -1.33 | 7.23E-03 |
| TCF25 | -1.32 | 3.11E-02 |
| MMAB | -1.32 | 2.22E-02 |
| GSE1 | -1.32 | 2.15E-02 |
| HINT3 | -1.32 | 3.18E-03 |
| COG2 | -1.32 | 2.29E-02 |
| TRAPPC6B | -1.32 | 4.15E-02 |
| MTHFS | -1.32 | 1.93E-02 |
| PXN | -1.32 | 4.68E-02 |
| NDC80 | -1.32 | 3.61E-02 |
| GDPGP1 | -1.32 | 7.54E-05 |
| H2AFY2 | -1.32 | 2.91E-03 |
| HIF1AN | -1.32 | 3.78E-02 |
| PREPL | -1.32 | 3.59E-02 |
| SPAG9 | -1.31 | 1.09E-02 |
| EHHADH | -1.31 | 3.38E-02 |
| ECD | -1.31 | 2.33E-02 |
| SESTD1 | -1.31 | 2.14E-02 |
| CENPM | -1.31 | 1.64E-02 |
| OVCA2 | -1.31 | 3.39E-02 |
| EXOC6 | -1.31 | 4.79E-02 |
| KIF21A | -1.31 | 3.40E-02 |
| | -1.31 | 4.39E-02 |
| FAIM | -1.31 | 3.99E-02 |
| NUDT16 | -1.31 | 7.03E-03 |
| SNX4 | -1.31 | 2.98E-04 |
| RPL31 | -1.31 | 2.94E-02 |
| CUX1 | -1.31 | 4.93E-02 |
| TPMT | -1.30 | 2.91E-02 |
| STXBP2 | -1.30 | 4.96E-02 |
| TXNL1 | -1.30 | 4.40E-02 |
| DCTN6 | -1.30 | 4.72E-02 |
| SOS1 | -1.30 | 4.05E-02 |
| CROCC | -1.30 | 3.90E-02 |
| NIT1 | -1.30 | 2.70E-02 |
| ASF1B | -1.30 | 1.80E-02 |
| MAN2B1 | -1.30 | 2.61E-02 |
| NCKIPSD | -1.30 | 4.40E-02 |
| FAM114A1 | -1.30 | 6.00E-03 |
| WASH2P | -1.30 | 3.01E-02 |
| SMU1 | -1.30 | 2.95E-02 |
| MRM1 | -1.30 | 1.09E-02 |
| ANXA11 | -1.30 | 3.88E-02 |
| SNX12 | -1.29 | 3.66E-02 |
| PPOX | -1.29 | 1.43E-02 |
| AGL | -1.29 | 4.71E-03 |
| VPS11 | -1.29 | 2.15E-02 |
| MIB1 | -1.29 | 4.90E-02 |
| GUK1 | -1.29 | 2.93E-02 |
| ISOC1 | -1.29 | 4.58E-02 |

| | | |
|---|---|---|
| ATXN7L3B | -1.09 | 6.15E-03 |
| COPE | -1.09 | 3.76E-02 |
| ILF2 | -1.09 | 1.79E-02 |
| VPS45 | -1.09 | 2.84E-02 |
| SRSF5 | -1.09 | 3.10E-02 |
| TRAPPC8 | -1.09 | 8.39E-03 |
| BIRC6 | -1.08 | 2.45E-02 |
| CACYBP | -1.08 | 3.24E-02 |
| SKIV2L2 | -1.08 | 2.67E-02 |
| DTYMK | -1.07 | 4.72E-02 |
| LARP7 | -1.07 | 2.53E-02 |
| VPS33A | -1.06 | 1.03E-03 |
| JMJD6 | -1.06 | 4.61E-02 |
| SMARCD2 | -1.05 | 2.94E-02 |
| POLR1B | -1.05 | 3.08E-02 |
| METTL15 | -1.05 | 3.42E-02 |
| IPO5 | -1.04 | 2.40E-02 |
| DDX39A | -1.04 | 3.08E-02 |
| SNRPD2 | -1.04 | 4.06E-02 |
| PREP | -1.04 | 1.90E-02 |
| DYNLRB1 | -1.03 | 2.78E-02 |
| ASPM | -1.01 | 4.42E-02 |
| UBE4B | 1.01 | 4.22E-02 |
| SEC24C | 1.03 | 4.75E-02 |
| NUP155 | 1.04 | 2.81E-02 |
| STRIP1 | 1.04 | 1.75E-02 |
| BAZ2A | 1.04 | 1.79E-02 |
| RPL27A | 1.04 | 3.19E-02 |
| COPB1 | 1.04 | 3.52E-02 |
| ELAC2 | 1.04 | 2.41E-02 |
| STAT2 | 1.04 | 9.29E-03 |
| TRRAP | 1.05 | 3.61E-02 |
| TFIP11 | 1.05 | 4.53E-02 |
| GDPGP1 | 1.05 | 1.05E-03 |
| PMS1 | 1.05 | 4.46E-02 |
| CIC | 1.06 | 1.89E-02 |
| HTT | 1.06 | 2.36E-02 |
| INTS3 | 1.06 | 4.55E-03 |
| PDS5A | 1.06 | 3.09E-02 |
| HSPA1L | 1.06 | 2.77E-02 |
| PRPF8 | 1.06 | 2.45E-02 |
| WDR82 | 1.07 | 3.69E-02 |
| ERH | 1.07 | 3.86E-02 |
| WDR5 | 1.08 | 3.39E-02 |
| API5 | 1.08 | 3.62E-05 |
| PSMD3 | 1.08 | 2.89E-02 |
| CHP1 | 1.08 | 1.34E-02 |
| GNB2 | 1.08 | 4.22E-02 |
| NRDE2 | 1.08 | 2.83E-02 |
| XPO7 | 1.09 | 3.17E-02 |
| HEATR2 | 1.09 | 1.47E-02 |
| OARD1 | 1.09 | 6.82E-03 |
| DPH1 | 1.09 | 3.91E-02 |
| PHC2 | 1.09 | 1.67E-02 |
| TRIT1 | 1.09 | 3.62E-02 |
| PACSIN2 | 1.09 | 4.37E-02 |
| AP3D1 | 1.09 | 2.24E-02 |
| BUB1 | 1.09 | 4.83E-02 |
| PTPRE | 1.10 | 1.19E-02 |
| PRPF38B | 1.10 | 1.62E-02 |
| NEK9 | 1.10 | 1.27E-02 |
| GCN1L1 | 1.10 | 6.70E-03 |
| EIF4EBP1 | 1.10 | 1.48E-02 |
| CARM1 | 1.10 | 3.60E-02 |
| AGPS | 1.10 | 4.37E-03 |
| BLM | 1.10 | 2.84E-02 |
| MAP1LC3B2 | 1.10 | 1.74E-03 |
| GBF1 | 1.10 | 4.04E-02 |

| | | |
|---|---|---|
| ATG7 | -1.13 | 1.8E-02 |
| GDI1 | -1.13 | 2.4E-02 |
| FAM21C | -1.13 | 3.8E-02 |
| ALDH9A1 | -1.13 | 3.5E-02 |
| KDM1A | -1.13 | 3.8E-02 |
| RAPGEF6 | -1.13 | 2.6E-03 |
| STRIP1 | -1.13 | 4.0E-02 |
| SEC31A | -1.12 | 2.7E-02 |
| KTI12 | -1.12 | 2.1E-02 |
| CHTOP | -1.12 | 1.8E-02 |
| PAPD5 | -1.12 | 7.4E-03 |
| MFN1 | -1.12 | 4.6E-02 |
| CDCA8 | -1.12 | 1.1E-02 |
| PFDN6 | -1.12 | 3.9E-02 |
| SREK1IP1 | -1.12 | 2.9E-02 |
| TSN | -1.11 | 4.5E-02 |
| RPS6 | -1.11 | 3.6E-03 |
| WDR77 | -1.11 | 2.3E-02 |
| NUP107 | -1.10 | 2.1E-02 |
| SUMF2 | -1.10 | 7.8E-05 |
| ISG15 | -1.10 | 1.5E-02 |
| CEP78 | -1.10 | 4.8E-02 |
| ZBTB21 | -1.10 | 2.9E-02 |
| CRELD2 | -1.10 | 2.1E-02 |
| DNAL1 | -1.09 | 9.9E-03 |
| SYMPK | -1.09 | 3.1E-03 |
| RAB3GAP2 | -1.08 | 3.0E-02 |
| SECISBP2L | -1.08 | 4.2E-02 |
| AIDA | -1.08 | 4.0E-02 |
| DICER1 | -1.06 | 5.0E-02 |
| MED22 | -1.06 | 4.6E-03 |
| CFL2 | -1.06 | 2.1E-02 |
| NTHL1 | -1.05 | 4.4E-03 |
| TOPBP1 | -1.05 | 3.7E-02 |
| BOLA1 | -1.04 | 3.8E-02 |
| RPS14 | -1.04 | 5.7E-03 |
| SFMBT1 | -1.04 | 2.3E-02 |
| USP28 | -1.04 | 5.4E-03 |
| UBE4B | -1.03 | 5.5E-03 |
| SMARCC2 | -1.03 | 1.4E-02 |
| DYNC1H1 | 1.04 | 3.7E-02 |
| ZZEF1 | 1.04 | 5.0E-03 |
| CTSL | 1.05 | 2.0E-02 |
| TIFA | 1.05 | 1.4E-02 |
| GAK | 1.06 | 6.9E-03 |
| COPB1 | 1.06 | 1.7E-02 |
| CSE1L | 1.07 | 4.4E-02 |
| RRS1 | 1.07 | 2.7E-03 |
| GTF3C3 | 1.08 | 3.3E-02 |
| GBA | 1.08 | 1.8E-02 |
| NAA30 | 1.08 | 3.0E-02 |
| DPH1 | 1.09 | 4.2E-02 |
| SNX4 | 1.09 | 1.4E-02 |
| RINT1 | 1.09 | 4.8E-02 |
| PPAN | 1.09 | 2.8E-02 |
| PES1 | 1.09 | 1.5E-02 |
| VPS16 | 1.09 | 2.1E-02 |
| PWP2 | 1.09 | 3.0E-02 |
| NOL6 | 1.10 | 8.1E-03 |
| GTF3C1 | 1.10 | 9.5E-03 |
| MYC | 1.10 | 1.1E-02 |
| NIPSNAP1 | 1.10 | 4.4E-02 |
| WDR46 | 1.10 | 1.1E-02 |
| DCUN1D5 | 1.10 | 1.1E-02 |
| PEX5 | 1.10 | 2.3E-02 |
| LYAR | 1.10 | 2.5E-02 |
| LARP7 | 1.11 | 3.7E-02 |
| EEF1D | 1.11 | 3.1E-02 |

| | | |
|---|---|---|
| TBL3 | -1.12 | 1.2E-02 |
| IPO7 | -1.12 | 4.6E-02 |
| SNTB2 | -1.12 | 1.2E-02 |
| MFN1 | -1.12 | 4.5E-02 |
| RPS14 | -1.12 | 5.1E-04 |
| AGL | -1.12 | 1.7E-03 |
| NRD1 | -1.12 | 3.4E-02 |
| HEATR2 | -1.12 | 2.3E-02 |
| OSBPL11 | -1.12 | 1.8E-02 |
| STAG1 | -1.12 | 2.7E-02 |
| KIAA1429 | -1.12 | 3.2E-02 |
| SIKE1 | -1.12 | 1.2E-02 |
| PRPF8 | -1.12 | 8.8E-03 |
| ARSA | -1.12 | 6.6E-03 |
| MRPL18 | -1.12 | 2.7E-02 |
| PGM3 | -1.12 | 1.3E-02 |
| NT5C3A | -1.12 | 3.0E-02 |
| GEMIN5 | -1.12 | 1.9E-02 |
| RPS11 | -1.11 | 1.7E-02 |
| RBM22 | -1.11 | 1.7E-02 |
| GARS | -1.11 | 4.4E-02 |
| MFN2 | -1.11 | 2.0E-02 |
| EEF1B2 | -1.11 | 1.3E-02 |
| RBBP4 | -1.11 | 3.3E-02 |
| MRPS18B | -1.11 | 4.8E-02 |
| RPS26P11 | -1.11 | 4.5E-02 |
| POLG | -1.11 | 1.7E-02 |
| IPO11 | -1.11 | 1.9E-02 |
| TERF2IP | -1.11 | 3.4E-02 |
| WDR12 | -1.11 | 4.4E-03 |
| SCLY | -1.10 | 4.5E-02 |
| ASCC1 | -1.10 | 4.3E-03 |
| MAP2K3 | -1.10 | 3.2E-02 |
| GMEB2 | -1.09 | 3.1E-02 |
| LSG1 | -1.09 | 5.0E-02 |
| WDR77 | -1.09 | 2.7E-02 |
| CDC6 | -1.09 | 1.3E-03 |
| NUP160 | -1.09 | 4.3E-02 |
| PRKCD | -1.09 | 7.3E-03 |
| PCNT | -1.09 | 3.7E-02 |
| CHMP1A | -1.08 | 1.3E-02 |
| NOL6 | -1.08 | 3.7E-02 |
| NUP93 | -1.08 | 4.6E-02 |
| RPS6 | -1.08 | 4.9E-02 |
| SNRPD3 | -1.08 | 2.7E-02 |
| USP15 | -1.08 | 4.3E-02 |
| HAT1 | -1.08 | 3.6E-02 |
| NF2 | -1.08 | 1.4E-02 |
| RPS13 | -1.08 | 2.9E-02 |
| EIF3L | -1.07 | 4.8E-02 |
| KDM4A | -1.07 | 2.3E-02 |
| RPL24 | -1.07 | 4.8E-02 |
| RPL7A | -1.07 | 4.8E-02 |
| STAT1 | -1.07 | 2.5E-02 |
| ARIH1 | -1.07 | 3.4E-03 |
| SPEG | -1.06 | 1.4E-02 |
| SKIV2L2 | -1.06 | 1.5E-02 |
| NPEPPS | -1.06 | 3.0E-03 |
| GPR56 | -1.06 | 3.4E-02 |
| TTK | -1.06 | 1.6E-02 |
| RPL18A | -1.06 | 3.6E-02 |
| POLD2 | -1.06 | 1.3E-02 |
| DVL1 | -1.06 | 3.9E-02 |
| HDAC3 | -1.05 | 4.9E-02 |
| PUS7L | -1.05 | 3.1E-03 |
| REXO2 | -1.05 | 1.4E-02 |
| IPO5 | -1.05 | 2.8E-02 |
| SRSF5 | -1.03 | 4.2E-02 |

| | | |
|---|---|---|
| SRA1 | -1.29 | 2.41E-02 |
| METTL2B | -1.29 | 4.13E-02 |
| DLG5 | -1.28 | 4.00E-02 |
| CEP44 | -1.28 | 6.37E-03 |
| ACACA | -1.28 | 1.18E-02 |
| HSDL1 | -1.28 | 4.97E-02 |
| C16orf87 | -1.28 | 3.98E-02 |
| UBE2E2 | -1.28 | 4.00E-02 |
| ZZEF1 | -1.28 | 1.00E-02 |
| GSTZ1 | -1.28 | 2.89E-02 |
| DIS3L2 | -1.28 | 2.99E-02 |
| ALDH9A1 | -1.28 | 2.19E-02 |
| DIAPH3 | -1.28 | 3.80E-02 |
| C6orf106 | -1.28 | 4.15E-02 |
| SMCHD1 | -1.28 | 4.46E-02 |
| DCTN3 | -1.28 | 3.03E-02 |
| TXNL4A | -1.28 | 2.15E-02 |
| ACSS2 | -1.28 | 4.96E-02 |
| DIP2B | -1.28 | 1.26E-02 |
| TNKS1BP1 | -1.27 | 4.50E-02 |
| ANKFY1 | -1.27 | 4.01E-02 |
| ARHGAP1 | -1.27 | 3.38E-02 |
| GNB5 | -1.27 | 4.07E-02 |
| AFTPH | -1.27 | 1.70E-02 |
| SH3GLB2 | -1.27 | 4.18E-02 |
| GOLIM4 | -1.27 | 4.10E-02 |
| MED15 | -1.26 | 3.28E-02 |
| CRLF3 | -1.26 | 3.26E-02 |
| CAPZA1 | -1.26 | 4.78E-02 |
| PRR5L | -1.26 | 1.36E-02 |
| SPPL2A | -1.26 | 3.10E-02 |
| FAM21C | -1.26 | 8.46E-04 |
| UEVLD | -1.26 | 8.62E-03 |
| SDCBP | -1.26 | 3.03E-02 |
| ELP5 | -1.26 | 9.30E-03 |
| PTRH1 | -1.26 | 2.71E-02 |
| EDC4 | -1.25 | 4.56E-02 |
| SH3PXD2B | -1.25 | 3.47E-02 |
| DCTN5 | -1.25 | 9.17E-03 |
| COG6 | -1.25 | 2.17E-02 |
| SEC14L1 | -1.25 | 2.43E-02 |
| TBC1D17 | -1.25 | 3.30E-02 |
| PGP | -1.25 | 3.15E-02 |
| TAX1BP1 | -1.25 | 3.23E-03 |
| RPL22L1 | -1.25 | 1.04E-03 |
| ACAP2 | -1.25 | 3.99E-02 |
| SNX17 | -1.25 | 2.87E-02 |
| PLCD3 | -1.24 | 3.91E-02 |
| VPS39 | -1.24 | 3.91E-02 |
| HEBP1 | -1.24 | 1.81E-02 |
| MKNK1 | -1.24 | 3.54E-02 |
| TSEN15 | -1.24 | 2.68E-02 |
| UBE2O | -1.24 | 3.24E-02 |
| FIGNL1 | -1.24 | 1.51E-02 |
| MON2 | -1.24 | 3.02E-02 |
| HERC4 | -1.24 | 3.86E-03 |
| WDR6 | -1.23 | 1.08E-02 |
| DYNLRB1 | -1.23 | 1.26E-02 |
| FOCAD | -1.23 | 1.70E-02 |
| STAT6 | -1.23 | 3.72E-02 |
| STK32C | -1.23 | 3.85E-02 |
| HERC1 | -1.23 | 4.11E-03 |
| PRR11 | -1.23 | 1.74E-02 |
| FAM188A | -1.23 | 3.89E-02 |
| SEPT9 | -1.23 | 3.31E-02 |
| SMTN | -1.23 | 3.98E-02 |
| EXOSC3 | -1.23 | 3.76E-02 |
| TOP3A | -1.22 | 5.61E-03 |

| | | |
|---|---|---|
| RRP1B | 1.11 | 4.80E-02 |
| RRN3 | 1.11 | 6.86E-04 |
| KIF11 | 1.11 | 4.61E-02 |
| ECM29 | 1.11 | 1.70E-03 |
| DST | 1.11 | 3.95E-02 |
| RAB3GAP2 | 1.11 | 2.00E-02 |
| PROSC | 1.12 | 4.29E-02 |
| ABHD5 | 1.12 | 7.07E-03 |
| RNMT | 1.12 | 4.93E-02 |
| QARS | 1.12 | 4.25E-02 |
| SPATS2L | 1.12 | 4.76E-02 |
| PEAK1 | 1.13 | 4.24E-02 |
| DIP2B | 1.13 | 4.17E-02 |
| RMDN1 | 1.13 | 2.96E-02 |
| TIMP2 | 1.13 | 3.08E-02 |
| ARHGAP5 | 1.13 | 1.92E-02 |
| CDC6 | 1.13 | 4.33E-02 |
| DCXR | 1.13 | 3.27E-02 |
| APBB1 | 1.13 | 2.34E-02 |
| IQGAP3 | 1.13 | 4.71E-02 |
| SMG1 | 1.13 | 2.10E-02 |
| CCDC115 | 1.14 | 2.48E-02 |
| XPO5 | 1.14 | 6.55E-03 |
| SMC5 | 1.14 | 4.04E-02 |
| HAUS6 | 1.14 | 1.24E-02 |
| MPDZ | 1.14 | 1.60E-03 |
| WDR6 | 1.14 | 1.95E-02 |
| NOL11 | 1.14 | 3.15E-02 |
| PLA2G4B | 1.14 | 1.59E-02 |
| DYNC1H1 | 1.14 | 4.92E-03 |
| CHCHD2 | 1.14 | 5.09E-03 |
| ZW10 | 1.14 | 3.81E-02 |
| PTK2 | 1.15 | 2.03E-02 |
| HSCB | 1.15 | 3.94E-02 |
| HK2 | 1.15 | 2.14E-02 |
| PPIB | 1.15 | 2.78E-02 |
| FLII | 1.15 | 3.63E-02 |
| EPN2 | 1.15 | 4.55E-03 |
| ARF5 | 1.16 | 4.98E-02 |
| WDR3 | 1.16 | 2.12E-02 |
| STAG1 | 1.16 | 1.04E-02 |
| PLAA | 1.16 | 9.65E-03 |
| CLIP2 | 1.16 | 4.76E-02 |
| AQR | 1.16 | 1.52E-02 |
| SOGA1 | 1.16 | 3.82E-02 |
| DLGAP5 | 1.16 | 1.39E-03 |
| DDX47 | 1.16 | 1.80E-02 |
| DHX34 | 1.16 | 2.54E-04 |
| CENPL | 1.16 | 3.96E-02 |
| RBM15B | 1.16 | 4.11E-02 |
| PDIA3 | 1.16 | 9.61E-03 |
| SNX1 | 1.16 | 4.24E-02 |
| ARHGEF18 | 1.16 | 3.10E-02 |
| EFHD2 | 1.16 | 3.98E-02 |
| HMGA1 | 1.16 | 3.36E-02 |
| EHD1 | 1.16 | 4.16E-02 |
| METTL1 | 1.16 | 4.25E-02 |
| PDCD11 | 1.16 | 5.15E-03 |
| SAR1A | 1.17 | 2.39E-02 |
| CDCA8 | 1.17 | 1.57E-02 |
| RPP25 | 1.17 | 6.45E-03 |
| GUK1 | 1.17 | 1.09E-02 |
| PARG | 1.17 | 8.85E-03 |
| ERCC2 | 1.17 | 1.20E-02 |
| RCN2 | 1.17 | 2.96E-02 |
| RIF1 | 1.18 | 1.49E-02 |
| UNC45A | 1.18 | 3.31E-02 |
| RRAGB | 1.18 | 2.46E-02 |

| | | |
|---|---|---|
| DDX56 | 1.11 | 2.3E-02 |
| PRPF8 | 1.11 | 2.0E-02 |
| ABT1 | 1.11 | 1.7E-02 |
| NELFB | 1.11 | 2.7E-02 |
| STK25 | 1.11 | 3.7E-02 |
| DHX37 | 1.11 | 8.5E-03 |
| EXOC2 | 1.11 | 1.5E-02 |
| XRN2 | 1.11 | 4.9E-02 |
| LRWD1 | 1.12 | 4.6E-02 |
| AP3D1 | 1.12 | 4.9E-02 |
| CHCHD5 | 1.12 | 3.8E-02 |
| TBL1X | 1.12 | 2.9E-02 |
| DPH2 | 1.12 | 2.6E-03 |
| TTC27 | 1.12 | 1.8E-02 |
| USP39 | 1.12 | 4.4E-02 |
| POP7 | 1.13 | 1.5E-02 |
| FLAD1 | 1.13 | 4.8E-02 |
| NUDT4 | 1.13 | 1.8E-02 |
| TACC1 | 1.13 | 3.0E-02 |
| REXO4 | 1.13 | 3.1E-02 |
| MCRS1 | 1.13 | 3.9E-02 |
| LENG1 | 1.13 | 2.8E-03 |
| DXO | 1.13 | 4.4E-02 |
| FOSL2 | 1.14 | 4.2E-02 |
| SNX27 | 1.14 | 2.8E-02 |
| PYGB | 1.14 | 3.5E-02 |
| FSTL1 | 1.14 | 4.6E-02 |
| AGO2 | 1.14 | 1.2E-02 |
| PDZD8 | 1.14 | 6.6E-03 |
| TOP1 | 1.14 | 4.3E-02 |
| CMTR1 | 1.14 | 1.3E-02 |
| SMARCA5 | 1.14 | 3.9E-02 |
| MRPS31 | 1.14 | 3.9E-02 |
| ELAC2 | 1.14 | 1.0E-02 |
| BCL7A | 1.14 | 2.3E-02 |
| SCLY | 1.14 | 2.5E-02 |
| USO1 | 1.15 | 4.9E-02 |
| RANBP2 | 1.15 | 3.0E-02 |
| OCRL | 1.15 | 3.6E-02 |
| MTMR2 | 1.15 | 3.8E-02 |
| ASAP1 | 1.15 | 4.7E-02 |
| CSTF1 | 1.15 | 3.6E-02 |
| OARD1 | 1.15 | 6.1E-03 |
| RPL36 | 1.15 | 2.9E-02 |
| ARHGEF18 | 1.15 | 1.3E-02 |
| MSTO1 | 1.15 | 4.6E-02 |
| SIRT5 | 1.15 | 3.0E-02 |
| TNPO3 | 1.15 | 5.0E-03 |
| MAPK8 | 1.16 | 1.8E-02 |
| BUB1B | 1.16 | 3.0E-02 |
| MINA | 1.16 | 4.5E-02 |
| TTC5 | 1.16 | 4.3E-02 |
| MAFK | 1.16 | 4.0E-02 |
| TRMT5 | 1.16 | 3.5E-02 |
| FAM91A1 | 1.16 | 1.8E-02 |
| CARM1 | 1.16 | 4.5E-02 |
| EXOC3 | 1.16 | 1.4E-02 |
| GTF2H3 | 1.16 | 7.1E-03 |
| NCAPD2 | 1.16 | 4.3E-02 |
| FMNL1 | 1.16 | 5.4E-03 |
| DNAJC13 | 1.17 | 2.4E-02 |
| IPO7 | 1.17 | 2.5E-02 |
| DNAAF2 | 1.17 | 8.3E-03 |
| NAGLU | 1.17 | 8.6E-04 |
| OSBPL6 | 1.17 | 1.6E-02 |
| RSL24D1 | 1.17 | 2.0E-02 |
| FOXM1 | 1.17 | 6.9E-03 |
| ATG5 | 1.17 | 1.7E-02 |

| | | |
|---|---|---|
| SOGA1 | -1.03 | 2.5E-02 |
| SMARCB1 | -1.03 | 7.0E-03 |
| SEC31A | 1.02 | 3.7E-02 |
| API5 | 1.04 | 2.2E-02 |
| PMS1 | 1.04 | 4.9E-02 |
| ACOX1 | 1.04 | 3.6E-02 |
| PSMD13 | 1.05 | 1.6E-02 |
| INIP | 1.05 | 3.2E-02 |
| CNOT1 | 1.05 | 4.1E-02 |
| SEC24C | 1.05 | 2.8E-02 |
| COPB1 | 1.05 | 2.3E-02 |
| TRRAP | 1.05 | 2.8E-02 |
| RECQL | 1.06 | 3.5E-02 |
| HMGA1 | 1.06 | 4.0E-02 |
| ATPIF1 | 1.07 | 2.8E-02 |
| IGBP1 | 1.07 | 2.2E-02 |
| C19orf53 | 1.07 | 3.6E-02 |
| SAFB | 1.07 | 4.1E-02 |
| TBC1D13 | 1.08 | 6.0E-03 |
| TEX10 | 1.08 | 4.2E-02 |
| WDR5 | 1.08 | 5.9E-03 |
| SKIV2L | 1.08 | 3.9E-02 |
| SMARCD2 | 1.08 | 6.2E-03 |
| PGP | 1.08 | 9.4E-03 |
| VPS45 | 1.08 | 2.2E-02 |
| NCKAP1 | 1.08 | 3.3E-02 |
| AGPS | 1.08 | 2.5E-02 |
| CSTF1 | 1.08 | 2.5E-02 |
| PDE3A | 1.08 | 3.6E-03 |
| PPP2R2A | 1.08 | 4.0E-02 |
| MAP2K1 | 1.08 | 7.3E-03 |
| SF3B1 | 1.08 | 2.1E-02 |
| PSMA4 | 1.09 | 2.6E-02 |
| LAMTOR3 | 1.09 | 3.7E-02 |
| SNX24 | 1.09 | 9.3E-03 |
| PRKAB2 | 1.09 | 2.5E-02 |
| H1FX | 1.09 | 1.8E-02 |
| COPA | 1.09 | 7.7E-03 |
| SUMO1 | 1.09 | 4.8E-02 |
| MTMR1 | 1.09 | 2.6E-03 |
| SMC5 | 1.10 | 1.5E-02 |
| SNRPD2 | 1.10 | 1.4E-02 |
| DDX39A | 1.10 | 3.5E-02 |
| CYHR1 | 1.10 | 2.2E-02 |
| EIF2S2 | 1.10 | 5.0E-02 |
| HERC4 | 1.10 | 2.7E-02 |
| C8orf33 | 1.10 | 2.1E-02 |
| RMI1 | 1.10 | 5.0E-02 |
| CELF1 | 1.10 | 4.9E-02 |
| PPP2R5C | 1.10 | 1.8E-02 |
| PLAA | 1.10 | 1.4E-02 |
| PHC2 | 1.10 | 4.7E-02 |
| RPS17L | 1.10 | 3.9E-02 |
| NUDT16L1 | 1.10 | 2.6E-02 |
| PFDN1 | 1.11 | 1.0E-02 |
| KIAA1279 | 1.11 | 3.9E-02 |
| COPG1 | 1.11 | 3.7E-02 |
| MRPL16 | 1.11 | 3.0E-02 |
| HMGN1 | 1.11 | 3.2E-03 |
| CPNE3 | 1.11 | 1.7E-02 |
| ADSS | 1.11 | 1.3E-02 |
| UXT | 1.11 | 1.0E-02 |
| UBL4A | 1.11 | 1.8E-02 |
| UNC13B | 1.11 | 4.2E-02 |
| PHB | 1.11 | 4.3E-02 |
| ZC3H4 | 1.11 | 6.5E-03 |
| SAMD11 | 1.11 | 1.7E-02 |
| ZFYVE28 | 1.12 | 2.9E-02 |

| | | |
|---|---|---|
| ASF1A | -1.22 | 4.17E-02 |
| KIF13A | -1.22 | 4.18E-02 |
| WDR11 | -1.22 | 4.40E-02 |
| RRP1B | -1.22 | 3.27E-03 |
| EXOSC8 | -1.22 | 3.93E-02 |
| EFNA5 | -1.21 | 3.69E-02 |
| TOR3A | -1.21 | 3.33E-02 |
| GBF1 | -1.21 | 1.19E-02 |
| EXOC5 | -1.21 | 9.68E-03 |
| VAT1 | -1.21 | 4.35E-02 |
| DCXR | -1.21 | 2.58E-02 |
| SEH1L | -1.21 | 5.22E-03 |
| DNAJC13 | -1.21 | 1.52E-02 |
| NCDN | -1.21 | 2.35E-02 |
| TERF2IP | -1.21 | 1.38E-02 |
| VPS45 | -1.21 | 2.97E-02 |
| MGEA5 | -1.21 | 3.42E-02 |
| PPIE | -1.21 | 4.33E-02 |
| DHRS1 | -1.20 | 3.12E-02 |
| FTO | -1.20 | 3.98E-02 |
| RRN3 | -1.20 | 2.35E-02 |
| SPG21 | -1.20 | 1.65E-02 |
| HELLS | -1.20 | 4.25E-02 |
| MAP3K2 | -1.20 | 4.80E-02 |
| COG1 | -1.20 | 9.85E-04 |
| CCDC115 | -1.20 | 1.55E-02 |
| ME3 | -1.20 | 3.12E-02 |
| SUPT6H | -1.20 | 4.90E-02 |
| UBR1 | -1.20 | 1.52E-02 |
| TCEB3 | -1.19 | 2.01E-02 |
| GNB4 | -1.19 | 2.04E-02 |
| IQGAP1 | -1.19 | 4.92E-02 |
| THOC5 | -1.19 | 4.61E-02 |
| TBC1D24 | -1.19 | 3.57E-02 |
| MAPK3 | -1.19 | 4.70E-02 |
| MAEA | -1.19 | 4.89E-02 |
| IK | -1.18 | 1.64E-02 |
| AP3M1 | -1.18 | 3.81E-02 |
| AP1B1 | -1.18 | 1.15E-02 |
| FRYL | -1.18 | 3.17E-02 |
| BARD1 | -1.18 | 4.45E-02 |
| FLII | -1.18 | 4.78E-02 |
| MKL2 | -1.18 | 1.23E-02 |
| PSMC3IP | -1.18 | 2.78E-02 |
| IPO9 | -1.18 | 4.89E-02 |
| UBE4B | -1.18 | 1.71E-02 |
| RABGAP1 | -1.18 | 3.49E-02 |
| SMG5 | -1.18 | 9.91E-03 |
| BRCC3 | -1.18 | 8.13E-04 |
| PFAS | -1.17 | 4.25E-02 |
| PYROXD1 | -1.17 | 2.85E-02 |
| DHRS4 | -1.17 | 1.30E-02 |
| SRGAP2 | -1.17 | 3.76E-02 |
| IFRD1 | -1.17 | 5.25E-03 |
| STAT1 | -1.16 | 2.67E-03 |
| RNF213 | -1.16 | 2.59E-02 |
| USP48 | -1.16 | 3.44E-02 |
| NEK9 | -1.16 | 4.36E-03 |
| URGCP | -1.16 | 2.56E-02 |
| AMACR | -1.16 | 4.04E-02 |
| EEF1D | -1.16 | 3.52E-03 |
| ASXL1 | -1.16 | 3.14E-02 |
| KLHL13 | -1.16 | 2.22E-02 |
| MEIS1 | -1.15 | 3.75E-02 |
| CTSZ | -1.15 | 4.97E-02 |
| RMI1 | -1.15 | 4.74E-02 |
| KBTBD4 | -1.15 | 1.84E-02 |
| GTPBP8 | -1.15 | 4.30E-03 |

| | | |
|---|---|---|
| NCAPG2 | 1.18 | 3.09E-02 |
| ARPC4 | 1.18 | 4.57E-02 |
| SETD8 | 1.18 | 3.31E-03 |
| HPDL | 1.18 | 3.53E-02 |
| CBLB | 1.18 | 2.05E-02 |
| APTX | 1.18 | 4.80E-02 |
| KIF23 | 1.18 | 4.01E-02 |
| POLE | 1.18 | 3.93E-02 |
| CD3EAP | 1.18 | 3.65E-03 |
| POLR1E | 1.18 | 2.90E-02 |
| ARHGEF10 | 1.18 | 4.95E-02 |
| MYO10 | 1.18 | 4.19E-02 |
| ARF4 | 1.18 | 1.26E-02 |
| ARPC2 | 1.18 | 2.50E-02 |
| FHL1 | 1.18 | 2.74E-03 |
| FAM160A2 | 1.18 | 2.09E-02 |
| CNNM3 | 1.19 | 3.29E-04 |
| ERCC3 | 1.19 | 2.52E-02 |
| IPO7 | 1.19 | 1.61E-02 |
| NUCB2 | 1.19 | 4.93E-02 |
| NCAPD3 | 1.19 | 2.80E-02 |
| STIL | 1.19 | 3.15E-03 |
| PELP1 | 1.19 | 1.60E-02 |
| XPOT | 1.19 | 2.80E-03 |
| TBL3 | 1.19 | 1.01E-02 |
| MAP2K1 | 1.19 | 8.85E-03 |
| TRIM16 | 1.19 | 1.13E-02 |
| PRC1 | 1.19 | 2.48E-02 |
| NEDD4 | 1.19 | 4.61E-02 |
| ADD1 | 1.19 | 6.45E-03 |
| ACSL4 | 1.19 | 6.71E-03 |
| KNTC1 | 1.19 | 2.28E-02 |
| RBM39 | 1.19 | 3.51E-02 |
| IGF2BP1 | 1.20 | 3.49E-02 |
| GPN1 | 1.20 | 8.45E-03 |
| PRR11 | 1.20 | 1.81E-03 |
| DOCK10 | 1.20 | 9.23E-03 |
| PPP4R1 | 1.20 | 3.05E-03 |
| FOXC2 | 1.20 | 4.02E-02 |
| ARIH1 | 1.20 | 2.58E-02 |
| GTF3C4 | 1.20 | 3.39E-02 |
| FAM105B | 1.21 | 2.90E-02 |
| HAUS2 | 1.21 | 1.12E-02 |
| ACTR3 | 1.21 | 2.22E-02 |
| NT5C | 1.21 | 3.65E-02 |
| MACF1 | 1.21 | 4.81E-02 |
| GTF2H4 | 1.21 | 3.39E-02 |
| CCZ1 | 1.21 | 3.86E-02 |
| EIF2AK4 | 1.21 | 2.74E-02 |
| LCMT2 | 1.21 | 7.33E-03 |
| PFKP | 1.21 | 3.55E-03 |
| CLIC4 | 1.22 | 4.62E-02 |
| NGRN | 1.22 | 4.02E-03 |
| ACY1 | 1.22 | 9.03E-03 |
| SEMA3C | 1.22 | 2.76E-02 |
| MTG1 | 1.22 | 3.26E-02 |
| POLG2 | 1.22 | 1.94E-02 |
| LYN | 1.22 | 5.00E-02 |
| ADD3 | 1.22 | 1.83E-02 |
| FAM129B | 1.22 | 2.44E-02 |
| LTN1 | 1.22 | 4.04E-02 |
| COASY | 1.22 | 4.63E-02 |
| AP3S1 | 1.22 | 2.44E-03 |
| SEC14L1 | 1.22 | 2.18E-02 |
| TK1 | 1.22 | 4.55E-02 |
| PXK | 1.23 | 2.00E-04 |
| APEX1 | 1.23 | 1.35E-02 |
| ERP29 | 1.23 | 6.90E-03 |

| | | |
|---|---|---|
| PPP4R1 | 1.17 | 1.9E-02 |
| PPM1B | 1.17 | 2.5E-02 |
| NUDT16 | 1.18 | 4.6E-02 |
| IARS | 1.18 | 4.6E-02 |
| RB1CC1 | 1.18 | 1.4E-02 |
| LSM1 | 1.18 | 1.5E-02 |
| ECD | 1.18 | 4.4E-02 |
| AJUBA | 1.18 | 1.1E-02 |
| CAMK1D | 1.18 | 4.1E-02 |
| TFCP2 | 1.18 | 3.0E-02 |
| RPL39 | 1.18 | 4.0E-02 |
| PUM2 | 1.18 | 2.8E-02 |
| MED17 | 1.18 | 2.4E-02 |
| CELF1 | 1.19 | 4.4E-02 |
| WDR35 | 1.19 | 2.7E-02 |
| WIPI2 | 1.19 | 2.3E-02 |
| PYGO2 | 1.19 | 3.7E-02 |
| MYO5A | 1.19 | 1.5E-02 |
| CENPB | 1.19 | 2.4E-02 |
| NOL11 | 1.19 | 3.4E-02 |
| TOLLIP | 1.19 | 4.3E-02 |
| PHB | 1.19 | 6.9E-03 |
| BAZ2A | 1.19 | 4.5E-02 |
| POLR1B | 1.19 | 1.1E-02 |
| TACC2 | 1.19 | 1.5E-02 |
| EXOC6 | 1.19 | 4.3E-02 |
| TNRC6B | 1.19 | 4.9E-02 |
| FNBP1L | 1.19 | 3.2E-02 |
| PRKCG | 1.19 | 4.4E-02 |
| CARD6 | 1.20 | 1.9E-02 |
| BDH2 | 1.20 | 4.5E-02 |
| CCDC137 | 1.20 | 1.7E-02 |
| MID1 | 1.20 | 1.8E-02 |
| PROSER2 | 1.20 | 4.6E-03 |
| SERPINB8 | 1.20 | 4.7E-02 |
| KIAA0020 | 1.20 | 1.2E-02 |
| CYTH3 | 1.20 | 1.1E-02 |
| COMMD8 | 1.20 | 1.3E-02 |
| CGA | 1.20 | 2.8E-02 |
| CSTF3 | 1.20 | 2.0E-02 |
| GTF2H4 | 1.20 | 4.0E-02 |
| ALDOC | 1.20 | 2.0E-02 |
| RBM19 | 1.20 | 4.6E-02 |
| SEPT11 | 1.20 | 3.3E-02 |
| CRLF3 | 1.21 | 4.0E-02 |
| ISCA2 | 1.21 | 2.6E-02 |
| ZC3H15 | 1.21 | 1.3E-02 |
| DUSP3 | 1.21 | 1.2E-02 |
| LSM14B | 1.21 | 4.6E-02 |
| DDX10 | 1.21 | 4.5E-02 |
| AIFM2 | 1.21 | 1.1E-02 |
| RIPK2 | 1.21 | 3.3E-04 |
| PCGF2 | 1.21 | 1.1E-02 |
| WDR6 | 1.21 | 1.9E-02 |
| UBA5 | 1.21 | 4.0E-02 |
| UCHL3 | 1.21 | 3.4E-02 |
| APBB1 | 1.21 | 1.5E-02 |
| TIMM13 | 1.21 | 1.0E-02 |
| HSPE1 | 1.22 | 4.5E-02 |
| HAUS2 | 1.22 | 2.9E-02 |
| PRKRIR | 1.22 | 1.8E-02 |
| PANK2 | 1.22 | 3.3E-02 |
| SAAL1 | 1.22 | 1.4E-02 |
| MRPL53 | 1.22 | 5.0E-02 |
| PNO1 | 1.22 | 3.5E-02 |
| DUSP12 | 1.22 | 1.6E-02 |
| LRCH1 | 1.22 | 3.4E-02 |
| DDX6 | 1.22 | 1.7E-02 |

| | | |
|---|---|---|
| RPS21 | 1.12 | 1.7E-02 |
| MUC16 | 1.12 | 3.5E-02 |
| GTF3C1 | 1.12 | 5.9E-03 |
| CRNKL1 | 1.12 | 1.1E-02 |
| THOC2 | 1.12 | 3.9E-02 |
| PPP2R1A | 1.12 | 5.0E-02 |
| POLR3B | 1.12 | 2.5E-02 |
| GCN1L1 | 1.12 | 4.3E-03 |
| DYNC1H1 | 1.12 | 1.1E-02 |
| CASK | 1.12 | 4.2E-02 |
| LYSMD1 | 1.12 | 3.3E-02 |
| USP7 | 1.12 | 2.4E-02 |
| PSMA3 | 1.12 | 2.2E-02 |
| PDS5A | 1.12 | 9.7E-03 |
| FAM160A2 | 1.12 | 1.7E-02 |
| WASF1 | 1.12 | 2.3E-02 |
| KDM1A | 1.12 | 1.7E-02 |
| ISG15 | 1.13 | 2.1E-03 |
| TUBA4A | 1.13 | 4.5E-02 |
| TSN | 1.13 | 3.0E-02 |
| METAP1 | 1.13 | 4.7E-02 |
| RAPGEF6 | 1.13 | 1.1E-03 |
| FKBP5 | 1.13 | 1.1E-05 |
| NAE1 | 1.13 | 1.9E-02 |
| RCN1 | 1.13 | 3.0E-02 |
| PPP1R11 | 1.13 | 4.4E-02 |
| WAC | 1.13 | 3.5E-03 |
| MSH2 | 1.13 | 1.2E-02 |
| SF3B5 | 1.13 | 4.3E-02 |
| CORO7 | 1.13 | 1.3E-02 |
| ATP6V1F | 1.13 | 3.0E-02 |
| DLGAP4 | 1.13 | 2.4E-04 |
| CCNT1 | 1.13 | 4.6E-02 |
| BPNT1 | 1.13 | 3.1E-02 |
| NFIC | 1.14 | 3.2E-02 |
| FAM177A1 | 1.14 | 2.4E-02 |
| MSTO1 | 1.14 | 4.7E-02 |
| GTF3C3 | 1.14 | 1.9E-02 |
| KLHL13 | 1.14 | 1.3E-02 |
| SAMD4B | 1.14 | 9.1E-03 |
| AP1G1 | 1.14 | 3.5E-02 |
| SAR1A | 1.14 | 3.3E-02 |
| AGO2 | 1.14 | 2.0E-02 |
| RRBP1 | 1.14 | 4.9E-02 |
| PRKAG1 | 1.14 | 4.0E-03 |
| NUMB | 1.14 | 3.9E-02 |
| STAG2 | 1.14 | 4.7E-02 |
| HEXA | 1.14 | 1.4E-02 |
| TBC1D15 | 1.14 | 1.3E-02 |
| ERH | 1.14 | 1.5E-02 |
| CASP7 | 1.14 | 5.0E-02 |
| GAK | 1.14 | 8.1E-03 |
| AP2B1 | 1.14 | 2.5E-02 |
| DIS3 | 1.15 | 1.2E-02 |
| ECM29 | 1.15 | 1.9E-02 |
| ERCC6L | 1.15 | 4.0E-02 |
| TUBGCP3 | 1.15 | 4.0E-03 |
| BOLA1 | 1.15 | 1.4E-03 |
| PPOX | 1.15 | 4.1E-02 |
| HNRNPUL2 | 1.15 | 4.3E-02 |
| GYS1 | 1.15 | 5.0E-02 |
| FEN1 | 1.15 | 4.0E-02 |
| MARS | 1.15 | 2.1E-02 |
| MSH6 | 1.15 | 2.2E-02 |
| JUNB | 1.16 | 5.8E-03 |
| CBWD1 | 1.16 | 2.7E-02 |
| IGF2BP1 | 1.16 | 2.1E-02 |
| MSH3 | 1.16 | 1.1E-02 |

| | | |
|---|---|---|
| SEC31A | -1.15 | 1.57E-02 |
| ASCC1 | -1.15 | 3.50E-02 |
| MLH1 | -1.15 | 3.26E-02 |
| PRKAB2 | -1.15 | 4.33E-02 |
| ASNA1 | -1.15 | 3.27E-02 |
| DOCK10 | -1.15 | 2.89E-02 |
| SKIV2L | -1.14 | 3.84E-02 |
| SEC24C | -1.14 | 1.27E-02 |
| C7orf25 | -1.14 | 2.77E-02 |
| CARKD | -1.14 | 1.81E-02 |
| KIAA1033 | -1.14 | 2.51E-02 |
| CYFIP2 | -1.13 | 2.26E-02 |
| KIFAP3 | -1.13 | 2.51E-03 |
| GTF2IRD1 | -1.13 | 2.81E-02 |
| NF1 | -1.13 | 9.38E-03 |
| RPL35A | -1.13 | 3.15E-02 |
| EXOSC5 | -1.13 | 1.65E-02 |
| TAF6L | -1.13 | 4.10E-02 |
| BRF1 | -1.13 | 4.30E-02 |
| AP3D1 | -1.12 | 2.20E-02 |
| EXOC6B | -1.12 | 3.10E-02 |
| TTC27 | -1.12 | 2.24E-02 |
| FMNL1 | -1.12 | 3.30E-02 |
| TANGO6 | -1.12 | 1.39E-02 |
| YWHAB | -1.12 | 2.82E-02 |
| CASP8 | -1.11 | 1.49E-02 |
| RAPH1 | -1.11 | 3.37E-02 |
| RAB3GAP2 | -1.10 | 1.84E-02 |
| LTN1 | -1.10 | 4.35E-02 |
| LCMT2 | -1.10 | 4.89E-02 |
| AKR1B1 | -1.09 | 8.13E-03 |
| CD109 | -1.09 | 4.03E-02 |
| DYNC1H1 | -1.09 | 9.00E-03 |
| PARP4 | -1.08 | 1.30E-02 |
| STRIP1 | -1.08 | 1.74E-02 |
| GCN1L1 | -1.08 | 1.49E-02 |
| MTA3 | -1.07 | 4.62E-02 |
| C8orf33 | -1.07 | 2.81E-02 |
| ELP2 | -1.06 | 1.36E-02 |
| INTS3 | -1.05 | 3.63E-03 |
| RMND5A | -1.05 | 1.78E-02 |
| C19orf53 | -1.05 | 3.07E-02 |
| LRBA | -1.05 | 1.80E-02 |
| ASPM | -1.04 | 6.65E-03 |
| DPH5 | -1.03 | 6.31E-04 |
| FANCI | -1.03 | 2.94E-03 |
| USP24 | 1.01 | 3.08E-03 |
| PDE3A | 1.04 | 3.56E-02 |
| TFCP2 | 1.05 | 2.47E-02 |
| PPP2R5C | 1.05 | 6.89E-03 |
| WDR3 | 1.07 | 2.56E-02 |
| TUBGCP3 | 1.08 | 4.49E-02 |
| API5 | 1.08 | 4.35E-03 |
| CNOT1 | 1.09 | 2.39E-02 |
| GTF3C1 | 1.09 | 2.74E-02 |
| VAC14 | 1.09 | 5.00E-02 |
| ZFYVE28 | 1.09 | 3.97E-02 |
| KIAA1429 | 1.09 | 4.64E-02 |
| IPO11 | 1.10 | 3.44E-02 |
| TEX10 | 1.10 | 1.33E-02 |
| DIEXF | 1.10 | 3.76E-02 |
| BTAF1 | 1.10 | 3.33E-02 |
| OSBP2 | 1.10 | 3.26E-02 |
| TBC1D2B | 1.10 | 2.27E-02 |
| C4A | 1.10 | 4.34E-02 |
| SRP9 | 1.10 | 4.21E-02 |
| NUP155 | 1.11 | 1.04E-02 |
| NIPSNAP1 | 1.11 | 4.72E-02 |

| | | |
|---|---|---|
| MYO5A | 1.23 | 4.61E-02 |
| RRBP1 | 1.23 | 1.15E-02 |
| PLK1 | 1.23 | 4.30E-02 |
| MRM1 | 1.23 | 2.13E-02 |
| TEX10 | 1.23 | 1.69E-02 |
| EML4 | 1.23 | 2.08E-02 |
| CYTH3 | 1.23 | 2.15E-02 |
| PGM3 | 1.23 | 2.30E-02 |
| FHOD1 | 1.23 | 2.74E-02 |
| TSEN15 | 1.23 | 1.30E-02 |
| STAMBPL1 | 1.24 | 1.26E-02 |
| AK4 | 1.24 | 2.65E-02 |
| NFIX | 1.24 | 7.40E-03 |
| DSP | 1.24 | 4.03E-02 |
| HPS5 | 1.24 | 3.34E-02 |
| DOCK4 | 1.24 | 4.23E-02 |
| TMPO | 1.24 | 8.73E-03 |
| CAP1 | 1.24 | 2.39E-02 |
| TTLL12 | 1.24 | 1.88E-02 |
| NF2 | 1.24 | 6.58E-04 |
| CRLF1 | 1.24 | 4.69E-02 |
| EYA3 | 1.24 | 2.84E-02 |
| MYO1C | 1.24 | 3.80E-02 |
| ELP2 | 1.24 | 1.61E-02 |
| FANCI | 1.24 | 4.05E-03 |
| BCR | 1.24 | 2.24E-02 |
| DAPK3 | 1.24 | 2.56E-02 |
| UNC13D | 1.25 | 1.03E-02 |
| ZC3HAV1L | 1.25 | 2.57E-02 |
| CNOT7 | 1.25 | 1.83E-02 |
| CRYZ | 1.25 | 1.34E-03 |
| MGME1 | 1.25 | 1.96E-03 |
| SNCG | 1.25 | 2.76E-02 |
| IFRD1 | 1.25 | 1.75E-02 |
| QDPR | 1.25 | 1.68E-02 |
| PWP2 | 1.25 | 3.06E-03 |
| PLEK2 | 1.25 | 4.89E-02 |
| TRAF2 | 1.25 | 1.28E-02 |
| PARD3 | 1.25 | 4.65E-02 |
| TDG | 1.25 | 3.63E-02 |
| DVL1 | 1.25 | 2.03E-02 |
| CD97 | 1.25 | 7.64E-04 |
| MYL9 | 1.25 | 2.35E-02 |
| SMTN | 1.26 | 2.67E-02 |
| ABHD14B | 1.26 | 3.92E-02 |
| PALLD | 1.26 | 3.65E-02 |
| PDLIM5 | 1.26 | 3.04E-02 |
| JUP | 1.26 | 1.07E-02 |
| CLK1 | 1.26 | 6.35E-03 |
| CRTAP | 1.26 | 3.57E-02 |
| YARS | 1.26 | 1.66E-02 |
| CDC20 | 1.26 | 4.49E-02 |
| ACTR2 | 1.26 | 3.05E-03 |
| IARS | 1.26 | 7.53E-03 |
| CIT | 1.26 | 3.63E-02 |
| LIMS1 | 1.27 | 2.58E-02 |
| TRIP10 | 1.27 | 4.96E-02 |
| CPOX | 1.27 | 1.43E-02 |
| PXN | 1.27 | 2.77E-02 |
| ACOT2 | 1.27 | 4.44E-02 |
| CAPN2 | 1.28 | 6.54E-03 |
| ANP32A | 1.28 | 1.98E-02 |
| C9orf64 | 1.28 | 1.83E-02 |
| SERF2 | 1.28 | 1.99E-02 |
| IVD | 1.28 | 4.06E-02 |
| IL18 | 1.28 | 6.03E-03 |
| AAGAB | 1.28 | 9.90E-03 |
| ADAL | 1.28 | 4.44E-02 |

| | | |
|---|---|---|
| MTMR10 | 1.22 | 4.4E-02 |
| NHLRC2 | 1.22 | 3.8E-02 |
| OTUD4 | 1.22 | 5.0E-02 |
| MPDZ | 1.22 | 2.1E-02 |
| CTSZ | 1.22 | 4.7E-03 |
| MDC1 | 1.22 | 2.9E-02 |
| NDUFAF2 | 1.23 | 4.9E-02 |
| CHRAC1 | 1.23 | 3.8E-02 |
| DAPK3 | 1.23 | 2.9E-02 |
| MTMR6 | 1.23 | 3.0E-02 |
| ENSA | 1.23 | 4.7E-02 |
| NPM3 | 1.23 | 4.1E-02 |
| COMMD6 | 1.23 | 7.0E-03 |
| SPECC1 | 1.23 | 1.8E-02 |
| PLK1 | 1.23 | 4.1E-02 |
| FAM129B | 1.23 | 3.4E-02 |
| TRIO | 1.23 | 3.8E-02 |
| KIF5A | 1.23 | 3.0E-02 |
| GTF3C4 | 1.23 | 2.6E-02 |
| TTK | 1.23 | 1.7E-02 |
| EHBP1L1 | 1.24 | 2.8E-02 |
| CHD5 | 1.24 | 2.7E-02 |
| COMMD9 | 1.24 | 7.0E-05 |
| RAB11FIP1 | 1.24 | 1.2E-02 |
| ALG2 | 1.24 | 4.7E-02 |
| SAFB | 1.24 | 2.4E-02 |
| PAK1 | 1.24 | 3.2E-02 |
| XPO5 | 1.24 | 1.5E-02 |
| INO80B | 1.24 | 3.2E-02 |
| PRKRIP1 | 1.24 | 3.8E-02 |
| ZADH2 | 1.24 | 1.9E-02 |
| HMG20A | 1.24 | 4.6E-02 |
| ERI1 | 1.25 | 4.4E-02 |
| C6orf211 | 1.25 | 1.5E-02 |
| PYCR1 | 1.25 | 2.8E-02 |
| CACTIN | 1.25 | 3.5E-02 |
| EPN2 | 1.25 | 1.7E-02 |
| CBLB | 1.25 | 1.3E-02 |
| MYO10 | 1.25 | 2.8E-02 |
| HEATR6 | 1.25 | 4.8E-02 |
| BLM | 1.25 | 5.1E-03 |
| ARHGAP29 | 1.25 | 1.3E-02 |
| RNASEH2B | 1.25 | 3.7E-02 |
| MLLT4 | 1.25 | 4.3E-02 |
| DHX34 | 1.26 | 6.8E-04 |
| ARHGAP10 | 1.26 | 4.6E-02 |
| MCC | 1.27 | 5.0E-03 |
| MED24 | 1.27 | 4.3E-02 |
| UBR2 | 1.27 | 3.8E-03 |
| PROCR | 1.27 | 1.4E-02 |
| RPRD1B | 1.27 | 4.8E-02 |
| MYBBP1A | 1.27 | 2.3E-02 |
| DDX18 | 1.27 | 4.9E-02 |
| ZNF259 | 1.27 | 4.8E-02 |
| ARMC9 | 1.27 | 5.4E-03 |
| CDC42EP1 | 1.28 | 2.3E-02 |
| RAPH1 | 1.28 | 5.5E-03 |
| POLR1A | 1.28 | 2.6E-03 |
| NLE1 | 1.28 | 2.9E-02 |
| PARVA | 1.28 | 6.2E-03 |
| NF2 | 1.28 | 1.1E-03 |
| DDX21 | 1.28 | 1.5E-02 |
| PICALM | 1.28 | 3.4E-03 |
| MORC2 | 1.29 | 4.9E-03 |
| MYO1B | 1.29 | 1.7E-03 |
| VCL | 1.29 | 4.7E-02 |
| POLG2 | 1.29 | 1.5E-02 |
| LIG4 | 1.29 | 4.0E-02 |

| | | |
|---|---|---|
| AP2A1 | 1.16 | 4.0E-02 |
| SOD2 | 1.16 | 3.8E-02 |
| SPTAN1 | 1.16 | 2.7E-02 |
| CACYBP | 1.16 | 8.8E-03 |
| PACSIN3 | 1.16 | 3.3E-02 |
| DUT | 1.16 | 1.6E-02 |
| EML3 | 1.16 | 2.2E-02 |
| PPP1R7 | 1.16 | 3.3E-02 |
| SPTBN1 | 1.16 | 2.5E-02 |
| TSEN15 | 1.16 | 3.5E-02 |
| PRCP | 1.16 | 2.5E-02 |
| PWWP2A | 1.16 | 1.4E-02 |
| HRSP12 | 1.16 | 4.5E-02 |
| IDH1 | 1.16 | 1.9E-02 |
| MGEA5 | 1.16 | 2.3E-02 |
| NIT1 | 1.17 | 2.1E-02 |
| HYI | 1.17 | 4.0E-02 |
| IARS | 1.17 | 2.0E-02 |
| KIF14 | 1.17 | 4.5E-02 |
| C19orf43 | 1.17 | 2.2E-02 |
| GGA2 | 1.17 | 3.4E-02 |
| DHRS1 | 1.17 | 1.8E-02 |
| METTL18 | 1.17 | 7.6E-03 |
| AP3S1 | 1.17 | 4.9E-02 |
| UBE2A | 1.17 | 2.5E-02 |
| CPEB3 | 1.17 | 1.8E-03 |
| FAM105B | 1.17 | 2.4E-02 |
| DAP | 1.17 | 2.5E-02 |
| C15orf39 | 1.17 | 2.4E-02 |
| LIN54 | 1.17 | 8.3E-03 |
| FAM129B | 1.17 | 6.9E-03 |
| RAB3GAP2 | 1.17 | 1.0E-03 |
| TXNDC5 | 1.18 | 4.6E-03 |
| AP3D1 | 1.18 | 1.7E-02 |
| PGM2 | 1.18 | 8.9E-03 |
| SNX1 | 1.18 | 3.1E-02 |
| MICU2 | 1.18 | 4.3E-02 |
| CHTOP | 1.18 | 2.2E-03 |
| RNASET2 | 1.18 | 5.6E-04 |
| AJUBA | 1.18 | 9.3E-03 |
| HNRNPU | 1.18 | 2.8E-02 |
| DPP9 | 1.18 | 3.7E-02 |
| PPP4C | 1.18 | 2.0E-02 |
| RPRD1B | 1.18 | 3.4E-02 |
| PHIP | 1.18 | 5.2E-03 |
| FIBP | 1.18 | 4.8E-02 |
| NIPBL | 1.18 | 9.0E-03 |
| MBD1 | 1.18 | 6.8E-03 |
| RASA1 | 1.18 | 8.7E-03 |
| PRC1 | 1.18 | 2.6E-02 |
| ACOT8 | 1.18 | 6.0E-03 |
| CD109 | 1.18 | 2.1E-03 |
| PICALM | 1.18 | 8.7E-03 |
| CCNA2 | 1.19 | 3.1E-02 |
| PDS5B | 1.19 | 2.7E-02 |
| ARHGAP5 | 1.19 | 3.5E-03 |
| AURKA | 1.19 | 1.2E-02 |
| SYMPK | 1.19 | 2.4E-03 |
| TRAPPC8 | 1.19 | 1.5E-03 |
| EFTUD1 | 1.19 | 3.4E-02 |
| MTMR2 | 1.19 | 2.2E-02 |
| KCTD3 | 1.19 | 3.0E-02 |
| ADD3 | 1.19 | 2.9E-02 |
| PRPF39 | 1.19 | 2.5E-03 |
| CAST | 1.19 | 4.8E-02 |
| PFKL | 1.19 | 3.8E-02 |
| KIF4A | 1.19 | 1.5E-02 |
| NTPCR | 1.19 | 4.3E-02 |

| | | |
|---|---|---|
| TNPO3 | 1.11 | 2.46E-03 |
| RANBP2 | 1.11 | 4.74E-02 |
| SRSF5 | 1.11 | 3.98E-02 |
| RBM25 | 1.11 | 3.17E-02 |
| KAT8 | 1.11 | 9.25E-03 |
| CENPC | 1.12 | 3.39E-02 |
| PFDN6 | 1.12 | 2.64E-02 |
| SMC5 | 1.12 | 1.37E-02 |
| TRRAP | 1.12 | 6.69E-03 |
| ABL1 | 1.12 | 4.99E-02 |
| DDX18 | 1.13 | 3.87E-03 |
| CWC22 | 1.13 | 3.20E-02 |
| TRIM25 | 1.13 | 5.83E-03 |
| MYCBP2 | 1.13 | 3.95E-02 |
| LAS1L | 1.13 | 4.40E-02 |
| SMG1 | 1.13 | 2.77E-03 |
| AURKA | 1.14 | 7.74E-03 |
| RPP30 | 1.14 | 4.49E-02 |
| SYTL4 | 1.14 | 1.94E-02 |
| USP9X | 1.14 | 7.37E-03 |
| UPF1 | 1.14 | 4.77E-02 |
| LIN54 | 1.14 | 1.52E-02 |
| NOC2L | 1.14 | 3.40E-02 |
| POLE | 1.14 | 1.43E-02 |
| CARM1 | 1.14 | 1.83E-02 |
| FOXK1 | 1.15 | 4.58E-02 |
| USP39 | 1.15 | 2.02E-02 |
| SMARCC2 | 1.15 | 4.80E-02 |
| HDAC3 | 1.15 | 6.01E-03 |
| TTI2 | 1.15 | 2.69E-02 |
| IKBKB | 1.15 | 2.03E-02 |
| TOPBP1 | 1.15 | 1.75E-02 |
| PAPOLA | 1.15 | 2.71E-02 |
| NDUFAF7 | 1.15 | 1.93E-02 |
| HUS1 | 1.15 | 1.35E-02 |
| GSTK1 | 1.15 | 5.97E-03 |
| XRN2 | 1.15 | 4.03E-02 |
| POLA1 | 1.15 | 1.56E-02 |
| ARHGEF18 | 1.15 | 1.96E-02 |
| CLP1 | 1.16 | 1.21E-02 |
| GSK3A | 1.16 | 4.99E-03 |
| PDZD11 | 1.16 | 1.49E-02 |
| SBF1 | 1.16 | 6.26E-03 |
| UTP6 | 1.16 | 1.35E-02 |
| CXorf56 | 1.16 | 4.73E-02 |
| NPEPL1 | 1.17 | 2.35E-03 |
| PAPD5 | 1.17 | 6.68E-03 |
| GLMN | 1.17 | 3.35E-02 |
| ARF5 | 1.17 | 2.21E-02 |
| RIOK1 | 1.17 | 3.56E-02 |
| TEX30 | 1.17 | 4.09E-02 |
| NFIX | 1.17 | 4.13E-02 |
| NCAPD2 | 1.17 | 2.82E-02 |
| NGRN | 1.17 | 2.31E-02 |
| POP7 | 1.17 | 4.94E-02 |
| SLFN5 | 1.17 | 2.57E-04 |
| LARS | 1.18 | 1.74E-02 |
| KDM1A | 1.18 | 2.80E-03 |
| ECHDC1 | 1.18 | 1.64E-02 |
| CARD6 | 1.18 | 4.27E-02 |
| JUND | 1.18 | 4.36E-02 |
| ISG15 | 1.18 | 5.24E-03 |
| SAFB | 1.18 | 2.75E-02 |
| POLR3B | 1.18 | 4.54E-02 |
| MRPS11 | 1.19 | 4.87E-02 |
| RALGAPB | 1.19 | 8.70E-03 |
| CAST | 1.19 | 4.68E-02 |
| DDX31 | 1.19 | 1.33E-02 |

| | | |
|---|---|---|
| RBMS2 | 1.29 | 4.38E-02 |
| DNAJC17 | 1.29 | 1.03E-02 |
| FRYL | 1.29 | 1.50E-02 |
| CARS | 1.29 | 1.82E-02 |
| CORO1C | 1.29 | 2.23E-02 |
| BAP18 | 1.29 | 1.98E-02 |
| VAT1L | 1.29 | 3.00E-02 |
| SNX18 | 1.29 | 3.24E-02 |
| RAB3IL1 | 1.29 | 2.70E-03 |
| ALG2 | 1.30 | 1.08E-02 |
| TAGLN | 1.30 | 2.46E-02 |
| APP | 1.30 | 3.34E-02 |
| TXNDC5 | 1.30 | 7.37E-03 |
| MDN1 | 1.30 | 2.41E-02 |
| GBP1 | 1.30 | 2.82E-02 |
| ISG15 | 1.30 | 2.10E-03 |
| INF2 | 1.30 | 6.89E-04 |
| NOL6 | 1.30 | 8.12E-04 |
| PPP1R18 | 1.30 | 2.37E-02 |
| TJP1 | 1.30 | 2.85E-02 |
| SORD | 1.30 | 1.42E-02 |
| CSK | 1.30 | 1.76E-02 |
| TGFB1I1 | 1.30 | 3.79E-02 |
| ALDH1A2 | 1.30 | 1.20E-02 |
| COTL1 | 1.31 | 2.12E-02 |
| LSMD1 | 1.31 | 2.41E-02 |
| ZFYVE21 | 1.31 | 4.41E-02 |
| CDK6 | 1.31 | 2.19E-03 |
| C4A | 1.31 | 4.97E-02 |
| MEMO1 | 1.31 | 5.93E-03 |
| VASP | 1.31 | 4.73E-02 |
| RAB11FIP5 | 1.31 | 3.97E-02 |
| VARS2 | 1.32 | 4.72E-02 |
| KCTD7 | 1.32 | 1.44E-02 |
| IFRD2 | 1.32 | 4.75E-02 |
| IQGAP1 | 1.32 | 1.49E-02 |
| TRAPPC9 | 1.32 | 4.80E-02 |
| C15orf39 | 1.32 | 6.37E-03 |
| KIAA0020 | 1.32 | 2.41E-02 |
| MICAL1 | 1.32 | 1.56E-02 |
| ARFGEF2 | 1.33 | 7.35E-03 |
| FSCN1 | 1.33 | 3.87E-02 |
| HACL1 | 1.34 | 1.07E-02 |
| LUM | 1.34 | 4.23E-02 |
| ISOC1 | 1.34 | 2.01E-02 |
| BCAR3 | 1.35 | 3.04E-02 |
| TPRG1L | 1.35 | 2.48E-02 |
| PITPNC1 | 1.35 | 4.29E-03 |
| CRMP1 | 1.35 | 3.05E-02 |
| SHB | 1.35 | 3.23E-02 |
| DIXDC1 | 1.35 | 3.48E-02 |
| MLLT3 | 1.35 | 4.81E-02 |
| RND3 | 1.35 | 1.47E-02 |
| TLN1 | 1.35 | 3.58E-02 |
| BAG1 | 1.35 | 2.75E-02 |
| UBR1 | 1.36 | 9.14E-03 |
| S100A2 | 1.36 | 2.65E-02 |
| KLF13 | 1.36 | 3.87E-02 |
| DUT | 1.36 | 2.03E-03 |
| PLCB4 | 1.36 | 3.21E-02 |
| TYMS | 1.36 | 5.41E-03 |
| EHD4 | 1.36 | 6.98E-03 |
| ANXA3 | 1.37 | 1.95E-02 |
| PTRH1 | 1.37 | 4.72E-03 |
| TLDC1 | 1.37 | 4.88E-02 |
| FAM120B | 1.37 | 1.81E-02 |
| CBS | 1.37 | 1.15E-02 |
| L3HYPDH | 1.37 | 2.86E-02 |

| | | |
|---|---|---|
| PPP1R18 | 1.29 | 4.1E-02 |
| SEC14L1 | 1.29 | 1.6E-02 |
| MYL6 | 1.29 | 4.7E-02 |
| DDX47 | 1.29 | 1.2E-02 |
| FOXK1 | 1.29 | 1.6E-02 |
| CNPY3 | 1.29 | 2.7E-02 |
| TRIM25 | 1.30 | 3.7E-02 |
| PUS1 | 1.30 | 4.7E-02 |
| EEF1B2 | 1.30 | 2.8E-02 |
| STK32C | 1.30 | 1.1E-02 |
| CARS | 1.30 | 4.5E-02 |
| MPRIP | 1.30 | 4.4E-02 |
| PEAK1 | 1.30 | 1.7E-02 |
| DOHH | 1.30 | 8.0E-03 |
| MAPKAPK3 | 1.30 | 3.7E-02 |
| METTL13 | 1.30 | 3.3E-02 |
| HMHA1 | 1.31 | 3.6E-02 |
| EML4 | 1.31 | 2.0E-02 |
| BAIAP2L1 | 1.31 | 4.5E-02 |
| ZCCHC8 | 1.31 | 3.8E-02 |
| SUV39H2 | 1.31 | 3.9E-02 |
| CLK1 | 1.31 | 4.0E-04 |
| NAV2 | 1.31 | 4.7E-02 |
| DLG5 | 1.31 | 2.2E-03 |
| PMM1 | 1.31 | 3.5E-02 |
| PTGES2 | 1.31 | 1.4E-02 |
| GNB1L | 1.32 | 3.9E-02 |
| AURKB | 1.32 | 5.0E-02 |
| STK10 | 1.32 | 3.7E-02 |
| POLR1E | 1.32 | 1.2E-02 |
| TJP2 | 1.32 | 3.1E-02 |
| UAP1 | 1.32 | 2.1E-02 |
| UBE2E3 | 1.32 | 4.1E-02 |
| ANKRD50 | 1.32 | 2.6E-02 |
| CDC6 | 1.32 | 1.5E-02 |
| PAPSS2 | 1.32 | 3.0E-02 |
| PXN | 1.33 | 4.5E-02 |
| MEX3C | 1.33 | 4.2E-02 |
| NFKBIB | 1.33 | 1.4E-02 |
| AKAP13 | 1.33 | 4.4E-02 |
| TLE3 | 1.33 | 4.7E-02 |
| PHC2 | 1.33 | 1.4E-02 |
| WDR62 | 1.33 | 2.4E-02 |
| IRS2 | 1.33 | 4.4E-02 |
| DUSP14 | 1.33 | 4.8E-03 |
| SP100 | 1.33 | 1.4E-02 |
| GNS | 1.33 | 1.4E-03 |
| MICU2 | 1.34 | 2.1E-02 |
| ABLIM1 | 1.34 | 2.9E-02 |
| DHX32 | 1.34 | 2.8E-02 |
| TADA3 | 1.34 | 2.0E-02 |
| DESI1 | 1.34 | 2.4E-02 |
| SRCAP | 1.34 | 1.1E-02 |
| CCDC124 | 1.34 | 2.4E-02 |
| EPS8 | 1.34 | 2.4E-02 |
| MTMR9 | 1.34 | 2.4E-02 |
| ZGPAT | 1.34 | 1.4E-02 |
| FAM118B | 1.34 | 4.4E-02 |
| RAB11FIP5 | 1.35 | 4.6E-02 |
| ANLN | 1.35 | 3.1E-02 |
| WWC1 | 1.35 | 3.1E-02 |
| TRIP12 | 1.35 | 1.0E-02 |
| MGME1 | 1.35 | 5.5E-03 |
| CKAP5 | 1.35 | 2.6E-02 |
| CAP1 | 1.35 | 3.3E-02 |
| CDK17 | 1.35 | 3.1E-02 |
| CNDP2 | 1.35 | 3.6E-02 |
| CRMP1 | 1.35 | 2.2E-02 |

| | | |
|---|---|---|
| INTS3 | 1.19 | 1.5E-02 |
| STK11IP | 1.19 | 3.4E-02 |
| USP9X | 1.19 | 2.5E-03 |
| ALG2 | 1.20 | 1.3E-02 |
| IREB2 | 1.20 | 3.7E-02 |
| ATP6V1A | 1.20 | 1.4E-02 |
| ALDH7A1 | 1.20 | 3.9E-02 |
| FAM45A | 1.20 | 1.6E-02 |
| FBXO22 | 1.20 | 3.5E-02 |
| FH | 1.20 | 1.3E-02 |
| RUVBL1 | 1.20 | 2.3E-02 |
| SIRT1 | 1.20 | 1.2E-02 |
| SMARCAL1 | 1.20 | 2.0E-02 |
| TBK1 | 1.20 | 2.8E-02 |
| SKP1 | 1.20 | 4.3E-02 |
| ATP6V0D1 | 1.20 | 3.2E-02 |
| RICTOR | 1.20 | 4.0E-02 |
| MYO1E | 1.20 | 2.3E-02 |
| PPP2R5A | 1.20 | 3.1E-02 |
| CAPNS1 | 1.20 | 2.7E-02 |
| CUEDC2 | 1.20 | 4.9E-02 |
| DGKA | 1.21 | 3.0E-02 |
| TAF6L | 1.21 | 2.8E-02 |
| UGP2 | 1.21 | 1.9E-02 |
| BRAF | 1.21 | 4.2E-02 |
| CRELD2 | 1.21 | 2.6E-02 |
| TOR1B | 1.21 | 4.0E-02 |
| RANBP6 | 1.21 | 4.0E-03 |
| ANXA8 | 1.21 | 3.9E-02 |
| SRGAP1 | 1.21 | 3.4E-02 |
| UBE2T | 1.21 | 6.5E-03 |
| ANP32A | 1.21 | 1.3E-02 |
| MAPRE2 | 1.21 | 3.0E-02 |
| HTT | 1.21 | 1.4E-03 |
| SH3BP5L | 1.21 | 8.9E-03 |
| AP1S3 | 1.21 | 3.4E-02 |
| FBXO4 | 1.21 | 4.9E-02 |
| MORF4L2 | 1.22 | 1.2E-03 |
| ARRDC1 | 1.22 | 4.2E-02 |
| NT5C2 | 1.22 | 2.2E-02 |
| HNRNPA1 | 1.22 | 3.5E-02 |
| RPRD1A | 1.22 | 4.6E-02 |
| LANCL2 | 1.22 | 2.1E-02 |
| OGT | 1.22 | 3.8E-02 |
| SERF2 | 1.22 | 2.8E-02 |
| TP53RK | 1.22 | 2.4E-02 |
| PDZD11 | 1.22 | 4.7E-02 |
| CMBL | 1.22 | 7.6E-03 |
| MEIS1 | 1.22 | 2.6E-02 |
| SPC25 | 1.22 | 2.8E-02 |
| SPPL2A | 1.22 | 3.6E-02 |
| PNP | 1.22 | 2.5E-02 |
| SNX12 | 1.22 | 1.5E-02 |
| APRT | 1.23 | 2.5E-02 |
| HNRNPLL | 1.23 | 3.5E-02 |
| PHF21A | 1.23 | 4.5E-02 |
| HUWE1 | 1.23 | 2.5E-02 |
| MLLT1 | 1.23 | 3.1E-02 |
| ASNA1 | 1.23 | 1.5E-02 |
| HLCS | 1.23 | 4.1E-02 |
| PYGO2 | 1.23 | 4.2E-02 |
| F8A1 | 1.23 | 1.4E-02 |
| LRPAP1 | 1.23 | 1.0E-02 |
| HAX1 | 1.23 | 2.2E-02 |
| SH3GLB2 | 1.23 | 4.1E-02 |
| ANKRD39 | 1.23 | 4.3E-02 |
| POGK | 1.23 | 2.9E-02 |
| ARFGEF2 | 1.23 | 3.9E-02 |

| | | |
|---|---|---|
| CDK3 | 1.19 | 2.47E-03 |
| KIAA0020 | 1.19 | 3.32E-02 |
| LYAR | 1.19 | 2.78E-02 |
| UFL1 | 1.19 | 2.27E-02 |
| SYNCRIP | 1.19 | 1.56E-02 |
| CELF1 | 1.19 | 3.16E-02 |
| SH3BP5L | 1.19 | 7.69E-03 |
| TXNDC5 | 1.19 | 2.55E-02 |
| NAA15 | 1.19 | 4.86E-02 |
| EYA3 | 1.20 | 2.21E-02 |
| NUDT3 | 1.20 | 4.07E-02 |
| OPLAH | 1.20 | 9.41E-03 |
| DCK | 1.20 | 3.25E-02 |
| ARHGEF1 | 1.20 | 3.41E-02 |
| GBE1 | 1.20 | 1.79E-03 |
| FAM91A1 | 1.20 | 2.29E-03 |
| DDX47 | 1.20 | 2.42E-02 |
| PARG | 1.20 | 4.04E-03 |
| MYO9B | 1.20 | 4.68E-02 |
| NELFCD | 1.20 | 2.92E-02 |
| FOLR1 | 1.20 | 2.30E-03 |
| PLCB1 | 1.20 | 4.73E-02 |
| PRKRIR | 1.21 | 1.20E-02 |
| WDR74 | 1.21 | 2.07E-02 |
| RPP14 | 1.21 | 1.50E-02 |
| IMP4 | 1.21 | 4.68E-02 |
| ERH | 1.21 | 2.37E-02 |
| TRAPPC9 | 1.21 | 3.61E-02 |
| DPH1 | 1.21 | 2.27E-02 |
| MIEF1 | 1.22 | 1.82E-02 |
| NDUFAF5 | 1.22 | 3.55E-02 |
| OARD1 | 1.22 | 4.55E-02 |
| AHRR | 1.22 | 3.30E-02 |
| PRPF39 | 1.22 | 4.40E-03 |
| RCL1 | 1.22 | 3.74E-02 |
| LYPLA1 | 1.22 | 3.05E-02 |
| KIF2C | 1.22 | 1.06E-02 |
| RDH14 | 1.22 | 5.40E-03 |
| CTNNBL1 | 1.22 | 3.57E-02 |
| FAM133A | 1.22 | 4.97E-02 |
| DPP9 | 1.23 | 3.97E-02 |
| RPRD1B | 1.23 | 1.35E-02 |
| NIP7 | 1.23 | 1.72E-02 |
| LTA4H | 1.23 | 2.77E-02 |
| AKAP8 | 1.23 | 4.14E-02 |
| SRCAP | 1.23 | 2.21E-02 |
| USP28 | 1.23 | 9.72E-03 |
| BUB1 | 1.23 | 1.24E-02 |
| HTATSF1 | 1.23 | 2.75E-02 |
| PUF60 | 1.24 | 2.12E-02 |
| NOP9 | 1.24 | 2.10E-02 |
| FAM129B | 1.24 | 2.28E-02 |
| WIZ | 1.24 | 3.13E-02 |
| MED16 | 1.24 | 3.77E-02 |
| FCF1 | 1.24 | 1.96E-02 |
| FARSB | 1.24 | 3.89E-02 |
| IPO8 | 1.24 | 3.66E-02 |
| GMEB2 | 1.24 | 5.12E-03 |
| XPO5 | 1.25 | 1.19E-02 |
| INTS4 | 1.25 | 3.15E-02 |
| MED24 | 1.25 | 8.42E-03 |
| MYO1B | 1.25 | 4.51E-02 |
| NOC4L | 1.25 | 4.07E-02 |
| ZNF644 | 1.25 | 8.53E-03 |
| PECR | 1.25 | 9.54E-03 |
| PEPD | 1.25 | 7.15E-03 |
| DUSP12 | 1.25 | 4.17E-02 |
| HPDL | 1.25 | 1.98E-02 |

| | | |
|---|---|---|
| SDC1 | 1.38 | 2.48E-02 |
| GLIPR2 | 1.38 | 2.84E-02 |
| ACTG1 | 1.38 | 3.49E-02 |
| APBB2 | 1.38 | 2.48E-02 |
| GFPT2 | 1.39 | 4.09E-03 |
| ANLN | 1.39 | 2.54E-02 |
| TRIM47 | 1.39 | 1.33E-02 |
| ITGB4 | 1.40 | 2.35E-02 |
| DPYD | 1.40 | 3.80E-03 |
| HMHA1 | 1.41 | 2.25E-03 |
| DCAF6 | 1.41 | 4.23E-02 |
| S100A3 | 1.41 | 2.02E-02 |
| SERPINH1 | 1.41 | 5.60E-03 |
| C11orf68 | 1.41 | 3.04E-03 |
| XYLB | 1.41 | 9.67E-03 |
| CDKN2A | 1.42 | 5.06E-03 |
| ACTN1 | 1.42 | 1.31E-02 |
| PTPRJ | 1.42 | 5.04E-03 |
| ACO1 | 1.42 | 1.65E-02 |
| FANCD2 | 1.42 | 1.10E-02 |
| RGS3 | 1.43 | 1.35E-02 |
| PTRF | 1.43 | 4.18E-03 |
| DNAJC25 | 1.43 | 4.75E-03 |
| DGKA | 1.44 | 1.66E-02 |
| LEPREL1 | 1.45 | 2.03E-02 |
| SUV39H2 | 1.45 | 1.86E-02 |
| SPATA20 | 1.45 | 3.62E-02 |
| SORBS2 | 1.45 | 3.15E-02 |
| IGSF3 | 1.46 | 4.65E-02 |
| EDIL3 | 1.46 | 4.23E-02 |
| NAV1 | 1.46 | 7.35E-03 |
| GNGT1 | 1.46 | 3.65E-02 |
| NFKB2 | 1.46 | 3.35E-02 |
| HTRA1 | 1.46 | 3.64E-02 |
| CCNB2 | 1.47 | 2.15E-02 |
| FLNB | 1.47 | 9.49E-03 |
| PRKCDBP | 1.47 | 3.45E-02 |
| PLXND1 | 1.47 | 2.18E-02 |
| HPD | 1.48 | 4.09E-03 |
| FMNL1 | 1.48 | 2.42E-03 |
| PDP1 | 1.48 | 3.76E-02 |
| ACLY | 1.48 | 5.83E-03 |
| PARVB | 1.49 | 2.61E-03 |
| FN1 | 1.49 | 6.21E-03 |
| LGALS1 | 1.50 | 4.63E-02 |
| PRUNE2 | 1.50 | 2.94E-02 |
| FBLN1 | 1.50 | 1.34E-02 |
| DAAM1 | 1.51 | 2.69E-02 |
| OSBPL6 | 1.51 | 3.09E-03 |
| IGFBP7 | 1.51 | 7.68E-03 |
| RRAS | 1.52 | 2.75E-02 |
| ENTPD5 | 1.53 | 4.62E-02 |
| DYNLT1 | 1.54 | 4.23E-02 |
| SPRYD4 | 1.55 | 4.59E-02 |
| TBC1D2 | 1.55 | 7.72E-03 |
| ACOX3 | 1.56 | 4.63E-02 |
| ASL | 1.57 | 7.22E-03 |
| NEK7 | 1.60 | 2.91E-03 |
| FRMPD1 | 1.60 | 1.76E-02 |
| ALDH1B1 | 1.61 | 2.87E-02 |
| EPB41L1 | 1.62 | 7.83E-03 |
| APOB | 1.62 | 1.15E-02 |
| FLNC | 1.62 | 7.01E-03 |
| NGEF | 1.63 | 4.93E-03 |
| ARHGEF28 | 1.63 | 3.53E-02 |
| GLS | 1.64 | 4.85E-02 |
| FMNL2 | 1.65 | 9.41E-03 |
| HDGFRP3 | 1.66 | 3.07E-03 |

| | | |
|---|---|---|
| CCDC59 | 1.35 | 4.7E-02 |
| C15orf39 | 1.35 | 1.7E-02 |
| PXK | 1.35 | 8.3E-04 |
| ERICH1 | 1.35 | 4.0E-02 |
| TGFB1I1 | 1.35 | 3.2E-02 |
| PREP | 1.35 | 4.5E-02 |
| CCDC134 | 1.36 | 1.5E-02 |
| RASAL2 | 1.36 | 1.3E-02 |
| NEXN | 1.36 | 1.7E-02 |
| HOOK2 | 1.36 | 4.9E-03 |
| STARD13 | 1.36 | 3.6E-02 |
| ZNF217 | 1.36 | 6.1E-03 |
| TRIM32 | 1.36 | 1.5E-02 |
| SHB | 1.36 | 2.2E-02 |
| YAP1 | 1.36 | 3.3E-02 |
| PEA15 | 1.36 | 1.0E-02 |
| HMGCS1 | 1.37 | 1.0E-02 |
| ABHD5 | 1.37 | 2.7E-02 |
| TK1 | 1.37 | 1.5E-02 |
| LGALSL | 1.37 | 8.6E-03 |
| GDA | 1.37 | 1.7E-02 |
| GSTT2 | 1.37 | 3.9E-02 |
| L3MBTL2 | 1.37 | 2.3E-02 |
| MYH14 | 1.37 | 3.8E-03 |
| GGA1 | 1.38 | 6.9E-03 |
| RHEB | 1.38 | 1.1E-02 |
| CCDC115 | 1.38 | 1.5E-03 |
| RBMS2 | 1.38 | 2.6E-02 |
| HYI | 1.38 | 4.6E-02 |
| ARHGAP1 | 1.38 | 3.5E-02 |
| IGF2BP1 | 1.38 | 5.2E-03 |
| MRPL12 | 1.38 | 4.2E-02 |
| CD3EAP | 1.38 | 2.6E-02 |
| HK2 | 1.38 | 8.6E-03 |
| SEMA3C | 1.38 | 2.2E-02 |
| MAP2K3 | 1.39 | 3.2E-02 |
| CORO1B | 1.39 | 7.8E-03 |
| SH3BGRL3 | 1.39 | 2.2E-02 |
| NF1 | 1.39 | 3.7E-02 |
| PLEKHO2 | 1.39 | 3.7E-02 |
| TRIM3 | 1.39 | 3.7E-02 |
| NEDD4L | 1.39 | 8.3E-03 |
| CAMK4 | 1.39 | 4.8E-02 |
| SNAP23 | 1.39 | 4.6E-02 |
| REXO2 | 1.39 | 3.5E-02 |
| CMSS1 | 1.40 | 2.1E-02 |
| FRMPD1 | 1.40 | 4.5E-02 |
| WASF1 | 1.40 | 8.0E-04 |
| ACLY | 1.40 | 4.0E-02 |
| DNPEP | 1.40 | 8.8E-03 |
| BOLA2 | 1.40 | 4.5E-02 |
| AGTPBP1 | 1.40 | 1.0E-03 |
| LMNB2 | 1.40 | 2.6E-02 |
| VAT1L | 1.41 | 2.6E-02 |
| DBN1 | 1.41 | 3.4E-02 |
| PDLIM5 | 1.41 | 3.2E-02 |
| ERCC6L | 1.41 | 1.2E-02 |
| NFKB2 | 1.41 | 1.8E-02 |
| GTSE1 | 1.41 | 1.1E-02 |
| CDC37 | 1.41 | 4.2E-02 |
| TANC2 | 1.41 | 1.0E-02 |
| ADO | 1.41 | 7.3E-03 |
| ZBTB7A | 1.41 | 3.5E-02 |
| SCRN1 | 1.41 | 3.6E-02 |
| TFPT | 1.41 | 4.6E-02 |
| EPB41L1 | 1.42 | 4.4E-02 |
| PTPN12 | 1.42 | 1.1E-02 |
| PDLIM3 | 1.42 | 2.4E-02 |

| | | |
|---|---|---|
| NSMCE4A | 1.23 | 4.5E-02 |
| NDUFS5 | 1.23 | 2.2E-02 |
| UFL1 | 1.23 | 5.5E-03 |
| NUTF2 | 1.23 | 4.5E-02 |
| XPOT | 1.23 | 1.5E-02 |
| RPS6KA4 | 1.23 | 4.6E-02 |
| ABI1 | 1.24 | 2.4E-02 |
| PITPNC1 | 1.24 | 2.7E-02 |
| PRCC | 1.24 | 1.5E-02 |
| SCPEP1 | 1.24 | 4.6E-02 |
| SDCBP | 1.24 | 1.6E-02 |
| ERP29 | 1.24 | 9.0E-03 |
| STAT6 | 1.24 | 3.2E-02 |
| LRBA | 1.24 | 1.5E-03 |
| EPRS | 1.24 | 7.1E-03 |
| HOOK2 | 1.24 | 1.1E-02 |
| INF2 | 1.24 | 1.2E-03 |
| VPS26A | 1.24 | 9.7E-03 |
| TSNAX | 1.24 | 1.9E-02 |
| ACY1 | 1.24 | 9.1E-03 |
| HTATSF1 | 1.24 | 3.2E-02 |
| MOCS2 | 1.24 | 3.8E-02 |
| NIT2 | 1.24 | 1.5E-02 |
| OSGEP | 1.24 | 3.9E-02 |
| EPPK1 | 1.24 | 3.2E-02 |
| PDE4D | 1.24 | 2.7E-02 |
| CTSL | 1.24 | 4.4E-03 |
| VBP1 | 1.24 | 2.2E-02 |
| CHML | 1.25 | 2.5E-02 |
| PDIA6 | 1.25 | 1.3E-02 |
| GNS | 1.25 | 1.9E-02 |
| IKBKB | 1.25 | 6.8E-03 |
| PTGR1 | 1.25 | 1.1E-02 |
| TSC22D3 | 1.25 | 2.7E-02 |
| SRCAP | 1.25 | 2.1E-02 |
| SPR | 1.25 | 9.2E-03 |
| TBC1D17 | 1.25 | 1.2E-02 |
| FOSL2 | 1.25 | 1.1E-02 |
| AP3B1 | 1.25 | 2.6E-02 |
| WDR13 | 1.25 | 5.0E-02 |
| SIPA1L1 | 1.25 | 3.7E-02 |
| DLG5 | 1.25 | 1.4E-03 |
| UBE2Q1 | 1.25 | 2.1E-02 |
| FBXO3 | 1.25 | 2.5E-03 |
| VPS11 | 1.25 | 2.4E-02 |
| NOL3 | 1.25 | 3.2E-02 |
| DUSP12 | 1.25 | 9.1E-03 |
| METTL13 | 1.26 | 4.2E-02 |
| RALGAPA1 | 1.26 | 6.5E-03 |
| FDPS | 1.26 | 3.6E-02 |
| CAPN2 | 1.26 | 1.0E-02 |
| NEU1 | 1.26 | 2.5E-02 |
| MANF | 1.26 | 3.3E-03 |
| SDE2 | 1.26 | 3.9E-02 |
| MSLN | 1.26 | 4.0E-02 |
| CAND1 | 1.26 | 1.5E-02 |
| FANCA | 1.26 | 5.1E-03 |
| ANXA3 | 1.26 | 9.1E-03 |
| C6orf211 | 1.26 | 1.3E-02 |
| SEC24B | 1.27 | 4.3E-02 |
| SMYD3 | 1.27 | 4.4E-02 |
| VPS39 | 1.27 | 2.3E-02 |
| NGEF | 1.27 | 4.4E-02 |
| S100A10 | 1.27 | 3.8E-02 |
| NFKB2 | 1.27 | 1.2E-03 |
| ESD | 1.27 | 3.9E-02 |
| TRNT1 | 1.27 | 1.6E-02 |
| PPT2 | 1.27 | 1.0E-02 |

| | | |
|---|---|---|
| INTS10 | 1.25 | 3.35E-02 |
| OXSM | 1.25 | 4.16E-02 |
| ABHD5 | 1.26 | 1.20E-02 |
| USP7 | 1.26 | 8.37E-03 |
| CASP9 | 1.26 | 4.35E-02 |
| TWISTNB | 1.26 | 2.26E-02 |
| CHMP7 | 1.26 | 4.23E-02 |
| EMSY | 1.26 | 2.48E-02 |
| SOGA1 | 1.26 | 3.01E-03 |
| NGDN | 1.26 | 3.23E-02 |
| RPS6KA4 | 1.26 | 8.62E-03 |
| CARS | 1.26 | 1.68E-02 |
| TOP2A | 1.27 | 4.07E-02 |
| HUWE1 | 1.27 | 2.69E-02 |
| SECISBP2L | 1.27 | 2.94E-03 |
| IRF2BPL | 1.27 | 3.38E-02 |
| POLR1B | 1.27 | 1.42E-02 |
| RICTOR | 1.27 | 3.08E-02 |
| PDCD11 | 1.27 | 1.05E-02 |
| EGLN1 | 1.27 | 1.32E-02 |
| DDX17 | 1.27 | 3.98E-02 |
| BLM | 1.27 | 3.25E-02 |
| NASP | 1.27 | 3.34E-02 |
| ZFP91 | 1.27 | 3.15E-02 |
| RYBP | 1.28 | 3.40E-02 |
| TBC1D9B | 1.28 | 1.31E-02 |
| DDX56 | 1.28 | 4.54E-02 |
| CCAR2 | 1.28 | 2.81E-02 |
| MINA | 1.28 | 2.08E-02 |
| RPP25 | 1.28 | 4.63E-04 |
| NSMCE4A | 1.28 | 4.24E-02 |
| PITPNM1 | 1.28 | 5.00E-03 |
| UTP14A | 1.28 | 4.61E-02 |
| DEK | 1.29 | 3.36E-02 |
| PGM2L1 | 1.29 | 1.50E-02 |
| SMARCA4 | 1.29 | 3.62E-02 |
| DDX21 | 1.29 | 1.18E-02 |
| MTRR | 1.29 | 1.29E-02 |
| ATPAF1 | 1.29 | 3.76E-02 |
| RASA1 | 1.30 | 4.55E-03 |
| ADO | 1.30 | 1.26E-02 |
| PDIA3 | 1.30 | 1.21E-03 |
| DDX24 | 1.30 | 8.62E-03 |
| MLKL | 1.30 | 4.29E-02 |
| CTU1 | 1.30 | 2.76E-02 |
| DHX32 | 1.30 | 3.90E-02 |
| POLR1A | 1.31 | 9.17E-03 |
| TCERG1 | 1.31 | 3.14E-02 |
| GNL3L | 1.31 | 2.13E-02 |
| RIF1 | 1.31 | 1.18E-02 |
| CPOX | 1.31 | 1.11E-02 |
| PLIN3 | 1.31 | 4.09E-02 |
| MCMBP | 1.31 | 2.05E-02 |
| TSHZ2 | 1.31 | 2.20E-02 |
| ZNF451 | 1.31 | 3.07E-02 |
| GPI | 1.31 | 3.22E-02 |
| ANXA10 | 1.31 | 4.97E-03 |
| SNX9 | 1.31 | 3.87E-02 |
| HEXB | 1.31 | 3.96E-02 |
| TTN | 1.31 | 4.24E-02 |
| LMCD1 | 1.32 | 4.82E-02 |
| UHRF1 | 1.32 | 1.58E-02 |
| MRE11A | 1.32 | 4.78E-02 |
| UTP15 | 1.32 | 3.24E-02 |
| ANGEL2 | 1.32 | 3.36E-02 |
| MAFG | 1.32 | 3.87E-02 |
| UBE2S | 1.32 | 1.25E-02 |
| PLEK2 | 1.32 | 4.57E-02 |

| | | |
|---|---|---|
| COL1A1 | 1.68 | 1.67E-02 |
| SLC38A1 | 1.68 | 4.56E-02 |
| SERPINE1 | 1.71 | 1.22E-02 |
| TNC | 1.71 | 1.30E-02 |
| TGFBI | 1.71 | 1.86E-02 |
| ITGB1 | 1.72 | 2.89E-02 |
| DGUOK | 1.74 | 7.88E-03 |
| PYCR1 | 1.75 | 4.97E-03 |
| C15orf52 | 1.76 | 1.44E-02 |
| FGF2 | 1.76 | 2.65E-02 |
| PSPH | 1.76 | 2.30E-03 |
| COL5A2 | 1.77 | 3.75E-02 |
| TMEM160 | 1.77 | 4.51E-02 |
| PTPRF | 1.77 | 5.26E-03 |
| DDB2 | 1.78 | 3.92E-03 |
| ASS1 | 1.81 | 2.76E-03 |
| SH3KBP1 | 1.82 | 1.48E-02 |
| SLC2A1 | 1.83 | 2.03E-02 |
| XPR1 | 1.87 | 4.89E-02 |
| GDA | 1.87 | 5.35E-03 |
| FOSL1 | 1.88 | 2.69E-02 |
| HYI | 1.90 | 1.34E-03 |
| PRSS23 | 1.91 | 1.55E-02 |
| CSPG4 | 1.95 | 3.03E-03 |
| LAMB1 | 1.96 | 2.06E-02 |
| ASNS | 1.96 | 2.33E-03 |
| RIN1 | 1.97 | 7.27E-03 |
| STC2 | 1.99 | 1.52E-03 |
| ITGA11 | 2.09 | 2.22E-02 |
| FAS | 2.11 | 2.53E-02 |
| CDCP1 | 2.13 | 9.07E-03 |
| TGM2 | 2.14 | 1.23E-03 |
| PCK2 | 2.16 | 3.21E-03 |
| DPYSL3 | 2.19 | 3.61E-03 |
| ITGA5 | 2.24 | 4.74E-03 |
| DMBT1 | 2.25 | 2.05E-02 |
| PTGS1 | 2.26 | 1.65E-02 |
| SLC38A2 | 2.28 | 1.45E-02 |
| BCAT1 | 2.31 | 6.47E-05 |
| GRAMD1A | 2.31 | 2.34E-03 |
| NES | 2.33 | 3.06E-03 |
| THBS1 | 2.40 | 5.13E-03 |
| SERPINE2 | 2.61 | 1.57E-03 |
| CD70 | 2.63 | 1.18E-02 |
| FAM32A | 2.64 | 1.80E-02 |
| THY1 | 2.65 | 1.99E-02 |
| CRABP2 | 2.65 | 5.15E-03 |
| EPHA2 | 2.70 | 5.47E-03 |
| F3 | 3.13 | 5.89E-03 |
| EPHA7 | 3.25 | 5.38E-03 |
| ANPEP | 5.02 | 7.08E-05 |

| | | |
|---|---|---|
| SNX9 | 1.42 | 1.1E-02 |
| ACO1 | 1.42 | 2.8E-02 |
| UXT | 1.42 | 2.0E-02 |
| PACS1 | 1.42 | 3.1E-02 |
| CNN2 | 1.43 | 2.3E-02 |
| ILK | 1.43 | 1.4E-02 |
| HDAC7 | 1.43 | 3.2E-02 |
| PCDH7 | 1.43 | 5.9E-03 |
| NCKAP5L | 1.43 | 3.2E-03 |
| ARFGAP1 | 1.44 | 2.8E-02 |
| ECHDC1 | 1.44 | 9.4E-03 |
| MSN | 1.44 | 4.2E-02 |
| IRF2BPL | 1.44 | 1.2E-02 |
| TBC1D23 | 1.44 | 1.1E-02 |
| CLPB | 1.44 | 7.0E-03 |
| PLCB4 | 1.44 | 2.4E-02 |
| ARPC3 | 1.44 | 1.5E-02 |
| HABP4 | 1.44 | 2.4E-02 |
| TJP1 | 1.44 | 2.5E-02 |
| RASSF8 | 1.44 | 1.1E-02 |
| NME7 | 1.44 | 3.9E-02 |
| DDAH2 | 1.44 | 2.8E-02 |
| LGMN | 1.45 | 2.9E-02 |
| RND3 | 1.45 | 2.2E-03 |
| CTNND1 | 1.45 | 9.0E-03 |
| UFM1 | 1.45 | 1.2E-02 |
| FSCN1 | 1.46 | 5.0E-02 |
| MVP | 1.46 | 1.2E-02 |
| CASP4 | 1.46 | 4.5E-02 |
| HMGA1 | 1.46 | 1.0E-03 |
| FAM103A1 | 1.47 | 2.5E-02 |
| RASA1 | 1.47 | 7.7E-03 |
| TES | 1.47 | 1.7E-02 |
| UBE2F | 1.47 | 1.9E-02 |
| PARD3 | 1.47 | 1.7E-02 |
| C7orf55 | 1.47 | 2.4E-02 |
| ABTB2 | 1.47 | 3.8E-03 |
| PLA2G4B | 1.47 | 1.7E-03 |
| PITPNC1 | 1.48 | 3.6E-02 |
| PTRF | 1.48 | 6.7E-03 |
| TTPAL | 1.48 | 2.1E-03 |
| ARHGAP18 | 1.48 | 2.1E-02 |
| PALM2 | 1.48 | 6.7E-03 |
| DLGAP4 | 1.49 | 1.8E-02 |
| XRCC4 | 1.49 | 3.0E-02 |
| ZEB1 | 1.49 | 2.3E-02 |
| CDKN2AIPN | 1.49 | 5.3E-03 |
| SPR | 1.49 | 3.7E-02 |
| LASP1 | 1.50 | 2.3E-02 |
| TMEM168 | 1.50 | 3.3E-02 |
| FHL3 | 1.50 | 1.5E-02 |
| ING5 | 1.50 | 4.3E-02 |
| PHLDB1 | 1.50 | 2.6E-02 |
| MED13L | 1.50 | 1.9E-02 |
| IGF2BP2 | 1.50 | 1.0E-02 |
| WIPF2 | 1.51 | 4.3E-02 |
| WDFY3 | 1.51 | 2.6E-02 |
| LYN | 1.51 | 4.8E-02 |
| JUP | 1.51 | 4.7E-03 |
| MAFF | 1.51 | 4.5E-02 |
| RRBP1 | 1.51 | 2.8E-03 |
| DHPS | 1.52 | 9.2E-03 |
| CHMP6 | 1.52 | 4.6E-02 |
| ACOT2 | 1.52 | 2.5E-02 |
| KCTD12 | 1.52 | 3.1E-02 |
| ITGA5 | 1.53 | 4.6E-02 |
| DDX58 | 1.53 | 3.3E-02 |
| URI1 | 1.53 | 2.6E-02 |

| | | |
|---|---|---|
| TIMM13 | 1.27 | 5.0E-03 |
| VPS36 | 1.27 | 4.4E-03 |
| MED12 | 1.27 | 3.5E-02 |
| PPA2 | 1.27 | 3.9E-02 |
| ENO3 | 1.27 | 4.2E-02 |
| C16orf62 | 1.27 | 3.1E-02 |
| C5orf51 | 1.27 | 1.2E-02 |
| USP11 | 1.27 | 3.3E-02 |
| GNPDA2 | 1.27 | 2.6E-02 |
| PSEN1 | 1.27 | 4.8E-02 |
| VPS16 | 1.27 | 4.8E-03 |
| BCL7A | 1.28 | 7.0E-03 |
| SUPT7L | 1.28 | 1.8E-02 |
| TIPRL | 1.28 | 1.6E-02 |
| AIDA | 1.28 | 3.4E-03 |
| FAM175B | 1.28 | 1.1E-02 |
| RBBP7 | 1.28 | 2.3E-02 |
| NAV1 | 1.28 | 2.0E-02 |
| MTMR6 | 1.28 | 2.1E-02 |
| SLFN5 | 1.28 | 9.6E-03 |
| CXXC1 | 1.28 | 2.1E-02 |
| CYFIP2 | 1.28 | 5.0E-03 |
| MTMR12 | 1.28 | 4.0E-02 |
| WDFY3 | 1.28 | 4.7E-02 |
| LANCL1 | 1.28 | 1.2E-02 |
| CTSB | 1.29 | 3.7E-02 |
| MMADHC | 1.29 | 2.3E-02 |
| ANXA4 | 1.29 | 2.5E-02 |
| PBX1 | 1.29 | 1.8E-02 |
| ZNF318 | 1.29 | 1.2E-02 |
| MAP2K6 | 1.29 | 2.6E-02 |
| KLHL42 | 1.29 | 5.4E-03 |
| STXBP2 | 1.29 | 1.9E-02 |
| MEPCE | 1.29 | 1.6E-02 |
| EPB41L1 | 1.29 | 2.4E-02 |
| LIMCH1 | 1.29 | 4.6E-02 |
| AHNAK2 | 1.30 | 3.3E-02 |
| IL18 | 1.30 | 1.4E-02 |
| NR4A1 | 1.30 | 4.5E-03 |
| HMG20B | 1.30 | 2.1E-02 |
| ATXN7 | 1.30 | 4.3E-02 |
| TJP2 | 1.30 | 7.5E-03 |
| PIN4 | 1.30 | 3.7E-02 |
| GSTK1 | 1.30 | 3.5E-03 |
| CENPF | 1.30 | 3.5E-02 |
| TBC1D9B | 1.30 | 7.2E-03 |
| GBA | 1.30 | 2.1E-04 |
| UTRN | 1.30 | 1.7E-02 |
| GMPR2 | 1.30 | 1.4E-02 |
| DOCK4 | 1.30 | 2.0E-02 |
| CARHSP1 | 1.31 | 2.1E-02 |
| COMMD7 | 1.31 | 4.3E-02 |
| WDR45 | 1.31 | 2.4E-02 |
| CSTB | 1.31 | 6.3E-04 |
| GOLGA1 | 1.31 | 2.8E-02 |
| ZMYM3 | 1.31 | 3.2E-02 |
| FAM120C | 1.31 | 1.3E-02 |
| C19orf66 | 1.31 | 2.6E-02 |
| MMAB | 1.31 | 1.4E-03 |
| ISOC1 | 1.31 | 3.9E-02 |
| KDELC1 | 1.31 | 2.7E-02 |
| GDI1 | 1.31 | 4.9E-04 |
| STC2 | 1.32 | 2.4E-02 |
| PRDX5 | 1.32 | 7.3E-03 |
| PPCS | 1.32 | 1.2E-02 |
| ADAR | 1.32 | 1.4E-02 |
| APEX1 | 1.32 | 3.9E-02 |
| SPG21 | 1.32 | 1.5E-02 |

| | | |
|---|---|---|
| DIAPH2 | 1.32 | 4.90E-02 |
| ESPL1 | 1.33 | 3.21E-02 |
| CHID1 | 1.33 | 1.37E-02 |
| NIPBL | 1.33 | 3.57E-02 |
| ASMTL | 1.33 | 3.23E-02 |
| ZFC3H1 | 1.33 | 3.23E-02 |
| AKT1 | 1.34 | 4.55E-02 |
| RRS1 | 1.34 | 3.47E-03 |
| TBC1D4 | 1.34 | 3.08E-02 |
| POLD1 | 1.34 | 2.11E-02 |
| CCDC25 | 1.34 | 2.31E-02 |
| SYPL1 | 1.34 | 3.47E-02 |
| IMP3 | 1.34 | 4.61E-03 |
| DDX54 | 1.34 | 5.91E-03 |
| POLR1E | 1.34 | 1.07E-02 |
| CCNB2 | 1.35 | 4.22E-02 |
| LTV1 | 1.35 | 1.53E-02 |
| HNRNPUL1 | 1.35 | 3.33E-03 |
| FBL | 1.35 | 1.33E-02 |
| SIPA1L1 | 1.35 | 3.29E-02 |
| SPRYD3 | 1.35 | 2.61E-02 |
| CDK6 | 1.35 | 1.55E-02 |
| CEP41 | 1.35 | 4.98E-02 |
| SERPINB1 | 1.35 | 2.70E-02 |
| SMARCC1 | 1.35 | 3.27E-02 |
| CENPB | 1.35 | 1.72E-02 |
| RAD50 | 1.36 | 2.41E-02 |
| HLTF | 1.36 | 2.19E-02 |
| BRD4 | 1.36 | 3.89E-02 |
| DDX41 | 1.36 | 4.08E-02 |
| C15orf39 | 1.37 | 5.64E-03 |
| DIMT1 | 1.37 | 3.07E-02 |
| UBA2 | 1.37 | 8.02E-03 |
| S100A10 | 1.37 | 3.66E-02 |
| NTHL1 | 1.37 | 2.07E-02 |
| TGM2 | 1.37 | 1.88E-02 |
| LRPAP1 | 1.37 | 3.47E-03 |
| BRD3 | 1.38 | 2.52E-02 |
| ADARB1 | 1.38 | 1.95E-02 |
| ZMYND11 | 1.38 | 4.55E-02 |
| MBLAC2 | 1.38 | 1.01E-02 |
| DDX39A | 1.38 | 3.20E-02 |
| YOD1 | 1.38 | 5.17E-03 |
| ADA | 1.38 | 4.70E-02 |
| GPX1 | 1.38 | 1.82E-02 |
| PHF2 | 1.38 | 4.28E-02 |
| CMSS1 | 1.38 | 4.04E-02 |
| SIL1 | 1.38 | 4.64E-02 |
| DKC1 | 1.39 | 6.34E-03 |
| SRM | 1.39 | 1.90E-02 |
| LSM6 | 1.39 | 2.05E-02 |
| RAB4A | 1.39 | 1.88E-02 |
| MED6 | 1.39 | 1.05E-02 |
| ZMIZ1 | 1.39 | 4.82E-02 |
| UTP23 | 1.40 | 6.97E-03 |
| PDK1 | 1.40 | 4.34E-03 |
| HEATR1 | 1.40 | 1.10E-02 |
| VRK1 | 1.40 | 2.66E-02 |
| NPM3 | 1.40 | 2.90E-02 |
| RSBN1 | 1.40 | 2.40E-02 |
| UBE2M | 1.40 | 1.56E-02 |
| TOP1 | 1.40 | 3.04E-02 |
| CRTAP | 1.41 | 2.11E-02 |
| CLK1 | 1.41 | 3.96E-02 |
| GEMIN5 | 1.41 | 4.38E-02 |
| CHD1 | 1.41 | 3.65E-02 |
| CASK | 1.41 | 4.64E-02 |
| GLUD1 | 1.41 | 3.37E-02 |

| | | |
|---|---|---|
| PLEK2 | 1.53 | 3.4E-02 |
| MICALL1 | 1.53 | 1.4E-03 |
| DIP2B | 1.53 | 3.8E-03 |
| UGP2 | 1.53 | 3.0E-03 |
| SIPA1 | 1.54 | 1.0E-02 |
| AFAP1 | 1.54 | 6.6E-03 |
| NFIX | 1.54 | 1.1E-03 |
| ADAT2 | 1.55 | 1.3E-02 |
| TTC38 | 1.55 | 3.5E-02 |
| VPS37B | 1.55 | 3.7E-02 |
| TTLL12 | 1.55 | 2.1E-02 |
| CHP1 | 1.55 | 1.1E-02 |
| OSBPL1A | 1.55 | 3.6E-02 |
| PFN2 | 1.55 | 2.8E-02 |
| FHL1 | 1.55 | 1.8E-02 |
| SRSF9 | 1.55 | 1.0E-02 |
| TIMP2 | 1.56 | 4.4E-03 |
| FGF2 | 1.56 | 4.0E-02 |
| BICC1 | 1.56 | 1.6E-02 |
| PPFIBP1 | 1.57 | 4.9E-02 |
| SH2B1 | 1.57 | 2.8E-02 |
| PIK3R1 | 1.58 | 2.5E-03 |
| L3MBTL3 | 1.58 | 2.7E-02 |
| ALKBH8 | 1.58 | 4.5E-02 |
| PHLDB2 | 1.58 | 1.2E-02 |
| MEF2A | 1.59 | 3.1E-02 |
| RPS5 | 1.59 | 3.2E-02 |
| SH3BP5L | 1.59 | 1.2E-02 |
| RCN3 | 1.59 | 1.2E-02 |
| IREB2 | 1.59 | 1.0E-02 |
| OSBPL3 | 1.59 | 2.6E-03 |
| DYNLT1 | 1.60 | 3.8E-02 |
| RAB11A | 1.60 | 2.3E-03 |
| ARPC1A | 1.60 | 1.8E-02 |
| CHCHD2 | 1.60 | 2.7E-02 |
| ZYX | 1.60 | 3.6E-02 |
| AKAP2 | 1.60 | 1.4E-02 |
| TBC1D4 | 1.60 | 1.1E-02 |
| RSU1 | 1.61 | 8.4E-03 |
| DUSP23 | 1.61 | 3.0E-03 |
| NEGR1 | 1.62 | 3.5E-02 |
| RAB3IL1 | 1.62 | 2.5E-03 |
| RGS10 | 1.62 | 2.1E-02 |
| ZSWIM8 | 1.63 | 2.3E-02 |
| FMNL2 | 1.63 | 2.3E-02 |
| ADD1 | 1.63 | 3.8E-02 |
| AKR1B1 | 1.63 | 7.3E-03 |
| SAR1A | 1.63 | 2.4E-02 |
| MYH9 | 1.64 | 2.5E-02 |
| MVB12A | 1.64 | 2.0E-02 |
| CD2AP | 1.64 | 2.7E-02 |
| ADD3 | 1.64 | 3.9E-03 |
| SELO | 1.64 | 3.2E-02 |
| PRKCDBP | 1.64 | 1.8E-02 |
| TXNDC5 | 1.64 | 5.1E-03 |
| RANGAP1 | 1.64 | 7.5E-03 |
| DTD1 | 1.65 | 1.8E-02 |
| PARVB | 1.65 | 4.7E-04 |
| FAM107B | 1.65 | 3.2E-02 |
| MARCKS | 1.65 | 8.9E-03 |
| LMCD1 | 1.66 | 1.1E-02 |
| PRR11 | 1.67 | 1.6E-02 |
| ACTN1 | 1.67 | 3.2E-02 |
| HACL1 | 1.68 | 1.4E-03 |
| APBB2 | 1.69 | 1.0E-02 |
| DOCK10 | 1.69 | 4.3E-04 |
| MYH11 | 1.70 | 2.4E-02 |
| BAP18 | 1.70 | 2.2E-04 |

| | | |
|---|---|---|
| FAM120A | 1.32 | 2.0E-02 |
| GGPS1 | 1.32 | 2.4E-02 |
| LCMT2 | 1.32 | 1.4E-02 |
| KIAA0947 | 1.32 | 3.7E-02 |
| NUCKS1 | 1.32 | 1.8E-02 |
| RPS6KA6 | 1.32 | 2.6E-02 |
| SMS | 1.32 | 1.2E-02 |
| IQGAP1 | 1.32 | 1.2E-02 |
| RABL3 | 1.32 | 2.8E-02 |
| LONP2 | 1.33 | 3.2E-02 |
| PFKFB2 | 1.33 | 2.2E-02 |
| ULK3 | 1.33 | 4.7E-02 |
| DTL | 1.33 | 4.4E-02 |
| GBE1 | 1.33 | 2.4E-03 |
| SNCG | 1.33 | 7.8E-03 |
| PYGB | 1.33 | 3.7E-03 |
| PDCD6 | 1.33 | 6.0E-03 |
| SERPINH1 | 1.33 | 1.3E-02 |
| GLRX | 1.33 | 2.1E-02 |
| ATE1 | 1.33 | 2.9E-02 |
| TUT1 | 1.33 | 1.5E-02 |
| ITGB4 | 1.33 | 3.5E-02 |
| AIP | 1.33 | 1.4E-02 |
| HDGF | 1.33 | 2.1E-02 |
| ABLIM1 | 1.34 | 3.2E-02 |
| GGH | 1.34 | 1.7E-02 |
| DNMBP | 1.34 | 4.0E-04 |
| TIGAR | 1.34 | 3.2E-02 |
| ENSA | 1.34 | 3.6E-02 |
| ANKRD50 | 1.34 | 7.3E-03 |
| MYD88 | 1.34 | 3.0E-02 |
| CNPY2 | 1.34 | 3.2E-03 |
| RIN1 | 1.34 | 2.4E-02 |
| FGD4 | 1.34 | 2.2E-02 |
| PRPS2 | 1.34 | 9.1E-03 |
| SORBS2 | 1.34 | 4.9E-02 |
| ARPIN | 1.34 | 4.5E-02 |
| ARHGEF2 | 1.35 | 2.5E-02 |
| ALDH5A1 | 1.35 | 3.4E-02 |
| STAT2 | 1.35 | 2.2E-02 |
| KDM5C | 1.35 | 5.5E-03 |
| OPTN | 1.35 | 1.1E-02 |
| BLOC1S1 | 1.35 | 1.4E-02 |
| SYTL4 | 1.35 | 3.3E-03 |
| ARAF | 1.35 | 1.6E-02 |
| CAMSAP2 | 1.35 | 2.9E-02 |
| TRAPPC2L | 1.35 | 1.2E-02 |
| TIMM9 | 1.35 | 2.1E-02 |
| PCBD1 | 1.35 | 7.9E-03 |
| C19orf10 | 1.35 | 1.6E-02 |
| PPIB | 1.35 | 5.4E-03 |
| DHX9 | 1.35 | 3.4E-02 |
| MYO10 | 1.36 | 1.3E-02 |
| CTSZ | 1.36 | 1.3E-02 |
| R3HCC1 | 1.36 | 3.5E-03 |
| NDUFA7 | 1.36 | 1.3E-02 |
| TRIM25 | 1.36 | 4.1E-03 |
| GRIPAP1 | 1.37 | 4.2E-02 |
| MTHFS | 1.37 | 2.2E-03 |
| GPX1 | 1.37 | 4.3E-03 |
| P4HTM | 1.37 | 2.6E-02 |
| LAP3 | 1.37 | 1.0E-03 |
| ACBD6 | 1.37 | 2.1E-02 |
| SRGAP2 | 1.37 | 1.9E-02 |
| RPRD2 | 1.37 | 4.4E-02 |
| TTC39C | 1.37 | 1.4E-02 |
| DDX41 | 1.37 | 1.7E-02 |
| VTN | 1.37 | 2.8E-03 |

| | | |
|---|---|---|
| XPO7 | 1.41 | 1.49E-02 |
| RDH13 | 1.42 | 3.89E-02 |
| ARAF | 1.43 | 2.31E-04 |
| NDUFB1 | 1.43 | 1.47E-03 |
| NBN | 1.43 | 3.56E-02 |
| DHX9 | 1.43 | 4.20E-02 |
| TRIM24 | 1.43 | 2.65E-02 |
| GPRIN1 | 1.43 | 3.12E-02 |
| PAIP1 | 1.44 | 3.61E-02 |
| BRD9 | 1.44 | 4.76E-02 |
| PHF6 | 1.44 | 3.94E-02 |
| MED8 | 1.44 | 3.57E-02 |
| ACOT13 | 1.44 | 2.52E-02 |
| PRPF38B | 1.44 | 6.08E-03 |
| PREP | 1.45 | 1.12E-03 |
| KMT2B | 1.45 | 1.03E-02 |
| BZW1 | 1.45 | 7.23E-03 |
| TARBP1 | 1.45 | 1.07E-02 |
| BCAR3 | 1.45 | 3.04E-02 |
| CNOT7 | 1.46 | 2.95E-02 |
| CLIP2 | 1.46 | 1.09E-03 |
| MYO10 | 1.46 | 2.28E-02 |
| PLEKHH2 | 1.46 | 9.27E-03 |
| ZNF217 | 1.47 | 1.66E-02 |
| ARHGAP12 | 1.47 | 1.01E-03 |
| TFAP4 | 1.47 | 3.75E-02 |
| LYN | 1.47 | 3.98E-02 |
| NR4A1 | 1.47 | 2.16E-03 |
| CD97 | 1.48 | 3.57E-02 |
| SSFA2 | 1.48 | 3.38E-02 |
| NOSIP | 1.48 | 4.19E-02 |
| PDS5A | 1.48 | 4.30E-03 |
| ZNF740 | 1.49 | 8.71E-03 |
| OSGEPL1 | 1.49 | 1.42E-02 |
| PUS7 | 1.50 | 8.80E-04 |
| TCOF1 | 1.50 | 2.95E-02 |
| GNB1L | 1.50 | 1.27E-02 |
| TRIM28 | 1.50 | 3.83E-02 |
| PXK | 1.51 | 3.43E-02 |
| NDUFA7 | 1.51 | 5.33E-03 |
| NDUFS4 | 1.52 | 2.08E-02 |
| KDM5C | 1.52 | 2.48E-02 |
| JUNB | 1.53 | 3.89E-02 |
| DHPS | 1.53 | 1.09E-02 |
| ECE2 | 1.53 | 3.89E-02 |
| MID1 | 1.54 | 7.16E-03 |
| UBE2D4 | 1.54 | 3.09E-02 |
| ALDH1L2 | 1.54 | 2.07E-02 |
| ZNF706 | 1.54 | 4.79E-02 |
| PRCP | 1.54 | 1.44E-03 |
| NDUFAF2 | 1.55 | 2.95E-02 |
| RIOK2 | 1.55 | 2.33E-02 |
| NSMF | 1.55 | 1.66E-02 |
| TPPP3 | 1.55 | 1.65E-02 |
| KSR1 | 1.56 | 1.14E-02 |
| RBM47 | 1.56 | 3.51E-03 |
| KDELC1 | 1.56 | 2.16E-02 |
| WRN | 1.56 | 1.25E-02 |
| SEMA3B | 1.57 | 1.83E-02 |
| UXT | 1.57 | 9.74E-03 |
| IRS2 | 1.57 | 4.14E-02 |
| CDK1 | 1.58 | 4.73E-02 |
| PAM | 1.58 | 3.51E-02 |
| CBX2 | 1.58 | 2.66E-02 |
| CCNB1 | 1.60 | 4.87E-03 |
| WHSC1L1 | 1.60 | 1.95E-02 |
| INA | 1.61 | 1.12E-02 |
| ZNF131 | 1.62 | 1.97E-02 |

| | | |
|---|---|---|
| INA | 1.70 | 6.0E-03 |
| GFER | 1.71 | 3.0E-03 |
| A2M | 1.71 | 7.3E-03 |
| ACSL4 | 1.71 | 1.9E-02 |
| SOX9 | 1.72 | 3.5E-02 |
| SLMAP | 1.73 | 2.5E-02 |
| ADSL | 1.73 | 9.6E-03 |
| CDK6 | 1.74 | 4.9E-04 |
| PLEKHA1 | 1.74 | 2.4E-02 |
| TAGLN | 1.74 | 5.2E-03 |
| PSPC1 | 1.75 | 1.1E-02 |
| CCNB2 | 1.76 | 9.1E-03 |
| IGFBP7 | 1.77 | 5.3E-03 |
| GTPBP2 | 1.78 | 2.7E-02 |
| UPP1 | 1.78 | 6.8E-03 |
| SEC14L2 | 1.79 | 9.6E-04 |
| NHSL1 | 1.80 | 1.4E-02 |
| LGALS1 | 1.81 | 1.9E-02 |
| PSTPIP2 | 1.81 | 1.3E-02 |
| MIEF1 | 1.83 | 1.7E-03 |
| PRKAB2 | 1.83 | 2.9E-04 |
| C4A | 1.84 | 8.3E-03 |
| SH3BP4 | 1.84 | 2.5E-03 |
| ITPK1 | 1.86 | 4.7E-02 |
| PTPRF | 1.86 | 3.4E-03 |
| HBA1 | 1.86 | 1.2E-02 |
| DPYSL3 | 1.88 | 9.1E-03 |
| VASP | 1.88 | 1.1E-02 |
| LTF | 1.89 | 4.6E-02 |
| SERPINH1 | 1.90 | 1.5E-03 |
| RIN1 | 1.92 | 7.0E-03 |
| CGN | 1.94 | 5.3E-03 |
| SACS | 1.94 | 1.7E-03 |
| STK17A | 1.95 | 2.0E-02 |
| CORO1C | 1.98 | 2.5E-02 |
| STK38L | 2.00 | 6.9E-03 |
| C15orf52 | 2.04 | 7.5E-03 |
| TNC | 2.05 | 1.1E-02 |
| NXN | 2.05 | 4.0E-03 |
| SLFN5 | 2.06 | 1.6E-04 |
| RIMKLB | 2.06 | 1.2E-03 |
| LAMA4 | 2.06 | 8.4E-03 |
| PDP1 | 2.07 | 2.1E-02 |
| EHD4 | 2.08 | 3.2E-03 |
| FLNC | 2.11 | 1.9E-02 |
| CTGF | 2.12 | 3.2E-02 |
| NES | 2.14 | 4.2E-03 |
| PZP | 2.14 | 2.9E-03 |
| SNX18 | 2.16 | 9.8E-03 |
| LACTB | 2.18 | 1.1E-02 |
| LIMA1 | 2.20 | 9.3E-03 |
| RBPMS | 2.23 | 9.2E-03 |
| MYL9 | 2.23 | 6.5E-03 |
| L3HYPDH | 2.25 | 4.7E-03 |
| TGFBI | 2.26 | 1.4E-02 |
| BCAT1 | 2.26 | 5.0E-03 |
| FOSL1 | 2.29 | 4.0E-03 |
| NT5E | 2.31 | 4.1E-03 |
| GMPR2 | 2.36 | 1.0E-03 |
| RPL22L1 | 2.37 | 2.0E-04 |
| STC2 | 2.39 | 3.6E-03 |
| LUM | 2.46 | 6.7E-03 |
| SH3KBP1 | 2.46 | 1.3E-02 |
| KSR1 | 2.48 | 1.2E-03 |
| TLR1 | 2.60 | 2.4E-02 |
| RFTN1 | 2.65 | 1.5E-03 |
| HDGFRP3 | 2.66 | 2.0E-03 |
| COL5A2 | 2.70 | 1.6E-02 |

| | | |
|---|---|---|
| RAB4A | 1.38 | 3.4E-02 |
| NUDT9 | 1.38 | 2.8E-02 |
| SRP9 | 1.38 | 2.4E-03 |
| HSD17B4 | 1.38 | 7.6E-03 |
| PTRHD1 | 1.38 | 4.3E-02 |
| CYLD | 1.38 | 4.1E-02 |
| MESDC2 | 1.38 | 1.2E-02 |
| GNPAT | 1.38 | 4.8E-02 |
| UBAP2L | 1.38 | 9.8E-03 |
| POFUT1 | 1.38 | 7.0E-03 |
| ADO | 1.38 | 1.5E-02 |
| PCBD2 | 1.39 | 3.5E-02 |
| PLA2G16 | 1.39 | 3.4E-02 |
| MCTS1 | 1.39 | 1.7E-02 |
| PCDH7 | 1.40 | 1.8E-02 |
| SSFA2 | 1.40 | 3.0E-02 |
| PLEKHA5 | 1.40 | 1.7E-02 |
| TIMM8A | 1.40 | 3.8E-02 |
| RABGGTA | 1.40 | 9.5E-03 |
| FLNB | 1.40 | 1.2E-02 |
| UHRF1BP1L | 1.40 | 2.7E-02 |
| IMPA2 | 1.40 | 2.8E-02 |
| UBA1 | 1.40 | 9.3E-03 |
| LRRC57 | 1.40 | 2.3E-02 |
| RNPEP | 1.40 | 2.3E-02 |
| CUL4B | 1.40 | 2.5E-02 |
| DCPS | 1.40 | 9.8E-03 |
| SS18 | 1.40 | 1.3E-02 |
| NEDD4L | 1.41 | 1.3E-02 |
| BEGAIN | 1.41 | 1.0E-02 |
| CALR | 1.41 | 3.6E-02 |
| RBM39 | 1.41 | 9.5E-03 |
| USP6NL | 1.41 | 8.4E-03 |
| CCDC102A | 1.41 | 3.4E-02 |
| MLKL | 1.41 | 1.2E-03 |
| NENF | 1.42 | 2.0E-02 |
| BRD3 | 1.42 | 8.2E-03 |
| IPO9 | 1.42 | 6.0E-03 |
| C9orf142 | 1.42 | 3.8E-02 |
| AKAP11 | 1.42 | 5.0E-02 |
| AUH | 1.42 | 4.0E-02 |
| NKAP | 1.42 | 3.5E-02 |
| CARD6 | 1.43 | 4.9E-03 |
| CCDC152 | 1.43 | 2.6E-03 |
| CKAP2 | 1.43 | 2.9E-02 |
| PARP4 | 1.43 | 1.6E-03 |
| GDPGP1 | 1.43 | 1.5E-02 |
| CHID1 | 1.43 | 3.6E-02 |
| SMYD2 | 1.43 | 1.9E-02 |
| PMVK | 1.43 | 3.4E-03 |
| HPRT1 | 1.43 | 4.1E-02 |
| FOLR1 | 1.43 | 4.3E-02 |
| DNAJC25 | 1.44 | 3.1E-02 |
| PLEK2 | 1.44 | 2.0E-02 |
| CPEB4 | 1.45 | 1.5E-02 |
| FOXC2 | 1.45 | 4.8E-02 |
| PDIA3 | 1.45 | 2.0E-03 |
| TAMM41 | 1.45 | 3.6E-02 |
| NR2C2AP | 1.45 | 4.2E-02 |
| ZNF687 | 1.45 | 5.0E-02 |
| IRS2 | 1.45 | 2.1E-02 |
| DLG3 | 1.45 | 3.4E-02 |
| NT5DC1 | 1.45 | 3.6E-03 |
| NUCB2 | 1.45 | 1.1E-02 |
| PRPS1 | 1.46 | 4.2E-03 |
| ANXA10 | 1.46 | 3.9E-04 |
| CNPY3 | 1.46 | 6.1E-03 |
| KIAA1598 | 1.46 | 1.5E-02 |

| | | |
|---|---|---|
| GAA | 1.63 | 2.52E-02 |
| KLF5 | 1.63 | 2.86E-02 |
| ERI1 | 1.63 | 6.76E-03 |
| LAMC1 | 1.63 | 4.88E-02 |
| R3HCC1 | 1.64 | 3.48E-03 |
| PHF8 | 1.65 | 6.47E-03 |
| BCL3 | 1.65 | 4.32E-02 |
| NKAP | 1.65 | 1.69E-02 |
| RXRA | 1.67 | 1.77E-02 |
| RIN1 | 1.67 | 9.46E-03 |
| ERCC6L | 1.68 | 1.43E-02 |
| CAB39 | 1.68 | 1.35E-03 |
| TRPS1 | 1.68 | 4.68E-02 |
| ZBTB33 | 1.70 | 1.29E-02 |
| SEMA4B | 1.70 | 3.66E-02 |
| STXBP5 | 1.70 | 1.76E-02 |
| NKRF | 1.70 | 1.44E-02 |
| FHIT | 1.71 | 1.70E-02 |
| ELL2 | 1.71 | 1.07E-02 |
| STC2 | 1.71 | 1.93E-03 |
| NHS | 1.71 | 3.72E-02 |
| NUDT19 | 1.71 | 1.60E-02 |
| SERPINB6 | 1.71 | 4.61E-02 |
| ARHGEF16 | 1.72 | 9.04E-03 |
| ERO1L | 1.72 | 4.03E-02 |
| KIAA0947 | 1.72 | 1.71E-02 |
| EDIL3 | 1.73 | 2.67E-02 |
| PLEKHA1 | 1.73 | 2.90E-02 |
| PUS1 | 1.73 | 5.42E-05 |
| AGTRAP | 1.74 | 3.55E-02 |
| RREB1 | 1.74 | 1.72E-02 |
| CLYBL | 1.74 | 2.28E-02 |
| FBXO4 | 1.74 | 1.37E-02 |
| PLEKHA5 | 1.75 | 5.20E-03 |
| GGH | 1.75 | 1.03E-03 |
| UBR5 | 1.75 | 2.10E-03 |
| GNL2 | 1.76 | 4.93E-02 |
| PGK2 | 1.76 | 2.62E-02 |
| PHC2 | 1.76 | 2.06E-02 |
| EFHD2 | 1.77 | 6.25E-03 |
| GGT1 | 1.79 | 1.11E-02 |
| MAFF | 1.79 | 4.94E-02 |
| COLGALT1 | 1.80 | 3.74E-02 |
| LEPREL1 | 1.81 | 8.35E-03 |
| SCAMP1 | 1.82 | 1.87E-02 |
| PITPNC1 | 1.82 | 4.86E-03 |
| TGFBI | 1.82 | 4.21E-03 |
| SLC5A6 | 1.83 | 4.32E-02 |
| PLOD1 | 1.83 | 2.98E-02 |
| SOWAHC | 1.83 | 1.34E-02 |
| ACYP2 | 1.84 | 4.66E-02 |
| IMPAD1 | 1.84 | 4.67E-02 |
| PDP1 | 1.85 | 9.13E-03 |
| TPBG | 1.85 | 4.45E-02 |
| PLIN2 | 1.85 | 4.57E-03 |
| KYNU | 1.87 | 8.19E-03 |
| FAM117B | 1.89 | 1.25E-02 |
| ANK3 | 1.89 | 4.62E-03 |
| CASP4 | 1.90 | 1.32E-02 |
| L3MBTL3 | 1.92 | 3.30E-02 |
| CYCS | 1.92 | 1.77E-02 |
| CTSC | 1.92 | 2.77E-02 |
| SBNO2 | 1.95 | 4.14E-02 |
| LDLR | 1.96 | 2.96E-02 |
| THEM4 | 1.97 | 1.22E-02 |
| ATP7A | 1.97 | 1.71E-02 |
| SLC38A2 | 1.98 | 4.97E-02 |
| FAT1 | 1.98 | 2.16E-02 |

| | | |
|---|---|---|
| INPP4B | 2.70 | 1.7E-02 |
| DOCK4 | 2.83 | 3.1E-03 |
| FBLIM1 | 2.83 | 7.0E-05 |
| EDIL3 | 2.83 | 2.6E-03 |
| TMOD2 | 2.83 | 1.2E-02 |
| DMBT1 | 2.83 | 4.9E-03 |
| HTRA1 | 2.84 | 5.3E-03 |
| PRSS23 | 2.87 | 4.2E-03 |
| GPR158 | 2.92 | 4.3E-02 |
| ITIH2 | 2.94 | 6.1E-03 |
| CNN1 | 2.96 | 1.8E-03 |
| LTBP1 | 2.97 | 2.0E-02 |
| THBS1 | 2.99 | 1.2E-02 |
| SERPINE1 | 3.10 | 3.2E-02 |
| NID1 | 3.17 | 2.6E-04 |
| LAMB1 | 3.37 | 1.7E-02 |
| COL1A1 | 3.49 | 6.1E-03 |
| TGM2 | 3.53 | 2.0E-04 |
| AKAP12 | 3.54 | 6.0E-03 |
| VTN | 3.55 | 2.1E-04 |
| HBD | 3.81 | 4.9E-04 |
| TESC | 3.83 | 6.9E-03 |
| FABP3 | 4.00 | 1.5E-03 |
| DCLK1 | 4.53 | 4.9E-03 |
| FN1 | 4.90 | 1.1E-03 |
| ANPEP | 5.91 | 6.8E-03 |
| ARHGDIB | 6.38 | 4.8E-03 |
| SERPINE2 | 6.84 | 5.6E-03 |

| | | |
|---|---|---|
| CLIC2 | 1.47 | 9.7E-03 |
| CHMP1B | 1.47 | 1.6E-02 |
| NPEPL1 | 1.47 | 8.6E-04 |
| HAGH | 1.47 | 7.1E-04 |
| SPECC1 | 1.47 | 1.1E-02 |
| LYPLAL1 | 1.48 | 4.0E-02 |
| GSTO1 | 1.48 | 1.8E-02 |
| GLA | 1.48 | 8.2E-03 |
| MDP1 | 1.48 | 1.4E-02 |
| DTD1 | 1.48 | 2.6E-02 |
| PDCD4 | 1.48 | 1.3E-02 |
| MSRB2 | 1.48 | 3.7E-02 |
| DPP7 | 1.48 | 1.7E-03 |
| HEXB | 1.49 | 1.0E-03 |
| CLU | 1.50 | 1.1E-02 |
| GUSB | 1.50 | 9.1E-03 |
| CMAS | 1.50 | 6.3E-03 |
| YOD1 | 1.50 | 4.6E-04 |
| SCAMP3 | 1.51 | 3.5E-02 |
| ACSL4 | 1.51 | 3.8E-03 |
| GM2A | 1.51 | 1.5E-03 |
| DPP3 | 1.51 | 4.0E-03 |
| OSTF1 | 1.52 | 4.5E-03 |
| GPATCH2 | 1.52 | 3.1E-02 |
| BZW1 | 1.52 | 5.5E-03 |
| L3HYPDH | 1.52 | 2.4E-02 |
| DCAF8 | 1.52 | 7.6E-03 |
| NAPRT1 | 1.53 | 1.0E-02 |
| S100A13 | 1.53 | 8.9E-03 |
| CSPG4 | 1.53 | 1.2E-02 |
| SERPINB1 | 1.53 | 1.6E-03 |
| SGK1 | 1.53 | 4.8E-03 |
| CRLF1 | 1.53 | 1.8E-02 |
| ALDH16A1 | 1.53 | 1.1E-02 |
| SSNA1 | 1.53 | 3.6E-02 |
| RCN2 | 1.53 | 1.2E-03 |
| VAT1L | 1.53 | 4.8E-02 |
| RUFY1 | 1.54 | 4.0E-03 |
| GAA | 1.54 | 5.0E-03 |
| ALDH9A1 | 1.55 | 1.5E-03 |
| PLIN3 | 1.55 | 1.3E-02 |
| PSMB8 | 1.55 | 4.7E-03 |
| TCEAL4 | 1.55 | 1.5E-02 |
| SERPINB8 | 1.55 | 9.3E-03 |
| BAG1 | 1.56 | 9.5E-03 |
| HS1BP3 | 1.56 | 7.8E-03 |
| RBM47 | 1.56 | 1.0E-02 |
| NPC2 | 1.56 | 1.8E-02 |
| NADSYN1 | 1.56 | 2.4E-02 |
| KLF4 | 1.57 | 2.3E-02 |
| SIAE | 1.57 | 4.5E-02 |
| KYNU | 1.57 | 1.0E-02 |
| UFC1 | 1.57 | 1.8E-02 |
| AGTPBP1 | 1.57 | 2.2E-03 |
| PPIL3 | 1.57 | 8.0E-03 |
| MVP | 1.58 | 5.1E-03 |
| DNAJC3 | 1.58 | 1.7E-02 |
| FLYWCH2 | 1.58 | 9.0E-03 |
| NDOR1 | 1.58 | 3.5E-03 |
| GUK1 | 1.59 | 8.3E-04 |
| PEPD | 1.60 | 1.2E-02 |
| THEM4 | 1.60 | 2.9E-02 |
| FAM160B1 | 1.60 | 2.0E-02 |
| DAK | 1.60 | 4.7E-03 |
| DUSP23 | 1.60 | 8.5E-04 |
| ZBED1 | 1.60 | 4.0E-03 |
| N6AMT1 | 1.61 | 6.8E-03 |
| DTX3L | 1.62 | 4.4E-02 |

| | | |
|---|---|---|
| BMP1 | 1.99 | 4.25E-02 |
| SLC44A1 | 1.99 | 3.23E-02 |
| SIAE | 2.00 | 7.61E-03 |
| COX6A1 | 2.01 | 4.95E-02 |
| SH3KBP1 | 2.01 | 3.99E-03 |
| H6PD | 2.04 | 3.01E-04 |
| FMNL2 | 2.04 | 1.63E-02 |
| MAOB | 2.08 | 7.23E-03 |
| PLP2 | 2.08 | 3.12E-02 |
| DTWD1 | 2.17 | 1.82E-02 |
| HK2 | 2.17 | 4.66E-02 |
| MAFK | 2.18 | 5.96E-03 |
| TGFB1 | 2.21 | 3.96E-02 |
| NAMPT | 2.22 | 4.10E-04 |
| LOXL2 | 2.25 | 3.61E-02 |
| NR2F2 | 2.31 | 4.02E-02 |
| PCDH7 | 2.32 | 1.62E-02 |
| FAM32A | 2.32 | 1.07E-02 |
| AKAP12 | 2.33 | 3.94E-03 |
| CGA | 2.34 | 1.09E-02 |
| SLC12A7 | 2.35 | 2.07E-02 |
| ABCB1 | 2.36 | 9.36E-03 |
| DUSP23 | 2.38 | 3.42E-02 |
| NOV | 2.38 | 3.65E-02 |
| CA2 | 2.39 | 1.66E-02 |
| NES | 2.39 | 5.23E-03 |
| ANPEP | 2.41 | 1.39E-02 |
| PLA2G4A | 2.58 | 1.63E-02 |
| TNC | 2.65 | 5.67E-03 |
| ACSL4 | 2.65 | 5.57E-03 |
| FAM120C | 2.72 | 7.05E-03 |
| ITIH2 | 2.73 | 5.33E-03 |
| ZNF185 | 2.73 | 1.09E-02 |
| PRSS23 | 2.75 | 1.53E-02 |
| LAMB1 | 2.80 | 6.86E-03 |
| DPYD | 3.00 | 6.38E-04 |
| DOCK4 | 3.10 | 2.61E-02 |
| CPM | 3.16 | 7.13E-03 |
| DMBT1 | 3.33 | 1.03E-02 |
| TESC | 3.40 | 6.00E-03 |
| SERPINE2 | 3.51 | 6.16E-03 |
| HBA1 | 3.52 | 1.12E-02 |
| DCLK1 | 3.68 | 4.34E-03 |
| NT5E | 3.70 | 1.73E-02 |
| IFI16 | 3.72 | 3.05E-03 |
| NID1 | 4.56 | 4.51E-04 |
| TMOD2 | 4.68 | 2.97E-03 |
| HTRA1 | 4.89 | 3.68E-04 |
| TIMP1 | 5.00 | 5.06E-03 |
| PTGS1 | 5.39 | 1.34E-03 |
| RCN3 | 5.56 | 4.72E-02 |
| ITGA2 | 5.94 | 1.88E-02 |
| SERPINB3 | 6.75 | 7.79E-03 |
| VTN | 7.76 | 1.64E-02 |
| FSTL4 | 8.78 | 1.32E-03 |
| HBD | 11.83 | 5.33E-04 |

| | | |
|---|---|---|
| DSP | 1.62 | 2.8E-03 |
| L1RE1 | 1.63 | 1.4E-02 |
| GORAB | 1.63 | 2.9E-02 |
| CD97 | 1.63 | 4.3E-04 |
| SERPINB6 | 1.64 | 4.2E-03 |
| MYO6 | 1.64 | 4.7E-04 |
| OSBPL6 | 1.64 | 1.1E-02 |
| OBSL1 | 1.64 | 8.1E-03 |
| HBD | 1.65 | 7.6E-03 |
| ADARB1 | 1.65 | 1.6E-02 |
| ETHE1 | 1.65 | 1.4E-02 |
| HPD | 1.66 | 4.3E-03 |
| SELENBP1 | 1.66 | 5.3E-03 |
| SUMF1 | 1.66 | 4.6E-02 |
| CHCHD5 | 1.67 | 1.5E-03 |
| DERA | 1.67 | 2.5E-03 |
| PARP9 | 1.67 | 1.3E-02 |
| B3GALTL | 1.68 | 1.4E-02 |
| S100A11 | 1.68 | 3.9E-03 |
| ABHD5 | 1.68 | 8.4E-05 |
| S100A14 | 1.69 | 2.1E-02 |
| CD46 | 1.70 | 2.8E-02 |
| TSTA3 | 1.71 | 1.1E-02 |
| NIPSNAP3A | 1.71 | 1.7E-02 |
| ARHGAP42 | 1.71 | 4.8E-03 |
| DDX60 | 1.71 | 3.5E-03 |
| POMP | 1.72 | 6.1E-03 |
| CTSA | 1.72 | 2.8E-02 |
| DNASE2 | 1.74 | 1.9E-03 |
| S100A16 | 1.74 | 4.4E-02 |
| C1orf85 | 1.75 | 8.9E-03 |
| CASP9 | 1.76 | 1.6E-02 |
| FYTTD1 | 1.76 | 5.2E-03 |
| PTGS1 | 1.77 | 4.3E-02 |
| SH3BGRL | 1.78 | 7.3E-03 |
| OCRL | 1.79 | 6.3E-03 |
| BROX | 1.79 | 5.6E-05 |
| SMPDL3A | 1.79 | 2.2E-03 |
| HSD17B11 | 1.80 | 3.5E-02 |
| ACADL | 1.80 | 4.4E-02 |
| GIPC3 | 1.80 | 1.3E-02 |
| B2M | 1.80 | 4.7E-02 |
| PSMB9 | 1.80 | 3.3E-03 |
| PLA2G4A | 1.80 | 3.0E-03 |
| RBM3 | 1.81 | 1.9E-03 |
| SERPINB5 | 1.83 | 3.8E-03 |
| CLIC3 | 1.84 | 1.0E-02 |
| CA2 | 1.85 | 1.0E-02 |
| TPP1 | 1.86 | 8.8E-03 |
| MCTP2 | 1.86 | 1.8E-02 |
| GLB1 | 1.88 | 9.4E-03 |
| PECR | 1.88 | 1.8E-03 |
| NEK7 | 1.90 | 1.1E-02 |
| SERPINB9 | 1.90 | 6.4E-03 |
| CTSC | 1.93 | 6.9E-04 |
| ENTPD5 | 1.93 | 9.6E-03 |
| SULT1A3 | 1.94 | 1.7E-03 |
| CAPG | 1.94 | 1.1E-02 |
| PYCARD | 1.97 | 1.0E-02 |
| CD55 | 2.01 | 3.2E-02 |
| RNASEL | 2.02 | 5.2E-03 |
| PLP2 | 2.02 | 2.3E-02 |
| GRAMD1A | 2.02 | 3.1E-03 |
| IFI16 | 2.02 | 1.0E-03 |
| ASS1 | 2.02 | 4.4E-03 |
| TCEAL1 | 2.03 | 1.4E-02 |
| DNAJC12 | 2.11 | 3.3E-02 |
| RIMKLB | 2.14 | 9.6E-03 |

| | | |
|---|---|---|
| POGLUT1 | 2.14 | 2.3E-02 |
| RAET1G | 2.16 | 3.6E-03 |
| PITPNM1 | 2.19 | 9.7E-03 |
| ACYP2 | 2.21 | 2.7E-03 |
| ABCB1 | 2.23 | 8.2E-03 |
| S100P | 2.31 | 2.3E-03 |
| NPR1 | 2.32 | 1.2E-02 |
| HNMT | 2.38 | 1.5E-03 |
| SH3KBP1 | 2.48 | 1.2E-03 |
| PHYH | 2.60 | 1.7E-03 |
| TESC | 2.83 | 2.9E-02 |
| MUC1 | 3.15 | 6.4E-03 |
| TIMP1 | 3.29 | 4.6E-03 |
| GDA | 3.40 | 4.1E-04 |
| RPL22L1 | 4.13 | 3.9E-03 |
| ARHGDIB | 4.51 | 2.0E-02 |
| SERPINB3 | 4.66 | 1.7E-03 |
| ADIRF | 5.51 | 2.0E-02 |
| CGA | 8.36 | 5.2E-05 |
