## Supplementary Table S3 for "Three ALS genes regulate expression of the MHC class II antigen presentation pathway"

**Table S2. GSEA negative in FUS KO line.** Gene set, size, False Discovery Rate (FDR q-val < 0.25) and normalized enrichment score (NES < -2) are listed. Immune gene sets are labeled in red. Gene sets shared by FUS, TAF15 and MATR3 KOs are highlighted in yellow. C5.bp.v7.0 database was used for GSEA.

| Gene set | Size | FDR q-val | NES |
|---|---|---|---|
| GO_INNATE_IMMUNE_RESPONSE | 77 | 0.00000000 | -3.0006218 |
| GO_DEFENSE_RESPONSE | 113 | 0.00091200 | -2.687138 |
| GO_RESPONSE_TO_INTERFERON_GAMMA | 20 | 0.00060800 | -2.6342564 |
| GO_ANTIGEN_PROCESSING_AND_PRESENTATION_OF_PEPTIDE_ANTIGEN | 26 | 0.00159744 | -2.5922835 |
| GO_CELLULAR_AMINO_ACID_METABOLIC_PROCESS | 36 | 0.00217036 | -2.5622177 |
| GO_OXIDATION_REDUCTION_PROCESS | 100 | 0.00225526 | -2.5361683 |
| GO_CELLULAR_RESPONSE_TO_XENOBIOTIC_STIMULUS | 17 | 0.00232802 | -2.5183525 |
| GO_CYTOKINE_MEDIATED_SIGNALING_PATHWAY | 60 | 0.00315446 | -2.487478 |
| GO_ANTIGEN_PROCESSING_AND_PRESENTATION | 30 | 0.00300055 | -2.4856365 |
| GO_IMMUNE_RESPONSE_REGULATING_SIGNALING_PATHWAY | 51 | 0.00334529 | -2.4727445 |
| GO_REGULATION_OF_IMMUNE_RESPONSE | 80 | 0.00393349 | -2.4593008 |
| GO_ACTIVATION_OF_IMMUNE_RESPONSE | 55 | 0.00584696 | -2.4144528 |
| GO_POSITIVE_REGULATION_OF_IMMUNE_RESPONSE | 68 | 0.00553757 | -2.409143 |
| GO_RESPONSE_TO_XENOBIOTIC_STIMULUS | 31 | 0.00514203 | -2.4084952 |
| GO_T_CELL_RECEPTOR_SIGNALING_PATHWAY | 17 | 0.00527432 | -2.3976939 |
| GO_IMMUNE_RESPONSE_REGULATING_CELL_SURFACE_RECEPTOR_SIGNALING_PATHWAY | 35 | 0.00752869 | -2.3553765 |
| GO_POSITIVE_REGULATION_OF_IMMUNE_SYSTEM_PROCESS | 79 | 0.01501998 | -2.2811222 |
| GO_ANTIGEN_RECEPTOR_MEDIATED_SIGNALING_PATHWAY | 21 | 0.02425738 | -2.21293 |
| GO_REGULATION_OF_INNATE_IMMUNE_RESPONSE | 42 | 0.02840877 | -2.1865716 |
| GO_CELLULAR_KETONE_METABOLIC_PROCESS | 24 | 0.02847156 | -2.1797576 |
| GO_ALPHA_AMINO_ACID_METABOLIC_PROCESS | 25 | 0.02736705 | -2.1782997 |
| GO_ACTIVATION_OF_INNATE_IMMUNE_RESPONSE | 33 | 0.03475300 | -2.1448014 |
| GO_ORGANIC_ACID_METABOLIC_PROCESS | 94 | 0.04130429 | -2.1170588 |
| GO_SMALL_MOLECULE_METABOLIC_PROCESS | 175 | 0.07019101 | -2.0343473 |
| GO_ANTIGEN_PROCESSING_AND_PRESENTATION_OF_PEPTIDE_OR_POLYSACCHARIDE_ANTIGEN_VIA_MHC_CLASS_II | 18 | 0.08486915 | -2.0024452 |
| GO_REGULATION_OF_IMMUNE_SYSTEM_PROCESS | 116 | 0.08181296 | -2.001703 |
| GO_REGULATION_OF_DEFENSE_RESPONSE | 61 | 0.07908427 | -2.0012279 |

**Table S2. GSEA negative in EWSR1 KO line.** Gene set name, size, False Discovery Rate (FDR q-val < 0.25) and normalized enrichment score (NES < -2) are listed. C5.bp.v7.0 database was used for GSEA.

| Gene Set | Size | FDR q-val | NES |
|---|---|---|---|
| GO_RIBOSOME_BIOGENESIS | 51 | 0.00000000 | -3.4037356 |
| GO_RIBONUCLEOPROTEIN_COMPLEX_BIOGENESIS | 74 | 0.00000000 | -3.1898916 |
| GO_RIBOSOMAL_LARGE_SUBUNIT_BIOGENESIS | 16 | 0.00196973 | -2.7003446 |
| GO_RRNA_METABOLIC_PROCESS | 38 | 0.00248230 | -2.623586 |
| GO_NUCLEAR_TRANSCRIBED_MRNA_CATABOLIC_PROCESS_NONSENSE_MEDIATED_DECAY | 16 | 0.00478446 | -2.4152641 |
| GO_MITOTIC_NUCLEAR_DIVISION | 42 | 0.01263519 | -2.229762 |
| GO_DEVELOPMENTAL_MATURATION | 21 | 0.02241550 | -2.0937681 |
| GO_NUCLEAR_CHROMOSOME_SEGREGATION | 33 | 0.03248555 | -2.0168114 |

**Table S2. GSEA negative in TAF15 KO line.** Gene set, size, False Discovery Rate (FDR q-val < 0.25) and normalized enrichment score (NES < -2) are listed. Immune gene sets are labeled in red. Gene sets shared by FUS, TAF15 and MATR3 are highlighted in yellow. C5.bp.v7.0 database was used for GSEA.

| Gene set | Size | FDR q-val | NES |
|---|---|---|---|
| GO_ANTIGEN_RECEPTOR_MEDIATED_SIGNALING_PATHWAY | 16 | 0.01255602 | -2.6114767 |
| GO_IMMUNE_RESPONSE_REGULATING_CELL_SURFACE_RECEPTOR_SIGNALING_PATHWAY | 28 | 0.05573051 | -2.4441483 |
| GO_INNATE_IMMUNE_RESPONSE | 69 | 0.06576295 | -2.3861315 |
| GO_SUPRAMOLECULAR_FIBER_ORGANIZATION | 85 | 0.05610438 | -2.3719552 |
| GO_ANTIGEN_PROCESSING_AND_PRESENTATION | 29 | 0.05008980 | -2.3598845 |
| GO_IMMUNE_RESPONSE_REGULATING_SIGNALING_PATHWAY | 35 | 0.04639125 | -2.3470206 |
| GO_CYTOSKELETON_ORGANIZATION | 155 | 0.03976392 | -2.3468869 |
| GO_ACTOMYOSIN_STRUCTURE_ORGANIZATION | 26 | 0.03575383 | -2.3435242 |
| GO_REGULATION_OF_CYSTEINE_TYPE_ENDOPEPTIDASE_ACTIVITY | 24 | 0.04013244 | -2.3254306 |
| GO_RESPONSE_TO_TEMPERATURE_STIMULUS | 26 | 0.03929509 | -2.3153644 |
| GO_ANTIGEN_PROCESSING_AND_PRESENTATION_OF_PEPTIDE_ANTIGEN | 24 | 0.04222152 | -2.2964225 |
| GO_MUSCLE_SYSTEM_PROCESS | 42 | 0.04295725 | -2.2852552 |
| GO_RESPONSE_TO_INTERFERON_GAMMA | 23 | 0.05123466 | -2.259336 |
| GO_REGULATION_OF_SUPRAMOLECULAR_FIBER_ORGANIZATION | 50 | 0.04805442 | -2.2583408 |
| GO_ACTIN_FILAMENT_BUNDLE_ORGANIZATION | 22 | 0.05854884 | -2.2245488 |
| GO_ACTIN_FILAMENT_BASED_PROCESS | 104 | 0.05693759 | -2.2189677 |
| GO_ACTIN_FILAMENT_ORGANIZATION | 54 | 0.06089318 | -2.2027233 |
| GO_ACTIVATION_OF_IMMUNE_RESPONSE | 41 | 0.06291314 | -2.192806 |
| GO_SIGNAL_TRANSDUCTION_BY_PROTEIN_PHOSPHORYLATION | 73 | 0.06254358 | -2.1867177 |
| GO_RESPONSE_TO_HEAT | 22 | 0.06398709 | -2.1763494 |
| GO_PROTEIN_HOMOOLIGOMERIZATION | 35 | 0.06547906 | -2.1663308 |
| GO_CELLULAR_RESPONSE_TO_HEAT | 16 | 0.07558033 | -2.1423497 |
| GO_POSITIVE_REGULATION_OF_IMMUNE_RESPONSE | 55 | 0.07351065 | -2.1396751 |
| GO_REGULATION_OF_CELLULAR_COMPONENT_BIOGENESIS | 92 | 0.07794090 | -2.1254222 |
| GO_CYTOKINE_MEDIATED_SIGNALING_PATHWAY | 57 | 0.07606001 | -2.123304 |
| GO_REGULATION_OF_ANATOMICAL_STRUCTURE_MORPHOGENESIS | 97 | 0.08376291 | -2.1045048 |
| GO_REGULATION_OF_CYTOSKELETON_ORGANIZATION | 65 | 0.09901235 | -2.0761282 |
| GO_NEGATIVE_REGULATION_OF_PROTEIN_COMPLEX_ASSEMBLY | 18 | 0.10942685 | -2.0558207 |
| GO_REGULATION_OF_IMMUNE_RESPONSE | 71 | 0.10607975 | -2.0554473 |
| GO_NEGATIVE_REGULATION_OF_SUPRAMOLECULAR_FIBER_ORGANIZATION | 25 | 0.10293122 | -2.054778 |
| GO_REGULATION_OF_CELL_SHAPE | 23 | 0.10069886 | -2.0533154 |
| GO_CELL_CELL_SIGNALING_BY_WNT | 30 | 0.10102225 | -2.0475688 |
| GO_ANTIGEN_PROCESSING_AND_PRESENTATION_OF_PEPTIDE_OR_POLYSACCHARIDE_ANTIGEN_VIA_MHC_CLASS_II | 20 | 0.10076575 | -2.0437331 |
| GO_NEGATIVE_REGULATION_OF_CELL_GROWTH | 20 | 0.09879983 | -2.0425258 |
| GO_MUSCLE_CONTRACTION | 34 | 0.09666532 | -2.0410233 |
| GO_SECOND_MESSENGER_MEDIATED_SIGNALING | 24 | 0.10116559 | -2.0297337 |
| GO_REGULATION_OF_ACTIN_FILAMENT_BASED_PROCESS | 48 | 0.09910867 | -2.0288262 |
| GO_REGULATION_OF_PROTEIN_COMPLEX_ASSEMBLY | 41 | 0.09960495 | -2.0242722 |
| GO_CANONICAL_WNT_SIGNALING_PATHWAY | 19 | 0.09969999 | -2.0203545 |
| GO_ESTABLISHMENT_OF_CELL_POLARITY | 15 | 0.09802551 | -2.0193837 |

| | | | |
|---|---|---|---|
| GO_ACTIN_POLYMERIZATION_OR_DEPOLYMERIZATION | 33 | 0.10528257 | -2.0054553 |
| GO_REGULATION_OF_ACTIN_FILAMENT_ORGANIZATION | 39 | 0.10376237 | -2.0042527 |
| GO_REGULATION_OF_PROTEIN_LOCALIZATION_TO_MEMBRANE | 22 | 0.10234126 | -2.002657 |

**Table S2. GSEA negative in MATR3 KO line.** Gene set, size, False Discovery Rate (FDR q-val <0.25) and normalized enrichment score (NES < -2) are listed. Immune gene sets are labeled in red. Gene sets shared by FUS, TAF15 and MATR3 are highlighted in yellow. C5.bp.v7.0 database was used for GSEA.

| Gene set | Size | FDR q-val | NES |
|---|---|---|---|
| GO_INNATE_IMMUNE_RESPONSE | 69 | 0.01759193 | -2.5339592 |
| GO_AMINE_METABOLIC_PROCESS | 16 | 0.22920977 | -2.1466372 |
| GO_POSITIVE_REGULATION_OF_IMMUNE_RESPONSE | 53 | 0.18053447 | -2.1403167 |
| GO_RESPONSE_TO_INTERFERON_GAMMA | 24 | 0.14973083 | -2.1356726 |
| GO_COFACTOR_METABOLIC_PROCESS | 45 | 0.13660790 | -2.1259844 |
| GO_POSITIVE_REGULATION_OF_IMMUNE_SYSTEM_PROCESS | 65 | 0.15737589 | -2.087914 |
| GO_CELLULAR_PROTEIN_CONTAINING_COMPLEX_ASSEMBLY | 72 | 0.13963939 | -2.0860527 |
| GO_ANTIGEN_RECEPTOR_MEDIATED_SIGNALING_PATHWAY | 22 | 0.15314719 | -2.0546396 |
| GO_T_CELL_RECEPTOR_SIGNALING_PATHWAY | 21 | 0.15408197 | -2.0403965 |
| GO_ANTIGEN_PROCESSING_AND_PRESENTATION_OF_PEPTIDE_ANTIGEN | 25 | 0.14306742 | -2.0375745 |
| GO_DEFENSE_RESPONSE | 101 | 0.14274974 | -2.0240035 |
| GO_RESPONSE_TO_INTERLEUKIN_1 | 23 | 0.13220713 | -2.02343 |
| GO_ELECTRON_TRANSPORT_CHAIN | 17 | 0.13436149 | -2.0085292 |
| GO_ANTIGEN_PROCESSING_AND_PRESENTATION | 27 | 0.12920770 | -2.0032337 |
