## Supplementary Table S4 for "Three ALS genes regulate expression of the MHC class II antigen presentation pathway"

**Table S4. GSEA positive in FUS KO line.** Gene set name, size, False Discovery Rate (FDR q-val <0.25) and normalized enrichment score (NES > 2) are listed. Immune gene sets are labeled in red. C5.bp.v7.0 database was used for GSEA.

| Gene set | Size | FDR q-val | NES |
|---|---|---|---|
| GO_EXTRACELLULAR_STRUCTURE_ORGANIZATION | 35 | 0.00000000 | 3.6859202 |
| GO_REGULATION_OF_CELL_ADHESION | 53 | 0.00000000 | 3.2261217 |
| GO_REGULATION_OF_CELL_SUBSTRATE_ADHESION | 27 | 0.00000000 | 2.9165444 |
| GO_BIOLOGICAL_ADHESION | 104 | 0.00000000 | 2.9000404 |
| GO_ACTIN_FILAMENT_BASED_PROCESS | 94 | 0.00046569 | 2.8045073 |
| GO_REGULATION_OF_CELLULAR_COMPONENT_MOVEMENT | 82 | 0.00097309 | 2.6370711 |
| GO_CELL_SUBSTRATE_ADHESION | 40 | 0.00150024 | 2.6149147 |
| GO_NEGATIVE_REGULATION_OF_CELL_ADHESION | 25 | 0.00448719 | 2.4551814 |
| GO_MULTI_MULTICELLULAR_ORGANISM_PROCESS | 19 | 0.00398861 | 2.4550517 |
| GO_ANATOMICAL_STRUCTURE_FORMATION_INVOLVED_IN_MORPHOGENESIS | 70 | 0.00406378 | 2.4396443 |
| GO_NEGATIVE_REGULATION_OF_PEPTIDASE_ACTIVITY | 24 | 0.00379253 | 2.4347742 |
| GO_CELL_MATRIX_ADHESION | 27 | 0.00538628 | 2.3847082 |
| GO_POSITIVE_REGULATION_OF_CELL_ADHESION | 27 | 0.00699845 | 2.3387094 |
| GO_CELL_MOTILITY | 117 | 0.01032177 | 2.2859507 |
| GO_WOUND_HEALING | 53 | 0.01054406 | 2.2751844 |
| GO_RESPONSE_TO_WOUNDING | 65 | 0.01032381 | 2.2725186 |
| GO_REGULATION_OF_CELL_MATRIX_ADHESION | 16 | 0.01047052 | 2.2595778 |
| GO_CELL_JUNCTION_ORGANIZATION | 38 | 0.01060945 | 2.2504504 |
| GO_REGULATION_OF_ACTIN_FILAMENT_BASED_PROCESS | 50 | 0.01485458 | 2.202184 |
| GO_LOCOMOTION | 133 | 0.01452088 | 2.1989582 |
| GO_COAGULATION | 34 | 0.01525565 | 2.1849203 |
| GO_NEGATIVE_REGULATION_OF_HYDROLASE_ACTIVITY | 38 | 0.01768820 | 2.1658938 |
| GO_NEGATIVE_REGULATION_OF_LOCOMOTION | 32 | 0.02054059 | 2.1389947 |
| GO_MYELOID_LEUKOCYTE_DIFFERENTIATION | 15 | 0.02083975 | 2.1324804 |
| GO_CELL_JUNCTION_ASSEMBLY | 34 | 0.02014817 | 2.1314719 |
| GO_ACTIN_FILAMENT_ORGANIZATION | 55 | 0.01986449 | 2.128688 |
| GO_NEGATIVE_REGULATION_OF_CELL_MOTILITY | 26 | 0.01948007 | 2.1247046 |
| GO_NEGATIVE_REGULATION_OF_MULTICELLULAR_ORGANISMAL_PROCESS | 90 | 0.02027449 | 2.1158328 |
| GO_NEGATIVE_REGULATION_OF_PROTEOLYSIS | 34 | 0.02577115 | 2.0812168 |
| GO_REPRODUCTIVE_SYSTEM_DEVELOPMENT | 19 | 0.03199566 | 2.0459886 |
| GO_GLIOGENESIS | 18 | 0.03148727 | 2.0443294 |
| GO_REGULATION_OF_BODY_FLUID_LEVELS | 44 | 0.03158241 | 2.0401454 |
| GO_EMBRYO_DEVELOPMENT | 63 | 0.03442579 | 2.0249493 |
| GO_CYTOSKELETON_ORGANIZATION | 154 | 0.03812958 | 2.002616 |
| GO_REGULATION_OF_PEPTIDASE_ACTIVITY | 43 | 0.03707262 | 2.0025105 |

**Table S4. GSEA positive in EWSR1 KO line.** Gene set, size, False Discovery Rate (FDR q-val <0.25) and normalized enrichment score (NES > 2) are listed. Immune gene sets are labeled in red. C5.bp.v7.0 database was used for GSEA.

| Gene set | Size | FDR q-val | NES |
|---|---|---|---|
| GO_MYELOID_LEUKOCYTE_MEDIATED_IMMUNITY | 79 | 0.02845459 | 2.4827263 |
| GO_LEUKOCYTE_MEDIATED_IMMUNITY | 95 | 0.02190341 | 2.4433587 |
| GO_IMMUNE_EFFECTOR_PROCESS | 126 | 0.02089711 | 2.4080465 |
| GO_MYELOID_LEUKOCYTE_ACTIVATION | 82 | 0.01719020 | 2.397132 |
| GO_RESPONSE_TO_VIRUS | 29 | 0.02239789 | 2.3349757 |
| GO_DEFENSE_RESPONSE_TO_VIRUS | 24 | 0.03186484 | 2.274665 |
| GO_CELL_ACTIVATION_INVOLVED_IN_IMMUNE_RESPONSE | 87 | 0.03038221 | 2.2573912 |
| GO_DEFENSE_RESPONSE_TO_OTHER_ORGANISM | 33 | 0.03350473 | 2.2233927 |
| GO_INNATE_IMMUNE_RESPONSE | 77 | 0.03349157 | 2.2076132 |
| GO_SMALL_MOLECULE_CATABOLIC_PROCESS | 57 | 0.04568381 | 2.1493092 |
| GO_REGULATION_OF_MULTI_ORGANISM_PROCESS | 38 | 0.04846210 | 2.1270251 |
| GO_VACUOLAR_TRANSPORT | 15 | 0.05387243 | 2.1014 |
| GO_ANTIGEN_PROCESSING_AND_PRESENTATION_OF_PEPTIDE_ANTIGEN | 27 | 0.05617986 | 2.0844245 |
| GO_VIRAL_LIFE_CYCLE | 28 | 0.08046717 | 2.0203848 |

**Table S4. GSEA positive in TAF15 KO line.** Gene set name, size, False Discovery Rate (FDR q-val <0.25) and normalized enrichment score (NES > 2) are listed. Immune gene sets are labeled in red. C5.bp.v7.0 database was used for GSEA.

| Gene set name | Size | FDR q-val | NES |
|---|---|---|---|
| GO_RIBONUCLEOPROTEIN_COMPLEX_BIOGENESIS | 58 | 0.00000000 | 3.2111974 |
| GO_RIBOSOME_BIOGENESIS | 47 | 0.00000000 | 3.0282516 |
| GO_NCRNA_METABOLIC_PROCESS | 64 | 0.00000000 | 2.9381225 |
| GO_NCRNA_PROCESSING | 59 | 0.00000000 | 2.902466 |
| GO_MULTI_MULTICELLULAR_ORGANISM_PROCESS | 25 | 0.00020156 | 2.8365731 |
| GO_RNA_METABOLIC_PROCESS | 116 | 0.00016797 | 2.823444 |
| GO_RRNA_METABOLIC_PROCESS | 40 | 0.00046876 | 2.7766144 |
| GO_EXTRACELLULAR_STRUCTURE_ORGANIZATION | 38 | 0.00133046 | 2.6083918 |
| GO_ORGANIC_ANION_TRANSPORT | 24 | 0.00286321 | 2.4985893 |
| GO_ANION_TRANSPORT | 27 | 0.00401584 | 2.451998 |
| GO_CHROMOSOME_ORGANIZATION | 123 | 0.01066266 | 2.3349226 |
| GO_RNA_SPLICING_VIA_TRANSESTERIFICATION_REACTIONS | 25 | 0.01938850 | 2.247905 |
| GO_MONOVALENT_INORGANIC_CATION_TRANSPORT | 22 | 0.01883351 | 2.2357714 |
| GO_RNA_EXPORT_FROM_NUCLEUS | 15 | 0.02472816 | 2.1926205 |
| GO_ION_TRANSPORT | 75 | 0.02584514 | 2.1751554 |
| GO_POSITIVE_REGULATION_OF_CELL_POPULATION_PROLIFERATION | 68 | 0.02550212 | 2.1687748 |
| GO_CARTILAGE_DEVELOPMENT | 15 | 0.02772710 | 2.1453285 |
| GO_POSITIVE_REGULATION_OF_CHROMOSOME_ORGANIZATION | 18 | 0.03081124 | 2.1209855 |
| GO_DNA_METABOLIC_PROCESS | 86 | 0.03137370 | 2.1112738 |
| GO_TELOMERE_ORGANIZATION | 21 | 0.03771757 | 2.0796552 |
| GO_MYELOID_LEUKOCYTE_DIFFERENTIATION | 19 | 0.03901365 | 2.0671132 |
| GO_CHROMATIN_ORGANIZATION | 77 | 0.04782771 | 2.0270483 |
| GO_MAMMARY_GLAND_DEVELOPMENT | 15 | 0.04807314 | 2.0176482 |
| GO_REGULATION_OF_CHROMOSOME_ORGANIZATION | 42 | 0.04689902 | 2.0142443 |
| GO_REPRODUCTIVE_SYSTEM_DEVELOPMENT | 31 | 0.04576272 | 2.0116198 |

**Table S4. GSEA positive in MATR3 KO line.** Gene set name, size, False Discovery Rate (FDR q-val <0.25) and normalized enrichment score (NES > 2) are listed. Immune gene sets are labeled in red. C5.bp.v7.0 database was used for GSEA.

| Gene set | Size | FDR q-val | NES |
|---|---|---|---|
| GO_BIOLOGICAL_ADHESION | 82 | 0.00000000 | 3.8074 |
| GO_EXTRACELLULAR_STRUCTURE_ORGANIZATION | 32 | 0.00000000 | 3.740174 |
| GO_CELL_SUBSTRATE_ADHESION | 37 | 0.00000000 | 3.6521194 |
| GO_CELL_JUNCTION_ASSEMBLY | 27 | 0.00000000 | 3.4387255 |
| GO_LOCOMOTION | 108 | 0.00000000 | 3.4379458 |
| GO_CELL_MOTILITY | 94 | 0.00000000 | 3.376674 |
| GO_CELL_JUNCTION_ORGANIZATION | 33 | 0.00000000 | 3.3555205 |
| GO_TUBE_DEVELOPMENT | 65 | 0.00000000 | 3.272315 |
| GO_REGULATION_OF_CELL_ADHESION | 51 | 0.00000000 | 3.1593573 |
| GO_REGULATION_OF_CELL_SUBSTRATE_ADHESION | 28 | 0.00000000 | 3.1544757 |
| GO_CARDIOVASCULAR_SYSTEM_DEVELOPMENT | 48 | 0.00000000 | 3.0204344 |
| GO_ANATOMICAL_STRUCTURE_FORMATION_INVOLVED_IN_MORPHOGENESIS | 66 | 0.00000000 | 3.005266 |
| GO_TUBE_MORPHOGENESIS | 51 | 0.00000000 | 3.0014138 |
| GO_REGULATION_OF_CELLULAR_COMPONENT_MOVEMENT | 62 | 0.00000000 | 2.9624763 |
| GO_CELL_MATRIX_ADHESION | 26 | 0.00009069 | 2.8586621 |
| GO_BLOOD_VESSEL_MORPHOGENESIS | 41 | 0.00008502 | 2.8471246 |
| GO_RESPONSE_TO_MECHANICAL_STIMULUS | 17 | 0.00008002 | 2.8430831 |
| GO_TAXIS | 28 | 0.00007558 | 2.8426826 |
| GO_REGULATION_OF_CELL_DEVELOPMENT | 57 | 0.00014729 | 2.803229 |
| GO_NEGATIVE_REGULATION_OF_DEVELOPMENTAL_PROCESS | 49 | 0.00013992 | 2.7866838 |
| GO_NEGATIVE_REGULATION_OF_CELL_ADHESION | 19 | 0.00013326 | 2.7527258 |
| GO_CELL_MORPHOGENESIS_INVOLVED_IN_DIFFERENTIATION | 54 | 0.00012720 | 2.7330213 |
| GO_ACTIN_FILAMENT_BASED_PROCESS | 65 | 0.00012167 | 2.7159915 |
| GO_POSITIVE_REGULATION_OF_DEVELOPMENTAL_PROCESS | 74 | 0.00011660 | 2.6974292 |
| GO_NEUROGENESIS | 98 | 0.00022214 | 2.664955 |
| GO_EMBRYONIC_MORPHOGENESIS | 23 | 0.00054876 | 2.6016457 |
| GO_SYNAPSE_ORGANIZATION | 21 | 0.00052843 | 2.5974023 |
| GO_CELLULAR_COMPONENT_MORPHOGENESIS | 85 | 0.00066410 | 2.5595744 |
| GO_RESPONSE_TO_WOUNDING | 44 | 0.00084072 | 2.540903 |
| GO_CIRCULATORY_SYSTEM_DEVELOPMENT | 64 | 0.00085536 | 2.5336614 |
| GO_NEGATIVE_REGULATION_OF_MULTICELLULAR_ORGANISMAL_PROCESS | 63 | 0.00109084 | 2.5139692 |
| GO_ERK1_AND_ERK2_CASCADE | 15 | 0.00105675 | 2.5136461 |
| GO_PHAGOCYTOSIS | 24 | 0.00102473 | 2.5087564 |
| GO_REGULATION_OF_CELL_MORPHOGENESIS_INVOLVED_IN_DIFFERENTIATION | 25 | 0.00099459 | 2.5073178 |
| GO_EMBRYO_DEVELOPMENT | 56 | 0.00112199 | 2.4968815 |
| GO_CELL_PART_MORPHOGENESIS | 48 | 0.00109082 | 2.495469 |
| GO_REGULATION_OF_NEURON_DIFFERENTIATION | 42 | 0.00109895 | 2.4897158 |
| GO_POSITIVE_REGULATION_OF_CELL_ADHESION | 27 | 0.00110861 | 2.4888318 |
| GO_REGULATION_OF_CELL_MORPHOGENESIS | 42 | 0.00129257 | 2.475817 |
| GO_CELL_SUBSTRATE_JUNCTION_ASSEMBLY | 18 | 0.00126026 | 2.4721944 |

| | | | |
|---|---|---|---|
| GO_NEURON_DIFFERENTIATION | 82 | 0.00146520 | 2.4514828 |
| GO_REGULATION_OF_CELL_PROJECTION_ORGANIZATION | 49 | 0.00153647 | 2.447087 |
| GO_REGULATION_OF_PEPTIDYL_TYROSINE_PHOSPHORYLATION | 19 | 0.00150074 | 2.446442 |
| GO_LEUKOCYTE_MIGRATION | 16 | 0.00146663 | 2.4454727 |
| GO_REGULATION_OF_VASCULATURE_DEVELOPMENT | 21 | 0.00152906 | 2.4396653 |
| GO_REGULATION_OF_CELL_MATRIX_ADHESION | 17 | 0.00175695 | 2.4130304 |
| GO_POSITIVE_REGULATION_OF_CELL_DEVELOPMENT | 28 | 0.00171957 | 2.4109375 |
| GO_POSITIVE_REGULATION_OF_CELLULAR_COMPONENT_BIOGENESIS | 35 | 0.00171301 | 2.4076817 |
| GO_AXON_DEVELOPMENT | 32 | 0.00167805 | 2.406844 |
| GO_REGULATION_OF_ANATOMICAL_STRUCTURE_MORPHOGENESIS | 86 | 0.00195372 | 2.3901455 |
| GO_NEURON_DEVELOPMENT | 70 | 0.00199665 | 2.3890295 |
| GO_NEURON_PROJECTION_GUIDANCE | 15 | 0.00204070 | 2.3779175 |
| GO_WOUND_HEALING | 38 | 0.00202959 | 2.3717127 |
| GO_NEGATIVE_REGULATION_OF_CELL_MOTILITY | 18 | 0.00209281 | 2.3678942 |
| GO_NEGATIVE_REGULATION_OF_LOCOMOTION | 20 | 0.00208054 | 2.3658898 |
| GO_REGULATION_OF_NERVOUS_SYSTEM_DEVELOPMENT | 50 | 0.00224689 | 2.3545246 |
| GO_CELL_MORPHOGENESIS_INVOLVED_IN_NEURON_DIFFERENTIATION | 38 | 0.00220747 | 2.3539891 |
| GO_CELL_PROJECTION_ORGANIZATION | 94 | 0.00224031 | 2.3488207 |
| GO_ORGANIC_ACID_BIOSYNTHETIC_PROCESS | 24 | 0.00224813 | 2.34642 |
| GO_CYTOSKELETON_ORGANIZATION | 111 | 0.00221066 | 2.3444626 |
| GO_TRANSMEMBRANE_RECEPTOR_PROTEIN_TYROSINE_KINASE_SIGNALING_PATHWAY | 52 | 0.00228931 | 2.3351343 |
| GO_ENZYME_LINKED_RECEPTOR_PROTEIN_SIGNALING_PATHWAY | 62 | 0.00256801 | 2.3200638 |
| GO_NEGATIVE_REGULATION_OF_CELL_DIFFERENTIATION | 41 | 0.00268231 | 2.3079326 |
| GO_REGULATION_OF_CELLULAR_COMPONENT_BIOGENESIS | 66 | 0.00279110 | 2.2989564 |
| GO_CELL_CELL_ADHESION | 36 | 0.00296390 | 2.287594 |
| GO_REGULATION_OF_NEURON_PROJECTION_DEVELOPMENT | 36 | 0.00333625 | 2.2667968 |
| GO_POSITIVE_REGULATION_OF_NERVOUS_SYSTEM_DEVELOPMENT | 22 | 0.00452171 | 2.2236817 |
| GO_ADHERENS_JUNCTION_ORGANIZATION | 18 | 0.00468310 | 2.218263 |
| GO_PEPTIDYL_TYROSINE_MODIFICATION | 28 | 0.00539489 | 2.1945512 |
| GO_REGULATION_OF_MAPK_CASCADE | 37 | 0.00557408 | 2.1869726 |
| GO_NEGATIVE_REGULATION_OF_CELL_DEVELOPMENT | 25 | 0.00596092 | 2.1707952 |
| GO_MULTI_MULTICELLULAR_ORGANISM_PROCESS | 21 | 0.00603352 | 2.1664622 |
| GO_POSITIVE_REGULATION_OF_LOCOMOTION | 32 | 0.00599149 | 2.165709 |
| GO_DEVELOPMENTAL_CELL_GROWTH | 18 | 0.00836984 | 2.1187744 |
| GO_REGULATION_OF_CELL_DIFFERENTIATION | 109 | 0.01087425 | 2.0792742 |
| GO_EPITHELIAL_CELL_PROLIFERATION | 16 | 0.01078771 | 2.0785904 |
| GO_POSITIVE_REGULATION_OF_MULTICELLULAR_ORGANISMAL_PROCESS | 77 | 0.01275046 | 2.0450432 |
