## Supplementary Table S5 for "Three ALS genes regulate expression of the MHC class II antigen presentation pathway"

**Table S5. Shared proteins dysregulated in FUS, TAF15, MATR3 and EWSR1 KO HeLa lines from quantitative mass spectrometry data.**
Proteins with a fold change >1.5 and a p-value < 0.05 are listed. Green and red highlighing indicate down and upregulated proteins, respectively.

| Protein | Fold Change | | | | p-value | | | |
|---|---|---|---|---|---|---|---|---|
| | FUS KO | TAF15 KO | MATR3 KO | EWSR1 KO | FUS KO | TAF15 KO | MATR3 KO | EWSR1 KO |
| **Downregulated** | | | | | | | | |
| HLA-DRA | -13.0 | -17.4 | -9.6 | 1.3 | 0.018 | 0.018 | 0.020 | 0.018 |
| HLA-DRB1 | -10.4 | -11.1 | -8.3 | 1.3 | 0.012 | 0.012 | 0.013 | 0.012 |
| TYRO3 | -6.7 | -2.6 | -2.8 | -1.3 | 0.004 | 0.008 | 0.006 | 0.039 |
| S100P | -4.9 | -9.3 | -3.2 | 2.3 | 0.002 | 0.002 | 0.003 | 0.002 |
| PSMB9 | -4.5 | -4.5 | -3.0 | 1.8 | 0.004 | 0.000 | 0.011 | 0.003 |
| PSMB10 | -4.2 | -3.8 | -2.5 | 1.1 | 0.002 | 0.005 | 0.006 | 0.080 |
| LXN | -3.8 | -3.0 | -2.2 | 1.1 | 0.002 | 0.000 | 0.008 | 0.148 |
| TRIM29 | -3.8 | -2.9 | -2.3 | -1.7 | 0.002 | 0.007 | 0.005 | 0.010 |
| DAB2 | -3.7 | -2.4 | -2.5 | -1.1 | 0.011 | 0.018 | 0.016 | 0.238 |
| SAMD11 | -3.6 | -1.7 | -2.6 | 1.1 | 0.000 | 0.045 | 0.001 | 0.017 |
| SYNPO2 | -3.1 | -6.0 | -2.2 | -4.4 | 0.027 | 0.018 | 0.041 | 0.020 |
| ZBED1 | -2.7 | -1.8 | -1.9 | 1.6 | 0.001 | 0.004 | 0.002 | 0.004 |
| MYPN | -2.7 | -2.6 | -2.2 | 1.1 | 0.014 | 0.019 | 0.021 | 0.296 |
| FN3K | -2.6 | -2.2 | -2.0 | 1.0 | 0.008 | 0.011 | 0.008 | 0.930 |
| PSMB8 | -2.3 | -1.9 | -1.6 | 1.6 | 0.007 | 0.006 | 0.018 | 0.005 |
| ALDH5A1 | -2.1 | -1.9 | -2.7 | 1.3 | 0.022 | 0.019 | 0.011 | 0.034 |
| CLU | -2.1 | -1.5 | -1.5 | 1.5 | 0.011 | 0.024 | 0.021 | 0.011 |
| PTER | -2.1 | -1.6 | -1.6 | -1.8 | 0.008 | 0.013 | 0.016 | 0.011 |
| CSRP2 | -2.0 | -1.8 | -2.2 | -2.5 | 0.009 | 0.018 | 0.006 | 0.003 |
| MAP7 | -2.0 | -1.7 | -1.6 | 1.0 | 0.016 | 0.022 | 0.018 | 0.454 |
| RNASEL | -1.9 | -1.6 | -2.0 | 2.0 | 0.004 | 0.040 | 0.020 | 0.005 |
| RDX | -1.9 | -1.7 | -1.5 | 1.1 | 0.005 | 0.004 | 0.010 | 0.108 |
| PIR | -1.8 | -1.8 | -1.7 | 1.0 | 0.010 | 0.012 | 0.008 | 0.830 |
| CDKN2C | -1.8 | -1.8 | -1.6 | -1.1 | 0.026 | 0.029 | 0.033 | 0.561 |
| ALDH5A1B | -1.7 | -1.9 | -1.5 | -1.2 | 0.014 | 0.011 | 0.016 | 0.082 |
| KCTD15 | -1.6 | -2.5 | -1.5 | -1.3 | 0.004 | 0.003 | 0.005 | 0.019 |
| TSC22D3 | -1.6 | -2.0 | -1.5 | 1.2 | 0.003 | 0.025 | 0.001 | 0.027 |
| TMOD1 | -1.6 | -1.9 | -1.8 | 1.1 | 0.023 | 0.014 | 0.017 | 0.249 |

| **Upregulated** | | | | | | | | |
|---|---|---|---|---|---|---|---|---|
| | FUS KO | TAF15 KO | MATR3 KO | EWSR1 KO | FUS KO | TAF15 KO | MATR3 KO | EWSR1 KO |
| SERPINE2 | 6.8 | 3.5 | 2.6 | 1.0 | 0.006 | 0.006 | 0.002 | 0.414 |
| ANPEP | 5.9 | 2.4 | 5.0 | 1.1 | 0.007 | 0.014 | 0.000 | 0.087 |
| LAMB1 | 3.4 | 2.8 | 2.0 | -1.2 | 0.017 | 0.007 | 0.021 | 0.333 |
| PRSS23 | 2.9 | 2.8 | 1.9 | 1.1 | 0.004 | 0.015 | 0.015 | 0.444 |
| DMBT1 | 2.8 | 3.3 | 2.2 | 1.3 | 0.005 | 0.010 | 0.021 | 0.121 |

| | | | | | | | | |
|---|---|---|---|---|---|---|---|---|
| SH3KBP1 | 2.5 | 2.0 | 1.8 | 2.5 | 0.013 | 0.004 | 0.015 | 0.001 |
| STC2 | 2.4 | 1.7 | 2.0 | 1.3 | 0.004 | 0.002 | 0.002 | 0.024 |
| TGFBI | 2.3 | 1.8 | 1.7 | -2.7 | 0.014 | 0.004 | 0.019 | 0.007 |
| NES | 2.1 | 2.4 | 2.3 | -2.1 | 0.004 | 0.005 | 0.003 | 0.012 |
| TNC | 2.0 | 2.6 | 1.7 | -2.2 | 0.011 | 0.006 | 0.013 | 0.021 |
| RIN1 | 1.9 | 1.7 | 2.0 | 1.3 | 0.007 | 0.009 | 0.007 | 0.024 |
| FMNL2 | 1.6 | 2.0 | 1.6 | 1.1 | 0.023 | 0.016 | 0.009 | 0.479 |
