## Supplementary Table S6 for "Three ALS genes regulate expression of the MHC class II antigen presentation pathway"

| Table S6. Sequences of primers. | |
|---|---|
| **Name** | **Sequence (5->3')** |
| GAPDH-F | GTCAAGGCTGAGAACGGGAA |
| GAPDH-R | AAATGAGCCCCAGCCTTCTC |
| HLA-A-F | GGAGCAGAGATACACCTGCC |
| HLA-A-R | GTGAGGGACACATCAGAGCC |
| HLA-B-F | GGAGAAGAGCAGAGATACACATGC |
| HLA-B-R | AGCGACCACAGCTCCGATG |
| HLA-DRA-F | AAGGGATTGCGCAAAAGCAA |
| HLA-DRA-R | TGCTTTCACTGAGGTCAAGG |
| HLA-DRB1-F | TGGGTGGAGGGGTTCATAGT |
| HLA-DRB1-R | AAGTATCTGTCCAGGAACCGTG |
| CD74-F | GAGTGGCCTTCTGTGGACGA |
| CD74-R | GGAGATAAGGTCGCGCTGGT |
| CIITA-ex7-F | CCTAGTGGGACCAGTGAGCG |
| CIITA-ex8-R | AGCCTCAGAGATTTGCCAGAG |
| NLRC5-ex3-F | GTTCTTCCTCCCCAACACGG |
| NLRC5-ex3-R | ATGAAAGACTGCCAGGTGTCC |
| NANOG-F | TCCTTCCTCCATGGATCTGC |
| NANOG-R | TCTGCTGGAGGCTGAGGTAT |
| CD43-1-F | GTGGTAAGCCCAGACGCTC |
| CD43-1-R | GGCACCAATGGAAGTCCAAA |
| CD34-F | CCTCAGTGTCTACTGCTGGTCT |
| CD34-R | GGAATAGCTCTGGTGGCTTGCA |
| FUS-F | AGCAGTTCTCAGAGCAGCAG |
| FUS-R | CACTGCCACCACCACTACTC |
| TAF15-F | TACGGTCAGTCTGGGGGTGA |
| TAF15-R | TGTCCATAACTGGAGTAACCGC |
| MATR3-F | GGGCATCCTTCACCCATCTGA |
| MATR3-R | AGCTGAGAACCAGCAGACAACT |
